# The pelvic organs receive no parasympathetic innervation

**DOI:** 10.1101/2023.07.11.548500

**Authors:** Margaux Sivori, Bowen Dempsey, Zoubida Chettouh, Franck Boismoreau, Maïlys Ayerdi, Annaliese Nucharee Eymael, Sylvain Baulande, Sonia Lameiras, Fanny Coulpier, Olivier Delattre, Hermann Rohrer, Olivier Mirabeau, Jean-François Brunet

**Affiliations:** Institut de Biologie de l’ENS (IBENS), Inserm, CNRS, École normale supérieure, PSL Research University, Paris, France; Faculty of Medicine, Health & Human Sciences, Macquarie University, Macquarie Park, NSW, Australia; Institut Curie, PSL University, ICGex Next-Generation Sequencing Platform, 75005 Paris, France; GenomiqueENS, Institut de Biologie de l’ENS (IBENS), Département de biologie, École normale supérieure, CNRS, INSERM, Université PSL, 75005 Paris, France; Inserm U955, Mondor Institute for Biomedical Research (IMRB), Creteil, France; Institut Curie, Inserm U830, PSL Research University, Diversity and Plasticity of Childhood Tumors Lab, Paris, France; Institute of Clinical Neuroanatomy, Dr. Senckenberg Anatomy, Neuroscience Center, Goethe University, Frankfurt/M, Germany; Institut Pasteur, Université Paris Cité, Bioinformatics and Biostatistics Hub, 75015, Paris, France

**Keywords:** Autonomic nervous system, neuron types, transcriptomics, neural circuits

## Abstract

The pelvic organs (bladder, rectum and sex organs) have been represented for a century as receiving autonomic innervation from two pathways — lumbar sympathetic and sacral parasympathetic — by way of a shared relay, the pelvic ganglion, conceived as an assemblage of sympathetic and parasympathetic neurons. Using single cell RNA sequencing, we find that the mouse pelvic ganglion is made of four classes of neurons, distinct from both sympathetic and parasympathetic ones, albeit with a kinship to the former, but not the latter, through a complex genetic signature. We also show that spinal lumbar preganglionic neurons synapse in the pelvic ganglion onto equal numbers of noradrenergic and cholinergic cells, both of which therefore serve as sympathetic relays. Thus, the pelvic viscera receive no innervation from parasympathetic or typical sympathetic neurons, but instead from a divergent tail end of the sympathetic chains, in charge of its idiosyncratic functions.

## Introduction

The pelvic ganglion is a collection of autonomic neurons, close to the walls of the bladder, organized as a loose ganglionated plexus (as in humans) or a bona fide ganglion (as in mouse). It receives input from preganglionic neurons of the throracolumbar intermediate lateral motor column, a sympathetic pathway that travels through the hypogastric nerve. A second input is from the so-called “sacral parasympathetic nucleus” that projects through the pelvic nerve. The assignment of these two inputs to different divisions of the autonomic nervous system by John Langley (1, 2), largely based on physiological observation on external genitals which were never generally accepted (reviewed in (*3*), has led, in turn, to propose that the pelvic ganglion, targeted by both pathways, is of a mixed sympathetic/parasympathetic nature (*4*). Of note, this unique case of anatomic promiscuity between the two types of neurons poses a challenge for the schematic representation of the autonomic nervous system, so that the pelvic ganglion is most often omitted from such representations (*3*). A duality of the pelvic ganglion was also suggested by the coexistence of noradrenergic and cholinergic neurons, which elsewhere in the autonomic nervous system form, respectively, the vast majority of sympathetic ganglionic cells and the totality of parasympathetic ones. Here we directly explore the composition of the mouse pelvic ganglion in cell types, using single-cell transcriptomics, and compare it to sympathetic and parasympathetic ganglia.

## Results

We isolated cells from several autonomic ganglia of postnatal day 5 mice: the stellate ganglion and the lumbar chain (both belonging to the paravertebral sympathetic chain), the coeliac+mesenteric ganglia (belonging to the prevertebral ganglia), the sphenopalatine ganglion (parasympathetic) and the pelvic ganglion, and processed them for single-cell RNA sequencing (cf Material and Methods). Neuronal cells segregated in three large ensembles (Fig. 1A; Fig. S1): one that contained all sympathetic neurons, one that contained all parasympathetic neurons, and the third that contained most pelvic neurons — except one subset that segregated close to the sympathetic cluster. Thus, no pelvic neuron segregates with parasympathetic neurons, but the great majority of them do not segregate with sympathetic neurons either.

**Figure 1.**
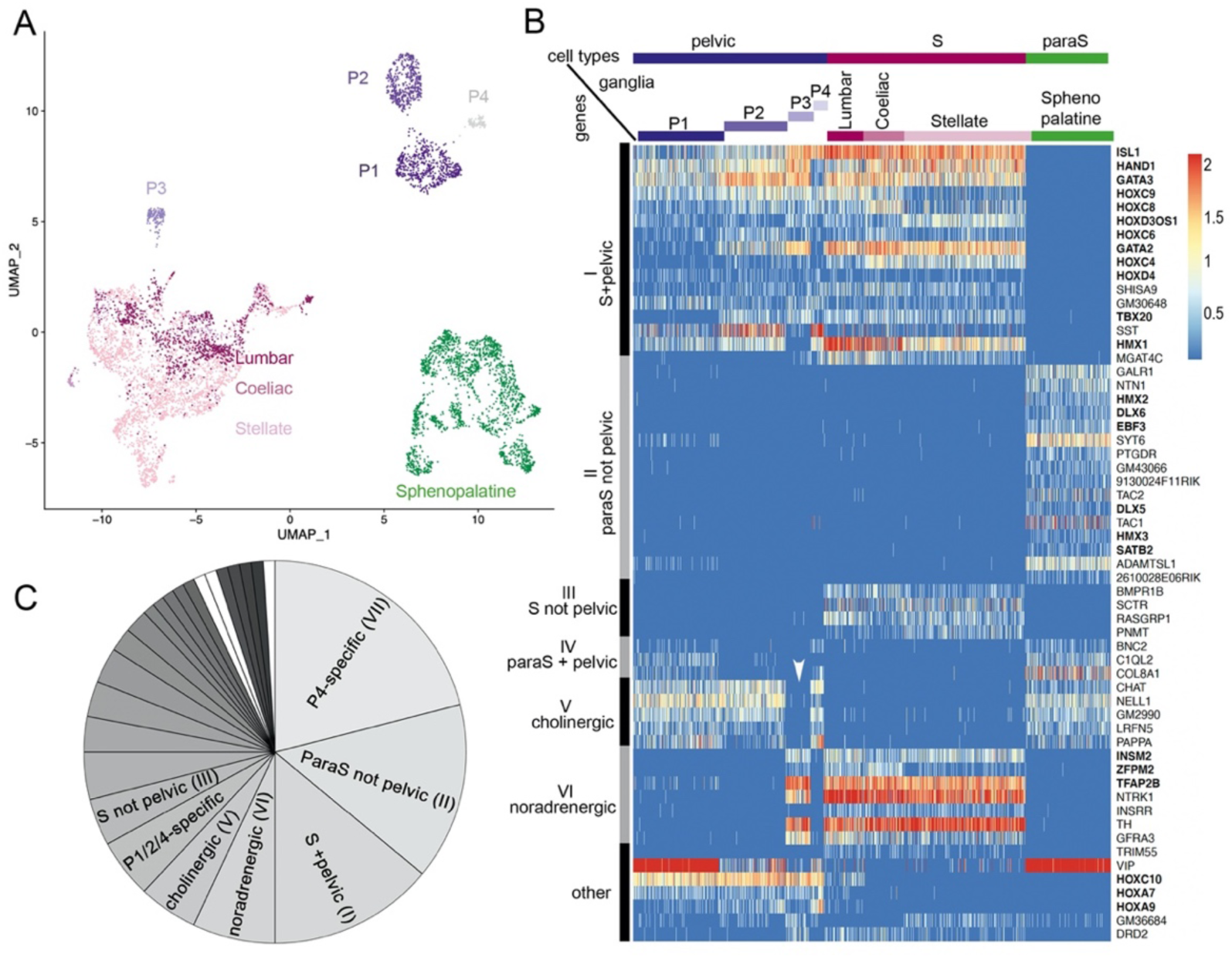
The pelvic ganglion does not contain parasympathetic neurons and is made of sympathetic-like neurons. (A) UMAP of cells isolated from 3 sympathetic ganglia (lumbar, stellate and celiac), a parasympathetic ganglion (sphenopalatine) and the pelvic ganglion dissected from postnatal day 5 mice. The pelvic ganglion is sharply divided into 4 clusters (P1-4), none of which co-segregates with sympathetic or parasympathetic neurons. (B) Heatmap of the highest scoring 100 genes in an all-versus-all comparison of their dichotomized expression pattern among the 4 ganglia and 4 pelvic clusters (see Material and Methods), excluding genes specific to the pelvic ganglion (shown in Fig. S2), and keeping only the top-scoring comparison for genes that appear twice. For overall legibility of the figure, the three largest cell groups (lumbar, stellate and sphenopalatine) are subsampled and genes are ordered by expression pattern (designated on the left), rather than score. “cholinergic” and “noradrenergic” genes are those that are coregulated with ChAT or Th, regardless of known function. “Other“ refers to various groupings that split sympathetic ganglia and are thus not informative as to a sympathetic or parasympathetic identity. Transcription factors are indicated in bold face. White arrowhead: pelvic P3 cluster; S, sympathetic; ParaS, parasympathetic. (C) Pie chart of the top 100 genes, counted by expression pattern in the all-versus-all comparison. Genes specific for the P4 cluster dominate (see heatmap in Fig. S1), followed by those which are “parasympathetic-not-pelvic” and “sympathetic-and-pelvic” (seen in 1B). The three genes marked in white are the only ones that are compatible with the current dogma of a mixed sympathetic/parasympathetic pelvic ganglion, by being expressed in the sphenopalatine ganglion and a subset of pelvic clusters (other than the full complement of cholinergic ones).

The separation, on the UMAP, of most pelvic from all other ganglionic cells contrasts with the suite of 5 developmental transcription factors that we previously reported as differentially expressed between the sympathetic and pelvic ganglia on one hand, and parasympathetic ganglia on the other (*5*). To explore this conundrum, we searched for more genes that would help place the pelvic ganglion relative to the sympatho-parasympathetic dichotomy: additional genes that would put the pelvic ganglion in the sympathetic category (as per our previous findings), genes that would put it in a class by itself (as the UMAP suggests), or genes that would split pelvic neurons into parasympathetic-like and sympathetic-like clusters (as the current dogma implies).

In an unbiased approach, we sought genes expressed in a higher proportion of cells in any set of ganglia compared to its complementary set (see Material and Methods). To do justice to the classical notion that the pelvic ganglion is heterogeneous, i.e. mixed sympatho/parasympathetic, we treated the clusters of pelvic ganglionic cells, 4 in our conservative estimate (P1-4) (Fig. 1A; Fig. S1) as 4 ganglia, alongside the 4 other ganglia (sphenopalatine, stellate, coeliac and lumbar) — i.e. we made (2^8^-2) = 254 comparisons and analyzed the 100 “top” genes, i.e. with the highest discrimination score, irrespective of the comparison scored (Table S1, S2, S3).

The vast majority of the top 100 genes fell into 7 of the 254 possible dichotomized expression patterns, visualized on a heatmap (patterns I-VI, Fig. 1B,C; pattern VII, Fig. S2) where cells are grouped by ganglion, and genes by expression pattern (i.e. disregarding their score). These 7 patterns can be consolidated into 3, among which only the first is informative about a sympathetic or parasympathetic identity of pelvic neurons:

i. Patterns I-IV comprise 39 genes with an opposite status in sympathetic and parasympathetic cells, among which 20 (I+III) are sympathetic-specific and 19 (II+IV) parasympathetic-specific (Fig. 1B,C). Those which are also expressed in pelvic clusters (I and IV) argue for their sympathetic (conversely parasympathetic) identity, and against the opposite identity; those which are not expressed in pelvic clusters (II and III), argue solely against such an identity. Overall, 32 genes (I+II) argue against a parasympathetic identity of all pelvic clusters, among which 16 genes (I) argue for their sympathetic identity; 7 genes (III+IV) argue against a sympathetic identity of all clusters, among which 3 genes (IV) argue for a parasympathetic identity of P1 or P4 (which are otherwise not parasympathetic by the criterion of 31 and 32 genes, respectively). We verified expression of 7 sympathetic and 7 parasympathetic markers by in situ hybridization on the pelvic ganglion at E16.5 (Fig. 2).
ii. Patterns V+VI comprise 12 genes that correlate with neurotransmitter phenotype (cholinergic or noradrenergic) by being expressed in the sphenopalatine ganglion and P1, P2 and P4 (5 genes (V), including ChAT) or, conversely, in all sympathetic ganglia and P3 (7 genes (VI), including Th) (Fig. 1B,C). These genes point to noradrenergic and cholinergic “synexpression groups”(*6*) broader than the defining biosynthetic enzymes and transporters. Neurotransmitter phenotype has been suggested to largely coincide with origin of input, lumbar or sacral (*7*) which currently defines, respectively, sympathetic versus parasympathetic identity (*8*). However, we find that the lumbar sympathetic pathway targets both cholinergic and noradrenergic pelvic ganglionic cells (in a proportion than reflects their relative abundance), by anterograde tracing (Fig. 3). Thus, noradrenergic and cholinergic genes are excluded from a debate on the sympathetic or parasympathetic identity of pelvic neurons.
iii. Pattern VII involves 42 genes that place all, or some pelvic clusters in a class by themselves (Fig. S2), thus neither sympathetic nor parasympathetic (as does the whole transcriptome, as evidenced by the UMAP (Fig. 1A)) by being expressed, or not expressed, exclusively in them. The cholinergic cluster P4 was particularly rich in genes with such idiosyncratic expression states (20 genes). Fig. S3 provides the 5 top genes of each pelvic cluster.

**Figure 2.**
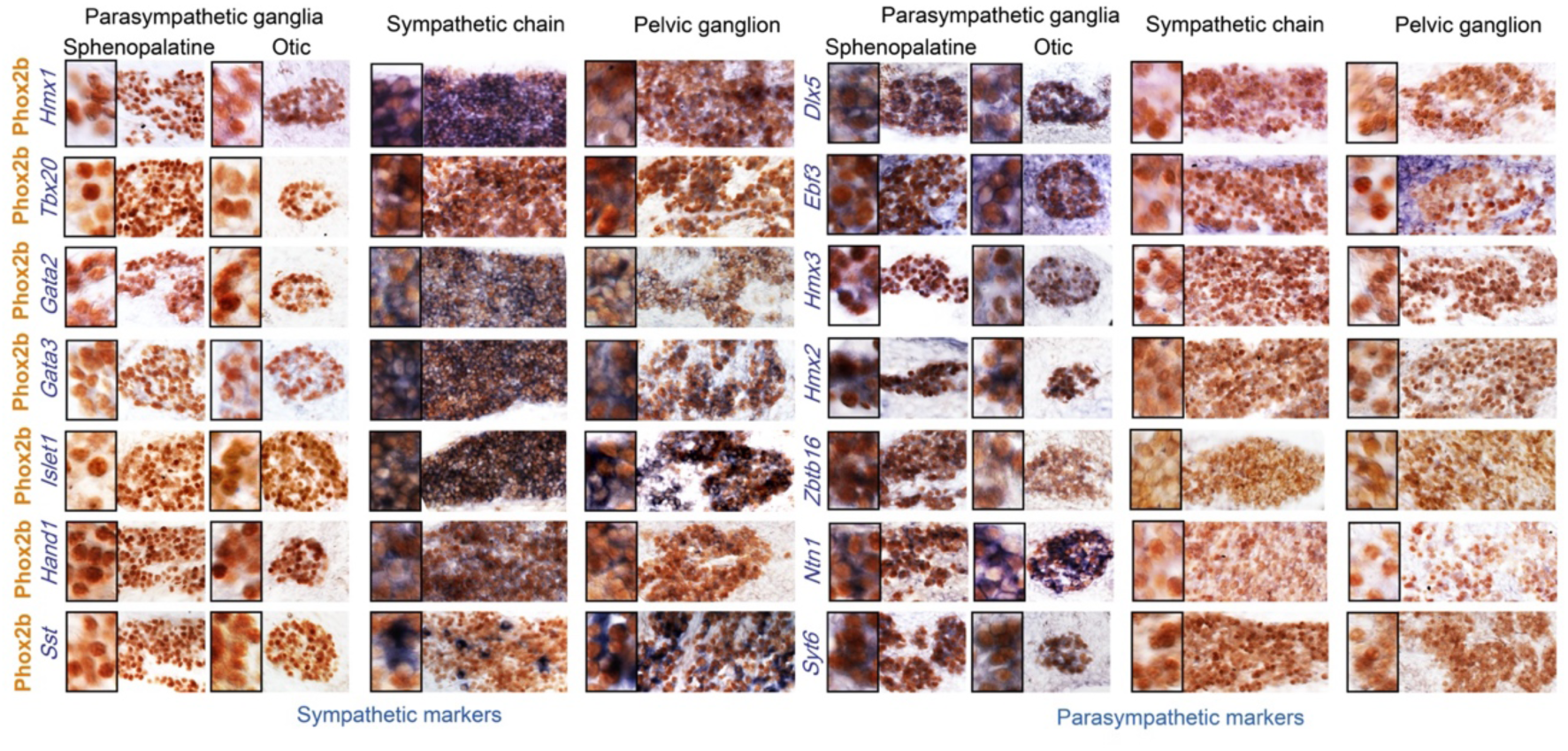
Pelvic ganglion cells express sympathetic but not parasympathetic markers. Combined immunohistochemistry for Phox2b and in situ hybridization for 7 sympathetic markers including 6 transcription factors (left panels) or 7 parasympathetic markers including 5 transcription factors (right panels), in two parasympathetic ganglia (sphenopalatine and otic), the lumbar sympathetic chain, and the pelvic ganglion, at low and high magnifications (inset on the left) in E16.5 embryos. Ebf3 is expressed in both, the parasympathetic ganglia and the mesenchyme surrounding all ganglia. Sst is expressed in a salt and pepper fashion. Zbtb16, a zinc-finger transcriptional repressor, appeared after the 100 highest scorer gene of our screen, but was spotted as expressed in the sphenopalatine in Genepaint. Some transcription factors detected by the RNA-Seq screen at P5 (Satb2, Dlx6) were expressed below the detection limit by in situ hybridization at E16.5.

**Figure 3.**
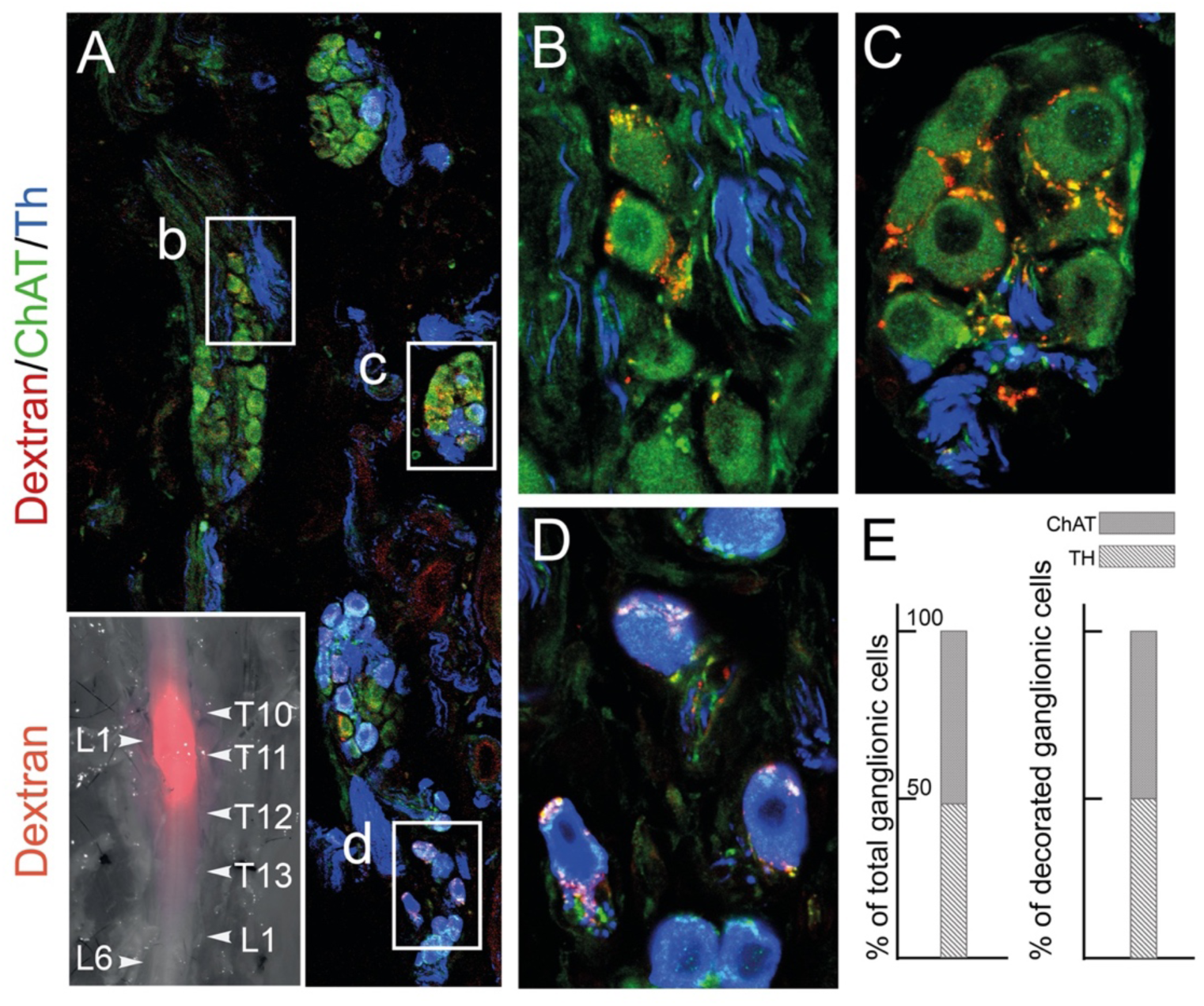
The lumbar outflow targets both cholinergic and noradrenergic pelvic ganglionic cells. (A-E) Section (A, low magnification, B-D high magnifications of selected regions) through the pelvic ganglion of an adult male mouse stereotactically injected with Dextran at the L1 level of the lumbar spinal cord (inset) and showing dextran filled boutons decorating both ChAT+ (B-C) and Th+ cells (D). Whether they are filled by Dextran or not, cholinergic boutons (green), presumably from spinal preganglionics (lumbar or sacral), are present on most cells. In the inset, levels of the vertebral column are indicated on the right, levels of the spinal cord on the left. (E) Quantification of Th and ChAT cells among total or bouton-decorated ganglionic cells. ChAT+ cells represent 51% of total cells and 50% of decorated cells (for a total of 3186 counted cells, among which 529 decorated cells, on 48 sections in 4 mice).

In the aggregate, the pelvic ganglion is best described as a divergent sympathetic ganglion, devoid of parasympathetic neurons. The fact that few of the top 100 genes (4 genes, III) marked sympathetic neurons to the exclusion of pelvic ones (Fig. 1B), while many (40 genes, VII) did the opposite (Fig. S2), argues that the pelvic identity is an evolutionary elaboration of a more generic, presumably more ancient, sympathetic one.

A third of the top 100 genes (32 genes) are transcription factors: 6 define a parasympathetic/non{sympathetic+pelvic} identity (Hmx2, Hmx3, Dlx5, Dlx6, Ebf3, Satb2); 12 define a {pelvic+sympathetic}/non-parasympathetic identity (Isl1, Gata2, Gata3, Hand1, Hmx1, Tbx20 plus 6 Hox genes); 3 correlate with the noradrenergic phenotype (Tfap2b, Insm2 and Zfpm2); 8 (including 6 Hox genes) are specific to the pelvic ganglion or some of its clusters (Fig. S2) and 3 Hox genes are specific to the pelvic+lumbar ganglia (Fig. 1B). Nothing is known of the function of the 6 parasympathetic transcription factors, but most of the non-Hox {sympathetic+pelvic} ones are implicated in sympathetic differentiation: Islet1 (*9*), Hand1(*10*), Gata2(*11*), Gata3 (*12*) and Hmx1(*13*).

Expression of a shared core of 12 transcription factors by pelvic and sympathetic ganglia, yet of divergent transcriptomes overall, logically calls for an additional layer of transcriptional control to modify the output of the 12 sympathetic transcription factors. Obvious candidates are the 6 Hox genes restricted to the pelvic ganglion (Hoxd9, Hoxa10, Hoxa5, Hoxd10, Hoxc11 and Hoxb3) (Fig. S2) and 6 more beyond the 100 top genes, mostly Hoxb paralogues (Fig. S4). Given that the parasympathetic and sympathetic ganglia are deployed according to a rostro-caudal pattern (parasympathetic ganglia in register with cranial nerves, sympathetic ganglia with thoraco-lumbar nerves, and the pelvic ganglion with lumbar and sacral nerves (Fig. 4), the entire taxonomy of autonomic ganglia could be a developmental readout of Hox genes (whose multiplicity makes this conjecture hard to test). A role for Hox genes in determining types of autonomic neurons would be reminiscent of their specification of other neuronal subtypes in Drosophila(*14*), C. elegans (*15*) and Mus (*16*).

**Figure 4:**
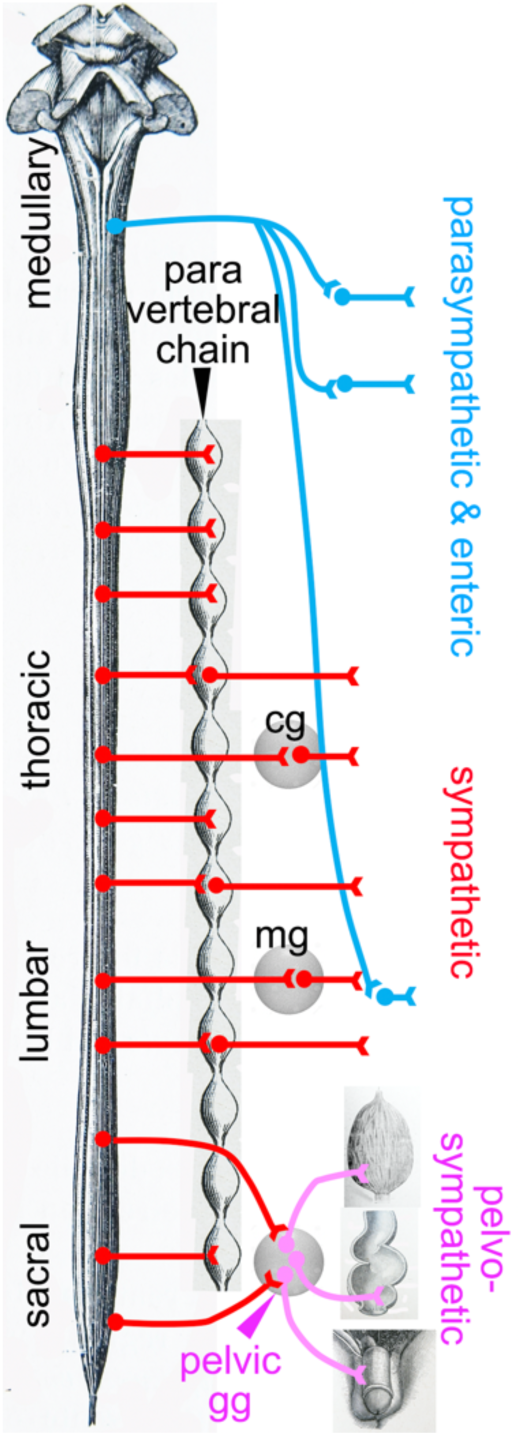
Deployment of the divisions of the autonomic nervous system on the rostro-caudal axis. Cg, celiac ganglion; mg, mesenteric ganglion; pelvic gg, pelvic ganglion. Only the target organs of the pelvo-sympathetic pathway are represented. The pelvic ganglion is shown with its lumbar input (through the hypogastric nerve), and sacral input (through the pelvic nerve).

## Discussion

The notion that the pelvic ganglion is a caudal elaboration of the sympathetic ganglionic chains echoes the situation of its preganglionic neurons, in the lumbar and sacral spinal cord. We showed that lumbar and sacral preganglionics are indistinguishable by several criteria, including the expression state of 6 transcription factors (Phox2a, Phox2b, Tbx3, Tbx2, Tbx20, FoxP1) and their dependency on the bHLH gene Olig2, in addition to their well-known location (the intermediate lateral column) and ventral exit point of their axon (*5*). Recent surveys of spinal preganglionic neurons by single-cell transcriptomics (*17*, *18*) discovered many subtypes, most of them evenly distributed from thoracic to sacral levels, and a few enriched at, or specific to the sacral level, confirming that the sacral preganglionic neurons are sympathetic, yet represent a caudal modification of the thoracolumbar intermediate lateral column.

In conclusion, the genetic signatures of neither pre-nor post-ganglionic neurons of the sacral autonomic outflow support its century-old assignment to the parasympathetic division of the visceral nervous system. Importantly, we have argued (*3*) that the supposed parasympathetic identity was unnecessary at best to understand the physiology of the pelvis, at worst incongruent with it, at least concerning sex organs and the bladder. The antagonism between an anti-erectile lumbar and a pro-erectile sacral autonomic pathways on the blood vessels of the external genitals — the key argument that Langley advanced for making the sacral pathway parasympathetic, even before he coined the term (*2*)— was repeatedly challenged by the evidence of a lumbo-sacral pro-erectile synergy (reviewed in (*3*)). And experimental support for a lumbo-sacral antagonism on the bladder is scant, despite common perception from textbooks and reviews (reviewed in (*3*)). It is thus remarkable that the cell type-based anatomy that we uncover in no way contradicts the physiology. There is no need to suppose that some sympathetic-like neurons in the pelvic region display a type of physiology common with that of parasympathetic ones in the head or thorax, which would deserve a common name. The classical parasympathetic label is thus best dropped and replaced by the genetic (i.e. embryological and evolutionary) notion of a sympathetic one, modified to suit the unique demands of pelvic physiology. The division of the autonomic nervous system that encompasses the pelvic ganglion and both its lumbar and sacral afferents (which could be termed “pelvo-sympathetic” and includes whatever antagonistic pathways operate in the region —e.g. on genital blood vessels) is now ripe for finer grain analysis, anatomical and physiological, at the level of neuron types, defined by their gene expression patterns.

## Materials and Methods

### Mice

Phox2b::Cre (*19*): a BAC transgenic line expressing Cre under the control of the Phox2b promoter. Rosa^lox-stop-lox-tdTomato^ (Rosa^tdT^)(*20*): Knock in line expressing the reporter gene tdTomato from the Rosa locus in a Cre-dependent manner.

### Obtainment of ganglionic cells

Sphenopalatine, stellate, coeliac, lumbar and pelvic ganglia were dissected from Phox2bCre;Rosa^tdT^ P5 pups representing both sexes and placed in artificial cerebrospinal fluid (aCSF) oxygenated with carbogen (4°C). Fat tissue was carefully removed and nerves emanating from the ganglia were cut. Ganglia were transferred into a 1.5ml Eppendorf tube containing 1ml PBS-Glucose (1mg/ml Glucose), 20 µl Papain solution (Worthington LS003126; 25.4 Units/mg) and 20 µl DNAse (2mg/ml in PBS) and incubated at 37°C for 15min. The ganglia were collected by centrifugation (300g, 1min), the supernatant replaced by 1ml PBS-Glucose supplemented with 50 µl Collagenase/Dispase (Worthington CLS-1 345U/mg; Dispase II Roche 1.2U/mg; 80mg Collagenase and 92mg Dispase II dissolved in 1ml PBS-Glucose) and 20 µl DNAse solution and the ganglia incubated at 37°C for 8min. After collecting the ganglia by centrifugation for 3min at 300g they were dissociated in 1ml PBS-Glucose supplemented with 0.04% bovine serum albumin (BSA) and DNAse (20µl) by trituration, using a fire-polished, siliconized Pasteur pipette. The cell suspension was then filtered through a 40µm cell strainer. To eliminate cell debris, the cell suspension was centrifuged through a density step gradient by overlaying the cell suspension onto 1ml OptiPrep solution (80 µl OptiPrep (Sigma), 900µl PBS supplemented with 0.04%BSA, 20µl DNAse) for 15 min, 100g at 5°C. After removal of the supernatant from the soft cell pellet, the cells were suspended in 100µl PBS-Glucose/0.04%BSA and collected again in 500µl Eppendorf tube for 15 min, 100g at 5°C. The supernatant was carefully removed (under control by fluorescence microscope). After addition of 40 µl PBS-Glucose/0.04%BSA the cell density was adjusted to 1000 cells/ml and transferred to the 10xGenomics platform.

### Library construction and Sequencing

Single cell RNA-Seq was performed in two separate experimental rounds, one for the stellate, sphenopalatine and pelvic ganglia (pelvic_1) performed at the École normale supérieure GenomiqueENS core facility (Paris, France), and one for the celiac, lumbar and pelvic ganglia (pelvic_2) performed at the ICGex NGS platform of the Institut Curie (Paris, France). Cellular suspensions (10000 cells for the first round, 5300 cells for the second) were loaded on a 10X Chromium instrument (10X Genomics) to generate single-cell GEMs (5000 for the first round, 3000 for the second). Single-cell RNA-Seq libraries were prepared using Chromium Single Cell 3’ Reagent Kit (v2 for the first round, v3 for the second) (10X Genomics) according to manufacturer’s protocol based on the 10X GEMCode proprietary technology. Briefly, the initial step consisted in performing an emulsion where individual cells were isolated into droplets together with gel beads coated with unique primers bearing 10X cell barcodes, UMI (unique molecular identifiers) and poly(dT) sequences. Reverse transcription reactions were applied to generate barcoded full-length cDNA followed by disruption of the emulsions using the recovery agent and the cDNA was cleaned up with DynaBeads MyOne Silane Beads (Thermo Fisher Scientific). Bulk cDNA was amplified using a GeneAmp PCR System 9700 with 96-Well Gold Sample Block Module (Applied Biosystems) (98°C for 3min; 12 cycles: 98°C for 15s, 63°C for 20s, and 72°C for 1min; 72°C for 1min; held at 4°C). The amplified cDNA product was cleaned up with the SPRI select Reagent Kit (Beckman Coulter). Indexed sequencing libraries were constructed using the reagents from the Chromium Single Cell 3’ Reagent Kit v3, in several steps: (1) fragmentation, end repair and A-tailing; (2) size selection with SPRI select; (3) adaptor ligation; (4) post ligation cleanup with SPRI select; (5) sample index PCR and cleanup with SPRI select beads (with 12 to 14 PCR cycles depending on the samples). Individual library quantification and quality assessment were performed using Qubit fluorometric assay (Invitrogen) with dsDNA HS (High Sensitivity) Assay Kit and Bioanalyzer Agilent 2100 using a High Sensitivity DNA chip (Agilent Genomics). Indexed libraries were then equimolarly pooled and quantified by qPCR using the KAPA library quantification kit (Roche). Sequencing was performed on a NextSeq 500 device (Illumina) for the first round and a NovaSeq 6000 (Illumina) for the second, targeting around 400M clusters per sample and using paired-end (26/57bp for the first round, 28×91bp for the second).

### Bioinformatic analysis

For each of the 6 samples (pelvic_1, stellate, sphenopalatine, pelvic_2, coeliac and lumbar) we performed demultiplexing, barcode processing, and gene counting using the Cell Ranger software (v. 6.0.1). The ‘filtered_feature_bc_matrix’ files (uploaded at https://www.ncbi.nlm.nih.gov/geo/query/acc.cgi?acc=GSE232789) were used as our starting points for defining cells. For each dataset, only droplets that expressed more than 1500 genes, less than 11000 genes and a percentage of mitochondrial genes below 15% were retained. This resulted in the selection of 4404 pelvic_1 cells, 7225 stellate cells, 4630 sphenopalatine cells, 1643 pelvic_2 cells, 1428 coeliac and 2120 lumbar cells.

We used Seurat version 3 (*21*) to read, manipulate, assemble, and normalize (*22*) the datasets. Specifically, we concatenated cells from all 6 datasets and normalized the data using the sctransform (SCT) method for which we fitted a Gamma-Poisson Generalized Linear Model (“glmGamPoi” option). Using the Seurat framework, we then performed a PCA on the normalized dataset.

Next, we integrated all 6 datasets using Batchelor (*23*) (a strategy for batch correction based on the detection of mutual nearest neighbors), and we visualized all cells, including neurons, in 2 dimensions using the Uniform Manifold Approximation and Projection method (*24*) on the first 50 components from the PCA.

We selected neurons based on their mean expression of a set of neuronal marker genes (Stmn2, Stmn3, Gap43 and Tubb3). Specifically, cells were classified as neurons if their mean marker SCT-normalized gene expression exceeded a threshold of 3, that best separated the bimodal distribution of the higher and lower values of the mean expression of neuronal markers. Likewise, we excluded glial-containing doublets using the following markers: Plp1, Ttyh1, Fabp7, Cryab and Mal. The number of neurons thus selected was 862 for Pelvic_1, 2689 for the stellate, 1857 for the sphenopalatine, 361 for Pelvic_2, 236 for celiac and 925 for the lumbar chain.

We then clustered pelvic neurons based on the first two components from the UMAP generated from the 50 batch-corrected components, using the graph-based clustering method Louvain (Blondel et al 2008) implemented in the Seurat framework, with a resolution of 0.3. This procedure defined 24 clusters and split the pelvic ganglion into four clusters 1, 7, 15 and 19 (Fig. S1B), that we renamed P1, P2, P3, P4 (Fig. 1A). The efficiency of batch correction is attested by the equal contribution from both batches of pelvic ganglia to all four defined pelvic clusters (P1-4) (Fig. S1A), and the comparable expression level of the top 100 genes across both pelvic batches in the violin plots (Table S2).

To search in an unbiased manner for gene expression similarities between pelvic neurons and either sympathetic or parasympathetic ones, we systematically compared the expression of every gene of the dataset across all possible splits among ganglia, i.e. between every subset of ganglia (“subset_1”), and its complementary subset (“subset_2”). Because we treated each of the 4 pelvic clusters as a ganglion, there were 2^8=256 such splits (2 of them, defined by “all_8_groups vs none” and “none vs all_8_groups”, being meaningless). For every split and every gene, we devised a score that would best reflect how higher the proportion is of cells expressing the gene (i.e. one read of the gene or more) in subset_1 than in subset_2. We used a metric based on the product of the proportions of cells expressing the gene in each ganglion of subset_1 or subset_2 (rather than throughout subset_1 or subset_2), to give an equal weight to each ganglion (irrespective of its size), and to penalize splits that contain outliers (i.e. ganglia that contain much fewer cells positive for that gene than other ganglia in subset_1, or ganglia that contain much more cells positive for that gene than other ganglia in subset_2).

Specifically, for a given split and a given gene, the score (here defined by the variable score, in the R programming language) is defined as:

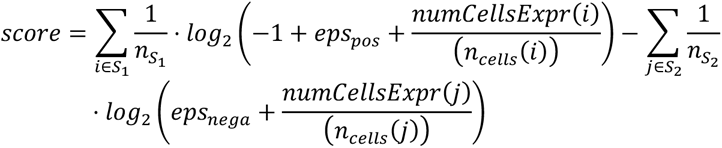

where:

- S_1_ and S_2_ are the ganglion subsets subset_1 and subset_2 as described above.
- n_S1_ and n_S2_ are the number of ganglion clusters that belong to subset_1 and subset_2.
- 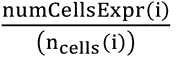 and 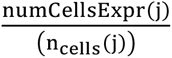 are the proportion of cells from ganglion or pelvic cluster i and j (i belonging to subset_1 and j to subset_2) which express the given gene.
- eps_pos_=0.9 and eps_nega_=0.02 are parameters controlling a tradeoff between favoring genes expressed in subset_1 and penalizing genes expressed in subset_2.
- Note that when eps_pos_=0.9 (the parameter value used here) the quantity 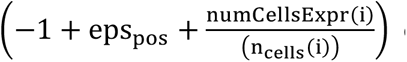 can be negative, in which case the expression becomes equal to -Inf and the gene gets the lowest possible score for that given split. This means that with these settings all dichotomies involving a ganglion or pelvic cluster in which less than 10% of cells express the gene (p=0.1) will get the minimal possible score.

### In vivo gene expression

#### Tissue preparation

Embryo sections. E16.5 WT embryos were freshly dissected, washed with 1X PBS and fixed overnight in 4% PFA at 4°C. Tissues were washed a few times in 1X PBS and treated in 15% sucrose to cryopreserve the tissues for frozen tissue sections. Tissues were then embedded in the Optimal Cutting Temperature (OCT) compound and snap frozen. Tissues were cut at 14 µm with cryostat. Tissue sections were stored at -80°C until use for staining.

#### In situ hybridization

Frozen tissue sections were washed in 5X SSC buffer 15min at RT and treated in the pre-hybridization solution (50% Formamide, 5X SSC buffer and 40 µg/mL Herring sperm DNA in H_2_O) for 1h at 60°C. Then, slides were put in the hybridization solution (50% Formamide, 5X SSC buffer, 5X Denhardt’s, 500 µg/mL Herring sperm DNA, 250 µg/mL Yeast RNA and 1mM DTT in H_2_O) containing the probe (100 ng/mL), at least 2 overnights at 60°C. Slides were washed two times in 5X SSC buffer (5min) and two times in 0.2X SSC buffer (30min) at 70°C, and then, three times in TBS at RT (10min). Then, tissues were put in the blocking solution (TBS + 10% FCS) for 1h in the dark, at RT and in humid atmosphere (250 µL/slide) and incubated 1h with the primary antibody (anti-DIG) diluted 1/200 in blocking solution (250 µL/slide). Then, slides were washed again three times in TBS (10min) and treated 5min in the AP buffer solution (100 nM Tris pH9.5, 50nM MgCl_2_ and 100 nM NaCl in H_2_O). The revelation was made by the NBT-BCIP solution (Sigma) in the dark (250 µL/slide). The reaction was stopped in PBS-Tween (PBST).

#### Immunohistochemistry

After in situ hybridization, frozen tissues were fixed 15 min in 4% PFA at RT and washed in PBST. Then, tissues were incubated in PBST + 10% FCS in the dark for 1h (500 µL/slide, without coverslip) and incubated overnight with the primary antibody in the same solution at 4°C (250 µL/slide). Slides were washed in PBST three times (10min) and incubated for 2h at RT with the secondary antibody in the same solution again. Tissues were washed three times in PBST and then incubated at RT with avidin/biotin solution diluted 1/100 in 1X PBS. Then, tissues were washed three times and the revelation was made in DAB solution containing 3,3’-Diaminobenzidine (DAB) and urea in H_2_O (Sigma). The reaction was stopped in PBST, and slides were washed in PBST and in ultra-pure H_2_O. Slides were mounted in Aquatex mounting medium (Merck).

#### Imaging

Tissues processed by an in situ hybridization and immunohistochemistry were photographed on a Leica bright field microscope with a 40X oil immersion objective. Images were then treated by Photoshop v. 24.1.0.

### Anterograde tracing

#### Tracer injections

Surgeries were conducted under aseptic conditions using a small animal digital stereotaxic instrument (David Kopf Instruments). Male, C57bl mice (2-4 months old) were anesthetized with isoflurane (3.5% at 1 l/min for induction and 2–3% at 0.3 l/min for maintenance). Carprofen (0.5 mg/kg) was administered subcutaneously for analgesia before surgery. A feed-back-controlled heating pad was used to maintain the animal temperature at 36 °C. Anesthetized animals were shaved along the spine and placed in a stereotaxic frame. The skin overlying the spine was sterilized with alternating scrubs of Vetadine and 80% (W/V) ethanol and a 100 µl injection of lidocaine (2%) was made subcutaneously along the spine before an incision was made from the iliac crest to the T9 vertebral spinous process. The T10 and T11 spinous processes were identified by counting rostrally from the L6 spinous process (identified relative to the iliac crest). Two small rostro-caudal incisions were made on each side of the vertebral column through the superficial-most layer of muscle. Parallel clamps were then placed within these incisions until firm contact was made with the transverse process of the T11 vertebra. The T11 vertebra was then raised upwards via the clamps until no respiratory movement was observed. The muscles connecting the T10 spinous process to transverse processes of caudal vertebra were dissected to expose the T10/T11 intervertebral space. A lateral incision was made across the dura to expose the underlying L1 spinal segment and 150nL injections of 4% lysine fixable, tetramethylrhodamine or Alexa-488 conjugated dextran, 3000MW (Thermofisher) were made bilaterally, into the intermediolateral nucleus (IML) and intermedio-medial nucleus (IMM) (0.600mm lateral, 0.600mm deep and 0.150mm lateral and 0.700mm deep) using a narrow-tapered glass pipette and a Nanoject III injector (Drummond Scientific). Injections were made at 5nL per second and left in place for 5 minutes following injection to prevent dextran leakage from the injection site. The pipette was then retracted, and the intervertebral space bathed with sterile physiological saline. A small piece of sterile gel foam hemostat (Pfizer) was then placed within the intervertebral space and the incision overlying the spine was sutured closed.

#### Histology

7-14 days after injection the mice were intracardially perfused with cold PBS until exsanguinated and subsequently fixed by perfusion with cold 4% PFA until the carcass became stiff. The bladder, prostate and pelvic ganglia were immediately dissected as an intact bloc and post fixed in 4% PFA overnight at 4 degrees. The intact spinal column was also dissected out and the dorsal surface of the spinal cord exposed before post fixation overnight in 4% PFA at 4 degrees. The fixed tissues were then rinsed in PBS 3× 30 minutes the following day. The “bladder block” was then cryopreserved in 15% sucrose (W/V) in PBS until non buoyant. The dorsal aspect of the injection site was imaged in situ using and fluorescence stereoscope. The spinal cord was then dissected out and cryopreserved as described above.

The bladder block was sectioned on a cryostat in a sagittal orientation to capture serial sections of the entire pelvic ganglion. Sections were cut at a thickness of 30µm and collected as a 1 in 4 series on Superfrost Plus glass slides. The spinal cord was sectioned coronally at a thickness of 60µm as a 1 in 2 series. On slide immunohistochemistry was performed against tyrosine hydroxylase (TH) and choline acetyltransferase (ChAT) for the bladder block sections, primary incubation: 4-6 hours at room temperature, secondary incubation: 2 hours at room temperature, with 3×5 minute washes in PBS after each incubation. Sections were then mounted with dako fluorescence mounting medium and cover slipped.

#### Imaging

Sections were imaged on a Leica Stellaris 5 confocal microscope (Leica microsystems). Images of the pelvic ganglia were captured as Z-stacks with 1µm interslice distances, with a 20x objective at 2000mp resolution.

#### Counting

All ganglionic cells, and cells surrounded by dextran labelled varicosities were identified as either ChAT or TH positive and counted, on a total of 4 animals, 48 sections and 3186 cells.

#### Probes

Primers were designed to amplify by PCR probe templates for the following genes:

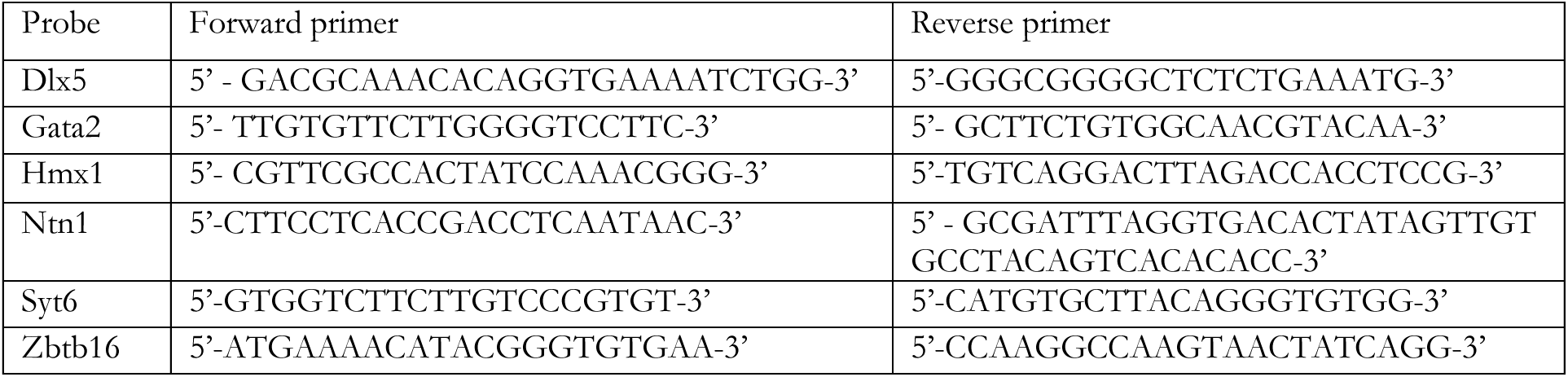

The PCR fragment were ligated into a pGEM-T Easy Vector System (Promega), transformed into chemically competent cells and sequenced. The other plasmid templates were Ebf3 (gift of S. Garel), Gata3 (gift of JD Engel), Hand1 (gift of P. Cserjesi), Hmx2 (gift of E.E. Turner), Hmx3 (gift of S. Mansour), Islet1 (*25*) and Tbx20 (*26*), Sst (Clone Image ID #4981984).

Plasmids were digested by restriction enzymes and purified using a DNA clean & concentrator kit (Zymo research). Antisense probes were synthesized with RNA polymerases and a DIG RNA labeling mix, and purified by the ProbeQuant G-50 micro columns kit (GE Healthcare). Probes were stored at -20°C.

#### Antibodies

Primary antibodies were α-Phox2b rabbit (1:500 or 1:1000, Pattyn et al., 1997). α-Th (Invitrogen: OPA1-04050, 1:1000) and α-choline acetyltransferase (ChAT) (Thermofisher: PA1-9027, 1:100).

Secondary antibodies were goat α-rabbit (PK-4005, Vector Laboratories), donkey anti-goat 647 (A-21447, Thermofisher), donkey anti-rabbit 488 (A-21206, Thermofisher) and donkey α-rabbit Cy3 (711-165-152, Jackson).

## Acknowledgments

We thank Nicolas Narboux-Nême for help with the analysis of Dlx5 expression, Sonia Garel for the gift of the Ebf3 probe, Amandine Delecourt and Gwendoline Firmin for help with the mouse colonies. The Brunet laboratory is supported by INSERM, CNRS, ANR-19-CE16-0029-01, ANR-17-CE16-0006-01, FRM EQU202003010297. M.A. is a recipient of a doctoral fellowship from Labex MemoLife. B.D. was funded by grant APP2001128 from the National Health & Medical Research Council, Australia. H.R. has been supported by a grant from the Wilhelm Sander Foundation. High-throughput sequencing was performed by i) the GenomiqueENS core facility supported by the France Génomique national infrastructure, funded as part of the “Investissements d’Avenir” program managed by the Agence Nationale de la Recherche (contract ANR-10-INBS-0009); ii) the ICGex NGS platform of the Institut Curie supported by the grants ANR-10-EQPX-03 (Equipex) and ANR-10-INBS-09-08 (France Génomique Consortium) from the Agence Nationale de la Recherche (“Investissements d’Avenir” program), by the ITMO-Cancer Aviesan (Plan Cancer III) and by the SiRIC-Curie program (SiRIC Grant INCa-DGOS-465 and INCa-DGOS-Inserm_12554). Data management, quality control and primary analysis were performed by the Bioinformatics platform of the Institut Curie.

---

**Fig. S1.**
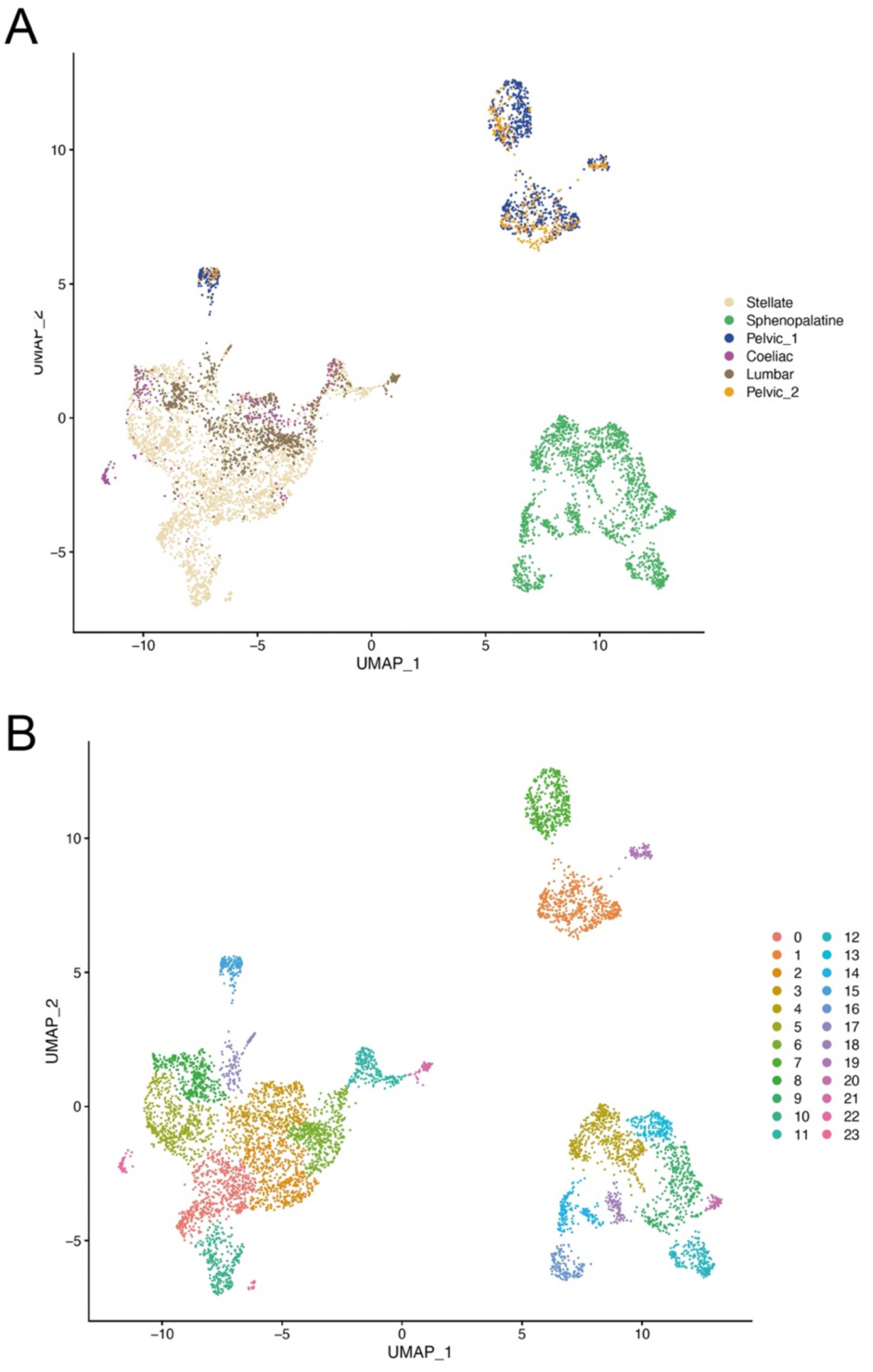
Uniform Manifold Approximation and Projection (UMAP) of all ganglionic neurons. (A) UMAP of neurons where the sample origin of cells is color-coded to show that both samples of the pelvic ganglion contribute to each of the P1-4 pelvic clusters. (B) UMAP of neurons where the clusters as defined by Seurat are color-coded. Clusters 1, 7, 15 and 19 correspond to ganglion clusters P1, P2, P3, P4 in the text.

**Fig. S2.**
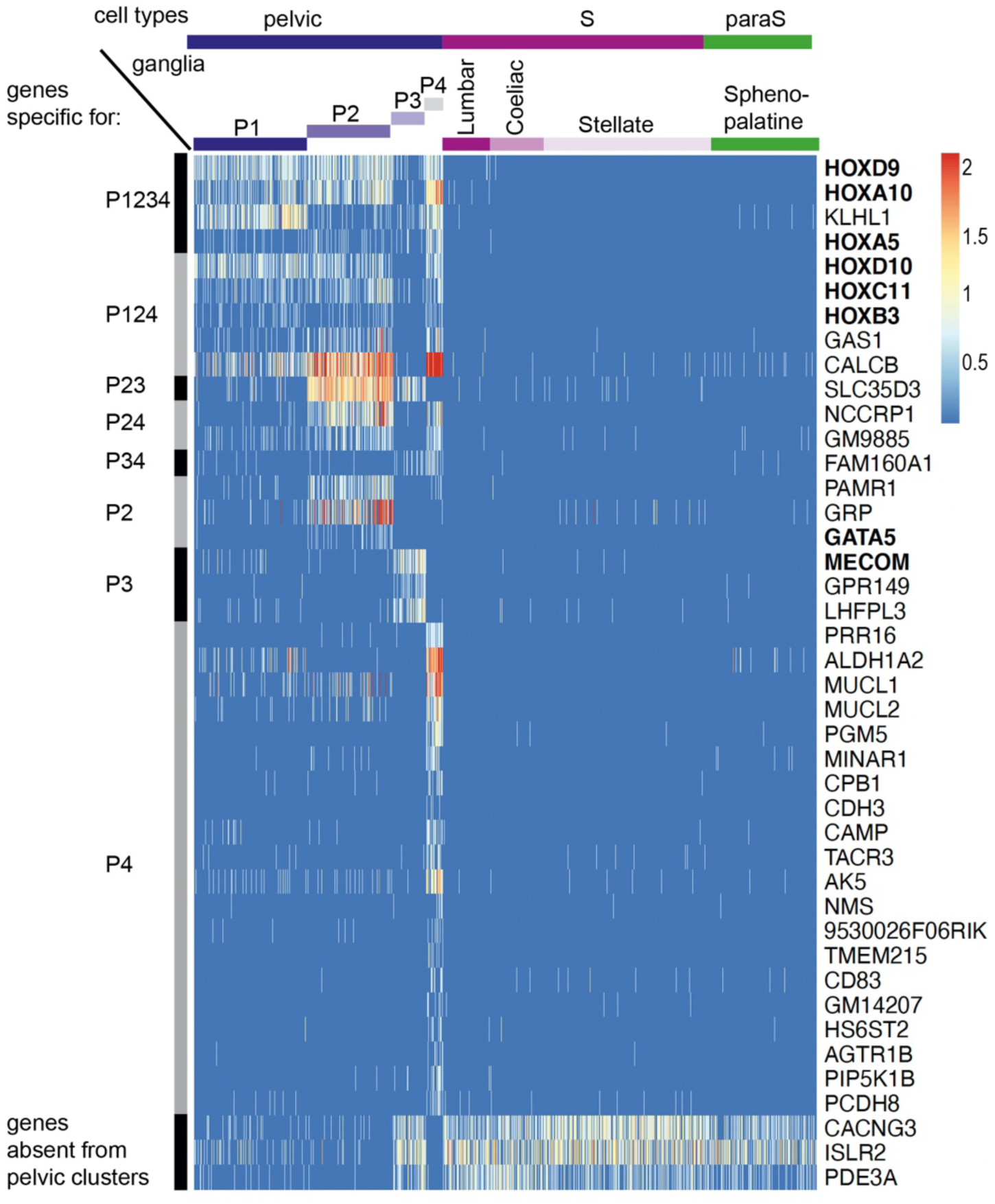
Genes specific for pelvic ganglionic cells among the top 100 genes of an all-versus-all comparison. These 42 genes correspond to pattern VII of main text. Transcription factors are indicated in bold face.

**Fig. S3:**
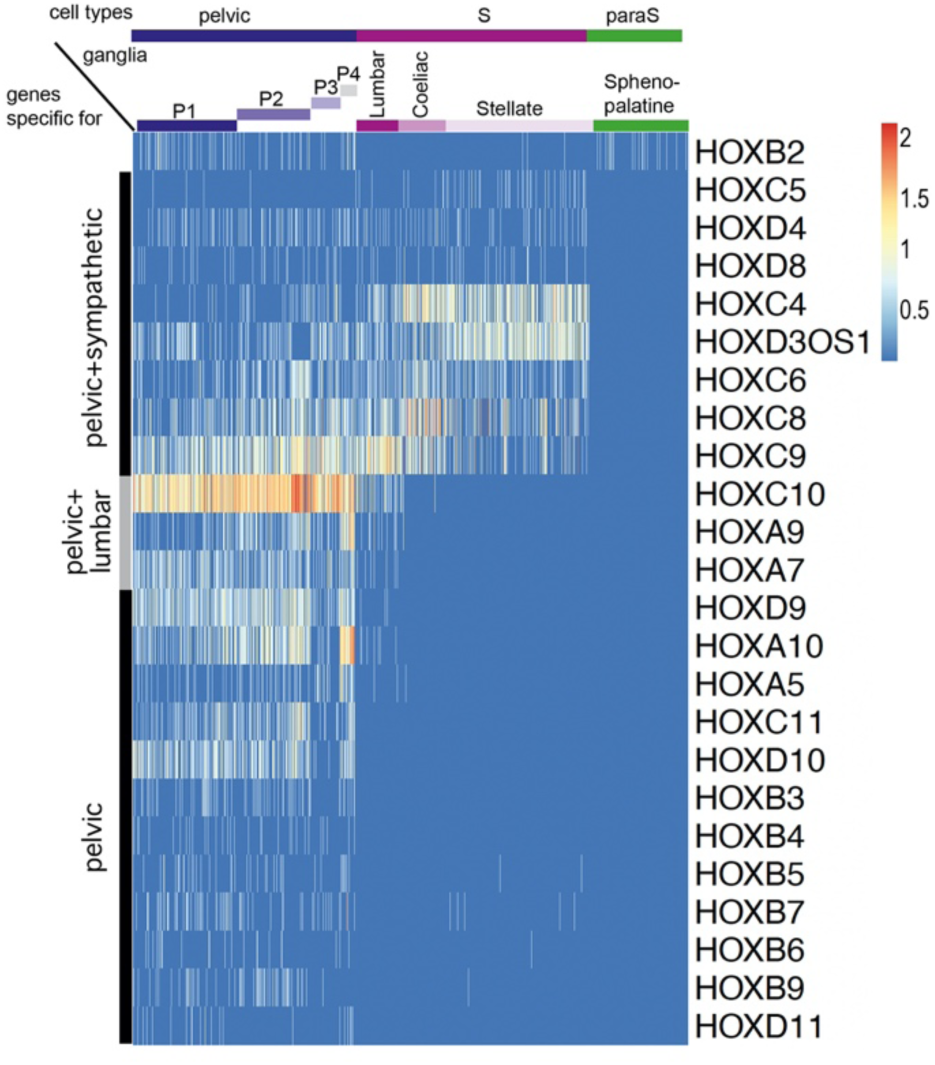
Expression of all Hox genes captured by the single-cell RNA-Seq data set. Apart from Hoxb2, all Hox genes are excluded from the sphenopalatine and are expressed either in all sympathetic and pelvic cells (8 genes), caudal sympathetic and pelvic cells (3 genes), or only in pelvic cells (12 genes). Hoxb2 appears in 177^th^ position as {P124/Sphenopalatine} versus {P3/Lumbar/Coeliac/Stellate}, thus as a cholinergic gene in the dichotomized comparison).

**Fig. S4:**
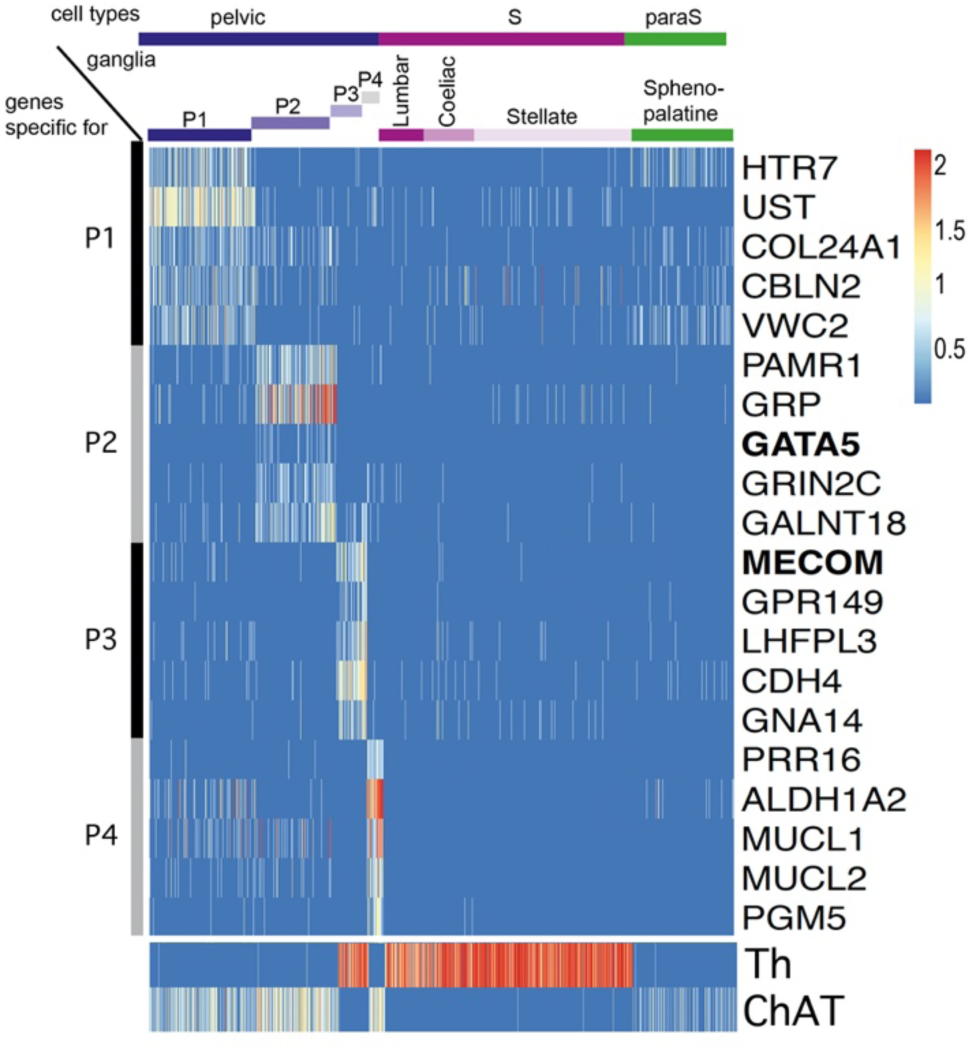
Five top genes for each of the 4 individual pelvic clusters in an all-versus-all comparison. P1 appears as the cluster the least sharply defined by specific genes. The only two transcription factors (indicated in bold face) among these top genes are Gata5, expressed in P2 and MECOM expressed in P3. Noradrenergic and cholinergic cells are indicated by Th and ChAT expression in the lower panel.

## Notes

https://www.ncbi.nlm.nih.gov/geo/query/acc.cgi?acc=GSE232789)

### Competing Interest Statement

The authors have declared no competing interest.

### Summary of Updates

- The legends of Fig. S3 and S4 were interchanged, this is now corrected. -In the material and methods it is now stated that both sexes are represented in the biological samples. -In the legend of figure 1 a reference to a supplementary figure has been corrected. -A few minor stylistic changes have been made.

