## Supplementary material for "The pelvic organs receive no parasympathetic innervation": Table S1

| gene | score | group.compar.string | P1 | P2 | P3 | P4 | Lumbar | Coeliac | Stellate | Sphenopalatine |
|---|---|---|---|---|---|---|---|---|---|---|
| ISL1 | 5.36 | P1_P2_P3_P4_Lumbar_Coeliac_Stellate_VS_Sphenopalatine | 1 | 1 | 1 | 1 | 1 | 1 | 1 | 0 |
| HAND1 | 5.29 | P1_P2_P3_P4_Lumbar_Coeliac_Stellate_VS_Sphenopalatine | 1 | 1 | 1 | 1 | 1 | 1 | 1 | 0 |
| HOXC10 | 5.16 | P1_P2_P3_P4_Lumbar_VS_Coeliac_Stellate_Sphenopalatine | 1 | 1 | 1 | 1 | 1 | 0 | 0 | 0 |
| GATA3 | 5.11 | P1_P2_P3_P4_Lumbar_Coeliac_Stellate_VS_Sphenopalatine | 1 | 1 | 1 | 1 | 1 | 1 | 1 | 0 |
| HOXC9 | 4.87 | P1_P2_P3_P4_Lumbar_Coeliac_Stellate_VS_Sphenopalatine | 1 | 1 | 1 | 1 | 1 | 1 | 1 | 0 |
| HOXC8 | 4.87 | P1_P2_P3_P4_Lumbar_Coeliac_Stellate_VS_Sphenopalatine | 1 | 1 | 1 | 1 | 1 | 1 | 1 | 0 |
| GALR1 | 4.82 | Sphenopalatine_VS_P1_P2_P3_P4_Lumbar_Coeliac_Stellate | 0 | 0 | 0 | 0 | 0 | 0 | 0 | 1 |
| HOXD3OS1 | 4.77 | P1_P2_P3_P4_Lumbar_Coeliac_Stellate_VS_Sphenopalatine | 1 | 1 | 1 | 1 | 1 | 1 | 1 | 0 |
| PRR16 | 4.59 | P4_VS_P1_P2_P3_Lumbar_Coeliac_Stellate_Sphenopalatine | 0 | 0 | 0 | 1 | 0 | 0 | 0 | 0 |
| HOXC6 | 4.59 | P1_P2_P3_P4_Lumbar_Coeliac_Stellate_VS_Sphenopalatine | 1 | 1 | 1 | 1 | 1 | 1 | 1 | 0 |
| GATA2 | 4.58 | P1_P2_P3_P4_Lumbar_Coeliac_Stellate_VS_Sphenopalatine | 1 | 1 | 1 | 1 | 1 | 1 | 1 | 0 |
| HOXC4 | 4.54 | P1_P2_P3_P4_Lumbar_Coeliac_Stellate_VS_Sphenopalatine | 1 | 1 | 1 | 1 | 1 | 1 | 1 | 0 |
| NTN1 | 4.52 | Sphenopalatine_VS_P1_P2_P3_P4_Lumbar_Coeliac_Stellate | 0 | 0 | 0 | 0 | 0 | 0 | 0 | 1 |
| HOXD10 | 4.5 | P1_P2_P4_VS_P3_Lumbar_Coeliac_Stellate_Sphenopalatine | 1 | 1 | 0 | 1 | 0 | 0 | 0 | 0 |
| HMX2 | 4.3 | Sphenopalatine_VS_P1_P2_P3_P4_Lumbar_Coeliac_Stellate | 0 | 0 | 0 | 0 | 0 | 0 | 0 | 1 |
| DLX6 | 4.29 | Sphenopalatine_VS_P1_P2_P3_P4_Lumbar_Coeliac_Stellate | 0 | 0 | 0 | 0 | 0 | 0 | 0 | 1 |
| ALDH1A2 | 4.28 | P4_VS_P1_P2_P3_Lumbar_Coeliac_Stellate_Sphenopalatine | 0 | 0 | 0 | 1 | 0 | 0 | 0 | 0 |
| CHAT | 4.17 | P1_P2_P4_Sphenopalatine_VS_P3_Lumbar_Coeliac_Stellate | 1 | 1 | 0 | 1 | 0 | 0 | 0 | 1 |
| HMX1 | 4.15 | P1_P2_P4_Lumbar_Coeliac_Stellate_VS_P3_Sphenopalatine | 1 | 1 | 0 | 1 | 1 | 1 | 1 | 0 |
| EBF3 | 4.13 | Sphenopalatine_VS_P1_P2_P3_P4_Lumbar_Coeliac_Stellate | 0 | 0 | 0 | 0 | 0 | 0 | 0 | 1 |
| HOXD9 | 4.12 | P1_P2_P3_P4_VS_Lumbar_Coeliac_Stellate_Sphenopalatine | 1 | 1 | 1 | 1 | 0 | 0 | 0 | 0 |
| HOXA7 | 4.1 | P1_P2_P3_P4_Lumbar_VS_Coeliac_Stellate_Sphenopalatine | 1 | 1 | 1 | 1 | 1 | 0 | 0 | 0 |
| MUCL1 | 4.06 | P4_VS_P1_P2_P3_Lumbar_Coeliac_Stellate_Sphenopalatine | 0 | 0 | 0 | 1 | 0 | 0 | 0 | 0 |
| HOXA9 | 4.04 | P1_P2_P3_P4_Lumbar_VS_Coeliac_Stellate_Sphenopalatine | 1 | 1 | 1 | 1 | 1 | 0 | 0 | 0 |
| SYT6 | 4.04 | Sphenopalatine_VS_P1_P2_P3_P4_Lumbar_Coeliac_Stellate | 0 | 0 | 0 | 0 | 0 | 0 | 0 | 1 |
| MUCL2 | 3.99 | P4_VS_P1_P2_P3_Lumbar_Coeliac_Stellate_Sphenopalatine | 0 | 0 | 0 | 1 | 0 | 0 | 0 | 0 |
| PGM5 | 3.99 | P4_VS_P1_P2_P3_Lumbar_Coeliac_Stellate_Sphenopalatine | 0 | 0 | 0 | 1 | 0 | 0 | 0 | 0 |
| NCCRP1 | 3.99 | P2_P4_VS_P1_P3_Lumbar_Coeliac_Stellate_Sphenopalatine | 0 | 1 | 0 | 1 | 0 | 0 | 0 | 0 |
| CACNG3 | 3.98 | P1_P2_P3_Lumbar_Coeliac_Stellate_Sphenopalatine_VS_P4 | 1 | 1 | 1 | 0 | 1 | 1 | 1 | 1 |

| | | | | | | | | | | |
|---|---|---|---|---|---|---|---|---|---|---|
| INSM2 | 3.96 | P3_Lumbar_Coeliac_Stellate_VS_P1_P2_P4_Sphenopalatine | 0 | 0 | 1 | 0 | 1 | 1 | 1 | 0 |
| PTGDR | 3.93 | Sphenopalatine_VS_P1_P2_P3_P4_Lumbar_Coeliac_Stellate | 0 | 0 | 0 | 0 | 0 | 0 | 0 | 1 |
| MECOM | 3.92 | P3_VS_P1_P2_P4_Lumbar_Coeliac_Stellate_Sphenopalatine | 0 | 0 | 1 | 0 | 0 | 0 | 0 | 0 |
| GM43066 | 3.9 | Sphenopalatine_VS_P1_P2_P3_P4_Lumbar_Coeliac_Stellate | 0 | 0 | 0 | 0 | 0 | 0 | 0 | 1 |
| ZFPM2 | 3.89 | P3_Lumbar_Coeliac_Stellate_VS_P1_P2_P4_Sphenopalatine | 0 | 0 | 1 | 0 | 1 | 1 | 1 | 0 |
| 9130024F11I | 3.88 | Sphenopalatine_VS_P1_P2_P3_P4_Lumbar_Coeliac_Stellate | 0 | 0 | 0 | 0 | 0 | 0 | 0 | 1 |
| HOXA10 | 3.87 | P1_P2_P3_P4_VS_Lumbar_Coeliac_Stellate_Sphenopalatine | 1 | 1 | 1 | 1 | 0 | 0 | 0 | 0 |
| HOXC11 | 3.85 | P1_P2_P4_VS_P3_Lumbar_Coeliac_Stellate_Sphenopalatine | 1 | 1 | 0 | 1 | 0 | 0 | 0 | 0 |
| HOXD4 | 3.84 | P1_P2_P3_P4_Lumbar_Coeliac_Stellate_VS_Sphenopalatine | 1 | 1 | 1 | 1 | 1 | 1 | 1 | 0 |
| NELL1 | 3.82 | P1_P2_P4_Sphenopalatine_VS_P3_Lumbar_Coeliac_Stellate | 1 | 1 | 0 | 1 | 0 | 0 | 0 | 1 |
| TAC2 | 3.76 | Sphenopalatine_VS_P1_P2_P3_P4_Lumbar_Coeliac_Stellate | 0 | 0 | 0 | 0 | 0 | 0 | 0 | 1 |
| SHISA9 | 3.73 | P1_P2_P3_P4_Lumbar_Coeliac_Stellate_VS_Sphenopalatine | 1 | 1 | 1 | 1 | 1 | 1 | 1 | 0 |
| BMPR1B | 3.72 | Lumbar_Coeliac_Stellate_VS_P1_P2_P3_P4_Sphenopalatine | 0 | 0 | 0 | 0 | 1 | 1 | 1 | 0 |
| DLX5 | 3.69 | Sphenopalatine_VS_P1_P2_P3_P4_Lumbar_Coeliac_Stellate | 0 | 0 | 0 | 0 | 0 | 0 | 0 | 1 |
| MINAR1 | 3.65 | P4_VS_P1_P2_P3_Lumbar_Coeliac_Stellate_Sphenopalatine | 0 | 0 | 0 | 1 | 0 | 0 | 0 | 0 |
| GPR149 | 3.64 | P3_VS_P1_P2_P4_Lumbar_Coeliac_Stellate_Sphenopalatine | 0 | 0 | 1 | 0 | 0 | 0 | 0 | 0 |
| SCTR | 3.64 | Lumbar_Coeliac_Stellate_VS_P1_P2_P3_P4_Sphenopalatine | 0 | 0 | 0 | 0 | 1 | 1 | 1 | 0 |
| CPB1 | 3.61 | P4_VS_P1_P2_P3_Lumbar_Coeliac_Stellate_Sphenopalatine | 0 | 0 | 0 | 1 | 0 | 0 | 0 | 0 |
| CDH3 | 3.6 | P4_VS_P1_P2_P3_Lumbar_Coeliac_Stellate_Sphenopalatine | 0 | 0 | 0 | 1 | 0 | 0 | 0 | 0 |
| TAC1 | 3.6 | Sphenopalatine_VS_P1_P2_P3_P4_Lumbar_Coeliac_Stellate | 0 | 0 | 0 | 0 | 0 | 0 | 0 | 1 |
| HMX3 | 3.59 | Sphenopalatine_VS_P1_P2_P3_P4_Lumbar_Coeliac_Stellate | 0 | 0 | 0 | 0 | 0 | 0 | 0 | 1 |
| TFAP2B | 3.58 | P3_Lumbar_Coeliac_Stellate_VS_P1_P2_P4_Sphenopalatine | 0 | 0 | 1 | 0 | 1 | 1 | 1 | 0 |
| NTRK1 | 3.56 | P3_Lumbar_Coeliac_Stellate_VS_P1_P2_P4_Sphenopalatine | 0 | 0 | 1 | 0 | 1 | 1 | 1 | 0 |
| RASGRP1 | 3.5 | Lumbar_Coeliac_Stellate_VS_P1_P2_P3_P4_Sphenopalatine | 0 | 0 | 0 | 0 | 1 | 1 | 1 | 0 |
| PAMR1 | 3.49 | P2_VS_P1_P3_P4_Lumbar_Coeliac_Stellate_Sphenopalatine | 0 | 1 | 0 | 0 | 0 | 0 | 0 | 0 |
| INSRR | 3.49 | P3_Lumbar_Coeliac_Stellate_VS_P1_P2_P4_Sphenopalatine | 0 | 0 | 1 | 0 | 1 | 1 | 1 | 0 |
| SLC35D3 | 3.48 | P2_P3_VS_P1_P4_Lumbar_Coeliac_Stellate_Sphenopalatine | 0 | 1 | 1 | 0 | 0 | 0 | 0 | 0 |
| KLHL1 | 3.48 | P1_P2_P3_P4_VS_Lumbar_Coeliac_Stellate_Sphenopalatine | 1 | 1 | 1 | 1 | 0 | 0 | 0 | 0 |
| GM2990 | 3.47 | P1_P2_P4_Sphenopalatine_VS_P3_Lumbar_Coeliac_Stellate | 1 | 1 | 0 | 1 | 0 | 0 | 0 | 1 |
| CAMP | 3.44 | P4_VS_P1_P2_P3_Lumbar_Coeliac_Stellate_Sphenopalatine | 0 | 0 | 0 | 1 | 0 | 0 | 0 | 0 |

| | | | | | | | | | | |
|---|---|---|---|---|---|---|---|---|---|---|
| SATB2 | 3.44 | Sphenopalatine_VS_P1_P2_P3_P4_Lumbar_Coeliac_Stellate | 0 | 0 | 0 | 0 | 0 | 0 | 0 | 1 |
| TH | 3.43 | P3_Lumbar_Coeliac_Stellate_VS_P1_P2_P4_Sphenopalatine | 0 | 0 | 1 | 0 | 1 | 1 | 1 | 0 |
| GRP | 3.41 | P2_VS_P1_P3_P4_Lumbar_Coeliac_Stellate_Sphenopalatine | 0 | 1 | 0 | 0 | 0 | 0 | 0 | 0 |
| ADAMTSL1 | 3.41 | Sphenopalatine_VS_P1_P2_P3_P4_Lumbar_Coeliac_Stellate | 0 | 0 | 0 | 0 | 0 | 0 | 0 | 1 |
| BNC2 | 3.39 | P1_P4_Sphenopalatine_VS_P2_P3_Lumbar_Coeliac_Stellate | 1 | 0 | 0 | 1 | 0 | 0 | 0 | 1 |
| TACR3 | 3.38 | P4_VS_P1_P2_P3_Lumbar_Coeliac_Stellate_Sphenopalatine | 0 | 0 | 0 | 1 | 0 | 0 | 0 | 0 |
| C1QL2 | 3.37 | P1_Sphenopalatine_VS_P2_P3_P4_Lumbar_Coeliac_Stellate | 1 | 0 | 0 | 0 | 0 | 0 | 0 | 1 |
| GM36684 | 3.35 | P1_P2_P3_P4_Stellate_Sphenopalatine_VS_Lumbar_Coeliac | 1 | 1 | 1 | 1 | 0 | 0 | 1 | 1 |
| ISLR2 | 3.35 | P1_P2_P3_Lumbar_Coeliac_Stellate_Sphenopalatine_VS_P4 | 1 | 1 | 1 | 0 | 1 | 1 | 1 | 1 |
| GM30648 | 3.33 | P1_P2_P3_P4_Lumbar_Coeliac_Stellate_VS_Sphenopalatine | 1 | 1 | 1 | 1 | 1 | 1 | 1 | 0 |
| HOXB3 | 3.32 | P1_P2_P4_VS_P3_Lumbar_Coeliac_Stellate_Sphenopalatine | 1 | 1 | 0 | 1 | 0 | 0 | 0 | 0 |
| GAS1 | 3.32 | P1_P2_P4_VS_P3_Lumbar_Coeliac_Stellate_Sphenopalatine | 1 | 1 | 0 | 1 | 0 | 0 | 0 | 0 |
| HOXA5 | 3.3 | P1_P2_P3_P4_VS_Lumbar_Coeliac_Stellate_Sphenopalatine | 1 | 1 | 1 | 1 | 0 | 0 | 0 | 0 |
| TBX20 | 3.29 | P1_P2_P3_P4_Lumbar_Coeliac_Stellate_VS_Sphenopalatine | 1 | 1 | 1 | 1 | 1 | 1 | 1 | 0 |
| AK5 | 3.27 | P4_VS_P1_P2_P3_Lumbar_Coeliac_Stellate_Sphenopalatine | 0 | 0 | 0 | 1 | 0 | 0 | 0 | 0 |
| SST | 3.27 | P1_P2_P3_P4_Lumbar_Coeliac_Stellate_VS_Sphenopalatine | 1 | 1 | 1 | 1 | 1 | 1 | 1 | 0 |
| COL8A1 | 3.27 | P4_Sphenopalatine_VS_P1_P2_P3_Lumbar_Coeliac_Stellate | 0 | 0 | 0 | 1 | 0 | 0 | 0 | 1 |
| NMS | 3.25 | P4_VS_P1_P2_P3_Lumbar_Coeliac_Stellate_Sphenopalatine | 0 | 0 | 0 | 1 | 0 | 0 | 0 | 0 |
| 9530026F06I | 3.24 | P4_VS_P1_P2_P3_Lumbar_Coeliac_Stellate_Sphenopalatine | 0 | 0 | 0 | 1 | 0 | 0 | 0 | 0 |
| CALCB | 3.22 | P1_P2_P4_VS_P3_Lumbar_Coeliac_Stellate_Sphenopalatine | 1 | 1 | 0 | 1 | 0 | 0 | 0 | 0 |
| FAM160A1 | 3.22 | P3_P4_VS_P1_P2_Lumbar_Coeliac_Stellate_Sphenopalatine | 0 | 0 | 1 | 1 | 0 | 0 | 0 | 0 |
| TRIM55 | 3.22 | Lumbar_Coeliac_VS_P1_P2_P3_P4_Stellate_Sphenopalatine | 0 | 0 | 0 | 0 | 1 | 1 | 0 | 0 |
| TMEM215 | 3.2 | P4_VS_P1_P2_P3_Lumbar_Coeliac_Stellate_Sphenopalatine | 0 | 0 | 0 | 1 | 0 | 0 | 0 | 0 |
| CD83 | 3.19 | P4_VS_P1_P2_P3_Lumbar_Coeliac_Stellate_Sphenopalatine | 0 | 0 | 0 | 1 | 0 | 0 | 0 | 0 |
| PNMT | 3.19 | Lumbar_Coeliac_Stellate_VS_P1_P2_P3_P4_Sphenopalatine | 0 | 0 | 0 | 0 | 1 | 1 | 1 | 0 |
| GFRA3 | 3.19 | P3_Lumbar_Coeliac_Stellate_VS_P1_P2_P4_Sphenopalatine | 0 | 0 | 1 | 0 | 1 | 1 | 1 | 0 |
| LHFPL3 | 3.18 | P3_VS_P1_P2_P4_Lumbar_Coeliac_Stellate_Sphenopalatine | 0 | 0 | 1 | 0 | 0 | 0 | 0 | 0 |
| GM14207 | 3.15 | P4_VS_P1_P2_P3_Lumbar_Coeliac_Stellate_Sphenopalatine | 0 | 0 | 0 | 1 | 0 | 0 | 0 | 0 |
| LRFN5 | 3.14 | P1_P2_P4_Sphenopalatine_VS_P3_Lumbar_Coeliac_Stellate | 1 | 1 | 0 | 1 | 0 | 0 | 0 | 1 |
| GM9885 | 3.11 | P2_P4_VS_P1_P3_Lumbar_Coeliac_Stellate_Sphenopalatine | 0 | 1 | 0 | 1 | 0 | 0 | 0 | 0 |

| | | | | | | | | | | |
|---|---|---|---|---|---|---|---|---|---|---|
| GATA5 | 3.09 | P2_VS_P1_P3_P4_Lumbar_Coeliac_Stellate_Sphenopalatine | 0 | 1 | 0 | 0 | 0 | 0 | 0 | 0 |
| PAPPA | 3.09 | P1_P2_P4_Sphenopalatine_VS_P3_Lumbar_Coeliac_Stellate | 1 | 1 | 0 | 1 | 0 | 0 | 0 | 1 |
| PDE3A | 3.09 | P1_P3_P4_Lumbar_Coeliac_Stellate_Sphenopalatine_VS_P2 | 1 | 0 | 1 | 1 | 1 | 1 | 1 | 1 |
| DRD2 | 3.08 | P3_Lumbar_Coeliac_VS_P1_P2_P4_Stellate_Sphenopalatine | 0 | 0 | 1 | 0 | 1 | 1 | 0 | 0 |
| MGAT4C | 3.08 | P3_P4_Lumbar_Coeliac_Stellate_VS_P1_P2_Sphenopalatine | 0 | 0 | 1 | 1 | 1 | 1 | 1 | 0 |
| HS6ST2 | 3.07 | P4_VS_P1_P2_P3_Lumbar_Coeliac_Stellate_Sphenopalatine | 0 | 0 | 0 | 1 | 0 | 0 | 0 | 0 |
| AGTR1B | 3.06 | P4_VS_P1_P2_P3_Lumbar_Coeliac_Stellate_Sphenopalatine | 0 | 0 | 0 | 1 | 0 | 0 | 0 | 0 |
| PIP5K1B | 3.06 | P4_VS_P1_P2_P3_Lumbar_Coeliac_Stellate_Sphenopalatine | 0 | 0 | 0 | 1 | 0 | 0 | 0 | 0 |
| PCDH8 | 3.05 | P4_VS_P1_P2_P3_Lumbar_Coeliac_Stellate_Sphenopalatine | 0 | 0 | 0 | 1 | 0 | 0 | 0 | 0 |
| 2610028E06I | 3.05 | Sphenopalatine_VS_P1_P2_P3_P4_Lumbar_Coeliac_Stellate | 0 | 0 | 0 | 0 | 0 | 0 | 0 | 1 |
| VIP | 3.05 | P1_P2_P3_P4_Lumbar_Stellate_Sphenopalatine_VS_Coeliac | 1 | 1 | 1 | 1 | 1 | 0 | 1 | 1 |
| CDH4 | 3.04 | P3_VS_P1_P2_P4_Lumbar_Coeliac_Stellate_Sphenopalatine | 0 | 0 | 1 | 0 | 0 | 0 | 0 | 0 |
| HOTAIRM1 | 3.04 | P4_VS_P1_P2_P3_Lumbar_Coeliac_Stellate_Sphenopalatine | 0 | 0 | 0 | 1 | 0 | 0 | 0 | 0 |
| RAI14 | 3.03 | P4_VS_P1_P2_P3_Lumbar_Coeliac_Stellate_Sphenopalatine | 0 | 0 | 0 | 1 | 0 | 0 | 0 | 0 |
| PYY | 3.03 | Sphenopalatine_VS_P1_P2_P3_P4_Lumbar_Coeliac_Stellate | 0 | 0 | 0 | 0 | 0 | 0 | 0 | 1 |
| 2900079G21 | 3.02 | Sphenopalatine_VS_P1_P2_P3_P4_Lumbar_Coeliac_Stellate | 0 | 0 | 0 | 0 | 0 | 0 | 0 | 1 |
| GNA14 | 3.01 | P3_VS_P1_P2_P4_Lumbar_Coeliac_Stellate_Sphenopalatine | 0 | 0 | 1 | 0 | 0 | 0 | 0 | 0 |
| HCRTR2 | 3.01 | P4_VS_P1_P2_P3_Lumbar_Coeliac_Stellate_Sphenopalatine | 0 | 0 | 0 | 1 | 0 | 0 | 0 | 0 |
| PALMD | 3 | P1_P2_P3_Lumbar_Coeliac_Stellate_Sphenopalatine_VS_P4 | 1 | 1 | 1 | 0 | 1 | 1 | 1 | 1 |
| AA387883 | 2.99 | P4_VS_P1_P2_P3_Lumbar_Coeliac_Stellate_Sphenopalatine | 0 | 0 | 0 | 1 | 0 | 0 | 0 | 0 |
| KCNIP4 | 2.97 | P1_P4_Lumbar_Coeliac_Stellate_Sphenopalatine_VS_P2_P3 | 1 | 0 | 0 | 1 | 1 | 1 | 1 | 1 |
| CTXN3 | 2.95 | P4_VS_P1_P2_P3_Lumbar_Coeliac_Stellate_Sphenopalatine | 0 | 0 | 0 | 1 | 0 | 0 | 0 | 0 |
| SOX9 | 2.94 | P4_VS_P1_P2_P3_Lumbar_Coeliac_Stellate_Sphenopalatine | 0 | 0 | 0 | 1 | 0 | 0 | 0 | 0 |
| CALCA | 2.93 | P4_VS_P1_P2_P3_Lumbar_Coeliac_Stellate_Sphenopalatine | 0 | 0 | 0 | 1 | 0 | 0 | 0 | 0 |
| CD79A | 2.92 | P3_VS_P1_P2_P4_Lumbar_Coeliac_Stellate_Sphenopalatine | 0 | 0 | 1 | 0 | 0 | 0 | 0 | 0 |
| CYP26B1 | 2.92 | P4_VS_P1_P2_P3_Lumbar_Coeliac_Stellate_Sphenopalatine | 0 | 0 | 0 | 1 | 0 | 0 | 0 | 0 |
| TNFSF13B | 2.92 | P4_VS_P1_P2_P3_Lumbar_Coeliac_Stellate_Sphenopalatine | 0 | 0 | 0 | 1 | 0 | 0 | 0 | 0 |
| TAFA1 | 2.92 | P1_P2_P3_P4_Sphenopalatine_VS_Lumbar_Coeliac_Stellate | 1 | 1 | 1 | 1 | 0 | 0 | 0 | 1 |
| GRIN2C | 2.91 | P2_VS_P1_P3_P4_Lumbar_Coeliac_Stellate_Sphenopalatine | 0 | 1 | 0 | 0 | 0 | 0 | 0 | 0 |
| HOXD11 | 2.91 | P4_VS_P1_P2_P3_Lumbar_Coeliac_Stellate_Sphenopalatine | 0 | 0 | 0 | 1 | 0 | 0 | 0 | 0 |

| | | | | | | | | | | |
|---|---|---|---|---|---|---|---|---|---|---|
| PDE1A | 2.91 | P4_VS_P1_P2_P3_Lumbar_Coeliac_Stellate_Sphenopalatine | 0 | 0 | 0 | 1 | 0 | 0 | 0 | 0 |
| NDST4 | 2.91 | P1_P4_VS_P2_P3_Lumbar_Coeliac_Stellate_Sphenopalatine | 1 | 0 | 0 | 1 | 0 | 0 | 0 | 0 |
| ST6GALNAC5 | 2.91 | P1_P4_Lumbar_Coeliac_Stellate_Sphenopalatine_VS_P2_P3 | 1 | 0 | 0 | 1 | 1 | 1 | 1 | 1 |
| PARD3B | 2.9 | P2_P4_VS_P1_P3_Lumbar_Coeliac_Stellate_Sphenopalatine | 0 | 1 | 0 | 1 | 0 | 0 | 0 | 0 |
| HOXB7 | 2.9 | P1_P2_P4_VS_P3_Lumbar_Coeliac_Stellate_Sphenopalatine | 1 | 1 | 0 | 1 | 0 | 0 | 0 | 0 |
| RSPO2 | 2.9 | P1_P4_Sphenopalatine_VS_P2_P3_Lumbar_Coeliac_Stellate | 1 | 0 | 0 | 1 | 0 | 0 | 0 | 1 |
| RBFOX3 | 2.9 | P1_P2_P3_P4_Sphenopalatine_VS_Lumbar_Coeliac_Stellate | 1 | 1 | 1 | 1 | 0 | 0 | 0 | 1 |
| GALNT18 | 2.89 | P2_VS_P1_P3_P4_Lumbar_Coeliac_Stellate_Sphenopalatine | 0 | 1 | 0 | 0 | 0 | 0 | 0 | 0 |
| RGMA | 2.89 | P4_VS_P1_P2_P3_Lumbar_Coeliac_Stellate_Sphenopalatine | 0 | 0 | 0 | 1 | 0 | 0 | 0 | 0 |
| ECEL1 | 2.89 | P1_P2_P3_P4_Lumbar_Coeliac_Stellate_VS_Sphenopalatine | 1 | 1 | 1 | 1 | 1 | 1 | 1 | 0 |
| GLT28D2 | 2.88 | P3_P4_VS_P1_P2_Lumbar_Coeliac_Stellate_Sphenopalatine | 0 | 0 | 1 | 1 | 0 | 0 | 0 | 0 |
| CDH13 | 2.88 | Sphenopalatine_VS_P1_P2_P3_P4_Lumbar_Coeliac_Stellate | 0 | 0 | 0 | 0 | 0 | 0 | 0 | 1 |
| CHST1 | 2.87 | P2_P3_P4_Lumbar_Coeliac_Stellate_VS_P1_Sphenopalatine | 0 | 1 | 1 | 1 | 1 | 1 | 1 | 0 |
| TLX3 | 2.86 | P3_VS_P1_P2_P4_Lumbar_Coeliac_Stellate_Sphenopalatine | 0 | 0 | 1 | 0 | 0 | 0 | 0 | 0 |
| HOXD8 | 2.86 | P1_P2_P3_P4_Lumbar_Coeliac_Stellate_VS_Sphenopalatine | 1 | 1 | 1 | 1 | 1 | 1 | 1 | 0 |
| UGT8A | 2.83 | P4_VS_P1_P2_P3_Lumbar_Coeliac_Stellate_Sphenopalatine | 0 | 0 | 0 | 1 | 0 | 0 | 0 | 0 |
| SOX6 | 2.82 | Coeliac_VS_P1_P2_P3_P4_Lumbar_Stellate_Sphenopalatine | 0 | 0 | 0 | 0 | 0 | 1 | 0 | 0 |
| ADRA2C | 2.82 | P1_P2_P3_P4_Lumbar_Coeliac_Stellate_VS_Sphenopalatine | 1 | 1 | 1 | 1 | 1 | 1 | 1 | 0 |
| ERICH2 | 2.81 | P4_VS_P1_P2_P3_Lumbar_Coeliac_Stellate_Sphenopalatine | 0 | 0 | 0 | 1 | 0 | 0 | 0 | 0 |
| SLC16A12 | 2.8 | P4_VS_P1_P2_P3_Lumbar_Coeliac_Stellate_Sphenopalatine | 0 | 0 | 0 | 1 | 0 | 0 | 0 | 0 |
| VSTM2A | 2.8 | Sphenopalatine_VS_P1_P2_P3_P4_Lumbar_Coeliac_Stellate | 0 | 0 | 0 | 0 | 0 | 0 | 0 | 1 |
| SLC6A2 | 2.8 | P3_Lumbar_Coeliac_Stellate_Sphenopalatine_VS_P1_P2_P4 | 0 | 0 | 1 | 0 | 1 | 1 | 1 | 1 |
| LMO7 | 2.79 | P3_VS_P1_P2_P4_Lumbar_Coeliac_Stellate_Sphenopalatine | 0 | 0 | 1 | 0 | 0 | 0 | 0 | 0 |
| GABRG3 | 2.79 | P1_Sphenopalatine_VS_P2_P3_P4_Lumbar_Coeliac_Stellate | 1 | 0 | 0 | 0 | 0 | 0 | 0 | 1 |
| KCNH5 | 2.78 | P4_VS_P1_P2_P3_Lumbar_Coeliac_Stellate_Sphenopalatine | 0 | 0 | 0 | 1 | 0 | 0 | 0 | 0 |
| THSD4 | 2.78 | P1_P2_P4_VS_P3_Lumbar_Coeliac_Stellate_Sphenopalatine | 1 | 1 | 0 | 1 | 0 | 0 | 0 | 0 |
| HTR7 | 2.76 | P1_VS_P2_P3_P4_Lumbar_Coeliac_Stellate_Sphenopalatine | 1 | 0 | 0 | 0 | 0 | 0 | 0 | 0 |
| OMP | 2.76 | Lumbar_Coeliac_Stellate_VS_P1_P2_P3_P4_Sphenopalatine | 0 | 0 | 0 | 0 | 1 | 1 | 1 | 0 |
| GRM7 | 2.76 | P1_P3_Lumbar_Coeliac_Stellate_Sphenopalatine_VS_P2_P4 | 1 | 0 | 1 | 0 | 1 | 1 | 1 | 1 |
| IFIH1 | 2.75 | P4_VS_P1_P2_P3_Lumbar_Coeliac_Stellate_Sphenopalatine | 0 | 0 | 0 | 1 | 0 | 0 | 0 | 0 |

| | | | | | | | | | | |
|---|---|---|---|---|---|---|---|---|---|---|
| CADPS2 | 2.74 | Sphenopalatine_VS_P1_P2_P3_P4_Lumbar_Coeliac_Stellate | 0 | 0 | 0 | 0 | 0 | 0 | 0 | 1 |
| GADL1 | 2.74 | Sphenopalatine_VS_P1_P2_P3_P4_Lumbar_Coeliac_Stellate | 0 | 0 | 0 | 0 | 0 | 0 | 0 | 1 |
| PENK | 2.73 | Sphenopalatine_VS_P1_P2_P3_P4_Lumbar_Coeliac_Stellate | 0 | 0 | 0 | 0 | 0 | 0 | 0 | 1 |
| CBLN4 | 2.73 | P1_P2_Sphenopalatine_VS_P3_P4_Lumbar_Coeliac_Stellate | 1 | 1 | 0 | 0 | 0 | 0 | 0 | 1 |
| PROM1 | 2.73 | P1_Lumbar_Coeliac_Stellate_Sphenopalatine_VS_P2_P3_P4 | 1 | 0 | 0 | 0 | 1 | 1 | 1 | 1 |
| AJAP1 | 2.71 | P2_P4_VS_P1_P3_Lumbar_Coeliac_Stellate_Sphenopalatine | 0 | 1 | 0 | 1 | 0 | 0 | 0 | 0 |
| CCND1 | 2.71 | P1_P2_P3_P4_Lumbar_Coeliac_Stellate_VS_Sphenopalatine | 1 | 1 | 1 | 1 | 1 | 1 | 1 | 0 |
| SYT10 | 2.7 | P2_VS_P1_P3_P4_Lumbar_Coeliac_Stellate_Sphenopalatine | 0 | 1 | 0 | 0 | 0 | 0 | 0 | 0 |
| NNMT | 2.7 | Sphenopalatine_VS_P1_P2_P3_P4_Lumbar_Coeliac_Stellate | 0 | 0 | 0 | 0 | 0 | 0 | 0 | 1 |
| C1QTNF7 | 2.7 | P1_P2_P3_P4_Lumbar_Coeliac_Sphenopalatine_VS_Stellate | 1 | 1 | 1 | 1 | 1 | 1 | 0 | 1 |
| GM53 | 2.69 | P1_P2_VS_P3_P4_Lumbar_Coeliac_Stellate_Sphenopalatine | 1 | 1 | 0 | 0 | 0 | 0 | 0 | 0 |
| IGFBP3 | 2.68 | P4_VS_P1_P2_P3_Lumbar_Coeliac_Stellate_Sphenopalatine | 0 | 0 | 0 | 1 | 0 | 0 | 0 | 0 |
| SNHG18 | 2.68 | P4_VS_P1_P2_P3_Lumbar_Coeliac_Stellate_Sphenopalatine | 0 | 0 | 0 | 1 | 0 | 0 | 0 | 0 |
| MYL1 | 2.68 | P1_P2_P3_P4_Lumbar_Coeliac_Stellate_VS_Sphenopalatine | 1 | 1 | 1 | 1 | 1 | 1 | 1 | 0 |
| GM8113 | 2.68 | Sphenopalatine_VS_P1_P2_P3_P4_Lumbar_Coeliac_Stellate | 0 | 0 | 0 | 0 | 0 | 0 | 0 | 1 |
| COLQ | 2.68 | P1_P2_P4_Sphenopalatine_VS_P3_Lumbar_Coeliac_Stellate | 1 | 1 | 0 | 1 | 0 | 0 | 0 | 1 |
| MME | 2.67 | P3_VS_P1_P2_P4_Lumbar_Coeliac_Stellate_Sphenopalatine | 0 | 0 | 1 | 0 | 0 | 0 | 0 | 0 |
| AGTR2 | 2.67 | P3_VS_P1_P2_P4_Lumbar_Coeliac_Stellate_Sphenopalatine | 0 | 0 | 1 | 0 | 0 | 0 | 0 | 0 |
| SCGN | 2.67 | Sphenopalatine_VS_P1_P2_P3_P4_Lumbar_Coeliac_Stellate | 0 | 0 | 0 | 0 | 0 | 0 | 0 | 1 |
| TNFRSF1A | 2.67 | P1_Sphenopalatine_VS_P2_P3_P4_Lumbar_Coeliac_Stellate | 1 | 0 | 0 | 0 | 0 | 0 | 0 | 1 |
| SLC18A1 | 2.66 | P1_P2_P3_Lumbar_Coeliac_Stellate_VS_P4_Sphenopalatine | 1 | 1 | 1 | 0 | 1 | 1 | 1 | 0 |
| GDA | 2.64 | P4_Lumbar_Stellate_VS_P1_P2_P3_Coeliac_Sphenopalatine | 0 | 0 | 0 | 1 | 1 | 0 | 1 | 0 |
| ADAMTS2 | 2.64 | P1_P2_Sphenopalatine_VS_P3_P4_Lumbar_Coeliac_Stellate | 1 | 1 | 0 | 0 | 0 | 0 | 0 | 1 |
| APOD | 2.63 | P4_VS_P1_P2_P3_Lumbar_Coeliac_Stellate_Sphenopalatine | 0 | 0 | 0 | 1 | 0 | 0 | 0 | 0 |
| HOXB2 | 2.63 | P1_P2_P4_Sphenopalatine_VS_P3_Lumbar_Coeliac_Stellate | 1 | 1 | 0 | 1 | 0 | 0 | 0 | 1 |
| KIRREL3 | 2.63 | P1_P2_P4_Lumbar_Coeliac_Stellate_Sphenopalatine_VS_P3 | 1 | 1 | 0 | 1 | 1 | 1 | 1 | 1 |
| HOXB9 | 2.62 | P2_VS_P1_P3_P4_Lumbar_Coeliac_Stellate_Sphenopalatine | 0 | 1 | 0 | 0 | 0 | 0 | 0 | 0 |
| BOC | 2.62 | P2_VS_P1_P3_P4_Lumbar_Coeliac_Stellate_Sphenopalatine | 0 | 1 | 0 | 0 | 0 | 0 | 0 | 0 |
| LAMC2 | 2.62 | Sphenopalatine_VS_P1_P2_P3_P4_Lumbar_Coeliac_Stellate | 0 | 0 | 0 | 0 | 0 | 0 | 0 | 1 |
| PTHLH | 2.61 | Lumbar_Stellate_VS_P1_P2_P3_P4_Coeliac_Sphenopalatine | 0 | 0 | 0 | 0 | 1 | 0 | 1 | 0 |

| | | | | | | | | | | |
|---|---|---|---|---|---|---|---|---|---|---|
| AGTR1A | 2.61 | P3_P4_Lumbar_Coeliac_Stellate_VS_P1_P2_Sphenopalatine | 0 | 0 | 1 | 1 | 1 | 1 | 1 | 0 |
| NOS1 | 2.61 | P1_P2_P3_P4_Sphenopalatine_VS_Lumbar_Coeliac_Stellate | 1 | 1 | 1 | 1 | 0 | 0 | 0 | 1 |
| GRID1 | 2.6 | P4_VS_P1_P2_P3_Lumbar_Coeliac_Stellate_Sphenopalatine | 0 | 0 | 0 | 1 | 0 | 0 | 0 | 0 |
| HOXB5 | 2.6 | P1_P2_P4_VS_P3_Lumbar_Coeliac_Stellate_Sphenopalatine | 1 | 1 | 0 | 1 | 0 | 0 | 0 | 0 |
| SKAP1 | 2.59 | P4_VS_P1_P2_P3_Lumbar_Coeliac_Stellate_Sphenopalatine | 0 | 0 | 0 | 1 | 0 | 0 | 0 | 0 |
| MEGF10 | 2.59 | P4_VS_P1_P2_P3_Lumbar_Coeliac_Stellate_Sphenopalatine | 0 | 0 | 0 | 1 | 0 | 0 | 0 | 0 |
| BAIAP2L1 | 2.58 | P2_VS_P1_P3_P4_Lumbar_Coeliac_Stellate_Sphenopalatine | 0 | 1 | 0 | 0 | 0 | 0 | 0 | 0 |
| SLC38A11 | 2.58 | P3_VS_P1_P2_P4_Lumbar_Coeliac_Stellate_Sphenopalatine | 0 | 0 | 1 | 0 | 0 | 0 | 0 | 0 |
| SHOX2 | 2.58 | Coeliac_VS_P1_P2_P3_P4_Lumbar_Stellate_Sphenopalatine | 0 | 0 | 0 | 0 | 0 | 1 | 0 | 0 |
| PRKCB | 2.58 | P3_Lumbar_Coeliac_Stellate_VS_P1_P2_P4_Sphenopalatine | 0 | 0 | 1 | 0 | 1 | 1 | 1 | 0 |
| HSPB7 | 2.58 | Sphenopalatine_VS_P1_P2_P3_P4_Lumbar_Coeliac_Stellate | 0 | 0 | 0 | 0 | 0 | 0 | 0 | 1 |
| CAV3 | 2.58 | Sphenopalatine_VS_P1_P2_P3_P4_Lumbar_Coeliac_Stellate | 0 | 0 | 0 | 0 | 0 | 0 | 0 | 1 |
| IGF2 | 2.57 | Sphenopalatine_VS_P1_P2_P3_P4_Lumbar_Coeliac_Stellate | 0 | 0 | 0 | 0 | 0 | 0 | 0 | 1 |
| ARPP21 | 2.57 | Sphenopalatine_VS_P1_P2_P3_P4_Lumbar_Coeliac_Stellate | 0 | 0 | 0 | 0 | 0 | 0 | 0 | 1 |
| LRP1B | 2.56 | P3_VS_P1_P2_P4_Lumbar_Coeliac_Stellate_Sphenopalatine | 0 | 0 | 1 | 0 | 0 | 0 | 0 | 0 |
| CGNL1 | 2.56 | P4_VS_P1_P2_P3_Lumbar_Coeliac_Stellate_Sphenopalatine | 0 | 0 | 0 | 1 | 0 | 0 | 0 | 0 |
| GM27164 | 2.55 | P4_VS_P1_P2_P3_Lumbar_Coeliac_Stellate_Sphenopalatine | 0 | 0 | 0 | 1 | 0 | 0 | 0 | 0 |
| HOXB4 | 2.55 | P4_VS_P1_P2_P3_Lumbar_Coeliac_Stellate_Sphenopalatine | 0 | 0 | 0 | 1 | 0 | 0 | 0 | 0 |
| EPHA7 | 2.55 | Sphenopalatine_VS_P1_P2_P3_P4_Lumbar_Coeliac_Stellate | 0 | 0 | 0 | 0 | 0 | 0 | 0 | 1 |
| HES1 | 2.55 | P1_P2_P4_Sphenopalatine_VS_P3_Lumbar_Coeliac_Stellate | 1 | 1 | 0 | 1 | 0 | 0 | 0 | 1 |
| UST | 2.54 | P1_VS_P2_P3_P4_Lumbar_Coeliac_Stellate_Sphenopalatine | 1 | 0 | 0 | 0 | 0 | 0 | 0 | 0 |
| HOXB6 | 2.54 | P4_VS_P1_P2_P3_Lumbar_Coeliac_Stellate_Sphenopalatine | 0 | 0 | 0 | 1 | 0 | 0 | 0 | 0 |
| NPTX2 | 2.54 | P1_P2_P4_VS_P3_Lumbar_Coeliac_Stellate_Sphenopalatine | 1 | 1 | 0 | 1 | 0 | 0 | 0 | 0 |
| ADAMTS19 | 2.53 | P3_Coeliac_VS_P1_P2_P4_Lumbar_Stellate_Sphenopalatine | 0 | 0 | 1 | 0 | 0 | 1 | 0 | 0 |
| NRN1 | 2.53 | P1_P2_P3_P4_Lumbar_Coeliac_Stellate_VS_Sphenopalatine | 1 | 1 | 1 | 1 | 1 | 1 | 1 | 0 |
| SHISAL2A | 2.53 | P1_P2_P3_Lumbar_Coeliac_Stellate_Sphenopalatine_VS_P4 | 1 | 1 | 1 | 0 | 1 | 1 | 1 | 1 |
| DNAH5 | 2.52 | P4_VS_P1_P2_P3_Lumbar_Coeliac_Stellate_Sphenopalatine | 0 | 0 | 0 | 1 | 0 | 0 | 0 | 0 |
| SEMA5A | 2.52 | P4_VS_P1_P2_P3_Lumbar_Coeliac_Stellate_Sphenopalatine | 0 | 0 | 0 | 1 | 0 | 0 | 0 | 0 |
| PDYN | 2.52 | Lumbar_VS_P1_P2_P3_P4_Coeliac_Stellate_Sphenopalatine | 0 | 0 | 0 | 0 | 1 | 0 | 0 | 0 |
| RALYL | 2.52 | P1_P3_P4_Lumbar_Coeliac_Stellate_Sphenopalatine_VS_P2 | 1 | 0 | 1 | 1 | 1 | 1 | 1 | 1 |

| | | | | | | | | | | |
|---|---|---|---|---|---|---|---|---|---|---|
| BMP7 | 2.51 | P4_VS_P1_P2_P3_Lumbar_Coeliac_Stellate_Sphenopalatine | 0 | 0 | 0 | 1 | 0 | 0 | 0 | 0 |
| HTRA4 | 2.51 | P4_VS_P1_P2_P3_Lumbar_Coeliac_Stellate_Sphenopalatine | 0 | 0 | 0 | 1 | 0 | 0 | 0 | 0 |
| SSU2 | 2.51 | P1_P2_P3_Lumbar_Coeliac_Stellate_Sphenopalatine_VS_P4 | 1 | 1 | 1 | 0 | 1 | 1 | 1 | 1 |
| KCNMB2 | 2.5 | Lumbar_VS_P1_P2_P3_P4_Coeliac_Stellate_Sphenopalatine | 0 | 0 | 0 | 0 | 1 | 0 | 0 | 0 |
| A330102I10F | 2.5 | P1_P2_P4_Lumbar_Coeliac_Stellate_Sphenopalatine_VS_P3 | 1 | 1 | 0 | 1 | 1 | 1 | 1 | 1 |
| WHRN | 2.49 | P4_VS_P1_P2_P3_Lumbar_Coeliac_Stellate_Sphenopalatine | 0 | 0 | 0 | 1 | 0 | 0 | 0 | 0 |
| C2CD4B | 2.49 | Lumbar_Coeliac_VS_P1_P2_P3_P4_Stellate_Sphenopalatine | 0 | 0 | 0 | 0 | 1 | 1 | 0 | 0 |
| MARCH1 | 2.48 | P2_VS_P1_P3_P4_Lumbar_Coeliac_Stellate_Sphenopalatine | 0 | 1 | 0 | 0 | 0 | 0 | 0 | 0 |
| BRINP3 | 2.48 | P4_VS_P1_P2_P3_Lumbar_Coeliac_Stellate_Sphenopalatine | 0 | 0 | 0 | 1 | 0 | 0 | 0 | 0 |
| GCGR | 2.47 | P2_VS_P1_P3_P4_Lumbar_Coeliac_Stellate_Sphenopalatine | 0 | 1 | 0 | 0 | 0 | 0 | 0 | 0 |
| IL19 | 2.47 | P4_VS_P1_P2_P3_Lumbar_Coeliac_Stellate_Sphenopalatine | 0 | 0 | 0 | 1 | 0 | 0 | 0 | 0 |
| RPRML | 2.47 | P4_VS_P1_P2_P3_Lumbar_Coeliac_Stellate_Sphenopalatine | 0 | 0 | 0 | 1 | 0 | 0 | 0 | 0 |
| DNASE1L3 | 2.47 | Lumbar_VS_P1_P2_P3_P4_Coeliac_Stellate_Sphenopalatine | 0 | 0 | 0 | 0 | 1 | 0 | 0 | 0 |
| OLFR78 | 2.47 | P1_P2_P4_Lumbar_Coeliac_Stellate_Sphenopalatine_VS_P3 | 1 | 1 | 0 | 1 | 1 | 1 | 1 | 1 |
| INHBB | 2.46 | P2_VS_P1_P3_P4_Lumbar_Coeliac_Stellate_Sphenopalatine | 0 | 1 | 0 | 0 | 0 | 0 | 0 | 0 |
| SLC1A2 | 2.46 | P1_P2_P4_VS_P3_Lumbar_Coeliac_Stellate_Sphenopalatine | 1 | 1 | 0 | 1 | 0 | 0 | 0 | 0 |
| CHST8 | 2.46 | P1_P2_P3_P4_Lumbar_Coeliac_Stellate_VS_Sphenopalatine | 1 | 1 | 1 | 1 | 1 | 1 | 1 | 0 |
| CDH1 | 2.45 | P4_VS_P1_P2_P3_Lumbar_Coeliac_Stellate_Sphenopalatine | 0 | 0 | 0 | 1 | 0 | 0 | 0 | 0 |
| MYO18B | 2.44 | P4_VS_P1_P2_P3_Lumbar_Coeliac_Stellate_Sphenopalatine | 0 | 0 | 0 | 1 | 0 | 0 | 0 | 0 |
| VCAN | 2.44 | P4_VS_P1_P2_P3_Lumbar_Coeliac_Stellate_Sphenopalatine | 0 | 0 | 0 | 1 | 0 | 0 | 0 | 0 |
| CACNG2 | 2.44 | P1_P2_P4_Lumbar_Coeliac_Stellate_Sphenopalatine_VS_P3 | 1 | 1 | 0 | 1 | 1 | 1 | 1 | 1 |
| SLC38A4 | 2.43 | P3_VS_P1_P2_P4_Lumbar_Coeliac_Stellate_Sphenopalatine | 0 | 0 | 1 | 0 | 0 | 0 | 0 | 0 |
| FGF10 | 2.43 | Lumbar_Coeliac_Stellate_VS_P1_P2_P3_P4_Sphenopalatine | 0 | 0 | 0 | 0 | 1 | 1 | 1 | 0 |
| KITL | 2.43 | P3_Lumbar_Coeliac_Stellate_VS_P1_P2_P4_Sphenopalatine | 0 | 0 | 1 | 0 | 1 | 1 | 1 | 0 |
| GM15270 | 2.42 | P4_VS_P1_P2_P3_Lumbar_Coeliac_Stellate_Sphenopalatine | 0 | 0 | 0 | 1 | 0 | 0 | 0 | 0 |
| LRRTM2 | 2.42 | P4_VS_P1_P2_P3_Lumbar_Coeliac_Stellate_Sphenopalatine | 0 | 0 | 0 | 1 | 0 | 0 | 0 | 0 |
| FBP2 | 2.42 | Lumbar_Coeliac_VS_P1_P2_P3_P4_Stellate_Sphenopalatine | 0 | 0 | 0 | 0 | 1 | 1 | 0 | 0 |
| C1QL1 | 2.41 | P3_Lumbar_Coeliac_Stellate_VS_P1_P2_P4_Sphenopalatine | 0 | 0 | 1 | 0 | 1 | 1 | 1 | 0 |
| WNT4 | 2.4 | P4_VS_P1_P2_P3_Lumbar_Coeliac_Stellate_Sphenopalatine | 0 | 0 | 0 | 1 | 0 | 0 | 0 | 0 |
| LDB2 | 2.4 | P4_VS_P1_P2_P3_Lumbar_Coeliac_Stellate_Sphenopalatine | 0 | 0 | 0 | 1 | 0 | 0 | 0 | 0 |

| | | | | | | | | | | |
|---|---|---|---|---|---|---|---|---|---|---|
| PTH1R | 2.4 | P1_P2_P4_Sphenopalatine_VS_P3_Lumbar_Coeliac_Stellate | 1 | 1 | 0 | 1 | 0 | 0 | 0 | 1 |
| SMCO3 | 2.39 | P1_P4_VS_P2_P3_Lumbar_Coeliac_Stellate_Sphenopalatine | 1 | 0 | 0 | 1 | 0 | 0 | 0 | 0 |
| CAMK2N1 | 2.39 | P1_P2_P3_P4_Lumbar_Coeliac_Sphenopalatine_VS_Stellate | 1 | 1 | 1 | 1 | 1 | 1 | 0 | 1 |
| GM29374 | 2.38 | Lumbar_Coeliac_VS_P1_P2_P3_P4_Stellate_Sphenopalatine | 0 | 0 | 0 | 0 | 1 | 1 | 0 | 0 |
| C1QTNF6 | 2.38 | P1_P2_P4_Sphenopalatine_VS_P3_Lumbar_Coeliac_Stellate | 1 | 1 | 0 | 1 | 0 | 0 | 0 | 1 |
| PEG12 | 2.37 | P2_VS_P1_P3_P4_Lumbar_Coeliac_Stellate_Sphenopalatine | 0 | 1 | 0 | 0 | 0 | 0 | 0 | 0 |
| LGALS7 | 2.37 | P3_Lumbar_Coeliac_Stellate_VS_P1_P2_P4_Sphenopalatine | 0 | 0 | 1 | 0 | 1 | 1 | 1 | 0 |
| RPP25 | 2.37 | P3_Lumbar_Coeliac_Stellate_VS_P1_P2_P4_Sphenopalatine | 0 | 0 | 1 | 0 | 1 | 1 | 1 | 0 |
| TCERG1L | 2.36 | P2_P4_VS_P1_P3_Lumbar_Coeliac_Stellate_Sphenopalatine | 0 | 1 | 0 | 1 | 0 | 0 | 0 | 0 |
| COL25A1 | 2.36 | P3_Coeliac_VS_P1_P2_P4_Lumbar_Stellate_Sphenopalatine | 0 | 0 | 1 | 0 | 0 | 1 | 0 | 0 |
| SLC5A7 | 2.36 | P1_P2_P3_P4_Sphenopalatine_VS_Lumbar_Coeliac_Stellate | 1 | 1 | 1 | 1 | 0 | 0 | 0 | 1 |
| CACNB2 | 2.35 | P4_VS_P1_P2_P3_Lumbar_Coeliac_Stellate_Sphenopalatine | 0 | 0 | 0 | 1 | 0 | 0 | 0 | 0 |
| TRABD2B | 2.35 | P4_VS_P1_P2_P3_Lumbar_Coeliac_Stellate_Sphenopalatine | 0 | 0 | 0 | 1 | 0 | 0 | 0 | 0 |
| TNS3 | 2.35 | P1_P2_P4_VS_P3_Lumbar_Coeliac_Stellate_Sphenopalatine | 1 | 1 | 0 | 1 | 0 | 0 | 0 | 0 |
| DKK1 | 2.35 | P1_P2_Lumbar_Coeliac_Stellate_VS_P3_P4_Sphenopalatine | 1 | 1 | 0 | 0 | 1 | 1 | 1 | 0 |
| RORB | 2.35 | P3_P4_Lumbar_Coeliac_Stellate_VS_P1_P2_Sphenopalatine | 0 | 0 | 1 | 1 | 1 | 1 | 1 | 0 |
| SMAD6 | 2.34 | P2_VS_P1_P3_P4_Lumbar_Coeliac_Stellate_Sphenopalatine | 0 | 1 | 0 | 0 | 0 | 0 | 0 | 0 |
| RSAD2 | 2.34 | P4_VS_P1_P2_P3_Lumbar_Coeliac_Stellate_Sphenopalatine | 0 | 0 | 0 | 1 | 0 | 0 | 0 | 0 |
| NPR3 | 2.34 | P4_VS_P1_P2_P3_Lumbar_Coeliac_Stellate_Sphenopalatine | 0 | 0 | 0 | 1 | 0 | 0 | 0 | 0 |
| ANTXR2 | 2.34 | P1_P2_P3_P4_Lumbar_Coeliac_Stellate_VS_Sphenopalatine | 1 | 1 | 1 | 1 | 1 | 1 | 1 | 0 |
| IGSF9 | 2.33 | Lumbar_Coeliac_VS_P1_P2_P3_P4_Stellate_Sphenopalatine | 0 | 0 | 0 | 0 | 1 | 1 | 0 | 0 |
| FAM163B | 2.33 | P3_Lumbar_Coeliac_Stellate_VS_P1_P2_P4_Sphenopalatine | 0 | 0 | 1 | 0 | 1 | 1 | 1 | 0 |
| DIO2 | 2.33 | P4_Sphenopalatine_VS_P1_P2_P3_Lumbar_Coeliac_Stellate | 0 | 0 | 0 | 1 | 0 | 0 | 0 | 1 |
| PROX1OS | 2.32 | P3_VS_P1_P2_P4_Lumbar_Coeliac_Stellate_Sphenopalatine | 0 | 0 | 1 | 0 | 0 | 0 | 0 | 0 |
| ANO2 | 2.32 | P2_P3_VS_P1_P4_Lumbar_Coeliac_Stellate_Sphenopalatine | 0 | 1 | 1 | 0 | 0 | 0 | 0 | 0 |
| MAL2 | 2.32 | P1_P3_P4_Lumbar_Coeliac_Stellate_Sphenopalatine_VS_P2 | 1 | 0 | 1 | 1 | 1 | 1 | 1 | 1 |
| PCP4L1 | 2.31 | P1_P2_VS_P3_P4_Lumbar_Coeliac_Stellate_Sphenopalatine | 1 | 1 | 0 | 0 | 0 | 0 | 0 | 0 |
| DSCAML1 | 2.31 | Sphenopalatine_VS_P1_P2_P3_P4_Lumbar_Coeliac_Stellate | 0 | 0 | 0 | 0 | 0 | 0 | 0 | 1 |
| NWD2 | 2.31 | P1_Sphenopalatine_VS_P2_P3_P4_Lumbar_Coeliac_Stellate | 1 | 0 | 0 | 0 | 0 | 0 | 0 | 1 |
| SLC18A3 | 2.31 | P1_P2_P4_Sphenopalatine_VS_P3_Lumbar_Coeliac_Stellate | 1 | 1 | 0 | 1 | 0 | 0 | 0 | 1 |

| | | | | | | | | | | |
|---|---|---|---|---|---|---|---|---|---|---|
| ZFP536 | 2.31 | P1_P2_P3_P4_Sphenopalatine_VS_Lumbar_Coeliac_Stellate | 1 | 1 | 1 | 1 | 0 | 0 | 0 | 1 |
| S100A4 | 2.3 | P3_VS_P1_P2_P4_Lumbar_Coeliac_Stellate_Sphenopalatine | 0 | 0 | 1 | 0 | 0 | 0 | 0 | 0 |
| TGFB2 | 2.3 | P4_VS_P1_P2_P3_Lumbar_Coeliac_Stellate_Sphenopalatine | 0 | 0 | 0 | 1 | 0 | 0 | 0 | 0 |
| TANC1 | 2.3 | P1_P2_P4_VS_P3_Lumbar_Coeliac_Stellate_Sphenopalatine | 1 | 1 | 0 | 1 | 0 | 0 | 0 | 0 |
| GALNTL6 | 2.3 | Lumbar_Coeliac_VS_P1_P2_P3_P4_Stellate_Sphenopalatine | 0 | 0 | 0 | 0 | 1 | 1 | 0 | 0 |
| PCDH15 | 2.3 | Sphenopalatine_VS_P1_P2_P3_P4_Lumbar_Coeliac_Stellate | 0 | 0 | 0 | 0 | 0 | 0 | 0 | 1 |
| MPPED1 | 2.29 | Lumbar_VS_P1_P2_P3_P4_Coeliac_Stellate_Sphenopalatine | 0 | 0 | 0 | 0 | 1 | 0 | 0 | 0 |
| RPH3A | 2.29 | P2_P3_Lumbar_Coeliac_Stellate_VS_P1_P4_Sphenopalatine | 0 | 1 | 1 | 0 | 1 | 1 | 1 | 0 |
| ADCY2 | 2.29 | P1_P2_P4_Sphenopalatine_VS_P3_Lumbar_Coeliac_Stellate | 1 | 1 | 0 | 1 | 0 | 0 | 0 | 1 |
| NPY2R | 2.29 | P1_P2_P3_P4_Coeliac_Sphenopalatine_VS_Lumbar_Stellate | 1 | 1 | 1 | 1 | 0 | 1 | 0 | 1 |
| SUSD5 | 2.29 | P1_P2_P4_Lumbar_Coeliac_Stellate_Sphenopalatine_VS_P3 | 1 | 1 | 0 | 1 | 1 | 1 | 1 | 1 |
| EPHB1 | 2.28 | P1_P4_VS_P2_P3_Lumbar_Coeliac_Stellate_Sphenopalatine | 1 | 0 | 0 | 1 | 0 | 0 | 0 | 0 |
| PRKG1 | 2.28 | P1_P2_P3_P4_VS_Lumbar_Coeliac_Stellate_Sphenopalatine | 1 | 1 | 1 | 1 | 0 | 0 | 0 | 0 |
| MASP1 | 2.28 | P1_P2_P4_Sphenopalatine_VS_P3_Lumbar_Coeliac_Stellate | 1 | 1 | 0 | 1 | 0 | 0 | 0 | 1 |
| ADAMTS14 | 2.26 | P2_VS_P1_P3_P4_Lumbar_Coeliac_Stellate_Sphenopalatine | 0 | 1 | 0 | 0 | 0 | 0 | 0 | 0 |
| LEPR | 2.25 | Lumbar_VS_P1_P2_P3_P4_Coeliac_Stellate_Sphenopalatine | 0 | 0 | 0 | 0 | 1 | 0 | 0 | 0 |
| CACNG5 | 2.25 | P1_P2_P3_Lumbar_Coeliac_Stellate_VS_P4_Sphenopalatine | 1 | 1 | 1 | 0 | 1 | 1 | 1 | 0 |
| RXFP1 | 2.25 | P4_Lumbar_Coeliac_Stellate_VS_P1_P2_P3_Sphenopalatine | 0 | 0 | 0 | 1 | 1 | 1 | 1 | 0 |
| GM11789 | 2.24 | P2_VS_P1_P3_P4_Lumbar_Coeliac_Stellate_Sphenopalatine | 0 | 1 | 0 | 0 | 0 | 0 | 0 | 0 |
| GULO | 2.24 | P3_VS_P1_P2_P4_Lumbar_Coeliac_Stellate_Sphenopalatine | 0 | 0 | 1 | 0 | 0 | 0 | 0 | 0 |
| MMP17 | 2.24 | P2_P3_VS_P1_P4_Lumbar_Coeliac_Stellate_Sphenopalatine | 0 | 1 | 1 | 0 | 0 | 0 | 0 | 0 |
| NXPH1 | 2.23 | P4_VS_P1_P2_P3_Lumbar_Coeliac_Stellate_Sphenopalatine | 0 | 0 | 0 | 1 | 0 | 0 | 0 | 0 |
| MDGA1 | 2.23 | P4_VS_P1_P2_P3_Lumbar_Coeliac_Stellate_Sphenopalatine | 0 | 0 | 0 | 1 | 0 | 0 | 0 | 0 |
| SERPINB7 | 2.22 | Sphenopalatine_VS_P1_P2_P3_P4_Lumbar_Coeliac_Stellate | 0 | 0 | 0 | 0 | 0 | 0 | 0 | 1 |
| SLC24A2 | 2.22 | P1_P2_P4_Sphenopalatine_VS_P3_Lumbar_Coeliac_Stellate | 1 | 1 | 0 | 1 | 0 | 0 | 0 | 1 |
| GM12166 | 2.22 | P1_P2_P3_P4_Stellate_Sphenopalatine_VS_Lumbar_Coeliac | 1 | 1 | 1 | 1 | 0 | 0 | 1 | 1 |
| HSD17B14 | 2.21 | P1_P2_P3_Lumbar_Coeliac_Stellate_VS_P4_Sphenopalatine | 1 | 1 | 1 | 0 | 1 | 1 | 1 | 0 |
| HOXA6 | 2.2 | P4_VS_P1_P2_P3_Lumbar_Coeliac_Stellate_Sphenopalatine | 0 | 0 | 0 | 1 | 0 | 0 | 0 | 0 |
| GM48957 | 2.2 | Lumbar_Coeliac_VS_P1_P2_P3_P4_Stellate_Sphenopalatine | 0 | 0 | 0 | 0 | 1 | 1 | 0 | 0 |
| ARHGAP20 | 2.2 | P3_P4_Lumbar_Coeliac_VS_P1_P2_Stellate_Sphenopalatine | 0 | 0 | 1 | 1 | 1 | 1 | 0 | 0 |

| | | | | | | | | | | |
|---|---|---|---|---|---|---|---|---|---|---|
| FAM20A | 2.2 | Lumbar_Coeliac_Stellate_VS_P1_P2_P3_P4_Sphenopalatine | 0 | 0 | 0 | 0 | 1 | 1 | 1 | 0 |
| STK17B | 2.2 | P3_Lumbar_Coeliac_Stellate_VS_P1_P2_P4_Sphenopalatine | 0 | 0 | 1 | 0 | 1 | 1 | 1 | 0 |
| PDE5A | 2.2 | P1_P4_Sphenopalatine_VS_P2_P3_Lumbar_Coeliac_Stellate | 1 | 0 | 0 | 1 | 0 | 0 | 0 | 1 |
| BC030500 | 2.2 | P3_Lumbar_Coeliac_Stellate_Sphenopalatine_VS_P1_P2_P4 | 0 | 0 | 1 | 0 | 1 | 1 | 1 | 1 |
| GM2464 | 2.2 | P1_P3_P4_Lumbar_Coeliac_Stellate_Sphenopalatine_VS_P2 | 1 | 0 | 1 | 1 | 1 | 1 | 1 | 1 |
| NOG | 2.19 | P2_VS_P1_P3_P4_Lumbar_Coeliac_Stellate_Sphenopalatine | 0 | 1 | 0 | 0 | 0 | 0 | 0 | 0 |
| NETO1 | 2.19 | P2_VS_P1_P3_P4_Lumbar_Coeliac_Stellate_Sphenopalatine | 0 | 1 | 0 | 0 | 0 | 0 | 0 | 0 |
| TMEM163 | 2.19 | P4_VS_P1_P2_P3_Lumbar_Coeliac_Stellate_Sphenopalatine | 0 | 0 | 0 | 1 | 0 | 0 | 0 | 0 |
| SYT13 | 2.19 | Sphenopalatine_VS_P1_P2_P3_P4_Lumbar_Coeliac_Stellate | 0 | 0 | 0 | 0 | 0 | 0 | 0 | 1 |
| 2610307P16I | 2.19 | P1_P2_P3_Lumbar_Coeliac_Stellate_Sphenopalatine_VS_P4 | 1 | 1 | 1 | 0 | 1 | 1 | 1 | 1 |
| LMO1 | 2.19 | P1_P3_P4_Lumbar_Coeliac_Stellate_Sphenopalatine_VS_P2 | 1 | 0 | 1 | 1 | 1 | 1 | 1 | 1 |
| CNTN6 | 2.18 | P4_VS_P1_P2_P3_Lumbar_Coeliac_Stellate_Sphenopalatine | 0 | 0 | 0 | 1 | 0 | 0 | 0 | 0 |
| GHR | 2.18 | P4_VS_P1_P2_P3_Lumbar_Coeliac_Stellate_Sphenopalatine | 0 | 0 | 0 | 1 | 0 | 0 | 0 | 0 |
| VSTM2B | 2.18 | Lumbar_VS_P1_P2_P3_P4_Coeliac_Stellate_Sphenopalatine | 0 | 0 | 0 | 0 | 1 | 0 | 0 | 0 |
| STUM | 2.18 | P2_Sphenopalatine_VS_P1_P3_P4_Lumbar_Coeliac_Stellate | 0 | 1 | 0 | 0 | 0 | 0 | 0 | 1 |
| GALNT3 | 2.17 | Lumbar_Coeliac_VS_P1_P2_P3_P4_Stellate_Sphenopalatine | 0 | 0 | 0 | 0 | 1 | 1 | 0 | 0 |
| SOX13 | 2.16 | P4_VS_P1_P2_P3_Lumbar_Coeliac_Stellate_Sphenopalatine | 0 | 0 | 0 | 1 | 0 | 0 | 0 | 0 |
| PLPP3 | 2.16 | P4_VS_P1_P2_P3_Lumbar_Coeliac_Stellate_Sphenopalatine | 0 | 0 | 0 | 1 | 0 | 0 | 0 | 0 |
| NR0B1 | 2.16 | P4_VS_P1_P2_P3_Lumbar_Coeliac_Stellate_Sphenopalatine | 0 | 0 | 0 | 1 | 0 | 0 | 0 | 0 |
| CLDN11 | 2.16 | P3_Lumbar_Coeliac_Stellate_VS_P1_P2_P4_Sphenopalatine | 0 | 0 | 1 | 0 | 1 | 1 | 1 | 0 |
| SVIL | 2.16 | P1_P2_P4_Sphenopalatine_VS_P3_Lumbar_Coeliac_Stellate | 1 | 1 | 0 | 1 | 0 | 0 | 0 | 1 |
| CACHD1 | 2.15 | P1_P2_P3_P4_Lumbar_Coeliac_Sphenopalatine_VS_Stellate | 1 | 1 | 1 | 1 | 1 | 1 | 0 | 1 |
| HCRTR1 | 2.15 | P1_P3_P4_Lumbar_Coeliac_Stellate_Sphenopalatine_VS_P2 | 1 | 0 | 1 | 1 | 1 | 1 | 1 | 1 |
| SIAH3 | 2.14 | P4_VS_P1_P2_P3_Lumbar_Coeliac_Stellate_Sphenopalatine | 0 | 0 | 0 | 1 | 0 | 0 | 0 | 0 |
| SORCS1 | 2.14 | P4_VS_P1_P2_P3_Lumbar_Coeliac_Stellate_Sphenopalatine | 0 | 0 | 0 | 1 | 0 | 0 | 0 | 0 |
| SLC30A10 | 2.14 | P1_P4_VS_P2_P3_Lumbar_Coeliac_Stellate_Sphenopalatine | 1 | 0 | 0 | 1 | 0 | 0 | 0 | 0 |
| GM19434 | 2.14 | P2_P4_VS_P1_P3_Lumbar_Coeliac_Stellate_Sphenopalatine | 0 | 1 | 0 | 1 | 0 | 0 | 0 | 0 |
| WNT5A | 2.14 | P2_P4_VS_P1_P3_Lumbar_Coeliac_Stellate_Sphenopalatine | 0 | 1 | 0 | 1 | 0 | 0 | 0 | 0 |
| MCC | 2.14 | P1_P2_P4_VS_P3_Lumbar_Coeliac_Stellate_Sphenopalatine | 1 | 1 | 0 | 1 | 0 | 0 | 0 | 0 |
| DSP | 2.14 | Coeliac_VS_P1_P2_P3_P4_Lumbar_Stellate_Sphenopalatine | 0 | 0 | 0 | 0 | 0 | 1 | 0 | 0 |

| | | | | | | | | | | |
|---|---|---|---|---|---|---|---|---|---|---|
| EGLN3 | 2.14 | P4_Lumbar_Coeliac_Stellate_VS_P1_P2_P3_Sphenopalatine | 0 | 0 | 0 | 1 | 1 | 1 | 1 | 0 |
| ARG1 | 2.14 | P3_P4_Lumbar_Coeliac_Stellate_VS_P1_P2_Sphenopalatine | 0 | 0 | 1 | 1 | 1 | 1 | 1 | 0 |
| H19 | 2.14 | Sphenopalatine_VS_P1_P2_P3_P4_Lumbar_Coeliac_Stellate | 0 | 0 | 0 | 0 | 0 | 0 | 0 | 1 |
| RGS4 | 2.14 | P1_P2_P3_Lumbar_Coeliac_Stellate_Sphenopalatine_VS_P4 | 1 | 1 | 1 | 0 | 1 | 1 | 1 | 1 |
| ADCY1 | 2.14 | P1_P2_P3_Lumbar_Coeliac_Stellate_Sphenopalatine_VS_P4 | 1 | 1 | 1 | 0 | 1 | 1 | 1 | 1 |
| KIT | 2.13 | P2_VS_P1_P3_P4_Lumbar_Coeliac_Stellate_Sphenopalatine | 0 | 1 | 0 | 0 | 0 | 0 | 0 | 0 |
| 1110032F04I | 2.13 | P4_VS_P1_P2_P3_Lumbar_Coeliac_Stellate_Sphenopalatine | 0 | 0 | 0 | 1 | 0 | 0 | 0 | 0 |
| AI115009 | 2.13 | P4_VS_P1_P2_P3_Lumbar_Coeliac_Stellate_Sphenopalatine | 0 | 0 | 0 | 1 | 0 | 0 | 0 | 0 |
| HOXA3 | 2.13 | P4_VS_P1_P2_P3_Lumbar_Coeliac_Stellate_Sphenopalatine | 0 | 0 | 0 | 1 | 0 | 0 | 0 | 0 |
| BMP4 | 2.13 | P1_P2_P3_P4_Lumbar_Coeliac_VS_Stellate_Sphenopalatine | 1 | 1 | 1 | 1 | 1 | 1 | 0 | 0 |
| TUNAR | 2.13 | Sphenopalatine_VS_P1_P2_P3_P4_Lumbar_Coeliac_Stellate | 0 | 0 | 0 | 0 | 0 | 0 | 0 | 1 |
| GM11549 | 2.13 | Lumbar_Stellate_Sphenopalatine_VS_P1_P2_P3_P4_Coeliac | 0 | 0 | 0 | 0 | 1 | 0 | 1 | 1 |
| COL24A1 | 2.12 | P1_VS_P2_P3_P4_Lumbar_Coeliac_Stellate_Sphenopalatine | 1 | 0 | 0 | 0 | 0 | 0 | 0 | 0 |
| SYTL4 | 2.12 | P3_VS_P1_P2_P4_Lumbar_Coeliac_Stellate_Sphenopalatine | 0 | 0 | 1 | 0 | 0 | 0 | 0 | 0 |
| NTRK3 | 2.12 | P1_P2_P4_VS_P3_Lumbar_Coeliac_Stellate_Sphenopalatine | 1 | 1 | 0 | 1 | 0 | 0 | 0 | 0 |
| GLB1L2 | 2.11 | P2_VS_P1_P3_P4_Lumbar_Coeliac_Stellate_Sphenopalatine | 0 | 1 | 0 | 0 | 0 | 0 | 0 | 0 |
| ADAMTSL2 | 2.11 | P3_VS_P1_P2_P4_Lumbar_Coeliac_Stellate_Sphenopalatine | 0 | 0 | 1 | 0 | 0 | 0 | 0 | 0 |
| CCKAR | 2.11 | Coeliac_VS_P1_P2_P3_P4_Lumbar_Stellate_Sphenopalatine | 0 | 0 | 0 | 0 | 0 | 1 | 0 | 0 |
| PPP1R17 | 2.11 | P3_Lumbar_Coeliac_Stellate_VS_P1_P2_P4_Sphenopalatine | 0 | 0 | 1 | 0 | 1 | 1 | 1 | 0 |
| RTN4RL1 | 2.11 | P1_P2_P4_Lumbar_Coeliac_Stellate_Sphenopalatine_VS_P3 | 1 | 1 | 0 | 1 | 1 | 1 | 1 | 1 |
| CBLN2 | 2.1 | P1_VS_P2_P3_P4_Lumbar_Coeliac_Stellate_Sphenopalatine | 1 | 0 | 0 | 0 | 0 | 0 | 0 | 0 |
| GM16105 | 2.1 | Lumbar_Coeliac_VS_P1_P2_P3_P4_Stellate_Sphenopalatine | 0 | 0 | 0 | 0 | 1 | 1 | 0 | 0 |
| ELN | 2.09 | P1_P2_P3_P4_VS_Lumbar_Coeliac_Stellate_Sphenopalatine | 1 | 1 | 1 | 1 | 0 | 0 | 0 | 0 |
| STK32B | 2.09 | P4_Lumbar_Coeliac_Stellate_VS_P1_P2_P3_Sphenopalatine | 0 | 0 | 0 | 1 | 1 | 1 | 1 | 0 |
| RAPGEF5 | 2.09 | P3_P4_Lumbar_Coeliac_Stellate_VS_P1_P2_Sphenopalatine | 0 | 0 | 1 | 1 | 1 | 1 | 1 | 0 |
| B230334C09I | 2.09 | Sphenopalatine_VS_P1_P2_P3_P4_Lumbar_Coeliac_Stellate | 0 | 0 | 0 | 0 | 0 | 0 | 0 | 1 |
| ASCL1 | 2.09 | Sphenopalatine_VS_P1_P2_P3_P4_Lumbar_Coeliac_Stellate | 0 | 0 | 0 | 0 | 0 | 0 | 0 | 1 |
| FAM122B | 2.09 | P1_P2_P3_P4_Sphenopalatine_VS_Lumbar_Coeliac_Stellate | 1 | 1 | 1 | 1 | 0 | 0 | 0 | 1 |
| HBA-A2 | 2.09 | P1_P2_P3_P4_Stellate_Sphenopalatine_VS_Lumbar_Coeliac | 1 | 1 | 1 | 1 | 0 | 0 | 1 | 1 |
| HOPX | 2.08 | P4_VS_P1_P2_P3_Lumbar_Coeliac_Stellate_Sphenopalatine | 0 | 0 | 0 | 1 | 0 | 0 | 0 | 0 |

| | | | | | | | | | | |
|---|---|---|---|---|---|---|---|---|---|---|
| LRFN2 | 2.08 | P4_VS_P1_P2_P3_Lumbar_Coeliac_Stellate_Sphenopalatine | 0 | 0 | 0 | 1 | 0 | 0 | 0 | 0 |
| ROBO1 | 2.08 | P3_P4_VS_P1_P2_Lumbar_Coeliac_Stellate_Sphenopalatine | 0 | 0 | 1 | 1 | 0 | 0 | 0 | 0 |
| ADCYAP1 | 2.08 | P1_P2_P4_Sphenopalatine_VS_P3_Lumbar_Coeliac_Stellate | 1 | 1 | 0 | 1 | 0 | 0 | 0 | 1 |
| CNTN4 | 2.07 | P4_Lumbar_VS_P1_P2_P3_Coeliac_Stellate_Sphenopalatine | 0 | 0 | 0 | 1 | 1 | 0 | 0 | 0 |
| 9230102O04 | 2.07 | P3_Lumbar_Coeliac_Stellate_VS_P1_P2_P4_Sphenopalatine | 0 | 0 | 1 | 0 | 1 | 1 | 1 | 0 |
| LSR | 2.07 | Sphenopalatine_VS_P1_P2_P3_P4_Lumbar_Coeliac_Stellate | 0 | 0 | 0 | 0 | 0 | 0 | 0 | 1 |
| CCL1 | 2.06 | P3_VS_P1_P2_P4_Lumbar_Coeliac_Stellate_Sphenopalatine | 0 | 0 | 1 | 0 | 0 | 0 | 0 | 0 |
| PXYLP1 | 2.06 | P1_P2_P3_Sphenopalatine_VS_P4_Lumbar_Coeliac_Stellate | 1 | 1 | 1 | 0 | 0 | 0 | 0 | 1 |
| KCNK3 | 2.06 | P1_P2_P3_P4_Lumbar_Coeliac_Sphenopalatine_VS_Stellate | 1 | 1 | 1 | 1 | 1 | 1 | 0 | 1 |
| CLMN | 2.05 | P1_P2_P3_P4_VS_Lumbar_Coeliac_Stellate_Sphenopalatine | 1 | 1 | 1 | 1 | 0 | 0 | 0 | 0 |
| VWC2 | 2.04 | P1_VS_P2_P3_P4_Lumbar_Coeliac_Stellate_Sphenopalatine | 1 | 0 | 0 | 0 | 0 | 0 | 0 | 0 |
| TMEM132B | 2.04 | P2_VS_P1_P3_P4_Lumbar_Coeliac_Stellate_Sphenopalatine | 0 | 1 | 0 | 0 | 0 | 0 | 0 | 0 |
| CES5A | 2.04 | P4_VS_P1_P2_P3_Lumbar_Coeliac_Stellate_Sphenopalatine | 0 | 0 | 0 | 1 | 0 | 0 | 0 | 0 |
| PROCR | 2.04 | P3_Lumbar_Coeliac_Stellate_VS_P1_P2_P4_Sphenopalatine | 0 | 0 | 1 | 0 | 1 | 1 | 1 | 0 |
| GM2832 | 2.04 | Sphenopalatine_VS_P1_P2_P3_P4_Lumbar_Coeliac_Stellate | 0 | 0 | 0 | 0 | 0 | 0 | 0 | 1 |
| LOX | 2.03 | P4_VS_P1_P2_P3_Lumbar_Coeliac_Stellate_Sphenopalatine | 0 | 0 | 0 | 1 | 0 | 0 | 0 | 0 |
| KCNK9 | 2.03 | P1_Lumbar_Coeliac_Sphenopalatine_VS_P2_P3_P4_Stellate | 1 | 0 | 0 | 0 | 1 | 1 | 0 | 1 |
| TMEM158 | 2.03 | P1_P2_P3_P4_Lumbar_Coeliac_Sphenopalatine_VS_Stellate | 1 | 1 | 1 | 1 | 1 | 1 | 0 | 1 |
| SMCO4 | 2.02 | P4_VS_P1_P2_P3_Lumbar_Coeliac_Stellate_Sphenopalatine | 0 | 0 | 0 | 1 | 0 | 0 | 0 | 0 |
| RARB | 2.02 | P4_VS_P1_P2_P3_Lumbar_Coeliac_Stellate_Sphenopalatine | 0 | 0 | 0 | 1 | 0 | 0 | 0 | 0 |
| HIST1H4D | 2.02 | Lumbar_Coeliac_VS_P1_P2_P3_P4_Stellate_Sphenopalatine | 0 | 0 | 0 | 0 | 1 | 1 | 0 | 0 |
| HIST1H1E | 2.02 | Lumbar_Coeliac_VS_P1_P2_P3_P4_Stellate_Sphenopalatine | 0 | 0 | 0 | 0 | 1 | 1 | 0 | 0 |
| STEAP2 | 2.01 | P4_VS_P1_P2_P3_Lumbar_Coeliac_Stellate_Sphenopalatine | 0 | 0 | 0 | 1 | 0 | 0 | 0 | 0 |
| FAM187B | 2.01 | Sphenopalatine_VS_P1_P2_P3_P4_Lumbar_Coeliac_Stellate | 0 | 0 | 0 | 0 | 0 | 0 | 0 | 1 |
| CNR1 | 2.01 | P1_P2_P3_P4_Lumbar_Coeliac_Sphenopalatine_VS_Stellate | 1 | 1 | 1 | 1 | 1 | 1 | 0 | 1 |
| PCDH11X | 2 | P1_VS_P2_P3_P4_Lumbar_Coeliac_Stellate_Sphenopalatine | 1 | 0 | 0 | 0 | 0 | 0 | 0 | 0 |
| CYP4B1 | 2 | P3_VS_P1_P2_P4_Lumbar_Coeliac_Stellate_Sphenopalatine | 0 | 0 | 1 | 0 | 0 | 0 | 0 | 0 |
| CAMKV | 2 | P3_VS_P1_P2_P4_Lumbar_Coeliac_Stellate_Sphenopalatine | 0 | 0 | 1 | 0 | 0 | 0 | 0 | 0 |
| SCN7A | 2 | P4_VS_P1_P2_P3_Lumbar_Coeliac_Stellate_Sphenopalatine | 0 | 0 | 0 | 1 | 0 | 0 | 0 | 0 |
| CSMD1 | 2 | P4_VS_P1_P2_P3_Lumbar_Coeliac_Stellate_Sphenopalatine | 0 | 0 | 0 | 1 | 0 | 0 | 0 | 0 |

| | | | | | | | | | | |
|---|---|---|---|---|---|---|---|---|---|---|
| PRKD1 | 2 | P4_VS_P1_P2_P3_Lumbar_Coeliac_Stellate_Sphenopalatine | 0 | 0 | 0 | 1 | 0 | 0 | 0 | 0 |
| CDK6 | 2 | P2_P4_VS_P1_P3_Lumbar_Coeliac_Stellate_Sphenopalatine | 0 | 1 | 0 | 1 | 0 | 0 | 0 | 0 |
| PROX1 | 2 | P1_P2_P3_Lumbar_Coeliac_VS_P4_Stellate_Sphenopalatine | 1 | 1 | 1 | 0 | 1 | 1 | 0 | 0 |
| PLD2 | 2 | P1_P2_P4_Sphenopalatine_VS_P3_Lumbar_Coeliac_Stellate | 1 | 1 | 0 | 1 | 0 | 0 | 0 | 1 |
| GPER1 | 1.99 | P3_VS_P1_P2_P4_Lumbar_Coeliac_Stellate_Sphenopalatine | 0 | 0 | 1 | 0 | 0 | 0 | 0 | 0 |
| KCNJ6 | 1.99 | P4_VS_P1_P2_P3_Lumbar_Coeliac_Stellate_Sphenopalatine | 0 | 0 | 0 | 1 | 0 | 0 | 0 | 0 |
| GAL | 1.99 | Lumbar_Coeliac_Stellate_VS_P1_P2_P3_P4_Sphenopalatine | 0 | 0 | 0 | 0 | 1 | 1 | 1 | 0 |
| DCC | 1.99 | P1_P2_P3_Lumbar_Coeliac_Stellate_Sphenopalatine_VS_P4 | 1 | 1 | 1 | 0 | 1 | 1 | 1 | 1 |
| HS3ST2 | 1.98 | P3_VS_P1_P2_P4_Lumbar_Coeliac_Stellate_Sphenopalatine | 0 | 0 | 1 | 0 | 0 | 0 | 0 | 0 |
| MGST1 | 1.98 | P4_VS_P1_P2_P3_Lumbar_Coeliac_Stellate_Sphenopalatine | 0 | 0 | 0 | 1 | 0 | 0 | 0 | 0 |
| AARD | 1.97 | P4_VS_P1_P2_P3_Lumbar_Coeliac_Stellate_Sphenopalatine | 0 | 0 | 0 | 1 | 0 | 0 | 0 | 0 |
| ADCY8 | 1.97 | P4_VS_P1_P2_P3_Lumbar_Coeliac_Stellate_Sphenopalatine | 0 | 0 | 0 | 1 | 0 | 0 | 0 | 0 |
| TRMT9B | 1.97 | P1_P2_P4_VS_P3_Lumbar_Coeliac_Stellate_Sphenopalatine | 1 | 1 | 0 | 1 | 0 | 0 | 0 | 0 |
| DKK2 | 1.97 | P3_Lumbar_Coeliac_VS_P1_P2_P4_Stellate_Sphenopalatine | 0 | 0 | 1 | 0 | 1 | 1 | 0 | 0 |
| ADORA1 | 1.97 | P1_P2_P3_P4_Sphenopalatine_VS_Lumbar_Coeliac_Stellate | 1 | 1 | 1 | 1 | 0 | 0 | 0 | 1 |
| FAM135B | 1.96 | P3_VS_P1_P2_P4_Lumbar_Coeliac_Stellate_Sphenopalatine | 0 | 0 | 1 | 0 | 0 | 0 | 0 | 0 |
| MAP3K5 | 1.96 | P4_VS_P1_P2_P3_Lumbar_Coeliac_Stellate_Sphenopalatine | 0 | 0 | 0 | 1 | 0 | 0 | 0 | 0 |
| UBTD2 | 1.96 | P1_P2_P3_P4_Lumbar_Coeliac_VS_Stellate_Sphenopalatine | 1 | 1 | 1 | 1 | 1 | 1 | 0 | 0 |
| CHN2 | 1.96 | P1_P4_Sphenopalatine_VS_P2_P3_Lumbar_Coeliac_Stellate | 1 | 0 | 0 | 1 | 0 | 0 | 0 | 1 |
| CAMK2A | 1.95 | P3_VS_P1_P2_P4_Lumbar_Coeliac_Stellate_Sphenopalatine | 0 | 0 | 1 | 0 | 0 | 0 | 0 | 0 |
| HHIP | 1.95 | P2_P4_VS_P1_P3_Lumbar_Coeliac_Stellate_Sphenopalatine | 0 | 1 | 0 | 1 | 0 | 0 | 0 | 0 |
| LHFPL2 | 1.95 | P2_P4_VS_P1_P3_Lumbar_Coeliac_Stellate_Sphenopalatine | 0 | 1 | 0 | 1 | 0 | 0 | 0 | 0 |
| CHST5 | 1.95 | Lumbar_VS_P1_P2_P3_P4_Coeliac_Stellate_Sphenopalatine | 0 | 0 | 0 | 0 | 1 | 0 | 0 | 0 |
| GM5468 | 1.95 | Lumbar_Coeliac_VS_P1_P2_P3_P4_Stellate_Sphenopalatine | 0 | 0 | 0 | 0 | 1 | 1 | 0 | 0 |
| NKAIN3 | 1.95 | P1_P3_P4_Lumbar_Sphenopalatine_VS_P2_Coeliac_Stellate | 1 | 0 | 1 | 1 | 1 | 0 | 0 | 1 |
| GM16499.1 | 1.95 | P1_P3_P4_Lumbar_Coeliac_Stellate_Sphenopalatine_VS_P2 | 1 | 0 | 1 | 1 | 1 | 1 | 1 | 1 |
| TENT5C | 1.94 | P4_VS_P1_P2_P3_Lumbar_Coeliac_Stellate_Sphenopalatine | 0 | 0 | 0 | 1 | 0 | 0 | 0 | 0 |
| MYLIP | 1.94 | P4_VS_P1_P2_P3_Lumbar_Coeliac_Stellate_Sphenopalatine | 0 | 0 | 0 | 1 | 0 | 0 | 0 | 0 |
| SYNDIG1 | 1.94 | P1_P2_Sphenopalatine_VS_P3_P4_Lumbar_Coeliac_Stellate | 1 | 1 | 0 | 0 | 0 | 0 | 0 | 1 |
| HBA-A1 | 1.94 | P1_P2_P3_P4_Coeliac_Stellate_Sphenopalatine_VS_Lumbar | 1 | 1 | 1 | 1 | 0 | 1 | 1 | 1 |

| | | | | | | | | | | |
|---|---|---|---|---|---|---|---|---|---|---|
| SP5 | 1.93 | P4_VS_P1_P2_P3_Lumbar_Coeliac_Stellate_Sphenopalatine | 0 | 0 | 0 | 1 | 0 | 0 | 0 | 0 |
| RARRES1 | 1.93 | P3_P4_Lumbar_Coeliac_Stellate_VS_P1_P2_Sphenopalatine | 0 | 0 | 1 | 1 | 1 | 1 | 1 | 0 |
| CDH6 | 1.93 | P1_P2_P4_Lumbar_Coeliac_Stellate_Sphenopalatine_VS_P3 | 1 | 1 | 0 | 1 | 1 | 1 | 1 | 1 |
| KLB | 1.92 | P4_VS_P1_P2_P3_Lumbar_Coeliac_Stellate_Sphenopalatine | 0 | 0 | 0 | 1 | 0 | 0 | 0 | 0 |
| GAD2 | 1.92 | Coeliac_VS_P1_P2_P3_P4_Lumbar_Stellate_Sphenopalatine | 0 | 0 | 0 | 0 | 0 | 1 | 0 | 0 |
| ZEB2 | 1.92 | P1_P2_P3_P4_Coeliac_Sphenopalatine_VS_Lumbar_Stellate | 1 | 1 | 1 | 1 | 0 | 1 | 0 | 1 |
| A830092H15 | 1.91 | P1_VS_P2_P3_P4_Lumbar_Coeliac_Stellate_Sphenopalatine | 1 | 0 | 0 | 0 | 0 | 0 | 0 | 0 |
| B3GAT1 | 1.91 | P2_P4_Lumbar_Coeliac_Stellate_VS_P1_P3_Sphenopalatine | 0 | 1 | 0 | 1 | 1 | 1 | 1 | 0 |
| DLL4 | 1.9 | P4_VS_P1_P2_P3_Lumbar_Coeliac_Stellate_Sphenopalatine | 0 | 0 | 0 | 1 | 0 | 0 | 0 | 0 |
| RGS6 | 1.9 | P4_VS_P1_P2_P3_Lumbar_Coeliac_Stellate_Sphenopalatine | 0 | 0 | 0 | 1 | 0 | 0 | 0 | 0 |
| ADGRG2 | 1.9 | Lumbar_VS_P1_P2_P3_P4_Coeliac_Stellate_Sphenopalatine | 0 | 0 | 0 | 0 | 1 | 0 | 0 | 0 |
| PRODH | 1.9 | Lumbar_Coeliac_VS_P1_P2_P3_P4_Stellate_Sphenopalatine | 0 | 0 | 0 | 0 | 1 | 1 | 0 | 0 |
| CFAP61 | 1.9 | Lumbar_Coeliac_Stellate_VS_P1_P2_P3_P4_Sphenopalatine | 0 | 0 | 0 | 0 | 1 | 1 | 1 | 0 |
| DMKN | 1.9 | Lumbar_Coeliac_Stellate_VS_P1_P2_P3_P4_Sphenopalatine | 0 | 0 | 0 | 0 | 1 | 1 | 1 | 0 |
| ITGA2 | 1.89 | P3_VS_P1_P2_P4_Lumbar_Coeliac_Stellate_Sphenopalatine | 0 | 0 | 1 | 0 | 0 | 0 | 0 | 0 |
| HSF4 | 1.88 | P4_VS_P1_P2_P3_Lumbar_Coeliac_Stellate_Sphenopalatine | 0 | 0 | 0 | 1 | 0 | 0 | 0 | 0 |
| GABRA1 | 1.88 | P2_P4_VS_P1_P3_Lumbar_Coeliac_Stellate_Sphenopalatine | 0 | 1 | 0 | 1 | 0 | 0 | 0 | 0 |
| CCDC85A | 1.88 | P1_P2_P3_P4_Lumbar_Coeliac_VS_Stellate_Sphenopalatine | 1 | 1 | 1 | 1 | 1 | 1 | 0 | 0 |
| SPINK8 | 1.87 | P4_VS_P1_P2_P3_Lumbar_Coeliac_Stellate_Sphenopalatine | 0 | 0 | 0 | 1 | 0 | 0 | 0 | 0 |
| BANF2OS | 1.87 | P3_Lumbar_Coeliac_Stellate_VS_P1_P2_P4_Sphenopalatine | 0 | 0 | 1 | 0 | 1 | 1 | 1 | 0 |
| TRF | 1.87 | P3_Lumbar_Coeliac_Stellate_VS_P1_P2_P4_Sphenopalatine | 0 | 0 | 1 | 0 | 1 | 1 | 1 | 0 |
| C030017B01I | 1.87 | P1_P2_P4_Lumbar_Coeliac_Stellate_VS_P3_Sphenopalatine | 1 | 1 | 0 | 1 | 1 | 1 | 1 | 0 |
| RNF182 | 1.87 | P1_P2_P3_P4_Lumbar_Coeliac_Sphenopalatine_VS_Stellate | 1 | 1 | 1 | 1 | 1 | 1 | 0 | 1 |
| NECAB2 | 1.86 | Lumbar_VS_P1_P2_P3_P4_Coeliac_Stellate_Sphenopalatine | 0 | 0 | 0 | 0 | 1 | 0 | 0 | 0 |
| JDP2 | 1.86 | P1_P2_P3_P4_Lumbar_Coeliac_VS_Stellate_Sphenopalatine | 1 | 1 | 1 | 1 | 1 | 1 | 0 | 0 |
| CNGA3 | 1.86 | Sphenopalatine_VS_P1_P2_P3_P4_Lumbar_Coeliac_Stellate | 0 | 0 | 0 | 0 | 0 | 0 | 0 | 1 |
| NKD2 | 1.86 | Sphenopalatine_VS_P1_P2_P3_P4_Lumbar_Coeliac_Stellate | 0 | 0 | 0 | 0 | 0 | 0 | 0 | 1 |
| PLPPR5 | 1.86 | P1_P3_Lumbar_Coeliac_Stellate_Sphenopalatine_VS_P2_P4 | 1 | 0 | 1 | 0 | 1 | 1 | 1 | 1 |
| CTTNBP2 | 1.86 | P1_P2_P4_Lumbar_Coeliac_Stellate_Sphenopalatine_VS_P3 | 1 | 1 | 0 | 1 | 1 | 1 | 1 | 1 |
| LRRC4C | 1.85 | P3_VS_P1_P2_P4_Lumbar_Coeliac_Stellate_Sphenopalatine | 0 | 0 | 1 | 0 | 0 | 0 | 0 | 0 |

| | | | | | | | | | | |
|---|---|---|---|---|---|---|---|---|---|---|
| JADE2 | 1.85 | P1_P2_P4_VS_P3_Lumbar_Coeliac_Stellate_Sphenopalatine | 1 | 1 | 0 | 1 | 0 | 0 | 0 | 0 |
| TGFBI | 1.85 | P1_Sphenopalatine_VS_P2_P3_P4_Lumbar_Coeliac_Stellate | 1 | 0 | 0 | 0 | 0 | 0 | 0 | 1 |
| SLC35F4 | 1.84 | P2_P4_VS_P1_P3_Lumbar_Coeliac_Stellate_Sphenopalatine | 0 | 1 | 0 | 1 | 0 | 0 | 0 | 0 |
| RGS5 | 1.84 | P3_Lumbar_Coeliac_Stellate_VS_P1_P2_P4_Sphenopalatine | 0 | 0 | 1 | 0 | 1 | 1 | 1 | 0 |
| KCNIP1 | 1.84 | Lumbar_Coeliac_Sphenopalatine_VS_P1_P2_P3_P4_Stellate | 0 | 0 | 0 | 0 | 1 | 1 | 0 | 1 |
| 4632428C04I | 1.83 | P1_P2_VS_P3_P4_Lumbar_Coeliac_Stellate_Sphenopalatine | 1 | 1 | 0 | 0 | 0 | 0 | 0 | 0 |
| GM26512 | 1.83 | P4_VS_P1_P2_P3_Lumbar_Coeliac_Stellate_Sphenopalatine | 0 | 0 | 0 | 1 | 0 | 0 | 0 | 0 |
| KRT9 | 1.83 | P4_Lumbar_VS_P1_P2_P3_Coeliac_Stellate_Sphenopalatine | 0 | 0 | 0 | 1 | 1 | 0 | 0 | 0 |
| CRLF1 | 1.82 | P4_VS_P1_P2_P3_Lumbar_Coeliac_Stellate_Sphenopalatine | 0 | 0 | 0 | 1 | 0 | 0 | 0 | 0 |
| NXPH4 | 1.82 | Lumbar_VS_P1_P2_P3_P4_Coeliac_Stellate_Sphenopalatine | 0 | 0 | 0 | 0 | 1 | 0 | 0 | 0 |
| NDUFA4L2 | 1.82 | Lumbar_Coeliac_VS_P1_P2_P3_P4_Stellate_Sphenopalatine | 0 | 0 | 0 | 0 | 1 | 1 | 0 | 0 |
| LMO4 | 1.82 | P1_P2_P4_Sphenopalatine_VS_P3_Lumbar_Coeliac_Stellate | 1 | 1 | 0 | 1 | 0 | 0 | 0 | 1 |
| PTPRK | 1.81 | P4_VS_P1_P2_P3_Lumbar_Coeliac_Stellate_Sphenopalatine | 0 | 0 | 0 | 1 | 0 | 0 | 0 | 0 |
| BDNF | 1.81 | P1_P2_P3_Lumbar_Coeliac_Stellate_VS_P4_Sphenopalatine | 1 | 1 | 1 | 0 | 1 | 1 | 1 | 0 |
| RNF128 | 1.81 | P1_P2_P3_P4_Sphenopalatine_VS_Lumbar_Coeliac_Stellate | 1 | 1 | 1 | 1 | 0 | 0 | 0 | 1 |
| NLGN1 | 1.81 | P1_P2_P4_Lumbar_Coeliac_Stellate_Sphenopalatine_VS_P3 | 1 | 1 | 0 | 1 | 1 | 1 | 1 | 1 |
| AQP1 | 1.8 | P3_VS_P1_P2_P4_Lumbar_Coeliac_Stellate_Sphenopalatine | 0 | 0 | 1 | 0 | 0 | 0 | 0 | 0 |
| PNLIP | 1.8 | P3_VS_P1_P2_P4_Lumbar_Coeliac_Stellate_Sphenopalatine | 0 | 0 | 1 | 0 | 0 | 0 | 0 | 0 |
| SCX | 1.8 | P4_VS_P1_P2_P3_Lumbar_Coeliac_Stellate_Sphenopalatine | 0 | 0 | 0 | 1 | 0 | 0 | 0 | 0 |
| DIRAS2 | 1.8 | Lumbar_Coeliac_VS_P1_P2_P3_P4_Stellate_Sphenopalatine | 0 | 0 | 0 | 0 | 1 | 1 | 0 | 0 |
| PPARA | 1.8 | Lumbar_Coeliac_VS_P1_P2_P3_P4_Stellate_Sphenopalatine | 0 | 0 | 0 | 0 | 1 | 1 | 0 | 0 |
| FLRT2 | 1.8 | P1_P2_P4_Lumbar_Sphenopalatine_VS_P3_Coeliac_Stellate | 1 | 1 | 0 | 1 | 1 | 0 | 0 | 1 |
| SCUBE1 | 1.8 | P1_P2_P3_P4_Lumbar_Coeliac_Sphenopalatine_VS_Stellate | 1 | 1 | 1 | 1 | 1 | 1 | 0 | 1 |
| SYNPO2 | 1.79 | Lumbar_VS_P1_P2_P3_P4_Coeliac_Stellate_Sphenopalatine | 0 | 0 | 0 | 0 | 1 | 0 | 0 | 0 |
| SEMA6A | 1.79 | Lumbar_VS_P1_P2_P3_P4_Coeliac_Stellate_Sphenopalatine | 0 | 0 | 0 | 0 | 1 | 0 | 0 | 0 |
| GM17167 | 1.79 | P1_P2_P3_P4_Lumbar_Coeliac_VS_Stellate_Sphenopalatine | 1 | 1 | 1 | 1 | 1 | 1 | 0 | 0 |
| DACH1 | 1.78 | P1_VS_P2_P3_P4_Lumbar_Coeliac_Stellate_Sphenopalatine | 1 | 0 | 0 | 0 | 0 | 0 | 0 | 0 |
| GM20559 | 1.78 | P3_VS_P1_P2_P4_Lumbar_Coeliac_Stellate_Sphenopalatine | 0 | 0 | 1 | 0 | 0 | 0 | 0 | 0 |
| ZFP703 | 1.78 | P4_VS_P1_P2_P3_Lumbar_Coeliac_Stellate_Sphenopalatine | 0 | 0 | 0 | 1 | 0 | 0 | 0 | 0 |
| PGR | 1.78 | P4_VS_P1_P2_P3_Lumbar_Coeliac_Stellate_Sphenopalatine | 0 | 0 | 0 | 1 | 0 | 0 | 0 | 0 |

| | | | | | | | | | | |
|---|---|---|---|---|---|---|---|---|---|---|
| KCNA5 | 1.78 | P1_P2_P4_Sphenopalatine_VS_P3_Lumbar_Coeliac_Stellate | 1 | 1 | 0 | 1 | 0 | 0 | 0 | 1 |
| XKR4 | 1.78 | P1_P2_P3_P4_Lumbar_Coeliac_Sphenopalatine_VS_Stellate | 1 | 1 | 1 | 1 | 1 | 1 | 0 | 1 |
| CEBPB | 1.78 | P1_P2_P3_P4_Lumbar_Coeliac_Sphenopalatine_VS_Stellate | 1 | 1 | 1 | 1 | 1 | 1 | 0 | 1 |
| NTRK2 | 1.77 | P4_VS_P1_P2_P3_Lumbar_Coeliac_Stellate_Sphenopalatine | 0 | 0 | 0 | 1 | 0 | 0 | 0 | 0 |
| SLC16A10 | 1.77 | Lumbar_Coeliac_VS_P1_P2_P3_P4_Stellate_Sphenopalatine | 0 | 0 | 0 | 0 | 1 | 1 | 0 | 0 |
| ARHGAP36 | 1.77 | P1_P2_P4_Lumbar_Coeliac_Stellate_VS_P3_Sphenopalatine | 1 | 1 | 0 | 1 | 1 | 1 | 1 | 0 |
| ESR1 | 1.76 | P2_P4_VS_P1_P3_Lumbar_Coeliac_Stellate_Sphenopalatine | 0 | 1 | 0 | 1 | 0 | 0 | 0 | 0 |
| BC049352 | 1.76 | Lumbar_VS_P1_P2_P3_P4_Coeliac_Stellate_Sphenopalatine | 0 | 0 | 0 | 0 | 1 | 0 | 0 | 0 |
| SYT2 | 1.76 | P1_P2_P3_Lumbar_Coeliac_Stellate_Sphenopalatine_VS_P4 | 1 | 1 | 1 | 0 | 1 | 1 | 1 | 1 |
| GAD1OS | 1.75 | P4_VS_P1_P2_P3_Lumbar_Coeliac_Stellate_Sphenopalatine | 0 | 0 | 0 | 1 | 0 | 0 | 0 | 0 |
| KCNAB1 | 1.75 | P4_Sphenopalatine_VS_P1_P2_P3_Lumbar_Coeliac_Stellate | 0 | 0 | 0 | 1 | 0 | 0 | 0 | 1 |
| CLSTN2 | 1.75 | P4_Sphenopalatine_VS_P1_P2_P3_Lumbar_Coeliac_Stellate | 0 | 0 | 0 | 1 | 0 | 0 | 0 | 1 |
| TIMP4 | 1.75 | P1_P2_P3_Lumbar_Coeliac_Stellate_Sphenopalatine_VS_P4 | 1 | 1 | 1 | 0 | 1 | 1 | 1 | 1 |
| 1700019D03I | 1.74 | P2_VS_P1_P3_P4_Lumbar_Coeliac_Stellate_Sphenopalatine | 0 | 1 | 0 | 0 | 0 | 0 | 0 | 0 |
| HIC1 | 1.74 | P4_VS_P1_P2_P3_Lumbar_Coeliac_Stellate_Sphenopalatine | 0 | 0 | 0 | 1 | 0 | 0 | 0 | 0 |
| CHST2 | 1.74 | P1_P2_P4_VS_P3_Lumbar_Coeliac_Stellate_Sphenopalatine | 1 | 1 | 0 | 1 | 0 | 0 | 0 | 0 |
| RGS9 | 1.74 | P1_P2_P3_P4_Lumbar_Coeliac_Sphenopalatine_VS_Stellate | 1 | 1 | 1 | 1 | 1 | 1 | 0 | 1 |
| TMIE | 1.73 | P3_VS_P1_P2_P4_Lumbar_Coeliac_Stellate_Sphenopalatine | 0 | 0 | 1 | 0 | 0 | 0 | 0 | 0 |
| BMP2 | 1.73 | Lumbar_VS_P1_P2_P3_P4_Coeliac_Stellate_Sphenopalatine | 0 | 0 | 0 | 0 | 1 | 0 | 0 | 0 |
| CDKL1 | 1.73 | Lumbar_Coeliac_VS_P1_P2_P3_P4_Stellate_Sphenopalatine | 0 | 0 | 0 | 0 | 1 | 1 | 0 | 0 |
| CPVL | 1.73 | Lumbar_Coeliac_Stellate_VS_P1_P2_P3_P4_Sphenopalatine | 0 | 0 | 0 | 0 | 1 | 1 | 1 | 0 |
| AIRN | 1.73 | P1_P2_P3_P4_Lumbar_Coeliac_Stellate_VS_Sphenopalatine | 1 | 1 | 1 | 1 | 1 | 1 | 1 | 0 |
| OLFR560 | 1.72 | P1_VS_P2_P3_P4_Lumbar_Coeliac_Stellate_Sphenopalatine | 1 | 0 | 0 | 0 | 0 | 0 | 0 | 0 |
| VLDLR | 1.72 | P2_P4_VS_P1_P3_Lumbar_Coeliac_Stellate_Sphenopalatine | 0 | 1 | 0 | 1 | 0 | 0 | 0 | 0 |
| SLC30A3 | 1.72 | Stellate_VS_P1_P2_P3_P4_Lumbar_Coeliac_Sphenopalatine | 0 | 0 | 0 | 0 | 0 | 0 | 1 | 0 |
| SORCS3 | 1.72 | P2_P3_Sphenopalatine_VS_P1_P4_Lumbar_Coeliac_Stellate | 0 | 1 | 1 | 0 | 0 | 0 | 0 | 1 |
| NACC2 | 1.72 | P1_P2_P4_Sphenopalatine_VS_P3_Lumbar_Coeliac_Stellate | 1 | 1 | 0 | 1 | 0 | 0 | 0 | 1 |
| ID4 | 1.72 | P1_P2_P4_Lumbar_Coeliac_Sphenopalatine_VS_P3_Stellate | 1 | 1 | 0 | 1 | 1 | 1 | 0 | 1 |
| FGF18 | 1.71 | P4_VS_P1_P2_P3_Lumbar_Coeliac_Stellate_Sphenopalatine | 0 | 0 | 0 | 1 | 0 | 0 | 0 | 0 |
| ADM | 1.71 | P3_P4_Lumbar_Coeliac_Stellate_VS_P1_P2_Sphenopalatine | 0 | 0 | 1 | 1 | 1 | 1 | 1 | 0 |

| | | | | | | | | | | |
|---|---|---|---|---|---|---|---|---|---|---|
| TMEM255B | 1.71 | P4_Sphenopalatine_VS_P1_P2_P3_Lumbar_Coeliac_Stellate | 0 | 0 | 0 | 1 | 0 | 0 | 0 | 1 |
| RGS8 | 1.7 | P2_VS_P1_P3_P4_Lumbar_Coeliac_Stellate_Sphenopalatine | 0 | 1 | 0 | 0 | 0 | 0 | 0 | 0 |
| TMEM150C | 1.7 | P4_VS_P1_P2_P3_Lumbar_Coeliac_Stellate_Sphenopalatine | 0 | 0 | 0 | 1 | 0 | 0 | 0 | 0 |
| CTNNA1 | 1.7 | P4_VS_P1_P2_P3_Lumbar_Coeliac_Stellate_Sphenopalatine | 0 | 0 | 0 | 1 | 0 | 0 | 0 | 0 |
| ROBO2 | 1.7 | P3_P4_VS_P1_P2_Lumbar_Coeliac_Stellate_Sphenopalatine | 0 | 0 | 1 | 1 | 0 | 0 | 0 | 0 |
| TSPAN33 | 1.7 | P1_P2_P3_P4_Lumbar_Coeliac_VS_Stellate_Sphenopalatine | 1 | 1 | 1 | 1 | 1 | 1 | 0 | 0 |
| PHYHIP | 1.7 | Lumbar_Coeliac_Stellate_VS_P1_P2_P3_P4_Sphenopalatine | 0 | 0 | 0 | 0 | 1 | 1 | 1 | 0 |
| GSG1L | 1.7 | Sphenopalatine_VS_P1_P2_P3_P4_Lumbar_Coeliac_Stellate | 0 | 0 | 0 | 0 | 0 | 0 | 0 | 1 |
| PLXDC2 | 1.7 | P1_Sphenopalatine_VS_P2_P3_P4_Lumbar_Coeliac_Stellate | 1 | 0 | 0 | 0 | 0 | 0 | 0 | 1 |
| PLXNA2 | 1.7 | P2_P4_Sphenopalatine_VS_P1_P3_Lumbar_Coeliac_Stellate | 0 | 1 | 0 | 1 | 0 | 0 | 0 | 1 |
| IGFBP5 | 1.7 | P1_P2_P3_P4_Lumbar_Coeliac_Sphenopalatine_VS_Stellate | 1 | 1 | 1 | 1 | 1 | 1 | 0 | 1 |
| CPNE4 | 1.7 | P1_P2_P3_P4_Lumbar_Coeliac_Sphenopalatine_VS_Stellate | 1 | 1 | 1 | 1 | 1 | 1 | 0 | 1 |
| APOLD1 | 1.69 | P3_VS_P1_P2_P4_Lumbar_Coeliac_Stellate_Sphenopalatine | 0 | 0 | 1 | 0 | 0 | 0 | 0 | 0 |
| GLDN | 1.69 | P3_VS_P1_P2_P4_Lumbar_Coeliac_Stellate_Sphenopalatine | 0 | 0 | 1 | 0 | 0 | 0 | 0 | 0 |
| D830030K20I | 1.69 | P3_VS_P1_P2_P4_Lumbar_Coeliac_Stellate_Sphenopalatine | 0 | 0 | 1 | 0 | 0 | 0 | 0 | 0 |
| CXCL12 | 1.69 | P4_VS_P1_P2_P3_Lumbar_Coeliac_Stellate_Sphenopalatine | 0 | 0 | 0 | 1 | 0 | 0 | 0 | 0 |
| SHB | 1.69 | Lumbar_VS_P1_P2_P3_P4_Coeliac_Stellate_Sphenopalatine | 0 | 0 | 0 | 0 | 1 | 0 | 0 | 0 |
| DBH | 1.69 | P3_Lumbar_Coeliac_Stellate_VS_P1_P2_P4_Sphenopalatine | 0 | 0 | 1 | 0 | 1 | 1 | 1 | 0 |
| ZFP503 | 1.69 | Sphenopalatine_VS_P1_P2_P3_P4_Lumbar_Coeliac_Stellate | 0 | 0 | 0 | 0 | 0 | 0 | 0 | 1 |
| FGF9 | 1.68 | P4_VS_P1_P2_P3_Lumbar_Coeliac_Stellate_Sphenopalatine | 0 | 0 | 0 | 1 | 0 | 0 | 0 | 0 |
| PRRT4 | 1.68 | Lumbar_Coeliac_VS_P1_P2_P3_P4_Stellate_Sphenopalatine | 0 | 0 | 0 | 0 | 1 | 1 | 0 | 0 |
| SPATS2L | 1.68 | P1_P2_P4_Lumbar_Coeliac_Sphenopalatine_VS_P3_Stellate | 1 | 1 | 0 | 1 | 1 | 1 | 0 | 1 |
| TMEM26 | 1.67 | P2_VS_P1_P3_P4_Lumbar_Coeliac_Stellate_Sphenopalatine | 0 | 1 | 0 | 0 | 0 | 0 | 0 | 0 |
| NEXMIF | 1.67 | Lumbar_Coeliac_VS_P1_P2_P3_P4_Stellate_Sphenopalatine | 0 | 0 | 0 | 0 | 1 | 1 | 0 | 0 |
| ZFPM1 | 1.67 | P1_P2_P3_P4_Lumbar_Coeliac_Stellate_VS_Sphenopalatine | 1 | 1 | 1 | 1 | 1 | 1 | 1 | 0 |
| ADAMTS20 | 1.67 | Sphenopalatine_VS_P1_P2_P3_P4_Lumbar_Coeliac_Stellate | 0 | 0 | 0 | 0 | 0 | 0 | 0 | 1 |
| CCKBR | 1.67 | P3_Sphenopalatine_VS_P1_P2_P4_Lumbar_Coeliac_Stellate | 0 | 0 | 1 | 0 | 0 | 0 | 0 | 1 |
| HOMER2 | 1.67 | P1_P2_P4_Sphenopalatine_VS_P3_Lumbar_Coeliac_Stellate | 1 | 1 | 0 | 1 | 0 | 0 | 0 | 1 |
| GAS7 | 1.67 | P1_P2_P4_Sphenopalatine_VS_P3_Lumbar_Coeliac_Stellate | 1 | 1 | 0 | 1 | 0 | 0 | 0 | 1 |
| PEX5L | 1.67 | P1_P2_P3_P4_Lumbar_Coeliac_Sphenopalatine_VS_Stellate | 1 | 1 | 1 | 1 | 1 | 1 | 0 | 1 |

| | | | | | | | | | | | |
|---|---|---|---|---|---|---|---|---|---|---|---|
| EYA1 | 1.67 | P1_P2_P3_Lumbar_Coeliac_Stellate_Sphenopalatine_VS_P4 | 1 | 1 | 1 | 0 | 1 | 1 | 1 | | 1 |
| KCND2 | 1.67 | P1_P3_P4_Lumbar_Coeliac_Stellate_Sphenopalatine_VS_P2 | 1 | 0 | 1 | 1 | 1 | 1 | 1 | | 1 |
| CCND2 | 1.66 | P4_VS_P1_P2_P3_Lumbar_Coeliac_Stellate_Sphenopalatine | 0 | 0 | 0 | 1 | 0 | 0 | 0 | | 0 |
| FFAR3 | 1.66 | Lumbar_Coeliac_VS_P1_P2_P3_P4_Stellate_Sphenopalatine | 0 | 0 | 0 | 0 | 1 | 1 | 0 | | 0 |
| THSD7B | 1.66 | P1_P2_P4_Lumbar_Coeliac_Stellate_VS_P3_Sphenopalatine | 1 | 1 | 0 | 1 | 1 | 1 | 1 | | 0 |
| NPY | 1.66 | P1_P2_P3_P4_Lumbar_Coeliac_Stellate_VS_Sphenopalatine | 1 | 1 | 1 | 1 | 1 | 1 | 1 | | 0 |
| COL12A1 | 1.66 | Sphenopalatine_VS_P1_P2_P3_P4_Lumbar_Coeliac_Stellate | 0 | 0 | 0 | 0 | 0 | 0 | 0 | | 1 |
| DSCAM | 1.66 | P1_P2_P3_P4_Lumbar_Coeliac_Sphenopalatine_VS_Stellate | 1 | 1 | 1 | 1 | 1 | 1 | 0 | | 1 |
| LAYN | 1.66 | P1_P2_P3_Lumbar_Coeliac_Stellate_Sphenopalatine_VS_P4 | 1 | 1 | 1 | 0 | 1 | 1 | 1 | | 1 |
| ZEB2OS | 1.65 | P2_VS_P1_P3_P4_Lumbar_Coeliac_Stellate_Sphenopalatine | 0 | 1 | 0 | 0 | 0 | 0 | 0 | | 0 |
| SDK2 | 1.65 | P1_P2_P4_Lumbar_VS_P3_Coeliac_Stellate_Sphenopalatine | 1 | 1 | 0 | 1 | 1 | 0 | 0 | | 0 |
| ABCC2 | 1.65 | Lumbar_Coeliac_VS_P1_P2_P3_P4_Stellate_Sphenopalatine | 0 | 0 | 0 | 0 | 1 | 1 | 0 | | 0 |
| RORC | 1.65 | P3_Sphenopalatine_VS_P1_P2_P4_Lumbar_Coeliac_Stellate | 0 | 0 | 1 | 0 | 0 | 0 | 0 | | 1 |
| CNTN2 | 1.65 | P1_P2_P3_P4_Lumbar_Coeliac_Sphenopalatine_VS_Stellate | 1 | 1 | 1 | 1 | 1 | 1 | 0 | | 1 |
| HTR2A | 1.64 | P2_VS_P1_P3_P4_Lumbar_Coeliac_Stellate_Sphenopalatine | 0 | 1 | 0 | 0 | 0 | 0 | 0 | | 0 |
| NPY4R | 1.64 | P4_VS_P1_P2_P3_Lumbar_Coeliac_Stellate_Sphenopalatine | 0 | 0 | 0 | 1 | 0 | 0 | 0 | | 0 |
| PPM1J | 1.64 | Lumbar_VS_P1_P2_P3_P4_Coeliac_Stellate_Sphenopalatine | 0 | 0 | 0 | 0 | 1 | 0 | 0 | | 0 |
| C130073E24I | 1.64 | P3_Lumbar_Coeliac_VS_P1_P2_P4_Stellate_Sphenopalatine | 0 | 0 | 1 | 0 | 1 | 1 | 0 | | 0 |
| KCTD16 | 1.64 | P1_P2_P4_Lumbar_Coeliac_Stellate_VS_P3_Sphenopalatine | 1 | 1 | 0 | 1 | 1 | 1 | 1 | | 0 |
| ITGA1 | 1.64 | P3_P4_Lumbar_Coeliac_Stellate_VS_P1_P2_Sphenopalatine | 0 | 0 | 1 | 1 | 1 | 1 | 1 | | 0 |
| SOX5OS2 | 1.64 | P2_P3_P4_Lumbar_Coeliac_Stellate_VS_P1_Sphenopalatine | 0 | 1 | 1 | 1 | 1 | 1 | 1 | | 0 |
| VAV2 | 1.64 | P1_P2_P3_P4_Lumbar_Coeliac_Stellate_VS_Sphenopalatine | 1 | 1 | 1 | 1 | 1 | 1 | 1 | | 0 |
| sept-04 | 1.64 | P1_P2_P3_Lumbar_Coeliac_Stellate_Sphenopalatine_VS_P4 | 1 | 1 | 1 | 0 | 1 | 1 | 1 | | 1 |
| FLRT1 | 1.64 | P1_P2_P4_Lumbar_Coeliac_Stellate_Sphenopalatine_VS_P3 | 1 | 1 | 0 | 1 | 1 | 1 | 1 | | 1 |
| ANGPT2 | 1.63 | P4_VS_P1_P2_P3_Lumbar_Coeliac_Stellate_Sphenopalatine | 0 | 0 | 0 | 1 | 0 | 0 | 0 | | 0 |
| ZFP146 | 1.63 | Lumbar_Coeliac_VS_P1_P2_P3_P4_Stellate_Sphenopalatine | 0 | 0 | 0 | 0 | 1 | 1 | 0 | | 0 |
| CCNJL | 1.63 | P3_Lumbar_Coeliac_VS_P1_P2_P4_Stellate_Sphenopalatine | 0 | 0 | 1 | 0 | 1 | 1 | 0 | | 0 |
| ZIM1 | 1.63 | Sphenopalatine_VS_P1_P2_P3_P4_Lumbar_Coeliac_Stellate | 0 | 0 | 0 | 0 | 0 | 0 | 0 | | 1 |
| ZBTB16 | 1.63 | P4_Sphenopalatine_VS_P1_P2_P3_Lumbar_Coeliac_Stellate | 0 | 0 | 0 | 1 | 0 | 0 | 0 | | 1 |
| HBB-BS | 1.63 | P1_P2_P3_P4_Coeliac_Stellate_Sphenopalatine_VS_Lumbar | 1 | 1 | 1 | 1 | 0 | 1 | 1 | | 1 |

| | | | | | | | | | | |
|---|---|---|---|---|---|---|---|---|---|---|
| PTGER4 | 1.62 | P1_P2_P4_VS_P3_Lumbar_Coeliac_Stellate_Sphenopalatine | 1 | 1 | 0 | 1 | 0 | 0 | 0 | 0 |
| GM3728 | 1.62 | Lumbar_VS_P1_P2_P3_P4_Coeliac_Stellate_Sphenopalatine | 0 | 0 | 0 | 0 | 1 | 0 | 0 | 0 |
| UNC5D | 1.62 | P4_Lumbar_VS_P1_P2_P3_Coeliac_Stellate_Sphenopalatine | 0 | 0 | 0 | 1 | 1 | 0 | 0 | 0 |
| TOX3 | 1.62 | P4_Lumbar_VS_P1_P2_P3_Coeliac_Stellate_Sphenopalatine | 0 | 0 | 0 | 1 | 1 | 0 | 0 | 0 |
| PAG1 | 1.62 | P1_P2_P3_P4_Lumbar_Coeliac_Stellate_VS_Sphenopalatine | 1 | 1 | 1 | 1 | 1 | 1 | 1 | 0 |
| AMER2 | 1.62 | P1_P2_P3_P4_Lumbar_Coeliac_Stellate_VS_Sphenopalatine | 1 | 1 | 1 | 1 | 1 | 1 | 1 | 0 |
| CPNE7 | 1.62 | P1_Sphenopalatine_VS_P2_P3_P4_Lumbar_Coeliac_Stellate | 1 | 0 | 0 | 0 | 0 | 0 | 0 | 1 |
| PTGDS | 1.62 | P3_Sphenopalatine_VS_P1_P2_P4_Lumbar_Coeliac_Stellate | 0 | 0 | 1 | 0 | 0 | 0 | 0 | 1 |
| MFNG | 1.62 | P3_Sphenopalatine_VS_P1_P2_P4_Lumbar_Coeliac_Stellate | 0 | 0 | 1 | 0 | 0 | 0 | 0 | 1 |
| PBX3 | 1.62 | P1_P2_P4_Lumbar_Coeliac_Sphenopalatine_VS_P3_Stellate | 1 | 1 | 0 | 1 | 1 | 1 | 0 | 1 |
| ST3GAL6 | 1.62 | Lumbar_Coeliac_Stellate_Sphenopalatine_VS_P1_P2_P3_P4 | 0 | 0 | 0 | 0 | 1 | 1 | 1 | 1 |
| RASGEF1B | 1.62 | P1_P4_Lumbar_Coeliac_Stellate_Sphenopalatine_VS_P2_P3 | 1 | 0 | 0 | 1 | 1 | 1 | 1 | 1 |
| TM4SF1 | 1.62 | P1_P3_P4_Lumbar_Coeliac_Stellate_Sphenopalatine_VS_P2 | 1 | 0 | 1 | 1 | 1 | 1 | 1 | 1 |
| PRSS12 | 1.61 | P1_VS_P2_P3_P4_Lumbar_Coeliac_Stellate_Sphenopalatine | 1 | 0 | 0 | 0 | 0 | 0 | 0 | 0 |
| PIEZO2 | 1.61 | P1_VS_P2_P3_P4_Lumbar_Coeliac_Stellate_Sphenopalatine | 1 | 0 | 0 | 0 | 0 | 0 | 0 | 0 |
| SNCAIP | 1.61 | P4_VS_P1_P2_P3_Lumbar_Coeliac_Stellate_Sphenopalatine | 0 | 0 | 0 | 1 | 0 | 0 | 0 | 0 |
| GDF5 | 1.61 | P1_P2_P4_VS_P3_Lumbar_Coeliac_Stellate_Sphenopalatine | 1 | 1 | 0 | 1 | 0 | 0 | 0 | 0 |
| CBLN1 | 1.61 | P1_P2_P4_VS_P3_Lumbar_Coeliac_Stellate_Sphenopalatine | 1 | 1 | 0 | 1 | 0 | 0 | 0 | 0 |
| IGSF1 | 1.61 | Lumbar_Coeliac_VS_P1_P2_P3_P4_Stellate_Sphenopalatine | 0 | 0 | 0 | 0 | 1 | 1 | 0 | 0 |
| HMGA1B | 1.61 | P1_P2_P3_P4_Stellate_Sphenopalatine_VS_Lumbar_Coeliac | 1 | 1 | 1 | 1 | 0 | 0 | 1 | 1 |
| 1500009C09I | 1.61 | P1_P2_P3_Lumbar_Coeliac_Stellate_Sphenopalatine_VS_P4 | 1 | 1 | 1 | 0 | 1 | 1 | 1 | 1 |
| LRRTM1 | 1.6 | P4_VS_P1_P2_P3_Lumbar_Coeliac_Stellate_Sphenopalatine | 0 | 0 | 0 | 1 | 0 | 0 | 0 | 0 |
| LRRTM3 | 1.6 | P4_VS_P1_P2_P3_Lumbar_Coeliac_Stellate_Sphenopalatine | 0 | 0 | 0 | 1 | 0 | 0 | 0 | 0 |
| TMEM132E | 1.6 | P1_P2_P3_P4_VS_Lumbar_Coeliac_Stellate_Sphenopalatine | 1 | 1 | 1 | 1 | 0 | 0 | 0 | 0 |
| KLHL11 | 1.6 | P1_P2_P3_P4_Lumbar_Coeliac_VS_Stellate_Sphenopalatine | 1 | 1 | 1 | 1 | 1 | 1 | 0 | 0 |
| GOLM1 | 1.6 | P1_P2_P3_P4_Lumbar_Coeliac_Stellate_VS_Sphenopalatine | 1 | 1 | 1 | 1 | 1 | 1 | 1 | 0 |
| KY | 1.6 | Sphenopalatine_VS_P1_P2_P3_P4_Lumbar_Coeliac_Stellate | 0 | 0 | 0 | 0 | 0 | 0 | 0 | 1 |
| DAZL | 1.59 | P1_P3_VS_P2_P4_Lumbar_Coeliac_Stellate_Sphenopalatine | 1 | 0 | 1 | 0 | 0 | 0 | 0 | 0 |
| TBC1D1 | 1.59 | P4_VS_P1_P2_P3_Lumbar_Coeliac_Stellate_Sphenopalatine | 0 | 0 | 0 | 1 | 0 | 0 | 0 | 0 |
| GM16070 | 1.59 | Lumbar_VS_P1_P2_P3_P4_Coeliac_Stellate_Sphenopalatine | 0 | 0 | 0 | 0 | 1 | 0 | 0 | 0 |

| | | | | | | | | | | |
|---|---|---|---|---|---|---|---|---|---|---|
| KLHL29 | 1.59 | Lumbar_Coeliac_VS_P1_P2_P3_P4_Stellate_Sphenopalatine | 0 | 0 | 0 | 0 | 1 | 1 | 0 | 0 |
| PTGER2 | 1.59 | Sphenopalatine_VS_P1_P2_P3_P4_Lumbar_Coeliac_Stellate | 0 | 0 | 0 | 0 | 0 | 0 | 0 | 1 |
| GFRA1 | 1.59 | P1_Sphenopalatine_VS_P2_P3_P4_Lumbar_Coeliac_Stellate | 1 | 0 | 0 | 0 | 0 | 0 | 0 | 1 |
| KCNA4 | 1.59 | P1_P2_P4_Sphenopalatine_VS_P3_Lumbar_Coeliac_Stellate | 1 | 1 | 0 | 1 | 0 | 0 | 0 | 1 |
| HTR4 | 1.58 | P4_VS_P1_P2_P3_Lumbar_Coeliac_Stellate_Sphenopalatine | 0 | 0 | 0 | 1 | 0 | 0 | 0 | 0 |
| PRELID2 | 1.58 | Lumbar_Coeliac_VS_P1_P2_P3_P4_Stellate_Sphenopalatine | 0 | 0 | 0 | 0 | 1 | 1 | 0 | 0 |
| LHFP | 1.58 | Sphenopalatine_VS_P1_P2_P3_P4_Lumbar_Coeliac_Stellate | 0 | 0 | 0 | 0 | 0 | 0 | 0 | 1 |
| RADX | 1.58 | Sphenopalatine_VS_P1_P2_P3_P4_Lumbar_Coeliac_Stellate | 0 | 0 | 0 | 0 | 0 | 0 | 0 | 1 |
| 9330158H04 | 1.58 | P1_P3_P4_Lumbar_Coeliac_Stellate_Sphenopalatine_VS_P2 | 1 | 0 | 1 | 1 | 1 | 1 | 1 | 1 |
| ACKR3 | 1.57 | P2_VS_P1_P3_P4_Lumbar_Coeliac_Stellate_Sphenopalatine | 0 | 1 | 0 | 0 | 0 | 0 | 0 | 0 |
| CHRM3 | 1.57 | P1_P2_P3_VS_P4_Lumbar_Coeliac_Stellate_Sphenopalatine | 1 | 1 | 1 | 0 | 0 | 0 | 0 | 0 |
| FCMR | 1.57 | P4_VS_P1_P2_P3_Lumbar_Coeliac_Stellate_Sphenopalatine | 0 | 0 | 0 | 1 | 0 | 0 | 0 | 0 |
| GASK1B | 1.57 | P4_VS_P1_P2_P3_Lumbar_Coeliac_Stellate_Sphenopalatine | 0 | 0 | 0 | 1 | 0 | 0 | 0 | 0 |
| GM40383 | 1.57 | P4_VS_P1_P2_P3_Lumbar_Coeliac_Stellate_Sphenopalatine | 0 | 0 | 0 | 1 | 0 | 0 | 0 | 0 |
| RRAGB | 1.57 | Lumbar_Coeliac_VS_P1_P2_P3_P4_Stellate_Sphenopalatine | 0 | 0 | 0 | 0 | 1 | 1 | 0 | 0 |
| CDH8 | 1.57 | P1_Lumbar_Coeliac_VS_P2_P3_P4_Stellate_Sphenopalatine | 1 | 0 | 0 | 0 | 1 | 1 | 0 | 0 |
| MKX | 1.57 | Sphenopalatine_VS_P1_P2_P3_P4_Lumbar_Coeliac_Stellate | 0 | 0 | 0 | 0 | 0 | 0 | 0 | 1 |
| KLF3 | 1.57 | P1_P2_P3_P4_Lumbar_Coeliac_Sphenopalatine_VS_Stellate | 1 | 1 | 1 | 1 | 1 | 1 | 0 | 1 |
| LUZP2 | 1.56 | P4_VS_P1_P2_P3_Lumbar_Coeliac_Stellate_Sphenopalatine | 0 | 0 | 0 | 1 | 0 | 0 | 0 | 0 |
| XKR6 | 1.56 | P1_P2_P3_P4_Lumbar_Coeliac_VS_Stellate_Sphenopalatine | 1 | 1 | 1 | 1 | 1 | 1 | 0 | 0 |
| PHACTR2 | 1.56 | P1_P2_P3_P4_Lumbar_Coeliac_Stellate_VS_Sphenopalatine | 1 | 1 | 1 | 1 | 1 | 1 | 1 | 0 |
| DLK1 | 1.56 | Sphenopalatine_VS_P1_P2_P3_P4_Lumbar_Coeliac_Stellate | 0 | 0 | 0 | 0 | 0 | 0 | 0 | 1 |
| EPB41L4A | 1.56 | P1_P2_P4_Lumbar_Coeliac_Stellate_Sphenopalatine_VS_P3 | 1 | 1 | 0 | 1 | 1 | 1 | 1 | 1 |
| S100B | 1.55 | P4_VS_P1_P2_P3_Lumbar_Coeliac_Stellate_Sphenopalatine | 0 | 0 | 0 | 1 | 0 | 0 | 0 | 0 |
| NTN4 | 1.55 | Lumbar_VS_P1_P2_P3_P4_Coeliac_Stellate_Sphenopalatine | 0 | 0 | 0 | 0 | 1 | 0 | 0 | 0 |
| HIST1H2AE | 1.55 | Coeliac_VS_P1_P2_P3_P4_Lumbar_Stellate_Sphenopalatine | 0 | 0 | 0 | 0 | 0 | 1 | 0 | 0 |
| IFI27L2A | 1.55 | Sphenopalatine_VS_P1_P2_P3_P4_Lumbar_Coeliac_Stellate | 0 | 0 | 0 | 0 | 0 | 0 | 0 | 1 |
| GPR50 | 1.55 | Sphenopalatine_VS_P1_P2_P3_P4_Lumbar_Coeliac_Stellate | 0 | 0 | 0 | 0 | 0 | 0 | 0 | 1 |
| KLF4 | 1.55 | P1_P2_P3_P4_Lumbar_Coeliac_Sphenopalatine_VS_Stellate | 1 | 1 | 1 | 1 | 1 | 1 | 0 | 1 |
| TRIL | 1.55 | P1_P3_P4_Lumbar_Coeliac_Stellate_Sphenopalatine_VS_P2 | 1 | 0 | 1 | 1 | 1 | 1 | 1 | 1 |

| | | | | | | | | | | |
|---|---|---|---|---|---|---|---|---|---|---|
| DACT1 | 1.54 | P1_VS_P2_P3_P4_Lumbar_Coeliac_Stellate_Sphenopalatine | 1 | 0 | 0 | 0 | 0 | 0 | 0 | 0 |
| GM17634 | 1.54 | P4_VS_P1_P2_P3_Lumbar_Coeliac_Stellate_Sphenopalatine | 0 | 0 | 0 | 1 | 0 | 0 | 0 | 0 |
| S100A13 | 1.54 | P1_P2_P3_P4_Lumbar_Coeliac_Stellate_VS_Sphenopalatine | 1 | 1 | 1 | 1 | 1 | 1 | 1 | 0 |
| RBP1 | 1.54 | P1_P4_Sphenopalatine_VS_P2_P3_Lumbar_Coeliac_Stellate | 1 | 0 | 0 | 1 | 0 | 0 | 0 | 1 |
| EDIL3 | 1.54 | P1_P4_Sphenopalatine_VS_P2_P3_Lumbar_Coeliac_Stellate | 1 | 0 | 0 | 1 | 0 | 0 | 0 | 1 |
| ASTN2 | 1.54 | P1_P2_P3_P4_Lumbar_Coeliac_Sphenopalatine_VS_Stellate | 1 | 1 | 1 | 1 | 1 | 1 | 0 | 1 |
| ONECUT2 | 1.54 | P1_P2_P4_Lumbar_Coeliac_Stellate_Sphenopalatine_VS_P3 | 1 | 1 | 0 | 1 | 1 | 1 | 1 | 1 |
| ELMOD1 | 1.54 | P1_P3_P4_Lumbar_Coeliac_Stellate_Sphenopalatine_VS_P2 | 1 | 0 | 1 | 1 | 1 | 1 | 1 | 1 |
| F730043M19 | 1.53 | P2_P3_VS_P1_P4_Lumbar_Coeliac_Stellate_Sphenopalatine | 0 | 1 | 1 | 0 | 0 | 0 | 0 | 0 |
| GM20501 | 1.53 | P4_VS_P1_P2_P3_Lumbar_Coeliac_Stellate_Sphenopalatine | 0 | 0 | 0 | 1 | 0 | 0 | 0 | 0 |
| ADRB3 | 1.53 | P4_VS_P1_P2_P3_Lumbar_Coeliac_Stellate_Sphenopalatine | 0 | 0 | 0 | 1 | 0 | 0 | 0 | 0 |
| SGCZ | 1.53 | Lumbar_Coeliac_VS_P1_P2_P3_P4_Stellate_Sphenopalatine | 0 | 0 | 0 | 0 | 1 | 1 | 0 | 0 |
| CASK | 1.53 | P1_P2_P3_P4_Lumbar_Coeliac_Stellate_VS_Sphenopalatine | 1 | 1 | 1 | 1 | 1 | 1 | 1 | 0 |
| FN1 | 1.53 | Sphenopalatine_VS_P1_P2_P3_P4_Lumbar_Coeliac_Stellate | 0 | 0 | 0 | 0 | 0 | 0 | 0 | 1 |
| ITIH5 | 1.52 | P2_VS_P1_P3_P4_Lumbar_Coeliac_Stellate_Sphenopalatine | 0 | 1 | 0 | 0 | 0 | 0 | 0 | 0 |
| CAR12 | 1.52 | P3_VS_P1_P2_P4_Lumbar_Coeliac_Stellate_Sphenopalatine | 0 | 0 | 1 | 0 | 0 | 0 | 0 | 0 |
| RGCC | 1.52 | P4_VS_P1_P2_P3_Lumbar_Coeliac_Stellate_Sphenopalatine | 0 | 0 | 0 | 1 | 0 | 0 | 0 | 0 |
| H2-EB1 | 1.52 | P4_VS_P1_P2_P3_Lumbar_Coeliac_Stellate_Sphenopalatine | 0 | 0 | 0 | 1 | 0 | 0 | 0 | 0 |
| DDR2 | 1.52 | P1_P2_P4_VS_P3_Lumbar_Coeliac_Stellate_Sphenopalatine | 1 | 1 | 0 | 1 | 0 | 0 | 0 | 0 |
| NLRP6 | 1.52 | Lumbar_Coeliac_VS_P1_P2_P3_P4_Stellate_Sphenopalatine | 0 | 0 | 0 | 0 | 1 | 1 | 0 | 0 |
| FAM43A | 1.52 | P1_P2_P3_P4_Lumbar_Coeliac_Stellate_VS_Sphenopalatine | 1 | 1 | 1 | 1 | 1 | 1 | 1 | 0 |
| ASB2 | 1.52 | Sphenopalatine_VS_P1_P2_P3_P4_Lumbar_Coeliac_Stellate | 0 | 0 | 0 | 0 | 0 | 0 | 0 | 1 |
| PKIB | 1.51 | P2_VS_P1_P3_P4_Lumbar_Coeliac_Stellate_Sphenopalatine | 0 | 1 | 0 | 0 | 0 | 0 | 0 | 0 |
| B230312C02 | 1.51 | P3_VS_P1_P2_P4_Lumbar_Coeliac_Stellate_Sphenopalatine | 0 | 0 | 1 | 0 | 0 | 0 | 0 | 0 |
| HOTAIR | 1.51 | P4_VS_P1_P2_P3_Lumbar_Coeliac_Stellate_Sphenopalatine | 0 | 0 | 0 | 1 | 0 | 0 | 0 | 0 |
| GM29771 | 1.51 | P2_P4_VS_P1_P3_Lumbar_Coeliac_Stellate_Sphenopalatine | 0 | 1 | 0 | 1 | 0 | 0 | 0 | 0 |
| NID1 | 1.51 | P1_P2_P4_VS_P3_Lumbar_Coeliac_Stellate_Sphenopalatine | 1 | 1 | 0 | 1 | 0 | 0 | 0 | 0 |
| WWC1 | 1.51 | Lumbar_Coeliac_VS_P1_P2_P3_P4_Stellate_Sphenopalatine | 0 | 0 | 0 | 0 | 1 | 1 | 0 | 0 |
| TMEM100 | 1.51 | P1_Sphenopalatine_VS_P2_P3_P4_Lumbar_Coeliac_Stellate | 1 | 0 | 0 | 0 | 0 | 0 | 0 | 1 |
| HBB-BT | 1.51 | P1_P2_P3_P4_Sphenopalatine_VS_Lumbar_Coeliac_Stellate | 1 | 1 | 1 | 1 | 0 | 0 | 0 | 1 |

| | | | | | | | | | | |
|---|---|---|---|---|---|---|---|---|---|---|
| HCN2 | 1.51 | P1_P2_P3_P4_Lumbar_Coeliac_Sphenopalatine_VS_Stellate | 1 | 1 | 1 | 1 | 1 | 1 | 0 | 1 |
| TPD52L1 | 1.51 | P3_Lumbar_Coeliac_Stellate_Sphenopalatine_VS_P1_P2_P4 | 0 | 0 | 1 | 0 | 1 | 1 | 1 | 1 |
| PRKG2 | 1.51 | P1_P3_Lumbar_Coeliac_Stellate_Sphenopalatine_VS_P2_P4 | 1 | 0 | 1 | 0 | 1 | 1 | 1 | 1 |
| TMEM200C | 1.5 | P4_VS_P1_P2_P3_Lumbar_Coeliac_Stellate_Sphenopalatine | 0 | 0 | 0 | 1 | 0 | 0 | 0 | 0 |
| ARAP2 | 1.5 | Sphenopalatine_VS_P1_P2_P3_P4_Lumbar_Coeliac_Stellate | 0 | 0 | 0 | 0 | 0 | 0 | 0 | 1 |
| RGS7BP | 1.5 | P1_P3_P4_Lumbar_Coeliac_Stellate_Sphenopalatine_VS_P2 | 1 | 0 | 1 | 1 | 1 | 1 | 1 | 1 |
| CCN3 | 1.5 | P1_P3_P4_Lumbar_Coeliac_Stellate_Sphenopalatine_VS_P2 | 1 | 0 | 1 | 1 | 1 | 1 | 1 | 1 |
| SEMA3D | 1.49 | P3_VS_P1_P2_P4_Lumbar_Coeliac_Stellate_Sphenopalatine | 0 | 0 | 1 | 0 | 0 | 0 | 0 | 0 |
| OOEP | 1.49 | Lumbar_VS_P1_P2_P3_P4_Coeliac_Stellate_Sphenopalatine | 0 | 0 | 0 | 0 | 1 | 0 | 0 | 0 |
| ST8SIA4 | 1.49 | P1_P3_P4_Lumbar_Coeliac_Stellate_VS_P2_Sphenopalatine | 1 | 0 | 1 | 1 | 1 | 1 | 1 | 0 |
| EPHX1 | 1.49 | P1_P2_P3_P4_Lumbar_Coeliac_Stellate_VS_Sphenopalatine | 1 | 1 | 1 | 1 | 1 | 1 | 1 | 0 |
| PPARGC1A | 1.49 | Sphenopalatine_VS_P1_P2_P3_P4_Lumbar_Coeliac_Stellate | 0 | 0 | 0 | 0 | 0 | 0 | 0 | 1 |
| TCF7L2 | 1.49 | Sphenopalatine_VS_P1_P2_P3_P4_Lumbar_Coeliac_Stellate | 0 | 0 | 0 | 0 | 0 | 0 | 0 | 1 |
| PELI2 | 1.49 | P2_Sphenopalatine_VS_P1_P3_P4_Lumbar_Coeliac_Stellate | 0 | 1 | 0 | 0 | 0 | 0 | 0 | 1 |
| GFRA2 | 1.49 | P1_P2_P4_Sphenopalatine_VS_P3_Lumbar_Coeliac_Stellate | 1 | 1 | 0 | 1 | 0 | 0 | 0 | 1 |
| COL14A1 | 1.48 | P4_VS_P1_P2_P3_Lumbar_Coeliac_Stellate_Sphenopalatine | 0 | 0 | 0 | 1 | 0 | 0 | 0 | 0 |
| KCNV1 | 1.48 | P1_P3_P4_VS_P2_Lumbar_Coeliac_Stellate_Sphenopalatine | 1 | 0 | 1 | 1 | 0 | 0 | 0 | 0 |
| TSPAN2OS | 1.48 | Lumbar_Coeliac_Stellate_VS_P1_P2_P3_P4_Sphenopalatine | 0 | 0 | 0 | 0 | 1 | 1 | 1 | 0 |
| AMMECR1 | 1.48 | P1_P2_P3_P4_Lumbar_Coeliac_Stellate_VS_Sphenopalatine | 1 | 1 | 1 | 1 | 1 | 1 | 1 | 0 |
| NFIA | 1.48 | P1_P2_P4_Sphenopalatine_VS_P3_Lumbar_Coeliac_Stellate | 1 | 1 | 0 | 1 | 0 | 0 | 0 | 1 |
| RERG | 1.48 | P1_P2_P3_P4_Sphenopalatine_VS_Lumbar_Coeliac_Stellate | 1 | 1 | 1 | 1 | 0 | 0 | 0 | 1 |
| NPAS3 | 1.48 | Lumbar_Sphenopalatine_VS_P1_P2_P3_P4_Coeliac_Stellate | 0 | 0 | 0 | 0 | 1 | 0 | 0 | 1 |
| AK4 | 1.48 | P1_P2_P3_P4_Lumbar_Coeliac_Sphenopalatine_VS_Stellate | 1 | 1 | 1 | 1 | 1 | 1 | 0 | 1 |
| LARP6 | 1.48 | P1_P2_P3_P4_Lumbar_Coeliac_Sphenopalatine_VS_Stellate | 1 | 1 | 1 | 1 | 1 | 1 | 0 | 1 |
| CHODL | 1.48 | P1_P2_P4_Lumbar_Coeliac_Stellate_Sphenopalatine_VS_P3 | 1 | 1 | 0 | 1 | 1 | 1 | 1 | 1 |
| NOTCH1 | 1.47 | P4_VS_P1_P2_P3_Lumbar_Coeliac_Stellate_Sphenopalatine | 0 | 0 | 0 | 1 | 0 | 0 | 0 | 0 |
| FOXD3 | 1.47 | P4_VS_P1_P2_P3_Lumbar_Coeliac_Stellate_Sphenopalatine | 0 | 0 | 0 | 1 | 0 | 0 | 0 | 0 |
| RADIL | 1.47 | P4_VS_P1_P2_P3_Lumbar_Coeliac_Stellate_Sphenopalatine | 0 | 0 | 0 | 1 | 0 | 0 | 0 | 0 |
| MIR99AHG | 1.47 | P3_P4_VS_P1_P2_Lumbar_Coeliac_Stellate_Sphenopalatine | 0 | 0 | 1 | 1 | 0 | 0 | 0 | 0 |
| LRRC36 | 1.47 | Lumbar_VS_P1_P2_P3_P4_Coeliac_Stellate_Sphenopalatine | 0 | 0 | 0 | 0 | 1 | 0 | 0 | 0 |

| | | | | | | | | | | |
|---|---|---|---|---|---|---|---|---|---|---|
| ADRA2B | 1.47 | Lumbar_Coeliac_Stellate_VS_P1_P2_P3_P4_Sphenopalatine | 0 | 0 | 0 | 0 | 1 | 1 | 1 | 0 |
| SPATA13 | 1.47 | P3_Lumbar_Coeliac_Stellate_VS_P1_P2_P4_Sphenopalatine | 0 | 0 | 1 | 0 | 1 | 1 | 1 | 0 |
| AC109619.1 | 1.47 | P1_P2_P4_Lumbar_Coeliac_Stellate_Sphenopalatine_VS_P3 | 1 | 1 | 0 | 1 | 1 | 1 | 1 | 1 |
| NRP1 | 1.47 | P1_P3_P4_Lumbar_Coeliac_Stellate_Sphenopalatine_VS_P2 | 1 | 0 | 1 | 1 | 1 | 1 | 1 | 1 |
| ALDH1L1 | 1.46 | P2_VS_P1_P3_P4_Lumbar_Coeliac_Stellate_Sphenopalatine | 0 | 1 | 0 | 0 | 0 | 0 | 0 | 0 |
| IRS1 | 1.46 | P3_VS_P1_P2_P4_Lumbar_Coeliac_Stellate_Sphenopalatine | 0 | 0 | 1 | 0 | 0 | 0 | 0 | 0 |
| EPHX2 | 1.46 | P3_VS_P1_P2_P4_Lumbar_Coeliac_Stellate_Sphenopalatine | 0 | 0 | 1 | 0 | 0 | 0 | 0 | 0 |
| CDHR2 | 1.46 | P4_VS_P1_P2_P3_Lumbar_Coeliac_Stellate_Sphenopalatine | 0 | 0 | 0 | 1 | 0 | 0 | 0 | 0 |
| MATN2 | 1.46 | P4_VS_P1_P2_P3_Lumbar_Coeliac_Stellate_Sphenopalatine | 0 | 0 | 0 | 1 | 0 | 0 | 0 | 0 |
| GM5535 | 1.46 | Coeliac_VS_P1_P2_P3_P4_Lumbar_Stellate_Sphenopalatine | 0 | 0 | 0 | 0 | 0 | 1 | 0 | 0 |
| NCK2 | 1.46 | Lumbar_Coeliac_VS_P1_P2_P3_P4_Stellate_Sphenopalatine | 0 | 0 | 0 | 0 | 1 | 1 | 0 | 0 |
| GM38085 | 1.46 | Lumbar_Coeliac_VS_P1_P2_P3_P4_Stellate_Sphenopalatine | 0 | 0 | 0 | 0 | 1 | 1 | 0 | 0 |
| CHD7 | 1.46 | P3_Lumbar_Coeliac_VS_P1_P2_P4_Stellate_Sphenopalatine | 0 | 0 | 1 | 0 | 1 | 1 | 0 | 0 |
| SYNC | 1.46 | P1_P2_P3_P4_Lumbar_Coeliac_VS_Stellate_Sphenopalatine | 1 | 1 | 1 | 1 | 1 | 1 | 0 | 0 |
| PTPN3 | 1.46 | P1_P2_P3_P4_Sphenopalatine_VS_Lumbar_Coeliac_Stellate | 1 | 1 | 1 | 1 | 0 | 0 | 0 | 1 |
| KSR2 | 1.46 | P1_P2_P3_P4_Lumbar_Coeliac_Sphenopalatine_VS_Stellate | 1 | 1 | 1 | 1 | 1 | 1 | 0 | 1 |
| MGAT5B | 1.46 | P1_P2_P3_P4_Lumbar_Coeliac_Sphenopalatine_VS_Stellate | 1 | 1 | 1 | 1 | 1 | 1 | 0 | 1 |
| AREG | 1.45 | P1_VS_P2_P3_P4_Lumbar_Coeliac_Stellate_Sphenopalatine | 1 | 0 | 0 | 0 | 0 | 0 | 0 | 0 |
| SLC43A3 | 1.45 | P4_VS_P1_P2_P3_Lumbar_Coeliac_Stellate_Sphenopalatine | 0 | 0 | 0 | 1 | 0 | 0 | 0 | 0 |
| EDN3 | 1.45 | P4_VS_P1_P2_P3_Lumbar_Coeliac_Stellate_Sphenopalatine | 0 | 0 | 0 | 1 | 0 | 0 | 0 | 0 |
| A330048O09 | 1.45 | P4_VS_P1_P2_P3_Lumbar_Coeliac_Stellate_Sphenopalatine | 0 | 0 | 0 | 1 | 0 | 0 | 0 | 0 |
| LDLRAD4 | 1.45 | P4_VS_P1_P2_P3_Lumbar_Coeliac_Stellate_Sphenopalatine | 0 | 0 | 0 | 1 | 0 | 0 | 0 | 0 |
| SETBP1 | 1.45 | P4_VS_P1_P2_P3_Lumbar_Coeliac_Stellate_Sphenopalatine | 0 | 0 | 0 | 1 | 0 | 0 | 0 | 0 |
| ANKRD44 | 1.45 | P2_P4_VS_P1_P3_Lumbar_Coeliac_Stellate_Sphenopalatine | 0 | 1 | 0 | 1 | 0 | 0 | 0 | 0 |
| GPR179 | 1.45 | P3_Coeliac_VS_P1_P2_P4_Lumbar_Stellate_Sphenopalatine | 0 | 0 | 1 | 0 | 0 | 1 | 0 | 0 |
| PABPC5 | 1.45 | Lumbar_Coeliac_VS_P1_P2_P3_P4_Stellate_Sphenopalatine | 0 | 0 | 0 | 0 | 1 | 1 | 0 | 0 |
| TGFA | 1.45 | P3_Lumbar_Coeliac_Stellate_VS_P1_P2_P4_Sphenopalatine | 0 | 0 | 1 | 0 | 1 | 1 | 1 | 0 |
| GALNT9 | 1.45 | Sphenopalatine_VS_P1_P2_P3_P4_Lumbar_Coeliac_Stellate | 0 | 0 | 0 | 0 | 0 | 0 | 0 | 1 |
| CEP112 | 1.45 | Sphenopalatine_VS_P1_P2_P3_P4_Lumbar_Coeliac_Stellate | 0 | 0 | 0 | 0 | 0 | 0 | 0 | 1 |
| CLSTN3 | 1.45 | P1_P2_P3_P4_Lumbar_Coeliac_Sphenopalatine_VS_Stellate | 1 | 1 | 1 | 1 | 1 | 1 | 0 | 1 |

| | | | | | | | | | | |
|---|---|---|---|---|---|---|---|---|---|---|
| FRRS1 | 1.44 | Lumbar_Coeliac_VS_P1_P2_P3_P4_Stellate_Sphenopalatine | 0 | 0 | 0 | 0 | 1 | 1 | 0 | 0 |
| MT-ND6 | 1.44 | Lumbar_Coeliac_VS_P1_P2_P3_P4_Stellate_Sphenopalatine | 0 | 0 | 0 | 0 | 1 | 1 | 0 | 0 |
| BTBD3 | 1.44 | P1_P2_P3_P4_Lumbar_Coeliac_VS_Stellate_Sphenopalatine | 1 | 1 | 1 | 1 | 1 | 1 | 0 | 0 |
| PTGS1 | 1.44 | P3_Lumbar_Coeliac_Stellate_VS_P1_P2_P4_Sphenopalatine | 0 | 0 | 1 | 0 | 1 | 1 | 1 | 0 |
| MYCN | 1.44 | P3_Lumbar_Coeliac_Stellate_VS_P1_P2_P4_Sphenopalatine | 0 | 0 | 1 | 0 | 1 | 1 | 1 | 0 |
| SCN5A | 1.44 | Sphenopalatine_VS_P1_P2_P3_P4_Lumbar_Coeliac_Stellate | 0 | 0 | 0 | 0 | 0 | 0 | 0 | 1 |
| LY6E | 1.44 | P1_P2_P3_P4_Coeliac_Sphenopalatine_VS_Lumbar_Stellate | 1 | 1 | 1 | 1 | 0 | 1 | 0 | 1 |
| CACNA1C | 1.44 | P1_P2_P3_P4_Lumbar_Coeliac_Sphenopalatine_VS_Stellate | 1 | 1 | 1 | 1 | 1 | 1 | 0 | 1 |
| GM30003 | 1.43 | P2_VS_P1_P3_P4_Lumbar_Coeliac_Stellate_Sphenopalatine | 0 | 1 | 0 | 0 | 0 | 0 | 0 | 0 |
| RBM8A2 | 1.43 | P4_VS_P1_P2_P3_Lumbar_Coeliac_Stellate_Sphenopalatine | 0 | 0 | 0 | 1 | 0 | 0 | 0 | 0 |
| 6530402F18I | 1.43 | P4_VS_P1_P2_P3_Lumbar_Coeliac_Stellate_Sphenopalatine | 0 | 0 | 0 | 1 | 0 | 0 | 0 | 0 |
| METRN | 1.43 | P4_VS_P1_P2_P3_Lumbar_Coeliac_Stellate_Sphenopalatine | 0 | 0 | 0 | 1 | 0 | 0 | 0 | 0 |
| PTGFRN | 1.43 | P2_P4_VS_P1_P3_Lumbar_Coeliac_Stellate_Sphenopalatine | 0 | 1 | 0 | 1 | 0 | 0 | 0 | 0 |
| CYP2S1 | 1.43 | Sphenopalatine_VS_P1_P2_P3_P4_Lumbar_Coeliac_Stellate | 0 | 0 | 0 | 0 | 0 | 0 | 0 | 1 |
| LYNX1 | 1.43 | P1_P2_Sphenopalatine_VS_P3_P4_Lumbar_Coeliac_Stellate | 1 | 1 | 0 | 0 | 0 | 0 | 0 | 1 |
| SELENOV | 1.43 | P1_P2_P3_P4_Sphenopalatine_VS_Lumbar_Coeliac_Stellate | 1 | 1 | 1 | 1 | 0 | 0 | 0 | 1 |
| MAGI3 | 1.43 | P1_P2_P3_P4_Lumbar_Coeliac_Sphenopalatine_VS_Stellate | 1 | 1 | 1 | 1 | 1 | 1 | 0 | 1 |
| ANO8 | 1.43 | P1_P2_P3_P4_Lumbar_Coeliac_Sphenopalatine_VS_Stellate | 1 | 1 | 1 | 1 | 1 | 1 | 0 | 1 |
| ADCY5 | 1.43 | P1_P2_P3_P4_Lumbar_Coeliac_Sphenopalatine_VS_Stellate | 1 | 1 | 1 | 1 | 1 | 1 | 0 | 1 |
| NAV2 | 1.42 | P2_VS_P1_P3_P4_Lumbar_Coeliac_Stellate_Sphenopalatine | 0 | 1 | 0 | 0 | 0 | 0 | 0 | 0 |
| A330084C13I | 1.42 | P1_P2_VS_P3_P4_Lumbar_Coeliac_Stellate_Sphenopalatine | 1 | 1 | 0 | 0 | 0 | 0 | 0 | 0 |
| AI593442 | 1.42 | P4_VS_P1_P2_P3_Lumbar_Coeliac_Stellate_Sphenopalatine | 0 | 0 | 0 | 1 | 0 | 0 | 0 | 0 |
| ERBB3 | 1.42 | P4_VS_P1_P2_P3_Lumbar_Coeliac_Stellate_Sphenopalatine | 0 | 0 | 0 | 1 | 0 | 0 | 0 | 0 |
| VWA7 | 1.42 | P4_VS_P1_P2_P3_Lumbar_Coeliac_Stellate_Sphenopalatine | 0 | 0 | 0 | 1 | 0 | 0 | 0 | 0 |
| SPON1 | 1.42 | Sphenopalatine_VS_P1_P2_P3_P4_Lumbar_Coeliac_Stellate | 0 | 0 | 0 | 0 | 0 | 0 | 0 | 1 |
| KCND3 | 1.41 | P2_VS_P1_P3_P4_Lumbar_Coeliac_Stellate_Sphenopalatine | 0 | 1 | 0 | 0 | 0 | 0 | 0 | 0 |
| SLC35F1 | 1.41 | P3_VS_P1_P2_P4_Lumbar_Coeliac_Stellate_Sphenopalatine | 0 | 0 | 1 | 0 | 0 | 0 | 0 | 0 |
| AEBP1 | 1.41 | P3_VS_P1_P2_P4_Lumbar_Coeliac_Stellate_Sphenopalatine | 0 | 0 | 1 | 0 | 0 | 0 | 0 | 0 |
| DIAPH3 | 1.41 | P3_VS_P1_P2_P4_Lumbar_Coeliac_Stellate_Sphenopalatine | 0 | 0 | 1 | 0 | 0 | 0 | 0 | 0 |
| EFHD1 | 1.41 | P4_VS_P1_P2_P3_Lumbar_Coeliac_Stellate_Sphenopalatine | 0 | 0 | 0 | 1 | 0 | 0 | 0 | 0 |

| | | | | | | | | | | |
|---|---|---|---|---|---|---|---|---|---|---|
| TNR | 1.41 | P1_P2_P3_P4_VS_Lumbar_Coeliac_Stellate_Sphenopalatine | 1 | 1 | 1 | 1 | 0 | 0 | 0 | 0 |
| GM5136 | 1.41 | P1_P2_P3_P4_Lumbar_Coeliac_Stellate_VS_Sphenopalatine | 1 | 1 | 1 | 1 | 1 | 1 | 1 | 0 |
| NELL2 | 1.41 | P1_Sphenopalatine_VS_P2_P3_P4_Lumbar_Coeliac_Stellate | 1 | 0 | 0 | 0 | 0 | 0 | 0 | 1 |
| PHLDA1 | 1.41 | P1_P2_P3_P4_Sphenopalatine_VS_Lumbar_Coeliac_Stellate | 1 | 1 | 1 | 1 | 0 | 0 | 0 | 1 |
| PDPN | 1.4 | P1_VS_P2_P3_P4_Lumbar_Coeliac_Stellate_Sphenopalatine | 1 | 0 | 0 | 0 | 0 | 0 | 0 | 0 |
| GM48992 | 1.4 | P1_VS_P2_P3_P4_Lumbar_Coeliac_Stellate_Sphenopalatine | 1 | 0 | 0 | 0 | 0 | 0 | 0 | 0 |
| PHF21B | 1.4 | P1_VS_P2_P3_P4_Lumbar_Coeliac_Stellate_Sphenopalatine | 1 | 0 | 0 | 0 | 0 | 0 | 0 | 0 |
| ARHGAP29 | 1.4 | P1_P2_VS_P3_P4_Lumbar_Coeliac_Stellate_Sphenopalatine | 1 | 1 | 0 | 0 | 0 | 0 | 0 | 0 |
| FRMPD3 | 1.4 | P4_VS_P1_P2_P3_Lumbar_Coeliac_Stellate_Sphenopalatine | 0 | 0 | 0 | 1 | 0 | 0 | 0 | 0 |
| C230038L03I | 1.4 | Lumbar_VS_P1_P2_P3_P4_Coeliac_Stellate_Sphenopalatine | 0 | 0 | 0 | 0 | 1 | 0 | 0 | 0 |
| STRA6 | 1.4 | P3_Lumbar_Coeliac_VS_P1_P2_P4_Stellate_Sphenopalatine | 0 | 0 | 1 | 0 | 1 | 1 | 0 | 0 |
| FAM177A | 1.4 | P4_Lumbar_Coeliac_VS_P1_P2_P3_Stellate_Sphenopalatine | 0 | 0 | 0 | 1 | 1 | 1 | 0 | 0 |
| ARNT2 | 1.4 | P1_P2_P3_P4_Lumbar_Coeliac_Stellate_VS_Sphenopalatine | 1 | 1 | 1 | 1 | 1 | 1 | 1 | 0 |
| TSPAN2 | 1.4 | P1_P2_P3_P4_Lumbar_Coeliac_Sphenopalatine_VS_Stellate | 1 | 1 | 1 | 1 | 1 | 1 | 0 | 1 |
| TMC3 | 1.4 | P1_P3_P4_Lumbar_Coeliac_Stellate_Sphenopalatine_VS_P2 | 1 | 0 | 1 | 1 | 1 | 1 | 1 | 1 |
| MAP3K20 | 1.39 | P4_VS_P1_P2_P3_Lumbar_Coeliac_Stellate_Sphenopalatine | 0 | 0 | 0 | 1 | 0 | 0 | 0 | 0 |
| PLATR14 | 1.39 | P4_VS_P1_P2_P3_Lumbar_Coeliac_Stellate_Sphenopalatine | 0 | 0 | 0 | 1 | 0 | 0 | 0 | 0 |
| ELFN1 | 1.39 | P2_P3_P4_VS_P1_Lumbar_Coeliac_Stellate_Sphenopalatine | 0 | 1 | 1 | 1 | 0 | 0 | 0 | 0 |
| ACAD10 | 1.39 | Coeliac_VS_P1_P2_P3_P4_Lumbar_Stellate_Sphenopalatine | 0 | 0 | 0 | 0 | 0 | 1 | 0 | 0 |
| ZFP384 | 1.39 | Lumbar_Coeliac_VS_P1_P2_P3_P4_Stellate_Sphenopalatine | 0 | 0 | 0 | 0 | 1 | 1 | 0 | 0 |
| ZFP609 | 1.39 | Lumbar_Coeliac_VS_P1_P2_P3_P4_Stellate_Sphenopalatine | 0 | 0 | 0 | 0 | 1 | 1 | 0 | 0 |
| A630089N07 | 1.39 | Lumbar_Coeliac_VS_P1_P2_P3_P4_Stellate_Sphenopalatine | 0 | 0 | 0 | 0 | 1 | 1 | 0 | 0 |
| PLK2 | 1.39 | P1_P3_Lumbar_Coeliac_Stellate_VS_P2_P4_Sphenopalatine | 1 | 0 | 1 | 0 | 1 | 1 | 1 | 0 |
| MCTP1 | 1.39 | P1_P2_P3_P4_Lumbar_Coeliac_Stellate_VS_Sphenopalatine | 1 | 1 | 1 | 1 | 1 | 1 | 1 | 0 |
| NECAB1 | 1.39 | P1_P2_P3_P4_Lumbar_Coeliac_Sphenopalatine_VS_Stellate | 1 | 1 | 1 | 1 | 1 | 1 | 0 | 1 |
| WTIP | 1.39 | P1_P2_P3_P4_Lumbar_Coeliac_Sphenopalatine_VS_Stellate | 1 | 1 | 1 | 1 | 1 | 1 | 0 | 1 |
| RGS3 | 1.39 | P1_P2_P3_Lumbar_Coeliac_Stellate_Sphenopalatine_VS_P4 | 1 | 1 | 1 | 0 | 1 | 1 | 1 | 1 |
| PTPRU | 1.38 | P4_VS_P1_P2_P3_Lumbar_Coeliac_Stellate_Sphenopalatine | 0 | 0 | 0 | 1 | 0 | 0 | 0 | 0 |
| A930012L18I | 1.38 | P1_P2_P4_VS_P3_Lumbar_Coeliac_Stellate_Sphenopalatine | 1 | 1 | 0 | 1 | 0 | 0 | 0 | 0 |
| ZFP955A | 1.38 | Lumbar_Coeliac_VS_P1_P2_P3_P4_Stellate_Sphenopalatine | 0 | 0 | 0 | 0 | 1 | 1 | 0 | 0 |

| | | | | | | | | | | |
|---|---|---|---|---|---|---|---|---|---|---|
| OTUD7A | 1.38 | P1_P2_P3_P4_Lumbar_Coeliac_VS_Stellate_Sphenopalatine | 1 | 1 | 1 | 1 | 1 | 1 | 0 | 0 |
| RAPSN | 1.38 | Lumbar_Coeliac_Stellate_VS_P1_P2_P3_P4_Sphenopalatine | 0 | 0 | 0 | 0 | 1 | 1 | 1 | 0 |
| GM15688 | 1.38 | Sphenopalatine_VS_P1_P2_P3_P4_Lumbar_Coeliac_Stellate | 0 | 0 | 0 | 0 | 0 | 0 | 0 | 1 |
| GRIK2 | 1.38 | P1_P2_P3_P4_Lumbar_Coeliac_Sphenopalatine_VS_Stellate | 1 | 1 | 1 | 1 | 1 | 1 | 0 | 1 |
| ADAMTS5 | 1.38 | P1_P2_P3_P4_Coeliac_Stellate_Sphenopalatine_VS_Lumbar | 1 | 1 | 1 | 1 | 0 | 1 | 1 | 1 |
| CPNE5 | 1.38 | P1_P2_P3_Lumbar_Coeliac_Stellate_Sphenopalatine_VS_P4 | 1 | 1 | 1 | 0 | 1 | 1 | 1 | 1 |
| NKD1 | 1.37 | P2_VS_P1_P3_P4_Lumbar_Coeliac_Stellate_Sphenopalatine | 0 | 1 | 0 | 0 | 0 | 0 | 0 | 0 |
| UNC5B | 1.37 | P1_P2_P3_Lumbar_Coeliac_VS_P4_Stellate_Sphenopalatine | 1 | 1 | 1 | 0 | 1 | 1 | 0 | 0 |
| SEMA3C | 1.37 | P1_P2_P3_P4_Lumbar_Coeliac_Stellate_VS_Sphenopalatine | 1 | 1 | 1 | 1 | 1 | 1 | 1 | 0 |
| P2RX2 | 1.37 | P1_P2_P3_P4_Sphenopalatine_VS_Lumbar_Coeliac_Stellate | 1 | 1 | 1 | 1 | 0 | 0 | 0 | 1 |
| TBX3OS1 | 1.37 | P1_P2_P3_Coeliac_Sphenopalatine_VS_P4_Lumbar_Stellate | 1 | 1 | 1 | 0 | 0 | 1 | 0 | 1 |
| SLC4A4 | 1.37 | P1_P2_Lumbar_Coeliac_Sphenopalatine_VS_P3_P4_Stellate | 1 | 1 | 0 | 0 | 1 | 1 | 0 | 1 |
| HFM1 | 1.36 | P4_VS_P1_P2_P3_Lumbar_Coeliac_Stellate_Sphenopalatine | 0 | 0 | 0 | 1 | 0 | 0 | 0 | 0 |
| APH1C | 1.36 | P4_VS_P1_P2_P3_Lumbar_Coeliac_Stellate_Sphenopalatine | 0 | 0 | 0 | 1 | 0 | 0 | 0 | 0 |
| SLITRK5 | 1.36 | P1_P4_VS_P2_P3_Lumbar_Coeliac_Stellate_Sphenopalatine | 1 | 0 | 0 | 1 | 0 | 0 | 0 | 0 |
| ATP8B2 | 1.36 | Lumbar_Coeliac_VS_P1_P2_P3_P4_Stellate_Sphenopalatine | 0 | 0 | 0 | 0 | 1 | 1 | 0 | 0 |
| GPR68 | 1.36 | P3_Lumbar_Coeliac_VS_P1_P2_P4_Stellate_Sphenopalatine | 0 | 0 | 1 | 0 | 1 | 1 | 0 | 0 |
| LZTS3 | 1.36 | P1_P2_P3_P4_Lumbar_Coeliac_VS_Stellate_Sphenopalatine | 1 | 1 | 1 | 1 | 1 | 1 | 0 | 0 |
| A230006K03 | 1.36 | P1_P2_P4_Lumbar_Coeliac_Sphenopalatine_VS_P3_Stellate | 1 | 1 | 0 | 1 | 1 | 1 | 0 | 1 |
| CACNA2D3 | 1.36 | P2_Lumbar_Coeliac_Stellate_Sphenopalatine_VS_P1_P3_P4 | 0 | 1 | 0 | 0 | 1 | 1 | 1 | 1 |
| TMEM44 | 1.36 | P1_P2_P3_Lumbar_Coeliac_Stellate_Sphenopalatine_VS_P4 | 1 | 1 | 1 | 0 | 1 | 1 | 1 | 1 |
| LRAT | 1.35 | P4_VS_P1_P2_P3_Lumbar_Coeliac_Stellate_Sphenopalatine | 0 | 0 | 0 | 1 | 0 | 0 | 0 | 0 |
| STON2 | 1.35 | P4_VS_P1_P2_P3_Lumbar_Coeliac_Stellate_Sphenopalatine | 0 | 0 | 0 | 1 | 0 | 0 | 0 | 0 |
| CSMD3 | 1.35 | P4_VS_P1_P2_P3_Lumbar_Coeliac_Stellate_Sphenopalatine | 0 | 0 | 0 | 1 | 0 | 0 | 0 | 0 |
| TMEM255A | 1.35 | P4_VS_P1_P2_P3_Lumbar_Coeliac_Stellate_Sphenopalatine | 0 | 0 | 0 | 1 | 0 | 0 | 0 | 0 |
| CHRNA7 | 1.35 | Coeliac_VS_P1_P2_P3_P4_Lumbar_Stellate_Sphenopalatine | 0 | 0 | 0 | 0 | 0 | 1 | 0 | 0 |
| DPY19L1 | 1.35 | P1_P2_P3_P4_Lumbar_Coeliac_Stellate_VS_Sphenopalatine | 1 | 1 | 1 | 1 | 1 | 1 | 1 | 0 |
| CMBL | 1.35 | P1_P2_P3_P4_Lumbar_Coeliac_Stellate_VS_Sphenopalatine | 1 | 1 | 1 | 1 | 1 | 1 | 1 | 0 |
| IGF2BP2 | 1.35 | P1_P2_P3_P4_Sphenopalatine_VS_Lumbar_Coeliac_Stellate | 1 | 1 | 1 | 1 | 0 | 0 | 0 | 1 |
| FOSB | 1.35 | P1_P2_P3_P4_Lumbar_Coeliac_Sphenopalatine_VS_Stellate | 1 | 1 | 1 | 1 | 1 | 1 | 0 | 1 |

| | | | | | | | | | | |
|---|---|---|---|---|---|---|---|---|---|---|
| TPBGL | 1.35 | P1_P2_P3_P4_Lumbar_Stellate_Sphenopalatine_VS_Coeliac | 1 | 1 | 1 | 1 | 1 | 0 | 1 | 1 |
| GM43445 | 1.34 | P1_VS_P2_P3_P4_Lumbar_Coeliac_Stellate_Sphenopalatine | 1 | 0 | 0 | 0 | 0 | 0 | 0 | 0 |
| AR | 1.34 | P3_VS_P1_P2_P4_Lumbar_Coeliac_Stellate_Sphenopalatine | 0 | 0 | 1 | 0 | 0 | 0 | 0 | 0 |
| IGFBP7 | 1.34 | P1_P2_P3_VS_P4_Lumbar_Coeliac_Stellate_Sphenopalatine | 1 | 1 | 1 | 0 | 0 | 0 | 0 | 0 |
| TRIM16 | 1.34 | P4_VS_P1_P2_P3_Lumbar_Coeliac_Stellate_Sphenopalatine | 0 | 0 | 0 | 1 | 0 | 0 | 0 | 0 |
| ZCCHC24 | 1.34 | P4_VS_P1_P2_P3_Lumbar_Coeliac_Stellate_Sphenopalatine | 0 | 0 | 0 | 1 | 0 | 0 | 0 | 0 |
| TIAM2 | 1.34 | P2_P4_VS_P1_P3_Lumbar_Coeliac_Stellate_Sphenopalatine | 0 | 1 | 0 | 1 | 0 | 0 | 0 | 0 |
| ZFP84 | 1.34 | Lumbar_Coeliac_VS_P1_P2_P3_P4_Stellate_Sphenopalatine | 0 | 0 | 0 | 0 | 1 | 1 | 0 | 0 |
| GALNT6 | 1.34 | Lumbar_Coeliac_VS_P1_P2_P3_P4_Stellate_Sphenopalatine | 0 | 0 | 0 | 0 | 1 | 1 | 0 | 0 |
| VWC2L | 1.34 | P4_Lumbar_Coeliac_VS_P1_P2_P3_Stellate_Sphenopalatine | 0 | 0 | 0 | 1 | 1 | 1 | 0 | 0 |
| BRINP2 | 1.34 | P3_Lumbar_Coeliac_Stellate_VS_P1_P2_P4_Sphenopalatine | 0 | 0 | 1 | 0 | 1 | 1 | 1 | 0 |
| ENOX2 | 1.33 | P4_VS_P1_P2_P3_Lumbar_Coeliac_Stellate_Sphenopalatine | 0 | 0 | 0 | 1 | 0 | 0 | 0 | 0 |
| PLCB1 | 1.33 | Sphenopalatine_VS_P1_P2_P3_P4_Lumbar_Coeliac_Stellate | 0 | 0 | 0 | 0 | 0 | 0 | 0 | 1 |
| INSYN2B | 1.33 | P1_Sphenopalatine_VS_P2_P3_P4_Lumbar_Coeliac_Stellate | 1 | 0 | 0 | 0 | 0 | 0 | 0 | 1 |
| IER5L | 1.33 | P1_P2_P3_P4_Lumbar_Coeliac_Sphenopalatine_VS_Stellate | 1 | 1 | 1 | 1 | 1 | 1 | 0 | 1 |
| 9130008F23I | 1.32 | P4_VS_P1_P2_P3_Lumbar_Coeliac_Stellate_Sphenopalatine | 0 | 0 | 0 | 1 | 0 | 0 | 0 | 0 |
| DLK2 | 1.32 | Lumbar_VS_P1_P2_P3_P4_Coeliac_Stellate_Sphenopalatine | 0 | 0 | 0 | 0 | 1 | 0 | 0 | 0 |
| TSHR | 1.32 | P1_Lumbar_VS_P2_P3_P4_Coeliac_Stellate_Sphenopalatine | 1 | 0 | 0 | 0 | 1 | 0 | 0 | 0 |
| GM37529 | 1.32 | Lumbar_Coeliac_VS_P1_P2_P3_P4_Stellate_Sphenopalatine | 0 | 0 | 0 | 0 | 1 | 1 | 0 | 0 |
| GPC2 | 1.32 | Lumbar_Coeliac_VS_P1_P2_P3_P4_Stellate_Sphenopalatine | 0 | 0 | 0 | 0 | 1 | 1 | 0 | 0 |
| CHSY3 | 1.32 | Lumbar_Coeliac_VS_P1_P2_P3_P4_Stellate_Sphenopalatine | 0 | 0 | 0 | 0 | 1 | 1 | 0 | 0 |
| JHY | 1.32 | P1_P3_P4_Lumbar_Coeliac_VS_P2_Stellate_Sphenopalatine | 1 | 0 | 1 | 1 | 1 | 1 | 0 | 0 |
| 1700011L22I | 1.32 | P1_P2_P3_P4_Lumbar_Coeliac_VS_Stellate_Sphenopalatine | 1 | 1 | 1 | 1 | 1 | 1 | 0 | 0 |
| SMTNL2 | 1.32 | P3_P4_Lumbar_Coeliac_Stellate_VS_P1_P2_Sphenopalatine | 0 | 0 | 1 | 1 | 1 | 1 | 1 | 0 |
| VASH2 | 1.32 | P1_P3_Lumbar_Coeliac_Stellate_Sphenopalatine_VS_P2_P4 | 1 | 0 | 1 | 0 | 1 | 1 | 1 | 1 |
| ZC3HAV1 | 1.32 | P1_P2_P4_Lumbar_Coeliac_Stellate_Sphenopalatine_VS_P3 | 1 | 1 | 0 | 1 | 1 | 1 | 1 | 1 |
| CYP1B1 | 1.31 | P1_VS_P2_P3_P4_Lumbar_Coeliac_Stellate_Sphenopalatine | 1 | 0 | 0 | 0 | 0 | 0 | 0 | 0 |
| NR4A3 | 1.31 | P2_VS_P1_P3_P4_Lumbar_Coeliac_Stellate_Sphenopalatine | 0 | 1 | 0 | 0 | 0 | 0 | 0 | 0 |
| FNDC5 | 1.31 | P2_VS_P1_P3_P4_Lumbar_Coeliac_Stellate_Sphenopalatine | 0 | 1 | 0 | 0 | 0 | 0 | 0 | 0 |
| ADAM12 | 1.31 | P2_VS_P1_P3_P4_Lumbar_Coeliac_Stellate_Sphenopalatine | 0 | 1 | 0 | 0 | 0 | 0 | 0 | 0 |

| | | | | | | | | | | |
|---|---|---|---|---|---|---|---|---|---|---|
| ARHGAP31 | 1.31 | P2_VS_P1_P3_P4_Lumbar_Coeliac_Stellate_Sphenopalatine | 0 | 1 | 0 | 0 | 0 | 0 | 0 | 0 |
| SERPINE2 | 1.31 | P1_P2_P4_VS_P3_Lumbar_Coeliac_Stellate_Sphenopalatine | 1 | 1 | 0 | 1 | 0 | 0 | 0 | 0 |
| FGGY | 1.31 | Lumbar_Coeliac_VS_P1_P2_P3_P4_Stellate_Sphenopalatine | 0 | 0 | 0 | 0 | 1 | 1 | 0 | 0 |
| GNB3 | 1.31 | P3_Lumbar_Coeliac_Stellate_VS_P1_P2_P4_Sphenopalatine | 0 | 0 | 1 | 0 | 1 | 1 | 1 | 0 |
| ACVR2B | 1.31 | P1_P2_P3_P4_Lumbar_Coeliac_Sphenopalatine_VS_Stellate | 1 | 1 | 1 | 1 | 1 | 1 | 0 | 1 |
| ZBTB7C | 1.31 | P1_P2_P3_P4_Lumbar_Coeliac_Sphenopalatine_VS_Stellate | 1 | 1 | 1 | 1 | 1 | 1 | 0 | 1 |
| FAM183B | 1.3 | P4_VS_P1_P2_P3_Lumbar_Coeliac_Stellate_Sphenopalatine | 0 | 0 | 0 | 1 | 0 | 0 | 0 | 0 |
| SINHCAF | 1.3 | Lumbar_Coeliac_VS_P1_P2_P3_P4_Stellate_Sphenopalatine | 0 | 0 | 0 | 0 | 1 | 1 | 0 | 0 |
| LMNB1 | 1.3 | Lumbar_Coeliac_VS_P1_P2_P3_P4_Stellate_Sphenopalatine | 0 | 0 | 0 | 0 | 1 | 1 | 0 | 0 |
| FRMPD4 | 1.3 | P2_P3_P4_Lumbar_Coeliac_VS_P1_Stellate_Sphenopalatine | 0 | 1 | 1 | 1 | 1 | 1 | 0 | 0 |
| TGFB1 | 1.3 | P2_P3_Lumbar_Coeliac_Stellate_VS_P1_P4_Sphenopalatine | 0 | 1 | 1 | 0 | 1 | 1 | 1 | 0 |
| PRAG1 | 1.3 | Sphenopalatine_VS_P1_P2_P3_P4_Lumbar_Coeliac_Stellate | 0 | 0 | 0 | 0 | 0 | 0 | 0 | 1 |
| MAPKAPK2 | 1.3 | P1_P2_P3_P4_Lumbar_Coeliac_Sphenopalatine_VS_Stellate | 1 | 1 | 1 | 1 | 1 | 1 | 0 | 1 |
| CREB5 | 1.3 | P1_P2_P3_P4_Lumbar_Coeliac_Sphenopalatine_VS_Stellate | 1 | 1 | 1 | 1 | 1 | 1 | 0 | 1 |
| DAPK3 | 1.3 | P1_P2_P3_P4_Lumbar_Coeliac_Sphenopalatine_VS_Stellate | 1 | 1 | 1 | 1 | 1 | 1 | 0 | 1 |
| OTULIN | 1.3 | P1_P2_P3_P4_Lumbar_Coeliac_Sphenopalatine_VS_Stellate | 1 | 1 | 1 | 1 | 1 | 1 | 0 | 1 |
| ADRA1B | 1.29 | P2_VS_P1_P3_P4_Lumbar_Coeliac_Stellate_Sphenopalatine | 0 | 1 | 0 | 0 | 0 | 0 | 0 | 0 |
| GALM | 1.29 | P2_VS_P1_P3_P4_Lumbar_Coeliac_Stellate_Sphenopalatine | 0 | 1 | 0 | 0 | 0 | 0 | 0 | 0 |
| PKDCC | 1.29 | P2_VS_P1_P3_P4_Lumbar_Coeliac_Stellate_Sphenopalatine | 0 | 1 | 0 | 0 | 0 | 0 | 0 | 0 |
| MATN4 | 1.29 | P4_VS_P1_P2_P3_Lumbar_Coeliac_Stellate_Sphenopalatine | 0 | 0 | 0 | 1 | 0 | 0 | 0 | 0 |
| FOXO1 | 1.29 | P2_P4_VS_P1_P3_Lumbar_Coeliac_Stellate_Sphenopalatine | 0 | 1 | 0 | 1 | 0 | 0 | 0 | 0 |
| SLC4A10 | 1.29 | P3_P4_VS_P1_P2_Lumbar_Coeliac_Stellate_Sphenopalatine | 0 | 0 | 1 | 1 | 0 | 0 | 0 | 0 |
| HPDL | 1.29 | Lumbar_VS_P1_P2_P3_P4_Coeliac_Stellate_Sphenopalatine | 0 | 0 | 0 | 0 | 1 | 0 | 0 | 0 |
| ZFP605 | 1.29 | Lumbar_Coeliac_VS_P1_P2_P3_P4_Stellate_Sphenopalatine | 0 | 0 | 0 | 0 | 1 | 1 | 0 | 0 |
| ADGRG6 | 1.29 | P1_P4_Sphenopalatine_VS_P2_P3_Lumbar_Coeliac_Stellate | 1 | 0 | 0 | 1 | 0 | 0 | 0 | 1 |
| GM12122 | 1.29 | P1_P2_P3_P4_Lumbar_Coeliac_Sphenopalatine_VS_Stellate | 1 | 1 | 1 | 1 | 1 | 1 | 0 | 1 |
| GM14964 | 1.29 | P1_P2_P3_Lumbar_Coeliac_Stellate_Sphenopalatine_VS_P4 | 1 | 1 | 1 | 0 | 1 | 1 | 1 | 1 |
| EPS8 | 1.28 | P4_VS_P1_P2_P3_Lumbar_Coeliac_Stellate_Sphenopalatine | 0 | 0 | 0 | 1 | 0 | 0 | 0 | 0 |
| EDNRA | 1.28 | P4_VS_P1_P2_P3_Lumbar_Coeliac_Stellate_Sphenopalatine | 0 | 0 | 0 | 1 | 0 | 0 | 0 | 0 |
| ALDH3B1 | 1.28 | P3_P4_VS_P1_P2_Lumbar_Coeliac_Stellate_Sphenopalatine | 0 | 0 | 1 | 1 | 0 | 0 | 0 | 0 |

| | | | | | | | | | | |
|---|---|---|---|---|---|---|---|---|---|---|
| DEXI | 1.28 | P1_P2_P3_P4_Lumbar_Coeliac_VS_Stellate_Sphenopalatine | 1 | 1 | 1 | 1 | 1 | 1 | 0 | 0 |
| RASGRP2 | 1.28 | Sphenopalatine_VS_P1_P2_P3_P4_Lumbar_Coeliac_Stellate | 0 | 0 | 0 | 0 | 0 | 0 | 0 | 1 |
| VEGFA | 1.28 | P1_P2_P3_P4_Lumbar_Coeliac_Sphenopalatine_VS_Stellate | 1 | 1 | 1 | 1 | 1 | 1 | 0 | 1 |
| ATP2B3 | 1.28 | P1_P2_P3_P4_Lumbar_Coeliac_Sphenopalatine_VS_Stellate | 1 | 1 | 1 | 1 | 1 | 1 | 0 | 1 |
| FXYD5 | 1.28 | P1_P3_Lumbar_Coeliac_Stellate_Sphenopalatine_VS_P2_P4 | 1 | 0 | 1 | 0 | 1 | 1 | 1 | 1 |
| SULF2 | 1.28 | P1_P2_P4_Lumbar_Coeliac_Stellate_Sphenopalatine_VS_P3 | 1 | 1 | 0 | 1 | 1 | 1 | 1 | 1 |
| RNF207 | 1.27 | P4_VS_P1_P2_P3_Lumbar_Coeliac_Stellate_Sphenopalatine | 0 | 0 | 0 | 1 | 0 | 0 | 0 | 0 |
| B3GNT8 | 1.27 | P4_VS_P1_P2_P3_Lumbar_Coeliac_Stellate_Sphenopalatine | 0 | 0 | 0 | 1 | 0 | 0 | 0 | 0 |
| RUNX1T1 | 1.27 | Lumbar_VS_P1_P2_P3_P4_Coeliac_Stellate_Sphenopalatine | 0 | 0 | 0 | 0 | 1 | 0 | 0 | 0 |
| CXCR4 | 1.27 | Lumbar_Coeliac_VS_P1_P2_P3_P4_Stellate_Sphenopalatine | 0 | 0 | 0 | 0 | 1 | 1 | 0 | 0 |
| DLEU2 | 1.27 | Lumbar_Coeliac_VS_P1_P2_P3_P4_Stellate_Sphenopalatine | 0 | 0 | 0 | 0 | 1 | 1 | 0 | 0 |
| SPTB | 1.27 | P1_P2_P3_P4_Lumbar_Coeliac_VS_Stellate_Sphenopalatine | 1 | 1 | 1 | 1 | 1 | 1 | 0 | 0 |
| MAOB | 1.27 | P1_P2_P3_P4_Lumbar_Coeliac_Stellate_VS_Sphenopalatine | 1 | 1 | 1 | 1 | 1 | 1 | 1 | 0 |
| DOCK10 | 1.27 | Sphenopalatine_VS_P1_P2_P3_P4_Lumbar_Coeliac_Stellate | 0 | 0 | 0 | 0 | 0 | 0 | 0 | 1 |
| KCNK2 | 1.27 | P1_P2_P3_P4_Lumbar_Coeliac_Sphenopalatine_VS_Stellate | 1 | 1 | 1 | 1 | 1 | 1 | 0 | 1 |
| ARHGEF11 | 1.27 | P1_P2_P3_P4_Lumbar_Coeliac_Sphenopalatine_VS_Stellate | 1 | 1 | 1 | 1 | 1 | 1 | 0 | 1 |
| SCN1B | 1.27 | P1_P2_P3_P4_Lumbar_Coeliac_Sphenopalatine_VS_Stellate | 1 | 1 | 1 | 1 | 1 | 1 | 0 | 1 |
| PALLD | 1.27 | P1_P2_P3_P4_Lumbar_Coeliac_Sphenopalatine_VS_Stellate | 1 | 1 | 1 | 1 | 1 | 1 | 0 | 1 |
| EYA4 | 1.27 | P1_P2_P3_P4_Lumbar_Coeliac_Sphenopalatine_VS_Stellate | 1 | 1 | 1 | 1 | 1 | 1 | 0 | 1 |
| PTPRE | 1.27 | P1_P2_P4_Lumbar_Coeliac_Stellate_Sphenopalatine_VS_P3 | 1 | 1 | 0 | 1 | 1 | 1 | 1 | 1 |
| TLE3 | 1.26 | P2_VS_P1_P3_P4_Lumbar_Coeliac_Stellate_Sphenopalatine | 0 | 1 | 0 | 0 | 0 | 0 | 0 | 0 |
| RIPPLY2 | 1.26 | Coeliac_VS_P1_P2_P3_P4_Lumbar_Stellate_Sphenopalatine | 0 | 0 | 0 | 0 | 0 | 1 | 0 | 0 |
| DOK5 | 1.26 | P1_P2_P3_P4_Lumbar_Coeliac_Stellate_VS_Sphenopalatine | 1 | 1 | 1 | 1 | 1 | 1 | 1 | 0 |
| XAF1 | 1.26 | P1_Sphenopalatine_VS_P2_P3_P4_Lumbar_Coeliac_Stellate | 1 | 0 | 0 | 0 | 0 | 0 | 0 | 1 |
| RAB3B | 1.26 | P1_P2_P3_Lumbar_Coeliac_Stellate_Sphenopalatine_VS_P4 | 1 | 1 | 1 | 0 | 1 | 1 | 1 | 1 |
| PDE8B | 1.25 | P2_VS_P1_P3_P4_Lumbar_Coeliac_Stellate_Sphenopalatine | 0 | 1 | 0 | 0 | 0 | 0 | 0 | 0 |
| B3GALT5 | 1.25 | P2_VS_P1_P3_P4_Lumbar_Coeliac_Stellate_Sphenopalatine | 0 | 1 | 0 | 0 | 0 | 0 | 0 | 0 |
| EEPD1 | 1.25 | P4_VS_P1_P2_P3_Lumbar_Coeliac_Stellate_Sphenopalatine | 0 | 0 | 0 | 1 | 0 | 0 | 0 | 0 |
| CCN2 | 1.25 | P4_VS_P1_P2_P3_Lumbar_Coeliac_Stellate_Sphenopalatine | 0 | 0 | 0 | 1 | 0 | 0 | 0 | 0 |
| RAB37 | 1.25 | P4_VS_P1_P2_P3_Lumbar_Coeliac_Stellate_Sphenopalatine | 0 | 0 | 0 | 1 | 0 | 0 | 0 | 0 |

| | | | | | | | | | | |
|---|---|---|---|---|---|---|---|---|---|---|
| LPAR4 | 1.25 | P4_VS_P1_P2_P3_Lumbar_Coeliac_Stellate_Sphenopalatine | 0 | 0 | 0 | 1 | 0 | 0 | 0 | 0 |
| PHGDH | 1.25 | P1_Sphenopalatine_VS_P2_P3_P4_Lumbar_Coeliac_Stellate | 1 | 0 | 0 | 0 | 0 | 0 | 0 | 1 |
| SHISAL1 | 1.25 | P1_P2_P3_Sphenopalatine_VS_P4_Lumbar_Coeliac_Stellate | 1 | 1 | 1 | 0 | 0 | 0 | 0 | 1 |
| NPEPL1 | 1.25 | P1_P2_P4_Sphenopalatine_VS_P3_Lumbar_Coeliac_Stellate | 1 | 1 | 0 | 1 | 0 | 0 | 0 | 1 |
| CREBRF | 1.25 | P1_P2_P3_P4_Lumbar_Coeliac_Sphenopalatine_VS_Stellate | 1 | 1 | 1 | 1 | 1 | 1 | 0 | 1 |
| DYNC1I1 | 1.25 | P2_P3_P4_Lumbar_Coeliac_Stellate_Sphenopalatine_VS_P1 | 0 | 1 | 1 | 1 | 1 | 1 | 1 | 1 |
| COL16A1 | 1.24 | P4_VS_P1_P2_P3_Lumbar_Coeliac_Stellate_Sphenopalatine | 0 | 0 | 0 | 1 | 0 | 0 | 0 | 0 |
| IFIT1BL1 | 1.24 | P4_VS_P1_P2_P3_Lumbar_Coeliac_Stellate_Sphenopalatine | 0 | 0 | 0 | 1 | 0 | 0 | 0 | 0 |
| B3GNT5 | 1.24 | Lumbar_VS_P1_P2_P3_P4_Coeliac_Stellate_Sphenopalatine | 0 | 0 | 0 | 0 | 1 | 0 | 0 | 0 |
| CAR10 | 1.24 | P1_Sphenopalatine_VS_P2_P3_P4_Lumbar_Coeliac_Stellate | 1 | 0 | 0 | 0 | 0 | 0 | 0 | 1 |
| GM40841 | 1.24 | P1_P2_P3_P4_Lumbar_Coeliac_Sphenopalatine_VS_Stellate | 1 | 1 | 1 | 1 | 1 | 1 | 0 | 1 |
| CHRM2 | 1.24 | P1_P3_P4_Lumbar_Coeliac_Stellate_Sphenopalatine_VS_P2 | 1 | 0 | 1 | 1 | 1 | 1 | 1 | 1 |
| VSNL1 | 1.23 | P4_VS_P1_P2_P3_Lumbar_Coeliac_Stellate_Sphenopalatine | 0 | 0 | 0 | 1 | 0 | 0 | 0 | 0 |
| MYO1B | 1.23 | P1_P2_P3_P4_Lumbar_Coeliac_Stellate_VS_Sphenopalatine | 1 | 1 | 1 | 1 | 1 | 1 | 1 | 0 |
| DGKK | 1.23 | Lumbar_Coeliac_Sphenopalatine_VS_P1_P2_P3_P4_Stellate | 0 | 0 | 0 | 0 | 1 | 1 | 0 | 1 |
| EEF1A2 | 1.23 | P1_P2_P3_P4_Lumbar_Coeliac_Sphenopalatine_VS_Stellate | 1 | 1 | 1 | 1 | 1 | 1 | 0 | 1 |
| CASZ1 | 1.23 | P1_P2_P4_Lumbar_Coeliac_Stellate_Sphenopalatine_VS_P3 | 1 | 1 | 0 | 1 | 1 | 1 | 1 | 1 |
| REPS2 | 1.23 | P1_P2_P4_Lumbar_Coeliac_Stellate_Sphenopalatine_VS_P3 | 1 | 1 | 0 | 1 | 1 | 1 | 1 | 1 |
| FNDC1 | 1.22 | P2_VS_P1_P3_P4_Lumbar_Coeliac_Stellate_Sphenopalatine | 0 | 1 | 0 | 0 | 0 | 0 | 0 | 0 |
| FBN2 | 1.22 | P2_VS_P1_P3_P4_Lumbar_Coeliac_Stellate_Sphenopalatine | 0 | 1 | 0 | 0 | 0 | 0 | 0 | 0 |
| A2ML1 | 1.22 | P4_VS_P1_P2_P3_Lumbar_Coeliac_Stellate_Sphenopalatine | 0 | 0 | 0 | 1 | 0 | 0 | 0 | 0 |
| MAML2 | 1.22 | P4_VS_P1_P2_P3_Lumbar_Coeliac_Stellate_Sphenopalatine | 0 | 0 | 0 | 1 | 0 | 0 | 0 | 0 |
| ACSBG1 | 1.22 | P4_VS_P1_P2_P3_Lumbar_Coeliac_Stellate_Sphenopalatine | 0 | 0 | 0 | 1 | 0 | 0 | 0 | 0 |
| SOX1OT | 1.22 | P1_P4_VS_P2_P3_Lumbar_Coeliac_Stellate_Sphenopalatine | 1 | 0 | 0 | 1 | 0 | 0 | 0 | 0 |
| TESCL | 1.22 | Lumbar_Coeliac_VS_P1_P2_P3_P4_Stellate_Sphenopalatine | 0 | 0 | 0 | 0 | 1 | 1 | 0 | 0 |
| ATP7A | 1.22 | P1_P2_P3_P4_Lumbar_Coeliac_Stellate_VS_Sphenopalatine | 1 | 1 | 1 | 1 | 1 | 1 | 1 | 0 |
| ITPKB | 1.22 | P1_P2_P3_P4_Sphenopalatine_VS_Lumbar_Coeliac_Stellate | 1 | 1 | 1 | 1 | 0 | 0 | 0 | 1 |
| PPP6R1 | 1.22 | P1_P2_P3_P4_Lumbar_Coeliac_Sphenopalatine_VS_Stellate | 1 | 1 | 1 | 1 | 1 | 1 | 0 | 1 |
| RAD50 | 1.22 | P1_P2_P3_P4_Lumbar_Coeliac_Sphenopalatine_VS_Stellate | 1 | 1 | 1 | 1 | 1 | 1 | 0 | 1 |
| L3MBTL4 | 1.22 | P1_P3_P4_Lumbar_Coeliac_Stellate_Sphenopalatine_VS_P2 | 1 | 0 | 1 | 1 | 1 | 1 | 1 | 1 |

| | | | | | | | | | | |
|---|---|---|---|---|---|---|---|---|---|---|
| PRKCH | 1.21 | P2_VS_P1_P3_P4_Lumbar_Coeliac_Stellate_Sphenopalatine | 0 | 1 | 0 | 0 | 0 | 0 | 0 | 0 |
| GM30074 | 1.21 | P3_VS_P1_P2_P4_Lumbar_Coeliac_Stellate_Sphenopalatine | 0 | 0 | 1 | 0 | 0 | 0 | 0 | 0 |
| SYNDIG1L | 1.21 | P3_VS_P1_P2_P4_Lumbar_Coeliac_Stellate_Sphenopalatine | 0 | 0 | 1 | 0 | 0 | 0 | 0 | 0 |
| GM43154 | 1.21 | P4_VS_P1_P2_P3_Lumbar_Coeliac_Stellate_Sphenopalatine | 0 | 0 | 0 | 1 | 0 | 0 | 0 | 0 |
| INSC | 1.21 | P4_VS_P1_P2_P3_Lumbar_Coeliac_Stellate_Sphenopalatine | 0 | 0 | 0 | 1 | 0 | 0 | 0 | 0 |
| GPRC5B | 1.21 | P4_VS_P1_P2_P3_Lumbar_Coeliac_Stellate_Sphenopalatine | 0 | 0 | 0 | 1 | 0 | 0 | 0 | 0 |
| MPP7 | 1.21 | Lumbar_Coeliac_VS_P1_P2_P3_P4_Stellate_Sphenopalatine | 0 | 0 | 0 | 0 | 1 | 1 | 0 | 0 |
| LRRC7 | 1.21 | Sphenopalatine_VS_P1_P2_P3_P4_Lumbar_Coeliac_Stellate | 0 | 0 | 0 | 0 | 0 | 0 | 0 | 1 |
| B3GLCT | 1.21 | P1_P2_P3_P4_Sphenopalatine_VS_Lumbar_Coeliac_Stellate | 1 | 1 | 1 | 1 | 0 | 0 | 0 | 1 |
| COTL1 | 1.21 | P1_P2_P3_P4_Coeliac_Sphenopalatine_VS_Lumbar_Stellate | 1 | 1 | 1 | 1 | 0 | 1 | 0 | 1 |
| SMAD7 | 1.21 | P1_P2_P3_P4_Lumbar_Coeliac_Sphenopalatine_VS_Stellate | 1 | 1 | 1 | 1 | 1 | 1 | 0 | 1 |
| CARTPT | 1.21 | P1_P2_P3_P4_Coeliac_Stellate_Sphenopalatine_VS_Lumbar | 1 | 1 | 1 | 1 | 0 | 1 | 1 | 1 |
| OCIAD2 | 1.21 | P3_P4_Lumbar_Coeliac_Stellate_Sphenopalatine_VS_P1_P2 | 0 | 0 | 1 | 1 | 1 | 1 | 1 | 1 |
| BRINP1 | 1.2 | P2_P3_VS_P1_P4_Lumbar_Coeliac_Stellate_Sphenopalatine | 0 | 1 | 1 | 0 | 0 | 0 | 0 | 0 |
| ATP1A2 | 1.2 | P4_VS_P1_P2_P3_Lumbar_Coeliac_Stellate_Sphenopalatine | 0 | 0 | 0 | 1 | 0 | 0 | 0 | 0 |
| NPPC | 1.2 | Coeliac_VS_P1_P2_P3_P4_Lumbar_Stellate_Sphenopalatine | 0 | 0 | 0 | 0 | 0 | 1 | 0 | 0 |
| GM45321 | 1.2 | Coeliac_VS_P1_P2_P3_P4_Lumbar_Stellate_Sphenopalatine | 0 | 0 | 0 | 0 | 0 | 1 | 0 | 0 |
| SLC16A9 | 1.2 | Lumbar_Coeliac_VS_P1_P2_P3_P4_Stellate_Sphenopalatine | 0 | 0 | 0 | 0 | 1 | 1 | 0 | 0 |
| GM28221 | 1.2 | Lumbar_Coeliac_VS_P1_P2_P3_P4_Stellate_Sphenopalatine | 0 | 0 | 0 | 0 | 1 | 1 | 0 | 0 |
| SAMD11 | 1.2 | P3_Lumbar_Coeliac_Stellate_VS_P1_P2_P4_Sphenopalatine | 0 | 0 | 1 | 0 | 1 | 1 | 1 | 0 |
| SOX2OT | 1.2 | Sphenopalatine_VS_P1_P2_P3_P4_Lumbar_Coeliac_Stellate | 0 | 0 | 0 | 0 | 0 | 0 | 0 | 1 |
| CDH10 | 1.2 | Sphenopalatine_VS_P1_P2_P3_P4_Lumbar_Coeliac_Stellate | 0 | 0 | 0 | 0 | 0 | 0 | 0 | 1 |
| KAZN | 1.2 | P1_P2_Sphenopalatine_VS_P3_P4_Lumbar_Coeliac_Stellate | 1 | 1 | 0 | 0 | 0 | 0 | 0 | 1 |
| CNTN5 | 1.2 | P1_P2_P3_Sphenopalatine_VS_P4_Lumbar_Coeliac_Stellate | 1 | 1 | 1 | 0 | 0 | 0 | 0 | 1 |
| HTR3A | 1.2 | P1_P2_P3_P4_Sphenopalatine_VS_Lumbar_Coeliac_Stellate | 1 | 1 | 1 | 1 | 0 | 0 | 0 | 1 |
| SH3BP4 | 1.2 | P1_P2_P3_P4_Lumbar_Coeliac_Sphenopalatine_VS_Stellate | 1 | 1 | 1 | 1 | 1 | 1 | 0 | 1 |
| WHAMM | 1.2 | P1_P2_P3_P4_Lumbar_Coeliac_Sphenopalatine_VS_Stellate | 1 | 1 | 1 | 1 | 1 | 1 | 0 | 1 |
| SESN3 | 1.2 | P1_P2_P3_P4_Lumbar_Coeliac_Sphenopalatine_VS_Stellate | 1 | 1 | 1 | 1 | 1 | 1 | 0 | 1 |
| RASGRF2 | 1.2 | P1_P2_P3_P4_Lumbar_Coeliac_Sphenopalatine_VS_Stellate | 1 | 1 | 1 | 1 | 1 | 1 | 0 | 1 |
| RYR3 | 1.19 | P1_VS_P2_P3_P4_Lumbar_Coeliac_Stellate_Sphenopalatine | 1 | 0 | 0 | 0 | 0 | 0 | 0 | 0 |

| | | | | | | | | | | |
|---|---|---|---|---|---|---|---|---|---|---|
| FAM129A | 1.19 | P4_VS_P1_P2_P3_Lumbar_Coeliac_Stellate_Sphenopalatine | 0 | 0 | 0 | 1 | 0 | 0 | 0 | 0 |
| IRF5 | 1.19 | P4_VS_P1_P2_P3_Lumbar_Coeliac_Stellate_Sphenopalatine | 0 | 0 | 0 | 1 | 0 | 0 | 0 | 0 |
| DBPHT2 | 1.19 | P3_P4_VS_P1_P2_Lumbar_Coeliac_Stellate_Sphenopalatine | 0 | 0 | 1 | 1 | 0 | 0 | 0 | 0 |
| 6030407O03 | 1.19 | Coeliac_VS_P1_P2_P3_P4_Lumbar_Stellate_Sphenopalatine | 0 | 0 | 0 | 0 | 0 | 1 | 0 | 0 |
| FAXC | 1.19 | Lumbar_Coeliac_VS_P1_P2_P3_P4_Stellate_Sphenopalatine | 0 | 0 | 0 | 0 | 1 | 1 | 0 | 0 |
| ZBTB12 | 1.19 | Lumbar_Coeliac_VS_P1_P2_P3_P4_Stellate_Sphenopalatine | 0 | 0 | 0 | 0 | 1 | 1 | 0 | 0 |
| TENM3 | 1.19 | P3_Lumbar_Coeliac_VS_P1_P2_P4_Stellate_Sphenopalatine | 0 | 0 | 1 | 0 | 1 | 1 | 0 | 0 |
| GPC3 | 1.19 | Sphenopalatine_VS_P1_P2_P3_P4_Lumbar_Coeliac_Stellate | 0 | 0 | 0 | 0 | 0 | 0 | 0 | 1 |
| ALDH1B1 | 1.19 | P1_P3_Lumbar_Coeliac_Sphenopalatine_VS_P2_P4_Stellate | 1 | 0 | 1 | 0 | 1 | 1 | 0 | 1 |
| ZMAT4 | 1.19 | P1_P3_P4_Lumbar_Coeliac_Stellate_Sphenopalatine_VS_P2 | 1 | 0 | 1 | 1 | 1 | 1 | 1 | 1 |
| GM26802 | 1.18 | P4_VS_P1_P2_P3_Lumbar_Coeliac_Stellate_Sphenopalatine | 0 | 0 | 0 | 1 | 0 | 0 | 0 | 0 |
| HLF | 1.18 | Coeliac_VS_P1_P2_P3_P4_Lumbar_Stellate_Sphenopalatine | 0 | 0 | 0 | 0 | 0 | 1 | 0 | 0 |
| CAD | 1.18 | Lumbar_Coeliac_VS_P1_P2_P3_P4_Stellate_Sphenopalatine | 0 | 0 | 0 | 0 | 1 | 1 | 0 | 0 |
| PPP1R36 | 1.18 | Lumbar_Coeliac_VS_P1_P2_P3_P4_Stellate_Sphenopalatine | 0 | 0 | 0 | 0 | 1 | 1 | 0 | 0 |
| NRG2 | 1.18 | Lumbar_Coeliac_VS_P1_P2_P3_P4_Stellate_Sphenopalatine | 0 | 0 | 0 | 0 | 1 | 1 | 0 | 0 |
| FAM189A2 | 1.18 | P1_P2_P3_P4_Lumbar_Coeliac_Stellate_VS_Sphenopalatine | 1 | 1 | 1 | 1 | 1 | 1 | 1 | 0 |
| RAMP3 | 1.18 | P1_P2_P3_Lumbar_Sphenopalatine_VS_P4_Coeliac_Stellate | 1 | 1 | 1 | 0 | 1 | 0 | 0 | 1 |
| FJX1 | 1.18 | P1_P2_P3_P4_Lumbar_Coeliac_Sphenopalatine_VS_Stellate | 1 | 1 | 1 | 1 | 1 | 1 | 0 | 1 |
| AI504432 | 1.18 | P1_P2_P3_P4_Lumbar_Coeliac_Sphenopalatine_VS_Stellate | 1 | 1 | 1 | 1 | 1 | 1 | 0 | 1 |
| ELFN2 | 1.18 | P1_P2_P3_P4_Lumbar_Coeliac_Sphenopalatine_VS_Stellate | 1 | 1 | 1 | 1 | 1 | 1 | 0 | 1 |
| PTCHD1 | 1.18 | P1_P2_P3_P4_Lumbar_Coeliac_Sphenopalatine_VS_Stellate | 1 | 1 | 1 | 1 | 1 | 1 | 0 | 1 |
| GRB10 | 1.18 | P1_P3_P4_Lumbar_Coeliac_Stellate_Sphenopalatine_VS_P2 | 1 | 0 | 1 | 1 | 1 | 1 | 1 | 1 |
| OLFM3 | 1.17 | P1_P3_VS_P2_P4_Lumbar_Coeliac_Stellate_Sphenopalatine | 1 | 0 | 1 | 0 | 0 | 0 | 0 | 0 |
| ARHGAP27 | 1.17 | P4_VS_P1_P2_P3_Lumbar_Coeliac_Stellate_Sphenopalatine | 0 | 0 | 0 | 1 | 0 | 0 | 0 | 0 |
| GREM1 | 1.17 | Lumbar_VS_P1_P2_P3_P4_Coeliac_Stellate_Sphenopalatine | 0 | 0 | 0 | 0 | 1 | 0 | 0 | 0 |
| OPCML | 1.17 | Lumbar_VS_P1_P2_P3_P4_Coeliac_Stellate_Sphenopalatine | 0 | 0 | 0 | 0 | 1 | 0 | 0 | 0 |
| PDLIM3 | 1.17 | Sphenopalatine_VS_P1_P2_P3_P4_Lumbar_Coeliac_Stellate | 0 | 0 | 0 | 0 | 0 | 0 | 0 | 1 |
| GM10605 | 1.17 | P1_P2_P3_P4_Lumbar_Stellate_Sphenopalatine_VS_Coeliac | 1 | 1 | 1 | 1 | 1 | 0 | 1 | 1 |
| DDC | 1.17 | P1_P2_P3_Lumbar_Coeliac_Stellate_Sphenopalatine_VS_P4 | 1 | 1 | 1 | 0 | 1 | 1 | 1 | 1 |
| LRRN3 | 1.17 | P1_P3_P4_Lumbar_Coeliac_Stellate_Sphenopalatine_VS_P2 | 1 | 0 | 1 | 1 | 1 | 1 | 1 | 1 |

| | | | | | | | | | | |
|---|---|---|---|---|---|---|---|---|---|---|
| 4833415N18 | 1.16 | P1_VS_P2_P3_P4_Lumbar_Coeliac_Stellate_Sphenopalatine | 1 | 0 | 0 | 0 | 0 | 0 | 0 | 0 |
| SLC12A5 | 1.16 | P2_VS_P1_P3_P4_Lumbar_Coeliac_Stellate_Sphenopalatine | 0 | 1 | 0 | 0 | 0 | 0 | 0 | 0 |
| POSTN | 1.16 | P4_VS_P1_P2_P3_Lumbar_Coeliac_Stellate_Sphenopalatine | 0 | 0 | 0 | 1 | 0 | 0 | 0 | 0 |
| WWTR1 | 1.16 | P4_VS_P1_P2_P3_Lumbar_Coeliac_Stellate_Sphenopalatine | 0 | 0 | 0 | 1 | 0 | 0 | 0 | 0 |
| BEND4 | 1.16 | P4_VS_P1_P2_P3_Lumbar_Coeliac_Stellate_Sphenopalatine | 0 | 0 | 0 | 1 | 0 | 0 | 0 | 0 |
| TENM1 | 1.16 | P4_VS_P1_P2_P3_Lumbar_Coeliac_Stellate_Sphenopalatine | 0 | 0 | 0 | 1 | 0 | 0 | 0 | 0 |
| SLC18A2 | 1.16 | P1_P2_P3_Lumbar_Coeliac_Stellate_VS_P4_Sphenopalatine | 1 | 1 | 1 | 0 | 1 | 1 | 1 | 0 |
| MSRB2 | 1.16 | P1_P2_P3_P4_Lumbar_Coeliac_Stellate_VS_Sphenopalatine | 1 | 1 | 1 | 1 | 1 | 1 | 1 | 0 |
| 2010204K13I | 1.16 | P1_P2_P3_P4_Lumbar_Coeliac_Stellate_VS_Sphenopalatine | 1 | 1 | 1 | 1 | 1 | 1 | 1 | 0 |
| UNC5C | 1.16 | P1_P2_P4_Lumbar_Coeliac_Sphenopalatine_VS_P3_Stellate | 1 | 1 | 0 | 1 | 1 | 1 | 0 | 1 |
| BMP2K | 1.16 | P1_P2_P3_P4_Lumbar_Coeliac_Sphenopalatine_VS_Stellate | 1 | 1 | 1 | 1 | 1 | 1 | 0 | 1 |
| STAT5B | 1.16 | P1_P2_P3_P4_Lumbar_Coeliac_Sphenopalatine_VS_Stellate | 1 | 1 | 1 | 1 | 1 | 1 | 0 | 1 |
| UTRN | 1.16 | P3_Lumbar_Coeliac_Stellate_Sphenopalatine_VS_P1_P2_P4 | 0 | 0 | 1 | 0 | 1 | 1 | 1 | 1 |
| GRB14 | 1.15 | P3_VS_P1_P2_P4_Lumbar_Coeliac_Stellate_Sphenopalatine | 0 | 0 | 1 | 0 | 0 | 0 | 0 | 0 |
| LGR4 | 1.15 | P4_VS_P1_P2_P3_Lumbar_Coeliac_Stellate_Sphenopalatine | 0 | 0 | 0 | 1 | 0 | 0 | 0 | 0 |
| TNC | 1.15 | P4_VS_P1_P2_P3_Lumbar_Coeliac_Stellate_Sphenopalatine | 0 | 0 | 0 | 1 | 0 | 0 | 0 | 0 |
| TULP2 | 1.15 | P4_VS_P1_P2_P3_Lumbar_Coeliac_Stellate_Sphenopalatine | 0 | 0 | 0 | 1 | 0 | 0 | 0 | 0 |
| ANK1 | 1.15 | P4_VS_P1_P2_P3_Lumbar_Coeliac_Stellate_Sphenopalatine | 0 | 0 | 0 | 1 | 0 | 0 | 0 | 0 |
| GPR119 | 1.15 | P4_VS_P1_P2_P3_Lumbar_Coeliac_Stellate_Sphenopalatine | 0 | 0 | 0 | 1 | 0 | 0 | 0 | 0 |
| PRR15 | 1.15 | P1_P4_VS_P2_P3_Lumbar_Coeliac_Stellate_Sphenopalatine | 1 | 0 | 0 | 1 | 0 | 0 | 0 | 0 |
| BMP1 | 1.15 | Lumbar_Coeliac_VS_P1_P2_P3_P4_Stellate_Sphenopalatine | 0 | 0 | 0 | 0 | 1 | 1 | 0 | 0 |
| PTGER3 | 1.15 | P4_Lumbar_Coeliac_VS_P1_P2_P3_Stellate_Sphenopalatine | 0 | 0 | 0 | 1 | 1 | 1 | 0 | 0 |
| CACNA1D | 1.15 | P1_P2_P3_P4_Lumbar_Coeliac_VS_Stellate_Sphenopalatine | 1 | 1 | 1 | 1 | 1 | 1 | 0 | 0 |
| GALNT16 | 1.15 | P2_P3_Lumbar_Coeliac_Stellate_VS_P1_P4_Sphenopalatine | 0 | 1 | 1 | 0 | 1 | 1 | 1 | 0 |
| NTNG1 | 1.15 | P1_P2_P3_P4_Lumbar_Coeliac_Stellate_VS_Sphenopalatine | 1 | 1 | 1 | 1 | 1 | 1 | 1 | 0 |
| NPY1R | 1.15 | P1_P2_P3_Sphenopalatine_VS_P4_Lumbar_Coeliac_Stellate | 1 | 1 | 1 | 0 | 0 | 0 | 0 | 1 |
| ADGRL2 | 1.15 | P1_P2_P3_P4_Lumbar_Coeliac_Sphenopalatine_VS_Stellate | 1 | 1 | 1 | 1 | 1 | 1 | 0 | 1 |
| ZBTB1 | 1.15 | P1_P2_P3_P4_Lumbar_Coeliac_Sphenopalatine_VS_Stellate | 1 | 1 | 1 | 1 | 1 | 1 | 0 | 1 |
| PLXNC1 | 1.15 | P2_P3_P4_Lumbar_Coeliac_Stellate_Sphenopalatine_VS_P1 | 0 | 1 | 1 | 1 | 1 | 1 | 1 | 1 |
| SOSTDC1 | 1.14 | P4_VS_P1_P2_P3_Lumbar_Coeliac_Stellate_Sphenopalatine | 0 | 0 | 0 | 1 | 0 | 0 | 0 | 0 |

| | | | | | | | | | | |
|---|---|---|---|---|---|---|---|---|---|---|
| TAFA2 | 1.14 | P1_P2_P4_VS_P3_Lumbar_Coeliac_Stellate_Sphenopalatine | 1 | 1 | 0 | 1 | 0 | 0 | 0 | 0 |
| GM47033 | 1.14 | Lumbar_Coeliac_VS_P1_P2_P3_P4_Stellate_Sphenopalatine | 0 | 0 | 0 | 0 | 1 | 1 | 0 | 0 |
| GM13889 | 1.14 | P1_P2_P3_P4_Lumbar_Coeliac_Stellate_VS_Sphenopalatine | 1 | 1 | 1 | 1 | 1 | 1 | 1 | 0 |
| CDH11 | 1.14 | P1_P4_Sphenopalatine_VS_P2_P3_Lumbar_Coeliac_Stellate | 1 | 0 | 0 | 1 | 0 | 0 | 0 | 1 |
| ASS1 | 1.14 | P1_P2_P3_P4_Sphenopalatine_VS_Lumbar_Coeliac_Stellate | 1 | 1 | 1 | 1 | 0 | 0 | 0 | 1 |
| AFF3 | 1.14 | P1_P2_P3_P4_Lumbar_Coeliac_Sphenopalatine_VS_Stellate | 1 | 1 | 1 | 1 | 1 | 1 | 0 | 1 |
| PYGL | 1.13 | Lumbar_VS_P1_P2_P3_P4_Coeliac_Stellate_Sphenopalatine | 0 | 0 | 0 | 0 | 1 | 0 | 0 | 0 |
| TMEM28 | 1.13 | Lumbar_VS_P1_P2_P3_P4_Coeliac_Stellate_Sphenopalatine | 0 | 0 | 0 | 0 | 1 | 0 | 0 | 0 |
| MITF | 1.13 | Coeliac_VS_P1_P2_P3_P4_Lumbar_Stellate_Sphenopalatine | 0 | 0 | 0 | 0 | 0 | 1 | 0 | 0 |
| PDE1B | 1.13 | P1_P2_P3_P4_Lumbar_Coeliac_Stellate_VS_Sphenopalatine | 1 | 1 | 1 | 1 | 1 | 1 | 1 | 0 |
| GPC6 | 1.13 | Lumbar_Sphenopalatine_VS_P1_P2_P3_P4_Coeliac_Stellate | 0 | 0 | 0 | 0 | 1 | 0 | 0 | 1 |
| FAT4 | 1.13 | P1_P2_P3_P4_Lumbar_Coeliac_Sphenopalatine_VS_Stellate | 1 | 1 | 1 | 1 | 1 | 1 | 0 | 1 |
| CNTLN | 1.13 | P1_P2_P3_P4_Lumbar_Coeliac_Sphenopalatine_VS_Stellate | 1 | 1 | 1 | 1 | 1 | 1 | 0 | 1 |
| KCNC1 | 1.13 | P1_P2_P3_P4_Lumbar_Coeliac_Sphenopalatine_VS_Stellate | 1 | 1 | 1 | 1 | 1 | 1 | 0 | 1 |
| PACS1 | 1.13 | P1_P2_P3_P4_Lumbar_Coeliac_Sphenopalatine_VS_Stellate | 1 | 1 | 1 | 1 | 1 | 1 | 0 | 1 |
| TRPM2 | 1.13 | P1_P2_P3_Lumbar_Coeliac_Stellate_Sphenopalatine_VS_P4 | 1 | 1 | 1 | 0 | 1 | 1 | 1 | 1 |
| RND3 | 1.13 | P1_P4_Lumbar_Coeliac_Stellate_Sphenopalatine_VS_P2_P3 | 1 | 0 | 0 | 1 | 1 | 1 | 1 | 1 |
| MAL | 1.12 | P4_VS_P1_P2_P3_Lumbar_Coeliac_Stellate_Sphenopalatine | 0 | 0 | 0 | 1 | 0 | 0 | 0 | 0 |
| MAF | 1.12 | P4_VS_P1_P2_P3_Lumbar_Coeliac_Stellate_Sphenopalatine | 0 | 0 | 0 | 1 | 0 | 0 | 0 | 0 |
| SFRP1 | 1.12 | P4_Lumbar_VS_P1_P2_P3_Coeliac_Stellate_Sphenopalatine | 0 | 0 | 0 | 1 | 1 | 0 | 0 | 0 |
| CALM4 | 1.12 | Coeliac_VS_P1_P2_P3_P4_Lumbar_Stellate_Sphenopalatine | 0 | 0 | 0 | 0 | 0 | 1 | 0 | 0 |
| DAB1 | 1.12 | P4_Coeliac_VS_P1_P2_P3_Lumbar_Stellate_Sphenopalatine | 0 | 0 | 0 | 1 | 0 | 1 | 0 | 0 |
| GM26984 | 1.12 | Lumbar_Coeliac_VS_P1_P2_P3_P4_Stellate_Sphenopalatine | 0 | 0 | 0 | 0 | 1 | 1 | 0 | 0 |
| IGFBP4 | 1.12 | Sphenopalatine_VS_P1_P2_P3_P4_Lumbar_Coeliac_Stellate | 0 | 0 | 0 | 0 | 0 | 0 | 0 | 1 |
| H1FX | 1.12 | P1_P2_P3_P4_Lumbar_Coeliac_Sphenopalatine_VS_Stellate | 1 | 1 | 1 | 1 | 1 | 1 | 0 | 1 |
| KIF13B | 1.12 | P1_P2_P3_P4_Lumbar_Coeliac_Sphenopalatine_VS_Stellate | 1 | 1 | 1 | 1 | 1 | 1 | 0 | 1 |
| CRISPLD1 | 1.12 | P1_P2_Lumbar_Coeliac_Stellate_Sphenopalatine_VS_P3_P4 | 1 | 1 | 0 | 0 | 1 | 1 | 1 | 1 |
| LAMA4 | 1.11 | P1_VS_P2_P3_P4_Lumbar_Coeliac_Stellate_Sphenopalatine | 1 | 0 | 0 | 0 | 0 | 0 | 0 | 0 |
| NPAS4 | 1.11 | P2_P4_VS_P1_P3_Lumbar_Coeliac_Stellate_Sphenopalatine | 0 | 1 | 0 | 1 | 0 | 0 | 0 | 0 |
| SPATA31D1A | 1.11 | Coeliac_VS_P1_P2_P3_P4_Lumbar_Stellate_Sphenopalatine | 0 | 0 | 0 | 0 | 0 | 1 | 0 | 0 |

| | | | | | | | | | | |
|---|---|---|---|---|---|---|---|---|---|---|
| DDX23 | 1.11 | Lumbar_Coeliac_VS_P1_P2_P3_P4_Stellate_Sphenopalatine | 0 | 0 | 0 | 0 | 1 | 1 | 0 | 0 |
| CXADR | 1.11 | Lumbar_Coeliac_VS_P1_P2_P3_P4_Stellate_Sphenopalatine | 0 | 0 | 0 | 0 | 1 | 1 | 0 | 0 |
| ZFP800 | 1.11 | P1_P2_P3_P4_Lumbar_Coeliac_VS_Stellate_Sphenopalatine | 1 | 1 | 1 | 1 | 1 | 1 | 0 | 0 |
| GM5148 | 1.11 | Sphenopalatine_VS_P1_P2_P3_P4_Lumbar_Coeliac_Stellate | 0 | 0 | 0 | 0 | 0 | 0 | 0 | 1 |
| LOXL1 | 1.11 | Sphenopalatine_VS_P1_P2_P3_P4_Lumbar_Coeliac_Stellate | 0 | 0 | 0 | 0 | 0 | 0 | 0 | 1 |
| GREB1L | 1.11 | Sphenopalatine_VS_P1_P2_P3_P4_Lumbar_Coeliac_Stellate | 0 | 0 | 0 | 0 | 0 | 0 | 0 | 1 |
| VWF | 1.11 | P1_Sphenopalatine_VS_P2_P3_P4_Lumbar_Coeliac_Stellate | 1 | 0 | 0 | 0 | 0 | 0 | 0 | 1 |
| VANGL1 | 1.11 | P1_P2_P4_Sphenopalatine_VS_P3_Lumbar_Coeliac_Stellate | 1 | 1 | 0 | 1 | 0 | 0 | 0 | 1 |
| CELSR1 | 1.1 | P3_VS_P1_P2_P4_Lumbar_Coeliac_Stellate_Sphenopalatine | 0 | 0 | 1 | 0 | 0 | 0 | 0 | 0 |
| PLD5 | 1.1 | P4_VS_P1_P2_P3_Lumbar_Coeliac_Stellate_Sphenopalatine | 0 | 0 | 0 | 1 | 0 | 0 | 0 | 0 |
| SLC7A2 | 1.1 | P4_VS_P1_P2_P3_Lumbar_Coeliac_Stellate_Sphenopalatine | 0 | 0 | 0 | 1 | 0 | 0 | 0 | 0 |
| LAMA2 | 1.1 | P4_VS_P1_P2_P3_Lumbar_Coeliac_Stellate_Sphenopalatine | 0 | 0 | 0 | 1 | 0 | 0 | 0 | 0 |
| MYO1A | 1.1 | Lumbar_VS_P1_P2_P3_P4_Coeliac_Stellate_Sphenopalatine | 0 | 0 | 0 | 0 | 1 | 0 | 0 | 0 |
| COCH | 1.1 | Lumbar_VS_P1_P2_P3_P4_Coeliac_Stellate_Sphenopalatine | 0 | 0 | 0 | 0 | 1 | 0 | 0 | 0 |
| HIST2H2AC | 1.1 | Coeliac_VS_P1_P2_P3_P4_Lumbar_Stellate_Sphenopalatine | 0 | 0 | 0 | 0 | 0 | 1 | 0 | 0 |
| ZFP976 | 1.1 | Lumbar_Coeliac_VS_P1_P2_P3_P4_Stellate_Sphenopalatine | 0 | 0 | 0 | 0 | 1 | 1 | 0 | 0 |
| LCORL | 1.1 | P1_P2_P3_P4_Lumbar_Coeliac_VS_Stellate_Sphenopalatine | 1 | 1 | 1 | 1 | 1 | 1 | 0 | 0 |
| NDP | 1.1 | Lumbar_Coeliac_Stellate_VS_P1_P2_P3_P4_Sphenopalatine | 0 | 0 | 0 | 0 | 1 | 1 | 1 | 0 |
| CIDEA | 1.1 | P1_P2_P3_Sphenopalatine_VS_P4_Lumbar_Coeliac_Stellate | 1 | 1 | 1 | 0 | 0 | 0 | 0 | 1 |
| MOSMO | 1.1 | P1_P2_P3_P4_Lumbar_Coeliac_Sphenopalatine_VS_Stellate | 1 | 1 | 1 | 1 | 1 | 1 | 0 | 1 |
| GPRIN2 | 1.1 | P1_P2_P3_P4_Lumbar_Coeliac_Sphenopalatine_VS_Stellate | 1 | 1 | 1 | 1 | 1 | 1 | 0 | 1 |
| KHSRP | 1.1 | P1_P2_P3_P4_Lumbar_Coeliac_Sphenopalatine_VS_Stellate | 1 | 1 | 1 | 1 | 1 | 1 | 0 | 1 |
| LRRTM4 | 1.1 | P1_P4_Lumbar_Stellate_Sphenopalatine_VS_P2_P3_Coeliac | 1 | 0 | 0 | 1 | 1 | 0 | 1 | 1 |
| CACNG4 | 1.1 | P1_P2_P3_Lumbar_Coeliac_Stellate_Sphenopalatine_VS_P4 | 1 | 1 | 1 | 0 | 1 | 1 | 1 | 1 |
| REM2 | 1.09 | P3_VS_P1_P2_P4_Lumbar_Coeliac_Stellate_Sphenopalatine | 0 | 0 | 1 | 0 | 0 | 0 | 0 | 0 |
| FIGN | 1.09 | P4_VS_P1_P2_P3_Lumbar_Coeliac_Stellate_Sphenopalatine | 0 | 0 | 0 | 1 | 0 | 0 | 0 | 0 |
| MMP14 | 1.09 | P4_VS_P1_P2_P3_Lumbar_Coeliac_Stellate_Sphenopalatine | 0 | 0 | 0 | 1 | 0 | 0 | 0 | 0 |
| GM45338 | 1.09 | P1_P2_P4_VS_P3_Lumbar_Coeliac_Stellate_Sphenopalatine | 1 | 1 | 0 | 1 | 0 | 0 | 0 | 0 |
| DOK7 | 1.09 | Lumbar_Coeliac_VS_P1_P2_P3_P4_Stellate_Sphenopalatine | 0 | 0 | 0 | 0 | 1 | 1 | 0 | 0 |
| MAP2K6 | 1.09 | Lumbar_Coeliac_VS_P1_P2_P3_P4_Stellate_Sphenopalatine | 0 | 0 | 0 | 0 | 1 | 1 | 0 | 0 |

| | | | | | | | | | | |
|---|---|---|---|---|---|---|---|---|---|---|
| ZFP758 | 1.09 | Lumbar_Coeliac_VS_P1_P2_P3_P4_Stellate_Sphenopalatine | 0 | 0 | 0 | 0 | 1 | 1 | 0 | 0 |
| TRIM66 | 1.09 | Sphenopalatine_VS_P1_P2_P3_P4_Lumbar_Coeliac_Stellate | 0 | 0 | 0 | 0 | 0 | 0 | 0 | 1 |
| SYNE1 | 1.09 | P1_P2_P3_P4_Lumbar_Coeliac_Sphenopalatine_VS_Stellate | 1 | 1 | 1 | 1 | 1 | 1 | 0 | 1 |
| MLH3 | 1.09 | P1_P2_P3_P4_Lumbar_Coeliac_Sphenopalatine_VS_Stellate | 1 | 1 | 1 | 1 | 1 | 1 | 0 | 1 |
| MBD2 | 1.09 | P1_P2_P3_P4_Lumbar_Coeliac_Sphenopalatine_VS_Stellate | 1 | 1 | 1 | 1 | 1 | 1 | 0 | 1 |
| TESC | 1.09 | P1_P3_P4_Lumbar_Coeliac_Stellate_Sphenopalatine_VS_P2 | 1 | 0 | 1 | 1 | 1 | 1 | 1 | 1 |
| DLX3 | 1.08 | P3_VS_P1_P2_P4_Lumbar_Coeliac_Stellate_Sphenopalatine | 0 | 0 | 1 | 0 | 0 | 0 | 0 | 0 |
| TGFBR3 | 1.08 | P4_VS_P1_P2_P3_Lumbar_Coeliac_Stellate_Sphenopalatine | 0 | 0 | 0 | 1 | 0 | 0 | 0 | 0 |
| CALML4 | 1.08 | P2_P4_VS_P1_P3_Lumbar_Coeliac_Stellate_Sphenopalatine | 0 | 1 | 0 | 1 | 0 | 0 | 0 | 0 |
| KCNN3 | 1.08 | Lumbar_VS_P1_P2_P3_P4_Coeliac_Stellate_Sphenopalatine | 0 | 0 | 0 | 0 | 1 | 0 | 0 | 0 |
| CHRNA4 | 1.08 | Coeliac_VS_P1_P2_P3_P4_Lumbar_Stellate_Sphenopalatine | 0 | 0 | 0 | 0 | 0 | 1 | 0 | 0 |
| SYNE2 | 1.08 | Coeliac_VS_P1_P2_P3_P4_Lumbar_Stellate_Sphenopalatine | 0 | 0 | 0 | 0 | 0 | 1 | 0 | 0 |
| EZH2 | 1.08 | Lumbar_Coeliac_VS_P1_P2_P3_P4_Stellate_Sphenopalatine | 0 | 0 | 0 | 0 | 1 | 1 | 0 | 0 |
| PRR18 | 1.08 | Lumbar_Coeliac_VS_P1_P2_P3_P4_Stellate_Sphenopalatine | 0 | 0 | 0 | 0 | 1 | 1 | 0 | 0 |
| SHROOM1 | 1.08 | P3_Lumbar_Coeliac_VS_P1_P2_P4_Stellate_Sphenopalatine | 0 | 0 | 1 | 0 | 1 | 1 | 0 | 0 |
| KCNJ3 | 1.08 | P1_P2_P3_P4_Sphenopalatine_VS_Lumbar_Coeliac_Stellate | 1 | 1 | 1 | 1 | 0 | 0 | 0 | 1 |
| PIP4K2A | 1.08 | P1_P2_P3_P4_Lumbar_Coeliac_Sphenopalatine_VS_Stellate | 1 | 1 | 1 | 1 | 1 | 1 | 0 | 1 |
| PIK3R3 | 1.08 | P1_P2_P3_P4_Lumbar_Coeliac_Sphenopalatine_VS_Stellate | 1 | 1 | 1 | 1 | 1 | 1 | 0 | 1 |
| SYNM | 1.08 | P1_P2_P3_P4_Lumbar_Coeliac_Sphenopalatine_VS_Stellate | 1 | 1 | 1 | 1 | 1 | 1 | 0 | 1 |
| FSTL1 | 1.08 | P1_P2_P3_P4_Lumbar_Coeliac_Sphenopalatine_VS_Stellate | 1 | 1 | 1 | 1 | 1 | 1 | 0 | 1 |
| GM26638 | 1.08 | P1_P2_P3_Lumbar_Coeliac_Stellate_Sphenopalatine_VS_P4 | 1 | 1 | 1 | 0 | 1 | 1 | 1 | 1 |
| GRIN3A | 1.07 | P2_VS_P1_P3_P4_Lumbar_Coeliac_Stellate_Sphenopalatine | 0 | 1 | 0 | 0 | 0 | 0 | 0 | 0 |
| GM31143 | 1.07 | P3_VS_P1_P2_P4_Lumbar_Coeliac_Stellate_Sphenopalatine | 0 | 0 | 1 | 0 | 0 | 0 | 0 | 0 |
| RIN2 | 1.07 | P4_VS_P1_P2_P3_Lumbar_Coeliac_Stellate_Sphenopalatine | 0 | 0 | 0 | 1 | 0 | 0 | 0 | 0 |
| BCHE | 1.07 | P4_VS_P1_P2_P3_Lumbar_Coeliac_Stellate_Sphenopalatine | 0 | 0 | 0 | 1 | 0 | 0 | 0 | 0 |
| PTPRZ1 | 1.07 | P4_VS_P1_P2_P3_Lumbar_Coeliac_Stellate_Sphenopalatine | 0 | 0 | 0 | 1 | 0 | 0 | 0 | 0 |
| PKP3 | 1.07 | P4_VS_P1_P2_P3_Lumbar_Coeliac_Stellate_Sphenopalatine | 0 | 0 | 0 | 1 | 0 | 0 | 0 | 0 |
| SYNPO | 1.07 | P4_VS_P1_P2_P3_Lumbar_Coeliac_Stellate_Sphenopalatine | 0 | 0 | 0 | 1 | 0 | 0 | 0 | 0 |
| D130040H23 | 1.07 | Coeliac_VS_P1_P2_P3_P4_Lumbar_Stellate_Sphenopalatine | 0 | 0 | 0 | 0 | 0 | 1 | 0 | 0 |
| E130102H24 | 1.07 | Lumbar_Coeliac_VS_P1_P2_P3_P4_Stellate_Sphenopalatine | 0 | 0 | 0 | 0 | 1 | 1 | 0 | 0 |

| | | | | | | | | | | |
|---|---|---|---|---|---|---|---|---|---|---|
| UTS2B | 1.07 | Stellate_VS_P1_P2_P3_P4_Lumbar_Coeliac_Sphenopalatine | 0 | 0 | 0 | 0 | 0 | 0 | 1 | 0 |
| PRKCD | 1.07 | P1_P2_P3_Lumbar_Coeliac_Stellate_VS_P4_Sphenopalatine | 1 | 1 | 1 | 0 | 1 | 1 | 1 | 0 |
| GNG8 | 1.07 | P1_Lumbar_Sphenopalatine_VS_P2_P3_P4_Coeliac_Stellate | 1 | 0 | 0 | 0 | 1 | 0 | 0 | 1 |
| TNRC18 | 1.07 | P1_P2_P3_P4_Lumbar_Coeliac_Sphenopalatine_VS_Stellate | 1 | 1 | 1 | 1 | 1 | 1 | 0 | 1 |
| LRRC75A | 1.07 | P1_P2_P3_P4_Lumbar_Coeliac_Sphenopalatine_VS_Stellate | 1 | 1 | 1 | 1 | 1 | 1 | 0 | 1 |
| ABCA8A | 1.06 | P4_VS_P1_P2_P3_Lumbar_Coeliac_Stellate_Sphenopalatine | 0 | 0 | 0 | 1 | 0 | 0 | 0 | 0 |
| EDNRB | 1.06 | P4_VS_P1_P2_P3_Lumbar_Coeliac_Stellate_Sphenopalatine | 0 | 0 | 0 | 1 | 0 | 0 | 0 | 0 |
| RTP1 | 1.06 | P4_VS_P1_P2_P3_Lumbar_Coeliac_Stellate_Sphenopalatine | 0 | 0 | 0 | 1 | 0 | 0 | 0 | 0 |
| GM49708 | 1.06 | P4_VS_P1_P2_P3_Lumbar_Coeliac_Stellate_Sphenopalatine | 0 | 0 | 0 | 1 | 0 | 0 | 0 | 0 |
| ACTA2 | 1.06 | P1_P2_P3_P4_VS_Lumbar_Coeliac_Stellate_Sphenopalatine | 1 | 1 | 1 | 1 | 0 | 0 | 0 | 0 |
| CEP350 | 1.06 | Lumbar_Coeliac_VS_P1_P2_P3_P4_Stellate_Sphenopalatine | 0 | 0 | 0 | 0 | 1 | 1 | 0 | 0 |
| 2310022B05I | 1.06 | Lumbar_Coeliac_VS_P1_P2_P3_P4_Stellate_Sphenopalatine | 0 | 0 | 0 | 0 | 1 | 1 | 0 | 0 |
| PREPL | 1.06 | Lumbar_Coeliac_VS_P1_P2_P3_P4_Stellate_Sphenopalatine | 0 | 0 | 0 | 0 | 1 | 1 | 0 | 0 |
| SLCO5A1 | 1.06 | P3_Lumbar_Coeliac_VS_P1_P2_P4_Stellate_Sphenopalatine | 0 | 0 | 1 | 0 | 1 | 1 | 0 | 0 |
| GM20721 | 1.06 | P1_P2_P3_P4_Lumbar_Coeliac_VS_Stellate_Sphenopalatine | 1 | 1 | 1 | 1 | 1 | 1 | 0 | 0 |
| SPHKAP | 1.06 | Sphenopalatine_VS_P1_P2_P3_P4_Lumbar_Coeliac_Stellate | 0 | 0 | 0 | 0 | 0 | 0 | 0 | 1 |
| LGR5 | 1.06 | Sphenopalatine_VS_P1_P2_P3_P4_Lumbar_Coeliac_Stellate | 0 | 0 | 0 | 0 | 0 | 0 | 0 | 1 |
| TOP1MT | 1.06 | P1_P2_P3_P4_Lumbar_Sphenopalatine_VS_Coeliac_Stellate | 1 | 1 | 1 | 1 | 1 | 0 | 0 | 1 |
| ZFYVE1 | 1.06 | P1_P2_P3_P4_Lumbar_Coeliac_Sphenopalatine_VS_Stellate | 1 | 1 | 1 | 1 | 1 | 1 | 0 | 1 |
| CBFA2T3 | 1.05 | P3_VS_P1_P2_P4_Lumbar_Coeliac_Stellate_Sphenopalatine | 0 | 0 | 1 | 0 | 0 | 0 | 0 | 0 |
| LRRC55 | 1.05 | P4_VS_P1_P2_P3_Lumbar_Coeliac_Stellate_Sphenopalatine | 0 | 0 | 0 | 1 | 0 | 0 | 0 | 0 |
| TTI1 | 1.05 | P4_VS_P1_P2_P3_Lumbar_Coeliac_Stellate_Sphenopalatine | 0 | 0 | 0 | 1 | 0 | 0 | 0 | 0 |
| CNGB1 | 1.05 | P4_VS_P1_P2_P3_Lumbar_Coeliac_Stellate_Sphenopalatine | 0 | 0 | 0 | 1 | 0 | 0 | 0 | 0 |
| SLC27A3 | 1.05 | Lumbar_VS_P1_P2_P3_P4_Coeliac_Stellate_Sphenopalatine | 0 | 0 | 0 | 0 | 1 | 0 | 0 | 0 |
| STK33 | 1.05 | Lumbar_VS_P1_P2_P3_P4_Coeliac_Stellate_Sphenopalatine | 0 | 0 | 0 | 0 | 1 | 0 | 0 | 0 |
| ATAD5 | 1.05 | Coeliac_VS_P1_P2_P3_P4_Lumbar_Stellate_Sphenopalatine | 0 | 0 | 0 | 0 | 0 | 1 | 0 | 0 |
| GM38393 | 1.05 | Lumbar_Coeliac_VS_P1_P2_P3_P4_Stellate_Sphenopalatine | 0 | 0 | 0 | 0 | 1 | 1 | 0 | 0 |
| PTK2B | 1.05 | P3_Lumbar_Coeliac_Stellate_VS_P1_P2_P4_Sphenopalatine | 0 | 0 | 1 | 0 | 1 | 1 | 1 | 0 |
| GCH1 | 1.05 | P1_P2_P3_P4_Lumbar_Coeliac_Stellate_VS_Sphenopalatine | 1 | 1 | 1 | 1 | 1 | 1 | 1 | 0 |
| KCNH2 | 1.05 | P1_P2_P3_P4_Lumbar_Coeliac_Sphenopalatine_VS_Stellate | 1 | 1 | 1 | 1 | 1 | 1 | 0 | 1 |

| | | | | | | | | | | |
|---|---|---|---|---|---|---|---|---|---|---|
| LMTK3 | 1.05 | P1_P2_P3_P4_Lumbar_Coeliac_Sphenopalatine_VS_Stellate | 1 | 1 | 1 | 1 | 1 | 1 | 0 | 1 |
| AY036118 | 1.05 | P1_P2_P3_P4_Lumbar_Coeliac_Sphenopalatine_VS_Stellate | 1 | 1 | 1 | 1 | 1 | 1 | 0 | 1 |
| FAM13B | 1.05 | P1_P2_P3_P4_Lumbar_Coeliac_Sphenopalatine_VS_Stellate | 1 | 1 | 1 | 1 | 1 | 1 | 0 | 1 |
| PLCE1 | 1.04 | P1_VS_P2_P3_P4_Lumbar_Coeliac_Stellate_Sphenopalatine | 1 | 0 | 0 | 0 | 0 | 0 | 0 | 0 |
| F2R | 1.04 | P2_VS_P1_P3_P4_Lumbar_Coeliac_Stellate_Sphenopalatine | 0 | 1 | 0 | 0 | 0 | 0 | 0 | 0 |
| STARD13 | 1.04 | P1_P3_VS_P2_P4_Lumbar_Coeliac_Stellate_Sphenopalatine | 1 | 0 | 1 | 0 | 0 | 0 | 0 | 0 |
| KNTC1 | 1.04 | P4_VS_P1_P2_P3_Lumbar_Coeliac_Stellate_Sphenopalatine | 0 | 0 | 0 | 1 | 0 | 0 | 0 | 0 |
| TTYH1 | 1.04 | P4_VS_P1_P2_P3_Lumbar_Coeliac_Stellate_Sphenopalatine | 0 | 0 | 0 | 1 | 0 | 0 | 0 | 0 |
| BDKRB2 | 1.04 | P4_VS_P1_P2_P3_Lumbar_Coeliac_Stellate_Sphenopalatine | 0 | 0 | 0 | 1 | 0 | 0 | 0 | 0 |
| PLCXD2 | 1.04 | P4_VS_P1_P2_P3_Lumbar_Coeliac_Stellate_Sphenopalatine | 0 | 0 | 0 | 1 | 0 | 0 | 0 | 0 |
| TIMELESS | 1.04 | Lumbar_VS_P1_P2_P3_P4_Coeliac_Stellate_Sphenopalatine | 0 | 0 | 0 | 0 | 1 | 0 | 0 | 0 |
| MEX3A | 1.04 | Lumbar_Coeliac_VS_P1_P2_P3_P4_Stellate_Sphenopalatine | 0 | 0 | 0 | 0 | 1 | 1 | 0 | 0 |
| FGD5 | 1.04 | Lumbar_Coeliac_VS_P1_P2_P3_P4_Stellate_Sphenopalatine | 0 | 0 | 0 | 0 | 1 | 1 | 0 | 0 |
| GM49700 | 1.04 | Lumbar_Coeliac_VS_P1_P2_P3_P4_Stellate_Sphenopalatine | 0 | 0 | 0 | 0 | 1 | 1 | 0 | 0 |
| E430024I08R | 1.04 | Lumbar_Coeliac_VS_P1_P2_P3_P4_Stellate_Sphenopalatine | 0 | 0 | 0 | 0 | 1 | 1 | 0 | 0 |
| TMEM74 | 1.04 | Lumbar_Coeliac_VS_P1_P2_P3_P4_Stellate_Sphenopalatine | 0 | 0 | 0 | 0 | 1 | 1 | 0 | 0 |
| PCDHA11.1 | 1.04 | Lumbar_Coeliac_VS_P1_P2_P3_P4_Stellate_Sphenopalatine | 0 | 0 | 0 | 0 | 1 | 1 | 0 | 0 |
| SOCS3 | 1.04 | P1_Sphenopalatine_VS_P2_P3_P4_Lumbar_Coeliac_Stellate | 1 | 0 | 0 | 0 | 0 | 0 | 0 | 1 |
| LPAR1 | 1.04 | P4_Sphenopalatine_VS_P1_P2_P3_Lumbar_Coeliac_Stellate | 0 | 0 | 0 | 1 | 0 | 0 | 0 | 1 |
| RAPGEF4 | 1.04 | P1_P2_P3_P4_Lumbar_Coeliac_Sphenopalatine_VS_Stellate | 1 | 1 | 1 | 1 | 1 | 1 | 0 | 1 |
| KCTD2 | 1.04 | P1_P2_P3_P4_Lumbar_Coeliac_Sphenopalatine_VS_Stellate | 1 | 1 | 1 | 1 | 1 | 1 | 0 | 1 |
| UTY | 1.04 | P1_P2_P3_P4_Lumbar_Coeliac_Sphenopalatine_VS_Stellate | 1 | 1 | 1 | 1 | 1 | 1 | 0 | 1 |
| FBXO2 | 1.04 | P1_P2_P3_P4_Stellate_Sphenopalatine_VS_Lumbar_Coeliac | 1 | 1 | 1 | 1 | 0 | 0 | 1 | 1 |
| KCNH1 | 1.04 | P1_P2_P3_Lumbar_Coeliac_Stellate_Sphenopalatine_VS_P4 | 1 | 1 | 1 | 0 | 1 | 1 | 1 | 1 |
| VSIG8 | 1.04 | P3_P4_Lumbar_Coeliac_Stellate_Sphenopalatine_VS_P1_P2 | 0 | 0 | 1 | 1 | 1 | 1 | 1 | 1 |
| SERPINA3G | 1.03 | P1_P2_VS_P3_P4_Lumbar_Coeliac_Stellate_Sphenopalatine | 1 | 1 | 0 | 0 | 0 | 0 | 0 | 0 |
| SSTR2 | 1.03 | P3_VS_P1_P2_P4_Lumbar_Coeliac_Stellate_Sphenopalatine | 0 | 0 | 1 | 0 | 0 | 0 | 0 | 0 |
| ENC1 | 1.03 | P3_VS_P1_P2_P4_Lumbar_Coeliac_Stellate_Sphenopalatine | 0 | 0 | 1 | 0 | 0 | 0 | 0 | 0 |
| OLFML2A | 1.03 | P4_VS_P1_P2_P3_Lumbar_Coeliac_Stellate_Sphenopalatine | 0 | 0 | 0 | 1 | 0 | 0 | 0 | 0 |
| CALN1 | 1.03 | P4_VS_P1_P2_P3_Lumbar_Coeliac_Stellate_Sphenopalatine | 0 | 0 | 0 | 1 | 0 | 0 | 0 | 0 |

| | | | | | | | | | | |
|---|---|---|---|---|---|---|---|---|---|---|
| COL26A1 | 1.03 | P4_VS_P1_P2_P3_Lumbar_Coeliac_Stellate_Sphenopalatine | 0 | 0 | 0 | 1 | 0 | 0 | 0 | 0 |
| GAB1 | 1.03 | P4_VS_P1_P2_P3_Lumbar_Coeliac_Stellate_Sphenopalatine | 0 | 0 | 0 | 1 | 0 | 0 | 0 | 0 |
| AMIGO2 | 1.03 | P4_VS_P1_P2_P3_Lumbar_Coeliac_Stellate_Sphenopalatine | 0 | 0 | 0 | 1 | 0 | 0 | 0 | 0 |
| NECTIN3 | 1.03 | P4_VS_P1_P2_P3_Lumbar_Coeliac_Stellate_Sphenopalatine | 0 | 0 | 0 | 1 | 0 | 0 | 0 | 0 |
| ABTB2 | 1.03 | P3_Lumbar_VS_P1_P2_P4_Coeliac_Stellate_Sphenopalatine | 0 | 0 | 1 | 0 | 1 | 0 | 0 | 0 |
| HR | 1.03 | Lumbar_Coeliac_VS_P1_P2_P3_P4_Stellate_Sphenopalatine | 0 | 0 | 0 | 0 | 1 | 1 | 0 | 0 |
| RTN4R | 1.03 | P3_P4_Lumbar_Coeliac_VS_P1_P2_Stellate_Sphenopalatine | 0 | 0 | 1 | 1 | 1 | 1 | 0 | 0 |
| CENPV | 1.03 | P1_P2_P3_P4_Lumbar_Coeliac_VS_Stellate_Sphenopalatine | 1 | 1 | 1 | 1 | 1 | 1 | 0 | 0 |
| PVR | 1.03 | P1_P2_P4_Sphenopalatine_VS_P3_Lumbar_Coeliac_Stellate | 1 | 1 | 0 | 1 | 0 | 0 | 0 | 1 |
| ITM2A | 1.03 | P1_P3_P4_Sphenopalatine_VS_P2_Lumbar_Coeliac_Stellate | 1 | 0 | 1 | 1 | 0 | 0 | 0 | 1 |
| GM47163 | 1.03 | P1_P2_P3_P4_Lumbar_Coeliac_Sphenopalatine_VS_Stellate | 1 | 1 | 1 | 1 | 1 | 1 | 0 | 1 |
| NHSL2 | 1.03 | P1_P2_P3_P4_Lumbar_Coeliac_Sphenopalatine_VS_Stellate | 1 | 1 | 1 | 1 | 1 | 1 | 0 | 1 |
| IFITM10 | 1.03 | P1_P2_P3_Lumbar_Coeliac_Stellate_Sphenopalatine_VS_P4 | 1 | 1 | 1 | 0 | 1 | 1 | 1 | 1 |
| NFATC2 | 1.02 | P3_VS_P1_P2_P4_Lumbar_Coeliac_Stellate_Sphenopalatine | 0 | 0 | 1 | 0 | 0 | 0 | 0 | 0 |
| GM11110 | 1.02 | P4_VS_P1_P2_P3_Lumbar_Coeliac_Stellate_Sphenopalatine | 0 | 0 | 0 | 1 | 0 | 0 | 0 | 0 |
| BC002189 | 1.02 | Lumbar_VS_P1_P2_P3_P4_Coeliac_Stellate_Sphenopalatine | 0 | 0 | 0 | 0 | 1 | 0 | 0 | 0 |
| PTPRM | 1.02 | Lumbar_VS_P1_P2_P3_P4_Coeliac_Stellate_Sphenopalatine | 0 | 0 | 0 | 0 | 1 | 0 | 0 | 0 |
| ASIC1 | 1.02 | Coeliac_VS_P1_P2_P3_P4_Lumbar_Stellate_Sphenopalatine | 0 | 0 | 0 | 0 | 0 | 1 | 0 | 0 |
| TMEM196 | 1.02 | P4_Coeliac_VS_P1_P2_P3_Lumbar_Stellate_Sphenopalatine | 0 | 0 | 0 | 1 | 0 | 1 | 0 | 0 |
| CTBP2 | 1.02 | Lumbar_Coeliac_VS_P1_P2_P3_P4_Stellate_Sphenopalatine | 0 | 0 | 0 | 0 | 1 | 1 | 0 | 0 |
| MTA3 | 1.02 | Lumbar_Coeliac_VS_P1_P2_P3_P4_Stellate_Sphenopalatine | 0 | 0 | 0 | 0 | 1 | 1 | 0 | 0 |
| GM43111 | 1.02 | P4_Lumbar_Coeliac_VS_P1_P2_P3_Stellate_Sphenopalatine | 0 | 0 | 0 | 1 | 1 | 1 | 0 | 0 |
| CABLES1 | 1.02 | P1_P4_Lumbar_Coeliac_VS_P2_P3_Stellate_Sphenopalatine | 1 | 0 | 0 | 1 | 1 | 1 | 0 | 0 |
| SHC3 | 1.02 | P1_P2_P3_P4_Lumbar_Coeliac_VS_Stellate_Sphenopalatine | 1 | 1 | 1 | 1 | 1 | 1 | 0 | 0 |
| TMEM132C | 1.02 | P1_P2_P3_P4_Lumbar_Coeliac_Stellate_VS_Sphenopalatine | 1 | 1 | 1 | 1 | 1 | 1 | 1 | 0 |
| NXN | 1.02 | P1_P2_P3_P4_Lumbar_Coeliac_Stellate_VS_Sphenopalatine | 1 | 1 | 1 | 1 | 1 | 1 | 1 | 0 |
| SLC44A5 | 1.02 | P1_P3_P4_Lumbar_Coeliac_Sphenopalatine_VS_P2_Stellate | 1 | 0 | 1 | 1 | 1 | 1 | 0 | 1 |
| NOS1AP | 1.02 | P1_P2_P3_P4_Lumbar_Coeliac_Sphenopalatine_VS_Stellate | 1 | 1 | 1 | 1 | 1 | 1 | 0 | 1 |
| IFFO2 | 1.02 | P1_P2_P3_P4_Lumbar_Coeliac_Sphenopalatine_VS_Stellate | 1 | 1 | 1 | 1 | 1 | 1 | 0 | 1 |
| A330076H08 | 1.02 | P1_P2_P3_P4_Lumbar_Coeliac_Sphenopalatine_VS_Stellate | 1 | 1 | 1 | 1 | 1 | 1 | 0 | 1 |

| | | | | | | | | | | |
|---|---|---|---|---|---|---|---|---|---|---|
| DLGAP2 | 1.02 | P1_P2_P3_P4_Lumbar_Coeliac_Sphenopalatine_VS_Stellate | 1 | 1 | 1 | 1 | 1 | 1 | 0 | 1 |
| SACS | 1.02 | P1_P2_P3_P4_Lumbar_Coeliac_Sphenopalatine_VS_Stellate | 1 | 1 | 1 | 1 | 1 | 1 | 0 | 1 |
| EP300 | 1.02 | P1_P2_P3_P4_Lumbar_Coeliac_Sphenopalatine_VS_Stellate | 1 | 1 | 1 | 1 | 1 | 1 | 0 | 1 |
| SMIM18 | 1.02 | P1_P4_Lumbar_Coeliac_Stellate_Sphenopalatine_VS_P2_P3 | 1 | 0 | 0 | 1 | 1 | 1 | 1 | 1 |
| A530058N18 | 1.02 | P1_P3_P4_Lumbar_Coeliac_Stellate_Sphenopalatine_VS_P2 | 1 | 0 | 1 | 1 | 1 | 1 | 1 | 1 |
| GALNT5 | 1.01 | P1_VS_P2_P3_P4_Lumbar_Coeliac_Stellate_Sphenopalatine | 1 | 0 | 0 | 0 | 0 | 0 | 0 | 0 |
| LGALS9 | 1.01 | P1_P2_VS_P3_P4_Lumbar_Coeliac_Stellate_Sphenopalatine | 1 | 1 | 0 | 0 | 0 | 0 | 0 | 0 |
| EPHB6 | 1.01 | P3_VS_P1_P2_P4_Lumbar_Coeliac_Stellate_Sphenopalatine | 0 | 0 | 1 | 0 | 0 | 0 | 0 | 0 |
| FAM92B | 1.01 | P4_VS_P1_P2_P3_Lumbar_Coeliac_Stellate_Sphenopalatine | 0 | 0 | 0 | 1 | 0 | 0 | 0 | 0 |
| 1700016P03I | 1.01 | P4_VS_P1_P2_P3_Lumbar_Coeliac_Stellate_Sphenopalatine | 0 | 0 | 0 | 1 | 0 | 0 | 0 | 0 |
| CDH18 | 1.01 | P4_VS_P1_P2_P3_Lumbar_Coeliac_Stellate_Sphenopalatine | 0 | 0 | 0 | 1 | 0 | 0 | 0 | 0 |
| CATSPERZ | 1.01 | P4_VS_P1_P2_P3_Lumbar_Coeliac_Stellate_Sphenopalatine | 0 | 0 | 0 | 1 | 0 | 0 | 0 | 0 |
| PLAAT3 | 1.01 | P4_VS_P1_P2_P3_Lumbar_Coeliac_Stellate_Sphenopalatine | 0 | 0 | 0 | 1 | 0 | 0 | 0 | 0 |
| G2E3 | 1.01 | Coeliac_VS_P1_P2_P3_P4_Lumbar_Stellate_Sphenopalatine | 0 | 0 | 0 | 0 | 0 | 1 | 0 | 0 |
| SORCS2 | 1.01 | Lumbar_Coeliac_VS_P1_P2_P3_P4_Stellate_Sphenopalatine | 0 | 0 | 0 | 0 | 1 | 1 | 0 | 0 |
| GM9801 | 1.01 | Lumbar_Coeliac_VS_P1_P2_P3_P4_Stellate_Sphenopalatine | 0 | 0 | 0 | 0 | 1 | 1 | 0 | 0 |
| ZFP711 | 1.01 | Lumbar_Coeliac_VS_P1_P2_P3_P4_Stellate_Sphenopalatine | 0 | 0 | 0 | 0 | 1 | 1 | 0 | 0 |
| NGB | 1.01 | P3_P4_Lumbar_Coeliac_Stellate_VS_P1_P2_Sphenopalatine | 0 | 0 | 1 | 1 | 1 | 1 | 1 | 0 |
| ADRA2A | 1.01 | P1_P2_P3_P4_Lumbar_Coeliac_Stellate_VS_Sphenopalatine | 1 | 1 | 1 | 1 | 1 | 1 | 1 | 0 |
| TRANK1 | 1.01 | Sphenopalatine_VS_P1_P2_P3_P4_Lumbar_Coeliac_Stellate | 0 | 0 | 0 | 0 | 0 | 0 | 0 | 1 |
| SEL1L3 | 1.01 | P1_P2_Sphenopalatine_VS_P3_P4_Lumbar_Coeliac_Stellate | 1 | 1 | 0 | 0 | 0 | 0 | 0 | 1 |
| PCDH17 | 1.01 | P1_P2_P3_P4_Lumbar_Sphenopalatine_VS_Coeliac_Stellate | 1 | 1 | 1 | 1 | 1 | 0 | 0 | 1 |
| LIX1L | 1.01 | P1_P2_P3_P4_Lumbar_Coeliac_Sphenopalatine_VS_Stellate | 1 | 1 | 1 | 1 | 1 | 1 | 0 | 1 |
| SPEN | 1.01 | P1_P2_P3_P4_Lumbar_Coeliac_Sphenopalatine_VS_Stellate | 1 | 1 | 1 | 1 | 1 | 1 | 0 | 1 |
| A230057D06 | 1.01 | P1_P2_P3_P4_Lumbar_Coeliac_Sphenopalatine_VS_Stellate | 1 | 1 | 1 | 1 | 1 | 1 | 0 | 1 |
| MCU | 1.01 | P1_P2_P3_P4_Lumbar_Coeliac_Sphenopalatine_VS_Stellate | 1 | 1 | 1 | 1 | 1 | 1 | 0 | 1 |
| GGTA1 | 1.01 | P1_P2_P3_Lumbar_Coeliac_Stellate_Sphenopalatine_VS_P4 | 1 | 1 | 1 | 0 | 1 | 1 | 1 | 1 |
| GSTM6 | 1.01 | P1_P2_P3_Lumbar_Coeliac_Stellate_Sphenopalatine_VS_P4 | 1 | 1 | 1 | 0 | 1 | 1 | 1 | 1 |
| STARD5 | 1 | P3_VS_P1_P2_P4_Lumbar_Coeliac_Stellate_Sphenopalatine | 0 | 0 | 1 | 0 | 0 | 0 | 0 | 0 |
| FKBP11 | 1 | P3_VS_P1_P2_P4_Lumbar_Coeliac_Stellate_Sphenopalatine | 0 | 0 | 1 | 0 | 0 | 0 | 0 | 0 |

| | | | | | | | | | | |
|---|---|---|---|---|---|---|---|---|---|---|
| NR2F2 | 1 | P4_VS_P1_P2_P3_Lumbar_Coeliac_Stellate_Sphenopalatine | 0 | 0 | 0 | 1 | 0 | 0 | 0 | 0 |
| TMEM71 | 1 | P1_P2_P4_VS_P3_Lumbar_Coeliac_Stellate_Sphenopalatine | 1 | 1 | 0 | 1 | 0 | 0 | 0 | 0 |
| SAP30L | 1 | P1_P2_P3_P4_Lumbar_Coeliac_VS_Stellate_Sphenopalatine | 1 | 1 | 1 | 1 | 1 | 1 | 0 | 0 |
| SNTG1 | 1 | P3_P4_Sphenopalatine_VS_P1_P2_Lumbar_Coeliac_Stellate | 0 | 0 | 1 | 1 | 0 | 0 | 0 | 1 |
| COL1A1 | 1 | P1_P2_P3_P4_Lumbar_Coeliac_Sphenopalatine_VS_Stellate | 1 | 1 | 1 | 1 | 1 | 1 | 0 | 1 |
| EFNB2 | 1 | P1_P2_P4_Lumbar_Coeliac_Stellate_Sphenopalatine_VS_P3 | 1 | 1 | 0 | 1 | 1 | 1 | 1 | 1 |
| PDE2A | 1 | P1_P3_P4_Lumbar_Coeliac_Stellate_Sphenopalatine_VS_P2 | 1 | 0 | 1 | 1 | 1 | 1 | 1 | 1 |
| GM20751 | 0.999 | P4_VS_P1_P2_P3_Lumbar_Coeliac_Stellate_Sphenopalatine | 0 | 0 | 0 | 1 | 0 | 0 | 0 | 0 |
| NRP2 | 0.999 | P1_P2_P3_P4_Lumbar_Coeliac_Stellate_VS_Sphenopalatine | 1 | 1 | 1 | 1 | 1 | 1 | 1 | 0 |
| GPR155 | 0.999 | P1_P2_P3_P4_Lumbar_Coeliac_Sphenopalatine_VS_Stellate | 1 | 1 | 1 | 1 | 1 | 1 | 0 | 1 |
| TJP2 | 0.998 | P4_VS_P1_P2_P3_Lumbar_Coeliac_Stellate_Sphenopalatine | 0 | 0 | 0 | 1 | 0 | 0 | 0 | 0 |
| SUSD4 | 0.998 | P1_P2_P3_P4_Lumbar_Coeliac_Sphenopalatine_VS_Stellate | 1 | 1 | 1 | 1 | 1 | 1 | 0 | 1 |
| ZFP458 | 0.997 | Lumbar_Coeliac_VS_P1_P2_P3_P4_Stellate_Sphenopalatine | 0 | 0 | 0 | 0 | 1 | 1 | 0 | 0 |
| CILP | 0.995 | P3_Coeliac_VS_P1_P2_P4_Lumbar_Stellate_Sphenopalatine | 0 | 0 | 1 | 0 | 0 | 1 | 0 | 0 |
| AU041133 | 0.995 | Lumbar_Coeliac_VS_P1_P2_P3_P4_Stellate_Sphenopalatine | 0 | 0 | 0 | 0 | 1 | 1 | 0 | 0 |
| CABP1 | 0.995 | P1_P2_P3_P4_Lumbar_Coeliac_VS_Stellate_Sphenopalatine | 1 | 1 | 1 | 1 | 1 | 1 | 0 | 0 |
| PTPRD | 0.995 | P1_P2_P4_Lumbar_Coeliac_Sphenopalatine_VS_P3_Stellate | 1 | 1 | 0 | 1 | 1 | 1 | 0 | 1 |
| FAM131C | 0.994 | Lumbar_VS_P1_P2_P3_P4_Coeliac_Stellate_Sphenopalatine | 0 | 0 | 0 | 0 | 1 | 0 | 0 | 0 |
| CAMK1D | 0.993 | P1_P2_P3_P4_Lumbar_Coeliac_Sphenopalatine_VS_Stellate | 1 | 1 | 1 | 1 | 1 | 1 | 0 | 1 |
| LARS2 | 0.993 | P1_P2_P3_P4_Lumbar_Coeliac_Sphenopalatine_VS_Stellate | 1 | 1 | 1 | 1 | 1 | 1 | 0 | 1 |
| SERHL | 0.992 | P2_VS_P1_P3_P4_Lumbar_Coeliac_Stellate_Sphenopalatine | 0 | 1 | 0 | 0 | 0 | 0 | 0 | 0 |
| AGRP | 0.992 | Lumbar_VS_P1_P2_P3_P4_Coeliac_Stellate_Sphenopalatine | 0 | 0 | 0 | 0 | 1 | 0 | 0 | 0 |
| GPRIN3 | 0.992 | Coeliac_VS_P1_P2_P3_P4_Lumbar_Stellate_Sphenopalatine | 0 | 0 | 0 | 0 | 0 | 1 | 0 | 0 |
| FAM184B | 0.992 | Lumbar_Coeliac_VS_P1_P2_P3_P4_Stellate_Sphenopalatine | 0 | 0 | 0 | 0 | 1 | 1 | 0 | 0 |
| RPS6KA2 | 0.991 | Lumbar_Coeliac_VS_P1_P2_P3_P4_Stellate_Sphenopalatine | 0 | 0 | 0 | 0 | 1 | 1 | 0 | 0 |
| WDR25 | 0.991 | Sphenopalatine_VS_P1_P2_P3_P4_Lumbar_Coeliac_Stellate | 0 | 0 | 0 | 0 | 0 | 0 | 0 | 1 |
| TMED8 | 0.99 | P4_VS_P1_P2_P3_Lumbar_Coeliac_Stellate_Sphenopalatine | 0 | 0 | 0 | 1 | 0 | 0 | 0 | 0 |
| JRKL | 0.99 | Lumbar_Coeliac_VS_P1_P2_P3_P4_Stellate_Sphenopalatine | 0 | 0 | 0 | 0 | 1 | 1 | 0 | 0 |
| HIVEP2 | 0.99 | P1_P2_P3_P4_Lumbar_Coeliac_Sphenopalatine_VS_Stellate | 1 | 1 | 1 | 1 | 1 | 1 | 0 | 1 |
| SRD5A1 | 0.99 | P1_P2_P3_P4_Lumbar_Coeliac_Sphenopalatine_VS_Stellate | 1 | 1 | 1 | 1 | 1 | 1 | 0 | 1 |

| | | | | | | | | | | |
|---|---|---|---|---|---|---|---|---|---|---|
| SLC7A3 | 0.99 | P1_P2_P3_P4_Lumbar_Coeliac_Sphenopalatine_VS_Stellate | 1 | 1 | 1 | 1 | 1 | 1 | 0 | 1 |
| DPP10 | 0.989 | P1_P4_Lumbar_VS_P2_P3_Coeliac_Stellate_Sphenopalatine | 1 | 0 | 0 | 1 | 1 | 0 | 0 | 0 |
| RBM46 | 0.988 | P4_VS_P1_P2_P3_Lumbar_Coeliac_Stellate_Sphenopalatine | 0 | 0 | 0 | 1 | 0 | 0 | 0 | 0 |
| NFIB | 0.988 | P1_P2_P3_P4_VS_Lumbar_Coeliac_Stellate_Sphenopalatine | 1 | 1 | 1 | 1 | 0 | 0 | 0 | 0 |
| ELMO1 | 0.988 | P1_P2_P3_P4_Sphenopalatine_VS_Lumbar_Coeliac_Stellate | 1 | 1 | 1 | 1 | 0 | 0 | 0 | 1 |
| FOSL2 | 0.985 | P4_Sphenopalatine_VS_P1_P2_P3_Lumbar_Coeliac_Stellate | 0 | 0 | 0 | 1 | 0 | 0 | 0 | 1 |
| MPZ | 0.984 | P4_VS_P1_P2_P3_Lumbar_Coeliac_Stellate_Sphenopalatine | 0 | 0 | 0 | 1 | 0 | 0 | 0 | 0 |
| PLEKHA4 | 0.984 | P4_VS_P1_P2_P3_Lumbar_Coeliac_Stellate_Sphenopalatine | 0 | 0 | 0 | 1 | 0 | 0 | 0 | 0 |
| ABHD4 | 0.984 | P4_VS_P1_P2_P3_Lumbar_Coeliac_Stellate_Sphenopalatine | 0 | 0 | 0 | 1 | 0 | 0 | 0 | 0 |
| ABCB1A | 0.983 | P4_Lumbar_Coeliac_VS_P1_P2_P3_Stellate_Sphenopalatine | 0 | 0 | 0 | 1 | 1 | 1 | 0 | 0 |
| CRISPLD2 | 0.983 | Sphenopalatine_VS_P1_P2_P3_P4_Lumbar_Coeliac_Stellate | 0 | 0 | 0 | 0 | 0 | 0 | 0 | 1 |
| ABCA8B | 0.983 | P1_P2_P3_P4_Lumbar_Coeliac_Sphenopalatine_VS_Stellate | 1 | 1 | 1 | 1 | 1 | 1 | 0 | 1 |
| YTHDC2 | 0.983 | P1_P2_P3_P4_Lumbar_Coeliac_Sphenopalatine_VS_Stellate | 1 | 1 | 1 | 1 | 1 | 1 | 0 | 1 |
| PRPS2 | 0.981 | P3_VS_P1_P2_P4_Lumbar_Coeliac_Stellate_Sphenopalatine | 0 | 0 | 1 | 0 | 0 | 0 | 0 | 0 |
| TM4SF4 | 0.98 | P1_P3_P4_Lumbar_Coeliac_Stellate_VS_P2_Sphenopalatine | 1 | 0 | 1 | 1 | 1 | 1 | 1 | 0 |
| B230209E15I | 0.98 | P1_P2_P3_P4_Lumbar_Coeliac_Sphenopalatine_VS_Stellate | 1 | 1 | 1 | 1 | 1 | 1 | 0 | 1 |
| ADAMTS1 | 0.978 | P4_VS_P1_P2_P3_Lumbar_Coeliac_Stellate_Sphenopalatine | 0 | 0 | 0 | 1 | 0 | 0 | 0 | 0 |
| PTPRJ | 0.973 | P3_VS_P1_P2_P4_Lumbar_Coeliac_Stellate_Sphenopalatine | 0 | 0 | 1 | 0 | 0 | 0 | 0 | 0 |
| ITPR1 | 0.973 | P3_P4_VS_P1_P2_Lumbar_Coeliac_Stellate_Sphenopalatine | 0 | 0 | 1 | 1 | 0 | 0 | 0 | 0 |
| MC4R | 0.972 | P4_VS_P1_P2_P3_Lumbar_Coeliac_Stellate_Sphenopalatine | 0 | 0 | 0 | 1 | 0 | 0 | 0 | 0 |
| RAB11FIP3 | 0.972 | P1_P2_P3_P4_Lumbar_Coeliac_Sphenopalatine_VS_Stellate | 1 | 1 | 1 | 1 | 1 | 1 | 0 | 1 |
| NOTCH2 | 0.971 | P4_VS_P1_P2_P3_Lumbar_Coeliac_Stellate_Sphenopalatine | 0 | 0 | 0 | 1 | 0 | 0 | 0 | 0 |
| PTPN13 | 0.971 | P4_VS_P1_P2_P3_Lumbar_Coeliac_Stellate_Sphenopalatine | 0 | 0 | 0 | 1 | 0 | 0 | 0 | 0 |
| GM38593 | 0.971 | P4_VS_P1_P2_P3_Lumbar_Coeliac_Stellate_Sphenopalatine | 0 | 0 | 0 | 1 | 0 | 0 | 0 | 0 |
| GM9866 | 0.971 | P1_P2_P4_Lumbar_Coeliac_Stellate_Sphenopalatine_VS_P3 | 1 | 1 | 0 | 1 | 1 | 1 | 1 | 1 |
| 2310001H17I | 0.97 | Sphenopalatine_VS_P1_P2_P3_P4_Lumbar_Coeliac_Stellate | 0 | 0 | 0 | 0 | 0 | 0 | 0 | 1 |
| FHOD3 | 0.97 | P1_P2_P3_P4_Lumbar_Coeliac_Sphenopalatine_VS_Stellate | 1 | 1 | 1 | 1 | 1 | 1 | 0 | 1 |
| KCNS2 | 0.969 | Lumbar_VS_P1_P2_P3_P4_Coeliac_Stellate_Sphenopalatine | 0 | 0 | 0 | 0 | 1 | 0 | 0 | 0 |
| SRCAP | 0.969 | Lumbar_Coeliac_VS_P1_P2_P3_P4_Stellate_Sphenopalatine | 0 | 0 | 0 | 0 | 1 | 1 | 0 | 0 |
| RAB27A | 0.969 | P3_P4_Lumbar_Coeliac_VS_P1_P2_Stellate_Sphenopalatine | 0 | 0 | 1 | 1 | 1 | 1 | 0 | 0 |

| | | | | | | | | | | |
|---|---|---|---|---|---|---|---|---|---|---|
| PQLC3 | 0.968 | P4_VS_P1_P2_P3_Lumbar_Coeliac_Stellate_Sphenopalatine | 0 | 0 | 0 | 1 | 0 | 0 | 0 | 0 |
| FBLN5 | 0.968 | P4_VS_P1_P2_P3_Lumbar_Coeliac_Stellate_Sphenopalatine | 0 | 0 | 0 | 1 | 0 | 0 | 0 | 0 |
| S100A1 | 0.967 | P1_P2_P3_P4_Lumbar_Coeliac_Stellate_VS_Sphenopalatine | 1 | 1 | 1 | 1 | 1 | 1 | 1 | 0 |
| MAGI2 | 0.967 | P1_P2_P3_P4_Lumbar_Coeliac_Sphenopalatine_VS_Stellate | 1 | 1 | 1 | 1 | 1 | 1 | 0 | 1 |
| KCNJ12 | 0.967 | P1_P2_P3_P4_Lumbar_Coeliac_Sphenopalatine_VS_Stellate | 1 | 1 | 1 | 1 | 1 | 1 | 0 | 1 |
| SHANK3 | 0.965 | Lumbar_Coeliac_VS_P1_P2_P3_P4_Stellate_Sphenopalatine | 0 | 0 | 0 | 0 | 1 | 1 | 0 | 0 |
| GLCE | 0.963 | P2_VS_P1_P3_P4_Lumbar_Coeliac_Stellate_Sphenopalatine | 0 | 1 | 0 | 0 | 0 | 0 | 0 | 0 |
| SKAP2 | 0.963 | P4_VS_P1_P2_P3_Lumbar_Coeliac_Stellate_Sphenopalatine | 0 | 0 | 0 | 1 | 0 | 0 | 0 | 0 |
| IL7 | 0.962 | P4_VS_P1_P2_P3_Lumbar_Coeliac_Stellate_Sphenopalatine | 0 | 0 | 0 | 1 | 0 | 0 | 0 | 0 |
| BRWD3 | 0.962 | Lumbar_Coeliac_VS_P1_P2_P3_P4_Stellate_Sphenopalatine | 0 | 0 | 0 | 0 | 1 | 1 | 0 | 0 |
| PCDH9 | 0.962 | P1_P2_P3_Lumbar_Coeliac_Stellate_Sphenopalatine_VS_P4 | 1 | 1 | 1 | 0 | 1 | 1 | 1 | 1 |
| P4HA3 | 0.961 | P1_P2_P3_P4_Lumbar_Coeliac_Sphenopalatine_VS_Stellate | 1 | 1 | 1 | 1 | 1 | 1 | 0 | 1 |
| LEMD2 | 0.961 | P1_P2_P3_P4_Lumbar_Coeliac_Sphenopalatine_VS_Stellate | 1 | 1 | 1 | 1 | 1 | 1 | 0 | 1 |
| ARHGAP24 | 0.96 | P4_VS_P1_P2_P3_Lumbar_Coeliac_Stellate_Sphenopalatine | 0 | 0 | 0 | 1 | 0 | 0 | 0 | 0 |
| MMD2 | 0.96 | P4_VS_P1_P2_P3_Lumbar_Coeliac_Stellate_Sphenopalatine | 0 | 0 | 0 | 1 | 0 | 0 | 0 | 0 |
| GM26532 | 0.96 | P1_P2_P3_P4_Lumbar_Coeliac_Sphenopalatine_VS_Stellate | 1 | 1 | 1 | 1 | 1 | 1 | 0 | 1 |
| 3010003L21I | 0.959 | Lumbar_VS_P1_P2_P3_P4_Coeliac_Stellate_Sphenopalatine | 0 | 0 | 0 | 0 | 1 | 0 | 0 | 0 |
| PLCL1 | 0.958 | Sphenopalatine_VS_P1_P2_P3_P4_Lumbar_Coeliac_Stellate | 0 | 0 | 0 | 0 | 0 | 0 | 0 | 1 |
| LIN7A | 0.958 | P1_P3_P4_Coeliac_Sphenopalatine_VS_P2_Lumbar_Stellate | 1 | 0 | 1 | 1 | 0 | 1 | 0 | 1 |
| KCTD6 | 0.958 | P1_P2_P3_P4_Lumbar_Coeliac_Sphenopalatine_VS_Stellate | 1 | 1 | 1 | 1 | 1 | 1 | 0 | 1 |
| EFCAB1 | 0.956 | P3_P4_Lumbar_Coeliac_VS_P1_P2_Stellate_Sphenopalatine | 0 | 0 | 1 | 1 | 1 | 1 | 0 | 0 |
| HECW2 | 0.953 | P1_P2_P3_P4_Lumbar_Coeliac_Sphenopalatine_VS_Stellate | 1 | 1 | 1 | 1 | 1 | 1 | 0 | 1 |
| PEAK1 | 0.952 | P1_P2_P3_P4_Lumbar_Coeliac_Sphenopalatine_VS_Stellate | 1 | 1 | 1 | 1 | 1 | 1 | 0 | 1 |
| GM5165 | 0.951 | Coeliac_VS_P1_P2_P3_P4_Lumbar_Stellate_Sphenopalatine | 0 | 0 | 0 | 0 | 0 | 1 | 0 | 0 |
| EFNB3 | 0.951 | P3_Lumbar_Coeliac_VS_P1_P2_P4_Stellate_Sphenopalatine | 0 | 0 | 1 | 0 | 1 | 1 | 0 | 0 |
| BCORL1 | 0.949 | P1_P2_P3_P4_Lumbar_Coeliac_Sphenopalatine_VS_Stellate | 1 | 1 | 1 | 1 | 1 | 1 | 0 | 1 |
| PLCH1 | 0.948 | P4_VS_P1_P2_P3_Lumbar_Coeliac_Stellate_Sphenopalatine | 0 | 0 | 0 | 1 | 0 | 0 | 0 | 0 |
| ATG9B | 0.948 | P4_VS_P1_P2_P3_Lumbar_Coeliac_Stellate_Sphenopalatine | 0 | 0 | 0 | 1 | 0 | 0 | 0 | 0 |
| RNASET2B | 0.948 | Coeliac_VS_P1_P2_P3_P4_Lumbar_Stellate_Sphenopalatine | 0 | 0 | 0 | 0 | 0 | 1 | 0 | 0 |
| CRIM1 | 0.947 | P1_VS_P2_P3_P4_Lumbar_Coeliac_Stellate_Sphenopalatine | 1 | 0 | 0 | 0 | 0 | 0 | 0 | 0 |

| | | | | | | | | | | |
|---|---|---|---|---|---|---|---|---|---|---|
| GM30382 | 0.947 | P1_P2_VS_P3_P4_Lumbar_Coeliac_Stellate_Sphenopalatine | 1 | 1 | 0 | 0 | 0 | 0 | 0 | 0 |
| SSC5D | 0.947 | P2_Sphenopalatine_VS_P1_P3_P4_Lumbar_Coeliac_Stellate | 0 | 1 | 0 | 0 | 0 | 0 | 0 | 1 |
| LARP1B | 0.947 | P1_P2_P3_P4_Lumbar_Coeliac_Sphenopalatine_VS_Stellate | 1 | 1 | 1 | 1 | 1 | 1 | 0 | 1 |
| PPP1R12C | 0.947 | P1_P2_P3_P4_Lumbar_Coeliac_Sphenopalatine_VS_Stellate | 1 | 1 | 1 | 1 | 1 | 1 | 0 | 1 |
| TMEM91 | 0.945 | P3_P4_Lumbar_Coeliac_Stellate_Sphenopalatine_VS_P1_P2 | 0 | 0 | 1 | 1 | 1 | 1 | 1 | 1 |
| SYT12 | 0.944 | P1_VS_P2_P3_P4_Lumbar_Coeliac_Stellate_Sphenopalatine | 1 | 0 | 0 | 0 | 0 | 0 | 0 | 0 |
| PAKAP | 0.944 | Lumbar_VS_P1_P2_P3_P4_Coeliac_Stellate_Sphenopalatine | 0 | 0 | 0 | 0 | 1 | 0 | 0 | 0 |
| CRYM | 0.944 | P4_Lumbar_Coeliac_Stellate_VS_P1_P2_P3_Sphenopalatine | 0 | 0 | 0 | 1 | 1 | 1 | 1 | 0 |
| SEZ6L | 0.944 | P1_P2_P3_P4_Lumbar_Coeliac_Sphenopalatine_VS_Stellate | 1 | 1 | 1 | 1 | 1 | 1 | 0 | 1 |
| CLIP4 | 0.944 | P1_P2_P3_P4_Lumbar_Coeliac_Sphenopalatine_VS_Stellate | 1 | 1 | 1 | 1 | 1 | 1 | 0 | 1 |
| BDH2 | 0.941 | P1_VS_P2_P3_P4_Lumbar_Coeliac_Stellate_Sphenopalatine | 1 | 0 | 0 | 0 | 0 | 0 | 0 | 0 |
| KAT6A | 0.941 | Lumbar_Coeliac_VS_P1_P2_P3_P4_Stellate_Sphenopalatine | 0 | 0 | 0 | 0 | 1 | 1 | 0 | 0 |
| IL13RA1 | 0.941 | P1_P2_Lumbar_Coeliac_Stellate_Sphenopalatine_VS_P3_P4 | 1 | 1 | 0 | 0 | 1 | 1 | 1 | 1 |
| 2310010J17F | 0.94 | P1_P2_P3_P4_Lumbar_Coeliac_Sphenopalatine_VS_Stellate | 1 | 1 | 1 | 1 | 1 | 1 | 0 | 1 |
| LPIN2 | 0.94 | P1_P2_P3_P4_Lumbar_Coeliac_Sphenopalatine_VS_Stellate | 1 | 1 | 1 | 1 | 1 | 1 | 0 | 1 |
| TMEM200A | 0.94 | P2_P3_P4_Lumbar_Coeliac_Stellate_Sphenopalatine_VS_P1 | 0 | 1 | 1 | 1 | 1 | 1 | 1 | 1 |
| MYH9 | 0.939 | P1_P2_P3_P4_Lumbar_Coeliac_Sphenopalatine_VS_Stellate | 1 | 1 | 1 | 1 | 1 | 1 | 0 | 1 |
| KLF5 | 0.938 | P3_Lumbar_VS_P1_P2_P4_Coeliac_Stellate_Sphenopalatine | 0 | 0 | 1 | 0 | 1 | 0 | 0 | 0 |
| TDRD7 | 0.938 | P1_P2_P3_P4_Lumbar_Coeliac_Sphenopalatine_VS_Stellate | 1 | 1 | 1 | 1 | 1 | 1 | 0 | 1 |
| ENTPD2 | 0.937 | P4_VS_P1_P2_P3_Lumbar_Coeliac_Stellate_Sphenopalatine | 0 | 0 | 0 | 1 | 0 | 0 | 0 | 0 |
| AXIN2 | 0.937 | Coeliac_VS_P1_P2_P3_P4_Lumbar_Stellate_Sphenopalatine | 0 | 0 | 0 | 0 | 0 | 1 | 0 | 0 |
| RIMKLB | 0.937 | P1_P3_P4_Lumbar_Coeliac_Sphenopalatine_VS_P2_Stellate | 1 | 0 | 1 | 1 | 1 | 1 | 0 | 1 |
| TRIM68 | 0.937 | P1_P2_P3_P4_Lumbar_Coeliac_Sphenopalatine_VS_Stellate | 1 | 1 | 1 | 1 | 1 | 1 | 0 | 1 |
| GM47018 | 0.936 | Sphenopalatine_VS_P1_P2_P3_P4_Lumbar_Coeliac_Stellate | 0 | 0 | 0 | 0 | 0 | 0 | 0 | 1 |
| MAP3K3 | 0.936 | P1_P2_P3_P4_Lumbar_Coeliac_Sphenopalatine_VS_Stellate | 1 | 1 | 1 | 1 | 1 | 1 | 0 | 1 |
| BHLHE40 | 0.935 | P3_VS_P1_P2_P4_Lumbar_Coeliac_Stellate_Sphenopalatine | 0 | 0 | 1 | 0 | 0 | 0 | 0 | 0 |
| LIMD1 | 0.934 | P2_VS_P1_P3_P4_Lumbar_Coeliac_Stellate_Sphenopalatine | 0 | 1 | 0 | 0 | 0 | 0 | 0 | 0 |
| PTPRG | 0.934 | P1_P2_P3_P4_VS_Lumbar_Coeliac_Stellate_Sphenopalatine | 1 | 1 | 1 | 1 | 0 | 0 | 0 | 0 |
| OLFML3 | 0.934 | P1_P4_Sphenopalatine_VS_P2_P3_Lumbar_Coeliac_Stellate | 1 | 0 | 0 | 1 | 0 | 0 | 0 | 1 |
| PPP1R1C | 0.932 | P3_VS_P1_P2_P4_Lumbar_Coeliac_Stellate_Sphenopalatine | 0 | 0 | 1 | 0 | 0 | 0 | 0 | 0 |

| | | | | | | | | | | |
|---|---|---|---|---|---|---|---|---|---|---|
| SH3BGR | 0.931 | P4_VS_P1_P2_P3_Lumbar_Coeliac_Stellate_Sphenopalatine | 0 | 0 | 0 | 1 | 0 | 0 | 0 | 0 |
| MMP2 | 0.931 | P3_Lumbar_Coeliac_Stellate_VS_P1_P2_P4_Sphenopalatine | 0 | 0 | 1 | 0 | 1 | 1 | 1 | 0 |
| FAM184A | 0.931 | P1_P2_P3_P4_Lumbar_Coeliac_Sphenopalatine_VS_Stellate | 1 | 1 | 1 | 1 | 1 | 1 | 0 | 1 |
| TMEM161B | 0.931 | P1_P2_P3_P4_Lumbar_Coeliac_Sphenopalatine_VS_Stellate | 1 | 1 | 1 | 1 | 1 | 1 | 0 | 1 |
| CPEB3 | 0.931 | P1_P2_P3_P4_Lumbar_Coeliac_Sphenopalatine_VS_Stellate | 1 | 1 | 1 | 1 | 1 | 1 | 0 | 1 |
| ADRB2 | 0.931 | P1_P4_Lumbar_Coeliac_Stellate_Sphenopalatine_VS_P2_P3 | 1 | 0 | 0 | 1 | 1 | 1 | 1 | 1 |
| BAMBI | 0.93 | P2_VS_P1_P3_P4_Lumbar_Coeliac_Stellate_Sphenopalatine | 0 | 1 | 0 | 0 | 0 | 0 | 0 | 0 |
| SYT14 | 0.928 | P1_P2_P3_P4_Lumbar_Coeliac_Sphenopalatine_VS_Stellate | 1 | 1 | 1 | 1 | 1 | 1 | 0 | 1 |
| SLIT2 | 0.928 | P1_P2_P3_P4_Lumbar_Coeliac_Sphenopalatine_VS_Stellate | 1 | 1 | 1 | 1 | 1 | 1 | 0 | 1 |
| HSPG2 | 0.927 | P4_VS_P1_P2_P3_Lumbar_Coeliac_Stellate_Sphenopalatine | 0 | 0 | 0 | 1 | 0 | 0 | 0 | 0 |
| ITGB6 | 0.926 | P2_VS_P1_P3_P4_Lumbar_Coeliac_Stellate_Sphenopalatine | 0 | 1 | 0 | 0 | 0 | 0 | 0 | 0 |
| HAAO | 0.925 | P4_VS_P1_P2_P3_Lumbar_Coeliac_Stellate_Sphenopalatine | 0 | 0 | 0 | 1 | 0 | 0 | 0 | 0 |
| TMEM54 | 0.924 | P2_P3_P4_VS_P1_Lumbar_Coeliac_Stellate_Sphenopalatine | 0 | 1 | 1 | 1 | 0 | 0 | 0 | 0 |
| CDK5RAP2 | 0.924 | Lumbar_Coeliac_VS_P1_P2_P3_P4_Stellate_Sphenopalatine | 0 | 0 | 0 | 0 | 1 | 1 | 0 | 0 |
| GM42941 | 0.923 | P1_P2_P3_P4_Lumbar_Coeliac_VS_Stellate_Sphenopalatine | 1 | 1 | 1 | 1 | 1 | 1 | 0 | 0 |
| BACE2 | 0.923 | P1_P2_P3_Sphenopalatine_VS_P4_Lumbar_Coeliac_Stellate | 1 | 1 | 1 | 0 | 0 | 0 | 0 | 1 |
| IQSEC1 | 0.923 | P1_P2_P3_P4_Lumbar_Coeliac_Sphenopalatine_VS_Stellate | 1 | 1 | 1 | 1 | 1 | 1 | 0 | 1 |
| EPB41L4B | 0.922 | P1_P2_P3_P4_Lumbar_Coeliac_Sphenopalatine_VS_Stellate | 1 | 1 | 1 | 1 | 1 | 1 | 0 | 1 |
| TMTC4 | 0.922 | P1_P2_P3_P4_Lumbar_Coeliac_Sphenopalatine_VS_Stellate | 1 | 1 | 1 | 1 | 1 | 1 | 0 | 1 |
| CRNKL1 | 0.921 | Lumbar_Coeliac_VS_P1_P2_P3_P4_Stellate_Sphenopalatine | 0 | 0 | 0 | 0 | 1 | 1 | 0 | 0 |
| NANOS3 | 0.921 | P1_P2_P3_P4_Lumbar_Coeliac_Sphenopalatine_VS_Stellate | 1 | 1 | 1 | 1 | 1 | 1 | 0 | 1 |
| SORL1 | 0.92 | P1_P2_P4_VS_P3_Lumbar_Coeliac_Stellate_Sphenopalatine | 1 | 1 | 0 | 1 | 0 | 0 | 0 | 0 |
| FGD3 | 0.92 | Lumbar_VS_P1_P2_P3_P4_Coeliac_Stellate_Sphenopalatine | 0 | 0 | 0 | 0 | 1 | 0 | 0 | 0 |
| HEATR5B | 0.92 | Lumbar_Coeliac_VS_P1_P2_P3_P4_Stellate_Sphenopalatine | 0 | 0 | 0 | 0 | 1 | 1 | 0 | 0 |
| OFD1 | 0.92 | Lumbar_Coeliac_VS_P1_P2_P3_P4_Stellate_Sphenopalatine | 0 | 0 | 0 | 0 | 1 | 1 | 0 | 0 |
| BCL9 | 0.92 | P1_P2_P3_P4_Lumbar_Coeliac_Sphenopalatine_VS_Stellate | 1 | 1 | 1 | 1 | 1 | 1 | 0 | 1 |
| SLC9A3R1 | 0.919 | P4_VS_P1_P2_P3_Lumbar_Coeliac_Stellate_Sphenopalatine | 0 | 0 | 0 | 1 | 0 | 0 | 0 | 0 |
| PCSK5 | 0.919 | P4_VS_P1_P2_P3_Lumbar_Coeliac_Stellate_Sphenopalatine | 0 | 0 | 0 | 1 | 0 | 0 | 0 | 0 |
| SLC16A11 | 0.919 | P4_Lumbar_VS_P1_P2_P3_Coeliac_Stellate_Sphenopalatine | 0 | 0 | 0 | 1 | 1 | 0 | 0 | 0 |
| DHX8 | 0.919 | P1_P2_P3_P4_Lumbar_Coeliac_Sphenopalatine_VS_Stellate | 1 | 1 | 1 | 1 | 1 | 1 | 0 | 1 |

| | | | | | | | | | | |
|---|---|---|---|---|---|---|---|---|---|---|
| MAST4 | 0.918 | P1_P2_P4_Sphenopalatine_VS_P3_Lumbar_Coeliac_Stellate | 1 | 1 | 0 | 1 | 0 | 0 | 0 | 1 |
| MORC3 | 0.917 | P1_P2_P3_P4_Lumbar_Coeliac_Sphenopalatine_VS_Stellate | 1 | 1 | 1 | 1 | 1 | 1 | 0 | 1 |
| DQX1 | 0.916 | Coeliac_VS_P1_P2_P3_P4_Lumbar_Stellate_Sphenopalatine | 0 | 0 | 0 | 0 | 0 | 1 | 0 | 0 |
| HMGN5 | 0.916 | P3_Lumbar_Coeliac_VS_P1_P2_P4_Stellate_Sphenopalatine | 0 | 0 | 1 | 0 | 1 | 1 | 0 | 0 |
| CASP8AP2 | 0.915 | Lumbar_Coeliac_VS_P1_P2_P3_P4_Stellate_Sphenopalatine | 0 | 0 | 0 | 0 | 1 | 1 | 0 | 0 |
| SPRED1 | 0.915 | P1_P2_P3_P4_Lumbar_Coeliac_Sphenopalatine_VS_Stellate | 1 | 1 | 1 | 1 | 1 | 1 | 0 | 1 |
| PDCD7 | 0.915 | P1_P2_P3_P4_Lumbar_Coeliac_Sphenopalatine_VS_Stellate | 1 | 1 | 1 | 1 | 1 | 1 | 0 | 1 |
| FZD5 | 0.913 | P1_P2_P3_P4_Lumbar_Coeliac_VS_Stellate_Sphenopalatine | 1 | 1 | 1 | 1 | 1 | 1 | 0 | 0 |
| SQOR | 0.912 | Sphenopalatine_VS_P1_P2_P3_P4_Lumbar_Coeliac_Stellate | 0 | 0 | 0 | 0 | 0 | 0 | 0 | 1 |
| CDYL | 0.911 | Lumbar_Coeliac_VS_P1_P2_P3_P4_Stellate_Sphenopalatine | 0 | 0 | 0 | 0 | 1 | 1 | 0 | 0 |
| ASAP2 | 0.911 | Coeliac_Sphenopalatine_VS_P1_P2_P3_P4_Lumbar_Stellate | 0 | 0 | 0 | 0 | 0 | 1 | 0 | 1 |
| LRRC4 | 0.91 | P4_VS_P1_P2_P3_Lumbar_Coeliac_Stellate_Sphenopalatine | 0 | 0 | 0 | 1 | 0 | 0 | 0 | 0 |
| PDPR | 0.91 | P1_P2_P3_P4_Lumbar_Coeliac_VS_Stellate_Sphenopalatine | 1 | 1 | 1 | 1 | 1 | 1 | 0 | 0 |
| GM10638 | 0.909 | P1_P2_P3_P4_Lumbar_Coeliac_VS_Stellate_Sphenopalatine | 1 | 1 | 1 | 1 | 1 | 1 | 0 | 0 |
| MAPK4 | 0.909 | P1_P2_P3_P4_Lumbar_Coeliac_VS_Stellate_Sphenopalatine | 1 | 1 | 1 | 1 | 1 | 1 | 0 | 0 |
| ARHGEF10L | 0.909 | P1_P2_P3_P4_Lumbar_Coeliac_Sphenopalatine_VS_Stellate | 1 | 1 | 1 | 1 | 1 | 1 | 0 | 1 |
| CYP4V3 | 0.908 | P2_VS_P1_P3_P4_Lumbar_Coeliac_Stellate_Sphenopalatine | 0 | 1 | 0 | 0 | 0 | 0 | 0 | 0 |
| SGTB | 0.908 | P1_P2_P3_P4_Lumbar_Coeliac_VS_Stellate_Sphenopalatine | 1 | 1 | 1 | 1 | 1 | 1 | 0 | 0 |
| RHOBTB3 | 0.907 | P1_P2_P3_P4_Lumbar_Coeliac_VS_Stellate_Sphenopalatine | 1 | 1 | 1 | 1 | 1 | 1 | 0 | 0 |
| MAP3K1 | 0.907 | P1_P2_P3_P4_Lumbar_Coeliac_Sphenopalatine_VS_Stellate | 1 | 1 | 1 | 1 | 1 | 1 | 0 | 1 |
| GM28523 | 0.906 | P3_Lumbar_Coeliac_VS_P1_P2_P4_Stellate_Sphenopalatine | 0 | 0 | 1 | 0 | 1 | 1 | 0 | 0 |
| MYL4 | 0.906 | P3_P4_Lumbar_Coeliac_Stellate_VS_P1_P2_Sphenopalatine | 0 | 0 | 1 | 1 | 1 | 1 | 1 | 0 |
| TRP53I11 | 0.906 | P1_Lumbar_Sphenopalatine_VS_P2_P3_P4_Coeliac_Stellate | 1 | 0 | 0 | 0 | 1 | 0 | 0 | 1 |
| MARCH9 | 0.906 | P1_P2_P3_P4_Lumbar_Coeliac_Sphenopalatine_VS_Stellate | 1 | 1 | 1 | 1 | 1 | 1 | 0 | 1 |
| UPRT | 0.905 | Lumbar_Coeliac_VS_P1_P2_P3_P4_Stellate_Sphenopalatine | 0 | 0 | 0 | 0 | 1 | 1 | 0 | 0 |
| NETO2 | 0.905 | P1_P2_P3_P4_Lumbar_Coeliac_Stellate_VS_Sphenopalatine | 1 | 1 | 1 | 1 | 1 | 1 | 1 | 0 |
| BRSK2 | 0.904 | P1_P2_P3_P4_Lumbar_Coeliac_Sphenopalatine_VS_Stellate | 1 | 1 | 1 | 1 | 1 | 1 | 0 | 1 |
| DIAPH1 | 0.904 | P1_P2_P3_P4_Lumbar_Coeliac_Sphenopalatine_VS_Stellate | 1 | 1 | 1 | 1 | 1 | 1 | 0 | 1 |
| STAT6 | 0.904 | P1_P2_P4_Lumbar_Coeliac_Stellate_Sphenopalatine_VS_P3 | 1 | 1 | 0 | 1 | 1 | 1 | 1 | 1 |
| CDA | 0.904 | P3_P4_Lumbar_Coeliac_Stellate_Sphenopalatine_VS_P1_P2 | 0 | 0 | 1 | 1 | 1 | 1 | 1 | 1 |

| | | | | | | | | | | |
|---|---|---|---|---|---|---|---|---|---|---|
| RARA | 0.903 | P1_P2_P3_P4_Lumbar_Coeliac_Sphenopalatine_VS_Stellate | 1 | 1 | 1 | 1 | 1 | 1 | 0 | 1 |
| NOC3L | 0.903 | P1_P2_P3_P4_Lumbar_Coeliac_Sphenopalatine_VS_Stellate | 1 | 1 | 1 | 1 | 1 | 1 | 0 | 1 |
| CMTM5 | 0.902 | P4_VS_P1_P2_P3_Lumbar_Coeliac_Stellate_Sphenopalatine | 0 | 0 | 0 | 1 | 0 | 0 | 0 | 0 |
| PGGHG | 0.902 | P1_P2_P3_Lumbar_Coeliac_Stellate_VS_P4_Sphenopalatine | 1 | 1 | 1 | 0 | 1 | 1 | 1 | 0 |
| RBM26 | 0.902 | P1_P2_P3_P4_Lumbar_Coeliac_Sphenopalatine_VS_Stellate | 1 | 1 | 1 | 1 | 1 | 1 | 0 | 1 |
| SMC2OS | 0.901 | Lumbar_Coeliac_VS_P1_P2_P3_P4_Stellate_Sphenopalatine | 0 | 0 | 0 | 0 | 1 | 1 | 0 | 0 |
| CALB1 | 0.901 | P1_P2_P3_Lumbar_Coeliac_Sphenopalatine_VS_P4_Stellate | 1 | 1 | 1 | 0 | 1 | 1 | 0 | 1 |
| RASL11A | 0.901 | P1_P2_P3_P4_Lumbar_Coeliac_Sphenopalatine_VS_Stellate | 1 | 1 | 1 | 1 | 1 | 1 | 0 | 1 |
| GM15860 | 0.899 | P4_VS_P1_P2_P3_Lumbar_Coeliac_Stellate_Sphenopalatine | 0 | 0 | 0 | 1 | 0 | 0 | 0 | 0 |
| IL4RA | 0.899 | P4_VS_P1_P2_P3_Lumbar_Coeliac_Stellate_Sphenopalatine | 0 | 0 | 0 | 1 | 0 | 0 | 0 | 0 |
| ZFP746 | 0.899 | P1_P2_P3_P4_Lumbar_Coeliac_Sphenopalatine_VS_Stellate | 1 | 1 | 1 | 1 | 1 | 1 | 0 | 1 |
| TAF5 | 0.898 | Lumbar_Coeliac_VS_P1_P2_P3_P4_Stellate_Sphenopalatine | 0 | 0 | 0 | 0 | 1 | 1 | 0 | 0 |
| SAA2 | 0.897 | Coeliac_VS_P1_P2_P3_P4_Lumbar_Stellate_Sphenopalatine | 0 | 0 | 0 | 0 | 0 | 1 | 0 | 0 |
| SLC27A6 | 0.897 | P4_Lumbar_Coeliac_VS_P1_P2_P3_Stellate_Sphenopalatine | 0 | 0 | 0 | 1 | 1 | 1 | 0 | 0 |
| ITGB8 | 0.896 | P4_VS_P1_P2_P3_Lumbar_Coeliac_Stellate_Sphenopalatine | 0 | 0 | 0 | 1 | 0 | 0 | 0 | 0 |
| ZFP36L1 | 0.895 | P4_VS_P1_P2_P3_Lumbar_Coeliac_Stellate_Sphenopalatine | 0 | 0 | 0 | 1 | 0 | 0 | 0 | 0 |
| MRTFB | 0.895 | P1_P2_P3_P4_Lumbar_Coeliac_Sphenopalatine_VS_Stellate | 1 | 1 | 1 | 1 | 1 | 1 | 0 | 1 |
| KCNH7 | 0.895 | P4_Lumbar_Coeliac_Stellate_Sphenopalatine_VS_P1_P2_P3 | 0 | 0 | 0 | 1 | 1 | 1 | 1 | 1 |
| TSHZ1 | 0.895 | P1_P2_P4_Lumbar_Coeliac_Stellate_Sphenopalatine_VS_P3 | 1 | 1 | 0 | 1 | 1 | 1 | 1 | 1 |
| ADGRE1 | 0.894 | P1_P2_P3_P4_Coeliac_Stellate_Sphenopalatine_VS_Lumbar | 1 | 1 | 1 | 1 | 0 | 1 | 1 | 1 |
| OGN | 0.892 | P4_VS_P1_P2_P3_Lumbar_Coeliac_Stellate_Sphenopalatine | 0 | 0 | 0 | 1 | 0 | 0 | 0 | 0 |
| RNF152 | 0.892 | P2_P4_VS_P1_P3_Lumbar_Coeliac_Stellate_Sphenopalatine | 0 | 1 | 0 | 1 | 0 | 0 | 0 | 0 |
| CUX2 | 0.892 | P1_P2_P3_P4_Lumbar_Coeliac_Sphenopalatine_VS_Stellate | 1 | 1 | 1 | 1 | 1 | 1 | 0 | 1 |
| KNDC1 | 0.892 | P1_P2_P3_P4_Lumbar_Coeliac_Sphenopalatine_VS_Stellate | 1 | 1 | 1 | 1 | 1 | 1 | 0 | 1 |
| STARD10 | 0.89 | P1_P2_P3_P4_Lumbar_Coeliac_Sphenopalatine_VS_Stellate | 1 | 1 | 1 | 1 | 1 | 1 | 0 | 1 |
| SH3KBP1 | 0.89 | P1_P2_P3_P4_Lumbar_Coeliac_Sphenopalatine_VS_Stellate | 1 | 1 | 1 | 1 | 1 | 1 | 0 | 1 |
| SLC9A9 | 0.889 | P1_P2_P3_P4_Lumbar_Coeliac_Sphenopalatine_VS_Stellate | 1 | 1 | 1 | 1 | 1 | 1 | 0 | 1 |
| UBE2QL1 | 0.888 | P1_P2_Lumbar_Coeliac_Sphenopalatine_VS_P3_P4_Stellate | 1 | 1 | 0 | 0 | 1 | 1 | 0 | 1 |
| LACTB2 | 0.885 | Lumbar_VS_P1_P2_P3_P4_Coeliac_Stellate_Sphenopalatine | 0 | 0 | 0 | 0 | 1 | 0 | 0 | 0 |
| MAP7 | 0.885 | P1_P2_P3_P4_Lumbar_Coeliac_VS_Stellate_Sphenopalatine | 1 | 1 | 1 | 1 | 1 | 1 | 0 | 0 |

| | | | | | | | | | | |
|---|---|---|---|---|---|---|---|---|---|---|
| SAMD9L | 0.884 | P3_VS_P1_P2_P4_Lumbar_Coeliac_Stellate_Sphenopalatine | 0 | 0 | 1 | 0 | 0 | 0 | 0 | 0 |
| CEP290 | 0.884 | Lumbar_Coeliac_VS_P1_P2_P3_P4_Stellate_Sphenopalatine | 0 | 0 | 0 | 0 | 1 | 1 | 0 | 0 |
| PPP1R3F | 0.884 | P1_P2_P3_P4_Lumbar_Coeliac_Sphenopalatine_VS_Stellate | 1 | 1 | 1 | 1 | 1 | 1 | 0 | 1 |
| SHISA2 | 0.883 | P3_Lumbar_Coeliac_Stellate_VS_P1_P2_P4_Sphenopalatine | 0 | 0 | 1 | 0 | 1 | 1 | 1 | 0 |
| ZDHHC5 | 0.883 | P1_P2_P3_P4_Lumbar_Coeliac_Sphenopalatine_VS_Stellate | 1 | 1 | 1 | 1 | 1 | 1 | 0 | 1 |
| CLCN5 | 0.883 | P1_P2_P3_P4_Lumbar_Coeliac_Sphenopalatine_VS_Stellate | 1 | 1 | 1 | 1 | 1 | 1 | 0 | 1 |
| CC2D2A | 0.882 | Coeliac_VS_P1_P2_P3_P4_Lumbar_Stellate_Sphenopalatine | 0 | 0 | 0 | 0 | 0 | 1 | 0 | 0 |
| CRLS1 | 0.882 | P1_P2_P3_P4_Lumbar_Coeliac_Sphenopalatine_VS_Stellate | 1 | 1 | 1 | 1 | 1 | 1 | 0 | 1 |
| PIK3CB | 0.882 | P1_P2_P3_P4_Lumbar_Coeliac_Sphenopalatine_VS_Stellate | 1 | 1 | 1 | 1 | 1 | 1 | 0 | 1 |
| ITGAV | 0.881 | P4_VS_P1_P2_P3_Lumbar_Coeliac_Stellate_Sphenopalatine | 0 | 0 | 0 | 1 | 0 | 0 | 0 | 0 |
| NSUN6 | 0.881 | Lumbar_Coeliac_VS_P1_P2_P3_P4_Stellate_Sphenopalatine | 0 | 0 | 0 | 0 | 1 | 1 | 0 | 0 |
| KCNN2 | 0.881 | P1_P2_P3_P4_Lumbar_Coeliac_Sphenopalatine_VS_Stellate | 1 | 1 | 1 | 1 | 1 | 1 | 0 | 1 |
| ARSB | 0.88 | P4_VS_P1_P2_P3_Lumbar_Coeliac_Stellate_Sphenopalatine | 0 | 0 | 0 | 1 | 0 | 0 | 0 | 0 |
| BEGAIN | 0.88 | Sphenopalatine_VS_P1_P2_P3_P4_Lumbar_Coeliac_Stellate | 0 | 0 | 0 | 0 | 0 | 0 | 0 | 1 |
| ENTPD7 | 0.879 | Lumbar_Coeliac_VS_P1_P2_P3_P4_Stellate_Sphenopalatine | 0 | 0 | 0 | 0 | 1 | 1 | 0 | 0 |
| PRKAB2 | 0.879 | P1_P2_P3_P4_Lumbar_Coeliac_Sphenopalatine_VS_Stellate | 1 | 1 | 1 | 1 | 1 | 1 | 0 | 1 |
| SYN3 | 0.878 | P1_P2_P3_P4_Lumbar_Coeliac_Sphenopalatine_VS_Stellate | 1 | 1 | 1 | 1 | 1 | 1 | 0 | 1 |
| TCF20 | 0.878 | P1_P2_P3_P4_Lumbar_Coeliac_Sphenopalatine_VS_Stellate | 1 | 1 | 1 | 1 | 1 | 1 | 0 | 1 |
| OPTC | 0.877 | P1_VS_P2_P3_P4_Lumbar_Coeliac_Stellate_Sphenopalatine | 1 | 0 | 0 | 0 | 0 | 0 | 0 | 0 |
| SLC22A4 | 0.877 | Lumbar_Coeliac_VS_P1_P2_P3_P4_Stellate_Sphenopalatine | 0 | 0 | 0 | 0 | 1 | 1 | 0 | 0 |
| TNKS1BP1 | 0.876 | P1_P2_P3_P4_Lumbar_Coeliac_Sphenopalatine_VS_Stellate | 1 | 1 | 1 | 1 | 1 | 1 | 0 | 1 |
| PLEKHA3 | 0.875 | P1_P2_P3_P4_Lumbar_Coeliac_Sphenopalatine_VS_Stellate | 1 | 1 | 1 | 1 | 1 | 1 | 0 | 1 |
| TSPAN9 | 0.875 | Lumbar_Coeliac_Stellate_Sphenopalatine_VS_P1_P2_P3_P4 | 0 | 0 | 0 | 0 | 1 | 1 | 1 | 1 |
| ALK | 0.874 | P1_P3_P4_Lumbar_Sphenopalatine_VS_P2_Coeliac_Stellate | 1 | 0 | 1 | 1 | 1 | 0 | 0 | 1 |
| PTPRT | 0.873 | P2_VS_P1_P3_P4_Lumbar_Coeliac_Stellate_Sphenopalatine | 0 | 1 | 0 | 0 | 0 | 0 | 0 | 0 |
| KLHL5 | 0.873 | P3_P4_Lumbar_Coeliac_VS_P1_P2_Stellate_Sphenopalatine | 0 | 0 | 1 | 1 | 1 | 1 | 0 | 0 |
| PDS5B | 0.873 | P1_P2_P3_P4_Lumbar_Coeliac_Sphenopalatine_VS_Stellate | 1 | 1 | 1 | 1 | 1 | 1 | 0 | 1 |
| NLRX1 | 0.872 | P3_VS_P1_P2_P4_Lumbar_Coeliac_Stellate_Sphenopalatine | 0 | 0 | 1 | 0 | 0 | 0 | 0 | 0 |
| UTP14B | 0.872 | P1_P2_P3_P4_Lumbar_Coeliac_Sphenopalatine_VS_Stellate | 1 | 1 | 1 | 1 | 1 | 1 | 0 | 1 |
| NIN | 0.872 | P1_P2_P3_P4_Lumbar_Coeliac_Sphenopalatine_VS_Stellate | 1 | 1 | 1 | 1 | 1 | 1 | 0 | 1 |

| | | | | | | | | | | |
|---|---|---|---|---|---|---|---|---|---|---|
| ZDHHC18 | 0.871 | P1_P2_P3_P4_Lumbar_Coeliac_Sphenopalatine_VS_Stellate | 1 | 1 | 1 | 1 | 1 | 1 | 0 | 1 |
| JAZF1 | 0.871 | P1_P2_P3_P4_Lumbar_Coeliac_Sphenopalatine_VS_Stellate | 1 | 1 | 1 | 1 | 1 | 1 | 0 | 1 |
| GABRG2 | 0.871 | P1_P2_P3_P4_Lumbar_Coeliac_Sphenopalatine_VS_Stellate | 1 | 1 | 1 | 1 | 1 | 1 | 0 | 1 |
| SERPINH1 | 0.87 | P4_VS_P1_P2_P3_Lumbar_Coeliac_Stellate_Sphenopalatine | 0 | 0 | 0 | 1 | 0 | 0 | 0 | 0 |
| CD44 | 0.87 | P1_P2_P4_Lumbar_Coeliac_Sphenopalatine_VS_P3_Stellate | 1 | 1 | 0 | 1 | 1 | 1 | 0 | 1 |
| LYPD1 | 0.869 | P4_VS_P1_P2_P3_Lumbar_Coeliac_Stellate_Sphenopalatine | 0 | 0 | 0 | 1 | 0 | 0 | 0 | 0 |
| CCBE1 | 0.869 | Sphenopalatine_VS_P1_P2_P3_P4_Lumbar_Coeliac_Stellate | 0 | 0 | 0 | 0 | 0 | 0 | 0 | 1 |
| BCL9L | 0.869 | P1_P2_P3_P4_Lumbar_Coeliac_Sphenopalatine_VS_Stellate | 1 | 1 | 1 | 1 | 1 | 1 | 0 | 1 |
| SYNJ2 | 0.868 | P4_VS_P1_P2_P3_Lumbar_Coeliac_Stellate_Sphenopalatine | 0 | 0 | 0 | 1 | 0 | 0 | 0 | 0 |
| SV2B | 0.867 | P1_Sphenopalatine_VS_P2_P3_P4_Lumbar_Coeliac_Stellate | 1 | 0 | 0 | 0 | 0 | 0 | 0 | 1 |
| NRXN3 | 0.867 | P1_P2_P4_Lumbar_Sphenopalatine_VS_P3_Coeliac_Stellate | 1 | 1 | 0 | 1 | 1 | 0 | 0 | 1 |
| PRTG | 0.866 | P1_VS_P2_P3_P4_Lumbar_Coeliac_Stellate_Sphenopalatine | 1 | 0 | 0 | 0 | 0 | 0 | 0 | 0 |
| PLXNB1 | 0.866 | P1_P2_P3_P4_Lumbar_Coeliac_Stellate_VS_Sphenopalatine | 1 | 1 | 1 | 1 | 1 | 1 | 1 | 0 |
| IL6ST | 0.863 | P1_P2_P3_P4_Lumbar_Coeliac_Sphenopalatine_VS_Stellate | 1 | 1 | 1 | 1 | 1 | 1 | 0 | 1 |
| ISYNA1 | 0.863 | P1_P3_P4_Lumbar_Coeliac_Stellate_Sphenopalatine_VS_P2 | 1 | 0 | 1 | 1 | 1 | 1 | 1 | 1 |
| GPSM3 | 0.862 | P3_VS_P1_P2_P4_Lumbar_Coeliac_Stellate_Sphenopalatine | 0 | 0 | 1 | 0 | 0 | 0 | 0 | 0 |
| MB21D2 | 0.862 | P1_P2_P3_P4_Lumbar_Coeliac_Sphenopalatine_VS_Stellate | 1 | 1 | 1 | 1 | 1 | 1 | 0 | 1 |
| PLCL2 | 0.861 | P1_P3_P4_Coeliac_Sphenopalatine_VS_P2_Lumbar_Stellate | 1 | 0 | 1 | 1 | 0 | 1 | 0 | 1 |
| CTSK | 0.86 | P4_VS_P1_P2_P3_Lumbar_Coeliac_Stellate_Sphenopalatine | 0 | 0 | 0 | 1 | 0 | 0 | 0 | 0 |
| CPA2 | 0.86 | P4_VS_P1_P2_P3_Lumbar_Coeliac_Stellate_Sphenopalatine | 0 | 0 | 0 | 1 | 0 | 0 | 0 | 0 |
| FAM107B | 0.86 | P1_P2_P4_Lumbar_Coeliac_VS_P3_Stellate_Sphenopalatine | 1 | 1 | 0 | 1 | 1 | 1 | 0 | 0 |
| DGKI | 0.86 | P1_P2_P3_P4_Lumbar_Coeliac_VS_Stellate_Sphenopalatine | 1 | 1 | 1 | 1 | 1 | 1 | 0 | 0 |
| NUP98 | 0.859 | P1_P2_P3_P4_Lumbar_Coeliac_Sphenopalatine_VS_Stellate | 1 | 1 | 1 | 1 | 1 | 1 | 0 | 1 |
| TFDP2 | 0.859 | P1_P2_P3_P4_Lumbar_Coeliac_Sphenopalatine_VS_Stellate | 1 | 1 | 1 | 1 | 1 | 1 | 0 | 1 |
| S100A6 | 0.858 | P3_VS_P1_P2_P4_Lumbar_Coeliac_Stellate_Sphenopalatine | 0 | 0 | 1 | 0 | 0 | 0 | 0 | 0 |
| NOL4 | 0.858 | Coeliac_VS_P1_P2_P3_P4_Lumbar_Stellate_Sphenopalatine | 0 | 0 | 0 | 0 | 0 | 1 | 0 | 0 |
| CITED1 | 0.858 | P3_Sphenopalatine_VS_P1_P2_P4_Lumbar_Coeliac_Stellate | 0 | 0 | 1 | 0 | 0 | 0 | 0 | 1 |
| SIRT1 | 0.858 | P1_P2_P3_P4_Lumbar_Coeliac_Sphenopalatine_VS_Stellate | 1 | 1 | 1 | 1 | 1 | 1 | 0 | 1 |
| LONRF1 | 0.857 | P1_P2_P3_P4_Lumbar_Coeliac_Sphenopalatine_VS_Stellate | 1 | 1 | 1 | 1 | 1 | 1 | 0 | 1 |
| KLHDC8A | 0.856 | Sphenopalatine_VS_P1_P2_P3_P4_Lumbar_Coeliac_Stellate | 0 | 0 | 0 | 0 | 0 | 0 | 0 | 1 |

| | | | | | | | | | | |
|---|---|---|---|---|---|---|---|---|---|---|
| ZFP40 | 0.856 | P1_P2_P3_P4_Lumbar_Coeliac_Sphenopalatine_VS_Stellate | 1 | 1 | 1 | 1 | 1 | 1 | 0 | 1 |
| ASIC2 | 0.855 | P1_P2_P3_P4_Lumbar_Coeliac_Stellate_VS_Sphenopalatine | 1 | 1 | 1 | 1 | 1 | 1 | 1 | 0 |
| GM47512 | 0.854 | Lumbar_Coeliac_VS_P1_P2_P3_P4_Stellate_Sphenopalatine | 0 | 0 | 0 | 0 | 1 | 1 | 0 | 0 |
| PPP2R5A | 0.854 | P1_P2_P3_P4_Lumbar_Coeliac_Sphenopalatine_VS_Stellate | 1 | 1 | 1 | 1 | 1 | 1 | 0 | 1 |
| SH3D19 | 0.854 | P1_P2_P3_P4_Lumbar_Coeliac_Sphenopalatine_VS_Stellate | 1 | 1 | 1 | 1 | 1 | 1 | 0 | 1 |
| EID2 | 0.854 | P1_P2_P3_P4_Lumbar_Coeliac_Sphenopalatine_VS_Stellate | 1 | 1 | 1 | 1 | 1 | 1 | 0 | 1 |
| HDDC3 | 0.854 | P1_P2_P3_P4_Lumbar_Coeliac_Sphenopalatine_VS_Stellate | 1 | 1 | 1 | 1 | 1 | 1 | 0 | 1 |
| GM43113 | 0.853 | Lumbar_VS_P1_P2_P3_P4_Coeliac_Stellate_Sphenopalatine | 0 | 0 | 0 | 0 | 1 | 0 | 0 | 0 |
| FBXL7 | 0.853 | P4_Lumbar_Coeliac_VS_P1_P2_P3_Stellate_Sphenopalatine | 0 | 0 | 0 | 1 | 1 | 1 | 0 | 0 |
| FBLL1 | 0.853 | P1_P2_P3_P4_Lumbar_Coeliac_Sphenopalatine_VS_Stellate | 1 | 1 | 1 | 1 | 1 | 1 | 0 | 1 |
| EGR1 | 0.853 | P1_P2_P3_P4_Lumbar_Coeliac_Sphenopalatine_VS_Stellate | 1 | 1 | 1 | 1 | 1 | 1 | 0 | 1 |
| TNNT2 | 0.852 | P3_VS_P1_P2_P4_Lumbar_Coeliac_Stellate_Sphenopalatine | 0 | 0 | 1 | 0 | 0 | 0 | 0 | 0 |
| GM37494 | 0.852 | Lumbar_Coeliac_VS_P1_P2_P3_P4_Stellate_Sphenopalatine | 0 | 0 | 0 | 0 | 1 | 1 | 0 | 0 |
| C1GALT1 | 0.852 | P1_P2_P4_Lumbar_Coeliac_VS_P3_Stellate_Sphenopalatine | 1 | 1 | 0 | 1 | 1 | 1 | 0 | 0 |
| CHST15 | 0.852 | P1_P2_P3_P4_Lumbar_Coeliac_VS_Stellate_Sphenopalatine | 1 | 1 | 1 | 1 | 1 | 1 | 0 | 0 |
| IGSF5 | 0.85 | P1_VS_P2_P3_P4_Lumbar_Coeliac_Stellate_Sphenopalatine | 1 | 0 | 0 | 0 | 0 | 0 | 0 | 0 |
| ZFP53 | 0.849 | Lumbar_Coeliac_VS_P1_P2_P3_P4_Stellate_Sphenopalatine | 0 | 0 | 0 | 0 | 1 | 1 | 0 | 0 |
| E130308A19I | 0.849 | P1_P2_P3_P4_Lumbar_Coeliac_Sphenopalatine_VS_Stellate | 1 | 1 | 1 | 1 | 1 | 1 | 0 | 1 |
| BGN | 0.848 | P4_VS_P1_P2_P3_Lumbar_Coeliac_Stellate_Sphenopalatine | 0 | 0 | 0 | 1 | 0 | 0 | 0 | 0 |
| L3MBTL3 | 0.848 | Lumbar_VS_P1_P2_P3_P4_Coeliac_Stellate_Sphenopalatine | 0 | 0 | 0 | 0 | 1 | 0 | 0 | 0 |
| ADCY7 | 0.848 | Sphenopalatine_VS_P1_P2_P3_P4_Lumbar_Coeliac_Stellate | 0 | 0 | 0 | 0 | 0 | 0 | 0 | 1 |
| ZFYVE16 | 0.848 | P1_P2_P3_P4_Lumbar_Coeliac_Sphenopalatine_VS_Stellate | 1 | 1 | 1 | 1 | 1 | 1 | 0 | 1 |
| SDC4 | 0.847 | P4_VS_P1_P2_P3_Lumbar_Coeliac_Stellate_Sphenopalatine | 0 | 0 | 0 | 1 | 0 | 0 | 0 | 0 |
| SASS6 | 0.847 | Lumbar_Coeliac_VS_P1_P2_P3_P4_Stellate_Sphenopalatine | 0 | 0 | 0 | 0 | 1 | 1 | 0 | 0 |
| XIST | 0.847 | Stellate_VS_P1_P2_P3_P4_Lumbar_Coeliac_Sphenopalatine | 0 | 0 | 0 | 0 | 0 | 0 | 1 | 0 |
| LXN | 0.847 | P1_P3_P4_Lumbar_Coeliac_Stellate_Sphenopalatine_VS_P2 | 1 | 0 | 1 | 1 | 1 | 1 | 1 | 1 |
| ZFP60 | 0.846 | Lumbar_Coeliac_VS_P1_P2_P3_P4_Stellate_Sphenopalatine | 0 | 0 | 0 | 0 | 1 | 1 | 0 | 0 |
| MGAT4A | 0.846 | P1_P2_P3_P4_Lumbar_Coeliac_Sphenopalatine_VS_Stellate | 1 | 1 | 1 | 1 | 1 | 1 | 0 | 1 |
| SCRT1 | 0.846 | P1_P2_P3_P4_Lumbar_Coeliac_Sphenopalatine_VS_Stellate | 1 | 1 | 1 | 1 | 1 | 1 | 0 | 1 |
| TOX2 | 0.846 | P3_P4_Lumbar_Coeliac_Stellate_Sphenopalatine_VS_P1_P2 | 0 | 0 | 1 | 1 | 1 | 1 | 1 | 1 |

| | | | | | | | | | | |
|---|---|---|---|---|---|---|---|---|---|---|
| PREX1 | 0.845 | Lumbar_Coeliac_VS_P1_P2_P3_P4_Stellate_Sphenopalatine | 0 | 0 | 0 | 0 | 1 | 1 | 0 | 0 |
| PRDM2 | 0.845 | P1_P2_P3_P4_Lumbar_Coeliac_Sphenopalatine_VS_Stellate | 1 | 1 | 1 | 1 | 1 | 1 | 0 | 1 |
| ST5 | 0.844 | P4_VS_P1_P2_P3_Lumbar_Coeliac_Stellate_Sphenopalatine | 0 | 0 | 0 | 1 | 0 | 0 | 0 | 0 |
| CCDC39 | 0.844 | P4_Lumbar_Coeliac_VS_P1_P2_P3_Stellate_Sphenopalatine | 0 | 0 | 0 | 1 | 1 | 1 | 0 | 0 |
| TGFB3 | 0.844 | P2_P3_P4_Lumbar_Coeliac_Stellate_VS_P1_Sphenopalatine | 0 | 1 | 1 | 1 | 1 | 1 | 1 | 0 |
| FASTKD1 | 0.844 | P1_P2_P3_P4_Lumbar_Coeliac_Sphenopalatine_VS_Stellate | 1 | 1 | 1 | 1 | 1 | 1 | 0 | 1 |
| STXBP5L | 0.843 | Lumbar_Coeliac_VS_P1_P2_P3_P4_Stellate_Sphenopalatine | 0 | 0 | 0 | 0 | 1 | 1 | 0 | 0 |
| FRAS1 | 0.842 | P4_VS_P1_P2_P3_Lumbar_Coeliac_Stellate_Sphenopalatine | 0 | 0 | 0 | 1 | 0 | 0 | 0 | 0 |
| BCAN | 0.841 | P4_VS_P1_P2_P3_Lumbar_Coeliac_Stellate_Sphenopalatine | 0 | 0 | 0 | 1 | 0 | 0 | 0 | 0 |
| AKAP7 | 0.841 | P1_P2_P3_P4_Lumbar_Coeliac_Stellate_VS_Sphenopalatine | 1 | 1 | 1 | 1 | 1 | 1 | 1 | 0 |
| PPP4R4 | 0.84 | Lumbar_Coeliac_VS_P1_P2_P3_P4_Stellate_Sphenopalatine | 0 | 0 | 0 | 0 | 1 | 1 | 0 | 0 |
| HLTF | 0.84 | P1_P2_P3_P4_Lumbar_Coeliac_Sphenopalatine_VS_Stellate | 1 | 1 | 1 | 1 | 1 | 1 | 0 | 1 |
| ATP11C | 0.84 | P1_P2_P3_P4_Lumbar_Coeliac_Sphenopalatine_VS_Stellate | 1 | 1 | 1 | 1 | 1 | 1 | 0 | 1 |
| SERPINA3N | 0.839 | P1_P2_P4_Sphenopalatine_VS_P3_Lumbar_Coeliac_Stellate | 1 | 1 | 0 | 1 | 0 | 0 | 0 | 1 |
| TENM4 | 0.838 | P1_P2_P3_P4_Lumbar_Coeliac_VS_Stellate_Sphenopalatine | 1 | 1 | 1 | 1 | 1 | 1 | 0 | 0 |
| RNF168 | 0.836 | P1_P2_P3_P4_Lumbar_Coeliac_Sphenopalatine_VS_Stellate | 1 | 1 | 1 | 1 | 1 | 1 | 0 | 1 |
| RUNX1 | 0.834 | Sphenopalatine_VS_P1_P2_P3_P4_Lumbar_Coeliac_Stellate | 0 | 0 | 0 | 0 | 0 | 0 | 0 | 1 |
| BCLAF3 | 0.833 | Lumbar_Coeliac_VS_P1_P2_P3_P4_Stellate_Sphenopalatine | 0 | 0 | 0 | 0 | 1 | 1 | 0 | 0 |
| CREBZF | 0.833 | P1_P2_P3_P4_Lumbar_Coeliac_Sphenopalatine_VS_Stellate | 1 | 1 | 1 | 1 | 1 | 1 | 0 | 1 |
| CNN2 | 0.832 | P4_VS_P1_P2_P3_Lumbar_Coeliac_Stellate_Sphenopalatine | 0 | 0 | 0 | 1 | 0 | 0 | 0 | 0 |
| HS3ST5 | 0.832 | P1_P2_P3_Lumbar_Coeliac_Stellate_Sphenopalatine_VS_P4 | 1 | 1 | 1 | 0 | 1 | 1 | 1 | 1 |
| KLC3 | 0.831 | Lumbar_VS_P1_P2_P3_P4_Coeliac_Stellate_Sphenopalatine | 0 | 0 | 0 | 0 | 1 | 0 | 0 | 0 |
| KCNJ11 | 0.831 | Sphenopalatine_VS_P1_P2_P3_P4_Lumbar_Coeliac_Stellate | 0 | 0 | 0 | 0 | 0 | 0 | 0 | 1 |
| MPDZ | 0.831 | P1_P2_P3_P4_Lumbar_Coeliac_Sphenopalatine_VS_Stellate | 1 | 1 | 1 | 1 | 1 | 1 | 0 | 1 |
| TMEM119 | 0.83 | P2_VS_P1_P3_P4_Lumbar_Coeliac_Stellate_Sphenopalatine | 0 | 1 | 0 | 0 | 0 | 0 | 0 | 0 |
| GCA | 0.829 | P4_VS_P1_P2_P3_Lumbar_Coeliac_Stellate_Sphenopalatine | 0 | 0 | 0 | 1 | 0 | 0 | 0 | 0 |
| TXNIP | 0.829 | P4_VS_P1_P2_P3_Lumbar_Coeliac_Stellate_Sphenopalatine | 0 | 0 | 0 | 1 | 0 | 0 | 0 | 0 |
| TMEM132A | 0.829 | P3_Lumbar_Coeliac_VS_P1_P2_P4_Stellate_Sphenopalatine | 0 | 0 | 1 | 0 | 1 | 1 | 0 | 0 |
| SLC25A28 | 0.829 | P1_P2_P3_P4_Lumbar_Coeliac_Sphenopalatine_VS_Stellate | 1 | 1 | 1 | 1 | 1 | 1 | 0 | 1 |
| S100G | 0.828 | Lumbar_Stellate_VS_P1_P2_P3_P4_Coeliac_Sphenopalatine | 0 | 0 | 0 | 0 | 1 | 0 | 1 | 0 |

| | | | | | | | | | | |
|---|---|---|---|---|---|---|---|---|---|---|
| CNOT6L | 0.828 | P1_P2_P3_P4_Lumbar_Coeliac_Sphenopalatine_VS_Stellate | 1 | 1 | 1 | 1 | 1 | 1 | 0 | 1 |
| SLC9A1 | 0.827 | P1_P2_P3_P4_Lumbar_Coeliac_Sphenopalatine_VS_Stellate | 1 | 1 | 1 | 1 | 1 | 1 | 0 | 1 |
| FAM126A | 0.827 | P1_P2_P3_P4_Lumbar_Coeliac_Sphenopalatine_VS_Stellate | 1 | 1 | 1 | 1 | 1 | 1 | 0 | 1 |
| SOX10 | 0.826 | P4_VS_P1_P2_P3_Lumbar_Coeliac_Stellate_Sphenopalatine | 0 | 0 | 0 | 1 | 0 | 0 | 0 | 0 |
| ZFX | 0.826 | P1_P2_P3_P4_Lumbar_Coeliac_Sphenopalatine_VS_Stellate | 1 | 1 | 1 | 1 | 1 | 1 | 0 | 1 |
| GM26759 | 0.822 | P2_VS_P1_P3_P4_Lumbar_Coeliac_Stellate_Sphenopalatine | 0 | 1 | 0 | 0 | 0 | 0 | 0 | 0 |
| LRCH3 | 0.822 | P1_P2_P3_P4_Lumbar_Coeliac_VS_Stellate_Sphenopalatine | 1 | 1 | 1 | 1 | 1 | 1 | 0 | 0 |
| SOGA1 | 0.822 | P1_P2_P3_P4_Lumbar_Coeliac_Sphenopalatine_VS_Stellate | 1 | 1 | 1 | 1 | 1 | 1 | 0 | 1 |
| CDC25A | 0.822 | P1_P2_P3_P4_Lumbar_Coeliac_Sphenopalatine_VS_Stellate | 1 | 1 | 1 | 1 | 1 | 1 | 0 | 1 |
| BTAF1 | 0.822 | P1_P2_P3_P4_Lumbar_Coeliac_Sphenopalatine_VS_Stellate | 1 | 1 | 1 | 1 | 1 | 1 | 0 | 1 |
| SLC7A7 | 0.822 | P3_P4_Lumbar_Coeliac_Stellate_Sphenopalatine_VS_P1_P2 | 0 | 0 | 1 | 1 | 1 | 1 | 1 | 1 |
| USP29 | 0.821 | P1_P2_P3_P4_Lumbar_Coeliac_Sphenopalatine_VS_Stellate | 1 | 1 | 1 | 1 | 1 | 1 | 0 | 1 |
| 2410018L13I | 0.821 | P1_P2_P3_P4_Lumbar_Coeliac_Sphenopalatine_VS_Stellate | 1 | 1 | 1 | 1 | 1 | 1 | 0 | 1 |
| PLEKHM2 | 0.819 | P1_P2_P3_P4_Lumbar_Coeliac_Sphenopalatine_VS_Stellate | 1 | 1 | 1 | 1 | 1 | 1 | 0 | 1 |
| 5330417C22I | 0.818 | P4_VS_P1_P2_P3_Lumbar_Coeliac_Stellate_Sphenopalatine | 0 | 0 | 0 | 1 | 0 | 0 | 0 | 0 |
| EVPL | 0.818 | Coeliac_VS_P1_P2_P3_P4_Lumbar_Stellate_Sphenopalatine | 0 | 0 | 0 | 0 | 0 | 1 | 0 | 0 |
| CSMD2 | 0.818 | P1_P2_P3_P4_Lumbar_Coeliac_VS_Stellate_Sphenopalatine | 1 | 1 | 1 | 1 | 1 | 1 | 0 | 0 |
| TBC1D8 | 0.818 | P1_P2_P3_P4_Lumbar_Coeliac_Sphenopalatine_VS_Stellate | 1 | 1 | 1 | 1 | 1 | 1 | 0 | 1 |
| MLLT3 | 0.817 | P4_VS_P1_P2_P3_Lumbar_Coeliac_Stellate_Sphenopalatine | 0 | 0 | 0 | 1 | 0 | 0 | 0 | 0 |
| PRDM16 | 0.817 | P4_VS_P1_P2_P3_Lumbar_Coeliac_Stellate_Sphenopalatine | 0 | 0 | 0 | 1 | 0 | 0 | 0 | 0 |
| HSPA12B | 0.817 | Lumbar_VS_P1_P2_P3_P4_Coeliac_Stellate_Sphenopalatine | 0 | 0 | 0 | 0 | 1 | 0 | 0 | 0 |
| CBLB | 0.817 | P1_P2_P3_P4_Lumbar_Coeliac_VS_Stellate_Sphenopalatine | 1 | 1 | 1 | 1 | 1 | 1 | 0 | 0 |
| BTBD7 | 0.817 | P1_P2_P3_P4_Lumbar_Coeliac_Sphenopalatine_VS_Stellate | 1 | 1 | 1 | 1 | 1 | 1 | 0 | 1 |
| CDK13 | 0.817 | P1_P2_P3_P4_Lumbar_Coeliac_Sphenopalatine_VS_Stellate | 1 | 1 | 1 | 1 | 1 | 1 | 0 | 1 |
| RPRM | 0.817 | P1_P4_Lumbar_Coeliac_Stellate_Sphenopalatine_VS_P2_P3 | 1 | 0 | 0 | 1 | 1 | 1 | 1 | 1 |
| 9530082P21I | 0.816 | P1_P4_Lumbar_Coeliac_VS_P2_P3_Stellate_Sphenopalatine | 1 | 0 | 0 | 1 | 1 | 1 | 0 | 0 |
| QPCT | 0.816 | P1_P2_P3_P4_Lumbar_Coeliac_Stellate_VS_Sphenopalatine | 1 | 1 | 1 | 1 | 1 | 1 | 1 | 0 |
| PIGZ | 0.815 | P4_Lumbar_Coeliac_VS_P1_P2_P3_Stellate_Sphenopalatine | 0 | 0 | 0 | 1 | 1 | 1 | 0 | 0 |
| FHL2 | 0.814 | Lumbar_VS_P1_P2_P3_P4_Coeliac_Stellate_Sphenopalatine | 0 | 0 | 0 | 0 | 1 | 0 | 0 | 0 |
| GM49359 | 0.814 | Lumbar_Coeliac_VS_P1_P2_P3_P4_Stellate_Sphenopalatine | 0 | 0 | 0 | 0 | 1 | 1 | 0 | 0 |

| | | | | | | | | | | |
|---|---|---|---|---|---|---|---|---|---|---|
| FAM222A | 0.814 | P1_P2_P3_P4_Lumbar_Coeliac_Stellate_VS_Sphenopalatine | 1 | 1 | 1 | 1 | 1 | 1 | 1 | 0 |
| APOL7D | 0.814 | Sphenopalatine_VS_P1_P2_P3_P4_Lumbar_Coeliac_Stellate | 0 | 0 | 0 | 0 | 0 | 0 | 0 | 1 |
| SESTD1 | 0.814 | P1_P2_P4_Lumbar_Coeliac_Sphenopalatine_VS_P3_Stellate | 1 | 1 | 0 | 1 | 1 | 1 | 0 | 1 |
| DACT2 | 0.813 | P4_Lumbar_VS_P1_P2_P3_Coeliac_Stellate_Sphenopalatine | 0 | 0 | 0 | 1 | 1 | 0 | 0 | 0 |
| MYL6B | 0.813 | P1_P2_P3_Lumbar_Coeliac_VS_P4_Stellate_Sphenopalatine | 1 | 1 | 1 | 0 | 1 | 1 | 0 | 0 |
| 1700095J03F | 0.812 | Lumbar_VS_P1_P2_P3_P4_Coeliac_Stellate_Sphenopalatine | 0 | 0 | 0 | 0 | 1 | 0 | 0 | 0 |
| ZSWIM6 | 0.812 | P1_P2_P3_P4_Lumbar_Coeliac_Sphenopalatine_VS_Stellate | 1 | 1 | 1 | 1 | 1 | 1 | 0 | 1 |
| H2-M3 | 0.811 | P4_VS_P1_P2_P3_Lumbar_Coeliac_Stellate_Sphenopalatine | 0 | 0 | 0 | 1 | 0 | 0 | 0 | 0 |
| ZFP74 | 0.811 | P1_P2_P3_P4_Lumbar_Coeliac_Sphenopalatine_VS_Stellate | 1 | 1 | 1 | 1 | 1 | 1 | 0 | 1 |
| ZBTB44 | 0.811 | P1_P2_P3_P4_Lumbar_Coeliac_Sphenopalatine_VS_Stellate | 1 | 1 | 1 | 1 | 1 | 1 | 0 | 1 |
| TMEM51 | 0.811 | P1_P2_P3_Lumbar_Coeliac_Stellate_Sphenopalatine_VS_P4 | 1 | 1 | 1 | 0 | 1 | 1 | 1 | 1 |
| FIBCD1 | 0.81 | P2_P4_VS_P1_P3_Lumbar_Coeliac_Stellate_Sphenopalatine | 0 | 1 | 0 | 1 | 0 | 0 | 0 | 0 |
| ATM | 0.81 | Lumbar_Coeliac_VS_P1_P2_P3_P4_Stellate_Sphenopalatine | 0 | 0 | 0 | 0 | 1 | 1 | 0 | 0 |
| CNOT3 | 0.81 | P1_P2_P3_P4_Lumbar_Coeliac_Sphenopalatine_VS_Stellate | 1 | 1 | 1 | 1 | 1 | 1 | 0 | 1 |
| PRSS35 | 0.809 | P3_VS_P1_P2_P4_Lumbar_Coeliac_Stellate_Sphenopalatine | 0 | 0 | 1 | 0 | 0 | 0 | 0 | 0 |
| ATF3 | 0.809 | P4_VS_P1_P2_P3_Lumbar_Coeliac_Stellate_Sphenopalatine | 0 | 0 | 0 | 1 | 0 | 0 | 0 | 0 |
| D430041D05 | 0.809 | Lumbar_VS_P1_P2_P3_P4_Coeliac_Stellate_Sphenopalatine | 0 | 0 | 0 | 0 | 1 | 0 | 0 | 0 |
| MOCS3 | 0.809 | P1_P2_P3_P4_Lumbar_Coeliac_Sphenopalatine_VS_Stellate | 1 | 1 | 1 | 1 | 1 | 1 | 0 | 1 |
| CLIC4 | 0.809 | P1_P2_P3_P4_Lumbar_Coeliac_Sphenopalatine_VS_Stellate | 1 | 1 | 1 | 1 | 1 | 1 | 0 | 1 |
| NR1D2 | 0.809 | P1_P2_P3_P4_Lumbar_Coeliac_Sphenopalatine_VS_Stellate | 1 | 1 | 1 | 1 | 1 | 1 | 0 | 1 |
| TBX3 | 0.808 | P1_P2_P3_Coeliac_Sphenopalatine_VS_P4_Lumbar_Stellate | 1 | 1 | 1 | 0 | 0 | 1 | 0 | 1 |
| SMARCAL1 | 0.807 | Lumbar_Coeliac_VS_P1_P2_P3_P4_Stellate_Sphenopalatine | 0 | 0 | 0 | 0 | 1 | 1 | 0 | 0 |
| RNF144A | 0.807 | P1_P2_P3_P4_Lumbar_Coeliac_Stellate_VS_Sphenopalatine | 1 | 1 | 1 | 1 | 1 | 1 | 1 | 0 |
| SAMD1 | 0.807 | P1_P2_P3_P4_Lumbar_Coeliac_Sphenopalatine_VS_Stellate | 1 | 1 | 1 | 1 | 1 | 1 | 0 | 1 |
| SPACA6 | 0.807 | P1_P2_P3_P4_Lumbar_Coeliac_Sphenopalatine_VS_Stellate | 1 | 1 | 1 | 1 | 1 | 1 | 0 | 1 |
| LTBP3 | 0.807 | P1_P2_P3_P4_Lumbar_Coeliac_Sphenopalatine_VS_Stellate | 1 | 1 | 1 | 1 | 1 | 1 | 0 | 1 |
| PLOD1 | 0.804 | P4_VS_P1_P2_P3_Lumbar_Coeliac_Stellate_Sphenopalatine | 0 | 0 | 0 | 1 | 0 | 0 | 0 | 0 |
| NRIP1 | 0.804 | P1_P2_P3_P4_Lumbar_Coeliac_Sphenopalatine_VS_Stellate | 1 | 1 | 1 | 1 | 1 | 1 | 0 | 1 |
| FANCF | 0.803 | Lumbar_VS_P1_P2_P3_P4_Coeliac_Stellate_Sphenopalatine | 0 | 0 | 0 | 0 | 1 | 0 | 0 | 0 |
| ST8SIA2 | 0.803 | Lumbar_Coeliac_VS_P1_P2_P3_P4_Stellate_Sphenopalatine | 0 | 0 | 0 | 0 | 1 | 1 | 0 | 0 |

| | | | | | | | | | | |
|---|---|---|---|---|---|---|---|---|---|---|
| FBLN1 | 0.803 | P1_P2_Lumbar_Coeliac_VS_P3_P4_Stellate_Sphenopalatine | 1 | 1 | 0 | 0 | 1 | 1 | 0 | 0 |
| CUL4B | 0.803 | P1_P2_P3_P4_Lumbar_Coeliac_Sphenopalatine_VS_Stellate | 1 | 1 | 1 | 1 | 1 | 1 | 0 | 1 |
| DPYD | 0.802 | P1_P2_P3_P4_Lumbar_Coeliac_Sphenopalatine_VS_Stellate | 1 | 1 | 1 | 1 | 1 | 1 | 0 | 1 |
| IQSEC2 | 0.802 | P1_P2_P3_P4_Lumbar_Coeliac_Sphenopalatine_VS_Stellate | 1 | 1 | 1 | 1 | 1 | 1 | 0 | 1 |
| LRBA | 0.801 | P1_P2_P3_P4_Lumbar_Coeliac_Sphenopalatine_VS_Stellate | 1 | 1 | 1 | 1 | 1 | 1 | 0 | 1 |
| DISP3 | 0.8 | P3_VS_P1_P2_P4_Lumbar_Coeliac_Stellate_Sphenopalatine | 0 | 0 | 1 | 0 | 0 | 0 | 0 | 0 |
| PLEKHA7 | 0.8 | Lumbar_VS_P1_P2_P3_P4_Coeliac_Stellate_Sphenopalatine | 0 | 0 | 0 | 0 | 1 | 0 | 0 | 0 |
| RIMS1 | 0.8 | P3_P4_Lumbar_Coeliac_VS_P1_P2_Stellate_Sphenopalatine | 0 | 0 | 1 | 1 | 1 | 1 | 0 | 0 |
| TOB2 | 0.8 | P1_P2_P3_P4_Lumbar_Coeliac_Sphenopalatine_VS_Stellate | 1 | 1 | 1 | 1 | 1 | 1 | 0 | 1 |
| SLK | 0.8 | P1_P2_P3_P4_Lumbar_Coeliac_Sphenopalatine_VS_Stellate | 1 | 1 | 1 | 1 | 1 | 1 | 0 | 1 |
| DCN | 0.798 | P1_P2_P3_P4_Coeliac_Sphenopalatine_VS_Lumbar_Stellate | 1 | 1 | 1 | 1 | 0 | 1 | 0 | 1 |
| CSRNP1 | 0.797 | P1_P2_P4_Sphenopalatine_VS_P3_Lumbar_Coeliac_Stellate | 1 | 1 | 0 | 1 | 0 | 0 | 0 | 1 |
| ERBIN | 0.797 | P1_P2_P3_P4_Lumbar_Coeliac_Sphenopalatine_VS_Stellate | 1 | 1 | 1 | 1 | 1 | 1 | 0 | 1 |
| ETL4 | 0.796 | P1_Lumbar_VS_P2_P3_P4_Coeliac_Stellate_Sphenopalatine | 1 | 0 | 0 | 0 | 1 | 0 | 0 | 0 |
| CEMIP2 | 0.796 | P4_Lumbar_Coeliac_VS_P1_P2_P3_Stellate_Sphenopalatine | 0 | 0 | 0 | 1 | 1 | 1 | 0 | 0 |
| SLIT3 | 0.796 | P1_P2_P3_P4_Lumbar_Coeliac_Sphenopalatine_VS_Stellate | 1 | 1 | 1 | 1 | 1 | 1 | 0 | 1 |
| PDCD11 | 0.796 | P1_P2_P3_P4_Lumbar_Coeliac_Sphenopalatine_VS_Stellate | 1 | 1 | 1 | 1 | 1 | 1 | 0 | 1 |
| GLP1R | 0.795 | P3_VS_P1_P2_P4_Lumbar_Coeliac_Stellate_Sphenopalatine | 0 | 0 | 1 | 0 | 0 | 0 | 0 | 0 |
| NOS3 | 0.795 | P4_VS_P1_P2_P3_Lumbar_Coeliac_Stellate_Sphenopalatine | 0 | 0 | 0 | 1 | 0 | 0 | 0 | 0 |
| S1PR3 | 0.795 | P4_VS_P1_P2_P3_Lumbar_Coeliac_Stellate_Sphenopalatine | 0 | 0 | 0 | 1 | 0 | 0 | 0 | 0 |
| INO80 | 0.795 | P1_P2_P3_P4_Lumbar_Coeliac_Sphenopalatine_VS_Stellate | 1 | 1 | 1 | 1 | 1 | 1 | 0 | 1 |
| NBL1 | 0.795 | P1_P2_P3_Lumbar_Coeliac_Stellate_Sphenopalatine_VS_P4 | 1 | 1 | 1 | 0 | 1 | 1 | 1 | 1 |
| TRPM3 | 0.794 | P4_Lumbar_VS_P1_P2_P3_Coeliac_Stellate_Sphenopalatine | 0 | 0 | 0 | 1 | 1 | 0 | 0 | 0 |
| ZFP618 | 0.793 | P1_P2_P3_P4_Lumbar_Coeliac_Sphenopalatine_VS_Stellate | 1 | 1 | 1 | 1 | 1 | 1 | 0 | 1 |
| ETS2 | 0.793 | P1_P2_P3_P4_Lumbar_Coeliac_Sphenopalatine_VS_Stellate | 1 | 1 | 1 | 1 | 1 | 1 | 0 | 1 |
| PCYT1B | 0.793 | P1_P2_P3_P4_Lumbar_Coeliac_Sphenopalatine_VS_Stellate | 1 | 1 | 1 | 1 | 1 | 1 | 0 | 1 |
| DUSP6 | 0.792 | P3_VS_P1_P2_P4_Lumbar_Coeliac_Stellate_Sphenopalatine | 0 | 0 | 1 | 0 | 0 | 0 | 0 | 0 |
| EBF1 | 0.792 | P1_P2_P3_P4_Lumbar_Coeliac_Stellate_VS_Sphenopalatine | 1 | 1 | 1 | 1 | 1 | 1 | 1 | 0 |
| DOCK4 | 0.792 | P1_P2_P3_P4_Lumbar_Coeliac_Stellate_VS_Sphenopalatine | 1 | 1 | 1 | 1 | 1 | 1 | 1 | 0 |
| USP6NL | 0.792 | P1_P2_P3_P4_Lumbar_Coeliac_Sphenopalatine_VS_Stellate | 1 | 1 | 1 | 1 | 1 | 1 | 0 | 1 |

| | | | | | | | | | | |
|---|---|---|---|---|---|---|---|---|---|---|
| MYCL | 0.792 | P1_P2_P3_Lumbar_Coeliac_Stellate_Sphenopalatine_VS_P4 | 1 | 1 | 1 | 0 | 1 | 1 | 1 | 1 |
| RBFOX1 | 0.792 | P1_P2_P3_Lumbar_Coeliac_Stellate_Sphenopalatine_VS_P4 | 1 | 1 | 1 | 0 | 1 | 1 | 1 | 1 |
| GPX8 | 0.791 | P4_VS_P1_P2_P3_Lumbar_Coeliac_Stellate_Sphenopalatine | 0 | 0 | 0 | 1 | 0 | 0 | 0 | 0 |
| PACRG | 0.791 | P3_P4_Coeliac_VS_P1_P2_Lumbar_Stellate_Sphenopalatine | 0 | 0 | 1 | 1 | 0 | 1 | 0 | 0 |
| ZFP1 | 0.791 | Lumbar_Coeliac_VS_P1_P2_P3_P4_Stellate_Sphenopalatine | 0 | 0 | 0 | 0 | 1 | 1 | 0 | 0 |
| TCAIM | 0.791 | P1_P2_P3_P4_Lumbar_Coeliac_Sphenopalatine_VS_Stellate | 1 | 1 | 1 | 1 | 1 | 1 | 0 | 1 |
| HIST1H4H | 0.79 | Coeliac_VS_P1_P2_P3_P4_Lumbar_Stellate_Sphenopalatine | 0 | 0 | 0 | 0 | 0 | 1 | 0 | 0 |
| HMCN2 | 0.789 | P1_Sphenopalatine_VS_P2_P3_P4_Lumbar_Coeliac_Stellate | 1 | 0 | 0 | 0 | 0 | 0 | 0 | 1 |
| ANKRD26 | 0.789 | P1_P2_P3_P4_Lumbar_Coeliac_Sphenopalatine_VS_Stellate | 1 | 1 | 1 | 1 | 1 | 1 | 0 | 1 |
| LDOC1 | 0.788 | P3_VS_P1_P2_P4_Lumbar_Coeliac_Stellate_Sphenopalatine | 0 | 0 | 1 | 0 | 0 | 0 | 0 | 0 |
| GM45159 | 0.788 | P4_Lumbar_Coeliac_VS_P1_P2_P3_Stellate_Sphenopalatine | 0 | 0 | 0 | 1 | 1 | 1 | 0 | 0 |
| COX6A2 | 0.788 | Sphenopalatine_VS_P1_P2_P3_P4_Lumbar_Coeliac_Stellate | 0 | 0 | 0 | 0 | 0 | 0 | 0 | 1 |
| LYSMD3 | 0.787 | Lumbar_Coeliac_VS_P1_P2_P3_P4_Stellate_Sphenopalatine | 0 | 0 | 0 | 0 | 1 | 1 | 0 | 0 |
| NCAPD3 | 0.787 | P1_P2_P3_P4_Lumbar_Coeliac_Sphenopalatine_VS_Stellate | 1 | 1 | 1 | 1 | 1 | 1 | 0 | 1 |
| TMTC2 | 0.786 | P4_VS_P1_P2_P3_Lumbar_Coeliac_Stellate_Sphenopalatine | 0 | 0 | 0 | 1 | 0 | 0 | 0 | 0 |
| SCRT2 | 0.786 | Lumbar_VS_P1_P2_P3_P4_Coeliac_Stellate_Sphenopalatine | 0 | 0 | 0 | 0 | 1 | 0 | 0 | 0 |
| CLU | 0.786 | P1_P2_P3_P4_Coeliac_Sphenopalatine_VS_Lumbar_Stellate | 1 | 1 | 1 | 1 | 0 | 1 | 0 | 1 |
| SAMHD1 | 0.786 | P1_P2_P3_P4_Lumbar_Coeliac_Sphenopalatine_VS_Stellate | 1 | 1 | 1 | 1 | 1 | 1 | 0 | 1 |
| GPN3 | 0.785 | Lumbar_Coeliac_VS_P1_P2_P3_P4_Stellate_Sphenopalatine | 0 | 0 | 0 | 0 | 1 | 1 | 0 | 0 |
| IL17D | 0.785 | Lumbar_Coeliac_VS_P1_P2_P3_P4_Stellate_Sphenopalatine | 0 | 0 | 0 | 0 | 1 | 1 | 0 | 0 |
| FLRT3 | 0.785 | P1_P2_P4_Lumbar_Coeliac_VS_P3_Stellate_Sphenopalatine | 1 | 1 | 0 | 1 | 1 | 1 | 0 | 0 |
| ACOT1 | 0.784 | P4_VS_P1_P2_P3_Lumbar_Coeliac_Stellate_Sphenopalatine | 0 | 0 | 0 | 1 | 0 | 0 | 0 | 0 |
| GM2694 | 0.784 | P1_P2_P4_Lumbar_VS_P3_Coeliac_Stellate_Sphenopalatine | 1 | 1 | 0 | 1 | 1 | 0 | 0 | 0 |
| NRG3 | 0.784 | P1_P4_Lumbar_Coeliac_VS_P2_P3_Stellate_Sphenopalatine | 1 | 0 | 0 | 1 | 1 | 1 | 0 | 0 |
| ASCC3 | 0.783 | P1_P2_P3_P4_Lumbar_Coeliac_Sphenopalatine_VS_Stellate | 1 | 1 | 1 | 1 | 1 | 1 | 0 | 1 |
| HACL1 | 0.782 | P4_VS_P1_P2_P3_Lumbar_Coeliac_Stellate_Sphenopalatine | 0 | 0 | 0 | 1 | 0 | 0 | 0 | 0 |
| TMEM268 | 0.782 | P2_P4_VS_P1_P3_Lumbar_Coeliac_Stellate_Sphenopalatine | 0 | 1 | 0 | 1 | 0 | 0 | 0 | 0 |
| TUBB6 | 0.782 | P1_P2_P3_P4_Sphenopalatine_VS_Lumbar_Coeliac_Stellate | 1 | 1 | 1 | 1 | 0 | 0 | 0 | 1 |
| ADAM9 | 0.782 | P1_P2_P3_P4_Lumbar_Coeliac_Sphenopalatine_VS_Stellate | 1 | 1 | 1 | 1 | 1 | 1 | 0 | 1 |
| PLPPR1 | 0.782 | P1_P2_P3_Lumbar_Coeliac_Stellate_Sphenopalatine_VS_P4 | 1 | 1 | 1 | 0 | 1 | 1 | 1 | 1 |

| | | | | | | | | | | | |
|---|---|---|---|---|---|---|---|---|---|---|---|
| MEX3B | 0.781 | Lumbar_Coeliac_VS_P1_P2_P3_P4_Stellate_Sphenopalatine | 0 | 0 | 0 | 0 | 1 | 1 | 0 | | 0 |
| TRIM62 | 0.781 | P1_P2_P3_P4_Lumbar_Coeliac_Sphenopalatine_VS_Stellate | 1 | 1 | 1 | 1 | 1 | 1 | 0 | | 1 |
| GIGYF1 | 0.781 | P1_P2_P3_P4_Lumbar_Coeliac_Sphenopalatine_VS_Stellate | 1 | 1 | 1 | 1 | 1 | 1 | 0 | | 1 |
| GAREM1 | 0.78 | P1_VS_P2_P3_P4_Lumbar_Coeliac_Stellate_Sphenopalatine | 1 | 0 | 0 | 0 | 0 | 0 | 0 | | 0 |
| RDH10 | 0.78 | P4_VS_P1_P2_P3_Lumbar_Coeliac_Stellate_Sphenopalatine | 0 | 0 | 0 | 1 | 0 | 0 | 0 | | 0 |
| 4930509H03 | 0.78 | Lumbar_Coeliac_VS_P1_P2_P3_P4_Stellate_Sphenopalatine | 0 | 0 | 0 | 0 | 1 | 1 | 0 | | 0 |
| PSG16 | 0.78 | Lumbar_Coeliac_VS_P1_P2_P3_P4_Stellate_Sphenopalatine | 0 | 0 | 0 | 0 | 1 | 1 | 0 | | 0 |
| NFKB2 | 0.78 | Lumbar_Coeliac_VS_P1_P2_P3_P4_Stellate_Sphenopalatine | 0 | 0 | 0 | 0 | 1 | 1 | 0 | | 0 |
| FAM126B | 0.78 | P1_P2_P3_P4_Lumbar_Coeliac_Sphenopalatine_VS_Stellate | 1 | 1 | 1 | 1 | 1 | 1 | 0 | | 1 |
| SLC12A2 | 0.779 | P1_P2_P3_P4_Lumbar_Coeliac_Sphenopalatine_VS_Stellate | 1 | 1 | 1 | 1 | 1 | 1 | 0 | | 1 |
| INSYN1 | 0.779 | P1_P3_P4_Lumbar_Coeliac_Stellate_Sphenopalatine_VS_P2 | 1 | 0 | 1 | 1 | 1 | 1 | 1 | | 1 |
| GM45847 | 0.778 | P4_VS_P1_P2_P3_Lumbar_Coeliac_Stellate_Sphenopalatine | 0 | 0 | 0 | 1 | 0 | 0 | 0 | | 0 |
| 2900026A02 | 0.778 | Lumbar_Coeliac_VS_P1_P2_P3_P4_Stellate_Sphenopalatine | 0 | 0 | 0 | 0 | 1 | 1 | 0 | | 0 |
| SRGAP2 | 0.778 | P1_P2_P3_P4_Lumbar_Coeliac_Sphenopalatine_VS_Stellate | 1 | 1 | 1 | 1 | 1 | 1 | 0 | | 1 |
| TTC12 | 0.777 | Lumbar_Coeliac_VS_P1_P2_P3_P4_Stellate_Sphenopalatine | 0 | 0 | 0 | 0 | 1 | 1 | 0 | | 0 |
| AKAP5 | 0.777 | P1_P3_Lumbar_Coeliac_Sphenopalatine_VS_P2_P4_Stellate | 1 | 0 | 1 | 0 | 1 | 1 | 0 | | 1 |
| CDR1OS | 0.777 | P1_P2_P3_P4_Lumbar_Coeliac_Sphenopalatine_VS_Stellate | 1 | 1 | 1 | 1 | 1 | 1 | 0 | | 1 |
| NFIX | 0.776 | P4_VS_P1_P2_P3_Lumbar_Coeliac_Stellate_Sphenopalatine | 0 | 0 | 0 | 1 | 0 | 0 | 0 | | 0 |
| CACNA1E | 0.776 | P2_P4_VS_P1_P3_Lumbar_Coeliac_Stellate_Sphenopalatine | 0 | 1 | 0 | 1 | 0 | 0 | 0 | | 0 |
| RASAL2 | 0.776 | P1_P2_P3_P4_Lumbar_Coeliac_Sphenopalatine_VS_Stellate | 1 | 1 | 1 | 1 | 1 | 1 | 0 | | 1 |
| RWDD2A | 0.776 | P1_P2_P3_P4_Lumbar_Coeliac_Sphenopalatine_VS_Stellate | 1 | 1 | 1 | 1 | 1 | 1 | 0 | | 1 |
| NUFIP2 | 0.776 | P1_P2_P3_P4_Lumbar_Coeliac_Sphenopalatine_VS_Stellate | 1 | 1 | 1 | 1 | 1 | 1 | 0 | | 1 |
| MDGA2 | 0.776 | P1_P2_P3_P4_Lumbar_Coeliac_Sphenopalatine_VS_Stellate | 1 | 1 | 1 | 1 | 1 | 1 | 0 | | 1 |
| HOXA4 | 0.775 | P4_VS_P1_P2_P3_Lumbar_Coeliac_Stellate_Sphenopalatine | 0 | 0 | 0 | 1 | 0 | 0 | 0 | | 0 |
| sept-05 | 0.775 | Lumbar_Coeliac_VS_P1_P2_P3_P4_Stellate_Sphenopalatine | 0 | 0 | 0 | 0 | 1 | 1 | 0 | | 0 |
| HIP1 | 0.775 | P1_P2_P3_P4_Lumbar_Coeliac_Stellate_VS_Sphenopalatine | 1 | 1 | 1 | 1 | 1 | 1 | 1 | | 0 |
| EPHX4 | 0.774 | P3_Lumbar_Coeliac_Stellate_VS_P1_P2_P4_Sphenopalatine | 0 | 0 | 1 | 0 | 1 | 1 | 1 | | 0 |
| DNAJB12 | 0.774 | P1_P2_P3_P4_Lumbar_Coeliac_Sphenopalatine_VS_Stellate | 1 | 1 | 1 | 1 | 1 | 1 | 0 | | 1 |
| ETV1 | 0.773 | P1_P3_VS_P2_P4_Lumbar_Coeliac_Stellate_Sphenopalatine | 1 | 0 | 1 | 0 | 0 | 0 | 0 | | 0 |
| GM9895 | 0.773 | Lumbar_Coeliac_VS_P1_P2_P3_P4_Stellate_Sphenopalatine | 0 | 0 | 0 | 0 | 1 | 1 | 0 | | 0 |

| | | | | | | | | | | |
|---|---|---|---|---|---|---|---|---|---|---|
| TEX10 | 0.773 | P1_P2_P3_P4_Lumbar_Coeliac_Sphenopalatine_VS_Stellate | 1 | 1 | 1 | 1 | 1 | 1 | 0 | 1 |
| MAN2A2 | 0.773 | P1_P2_P3_P4_Lumbar_Coeliac_Sphenopalatine_VS_Stellate | 1 | 1 | 1 | 1 | 1 | 1 | 0 | 1 |
| RNF180 | 0.772 | P3_VS_P1_P2_P4_Lumbar_Coeliac_Stellate_Sphenopalatine | 0 | 0 | 1 | 0 | 0 | 0 | 0 | 0 |
| GPR137C | 0.772 | P1_P2_P3_P4_Lumbar_Coeliac_Sphenopalatine_VS_Stellate | 1 | 1 | 1 | 1 | 1 | 1 | 0 | 1 |
| LCA5 | 0.771 | Lumbar_Coeliac_VS_P1_P2_P3_P4_Stellate_Sphenopalatine | 0 | 0 | 0 | 0 | 1 | 1 | 0 | 0 |
| FILIP1 | 0.771 | P1_P2_Lumbar_Coeliac_Sphenopalatine_VS_P3_P4_Stellate | 1 | 1 | 0 | 0 | 1 | 1 | 0 | 1 |
| GM4673 | 0.77 | Lumbar_Coeliac_VS_P1_P2_P3_P4_Stellate_Sphenopalatine | 0 | 0 | 0 | 0 | 1 | 1 | 0 | 0 |
| SLC2A12 | 0.77 | P4_Lumbar_Coeliac_VS_P1_P2_P3_Stellate_Sphenopalatine | 0 | 0 | 0 | 1 | 1 | 1 | 0 | 0 |
| PDIA5 | 0.769 | Lumbar_VS_P1_P2_P3_P4_Coeliac_Stellate_Sphenopalatine | 0 | 0 | 0 | 0 | 1 | 0 | 0 | 0 |
| CEP162 | 0.769 | Lumbar_Coeliac_VS_P1_P2_P3_P4_Stellate_Sphenopalatine | 0 | 0 | 0 | 0 | 1 | 1 | 0 | 0 |
| CCDC160 | 0.768 | Coeliac_VS_P1_P2_P3_P4_Lumbar_Stellate_Sphenopalatine | 0 | 0 | 0 | 0 | 0 | 1 | 0 | 0 |
| GM9962 | 0.768 | Sphenopalatine_VS_P1_P2_P3_P4_Lumbar_Coeliac_Stellate | 0 | 0 | 0 | 0 | 0 | 0 | 0 | 1 |
| RASL11B | 0.767 | P4_VS_P1_P2_P3_Lumbar_Coeliac_Stellate_Sphenopalatine | 0 | 0 | 0 | 1 | 0 | 0 | 0 | 0 |
| HSD17B4 | 0.767 | P4_Lumbar_Coeliac_VS_P1_P2_P3_Stellate_Sphenopalatine | 0 | 0 | 0 | 1 | 1 | 1 | 0 | 0 |
| GM42937 | 0.767 | P1_P2_P3_P4_Lumbar_Coeliac_VS_Stellate_Sphenopalatine | 1 | 1 | 1 | 1 | 1 | 1 | 0 | 0 |
| ZBTB11 | 0.767 | P1_P2_P3_P4_Lumbar_Coeliac_Sphenopalatine_VS_Stellate | 1 | 1 | 1 | 1 | 1 | 1 | 0 | 1 |
| KIFC3 | 0.766 | P4_VS_P1_P2_P3_Lumbar_Coeliac_Stellate_Sphenopalatine | 0 | 0 | 0 | 1 | 0 | 0 | 0 | 0 |
| SPTY2D1 | 0.766 | P1_P2_P3_P4_Lumbar_Coeliac_Sphenopalatine_VS_Stellate | 1 | 1 | 1 | 1 | 1 | 1 | 0 | 1 |
| AMPD2 | 0.765 | P1_P2_P3_P4_Lumbar_Coeliac_Sphenopalatine_VS_Stellate | 1 | 1 | 1 | 1 | 1 | 1 | 0 | 1 |
| GPBP1L1 | 0.765 | P1_P2_P3_P4_Lumbar_Coeliac_Sphenopalatine_VS_Stellate | 1 | 1 | 1 | 1 | 1 | 1 | 0 | 1 |
| BRD1 | 0.765 | P1_P2_P3_P4_Lumbar_Coeliac_Sphenopalatine_VS_Stellate | 1 | 1 | 1 | 1 | 1 | 1 | 0 | 1 |
| RNF219 | 0.764 | Lumbar_Coeliac_VS_P1_P2_P3_P4_Stellate_Sphenopalatine | 0 | 0 | 0 | 0 | 1 | 1 | 0 | 0 |
| FER | 0.764 | Lumbar_Coeliac_VS_P1_P2_P3_P4_Stellate_Sphenopalatine | 0 | 0 | 0 | 0 | 1 | 1 | 0 | 0 |
| PCDH7 | 0.764 | P1_P2_P4_Lumbar_Coeliac_Stellate_VS_P3_Sphenopalatine | 1 | 1 | 0 | 1 | 1 | 1 | 1 | 0 |
| PMEPA1 | 0.763 | P1_P2_P4_Lumbar_Coeliac_Sphenopalatine_VS_P3_Stellate | 1 | 1 | 0 | 1 | 1 | 1 | 0 | 1 |
| DUSP5 | 0.762 | P2_VS_P1_P3_P4_Lumbar_Coeliac_Stellate_Sphenopalatine | 0 | 1 | 0 | 0 | 0 | 0 | 0 | 0 |
| NEBL | 0.762 | Lumbar_VS_P1_P2_P3_P4_Coeliac_Stellate_Sphenopalatine | 0 | 0 | 0 | 0 | 1 | 0 | 0 | 0 |
| TNNC1 | 0.762 | Lumbar_VS_P1_P2_P3_P4_Coeliac_Stellate_Sphenopalatine | 0 | 0 | 0 | 0 | 1 | 0 | 0 | 0 |
| RIF1 | 0.762 | Lumbar_Coeliac_VS_P1_P2_P3_P4_Stellate_Sphenopalatine | 0 | 0 | 0 | 0 | 1 | 1 | 0 | 0 |
| ATAD2B | 0.762 | Lumbar_Coeliac_VS_P1_P2_P3_P4_Stellate_Sphenopalatine | 0 | 0 | 0 | 0 | 1 | 1 | 0 | 0 |

| | | | | | | | | | | |
|---|---|---|---|---|---|---|---|---|---|---|
| N4BP1 | 0.762 | P1_P2_P3_P4_Lumbar_Coeliac_Sphenopalatine_VS_Stellate | 1 | 1 | 1 | 1 | 1 | 1 | 0 | 1 |
| ZFP608 | 0.761 | P1_P2_P3_P4_Lumbar_Coeliac_Sphenopalatine_VS_Stellate | 1 | 1 | 1 | 1 | 1 | 1 | 0 | 1 |
| ZFP57 | 0.76 | Lumbar_Coeliac_VS_P1_P2_P3_P4_Stellate_Sphenopalatine | 0 | 0 | 0 | 0 | 1 | 1 | 0 | 0 |
| COL1A2 | 0.76 | P1_P2_P3_P4_Coeliac_Sphenopalatine_VS_Lumbar_Stellate | 1 | 1 | 1 | 1 | 0 | 1 | 0 | 1 |
| CDK8 | 0.76 | P1_P2_P3_P4_Lumbar_Coeliac_Sphenopalatine_VS_Stellate | 1 | 1 | 1 | 1 | 1 | 1 | 0 | 1 |
| ZFP646 | 0.76 | P1_P2_P3_P4_Lumbar_Coeliac_Sphenopalatine_VS_Stellate | 1 | 1 | 1 | 1 | 1 | 1 | 0 | 1 |
| NMRAL1 | 0.759 | Lumbar_Coeliac_VS_P1_P2_P3_P4_Stellate_Sphenopalatine | 0 | 0 | 0 | 0 | 1 | 1 | 0 | 0 |
| TACC2 | 0.759 | P1_P2_P3_P4_Lumbar_Coeliac_Stellate_VS_Sphenopalatine | 1 | 1 | 1 | 1 | 1 | 1 | 1 | 0 |
| IFIT2 | 0.759 | P1_P2_P3_P4_Lumbar_Coeliac_Sphenopalatine_VS_Stellate | 1 | 1 | 1 | 1 | 1 | 1 | 0 | 1 |
| FAM98B | 0.758 | Lumbar_Coeliac_VS_P1_P2_P3_P4_Stellate_Sphenopalatine | 0 | 0 | 0 | 0 | 1 | 1 | 0 | 0 |
| YES1 | 0.758 | Lumbar_Coeliac_VS_P1_P2_P3_P4_Stellate_Sphenopalatine | 0 | 0 | 0 | 0 | 1 | 1 | 0 | 0 |
| ARL14EP | 0.758 | P1_P2_P3_P4_Lumbar_Coeliac_Sphenopalatine_VS_Stellate | 1 | 1 | 1 | 1 | 1 | 1 | 0 | 1 |
| PHACTR4 | 0.757 | P1_P2_P3_P4_Lumbar_Coeliac_Sphenopalatine_VS_Stellate | 1 | 1 | 1 | 1 | 1 | 1 | 0 | 1 |
| BICDL1 | 0.757 | P1_P2_P3_P4_Lumbar_Coeliac_Sphenopalatine_VS_Stellate | 1 | 1 | 1 | 1 | 1 | 1 | 0 | 1 |
| TECPR2 | 0.757 | P1_P2_P3_P4_Lumbar_Coeliac_Sphenopalatine_VS_Stellate | 1 | 1 | 1 | 1 | 1 | 1 | 0 | 1 |
| RASIP1 | 0.756 | Lumbar_VS_P1_P2_P3_P4_Coeliac_Stellate_Sphenopalatine | 0 | 0 | 0 | 0 | 1 | 0 | 0 | 0 |
| ZFP592 | 0.756 | P1_P2_P3_P4_Lumbar_Coeliac_Sphenopalatine_VS_Stellate | 1 | 1 | 1 | 1 | 1 | 1 | 0 | 1 |
| TAFA5 | 0.756 | P1_P2_P3_P4_Lumbar_Coeliac_Sphenopalatine_VS_Stellate | 1 | 1 | 1 | 1 | 1 | 1 | 0 | 1 |
| SP1 | 0.756 | P1_P2_P3_P4_Lumbar_Coeliac_Sphenopalatine_VS_Stellate | 1 | 1 | 1 | 1 | 1 | 1 | 0 | 1 |
| ELP4 | 0.755 | Lumbar_Coeliac_VS_P1_P2_P3_P4_Stellate_Sphenopalatine | 0 | 0 | 0 | 0 | 1 | 1 | 0 | 0 |
| GALNT7 | 0.755 | P1_P2_P3_P4_Lumbar_Coeliac_Sphenopalatine_VS_Stellate | 1 | 1 | 1 | 1 | 1 | 1 | 0 | 1 |
| FAM76B | 0.754 | Lumbar_Coeliac_VS_P1_P2_P3_P4_Stellate_Sphenopalatine | 0 | 0 | 0 | 0 | 1 | 1 | 0 | 0 |
| SGK1 | 0.753 | Lumbar_VS_P1_P2_P3_P4_Coeliac_Stellate_Sphenopalatine | 0 | 0 | 0 | 0 | 1 | 0 | 0 | 0 |
| ZBTB43 | 0.753 | Lumbar_Coeliac_VS_P1_P2_P3_P4_Stellate_Sphenopalatine | 0 | 0 | 0 | 0 | 1 | 1 | 0 | 0 |
| WDR44 | 0.753 | P1_P2_P3_P4_Lumbar_Coeliac_Sphenopalatine_VS_Stellate | 1 | 1 | 1 | 1 | 1 | 1 | 0 | 1 |
| AFF2 | 0.752 | P2_VS_P1_P3_P4_Lumbar_Coeliac_Stellate_Sphenopalatine | 0 | 1 | 0 | 0 | 0 | 0 | 0 | 0 |
| 4732440D04 | 0.752 | Lumbar_Coeliac_VS_P1_P2_P3_P4_Stellate_Sphenopalatine | 0 | 0 | 0 | 0 | 1 | 1 | 0 | 0 |
| TEX29 | 0.751 | P3_P4_VS_P1_P2_Lumbar_Coeliac_Stellate_Sphenopalatine | 0 | 0 | 1 | 1 | 0 | 0 | 0 | 0 |
| NAB1 | 0.751 | P1_P2_P3_P4_Lumbar_Coeliac_Sphenopalatine_VS_Stellate | 1 | 1 | 1 | 1 | 1 | 1 | 0 | 1 |
| ZFYVE9 | 0.751 | P1_P2_P3_P4_Lumbar_Coeliac_Sphenopalatine_VS_Stellate | 1 | 1 | 1 | 1 | 1 | 1 | 0 | 1 |

| | | | | | | | | | | |
|---|---|---|---|---|---|---|---|---|---|---|
| CYFIP1 | 0.751 | P1_P2_P3_P4_Lumbar_Coeliac_Sphenopalatine_VS_Stellate | 1 | 1 | 1 | 1 | 1 | 1 | 0 | 1 |
| TNNT1 | 0.751 | P1_P2_P3_Lumbar_Coeliac_Stellate_Sphenopalatine_VS_P4 | 1 | 1 | 1 | 0 | 1 | 1 | 1 | 1 |
| SPATA5 | 0.75 | Lumbar_Coeliac_VS_P1_P2_P3_P4_Stellate_Sphenopalatine | 0 | 0 | 0 | 0 | 1 | 1 | 0 | 0 |
| SBNO2 | 0.75 | P1_Sphenopalatine_VS_P2_P3_P4_Lumbar_Coeliac_Stellate | 1 | 0 | 0 | 0 | 0 | 0 | 0 | 1 |
| SPIRE2 | 0.75 | P1_P2_P3_P4_Lumbar_Coeliac_Sphenopalatine_VS_Stellate | 1 | 1 | 1 | 1 | 1 | 1 | 0 | 1 |
| EZH1 | 0.75 | P1_P2_P3_P4_Lumbar_Coeliac_Sphenopalatine_VS_Stellate | 1 | 1 | 1 | 1 | 1 | 1 | 0 | 1 |
| DSG2 | 0.749 | Lumbar_VS_P1_P2_P3_P4_Coeliac_Stellate_Sphenopalatine | 0 | 0 | 0 | 0 | 1 | 0 | 0 | 0 |
| TNS1 | 0.749 | P1_P2_P3_P4_Lumbar_Coeliac_Sphenopalatine_VS_Stellate | 1 | 1 | 1 | 1 | 1 | 1 | 0 | 1 |
| AGO3 | 0.749 | P1_P2_P3_P4_Lumbar_Coeliac_Sphenopalatine_VS_Stellate | 1 | 1 | 1 | 1 | 1 | 1 | 0 | 1 |
| PGM3 | 0.748 | Lumbar_Coeliac_VS_P1_P2_P3_P4_Stellate_Sphenopalatine | 0 | 0 | 0 | 0 | 1 | 1 | 0 | 0 |
| EDRF1 | 0.748 | P4_Lumbar_Coeliac_VS_P1_P2_P3_Stellate_Sphenopalatine | 0 | 0 | 0 | 1 | 1 | 1 | 0 | 0 |
| GM17066 | 0.748 | P1_P2_P3_P4_Lumbar_Coeliac_Sphenopalatine_VS_Stellate | 1 | 1 | 1 | 1 | 1 | 1 | 0 | 1 |
| SCRN2 | 0.747 | Lumbar_VS_P1_P2_P3_P4_Coeliac_Stellate_Sphenopalatine | 0 | 0 | 0 | 0 | 1 | 0 | 0 | 0 |
| BACH1 | 0.747 | Lumbar_Coeliac_VS_P1_P2_P3_P4_Stellate_Sphenopalatine | 0 | 0 | 0 | 0 | 1 | 1 | 0 | 0 |
| LGI2 | 0.747 | P1_P2_P3_P4_Lumbar_Coeliac_Sphenopalatine_VS_Stellate | 1 | 1 | 1 | 1 | 1 | 1 | 0 | 1 |
| MED13 | 0.746 | P1_P2_P3_P4_Lumbar_Coeliac_Sphenopalatine_VS_Stellate | 1 | 1 | 1 | 1 | 1 | 1 | 0 | 1 |
| DOT1L | 0.745 | P1_P2_P3_P4_Lumbar_Coeliac_Sphenopalatine_VS_Stellate | 1 | 1 | 1 | 1 | 1 | 1 | 0 | 1 |
| RAB12 | 0.745 | P1_P2_P3_P4_Lumbar_Coeliac_Sphenopalatine_VS_Stellate | 1 | 1 | 1 | 1 | 1 | 1 | 0 | 1 |
| PHKA2 | 0.745 | P1_P2_P3_P4_Lumbar_Coeliac_Sphenopalatine_VS_Stellate | 1 | 1 | 1 | 1 | 1 | 1 | 0 | 1 |
| GM35147 | 0.744 | P2_VS_P1_P3_P4_Lumbar_Coeliac_Stellate_Sphenopalatine | 0 | 1 | 0 | 0 | 0 | 0 | 0 | 0 |
| ETV6 | 0.744 | P1_P2_P3_P4_Lumbar_Coeliac_VS_Stellate_Sphenopalatine | 1 | 1 | 1 | 1 | 1 | 1 | 0 | 0 |
| ELOA | 0.744 | P1_P2_P3_P4_Lumbar_Coeliac_Sphenopalatine_VS_Stellate | 1 | 1 | 1 | 1 | 1 | 1 | 0 | 1 |
| MNT | 0.744 | P1_P2_P3_P4_Lumbar_Coeliac_Sphenopalatine_VS_Stellate | 1 | 1 | 1 | 1 | 1 | 1 | 0 | 1 |
| NEAT1 | 0.744 | P1_P2_P3_P4_Lumbar_Coeliac_Sphenopalatine_VS_Stellate | 1 | 1 | 1 | 1 | 1 | 1 | 0 | 1 |
| STK26 | 0.743 | Coeliac_VS_P1_P2_P3_P4_Lumbar_Stellate_Sphenopalatine | 0 | 0 | 0 | 0 | 0 | 1 | 0 | 0 |
| KCNT1 | 0.743 | P3_P4_Lumbar_Coeliac_Stellate_VS_P1_P2_Sphenopalatine | 0 | 0 | 1 | 1 | 1 | 1 | 1 | 0 |
| ZFP62 | 0.743 | P1_P2_P3_P4_Lumbar_Coeliac_Sphenopalatine_VS_Stellate | 1 | 1 | 1 | 1 | 1 | 1 | 0 | 1 |
| DBF4 | 0.742 | Lumbar_Coeliac_VS_P1_P2_P3_P4_Stellate_Sphenopalatine | 0 | 0 | 0 | 0 | 1 | 1 | 0 | 0 |
| ZFP830 | 0.742 | Lumbar_Coeliac_VS_P1_P2_P3_P4_Stellate_Sphenopalatine | 0 | 0 | 0 | 0 | 1 | 1 | 0 | 0 |
| MMP16 | 0.742 | P1_P2_P3_P4_Lumbar_Coeliac_Sphenopalatine_VS_Stellate | 1 | 1 | 1 | 1 | 1 | 1 | 0 | 1 |

| | | | | | | | | | | |
|---|---|---|---|---|---|---|---|---|---|---|
| PRR14L | 0.742 | P1_P2_P3_P4_Lumbar_Coeliac_Sphenopalatine_VS_Stellate | 1 | 1 | 1 | 1 | 1 | 1 | 0 | 1 |
| SCPEP1 | 0.742 | P1_P2_P3_P4_Lumbar_Coeliac_Sphenopalatine_VS_Stellate | 1 | 1 | 1 | 1 | 1 | 1 | 0 | 1 |
| GM4258 | 0.741 | P3_Lumbar_Coeliac_VS_P1_P2_P4_Stellate_Sphenopalatine | 0 | 0 | 1 | 0 | 1 | 1 | 0 | 0 |
| GRIP1 | 0.741 | P1_P2_P4_Lumbar_Coeliac_Sphenopalatine_VS_P3_Stellate | 1 | 1 | 0 | 1 | 1 | 1 | 0 | 1 |
| THEM6 | 0.741 | P1_P2_P3_P4_Lumbar_Coeliac_Sphenopalatine_VS_Stellate | 1 | 1 | 1 | 1 | 1 | 1 | 0 | 1 |
| RGS20 | 0.74 | Lumbar_VS_P1_P2_P3_P4_Coeliac_Stellate_Sphenopalatine | 0 | 0 | 0 | 0 | 1 | 0 | 0 | 0 |
| GJC1 | 0.74 | Lumbar_Coeliac_VS_P1_P2_P3_P4_Stellate_Sphenopalatine | 0 | 0 | 0 | 0 | 1 | 1 | 0 | 0 |
| PHLPP2 | 0.74 | P1_P2_P3_P4_Lumbar_Coeliac_Sphenopalatine_VS_Stellate | 1 | 1 | 1 | 1 | 1 | 1 | 0 | 1 |
| PAXBP1 | 0.739 | Lumbar_Coeliac_VS_P1_P2_P3_P4_Stellate_Sphenopalatine | 0 | 0 | 0 | 0 | 1 | 1 | 0 | 0 |
| GM10419 | 0.739 | P1_P2_P4_Sphenopalatine_VS_P3_Lumbar_Coeliac_Stellate | 1 | 1 | 0 | 1 | 0 | 0 | 0 | 1 |
| NEGR1 | 0.739 | P1_P2_P3_P4_Lumbar_Coeliac_Sphenopalatine_VS_Stellate | 1 | 1 | 1 | 1 | 1 | 1 | 0 | 1 |
| ARRB1 | 0.739 | P1_P2_P3_P4_Lumbar_Coeliac_Sphenopalatine_VS_Stellate | 1 | 1 | 1 | 1 | 1 | 1 | 0 | 1 |
| POLK | 0.739 | P1_P2_P3_P4_Lumbar_Coeliac_Sphenopalatine_VS_Stellate | 1 | 1 | 1 | 1 | 1 | 1 | 0 | 1 |
| RNF138RT1 | 0.738 | P3_Lumbar_Coeliac_VS_P1_P2_P4_Stellate_Sphenopalatine | 0 | 0 | 1 | 0 | 1 | 1 | 0 | 0 |
| AKAIN1 | 0.738 | P4_Lumbar_Coeliac_Stellate_VS_P1_P2_P3_Sphenopalatine | 0 | 0 | 0 | 1 | 1 | 1 | 1 | 0 |
| PCDH19 | 0.738 | P1_P4_Lumbar_Coeliac_Sphenopalatine_VS_P2_P3_Stellate | 1 | 0 | 0 | 1 | 1 | 1 | 0 | 1 |
| ZFP423 | 0.738 | P1_P2_P3_P4_Lumbar_Coeliac_Sphenopalatine_VS_Stellate | 1 | 1 | 1 | 1 | 1 | 1 | 0 | 1 |
| ETAA1 | 0.738 | P1_P2_P3_P4_Lumbar_Coeliac_Sphenopalatine_VS_Stellate | 1 | 1 | 1 | 1 | 1 | 1 | 0 | 1 |
| SDC1 | 0.737 | P1_VS_P2_P3_P4_Lumbar_Coeliac_Stellate_Sphenopalatine | 1 | 0 | 0 | 0 | 0 | 0 | 0 | 0 |
| GM19345 | 0.737 | Sphenopalatine_VS_P1_P2_P3_P4_Lumbar_Coeliac_Stellate | 0 | 0 | 0 | 0 | 0 | 0 | 0 | 1 |
| HIVEP3 | 0.737 | P1_P2_P3_P4_Lumbar_Coeliac_Sphenopalatine_VS_Stellate | 1 | 1 | 1 | 1 | 1 | 1 | 0 | 1 |
| ABCA5 | 0.737 | P1_P2_P3_P4_Lumbar_Coeliac_Sphenopalatine_VS_Stellate | 1 | 1 | 1 | 1 | 1 | 1 | 0 | 1 |
| ZDBF2 | 0.736 | P1_P2_P3_P4_Lumbar_Coeliac_Sphenopalatine_VS_Stellate | 1 | 1 | 1 | 1 | 1 | 1 | 0 | 1 |
| ATR | 0.736 | P1_P2_P3_P4_Lumbar_Coeliac_Sphenopalatine_VS_Stellate | 1 | 1 | 1 | 1 | 1 | 1 | 0 | 1 |
| CLVS2 | 0.735 | Lumbar_VS_P1_P2_P3_P4_Coeliac_Stellate_Sphenopalatine | 0 | 0 | 0 | 0 | 1 | 0 | 0 | 0 |
| MTERF3 | 0.735 | Coeliac_VS_P1_P2_P3_P4_Lumbar_Stellate_Sphenopalatine | 0 | 0 | 0 | 0 | 0 | 1 | 0 | 0 |
| KCNH8 | 0.735 | P1_P3_Lumbar_Coeliac_VS_P2_P4_Stellate_Sphenopalatine | 1 | 0 | 1 | 0 | 1 | 1 | 0 | 0 |
| KCNK4 | 0.735 | P4_Sphenopalatine_VS_P1_P2_P3_Lumbar_Coeliac_Stellate | 0 | 0 | 0 | 1 | 0 | 0 | 0 | 1 |
| LTB | 0.735 | P1_P2_Lumbar_Coeliac_Sphenopalatine_VS_P3_P4_Stellate | 1 | 1 | 0 | 0 | 1 | 1 | 0 | 1 |
| TAOK2 | 0.735 | P1_P2_P3_P4_Lumbar_Coeliac_Sphenopalatine_VS_Stellate | 1 | 1 | 1 | 1 | 1 | 1 | 0 | 1 |

| | | | | | | | | | | |
|---|---|---|---|---|---|---|---|---|---|---|
| XRN1 | 0.735 | P1_P2_P3_P4_Lumbar_Coeliac_Sphenopalatine_VS_Stellate | 1 | 1 | 1 | 1 | 1 | 1 | 0 | 1 |
| SOS2 | 0.735 | P1_P2_P3_P4_Lumbar_Coeliac_Sphenopalatine_VS_Stellate | 1 | 1 | 1 | 1 | 1 | 1 | 0 | 1 |
| 2310040G24 | 0.734 | Lumbar_VS_P1_P2_P3_P4_Coeliac_Stellate_Sphenopalatine | 0 | 0 | 0 | 0 | 1 | 0 | 0 | 0 |
| TASOR2 | 0.734 | Lumbar_Coeliac_VS_P1_P2_P3_P4_Stellate_Sphenopalatine | 0 | 0 | 0 | 0 | 1 | 1 | 0 | 0 |
| PARG | 0.734 | P1_P2_P3_P4_Lumbar_Coeliac_Sphenopalatine_VS_Stellate | 1 | 1 | 1 | 1 | 1 | 1 | 0 | 1 |
| KIF26A | 0.734 | P1_P2_P3_Lumbar_Coeliac_Stellate_Sphenopalatine_VS_P4 | 1 | 1 | 1 | 0 | 1 | 1 | 1 | 1 |
| RASD2 | 0.733 | P3_VS_P1_P2_P4_Lumbar_Coeliac_Stellate_Sphenopalatine | 0 | 0 | 1 | 0 | 0 | 0 | 0 | 0 |
| DUSP10 | 0.733 | P3_Lumbar_Coeliac_Stellate_VS_P1_P2_P4_Sphenopalatine | 0 | 0 | 1 | 0 | 1 | 1 | 1 | 0 |
| TBC1D12 | 0.733 | P1_P2_P3_P4_Lumbar_Coeliac_Sphenopalatine_VS_Stellate | 1 | 1 | 1 | 1 | 1 | 1 | 0 | 1 |
| LCOR | 0.732 | Lumbar_Coeliac_VS_P1_P2_P3_P4_Stellate_Sphenopalatine | 0 | 0 | 0 | 0 | 1 | 1 | 0 | 0 |
| STIM1 | 0.732 | P1_P2_P3_P4_Lumbar_Coeliac_Sphenopalatine_VS_Stellate | 1 | 1 | 1 | 1 | 1 | 1 | 0 | 1 |
| TLE2 | 0.731 | P1_P2_P3_Lumbar_Coeliac_Sphenopalatine_VS_P4_Stellate | 1 | 1 | 1 | 0 | 1 | 1 | 0 | 1 |
| HS3ST4 | 0.73 | P2_VS_P1_P3_P4_Lumbar_Coeliac_Stellate_Sphenopalatine | 0 | 1 | 0 | 0 | 0 | 0 | 0 | 0 |
| GRM4 | 0.73 | P2_VS_P1_P3_P4_Lumbar_Coeliac_Stellate_Sphenopalatine | 0 | 1 | 0 | 0 | 0 | 0 | 0 | 0 |
| WFS1 | 0.73 | P1_P2_P3_P4_Lumbar_Coeliac_Sphenopalatine_VS_Stellate | 1 | 1 | 1 | 1 | 1 | 1 | 0 | 1 |
| 2810032G03 | 0.729 | Lumbar_VS_P1_P2_P3_P4_Coeliac_Stellate_Sphenopalatine | 0 | 0 | 0 | 0 | 1 | 0 | 0 | 0 |
| PHLDB2 | 0.729 | Lumbar_VS_P1_P2_P3_P4_Coeliac_Stellate_Sphenopalatine | 0 | 0 | 0 | 0 | 1 | 0 | 0 | 0 |
| PPP1R12B | 0.729 | P1_P2_P3_P4_Lumbar_Coeliac_Sphenopalatine_VS_Stellate | 1 | 1 | 1 | 1 | 1 | 1 | 0 | 1 |
| DNAJC1 | 0.729 | P1_P2_P3_P4_Lumbar_Coeliac_Sphenopalatine_VS_Stellate | 1 | 1 | 1 | 1 | 1 | 1 | 0 | 1 |
| IL11RA1 | 0.728 | P1_P2_P3_P4_Sphenopalatine_VS_Lumbar_Coeliac_Stellate | 1 | 1 | 1 | 1 | 0 | 0 | 0 | 1 |
| PHLPP1 | 0.728 | P1_P2_P3_P4_Lumbar_Coeliac_Sphenopalatine_VS_Stellate | 1 | 1 | 1 | 1 | 1 | 1 | 0 | 1 |
| FNBP1 | 0.728 | P1_P2_P3_P4_Lumbar_Coeliac_Sphenopalatine_VS_Stellate | 1 | 1 | 1 | 1 | 1 | 1 | 0 | 1 |
| ZFP362 | 0.728 | P1_P2_P3_P4_Lumbar_Coeliac_Sphenopalatine_VS_Stellate | 1 | 1 | 1 | 1 | 1 | 1 | 0 | 1 |
| TJP1 | 0.728 | P1_P2_P3_P4_Lumbar_Coeliac_Sphenopalatine_VS_Stellate | 1 | 1 | 1 | 1 | 1 | 1 | 0 | 1 |
| GM43581 | 0.727 | P2_VS_P1_P3_P4_Lumbar_Coeliac_Stellate_Sphenopalatine | 0 | 1 | 0 | 0 | 0 | 0 | 0 | 0 |
| SMARCAD1 | 0.727 | Lumbar_Coeliac_VS_P1_P2_P3_P4_Stellate_Sphenopalatine | 0 | 0 | 0 | 0 | 1 | 1 | 0 | 0 |
| TNFSF12 | 0.727 | P1_P2_P3_P4_Lumbar_Coeliac_Sphenopalatine_VS_Stellate | 1 | 1 | 1 | 1 | 1 | 1 | 0 | 1 |
| COX6B2 | 0.726 | P2_VS_P1_P3_P4_Lumbar_Coeliac_Stellate_Sphenopalatine | 0 | 1 | 0 | 0 | 0 | 0 | 0 | 0 |
| WDR35 | 0.726 | P1_P2_P3_P4_Lumbar_Coeliac_Sphenopalatine_VS_Stellate | 1 | 1 | 1 | 1 | 1 | 1 | 0 | 1 |
| RPGRIP1L | 0.725 | Lumbar_Coeliac_VS_P1_P2_P3_P4_Stellate_Sphenopalatine | 0 | 0 | 0 | 0 | 1 | 1 | 0 | 0 |

| | | | | | | | | | | |
|---|---|---|---|---|---|---|---|---|---|---|
| 2610037D02I | 0.725 | P1_P2_P3_P4_Lumbar_Coeliac_Sphenopalatine_VS_Stellate | 1 | 1 | 1 | 1 | 1 | 1 | 0 | 1 |
| RASSF3 | 0.724 | P3_VS_P1_P2_P4_Lumbar_Coeliac_Stellate_Sphenopalatine | 0 | 0 | 1 | 0 | 0 | 0 | 0 | 0 |
| COL5A3 | 0.724 | Lumbar_VS_P1_P2_P3_P4_Coeliac_Stellate_Sphenopalatine | 0 | 0 | 0 | 0 | 1 | 0 | 0 | 0 |
| ZFP395 | 0.724 | P1_P2_P3_P4_Lumbar_Coeliac_VS_Stellate_Sphenopalatine | 1 | 1 | 1 | 1 | 1 | 1 | 0 | 0 |
| FOXN2 | 0.724 | P1_P2_P3_P4_Lumbar_Coeliac_VS_Stellate_Sphenopalatine | 1 | 1 | 1 | 1 | 1 | 1 | 0 | 0 |
| ASXL3 | 0.724 | P1_P2_P3_P4_Lumbar_Coeliac_Sphenopalatine_VS_Stellate | 1 | 1 | 1 | 1 | 1 | 1 | 0 | 1 |
| GRIN2D | 0.723 | P2_P4_VS_P1_P3_Lumbar_Coeliac_Stellate_Sphenopalatine | 0 | 1 | 0 | 1 | 0 | 0 | 0 | 0 |
| FGFR1 | 0.723 | P1_P2_P3_P4_Sphenopalatine_VS_Lumbar_Coeliac_Stellate | 1 | 1 | 1 | 1 | 0 | 0 | 0 | 1 |
| IARS2 | 0.723 | P1_P2_P3_P4_Lumbar_Coeliac_Sphenopalatine_VS_Stellate | 1 | 1 | 1 | 1 | 1 | 1 | 0 | 1 |
| TSC22D2 | 0.723 | P1_P2_P3_P4_Lumbar_Coeliac_Sphenopalatine_VS_Stellate | 1 | 1 | 1 | 1 | 1 | 1 | 0 | 1 |
| FRRS1L | 0.723 | P1_P2_P3_P4_Lumbar_Coeliac_Sphenopalatine_VS_Stellate | 1 | 1 | 1 | 1 | 1 | 1 | 0 | 1 |
| SLF1 | 0.723 | P1_P2_P3_P4_Lumbar_Coeliac_Sphenopalatine_VS_Stellate | 1 | 1 | 1 | 1 | 1 | 1 | 0 | 1 |
| SMC2 | 0.722 | Lumbar_Coeliac_VS_P1_P2_P3_P4_Stellate_Sphenopalatine | 0 | 0 | 0 | 0 | 1 | 1 | 0 | 0 |
| ZC3H12C | 0.722 | Lumbar_Coeliac_VS_P1_P2_P3_P4_Stellate_Sphenopalatine | 0 | 0 | 0 | 0 | 1 | 1 | 0 | 0 |
| CAVIN4 | 0.722 | P2_P3_Lumbar_Coeliac_VS_P1_P4_Stellate_Sphenopalatine | 0 | 1 | 1 | 0 | 1 | 1 | 0 | 0 |
| WDR92 | 0.721 | Lumbar_Coeliac_VS_P1_P2_P3_P4_Stellate_Sphenopalatine | 0 | 0 | 0 | 0 | 1 | 1 | 0 | 0 |
| SRBD1 | 0.721 | Lumbar_Coeliac_VS_P1_P2_P3_P4_Stellate_Sphenopalatine | 0 | 0 | 0 | 0 | 1 | 1 | 0 | 0 |
| RMND5A | 0.721 | P1_P2_P3_P4_Lumbar_Coeliac_Sphenopalatine_VS_Stellate | 1 | 1 | 1 | 1 | 1 | 1 | 0 | 1 |
| VWA5A | 0.721 | P1_P2_P3_P4_Lumbar_Coeliac_Sphenopalatine_VS_Stellate | 1 | 1 | 1 | 1 | 1 | 1 | 0 | 1 |
| 6720427I07R | 0.721 | P1_P2_P3_P4_Lumbar_Coeliac_Sphenopalatine_VS_Stellate | 1 | 1 | 1 | 1 | 1 | 1 | 0 | 1 |
| CTXN2 | 0.721 | P1_P3_P4_Lumbar_Coeliac_Stellate_Sphenopalatine_VS_P2 | 1 | 0 | 1 | 1 | 1 | 1 | 1 | 1 |
| TEX9 | 0.72 | Lumbar_Coeliac_VS_P1_P2_P3_P4_Stellate_Sphenopalatine | 0 | 0 | 0 | 0 | 1 | 1 | 0 | 0 |
| PAK7 | 0.719 | Lumbar_Coeliac_VS_P1_P2_P3_P4_Stellate_Sphenopalatine | 0 | 0 | 0 | 0 | 1 | 1 | 0 | 0 |
| TRPS1 | 0.719 | P1_P4_Lumbar_Coeliac_VS_P2_P3_Stellate_Sphenopalatine | 1 | 0 | 0 | 1 | 1 | 1 | 0 | 0 |
| BEND7 | 0.719 | P1_P2_P3_P4_Lumbar_Coeliac_Sphenopalatine_VS_Stellate | 1 | 1 | 1 | 1 | 1 | 1 | 0 | 1 |
| SMURF2 | 0.719 | P1_P2_P3_P4_Lumbar_Coeliac_Sphenopalatine_VS_Stellate | 1 | 1 | 1 | 1 | 1 | 1 | 0 | 1 |
| TENM2 | 0.719 | P1_P2_P4_Lumbar_Coeliac_Stellate_Sphenopalatine_VS_P3 | 1 | 1 | 0 | 1 | 1 | 1 | 1 | 1 |
| RAB3IL1 | 0.718 | Lumbar_VS_P1_P2_P3_P4_Coeliac_Stellate_Sphenopalatine | 0 | 0 | 0 | 0 | 1 | 0 | 0 | 0 |
| ZHX3 | 0.718 | P1_P2_P3_P4_Lumbar_Coeliac_Sphenopalatine_VS_Stellate | 1 | 1 | 1 | 1 | 1 | 1 | 0 | 1 |
| VPS33B | 0.718 | P1_P2_P3_P4_Lumbar_Coeliac_Sphenopalatine_VS_Stellate | 1 | 1 | 1 | 1 | 1 | 1 | 0 | 1 |

| | | | | | | | | | | |
|---|---|---|---|---|---|---|---|---|---|---|
| CCDC66 | 0.718 | P1_P2_P3_P4_Lumbar_Coeliac_Sphenopalatine_VS_Stellate | 1 | 1 | 1 | 1 | 1 | 1 | 0 | 1 |
| ESCO1 | 0.717 | Lumbar_Coeliac_VS_P1_P2_P3_P4_Stellate_Sphenopalatine | 0 | 0 | 0 | 0 | 1 | 1 | 0 | 0 |
| FGD1 | 0.717 | P1_P2_P3_P4_Lumbar_Coeliac_Sphenopalatine_VS_Stellate | 1 | 1 | 1 | 1 | 1 | 1 | 0 | 1 |
| TRIP10 | 0.716 | Lumbar_VS_P1_P2_P3_P4_Coeliac_Stellate_Sphenopalatine | 0 | 0 | 0 | 0 | 1 | 0 | 0 | 0 |
| RAB3GAP1 | 0.716 | P1_P2_P3_P4_Lumbar_Coeliac_Sphenopalatine_VS_Stellate | 1 | 1 | 1 | 1 | 1 | 1 | 0 | 1 |
| PRKDC | 0.716 | P1_P2_P3_P4_Lumbar_Coeliac_Sphenopalatine_VS_Stellate | 1 | 1 | 1 | 1 | 1 | 1 | 0 | 1 |
| CAMK4 | 0.716 | P1_P2_P3_P4_Lumbar_Coeliac_Sphenopalatine_VS_Stellate | 1 | 1 | 1 | 1 | 1 | 1 | 0 | 1 |
| PPFIA4 | 0.715 | P1_P2_P3_P4_Lumbar_Coeliac_Sphenopalatine_VS_Stellate | 1 | 1 | 1 | 1 | 1 | 1 | 0 | 1 |
| DLG1 | 0.715 | P1_P2_P3_P4_Lumbar_Coeliac_Sphenopalatine_VS_Stellate | 1 | 1 | 1 | 1 | 1 | 1 | 0 | 1 |
| PANK4 | 0.714 | P1_P2_P3_P4_Lumbar_Coeliac_Sphenopalatine_VS_Stellate | 1 | 1 | 1 | 1 | 1 | 1 | 0 | 1 |
| DERA | 0.713 | P3_P4_Lumbar_Coeliac_VS_P1_P2_Stellate_Sphenopalatine | 0 | 0 | 1 | 1 | 1 | 1 | 0 | 0 |
| GM16701.1 | 0.713 | P1_P2_P3_P4_Lumbar_Coeliac_VS_Stellate_Sphenopalatine | 1 | 1 | 1 | 1 | 1 | 1 | 0 | 0 |
| SORT1 | 0.713 | P1_P2_P3_P4_Lumbar_Coeliac_Sphenopalatine_VS_Stellate | 1 | 1 | 1 | 1 | 1 | 1 | 0 | 1 |
| CLEC2L | 0.713 | P1_P2_P3_P4_Lumbar_Coeliac_Sphenopalatine_VS_Stellate | 1 | 1 | 1 | 1 | 1 | 1 | 0 | 1 |
| SMG6 | 0.713 | P1_P2_P3_P4_Lumbar_Coeliac_Sphenopalatine_VS_Stellate | 1 | 1 | 1 | 1 | 1 | 1 | 0 | 1 |
| GJD2 | 0.713 | P1_P3_P4_Lumbar_Coeliac_Stellate_Sphenopalatine_VS_P2 | 1 | 0 | 1 | 1 | 1 | 1 | 1 | 1 |
| BDP1 | 0.712 | P1_P2_P3_P4_Lumbar_Coeliac_Sphenopalatine_VS_Stellate | 1 | 1 | 1 | 1 | 1 | 1 | 0 | 1 |
| USP3 | 0.711 | Lumbar_Coeliac_VS_P1_P2_P3_P4_Stellate_Sphenopalatine | 0 | 0 | 0 | 0 | 1 | 1 | 0 | 0 |
| ABCC9 | 0.711 | Sphenopalatine_VS_P1_P2_P3_P4_Lumbar_Coeliac_Stellate | 0 | 0 | 0 | 0 | 0 | 0 | 0 | 1 |
| DSEL | 0.711 | P1_P2_P3_P4_Lumbar_Coeliac_Sphenopalatine_VS_Stellate | 1 | 1 | 1 | 1 | 1 | 1 | 0 | 1 |
| FRMD3 | 0.711 | P1_P2_P3_P4_Lumbar_Coeliac_Sphenopalatine_VS_Stellate | 1 | 1 | 1 | 1 | 1 | 1 | 0 | 1 |
| ZFP36 | 0.71 | P1_P2_P4_VS_P3_Lumbar_Coeliac_Stellate_Sphenopalatine | 1 | 1 | 0 | 1 | 0 | 0 | 0 | 0 |
| TSHZ2 | 0.71 | P1_P2_P3_Lumbar_Coeliac_Sphenopalatine_VS_P4_Stellate | 1 | 1 | 1 | 0 | 1 | 1 | 0 | 1 |
| CPNE8 | 0.709 | P1_VS_P2_P3_P4_Lumbar_Coeliac_Stellate_Sphenopalatine | 1 | 0 | 0 | 0 | 0 | 0 | 0 | 0 |
| ZFP658 | 0.709 | Lumbar_Coeliac_VS_P1_P2_P3_P4_Stellate_Sphenopalatine | 0 | 0 | 0 | 0 | 1 | 1 | 0 | 0 |
| FBXO30 | 0.709 | Lumbar_Coeliac_VS_P1_P2_P3_P4_Stellate_Sphenopalatine | 0 | 0 | 0 | 0 | 1 | 1 | 0 | 0 |
| CHPT1 | 0.709 | P1_P2_P3_P4_Lumbar_Coeliac_VS_Stellate_Sphenopalatine | 1 | 1 | 1 | 1 | 1 | 1 | 0 | 0 |
| IPMK | 0.708 | P1_P2_P3_P4_Lumbar_Coeliac_Sphenopalatine_VS_Stellate | 1 | 1 | 1 | 1 | 1 | 1 | 0 | 1 |
| GCHFR | 0.707 | P3_VS_P1_P2_P4_Lumbar_Coeliac_Stellate_Sphenopalatine | 0 | 0 | 1 | 0 | 0 | 0 | 0 | 0 |
| JPH1 | 0.707 | Lumbar_Coeliac_VS_P1_P2_P3_P4_Stellate_Sphenopalatine | 0 | 0 | 0 | 0 | 1 | 1 | 0 | 0 |

| | | | | | | | | | | |
|---|---|---|---|---|---|---|---|---|---|---|
| FBXO41 | 0.707 | P1_P2_P3_P4_Lumbar_Coeliac_Sphenopalatine_VS_Stellate | 1 | 1 | 1 | 1 | 1 | 1 | 0 | 1 |
| NCKAP5L | 0.707 | P1_P2_P3_P4_Lumbar_Coeliac_Sphenopalatine_VS_Stellate | 1 | 1 | 1 | 1 | 1 | 1 | 0 | 1 |
| ZDHHC22 | 0.706 | P4_Sphenopalatine_VS_P1_P2_P3_Lumbar_Coeliac_Stellate | 0 | 0 | 0 | 1 | 0 | 0 | 0 | 1 |
| PPFIA3 | 0.706 | P1_P2_P3_P4_Lumbar_Coeliac_Sphenopalatine_VS_Stellate | 1 | 1 | 1 | 1 | 1 | 1 | 0 | 1 |
| TEAD1 | 0.705 | P1_P2_P3_P4_Lumbar_Coeliac_Sphenopalatine_VS_Stellate | 1 | 1 | 1 | 1 | 1 | 1 | 0 | 1 |
| WAPL | 0.705 | P1_P2_P3_P4_Lumbar_Coeliac_Sphenopalatine_VS_Stellate | 1 | 1 | 1 | 1 | 1 | 1 | 0 | 1 |
| A4GALT | 0.705 | P1_P2_P3_P4_Lumbar_Coeliac_Sphenopalatine_VS_Stellate | 1 | 1 | 1 | 1 | 1 | 1 | 0 | 1 |
| DDX3Y | 0.705 | P1_P2_P3_P4_Lumbar_Coeliac_Sphenopalatine_VS_Stellate | 1 | 1 | 1 | 1 | 1 | 1 | 0 | 1 |
| TATDN2 | 0.704 | P1_P2_P3_P4_Lumbar_Coeliac_Sphenopalatine_VS_Stellate | 1 | 1 | 1 | 1 | 1 | 1 | 0 | 1 |
| LZIC | 0.703 | Lumbar_Coeliac_VS_P1_P2_P3_P4_Stellate_Sphenopalatine | 0 | 0 | 0 | 0 | 1 | 1 | 0 | 0 |
| KCNMB4OS2 | 0.703 | Lumbar_Coeliac_VS_P1_P2_P3_P4_Stellate_Sphenopalatine | 0 | 0 | 0 | 0 | 1 | 1 | 0 | 0 |
| GPC1 | 0.703 | P1_P2_P3_P4_Lumbar_Coeliac_Sphenopalatine_VS_Stellate | 1 | 1 | 1 | 1 | 1 | 1 | 0 | 1 |
| HS2ST1 | 0.703 | P1_P2_P3_P4_Lumbar_Coeliac_Sphenopalatine_VS_Stellate | 1 | 1 | 1 | 1 | 1 | 1 | 0 | 1 |
| SIK3 | 0.703 | P1_P2_P3_P4_Lumbar_Coeliac_Sphenopalatine_VS_Stellate | 1 | 1 | 1 | 1 | 1 | 1 | 0 | 1 |
| SDC2 | 0.702 | P1_P2_P3_P4_Lumbar_Coeliac_Stellate_VS_Sphenopalatine | 1 | 1 | 1 | 1 | 1 | 1 | 1 | 0 |
| DCBLD2 | 0.702 | P1_P2_P4_Lumbar_Coeliac_Sphenopalatine_VS_P3_Stellate | 1 | 1 | 0 | 1 | 1 | 1 | 0 | 1 |
| WWP1 | 0.702 | P1_P2_P3_P4_Lumbar_Coeliac_Sphenopalatine_VS_Stellate | 1 | 1 | 1 | 1 | 1 | 1 | 0 | 1 |
| SEMA4C | 0.701 | Lumbar_VS_P1_P2_P3_P4_Coeliac_Stellate_Sphenopalatine | 0 | 0 | 0 | 0 | 1 | 0 | 0 | 0 |
| SOX5 | 0.701 | Lumbar_VS_P1_P2_P3_P4_Coeliac_Stellate_Sphenopalatine | 0 | 0 | 0 | 0 | 1 | 0 | 0 | 0 |
| FOXO6 | 0.701 | Coeliac_VS_P1_P2_P3_P4_Lumbar_Stellate_Sphenopalatine | 0 | 0 | 0 | 0 | 0 | 1 | 0 | 0 |
| KIN | 0.701 | Lumbar_Coeliac_VS_P1_P2_P3_P4_Stellate_Sphenopalatine | 0 | 0 | 0 | 0 | 1 | 1 | 0 | 0 |
| DDX55 | 0.701 | Lumbar_Coeliac_VS_P1_P2_P3_P4_Stellate_Sphenopalatine | 0 | 0 | 0 | 0 | 1 | 1 | 0 | 0 |
| RBM41 | 0.701 | Lumbar_Coeliac_VS_P1_P2_P3_P4_Stellate_Sphenopalatine | 0 | 0 | 0 | 0 | 1 | 1 | 0 | 0 |
| SPSB1 | 0.701 | P1_P2_Lumbar_Coeliac_Sphenopalatine_VS_P3_P4_Stellate | 1 | 1 | 0 | 0 | 1 | 1 | 0 | 1 |
| KAT6B | 0.701 | P1_P2_P3_P4_Lumbar_Coeliac_Sphenopalatine_VS_Stellate | 1 | 1 | 1 | 1 | 1 | 1 | 0 | 1 |
| SPARC | 0.7 | P1_P2_P3_P4_Coeliac_Stellate_Sphenopalatine_VS_Lumbar | 1 | 1 | 1 | 1 | 0 | 1 | 1 | 1 |
| RCN1 | 0.699 | P1_P2_P4_VS_P3_Lumbar_Coeliac_Stellate_Sphenopalatine | 1 | 1 | 0 | 1 | 0 | 0 | 0 | 0 |
| EML6 | 0.699 | P1_P2_P3_P4_Lumbar_Coeliac_Sphenopalatine_VS_Stellate | 1 | 1 | 1 | 1 | 1 | 1 | 0 | 1 |
| DMXL1 | 0.699 | P1_P2_P3_P4_Lumbar_Coeliac_Sphenopalatine_VS_Stellate | 1 | 1 | 1 | 1 | 1 | 1 | 0 | 1 |
| AI837181 | 0.699 | P1_P2_P3_P4_Lumbar_Coeliac_Sphenopalatine_VS_Stellate | 1 | 1 | 1 | 1 | 1 | 1 | 0 | 1 |

| | | | | | | | | | | |
|---|---|---|---|---|---|---|---|---|---|---|
| PPFIA1 | 0.698 | P1_P2_P3_P4_Lumbar_Coeliac_Sphenopalatine_VS_Stellate | 1 | 1 | 1 | 1 | 1 | 1 | 0 | 1 |
| LEMD3 | 0.698 | P1_P2_P3_P4_Lumbar_Coeliac_Sphenopalatine_VS_Stellate | 1 | 1 | 1 | 1 | 1 | 1 | 0 | 1 |
| ATP13A3 | 0.698 | P1_P2_P3_P4_Lumbar_Coeliac_Sphenopalatine_VS_Stellate | 1 | 1 | 1 | 1 | 1 | 1 | 0 | 1 |
| 2700049A03 | 0.697 | P4_Lumbar_Coeliac_VS_P1_P2_P3_Stellate_Sphenopalatine | 0 | 0 | 0 | 1 | 1 | 1 | 0 | 0 |
| COL4A2 | 0.697 | Sphenopalatine_VS_P1_P2_P3_P4_Lumbar_Coeliac_Stellate | 0 | 0 | 0 | 0 | 0 | 0 | 0 | 1 |
| DKK3 | 0.697 | P4_Sphenopalatine_VS_P1_P2_P3_Lumbar_Coeliac_Stellate | 0 | 0 | 0 | 1 | 0 | 0 | 0 | 1 |
| LUZP1 | 0.697 | P1_P2_P3_P4_Lumbar_Coeliac_Sphenopalatine_VS_Stellate | 1 | 1 | 1 | 1 | 1 | 1 | 0 | 1 |
| JMY | 0.697 | P1_P2_P3_P4_Lumbar_Coeliac_Sphenopalatine_VS_Stellate | 1 | 1 | 1 | 1 | 1 | 1 | 0 | 1 |
| NYNRIN | 0.697 | P1_P2_P3_P4_Lumbar_Coeliac_Sphenopalatine_VS_Stellate | 1 | 1 | 1 | 1 | 1 | 1 | 0 | 1 |
| CNTN1 | 0.697 | P2_P3_P4_Lumbar_Coeliac_Stellate_Sphenopalatine_VS_P1 | 0 | 1 | 1 | 1 | 1 | 1 | 1 | 1 |
| PROKR1 | 0.696 | P1_VS_P2_P3_P4_Lumbar_Coeliac_Stellate_Sphenopalatine | 1 | 0 | 0 | 0 | 0 | 0 | 0 | 0 |
| B230369F24 | 0.696 | Lumbar_Coeliac_VS_P1_P2_P3_P4_Stellate_Sphenopalatine | 0 | 0 | 0 | 0 | 1 | 1 | 0 | 0 |
| PPM1D | 0.696 | P1_P2_P3_P4_Lumbar_Coeliac_VS_Stellate_Sphenopalatine | 1 | 1 | 1 | 1 | 1 | 1 | 0 | 0 |
| HPCA | 0.696 | P1_P2_P3_P4_Lumbar_Coeliac_Stellate_VS_Sphenopalatine | 1 | 1 | 1 | 1 | 1 | 1 | 1 | 0 |
| LIFR | 0.696 | P1_P2_P3_P4_Lumbar_Coeliac_Stellate_VS_Sphenopalatine | 1 | 1 | 1 | 1 | 1 | 1 | 1 | 0 |
| USP24 | 0.696 | P1_P2_P3_P4_Lumbar_Coeliac_Sphenopalatine_VS_Stellate | 1 | 1 | 1 | 1 | 1 | 1 | 0 | 1 |
| ZFP354C | 0.696 | P1_P2_P3_P4_Lumbar_Coeliac_Sphenopalatine_VS_Stellate | 1 | 1 | 1 | 1 | 1 | 1 | 0 | 1 |
| ATXN3 | 0.696 | P1_P2_P3_P4_Lumbar_Coeliac_Sphenopalatine_VS_Stellate | 1 | 1 | 1 | 1 | 1 | 1 | 0 | 1 |
| ZBTB6 | 0.695 | Coeliac_VS_P1_P2_P3_P4_Lumbar_Stellate_Sphenopalatine | 0 | 0 | 0 | 0 | 0 | 1 | 0 | 0 |
| INSM1 | 0.695 | Coeliac_VS_P1_P2_P3_P4_Lumbar_Stellate_Sphenopalatine | 0 | 0 | 0 | 0 | 0 | 1 | 0 | 0 |
| SLC16A1 | 0.695 | P1_P2_P3_P4_Lumbar_Coeliac_Stellate_VS_Sphenopalatine | 1 | 1 | 1 | 1 | 1 | 1 | 1 | 0 |
| GALNT14 | 0.695 | P1_P2_P4_Lumbar_Coeliac_Sphenopalatine_VS_P3_Stellate | 1 | 1 | 0 | 1 | 1 | 1 | 0 | 1 |
| SLC22A15 | 0.695 | P1_P2_P3_P4_Lumbar_Coeliac_Sphenopalatine_VS_Stellate | 1 | 1 | 1 | 1 | 1 | 1 | 0 | 1 |
| KCNC3 | 0.695 | P1_P2_P3_P4_Lumbar_Coeliac_Sphenopalatine_VS_Stellate | 1 | 1 | 1 | 1 | 1 | 1 | 0 | 1 |
| 6430548M08 | 0.695 | P1_P2_P3_P4_Lumbar_Coeliac_Sphenopalatine_VS_Stellate | 1 | 1 | 1 | 1 | 1 | 1 | 0 | 1 |
| CDK12 | 0.695 | P1_P2_P3_P4_Lumbar_Coeliac_Sphenopalatine_VS_Stellate | 1 | 1 | 1 | 1 | 1 | 1 | 0 | 1 |
| GLCCI1 | 0.694 | P1_P2_P3_P4_Lumbar_Coeliac_Sphenopalatine_VS_Stellate | 1 | 1 | 1 | 1 | 1 | 1 | 0 | 1 |
| CEP295 | 0.694 | P1_P2_P3_P4_Lumbar_Coeliac_Sphenopalatine_VS_Stellate | 1 | 1 | 1 | 1 | 1 | 1 | 0 | 1 |
| CEP126 | 0.693 | Lumbar_VS_P1_P2_P3_P4_Coeliac_Stellate_Sphenopalatine | 0 | 0 | 0 | 0 | 1 | 0 | 0 | 0 |
| VGLL4 | 0.693 | Lumbar_Coeliac_VS_P1_P2_P3_P4_Stellate_Sphenopalatine | 0 | 0 | 0 | 0 | 1 | 1 | 0 | 0 |

| | | | | | | | | | | |
|---|---|---|---|---|---|---|---|---|---|---|
| GM16183 | 0.693 | Lumbar_Coeliac_VS_P1_P2_P3_P4_Stellate_Sphenopalatine | 0 | 0 | 0 | 0 | 1 | 1 | 0 | 0 |
| NIM1K | 0.693 | P1_P2_P3_P4_Lumbar_Coeliac_Stellate_VS_Sphenopalatine | 1 | 1 | 1 | 1 | 1 | 1 | 1 | 0 |
| VIRMA | 0.693 | P1_P2_P3_P4_Lumbar_Coeliac_Sphenopalatine_VS_Stellate | 1 | 1 | 1 | 1 | 1 | 1 | 0 | 1 |
| RBM11 | 0.693 | P1_P2_P3_P4_Lumbar_Coeliac_Sphenopalatine_VS_Stellate | 1 | 1 | 1 | 1 | 1 | 1 | 0 | 1 |
| PTAR1 | 0.693 | P1_P2_P3_P4_Lumbar_Coeliac_Sphenopalatine_VS_Stellate | 1 | 1 | 1 | 1 | 1 | 1 | 0 | 1 |
| SNX19 | 0.692 | P1_P2_P3_P4_Lumbar_Coeliac_Sphenopalatine_VS_Stellate | 1 | 1 | 1 | 1 | 1 | 1 | 0 | 1 |
| MT2 | 0.691 | P3_P4_Sphenopalatine_VS_P1_P2_Lumbar_Coeliac_Stellate | 0 | 0 | 1 | 1 | 0 | 0 | 0 | 1 |
| MINK1 | 0.691 | P1_P2_P3_P4_Lumbar_Coeliac_Sphenopalatine_VS_Stellate | 1 | 1 | 1 | 1 | 1 | 1 | 0 | 1 |
| B3GALT1 | 0.69 | P1_P2_P3_P4_Lumbar_Coeliac_Stellate_VS_Sphenopalatine | 1 | 1 | 1 | 1 | 1 | 1 | 1 | 0 |
| ENDOD1 | 0.69 | P1_P2_P3_P4_Lumbar_Coeliac_Sphenopalatine_VS_Stellate | 1 | 1 | 1 | 1 | 1 | 1 | 0 | 1 |
| HYOU1 | 0.69 | P1_P2_P3_P4_Lumbar_Coeliac_Sphenopalatine_VS_Stellate | 1 | 1 | 1 | 1 | 1 | 1 | 0 | 1 |
| NUCB2 | 0.688 | Lumbar_Coeliac_VS_P1_P2_P3_P4_Stellate_Sphenopalatine | 0 | 0 | 0 | 0 | 1 | 1 | 0 | 0 |
| A130014A01 | 0.688 | Lumbar_Coeliac_VS_P1_P2_P3_P4_Stellate_Sphenopalatine | 0 | 0 | 0 | 0 | 1 | 1 | 0 | 0 |
| PRKD3 | 0.688 | Lumbar_Coeliac_VS_P1_P2_P3_P4_Stellate_Sphenopalatine | 0 | 0 | 0 | 0 | 1 | 1 | 0 | 0 |
| TMEM176A | 0.688 | P1_P2_P3_P4_Lumbar_Coeliac_Sphenopalatine_VS_Stellate | 1 | 1 | 1 | 1 | 1 | 1 | 0 | 1 |
| CDKL5 | 0.687 | P1_P2_P3_P4_Lumbar_Coeliac_VS_Stellate_Sphenopalatine | 1 | 1 | 1 | 1 | 1 | 1 | 0 | 0 |
| KCNQ3 | 0.687 | P1_P2_P3_P4_Lumbar_Coeliac_Stellate_VS_Sphenopalatine | 1 | 1 | 1 | 1 | 1 | 1 | 1 | 0 |
| RPRD1B | 0.687 | P1_P2_P3_P4_Lumbar_Coeliac_Sphenopalatine_VS_Stellate | 1 | 1 | 1 | 1 | 1 | 1 | 0 | 1 |
| ATP11A | 0.687 | P1_P2_P3_P4_Lumbar_Coeliac_Sphenopalatine_VS_Stellate | 1 | 1 | 1 | 1 | 1 | 1 | 0 | 1 |
| LOR | 0.686 | P3_VS_P1_P2_P4_Lumbar_Coeliac_Stellate_Sphenopalatine | 0 | 0 | 1 | 0 | 0 | 0 | 0 | 0 |
| PLSCR4 | 0.686 | P1_P2_P3_P4_Coeliac_Sphenopalatine_VS_Lumbar_Stellate | 1 | 1 | 1 | 1 | 0 | 1 | 0 | 1 |
| MAP3K7 | 0.686 | P1_P2_P3_P4_Lumbar_Coeliac_Sphenopalatine_VS_Stellate | 1 | 1 | 1 | 1 | 1 | 1 | 0 | 1 |
| NR3C1 | 0.685 | P2_P4_VS_P1_P3_Lumbar_Coeliac_Stellate_Sphenopalatine | 0 | 1 | 0 | 1 | 0 | 0 | 0 | 0 |
| NPC1 | 0.685 | P1_P2_P3_P4_Lumbar_Coeliac_Sphenopalatine_VS_Stellate | 1 | 1 | 1 | 1 | 1 | 1 | 0 | 1 |
| TNFRSF12A | 0.684 | P1_P2_P4_VS_P3_Lumbar_Coeliac_Stellate_Sphenopalatine | 1 | 1 | 0 | 1 | 0 | 0 | 0 | 0 |
| MRE11A | 0.684 | Coeliac_VS_P1_P2_P3_P4_Lumbar_Stellate_Sphenopalatine | 0 | 0 | 0 | 0 | 0 | 1 | 0 | 0 |
| MPHOSPH10 | 0.684 | Lumbar_Coeliac_VS_P1_P2_P3_P4_Stellate_Sphenopalatine | 0 | 0 | 0 | 0 | 1 | 1 | 0 | 0 |
| HIGD1C | 0.684 | Lumbar_Coeliac_VS_P1_P2_P3_P4_Stellate_Sphenopalatine | 0 | 0 | 0 | 0 | 1 | 1 | 0 | 0 |
| TBC1D32 | 0.684 | P1_P2_P3_P4_Lumbar_Coeliac_VS_Stellate_Sphenopalatine | 1 | 1 | 1 | 1 | 1 | 1 | 0 | 0 |
| NAF1 | 0.684 | P1_P2_P3_P4_Lumbar_Coeliac_Sphenopalatine_VS_Stellate | 1 | 1 | 1 | 1 | 1 | 1 | 0 | 1 |

| | | | | | | | | | | |
|---|---|---|---|---|---|---|---|---|---|---|
| PRPF39 | 0.684 | P1_P2_P3_P4_Lumbar_Coeliac_Sphenopalatine_VS_Stellate | 1 | 1 | 1 | 1 | 1 | 1 | 0 | 1 |
| ANKRD52 | 0.683 | P4_Lumbar_Coeliac_VS_P1_P2_P3_Stellate_Sphenopalatine | 0 | 0 | 0 | 1 | 1 | 1 | 0 | 0 |
| ZFP955B | 0.683 | P1_P2_P3_P4_Lumbar_Coeliac_Sphenopalatine_VS_Stellate | 1 | 1 | 1 | 1 | 1 | 1 | 0 | 1 |
| CRTAC1 | 0.683 | P1_P2_P3_P4_Lumbar_Coeliac_Sphenopalatine_VS_Stellate | 1 | 1 | 1 | 1 | 1 | 1 | 0 | 1 |
| VPS13D | 0.682 | P1_P2_P3_P4_Lumbar_Coeliac_Sphenopalatine_VS_Stellate | 1 | 1 | 1 | 1 | 1 | 1 | 0 | 1 |
| FURIN | 0.682 | P1_P2_P3_P4_Lumbar_Coeliac_Sphenopalatine_VS_Stellate | 1 | 1 | 1 | 1 | 1 | 1 | 0 | 1 |
| MVP | 0.682 | P1_P2_P3_P4_Lumbar_Coeliac_Sphenopalatine_VS_Stellate | 1 | 1 | 1 | 1 | 1 | 1 | 0 | 1 |
| PCNT | 0.682 | P1_P2_P3_P4_Lumbar_Coeliac_Sphenopalatine_VS_Stellate | 1 | 1 | 1 | 1 | 1 | 1 | 0 | 1 |
| ANKHD1 | 0.682 | P1_P2_P3_P4_Lumbar_Coeliac_Sphenopalatine_VS_Stellate | 1 | 1 | 1 | 1 | 1 | 1 | 0 | 1 |
| SAMD5 | 0.681 | Lumbar_VS_P1_P2_P3_P4_Coeliac_Stellate_Sphenopalatine | 0 | 0 | 0 | 0 | 1 | 0 | 0 | 0 |
| BTG2 | 0.681 | P1_P2_P3_P4_Lumbar_Coeliac_Sphenopalatine_VS_Stellate | 1 | 1 | 1 | 1 | 1 | 1 | 0 | 1 |
| POU2F1 | 0.681 | P1_P2_P3_P4_Lumbar_Coeliac_Sphenopalatine_VS_Stellate | 1 | 1 | 1 | 1 | 1 | 1 | 0 | 1 |
| RANBP10 | 0.681 | P1_P2_P3_P4_Lumbar_Coeliac_Sphenopalatine_VS_Stellate | 1 | 1 | 1 | 1 | 1 | 1 | 0 | 1 |
| TACO1OS | 0.681 | P1_P2_P3_P4_Lumbar_Coeliac_Sphenopalatine_VS_Stellate | 1 | 1 | 1 | 1 | 1 | 1 | 0 | 1 |
| TRMT10C | 0.68 | Lumbar_Coeliac_VS_P1_P2_P3_P4_Stellate_Sphenopalatine | 0 | 0 | 0 | 0 | 1 | 1 | 0 | 0 |
| DEPDC7 | 0.678 | Sphenopalatine_VS_P1_P2_P3_P4_Lumbar_Coeliac_Stellate | 0 | 0 | 0 | 0 | 0 | 0 | 0 | 1 |
| MATK | 0.678 | P4_Sphenopalatine_VS_P1_P2_P3_Lumbar_Coeliac_Stellate | 0 | 0 | 0 | 1 | 0 | 0 | 0 | 1 |
| IMPA2 | 0.677 | P1_VS_P2_P3_P4_Lumbar_Coeliac_Stellate_Sphenopalatine | 1 | 0 | 0 | 0 | 0 | 0 | 0 | 0 |
| CCDC40 | 0.677 | P1_P2_P3_Lumbar_Coeliac_Stellate_Sphenopalatine_VS_P4 | 1 | 1 | 1 | 0 | 1 | 1 | 1 | 1 |
| PARP2 | 0.676 | Lumbar_Coeliac_VS_P1_P2_P3_P4_Stellate_Sphenopalatine | 0 | 0 | 0 | 0 | 1 | 1 | 0 | 0 |
| AFAP1L1 | 0.675 | P1_P3_VS_P2_P4_Lumbar_Coeliac_Stellate_Sphenopalatine | 1 | 0 | 1 | 0 | 0 | 0 | 0 | 0 |
| CDC42EP5 | 0.675 | P1_P2_P4_Lumbar_Coeliac_Stellate_Sphenopalatine_VS_P3 | 1 | 1 | 0 | 1 | 1 | 1 | 1 | 1 |
| GPR89 | 0.674 | Lumbar_Coeliac_VS_P1_P2_P3_P4_Stellate_Sphenopalatine | 0 | 0 | 0 | 0 | 1 | 1 | 0 | 0 |
| BVHT | 0.674 | Sphenopalatine_VS_P1_P2_P3_P4_Lumbar_Coeliac_Stellate | 0 | 0 | 0 | 0 | 0 | 0 | 0 | 1 |
| sept-02 | 0.673 | Coeliac_VS_P1_P2_P3_P4_Lumbar_Stellate_Sphenopalatine | 0 | 0 | 0 | 0 | 0 | 1 | 0 | 0 |
| RAP2B | 0.673 | Lumbar_Coeliac_VS_P1_P2_P3_P4_Stellate_Sphenopalatine | 0 | 0 | 0 | 0 | 1 | 1 | 0 | 0 |
| FSIP1 | 0.672 | Lumbar_Coeliac_VS_P1_P2_P3_P4_Stellate_Sphenopalatine | 0 | 0 | 0 | 0 | 1 | 1 | 0 | 0 |
| SPARCL1 | 0.672 | P1_P2_P3_P4_Coeliac_Sphenopalatine_VS_Lumbar_Stellate | 1 | 1 | 1 | 1 | 0 | 1 | 0 | 1 |
| MBOAT2 | 0.671 | P3_P4_Lumbar_Coeliac_Sphenopalatine_VS_P1_P2_Stellate | 0 | 0 | 1 | 1 | 1 | 1 | 0 | 1 |
| ZFP160 | 0.67 | Lumbar_Coeliac_VS_P1_P2_P3_P4_Stellate_Sphenopalatine | 0 | 0 | 0 | 0 | 1 | 1 | 0 | 0 |

| | | | | | | | | | | |
|---|---|---|---|---|---|---|---|---|---|---|
| GM47664 | 0.669 | P1_P2_VS_P3_P4_Lumbar_Coeliac_Stellate_Sphenopalatine | 1 | 1 | 0 | 0 | 0 | 0 | 0 | 0 |
| MTRF1L | 0.669 | Lumbar_Coeliac_VS_P1_P2_P3_P4_Stellate_Sphenopalatine | 0 | 0 | 0 | 0 | 1 | 1 | 0 | 0 |
| SLC8A3 | 0.669 | P1_P2_P4_Lumbar_Coeliac_VS_P3_Stellate_Sphenopalatine | 1 | 1 | 0 | 1 | 1 | 1 | 0 | 0 |
| ARHGEF40 | 0.669 | P1_P2_P4_Lumbar_Coeliac_Sphenopalatine_VS_P3_Stellate | 1 | 1 | 0 | 1 | 1 | 1 | 0 | 1 |
| MYT1 | 0.668 | P1_P2_P3_P4_Lumbar_Coeliac_VS_Stellate_Sphenopalatine | 1 | 1 | 1 | 1 | 1 | 1 | 0 | 0 |
| SH2B2 | 0.667 | Lumbar_Coeliac_VS_P1_P2_P3_P4_Stellate_Sphenopalatine | 0 | 0 | 0 | 0 | 1 | 1 | 0 | 0 |
| GM14703 | 0.667 | Lumbar_Coeliac_VS_P1_P2_P3_P4_Stellate_Sphenopalatine | 0 | 0 | 0 | 0 | 1 | 1 | 0 | 0 |
| LRCH2 | 0.667 | Lumbar_Coeliac_VS_P1_P2_P3_P4_Stellate_Sphenopalatine | 0 | 0 | 0 | 0 | 1 | 1 | 0 | 0 |
| CNTNAP1 | 0.667 | P1_P2_P4_Sphenopalatine_VS_P3_Lumbar_Coeliac_Stellate | 1 | 1 | 0 | 1 | 0 | 0 | 0 | 1 |
| PPP2CB | 0.666 | Lumbar_Coeliac_VS_P1_P2_P3_P4_Stellate_Sphenopalatine | 0 | 0 | 0 | 0 | 1 | 1 | 0 | 0 |
| HDAC9 | 0.666 | P1_P2_P3_P4_Coeliac_Sphenopalatine_VS_Lumbar_Stellate | 1 | 1 | 1 | 1 | 0 | 1 | 0 | 1 |
| CEP70 | 0.665 | Lumbar_VS_P1_P2_P3_P4_Coeliac_Stellate_Sphenopalatine | 0 | 0 | 0 | 0 | 1 | 0 | 0 | 0 |
| EMILIN1 | 0.665 | P3_Lumbar_Coeliac_VS_P1_P2_P4_Stellate_Sphenopalatine | 0 | 0 | 1 | 0 | 1 | 1 | 0 | 0 |
| ANKRD55 | 0.665 | P3_Lumbar_Coeliac_VS_P1_P2_P4_Stellate_Sphenopalatine | 0 | 0 | 1 | 0 | 1 | 1 | 0 | 0 |
| IFI27 | 0.665 | P1_P2_P4_Sphenopalatine_VS_P3_Lumbar_Coeliac_Stellate | 1 | 1 | 0 | 1 | 0 | 0 | 0 | 1 |
| MSRA | 0.664 | P1_P2_VS_P3_P4_Lumbar_Coeliac_Stellate_Sphenopalatine | 1 | 1 | 0 | 0 | 0 | 0 | 0 | 0 |
| FXYD1 | 0.663 | P4_Sphenopalatine_VS_P1_P2_P3_Lumbar_Coeliac_Stellate | 0 | 0 | 0 | 1 | 0 | 0 | 0 | 1 |
| CADM2 | 0.662 | Sphenopalatine_VS_P1_P2_P3_P4_Lumbar_Coeliac_Stellate | 0 | 0 | 0 | 0 | 0 | 0 | 0 | 1 |
| DOCK5 | 0.661 | Lumbar_Coeliac_Sphenopalatine_VS_P1_P2_P3_P4_Stellate | 0 | 0 | 0 | 0 | 1 | 1 | 0 | 1 |
| LMO3 | 0.659 | Coeliac_VS_P1_P2_P3_P4_Lumbar_Stellate_Sphenopalatine | 0 | 0 | 0 | 0 | 0 | 1 | 0 | 0 |
| CDKN1C | 0.659 | Coeliac_VS_P1_P2_P3_P4_Lumbar_Stellate_Sphenopalatine | 0 | 0 | 0 | 0 | 0 | 1 | 0 | 0 |
| MTERF2 | 0.659 | Lumbar_Coeliac_VS_P1_P2_P3_P4_Stellate_Sphenopalatine | 0 | 0 | 0 | 0 | 1 | 1 | 0 | 0 |
| KCNN1 | 0.658 | P3_VS_P1_P2_P4_Lumbar_Coeliac_Stellate_Sphenopalatine | 0 | 0 | 1 | 0 | 0 | 0 | 0 | 0 |
| GPR150 | 0.658 | Lumbar_VS_P1_P2_P3_P4_Coeliac_Stellate_Sphenopalatine | 0 | 0 | 0 | 0 | 1 | 0 | 0 | 0 |
| SIK2 | 0.658 | Lumbar_Coeliac_VS_P1_P2_P3_P4_Stellate_Sphenopalatine | 0 | 0 | 0 | 0 | 1 | 1 | 0 | 0 |
| ELOVL7 | 0.658 | P1_P2_P4_Lumbar_Coeliac_Sphenopalatine_VS_P3_Stellate | 1 | 1 | 0 | 1 | 1 | 1 | 0 | 1 |
| SORBS2 | 0.658 | P1_P3_P4_Lumbar_Coeliac_Sphenopalatine_VS_P2_Stellate | 1 | 0 | 1 | 1 | 1 | 1 | 0 | 1 |
| RCOR2 | 0.657 | Lumbar_Coeliac_VS_P1_P2_P3_P4_Stellate_Sphenopalatine | 0 | 0 | 0 | 0 | 1 | 1 | 0 | 0 |
| DYNLT1A | 0.656 | Lumbar_Coeliac_VS_P1_P2_P3_P4_Stellate_Sphenopalatine | 0 | 0 | 0 | 0 | 1 | 1 | 0 | 0 |
| ZFP516 | 0.655 | Lumbar_Coeliac_VS_P1_P2_P3_P4_Stellate_Sphenopalatine | 0 | 0 | 0 | 0 | 1 | 1 | 0 | 0 |

| | | | | | | | | | | |
|---|---|---|---|---|---|---|---|---|---|---|
| 5033430I15R | 0.655 | P1_P3_P4_Lumbar_Coeliac_Stellate_Sphenopalatine_VS_P2 | 1 | 0 | 1 | 1 | 1 | 1 | 1 | 1 |
| TSHZ3 | 0.654 | P4_Coeliac_VS_P1_P2_P3_Lumbar_Stellate_Sphenopalatine | 0 | 0 | 0 | 1 | 0 | 1 | 0 | 0 |
| 9430021M05 | 0.653 | P2_Coeliac_VS_P1_P3_P4_Lumbar_Stellate_Sphenopalatine | 0 | 1 | 0 | 0 | 0 | 1 | 0 | 0 |
| UNC13B | 0.653 | Lumbar_Coeliac_VS_P1_P2_P3_P4_Stellate_Sphenopalatine | 0 | 0 | 0 | 0 | 1 | 1 | 0 | 0 |
| TSPAN15 | 0.653 | P3_P4_Lumbar_Coeliac_Stellate_VS_P1_P2_Sphenopalatine | 0 | 0 | 1 | 1 | 1 | 1 | 1 | 0 |
| APOBEC3 | 0.651 | P1_VS_P2_P3_P4_Lumbar_Coeliac_Stellate_Sphenopalatine | 1 | 0 | 0 | 0 | 0 | 0 | 0 | 0 |
| DIS3 | 0.651 | Lumbar_Coeliac_VS_P1_P2_P3_P4_Stellate_Sphenopalatine | 0 | 0 | 0 | 0 | 1 | 1 | 0 | 0 |
| FMNL1 | 0.651 | P1_P2_P3_Lumbar_Coeliac_Stellate_Sphenopalatine_VS_P4 | 1 | 1 | 1 | 0 | 1 | 1 | 1 | 1 |
| ABCA6 | 0.65 | Coeliac_VS_P1_P2_P3_P4_Lumbar_Stellate_Sphenopalatine | 0 | 0 | 0 | 0 | 0 | 1 | 0 | 0 |
| PCSK2 | 0.65 | P3_Lumbar_Coeliac_VS_P1_P2_P4_Stellate_Sphenopalatine | 0 | 0 | 1 | 0 | 1 | 1 | 0 | 0 |
| CNTN3 | 0.649 | P1_VS_P2_P3_P4_Lumbar_Coeliac_Stellate_Sphenopalatine | 1 | 0 | 0 | 0 | 0 | 0 | 0 | 0 |
| SEZ6 | 0.649 | Lumbar_VS_P1_P2_P3_P4_Coeliac_Stellate_Sphenopalatine | 0 | 0 | 0 | 0 | 1 | 0 | 0 | 0 |
| ZFP882 | 0.649 | Coeliac_VS_P1_P2_P3_P4_Lumbar_Stellate_Sphenopalatine | 0 | 0 | 0 | 0 | 0 | 1 | 0 | 0 |
| PI4K2B | 0.649 | P1_P2_P3_P4_Lumbar_Coeliac_Stellate_VS_Sphenopalatine | 1 | 1 | 1 | 1 | 1 | 1 | 1 | 0 |
| HIST1H1C | 0.648 | P2_VS_P1_P3_P4_Lumbar_Coeliac_Stellate_Sphenopalatine | 0 | 1 | 0 | 0 | 0 | 0 | 0 | 0 |
| CDH22 | 0.648 | P3_Coeliac_VS_P1_P2_P4_Lumbar_Stellate_Sphenopalatine | 0 | 0 | 1 | 0 | 0 | 1 | 0 | 0 |
| DHX29 | 0.648 | Lumbar_Coeliac_VS_P1_P2_P3_P4_Stellate_Sphenopalatine | 0 | 0 | 0 | 0 | 1 | 1 | 0 | 0 |
| ZFP248 | 0.647 | Lumbar_Coeliac_VS_P1_P2_P3_P4_Stellate_Sphenopalatine | 0 | 0 | 0 | 0 | 1 | 1 | 0 | 0 |
| TMEM243 | 0.647 | P1_P2_P3_P4_Lumbar_Coeliac_VS_Stellate_Sphenopalatine | 1 | 1 | 1 | 1 | 1 | 1 | 0 | 0 |
| PERP | 0.647 | P3_P4_Lumbar_Coeliac_Stellate_Sphenopalatine_VS_P1_P2 | 0 | 0 | 1 | 1 | 1 | 1 | 1 | 1 |
| WDR3 | 0.646 | Lumbar_Coeliac_VS_P1_P2_P3_P4_Stellate_Sphenopalatine | 0 | 0 | 0 | 0 | 1 | 1 | 0 | 0 |
| B230307C23 | 0.645 | Lumbar_Coeliac_VS_P1_P2_P3_P4_Stellate_Sphenopalatine | 0 | 0 | 0 | 0 | 1 | 1 | 0 | 0 |
| CHD1 | 0.645 | Lumbar_Coeliac_VS_P1_P2_P3_P4_Stellate_Sphenopalatine | 0 | 0 | 0 | 0 | 1 | 1 | 0 | 0 |
| OPHN1 | 0.644 | Lumbar_Coeliac_VS_P1_P2_P3_P4_Stellate_Sphenopalatine | 0 | 0 | 0 | 0 | 1 | 1 | 0 | 0 |
| KHDRBS2 | 0.643 | P1_P4_VS_P2_P3_Lumbar_Coeliac_Stellate_Sphenopalatine | 1 | 0 | 0 | 1 | 0 | 0 | 0 | 0 |
| ZFP994 | 0.643 | Lumbar_Coeliac_VS_P1_P2_P3_P4_Stellate_Sphenopalatine | 0 | 0 | 0 | 0 | 1 | 1 | 0 | 0 |
| RBMX | 0.643 | Lumbar_Coeliac_VS_P1_P2_P3_P4_Stellate_Sphenopalatine | 0 | 0 | 0 | 0 | 1 | 1 | 0 | 0 |
| YPEL2 | 0.643 | P1_P2_P4_Lumbar_Coeliac_Stellate_Sphenopalatine_VS_P3 | 1 | 1 | 0 | 1 | 1 | 1 | 1 | 1 |
| GM28154 | 0.642 | Lumbar_VS_P1_P2_P3_P4_Coeliac_Stellate_Sphenopalatine | 0 | 0 | 0 | 0 | 1 | 0 | 0 | 0 |
| ARHGEF2 | 0.64 | Lumbar_Coeliac_VS_P1_P2_P3_P4_Stellate_Sphenopalatine | 0 | 0 | 0 | 0 | 1 | 1 | 0 | 0 |

| | | | | | | | | | | |
|---|---|---|---|---|---|---|---|---|---|---|
| N4BP2 | 0.639 | Lumbar_Coeliac_VS_P1_P2_P3_P4_Stellate_Sphenopalatine | 0 | 0 | 0 | 0 | 1 | 1 | 0 | 0 |
| RFC3 | 0.639 | Lumbar_Coeliac_VS_P1_P2_P3_P4_Stellate_Sphenopalatine | 0 | 0 | 0 | 0 | 1 | 1 | 0 | 0 |
| NXPE3 | 0.637 | P2_VS_P1_P3_P4_Lumbar_Coeliac_Stellate_Sphenopalatine | 0 | 1 | 0 | 0 | 0 | 0 | 0 | 0 |
| DOK6 | 0.637 | P2_Lumbar_Coeliac_VS_P1_P3_P4_Stellate_Sphenopalatine | 0 | 1 | 0 | 0 | 1 | 1 | 0 | 0 |
| CPQ | 0.637 | P1_P2_P3_P4_Lumbar_Coeliac_VS_Stellate_Sphenopalatine | 1 | 1 | 1 | 1 | 1 | 1 | 0 | 0 |
| NEDD4L | 0.637 | P1_P2_P4_Lumbar_Coeliac_Sphenopalatine_VS_P3_Stellate | 1 | 1 | 0 | 1 | 1 | 1 | 0 | 1 |
| TSPAN8 | 0.637 | P1_P3_P4_Lumbar_Coeliac_Stellate_Sphenopalatine_VS_P2 | 1 | 0 | 1 | 1 | 1 | 1 | 1 | 1 |
| PIK3C2A | 0.636 | P1_P2_P3_P4_Lumbar_Coeliac_VS_Stellate_Sphenopalatine | 1 | 1 | 1 | 1 | 1 | 1 | 0 | 0 |
| SAMD12 | 0.636 | P1_P2_P3_P4_Lumbar_Coeliac_Stellate_VS_Sphenopalatine | 1 | 1 | 1 | 1 | 1 | 1 | 1 | 0 |
| GRK5 | 0.634 | Lumbar_VS_P1_P2_P3_P4_Coeliac_Stellate_Sphenopalatine | 0 | 0 | 0 | 0 | 1 | 0 | 0 | 0 |
| SCLT1 | 0.634 | P4_Lumbar_Coeliac_VS_P1_P2_P3_Stellate_Sphenopalatine | 0 | 0 | 0 | 1 | 1 | 1 | 0 | 0 |
| GEM | 0.633 | P1_P4_VS_P2_P3_Lumbar_Coeliac_Stellate_Sphenopalatine | 1 | 0 | 0 | 1 | 0 | 0 | 0 | 0 |
| 5033428I22R | 0.633 | P1_P3_Lumbar_Coeliac_Stellate_Sphenopalatine_VS_P2_P4 | 1 | 0 | 1 | 0 | 1 | 1 | 1 | 1 |
| DGKG | 0.632 | P2_VS_P1_P3_P4_Lumbar_Coeliac_Stellate_Sphenopalatine | 0 | 1 | 0 | 0 | 0 | 0 | 0 | 0 |
| CDK14 | 0.632 | P1_P2_P3_P4_Lumbar_Coeliac_Stellate_VS_Sphenopalatine | 1 | 1 | 1 | 1 | 1 | 1 | 1 | 0 |
| GM13446 | 0.631 | P1_P2_VS_P3_P4_Lumbar_Coeliac_Stellate_Sphenopalatine | 1 | 1 | 0 | 0 | 0 | 0 | 0 | 0 |
| TCP11L2 | 0.631 | P1_P2_P3_P4_Lumbar_Coeliac_VS_Stellate_Sphenopalatine | 1 | 1 | 1 | 1 | 1 | 1 | 0 | 0 |
| FGF1 | 0.631 | P1_P2_P3_P4_Sphenopalatine_VS_Lumbar_Coeliac_Stellate | 1 | 1 | 1 | 1 | 0 | 0 | 0 | 1 |
| ENPP2 | 0.63 | P3_Lumbar_Coeliac_VS_P1_P2_P4_Stellate_Sphenopalatine | 0 | 0 | 1 | 0 | 1 | 1 | 0 | 0 |
| WRN | 0.63 | P1_P2_P3_P4_Lumbar_Coeliac_VS_Stellate_Sphenopalatine | 1 | 1 | 1 | 1 | 1 | 1 | 0 | 0 |
| PPAT | 0.63 | P1_P2_P3_P4_Lumbar_Coeliac_Stellate_VS_Sphenopalatine | 1 | 1 | 1 | 1 | 1 | 1 | 1 | 0 |
| THY1 | 0.63 | Sphenopalatine_VS_P1_P2_P3_P4_Lumbar_Coeliac_Stellate | 0 | 0 | 0 | 0 | 0 | 0 | 0 | 1 |
| PRTN3 | 0.63 | P1_P2_P3_Lumbar_Coeliac_Stellate_Sphenopalatine_VS_P4 | 1 | 1 | 1 | 0 | 1 | 1 | 1 | 1 |
| TRIOBP | 0.629 | P1_P2_P3_P4_Lumbar_Coeliac_VS_Stellate_Sphenopalatine | 1 | 1 | 1 | 1 | 1 | 1 | 0 | 0 |
| CPT1A | 0.628 | P1_P2_P4_Lumbar_Coeliac_Stellate_Sphenopalatine_VS_P3 | 1 | 1 | 0 | 1 | 1 | 1 | 1 | 1 |
| AGPAT2 | 0.627 | P1_P2_P3_Lumbar_Coeliac_Stellate_Sphenopalatine_VS_P4 | 1 | 1 | 1 | 0 | 1 | 1 | 1 | 1 |
| EHBP1L1 | 0.627 | P1_P3_P4_Lumbar_Coeliac_Stellate_Sphenopalatine_VS_P2 | 1 | 0 | 1 | 1 | 1 | 1 | 1 | 1 |
| GM15222 | 0.626 | P2_P3_Coeliac_VS_P1_P4_Lumbar_Stellate_Sphenopalatine | 0 | 1 | 1 | 0 | 0 | 1 | 0 | 0 |
| GP1BB | 0.626 | P1_P2_P3_P4_Lumbar_Coeliac_VS_Stellate_Sphenopalatine | 1 | 1 | 1 | 1 | 1 | 1 | 0 | 0 |
| NFE2L3 | 0.625 | Coeliac_VS_P1_P2_P3_P4_Lumbar_Stellate_Sphenopalatine | 0 | 0 | 0 | 0 | 0 | 1 | 0 | 0 |

| | | | | | | | | | | |
|---|---|---|---|---|---|---|---|---|---|---|
| GUCY1A2 | 0.625 | P1_P2_P3_P4_Lumbar_Coeliac_VS_Stellate_Sphenopalatine | 1 | 1 | 1 | 1 | 1 | 1 | 0 | 0 |
| sept-09 | 0.625 | P1_P2_P3_Lumbar_Coeliac_Stellate_Sphenopalatine_VS_P4 | 1 | 1 | 1 | 0 | 1 | 1 | 1 | 1 |
| A830018L16I | 0.625 | P1_P3_P4_Lumbar_Coeliac_Stellate_Sphenopalatine_VS_P2 | 1 | 0 | 1 | 1 | 1 | 1 | 1 | 1 |
| PI16 | 0.624 | P3_VS_P1_P2_P4_Lumbar_Coeliac_Stellate_Sphenopalatine | 0 | 0 | 1 | 0 | 0 | 0 | 0 | 0 |
| TXNRD3 | 0.624 | Lumbar_VS_P1_P2_P3_P4_Coeliac_Stellate_Sphenopalatine | 0 | 0 | 0 | 0 | 1 | 0 | 0 | 0 |
| ICAM2 | 0.623 | P3_VS_P1_P2_P4_Lumbar_Coeliac_Stellate_Sphenopalatine | 0 | 0 | 1 | 0 | 0 | 0 | 0 | 0 |
| NEURL1B | 0.623 | P3_Lumbar_Coeliac_VS_P1_P2_P4_Stellate_Sphenopalatine | 0 | 0 | 1 | 0 | 1 | 1 | 0 | 0 |
| RSPH1 | 0.623 | P3_P4_Lumbar_Coeliac_VS_P1_P2_Stellate_Sphenopalatine | 0 | 0 | 1 | 1 | 1 | 1 | 0 | 0 |
| EPB41 | 0.622 | P2_P3_P4_Lumbar_Coeliac_VS_P1_Stellate_Sphenopalatine | 0 | 1 | 1 | 1 | 1 | 1 | 0 | 0 |
| AMOTL1 | 0.621 | Sphenopalatine_VS_P1_P2_P3_P4_Lumbar_Coeliac_Stellate | 0 | 0 | 0 | 0 | 0 | 0 | 0 | 1 |
| HTR3B | 0.62 | P1_P2_P4_VS_P3_Lumbar_Coeliac_Stellate_Sphenopalatine | 1 | 1 | 0 | 1 | 0 | 0 | 0 | 0 |
| FGL1 | 0.619 | P2_VS_P1_P3_P4_Lumbar_Coeliac_Stellate_Sphenopalatine | 0 | 1 | 0 | 0 | 0 | 0 | 0 | 0 |
| CNTNAP5A | 0.619 | P1_P2_P3_Lumbar_Coeliac_Stellate_Sphenopalatine_VS_P4 | 1 | 1 | 1 | 0 | 1 | 1 | 1 | 1 |
| FAM124A | 0.618 | P2_P4_Lumbar_Coeliac_VS_P1_P3_Stellate_Sphenopalatine | 0 | 1 | 0 | 1 | 1 | 1 | 0 | 0 |
| ZFP667 | 0.617 | P1_P2_P3_P4_Lumbar_Coeliac_VS_Stellate_Sphenopalatine | 1 | 1 | 1 | 1 | 1 | 1 | 0 | 0 |
| PPARGC1B | 0.616 | P1_VS_P2_P3_P4_Lumbar_Coeliac_Stellate_Sphenopalatine | 1 | 0 | 0 | 0 | 0 | 0 | 0 | 0 |
| L3HYPDH | 0.612 | Lumbar_VS_P1_P2_P3_P4_Coeliac_Stellate_Sphenopalatine | 0 | 0 | 0 | 0 | 1 | 0 | 0 | 0 |
| INTS8 | 0.612 | P4_Lumbar_Coeliac_VS_P1_P2_P3_Stellate_Sphenopalatine | 0 | 0 | 0 | 1 | 1 | 1 | 0 | 0 |
| PYROXD1 | 0.611 | P1_P3_P4_Lumbar_Coeliac_VS_P2_Stellate_Sphenopalatine | 1 | 0 | 1 | 1 | 1 | 1 | 0 | 0 |
| TCFL5 | 0.61 | Lumbar_VS_P1_P2_P3_P4_Coeliac_Stellate_Sphenopalatine | 0 | 0 | 0 | 0 | 1 | 0 | 0 | 0 |
| SLC9B2 | 0.61 | P1_P2_P3_P4_Lumbar_Coeliac_Stellate_VS_Sphenopalatine | 1 | 1 | 1 | 1 | 1 | 1 | 1 | 0 |
| MRAP2 | 0.609 | P2_P4_VS_P1_P3_Lumbar_Coeliac_Stellate_Sphenopalatine | 0 | 1 | 0 | 1 | 0 | 0 | 0 | 0 |
| IKZF4 | 0.609 | P1_P2_P3_Lumbar_Coeliac_Sphenopalatine_VS_P4_Stellate | 1 | 1 | 1 | 0 | 1 | 1 | 0 | 1 |
| ZCCHC8 | 0.608 | Coeliac_VS_P1_P2_P3_P4_Lumbar_Stellate_Sphenopalatine | 0 | 0 | 0 | 0 | 0 | 1 | 0 | 0 |
| RARG | 0.607 | P2_P3_P4_VS_P1_Lumbar_Coeliac_Stellate_Sphenopalatine | 0 | 1 | 1 | 1 | 0 | 0 | 0 | 0 |
| RPIA | 0.607 | Coeliac_VS_P1_P2_P3_P4_Lumbar_Stellate_Sphenopalatine | 0 | 0 | 0 | 0 | 0 | 1 | 0 | 0 |
| CCDC85C | 0.607 | Coeliac_VS_P1_P2_P3_P4_Lumbar_Stellate_Sphenopalatine | 0 | 0 | 0 | 0 | 0 | 1 | 0 | 0 |
| PTPRN2 | 0.607 | P1_P2_P3_P4_Lumbar_Coeliac_VS_Stellate_Sphenopalatine | 1 | 1 | 1 | 1 | 1 | 1 | 0 | 0 |
| GADD45G | 0.607 | P1_P4_Sphenopalatine_VS_P2_P3_Lumbar_Coeliac_Stellate | 1 | 0 | 0 | 1 | 0 | 0 | 0 | 1 |
| ELK3 | 0.606 | Lumbar_VS_P1_P2_P3_P4_Coeliac_Stellate_Sphenopalatine | 0 | 0 | 0 | 0 | 1 | 0 | 0 | 0 |

| | | | | | | | | | | |
|---|---|---|---|---|---|---|---|---|---|---|
| SCN1A | 0.605 | P1_P2_P3_P4_Lumbar_Coeliac_VS_Stellate_Sphenopalatine | 1 | 1 | 1 | 1 | 1 | 1 | 0 | 0 |
| SHISA3 | 0.604 | P1_VS_P2_P3_P4_Lumbar_Coeliac_Stellate_Sphenopalatine | 1 | 0 | 0 | 0 | 0 | 0 | 0 | 0 |
| SLIT1 | 0.604 | Lumbar_VS_P1_P2_P3_P4_Coeliac_Stellate_Sphenopalatine | 0 | 0 | 0 | 0 | 1 | 0 | 0 | 0 |
| GM43623 | 0.604 | P4_Lumbar_Coeliac_VS_P1_P2_P3_Stellate_Sphenopalatine | 0 | 0 | 0 | 1 | 1 | 1 | 0 | 0 |
| CEP78 | 0.604 | P4_Lumbar_Coeliac_VS_P1_P2_P3_Stellate_Sphenopalatine | 0 | 0 | 0 | 1 | 1 | 1 | 0 | 0 |
| CTHRC1 | 0.604 | P3_P4_Lumbar_Coeliac_Stellate_Sphenopalatine_VS_P1_P2 | 0 | 0 | 1 | 1 | 1 | 1 | 1 | 1 |
| SLC5A3 | 0.603 | P1_P2_P3_P4_Lumbar_Coeliac_VS_Stellate_Sphenopalatine | 1 | 1 | 1 | 1 | 1 | 1 | 0 | 0 |
| SLC35G2 | 0.603 | P1_P2_P4_Lumbar_Coeliac_Sphenopalatine_VS_P3_Stellate | 1 | 1 | 0 | 1 | 1 | 1 | 0 | 1 |
| THG1L | 0.602 | P4_Lumbar_Coeliac_VS_P1_P2_P3_Stellate_Sphenopalatine | 0 | 0 | 0 | 1 | 1 | 1 | 0 | 0 |
| GRIA2 | 0.602 | P1_P3_P4_Lumbar_Coeliac_Stellate_Sphenopalatine_VS_P2 | 1 | 0 | 1 | 1 | 1 | 1 | 1 | 1 |
| BMS1 | 0.601 | P4_Lumbar_Coeliac_VS_P1_P2_P3_Stellate_Sphenopalatine | 0 | 0 | 0 | 1 | 1 | 1 | 0 | 0 |
| KDM3A | 0.601 | P1_P2_P3_P4_Lumbar_Coeliac_VS_Stellate_Sphenopalatine | 1 | 1 | 1 | 1 | 1 | 1 | 0 | 0 |
| AA467197 | 0.601 | P1_P2_P3_Lumbar_Coeliac_Stellate_VS_P4_Sphenopalatine | 1 | 1 | 1 | 0 | 1 | 1 | 1 | 0 |
| SPRY3 | 0.599 | P1_VS_P2_P3_P4_Lumbar_Coeliac_Stellate_Sphenopalatine | 1 | 0 | 0 | 0 | 0 | 0 | 0 | 0 |
| DYRK3 | 0.599 | P3_P4_Lumbar_Coeliac_VS_P1_P2_Stellate_Sphenopalatine | 0 | 0 | 1 | 1 | 1 | 1 | 0 | 0 |
| LRP6 | 0.599 | P1_P2_P3_P4_Lumbar_Coeliac_VS_Stellate_Sphenopalatine | 1 | 1 | 1 | 1 | 1 | 1 | 0 | 0 |
| KCNJ9 | 0.597 | P3_VS_P1_P2_P4_Lumbar_Coeliac_Stellate_Sphenopalatine | 0 | 0 | 1 | 0 | 0 | 0 | 0 | 0 |
| RMC1 | 0.597 | Lumbar_VS_P1_P2_P3_P4_Coeliac_Stellate_Sphenopalatine | 0 | 0 | 0 | 0 | 1 | 0 | 0 | 0 |
| RPH3AL | 0.596 | Coeliac_VS_P1_P2_P3_P4_Lumbar_Stellate_Sphenopalatine | 0 | 0 | 0 | 0 | 0 | 1 | 0 | 0 |
| GRIA1 | 0.596 | P2_P3_P4_Lumbar_Coeliac_VS_P1_Stellate_Sphenopalatine | 0 | 1 | 1 | 1 | 1 | 1 | 0 | 0 |
| SLX4IP | 0.595 | Lumbar_VS_P1_P2_P3_P4_Coeliac_Stellate_Sphenopalatine | 0 | 0 | 0 | 0 | 1 | 0 | 0 | 0 |
| RAD17 | 0.595 | P1_P2_P3_P4_Lumbar_Coeliac_VS_Stellate_Sphenopalatine | 1 | 1 | 1 | 1 | 1 | 1 | 0 | 0 |
| TNFRSF11A | 0.593 | P3_VS_P1_P2_P4_Lumbar_Coeliac_Stellate_Sphenopalatine | 0 | 0 | 1 | 0 | 0 | 0 | 0 | 0 |
| ARID3A | 0.592 | Coeliac_VS_P1_P2_P3_P4_Lumbar_Stellate_Sphenopalatine | 0 | 0 | 0 | 0 | 0 | 1 | 0 | 0 |
| MYO16 | 0.591 | P1_P2_P4_Lumbar_Coeliac_Sphenopalatine_VS_P3_Stellate | 1 | 1 | 0 | 1 | 1 | 1 | 0 | 1 |
| EIF5A2 | 0.59 | P2_P4_VS_P1_P3_Lumbar_Coeliac_Stellate_Sphenopalatine | 0 | 1 | 0 | 1 | 0 | 0 | 0 | 0 |
| BBS9 | 0.59 | P1_P2_P3_P4_Lumbar_Coeliac_VS_Stellate_Sphenopalatine | 1 | 1 | 1 | 1 | 1 | 1 | 0 | 0 |
| DUSP23 | 0.59 | P3_P4_Lumbar_Coeliac_Stellate_Sphenopalatine_VS_P1_P2 | 0 | 0 | 1 | 1 | 1 | 1 | 1 | 1 |
| GSTA3 | 0.589 | P2_P4_VS_P1_P3_Lumbar_Coeliac_Stellate_Sphenopalatine | 0 | 1 | 0 | 1 | 0 | 0 | 0 | 0 |
| MED6 | 0.589 | P4_Lumbar_Coeliac_VS_P1_P2_P3_Stellate_Sphenopalatine | 0 | 0 | 0 | 1 | 1 | 1 | 0 | 0 |

| | | | | | | | | | | |
|---|---|---|---|---|---|---|---|---|---|---|
| LRATD2 | 0.589 | Sphenopalatine_VS_P1_P2_P3_P4_Lumbar_Coeliac_Stellate | 0 | 0 | 0 | 0 | 0 | 0 | 0 | 1 |
| LTK | 0.588 | P1_VS_P2_P3_P4_Lumbar_Coeliac_Stellate_Sphenopalatine | 1 | 0 | 0 | 0 | 0 | 0 | 0 | 0 |
| DRAXIN | 0.587 | Lumbar_VS_P1_P2_P3_P4_Coeliac_Stellate_Sphenopalatine | 0 | 0 | 0 | 0 | 1 | 0 | 0 | 0 |
| APLF | 0.586 | P1_VS_P2_P3_P4_Lumbar_Coeliac_Stellate_Sphenopalatine | 1 | 0 | 0 | 0 | 0 | 0 | 0 | 0 |
| 9630028I04R | 0.586 | P3_VS_P1_P2_P4_Lumbar_Coeliac_Stellate_Sphenopalatine | 0 | 0 | 1 | 0 | 0 | 0 | 0 | 0 |
| D030025P21I | 0.586 | Lumbar_VS_P1_P2_P3_P4_Coeliac_Stellate_Sphenopalatine | 0 | 0 | 0 | 0 | 1 | 0 | 0 | 0 |
| COL3A1 | 0.586 | P1_P2_P3_P4_Coeliac_Sphenopalatine_VS_Lumbar_Stellate | 1 | 1 | 1 | 1 | 0 | 1 | 0 | 1 |
| STAT1 | 0.586 | P1_P2_P4_Lumbar_Coeliac_Sphenopalatine_VS_P3_Stellate | 1 | 1 | 0 | 1 | 1 | 1 | 0 | 1 |
| CST6 | 0.584 | P4_Sphenopalatine_VS_P1_P2_P3_Lumbar_Coeliac_Stellate | 0 | 0 | 0 | 1 | 0 | 0 | 0 | 1 |
| FRMD4B | 0.584 | P1_P2_P3_Lumbar_Coeliac_Sphenopalatine_VS_P4_Stellate | 1 | 1 | 1 | 0 | 1 | 1 | 0 | 1 |
| KDM6A | 0.583 | P1_P2_P3_P4_Lumbar_Coeliac_VS_Stellate_Sphenopalatine | 1 | 1 | 1 | 1 | 1 | 1 | 0 | 0 |
| LAPTM4B | 0.583 | P1_P2_P3_P4_Sphenopalatine_VS_Lumbar_Coeliac_Stellate | 1 | 1 | 1 | 1 | 0 | 0 | 0 | 1 |
| APEX2 | 0.582 | Lumbar_VS_P1_P2_P3_P4_Coeliac_Stellate_Sphenopalatine | 0 | 0 | 0 | 0 | 1 | 0 | 0 | 0 |
| ZBTB26 | 0.582 | Coeliac_VS_P1_P2_P3_P4_Lumbar_Stellate_Sphenopalatine | 0 | 0 | 0 | 0 | 0 | 1 | 0 | 0 |
| 2210408F21I | 0.582 | P1_P2_P3_P4_Sphenopalatine_VS_Lumbar_Coeliac_Stellate | 1 | 1 | 1 | 1 | 0 | 0 | 0 | 1 |
| ANKS1B | 0.582 | P1_P2_P4_Lumbar_Coeliac_Sphenopalatine_VS_P3_Stellate | 1 | 1 | 0 | 1 | 1 | 1 | 0 | 1 |
| ZKSCAN2 | 0.581 | Coeliac_VS_P1_P2_P3_P4_Lumbar_Stellate_Sphenopalatine | 0 | 0 | 0 | 0 | 0 | 1 | 0 | 0 |
| ARL13B | 0.581 | P1_P2_P3_P4_Lumbar_Coeliac_VS_Stellate_Sphenopalatine | 1 | 1 | 1 | 1 | 1 | 1 | 0 | 0 |
| ZFP770 | 0.58 | Lumbar_VS_P1_P2_P3_P4_Coeliac_Stellate_Sphenopalatine | 0 | 0 | 0 | 0 | 1 | 0 | 0 | 0 |
| DLEU7 | 0.58 | P1_P2_P3_Lumbar_Coeliac_Stellate_Sphenopalatine_VS_P4 | 1 | 1 | 1 | 0 | 1 | 1 | 1 | 1 |
| GK5 | 0.579 | Lumbar_VS_P1_P2_P3_P4_Coeliac_Stellate_Sphenopalatine | 0 | 0 | 0 | 0 | 1 | 0 | 0 | 0 |
| NKAIN4 | 0.579 | P4_Lumbar_Coeliac_VS_P1_P2_P3_Stellate_Sphenopalatine | 0 | 0 | 0 | 1 | 1 | 1 | 0 | 0 |
| EYA2 | 0.579 | Sphenopalatine_VS_P1_P2_P3_P4_Lumbar_Coeliac_Stellate | 0 | 0 | 0 | 0 | 0 | 0 | 0 | 1 |
| GMIP | 0.578 | Lumbar_VS_P1_P2_P3_P4_Coeliac_Stellate_Sphenopalatine | 0 | 0 | 0 | 0 | 1 | 0 | 0 | 0 |
| UNC5A | 0.578 | P3_Coeliac_VS_P1_P2_P4_Lumbar_Stellate_Sphenopalatine | 0 | 0 | 1 | 0 | 0 | 1 | 0 | 0 |
| ARMC2 | 0.578 | Lumbar_Coeliac_Stellate_VS_P1_P2_P3_P4_Sphenopalatine | 0 | 0 | 0 | 0 | 1 | 1 | 1 | 0 |
| HIST1H2AP | 0.578 | P1_P2_P3_P4_Coeliac_Stellate_Sphenopalatine_VS_Lumbar | 1 | 1 | 1 | 1 | 0 | 1 | 1 | 1 |
| EMID1 | 0.577 | P1_VS_P2_P3_P4_Lumbar_Coeliac_Stellate_Sphenopalatine | 1 | 0 | 0 | 0 | 0 | 0 | 0 | 0 |
| LGI3 | 0.577 | P3_VS_P1_P2_P4_Lumbar_Coeliac_Stellate_Sphenopalatine | 0 | 0 | 1 | 0 | 0 | 0 | 0 | 0 |
| ARMCX6 | 0.577 | P3_Coeliac_VS_P1_P2_P4_Lumbar_Stellate_Sphenopalatine | 0 | 0 | 1 | 0 | 0 | 1 | 0 | 0 |

| | | | | | | | | | | |
|---|---|---|---|---|---|---|---|---|---|---|
| EIF4E3 | 0.577 | P1_P2_P3_P4_Lumbar_Coeliac_Stellate_VS_Sphenopalatine | 1 | 1 | 1 | 1 | 1 | 1 | 1 | 0 |
| BEND5 | 0.577 | P1_P2_P3_Lumbar_Coeliac_Stellate_Sphenopalatine_VS_P4 | 1 | 1 | 1 | 0 | 1 | 1 | 1 | 1 |
| KCNA6 | 0.576 | P1_P2_P3_Sphenopalatine_VS_P4_Lumbar_Coeliac_Stellate | 1 | 1 | 1 | 0 | 0 | 0 | 0 | 1 |
| PARM1 | 0.574 | P1_P2_P3_P4_Coeliac_Sphenopalatine_VS_Lumbar_Stellate | 1 | 1 | 1 | 1 | 0 | 1 | 0 | 1 |
| ALDH1L2 | 0.572 | Lumbar_VS_P1_P2_P3_P4_Coeliac_Stellate_Sphenopalatine | 0 | 0 | 0 | 0 | 1 | 0 | 0 | 0 |
| 2010300C02I | 0.572 | P4_Lumbar_Coeliac_VS_P1_P2_P3_Stellate_Sphenopalatine | 0 | 0 | 0 | 1 | 1 | 1 | 0 | 0 |
| TRIB1 | 0.571 | P1_VS_P2_P3_P4_Lumbar_Coeliac_Stellate_Sphenopalatine | 1 | 0 | 0 | 0 | 0 | 0 | 0 | 0 |
| C030034I22R | 0.571 | Coeliac_VS_P1_P2_P3_P4_Lumbar_Stellate_Sphenopalatine | 0 | 0 | 0 | 0 | 0 | 1 | 0 | 0 |
| PXMP2 | 0.571 | P1_P2_P3_P4_Sphenopalatine_VS_Lumbar_Coeliac_Stellate | 1 | 1 | 1 | 1 | 0 | 0 | 0 | 1 |
| PPM1E | 0.571 | P1_P2_P4_Lumbar_Coeliac_Sphenopalatine_VS_P3_Stellate | 1 | 1 | 0 | 1 | 1 | 1 | 0 | 1 |
| DNAJC17 | 0.57 | Lumbar_VS_P1_P2_P3_P4_Coeliac_Stellate_Sphenopalatine | 0 | 0 | 0 | 0 | 1 | 0 | 0 | 0 |
| XPC | 0.57 | P1_P2_P3_P4_Lumbar_Coeliac_VS_Stellate_Sphenopalatine | 1 | 1 | 1 | 1 | 1 | 1 | 0 | 0 |
| MPP6 | 0.569 | P1_P2_P3_P4_Lumbar_Coeliac_VS_Stellate_Sphenopalatine | 1 | 1 | 1 | 1 | 1 | 1 | 0 | 0 |
| PLEKHB1 | 0.569 | P2_P4_Sphenopalatine_VS_P1_P3_Lumbar_Coeliac_Stellate | 0 | 1 | 0 | 1 | 0 | 0 | 0 | 1 |
| PWWP2B | 0.568 | P3_VS_P1_P2_P4_Lumbar_Coeliac_Stellate_Sphenopalatine | 0 | 0 | 1 | 0 | 0 | 0 | 0 | 0 |
| PRIMA1 | 0.568 | P1_P2_P3_P4_Lumbar_Sphenopalatine_VS_Coeliac_Stellate | 1 | 1 | 1 | 1 | 1 | 0 | 0 | 1 |
| SPRY1 | 0.567 | P1_VS_P2_P3_P4_Lumbar_Coeliac_Stellate_Sphenopalatine | 1 | 0 | 0 | 0 | 0 | 0 | 0 | 0 |
| 2310030G06 | 0.566 | P3_Lumbar_Coeliac_VS_P1_P2_P4_Stellate_Sphenopalatine | 0 | 0 | 1 | 0 | 1 | 1 | 0 | 0 |
| FDXR | 0.566 | P1_P2_P4_Lumbar_Coeliac_VS_P3_Stellate_Sphenopalatine | 1 | 1 | 0 | 1 | 1 | 1 | 0 | 0 |
| SCML4 | 0.566 | P1_P2_P3_P4_Lumbar_Coeliac_VS_Stellate_Sphenopalatine | 1 | 1 | 1 | 1 | 1 | 1 | 0 | 0 |
| MIPOL1 | 0.565 | P1_P2_P3_P4_Lumbar_Coeliac_VS_Stellate_Sphenopalatine | 1 | 1 | 1 | 1 | 1 | 1 | 0 | 0 |
| PEPD | 0.564 | P1_P2_P3_P4_Lumbar_Coeliac_VS_Stellate_Sphenopalatine | 1 | 1 | 1 | 1 | 1 | 1 | 0 | 0 |
| SNTB2 | 0.563 | P1_P2_P3_P4_Lumbar_Coeliac_VS_Stellate_Sphenopalatine | 1 | 1 | 1 | 1 | 1 | 1 | 0 | 0 |
| CALR3 | 0.562 | P3_VS_P1_P2_P4_Lumbar_Coeliac_Stellate_Sphenopalatine | 0 | 0 | 1 | 0 | 0 | 0 | 0 | 0 |
| ST6GAL1 | 0.562 | P1_P2_Sphenopalatine_VS_P3_P4_Lumbar_Coeliac_Stellate | 1 | 1 | 0 | 0 | 0 | 0 | 0 | 1 |
| NR4A1 | 0.562 | P1_P2_P3_P4_Coeliac_Stellate_Sphenopalatine_VS_Lumbar | 1 | 1 | 1 | 1 | 0 | 1 | 1 | 1 |
| CENPA | 0.561 | P2_VS_P1_P3_P4_Lumbar_Coeliac_Stellate_Sphenopalatine | 0 | 1 | 0 | 0 | 0 | 0 | 0 | 0 |
| APCDD1 | 0.561 | P2_VS_P1_P3_P4_Lumbar_Coeliac_Stellate_Sphenopalatine | 0 | 1 | 0 | 0 | 0 | 0 | 0 | 0 |
| NFE2L2 | 0.561 | P1_P2_P4_Lumbar_Coeliac_VS_P3_Stellate_Sphenopalatine | 1 | 1 | 0 | 1 | 1 | 1 | 0 | 0 |
| NTM | 0.561 | Sphenopalatine_VS_P1_P2_P3_P4_Lumbar_Coeliac_Stellate | 0 | 0 | 0 | 0 | 0 | 0 | 0 | 1 |

| | | | | | | | | | | |
|---|---|---|---|---|---|---|---|---|---|---|
| TCEAL6 | 0.56 | P3_VS_P1_P2_P4_Lumbar_Coeliac_Stellate_Sphenopalatine | 0 | 0 | 1 | 0 | 0 | 0 | 0 | 0 |
| ZSCAN2 | 0.56 | Coeliac_VS_P1_P2_P3_P4_Lumbar_Stellate_Sphenopalatine | 0 | 0 | 0 | 0 | 0 | 1 | 0 | 0 |
| GM32352 | 0.56 | Sphenopalatine_VS_P1_P2_P3_P4_Lumbar_Coeliac_Stellate | 0 | 0 | 0 | 0 | 0 | 0 | 0 | 1 |
| GM37041 | 0.559 | P1_VS_P2_P3_P4_Lumbar_Coeliac_Stellate_Sphenopalatine | 1 | 0 | 0 | 0 | 0 | 0 | 0 | 0 |
| PPP2R3A | 0.559 | P1_P2_P4_Lumbar_Coeliac_VS_P3_Stellate_Sphenopalatine | 1 | 1 | 0 | 1 | 1 | 1 | 0 | 0 |
| SIAH2 | 0.559 | P1_P2_P3_P4_Lumbar_Coeliac_VS_Stellate_Sphenopalatine | 1 | 1 | 1 | 1 | 1 | 1 | 0 | 0 |
| 4930452B06I | 0.558 | P1_P4_VS_P2_P3_Lumbar_Coeliac_Stellate_Sphenopalatine | 1 | 0 | 0 | 1 | 0 | 0 | 0 | 0 |
| RLF | 0.558 | P1_P2_P3_P4_Lumbar_Coeliac_VS_Stellate_Sphenopalatine | 1 | 1 | 1 | 1 | 1 | 1 | 0 | 0 |
| GM8773 | 0.558 | P1_P2_P3_P4_Lumbar_Coeliac_Stellate_VS_Sphenopalatine | 1 | 1 | 1 | 1 | 1 | 1 | 1 | 0 |
| SHLD2 | 0.557 | Coeliac_VS_P1_P2_P3_P4_Lumbar_Stellate_Sphenopalatine | 0 | 0 | 0 | 0 | 0 | 1 | 0 | 0 |
| CPNE2 | 0.557 | P1_P2_P3_Lumbar_Coeliac_Stellate_VS_P4_Sphenopalatine | 1 | 1 | 1 | 0 | 1 | 1 | 1 | 0 |
| MYRIP | 0.556 | P1_P2_P4_VS_P3_Lumbar_Coeliac_Stellate_Sphenopalatine | 1 | 1 | 0 | 1 | 0 | 0 | 0 | 0 |
| TRMO | 0.556 | Coeliac_VS_P1_P2_P3_P4_Lumbar_Stellate_Sphenopalatine | 0 | 0 | 0 | 0 | 0 | 1 | 0 | 0 |
| MT1 | 0.556 | P3_P4_Sphenopalatine_VS_P1_P2_Lumbar_Coeliac_Stellate | 0 | 0 | 1 | 1 | 0 | 0 | 0 | 1 |
| ARHGAP42 | 0.555 | P1_P2_VS_P3_P4_Lumbar_Coeliac_Stellate_Sphenopalatine | 1 | 1 | 0 | 0 | 0 | 0 | 0 | 0 |
| GM15513 | 0.554 | Coeliac_VS_P1_P2_P3_P4_Lumbar_Stellate_Sphenopalatine | 0 | 0 | 0 | 0 | 0 | 1 | 0 | 0 |
| PDZRN3 | 0.553 | P4_Coeliac_VS_P1_P2_P3_Lumbar_Stellate_Sphenopalatine | 0 | 0 | 0 | 1 | 0 | 1 | 0 | 0 |
| GXYLT1 | 0.553 | P4_Lumbar_Coeliac_VS_P1_P2_P3_Stellate_Sphenopalatine | 0 | 0 | 0 | 1 | 1 | 1 | 0 | 0 |
| MTUS2 | 0.553 | P1_P2_P3_Lumbar_Coeliac_Stellate_Sphenopalatine_VS_P4 | 1 | 1 | 1 | 0 | 1 | 1 | 1 | 1 |
| SMOC1 | 0.552 | P2_Lumbar_VS_P1_P3_P4_Coeliac_Stellate_Sphenopalatine | 0 | 1 | 0 | 0 | 1 | 0 | 0 | 0 |
| FIGNL2 | 0.55 | P2_VS_P1_P3_P4_Lumbar_Coeliac_Stellate_Sphenopalatine | 0 | 1 | 0 | 0 | 0 | 0 | 0 | 0 |
| ZFP933 | 0.55 | Coeliac_VS_P1_P2_P3_P4_Lumbar_Stellate_Sphenopalatine | 0 | 0 | 0 | 0 | 0 | 1 | 0 | 0 |
| ELF1 | 0.55 | Coeliac_VS_P1_P2_P3_P4_Lumbar_Stellate_Sphenopalatine | 0 | 0 | 0 | 0 | 0 | 1 | 0 | 0 |
| COX15 | 0.55 | Coeliac_VS_P1_P2_P3_P4_Lumbar_Stellate_Sphenopalatine | 0 | 0 | 0 | 0 | 0 | 1 | 0 | 0 |
| STAM2 | 0.55 | P1_P2_P3_P4_Lumbar_Coeliac_VS_Stellate_Sphenopalatine | 1 | 1 | 1 | 1 | 1 | 1 | 0 | 0 |
| GM5532 | 0.55 | P4_Lumbar_Coeliac_Stellate_VS_P1_P2_P3_Sphenopalatine | 0 | 0 | 0 | 1 | 1 | 1 | 1 | 0 |
| KLHL32 | 0.549 | P3_VS_P1_P2_P4_Lumbar_Coeliac_Stellate_Sphenopalatine | 0 | 0 | 1 | 0 | 0 | 0 | 0 | 0 |
| GOLIM4 | 0.549 | P1_P2_P3_P4_Lumbar_Coeliac_VS_Stellate_Sphenopalatine | 1 | 1 | 1 | 1 | 1 | 1 | 0 | 0 |
| GM3636 | 0.549 | P1_P2_P3_Lumbar_Coeliac_Stellate_Sphenopalatine_VS_P4 | 1 | 1 | 1 | 0 | 1 | 1 | 1 | 1 |
| CCDC51 | 0.548 | P4_Lumbar_Coeliac_VS_P1_P2_P3_Stellate_Sphenopalatine | 0 | 0 | 0 | 1 | 1 | 1 | 0 | 0 |

| | | | | | | | | | | |
|---|---|---|---|---|---|---|---|---|---|---|
| TMTC1 | 0.548 | P1_P2_P4_Lumbar_Sphenopalatine_VS_P3_Coeliac_Stellate | 1 | 1 | 0 | 1 | 1 | 0 | 0 | 1 |
| LINGO2 | 0.548 | P2_P3_P4_Lumbar_Coeliac_Stellate_Sphenopalatine_VS_P1 | 0 | 1 | 1 | 1 | 1 | 1 | 1 | 1 |
| CCDC122 | 0.547 | P2_P3_P4_VS_P1_Lumbar_Coeliac_Stellate_Sphenopalatine | 0 | 1 | 1 | 1 | 0 | 0 | 0 | 0 |
| SLC8A2 | 0.547 | Sphenopalatine_VS_P1_P2_P3_P4_Lumbar_Coeliac_Stellate | 0 | 0 | 0 | 0 | 0 | 0 | 0 | 1 |
| ERCC4 | 0.546 | P4_Lumbar_Coeliac_VS_P1_P2_P3_Stellate_Sphenopalatine | 0 | 0 | 0 | 1 | 1 | 1 | 0 | 0 |
| MLLT1 | 0.545 | P3_Lumbar_Coeliac_VS_P1_P2_P4_Stellate_Sphenopalatine | 0 | 0 | 1 | 0 | 1 | 1 | 0 | 0 |
| MAP3K4 | 0.545 | P4_Lumbar_Coeliac_VS_P1_P2_P3_Stellate_Sphenopalatine | 0 | 0 | 0 | 1 | 1 | 1 | 0 | 0 |
| KCNA2 | 0.545 | P4_Sphenopalatine_VS_P1_P2_P3_Lumbar_Coeliac_Stellate | 0 | 0 | 0 | 1 | 0 | 0 | 0 | 1 |
| EMP3 | 0.545 | P1_P3_P4_Sphenopalatine_VS_P2_Lumbar_Coeliac_Stellate | 1 | 0 | 1 | 1 | 0 | 0 | 0 | 1 |
| DCAF1 | 0.544 | Coeliac_VS_P1_P2_P3_P4_Lumbar_Stellate_Sphenopalatine | 0 | 0 | 0 | 0 | 0 | 1 | 0 | 0 |
| COL23A1 | 0.544 | P2_Coeliac_VS_P1_P3_P4_Lumbar_Stellate_Sphenopalatine | 0 | 1 | 0 | 0 | 0 | 1 | 0 | 0 |
| GM26549 | 0.543 | Coeliac_VS_P1_P2_P3_P4_Lumbar_Stellate_Sphenopalatine | 0 | 0 | 0 | 0 | 0 | 1 | 0 | 0 |
| FAM13C | 0.543 | P1_P4_Lumbar_Coeliac_Sphenopalatine_VS_P2_P3_Stellate | 1 | 0 | 0 | 1 | 1 | 1 | 0 | 1 |
| CALD1 | 0.543 | P1_P2_P4_Lumbar_Coeliac_Sphenopalatine_VS_P3_Stellate | 1 | 1 | 0 | 1 | 1 | 1 | 0 | 1 |
| LRP1 | 0.543 | P1_P2_P4_Lumbar_Coeliac_Sphenopalatine_VS_P3_Stellate | 1 | 1 | 0 | 1 | 1 | 1 | 0 | 1 |
| TOX | 0.542 | P1_Lumbar_Coeliac_Stellate_Sphenopalatine_VS_P2_P3_P4 | 1 | 0 | 0 | 0 | 1 | 1 | 1 | 1 |
| MLXIP | 0.541 | P4_Lumbar_VS_P1_P2_P3_Coeliac_Stellate_Sphenopalatine | 0 | 0 | 0 | 1 | 1 | 0 | 0 | 0 |
| MICAL2 | 0.541 | P1_P3_Lumbar_Coeliac_Sphenopalatine_VS_P2_P4_Stellate | 1 | 0 | 1 | 0 | 1 | 1 | 0 | 1 |
| SPIN2C | 0.54 | P1_P3_P4_Lumbar_Coeliac_VS_P2_Stellate_Sphenopalatine | 1 | 0 | 1 | 1 | 1 | 1 | 0 | 0 |
| DPP8 | 0.54 | P1_P2_P3_P4_Lumbar_Coeliac_VS_Stellate_Sphenopalatine | 1 | 1 | 1 | 1 | 1 | 1 | 0 | 0 |
| UBE2L6 | 0.54 | P1_P2_P3_P4_Sphenopalatine_VS_Lumbar_Coeliac_Stellate | 1 | 1 | 1 | 1 | 0 | 0 | 0 | 1 |
| TRAF3IP1 | 0.539 | Coeliac_VS_P1_P2_P3_P4_Lumbar_Stellate_Sphenopalatine | 0 | 0 | 0 | 0 | 0 | 1 | 0 | 0 |
| BMP3 | 0.539 | P1_P2_P3_P4_Lumbar_Coeliac_VS_Stellate_Sphenopalatine | 1 | 1 | 1 | 1 | 1 | 1 | 0 | 0 |
| ERICH3 | 0.538 | P4_Lumbar_VS_P1_P2_P3_Coeliac_Stellate_Sphenopalatine | 0 | 0 | 0 | 1 | 1 | 0 | 0 | 0 |
| INTS6L | 0.537 | P4_Lumbar_Coeliac_VS_P1_P2_P3_Stellate_Sphenopalatine | 0 | 0 | 0 | 1 | 1 | 1 | 0 | 0 |
| 1700023F06I | 0.537 | P1_P2_P3_P4_Sphenopalatine_VS_Lumbar_Coeliac_Stellate | 1 | 1 | 1 | 1 | 0 | 0 | 0 | 1 |
| TMOD1 | 0.537 | P1_P2_P3_Lumbar_Coeliac_Stellate_Sphenopalatine_VS_P4 | 1 | 1 | 1 | 0 | 1 | 1 | 1 | 1 |
| AMPD3 | 0.537 | P3_P4_Lumbar_Coeliac_Stellate_Sphenopalatine_VS_P1_P2 | 0 | 0 | 1 | 1 | 1 | 1 | 1 | 1 |
| PDE4A | 0.536 | P2_VS_P1_P3_P4_Lumbar_Coeliac_Stellate_Sphenopalatine | 0 | 1 | 0 | 0 | 0 | 0 | 0 | 0 |
| PABPC1L | 0.536 | Coeliac_VS_P1_P2_P3_P4_Lumbar_Stellate_Sphenopalatine | 0 | 0 | 0 | 0 | 0 | 1 | 0 | 0 |

| | | | | | | | | | | |
|---|---|---|---|---|---|---|---|---|---|---|
| TIPARP | 0.536 | Coeliac_VS_P1_P2_P3_P4_Lumbar_Stellate_Sphenopalatine | 0 | 0 | 0 | 0 | 0 | 1 | 0 | 0 |
| SHTN1 | 0.536 | P1_P2_P3_Lumbar_Coeliac_Sphenopalatine_VS_P4_Stellate | 1 | 1 | 1 | 0 | 1 | 1 | 0 | 1 |
| TRAF3IP2 | 0.535 | P4_Lumbar_Coeliac_VS_P1_P2_P3_Stellate_Sphenopalatine | 0 | 0 | 0 | 1 | 1 | 1 | 0 | 0 |
| YAP1 | 0.534 | Coeliac_VS_P1_P2_P3_P4_Lumbar_Stellate_Sphenopalatine | 0 | 0 | 0 | 0 | 0 | 1 | 0 | 0 |
| AMT | 0.534 | Coeliac_VS_P1_P2_P3_P4_Lumbar_Stellate_Sphenopalatine | 0 | 0 | 0 | 0 | 0 | 1 | 0 | 0 |
| GLMN | 0.533 | P4_Lumbar_Coeliac_VS_P1_P2_P3_Stellate_Sphenopalatine | 0 | 0 | 0 | 1 | 1 | 1 | 0 | 0 |
| BIRC2 | 0.532 | P4_Lumbar_Coeliac_VS_P1_P2_P3_Stellate_Sphenopalatine | 0 | 0 | 0 | 1 | 1 | 1 | 0 | 0 |
| XRCC4 | 0.532 | P4_Lumbar_Coeliac_VS_P1_P2_P3_Stellate_Sphenopalatine | 0 | 0 | 0 | 1 | 1 | 1 | 0 | 0 |
| ARMH4 | 0.532 | P2_P4_Lumbar_Coeliac_VS_P1_P3_Stellate_Sphenopalatine | 0 | 1 | 0 | 1 | 1 | 1 | 0 | 0 |
| RAB8B | 0.532 | P1_P2_P3_P4_Lumbar_Coeliac_VS_Stellate_Sphenopalatine | 1 | 1 | 1 | 1 | 1 | 1 | 0 | 0 |
| TMEM108 | 0.532 | P1_P3_Lumbar_Coeliac_Stellate_Sphenopalatine_VS_P2_P4 | 1 | 0 | 1 | 0 | 1 | 1 | 1 | 1 |
| MAGEL2 | 0.531 | P1_VS_P2_P3_P4_Lumbar_Coeliac_Stellate_Sphenopalatine | 1 | 0 | 0 | 0 | 0 | 0 | 0 | 0 |
| WSCD1 | 0.531 | P1_VS_P2_P3_P4_Lumbar_Coeliac_Stellate_Sphenopalatine | 1 | 0 | 0 | 0 | 0 | 0 | 0 | 0 |
| TRP53BP2 | 0.531 | Coeliac_VS_P1_P2_P3_P4_Lumbar_Stellate_Sphenopalatine | 0 | 0 | 0 | 0 | 0 | 1 | 0 | 0 |
| TAPBP | 0.531 | P1_P2_P3_P4_Sphenopalatine_VS_Lumbar_Coeliac_Stellate | 1 | 1 | 1 | 1 | 0 | 0 | 0 | 1 |
| BICRA | 0.53 | Coeliac_VS_P1_P2_P3_P4_Lumbar_Stellate_Sphenopalatine | 0 | 0 | 0 | 0 | 0 | 1 | 0 | 0 |
| RFX3 | 0.53 | Coeliac_VS_P1_P2_P3_P4_Lumbar_Stellate_Sphenopalatine | 0 | 0 | 0 | 0 | 0 | 1 | 0 | 0 |
| PAK6 | 0.53 | P3_Lumbar_Coeliac_VS_P1_P2_P4_Stellate_Sphenopalatine | 0 | 0 | 1 | 0 | 1 | 1 | 0 | 0 |
| 2610507B11I | 0.53 | P1_P2_P4_Lumbar_Coeliac_Sphenopalatine_VS_P3_Stellate | 1 | 1 | 0 | 1 | 1 | 1 | 0 | 1 |
| IGSF21 | 0.529 | P1_P2_P3_Lumbar_Coeliac_Stellate_Sphenopalatine_VS_P4 | 1 | 1 | 1 | 0 | 1 | 1 | 1 | 1 |
| ALKAL2 | 0.528 | P3_VS_P1_P2_P4_Lumbar_Coeliac_Stellate_Sphenopalatine | 0 | 0 | 1 | 0 | 0 | 0 | 0 | 0 |
| FAM181B | 0.528 | P3_P4_Lumbar_Coeliac_VS_P1_P2_Stellate_Sphenopalatine | 0 | 0 | 1 | 1 | 1 | 1 | 0 | 0 |
| DNAJC21 | 0.528 | P1_P2_P3_P4_Lumbar_Coeliac_VS_Stellate_Sphenopalatine | 1 | 1 | 1 | 1 | 1 | 1 | 0 | 0 |
| AVIL | 0.528 | Sphenopalatine_VS_P1_P2_P3_P4_Lumbar_Coeliac_Stellate | 0 | 0 | 0 | 0 | 0 | 0 | 0 | 1 |
| TSPAN12 | 0.527 | P2_P3_VS_P1_P4_Lumbar_Coeliac_Stellate_Sphenopalatine | 0 | 1 | 1 | 0 | 0 | 0 | 0 | 0 |
| ETAA1OS | 0.527 | P2_P3_P4_VS_P1_Lumbar_Coeliac_Stellate_Sphenopalatine | 0 | 1 | 1 | 1 | 0 | 0 | 0 | 0 |
| CENPH | 0.526 | Coeliac_VS_P1_P2_P3_P4_Lumbar_Stellate_Sphenopalatine | 0 | 0 | 0 | 0 | 0 | 1 | 0 | 0 |
| UFL1 | 0.526 | P4_Lumbar_Coeliac_VS_P1_P2_P3_Stellate_Sphenopalatine | 0 | 0 | 0 | 1 | 1 | 1 | 0 | 0 |
| CCDC82 | 0.526 | P4_Lumbar_Coeliac_VS_P1_P2_P3_Stellate_Sphenopalatine | 0 | 0 | 0 | 1 | 1 | 1 | 0 | 0 |
| QK | 0.526 | P4_Lumbar_Coeliac_VS_P1_P2_P3_Stellate_Sphenopalatine | 0 | 0 | 0 | 1 | 1 | 1 | 0 | 0 |

| | | | | | | | | | | |
|---|---|---|---|---|---|---|---|---|---|---|
| PUS7 | 0.526 | P1_P2_P3_P4_Lumbar_Coeliac_VS_Stellate_Sphenopalatine | 1 | 1 | 1 | 1 | 1 | 1 | 0 | 0 |
| KBTBD11 | 0.526 | P1_P2_P3_P4_Lumbar_Coeliac_VS_Stellate_Sphenopalatine | 1 | 1 | 1 | 1 | 1 | 1 | 0 | 0 |
| PAQR8 | 0.525 | P1_P2_P3_P4_Lumbar_VS_Coeliac_Stellate_Sphenopalatine | 1 | 1 | 1 | 1 | 1 | 0 | 0 | 0 |
| MIR670HG | 0.525 | P3_Sphenopalatine_VS_P1_P2_P4_Lumbar_Coeliac_Stellate | 0 | 0 | 1 | 0 | 0 | 0 | 0 | 1 |
| ARHGAP10 | 0.525 | P1_P3_P4_Lumbar_Coeliac_Sphenopalatine_VS_P2_Stellate | 1 | 0 | 1 | 1 | 1 | 1 | 0 | 1 |
| ZFP606 | 0.524 | Coeliac_VS_P1_P2_P3_P4_Lumbar_Stellate_Sphenopalatine | 0 | 0 | 0 | 0 | 0 | 1 | 0 | 0 |
| MSH6 | 0.524 | Coeliac_VS_P1_P2_P3_P4_Lumbar_Stellate_Sphenopalatine | 0 | 0 | 0 | 0 | 0 | 1 | 0 | 0 |
| SERPING1 | 0.524 | Sphenopalatine_VS_P1_P2_P3_P4_Lumbar_Coeliac_Stellate | 0 | 0 | 0 | 0 | 0 | 0 | 0 | 1 |
| HSPA2 | 0.524 | P1_P2_P4_Lumbar_Coeliac_Sphenopalatine_VS_P3_Stellate | 1 | 1 | 0 | 1 | 1 | 1 | 0 | 1 |
| DOC2A | 0.523 | P3_VS_P1_P2_P4_Lumbar_Coeliac_Stellate_Sphenopalatine | 0 | 0 | 1 | 0 | 0 | 0 | 0 | 0 |
| SLC1A6 | 0.523 | P3_Lumbar_Coeliac_VS_P1_P2_P4_Stellate_Sphenopalatine | 0 | 0 | 1 | 0 | 1 | 1 | 0 | 0 |
| ZMYND19 | 0.523 | P1_P2_P3_P4_Lumbar_Coeliac_VS_Stellate_Sphenopalatine | 1 | 1 | 1 | 1 | 1 | 1 | 0 | 0 |
| RAD51D | 0.523 | P1_P2_P3_P4_Lumbar_Coeliac_VS_Stellate_Sphenopalatine | 1 | 1 | 1 | 1 | 1 | 1 | 0 | 0 |
| TBC1D4 | 0.521 | P1_Lumbar_Coeliac_VS_P2_P3_P4_Stellate_Sphenopalatine | 1 | 0 | 0 | 0 | 1 | 1 | 0 | 0 |
| FMN1 | 0.519 | P1_P2_P3_P4_Lumbar_Coeliac_VS_Stellate_Sphenopalatine | 1 | 1 | 1 | 1 | 1 | 1 | 0 | 0 |
| DUSP7 | 0.518 | P1_P2_P4_Lumbar_Coeliac_VS_P3_Stellate_Sphenopalatine | 1 | 1 | 0 | 1 | 1 | 1 | 0 | 0 |
| DUSP16 | 0.518 | P1_P2_P3_P4_Lumbar_Coeliac_VS_Stellate_Sphenopalatine | 1 | 1 | 1 | 1 | 1 | 1 | 0 | 0 |
| LIG4 | 0.517 | Coeliac_VS_P1_P2_P3_P4_Lumbar_Stellate_Sphenopalatine | 0 | 0 | 0 | 0 | 0 | 1 | 0 | 0 |
| CRABP1 | 0.517 | P2_P3_P4_Coeliac_Stellate_VS_P1_Lumbar_Sphenopalatine | 0 | 1 | 1 | 1 | 0 | 1 | 1 | 0 |
| PDLIM5 | 0.516 | P1_VS_P2_P3_P4_Lumbar_Coeliac_Stellate_Sphenopalatine | 1 | 0 | 0 | 0 | 0 | 0 | 0 | 0 |
| C630043F03I | 0.516 | P3_VS_P1_P2_P4_Lumbar_Coeliac_Stellate_Sphenopalatine | 0 | 0 | 1 | 0 | 0 | 0 | 0 | 0 |
| SLC26A4 | 0.516 | P3_VS_P1_P2_P4_Lumbar_Coeliac_Stellate_Sphenopalatine | 0 | 0 | 1 | 0 | 0 | 0 | 0 | 0 |
| PIR | 0.516 | P1_P2_P3_P4_Lumbar_Coeliac_Stellate_VS_Sphenopalatine | 1 | 1 | 1 | 1 | 1 | 1 | 1 | 0 |
| POLD3 | 0.515 | P1_P2_P3_P4_Lumbar_Coeliac_VS_Stellate_Sphenopalatine | 1 | 1 | 1 | 1 | 1 | 1 | 0 | 0 |
| PANK1 | 0.515 | P1_P2_P3_P4_Lumbar_Coeliac_VS_Stellate_Sphenopalatine | 1 | 1 | 1 | 1 | 1 | 1 | 0 | 0 |
| TRAF3IP3 | 0.515 | Sphenopalatine_VS_P1_P2_P3_P4_Lumbar_Coeliac_Stellate | 0 | 0 | 0 | 0 | 0 | 0 | 0 | 1 |
| RNF122 | 0.515 | P1_Lumbar_Coeliac_Sphenopalatine_VS_P2_P3_P4_Stellate | 1 | 0 | 0 | 0 | 1 | 1 | 0 | 1 |
| SGMS2 | 0.514 | P3_VS_P1_P2_P4_Lumbar_Coeliac_Stellate_Sphenopalatine | 0 | 0 | 1 | 0 | 0 | 0 | 0 | 0 |
| PFKFB3 | 0.514 | P1_Lumbar_VS_P2_P3_P4_Coeliac_Stellate_Sphenopalatine | 1 | 0 | 0 | 0 | 1 | 0 | 0 | 0 |
| TMEM39B | 0.514 | P4_Lumbar_Coeliac_VS_P1_P2_P3_Stellate_Sphenopalatine | 0 | 0 | 0 | 1 | 1 | 1 | 0 | 0 |

| | | | | | | | | | | |
|---|---|---|---|---|---|---|---|---|---|---|
| NR2C1 | 0.513 | P4_Lumbar_Coeliac_VS_P1_P2_P3_Stellate_Sphenopalatine | 0 | 0 | 0 | 1 | 1 | 1 | 0 | 0 |
| SLX4 | 0.512 | Coeliac_VS_P1_P2_P3_P4_Lumbar_Stellate_Sphenopalatine | 0 | 0 | 0 | 0 | 0 | 1 | 0 | 0 |
| GM9993 | 0.512 | Coeliac_VS_P1_P2_P3_P4_Lumbar_Stellate_Sphenopalatine | 0 | 0 | 0 | 0 | 0 | 1 | 0 | 0 |
| KIF16B | 0.512 | P1_P2_P3_P4_Lumbar_Coeliac_VS_Stellate_Sphenopalatine | 1 | 1 | 1 | 1 | 1 | 1 | 0 | 0 |
| TUBGCP3 | 0.512 | P1_P2_P3_P4_Lumbar_Coeliac_VS_Stellate_Sphenopalatine | 1 | 1 | 1 | 1 | 1 | 1 | 0 | 0 |
| PWWP2A | 0.512 | P1_P2_P3_P4_Lumbar_Coeliac_VS_Stellate_Sphenopalatine | 1 | 1 | 1 | 1 | 1 | 1 | 0 | 0 |
| TMEM240 | 0.511 | P1_P2_P3_P4_Lumbar_Coeliac_VS_Stellate_Sphenopalatine | 1 | 1 | 1 | 1 | 1 | 1 | 0 | 0 |
| H2-K1 | 0.511 | P1_P2_P3_P4_Lumbar_Stellate_Sphenopalatine_VS_Coeliac | 1 | 1 | 1 | 1 | 1 | 0 | 1 | 1 |
| SHD | 0.511 | P1_P2_P3_Lumbar_Coeliac_Stellate_Sphenopalatine_VS_P4 | 1 | 1 | 1 | 0 | 1 | 1 | 1 | 1 |
| TSPAN4 | 0.511 | P1_P3_P4_Lumbar_Coeliac_Stellate_Sphenopalatine_VS_P2 | 1 | 0 | 1 | 1 | 1 | 1 | 1 | 1 |
| PPP2R1B | 0.511 | P1_P3_P4_Lumbar_Coeliac_Stellate_Sphenopalatine_VS_P2 | 1 | 0 | 1 | 1 | 1 | 1 | 1 | 1 |
| GM19412 | 0.51 | P3_P4_VS_P1_P2_Lumbar_Coeliac_Stellate_Sphenopalatine | 0 | 0 | 1 | 1 | 0 | 0 | 0 | 0 |
| FGF14 | 0.509 | P3_Lumbar_Coeliac_VS_P1_P2_P4_Stellate_Sphenopalatine | 0 | 0 | 1 | 0 | 1 | 1 | 0 | 0 |
| KLHL33 | 0.508 | Coeliac_VS_P1_P2_P3_P4_Lumbar_Stellate_Sphenopalatine | 0 | 0 | 0 | 0 | 0 | 1 | 0 | 0 |
| TLE1 | 0.508 | P1_P3_P4_Coeliac_VS_P2_Lumbar_Stellate_Sphenopalatine | 1 | 0 | 1 | 1 | 0 | 1 | 0 | 0 |
| CDC40 | 0.508 | P4_Lumbar_Coeliac_VS_P1_P2_P3_Stellate_Sphenopalatine | 0 | 0 | 0 | 1 | 1 | 1 | 0 | 0 |
| MSANTD3 | 0.508 | P1_P2_P3_P4_Lumbar_Coeliac_VS_Stellate_Sphenopalatine | 1 | 1 | 1 | 1 | 1 | 1 | 0 | 0 |
| STK32A | 0.508 | P1_P2_P3_P4_Lumbar_Coeliac_Stellate_VS_Sphenopalatine | 1 | 1 | 1 | 1 | 1 | 1 | 1 | 0 |
| ELF2 | 0.507 | Coeliac_VS_P1_P2_P3_P4_Lumbar_Stellate_Sphenopalatine | 0 | 0 | 0 | 0 | 0 | 1 | 0 | 0 |
| MANEAL | 0.507 | P4_Lumbar_Coeliac_VS_P1_P2_P3_Stellate_Sphenopalatine | 0 | 0 | 0 | 1 | 1 | 1 | 0 | 0 |
| GOLGA1 | 0.507 | P1_P2_P3_P4_Lumbar_Coeliac_VS_Stellate_Sphenopalatine | 1 | 1 | 1 | 1 | 1 | 1 | 0 | 0 |
| RYBP | 0.507 | P1_P2_P3_P4_Lumbar_Coeliac_VS_Stellate_Sphenopalatine | 1 | 1 | 1 | 1 | 1 | 1 | 0 | 0 |
| PIBF1 | 0.507 | P1_P2_P3_P4_Lumbar_Coeliac_VS_Stellate_Sphenopalatine | 1 | 1 | 1 | 1 | 1 | 1 | 0 | 0 |
| PCCA | 0.507 | P1_P2_P3_P4_Lumbar_Coeliac_VS_Stellate_Sphenopalatine | 1 | 1 | 1 | 1 | 1 | 1 | 0 | 0 |
| CHIC1 | 0.507 | P1_P2_P3_P4_Lumbar_Coeliac_VS_Stellate_Sphenopalatine | 1 | 1 | 1 | 1 | 1 | 1 | 0 | 0 |
| BAIAP2 | 0.506 | P1_P2_P3_P4_Lumbar_Coeliac_VS_Stellate_Sphenopalatine | 1 | 1 | 1 | 1 | 1 | 1 | 0 | 0 |
| EFNA1 | 0.506 | P1_P2_P3_P4_Lumbar_Coeliac_Stellate_VS_Sphenopalatine | 1 | 1 | 1 | 1 | 1 | 1 | 1 | 0 |
| HRK | 0.506 | P4_Lumbar_Coeliac_Sphenopalatine_VS_P1_P2_P3_Stellate | 0 | 0 | 0 | 1 | 1 | 1 | 0 | 1 |
| CYB561 | 0.506 | P1_P3_P4_Lumbar_Coeliac_Stellate_Sphenopalatine_VS_P2 | 1 | 0 | 1 | 1 | 1 | 1 | 1 | 1 |
| CCDC170 | 0.505 | Coeliac_VS_P1_P2_P3_P4_Lumbar_Stellate_Sphenopalatine | 0 | 0 | 0 | 0 | 0 | 1 | 0 | 0 |

| | | | | | | | | | | |
|---|---|---|---|---|---|---|---|---|---|---|
| RNF215 | 0.505 | P1_P2_P3_P4_Lumbar_Coeliac_VS_Stellate_Sphenopalatine | 1 | 1 | 1 | 1 | 1 | 1 | 0 | 0 |
| MBP | 0.505 | P1_P2_P4_Lumbar_Coeliac_Sphenopalatine_VS_P3_Stellate | 1 | 1 | 0 | 1 | 1 | 1 | 0 | 1 |
| UTP14A | 0.504 | P1_P2_P3_P4_Lumbar_Coeliac_VS_Stellate_Sphenopalatine | 1 | 1 | 1 | 1 | 1 | 1 | 0 | 0 |
| KDM5D | 0.504 | P1_P2_P3_P4_Lumbar_Coeliac_VS_Stellate_Sphenopalatine | 1 | 1 | 1 | 1 | 1 | 1 | 0 | 0 |
| RET | 0.503 | P1_P2_P3_P4_Sphenopalatine_VS_Lumbar_Coeliac_Stellate | 1 | 1 | 1 | 1 | 0 | 0 | 0 | 1 |
| BRIX1 | 0.502 | P4_Lumbar_Coeliac_VS_P1_P2_P3_Stellate_Sphenopalatine | 0 | 0 | 0 | 1 | 1 | 1 | 0 | 0 |
| AP5M1 | 0.5 | P1_P2_P3_P4_Lumbar_Coeliac_VS_Stellate_Sphenopalatine | 1 | 1 | 1 | 1 | 1 | 1 | 0 | 0 |
| MTM1 | 0.5 | P1_P2_P3_P4_Lumbar_Coeliac_VS_Stellate_Sphenopalatine | 1 | 1 | 1 | 1 | 1 | 1 | 0 | 0 |
| TMEM250-PS | 0.5 | P1_P2_P3_P4_Sphenopalatine_VS_Lumbar_Coeliac_Stellate | 1 | 1 | 1 | 1 | 0 | 0 | 0 | 1 |
| GLTP | 0.499 | P1_P2_P3_P4_Lumbar_Coeliac_VS_Stellate_Sphenopalatine | 1 | 1 | 1 | 1 | 1 | 1 | 0 | 0 |
| ZFP277 | 0.499 | P1_P2_P3_P4_Lumbar_Coeliac_VS_Stellate_Sphenopalatine | 1 | 1 | 1 | 1 | 1 | 1 | 0 | 0 |
| EHD2 | 0.499 | P1_P2_P4_Sphenopalatine_VS_P3_Lumbar_Coeliac_Stellate | 1 | 1 | 0 | 1 | 0 | 0 | 0 | 1 |
| TRIM33 | 0.498 | P1_P2_P3_P4_Lumbar_Coeliac_VS_Stellate_Sphenopalatine | 1 | 1 | 1 | 1 | 1 | 1 | 0 | 0 |
| CNNM3 | 0.498 | P1_P2_P4_Lumbar_Coeliac_Sphenopalatine_VS_P3_Stellate | 1 | 1 | 0 | 1 | 1 | 1 | 0 | 1 |
| NIPAL2 | 0.497 | P4_Lumbar_Coeliac_VS_P1_P2_P3_Stellate_Sphenopalatine | 0 | 0 | 0 | 1 | 1 | 1 | 0 | 0 |
| ZFYVE28 | 0.497 | P1_P2_P3_P4_Lumbar_Coeliac_VS_Stellate_Sphenopalatine | 1 | 1 | 1 | 1 | 1 | 1 | 0 | 0 |
| ALAS1 | 0.496 | P4_Lumbar_Coeliac_VS_P1_P2_P3_Stellate_Sphenopalatine | 0 | 0 | 0 | 1 | 1 | 1 | 0 | 0 |
| GCNT2 | 0.496 | P1_P2_P3_Lumbar_Coeliac_Stellate_Sphenopalatine_VS_P4 | 1 | 1 | 1 | 0 | 1 | 1 | 1 | 1 |
| ZUP1 | 0.495 | P1_P2_P3_P4_Lumbar_Coeliac_VS_Stellate_Sphenopalatine | 1 | 1 | 1 | 1 | 1 | 1 | 0 | 0 |
| TMEM47 | 0.495 | P1_P2_P3_P4_Lumbar_Coeliac_VS_Stellate_Sphenopalatine | 1 | 1 | 1 | 1 | 1 | 1 | 0 | 0 |
| VAMP1 | 0.495 | P1_P2_P3_P4_Sphenopalatine_VS_Lumbar_Coeliac_Stellate | 1 | 1 | 1 | 1 | 0 | 0 | 0 | 1 |
| ZBTB8A | 0.494 | P1_P2_P3_P4_Lumbar_Coeliac_VS_Stellate_Sphenopalatine | 1 | 1 | 1 | 1 | 1 | 1 | 0 | 0 |
| FAM241A | 0.493 | P1_P2_P3_Lumbar_Coeliac_VS_P4_Stellate_Sphenopalatine | 1 | 1 | 1 | 0 | 1 | 1 | 0 | 0 |
| PRICKLE2 | 0.492 | P2_P4_VS_P1_P3_Lumbar_Coeliac_Stellate_Sphenopalatine | 0 | 1 | 0 | 1 | 0 | 0 | 0 | 0 |
| NAAA | 0.492 | P3_P4_Lumbar_VS_P1_P2_Coeliac_Stellate_Sphenopalatine | 0 | 0 | 1 | 1 | 1 | 0 | 0 | 0 |
| NBN | 0.492 | P3_Lumbar_Coeliac_VS_P1_P2_P4_Stellate_Sphenopalatine | 0 | 0 | 1 | 0 | 1 | 1 | 0 | 0 |
| PSD3 | 0.492 | P2_P3_Lumbar_Coeliac_Sphenopalatine_VS_P1_P4_Stellate | 0 | 1 | 1 | 0 | 1 | 1 | 0 | 1 |
| CCDC25 | 0.492 | P1_P3_P4_Lumbar_Coeliac_Sphenopalatine_VS_P2_Stellate | 1 | 0 | 1 | 1 | 1 | 1 | 0 | 1 |
| TPCN1 | 0.491 | P2_P4_VS_P1_P3_Lumbar_Coeliac_Stellate_Sphenopalatine | 0 | 1 | 0 | 1 | 0 | 0 | 0 | 0 |
| WDR60 | 0.491 | P4_Lumbar_Coeliac_VS_P1_P2_P3_Stellate_Sphenopalatine | 0 | 0 | 0 | 1 | 1 | 1 | 0 | 0 |

| | | | | | | | | | | |
|---|---|---|---|---|---|---|---|---|---|---|
| CWC22 | 0.491 | P1_P4_Lumbar_Coeliac_VS_P2_P3_Stellate_Sphenopalatine | 1 | 0 | 0 | 1 | 1 | 1 | 0 | 0 |
| PELO | 0.491 | P1_P2_P3_P4_Lumbar_Coeliac_Stellate_VS_Sphenopalatine | 1 | 1 | 1 | 1 | 1 | 1 | 1 | 0 |
| TES | 0.491 | P1_P2_P3_Lumbar_Coeliac_Stellate_Sphenopalatine_VS_P4 | 1 | 1 | 1 | 0 | 1 | 1 | 1 | 1 |
| NOVA1 | 0.49 | P4_Lumbar_Coeliac_VS_P1_P2_P3_Stellate_Sphenopalatine | 0 | 0 | 0 | 1 | 1 | 1 | 0 | 0 |
| GALNT2 | 0.49 | P1_P2_P3_P4_Lumbar_Coeliac_VS_Stellate_Sphenopalatine | 1 | 1 | 1 | 1 | 1 | 1 | 0 | 0 |
| RANBP2 | 0.49 | P1_P2_P3_P4_Lumbar_Coeliac_VS_Stellate_Sphenopalatine | 1 | 1 | 1 | 1 | 1 | 1 | 0 | 0 |
| MILL2 | 0.489 | P1_P2_P3_Lumbar_Coeliac_VS_P4_Stellate_Sphenopalatine | 1 | 1 | 1 | 0 | 1 | 1 | 0 | 0 |
| RBBP8 | 0.489 | P1_P2_P3_P4_Lumbar_Coeliac_VS_Stellate_Sphenopalatine | 1 | 1 | 1 | 1 | 1 | 1 | 0 | 0 |
| GM43462 | 0.488 | P4_Lumbar_Coeliac_VS_P1_P2_P3_Stellate_Sphenopalatine | 0 | 0 | 0 | 1 | 1 | 1 | 0 | 0 |
| CTDSPL2 | 0.488 | P1_P2_P3_P4_Lumbar_Coeliac_VS_Stellate_Sphenopalatine | 1 | 1 | 1 | 1 | 1 | 1 | 0 | 0 |
| GNASAS1 | 0.488 | P1_P2_P4_Lumbar_Coeliac_Sphenopalatine_VS_P3_Stellate | 1 | 1 | 0 | 1 | 1 | 1 | 0 | 1 |
| GM31107 | 0.487 | P3_P4_VS_P1_P2_Lumbar_Coeliac_Stellate_Sphenopalatine | 0 | 0 | 1 | 1 | 0 | 0 | 0 | 0 |
| KANSL1L | 0.487 | P1_P2_P3_P4_Lumbar_Coeliac_VS_Stellate_Sphenopalatine | 1 | 1 | 1 | 1 | 1 | 1 | 0 | 0 |
| RNF24 | 0.486 | P4_Lumbar_Coeliac_VS_P1_P2_P3_Stellate_Sphenopalatine | 0 | 0 | 0 | 1 | 1 | 1 | 0 | 0 |
| PELP1 | 0.486 | P4_Lumbar_Coeliac_VS_P1_P2_P3_Stellate_Sphenopalatine | 0 | 0 | 0 | 1 | 1 | 1 | 0 | 0 |
| CCDC71L | 0.486 | P4_Lumbar_Coeliac_VS_P1_P2_P3_Stellate_Sphenopalatine | 0 | 0 | 0 | 1 | 1 | 1 | 0 | 0 |
| CACNB4 | 0.486 | P1_P2_P4_Lumbar_Coeliac_VS_P3_Stellate_Sphenopalatine | 1 | 1 | 0 | 1 | 1 | 1 | 0 | 0 |
| SETX | 0.486 | P1_P2_P3_P4_Lumbar_Coeliac_VS_Stellate_Sphenopalatine | 1 | 1 | 1 | 1 | 1 | 1 | 0 | 0 |
| CDKN1B | 0.486 | P1_P2_P3_P4_Lumbar_Coeliac_VS_Stellate_Sphenopalatine | 1 | 1 | 1 | 1 | 1 | 1 | 0 | 0 |
| ABCD2 | 0.485 | P3_P4_VS_P1_P2_Lumbar_Coeliac_Stellate_Sphenopalatine | 0 | 0 | 1 | 1 | 0 | 0 | 0 | 0 |
| MCPH1 | 0.485 | P1_P2_P3_P4_Lumbar_Coeliac_VS_Stellate_Sphenopalatine | 1 | 1 | 1 | 1 | 1 | 1 | 0 | 0 |
| KAT2B | 0.485 | P1_P2_P3_P4_Lumbar_Coeliac_VS_Stellate_Sphenopalatine | 1 | 1 | 1 | 1 | 1 | 1 | 0 | 0 |
| PLCB3 | 0.484 | P2_VS_P1_P3_P4_Lumbar_Coeliac_Stellate_Sphenopalatine | 0 | 1 | 0 | 0 | 0 | 0 | 0 | 0 |
| PRIM1 | 0.484 | P3_Lumbar_Coeliac_VS_P1_P2_P4_Stellate_Sphenopalatine | 0 | 0 | 1 | 0 | 1 | 1 | 0 | 0 |
| CHD2 | 0.484 | P1_P2_P3_P4_Lumbar_Coeliac_VS_Stellate_Sphenopalatine | 1 | 1 | 1 | 1 | 1 | 1 | 0 | 0 |
| DMD | 0.484 | P1_P4_Sphenopalatine_VS_P2_P3_Lumbar_Coeliac_Stellate | 1 | 0 | 0 | 1 | 0 | 0 | 0 | 1 |
| PPIL4 | 0.483 | P1_P2_P3_P4_Lumbar_Coeliac_VS_Stellate_Sphenopalatine | 1 | 1 | 1 | 1 | 1 | 1 | 0 | 0 |
| TNK2 | 0.483 | P1_P2_P3_P4_Lumbar_Sphenopalatine_VS_Coeliac_Stellate | 1 | 1 | 1 | 1 | 1 | 0 | 0 | 1 |
| STOM | 0.483 | P1_P2_P3_Lumbar_Coeliac_Stellate_Sphenopalatine_VS_P4 | 1 | 1 | 1 | 0 | 1 | 1 | 1 | 1 |
| OSGIN2 | 0.482 | P3_VS_P1_P2_P4_Lumbar_Coeliac_Stellate_Sphenopalatine | 0 | 0 | 1 | 0 | 0 | 0 | 0 | 0 |

| | | | | | | | | | | |
|---|---|---|---|---|---|---|---|---|---|---|
| TACO1 | 0.482 | P4_Lumbar_Coeliac_VS_P1_P2_P3_Stellate_Sphenopalatine | 0 | 0 | 0 | 1 | 1 | 1 | 0 | 0 |
| KDM4C | 0.482 | P1_P2_P3_P4_Lumbar_Coeliac_VS_Stellate_Sphenopalatine | 1 | 1 | 1 | 1 | 1 | 1 | 0 | 0 |
| ZFP507 | 0.482 | P1_P2_P3_P4_Lumbar_Coeliac_VS_Stellate_Sphenopalatine | 1 | 1 | 1 | 1 | 1 | 1 | 0 | 0 |
| E2F3 | 0.482 | P1_P2_P3_P4_Lumbar_Coeliac_VS_Stellate_Sphenopalatine | 1 | 1 | 1 | 1 | 1 | 1 | 0 | 0 |
| ZFP467 | 0.482 | P1_P2_P4_Lumbar_Coeliac_Sphenopalatine_VS_P3_Stellate | 1 | 1 | 0 | 1 | 1 | 1 | 0 | 1 |
| FAM229B | 0.481 | P1_P2_P3_P4_Sphenopalatine_VS_Lumbar_Coeliac_Stellate | 1 | 1 | 1 | 1 | 0 | 0 | 0 | 1 |
| PER1 | 0.48 | P2_P4_VS_P1_P3_Lumbar_Coeliac_Stellate_Sphenopalatine | 0 | 1 | 0 | 1 | 0 | 0 | 0 | 0 |
| TTC21B | 0.48 | P1_Lumbar_Coeliac_VS_P2_P3_P4_Stellate_Sphenopalatine | 1 | 0 | 0 | 0 | 1 | 1 | 0 | 0 |
| XRCC5 | 0.48 | P1_P2_P3_P4_Lumbar_Coeliac_Stellate_VS_Sphenopalatine | 1 | 1 | 1 | 1 | 1 | 1 | 1 | 0 |
| KMT2B | 0.479 | P2_Lumbar_Coeliac_VS_P1_P3_P4_Stellate_Sphenopalatine | 0 | 1 | 0 | 0 | 1 | 1 | 0 | 0 |
| RPS6KA6 | 0.479 | P2_P3_P4_Lumbar_Coeliac_VS_P1_Stellate_Sphenopalatine | 0 | 1 | 1 | 1 | 1 | 1 | 0 | 0 |
| LRRCC1 | 0.479 | P1_P2_P3_P4_Lumbar_Coeliac_VS_Stellate_Sphenopalatine | 1 | 1 | 1 | 1 | 1 | 1 | 0 | 0 |
| SUSD2 | 0.478 | P1_P2_P3_P4_VS_Lumbar_Coeliac_Stellate_Sphenopalatine | 1 | 1 | 1 | 1 | 0 | 0 | 0 | 0 |
| TMEM35A | 0.478 | P3_Lumbar_Coeliac_Stellate_VS_P1_P2_P4_Sphenopalatine | 0 | 0 | 1 | 0 | 1 | 1 | 1 | 0 |
| IFIT1 | 0.478 | P1_P4_Sphenopalatine_VS_P2_P3_Lumbar_Coeliac_Stellate | 1 | 0 | 0 | 1 | 0 | 0 | 0 | 1 |
| MAN1A | 0.478 | P1_P2_P3_P4_Coeliac_Sphenopalatine_VS_Lumbar_Stellate | 1 | 1 | 1 | 1 | 0 | 1 | 0 | 1 |
| D130043K22 | 0.477 | P4_Lumbar_VS_P1_P2_P3_Coeliac_Stellate_Sphenopalatine | 0 | 0 | 0 | 1 | 1 | 0 | 0 | 0 |
| ZBTB37 | 0.477 | P2_Lumbar_Coeliac_VS_P1_P3_P4_Stellate_Sphenopalatine | 0 | 1 | 0 | 0 | 1 | 1 | 0 | 0 |
| TNNI1 | 0.477 | P3_Lumbar_Coeliac_VS_P1_P2_P4_Stellate_Sphenopalatine | 0 | 0 | 1 | 0 | 1 | 1 | 0 | 0 |
| CEP250 | 0.477 | P4_Lumbar_Coeliac_VS_P1_P2_P3_Stellate_Sphenopalatine | 0 | 0 | 0 | 1 | 1 | 1 | 0 | 0 |
| LACTB | 0.477 | P1_P3_P4_Lumbar_Coeliac_VS_P2_Stellate_Sphenopalatine | 1 | 0 | 1 | 1 | 1 | 1 | 0 | 0 |
| PNPT1 | 0.476 | P1_P2_P3_P4_Lumbar_Coeliac_VS_Stellate_Sphenopalatine | 1 | 1 | 1 | 1 | 1 | 1 | 0 | 0 |
| UGGT2 | 0.476 | P1_P2_P3_P4_Lumbar_Coeliac_VS_Stellate_Sphenopalatine | 1 | 1 | 1 | 1 | 1 | 1 | 0 | 0 |
| CDC42EP3 | 0.476 | P1_P2_P3_P4_Lumbar_Coeliac_Stellate_VS_Sphenopalatine | 1 | 1 | 1 | 1 | 1 | 1 | 1 | 0 |
| JAG1 | 0.476 | Sphenopalatine_VS_P1_P2_P3_P4_Lumbar_Coeliac_Stellate | 0 | 0 | 0 | 0 | 0 | 0 | 0 | 1 |
| PLS3 | 0.476 | P1_P2_P3_P4_Sphenopalatine_VS_Lumbar_Coeliac_Stellate | 1 | 1 | 1 | 1 | 0 | 0 | 0 | 1 |
| PLPP4 | 0.475 | P2_VS_P1_P3_P4_Lumbar_Coeliac_Stellate_Sphenopalatine | 0 | 1 | 0 | 0 | 0 | 0 | 0 | 0 |
| NR4A2 | 0.475 | P3_P4_VS_P1_P2_Lumbar_Coeliac_Stellate_Sphenopalatine | 0 | 0 | 1 | 1 | 0 | 0 | 0 | 0 |
| RRAD | 0.475 | P3_P4_VS_P1_P2_Lumbar_Coeliac_Stellate_Sphenopalatine | 0 | 0 | 1 | 1 | 0 | 0 | 0 | 0 |
| ACAD9 | 0.475 | P1_Lumbar_Coeliac_VS_P2_P3_P4_Stellate_Sphenopalatine | 1 | 0 | 0 | 0 | 1 | 1 | 0 | 0 |

| | | | | | | | | | | |
|---|---|---|---|---|---|---|---|---|---|---|
| TUBGCP2 | 0.475 | P4_Lumbar_Coeliac_VS_P1_P2_P3_Stellate_Sphenopalatine | 0 | 0 | 0 | 1 | 1 | 1 | 0 | 0 |
| LEO1 | 0.475 | P4_Lumbar_Coeliac_VS_P1_P2_P3_Stellate_Sphenopalatine | 0 | 0 | 0 | 1 | 1 | 1 | 0 | 0 |
| ORC3 | 0.475 | P1_P2_P3_P4_Lumbar_Coeliac_VS_Stellate_Sphenopalatine | 1 | 1 | 1 | 1 | 1 | 1 | 0 | 0 |
| MTHFD1 | 0.475 | P1_P2_P3_P4_Lumbar_Coeliac_VS_Stellate_Sphenopalatine | 1 | 1 | 1 | 1 | 1 | 1 | 0 | 0 |
| UNC79 | 0.475 | P1_P2_P3_P4_Lumbar_Coeliac_VS_Stellate_Sphenopalatine | 1 | 1 | 1 | 1 | 1 | 1 | 0 | 0 |
| SLC22A23 | 0.474 | P1_P4_VS_P2_P3_Lumbar_Coeliac_Stellate_Sphenopalatine | 1 | 0 | 0 | 1 | 0 | 0 | 0 | 0 |
| ADGRG1 | 0.474 | P3_P4_Lumbar_Coeliac_VS_P1_P2_Stellate_Sphenopalatine | 0 | 0 | 1 | 1 | 1 | 1 | 0 | 0 |
| CBLL1 | 0.474 | P1_P2_P3_P4_Lumbar_Coeliac_VS_Stellate_Sphenopalatine | 1 | 1 | 1 | 1 | 1 | 1 | 0 | 0 |
| FANCM | 0.474 | P1_P2_P3_P4_Lumbar_Coeliac_VS_Stellate_Sphenopalatine | 1 | 1 | 1 | 1 | 1 | 1 | 0 | 0 |
| DRC1 | 0.473 | P3_VS_P1_P2_P4_Lumbar_Coeliac_Stellate_Sphenopalatine | 0 | 0 | 1 | 0 | 0 | 0 | 0 | 0 |
| HSPA1B | 0.473 | P4_Lumbar_Coeliac_VS_P1_P2_P3_Stellate_Sphenopalatine | 0 | 0 | 0 | 1 | 1 | 1 | 0 | 0 |
| FGF11 | 0.472 | P1_P2_P4_Sphenopalatine_VS_P3_Lumbar_Coeliac_Stellate | 1 | 1 | 0 | 1 | 0 | 0 | 0 | 1 |
| NRIP3 | 0.471 | P2_VS_P1_P3_P4_Lumbar_Coeliac_Stellate_Sphenopalatine | 0 | 1 | 0 | 0 | 0 | 0 | 0 | 0 |
| TIMP3 | 0.471 | P2_P4_VS_P1_P3_Lumbar_Coeliac_Stellate_Sphenopalatine | 0 | 1 | 0 | 1 | 0 | 0 | 0 | 0 |
| FST | 0.471 | P1_P3_P4_Lumbar_Coeliac_Stellate_Sphenopalatine_VS_P2 | 1 | 0 | 1 | 1 | 1 | 1 | 1 | 1 |
| PPM1K | 0.47 | P2_VS_P1_P3_P4_Lumbar_Coeliac_Stellate_Sphenopalatine | 0 | 1 | 0 | 0 | 0 | 0 | 0 | 0 |
| UTP23 | 0.47 | P1_Lumbar_Coeliac_VS_P2_P3_P4_Stellate_Sphenopalatine | 1 | 0 | 0 | 0 | 1 | 1 | 0 | 0 |
| ERCC5 | 0.47 | P4_Lumbar_Coeliac_VS_P1_P2_P3_Stellate_Sphenopalatine | 0 | 0 | 0 | 1 | 1 | 1 | 0 | 0 |
| ASAH2 | 0.469 | P3_VS_P1_P2_P4_Lumbar_Coeliac_Stellate_Sphenopalatine | 0 | 0 | 1 | 0 | 0 | 0 | 0 | 0 |
| ACE | 0.469 | P1_P2_P3_Lumbar_Coeliac_Stellate_Sphenopalatine_VS_P4 | 1 | 1 | 1 | 0 | 1 | 1 | 1 | 1 |
| KIF22 | 0.468 | P1_P2_P4_Sphenopalatine_VS_P3_Lumbar_Coeliac_Stellate | 1 | 1 | 0 | 1 | 0 | 0 | 0 | 1 |
| HPCAL1 | 0.468 | P1_P2_P3_Lumbar_Coeliac_Stellate_Sphenopalatine_VS_P4 | 1 | 1 | 1 | 0 | 1 | 1 | 1 | 1 |
| 2210408I21R | 0.467 | P4_Lumbar_Coeliac_VS_P1_P2_P3_Stellate_Sphenopalatine | 0 | 0 | 0 | 1 | 1 | 1 | 0 | 0 |
| ZFP518A | 0.467 | P4_Lumbar_Coeliac_VS_P1_P2_P3_Stellate_Sphenopalatine | 0 | 0 | 0 | 1 | 1 | 1 | 0 | 0 |
| NME7 | 0.466 | P3_Lumbar_Coeliac_VS_P1_P2_P4_Stellate_Sphenopalatine | 0 | 0 | 1 | 0 | 1 | 1 | 0 | 0 |
| GPX3 | 0.465 | P4_Stellate_VS_P1_P2_P3_Lumbar_Coeliac_Sphenopalatine | 0 | 0 | 0 | 1 | 0 | 0 | 1 | 0 |
| SBK1 | 0.465 | P1_P2_P3_Lumbar_Coeliac_Stellate_Sphenopalatine_VS_P4 | 1 | 1 | 1 | 0 | 1 | 1 | 1 | 1 |
| STK4 | 0.463 | P4_Lumbar_Coeliac_VS_P1_P2_P3_Stellate_Sphenopalatine | 0 | 0 | 0 | 1 | 1 | 1 | 0 | 0 |
| TCEANC2 | 0.463 | P4_Lumbar_Coeliac_VS_P1_P2_P3_Stellate_Sphenopalatine | 0 | 0 | 0 | 1 | 1 | 1 | 0 | 0 |
| GPM6B | 0.462 | P2_P4_VS_P1_P3_Lumbar_Coeliac_Stellate_Sphenopalatine | 0 | 1 | 0 | 1 | 0 | 0 | 0 | 0 |

| | | | | | | | | | | |
|---|---|---|---|---|---|---|---|---|---|---|
| TRMT2B | 0.462 | P2_Lumbar_Coeliac_VS_P1_P3_P4_Stellate_Sphenopalatine | 0 | 1 | 0 | 0 | 1 | 1 | 0 | 0 |
| SLC24A3 | 0.462 | P2_P4_Sphenopalatine_VS_P1_P3_Lumbar_Coeliac_Stellate | 0 | 1 | 0 | 1 | 0 | 0 | 0 | 1 |
| STK32C | 0.462 | P1_P2_P4_Sphenopalatine_VS_P3_Lumbar_Coeliac_Stellate | 1 | 1 | 0 | 1 | 0 | 0 | 0 | 1 |
| GPR161 | 0.461 | P1_P2_P3_Lumbar_Coeliac_VS_P4_Stellate_Sphenopalatine | 1 | 1 | 1 | 0 | 1 | 1 | 0 | 0 |
| TAF1B | 0.461 | P4_Lumbar_Coeliac_VS_P1_P2_P3_Stellate_Sphenopalatine | 0 | 0 | 0 | 1 | 1 | 1 | 0 | 0 |
| IRF6 | 0.46 | P1_P2_VS_P3_P4_Lumbar_Coeliac_Stellate_Sphenopalatine | 1 | 1 | 0 | 0 | 0 | 0 | 0 | 0 |
| SAMD4 | 0.46 | P1_P2_P3_Lumbar_Coeliac_Stellate_Sphenopalatine_VS_P4 | 1 | 1 | 1 | 0 | 1 | 1 | 1 | 1 |
| CEP85L | 0.459 | P1_P2_P4_Lumbar_Coeliac_VS_P3_Stellate_Sphenopalatine | 1 | 1 | 0 | 1 | 1 | 1 | 0 | 0 |
| EFCAB10 | 0.458 | P3_P4_Lumbar_Coeliac_Stellate_VS_P1_P2_Sphenopalatine | 0 | 0 | 1 | 1 | 1 | 1 | 1 | 0 |
| GM3294 | 0.458 | P3_Sphenopalatine_VS_P1_P2_P4_Lumbar_Coeliac_Stellate | 0 | 0 | 1 | 0 | 0 | 0 | 0 | 1 |
| MPPED2 | 0.458 | P1_P2_P3_Lumbar_Coeliac_Stellate_Sphenopalatine_VS_P4 | 1 | 1 | 1 | 0 | 1 | 1 | 1 | 1 |
| ZFP36L2 | 0.457 | P1_P2_P4_Lumbar_Coeliac_VS_P3_Stellate_Sphenopalatine | 1 | 1 | 0 | 1 | 1 | 1 | 0 | 0 |
| RASSF8 | 0.456 | P2_VS_P1_P3_P4_Lumbar_Coeliac_Stellate_Sphenopalatine | 0 | 1 | 0 | 0 | 0 | 0 | 0 | 0 |
| ADCYAP1R1 | 0.455 | P1_P2_P3_Lumbar_Coeliac_Stellate_Sphenopalatine_VS_P4 | 1 | 1 | 1 | 0 | 1 | 1 | 1 | 1 |
| NHS | 0.454 | P1_P3_P4_Lumbar_Coeliac_Stellate_Sphenopalatine_VS_P2 | 1 | 0 | 1 | 1 | 1 | 1 | 1 | 1 |
| PRKCI | 0.453 | P2_VS_P1_P3_P4_Lumbar_Coeliac_Stellate_Sphenopalatine | 0 | 1 | 0 | 0 | 0 | 0 | 0 | 0 |
| SHROOM2 | 0.452 | P3_VS_P1_P2_P4_Lumbar_Coeliac_Stellate_Sphenopalatine | 0 | 0 | 1 | 0 | 0 | 0 | 0 | 0 |
| FKBP9 | 0.452 | P2_P4_VS_P1_P3_Lumbar_Coeliac_Stellate_Sphenopalatine | 0 | 1 | 0 | 1 | 0 | 0 | 0 | 0 |
| BAZ2A | 0.452 | P2_Lumbar_Coeliac_VS_P1_P3_P4_Stellate_Sphenopalatine | 0 | 1 | 0 | 0 | 1 | 1 | 0 | 0 |
| SF3A2 | 0.451 | P4_Lumbar_Coeliac_VS_P1_P2_P3_Stellate_Sphenopalatine | 0 | 0 | 0 | 1 | 1 | 1 | 0 | 0 |
| PPM1H | 0.451 | P1_P2_P4_Sphenopalatine_VS_P3_Lumbar_Coeliac_Stellate | 1 | 1 | 0 | 1 | 0 | 0 | 0 | 1 |
| FGF13 | 0.451 | P1_P2_P3_P4_Lumbar_Sphenopalatine_VS_Coeliac_Stellate | 1 | 1 | 1 | 1 | 1 | 0 | 0 | 1 |
| ZBTB46 | 0.451 | P1_P2_P4_Lumbar_Coeliac_Sphenopalatine_VS_P3_Stellate | 1 | 1 | 0 | 1 | 1 | 1 | 0 | 1 |
| TXNDC16 | 0.451 | P1_P2_P4_Lumbar_Coeliac_Sphenopalatine_VS_P3_Stellate | 1 | 1 | 0 | 1 | 1 | 1 | 0 | 1 |
| RP2 | 0.45 | P3_Lumbar_Coeliac_VS_P1_P2_P4_Stellate_Sphenopalatine | 0 | 0 | 1 | 0 | 1 | 1 | 0 | 0 |
| NEK3 | 0.45 | P4_Lumbar_Coeliac_VS_P1_P2_P3_Stellate_Sphenopalatine | 0 | 0 | 0 | 1 | 1 | 1 | 0 | 0 |
| RAMP1 | 0.45 | P1_P2_P3_P4_Lumbar_Coeliac_Stellate_VS_Sphenopalatine | 1 | 1 | 1 | 1 | 1 | 1 | 1 | 0 |
| METTL16 | 0.449 | P4_Lumbar_Coeliac_VS_P1_P2_P3_Stellate_Sphenopalatine | 0 | 0 | 0 | 1 | 1 | 1 | 0 | 0 |
| TTC28 | 0.448 | P4_Lumbar_Coeliac_VS_P1_P2_P3_Stellate_Sphenopalatine | 0 | 0 | 0 | 1 | 1 | 1 | 0 | 0 |
| KCTD12 | 0.448 | P1_P2_Lumbar_Coeliac_Stellate_Sphenopalatine_VS_P3_P4 | 1 | 1 | 0 | 0 | 1 | 1 | 1 | 1 |

| | | | | | | | | | | |
|---|---|---|---|---|---|---|---|---|---|---|
| PDP2 | 0.447 | P2_VS_P1_P3_P4_Lumbar_Coeliac_Stellate_Sphenopalatine | 0 | 1 | 0 | 0 | 0 | 0 | 0 | 0 |
| OMG | 0.447 | P1_P2_P3_P4_Sphenopalatine_VS_Lumbar_Coeliac_Stellate | 1 | 1 | 1 | 1 | 0 | 0 | 0 | 1 |
| PDE4D | 0.447 | P1_P4_Lumbar_Coeliac_Sphenopalatine_VS_P2_P3_Stellate | 1 | 0 | 0 | 1 | 1 | 1 | 0 | 1 |
| IGF2BP3 | 0.446 | Lumbar_Coeliac_Sphenopalatine_VS_P1_P2_P3_P4_Stellate | 0 | 0 | 0 | 0 | 1 | 1 | 0 | 1 |
| ERICH1 | 0.445 | P1_P2_Lumbar_Coeliac_VS_P3_P4_Stellate_Sphenopalatine | 1 | 1 | 0 | 0 | 1 | 1 | 0 | 0 |
| GPR158 | 0.444 | P2_P4_VS_P1_P3_Lumbar_Coeliac_Stellate_Sphenopalatine | 0 | 1 | 0 | 1 | 0 | 0 | 0 | 0 |
| STRN | 0.444 | P1_P2_P4_Lumbar_Coeliac_VS_P3_Stellate_Sphenopalatine | 1 | 1 | 0 | 1 | 1 | 1 | 0 | 0 |
| SUPT16 | 0.443 | P1_Lumbar_Coeliac_VS_P2_P3_P4_Stellate_Sphenopalatine | 1 | 0 | 0 | 0 | 1 | 1 | 0 | 0 |
| SLC4A7 | 0.443 | P3_Lumbar_Coeliac_VS_P1_P2_P4_Stellate_Sphenopalatine | 0 | 0 | 1 | 0 | 1 | 1 | 0 | 0 |
| FAM50A | 0.443 | P4_Lumbar_Coeliac_VS_P1_P2_P3_Stellate_Sphenopalatine | 0 | 0 | 0 | 1 | 1 | 1 | 0 | 0 |
| FAIM2 | 0.443 | P1_P2_P3_P4_Lumbar_Coeliac_Stellate_VS_Sphenopalatine | 1 | 1 | 1 | 1 | 1 | 1 | 1 | 0 |
| IGSF10 | 0.443 | P1_P3_P4_Lumbar_Coeliac_Sphenopalatine_VS_P2_Stellate | 1 | 0 | 1 | 1 | 1 | 1 | 0 | 1 |
| ZC3H18 | 0.442 | P4_Lumbar_Coeliac_VS_P1_P2_P3_Stellate_Sphenopalatine | 0 | 0 | 0 | 1 | 1 | 1 | 0 | 0 |
| SMAD9 | 0.442 | P1_P2_P3_P4_Coeliac_Sphenopalatine_VS_Lumbar_Stellate | 1 | 1 | 1 | 1 | 0 | 1 | 0 | 1 |
| VRK1 | 0.441 | P1_Lumbar_Coeliac_VS_P2_P3_P4_Stellate_Sphenopalatine | 1 | 0 | 0 | 0 | 1 | 1 | 0 | 0 |
| DNAJB5 | 0.441 | P3_Lumbar_Coeliac_VS_P1_P2_P4_Stellate_Sphenopalatine | 0 | 0 | 1 | 0 | 1 | 1 | 0 | 0 |
| MRPS22 | 0.441 | P4_Lumbar_Coeliac_VS_P1_P2_P3_Stellate_Sphenopalatine | 0 | 0 | 0 | 1 | 1 | 1 | 0 | 0 |
| PRRT1 | 0.44 | P2_P3_P4_VS_P1_Lumbar_Coeliac_Stellate_Sphenopalatine | 0 | 1 | 1 | 1 | 0 | 0 | 0 | 0 |
| SUN1 | 0.44 | P4_Lumbar_Coeliac_VS_P1_P2_P3_Stellate_Sphenopalatine | 0 | 0 | 0 | 1 | 1 | 1 | 0 | 0 |
| DYSF | 0.439 | P4_Lumbar_VS_P1_P2_P3_Coeliac_Stellate_Sphenopalatine | 0 | 0 | 0 | 1 | 1 | 0 | 0 | 0 |
| SCAF8 | 0.439 | P1_P2_P4_Lumbar_Coeliac_VS_P3_Stellate_Sphenopalatine | 1 | 1 | 0 | 1 | 1 | 1 | 0 | 0 |
| HUNK | 0.439 | P1_P2_P4_Lumbar_Coeliac_Sphenopalatine_VS_P3_Stellate | 1 | 1 | 0 | 1 | 1 | 1 | 0 | 1 |
| SCAMP2 | 0.438 | P4_Lumbar_Coeliac_VS_P1_P2_P3_Stellate_Sphenopalatine | 0 | 0 | 0 | 1 | 1 | 1 | 0 | 0 |
| RARS | 0.438 | P4_Lumbar_Coeliac_VS_P1_P2_P3_Stellate_Sphenopalatine | 0 | 0 | 0 | 1 | 1 | 1 | 0 | 0 |
| ATRIP | 0.438 | P1_P3_P4_Lumbar_Coeliac_VS_P2_Stellate_Sphenopalatine | 1 | 0 | 1 | 1 | 1 | 1 | 0 | 0 |
| DENND2A | 0.438 | P1_P2_P3_P4_Lumbar_Coeliac_Stellate_VS_Sphenopalatine | 1 | 1 | 1 | 1 | 1 | 1 | 1 | 0 |
| ANKRD50 | 0.437 | P1_P2_P4_Lumbar_Coeliac_Sphenopalatine_VS_P3_Stellate | 1 | 1 | 0 | 1 | 1 | 1 | 0 | 1 |
| 4930578M01 | 0.436 | P3_Lumbar_Coeliac_VS_P1_P2_P4_Stellate_Sphenopalatine | 0 | 0 | 1 | 0 | 1 | 1 | 0 | 0 |
| WRAP53 | 0.436 | P1_P4_Lumbar_Coeliac_VS_P2_P3_Stellate_Sphenopalatine | 1 | 0 | 0 | 1 | 1 | 1 | 0 | 0 |
| TMEM229A | 0.435 | P3_P4_VS_P1_P2_Lumbar_Coeliac_Stellate_Sphenopalatine | 0 | 0 | 1 | 1 | 0 | 0 | 0 | 0 |

| | | | | | | | | | | |
|---|---|---|---|---|---|---|---|---|---|---|
| SMUG1 | 0.435 | P2_P3_Lumbar_Coeliac_VS_P1_P4_Stellate_Sphenopalatine | 0 | 1 | 1 | 0 | 1 | 1 | 0 | 0 |
| ABHD17C | 0.435 | P1_P2_P3_P4_Lumbar_Coeliac_Stellate_VS_Sphenopalatine | 1 | 1 | 1 | 1 | 1 | 1 | 1 | 0 |
| DOC2B | 0.435 | P1_P2_P3_Lumbar_Coeliac_Stellate_Sphenopalatine_VS_P4 | 1 | 1 | 1 | 0 | 1 | 1 | 1 | 1 |
| SLC39A11 | 0.434 | P3_VS_P1_P2_P4_Lumbar_Coeliac_Stellate_Sphenopalatine | 0 | 0 | 1 | 0 | 0 | 0 | 0 | 0 |
| FGD4 | 0.434 | P4_Lumbar_Coeliac_VS_P1_P2_P3_Stellate_Sphenopalatine | 0 | 0 | 0 | 1 | 1 | 1 | 0 | 0 |
| KDM3B | 0.434 | P4_Lumbar_Coeliac_VS_P1_P2_P3_Stellate_Sphenopalatine | 0 | 0 | 0 | 1 | 1 | 1 | 0 | 0 |
| LIMCH1 | 0.434 | Sphenopalatine_VS_P1_P2_P3_P4_Lumbar_Coeliac_Stellate | 0 | 0 | 0 | 0 | 0 | 0 | 0 | 1 |
| GRID2 | 0.434 | P1_P2_P4_Lumbar_Coeliac_Sphenopalatine_VS_P3_Stellate | 1 | 1 | 0 | 1 | 1 | 1 | 0 | 1 |
| FBXO16 | 0.433 | P3_Lumbar_Coeliac_VS_P1_P2_P4_Stellate_Sphenopalatine | 0 | 0 | 1 | 0 | 1 | 1 | 0 | 0 |
| ADGRL3 | 0.433 | P1_P3_P4_Lumbar_Coeliac_Stellate_Sphenopalatine_VS_P2 | 1 | 0 | 1 | 1 | 1 | 1 | 1 | 1 |
| AMOTL2 | 0.432 | P1_P2_VS_P3_P4_Lumbar_Coeliac_Stellate_Sphenopalatine | 1 | 1 | 0 | 0 | 0 | 0 | 0 | 0 |
| ROM1 | 0.432 | Sphenopalatine_VS_P1_P2_P3_P4_Lumbar_Coeliac_Stellate | 0 | 0 | 0 | 0 | 0 | 0 | 0 | 1 |
| CLDN34C1 | 0.431 | P4_Lumbar_Coeliac_VS_P1_P2_P3_Stellate_Sphenopalatine | 0 | 0 | 0 | 1 | 1 | 1 | 0 | 0 |
| EXT1 | 0.43 | P3_P4_Lumbar_Coeliac_VS_P1_P2_Stellate_Sphenopalatine | 0 | 0 | 1 | 1 | 1 | 1 | 0 | 0 |
| CELF2 | 0.429 | P4_Sphenopalatine_VS_P1_P2_P3_Lumbar_Coeliac_Stellate | 0 | 0 | 0 | 1 | 0 | 0 | 0 | 1 |
| TRIM21 | 0.429 | P1_P2_P4_Lumbar_Coeliac_Sphenopalatine_VS_P3_Stellate | 1 | 1 | 0 | 1 | 1 | 1 | 0 | 1 |
| EPB41L2 | 0.429 | P1_P2_P4_Lumbar_Coeliac_Sphenopalatine_VS_P3_Stellate | 1 | 1 | 0 | 1 | 1 | 1 | 0 | 1 |
| UCP2 | 0.429 | P1_P3_P4_Lumbar_Coeliac_Sphenopalatine_VS_P2_Stellate | 1 | 0 | 1 | 1 | 1 | 1 | 0 | 1 |
| ACAP2 | 0.428 | P1_P2_P4_Lumbar_Coeliac_Sphenopalatine_VS_P3_Stellate | 1 | 1 | 0 | 1 | 1 | 1 | 0 | 1 |
| SERTAD4 | 0.427 | P2_Lumbar_Coeliac_VS_P1_P3_P4_Stellate_Sphenopalatine | 0 | 1 | 0 | 0 | 1 | 1 | 0 | 0 |
| CLMP | 0.427 | P1_P2_P3_Lumbar_Coeliac_Stellate_Sphenopalatine_VS_P4 | 1 | 1 | 1 | 0 | 1 | 1 | 1 | 1 |
| QSER1 | 0.426 | P1_Lumbar_Coeliac_VS_P2_P3_P4_Stellate_Sphenopalatine | 1 | 0 | 0 | 0 | 1 | 1 | 0 | 0 |
| WDR20 | 0.426 | P1_Lumbar_Coeliac_VS_P2_P3_P4_Stellate_Sphenopalatine | 1 | 0 | 0 | 0 | 1 | 1 | 0 | 0 |
| TSPAN5 | 0.426 | P4_Lumbar_Coeliac_VS_P1_P2_P3_Stellate_Sphenopalatine | 0 | 0 | 0 | 1 | 1 | 1 | 0 | 0 |
| ACADS | 0.426 | P1_P2_P3_P4_Lumbar_Coeliac_Stellate_VS_Sphenopalatine | 1 | 1 | 1 | 1 | 1 | 1 | 1 | 0 |
| GALNT13 | 0.426 | P3_Sphenopalatine_VS_P1_P2_P4_Lumbar_Coeliac_Stellate | 0 | 0 | 1 | 0 | 0 | 0 | 0 | 1 |
| HIST1H2BC | 0.425 | P2_VS_P1_P3_P4_Lumbar_Coeliac_Stellate_Sphenopalatine | 0 | 1 | 0 | 0 | 0 | 0 | 0 | 0 |
| SENP1 | 0.425 | P3_Lumbar_Coeliac_VS_P1_P2_P4_Stellate_Sphenopalatine | 0 | 0 | 1 | 0 | 1 | 1 | 0 | 0 |
| TASOR | 0.425 | P4_Lumbar_Coeliac_VS_P1_P2_P3_Stellate_Sphenopalatine | 0 | 0 | 0 | 1 | 1 | 1 | 0 | 0 |
| ITGA6 | 0.425 | Lumbar_Sphenopalatine_VS_P1_P2_P3_P4_Coeliac_Stellate | 0 | 0 | 0 | 0 | 1 | 0 | 0 | 1 |

| | | | | | | | | | | |
|---|---|---|---|---|---|---|---|---|---|---|
| SLC2A13 | 0.424 | P3_VS_P1_P2_P4_Lumbar_Coeliac_Stellate_Sphenopalatine | 0 | 0 | 1 | 0 | 0 | 0 | 0 | 0 |
| sept-10 | 0.424 | P1_P2_P4_Lumbar_Coeliac_VS_P3_Stellate_Sphenopalatine | 1 | 1 | 0 | 1 | 1 | 1 | 0 | 0 |
| CAPN9 | 0.424 | P4_Lumbar_Coeliac_Stellate_VS_P1_P2_P3_Sphenopalatine | 0 | 0 | 0 | 1 | 1 | 1 | 1 | 0 |
| CCDC77 | 0.423 | P4_Lumbar_Coeliac_VS_P1_P2_P3_Stellate_Sphenopalatine | 0 | 0 | 0 | 1 | 1 | 1 | 0 | 0 |
| SLITRK1 | 0.423 | P1_P4_Lumbar_Coeliac_VS_P2_P3_Stellate_Sphenopalatine | 1 | 0 | 0 | 1 | 1 | 1 | 0 | 0 |
| MLYCD | 0.422 | P3_Lumbar_Coeliac_VS_P1_P2_P4_Stellate_Sphenopalatine | 0 | 0 | 1 | 0 | 1 | 1 | 0 | 0 |
| DENND4C | 0.422 | P4_Lumbar_Coeliac_VS_P1_P2_P3_Stellate_Sphenopalatine | 0 | 0 | 0 | 1 | 1 | 1 | 0 | 0 |
| SPICE1 | 0.421 | P4_Lumbar_Coeliac_VS_P1_P2_P3_Stellate_Sphenopalatine | 0 | 0 | 0 | 1 | 1 | 1 | 0 | 0 |
| DYNC2LI1 | 0.421 | P4_Lumbar_Coeliac_VS_P1_P2_P3_Stellate_Sphenopalatine | 0 | 0 | 0 | 1 | 1 | 1 | 0 | 0 |
| HACE1 | 0.421 | P1_P2_P4_Lumbar_Coeliac_Sphenopalatine_VS_P3_Stellate | 1 | 1 | 0 | 1 | 1 | 1 | 0 | 1 |
| SIPA1L1 | 0.42 | P1_VS_P2_P3_P4_Lumbar_Coeliac_Stellate_Sphenopalatine | 1 | 0 | 0 | 0 | 0 | 0 | 0 | 0 |
| MLF1 | 0.42 | P3_VS_P1_P2_P4_Lumbar_Coeliac_Stellate_Sphenopalatine | 0 | 0 | 1 | 0 | 0 | 0 | 0 | 0 |
| GM19409 | 0.42 | P2_P4_VS_P1_P3_Lumbar_Coeliac_Stellate_Sphenopalatine | 0 | 1 | 0 | 1 | 0 | 0 | 0 | 0 |
| EVA1C | 0.42 | P3_P4_VS_P1_P2_Lumbar_Coeliac_Stellate_Sphenopalatine | 0 | 0 | 1 | 1 | 0 | 0 | 0 | 0 |
| TMC6 | 0.42 | P1_P2_P3_P4_Sphenopalatine_VS_Lumbar_Coeliac_Stellate | 1 | 1 | 1 | 1 | 0 | 0 | 0 | 1 |
| GNPTAB | 0.419 | P1_P2_P3_Lumbar_Coeliac_Stellate_Sphenopalatine_VS_P4 | 1 | 1 | 1 | 0 | 1 | 1 | 1 | 1 |
| FAM102A | 0.418 | P2_P4_VS_P1_P3_Lumbar_Coeliac_Stellate_Sphenopalatine | 0 | 1 | 0 | 1 | 0 | 0 | 0 | 0 |
| GABBR2 | 0.418 | P1_P2_P4_Lumbar_VS_P3_Coeliac_Stellate_Sphenopalatine | 1 | 1 | 0 | 1 | 1 | 0 | 0 | 0 |
| THOC5 | 0.418 | P1_Lumbar_Coeliac_VS_P2_P3_P4_Stellate_Sphenopalatine | 1 | 0 | 0 | 0 | 1 | 1 | 0 | 0 |
| HPS6 | 0.418 | P1_P4_Lumbar_Coeliac_VS_P2_P3_Stellate_Sphenopalatine | 1 | 0 | 0 | 1 | 1 | 1 | 0 | 0 |
| TMEM131L | 0.417 | P1_Lumbar_Coeliac_VS_P2_P3_P4_Stellate_Sphenopalatine | 1 | 0 | 0 | 0 | 1 | 1 | 0 | 0 |
| DNTTIP2 | 0.417 | P1_Lumbar_Coeliac_VS_P2_P3_P4_Stellate_Sphenopalatine | 1 | 0 | 0 | 0 | 1 | 1 | 0 | 0 |
| DKC1 | 0.417 | P3_Lumbar_Coeliac_VS_P1_P2_P4_Stellate_Sphenopalatine | 0 | 0 | 1 | 0 | 1 | 1 | 0 | 0 |
| ROCK2 | 0.417 | P4_Lumbar_Coeliac_VS_P1_P2_P3_Stellate_Sphenopalatine | 0 | 0 | 0 | 1 | 1 | 1 | 0 | 0 |
| FAM120C | 0.417 | P4_Lumbar_Coeliac_VS_P1_P2_P3_Stellate_Sphenopalatine | 0 | 0 | 0 | 1 | 1 | 1 | 0 | 0 |
| KMT5A | 0.417 | P1_P2_P4_Lumbar_Coeliac_VS_P3_Stellate_Sphenopalatine | 1 | 1 | 0 | 1 | 1 | 1 | 0 | 0 |
| AEBP2 | 0.417 | P1_P2_P4_Lumbar_Coeliac_Sphenopalatine_VS_P3_Stellate | 1 | 1 | 0 | 1 | 1 | 1 | 0 | 1 |
| BACH2 | 0.416 | P4_Lumbar_Coeliac_VS_P1_P2_P3_Stellate_Sphenopalatine | 0 | 0 | 0 | 1 | 1 | 1 | 0 | 0 |
| EFL1 | 0.416 | P4_Lumbar_Coeliac_VS_P1_P2_P3_Stellate_Sphenopalatine | 0 | 0 | 0 | 1 | 1 | 1 | 0 | 0 |
| DIAPH2 | 0.416 | P4_Lumbar_Coeliac_VS_P1_P2_P3_Stellate_Sphenopalatine | 0 | 0 | 0 | 1 | 1 | 1 | 0 | 0 |

| | | | | | | | | | | |
|---|---|---|---|---|---|---|---|---|---|---|
| GM42517 | 0.416 | P1_P2_P4_Lumbar_Sphenopalatine_VS_P3_Coeliac_Stellate | 1 | 1 | 0 | 1 | 1 | 0 | 0 | 1 |
| IL18 | 0.415 | P3_VS_P1_P2_P4_Lumbar_Coeliac_Stellate_Sphenopalatine | 0 | 0 | 1 | 0 | 0 | 0 | 0 | 0 |
| HSPB1 | 0.415 | P1_P4_VS_P2_P3_Lumbar_Coeliac_Stellate_Sphenopalatine | 1 | 0 | 0 | 1 | 0 | 0 | 0 | 0 |
| ARL15 | 0.415 | P1_Lumbar_Coeliac_VS_P2_P3_P4_Stellate_Sphenopalatine | 1 | 0 | 0 | 0 | 1 | 1 | 0 | 0 |
| IFT74 | 0.415 | P4_Lumbar_Coeliac_VS_P1_P2_P3_Stellate_Sphenopalatine | 0 | 0 | 0 | 1 | 1 | 1 | 0 | 0 |
| AHDC1 | 0.415 | P1_P3_P4_Lumbar_Coeliac_Sphenopalatine_VS_P2_Stellate | 1 | 0 | 1 | 1 | 1 | 1 | 0 | 1 |
| ZMYM1 | 0.414 | P4_Lumbar_Coeliac_VS_P1_P2_P3_Stellate_Sphenopalatine | 0 | 0 | 0 | 1 | 1 | 1 | 0 | 0 |
| SMC5 | 0.414 | P4_Lumbar_Coeliac_VS_P1_P2_P3_Stellate_Sphenopalatine | 0 | 0 | 0 | 1 | 1 | 1 | 0 | 0 |
| SMPD2 | 0.414 | P3_P4_Lumbar_Coeliac_VS_P1_P2_Stellate_Sphenopalatine | 0 | 0 | 1 | 1 | 1 | 1 | 0 | 0 |
| PEG10 | 0.414 | Sphenopalatine_VS_P1_P2_P3_P4_Lumbar_Coeliac_Stellate | 0 | 0 | 0 | 0 | 0 | 0 | 0 | 1 |
| NME4 | 0.414 | P1_P2_P3_Lumbar_Coeliac_Stellate_Sphenopalatine_VS_P4 | 1 | 1 | 1 | 0 | 1 | 1 | 1 | 1 |
| IL10RB | 0.413 | P2_P4_VS_P1_P3_Lumbar_Coeliac_Stellate_Sphenopalatine | 0 | 1 | 0 | 1 | 0 | 0 | 0 | 0 |
| ZFP442 | 0.413 | P4_Coeliac_VS_P1_P2_P3_Lumbar_Stellate_Sphenopalatine | 0 | 0 | 0 | 1 | 0 | 1 | 0 | 0 |
| CBFB | 0.413 | P1_P2_P3_P4_Coeliac_Sphenopalatine_VS_Lumbar_Stellate | 1 | 1 | 1 | 1 | 0 | 1 | 0 | 1 |
| RNF150 | 0.412 | P4_Lumbar_Coeliac_VS_P1_P2_P3_Stellate_Sphenopalatine | 0 | 0 | 0 | 1 | 1 | 1 | 0 | 0 |
| MSANTD2 | 0.412 | P4_Lumbar_Coeliac_VS_P1_P2_P3_Stellate_Sphenopalatine | 0 | 0 | 0 | 1 | 1 | 1 | 0 | 0 |
| PDE8A | 0.412 | P2_P4_Sphenopalatine_VS_P1_P3_Lumbar_Coeliac_Stellate | 0 | 1 | 0 | 1 | 0 | 0 | 0 | 1 |
| SYBU | 0.411 | P3_VS_P1_P2_P4_Lumbar_Coeliac_Stellate_Sphenopalatine | 0 | 0 | 1 | 0 | 0 | 0 | 0 | 0 |
| CRLF2 | 0.411 | P1_P2_P3_P4_Lumbar_Sphenopalatine_VS_Coeliac_Stellate | 1 | 1 | 1 | 1 | 1 | 0 | 0 | 1 |
| CAVIN3 | 0.411 | P1_P3_P4_Lumbar_Coeliac_Sphenopalatine_VS_P2_Stellate | 1 | 0 | 1 | 1 | 1 | 1 | 0 | 1 |
| HYAL2 | 0.41 | P4_Lumbar_Coeliac_VS_P1_P2_P3_Stellate_Sphenopalatine | 0 | 0 | 0 | 1 | 1 | 1 | 0 | 0 |
| TMEM151B | 0.41 | P1_P2_P3_Lumbar_Coeliac_Stellate_Sphenopalatine_VS_P4 | 1 | 1 | 1 | 0 | 1 | 1 | 1 | 1 |
| GM13205 | 0.409 | P3_VS_P1_P2_P4_Lumbar_Coeliac_Stellate_Sphenopalatine | 0 | 0 | 1 | 0 | 0 | 0 | 0 | 0 |
| DHX38 | 0.409 | P4_Lumbar_Coeliac_VS_P1_P2_P3_Stellate_Sphenopalatine | 0 | 0 | 0 | 1 | 1 | 1 | 0 | 0 |
| AATF | 0.409 | P4_Lumbar_Coeliac_VS_P1_P2_P3_Stellate_Sphenopalatine | 0 | 0 | 0 | 1 | 1 | 1 | 0 | 0 |
| OSBP | 0.409 | P1_P2_P4_Lumbar_Coeliac_Sphenopalatine_VS_P3_Stellate | 1 | 1 | 0 | 1 | 1 | 1 | 0 | 1 |
| GRIA3 | 0.408 | P1_P3_Lumbar_Coeliac_VS_P2_P4_Stellate_Sphenopalatine | 1 | 0 | 1 | 0 | 1 | 1 | 0 | 0 |
| BEND6 | 0.408 | P1_P2_P3_P4_Sphenopalatine_VS_Lumbar_Coeliac_Stellate | 1 | 1 | 1 | 1 | 0 | 0 | 0 | 1 |
| ME1 | 0.408 | P1_P2_P3_P4_Sphenopalatine_VS_Lumbar_Coeliac_Stellate | 1 | 1 | 1 | 1 | 0 | 0 | 0 | 1 |
| MSC | 0.407 | P1_VS_P2_P3_P4_Lumbar_Coeliac_Stellate_Sphenopalatine | 1 | 0 | 0 | 0 | 0 | 0 | 0 | 0 |

| | | | | | | | | | | |
|---|---|---|---|---|---|---|---|---|---|---|
| SAV1 | 0.407 | P1_Lumbar_Coeliac_VS_P2_P3_P4_Stellate_Sphenopalatine | 1 | 0 | 0 | 0 | 1 | 1 | 0 | 0 |
| ZFP652 | 0.407 | P2_Lumbar_Coeliac_VS_P1_P3_P4_Stellate_Sphenopalatine | 0 | 1 | 0 | 0 | 1 | 1 | 0 | 0 |
| B3GNT2 | 0.407 | P3_Lumbar_Coeliac_VS_P1_P2_P4_Stellate_Sphenopalatine | 0 | 0 | 1 | 0 | 1 | 1 | 0 | 0 |
| PCDHAC2 | 0.406 | P2_VS_P1_P3_P4_Lumbar_Coeliac_Stellate_Sphenopalatine | 0 | 1 | 0 | 0 | 0 | 0 | 0 | 0 |
| NHLH2 | 0.405 | P1_VS_P2_P3_P4_Lumbar_Coeliac_Stellate_Sphenopalatine | 1 | 0 | 0 | 0 | 0 | 0 | 0 | 0 |
| ARHGAP32 | 0.405 | P1_P2_P4_Lumbar_Coeliac_Sphenopalatine_VS_P3_Stellate | 1 | 1 | 0 | 1 | 1 | 1 | 0 | 1 |
| SASH1 | 0.404 | P2_P4_VS_P1_P3_Lumbar_Coeliac_Stellate_Sphenopalatine | 0 | 1 | 0 | 1 | 0 | 0 | 0 | 0 |
| SLC24A5 | 0.404 | P1_Lumbar_Coeliac_VS_P2_P3_P4_Stellate_Sphenopalatine | 1 | 0 | 0 | 0 | 1 | 1 | 0 | 0 |
| PSD2 | 0.404 | P3_Lumbar_Coeliac_VS_P1_P2_P4_Stellate_Sphenopalatine | 0 | 0 | 1 | 0 | 1 | 1 | 0 | 0 |
| SPRY4 | 0.404 | P1_P3_Lumbar_Coeliac_VS_P2_P4_Stellate_Sphenopalatine | 1 | 0 | 1 | 0 | 1 | 1 | 0 | 0 |
| RORA | 0.404 | P1_P2_P3_Lumbar_Coeliac_VS_P4_Stellate_Sphenopalatine | 1 | 1 | 1 | 0 | 1 | 1 | 0 | 0 |
| SLC31A1 | 0.404 | P3_Lumbar_Coeliac_Stellate_VS_P1_P2_P4_Sphenopalatine | 0 | 0 | 1 | 0 | 1 | 1 | 1 | 0 |
| GM10503 | 0.404 | Sphenopalatine_VS_P1_P2_P3_P4_Lumbar_Coeliac_Stellate | 0 | 0 | 0 | 0 | 0 | 0 | 0 | 1 |
| RNASE4 | 0.404 | P1_Lumbar_Coeliac_Sphenopalatine_VS_P2_P3_P4_Stellate | 1 | 0 | 0 | 0 | 1 | 1 | 0 | 1 |
| FAM160B1 | 0.404 | P1_P2_P4_Lumbar_Coeliac_Sphenopalatine_VS_P3_Stellate | 1 | 1 | 0 | 1 | 1 | 1 | 0 | 1 |
| SCG2 | 0.404 | P1_P2_P3_Lumbar_Coeliac_Stellate_Sphenopalatine_VS_P4 | 1 | 1 | 1 | 0 | 1 | 1 | 1 | 1 |
| PPWD1 | 0.403 | P3_P4_Lumbar_Coeliac_VS_P1_P2_Stellate_Sphenopalatine | 0 | 0 | 1 | 1 | 1 | 1 | 0 | 0 |
| SFT2D3 | 0.403 | P1_P3_P4_Lumbar_Coeliac_VS_P2_Stellate_Sphenopalatine | 1 | 0 | 1 | 1 | 1 | 1 | 0 | 0 |
| CCDC186 | 0.403 | P1_P2_P4_Lumbar_Coeliac_Sphenopalatine_VS_P3_Stellate | 1 | 1 | 0 | 1 | 1 | 1 | 0 | 1 |
| LGALS3 | 0.402 | P1_P3_Sphenopalatine_VS_P2_P4_Lumbar_Coeliac_Stellate | 1 | 0 | 1 | 0 | 0 | 0 | 0 | 1 |
| LITAF | 0.402 | P1_P3_P4_Sphenopalatine_VS_P2_Lumbar_Coeliac_Stellate | 1 | 0 | 1 | 1 | 0 | 0 | 0 | 1 |
| GM44644 | 0.402 | P4_Lumbar_Sphenopalatine_VS_P1_P2_P3_Coeliac_Stellate | 0 | 0 | 0 | 1 | 1 | 0 | 0 | 1 |
| SETD1A | 0.402 | P1_P2_P4_Lumbar_Coeliac_Sphenopalatine_VS_P3_Stellate | 1 | 1 | 0 | 1 | 1 | 1 | 0 | 1 |
| PTPN4 | 0.401 | P3_Lumbar_Coeliac_VS_P1_P2_P4_Stellate_Sphenopalatine | 0 | 0 | 1 | 0 | 1 | 1 | 0 | 0 |
| DLL3 | 0.401 | P1_P2_P3_P4_Lumbar_Sphenopalatine_VS_Coeliac_Stellate | 1 | 1 | 1 | 1 | 1 | 0 | 0 | 1 |
| DBR1 | 0.4 | P1_Lumbar_Coeliac_VS_P2_P3_P4_Stellate_Sphenopalatine | 1 | 0 | 0 | 0 | 1 | 1 | 0 | 0 |
| MOGS | 0.4 | P1_P4_Lumbar_Coeliac_VS_P2_P3_Stellate_Sphenopalatine | 1 | 0 | 0 | 1 | 1 | 1 | 0 | 0 |
| SERTAD2 | 0.399 | P1_P2_P4_Lumbar_Coeliac_Sphenopalatine_VS_P3_Stellate | 1 | 1 | 0 | 1 | 1 | 1 | 0 | 1 |
| LTBR | 0.398 | P1_P2_P3_P4_Lumbar_Sphenopalatine_VS_Coeliac_Stellate | 1 | 1 | 1 | 1 | 1 | 0 | 0 | 1 |
| TOPBP1 | 0.397 | P3_Lumbar_Coeliac_VS_P1_P2_P4_Stellate_Sphenopalatine | 0 | 0 | 1 | 0 | 1 | 1 | 0 | 0 |

| | | | | | | | | | | |
|---|---|---|---|---|---|---|---|---|---|---|
| MYBBP1A | 0.397 | P1_P4_Lumbar_Coeliac_VS_P2_P3_Stellate_Sphenopalatine | 1 | 0 | 0 | 1 | 1 | 1 | 0 | 0 |
| WDR53 | 0.397 | P1_P3_P4_Lumbar_Coeliac_VS_P2_Stellate_Sphenopalatine | 1 | 0 | 1 | 1 | 1 | 1 | 0 | 0 |
| TMCC3 | 0.397 | P1_P2_P4_Lumbar_Coeliac_Stellate_VS_P3_Sphenopalatine | 1 | 1 | 0 | 1 | 1 | 1 | 1 | 0 |
| GM28791 | 0.396 | P4_Coeliac_VS_P1_P2_P3_Lumbar_Stellate_Sphenopalatine | 0 | 0 | 0 | 1 | 0 | 1 | 0 | 0 |
| BTRC | 0.396 | P1_P2_P4_Lumbar_Coeliac_VS_P3_Stellate_Sphenopalatine | 1 | 1 | 0 | 1 | 1 | 1 | 0 | 0 |
| MAVS | 0.395 | P1_P4_VS_P2_P3_Lumbar_Coeliac_Stellate_Sphenopalatine | 1 | 0 | 0 | 1 | 0 | 0 | 0 | 0 |
| GM43305 | 0.395 | P3_P4_VS_P1_P2_Lumbar_Coeliac_Stellate_Sphenopalatine | 0 | 0 | 1 | 1 | 0 | 0 | 0 | 0 |
| ARHGAP17 | 0.395 | P1_P2_P4_Lumbar_Coeliac_VS_P3_Stellate_Sphenopalatine | 1 | 1 | 0 | 1 | 1 | 1 | 0 | 0 |
| ZFP945 | 0.395 | P1_P2_P4_Lumbar_Coeliac_VS_P3_Stellate_Sphenopalatine | 1 | 1 | 0 | 1 | 1 | 1 | 0 | 0 |
| SDCCAG8 | 0.395 | P3_P4_Lumbar_Coeliac_VS_P1_P2_Stellate_Sphenopalatine | 0 | 0 | 1 | 1 | 1 | 1 | 0 | 0 |
| USF3 | 0.395 | P1_P3_P4_Lumbar_Coeliac_VS_P2_Stellate_Sphenopalatine | 1 | 0 | 1 | 1 | 1 | 1 | 0 | 0 |
| SORD | 0.395 | P1_P2_P3_P4_Lumbar_Coeliac_Stellate_VS_Sphenopalatine | 1 | 1 | 1 | 1 | 1 | 1 | 1 | 0 |
| UBR2 | 0.395 | P1_P2_P4_Lumbar_Coeliac_Sphenopalatine_VS_P3_Stellate | 1 | 1 | 0 | 1 | 1 | 1 | 0 | 1 |
| RGL1 | 0.394 | P2_VS_P1_P3_P4_Lumbar_Coeliac_Stellate_Sphenopalatine | 0 | 1 | 0 | 0 | 0 | 0 | 0 | 0 |
| KLHDC4 | 0.394 | P3_Lumbar_Coeliac_VS_P1_P2_P4_Stellate_Sphenopalatine | 0 | 0 | 1 | 0 | 1 | 1 | 0 | 0 |
| ARID3B | 0.394 | P2_P4_Lumbar_Coeliac_VS_P1_P3_Stellate_Sphenopalatine | 0 | 1 | 0 | 1 | 1 | 1 | 0 | 0 |
| TENT4B | 0.394 | P1_P2_P4_Lumbar_Coeliac_VS_P3_Stellate_Sphenopalatine | 1 | 1 | 0 | 1 | 1 | 1 | 0 | 0 |
| SHMT2 | 0.393 | P4_Lumbar_VS_P1_P2_P3_Coeliac_Stellate_Sphenopalatine | 0 | 0 | 0 | 1 | 1 | 0 | 0 | 0 |
| RNF113A1 | 0.393 | P4_Lumbar_VS_P1_P2_P3_Coeliac_Stellate_Sphenopalatine | 0 | 0 | 0 | 1 | 1 | 0 | 0 | 0 |
| LIN28B | 0.393 | P4_Coeliac_VS_P1_P2_P3_Lumbar_Stellate_Sphenopalatine | 0 | 0 | 0 | 1 | 0 | 1 | 0 | 0 |
| SEMA4B | 0.393 | P3_Lumbar_Coeliac_VS_P1_P2_P4_Stellate_Sphenopalatine | 0 | 0 | 1 | 0 | 1 | 1 | 0 | 0 |
| MAP3K13 | 0.393 | P3_Lumbar_Coeliac_VS_P1_P2_P4_Stellate_Sphenopalatine | 0 | 0 | 1 | 0 | 1 | 1 | 0 | 0 |
| GM9916 | 0.393 | P1_P3_P4_Lumbar_Coeliac_VS_P2_Stellate_Sphenopalatine | 1 | 0 | 1 | 1 | 1 | 1 | 0 | 0 |
| CTLA2A | 0.393 | Sphenopalatine_VS_P1_P2_P3_P4_Lumbar_Coeliac_Stellate | 0 | 0 | 0 | 0 | 0 | 0 | 0 | 1 |
| ADGRB2 | 0.392 | P2_P4_VS_P1_P3_Lumbar_Coeliac_Stellate_Sphenopalatine | 0 | 1 | 0 | 1 | 0 | 0 | 0 | 0 |
| NUBPL | 0.392 | P4_Lumbar_VS_P1_P2_P3_Coeliac_Stellate_Sphenopalatine | 0 | 0 | 0 | 1 | 1 | 0 | 0 | 0 |
| EIF2AK3 | 0.392 | P1_P2_P4_Lumbar_Coeliac_VS_P3_Stellate_Sphenopalatine | 1 | 1 | 0 | 1 | 1 | 1 | 0 | 0 |
| SNX18 | 0.392 | P1_P2_P3_P4_Lumbar_Coeliac_Stellate_VS_Sphenopalatine | 1 | 1 | 1 | 1 | 1 | 1 | 1 | 0 |
| BTG3 | 0.392 | P1_P2_P3_Lumbar_Coeliac_Sphenopalatine_VS_P4_Stellate | 1 | 1 | 1 | 0 | 1 | 1 | 0 | 1 |
| IMPDH1 | 0.392 | P1_P2_P4_Lumbar_Coeliac_Sphenopalatine_VS_P3_Stellate | 1 | 1 | 0 | 1 | 1 | 1 | 0 | 1 |

| | | | | | | | | | | |
|---|---|---|---|---|---|---|---|---|---|---|
| NUP133 | 0.39 | P1_P2_P4_Lumbar_Coeliac_VS_P3_Stellate_Sphenopalatine | 1 | 1 | 0 | 1 | 1 | 1 | 0 | 0 |
| PHF6 | 0.39 | P1_P2_P4_Lumbar_Coeliac_VS_P3_Stellate_Sphenopalatine | 1 | 1 | 0 | 1 | 1 | 1 | 0 | 0 |
| MARCH11 | 0.39 | P1_P3_P4_Lumbar_Coeliac_Stellate_VS_P2_Sphenopalatine | 1 | 0 | 1 | 1 | 1 | 1 | 1 | 0 |
| LRP3 | 0.39 | P1_P2_P4_Lumbar_Coeliac_Sphenopalatine_VS_P3_Stellate | 1 | 1 | 0 | 1 | 1 | 1 | 0 | 1 |
| MDK | 0.389 | P2_P4_VS_P1_P3_Lumbar_Coeliac_Stellate_Sphenopalatine | 0 | 1 | 0 | 1 | 0 | 0 | 0 | 0 |
| IKZF5 | 0.389 | P3_Lumbar_Coeliac_VS_P1_P2_P4_Stellate_Sphenopalatine | 0 | 0 | 1 | 0 | 1 | 1 | 0 | 0 |
| FASTKD2 | 0.389 | P1_P4_Lumbar_Coeliac_VS_P2_P3_Stellate_Sphenopalatine | 1 | 0 | 0 | 1 | 1 | 1 | 0 | 0 |
| RNF130 | 0.389 | P1_P2_P4_Lumbar_Coeliac_VS_P3_Stellate_Sphenopalatine | 1 | 1 | 0 | 1 | 1 | 1 | 0 | 0 |
| XAB2 | 0.389 | P1_P3_P4_Lumbar_Coeliac_VS_P2_Stellate_Sphenopalatine | 1 | 0 | 1 | 1 | 1 | 1 | 0 | 0 |
| IGF2R | 0.389 | P2_P3_P4_Lumbar_Coeliac_VS_P1_Stellate_Sphenopalatine | 0 | 1 | 1 | 1 | 1 | 1 | 0 | 0 |
| FAAH | 0.389 | P1_P3_P4_Lumbar_Coeliac_Stellate_Sphenopalatine_VS_P2 | 1 | 0 | 1 | 1 | 1 | 1 | 1 | 1 |
| NUP205 | 0.388 | P2_P4_Lumbar_Coeliac_VS_P1_P3_Stellate_Sphenopalatine | 0 | 1 | 0 | 1 | 1 | 1 | 0 | 0 |
| ELMSAN1 | 0.388 | P1_P2_P4_Lumbar_Coeliac_VS_P3_Stellate_Sphenopalatine | 1 | 1 | 0 | 1 | 1 | 1 | 0 | 0 |
| SLC35B3 | 0.388 | Lumbar_Coeliac_Stellate_VS_P1_P2_P3_P4_Sphenopalatine | 0 | 0 | 0 | 0 | 1 | 1 | 1 | 0 |
| AP1S2 | 0.388 | P1_P2_P3_Lumbar_Coeliac_Stellate_Sphenopalatine_VS_P4 | 1 | 1 | 1 | 0 | 1 | 1 | 1 | 1 |
| MOSPD1 | 0.387 | P1_VS_P2_P3_P4_Lumbar_Coeliac_Stellate_Sphenopalatine | 1 | 0 | 0 | 0 | 0 | 0 | 0 | 0 |
| SLC25A15 | 0.387 | P2_VS_P1_P3_P4_Lumbar_Coeliac_Stellate_Sphenopalatine | 0 | 1 | 0 | 0 | 0 | 0 | 0 | 0 |
| HSPA14 | 0.387 | P1_P4_Lumbar_Coeliac_VS_P2_P3_Stellate_Sphenopalatine | 1 | 0 | 0 | 1 | 1 | 1 | 0 | 0 |
| MID1 | 0.387 | P1_P2_Sphenopalatine_VS_P3_P4_Lumbar_Coeliac_Stellate | 1 | 1 | 0 | 0 | 0 | 0 | 0 | 1 |
| MRGPRE | 0.386 | P2_Lumbar_Coeliac_VS_P1_P3_P4_Stellate_Sphenopalatine | 0 | 1 | 0 | 0 | 1 | 1 | 0 | 0 |
| NUFIP1 | 0.386 | P3_Lumbar_Coeliac_VS_P1_P2_P4_Stellate_Sphenopalatine | 0 | 0 | 1 | 0 | 1 | 1 | 0 | 0 |
| FTX | 0.386 | P1_P2_P3_Lumbar_Coeliac_Sphenopalatine_VS_P4_Stellate | 1 | 1 | 1 | 0 | 1 | 1 | 0 | 1 |
| TRPM7 | 0.386 | P1_P2_P4_Lumbar_Coeliac_Sphenopalatine_VS_P3_Stellate | 1 | 1 | 0 | 1 | 1 | 1 | 0 | 1 |
| GM4419 | 0.385 | P3_VS_P1_P2_P4_Lumbar_Coeliac_Stellate_Sphenopalatine | 0 | 0 | 1 | 0 | 0 | 0 | 0 | 0 |
| SELENON | 0.385 | P1_P2_P4_Lumbar_Coeliac_Sphenopalatine_VS_P3_Stellate | 1 | 1 | 0 | 1 | 1 | 1 | 0 | 1 |
| LMTK2 | 0.385 | P1_P2_P4_Lumbar_Coeliac_Sphenopalatine_VS_P3_Stellate | 1 | 1 | 0 | 1 | 1 | 1 | 0 | 1 |
| GM16141 | 0.384 | P3_Coeliac_VS_P1_P2_P4_Lumbar_Stellate_Sphenopalatine | 0 | 0 | 1 | 0 | 0 | 1 | 0 | 0 |
| PMM2 | 0.384 | P3_Lumbar_Coeliac_VS_P1_P2_P4_Stellate_Sphenopalatine | 0 | 0 | 1 | 0 | 1 | 1 | 0 | 0 |
| CERS6 | 0.384 | P1_P3_P4_Lumbar_Coeliac_Sphenopalatine_VS_P2_Stellate | 1 | 0 | 1 | 1 | 1 | 1 | 0 | 1 |
| CIT | 0.383 | P2_P3_P4_Lumbar_Coeliac_VS_P1_Stellate_Sphenopalatine | 0 | 1 | 1 | 1 | 1 | 1 | 0 | 0 |

| | | | | | | | | | | |
|---|---|---|---|---|---|---|---|---|---|---|
| GSTM7 | 0.382 | P3_VS_P1_P2_P4_Lumbar_Coeliac_Stellate_Sphenopalatine | 0 | 0 | 1 | 0 | 0 | 0 | 0 | 0 |
| RFTN1 | 0.381 | P1_P3_VS_P2_P4_Lumbar_Coeliac_Stellate_Sphenopalatine | 1 | 0 | 1 | 0 | 0 | 0 | 0 | 0 |
| TBC1D15 | 0.381 | P1_Lumbar_Coeliac_VS_P2_P3_P4_Stellate_Sphenopalatine | 1 | 0 | 0 | 0 | 1 | 1 | 0 | 0 |
| PHC3 | 0.381 | P1_P2_P4_Lumbar_Coeliac_VS_P3_Stellate_Sphenopalatine | 1 | 1 | 0 | 1 | 1 | 1 | 0 | 0 |
| PLRG1 | 0.381 | P1_P3_P4_Lumbar_Coeliac_VS_P2_Stellate_Sphenopalatine | 1 | 0 | 1 | 1 | 1 | 1 | 0 | 0 |
| MAP4K5 | 0.381 | P1_P2_P4_Lumbar_Coeliac_Sphenopalatine_VS_P3_Stellate | 1 | 1 | 0 | 1 | 1 | 1 | 0 | 1 |
| GRIP2 | 0.38 | P2_VS_P1_P3_P4_Lumbar_Coeliac_Stellate_Sphenopalatine | 0 | 1 | 0 | 0 | 0 | 0 | 0 | 0 |
| AP4E1 | 0.38 | P1_P2_P4_Lumbar_Coeliac_VS_P3_Stellate_Sphenopalatine | 1 | 1 | 0 | 1 | 1 | 1 | 0 | 0 |
| TUBGCP5 | 0.38 | P1_P2_P4_Lumbar_Coeliac_VS_P3_Stellate_Sphenopalatine | 1 | 1 | 0 | 1 | 1 | 1 | 0 | 0 |
| ATF7 | 0.38 | P1_P2_P3_Lumbar_Coeliac_Sphenopalatine_VS_P4_Stellate | 1 | 1 | 1 | 0 | 1 | 1 | 0 | 1 |
| CREBL2 | 0.379 | P2_P4_VS_P1_P3_Lumbar_Coeliac_Stellate_Sphenopalatine | 0 | 1 | 0 | 1 | 0 | 0 | 0 | 0 |
| MCM3AP | 0.379 | P2_Lumbar_Coeliac_VS_P1_P3_P4_Stellate_Sphenopalatine | 0 | 1 | 0 | 0 | 1 | 1 | 0 | 0 |
| MED13L | 0.379 | P1_P2_P4_Lumbar_Coeliac_Sphenopalatine_VS_P3_Stellate | 1 | 1 | 0 | 1 | 1 | 1 | 0 | 1 |
| GM26782 | 0.379 | P1_P2_P3_P4_Stellate_Sphenopalatine_VS_Lumbar_Coeliac | 1 | 1 | 1 | 1 | 0 | 0 | 1 | 1 |
| 1110006O24 | 0.378 | P3_VS_P1_P2_P4_Lumbar_Coeliac_Stellate_Sphenopalatine | 0 | 0 | 1 | 0 | 0 | 0 | 0 | 0 |
| NOVA2 | 0.378 | P2_Lumbar_Coeliac_VS_P1_P3_P4_Stellate_Sphenopalatine | 0 | 1 | 0 | 0 | 1 | 1 | 0 | 0 |
| WNT3 | 0.378 | P3_Lumbar_Coeliac_VS_P1_P2_P4_Stellate_Sphenopalatine | 0 | 0 | 1 | 0 | 1 | 1 | 0 | 0 |
| EIF2AK4 | 0.378 | P2_P3_P4_Lumbar_Coeliac_VS_P1_Stellate_Sphenopalatine | 0 | 1 | 1 | 1 | 1 | 1 | 0 | 0 |
| NFATC3 | 0.377 | P2_P3_P4_Lumbar_Coeliac_VS_P1_Stellate_Sphenopalatine | 0 | 1 | 1 | 1 | 1 | 1 | 0 | 0 |
| OSBPL3 | 0.377 | P1_P2_P3_P4_Coeliac_Sphenopalatine_VS_Lumbar_Stellate | 1 | 1 | 1 | 1 | 0 | 1 | 0 | 1 |
| RPAP3 | 0.377 | P4_Lumbar_Coeliac_Sphenopalatine_VS_P1_P2_P3_Stellate | 0 | 0 | 0 | 1 | 1 | 1 | 0 | 1 |
| TESK1 | 0.377 | P1_P2_P4_Lumbar_Coeliac_Sphenopalatine_VS_P3_Stellate | 1 | 1 | 0 | 1 | 1 | 1 | 0 | 1 |
| FNDC3B | 0.376 | P3_P4_VS_P1_P2_Lumbar_Coeliac_Stellate_Sphenopalatine | 0 | 0 | 1 | 1 | 0 | 0 | 0 | 0 |
| USP35 | 0.376 | P3_Lumbar_Coeliac_VS_P1_P2_P4_Stellate_Sphenopalatine | 0 | 0 | 1 | 0 | 1 | 1 | 0 | 0 |
| AGA | 0.376 | P3_Lumbar_Coeliac_VS_P1_P2_P4_Stellate_Sphenopalatine | 0 | 0 | 1 | 0 | 1 | 1 | 0 | 0 |
| PCMTD2 | 0.376 | P1_P2_P4_Lumbar_Coeliac_VS_P3_Stellate_Sphenopalatine | 1 | 1 | 0 | 1 | 1 | 1 | 0 | 0 |
| PPM1F | 0.376 | P1_P2_P3_P4_Lumbar_Sphenopalatine_VS_Coeliac_Stellate | 1 | 1 | 1 | 1 | 1 | 0 | 0 | 1 |
| PLEKHA5 | 0.376 | P1_P2_P3_Lumbar_Coeliac_Sphenopalatine_VS_P4_Stellate | 1 | 1 | 1 | 0 | 1 | 1 | 0 | 1 |
| ZFP651 | 0.376 | P1_P2_P4_Lumbar_Coeliac_Sphenopalatine_VS_P3_Stellate | 1 | 1 | 0 | 1 | 1 | 1 | 0 | 1 |
| DNAJC27 | 0.375 | P2_P4_VS_P1_P3_Lumbar_Coeliac_Stellate_Sphenopalatine | 0 | 1 | 0 | 1 | 0 | 0 | 0 | 0 |

| | | | | | | | | | | |
|---|---|---|---|---|---|---|---|---|---|---|
| GM12371 | 0.375 | P3_P4_VS_P1_P2_Lumbar_Coeliac_Stellate_Sphenopalatine | 0 | 0 | 1 | 1 | 0 | 0 | 0 | 0 |
| FTSJ3 | 0.375 | P1_Lumbar_Coeliac_VS_P2_P3_P4_Stellate_Sphenopalatine | 1 | 0 | 0 | 0 | 1 | 1 | 0 | 0 |
| AXIN1 | 0.375 | P2_Lumbar_Coeliac_VS_P1_P3_P4_Stellate_Sphenopalatine | 0 | 1 | 0 | 0 | 1 | 1 | 0 | 0 |
| SPG11 | 0.375 | P3_Lumbar_Coeliac_VS_P1_P2_P4_Stellate_Sphenopalatine | 0 | 0 | 1 | 0 | 1 | 1 | 0 | 0 |
| NUMB | 0.375 | P1_P2_P4_Lumbar_Coeliac_VS_P3_Stellate_Sphenopalatine | 1 | 1 | 0 | 1 | 1 | 1 | 0 | 0 |
| PECR | 0.375 | P1_P3_P4_Lumbar_Coeliac_VS_P2_Stellate_Sphenopalatine | 1 | 0 | 1 | 1 | 1 | 1 | 0 | 0 |
| WIPF2 | 0.375 | P1_P2_P4_Lumbar_Coeliac_Sphenopalatine_VS_P3_Stellate | 1 | 1 | 0 | 1 | 1 | 1 | 0 | 1 |
| RBM27 | 0.375 | P1_P2_P4_Lumbar_Coeliac_Sphenopalatine_VS_P3_Stellate | 1 | 1 | 0 | 1 | 1 | 1 | 0 | 1 |
| MICAL1 | 0.374 | P2_Lumbar_Coeliac_VS_P1_P3_P4_Stellate_Sphenopalatine | 0 | 1 | 0 | 0 | 1 | 1 | 0 | 0 |
| ZFP760 | 0.374 | P1_P2_P4_Lumbar_Coeliac_VS_P3_Stellate_Sphenopalatine | 1 | 1 | 0 | 1 | 1 | 1 | 0 | 0 |
| NCAN | 0.374 | P1_P2_P3_Lumbar_Coeliac_Stellate_Sphenopalatine_VS_P4 | 1 | 1 | 1 | 0 | 1 | 1 | 1 | 1 |
| GPR176 | 0.373 | P3_VS_P1_P2_P4_Lumbar_Coeliac_Stellate_Sphenopalatine | 0 | 0 | 1 | 0 | 0 | 0 | 0 | 0 |
| OPRD1 | 0.373 | P3_VS_P1_P2_P4_Lumbar_Coeliac_Stellate_Sphenopalatine | 0 | 0 | 1 | 0 | 0 | 0 | 0 | 0 |
| CRYAB | 0.373 | P2_P4_VS_P1_P3_Lumbar_Coeliac_Stellate_Sphenopalatine | 0 | 1 | 0 | 1 | 0 | 0 | 0 | 0 |
| BRMS1 | 0.373 | P3_Lumbar_Coeliac_VS_P1_P2_P4_Stellate_Sphenopalatine | 0 | 0 | 1 | 0 | 1 | 1 | 0 | 0 |
| DDX10 | 0.373 | P1_P4_Lumbar_Coeliac_VS_P2_P3_Stellate_Sphenopalatine | 1 | 0 | 0 | 1 | 1 | 1 | 0 | 0 |
| TATDN3 | 0.373 | P1_P3_P4_Lumbar_Coeliac_VS_P2_Stellate_Sphenopalatine | 1 | 0 | 1 | 1 | 1 | 1 | 0 | 0 |
| LIMS2 | 0.373 | Lumbar_Coeliac_Stellate_VS_P1_P2_P3_P4_Sphenopalatine | 0 | 0 | 0 | 0 | 1 | 1 | 1 | 0 |
| AP4M1 | 0.373 | P1_P2_P3_Lumbar_Coeliac_Sphenopalatine_VS_P4_Stellate | 1 | 1 | 1 | 0 | 1 | 1 | 0 | 1 |
| LNX1 | 0.373 | P2_P3_P4_Lumbar_Coeliac_Sphenopalatine_VS_P1_Stellate | 0 | 1 | 1 | 1 | 1 | 1 | 0 | 1 |
| LONP2 | 0.372 | P1_P3_P4_Lumbar_Coeliac_VS_P2_Stellate_Sphenopalatine | 1 | 0 | 1 | 1 | 1 | 1 | 0 | 0 |
| NEURL1A | 0.372 | P1_P2_P3_P4_Coeliac_Sphenopalatine_VS_Lumbar_Stellate | 1 | 1 | 1 | 1 | 0 | 1 | 0 | 1 |
| D930028M1[illegible] | 0.372 | P1_P3_P4_Lumbar_Coeliac_Sphenopalatine_VS_P2_Stellate | 1 | 0 | 1 | 1 | 1 | 1 | 0 | 1 |
| CNN3 | 0.372 | P1_P2_P4_Lumbar_Coeliac_Stellate_Sphenopalatine_VS_P3 | 1 | 1 | 0 | 1 | 1 | 1 | 1 | 1 |
| SHISA6 | 0.371 | P1_VS_P2_P3_P4_Lumbar_Coeliac_Stellate_Sphenopalatine | 1 | 0 | 0 | 0 | 0 | 0 | 0 | 0 |
| DUSP2 | 0.37 | P1_VS_P2_P3_P4_Lumbar_Coeliac_Stellate_Sphenopalatine | 1 | 0 | 0 | 0 | 0 | 0 | 0 | 0 |
| TACC3 | 0.37 | P1_P4_VS_P2_P3_Lumbar_Coeliac_Stellate_Sphenopalatine | 1 | 0 | 0 | 1 | 0 | 0 | 0 | 0 |
| AMOT | 0.37 | P3_P4_VS_P1_P2_Lumbar_Coeliac_Stellate_Sphenopalatine | 0 | 0 | 1 | 1 | 0 | 0 | 0 | 0 |
| POGLUT3 | 0.37 | P1_P2_P4_Lumbar_Coeliac_VS_P3_Stellate_Sphenopalatine | 1 | 1 | 0 | 1 | 1 | 1 | 0 | 0 |
| FCHO2 | 0.37 | P1_P2_P4_Lumbar_Coeliac_VS_P3_Stellate_Sphenopalatine | 1 | 1 | 0 | 1 | 1 | 1 | 0 | 0 |

| | | | | | | | | | | |
|---|---|---|---|---|---|---|---|---|---|---|
| MTCL1 | 0.37 | P1_P2_P3_P4_Lumbar_Coeliac_Stellate_VS_Sphenopalatine | 1 | 1 | 1 | 1 | 1 | 1 | 1 | 0 |
| KDM5A | 0.37 | P1_P2_P4_Lumbar_Coeliac_Sphenopalatine_VS_P3_Stellate | 1 | 1 | 0 | 1 | 1 | 1 | 0 | 1 |
| AKAP13 | 0.37 | P1_P2_P4_Lumbar_Coeliac_Stellate_Sphenopalatine_VS_P3 | 1 | 1 | 0 | 1 | 1 | 1 | 1 | 1 |
| EBF4 | 0.369 | P2_Lumbar_Coeliac_VS_P1_P3_P4_Stellate_Sphenopalatine | 0 | 1 | 0 | 0 | 1 | 1 | 0 | 0 |
| AGGF1 | 0.369 | P2_Lumbar_Coeliac_VS_P1_P3_P4_Stellate_Sphenopalatine | 0 | 1 | 0 | 0 | 1 | 1 | 0 | 0 |
| COQ10A | 0.369 | P1_P3_P4_Lumbar_Coeliac_VS_P2_Stellate_Sphenopalatine | 1 | 0 | 1 | 1 | 1 | 1 | 0 | 0 |
| RPGR | 0.369 | P1_P3_P4_Lumbar_Coeliac_VS_P2_Stellate_Sphenopalatine | 1 | 0 | 1 | 1 | 1 | 1 | 0 | 0 |
| CSDC2 | 0.369 | P1_P2_P3_Lumbar_Coeliac_Stellate_Sphenopalatine_VS_P4 | 1 | 1 | 1 | 0 | 1 | 1 | 1 | 1 |
| PPP1R16B | 0.369 | P1_P2_P4_Lumbar_Coeliac_Stellate_Sphenopalatine_VS_P3 | 1 | 1 | 0 | 1 | 1 | 1 | 1 | 1 |
| USP27X | 0.368 | P3_VS_P1_P2_P4_Lumbar_Coeliac_Stellate_Sphenopalatine | 0 | 0 | 1 | 0 | 0 | 0 | 0 | 0 |
| NFKB1 | 0.368 | P4_Lumbar_VS_P1_P2_P3_Coeliac_Stellate_Sphenopalatine | 0 | 0 | 0 | 1 | 1 | 0 | 0 | 0 |
| HMGCLL1 | 0.368 | P3_Lumbar_Coeliac_VS_P1_P2_P4_Stellate_Sphenopalatine | 0 | 0 | 1 | 0 | 1 | 1 | 0 | 0 |
| GINS4 | 0.368 | P1_P3_P4_Lumbar_Coeliac_VS_P2_Stellate_Sphenopalatine | 1 | 0 | 1 | 1 | 1 | 1 | 0 | 0 |
| BOK | 0.368 | Sphenopalatine_VS_P1_P2_P3_P4_Lumbar_Coeliac_Stellate | 0 | 0 | 0 | 0 | 0 | 0 | 0 | 1 |
| TEX2 | 0.368 | P1_P2_P3_P4_Lumbar_Sphenopalatine_VS_Coeliac_Stellate | 1 | 1 | 1 | 1 | 1 | 0 | 0 | 1 |
| ZFP407 | 0.367 | P1_Lumbar_Coeliac_VS_P2_P3_P4_Stellate_Sphenopalatine | 1 | 0 | 0 | 0 | 1 | 1 | 0 | 0 |
| TRIP11 | 0.367 | P2_P4_Lumbar_Coeliac_VS_P1_P3_Stellate_Sphenopalatine | 0 | 1 | 0 | 1 | 1 | 1 | 0 | 0 |
| ZYX | 0.367 | P1_P2_P3_P4_Lumbar_Sphenopalatine_VS_Coeliac_Stellate | 1 | 1 | 1 | 1 | 1 | 0 | 0 | 1 |
| FNDC3A | 0.367 | P1_P2_P4_Lumbar_Coeliac_Sphenopalatine_VS_P3_Stellate | 1 | 1 | 0 | 1 | 1 | 1 | 0 | 1 |
| PGF | 0.367 | P1_P2_P3_Lumbar_Coeliac_Stellate_Sphenopalatine_VS_P4 | 1 | 1 | 1 | 0 | 1 | 1 | 1 | 1 |
| EFNB1 | 0.366 | P3_P4_Lumbar_Coeliac_VS_P1_P2_Stellate_Sphenopalatine | 0 | 0 | 1 | 1 | 1 | 1 | 0 | 0 |
| GALNT17 | 0.366 | P1_P2_Lumbar_Coeliac_Sphenopalatine_VS_P3_P4_Stellate | 1 | 1 | 0 | 0 | 1 | 1 | 0 | 1 |
| ABCC4 | 0.365 | P1_P2_P3_Lumbar_Coeliac_Stellate_VS_P4_Sphenopalatine | 1 | 1 | 1 | 0 | 1 | 1 | 1 | 0 |
| METTL14 | 0.365 | P1_P2_P4_Lumbar_Coeliac_Sphenopalatine_VS_P3_Stellate | 1 | 1 | 0 | 1 | 1 | 1 | 0 | 1 |
| SUSD6 | 0.365 | P1_P2_P4_Lumbar_Coeliac_Sphenopalatine_VS_P3_Stellate | 1 | 1 | 0 | 1 | 1 | 1 | 0 | 1 |
| HS6ST1 | 0.365 | P1_P3_P4_Lumbar_Coeliac_Sphenopalatine_VS_P2_Stellate | 1 | 0 | 1 | 1 | 1 | 1 | 0 | 1 |
| PCGF5 | 0.364 | P1_P2_P4_VS_P3_Lumbar_Coeliac_Stellate_Sphenopalatine | 1 | 1 | 0 | 1 | 0 | 0 | 0 | 0 |
| USP28 | 0.364 | P1_P2_Lumbar_Coeliac_VS_P3_P4_Stellate_Sphenopalatine | 1 | 1 | 0 | 0 | 1 | 1 | 0 | 0 |
| RCOR1 | 0.364 | P1_P2_P4_Lumbar_Coeliac_VS_P3_Stellate_Sphenopalatine | 1 | 1 | 0 | 1 | 1 | 1 | 0 | 0 |
| SPPL2A | 0.364 | P1_P2_P4_Lumbar_Coeliac_Sphenopalatine_VS_P3_Stellate | 1 | 1 | 0 | 1 | 1 | 1 | 0 | 1 |

| | | | | | | | | | | |
|---|---|---|---|---|---|---|---|---|---|---|
| MAN2A1 | 0.364 | P1_P3_P4_Lumbar_Coeliac_Stellate_Sphenopalatine_VS_P2 | 1 | 0 | 1 | 1 | 1 | 1 | 1 | 1 |
| AV356131 | 0.363 | P3_P4_Lumbar_Coeliac_VS_P1_P2_Stellate_Sphenopalatine | 0 | 0 | 1 | 1 | 1 | 1 | 0 | 0 |
| CTDSP2 | 0.363 | P1_P2_P4_Lumbar_Coeliac_Sphenopalatine_VS_P3_Stellate | 1 | 1 | 0 | 1 | 1 | 1 | 0 | 1 |
| GM32618 | 0.362 | P3_P4_VS_P1_P2_Lumbar_Coeliac_Stellate_Sphenopalatine | 0 | 0 | 1 | 1 | 0 | 0 | 0 | 0 |
| NFRKB | 0.362 | P2_Lumbar_Coeliac_VS_P1_P3_P4_Stellate_Sphenopalatine | 0 | 1 | 0 | 0 | 1 | 1 | 0 | 0 |
| 4930503L19F | 0.362 | P3_Lumbar_Coeliac_VS_P1_P2_P4_Stellate_Sphenopalatine | 0 | 0 | 1 | 0 | 1 | 1 | 0 | 0 |
| RNASEL | 0.362 | P1_P2_P4_Lumbar_Coeliac_VS_P3_Stellate_Sphenopalatine | 1 | 1 | 0 | 1 | 1 | 1 | 0 | 0 |
| MAFF | 0.362 | P1_P2_P3_P4_Lumbar_Coeliac_Stellate_VS_Sphenopalatine | 1 | 1 | 1 | 1 | 1 | 1 | 1 | 0 |
| PNPLA6 | 0.362 | P1_P2_P4_Lumbar_Coeliac_Sphenopalatine_VS_P3_Stellate | 1 | 1 | 0 | 1 | 1 | 1 | 0 | 1 |
| GTF2F1 | 0.362 | P1_P2_P4_Lumbar_Coeliac_Sphenopalatine_VS_P3_Stellate | 1 | 1 | 0 | 1 | 1 | 1 | 0 | 1 |
| FMNL2 | 0.361 | P2_Lumbar_Coeliac_VS_P1_P3_P4_Stellate_Sphenopalatine | 0 | 1 | 0 | 0 | 1 | 1 | 0 | 0 |
| MMP15 | 0.361 | P1_P2_P3_P4_Lumbar_Coeliac_Stellate_VS_Sphenopalatine | 1 | 1 | 1 | 1 | 1 | 1 | 1 | 0 |
| NUAK1 | 0.361 | P1_P2_P4_Lumbar_Coeliac_Stellate_Sphenopalatine_VS_P3 | 1 | 1 | 0 | 1 | 1 | 1 | 1 | 1 |
| GM45060 | 0.36 | P3_Coeliac_VS_P1_P2_P4_Lumbar_Stellate_Sphenopalatine | 0 | 0 | 1 | 0 | 0 | 1 | 0 | 0 |
| ZFP827 | 0.36 | P2_P3_P4_Lumbar_Coeliac_VS_P1_Stellate_Sphenopalatine | 0 | 1 | 1 | 1 | 1 | 1 | 0 | 0 |
| PDZD2 | 0.359 | P3_VS_P1_P2_P4_Lumbar_Coeliac_Stellate_Sphenopalatine | 0 | 0 | 1 | 0 | 0 | 0 | 0 | 0 |
| AP3B1 | 0.359 | P4_Lumbar_VS_P1_P2_P3_Coeliac_Stellate_Sphenopalatine | 0 | 0 | 0 | 1 | 1 | 0 | 0 | 0 |
| CCSER1 | 0.359 | P3_Lumbar_Coeliac_VS_P1_P2_P4_Stellate_Sphenopalatine | 0 | 0 | 1 | 0 | 1 | 1 | 0 | 0 |
| DPF1 | 0.359 | P1_P2_P3_Lumbar_Coeliac_Sphenopalatine_VS_P4_Stellate | 1 | 1 | 1 | 0 | 1 | 1 | 0 | 1 |
| CDC14B | 0.358 | P2_VS_P1_P3_P4_Lumbar_Coeliac_Stellate_Sphenopalatine | 0 | 1 | 0 | 0 | 0 | 0 | 0 | 0 |
| SLC12A7 | 0.358 | P2_VS_P1_P3_P4_Lumbar_Coeliac_Stellate_Sphenopalatine | 0 | 1 | 0 | 0 | 0 | 0 | 0 | 0 |
| TTR | 0.358 | P3_P4_Coeliac_VS_P1_P2_Lumbar_Stellate_Sphenopalatine | 0 | 0 | 1 | 1 | 0 | 1 | 0 | 0 |
| SHPRH | 0.358 | P2_P4_Lumbar_Coeliac_VS_P1_P3_Stellate_Sphenopalatine | 0 | 1 | 0 | 1 | 1 | 1 | 0 | 0 |
| LRP12 | 0.358 | P2_P4_Lumbar_Coeliac_VS_P1_P3_Stellate_Sphenopalatine | 0 | 1 | 0 | 1 | 1 | 1 | 0 | 0 |
| CARMIL1 | 0.358 | P1_P2_P3_P4_Lumbar_Sphenopalatine_VS_Coeliac_Stellate | 1 | 1 | 1 | 1 | 1 | 0 | 0 | 1 |
| PLXND1 | 0.358 | P1_P2_Lumbar_Coeliac_Sphenopalatine_VS_P3_P4_Stellate | 1 | 1 | 0 | 0 | 1 | 1 | 0 | 1 |
| DIP2B | 0.358 | P2_P3_P4_Lumbar_Coeliac_Sphenopalatine_VS_P1_Stellate | 0 | 1 | 1 | 1 | 1 | 1 | 0 | 1 |
| FILIP1L | 0.357 | P1_Coeliac_VS_P2_P3_P4_Lumbar_Stellate_Sphenopalatine | 1 | 0 | 0 | 0 | 0 | 1 | 0 | 0 |
| RCBTB2 | 0.357 | P1_Lumbar_Coeliac_VS_P2_P3_P4_Stellate_Sphenopalatine | 1 | 0 | 0 | 0 | 1 | 1 | 0 | 0 |
| GM27217 | 0.357 | P1_P3_Lumbar_Coeliac_VS_P2_P4_Stellate_Sphenopalatine | 1 | 0 | 1 | 0 | 1 | 1 | 0 | 0 |

| | | | | | | | | | |
|---|---|---|---|---|---|---|---|---|---|
| RHOU | 0.357 | P1_P2_P3_Lumbar_Coeliac_VS_P4_Stellate_Sphenopalatine | 1 | 1 | 1 | 0 | 1 | 1 | 0 | 0 |
| ZC3H8 | 0.357 | P1_P3_P4_Lumbar_Coeliac_VS_P2_Stellate_Sphenopalatine | 1 | 0 | 1 | 1 | 1 | 1 | 0 | 0 |
| KRBA1 | 0.357 | P1_P2_P4_Lumbar_Coeliac_Sphenopalatine_VS_P3_Stellate | 1 | 1 | 0 | 1 | 1 | 1 | 0 | 1 |
| CHST11 | 0.357 | P1_P2_P4_Lumbar_Coeliac_Sphenopalatine_VS_P3_Stellate | 1 | 1 | 0 | 1 | 1 | 1 | 0 | 1 |
| JMJD1C | 0.357 | P1_P3_P4_Lumbar_Coeliac_Sphenopalatine_VS_P2_Stellate | 1 | 0 | 1 | 1 | 1 | 1 | 0 | 1 |
| ARHGEF9 | 0.357 | P2_P3_P4_Lumbar_Coeliac_Sphenopalatine_VS_P1_Stellate | 0 | 1 | 1 | 1 | 1 | 1 | 0 | 1 |
| RASSF5 | 0.356 | P3_VS_P1_P2_P4_Lumbar_Coeliac_Stellate_Sphenopalatine | 0 | 0 | 1 | 0 | 0 | 0 | 0 | 0 |
| HSPA1A | 0.356 | P4_Lumbar_VS_P1_P2_P3_Coeliac_Stellate_Sphenopalatine | 0 | 0 | 0 | 1 | 1 | 0 | 0 | 0 |
| PRKX | 0.356 | P2_Lumbar_Coeliac_VS_P1_P3_P4_Stellate_Sphenopalatine | 0 | 1 | 0 | 0 | 1 | 1 | 0 | 0 |
| RBM10 | 0.356 | P3_Lumbar_Coeliac_VS_P1_P2_P4_Stellate_Sphenopalatine | 0 | 0 | 1 | 0 | 1 | 1 | 0 | 0 |
| PINX1 | 0.356 | P1_P3_P4_Lumbar_Coeliac_VS_P2_Stellate_Sphenopalatine | 1 | 0 | 1 | 1 | 1 | 1 | 0 | 0 |
| SOCS5 | 0.356 | P1_P2_P3_P4_Lumbar_Sphenopalatine_VS_Coeliac_Stellate | 1 | 1 | 1 | 1 | 1 | 0 | 0 | 1 |
| SCRIB | 0.356 | P1_P2_P4_Lumbar_Coeliac_Sphenopalatine_VS_P3_Stellate | 1 | 1 | 0 | 1 | 1 | 1 | 0 | 1 |
| RFFL | 0.355 | P3_VS_P1_P2_P4_Lumbar_Coeliac_Stellate_Sphenopalatine | 0 | 0 | 1 | 0 | 0 | 0 | 0 | 0 |
| DIPK1A | 0.355 | P4_Lumbar_VS_P1_P2_P3_Coeliac_Stellate_Sphenopalatine | 0 | 0 | 0 | 1 | 1 | 0 | 0 | 0 |
| P4HA1 | 0.355 | P4_Coeliac_VS_P1_P2_P3_Lumbar_Stellate_Sphenopalatine | 0 | 0 | 0 | 1 | 0 | 1 | 0 | 0 |
| PLEKHA8 | 0.355 | P1_Lumbar_Coeliac_VS_P2_P3_P4_Stellate_Sphenopalatine | 1 | 0 | 0 | 0 | 1 | 1 | 0 | 0 |
| MRPS31 | 0.355 | P1_Lumbar_Coeliac_VS_P2_P3_P4_Stellate_Sphenopalatine | 1 | 0 | 0 | 0 | 1 | 1 | 0 | 0 |
| LRPPRC | 0.355 | P2_P4_Lumbar_Coeliac_VS_P1_P3_Stellate_Sphenopalatine | 0 | 1 | 0 | 1 | 1 | 1 | 0 | 0 |
| MIPEP | 0.355 | P1_P3_P4_Lumbar_Coeliac_VS_P2_Stellate_Sphenopalatine | 1 | 0 | 1 | 1 | 1 | 1 | 0 | 0 |
| THAP12 | 0.355 | P2_P3_P4_Lumbar_Coeliac_VS_P1_Stellate_Sphenopalatine | 0 | 1 | 1 | 1 | 1 | 1 | 0 | 0 |
| COQ6 | 0.355 | P1_P2_P3_P4_Lumbar_Coeliac_Stellate_VS_Sphenopalatine | 1 | 1 | 1 | 1 | 1 | 1 | 1 | 0 |
| SH3PXD2A | 0.355 | P1_P2_P3_P4_Coeliac_Sphenopalatine_VS_Lumbar_Stellate | 1 | 1 | 1 | 1 | 0 | 1 | 0 | 1 |
| ZFP931 | 0.354 | P1_P2_P4_VS_P3_Lumbar_Coeliac_Stellate_Sphenopalatine | 1 | 1 | 0 | 1 | 0 | 0 | 0 | 0 |
| DGAT1 | 0.354 | P4_Lumbar_VS_P1_P2_P3_Coeliac_Stellate_Sphenopalatine | 0 | 0 | 0 | 1 | 1 | 0 | 0 | 0 |
| ABCG2 | 0.354 | P1_P2_P4_Lumbar_VS_P3_Coeliac_Stellate_Sphenopalatine | 1 | 1 | 0 | 1 | 1 | 0 | 0 | 0 |
| MPHOSPH9 | 0.354 | P3_Lumbar_Coeliac_VS_P1_P2_P4_Stellate_Sphenopalatine | 0 | 0 | 1 | 0 | 1 | 1 | 0 | 0 |
| RALB | 0.354 | P1_P2_P3_Lumbar_Coeliac_VS_P4_Stellate_Sphenopalatine | 1 | 1 | 1 | 0 | 1 | 1 | 0 | 0 |
| MCCC1 | 0.354 | P1_P2_P4_Lumbar_Coeliac_Sphenopalatine_VS_P3_Stellate | 1 | 1 | 0 | 1 | 1 | 1 | 0 | 1 |
| IFNAR2 | 0.354 | P1_P2_P4_Lumbar_Coeliac_Sphenopalatine_VS_P3_Stellate | 1 | 1 | 0 | 1 | 1 | 1 | 0 | 1 |

| | | | | | | | | | | |
|---|---|---|---|---|---|---|---|---|---|---|
| MYC | 0.354 | P1_P2_P4_Lumbar_Coeliac_Stellate_Sphenopalatine_VS_P3 | 1 | 1 | 0 | 1 | 1 | 1 | 1 | 1 |
| PSMG1 | 0.353 | P3_Lumbar_Coeliac_VS_P1_P2_P4_Stellate_Sphenopalatine | 0 | 0 | 1 | 0 | 1 | 1 | 0 | 0 |
| HADH | 0.353 | P1_P2_P3_P4_Lumbar_Coeliac_Stellate_VS_Sphenopalatine | 1 | 1 | 1 | 1 | 1 | 1 | 1 | 0 |
| IER5 | 0.353 | Sphenopalatine_VS_P1_P2_P3_P4_Lumbar_Coeliac_Stellate | 0 | 0 | 0 | 0 | 0 | 0 | 0 | 1 |
| RAB11FIP5 | 0.353 | P1_P2_P3_P4_Lumbar_Sphenopalatine_VS_Coeliac_Stellate | 1 | 1 | 1 | 1 | 1 | 0 | 0 | 1 |
| LRRN1 | 0.353 | P1_P3_Lumbar_Coeliac_Sphenopalatine_VS_P2_P4_Stellate | 1 | 0 | 1 | 0 | 1 | 1 | 0 | 1 |
| ZSCAN18 | 0.353 | P1_P2_P4_Lumbar_Coeliac_Sphenopalatine_VS_P3_Stellate | 1 | 1 | 0 | 1 | 1 | 1 | 0 | 1 |
| DNM3 | 0.353 | P2_P3_P4_Lumbar_Coeliac_Sphenopalatine_VS_P1_Stellate | 0 | 1 | 1 | 1 | 1 | 1 | 0 | 1 |
| DGKH | 0.353 | P1_P2_P4_Lumbar_Coeliac_Stellate_Sphenopalatine_VS_P3 | 1 | 1 | 0 | 1 | 1 | 1 | 1 | 1 |
| CHAC2 | 0.352 | P3_Lumbar_VS_P1_P2_P4_Coeliac_Stellate_Sphenopalatine | 0 | 0 | 1 | 0 | 1 | 0 | 0 | 0 |
| MBTPS2 | 0.352 | P3_Lumbar_Coeliac_VS_P1_P2_P4_Stellate_Sphenopalatine | 0 | 0 | 1 | 0 | 1 | 1 | 0 | 0 |
| ABCC8 | 0.352 | Sphenopalatine_VS_P1_P2_P3_P4_Lumbar_Coeliac_Stellate | 0 | 0 | 0 | 0 | 0 | 0 | 0 | 1 |
| E130008D07I | 0.352 | Sphenopalatine_VS_P1_P2_P3_P4_Lumbar_Coeliac_Stellate | 0 | 0 | 0 | 0 | 0 | 0 | 0 | 1 |
| HCN4 | 0.352 | Lumbar_Coeliac_Sphenopalatine_VS_P1_P2_P3_P4_Stellate | 0 | 0 | 0 | 0 | 1 | 1 | 0 | 1 |
| PTPDC1 | 0.352 | P1_P2_P4_Lumbar_Coeliac_Sphenopalatine_VS_P3_Stellate | 1 | 1 | 0 | 1 | 1 | 1 | 0 | 1 |
| IRF2BP1 | 0.352 | P1_P3_P4_Lumbar_Coeliac_Sphenopalatine_VS_P2_Stellate | 1 | 0 | 1 | 1 | 1 | 1 | 0 | 1 |
| AHR | 0.351 | P4_Lumbar_VS_P1_P2_P3_Coeliac_Stellate_Sphenopalatine | 0 | 0 | 0 | 1 | 1 | 0 | 0 | 0 |
| PDE7A | 0.351 | P3_Lumbar_Coeliac_VS_P1_P2_P4_Stellate_Sphenopalatine | 0 | 0 | 1 | 0 | 1 | 1 | 0 | 0 |
| RINT1 | 0.351 | P3_Lumbar_Coeliac_VS_P1_P2_P4_Stellate_Sphenopalatine | 0 | 0 | 1 | 0 | 1 | 1 | 0 | 0 |
| BNIP2 | 0.351 | P3_Lumbar_Coeliac_VS_P1_P2_P4_Stellate_Sphenopalatine | 0 | 0 | 1 | 0 | 1 | 1 | 0 | 0 |
| GALT | 0.351 | P1_P2_P3_P4_Lumbar_Sphenopalatine_VS_Coeliac_Stellate | 1 | 1 | 1 | 1 | 1 | 0 | 0 | 1 |
| SLC16A2 | 0.351 | P1_P2_P3_Lumbar_Coeliac_Sphenopalatine_VS_P4_Stellate | 1 | 1 | 1 | 0 | 1 | 1 | 0 | 1 |
| SERPINF1 | 0.351 | P1_P3_P4_Lumbar_Coeliac_Stellate_Sphenopalatine_VS_P2 | 1 | 0 | 1 | 1 | 1 | 1 | 1 | 1 |
| HMGXB4 | 0.35 | P1_Lumbar_Coeliac_VS_P2_P3_P4_Stellate_Sphenopalatine | 1 | 0 | 0 | 0 | 1 | 1 | 0 | 0 |
| PRKAR2B | 0.35 | P1_P3_Lumbar_Coeliac_VS_P2_P4_Stellate_Sphenopalatine | 1 | 0 | 1 | 0 | 1 | 1 | 0 | 0 |
| STAG2 | 0.35 | P2_P4_Lumbar_Coeliac_VS_P1_P3_Stellate_Sphenopalatine | 0 | 1 | 0 | 1 | 1 | 1 | 0 | 0 |
| MFAP3L | 0.35 | P1_P2_P4_Lumbar_Coeliac_Sphenopalatine_VS_P3_Stellate | 1 | 1 | 0 | 1 | 1 | 1 | 0 | 1 |
| KCTD9 | 0.349 | P2_VS_P1_P3_P4_Lumbar_Coeliac_Stellate_Sphenopalatine | 0 | 1 | 0 | 0 | 0 | 0 | 0 | 0 |
| GLOD4 | 0.349 | P1_P3_P4_Lumbar_Coeliac_VS_P2_Stellate_Sphenopalatine | 1 | 0 | 1 | 1 | 1 | 1 | 0 | 0 |
| 1500026H17I | 0.349 | P2_P3_P4_Lumbar_Coeliac_VS_P1_Stellate_Sphenopalatine | 0 | 1 | 1 | 1 | 1 | 1 | 0 | 0 |

| | | | | | | | | | | |
|---|---|---|---|---|---|---|---|---|---|---|
| DLX2 | 0.349 | Sphenopalatine_VS_P1_P2_P3_P4_Lumbar_Coeliac_Stellate | 0 | 0 | 0 | 0 | 0 | 0 | 0 | 1 |
| PHTF2 | 0.349 | P1_P2_P4_Lumbar_Coeliac_Sphenopalatine_VS_P3_Stellate | 1 | 1 | 0 | 1 | 1 | 1 | 0 | 1 |
| FKBP14 | 0.348 | P1_P2_P3_P4_Lumbar_Coeliac_Stellate_VS_Sphenopalatine | 1 | 1 | 1 | 1 | 1 | 1 | 1 | 0 |
| SLC6A4 | 0.348 | P1_Lumbar_Sphenopalatine_VS_P2_P3_P4_Coeliac_Stellate | 1 | 0 | 0 | 0 | 1 | 0 | 0 | 1 |
| RAB9B | 0.348 | P1_P2_P3_P4_Lumbar_Sphenopalatine_VS_Coeliac_Stellate | 1 | 1 | 1 | 1 | 1 | 0 | 0 | 1 |
| TOGARAM1 | 0.348 | P1_P2_P4_Lumbar_Coeliac_Sphenopalatine_VS_P3_Stellate | 1 | 1 | 0 | 1 | 1 | 1 | 0 | 1 |
| PPFIBP2 | 0.348 | P1_P3_P4_Lumbar_Coeliac_Stellate_Sphenopalatine_VS_P2 | 1 | 0 | 1 | 1 | 1 | 1 | 1 | 1 |
| ZFP397 | 0.347 | P1_Lumbar_Coeliac_VS_P2_P3_P4_Stellate_Sphenopalatine | 1 | 0 | 0 | 0 | 1 | 1 | 0 | 0 |
| NXPH3 | 0.347 | Sphenopalatine_VS_P1_P2_P3_P4_Lumbar_Coeliac_Stellate | 0 | 0 | 0 | 0 | 0 | 0 | 0 | 1 |
| STAT3 | 0.347 | P1_P2_P3_P4_Lumbar_Sphenopalatine_VS_Coeliac_Stellate | 1 | 1 | 1 | 1 | 1 | 0 | 0 | 1 |
| INF2 | 0.346 | P2_VS_P1_P3_P4_Lumbar_Coeliac_Stellate_Sphenopalatine | 0 | 1 | 0 | 0 | 0 | 0 | 0 | 0 |
| SMC4 | 0.346 | P1_P3_P4_Lumbar_Coeliac_VS_P2_Stellate_Sphenopalatine | 1 | 0 | 1 | 1 | 1 | 1 | 0 | 0 |
| PARVB | 0.346 | P1_P2_P3_Sphenopalatine_VS_P4_Lumbar_Coeliac_Stellate | 1 | 1 | 1 | 0 | 0 | 0 | 0 | 1 |
| CHRM1 | 0.346 | P1_P2_P4_Lumbar_Sphenopalatine_VS_P3_Coeliac_Stellate | 1 | 1 | 0 | 1 | 1 | 0 | 0 | 1 |
| TGFBRAP1 | 0.346 | P1_P2_P4_Lumbar_Coeliac_Sphenopalatine_VS_P3_Stellate | 1 | 1 | 0 | 1 | 1 | 1 | 0 | 1 |
| MTAP | 0.346 | P1_P2_P4_Lumbar_Coeliac_Sphenopalatine_VS_P3_Stellate | 1 | 1 | 0 | 1 | 1 | 1 | 0 | 1 |
| MFHAS1 | 0.346 | P1_P2_P4_Lumbar_Coeliac_Sphenopalatine_VS_P3_Stellate | 1 | 1 | 0 | 1 | 1 | 1 | 0 | 1 |
| DNAH1 | 0.345 | P3_Coeliac_VS_P1_P2_P4_Lumbar_Stellate_Sphenopalatine | 0 | 0 | 1 | 0 | 0 | 1 | 0 | 0 |
| ORC2 | 0.345 | P1_P3_P4_Lumbar_Coeliac_VS_P2_Stellate_Sphenopalatine | 1 | 0 | 1 | 1 | 1 | 1 | 0 | 0 |
| MAN1C1 | 0.345 | P1_P2_P3_P4_Lumbar_Coeliac_Stellate_VS_Sphenopalatine | 1 | 1 | 1 | 1 | 1 | 1 | 1 | 0 |
| MEPCE | 0.345 | P1_P2_P4_Lumbar_Coeliac_Sphenopalatine_VS_P3_Stellate | 1 | 1 | 0 | 1 | 1 | 1 | 0 | 1 |
| CACNA1A | 0.345 | P1_P2_P4_Lumbar_Coeliac_Sphenopalatine_VS_P3_Stellate | 1 | 1 | 0 | 1 | 1 | 1 | 0 | 1 |
| MRTFA | 0.345 | P1_P2_P4_Lumbar_Coeliac_Sphenopalatine_VS_P3_Stellate | 1 | 1 | 0 | 1 | 1 | 1 | 0 | 1 |
| PCX | 0.345 | P1_P2_P4_Lumbar_Coeliac_Stellate_Sphenopalatine_VS_P3 | 1 | 1 | 0 | 1 | 1 | 1 | 1 | 1 |
| YJEFN3 | 0.344 | P4_Lumbar_VS_P1_P2_P3_Coeliac_Stellate_Sphenopalatine | 0 | 0 | 0 | 1 | 1 | 0 | 0 | 0 |
| RAB27B | 0.344 | P4_Coeliac_VS_P1_P2_P3_Lumbar_Stellate_Sphenopalatine | 0 | 0 | 0 | 1 | 0 | 1 | 0 | 0 |
| ZBTB11OS1 | 0.344 | P1_P2_P3_P4_Sphenopalatine_VS_Lumbar_Coeliac_Stellate | 1 | 1 | 1 | 1 | 0 | 0 | 0 | 1 |
| GPX7 | 0.344 | P1_P2_P4_Lumbar_Coeliac_Stellate_Sphenopalatine_VS_P3 | 1 | 1 | 0 | 1 | 1 | 1 | 1 | 1 |
| PUS3 | 0.343 | P1_P4_Lumbar_Coeliac_VS_P2_P3_Stellate_Sphenopalatine | 1 | 0 | 0 | 1 | 1 | 1 | 0 | 0 |
| GPHN | 0.343 | P1_P4_Lumbar_Coeliac_VS_P2_P3_Stellate_Sphenopalatine | 1 | 0 | 0 | 1 | 1 | 1 | 0 | 0 |

| | | | | | | | | | | |
|---|---|---|---|---|---|---|---|---|---|---|
| MCRIP2 | 0.343 | P1_P2_P3_Lumbar_Coeliac_Sphenopalatine_VS_P4_Stellate | 1 | 1 | 1 | 0 | 1 | 1 | 0 | 1 |
| NAP1L2 | 0.343 | P1_P2_P4_Lumbar_Coeliac_Sphenopalatine_VS_P3_Stellate | 1 | 1 | 0 | 1 | 1 | 1 | 0 | 1 |
| GLIPR2 | 0.343 | P1_P2_P3_Lumbar_Coeliac_Stellate_Sphenopalatine_VS_P4 | 1 | 1 | 1 | 0 | 1 | 1 | 1 | 1 |
| PRKCA | 0.343 | P1_P2_P3_Lumbar_Coeliac_Stellate_Sphenopalatine_VS_P4 | 1 | 1 | 1 | 0 | 1 | 1 | 1 | 1 |
| SLC29A1 | 0.343 | P1_P2_P3_Lumbar_Coeliac_Stellate_Sphenopalatine_VS_P4 | 1 | 1 | 1 | 0 | 1 | 1 | 1 | 1 |
| AI429214 | 0.342 | P3_VS_P1_P2_P4_Lumbar_Coeliac_Stellate_Sphenopalatine | 0 | 0 | 1 | 0 | 0 | 0 | 0 | 0 |
| IFT81 | 0.342 | P1_P4_Lumbar_Coeliac_VS_P2_P3_Stellate_Sphenopalatine | 1 | 0 | 0 | 1 | 1 | 1 | 0 | 0 |
| STAG1 | 0.342 | P2_P4_Lumbar_Coeliac_VS_P1_P3_Stellate_Sphenopalatine | 0 | 1 | 0 | 1 | 1 | 1 | 0 | 0 |
| FKBP5 | 0.342 | P3_P4_Lumbar_Coeliac_VS_P1_P2_Stellate_Sphenopalatine | 0 | 0 | 1 | 1 | 1 | 1 | 0 | 0 |
| GCC2 | 0.342 | P1_P2_P4_Lumbar_Coeliac_Sphenopalatine_VS_P3_Stellate | 1 | 1 | 0 | 1 | 1 | 1 | 0 | 1 |
| BMF | 0.341 | P2_VS_P1_P3_P4_Lumbar_Coeliac_Stellate_Sphenopalatine | 0 | 1 | 0 | 0 | 0 | 0 | 0 | 0 |
| PRMT6 | 0.341 | P4_Lumbar_VS_P1_P2_P3_Coeliac_Stellate_Sphenopalatine | 0 | 0 | 0 | 1 | 1 | 0 | 0 | 0 |
| NUP93 | 0.34 | P1_P3_P4_Lumbar_Coeliac_VS_P2_Stellate_Sphenopalatine | 1 | 0 | 1 | 1 | 1 | 1 | 0 | 0 |
| VCL | 0.339 | P3_P4_VS_P1_P2_Lumbar_Coeliac_Stellate_Sphenopalatine | 0 | 0 | 1 | 1 | 0 | 0 | 0 | 0 |
| MICAL3 | 0.339 | P1_P2_P4_Lumbar_Coeliac_Sphenopalatine_VS_P3_Stellate | 1 | 1 | 0 | 1 | 1 | 1 | 0 | 1 |
| TDRP | 0.338 | P2_P3_P4_VS_P1_Lumbar_Coeliac_Stellate_Sphenopalatine | 0 | 1 | 1 | 1 | 0 | 0 | 0 | 0 |
| RRM2B | 0.338 | P2_P4_Lumbar_Coeliac_VS_P1_P3_Stellate_Sphenopalatine | 0 | 1 | 0 | 1 | 1 | 1 | 0 | 0 |
| ENPP4 | 0.338 | P2_P4_Lumbar_Coeliac_VS_P1_P3_Stellate_Sphenopalatine | 0 | 1 | 0 | 1 | 1 | 1 | 0 | 0 |
| KCNQ5 | 0.338 | P1_P2_P4_Lumbar_Coeliac_Sphenopalatine_VS_P3_Stellate | 1 | 1 | 0 | 1 | 1 | 1 | 0 | 1 |
| JAG2 | 0.338 | P1_P2_P4_Lumbar_Coeliac_Sphenopalatine_VS_P3_Stellate | 1 | 1 | 0 | 1 | 1 | 1 | 0 | 1 |
| MCFD2 | 0.337 | P1_P4_VS_P2_P3_Lumbar_Coeliac_Stellate_Sphenopalatine | 1 | 0 | 0 | 1 | 0 | 0 | 0 | 0 |
| MFSD10 | 0.337 | P1_P4_Lumbar_Coeliac_VS_P2_P3_Stellate_Sphenopalatine | 1 | 0 | 0 | 1 | 1 | 1 | 0 | 0 |
| RALGAPA2 | 0.337 | P2_P3_P4_Lumbar_Coeliac_VS_P1_Stellate_Sphenopalatine | 0 | 1 | 1 | 1 | 1 | 1 | 0 | 0 |
| BCAT1 | 0.337 | Lumbar_Coeliac_Stellate_VS_P1_P2_P3_P4_Sphenopalatine | 0 | 0 | 0 | 0 | 1 | 1 | 1 | 0 |
| ID1 | 0.337 | P1_P2_P3_Sphenopalatine_VS_P4_Lumbar_Coeliac_Stellate | 1 | 1 | 1 | 0 | 0 | 0 | 0 | 1 |
| PPP1R10 | 0.337 | P1_P2_P4_Lumbar_Coeliac_Sphenopalatine_VS_P3_Stellate | 1 | 1 | 0 | 1 | 1 | 1 | 0 | 1 |
| ADGRB3 | 0.337 | P1_P3_P4_Lumbar_Coeliac_Stellate_Sphenopalatine_VS_P2 | 1 | 0 | 1 | 1 | 1 | 1 | 1 | 1 |
| RUVBL1 | 0.336 | P1_P3_P4_Lumbar_Coeliac_VS_P2_Stellate_Sphenopalatine | 1 | 0 | 1 | 1 | 1 | 1 | 0 | 0 |
| EFNA3 | 0.336 | P1_P2_P3_P4_Lumbar_Coeliac_Stellate_VS_Sphenopalatine | 1 | 1 | 1 | 1 | 1 | 1 | 1 | 0 |
| RIMS2 | 0.336 | P1_P3_P4_Lumbar_Coeliac_Sphenopalatine_VS_P2_Stellate | 1 | 0 | 1 | 1 | 1 | 1 | 0 | 1 |

| | | | | | | | | | | |
|---|---|---|---|---|---|---|---|---|---|---|
| ADD3 | 0.336 | P1_P3_P4_Lumbar_Coeliac_Sphenopalatine_VS_P2_Stellate | 1 | 0 | 1 | 1 | 1 | 1 | 0 | 1 |
| MTIF3 | 0.335 | P1_P3_P4_Lumbar_Coeliac_VS_P2_Stellate_Sphenopalatine | 1 | 0 | 1 | 1 | 1 | 1 | 0 | 0 |
| MIR124A-1H | 0.335 | Lumbar_Coeliac_Sphenopalatine_VS_P1_P2_P3_P4_Stellate | 0 | 0 | 0 | 0 | 1 | 1 | 0 | 1 |
| BMPR1A | 0.335 | P1_P2_P4_Lumbar_Coeliac_Sphenopalatine_VS_P3_Stellate | 1 | 1 | 0 | 1 | 1 | 1 | 0 | 1 |
| BRIP1OS | 0.334 | P2_VS_P1_P3_P4_Lumbar_Coeliac_Stellate_Sphenopalatine | 0 | 1 | 0 | 0 | 0 | 0 | 0 | 0 |
| RESF1 | 0.334 | P1_Lumbar_Coeliac_VS_P2_P3_P4_Stellate_Sphenopalatine | 1 | 0 | 0 | 0 | 1 | 1 | 0 | 0 |
| PROSER2 | 0.334 | P1_P3_Lumbar_Coeliac_Stellate_VS_P2_P4_Sphenopalatine | 1 | 0 | 1 | 0 | 1 | 1 | 1 | 0 |
| PARP8 | 0.334 | P1_P2_P3_P4_Lumbar_Coeliac_Stellate_VS_Sphenopalatine | 1 | 1 | 1 | 1 | 1 | 1 | 1 | 0 |
| ZADH2 | 0.334 | P1_P2_P3_P4_Lumbar_Sphenopalatine_VS_Coeliac_Stellate | 1 | 1 | 1 | 1 | 1 | 0 | 0 | 1 |
| SH3RF1 | 0.334 | P1_P2_P4_Lumbar_Coeliac_Sphenopalatine_VS_P3_Stellate | 1 | 1 | 0 | 1 | 1 | 1 | 0 | 1 |
| N4BP2L1 | 0.333 | P1_P4_VS_P2_P3_Lumbar_Coeliac_Stellate_Sphenopalatine | 1 | 0 | 0 | 1 | 0 | 0 | 0 | 0 |
| NUP54 | 0.333 | P1_P4_Lumbar_Coeliac_VS_P2_P3_Stellate_Sphenopalatine | 1 | 0 | 0 | 1 | 1 | 1 | 0 | 0 |
| WASHC4 | 0.333 | P2_P4_Lumbar_Coeliac_VS_P1_P3_Stellate_Sphenopalatine | 0 | 1 | 0 | 1 | 1 | 1 | 0 | 0 |
| VANGL2 | 0.333 | P3_P4_Lumbar_Coeliac_VS_P1_P2_Stellate_Sphenopalatine | 0 | 0 | 1 | 1 | 1 | 1 | 0 | 0 |
| PPFIBP1 | 0.333 | P3_P4_Lumbar_Coeliac_VS_P1_P2_Stellate_Sphenopalatine | 0 | 0 | 1 | 1 | 1 | 1 | 0 | 0 |
| 2610027K06I | 0.333 | P1_P2_P4_Lumbar_Sphenopalatine_VS_P3_Coeliac_Stellate | 1 | 1 | 0 | 1 | 1 | 0 | 0 | 1 |
| SKIL | 0.333 | P1_P2_P4_Lumbar_Coeliac_Sphenopalatine_VS_P3_Stellate | 1 | 1 | 0 | 1 | 1 | 1 | 0 | 1 |
| NFXL1 | 0.332 | P2_Lumbar_Coeliac_VS_P1_P3_P4_Stellate_Sphenopalatine | 0 | 1 | 0 | 0 | 1 | 1 | 0 | 0 |
| LRRC42 | 0.332 | P3_P4_Lumbar_Coeliac_VS_P1_P2_Stellate_Sphenopalatine | 0 | 0 | 1 | 1 | 1 | 1 | 0 | 0 |
| CR1L | 0.332 | P1_P3_P4_Lumbar_Coeliac_VS_P2_Stellate_Sphenopalatine | 1 | 0 | 1 | 1 | 1 | 1 | 0 | 0 |
| BMI1 | 0.332 | P1_P3_P4_Lumbar_Coeliac_VS_P2_Stellate_Sphenopalatine | 1 | 0 | 1 | 1 | 1 | 1 | 0 | 0 |
| MSI2 | 0.332 | P1_P2_P3_Lumbar_Coeliac_Sphenopalatine_VS_P4_Stellate | 1 | 1 | 1 | 0 | 1 | 1 | 0 | 1 |
| SLC6A17 | 0.332 | P1_P2_P4_Lumbar_Coeliac_Sphenopalatine_VS_P3_Stellate | 1 | 1 | 0 | 1 | 1 | 1 | 0 | 1 |
| UTP18 | 0.331 | P1_Lumbar_Coeliac_VS_P2_P3_P4_Stellate_Sphenopalatine | 1 | 0 | 0 | 0 | 1 | 1 | 0 | 0 |
| OPLAH | 0.331 | P1_Lumbar_Coeliac_VS_P2_P3_P4_Stellate_Sphenopalatine | 1 | 0 | 0 | 0 | 1 | 1 | 0 | 0 |
| GLA | 0.331 | P1_P2_P3_Lumbar_Coeliac_VS_P4_Stellate_Sphenopalatine | 1 | 1 | 1 | 0 | 1 | 1 | 0 | 0 |
| PPHLN1 | 0.331 | P1_P4_Lumbar_Coeliac_VS_P2_P3_Stellate_Sphenopalatine | 1 | 0 | 0 | 1 | 1 | 1 | 0 | 0 |
| ST3GAL5 | 0.331 | P2_P4_Lumbar_Coeliac_VS_P1_P3_Stellate_Sphenopalatine | 0 | 1 | 0 | 1 | 1 | 1 | 0 | 0 |
| IGLON5 | 0.331 | P1_P2_P4_Lumbar_Coeliac_Sphenopalatine_VS_P3_Stellate | 1 | 1 | 0 | 1 | 1 | 1 | 0 | 1 |
| ARHGAP23 | 0.331 | P1_P2_P4_Lumbar_Coeliac_Sphenopalatine_VS_P3_Stellate | 1 | 1 | 0 | 1 | 1 | 1 | 0 | 1 |

| | | | | | | | | | | |
|---|---|---|---|---|---|---|---|---|---|---|
| RAI2 | 0.33 | P2_P4_VS_P1_P3_Lumbar_Coeliac_Stellate_Sphenopalatine | 0 | 1 | 0 | 1 | 0 | 0 | 0 | 0 |
| ZFP647 | 0.33 | P1_Lumbar_Coeliac_VS_P2_P3_P4_Stellate_Sphenopalatine | 1 | 0 | 0 | 0 | 1 | 1 | 0 | 0 |
| SUN2 | 0.33 | P2_P4_Lumbar_Coeliac_VS_P1_P3_Stellate_Sphenopalatine | 0 | 1 | 0 | 1 | 1 | 1 | 0 | 0 |
| TTC37 | 0.33 | P1_P3_P4_Lumbar_Coeliac_VS_P2_Stellate_Sphenopalatine | 1 | 0 | 1 | 1 | 1 | 1 | 0 | 0 |
| WDR46 | 0.33 | P1_P3_P4_Lumbar_Coeliac_VS_P2_Stellate_Sphenopalatine | 1 | 0 | 1 | 1 | 1 | 1 | 0 | 0 |
| TYRO3 | 0.33 | P1_P3_Sphenopalatine_VS_P2_P4_Lumbar_Coeliac_Stellate | 1 | 0 | 1 | 0 | 0 | 0 | 0 | 1 |
| TRIM56 | 0.33 | P1_P2_P4_Lumbar_Coeliac_Sphenopalatine_VS_P3_Stellate | 1 | 1 | 0 | 1 | 1 | 1 | 0 | 1 |
| CD59A | 0.33 | P1_P2_P3_P4_Stellate_Sphenopalatine_VS_Lumbar_Coeliac | 1 | 1 | 1 | 1 | 0 | 0 | 1 | 1 |
| 9330020H09I | 0.329 | P3_VS_P1_P2_P4_Lumbar_Coeliac_Stellate_Sphenopalatine | 0 | 0 | 1 | 0 | 0 | 0 | 0 | 0 |
| ENTPD3 | 0.329 | P1_P3_VS_P2_P4_Lumbar_Coeliac_Stellate_Sphenopalatine | 1 | 0 | 1 | 0 | 0 | 0 | 0 | 0 |
| PAKAP.1 | 0.329 | P2_P4_VS_P1_P3_Lumbar_Coeliac_Stellate_Sphenopalatine | 0 | 1 | 0 | 1 | 0 | 0 | 0 | 0 |
| HSPBAP1 | 0.329 | P1_Lumbar_Coeliac_VS_P2_P3_P4_Stellate_Sphenopalatine | 1 | 0 | 0 | 0 | 1 | 1 | 0 | 0 |
| PPP4R2 | 0.329 | P2_Lumbar_Coeliac_VS_P1_P3_P4_Stellate_Sphenopalatine | 0 | 1 | 0 | 0 | 1 | 1 | 0 | 0 |
| TXLNG | 0.329 | P3_P4_Lumbar_Coeliac_VS_P1_P2_Stellate_Sphenopalatine | 0 | 0 | 1 | 1 | 1 | 1 | 0 | 0 |
| TOPORS | 0.329 | P1_P3_P4_Lumbar_Coeliac_VS_P2_Stellate_Sphenopalatine | 1 | 0 | 1 | 1 | 1 | 1 | 0 | 0 |
| NASP | 0.329 | P1_P3_P4_Lumbar_Coeliac_VS_P2_Stellate_Sphenopalatine | 1 | 0 | 1 | 1 | 1 | 1 | 0 | 0 |
| COG1 | 0.329 | P1_P3_P4_Lumbar_Coeliac_VS_P2_Stellate_Sphenopalatine | 1 | 0 | 1 | 1 | 1 | 1 | 0 | 0 |
| TRIM9 | 0.329 | P1_P2_P3_Lumbar_Coeliac_Sphenopalatine_VS_P4_Stellate | 1 | 1 | 1 | 0 | 1 | 1 | 0 | 1 |
| OSBPL1A | 0.329 | P4_Lumbar_Coeliac_Sphenopalatine_VS_P1_P2_P3_Stellate | 0 | 0 | 0 | 1 | 1 | 1 | 0 | 1 |
| PIK3R4 | 0.329 | P1_P2_P4_Lumbar_Coeliac_Sphenopalatine_VS_P3_Stellate | 1 | 1 | 0 | 1 | 1 | 1 | 0 | 1 |
| PITRM1 | 0.329 | P1_P2_P4_Lumbar_Coeliac_Sphenopalatine_VS_P3_Stellate | 1 | 1 | 0 | 1 | 1 | 1 | 0 | 1 |
| HEY1 | 0.329 | P2_P3_P4_Lumbar_Coeliac_Stellate_Sphenopalatine_VS_P1 | 0 | 1 | 1 | 1 | 1 | 1 | 1 | 1 |
| AVEN | 0.328 | P4_Lumbar_VS_P1_P2_P3_Coeliac_Stellate_Sphenopalatine | 0 | 0 | 0 | 1 | 1 | 0 | 0 | 0 |
| RAP2C | 0.328 | P3_P4_Lumbar_Coeliac_VS_P1_P2_Stellate_Sphenopalatine | 0 | 0 | 1 | 1 | 1 | 1 | 0 | 0 |
| BAZ1B | 0.328 | P2_P3_P4_Lumbar_Coeliac_VS_P1_Stellate_Sphenopalatine | 0 | 1 | 1 | 1 | 1 | 1 | 0 | 0 |
| 2310015A10I | 0.328 | P4_Sphenopalatine_VS_P1_P2_P3_Lumbar_Coeliac_Stellate | 0 | 0 | 0 | 1 | 0 | 0 | 0 | 1 |
| RNF19A | 0.328 | P1_P2_P4_Lumbar_Coeliac_Sphenopalatine_VS_P3_Stellate | 1 | 1 | 0 | 1 | 1 | 1 | 0 | 1 |
| ZFP219 | 0.327 | P2_Lumbar_Coeliac_VS_P1_P3_P4_Stellate_Sphenopalatine | 0 | 1 | 0 | 0 | 1 | 1 | 0 | 0 |
| A730020E08I | 0.327 | P1_P3_Lumbar_Coeliac_VS_P2_P4_Stellate_Sphenopalatine | 1 | 0 | 1 | 0 | 1 | 1 | 0 | 0 |
| MIOS | 0.327 | P1_P4_Lumbar_Coeliac_VS_P2_P3_Stellate_Sphenopalatine | 1 | 0 | 0 | 1 | 1 | 1 | 0 | 0 |

| | | | | | | | | | | |
|---|---|---|---|---|---|---|---|---|---|---|
| PEX5 | 0.327 | P1_P2_P3_P4_Lumbar_Sphenopalatine_VS_Coeliac_Stellate | 1 | 1 | 1 | 1 | 1 | 0 | 0 | 1 |
| WWC2 | 0.327 | P1_P2_P3_P4_Lumbar_Sphenopalatine_VS_Coeliac_Stellate | 1 | 1 | 1 | 1 | 1 | 0 | 0 | 1 |
| JPT2 | 0.327 | P1_P3_P4_Lumbar_Coeliac_Stellate_Sphenopalatine_VS_P2 | 1 | 0 | 1 | 1 | 1 | 1 | 1 | 1 |
| SETDB1 | 0.326 | P2_P4_Lumbar_Coeliac_VS_P1_P3_Stellate_Sphenopalatine | 0 | 1 | 0 | 1 | 1 | 1 | 0 | 0 |
| PTBP1 | 0.326 | P2_P4_Lumbar_Coeliac_VS_P1_P3_Stellate_Sphenopalatine | 0 | 1 | 0 | 1 | 1 | 1 | 0 | 0 |
| GPR135 | 0.326 | P2_P4_Lumbar_Coeliac_VS_P1_P3_Stellate_Sphenopalatine | 0 | 1 | 0 | 1 | 1 | 1 | 0 | 0 |
| KDM4A | 0.326 | P1_P2_P4_Lumbar_Coeliac_Sphenopalatine_VS_P3_Stellate | 1 | 1 | 0 | 1 | 1 | 1 | 0 | 1 |
| KHNYN | 0.326 | P1_P2_P4_Lumbar_Coeliac_Sphenopalatine_VS_P3_Stellate | 1 | 1 | 0 | 1 | 1 | 1 | 0 | 1 |
| CLPX | 0.325 | P1_P4_Lumbar_Coeliac_VS_P2_P3_Stellate_Sphenopalatine | 1 | 0 | 0 | 1 | 1 | 1 | 0 | 0 |
| TTC8 | 0.325 | P2_P4_Lumbar_Coeliac_VS_P1_P3_Stellate_Sphenopalatine | 0 | 1 | 0 | 1 | 1 | 1 | 0 | 0 |
| DGKQ | 0.325 | P1_P2_P4_Lumbar_Coeliac_Sphenopalatine_VS_P3_Stellate | 1 | 1 | 0 | 1 | 1 | 1 | 0 | 1 |
| RAD23A | 0.325 | P1_P2_P4_Lumbar_Coeliac_Sphenopalatine_VS_P3_Stellate | 1 | 1 | 0 | 1 | 1 | 1 | 0 | 1 |
| PRPF6 | 0.324 | P1_P4_Lumbar_Coeliac_VS_P2_P3_Stellate_Sphenopalatine | 1 | 0 | 0 | 1 | 1 | 1 | 0 | 0 |
| WDR43 | 0.324 | P1_P4_Lumbar_Coeliac_VS_P2_P3_Stellate_Sphenopalatine | 1 | 0 | 0 | 1 | 1 | 1 | 0 | 0 |
| TMEM117 | 0.324 | P1_P2_P3_P4_Sphenopalatine_VS_Lumbar_Coeliac_Stellate | 1 | 1 | 1 | 1 | 0 | 0 | 0 | 1 |
| ST3GAL4 | 0.324 | P3_Lumbar_Coeliac_Sphenopalatine_VS_P1_P2_P4_Stellate | 0 | 0 | 1 | 0 | 1 | 1 | 0 | 1 |
| SUCO | 0.324 | P1_P2_P4_Lumbar_Coeliac_Sphenopalatine_VS_P3_Stellate | 1 | 1 | 0 | 1 | 1 | 1 | 0 | 1 |
| AKT3 | 0.324 | P1_P2_P4_Lumbar_Coeliac_Sphenopalatine_VS_P3_Stellate | 1 | 1 | 0 | 1 | 1 | 1 | 0 | 1 |
| CCDC74A | 0.323 | P2_P3_P4_VS_P1_Lumbar_Coeliac_Stellate_Sphenopalatine | 0 | 1 | 1 | 1 | 0 | 0 | 0 | 0 |
| ZDHHC14 | 0.323 | P4_Lumbar_VS_P1_P2_P3_Coeliac_Stellate_Sphenopalatine | 0 | 0 | 0 | 1 | 1 | 0 | 0 | 0 |
| PDE3B | 0.323 | P1_P4_Lumbar_Coeliac_VS_P2_P3_Stellate_Sphenopalatine | 1 | 0 | 0 | 1 | 1 | 1 | 0 | 0 |
| COIL | 0.323 | P1_P3_P4_Lumbar_Coeliac_VS_P2_Stellate_Sphenopalatine | 1 | 0 | 1 | 1 | 1 | 1 | 0 | 0 |
| TET1 | 0.323 | P2_P3_P4_Lumbar_Coeliac_VS_P1_Stellate_Sphenopalatine | 0 | 1 | 1 | 1 | 1 | 1 | 0 | 0 |
| PPP4R3A | 0.323 | P2_P3_P4_Lumbar_Coeliac_VS_P1_Stellate_Sphenopalatine | 0 | 1 | 1 | 1 | 1 | 1 | 0 | 0 |
| DTX3L | 0.323 | P1_P2_Sphenopalatine_VS_P3_P4_Lumbar_Coeliac_Stellate | 1 | 1 | 0 | 0 | 0 | 0 | 0 | 1 |
| GTF3C1 | 0.323 | P1_P2_P4_Lumbar_Coeliac_Sphenopalatine_VS_P3_Stellate | 1 | 1 | 0 | 1 | 1 | 1 | 0 | 1 |
| MPP5 | 0.323 | P1_P2_P4_Lumbar_Coeliac_Sphenopalatine_VS_P3_Stellate | 1 | 1 | 0 | 1 | 1 | 1 | 0 | 1 |
| DENND6A | 0.323 | P1_P2_P4_Lumbar_Coeliac_Sphenopalatine_VS_P3_Stellate | 1 | 1 | 0 | 1 | 1 | 1 | 0 | 1 |
| FAM167A | 0.322 | P3_VS_P1_P2_P4_Lumbar_Coeliac_Stellate_Sphenopalatine | 0 | 0 | 1 | 0 | 0 | 0 | 0 | 0 |
| SMIM19 | 0.322 | P1_P2_P3_P4_Lumbar_VS_Coeliac_Stellate_Sphenopalatine | 1 | 1 | 1 | 1 | 1 | 0 | 0 | 0 |

| | | | | | | | | | | |
|---|---|---|---|---|---|---|---|---|---|---|
| HAUS7 | 0.322 | P3_P4_Lumbar_Coeliac_VS_P1_P2_Stellate_Sphenopalatine | 0 | 0 | 1 | 1 | 1 | 1 | 0 | 0 |
| ZCCHC2 | 0.322 | P1_P3_P4_Lumbar_Coeliac_VS_P2_Stellate_Sphenopalatine | 1 | 0 | 1 | 1 | 1 | 1 | 0 | 0 |
| BMT2 | 0.322 | P2_P3_P4_Lumbar_Coeliac_VS_P1_Stellate_Sphenopalatine | 0 | 1 | 1 | 1 | 1 | 1 | 0 | 0 |
| 2610008E11I | 0.322 | P2_P3_P4_Lumbar_Coeliac_VS_P1_Stellate_Sphenopalatine | 0 | 1 | 1 | 1 | 1 | 1 | 0 | 0 |
| SAP30 | 0.322 | P1_P3_P4_Lumbar_Coeliac_Stellate_VS_P2_Sphenopalatine | 1 | 0 | 1 | 1 | 1 | 1 | 1 | 0 |
| LZTS1 | 0.322 | P1_P2_P3_Lumbar_Coeliac_Sphenopalatine_VS_P4_Stellate | 1 | 1 | 1 | 0 | 1 | 1 | 0 | 1 |
| MBD5 | 0.322 | P1_P2_P4_Lumbar_Coeliac_Sphenopalatine_VS_P3_Stellate | 1 | 1 | 0 | 1 | 1 | 1 | 0 | 1 |
| MTHFR | 0.322 | P1_P2_P4_Lumbar_Coeliac_Sphenopalatine_VS_P3_Stellate | 1 | 1 | 0 | 1 | 1 | 1 | 0 | 1 |
| ACADSB | 0.322 | P1_P2_P4_Lumbar_Coeliac_Sphenopalatine_VS_P3_Stellate | 1 | 1 | 0 | 1 | 1 | 1 | 0 | 1 |
| NAT8L | 0.321 | P2_P4_VS_P1_P3_Lumbar_Coeliac_Stellate_Sphenopalatine | 0 | 1 | 0 | 1 | 0 | 0 | 0 | 0 |
| PTPN2 | 0.321 | P1_P4_Lumbar_VS_P2_P3_Coeliac_Stellate_Sphenopalatine | 1 | 0 | 0 | 1 | 1 | 0 | 0 | 0 |
| RBM28 | 0.321 | P1_P4_Lumbar_Coeliac_VS_P2_P3_Stellate_Sphenopalatine | 1 | 0 | 0 | 1 | 1 | 1 | 0 | 0 |
| FUBP3 | 0.321 | P2_P4_Lumbar_Coeliac_VS_P1_P3_Stellate_Sphenopalatine | 0 | 1 | 0 | 1 | 1 | 1 | 0 | 0 |
| NOP14 | 0.321 | P1_P3_P4_Lumbar_Coeliac_VS_P2_Stellate_Sphenopalatine | 1 | 0 | 1 | 1 | 1 | 1 | 0 | 0 |
| NOC4L | 0.321 | P1_P3_P4_Lumbar_Coeliac_VS_P2_Stellate_Sphenopalatine | 1 | 0 | 1 | 1 | 1 | 1 | 0 | 0 |
| EPHA10 | 0.321 | Lumbar_Coeliac_Stellate_VS_P1_P2_P3_P4_Sphenopalatine | 0 | 0 | 0 | 0 | 1 | 1 | 1 | 0 |
| CORO7 | 0.321 | P1_P2_P4_Lumbar_Coeliac_Sphenopalatine_VS_P3_Stellate | 1 | 1 | 0 | 1 | 1 | 1 | 0 | 1 |
| PARD6G | 0.32 | P2_Lumbar_Coeliac_VS_P1_P3_P4_Stellate_Sphenopalatine | 0 | 1 | 0 | 0 | 1 | 1 | 0 | 0 |
| FAM78B | 0.32 | P1_P2_P3_Lumbar_Coeliac_VS_P4_Stellate_Sphenopalatine | 1 | 1 | 1 | 0 | 1 | 1 | 0 | 0 |
| GSPT2 | 0.32 | P1_P4_Lumbar_Coeliac_VS_P2_P3_Stellate_Sphenopalatine | 1 | 0 | 0 | 1 | 1 | 1 | 0 | 0 |
| WDR75 | 0.32 | P3_P4_Lumbar_Coeliac_VS_P1_P2_Stellate_Sphenopalatine | 0 | 0 | 1 | 1 | 1 | 1 | 0 | 0 |
| SRFBP1 | 0.32 | P3_P4_Lumbar_Coeliac_VS_P1_P2_Stellate_Sphenopalatine | 0 | 0 | 1 | 1 | 1 | 1 | 0 | 0 |
| RARS2 | 0.32 | P1_P3_P4_Lumbar_Coeliac_VS_P2_Stellate_Sphenopalatine | 1 | 0 | 1 | 1 | 1 | 1 | 0 | 0 |
| UHRF1BP1L | 0.32 | P2_P3_P4_Lumbar_Coeliac_VS_P1_Stellate_Sphenopalatine | 0 | 1 | 1 | 1 | 1 | 1 | 0 | 0 |
| USP11 | 0.32 | P2_P3_P4_Lumbar_Coeliac_Sphenopalatine_VS_P1_Stellate | 0 | 1 | 1 | 1 | 1 | 1 | 0 | 1 |
| ANKLE2 | 0.319 | P4_Coeliac_VS_P1_P2_P3_Lumbar_Stellate_Sphenopalatine | 0 | 0 | 0 | 1 | 0 | 1 | 0 | 0 |
| EML2 | 0.318 | P1_P2_P3_P4_Lumbar_Sphenopalatine_VS_Coeliac_Stellate | 1 | 1 | 1 | 1 | 1 | 0 | 0 | 1 |
| PITPNM3 | 0.318 | Coeliac_Sphenopalatine_VS_P1_P2_P3_P4_Lumbar_Stellate | 0 | 0 | 0 | 0 | 0 | 1 | 0 | 1 |
| FRS3 | 0.318 | Lumbar_Coeliac_Sphenopalatine_VS_P1_P2_P3_P4_Stellate | 0 | 0 | 0 | 0 | 1 | 1 | 0 | 1 |
| BID | 0.317 | P3_Lumbar_VS_P1_P2_P4_Coeliac_Stellate_Sphenopalatine | 0 | 0 | 1 | 0 | 1 | 0 | 0 | 0 |

| | | | | | | | | | | |
|---|---|---|---|---|---|---|---|---|---|---|
| EML4 | 0.317 | P1_P2_P3_P4_Lumbar_Coeliac_Stellate_VS_Sphenopalatine | 1 | 1 | 1 | 1 | 1 | 1 | 1 | 0 |
| TCOF1 | 0.317 | P1_P3_P4_Lumbar_Coeliac_Sphenopalatine_VS_P2_Stellate | 1 | 0 | 1 | 1 | 1 | 1 | 0 | 1 |
| GM26917 | 0.317 | P2_P3_P4_Lumbar_Coeliac_Sphenopalatine_VS_P1_Stellate | 0 | 1 | 1 | 1 | 1 | 1 | 0 | 1 |
| RBMS2 | 0.316 | P2_VS_P1_P3_P4_Lumbar_Coeliac_Stellate_Sphenopalatine | 0 | 1 | 0 | 0 | 0 | 0 | 0 | 0 |
| DCAF12L1 | 0.316 | P2_P4_VS_P1_P3_Lumbar_Coeliac_Stellate_Sphenopalatine | 0 | 1 | 0 | 1 | 0 | 0 | 0 | 0 |
| GM28050 | 0.316 | P2_P3_P4_Lumbar_Coeliac_Stellate_VS_P1_Sphenopalatine | 0 | 1 | 1 | 1 | 1 | 1 | 1 | 0 |
| ADAM11 | 0.316 | Sphenopalatine_VS_P1_P2_P3_P4_Lumbar_Coeliac_Stellate | 0 | 0 | 0 | 0 | 0 | 0 | 0 | 1 |
| GRAMD3 | 0.315 | P1_VS_P2_P3_P4_Lumbar_Coeliac_Stellate_Sphenopalatine | 1 | 0 | 0 | 0 | 0 | 0 | 0 | 0 |
| GM14326 | 0.315 | P2_P4_VS_P1_P3_Lumbar_Coeliac_Stellate_Sphenopalatine | 0 | 1 | 0 | 1 | 0 | 0 | 0 | 0 |
| RNPC3 | 0.315 | P1_Lumbar_Coeliac_VS_P2_P3_P4_Stellate_Sphenopalatine | 1 | 0 | 0 | 0 | 1 | 1 | 0 | 0 |
| CEP57L1 | 0.315 | P1_Lumbar_Coeliac_VS_P2_P3_P4_Stellate_Sphenopalatine | 1 | 0 | 0 | 0 | 1 | 1 | 0 | 0 |
| RAPGEF2 | 0.315 | P2_P3_P4_Lumbar_Coeliac_VS_P1_Stellate_Sphenopalatine | 0 | 1 | 1 | 1 | 1 | 1 | 0 | 0 |
| DCHS1 | 0.315 | P2_P3_P4_Lumbar_Coeliac_VS_P1_Stellate_Sphenopalatine | 0 | 1 | 1 | 1 | 1 | 1 | 0 | 0 |
| PACSIN1 | 0.315 | P1_P2_P3_P4_Sphenopalatine_VS_Lumbar_Coeliac_Stellate | 1 | 1 | 1 | 1 | 0 | 0 | 0 | 1 |
| DZIP1 | 0.315 | P1_P3_P4_Lumbar_Coeliac_Sphenopalatine_VS_P2_Stellate | 1 | 0 | 1 | 1 | 1 | 1 | 0 | 1 |
| CELA1 | 0.314 | P1_P3_P4_Sphenopalatine_VS_P2_Lumbar_Coeliac_Stellate | 1 | 0 | 1 | 1 | 0 | 0 | 0 | 1 |
| MBNL1 | 0.314 | P1_P2_P3_P4_Lumbar_Sphenopalatine_VS_Coeliac_Stellate | 1 | 1 | 1 | 1 | 1 | 0 | 0 | 1 |
| GSTM1 | 0.314 | P1_P2_P3_Lumbar_Coeliac_Stellate_Sphenopalatine_VS_P4 | 1 | 1 | 1 | 0 | 1 | 1 | 1 | 1 |
| ITPKA | 0.313 | P1_VS_P2_P3_P4_Lumbar_Coeliac_Stellate_Sphenopalatine | 1 | 0 | 0 | 0 | 0 | 0 | 0 | 0 |
| RAD54L2 | 0.313 | P2_P4_Lumbar_Coeliac_VS_P1_P3_Stellate_Sphenopalatine | 0 | 1 | 0 | 1 | 1 | 1 | 0 | 0 |
| KLF9 | 0.313 | P1_P2_P4_Sphenopalatine_VS_P3_Lumbar_Coeliac_Stellate | 1 | 1 | 0 | 1 | 0 | 0 | 0 | 1 |
| MEGF9 | 0.313 | P1_P2_P3_Lumbar_Coeliac_Sphenopalatine_VS_P4_Stellate | 1 | 1 | 1 | 0 | 1 | 1 | 0 | 1 |
| IL17RD | 0.312 | P2_Lumbar_Coeliac_VS_P1_P3_P4_Stellate_Sphenopalatine | 0 | 1 | 0 | 0 | 1 | 1 | 0 | 0 |
| ARNTL | 0.312 | P3_P4_Lumbar_Coeliac_VS_P1_P2_Stellate_Sphenopalatine | 0 | 0 | 1 | 1 | 1 | 1 | 0 | 0 |
| RNF216 | 0.312 | P2_P3_P4_Lumbar_Coeliac_VS_P1_Stellate_Sphenopalatine | 0 | 1 | 1 | 1 | 1 | 1 | 0 | 0 |
| TUBA4A | 0.312 | P4_Sphenopalatine_VS_P1_P2_P3_Lumbar_Coeliac_Stellate | 0 | 0 | 0 | 1 | 0 | 0 | 0 | 1 |
| ARRDC4 | 0.311 | P3_P4_VS_P1_P2_Lumbar_Coeliac_Stellate_Sphenopalatine | 0 | 0 | 1 | 1 | 0 | 0 | 0 | 0 |
| CRADD | 0.311 | P3_P4_VS_P1_P2_Lumbar_Coeliac_Stellate_Sphenopalatine | 0 | 0 | 1 | 1 | 0 | 0 | 0 | 0 |
| MCM7 | 0.311 | P1_Lumbar_Coeliac_VS_P2_P3_P4_Stellate_Sphenopalatine | 1 | 0 | 0 | 0 | 1 | 1 | 0 | 0 |
| RGS12 | 0.311 | P2_Lumbar_Coeliac_VS_P1_P3_P4_Stellate_Sphenopalatine | 0 | 1 | 0 | 0 | 1 | 1 | 0 | 0 |

| | | | | | | | | | | |
|---|---|---|---|---|---|---|---|---|---|---|
| PTCH1 | 0.311 | P2_Lumbar_Coeliac_VS_P1_P3_P4_Stellate_Sphenopalatine | 0 | 1 | 0 | 0 | 1 | 1 | 0 | 0 |
| PSAT1 | 0.311 | P1_P3_Lumbar_Coeliac_Stellate_Sphenopalatine_VS_P2_P4 | 1 | 0 | 1 | 0 | 1 | 1 | 1 | 1 |
| GPR162 | 0.311 | P1_P2_P3_Lumbar_Coeliac_Stellate_Sphenopalatine_VS_P4 | 1 | 1 | 1 | 0 | 1 | 1 | 1 | 1 |
| AIFM1 | 0.31 | P1_Lumbar_Coeliac_VS_P2_P3_P4_Stellate_Sphenopalatine | 1 | 0 | 0 | 0 | 1 | 1 | 0 | 0 |
| SLC45A4 | 0.31 | P1_P2_Lumbar_Coeliac_VS_P3_P4_Stellate_Sphenopalatine | 1 | 1 | 0 | 0 | 1 | 1 | 0 | 0 |
| KLHL13 | 0.31 | P1_P2_P3_P4_Lumbar_Coeliac_Stellate_VS_Sphenopalatine | 1 | 1 | 1 | 1 | 1 | 1 | 1 | 0 |
| ERF | 0.31 | P1_P2_P3_P4_Coeliac_Sphenopalatine_VS_Lumbar_Stellate | 1 | 1 | 1 | 1 | 0 | 1 | 0 | 1 |
| RIMS4 | 0.31 | P1_P2_P3_Lumbar_Coeliac_Sphenopalatine_VS_P4_Stellate | 1 | 1 | 1 | 0 | 1 | 1 | 0 | 1 |
| LSG1 | 0.309 | P1_Lumbar_Coeliac_VS_P2_P3_P4_Stellate_Sphenopalatine | 1 | 0 | 0 | 0 | 1 | 1 | 0 | 0 |
| WNK3 | 0.309 | P2_Lumbar_Coeliac_VS_P1_P3_P4_Stellate_Sphenopalatine | 0 | 1 | 0 | 0 | 1 | 1 | 0 | 0 |
| CFAP97 | 0.309 | P3_P4_Lumbar_Coeliac_VS_P1_P2_Stellate_Sphenopalatine | 0 | 0 | 1 | 1 | 1 | 1 | 0 | 0 |
| PLAGL1 | 0.309 | P3_P4_Lumbar_Coeliac_VS_P1_P2_Stellate_Sphenopalatine | 0 | 0 | 1 | 1 | 1 | 1 | 0 | 0 |
| HIVEP1 | 0.309 | P1_P3_P4_Lumbar_Coeliac_Sphenopalatine_VS_P2_Stellate | 1 | 0 | 1 | 1 | 1 | 1 | 0 | 1 |
| TTLL5 | 0.308 | P4_Lumbar_VS_P1_P2_P3_Coeliac_Stellate_Sphenopalatine | 0 | 0 | 0 | 1 | 1 | 0 | 0 | 0 |
| EFR3A | 0.308 | P1_P2_P3_P4_Lumbar_Coeliac_Stellate_VS_Sphenopalatine | 1 | 1 | 1 | 1 | 1 | 1 | 1 | 0 |
| CARM1 | 0.308 | P1_P2_P3_Lumbar_Coeliac_Sphenopalatine_VS_P4_Stellate | 1 | 1 | 1 | 0 | 1 | 1 | 0 | 1 |
| FZD3 | 0.307 | P1_P2_P3_Lumbar_Coeliac_VS_P4_Stellate_Sphenopalatine | 1 | 1 | 1 | 0 | 1 | 1 | 0 | 0 |
| GPALPP1 | 0.307 | P2_P3_P4_Lumbar_Coeliac_VS_P1_Stellate_Sphenopalatine | 0 | 1 | 1 | 1 | 1 | 1 | 0 | 0 |
| TCN2 | 0.307 | P1_P2_P3_P4_Lumbar_Coeliac_Stellate_VS_Sphenopalatine | 1 | 1 | 1 | 1 | 1 | 1 | 1 | 0 |
| UHRF1BP1 | 0.307 | P1_P2_Sphenopalatine_VS_P3_P4_Lumbar_Coeliac_Stellate | 1 | 1 | 0 | 0 | 0 | 0 | 0 | 1 |
| NUP214 | 0.307 | P1_P3_P4_Lumbar_Coeliac_Sphenopalatine_VS_P2_Stellate | 1 | 0 | 1 | 1 | 1 | 1 | 0 | 1 |
| AGFG2 | 0.306 | P4_Lumbar_VS_P1_P2_P3_Coeliac_Stellate_Sphenopalatine | 0 | 0 | 0 | 1 | 1 | 0 | 0 | 0 |
| CEP63 | 0.306 | P4_Lumbar_VS_P1_P2_P3_Coeliac_Stellate_Sphenopalatine | 0 | 0 | 0 | 1 | 1 | 0 | 0 | 0 |
| NBAS | 0.306 | P2_P3_P4_Lumbar_Coeliac_VS_P1_Stellate_Sphenopalatine | 0 | 1 | 1 | 1 | 1 | 1 | 0 | 0 |
| MAPK14 | 0.306 | P1_P2_P3_P4_Lumbar_Sphenopalatine_VS_Coeliac_Stellate | 1 | 1 | 1 | 1 | 1 | 0 | 0 | 1 |
| STK38L | 0.306 | P1_P3_P4_Lumbar_Coeliac_Sphenopalatine_VS_P2_Stellate | 1 | 0 | 1 | 1 | 1 | 1 | 0 | 1 |
| KCTD1 | 0.306 | P1_P3_P4_Lumbar_Coeliac_Sphenopalatine_VS_P2_Stellate | 1 | 0 | 1 | 1 | 1 | 1 | 0 | 1 |
| CALHM5 | 0.305 | P2_VS_P1_P3_P4_Lumbar_Coeliac_Stellate_Sphenopalatine | 0 | 1 | 0 | 0 | 0 | 0 | 0 | 0 |
| PEX3 | 0.305 | P3_P4_Lumbar_Coeliac_VS_P1_P2_Stellate_Sphenopalatine | 0 | 0 | 1 | 1 | 1 | 1 | 0 | 0 |
| AHNAK2 | 0.305 | P1_Lumbar_Sphenopalatine_VS_P2_P3_P4_Coeliac_Stellate | 1 | 0 | 0 | 0 | 1 | 0 | 0 | 1 |

| | | | | | | | | | | |
|---|---|---|---|---|---|---|---|---|---|---|
| PRKCE | 0.305 | P1_P2_P3_P4_Lumbar_Sphenopalatine_VS_Coeliac_Stellate | 1 | 1 | 1 | 1 | 1 | 0 | 0 | 1 |
| SENP5 | 0.305 | P2_P3_P4_Lumbar_Coeliac_Sphenopalatine_VS_P1_Stellate | 0 | 1 | 1 | 1 | 1 | 1 | 0 | 1 |
| SPNS2 | 0.304 | P4_Lumbar_VS_P1_P2_P3_Coeliac_Stellate_Sphenopalatine | 0 | 0 | 0 | 1 | 1 | 0 | 0 | 0 |
| VPS8 | 0.304 | P3_P4_Lumbar_Coeliac_VS_P1_P2_Stellate_Sphenopalatine | 0 | 0 | 1 | 1 | 1 | 1 | 0 | 0 |
| CNKSR2 | 0.304 | P3_P4_Lumbar_Coeliac_VS_P1_P2_Stellate_Sphenopalatine | 0 | 0 | 1 | 1 | 1 | 1 | 0 | 0 |
| XPO5 | 0.304 | P2_P3_P4_Lumbar_Coeliac_VS_P1_Stellate_Sphenopalatine | 0 | 1 | 1 | 1 | 1 | 1 | 0 | 0 |
| FOXP4 | 0.304 | Lumbar_Coeliac_Stellate_VS_P1_P2_P3_P4_Sphenopalatine | 0 | 0 | 0 | 0 | 1 | 1 | 1 | 0 |
| OXTR | 0.304 | Sphenopalatine_VS_P1_P2_P3_P4_Lumbar_Coeliac_Stellate | 0 | 0 | 0 | 0 | 0 | 0 | 0 | 1 |
| TMEM189 | 0.304 | P1_P2_P3_P4_Lumbar_Sphenopalatine_VS_Coeliac_Stellate | 1 | 1 | 1 | 1 | 1 | 0 | 0 | 1 |
| LRRN2 | 0.304 | P1_P2_P3_P4_Coeliac_Sphenopalatine_VS_Lumbar_Stellate | 1 | 1 | 1 | 1 | 0 | 1 | 0 | 1 |
| ARHGEF15 | 0.304 | P1_P3_P4_Lumbar_Coeliac_Stellate_Sphenopalatine_VS_P2 | 1 | 0 | 1 | 1 | 1 | 1 | 1 | 1 |
| RAB30 | 0.303 | P1_P2_P3_Lumbar_Coeliac_Sphenopalatine_VS_P4_Stellate | 1 | 1 | 1 | 0 | 1 | 1 | 0 | 1 |
| MFAP2 | 0.303 | P1_P3_P4_Lumbar_Coeliac_Stellate_Sphenopalatine_VS_P2 | 1 | 0 | 1 | 1 | 1 | 1 | 1 | 1 |
| TBC1D10A | 0.302 | P3_P4_VS_P1_P2_Lumbar_Coeliac_Stellate_Sphenopalatine | 0 | 0 | 1 | 1 | 0 | 0 | 0 | 0 |
| DDX41 | 0.302 | P1_P3_Lumbar_Coeliac_VS_P2_P4_Stellate_Sphenopalatine | 1 | 0 | 1 | 0 | 1 | 1 | 0 | 0 |
| NRCAM | 0.302 | P1_P2_P3_P4_Lumbar_Coeliac_Stellate_VS_Sphenopalatine | 1 | 1 | 1 | 1 | 1 | 1 | 1 | 0 |
| FGF7 | 0.302 | Sphenopalatine_VS_P1_P2_P3_P4_Lumbar_Coeliac_Stellate | 0 | 0 | 0 | 0 | 0 | 0 | 0 | 1 |
| B3GALT2 | 0.302 | Lumbar_Coeliac_Sphenopalatine_VS_P1_P2_P3_P4_Stellate | 0 | 0 | 0 | 0 | 1 | 1 | 0 | 1 |
| TRAF4 | 0.302 | P1_P2_P3_Lumbar_Coeliac_Stellate_Sphenopalatine_VS_P4 | 1 | 1 | 1 | 0 | 1 | 1 | 1 | 1 |
| LURAP1L | 0.301 | P3_VS_P1_P2_P4_Lumbar_Coeliac_Stellate_Sphenopalatine | 0 | 0 | 1 | 0 | 0 | 0 | 0 | 0 |
| CEP57 | 0.301 | P1_Lumbar_Coeliac_VS_P2_P3_P4_Stellate_Sphenopalatine | 1 | 0 | 0 | 0 | 1 | 1 | 0 | 0 |
| MKRN3 | 0.301 | P1_P3_Lumbar_Coeliac_VS_P2_P4_Stellate_Sphenopalatine | 1 | 0 | 1 | 0 | 1 | 1 | 0 | 0 |
| SP3 | 0.301 | P2_P3_P4_Lumbar_Coeliac_VS_P1_Stellate_Sphenopalatine | 0 | 1 | 1 | 1 | 1 | 1 | 0 | 0 |
| STBD1 | 0.301 | P4_Lumbar_Coeliac_Stellate_VS_P1_P2_P3_Sphenopalatine | 0 | 0 | 0 | 1 | 1 | 1 | 1 | 0 |
| PMS2 | 0.301 | P1_P3_P4_Lumbar_Coeliac_Sphenopalatine_VS_P2_Stellate | 1 | 0 | 1 | 1 | 1 | 1 | 0 | 1 |
| INSL6 | 0.3 | P3_VS_P1_P2_P4_Lumbar_Coeliac_Stellate_Sphenopalatine | 0 | 0 | 1 | 0 | 0 | 0 | 0 | 0 |
| 2310068J16F | 0.3 | P4_Lumbar_VS_P1_P2_P3_Coeliac_Stellate_Sphenopalatine | 0 | 0 | 0 | 1 | 1 | 0 | 0 | 0 |
| TMPO | 0.3 | P1_Lumbar_Coeliac_VS_P2_P3_P4_Stellate_Sphenopalatine | 1 | 0 | 0 | 0 | 1 | 1 | 0 | 0 |
| CPSF3 | 0.3 | P1_Lumbar_Coeliac_VS_P2_P3_P4_Stellate_Sphenopalatine | 1 | 0 | 0 | 0 | 1 | 1 | 0 | 0 |
| ZBTB21 | 0.3 | P1_Lumbar_Coeliac_VS_P2_P3_P4_Stellate_Sphenopalatine | 1 | 0 | 0 | 0 | 1 | 1 | 0 | 0 |

| | | | | | | | | | | |
|---|---|---|---|---|---|---|---|---|---|---|
| HERPUD2 | 0.3 | P2_P3_P4_Lumbar_Coeliac_VS_P1_Stellate_Sphenopalatine | 0 | 1 | 1 | 1 | 1 | 1 | 0 | 0 |
| GRAMD1B | 0.3 | P2_P3_P4_Lumbar_Coeliac_Sphenopalatine_VS_P1_Stellate | 0 | 1 | 1 | 1 | 1 | 1 | 0 | 1 |
| ST8SIA1 | 0.299 | P2_Lumbar_Coeliac_VS_P1_P3_P4_Stellate_Sphenopalatine | 0 | 1 | 0 | 0 | 1 | 1 | 0 | 0 |
| AHCTF1 | 0.299 | P2_P3_P4_Lumbar_Coeliac_VS_P1_Stellate_Sphenopalatine | 0 | 1 | 1 | 1 | 1 | 1 | 0 | 0 |
| TADA2B | 0.299 | P2_P3_P4_Lumbar_Coeliac_VS_P1_Stellate_Sphenopalatine | 0 | 1 | 1 | 1 | 1 | 1 | 0 | 0 |
| EFHC2 | 0.299 | P1_P2_P3_P4_Lumbar_Coeliac_Stellate_VS_Sphenopalatine | 1 | 1 | 1 | 1 | 1 | 1 | 1 | 0 |
| STAC | 0.299 | P1_P3_P4_Lumbar_Coeliac_Stellate_Sphenopalatine_VS_P2 | 1 | 0 | 1 | 1 | 1 | 1 | 1 | 1 |
| EMP2 | 0.298 | P2_VS_P1_P3_P4_Lumbar_Coeliac_Stellate_Sphenopalatine | 0 | 1 | 0 | 0 | 0 | 0 | 0 | 0 |
| IFT43 | 0.298 | P4_Lumbar_VS_P1_P2_P3_Coeliac_Stellate_Sphenopalatine | 0 | 0 | 0 | 1 | 1 | 0 | 0 | 0 |
| ECI1 | 0.298 | P4_Lumbar_VS_P1_P2_P3_Coeliac_Stellate_Sphenopalatine | 0 | 0 | 0 | 1 | 1 | 0 | 0 | 0 |
| TOE1 | 0.298 | P1_Lumbar_Coeliac_VS_P2_P3_P4_Stellate_Sphenopalatine | 1 | 0 | 0 | 0 | 1 | 1 | 0 | 0 |
| LTN1 | 0.298 | P2_P3_P4_Lumbar_Coeliac_VS_P1_Stellate_Sphenopalatine | 0 | 1 | 1 | 1 | 1 | 1 | 0 | 0 |
| INTS1 | 0.298 | P1_P3_P4_Lumbar_Coeliac_Sphenopalatine_VS_P2_Stellate | 1 | 0 | 1 | 1 | 1 | 1 | 0 | 1 |
| GGCT | 0.297 | P1_P2_P4_VS_P3_Lumbar_Coeliac_Stellate_Sphenopalatine | 1 | 1 | 0 | 1 | 0 | 0 | 0 | 0 |
| BRAT1 | 0.297 | P2_Lumbar_Coeliac_VS_P1_P3_P4_Stellate_Sphenopalatine | 0 | 1 | 0 | 0 | 1 | 1 | 0 | 0 |
| TADA2A | 0.297 | P1_P2_P3_Lumbar_Coeliac_VS_P4_Stellate_Sphenopalatine | 1 | 1 | 1 | 0 | 1 | 1 | 0 | 0 |
| TOR1AIP1 | 0.297 | P2_P3_P4_Lumbar_Coeliac_VS_P1_Stellate_Sphenopalatine | 0 | 1 | 1 | 1 | 1 | 1 | 0 | 0 |
| ECE1 | 0.297 | P2_P3_P4_Lumbar_Coeliac_VS_P1_Stellate_Sphenopalatine | 0 | 1 | 1 | 1 | 1 | 1 | 0 | 0 |
| SNHG9 | 0.297 | P1_P3_P4_Lumbar_Coeliac_Sphenopalatine_VS_P2_Stellate | 1 | 0 | 1 | 1 | 1 | 1 | 0 | 1 |
| SPA17 | 0.296 | P1_P3_P4_Lumbar_Coeliac_Sphenopalatine_VS_P2_Stellate | 1 | 0 | 1 | 1 | 1 | 1 | 0 | 1 |
| USP53 | 0.295 | P4_Lumbar_VS_P1_P2_P3_Coeliac_Stellate_Sphenopalatine | 0 | 0 | 0 | 1 | 1 | 0 | 0 | 0 |
| 4932441J04F | 0.295 | P4_Coeliac_VS_P1_P2_P3_Lumbar_Stellate_Sphenopalatine | 0 | 0 | 0 | 1 | 0 | 1 | 0 | 0 |
| DPF3 | 0.295 | P4_Coeliac_VS_P1_P2_P3_Lumbar_Stellate_Sphenopalatine | 0 | 0 | 0 | 1 | 0 | 1 | 0 | 0 |
| HDAC11 | 0.295 | P1_P2_P3_P4_Lumbar_Sphenopalatine_VS_Coeliac_Stellate | 1 | 1 | 1 | 1 | 1 | 0 | 0 | 1 |
| IRF9 | 0.295 | P1_P4_Lumbar_Coeliac_Sphenopalatine_VS_P2_P3_Stellate | 1 | 0 | 0 | 1 | 1 | 1 | 0 | 1 |
| TRPC4 | 0.294 | P2_P3_VS_P1_P4_Lumbar_Coeliac_Stellate_Sphenopalatine | 0 | 1 | 1 | 0 | 0 | 0 | 0 | 0 |
| IFI30 | 0.294 | P4_Lumbar_VS_P1_P2_P3_Coeliac_Stellate_Sphenopalatine | 0 | 0 | 0 | 1 | 1 | 0 | 0 | 0 |
| TBC1D5 | 0.294 | P4_Lumbar_VS_P1_P2_P3_Coeliac_Stellate_Sphenopalatine | 0 | 0 | 0 | 1 | 1 | 0 | 0 | 0 |
| MIER1 | 0.294 | P1_P2_P3_Lumbar_Coeliac_VS_P4_Stellate_Sphenopalatine | 1 | 1 | 1 | 0 | 1 | 1 | 0 | 0 |
| GFOD1 | 0.294 | P2_P3_P4_Lumbar_Coeliac_VS_P1_Stellate_Sphenopalatine | 0 | 1 | 1 | 1 | 1 | 1 | 0 | 0 |

| | | | | | | | | | | |
|---|---|---|---|---|---|---|---|---|---|---|
| RIMBP2 | 0.294 | P3_P4_Lumbar_Coeliac_Stellate_VS_P1_P2_Sphenopalatine | 0 | 0 | 1 | 1 | 1 | 1 | 1 | 0 |
| POMC | 0.294 | Sphenopalatine_VS_P1_P2_P3_P4_Lumbar_Coeliac_Stellate | 0 | 0 | 0 | 0 | 0 | 0 | 0 | 1 |
| NRBP2 | 0.294 | P1_P2_P3_P4_Lumbar_Sphenopalatine_VS_Coeliac_Stellate | 1 | 1 | 1 | 1 | 1 | 0 | 0 | 1 |
| PCM1 | 0.294 | P1_P3_P4_Lumbar_Coeliac_Sphenopalatine_VS_P2_Stellate | 1 | 0 | 1 | 1 | 1 | 1 | 0 | 1 |
| DIDO1 | 0.294 | P2_P3_P4_Lumbar_Coeliac_Sphenopalatine_VS_P1_Stellate | 0 | 1 | 1 | 1 | 1 | 1 | 0 | 1 |
| METTL3 | 0.293 | P4_Lumbar_VS_P1_P2_P3_Coeliac_Stellate_Sphenopalatine | 0 | 0 | 0 | 1 | 1 | 0 | 0 | 0 |
| LANCL3 | 0.293 | P4_Lumbar_VS_P1_P2_P3_Coeliac_Stellate_Sphenopalatine | 0 | 0 | 0 | 1 | 1 | 0 | 0 | 0 |
| DDX39 | 0.293 | P1_Lumbar_Coeliac_VS_P2_P3_P4_Stellate_Sphenopalatine | 1 | 0 | 0 | 0 | 1 | 1 | 0 | 0 |
| CDO1 | 0.293 | P1_P3_Lumbar_Coeliac_Stellate_VS_P2_P4_Sphenopalatine | 1 | 0 | 1 | 0 | 1 | 1 | 1 | 0 |
| CCDC28A | 0.292 | P4_Lumbar_VS_P1_P2_P3_Coeliac_Stellate_Sphenopalatine | 0 | 0 | 0 | 1 | 1 | 0 | 0 | 0 |
| MOB1B | 0.292 | P1_P2_P3_Lumbar_Coeliac_VS_P4_Stellate_Sphenopalatine | 1 | 1 | 1 | 0 | 1 | 1 | 0 | 0 |
| PCF11 | 0.292 | P2_P3_P4_Lumbar_Coeliac_VS_P1_Stellate_Sphenopalatine | 0 | 1 | 1 | 1 | 1 | 1 | 0 | 0 |
| FRAT2 | 0.292 | P1_P2_P3_P4_Lumbar_Coeliac_Stellate_VS_Sphenopalatine | 1 | 1 | 1 | 1 | 1 | 1 | 1 | 0 |
| SWSAP1 | 0.291 | P3_VS_P1_P2_P4_Lumbar_Coeliac_Stellate_Sphenopalatine | 0 | 0 | 1 | 0 | 0 | 0 | 0 | 0 |
| YPEL4 | 0.291 | P1_P2_P3_P4_Sphenopalatine_VS_Lumbar_Coeliac_Stellate | 1 | 1 | 1 | 1 | 0 | 0 | 0 | 1 |
| 2810468N07 | 0.291 | P1_P2_P3_P4_Sphenopalatine_VS_Lumbar_Coeliac_Stellate | 1 | 1 | 1 | 1 | 0 | 0 | 0 | 1 |
| CSF1 | 0.291 | P1_P2_P3_Lumbar_Coeliac_Sphenopalatine_VS_P4_Stellate | 1 | 1 | 1 | 0 | 1 | 1 | 0 | 1 |
| RNASET2A | 0.29 | P3_VS_P1_P2_P4_Lumbar_Coeliac_Stellate_Sphenopalatine | 0 | 0 | 1 | 0 | 0 | 0 | 0 | 0 |
| TFB1M | 0.29 | P1_P4_VS_P2_P3_Lumbar_Coeliac_Stellate_Sphenopalatine | 1 | 0 | 0 | 1 | 0 | 0 | 0 | 0 |
| ZFP518B | 0.29 | P2_P3_Lumbar_Coeliac_VS_P1_P4_Stellate_Sphenopalatine | 0 | 1 | 1 | 0 | 1 | 1 | 0 | 0 |
| GRHL3 | 0.289 | P1_VS_P2_P3_P4_Lumbar_Coeliac_Stellate_Sphenopalatine | 1 | 0 | 0 | 0 | 0 | 0 | 0 | 0 |
| SENP7 | 0.289 | P1_P3_P4_Lumbar_Coeliac_Sphenopalatine_VS_P2_Stellate | 1 | 0 | 1 | 1 | 1 | 1 | 0 | 1 |
| TAGAP1 | 0.288 | P3_VS_P1_P2_P4_Lumbar_Coeliac_Stellate_Sphenopalatine | 0 | 0 | 1 | 0 | 0 | 0 | 0 | 0 |
| KDM7A | 0.288 | P2_Lumbar_Coeliac_VS_P1_P3_P4_Stellate_Sphenopalatine | 0 | 1 | 0 | 0 | 1 | 1 | 0 | 0 |
| DEPTOR | 0.288 | P1_P3_P4_Lumbar_Coeliac_Stellate_VS_P2_Sphenopalatine | 1 | 0 | 1 | 1 | 1 | 1 | 1 | 0 |
| R74862 | 0.288 | P1_P2_P3_P4_Coeliac_Sphenopalatine_VS_Lumbar_Stellate | 1 | 1 | 1 | 1 | 0 | 1 | 0 | 1 |
| G0S2 | 0.287 | P1_P2_P3_P4_Sphenopalatine_VS_Lumbar_Coeliac_Stellate | 1 | 1 | 1 | 1 | 0 | 0 | 0 | 1 |
| FAM71E1 | 0.287 | P1_P3_P4_Lumbar_Coeliac_Sphenopalatine_VS_P2_Stellate | 1 | 0 | 1 | 1 | 1 | 1 | 0 | 1 |
| OPRL1 | 0.287 | P2_P3_P4_Lumbar_Coeliac_Stellate_Sphenopalatine_VS_P1 | 0 | 1 | 1 | 1 | 1 | 1 | 1 | 1 |
| SPSB4 | 0.286 | P1_P2_VS_P3_P4_Lumbar_Coeliac_Stellate_Sphenopalatine | 1 | 1 | 0 | 0 | 0 | 0 | 0 | 0 |

| | | | | | | | | | | |
|---|---|---|---|---|---|---|---|---|---|---|
| UTP25 | 0.286 | P2_Lumbar_Coeliac_VS_P1_P3_P4_Stellate_Sphenopalatine | 0 | 1 | 0 | 0 | 1 | 1 | 0 | 0 |
| PHRF1 | 0.286 | P1_P2_P3_Lumbar_Coeliac_VS_P4_Stellate_Sphenopalatine | 1 | 1 | 1 | 0 | 1 | 1 | 0 | 0 |
| BST2 | 0.286 | P1_P2_P3_P4_Lumbar_Sphenopalatine_VS_Coeliac_Stellate | 1 | 1 | 1 | 1 | 1 | 0 | 0 | 1 |
| DOCK3 | 0.286 | P1_P2_P3_P4_Lumbar_Sphenopalatine_VS_Coeliac_Stellate | 1 | 1 | 1 | 1 | 1 | 0 | 0 | 1 |
| ADGRB1 | 0.286 | P1_P2_P3_P4_Lumbar_Sphenopalatine_VS_Coeliac_Stellate | 1 | 1 | 1 | 1 | 1 | 0 | 0 | 1 |
| PPP1R18 | 0.286 | P1_P2_P3_Lumbar_Coeliac_Sphenopalatine_VS_P4_Stellate | 1 | 1 | 1 | 0 | 1 | 1 | 0 | 1 |
| DIPK2A | 0.285 | P1_P4_VS_P2_P3_Lumbar_Coeliac_Stellate_Sphenopalatine | 1 | 0 | 0 | 1 | 0 | 0 | 0 | 0 |
| KLF10 | 0.285 | P1_P2_P3_P4_Coeliac_VS_Lumbar_Stellate_Sphenopalatine | 1 | 1 | 1 | 1 | 0 | 1 | 0 | 0 |
| MSI1 | 0.285 | P2_Lumbar_Coeliac_VS_P1_P3_P4_Stellate_Sphenopalatine | 0 | 1 | 0 | 0 | 1 | 1 | 0 | 0 |
| APBA1 | 0.285 | P2_P3_P4_Lumbar_Coeliac_Sphenopalatine_VS_P1_Stellate | 0 | 1 | 1 | 1 | 1 | 1 | 0 | 1 |
| SH3BP5 | 0.285 | P1_P2_P4_Lumbar_Coeliac_Stellate_Sphenopalatine_VS_P3 | 1 | 1 | 0 | 1 | 1 | 1 | 1 | 1 |
| CCP110 | 0.284 | P2_Lumbar_Coeliac_VS_P1_P3_P4_Stellate_Sphenopalatine | 0 | 1 | 0 | 0 | 1 | 1 | 0 | 0 |
| ODF2L | 0.284 | P1_P3_Lumbar_Coeliac_VS_P2_P4_Stellate_Sphenopalatine | 1 | 0 | 1 | 0 | 1 | 1 | 0 | 0 |
| FAHD2A | 0.284 | P1_P2_P3_P4_Lumbar_Coeliac_Stellate_VS_Sphenopalatine | 1 | 1 | 1 | 1 | 1 | 1 | 1 | 0 |
| NECTIN1 | 0.284 | P1_P2_P3_P4_Lumbar_Sphenopalatine_VS_Coeliac_Stellate | 1 | 1 | 1 | 1 | 1 | 0 | 0 | 1 |
| ZFP449 | 0.283 | P4_Lumbar_VS_P1_P2_P3_Coeliac_Stellate_Sphenopalatine | 0 | 0 | 0 | 1 | 1 | 0 | 0 | 0 |
| GM17396 | 0.282 | P3_VS_P1_P2_P4_Lumbar_Coeliac_Stellate_Sphenopalatine | 0 | 0 | 1 | 0 | 0 | 0 | 0 | 0 |
| A730063M1[illegible] | 0.282 | P3_VS_P1_P2_P4_Lumbar_Coeliac_Stellate_Sphenopalatine | 0 | 0 | 1 | 0 | 0 | 0 | 0 | 0 |
| MYPOP | 0.282 | P1_P2_P3_P4_Sphenopalatine_VS_Lumbar_Coeliac_Stellate | 1 | 1 | 1 | 1 | 0 | 0 | 0 | 1 |
| DLGAP1 | 0.282 | P1_P2_P3_P4_Coeliac_Sphenopalatine_VS_Lumbar_Stellate | 1 | 1 | 1 | 1 | 0 | 1 | 0 | 1 |
| NOL6 | 0.281 | P2_P3_P4_Lumbar_Coeliac_Sphenopalatine_VS_P1_Stellate | 0 | 1 | 1 | 1 | 1 | 1 | 0 | 1 |
| PRR12 | 0.281 | P2_P3_P4_Lumbar_Coeliac_Sphenopalatine_VS_P1_Stellate | 0 | 1 | 1 | 1 | 1 | 1 | 0 | 1 |
| SCARB1 | 0.28 | P3_VS_P1_P2_P4_Lumbar_Coeliac_Stellate_Sphenopalatine | 0 | 0 | 1 | 0 | 0 | 0 | 0 | 0 |
| SCAF4 | 0.28 | P2_Lumbar_Coeliac_VS_P1_P3_P4_Stellate_Sphenopalatine | 0 | 1 | 0 | 0 | 1 | 1 | 0 | 0 |
| CLGN | 0.28 | P1_P2_Lumbar_Coeliac_VS_P3_P4_Stellate_Sphenopalatine | 1 | 1 | 0 | 0 | 1 | 1 | 0 | 0 |
| RNF149 | 0.28 | P1_P2_P3_P4_Lumbar_Coeliac_Stellate_VS_Sphenopalatine | 1 | 1 | 1 | 1 | 1 | 1 | 1 | 0 |
| GFRA4 | 0.28 | P1_P2_P4_Sphenopalatine_VS_P3_Lumbar_Coeliac_Stellate | 1 | 1 | 0 | 1 | 0 | 0 | 0 | 1 |
| ELL2 | 0.279 | P2_VS_P1_P3_P4_Lumbar_Coeliac_Stellate_Sphenopalatine | 0 | 1 | 0 | 0 | 0 | 0 | 0 | 0 |
| ALDOC | 0.279 | P1_P2_P3_P4_VS_Lumbar_Coeliac_Stellate_Sphenopalatine | 1 | 1 | 1 | 1 | 0 | 0 | 0 | 0 |
| SREBF1 | 0.279 | P4_Lumbar_VS_P1_P2_P3_Coeliac_Stellate_Sphenopalatine | 0 | 0 | 0 | 1 | 1 | 0 | 0 | 0 |

| | | | | | | | | | | |
|---|---|---|---|---|---|---|---|---|---|---|
| PHLDB1 | 0.279 | P2_Lumbar_Coeliac_VS_P1_P3_P4_Stellate_Sphenopalatine | 0 | 1 | 0 | 0 | 1 | 1 | 0 | 0 |
| ZFP106 | 0.279 | P2_P3_P4_Lumbar_Coeliac_Sphenopalatine_VS_P1_Stellate | 0 | 1 | 1 | 1 | 1 | 1 | 0 | 1 |
| DDX58 | 0.278 | P1_VS_P2_P3_P4_Lumbar_Coeliac_Stellate_Sphenopalatine | 1 | 0 | 0 | 0 | 0 | 0 | 0 | 0 |
| CENPC1 | 0.278 | P4_Lumbar_VS_P1_P2_P3_Coeliac_Stellate_Sphenopalatine | 0 | 0 | 0 | 1 | 1 | 0 | 0 | 0 |
| PIKFYVE | 0.278 | P2_Lumbar_Coeliac_VS_P1_P3_P4_Stellate_Sphenopalatine | 0 | 1 | 0 | 0 | 1 | 1 | 0 | 0 |
| EPHB2 | 0.278 | P2_Lumbar_Coeliac_VS_P1_P3_P4_Stellate_Sphenopalatine | 0 | 1 | 0 | 0 | 1 | 1 | 0 | 0 |
| PLATR25 | 0.278 | P2_Lumbar_Coeliac_VS_P1_P3_P4_Stellate_Sphenopalatine | 0 | 1 | 0 | 0 | 1 | 1 | 0 | 0 |
| SESN1 | 0.277 | P4_Coeliac_VS_P1_P2_P3_Lumbar_Stellate_Sphenopalatine | 0 | 0 | 0 | 1 | 0 | 1 | 0 | 0 |
| HMBOX1 | 0.277 | P1_P2_P4_Lumbar_Sphenopalatine_VS_P3_Coeliac_Stellate | 1 | 1 | 0 | 1 | 1 | 0 | 0 | 1 |
| SGPP2 | 0.277 | P1_P2_P3_P4_Lumbar_Stellate_Sphenopalatine_VS_Coeliac | 1 | 1 | 1 | 1 | 1 | 0 | 1 | 1 |
| TTC39B | 0.276 | P1_P4_VS_P2_P3_Lumbar_Coeliac_Stellate_Sphenopalatine | 1 | 0 | 0 | 1 | 0 | 0 | 0 | 0 |
| JOSD2 | 0.276 | P1_P2_P3_P4_Sphenopalatine_VS_Lumbar_Coeliac_Stellate | 1 | 1 | 1 | 1 | 0 | 0 | 0 | 1 |
| GNG4 | 0.276 | P1_P2_P3_P4_Sphenopalatine_VS_Lumbar_Coeliac_Stellate | 1 | 1 | 1 | 1 | 0 | 0 | 0 | 1 |
| SLC1A4 | 0.276 | P1_P2_P4_Lumbar_Sphenopalatine_VS_P3_Coeliac_Stellate | 1 | 1 | 0 | 1 | 1 | 0 | 0 | 1 |
| sept-08 | 0.276 | P1_P2_P3_P4_Coeliac_Sphenopalatine_VS_Lumbar_Stellate | 1 | 1 | 1 | 1 | 0 | 1 | 0 | 1 |
| BAALC | 0.276 | P2_P3_Lumbar_Coeliac_Stellate_Sphenopalatine_VS_P1_P4 | 0 | 1 | 1 | 0 | 1 | 1 | 1 | 1 |
| GCAT | 0.275 | P3_Lumbar_VS_P1_P2_P4_Coeliac_Stellate_Sphenopalatine | 0 | 0 | 1 | 0 | 1 | 0 | 0 | 0 |
| RCCD1 | 0.275 | P4_Coeliac_VS_P1_P2_P3_Lumbar_Stellate_Sphenopalatine | 0 | 0 | 0 | 1 | 0 | 1 | 0 | 0 |
| MCMBP | 0.275 | P1_P3_Lumbar_Coeliac_VS_P2_P4_Stellate_Sphenopalatine | 1 | 0 | 1 | 0 | 1 | 1 | 0 | 0 |
| ADGRA2 | 0.274 | P1_VS_P2_P3_P4_Lumbar_Coeliac_Stellate_Sphenopalatine | 1 | 0 | 0 | 0 | 0 | 0 | 0 | 0 |
| CABLES2 | 0.274 | P2_VS_P1_P3_P4_Lumbar_Coeliac_Stellate_Sphenopalatine | 0 | 1 | 0 | 0 | 0 | 0 | 0 | 0 |
| MED17 | 0.274 | P4_Lumbar_VS_P1_P2_P3_Coeliac_Stellate_Sphenopalatine | 0 | 0 | 0 | 1 | 1 | 0 | 0 | 0 |
| AC241534.1 | 0.274 | P4_Lumbar_VS_P1_P2_P3_Coeliac_Stellate_Sphenopalatine | 0 | 0 | 0 | 1 | 1 | 0 | 0 | 0 |
| VPS13C | 0.274 | P2_Lumbar_Coeliac_VS_P1_P3_P4_Stellate_Sphenopalatine | 0 | 1 | 0 | 0 | 1 | 1 | 0 | 0 |
| B020010K11I | 0.274 | Sphenopalatine_VS_P1_P2_P3_P4_Lumbar_Coeliac_Stellate | 0 | 0 | 0 | 0 | 0 | 0 | 0 | 1 |
| TMEM178B | 0.274 | P1_P2_P3_Lumbar_Coeliac_Stellate_Sphenopalatine_VS_P4 | 1 | 1 | 1 | 0 | 1 | 1 | 1 | 1 |
| GM42851 | 0.273 | Lumbar_Coeliac_Stellate_VS_P1_P2_P3_P4_Sphenopalatine | 0 | 0 | 0 | 0 | 1 | 1 | 1 | 0 |
| IMMP2L | 0.273 | P1_P2_P3_P4_Lumbar_Coeliac_Stellate_VS_Sphenopalatine | 1 | 1 | 1 | 1 | 1 | 1 | 1 | 0 |
| FMN2 | 0.273 | P2_P3_P4_Lumbar_Coeliac_Sphenopalatine_VS_P1_Stellate | 0 | 1 | 1 | 1 | 1 | 1 | 0 | 1 |
| LAG3 | 0.273 | P1_P2_P3_Lumbar_Coeliac_Stellate_Sphenopalatine_VS_P4 | 1 | 1 | 1 | 0 | 1 | 1 | 1 | 1 |

| | | | | | | | | | | |
|---|---|---|---|---|---|---|---|---|---|---|
| MCCC2 | 0.272 | P1_P4_Lumbar_VS_P2_P3_Coeliac_Stellate_Sphenopalatine | 1 | 0 | 0 | 1 | 1 | 0 | 0 | 0 |
| OSBPL11 | 0.272 | P2_Lumbar_Coeliac_VS_P1_P3_P4_Stellate_Sphenopalatine | 0 | 1 | 0 | 0 | 1 | 1 | 0 | 0 |
| ZBTB33 | 0.272 | P2_P3_Lumbar_Coeliac_VS_P1_P4_Stellate_Sphenopalatine | 0 | 1 | 1 | 0 | 1 | 1 | 0 | 0 |
| TRAF3 | 0.272 | P1_P2_P3_P4_Lumbar_Sphenopalatine_VS_Coeliac_Stellate | 1 | 1 | 1 | 1 | 1 | 0 | 0 | 1 |
| DOP1A | 0.272 | P1_P3_P4_Lumbar_Coeliac_Sphenopalatine_VS_P2_Stellate | 1 | 0 | 1 | 1 | 1 | 1 | 0 | 1 |
| WASF1 | 0.272 | P1_P2_P4_Lumbar_Coeliac_Stellate_Sphenopalatine_VS_P3 | 1 | 1 | 0 | 1 | 1 | 1 | 1 | 1 |
| ANK | 0.271 | P1_P2_P3_P4_Coeliac_Sphenopalatine_VS_Lumbar_Stellate | 1 | 1 | 1 | 1 | 0 | 1 | 0 | 1 |
| PYGO1 | 0.271 | P1_P2_P3_Lumbar_Coeliac_Sphenopalatine_VS_P4_Stellate | 1 | 1 | 1 | 0 | 1 | 1 | 0 | 1 |
| KMT5B | 0.27 | P2_Lumbar_Coeliac_VS_P1_P3_P4_Stellate_Sphenopalatine | 0 | 1 | 0 | 0 | 1 | 1 | 0 | 0 |
| EHD3 | 0.27 | P1_P2_P3_P4_Lumbar_Sphenopalatine_VS_Coeliac_Stellate | 1 | 1 | 1 | 1 | 1 | 0 | 0 | 1 |
| WIPI1 | 0.269 | P1_P2_P3_P4_Lumbar_VS_Coeliac_Stellate_Sphenopalatine | 1 | 1 | 1 | 1 | 1 | 0 | 0 | 0 |
| SEC22C | 0.269 | P4_Lumbar_Coeliac_Sphenopalatine_VS_P1_P2_P3_Stellate | 0 | 0 | 0 | 1 | 1 | 1 | 0 | 1 |
| CITED2 | 0.269 | P1_P3_P4_Lumbar_Coeliac_Sphenopalatine_VS_P2_Stellate | 1 | 0 | 1 | 1 | 1 | 1 | 0 | 1 |
| YEATS2 | 0.269 | P2_P3_P4_Lumbar_Coeliac_Sphenopalatine_VS_P1_Stellate | 0 | 1 | 1 | 1 | 1 | 1 | 0 | 1 |
| GM12258 | 0.268 | P2_Lumbar_Coeliac_VS_P1_P3_P4_Stellate_Sphenopalatine | 0 | 1 | 0 | 0 | 1 | 1 | 0 | 0 |
| RAB33A | 0.268 | Lumbar_Coeliac_Stellate_VS_P1_P2_P3_P4_Sphenopalatine | 0 | 0 | 0 | 0 | 1 | 1 | 1 | 0 |
| AMD1 | 0.268 | P1_P3_P4_Lumbar_Coeliac_Sphenopalatine_VS_P2_Stellate | 1 | 0 | 1 | 1 | 1 | 1 | 0 | 1 |
| NOL11 | 0.268 | P1_P3_P4_Lumbar_Coeliac_Sphenopalatine_VS_P2_Stellate | 1 | 0 | 1 | 1 | 1 | 1 | 0 | 1 |
| FCHSD2 | 0.268 | P2_P3_P4_Lumbar_Coeliac_Sphenopalatine_VS_P1_Stellate | 0 | 1 | 1 | 1 | 1 | 1 | 0 | 1 |
| INTS11 | 0.267 | P3_Lumbar_VS_P1_P2_P4_Coeliac_Stellate_Sphenopalatine | 0 | 0 | 1 | 0 | 1 | 0 | 0 | 0 |
| MXI1 | 0.267 | P2_Lumbar_Coeliac_VS_P1_P3_P4_Stellate_Sphenopalatine | 0 | 1 | 0 | 0 | 1 | 1 | 0 | 0 |
| SLF2 | 0.267 | P2_P3_P4_Lumbar_Coeliac_Sphenopalatine_VS_P1_Stellate | 0 | 1 | 1 | 1 | 1 | 1 | 0 | 1 |
| AGRN | 0.266 | P2_Lumbar_Coeliac_VS_P1_P3_P4_Stellate_Sphenopalatine | 0 | 1 | 0 | 0 | 1 | 1 | 0 | 0 |
| PUS1 | 0.266 | P1_P3_Lumbar_Coeliac_VS_P2_P4_Stellate_Sphenopalatine | 1 | 0 | 1 | 0 | 1 | 1 | 0 | 0 |
| OSBPL10 | 0.266 | P2_P3_Lumbar_Coeliac_VS_P1_P4_Stellate_Sphenopalatine | 0 | 1 | 1 | 0 | 1 | 1 | 0 | 0 |
| TDRD3 | 0.266 | P1_P2_P3_Lumbar_Coeliac_VS_P4_Stellate_Sphenopalatine | 1 | 1 | 1 | 0 | 1 | 1 | 0 | 0 |
| GSDME | 0.266 | P1_P3_P4_Lumbar_Coeliac_Sphenopalatine_VS_P2_Stellate | 1 | 0 | 1 | 1 | 1 | 1 | 0 | 1 |
| AKR1B8 | 0.266 | P1_P2_P3_Lumbar_Coeliac_Stellate_Sphenopalatine_VS_P4 | 1 | 1 | 1 | 0 | 1 | 1 | 1 | 1 |
| ZFP81 | 0.265 | P2_Lumbar_Coeliac_VS_P1_P3_P4_Stellate_Sphenopalatine | 0 | 1 | 0 | 0 | 1 | 1 | 0 | 0 |
| FAM172A | 0.265 | P1_P2_P3_Lumbar_Coeliac_VS_P4_Stellate_Sphenopalatine | 1 | 1 | 1 | 0 | 1 | 1 | 0 | 0 |

| | | | | | | | | | | |
|---|---|---|---|---|---|---|---|---|---|---|
| NEK7 | 0.265 | P1_P2_P3_P4_Lumbar_Sphenopalatine_VS_Coeliac_Stellate | 1 | 1 | 1 | 1 | 1 | 0 | 0 | 1 |
| CD2AP | 0.265 | P2_P3_P4_Lumbar_Coeliac_Sphenopalatine_VS_P1_Stellate | 0 | 1 | 1 | 1 | 1 | 1 | 0 | 1 |
| ORAI3 | 0.264 | P1_P4_VS_P2_P3_Lumbar_Coeliac_Stellate_Sphenopalatine | 1 | 0 | 0 | 1 | 0 | 0 | 0 | 0 |
| PRPF4 | 0.264 | P4_Coeliac_VS_P1_P2_P3_Lumbar_Stellate_Sphenopalatine | 0 | 0 | 0 | 1 | 0 | 1 | 0 | 0 |
| 6430590A07I | 0.264 | P2_Lumbar_Coeliac_VS_P1_P3_P4_Stellate_Sphenopalatine | 0 | 1 | 0 | 0 | 1 | 1 | 0 | 0 |
| BBX | 0.264 | P1_P2_P3_P4_Lumbar_Coeliac_Stellate_VS_Sphenopalatine | 1 | 1 | 1 | 1 | 1 | 1 | 1 | 0 |
| SLC7A8 | 0.264 | P1_P2_P3_P4_Lumbar_Sphenopalatine_VS_Coeliac_Stellate | 1 | 1 | 1 | 1 | 1 | 0 | 0 | 1 |
| PXK | 0.264 | P1_P3_P4_Lumbar_Coeliac_Sphenopalatine_VS_P2_Stellate | 1 | 0 | 1 | 1 | 1 | 1 | 0 | 1 |
| N4BP2OS | 0.263 | P3_VS_P1_P2_P4_Lumbar_Coeliac_Stellate_Sphenopalatine | 0 | 0 | 1 | 0 | 0 | 0 | 0 | 0 |
| PLSCR1 | 0.263 | P1_P2_Coeliac_Sphenopalatine_VS_P3_P4_Lumbar_Stellate | 1 | 1 | 0 | 0 | 0 | 1 | 0 | 1 |
| FSTL5 | 0.263 | P1_P3_P4_Lumbar_Coeliac_Sphenopalatine_VS_P2_Stellate | 1 | 0 | 1 | 1 | 1 | 1 | 0 | 1 |
| LBR | 0.262 | P4_Coeliac_VS_P1_P2_P3_Lumbar_Stellate_Sphenopalatine | 0 | 0 | 0 | 1 | 0 | 1 | 0 | 0 |
| SF3A3 | 0.262 | P1_P3_Lumbar_Coeliac_VS_P2_P4_Stellate_Sphenopalatine | 1 | 0 | 1 | 0 | 1 | 1 | 0 | 0 |
| ZFP553 | 0.262 | P1_P3_Lumbar_Coeliac_VS_P2_P4_Stellate_Sphenopalatine | 1 | 0 | 1 | 0 | 1 | 1 | 0 | 0 |
| SGMS1 | 0.262 | P2_P3_Lumbar_Coeliac_VS_P1_P4_Stellate_Sphenopalatine | 0 | 1 | 1 | 0 | 1 | 1 | 0 | 0 |
| TRPT1 | 0.262 | P1_P2_P3_P4_Sphenopalatine_VS_Lumbar_Coeliac_Stellate | 1 | 1 | 1 | 1 | 0 | 0 | 0 | 1 |
| CAMTA2 | 0.262 | P1_P2_P3_P4_Lumbar_Sphenopalatine_VS_Coeliac_Stellate | 1 | 1 | 1 | 1 | 1 | 0 | 0 | 1 |
| PKP4 | 0.262 | P1_P2_P3_P4_Coeliac_Sphenopalatine_VS_Lumbar_Stellate | 1 | 1 | 1 | 1 | 0 | 1 | 0 | 1 |
| AHNAK | 0.262 | P1_P4_Lumbar_Coeliac_Stellate_Sphenopalatine_VS_P2_P3 | 1 | 0 | 0 | 1 | 1 | 1 | 1 | 1 |
| ZFP280B | 0.261 | P2_VS_P1_P3_P4_Lumbar_Coeliac_Stellate_Sphenopalatine | 0 | 1 | 0 | 0 | 0 | 0 | 0 | 0 |
| 6330418K02I | 0.261 | P3_VS_P1_P2_P4_Lumbar_Coeliac_Stellate_Sphenopalatine | 0 | 0 | 1 | 0 | 0 | 0 | 0 | 0 |
| GM10575 | 0.261 | P3_VS_P1_P2_P4_Lumbar_Coeliac_Stellate_Sphenopalatine | 0 | 0 | 1 | 0 | 0 | 0 | 0 | 0 |
| PHKB | 0.261 | P4_Coeliac_VS_P1_P2_P3_Lumbar_Stellate_Sphenopalatine | 0 | 0 | 0 | 1 | 0 | 1 | 0 | 0 |
| SLC7A6 | 0.261 | P2_Lumbar_Coeliac_VS_P1_P3_P4_Stellate_Sphenopalatine | 0 | 1 | 0 | 0 | 1 | 1 | 0 | 0 |
| CHML | 0.261 | P1_P2_P3_Lumbar_Coeliac_Sphenopalatine_VS_P4_Stellate | 1 | 1 | 1 | 0 | 1 | 1 | 0 | 1 |
| AAR2 | 0.26 | P2_P3_Lumbar_Coeliac_VS_P1_P4_Stellate_Sphenopalatine | 0 | 1 | 1 | 0 | 1 | 1 | 0 | 0 |
| CD1D1 | 0.26 | P1_P3_P4_Lumbar_Coeliac_Sphenopalatine_VS_P2_Stellate | 1 | 0 | 1 | 1 | 1 | 1 | 0 | 1 |
| CRTC3 | 0.26 | P1_P3_P4_Lumbar_Coeliac_Sphenopalatine_VS_P2_Stellate | 1 | 0 | 1 | 1 | 1 | 1 | 0 | 1 |
| ID3 | 0.259 | P3_VS_P1_P2_P4_Lumbar_Coeliac_Stellate_Sphenopalatine | 0 | 0 | 1 | 0 | 0 | 0 | 0 | 0 |
| UFSP1 | 0.259 | P1_P2_P3_P4_Lumbar_Sphenopalatine_VS_Coeliac_Stellate | 1 | 1 | 1 | 1 | 1 | 0 | 0 | 1 |

| | | | | | | | | | | |
|---|---|---|---|---|---|---|---|---|---|---|
| NIPAL3 | 0.259 | P1_P2_P3_P4_Coeliac_Sphenopalatine_VS_Lumbar_Stellate | 1 | 1 | 1 | 1 | 0 | 1 | 0 | 1 |
| FAM129B | 0.259 | P1_P2_P3_Lumbar_Coeliac_Sphenopalatine_VS_P4_Stellate | 1 | 1 | 1 | 0 | 1 | 1 | 0 | 1 |
| PDCD4 | 0.259 | P1_P3_P4_Lumbar_Coeliac_Sphenopalatine_VS_P2_Stellate | 1 | 0 | 1 | 1 | 1 | 1 | 0 | 1 |
| RICTOR | 0.258 | P2_Lumbar_Coeliac_VS_P1_P3_P4_Stellate_Sphenopalatine | 0 | 1 | 0 | 0 | 1 | 1 | 0 | 0 |
| PCDH10 | 0.258 | Lumbar_Sphenopalatine_VS_P1_P2_P3_P4_Coeliac_Stellate | 0 | 0 | 0 | 0 | 1 | 0 | 0 | 1 |
| MSH2 | 0.258 | P1_P3_P4_Lumbar_Coeliac_Sphenopalatine_VS_P2_Stellate | 1 | 0 | 1 | 1 | 1 | 1 | 0 | 1 |
| RGS2 | 0.257 | P1_VS_P2_P3_P4_Lumbar_Coeliac_Stellate_Sphenopalatine | 1 | 0 | 0 | 0 | 0 | 0 | 0 | 0 |
| SLC31A2 | 0.257 | P3_Lumbar_VS_P1_P2_P4_Coeliac_Stellate_Sphenopalatine | 0 | 0 | 1 | 0 | 1 | 0 | 0 | 0 |
| GRWD1 | 0.257 | P1_P3_Lumbar_Coeliac_VS_P2_P4_Stellate_Sphenopalatine | 1 | 0 | 1 | 0 | 1 | 1 | 0 | 0 |
| MTRF1 | 0.257 | P1_P3_Lumbar_Coeliac_VS_P2_P4_Stellate_Sphenopalatine | 1 | 0 | 1 | 0 | 1 | 1 | 0 | 0 |
| XRCC6 | 0.257 | P1_P3_Lumbar_Coeliac_VS_P2_P4_Stellate_Sphenopalatine | 1 | 0 | 1 | 0 | 1 | 1 | 0 | 0 |
| SHF | 0.257 | P1_P2_P3_P4_Lumbar_Coeliac_Stellate_VS_Sphenopalatine | 1 | 1 | 1 | 1 | 1 | 1 | 1 | 0 |
| A930005H10 | 0.257 | P1_P2_P3_P4_Sphenopalatine_VS_Lumbar_Coeliac_Stellate | 1 | 1 | 1 | 1 | 0 | 0 | 0 | 1 |
| NDRG1 | 0.257 | P1_P2_P3_P4_Coeliac_Sphenopalatine_VS_Lumbar_Stellate | 1 | 1 | 1 | 1 | 0 | 1 | 0 | 1 |
| ZFP954 | 0.257 | P1_P2_P3_Lumbar_Coeliac_Sphenopalatine_VS_P4_Stellate | 1 | 1 | 1 | 0 | 1 | 1 | 0 | 1 |
| RPF1 | 0.256 | P4_Coeliac_VS_P1_P2_P3_Lumbar_Stellate_Sphenopalatine | 0 | 0 | 0 | 1 | 0 | 1 | 0 | 0 |
| SLAIN1 | 0.256 | P2_Lumbar_Coeliac_VS_P1_P3_P4_Stellate_Sphenopalatine | 0 | 1 | 0 | 0 | 1 | 1 | 0 | 0 |
| ALYREF | 0.256 | P1_P2_P3_P4_Lumbar_Coeliac_Stellate_VS_Sphenopalatine | 1 | 1 | 1 | 1 | 1 | 1 | 1 | 0 |
| KRT1 | 0.256 | Sphenopalatine_VS_P1_P2_P3_P4_Lumbar_Coeliac_Stellate | 0 | 0 | 0 | 0 | 0 | 0 | 0 | 1 |
| LRFN1 | 0.256 | P1_P2_P3_P4_Lumbar_Sphenopalatine_VS_Coeliac_Stellate | 1 | 1 | 1 | 1 | 1 | 0 | 0 | 1 |
| GPR137 | 0.256 | P1_P2_P3_P4_Lumbar_Sphenopalatine_VS_Coeliac_Stellate | 1 | 1 | 1 | 1 | 1 | 0 | 0 | 1 |
| KLHL23 | 0.256 | Lumbar_Coeliac_Sphenopalatine_VS_P1_P2_P3_P4_Stellate | 0 | 0 | 0 | 0 | 1 | 1 | 0 | 1 |
| GPR45 | 0.256 | P1_P3_P4_Lumbar_Coeliac_Sphenopalatine_VS_P2_Stellate | 1 | 0 | 1 | 1 | 1 | 1 | 0 | 1 |
| RPP40 | 0.256 | P1_P3_P4_Lumbar_Coeliac_Sphenopalatine_VS_P2_Stellate | 1 | 0 | 1 | 1 | 1 | 1 | 0 | 1 |
| ULK1 | 0.256 | P2_P3_P4_Lumbar_Coeliac_Sphenopalatine_VS_P1_Stellate | 0 | 1 | 1 | 1 | 1 | 1 | 0 | 1 |
| SMPD5 | 0.256 | P1_P3_P4_Lumbar_Coeliac_Stellate_Sphenopalatine_VS_P2 | 1 | 0 | 1 | 1 | 1 | 1 | 1 | 1 |
| PPCS | 0.255 | P1_P2_P4_Lumbar_VS_P3_Coeliac_Stellate_Sphenopalatine | 1 | 1 | 0 | 1 | 1 | 0 | 0 | 0 |
| AARS2 | 0.255 | P2_Lumbar_Coeliac_VS_P1_P3_P4_Stellate_Sphenopalatine | 0 | 1 | 0 | 0 | 1 | 1 | 0 | 0 |
| 1190005I06R | 0.255 | P1_P3_Lumbar_Coeliac_Sphenopalatine_VS_P2_P4_Stellate | 1 | 0 | 1 | 0 | 1 | 1 | 0 | 1 |
| 9930021J03F | 0.255 | P1_P3_P4_Lumbar_Coeliac_Sphenopalatine_VS_P2_Stellate | 1 | 0 | 1 | 1 | 1 | 1 | 0 | 1 |

| | | | | | | | | | | |
|---|---|---|---|---|---|---|---|---|---|---|
| DSTYK | 0.255 | P2_P3_P4_Lumbar_Coeliac_Sphenopalatine_VS_P1_Stellate | 0 | 1 | 1 | 1 | 1 | 1 | 0 | 1 |
| ETHE1 | 0.255 | P1_P3_P4_Lumbar_Coeliac_Stellate_Sphenopalatine_VS_P2 | 1 | 0 | 1 | 1 | 1 | 1 | 1 | 1 |
| THSD7A | 0.254 | P3_Lumbar_VS_P1_P2_P4_Coeliac_Stellate_Sphenopalatine | 0 | 0 | 1 | 0 | 1 | 0 | 0 | 0 |
| LTO1 | 0.254 | P2_Lumbar_Coeliac_VS_P1_P3_P4_Stellate_Sphenopalatine | 0 | 1 | 0 | 0 | 1 | 1 | 0 | 0 |
| ADCY6 | 0.254 | P2_Lumbar_Coeliac_VS_P1_P3_P4_Stellate_Sphenopalatine | 0 | 1 | 0 | 0 | 1 | 1 | 0 | 0 |
| PELI1 | 0.254 | P1_P2_P3_P4_Lumbar_Coeliac_Stellate_VS_Sphenopalatine | 1 | 1 | 1 | 1 | 1 | 1 | 1 | 0 |
| MIRG | 0.254 | P1_P2_P3_P4_Lumbar_Sphenopalatine_VS_Coeliac_Stellate | 1 | 1 | 1 | 1 | 1 | 0 | 0 | 1 |
| BCL2L13 | 0.254 | P1_P2_P3_Lumbar_Coeliac_Sphenopalatine_VS_P4_Stellate | 1 | 1 | 1 | 0 | 1 | 1 | 0 | 1 |
| SCHIP1 | 0.254 | P1_P2_P3_P4_Lumbar_Stellate_Sphenopalatine_VS_Coeliac | 1 | 1 | 1 | 1 | 1 | 0 | 1 | 1 |
| GAST | 0.253 | P3_P4_Coeliac_VS_P1_P2_Lumbar_Stellate_Sphenopalatine | 0 | 0 | 1 | 1 | 0 | 1 | 0 | 0 |
| SHKBP1 | 0.253 | P1_P3_Lumbar_Coeliac_VS_P2_P4_Stellate_Sphenopalatine | 1 | 0 | 1 | 0 | 1 | 1 | 0 | 0 |
| NTN3 | 0.252 | P1_P4_VS_P2_P3_Lumbar_Coeliac_Stellate_Sphenopalatine | 1 | 0 | 0 | 1 | 0 | 0 | 0 | 0 |
| SPATA24 | 0.252 | P1_P3_P4_Lumbar_VS_P2_Coeliac_Stellate_Sphenopalatine | 1 | 0 | 1 | 1 | 1 | 0 | 0 | 0 |
| TMSB15B1 | 0.252 | P1_P2_P3_P4_Coeliac_Sphenopalatine_VS_Lumbar_Stellate | 1 | 1 | 1 | 1 | 0 | 1 | 0 | 1 |
| FAM131B | 0.252 | P1_P2_P3_Lumbar_Coeliac_Sphenopalatine_VS_P4_Stellate | 1 | 1 | 1 | 0 | 1 | 1 | 0 | 1 |
| PCNX2 | 0.252 | P1_P2_P3_Lumbar_Coeliac_Sphenopalatine_VS_P4_Stellate | 1 | 1 | 1 | 0 | 1 | 1 | 0 | 1 |
| SLC12A6 | 0.252 | P2_P3_P4_Lumbar_Coeliac_Sphenopalatine_VS_P1_Stellate | 0 | 1 | 1 | 1 | 1 | 1 | 0 | 1 |
| ZFP738 | 0.252 | P2_P3_P4_Lumbar_Coeliac_Sphenopalatine_VS_P1_Stellate | 0 | 1 | 1 | 1 | 1 | 1 | 0 | 1 |
| RPS6KA3 | 0.251 | P4_Coeliac_VS_P1_P2_P3_Lumbar_Stellate_Sphenopalatine | 0 | 0 | 0 | 1 | 0 | 1 | 0 | 0 |
| SEC63 | 0.251 | P2_Lumbar_Coeliac_VS_P1_P3_P4_Stellate_Sphenopalatine | 0 | 1 | 0 | 0 | 1 | 1 | 0 | 0 |
| DENND1B | 0.251 | P1_P3_Lumbar_Coeliac_VS_P2_P4_Stellate_Sphenopalatine | 1 | 0 | 1 | 0 | 1 | 1 | 0 | 0 |
| KLHL2 | 0.251 | P1_P2_P3_Lumbar_Coeliac_Sphenopalatine_VS_P4_Stellate | 1 | 1 | 1 | 0 | 1 | 1 | 0 | 1 |
| HSF2 | 0.251 | P1_P2_P3_Lumbar_Coeliac_Sphenopalatine_VS_P4_Stellate | 1 | 1 | 1 | 0 | 1 | 1 | 0 | 1 |
| HAT1 | 0.251 | P1_P3_P4_Lumbar_Coeliac_Sphenopalatine_VS_P2_Stellate | 1 | 0 | 1 | 1 | 1 | 1 | 0 | 1 |
| POR | 0.251 | P1_P3_P4_Lumbar_Coeliac_Sphenopalatine_VS_P2_Stellate | 1 | 0 | 1 | 1 | 1 | 1 | 0 | 1 |
| FOXJ3 | 0.251 | P2_P3_P4_Lumbar_Coeliac_Sphenopalatine_VS_P1_Stellate | 0 | 1 | 1 | 1 | 1 | 1 | 0 | 1 |
| CERK | 0.251 | P2_P3_P4_Lumbar_Coeliac_Sphenopalatine_VS_P1_Stellate | 0 | 1 | 1 | 1 | 1 | 1 | 0 | 1 |
| DYNC2H1 | 0.25 | P1_P3_Lumbar_Coeliac_VS_P2_P4_Stellate_Sphenopalatine | 1 | 0 | 1 | 0 | 1 | 1 | 0 | 0 |
| CDK2AP1 | 0.25 | P1_P2_P3_Lumbar_Coeliac_VS_P4_Stellate_Sphenopalatine | 1 | 1 | 1 | 0 | 1 | 1 | 0 | 0 |
| CNST | 0.25 | P1_P2_P4_Lumbar_Coeliac_Stellate_VS_P3_Sphenopalatine | 1 | 1 | 0 | 1 | 1 | 1 | 1 | 0 |

| | | | | | | | | | | |
|---|---|---|---|---|---|---|---|---|---|---|
| CD55 | 0.25 | P3_P4_Lumbar_Coeliac_Sphenopalatine_VS_P1_P2_Stellate | 0 | 0 | 1 | 1 | 1 | 1 | 0 | 1 |
| NOL8 | 0.25 | P1_P3_P4_Lumbar_Coeliac_Sphenopalatine_VS_P2_Stellate | 1 | 0 | 1 | 1 | 1 | 1 | 0 | 1 |
| SFMBT2 | 0.249 | P1_VS_P2_P3_P4_Lumbar_Coeliac_Stellate_Sphenopalatine | 1 | 0 | 0 | 0 | 0 | 0 | 0 | 0 |
| MOV10 | 0.249 | P1_P4_VS_P2_P3_Lumbar_Coeliac_Stellate_Sphenopalatine | 1 | 0 | 0 | 1 | 0 | 0 | 0 | 0 |
| 0610009L18I | 0.249 | P3_P4_VS_P1_P2_Lumbar_Coeliac_Stellate_Sphenopalatine | 0 | 0 | 1 | 1 | 0 | 0 | 0 | 0 |
| PDE9A | 0.249 | P1_P2_P3_Lumbar_Coeliac_VS_P4_Stellate_Sphenopalatine | 1 | 1 | 1 | 0 | 1 | 1 | 0 | 0 |
| DPY19L4 | 0.249 | P2_P3_P4_Lumbar_Coeliac_Sphenopalatine_VS_P1_Stellate | 0 | 1 | 1 | 1 | 1 | 1 | 0 | 1 |
| ANKRD6 | 0.249 | P3_P4_Lumbar_Coeliac_Stellate_Sphenopalatine_VS_P1_P2 | 0 | 0 | 1 | 1 | 1 | 1 | 1 | 1 |
| CPNE6 | 0.248 | P1_P2_VS_P3_P4_Lumbar_Coeliac_Stellate_Sphenopalatine | 1 | 1 | 0 | 0 | 0 | 0 | 0 | 0 |
| PTPRR | 0.248 | P1_P2_P3_P4_Lumbar_Coeliac_Stellate_VS_Sphenopalatine | 1 | 1 | 1 | 1 | 1 | 1 | 1 | 0 |
| PAQR9 | 0.248 | P1_Sphenopalatine_VS_P2_P3_P4_Lumbar_Coeliac_Stellate | 1 | 0 | 0 | 0 | 0 | 0 | 0 | 1 |
| STXBP5 | 0.248 | P1_P2_P3_P4_Lumbar_Sphenopalatine_VS_Coeliac_Stellate | 1 | 1 | 1 | 1 | 1 | 0 | 0 | 1 |
| ZRSR1 | 0.248 | P1_P2_P3_P4_Coeliac_Sphenopalatine_VS_Lumbar_Stellate | 1 | 1 | 1 | 1 | 0 | 1 | 0 | 1 |
| ZFP90 | 0.248 | P1_P3_P4_Lumbar_Coeliac_Sphenopalatine_VS_P2_Stellate | 1 | 0 | 1 | 1 | 1 | 1 | 0 | 1 |
| FLII | 0.248 | P1_P3_P4_Lumbar_Coeliac_Sphenopalatine_VS_P2_Stellate | 1 | 0 | 1 | 1 | 1 | 1 | 0 | 1 |
| ZFP948 | 0.248 | P1_P3_P4_Lumbar_Coeliac_Sphenopalatine_VS_P2_Stellate | 1 | 0 | 1 | 1 | 1 | 1 | 0 | 1 |
| PHC2 | 0.248 | P2_P3_P4_Lumbar_Coeliac_Sphenopalatine_VS_P1_Stellate | 0 | 1 | 1 | 1 | 1 | 1 | 0 | 1 |
| ZFP335OS | 0.248 | P1_P2_P3_Lumbar_Coeliac_Stellate_Sphenopalatine_VS_P4 | 1 | 1 | 1 | 0 | 1 | 1 | 1 | 1 |
| PLPPR4 | 0.248 | P1_P3_P4_Lumbar_Coeliac_Stellate_Sphenopalatine_VS_P2 | 1 | 0 | 1 | 1 | 1 | 1 | 1 | 1 |
| CRPPA | 0.247 | P4_Coeliac_VS_P1_P2_P3_Lumbar_Stellate_Sphenopalatine | 0 | 0 | 0 | 1 | 0 | 1 | 0 | 0 |
| TECPR1 | 0.247 | P1_P2_P3_P4_Lumbar_Sphenopalatine_VS_Coeliac_Stellate | 1 | 1 | 1 | 1 | 1 | 0 | 0 | 1 |
| ACTA1 | 0.247 | P1_P3_P4_Lumbar_Coeliac_Sphenopalatine_VS_P2_Stellate | 1 | 0 | 1 | 1 | 1 | 1 | 0 | 1 |
| ZCCHC9 | 0.247 | P1_P3_P4_Lumbar_Coeliac_Sphenopalatine_VS_P2_Stellate | 1 | 0 | 1 | 1 | 1 | 1 | 0 | 1 |
| ZMYM4 | 0.247 | P2_P3_P4_Lumbar_Coeliac_Sphenopalatine_VS_P1_Stellate | 0 | 1 | 1 | 1 | 1 | 1 | 0 | 1 |
| EGFLAM | 0.246 | P1_VS_P2_P3_P4_Lumbar_Coeliac_Stellate_Sphenopalatine | 1 | 0 | 0 | 0 | 0 | 0 | 0 | 0 |
| GM8098 | 0.246 | P3_VS_P1_P2_P4_Lumbar_Coeliac_Stellate_Sphenopalatine | 0 | 0 | 1 | 0 | 0 | 0 | 0 | 0 |
| KCTD15 | 0.246 | P1_P3_Lumbar_Coeliac_VS_P2_P4_Stellate_Sphenopalatine | 1 | 0 | 1 | 0 | 1 | 1 | 0 | 0 |
| B4GALT5 | 0.246 | P1_P2_P4_Lumbar_Sphenopalatine_VS_P3_Coeliac_Stellate | 1 | 1 | 0 | 1 | 1 | 0 | 0 | 1 |
| PDK1 | 0.246 | P1_P2_P3_P4_Lumbar_Sphenopalatine_VS_Coeliac_Stellate | 1 | 1 | 1 | 1 | 1 | 0 | 0 | 1 |
| POLR2B | 0.246 | P1_P3_P4_Lumbar_Coeliac_Sphenopalatine_VS_P2_Stellate | 1 | 0 | 1 | 1 | 1 | 1 | 0 | 1 |

| | | | | | | | | | | |
|---|---|---|---|---|---|---|---|---|---|---|
| MARS | 0.246 | P1_P3_P4_Lumbar_Coeliac_Sphenopalatine_VS_P2_Stellate | 1 | 0 | 1 | 1 | 1 | 1 | 0 | 1 |
| ZER1 | 0.246 | P2_P3_P4_Lumbar_Coeliac_Sphenopalatine_VS_P1_Stellate | 0 | 1 | 1 | 1 | 1 | 1 | 0 | 1 |
| RRAS | 0.245 | P1_P2_P4_Lumbar_VS_P3_Coeliac_Stellate_Sphenopalatine | 1 | 1 | 0 | 1 | 1 | 0 | 0 | 0 |
| CYP4X1 | 0.245 | P1_P2_P3_P4_Lumbar_Sphenopalatine_VS_Coeliac_Stellate | 1 | 1 | 1 | 1 | 1 | 0 | 0 | 1 |
| WDR54 | 0.245 | P1_P3_P4_Lumbar_Coeliac_Sphenopalatine_VS_P2_Stellate | 1 | 0 | 1 | 1 | 1 | 1 | 0 | 1 |
| PPP1R13B | 0.245 | P1_P3_P4_Lumbar_Coeliac_Sphenopalatine_VS_P2_Stellate | 1 | 0 | 1 | 1 | 1 | 1 | 0 | 1 |
| TAF1 | 0.245 | P2_P3_P4_Lumbar_Coeliac_Sphenopalatine_VS_P1_Stellate | 0 | 1 | 1 | 1 | 1 | 1 | 0 | 1 |
| GM34030 | 0.245 | P1_P3_Lumbar_Coeliac_Stellate_Sphenopalatine_VS_P2_P4 | 1 | 0 | 1 | 0 | 1 | 1 | 1 | 1 |
| LRRFIP1 | 0.245 | P1_P2_P4_Lumbar_Coeliac_Stellate_Sphenopalatine_VS_P3 | 1 | 1 | 0 | 1 | 1 | 1 | 1 | 1 |
| SLC10A7 | 0.244 | P1_P2_P4_Lumbar_VS_P3_Coeliac_Stellate_Sphenopalatine | 1 | 1 | 0 | 1 | 1 | 0 | 0 | 0 |
| GCLC | 0.244 | P1_P2_P3_P4_Lumbar_Coeliac_Stellate_VS_Sphenopalatine | 1 | 1 | 1 | 1 | 1 | 1 | 1 | 0 |
| ATP2B4 | 0.244 | P1_P2_P3_P4_Coeliac_Sphenopalatine_VS_Lumbar_Stellate | 1 | 1 | 1 | 1 | 0 | 1 | 0 | 1 |
| CSPP1 | 0.244 | Lumbar_Coeliac_Sphenopalatine_VS_P1_P2_P3_P4_Stellate | 0 | 0 | 0 | 0 | 1 | 1 | 0 | 1 |
| GM11837 | 0.244 | P2_P3_P4_Lumbar_Coeliac_Sphenopalatine_VS_P1_Stellate | 0 | 1 | 1 | 1 | 1 | 1 | 0 | 1 |
| ABLIM3 | 0.244 | P1_P2_P3_P4_Lumbar_Stellate_Sphenopalatine_VS_Coeliac | 1 | 1 | 1 | 1 | 1 | 0 | 1 | 1 |
| CPPED1 | 0.243 | P1_P2_P3_P4_Lumbar_VS_Coeliac_Stellate_Sphenopalatine | 1 | 1 | 1 | 1 | 1 | 0 | 0 | 0 |
| UPF1 | 0.243 | P2_P3_P4_Lumbar_Coeliac_Sphenopalatine_VS_P1_Stellate | 0 | 1 | 1 | 1 | 1 | 1 | 0 | 1 |
| WDR48 | 0.243 | P2_P3_P4_Lumbar_Coeliac_Sphenopalatine_VS_P1_Stellate | 0 | 1 | 1 | 1 | 1 | 1 | 0 | 1 |
| VAMP8 | 0.242 | P3_P4_VS_P1_P2_Lumbar_Coeliac_Stellate_Sphenopalatine | 0 | 0 | 1 | 1 | 0 | 0 | 0 | 0 |
| KDM2A | 0.242 | P2_P3_P4_Lumbar_Coeliac_Sphenopalatine_VS_P1_Stellate | 0 | 1 | 1 | 1 | 1 | 1 | 0 | 1 |
| 9930104L06F | 0.241 | P4_Coeliac_VS_P1_P2_P3_Lumbar_Stellate_Sphenopalatine | 0 | 0 | 0 | 1 | 0 | 1 | 0 | 0 |
| CTNNBL1 | 0.241 | P1_P3_Lumbar_Coeliac_VS_P2_P4_Stellate_Sphenopalatine | 1 | 0 | 1 | 0 | 1 | 1 | 0 | 0 |
| IQSEC3 | 0.241 | P1_P2_P3_P4_Lumbar_Sphenopalatine_VS_Coeliac_Stellate | 1 | 1 | 1 | 1 | 1 | 0 | 0 | 1 |
| NEFH | 0.241 | P1_P2_P3_P4_Lumbar_Sphenopalatine_VS_Coeliac_Stellate | 1 | 1 | 1 | 1 | 1 | 0 | 0 | 1 |
| LAMC1 | 0.241 | P1_P2_P3_P4_Coeliac_Sphenopalatine_VS_Lumbar_Stellate | 1 | 1 | 1 | 1 | 0 | 1 | 0 | 1 |
| ADCY3 | 0.241 | P1_P2_P3_P4_Coeliac_Sphenopalatine_VS_Lumbar_Stellate | 1 | 1 | 1 | 1 | 0 | 1 | 0 | 1 |
| PAF1 | 0.241 | P1_P3_P4_Lumbar_Coeliac_Sphenopalatine_VS_P2_Stellate | 1 | 0 | 1 | 1 | 1 | 1 | 0 | 1 |
| NCOA6 | 0.241 | P2_P3_P4_Lumbar_Coeliac_Sphenopalatine_VS_P1_Stellate | 0 | 1 | 1 | 1 | 1 | 1 | 0 | 1 |
| ARID1B | 0.241 | P2_P3_P4_Lumbar_Coeliac_Sphenopalatine_VS_P1_Stellate | 0 | 1 | 1 | 1 | 1 | 1 | 0 | 1 |
| PLEKHH2 | 0.24 | P2_P3_VS_P1_P4_Lumbar_Coeliac_Stellate_Sphenopalatine | 0 | 1 | 1 | 0 | 0 | 0 | 0 | 0 |

| | | | | | | | | | | |
|---|---|---|---|---|---|---|---|---|---|---|
| MBTPS1 | 0.24 | P4_Coeliac_VS_P1_P2_P3_Lumbar_Stellate_Sphenopalatine | 0 | 0 | 0 | 1 | 0 | 1 | 0 | 0 |
| MIS12 | 0.24 | P1_P3_Lumbar_Coeliac_VS_P2_P4_Stellate_Sphenopalatine | 1 | 0 | 1 | 0 | 1 | 1 | 0 | 0 |
| HIPK2 | 0.24 | P1_P2_P3_P4_Lumbar_Coeliac_Stellate_VS_Sphenopalatine | 1 | 1 | 1 | 1 | 1 | 1 | 1 | 0 |
| ABLIM2 | 0.24 | P2_P4_Sphenopalatine_VS_P1_P3_Lumbar_Coeliac_Stellate | 0 | 1 | 0 | 1 | 0 | 0 | 0 | 1 |
| RAB3GAP2 | 0.24 | P1_P2_P3_P4_Lumbar_Sphenopalatine_VS_Coeliac_Stellate | 1 | 1 | 1 | 1 | 1 | 0 | 0 | 1 |
| FRMPD1 | 0.24 | P1_P2_P3_P4_Lumbar_Sphenopalatine_VS_Coeliac_Stellate | 1 | 1 | 1 | 1 | 1 | 0 | 0 | 1 |
| REV1 | 0.24 | P2_P3_P4_Lumbar_Coeliac_Sphenopalatine_VS_P1_Stellate | 0 | 1 | 1 | 1 | 1 | 1 | 0 | 1 |
| EGFL7 | 0.239 | P3_VS_P1_P2_P4_Lumbar_Coeliac_Stellate_Sphenopalatine | 0 | 0 | 1 | 0 | 0 | 0 | 0 | 0 |
| 2510046G10 | 0.239 | P3_VS_P1_P2_P4_Lumbar_Coeliac_Stellate_Sphenopalatine | 0 | 0 | 1 | 0 | 0 | 0 | 0 | 0 |
| SEC14L1 | 0.239 | P1_P2_P3_Lumbar_Coeliac_Sphenopalatine_VS_P4_Stellate | 1 | 1 | 1 | 0 | 1 | 1 | 0 | 1 |
| BICRAL | 0.239 | P2_P3_P4_Lumbar_Coeliac_Sphenopalatine_VS_P1_Stellate | 0 | 1 | 1 | 1 | 1 | 1 | 0 | 1 |
| SMYD3 | 0.239 | P1_P3_P4_Lumbar_Coeliac_Stellate_Sphenopalatine_VS_P2 | 1 | 0 | 1 | 1 | 1 | 1 | 1 | 1 |
| PSMB8 | 0.238 | P1_VS_P2_P3_P4_Lumbar_Coeliac_Stellate_Sphenopalatine | 1 | 0 | 0 | 0 | 0 | 0 | 0 | 0 |
| SDSL | 0.238 | P2_P3_VS_P1_P4_Lumbar_Coeliac_Stellate_Sphenopalatine | 0 | 1 | 1 | 0 | 0 | 0 | 0 | 0 |
| STX3 | 0.238 | P2_P3_P4_VS_P1_Lumbar_Coeliac_Stellate_Sphenopalatine | 0 | 1 | 1 | 1 | 0 | 0 | 0 | 0 |
| FNIP2 | 0.238 | P4_Coeliac_VS_P1_P2_P3_Lumbar_Stellate_Sphenopalatine | 0 | 0 | 0 | 1 | 0 | 1 | 0 | 0 |
| APPL2 | 0.238 | P4_Coeliac_VS_P1_P2_P3_Lumbar_Stellate_Sphenopalatine | 0 | 0 | 0 | 1 | 0 | 1 | 0 | 0 |
| GEMIN5 | 0.238 | P1_P3_Lumbar_Coeliac_VS_P2_P4_Stellate_Sphenopalatine | 1 | 0 | 1 | 0 | 1 | 1 | 0 | 0 |
| PDGFA | 0.238 | P4_Sphenopalatine_VS_P1_P2_P3_Lumbar_Coeliac_Stellate | 0 | 0 | 0 | 1 | 0 | 0 | 0 | 1 |
| CTDSPL | 0.238 | P1_P3_P4_Lumbar_Coeliac_Sphenopalatine_VS_P2_Stellate | 1 | 0 | 1 | 1 | 1 | 1 | 0 | 1 |
| RELB | 0.237 | P1_VS_P2_P3_P4_Lumbar_Coeliac_Stellate_Sphenopalatine | 1 | 0 | 0 | 0 | 0 | 0 | 0 | 0 |
| CAND2 | 0.237 | P3_VS_P1_P2_P4_Lumbar_Coeliac_Stellate_Sphenopalatine | 0 | 0 | 1 | 0 | 0 | 0 | 0 | 0 |
| PPP1R14C | 0.237 | P2_P4_Lumbar_VS_P1_P3_Coeliac_Stellate_Sphenopalatine | 0 | 1 | 0 | 1 | 1 | 0 | 0 | 0 |
| FBXW7 | 0.237 | P1_P2_P3_P4_Lumbar_Sphenopalatine_VS_Coeliac_Stellate | 1 | 1 | 1 | 1 | 1 | 0 | 0 | 1 |
| RNF217 | 0.237 | P1_P2_P3_Lumbar_Coeliac_Sphenopalatine_VS_P4_Stellate | 1 | 1 | 1 | 0 | 1 | 1 | 0 | 1 |
| PHC1 | 0.237 | P1_P3_P4_Lumbar_Coeliac_Sphenopalatine_VS_P2_Stellate | 1 | 0 | 1 | 1 | 1 | 1 | 0 | 1 |
| MIGA1 | 0.237 | P2_P3_P4_Lumbar_Coeliac_Sphenopalatine_VS_P1_Stellate | 0 | 1 | 1 | 1 | 1 | 1 | 0 | 1 |
| TMX3 | 0.237 | P2_P3_P4_Lumbar_Coeliac_Sphenopalatine_VS_P1_Stellate | 0 | 1 | 1 | 1 | 1 | 1 | 0 | 1 |
| TRNP1 | 0.236 | P1_P2_P3_P4_Lumbar_VS_Coeliac_Stellate_Sphenopalatine | 1 | 1 | 1 | 1 | 1 | 0 | 0 | 0 |
| SLC35D1 | 0.236 | P4_Coeliac_VS_P1_P2_P3_Lumbar_Stellate_Sphenopalatine | 0 | 0 | 0 | 1 | 0 | 1 | 0 | 0 |

| | | | | | | | | | | |
|---|---|---|---|---|---|---|---|---|---|---|
| VPS13A | 0.236 | P2_P3_Lumbar_Coeliac_VS_P1_P4_Stellate_Sphenopalatine | 0 | 1 | 1 | 0 | 1 | 1 | 0 | 0 |
| SPRY2 | 0.236 | P1_P4_Lumbar_Coeliac_Stellate_VS_P2_P3_Sphenopalatine | 1 | 0 | 0 | 1 | 1 | 1 | 1 | 0 |
| 9530077C05I | 0.236 | P1_P2_P3_P4_Lumbar_Coeliac_Stellate_VS_Sphenopalatine | 1 | 1 | 1 | 1 | 1 | 1 | 1 | 0 |
| ZFP780B | 0.235 | P1_P2_P3_Lumbar_Coeliac_VS_P4_Stellate_Sphenopalatine | 1 | 1 | 1 | 0 | 1 | 1 | 0 | 0 |
| DLG3 | 0.235 | P1_P3_P4_Lumbar_Coeliac_Sphenopalatine_VS_P2_Stellate | 1 | 0 | 1 | 1 | 1 | 1 | 0 | 1 |
| PNMAL1 | 0.235 | P2_P3_P4_Lumbar_Coeliac_Sphenopalatine_VS_P1_Stellate | 0 | 1 | 1 | 1 | 1 | 1 | 0 | 1 |
| ATXN2L | 0.235 | P2_P3_P4_Lumbar_Coeliac_Sphenopalatine_VS_P1_Stellate | 0 | 1 | 1 | 1 | 1 | 1 | 0 | 1 |
| RASD1 | 0.234 | Lumbar_Coeliac_Sphenopalatine_VS_P1_P2_P3_P4_Stellate | 0 | 0 | 0 | 0 | 1 | 1 | 0 | 1 |
| RTL5 | 0.234 | P3_P4_Lumbar_Coeliac_Sphenopalatine_VS_P1_P2_Stellate | 0 | 0 | 1 | 1 | 1 | 1 | 0 | 1 |
| CMSS1 | 0.234 | P1_P3_P4_Lumbar_Coeliac_Sphenopalatine_VS_P2_Stellate | 1 | 0 | 1 | 1 | 1 | 1 | 0 | 1 |
| SMC1A | 0.234 | P1_P3_P4_Lumbar_Coeliac_Sphenopalatine_VS_P2_Stellate | 1 | 0 | 1 | 1 | 1 | 1 | 0 | 1 |
| CTR9 | 0.234 | P2_P3_P4_Lumbar_Coeliac_Sphenopalatine_VS_P1_Stellate | 0 | 1 | 1 | 1 | 1 | 1 | 0 | 1 |
| GDF11 | 0.234 | P1_P2_P3_Lumbar_Coeliac_Stellate_Sphenopalatine_VS_P4 | 1 | 1 | 1 | 0 | 1 | 1 | 1 | 1 |
| KIZ | 0.233 | P4_Coeliac_VS_P1_P2_P3_Lumbar_Stellate_Sphenopalatine | 0 | 0 | 0 | 1 | 0 | 1 | 0 | 0 |
| UEVLD | 0.233 | P4_Coeliac_VS_P1_P2_P3_Lumbar_Stellate_Sphenopalatine | 0 | 0 | 0 | 1 | 0 | 1 | 0 | 0 |
| ACTN1 | 0.233 | P1_P4_Lumbar_Coeliac_Stellate_VS_P2_P3_Sphenopalatine | 1 | 0 | 0 | 1 | 1 | 1 | 1 | 0 |
| BC029722 | 0.233 | P1_P2_P3_P4_Sphenopalatine_VS_Lumbar_Coeliac_Stellate | 1 | 1 | 1 | 1 | 0 | 0 | 0 | 1 |
| SCN2B | 0.233 | P1_P2_P3_P4_Sphenopalatine_VS_Lumbar_Coeliac_Stellate | 1 | 1 | 1 | 1 | 0 | 0 | 0 | 1 |
| D230025D16 | 0.233 | P1_P2_P3_Lumbar_Coeliac_Sphenopalatine_VS_P4_Stellate | 1 | 1 | 1 | 0 | 1 | 1 | 0 | 1 |
| TARS | 0.233 | P1_P4_Lumbar_Coeliac_Sphenopalatine_VS_P2_P3_Stellate | 1 | 0 | 0 | 1 | 1 | 1 | 0 | 1 |
| DNAJC3 | 0.233 | P1_P3_P4_Lumbar_Coeliac_Sphenopalatine_VS_P2_Stellate | 1 | 0 | 1 | 1 | 1 | 1 | 0 | 1 |
| FIZ1 | 0.233 | P1_P2_P3_Lumbar_Coeliac_Stellate_Sphenopalatine_VS_P4 | 1 | 1 | 1 | 0 | 1 | 1 | 1 | 1 |
| PTPN5 | 0.233 | P1_P2_P3_Lumbar_Coeliac_Stellate_Sphenopalatine_VS_P4 | 1 | 1 | 1 | 0 | 1 | 1 | 1 | 1 |
| CCDC157 | 0.232 | P3_P4_Coeliac_VS_P1_P2_Lumbar_Stellate_Sphenopalatine | 0 | 0 | 1 | 1 | 0 | 1 | 0 | 0 |
| STX17 | 0.232 | P1_P2_P3_Lumbar_Coeliac_VS_P4_Stellate_Sphenopalatine | 1 | 1 | 1 | 0 | 1 | 1 | 0 | 0 |
| ICMT | 0.232 | P1_P2_P3_P4_Lumbar_Sphenopalatine_VS_Coeliac_Stellate | 1 | 1 | 1 | 1 | 1 | 0 | 0 | 1 |
| CHPF2 | 0.232 | P1_P2_P3_P4_Lumbar_Sphenopalatine_VS_Coeliac_Stellate | 1 | 1 | 1 | 1 | 1 | 0 | 0 | 1 |
| ZFP523 | 0.232 | P1_P2_P3_Lumbar_Coeliac_Sphenopalatine_VS_P4_Stellate | 1 | 1 | 1 | 0 | 1 | 1 | 0 | 1 |
| CXXC5 | 0.232 | P1_P2_P3_Lumbar_Coeliac_Sphenopalatine_VS_P4_Stellate | 1 | 1 | 1 | 0 | 1 | 1 | 0 | 1 |
| LRFN4 | 0.232 | P1_P2_P3_Lumbar_Coeliac_Sphenopalatine_VS_P4_Stellate | 1 | 1 | 1 | 0 | 1 | 1 | 0 | 1 |

| | | | | | | | | | | |
|---|---|---|---|---|---|---|---|---|---|---|
| MYSM1 | 0.232 | P1_P3_P4_Lumbar_Coeliac_Sphenopalatine_VS_P2_Stellate | 1 | 0 | 1 | 1 | 1 | 1 | 0 | 1 |
| BTBD8 | 0.231 | P1_P2_P3_Lumbar_Coeliac_VS_P4_Stellate_Sphenopalatine | 1 | 1 | 1 | 0 | 1 | 1 | 0 | 0 |
| CCL25 | 0.231 | P1_P2_P3_Lumbar_Coeliac_VS_P4_Stellate_Sphenopalatine | 1 | 1 | 1 | 0 | 1 | 1 | 0 | 0 |
| COL4A1 | 0.231 | P4_Sphenopalatine_VS_P1_P2_P3_Lumbar_Coeliac_Stellate | 0 | 0 | 0 | 1 | 0 | 0 | 0 | 1 |
| EXOSC10 | 0.231 | P1_P3_P4_Lumbar_Coeliac_Sphenopalatine_VS_P2_Stellate | 1 | 0 | 1 | 1 | 1 | 1 | 0 | 1 |
| FRS2 | 0.231 | P1_P3_P4_Lumbar_Coeliac_Sphenopalatine_VS_P2_Stellate | 1 | 0 | 1 | 1 | 1 | 1 | 0 | 1 |
| FAM193A | 0.231 | P2_P3_P4_Lumbar_Coeliac_Sphenopalatine_VS_P1_Stellate | 0 | 1 | 1 | 1 | 1 | 1 | 0 | 1 |
| COL4A3BP | 0.231 | P2_P3_P4_Lumbar_Coeliac_Sphenopalatine_VS_P1_Stellate | 0 | 1 | 1 | 1 | 1 | 1 | 0 | 1 |
| ARPC1B | 0.231 | P1_P3_P4_Lumbar_Coeliac_Stellate_Sphenopalatine_VS_P2 | 1 | 0 | 1 | 1 | 1 | 1 | 1 | 1 |
| RFX2 | 0.23 | P3_VS_P1_P2_P4_Lumbar_Coeliac_Stellate_Sphenopalatine | 0 | 0 | 1 | 0 | 0 | 0 | 0 | 0 |
| SPRED3 | 0.23 | P1_P2_P3_Lumbar_Coeliac_VS_P4_Stellate_Sphenopalatine | 1 | 1 | 1 | 0 | 1 | 1 | 0 | 0 |
| BCAS3 | 0.23 | P1_P2_P3_P4_Lumbar_Coeliac_Stellate_VS_Sphenopalatine | 1 | 1 | 1 | 1 | 1 | 1 | 1 | 0 |
| 5430416N02 | 0.23 | P1_P2_P3_Lumbar_Coeliac_Sphenopalatine_VS_P4_Stellate | 1 | 1 | 1 | 0 | 1 | 1 | 0 | 1 |
| NCOA3 | 0.23 | P2_P3_P4_Lumbar_Coeliac_Sphenopalatine_VS_P1_Stellate | 0 | 1 | 1 | 1 | 1 | 1 | 0 | 1 |
| NUMA1 | 0.23 | P2_P3_P4_Lumbar_Coeliac_Sphenopalatine_VS_P1_Stellate | 0 | 1 | 1 | 1 | 1 | 1 | 0 | 1 |
| TLE4 | 0.23 | P1_P2_P4_Lumbar_Coeliac_Stellate_Sphenopalatine_VS_P3 | 1 | 1 | 0 | 1 | 1 | 1 | 1 | 1 |
| METTL27 | 0.229 | P1_P2_P3_P4_VS_Lumbar_Coeliac_Stellate_Sphenopalatine | 1 | 1 | 1 | 1 | 0 | 0 | 0 | 0 |
| CDC34 | 0.229 | P1_P2_P3_P4_Lumbar_Sphenopalatine_VS_Coeliac_Stellate | 1 | 1 | 1 | 1 | 1 | 0 | 0 | 1 |
| TTC7B | 0.229 | P1_P2_P3_P4_Lumbar_Sphenopalatine_VS_Coeliac_Stellate | 1 | 1 | 1 | 1 | 1 | 0 | 0 | 1 |
| ARHGAP39 | 0.229 | P1_P2_P3_P4_Lumbar_Sphenopalatine_VS_Coeliac_Stellate | 1 | 1 | 1 | 1 | 1 | 0 | 0 | 1 |
| GTF2IRD1 | 0.229 | P2_P3_P4_Lumbar_Coeliac_Sphenopalatine_VS_P1_Stellate | 0 | 1 | 1 | 1 | 1 | 1 | 0 | 1 |
| CYLD | 0.229 | P2_P3_P4_Lumbar_Coeliac_Sphenopalatine_VS_P1_Stellate | 0 | 1 | 1 | 1 | 1 | 1 | 0 | 1 |
| 1700088E04I | 0.228 | P3_VS_P1_P2_P4_Lumbar_Coeliac_Stellate_Sphenopalatine | 0 | 0 | 1 | 0 | 0 | 0 | 0 | 0 |
| RILPL1 | 0.228 | P1_P2_P4_Lumbar_VS_P3_Coeliac_Stellate_Sphenopalatine | 1 | 1 | 0 | 1 | 1 | 0 | 0 | 0 |
| CD109 | 0.228 | Sphenopalatine_VS_P1_P2_P3_P4_Lumbar_Coeliac_Stellate | 0 | 0 | 0 | 0 | 0 | 0 | 0 | 1 |
| SLC35E1 | 0.228 | P1_P2_P3_P4_Lumbar_Sphenopalatine_VS_Coeliac_Stellate | 1 | 1 | 1 | 1 | 1 | 0 | 0 | 1 |
| ZFP629 | 0.228 | P1_P2_P3_Lumbar_Coeliac_Sphenopalatine_VS_P4_Stellate | 1 | 1 | 1 | 0 | 1 | 1 | 0 | 1 |
| LENG8 | 0.228 | P2_P4_Lumbar_Coeliac_Sphenopalatine_VS_P1_P3_Stellate | 0 | 1 | 0 | 1 | 1 | 1 | 0 | 1 |
| SPECC1L | 0.228 | P2_P4_Lumbar_Coeliac_Sphenopalatine_VS_P1_P3_Stellate | 0 | 1 | 0 | 1 | 1 | 1 | 0 | 1 |
| SLC39A10 | 0.228 | P2_P3_P4_Lumbar_Coeliac_Sphenopalatine_VS_P1_Stellate | 0 | 1 | 1 | 1 | 1 | 1 | 0 | 1 |

| | | | | | | | | | | |
|---|---|---|---|---|---|---|---|---|---|---|
| FAM234B | 0.227 | P2_P4_Lumbar_VS_P1_P3_Coeliac_Stellate_Sphenopalatine | 0 | 1 | 0 | 1 | 1 | 0 | 0 | 0 |
| ANKRD16 | 0.227 | P1_P2_Lumbar_Coeliac_VS_P3_P4_Stellate_Sphenopalatine | 1 | 1 | 0 | 0 | 1 | 1 | 0 | 0 |
| OSBPL8 | 0.227 | P1_P3_Lumbar_Coeliac_VS_P2_P4_Stellate_Sphenopalatine | 1 | 0 | 1 | 0 | 1 | 1 | 0 | 0 |
| GPR153 | 0.227 | P1_P2_P3_P4_Lumbar_Sphenopalatine_VS_Coeliac_Stellate | 1 | 1 | 1 | 1 | 1 | 0 | 0 | 1 |
| CWF19L2 | 0.227 | P1_P3_P4_Lumbar_Coeliac_Sphenopalatine_VS_P2_Stellate | 1 | 0 | 1 | 1 | 1 | 1 | 0 | 1 |
| FIG4 | 0.227 | P1_P3_P4_Lumbar_Coeliac_Sphenopalatine_VS_P2_Stellate | 1 | 0 | 1 | 1 | 1 | 1 | 0 | 1 |
| DENND1A | 0.227 | P2_P3_P4_Lumbar_Coeliac_Sphenopalatine_VS_P1_Stellate | 0 | 1 | 1 | 1 | 1 | 1 | 0 | 1 |
| SNRNP200 | 0.227 | P2_P3_P4_Lumbar_Coeliac_Sphenopalatine_VS_P1_Stellate | 0 | 1 | 1 | 1 | 1 | 1 | 0 | 1 |
| PKD2L1 | 0.226 | P1_VS_P2_P3_P4_Lumbar_Coeliac_Stellate_Sphenopalatine | 1 | 0 | 0 | 0 | 0 | 0 | 0 | 0 |
| GM14305 | 0.226 | P1_P2_P3_P4_VS_Lumbar_Coeliac_Stellate_Sphenopalatine | 1 | 1 | 1 | 1 | 0 | 0 | 0 | 0 |
| RNASEH2B | 0.226 | Lumbar_Coeliac_Stellate_VS_P1_P2_P3_P4_Sphenopalatine | 0 | 0 | 0 | 0 | 1 | 1 | 1 | 0 |
| BCL2 | 0.226 | P1_P2_P3_P4_Coeliac_Sphenopalatine_VS_Lumbar_Stellate | 1 | 1 | 1 | 1 | 0 | 1 | 0 | 1 |
| UGCG | 0.226 | P1_P3_P4_Lumbar_Coeliac_Sphenopalatine_VS_P2_Stellate | 1 | 0 | 1 | 1 | 1 | 1 | 0 | 1 |
| PRMT9 | 0.226 | P1_P3_P4_Lumbar_Coeliac_Sphenopalatine_VS_P2_Stellate | 1 | 0 | 1 | 1 | 1 | 1 | 0 | 1 |
| DDX21 | 0.226 | P1_P3_P4_Lumbar_Coeliac_Sphenopalatine_VS_P2_Stellate | 1 | 0 | 1 | 1 | 1 | 1 | 0 | 1 |
| MEF2A | 0.226 | P2_P3_P4_Lumbar_Coeliac_Sphenopalatine_VS_P1_Stellate | 0 | 1 | 1 | 1 | 1 | 1 | 0 | 1 |
| EDC4 | 0.226 | P2_P3_P4_Lumbar_Coeliac_Sphenopalatine_VS_P1_Stellate | 0 | 1 | 1 | 1 | 1 | 1 | 0 | 1 |
| SUZ12 | 0.226 | P2_P3_P4_Lumbar_Coeliac_Sphenopalatine_VS_P1_Stellate | 0 | 1 | 1 | 1 | 1 | 1 | 0 | 1 |
| CDR2L | 0.225 | P1_P2_P3_Lumbar_Coeliac_VS_P4_Stellate_Sphenopalatine | 1 | 1 | 1 | 0 | 1 | 1 | 0 | 0 |
| MED12 | 0.225 | P1_P2_P3_Lumbar_Coeliac_VS_P4_Stellate_Sphenopalatine | 1 | 1 | 1 | 0 | 1 | 1 | 0 | 0 |
| DUSP18 | 0.225 | P1_P2_P3_P4_Lumbar_Coeliac_Stellate_VS_Sphenopalatine | 1 | 1 | 1 | 1 | 1 | 1 | 1 | 0 |
| CLUH | 0.225 | P1_P2_P3_P4_Lumbar_Sphenopalatine_VS_Coeliac_Stellate | 1 | 1 | 1 | 1 | 1 | 0 | 0 | 1 |
| ZFP426 | 0.225 | P1_P2_P3_Lumbar_Coeliac_Sphenopalatine_VS_P4_Stellate | 1 | 1 | 1 | 0 | 1 | 1 | 0 | 1 |
| 4931406P16I | 0.225 | P1_P3_P4_Lumbar_Coeliac_Sphenopalatine_VS_P2_Stellate | 1 | 0 | 1 | 1 | 1 | 1 | 0 | 1 |
| TBL1XR1 | 0.225 | P2_P3_P4_Lumbar_Coeliac_Sphenopalatine_VS_P1_Stellate | 0 | 1 | 1 | 1 | 1 | 1 | 0 | 1 |
| SLC36A4 | 0.225 | P2_P3_P4_Lumbar_Coeliac_Sphenopalatine_VS_P1_Stellate | 0 | 1 | 1 | 1 | 1 | 1 | 0 | 1 |
| SRP54A | 0.225 | P2_P3_P4_Lumbar_Coeliac_Sphenopalatine_VS_P1_Stellate | 0 | 1 | 1 | 1 | 1 | 1 | 0 | 1 |
| CUL9 | 0.225 | P2_P3_P4_Lumbar_Coeliac_Sphenopalatine_VS_P1_Stellate | 0 | 1 | 1 | 1 | 1 | 1 | 0 | 1 |
| GM15706 | 0.224 | P1_P2_P3_P4_Lumbar_Coeliac_Stellate_VS_Sphenopalatine | 1 | 1 | 1 | 1 | 1 | 1 | 1 | 0 |
| PCNX | 0.224 | P1_P2_P3_P4_Lumbar_Coeliac_Stellate_VS_Sphenopalatine | 1 | 1 | 1 | 1 | 1 | 1 | 1 | 0 |

| | | | | | | | | | | |
|---|---|---|---|---|---|---|---|---|---|---|
| LRRC75B | 0.224 | P1_P2_P4_Lumbar_Sphenopalatine_VS_P3_Coeliac_Stellate | 1 | 1 | 0 | 1 | 1 | 0 | 0 | 1 |
| BCL2L2 | 0.224 | P1_P2_P3_P4_Lumbar_Sphenopalatine_VS_Coeliac_Stellate | 1 | 1 | 1 | 1 | 1 | 0 | 0 | 1 |
| HDAC6 | 0.224 | P2_P3_P4_Lumbar_Coeliac_Sphenopalatine_VS_P1_Stellate | 0 | 1 | 1 | 1 | 1 | 1 | 0 | 1 |
| MSH3 | 0.223 | P1_P2_Lumbar_Coeliac_VS_P3_P4_Stellate_Sphenopalatine | 1 | 1 | 0 | 0 | 1 | 1 | 0 | 0 |
| FHIT | 0.223 | P1_P2_P3_P4_Lumbar_Coeliac_Stellate_VS_Sphenopalatine | 1 | 1 | 1 | 1 | 1 | 1 | 1 | 0 |
| GOLGA2 | 0.223 | P2_P3_P4_Lumbar_Coeliac_Sphenopalatine_VS_P1_Stellate | 0 | 1 | 1 | 1 | 1 | 1 | 0 | 1 |
| PKD1 | 0.223 | P2_P3_P4_Lumbar_Coeliac_Sphenopalatine_VS_P1_Stellate | 0 | 1 | 1 | 1 | 1 | 1 | 0 | 1 |
| 6030458C11I | 0.222 | P1_P2_P3_Lumbar_Coeliac_VS_P4_Stellate_Sphenopalatine | 1 | 1 | 1 | 0 | 1 | 1 | 0 | 0 |
| FABP7 | 0.222 | Sphenopalatine_VS_P1_P2_P3_P4_Lumbar_Coeliac_Stellate | 0 | 0 | 0 | 0 | 0 | 0 | 0 | 1 |
| DHX57 | 0.222 | P2_P4_Lumbar_Coeliac_Sphenopalatine_VS_P1_P3_Stellate | 0 | 1 | 0 | 1 | 1 | 1 | 0 | 1 |
| IFT88 | 0.222 | P1_P3_P4_Lumbar_Coeliac_Sphenopalatine_VS_P2_Stellate | 1 | 0 | 1 | 1 | 1 | 1 | 0 | 1 |
| SMURF1 | 0.222 | P2_P3_P4_Lumbar_Coeliac_Sphenopalatine_VS_P1_Stellate | 0 | 1 | 1 | 1 | 1 | 1 | 0 | 1 |
| PPP4R3B | 0.222 | P2_P3_P4_Lumbar_Coeliac_Sphenopalatine_VS_P1_Stellate | 0 | 1 | 1 | 1 | 1 | 1 | 0 | 1 |
| ZFP697 | 0.222 | P1_P2_P3_Lumbar_Coeliac_Stellate_Sphenopalatine_VS_P4 | 1 | 1 | 1 | 0 | 1 | 1 | 1 | 1 |
| KCNMB4 | 0.221 | P3_Lumbar_VS_P1_P2_P4_Coeliac_Stellate_Sphenopalatine | 0 | 0 | 1 | 0 | 1 | 0 | 0 | 0 |
| POLRMT | 0.221 | P1_P2_Lumbar_Coeliac_VS_P3_P4_Stellate_Sphenopalatine | 1 | 1 | 0 | 0 | 1 | 1 | 0 | 0 |
| KLF8 | 0.221 | P1_P2_Lumbar_Coeliac_VS_P3_P4_Stellate_Sphenopalatine | 1 | 1 | 0 | 0 | 1 | 1 | 0 | 0 |
| MRPL39 | 0.221 | P1_P3_P4_Lumbar_Coeliac_Stellate_VS_P2_Sphenopalatine | 1 | 0 | 1 | 1 | 1 | 1 | 1 | 0 |
| RHBDL1 | 0.221 | P4_Sphenopalatine_VS_P1_P2_P3_Lumbar_Coeliac_Stellate | 0 | 0 | 0 | 1 | 0 | 0 | 0 | 1 |
| VPS37C | 0.221 | P1_P2_P3_P4_Sphenopalatine_VS_Lumbar_Coeliac_Stellate | 1 | 1 | 1 | 1 | 0 | 0 | 0 | 1 |
| GYS1 | 0.221 | P1_P2_P3_P4_Lumbar_Sphenopalatine_VS_Coeliac_Stellate | 1 | 1 | 1 | 1 | 1 | 0 | 0 | 1 |
| ZFP672 | 0.221 | P1_P2_P3_P4_Coeliac_Sphenopalatine_VS_Lumbar_Stellate | 1 | 1 | 1 | 1 | 0 | 1 | 0 | 1 |
| DDX56 | 0.221 | P1_P3_P4_Lumbar_Coeliac_Sphenopalatine_VS_P2_Stellate | 1 | 0 | 1 | 1 | 1 | 1 | 0 | 1 |
| FAM193B | 0.221 | P2_P3_P4_Lumbar_Coeliac_Sphenopalatine_VS_P1_Stellate | 0 | 1 | 1 | 1 | 1 | 1 | 0 | 1 |
| GM19531 | 0.22 | P3_VS_P1_P2_P4_Lumbar_Coeliac_Stellate_Sphenopalatine | 0 | 0 | 1 | 0 | 0 | 0 | 0 | 0 |
| SNX30 | 0.22 | P1_Lumbar_Coeliac_Sphenopalatine_VS_P2_P3_P4_Stellate | 1 | 0 | 0 | 0 | 1 | 1 | 0 | 1 |
| ACVR2A | 0.22 | P2_P4_Lumbar_Coeliac_Sphenopalatine_VS_P1_P3_Stellate | 0 | 1 | 0 | 1 | 1 | 1 | 0 | 1 |
| CSNK2A2 | 0.22 | P2_P4_Lumbar_Coeliac_Sphenopalatine_VS_P1_P3_Stellate | 0 | 1 | 0 | 1 | 1 | 1 | 0 | 1 |
| ARMCX1 | 0.22 | P3_P4_Lumbar_Coeliac_Sphenopalatine_VS_P1_P2_Stellate | 0 | 0 | 1 | 1 | 1 | 1 | 0 | 1 |
| ATP13A1 | 0.22 | P1_P3_P4_Lumbar_Coeliac_Sphenopalatine_VS_P2_Stellate | 1 | 0 | 1 | 1 | 1 | 1 | 0 | 1 |

| | | | | | | | | | | |
|---|---|---|---|---|---|---|---|---|---|---|
| ACBD3 | 0.22 | P2_P3_P4_Lumbar_Coeliac_Sphenopalatine_VS_P1_Stellate | 0 | 1 | 1 | 1 | 1 | 1 | 0 | 1 |
| FAH | 0.219 | P3_Lumbar_VS_P1_P2_P4_Coeliac_Stellate_Sphenopalatine | 0 | 0 | 1 | 0 | 1 | 0 | 0 | 0 |
| ABCC1 | 0.219 | P1_P2_Lumbar_Coeliac_VS_P3_P4_Stellate_Sphenopalatine | 1 | 1 | 0 | 0 | 1 | 1 | 0 | 0 |
| LHPP | 0.219 | P1_P2_P3_P4_Lumbar_Coeliac_Stellate_VS_Sphenopalatine | 1 | 1 | 1 | 1 | 1 | 1 | 1 | 0 |
| ARHGAP44 | 0.219 | P1_P2_P3_P4_Coeliac_Sphenopalatine_VS_Lumbar_Stellate | 1 | 1 | 1 | 1 | 0 | 1 | 0 | 1 |
| TMEM64 | 0.219 | P1_P2_P3_Lumbar_Coeliac_Sphenopalatine_VS_P4_Stellate | 1 | 1 | 1 | 0 | 1 | 1 | 0 | 1 |
| PARD3 | 0.219 | P1_P2_P3_Lumbar_Coeliac_Sphenopalatine_VS_P4_Stellate | 1 | 1 | 1 | 0 | 1 | 1 | 0 | 1 |
| PKN2 | 0.219 | P1_P3_P4_Lumbar_Coeliac_Sphenopalatine_VS_P2_Stellate | 1 | 0 | 1 | 1 | 1 | 1 | 0 | 1 |
| HEATR3 | 0.219 | P1_P3_P4_Lumbar_Coeliac_Sphenopalatine_VS_P2_Stellate | 1 | 0 | 1 | 1 | 1 | 1 | 0 | 1 |
| FGF12 | 0.219 | P1_P3_P4_Lumbar_Coeliac_Sphenopalatine_VS_P2_Stellate | 1 | 0 | 1 | 1 | 1 | 1 | 0 | 1 |
| IER2 | 0.218 | P1_P2_P3_P4_Lumbar_Sphenopalatine_VS_Coeliac_Stellate | 1 | 1 | 1 | 1 | 1 | 0 | 0 | 1 |
| TSPYL5 | 0.218 | P1_P2_P3_Lumbar_Coeliac_Sphenopalatine_VS_P4_Stellate | 1 | 1 | 1 | 0 | 1 | 1 | 0 | 1 |
| PGBD5 | 0.218 | P2_P4_Lumbar_Coeliac_Sphenopalatine_VS_P1_P3_Stellate | 0 | 1 | 0 | 1 | 1 | 1 | 0 | 1 |
| CIAO1 | 0.218 | P1_P3_P4_Lumbar_Coeliac_Sphenopalatine_VS_P2_Stellate | 1 | 0 | 1 | 1 | 1 | 1 | 0 | 1 |
| LRP8 | 0.218 | P1_P3_P4_Lumbar_Coeliac_Sphenopalatine_VS_P2_Stellate | 1 | 0 | 1 | 1 | 1 | 1 | 0 | 1 |
| RFC1 | 0.218 | P1_P3_P4_Lumbar_Coeliac_Sphenopalatine_VS_P2_Stellate | 1 | 0 | 1 | 1 | 1 | 1 | 0 | 1 |
| ZFP239 | 0.218 | P1_P3_P4_Lumbar_Coeliac_Sphenopalatine_VS_P2_Stellate | 1 | 0 | 1 | 1 | 1 | 1 | 0 | 1 |
| SIL1 | 0.217 | P3_P4_VS_P1_P2_Lumbar_Coeliac_Stellate_Sphenopalatine | 0 | 0 | 1 | 1 | 0 | 0 | 0 | 0 |
| ARFGAP3 | 0.217 | P2_P3_Lumbar_Coeliac_VS_P1_P4_Stellate_Sphenopalatine | 0 | 1 | 1 | 0 | 1 | 1 | 0 | 0 |
| GLRB | 0.217 | P1_P2_P3_P4_Sphenopalatine_VS_Lumbar_Coeliac_Stellate | 1 | 1 | 1 | 1 | 0 | 0 | 0 | 1 |
| TBX2 | 0.217 | P1_P2_P4_Lumbar_Sphenopalatine_VS_P3_Coeliac_Stellate | 1 | 1 | 0 | 1 | 1 | 0 | 0 | 1 |
| KIFC2 | 0.217 | P1_P2_P4_Lumbar_Sphenopalatine_VS_P3_Coeliac_Stellate | 1 | 1 | 0 | 1 | 1 | 0 | 0 | 1 |
| DIS3L | 0.217 | P1_P3_P4_Lumbar_Coeliac_Sphenopalatine_VS_P2_Stellate | 1 | 0 | 1 | 1 | 1 | 1 | 0 | 1 |
| RFX7 | 0.217 | P2_P3_P4_Lumbar_Coeliac_Sphenopalatine_VS_P1_Stellate | 0 | 1 | 1 | 1 | 1 | 1 | 0 | 1 |
| GM45051 | 0.216 | P3_VS_P1_P2_P4_Lumbar_Coeliac_Stellate_Sphenopalatine | 0 | 0 | 1 | 0 | 0 | 0 | 0 | 0 |
| 2510039O18 | 0.216 | P1_P2_P3_P4_Lumbar_VS_Coeliac_Stellate_Sphenopalatine | 1 | 1 | 1 | 1 | 1 | 0 | 0 | 0 |
| MAFK | 0.216 | P1_P2_P3_P4_Lumbar_Coeliac_Stellate_VS_Sphenopalatine | 1 | 1 | 1 | 1 | 1 | 1 | 1 | 0 |
| PLIN2 | 0.216 | P1_P4_Lumbar_Coeliac_Sphenopalatine_VS_P2_P3_Stellate | 1 | 0 | 0 | 1 | 1 | 1 | 0 | 1 |
| RPP25L | 0.216 | P1_P3_P4_Lumbar_Coeliac_Sphenopalatine_VS_P2_Stellate | 1 | 0 | 1 | 1 | 1 | 1 | 0 | 1 |
| MRM3 | 0.216 | P1_P3_P4_Lumbar_Coeliac_Sphenopalatine_VS_P2_Stellate | 1 | 0 | 1 | 1 | 1 | 1 | 0 | 1 |

| | | | | | | | | | | |
|---|---|---|---|---|---|---|---|---|---|---|
| DVL1 | 0.216 | P2_P3_P4_Lumbar_Coeliac_Sphenopalatine_VS_P1_Stellate | 0 | 1 | 1 | 1 | 1 | 1 | 0 | 1 |
| TRAPPC11 | 0.216 | P2_P3_P4_Lumbar_Coeliac_Sphenopalatine_VS_P1_Stellate | 0 | 1 | 1 | 1 | 1 | 1 | 0 | 1 |
| E2F6 | 0.215 | P1_P2_P4_Lumbar_VS_P3_Coeliac_Stellate_Sphenopalatine | 1 | 1 | 0 | 1 | 1 | 0 | 0 | 0 |
| PCSK2OS1 | 0.215 | P2_P3_Lumbar_Coeliac_VS_P1_P4_Stellate_Sphenopalatine | 0 | 1 | 1 | 0 | 1 | 1 | 0 | 0 |
| DUSP15 | 0.215 | Sphenopalatine_VS_P1_P2_P3_P4_Lumbar_Coeliac_Stellate | 0 | 0 | 0 | 0 | 0 | 0 | 0 | 1 |
| JUNB | 0.215 | P1_P2_P3_P4_Coeliac_Sphenopalatine_VS_Lumbar_Stellate | 1 | 1 | 1 | 1 | 0 | 1 | 0 | 1 |
| NCK1 | 0.215 | P1_P3_P4_Lumbar_Coeliac_Sphenopalatine_VS_P2_Stellate | 1 | 0 | 1 | 1 | 1 | 1 | 0 | 1 |
| POC1B | 0.215 | P1_P3_P4_Lumbar_Coeliac_Sphenopalatine_VS_P2_Stellate | 1 | 0 | 1 | 1 | 1 | 1 | 0 | 1 |
| ACVR1 | 0.215 | P2_P3_P4_Lumbar_Coeliac_Sphenopalatine_VS_P1_Stellate | 0 | 1 | 1 | 1 | 1 | 1 | 0 | 1 |
| MEF2D | 0.215 | P2_P3_P4_Lumbar_Coeliac_Sphenopalatine_VS_P1_Stellate | 0 | 1 | 1 | 1 | 1 | 1 | 0 | 1 |
| ZCCHC7 | 0.215 | P2_P3_P4_Lumbar_Coeliac_Sphenopalatine_VS_P1_Stellate | 0 | 1 | 1 | 1 | 1 | 1 | 0 | 1 |
| DENND4A | 0.215 | P2_P3_P4_Lumbar_Coeliac_Sphenopalatine_VS_P1_Stellate | 0 | 1 | 1 | 1 | 1 | 1 | 0 | 1 |
| TRAPPC10 | 0.215 | P2_P3_P4_Lumbar_Coeliac_Sphenopalatine_VS_P1_Stellate | 0 | 1 | 1 | 1 | 1 | 1 | 0 | 1 |
| TMEM86A | 0.214 | P3_VS_P1_P2_P4_Lumbar_Coeliac_Stellate_Sphenopalatine | 0 | 0 | 1 | 0 | 0 | 0 | 0 | 0 |
| HIPK3 | 0.214 | P1_P2_P3_P4_Coeliac_Sphenopalatine_VS_Lumbar_Stellate | 1 | 1 | 1 | 1 | 0 | 1 | 0 | 1 |
| TACC1 | 0.214 | P1_P2_P3_P4_Coeliac_Sphenopalatine_VS_Lumbar_Stellate | 1 | 1 | 1 | 1 | 0 | 1 | 0 | 1 |
| B3GNT9 | 0.214 | P1_P2_P3_Lumbar_Coeliac_Sphenopalatine_VS_P4_Stellate | 1 | 1 | 1 | 0 | 1 | 1 | 0 | 1 |
| AQR | 0.214 | P1_P3_P4_Lumbar_Coeliac_Sphenopalatine_VS_P2_Stellate | 1 | 0 | 1 | 1 | 1 | 1 | 0 | 1 |
| GTPBP3 | 0.214 | P1_P3_P4_Lumbar_Coeliac_Sphenopalatine_VS_P2_Stellate | 1 | 0 | 1 | 1 | 1 | 1 | 0 | 1 |
| INTS9 | 0.214 | P1_P3_P4_Lumbar_Coeliac_Sphenopalatine_VS_P2_Stellate | 1 | 0 | 1 | 1 | 1 | 1 | 0 | 1 |
| SCAF1 | 0.214 | P2_P3_P4_Lumbar_Coeliac_Sphenopalatine_VS_P1_Stellate | 0 | 1 | 1 | 1 | 1 | 1 | 0 | 1 |
| PI4K2A | 0.214 | P2_P3_P4_Lumbar_Coeliac_Sphenopalatine_VS_P1_Stellate | 0 | 1 | 1 | 1 | 1 | 1 | 0 | 1 |
| PDZD8 | 0.214 | P2_P3_P4_Lumbar_Coeliac_Sphenopalatine_VS_P1_Stellate | 0 | 1 | 1 | 1 | 1 | 1 | 0 | 1 |
| GSTM4 | 0.214 | P1_P3_P4_Lumbar_Coeliac_Stellate_Sphenopalatine_VS_P2 | 1 | 0 | 1 | 1 | 1 | 1 | 1 | 1 |
| ZFP37 | 0.213 | P2_P3_Lumbar_Coeliac_VS_P1_P4_Stellate_Sphenopalatine | 0 | 1 | 1 | 0 | 1 | 1 | 0 | 0 |
| TMEM97 | 0.213 | P1_P3_P4_Lumbar_Coeliac_Stellate_VS_P2_Sphenopalatine | 1 | 0 | 1 | 1 | 1 | 1 | 1 | 0 |
| ANKRD39 | 0.213 | P1_P2_P3_P4_Lumbar_Sphenopalatine_VS_Coeliac_Stellate | 1 | 1 | 1 | 1 | 1 | 0 | 0 | 1 |
| HSPA12A | 0.213 | P1_P2_P3_P4_Lumbar_Sphenopalatine_VS_Coeliac_Stellate | 1 | 1 | 1 | 1 | 1 | 0 | 0 | 1 |
| FAM49A | 0.213 | P2_P3_P4_Lumbar_Coeliac_Sphenopalatine_VS_P1_Stellate | 0 | 1 | 1 | 1 | 1 | 1 | 0 | 1 |
| KPNA1 | 0.213 | P2_P3_P4_Lumbar_Coeliac_Sphenopalatine_VS_P1_Stellate | 0 | 1 | 1 | 1 | 1 | 1 | 0 | 1 |

| | | | | | | | | | | |
|---|---|---|---|---|---|---|---|---|---|---|
| RIIAD1 | 0.212 | P1_P2_P3_P4_Sphenopalatine_VS_Lumbar_Coeliac_Stellate | 1 | 1 | 1 | 1 | 0 | 0 | 0 | 1 |
| RAPGEF1 | 0.212 | P1_P2_P3_P4_Lumbar_Sphenopalatine_VS_Coeliac_Stellate | 1 | 1 | 1 | 1 | 1 | 0 | 0 | 1 |
| MEX3D | 0.212 | P1_P2_P3_Lumbar_Coeliac_Sphenopalatine_VS_P4_Stellate | 1 | 1 | 1 | 0 | 1 | 1 | 0 | 1 |
| UBP1 | 0.212 | P2_P3_P4_Lumbar_Coeliac_Sphenopalatine_VS_P1_Stellate | 0 | 1 | 1 | 1 | 1 | 1 | 0 | 1 |
| MYH14 | 0.211 | P2_P3_P4_VS_P1_Lumbar_Coeliac_Stellate_Sphenopalatine | 0 | 1 | 1 | 1 | 0 | 0 | 0 | 0 |
| FUCA1 | 0.211 | P3_Lumbar_VS_P1_P2_P4_Coeliac_Stellate_Sphenopalatine | 0 | 0 | 1 | 0 | 1 | 0 | 0 | 0 |
| LPIN1 | 0.211 | P1_P2_P3_P4_Lumbar_VS_Coeliac_Stellate_Sphenopalatine | 1 | 1 | 1 | 1 | 1 | 0 | 0 | 0 |
| RRAGD | 0.211 | P2_P3_Lumbar_Coeliac_VS_P1_P4_Stellate_Sphenopalatine | 0 | 1 | 1 | 0 | 1 | 1 | 0 | 0 |
| ARHGEF4 | 0.211 | P1_P2_P3_P4_Lumbar_Sphenopalatine_VS_Coeliac_Stellate | 1 | 1 | 1 | 1 | 1 | 0 | 0 | 1 |
| SIGMAR1 | 0.211 | P1_P2_P3_P4_Lumbar_Sphenopalatine_VS_Coeliac_Stellate | 1 | 1 | 1 | 1 | 1 | 0 | 0 | 1 |
| MIR22HG | 0.211 | P1_P2_P3_P4_Lumbar_Sphenopalatine_VS_Coeliac_Stellate | 1 | 1 | 1 | 1 | 1 | 0 | 0 | 1 |
| ZNHIT6 | 0.211 | Lumbar_Coeliac_Sphenopalatine_VS_P1_P2_P3_P4_Stellate | 0 | 0 | 0 | 0 | 1 | 1 | 0 | 1 |
| NFIL3 | 0.211 | P1_Lumbar_Coeliac_Sphenopalatine_VS_P2_P3_P4_Stellate | 1 | 0 | 0 | 0 | 1 | 1 | 0 | 1 |
| ATF6 | 0.211 | P2_P3_P4_Lumbar_Coeliac_Sphenopalatine_VS_P1_Stellate | 0 | 1 | 1 | 1 | 1 | 1 | 0 | 1 |
| GOLPH3L | 0.211 | P2_P3_P4_Lumbar_Coeliac_Sphenopalatine_VS_P1_Stellate | 0 | 1 | 1 | 1 | 1 | 1 | 0 | 1 |
| RPRD2 | 0.211 | P2_P3_P4_Lumbar_Coeliac_Sphenopalatine_VS_P1_Stellate | 0 | 1 | 1 | 1 | 1 | 1 | 0 | 1 |
| ITSN1 | 0.211 | P2_P3_P4_Lumbar_Coeliac_Sphenopalatine_VS_P1_Stellate | 0 | 1 | 1 | 1 | 1 | 1 | 0 | 1 |
| SYVN1 | 0.211 | P2_P3_P4_Lumbar_Coeliac_Sphenopalatine_VS_P1_Stellate | 0 | 1 | 1 | 1 | 1 | 1 | 0 | 1 |
| SLC16A7 | 0.21 | P1_P2_P4_Lumbar_VS_P3_Coeliac_Stellate_Sphenopalatine | 1 | 1 | 0 | 1 | 1 | 0 | 0 | 0 |
| FCF1 | 0.21 | P1_P3_P4_Lumbar_Coeliac_Stellate_VS_P2_Sphenopalatine | 1 | 0 | 1 | 1 | 1 | 1 | 1 | 0 |
| CPT2 | 0.21 | P1_P2_P3_P4_Lumbar_Coeliac_Stellate_VS_Sphenopalatine | 1 | 1 | 1 | 1 | 1 | 1 | 1 | 0 |
| LRRC20 | 0.21 | P1_P2_P3_P4_Lumbar_Sphenopalatine_VS_Coeliac_Stellate | 1 | 1 | 1 | 1 | 1 | 0 | 0 | 1 |
| ZFR2 | 0.21 | P1_P2_P3_P4_Lumbar_Sphenopalatine_VS_Coeliac_Stellate | 1 | 1 | 1 | 1 | 1 | 0 | 0 | 1 |
| MGA | 0.21 | P1_P4_Lumbar_Coeliac_Sphenopalatine_VS_P2_P3_Stellate | 1 | 0 | 0 | 1 | 1 | 1 | 0 | 1 |
| ZC3H13 | 0.21 | P2_P3_P4_Lumbar_Coeliac_Sphenopalatine_VS_P1_Stellate | 0 | 1 | 1 | 1 | 1 | 1 | 0 | 1 |
| GDPD5 | 0.209 | P2_P4_Lumbar_VS_P1_P3_Coeliac_Stellate_Sphenopalatine | 0 | 1 | 0 | 1 | 1 | 0 | 0 | 0 |
| PHLDA3 | 0.209 | P1_P2_P3_P4_Lumbar_Sphenopalatine_VS_Coeliac_Stellate | 1 | 1 | 1 | 1 | 1 | 0 | 0 | 1 |
| CRY2 | 0.209 | P1_P2_P3_P4_Lumbar_Sphenopalatine_VS_Coeliac_Stellate | 1 | 1 | 1 | 1 | 1 | 0 | 0 | 1 |
| AMIGO1 | 0.209 | P1_P2_P3_P4_Coeliac_Sphenopalatine_VS_Lumbar_Stellate | 1 | 1 | 1 | 1 | 0 | 1 | 0 | 1 |
| INO80D | 0.209 | P1_P2_P3_Lumbar_Coeliac_Sphenopalatine_VS_P4_Stellate | 1 | 1 | 1 | 0 | 1 | 1 | 0 | 1 |

| | | | | | | | | | | |
|---|---|---|---|---|---|---|---|---|---|---|
| ZDHHC8 | 0.209 | P2_P3_P4_Lumbar_Coeliac_Sphenopalatine_VS_P1_Stellate | 0 | 1 | 1 | 1 | 1 | 1 | 0 | 1 |
| 2700081O15 | 0.209 | P2_P3_P4_Lumbar_Coeliac_Sphenopalatine_VS_P1_Stellate | 0 | 1 | 1 | 1 | 1 | 1 | 0 | 1 |
| HIF1A | 0.208 | P2_P4_Lumbar_VS_P1_P3_Coeliac_Stellate_Sphenopalatine | 0 | 1 | 0 | 1 | 1 | 0 | 0 | 0 |
| ZFP932 | 0.208 | P1_P2_P3_P4_Coeliac_VS_Lumbar_Stellate_Sphenopalatine | 1 | 1 | 1 | 1 | 0 | 1 | 0 | 0 |
| MAP2K3 | 0.208 | P1_P2_P3_P4_Lumbar_Sphenopalatine_VS_Coeliac_Stellate | 1 | 1 | 1 | 1 | 1 | 0 | 0 | 1 |
| HDAC4 | 0.208 | P2_P3_P4_Lumbar_Coeliac_Sphenopalatine_VS_P1_Stellate | 0 | 1 | 1 | 1 | 1 | 1 | 0 | 1 |
| GLG1 | 0.208 | P2_P3_P4_Lumbar_Coeliac_Sphenopalatine_VS_P1_Stellate | 0 | 1 | 1 | 1 | 1 | 1 | 0 | 1 |
| PIK3C3 | 0.208 | P2_P3_P4_Lumbar_Coeliac_Sphenopalatine_VS_P1_Stellate | 0 | 1 | 1 | 1 | 1 | 1 | 0 | 1 |
| ASL | 0.208 | P1_P2_P3_Lumbar_Coeliac_Stellate_Sphenopalatine_VS_P4 | 1 | 1 | 1 | 0 | 1 | 1 | 1 | 1 |
| GM19938 | 0.207 | P2_P4_Lumbar_VS_P1_P3_Coeliac_Stellate_Sphenopalatine | 0 | 1 | 0 | 1 | 1 | 0 | 0 | 0 |
| ETV5 | 0.207 | P1_P2_P3_P4_Lumbar_Coeliac_Stellate_VS_Sphenopalatine | 1 | 1 | 1 | 1 | 1 | 1 | 1 | 0 |
| IGIP | 0.207 | P1_P2_P3_P4_Lumbar_Sphenopalatine_VS_Coeliac_Stellate | 1 | 1 | 1 | 1 | 1 | 0 | 0 | 1 |
| LRIG2 | 0.207 | P2_P3_P4_Lumbar_Coeliac_Sphenopalatine_VS_P1_Stellate | 0 | 1 | 1 | 1 | 1 | 1 | 0 | 1 |
| DDI2 | 0.207 | P2_P3_P4_Lumbar_Coeliac_Sphenopalatine_VS_P1_Stellate | 0 | 1 | 1 | 1 | 1 | 1 | 0 | 1 |
| EGLN1 | 0.207 | P2_P3_P4_Lumbar_Coeliac_Sphenopalatine_VS_P1_Stellate | 0 | 1 | 1 | 1 | 1 | 1 | 0 | 1 |
| COBL | 0.207 | P2_P3_P4_Lumbar_Coeliac_Sphenopalatine_VS_P1_Stellate | 0 | 1 | 1 | 1 | 1 | 1 | 0 | 1 |
| CENPX | 0.207 | P1_P2_P3_Lumbar_Coeliac_Stellate_Sphenopalatine_VS_P4 | 1 | 1 | 1 | 0 | 1 | 1 | 1 | 1 |
| GM14295 | 0.206 | P1_P2_P3_P4_VS_Lumbar_Coeliac_Stellate_Sphenopalatine | 1 | 1 | 1 | 1 | 0 | 0 | 0 | 0 |
| NT5DC2 | 0.206 | P3_Lumbar_Coeliac_Stellate_VS_P1_P2_P4_Sphenopalatine | 0 | 0 | 1 | 0 | 1 | 1 | 1 | 0 |
| ADAMTS8 | 0.206 | Sphenopalatine_VS_P1_P2_P3_P4_Lumbar_Coeliac_Stellate | 0 | 0 | 0 | 0 | 0 | 0 | 0 | 1 |
| NPTX1 | 0.206 | Sphenopalatine_VS_P1_P2_P3_P4_Lumbar_Coeliac_Stellate | 0 | 0 | 0 | 0 | 0 | 0 | 0 | 1 |
| PITPNM2 | 0.206 | P1_P2_P3_P4_Lumbar_Sphenopalatine_VS_Coeliac_Stellate | 1 | 1 | 1 | 1 | 1 | 0 | 0 | 1 |
| CPD | 0.206 | P2_P3_P4_Lumbar_Coeliac_Sphenopalatine_VS_P1_Stellate | 0 | 1 | 1 | 1 | 1 | 1 | 0 | 1 |
| IGSF8 | 0.205 | P3_P4_Lumbar_VS_P1_P2_Coeliac_Stellate_Sphenopalatine | 0 | 0 | 1 | 1 | 1 | 0 | 0 | 0 |
| R3HDML | 0.205 | Sphenopalatine_VS_P1_P2_P3_P4_Lumbar_Coeliac_Stellate | 0 | 0 | 0 | 0 | 0 | 0 | 0 | 1 |
| GGACT | 0.205 | P1_P2_P3_P4_Sphenopalatine_VS_Lumbar_Coeliac_Stellate | 1 | 1 | 1 | 1 | 0 | 0 | 0 | 1 |
| UPP2 | 0.205 | P2_P3_P4_Lumbar_Coeliac_Sphenopalatine_VS_P1_Stellate | 0 | 1 | 1 | 1 | 1 | 1 | 0 | 1 |
| OXSR1 | 0.205 | P2_P3_P4_Lumbar_Coeliac_Sphenopalatine_VS_P1_Stellate | 0 | 1 | 1 | 1 | 1 | 1 | 0 | 1 |
| ZFP445 | 0.205 | P2_P3_P4_Lumbar_Coeliac_Sphenopalatine_VS_P1_Stellate | 0 | 1 | 1 | 1 | 1 | 1 | 0 | 1 |
| NHSL1 | 0.204 | Lumbar_Coeliac_Sphenopalatine_VS_P1_P2_P3_P4_Stellate | 0 | 0 | 0 | 0 | 1 | 1 | 0 | 1 |

| | | | | | | | | | | |
|---|---|---|---|---|---|---|---|---|---|---|
| ARHGEF18 | 0.204 | P2_P3_P4_Lumbar_Coeliac_Sphenopalatine_VS_P1_Stellate | 0 | 1 | 1 | 1 | 1 | 1 | 0 | 1 |
| ABCG4 | 0.204 | P2_P3_P4_Lumbar_Coeliac_Sphenopalatine_VS_P1_Stellate | 0 | 1 | 1 | 1 | 1 | 1 | 0 | 1 |
| ATP1B2 | 0.203 | Sphenopalatine_VS_P1_P2_P3_P4_Lumbar_Coeliac_Stellate | 0 | 0 | 0 | 0 | 0 | 0 | 0 | 1 |
| UNC80 | 0.203 | P1_P2_P3_P4_Lumbar_Sphenopalatine_VS_Coeliac_Stellate | 1 | 1 | 1 | 1 | 1 | 0 | 0 | 1 |
| IFFO1 | 0.203 | P1_P2_P3_P4_Lumbar_Sphenopalatine_VS_Coeliac_Stellate | 1 | 1 | 1 | 1 | 1 | 0 | 0 | 1 |
| PTPRO | 0.203 | P1_P2_P3_P4_Lumbar_Sphenopalatine_VS_Coeliac_Stellate | 1 | 1 | 1 | 1 | 1 | 0 | 0 | 1 |
| TOM1 | 0.203 | P1_P2_P3_P4_Lumbar_Sphenopalatine_VS_Coeliac_Stellate | 1 | 1 | 1 | 1 | 1 | 0 | 0 | 1 |
| THAP2 | 0.203 | P1_P2_P3_Lumbar_Coeliac_Sphenopalatine_VS_P4_Stellate | 1 | 1 | 1 | 0 | 1 | 1 | 0 | 1 |
| H2-T22 | 0.202 | P1_P3_VS_P2_P4_Lumbar_Coeliac_Stellate_Sphenopalatine | 1 | 0 | 1 | 0 | 0 | 0 | 0 | 0 |
| SNHG10 | 0.202 | P3_P4_VS_P1_P2_Lumbar_Coeliac_Stellate_Sphenopalatine | 0 | 0 | 1 | 1 | 0 | 0 | 0 | 0 |
| PER3 | 0.202 | P2_P3_P4_VS_P1_Lumbar_Coeliac_Stellate_Sphenopalatine | 0 | 1 | 1 | 1 | 0 | 0 | 0 | 0 |
| PALM3 | 0.202 | P1_P2_P3_P4_VS_Lumbar_Coeliac_Stellate_Sphenopalatine | 1 | 1 | 1 | 1 | 0 | 0 | 0 | 0 |
| D630045J12I | 0.202 | P2_P3_Lumbar_Coeliac_VS_P1_P4_Stellate_Sphenopalatine | 0 | 1 | 1 | 0 | 1 | 1 | 0 | 0 |
| EML1 | 0.202 | P1_P2_P3_P4_Lumbar_Coeliac_Stellate_VS_Sphenopalatine | 1 | 1 | 1 | 1 | 1 | 1 | 1 | 0 |
| SLC5A5 | 0.202 | P1_P2_P3_P4_Sphenopalatine_VS_Lumbar_Coeliac_Stellate | 1 | 1 | 1 | 1 | 0 | 0 | 0 | 1 |
| ST8SIA3 | 0.202 | P1_P2_P3_P4_Lumbar_Sphenopalatine_VS_Coeliac_Stellate | 1 | 1 | 1 | 1 | 1 | 0 | 0 | 1 |
| ANKRD29 | 0.202 | P2_P3_P4_Lumbar_Coeliac_Stellate_Sphenopalatine_VS_P1 | 0 | 1 | 1 | 1 | 1 | 1 | 1 | 1 |
| SORBS1 | 0.201 | P2_P4_Lumbar_VS_P1_P3_Coeliac_Stellate_Sphenopalatine | 0 | 1 | 0 | 1 | 1 | 0 | 0 | 0 |
| ZFP354A | 0.201 | P2_P3_Lumbar_Coeliac_VS_P1_P4_Stellate_Sphenopalatine | 0 | 1 | 1 | 0 | 1 | 1 | 0 | 0 |
| ADAP1 | 0.201 | P1_P2_P3_P4_Lumbar_Sphenopalatine_VS_Coeliac_Stellate | 1 | 1 | 1 | 1 | 1 | 0 | 0 | 1 |
| CEBPD | 0.201 | P1_P2_P3_P4_Coeliac_Sphenopalatine_VS_Lumbar_Stellate | 1 | 1 | 1 | 1 | 0 | 1 | 0 | 1 |
| PHF3 | 0.201 | P1_P2_P3_Lumbar_Coeliac_Sphenopalatine_VS_P4_Stellate | 1 | 1 | 1 | 0 | 1 | 1 | 0 | 1 |
| ABRACL | 0.201 | P1_P2_P3_Lumbar_Coeliac_Stellate_Sphenopalatine_VS_P4 | 1 | 1 | 1 | 0 | 1 | 1 | 1 | 1 |
| ATPAF2 | 0.2 | P2_P3_P4_VS_P1_Lumbar_Coeliac_Stellate_Sphenopalatine | 0 | 1 | 1 | 1 | 0 | 0 | 0 | 0 |
| ZC3H14 | 0.2 | P3_P4_Lumbar_VS_P1_P2_Coeliac_Stellate_Sphenopalatine | 0 | 0 | 1 | 1 | 1 | 0 | 0 | 0 |
| HERC3 | 0.2 | P1_P2_P3_P4_Lumbar_Sphenopalatine_VS_Coeliac_Stellate | 1 | 1 | 1 | 1 | 1 | 0 | 0 | 1 |
| EXOC6 | 0.2 | P1_P2_P3_P4_Lumbar_Sphenopalatine_VS_Coeliac_Stellate | 1 | 1 | 1 | 1 | 1 | 0 | 0 | 1 |
| AI480526 | 0.2 | P2_P4_Lumbar_Coeliac_Sphenopalatine_VS_P1_P3_Stellate | 0 | 1 | 0 | 1 | 1 | 1 | 0 | 1 |
| ZFP462 | 0.2 | P1_P2_P4_Lumbar_Coeliac_Stellate_Sphenopalatine_VS_P3 | 1 | 1 | 0 | 1 | 1 | 1 | 1 | 1 |
| HTATIP2 | 0.199 | P2_P3_P4_VS_P1_Lumbar_Coeliac_Stellate_Sphenopalatine | 0 | 1 | 1 | 1 | 0 | 0 | 0 | 0 |

| | | | | | | | | | | |
|---|---|---|---|---|---|---|---|---|---|---|
| SH3PXD2B | 0.199 | P1_P2_P3_P4_Lumbar_Coeliac_Stellate_VS_Sphenopalatine | 1 | 1 | 1 | 1 | 1 | 1 | 1 | 0 |
| GAS2L1 | 0.199 | P1_P2_P3_P4_Sphenopalatine_VS_Lumbar_Coeliac_Stellate | 1 | 1 | 1 | 1 | 0 | 0 | 0 | 1 |
| PGAP1 | 0.199 | P1_P2_P3_P4_Lumbar_Sphenopalatine_VS_Coeliac_Stellate | 1 | 1 | 1 | 1 | 1 | 0 | 0 | 1 |
| SYNGR1 | 0.199 | P1_P2_P3_P4_Lumbar_Sphenopalatine_VS_Coeliac_Stellate | 1 | 1 | 1 | 1 | 1 | 0 | 0 | 1 |
| EEA1 | 0.199 | P1_P4_Lumbar_Coeliac_Sphenopalatine_VS_P2_P3_Stellate | 1 | 0 | 0 | 1 | 1 | 1 | 0 | 1 |
| GM10260 | 0.199 | P1_P2_P3_P4_Stellate_Sphenopalatine_VS_Lumbar_Coeliac | 1 | 1 | 1 | 1 | 0 | 0 | 1 | 1 |
| AW554918 | 0.198 | P2_P3_Lumbar_Coeliac_VS_P1_P4_Stellate_Sphenopalatine | 0 | 1 | 1 | 0 | 1 | 1 | 0 | 0 |
| PANX2 | 0.198 | P4_Sphenopalatine_VS_P1_P2_P3_Lumbar_Coeliac_Stellate | 0 | 0 | 0 | 1 | 0 | 0 | 0 | 1 |
| ZFP180 | 0.198 | P1_P2_P3_P4_Lumbar_Sphenopalatine_VS_Coeliac_Stellate | 1 | 1 | 1 | 1 | 1 | 0 | 0 | 1 |
| TMEM229B | 0.198 | P1_P2_P3_P4_Lumbar_Sphenopalatine_VS_Coeliac_Stellate | 1 | 1 | 1 | 1 | 1 | 0 | 0 | 1 |
| EXOC2 | 0.198 | P1_P2_P3_Lumbar_Coeliac_Sphenopalatine_VS_P4_Stellate | 1 | 1 | 1 | 0 | 1 | 1 | 0 | 1 |
| ZFP512B | 0.198 | P2_P4_Lumbar_Coeliac_Sphenopalatine_VS_P1_P3_Stellate | 0 | 1 | 0 | 1 | 1 | 1 | 0 | 1 |
| SPINT2 | 0.198 | P1_P3_P4_Lumbar_Coeliac_Stellate_Sphenopalatine_VS_P2 | 1 | 0 | 1 | 1 | 1 | 1 | 1 | 1 |
| JAKMIP3 | 0.197 | P3_P4_VS_P1_P2_Lumbar_Coeliac_Stellate_Sphenopalatine | 0 | 0 | 1 | 1 | 0 | 0 | 0 | 0 |
| GGH | 0.196 | P3_Lumbar_VS_P1_P2_P4_Coeliac_Stellate_Sphenopalatine | 0 | 0 | 1 | 0 | 1 | 0 | 0 | 0 |
| ITGA3 | 0.196 | P1_P2_P3_P4_Lumbar_Sphenopalatine_VS_Coeliac_Stellate | 1 | 1 | 1 | 1 | 1 | 0 | 0 | 1 |
| FARP1 | 0.196 | P1_P2_P3_Lumbar_Coeliac_Sphenopalatine_VS_P4_Stellate | 1 | 1 | 1 | 0 | 1 | 1 | 0 | 1 |
| GTF2B | 0.196 | P1_P4_Lumbar_Coeliac_Sphenopalatine_VS_P2_P3_Stellate | 1 | 0 | 0 | 1 | 1 | 1 | 0 | 1 |
| SCN8A | 0.195 | P2_P4_Lumbar_VS_P1_P3_Coeliac_Stellate_Sphenopalatine | 0 | 1 | 0 | 1 | 1 | 0 | 0 | 0 |
| CCDC149 | 0.195 | P1_P2_P3_P4_Coeliac_Sphenopalatine_VS_Lumbar_Stellate | 1 | 1 | 1 | 1 | 0 | 1 | 0 | 1 |
| CACNG7 | 0.195 | P1_P2_P3_P4_Coeliac_Sphenopalatine_VS_Lumbar_Stellate | 1 | 1 | 1 | 1 | 0 | 1 | 0 | 1 |
| CASTOR2 | 0.195 | P2_Lumbar_Coeliac_Sphenopalatine_VS_P1_P3_P4_Stellate | 0 | 1 | 0 | 0 | 1 | 1 | 0 | 1 |
| ABHD2 | 0.194 | P1_P2_P3_P4_Coeliac_VS_Lumbar_Stellate_Sphenopalatine | 1 | 1 | 1 | 1 | 0 | 1 | 0 | 0 |
| SUMF1 | 0.194 | P1_P2_P3_P4_Sphenopalatine_VS_Lumbar_Coeliac_Stellate | 1 | 1 | 1 | 1 | 0 | 0 | 0 | 1 |
| EHD4 | 0.194 | P1_P2_P4_Lumbar_Sphenopalatine_VS_P3_Coeliac_Stellate | 1 | 1 | 0 | 1 | 1 | 0 | 0 | 1 |
| CDC42EP4 | 0.194 | P1_P2_P3_Lumbar_Coeliac_Sphenopalatine_VS_P4_Stellate | 1 | 1 | 1 | 0 | 1 | 1 | 0 | 1 |
| MAPK6 | 0.194 | P3_P4_Lumbar_Coeliac_Sphenopalatine_VS_P1_P2_Stellate | 0 | 0 | 1 | 1 | 1 | 1 | 0 | 1 |
| AI506816 | 0.193 | P1_Lumbar_VS_P2_P3_P4_Coeliac_Stellate_Sphenopalatine | 1 | 0 | 0 | 0 | 1 | 0 | 0 | 0 |
| GFPT1 | 0.193 | P2_P3_Lumbar_Coeliac_VS_P1_P4_Stellate_Sphenopalatine | 0 | 1 | 1 | 0 | 1 | 1 | 0 | 0 |
| SERPINI1 | 0.193 | P1_P3_P4_Lumbar_Coeliac_Stellate_VS_P2_Sphenopalatine | 1 | 0 | 1 | 1 | 1 | 1 | 1 | 0 |

| | | | | | | | | | | |
|---|---|---|---|---|---|---|---|---|---|---|
| STAMBPL1 | 0.193 | P1_P2_P3_P4_Lumbar_Coeliac_Stellate_VS_Sphenopalatine | 1 | 1 | 1 | 1 | 1 | 1 | 1 | 0 |
| ACVR1B | 0.193 | P1_P2_P3_P4_Lumbar_Sphenopalatine_VS_Coeliac_Stellate | 1 | 1 | 1 | 1 | 1 | 0 | 0 | 1 |
| ATL3 | 0.193 | P1_P2_P3_P4_Lumbar_Sphenopalatine_VS_Coeliac_Stellate | 1 | 1 | 1 | 1 | 1 | 0 | 0 | 1 |
| KLHL8 | 0.193 | P1_P2_P3_Lumbar_Coeliac_Sphenopalatine_VS_P4_Stellate | 1 | 1 | 1 | 0 | 1 | 1 | 0 | 1 |
| EEF2K | 0.193 | P1_P2_P3_Lumbar_Coeliac_Stellate_Sphenopalatine_VS_P4 | 1 | 1 | 1 | 0 | 1 | 1 | 1 | 1 |
| FXYD2 | 0.193 | P1_P3_P4_Lumbar_Coeliac_Stellate_Sphenopalatine_VS_P2 | 1 | 0 | 1 | 1 | 1 | 1 | 1 | 1 |
| GM11361 | 0.192 | P3_VS_P1_P2_P4_Lumbar_Coeliac_Stellate_Sphenopalatine | 0 | 0 | 1 | 0 | 0 | 0 | 0 | 0 |
| PPIE | 0.192 | P3_Lumbar_VS_P1_P2_P4_Coeliac_Stellate_Sphenopalatine | 0 | 0 | 1 | 0 | 1 | 0 | 0 | 0 |
| PIP4K2C | 0.192 | P1_P2_P3_P4_Lumbar_Sphenopalatine_VS_Coeliac_Stellate | 1 | 1 | 1 | 1 | 1 | 0 | 0 | 1 |
| ARRB2 | 0.192 | P1_P2_P3_P4_Lumbar_Sphenopalatine_VS_Coeliac_Stellate | 1 | 1 | 1 | 1 | 1 | 0 | 0 | 1 |
| SOBP | 0.192 | P1_P2_P3_Lumbar_Coeliac_Sphenopalatine_VS_P4_Stellate | 1 | 1 | 1 | 0 | 1 | 1 | 0 | 1 |
| LRRC3 | 0.192 | P1_P2_P3_Lumbar_Coeliac_Sphenopalatine_VS_P4_Stellate | 1 | 1 | 1 | 0 | 1 | 1 | 0 | 1 |
| GADD45B | 0.191 | P3_VS_P1_P2_P4_Lumbar_Coeliac_Stellate_Sphenopalatine | 0 | 0 | 1 | 0 | 0 | 0 | 0 | 0 |
| SLC30A4 | 0.191 | P2_P3_Coeliac_VS_P1_P4_Lumbar_Stellate_Sphenopalatine | 0 | 1 | 1 | 0 | 0 | 1 | 0 | 0 |
| SEC23IP | 0.191 | P2_P3_Lumbar_Coeliac_VS_P1_P4_Stellate_Sphenopalatine | 0 | 1 | 1 | 0 | 1 | 1 | 0 | 0 |
| FBXW8 | 0.191 | P1_P2_P3_P4_Lumbar_Sphenopalatine_VS_Coeliac_Stellate | 1 | 1 | 1 | 1 | 1 | 0 | 0 | 1 |
| OSBPL5 | 0.191 | P1_P2_P3_P4_Lumbar_Sphenopalatine_VS_Coeliac_Stellate | 1 | 1 | 1 | 1 | 1 | 0 | 0 | 1 |
| RNF165 | 0.191 | P2_P4_Lumbar_Coeliac_Sphenopalatine_VS_P1_P3_Stellate | 0 | 1 | 0 | 1 | 1 | 1 | 0 | 1 |
| TRIQK | 0.19 | P3_P4_Lumbar_VS_P1_P2_Coeliac_Stellate_Sphenopalatine | 0 | 0 | 1 | 1 | 1 | 0 | 0 | 0 |
| RYK | 0.19 | P1_P2_P3_P4_Lumbar_Coeliac_Stellate_VS_Sphenopalatine | 1 | 1 | 1 | 1 | 1 | 1 | 1 | 0 |
| FAM219B | 0.19 | P1_P2_P3_P4_Lumbar_Sphenopalatine_VS_Coeliac_Stellate | 1 | 1 | 1 | 1 | 1 | 0 | 0 | 1 |
| EMB | 0.19 | P1_P2_P3_P4_Lumbar_Sphenopalatine_VS_Coeliac_Stellate | 1 | 1 | 1 | 1 | 1 | 0 | 0 | 1 |
| EPC2 | 0.19 | P1_P2_P3_Lumbar_Coeliac_Sphenopalatine_VS_P4_Stellate | 1 | 1 | 1 | 0 | 1 | 1 | 0 | 1 |
| NPAT | 0.189 | P2_P3_Lumbar_Coeliac_VS_P1_P4_Stellate_Sphenopalatine | 0 | 1 | 1 | 0 | 1 | 1 | 0 | 0 |
| SYPL | 0.189 | P1_P2_P3_P4_Lumbar_Sphenopalatine_VS_Coeliac_Stellate | 1 | 1 | 1 | 1 | 1 | 0 | 0 | 1 |
| TBC1D9B | 0.189 | P1_P2_P3_P4_Coeliac_Sphenopalatine_VS_Lumbar_Stellate | 1 | 1 | 1 | 1 | 0 | 1 | 0 | 1 |
| RDH13 | 0.189 | P1_P2_P3_Lumbar_Coeliac_Sphenopalatine_VS_P4_Stellate | 1 | 1 | 1 | 0 | 1 | 1 | 0 | 1 |
| SLC43A2 | 0.189 | P2_P4_Lumbar_Coeliac_Sphenopalatine_VS_P1_P3_Stellate | 0 | 1 | 0 | 1 | 1 | 1 | 0 | 1 |
| MAB21L1 | 0.188 | P3_VS_P1_P2_P4_Lumbar_Coeliac_Stellate_Sphenopalatine | 0 | 0 | 1 | 0 | 0 | 0 | 0 | 0 |
| DGKB | 0.188 | P1_P2_P4_Coeliac_VS_P3_Lumbar_Stellate_Sphenopalatine | 1 | 1 | 0 | 1 | 0 | 1 | 0 | 0 |

| | | | | | | | | | | |
|---|---|---|---|---|---|---|---|---|---|---|
| CTCF | 0.188 | P2_P3_Lumbar_Coeliac_VS_P1_P4_Stellate_Sphenopalatine | 0 | 1 | 1 | 0 | 1 | 1 | 0 | 0 |
| ICE1 | 0.188 | P1_P2_P3_Lumbar_Coeliac_Sphenopalatine_VS_P4_Stellate | 1 | 1 | 1 | 0 | 1 | 1 | 0 | 1 |
| ADAMTS10 | 0.188 | P1_P4_Lumbar_Coeliac_Sphenopalatine_VS_P2_P3_Stellate | 1 | 0 | 0 | 1 | 1 | 1 | 0 | 1 |
| COQ3 | 0.187 | P3_P4_Lumbar_VS_P1_P2_Coeliac_Stellate_Sphenopalatine | 0 | 0 | 1 | 1 | 1 | 0 | 0 | 0 |
| OSCP1 | 0.187 | P3_P4_Lumbar_VS_P1_P2_Coeliac_Stellate_Sphenopalatine | 0 | 0 | 1 | 1 | 1 | 0 | 0 | 0 |
| CDKN2AIPNL | 0.187 | P1_P2_P3_P4_Lumbar_Sphenopalatine_VS_Coeliac_Stellate | 1 | 1 | 1 | 1 | 1 | 0 | 0 | 1 |
| MPP2 | 0.187 | P1_P2_P3_P4_Lumbar_Sphenopalatine_VS_Coeliac_Stellate | 1 | 1 | 1 | 1 | 1 | 0 | 0 | 1 |
| TBC1D22A | 0.187 | P1_P2_P3_P4_Coeliac_Sphenopalatine_VS_Lumbar_Stellate | 1 | 1 | 1 | 1 | 0 | 1 | 0 | 1 |
| 2900093K20I | 0.187 | P3_P4_Lumbar_Coeliac_Sphenopalatine_VS_P1_P2_Stellate | 0 | 0 | 1 | 1 | 1 | 1 | 0 | 1 |
| DDIT4 | 0.186 | P1_VS_P2_P3_P4_Lumbar_Coeliac_Stellate_Sphenopalatine | 1 | 0 | 0 | 0 | 0 | 0 | 0 | 0 |
| RAB36 | 0.186 | P2_P4_Coeliac_VS_P1_P3_Lumbar_Stellate_Sphenopalatine | 0 | 1 | 0 | 1 | 0 | 1 | 0 | 0 |
| LYSMD2 | 0.186 | P1_P2_P3_P4_Sphenopalatine_VS_Lumbar_Coeliac_Stellate | 1 | 1 | 1 | 1 | 0 | 0 | 0 | 1 |
| BSN | 0.186 | P1_P2_P3_P4_Lumbar_Sphenopalatine_VS_Coeliac_Stellate | 1 | 1 | 1 | 1 | 1 | 0 | 0 | 1 |
| ANXA2 | 0.186 | P1_P2_P3_P4_Lumbar_Stellate_Sphenopalatine_VS_Coeliac | 1 | 1 | 1 | 1 | 1 | 0 | 1 | 1 |
| PCED1B | 0.185 | P3_P4_VS_P1_P2_Lumbar_Coeliac_Stellate_Sphenopalatine | 0 | 0 | 1 | 1 | 0 | 0 | 0 | 0 |
| WWP2 | 0.185 | P2_P4_Lumbar_VS_P1_P3_Coeliac_Stellate_Sphenopalatine | 0 | 1 | 0 | 1 | 1 | 0 | 0 | 0 |
| BAG2 | 0.185 | P1_P2_P3_P4_Lumbar_Coeliac_Stellate_VS_Sphenopalatine | 1 | 1 | 1 | 1 | 1 | 1 | 1 | 0 |
| ZFP804A | 0.185 | P1_P2_P3_P4_Lumbar_Sphenopalatine_VS_Coeliac_Stellate | 1 | 1 | 1 | 1 | 1 | 0 | 0 | 1 |
| SERPINB9 | 0.185 | P1_P2_P3_P4_Coeliac_Sphenopalatine_VS_Lumbar_Stellate | 1 | 1 | 1 | 1 | 0 | 1 | 0 | 1 |
| TMEM169 | 0.185 | Lumbar_Coeliac_Sphenopalatine_VS_P1_P2_P3_P4_Stellate | 0 | 0 | 0 | 0 | 1 | 1 | 0 | 1 |
| TBC1D24 | 0.185 | P1_P2_P3_Lumbar_Coeliac_Sphenopalatine_VS_P4_Stellate | 1 | 1 | 1 | 0 | 1 | 1 | 0 | 1 |
| FAM110B | 0.185 | P1_P2_P4_Lumbar_Coeliac_Stellate_Sphenopalatine_VS_P3 | 1 | 1 | 0 | 1 | 1 | 1 | 1 | 1 |
| PPIC | 0.184 | P3_P4_VS_P1_P2_Lumbar_Coeliac_Stellate_Sphenopalatine | 0 | 0 | 1 | 1 | 0 | 0 | 0 | 0 |
| CREG1 | 0.184 | P3_P4_Lumbar_VS_P1_P2_Coeliac_Stellate_Sphenopalatine | 0 | 0 | 1 | 1 | 1 | 0 | 0 | 0 |
| SLC25A20 | 0.184 | P1_P2_P3_P4_Lumbar_Coeliac_Stellate_VS_Sphenopalatine | 1 | 1 | 1 | 1 | 1 | 1 | 1 | 0 |
| SIPA1L2 | 0.184 | P4_Lumbar_Sphenopalatine_VS_P1_P2_P3_Coeliac_Stellate | 0 | 0 | 0 | 1 | 1 | 0 | 0 | 1 |
| INTS3 | 0.184 | P1_P2_P3_Lumbar_Coeliac_Sphenopalatine_VS_P4_Stellate | 1 | 1 | 1 | 0 | 1 | 1 | 0 | 1 |
| PLOD2 | 0.184 | P1_P2_P3_Lumbar_Coeliac_Sphenopalatine_VS_P4_Stellate | 1 | 1 | 1 | 0 | 1 | 1 | 0 | 1 |
| KLHL18 | 0.184 | P1_P2_P3_Lumbar_Coeliac_Sphenopalatine_VS_P4_Stellate | 1 | 1 | 1 | 0 | 1 | 1 | 0 | 1 |
| HECA | 0.184 | P2_P4_Lumbar_Coeliac_Sphenopalatine_VS_P1_P3_Stellate | 0 | 1 | 0 | 1 | 1 | 1 | 0 | 1 |

| | | | | | | | | | | |
|---|---|---|---|---|---|---|---|---|---|---|
| ELMO2 | 0.183 | P1_P2_P3_P4_Lumbar_Sphenopalatine_VS_Coeliac_Stellate | 1 | 1 | 1 | 1 | 1 | 0 | 0 | 1 |
| MON2 | 0.183 | P1_P2_P3_Lumbar_Coeliac_Sphenopalatine_VS_P4_Stellate | 1 | 1 | 1 | 0 | 1 | 1 | 0 | 1 |
| RBMX2 | 0.183 | P1_P4_Lumbar_Coeliac_Sphenopalatine_VS_P2_P3_Stellate | 1 | 0 | 0 | 1 | 1 | 1 | 0 | 1 |
| KCNK1 | 0.183 | P1_P2_P3_Lumbar_Coeliac_Stellate_Sphenopalatine_VS_P4 | 1 | 1 | 1 | 0 | 1 | 1 | 1 | 1 |
| BOP1 | 0.182 | P1_P4_Lumbar_VS_P2_P3_Coeliac_Stellate_Sphenopalatine | 1 | 0 | 0 | 1 | 1 | 0 | 0 | 0 |
| RHOQ | 0.182 | P1_P2_P3_P4_Lumbar_VS_Coeliac_Stellate_Sphenopalatine | 1 | 1 | 1 | 1 | 1 | 0 | 0 | 0 |
| POLD2 | 0.182 | P1_P2_P3_P4_Lumbar_Coeliac_Stellate_VS_Sphenopalatine | 1 | 1 | 1 | 1 | 1 | 1 | 1 | 0 |
| TCEA2 | 0.182 | P1_P2_P3_P4_Sphenopalatine_VS_Lumbar_Coeliac_Stellate | 1 | 1 | 1 | 1 | 0 | 0 | 0 | 1 |
| CBX4 | 0.182 | P2_Lumbar_Coeliac_Sphenopalatine_VS_P1_P3_P4_Stellate | 0 | 1 | 0 | 0 | 1 | 1 | 0 | 1 |
| POLR3D | 0.182 | P3_Lumbar_Coeliac_Sphenopalatine_VS_P1_P2_P4_Stellate | 0 | 0 | 1 | 0 | 1 | 1 | 0 | 1 |
| TMEM56 | 0.182 | P1_P4_Lumbar_Coeliac_Sphenopalatine_VS_P2_P3_Stellate | 1 | 0 | 0 | 1 | 1 | 1 | 0 | 1 |
| FUT8 | 0.182 | P1_P3_P4_Lumbar_Coeliac_Stellate_Sphenopalatine_VS_P2 | 1 | 0 | 1 | 1 | 1 | 1 | 1 | 1 |
| IFT80 | 0.181 | P2_P3_P4_Coeliac_VS_P1_Lumbar_Stellate_Sphenopalatine | 0 | 1 | 1 | 1 | 0 | 1 | 0 | 0 |
| FYN | 0.181 | P4_Lumbar_Coeliac_Sphenopalatine_VS_P1_P2_P3_Stellate | 0 | 0 | 0 | 1 | 1 | 1 | 0 | 1 |
| AI413582 | 0.181 | P1_P2_P3_P4_Lumbar_Stellate_Sphenopalatine_VS_Coeliac | 1 | 1 | 1 | 1 | 1 | 0 | 1 | 1 |
| RGMB | 0.181 | P3_P4_Lumbar_Coeliac_Stellate_Sphenopalatine_VS_P1_P2 | 0 | 0 | 1 | 1 | 1 | 1 | 1 | 1 |
| MACROD1 | 0.18 | P3_Coeliac_VS_P1_P2_P4_Lumbar_Stellate_Sphenopalatine | 0 | 0 | 1 | 0 | 0 | 1 | 0 | 0 |
| NFKBIA | 0.18 | P1_P2_P3_P4_Coeliac_VS_Lumbar_Stellate_Sphenopalatine | 1 | 1 | 1 | 1 | 0 | 1 | 0 | 0 |
| RAB26OS | 0.18 | P3_Sphenopalatine_VS_P1_P2_P4_Lumbar_Coeliac_Stellate | 0 | 0 | 1 | 0 | 0 | 0 | 0 | 1 |
| GRIK4 | 0.18 | P4_Sphenopalatine_VS_P1_P2_P3_Lumbar_Coeliac_Stellate | 0 | 0 | 0 | 1 | 0 | 0 | 0 | 1 |
| C530008M17 | 0.18 | P4_Lumbar_Coeliac_Sphenopalatine_VS_P1_P2_P3_Stellate | 0 | 0 | 0 | 1 | 1 | 1 | 0 | 1 |
| TANK | 0.179 | P3_Lumbar_VS_P1_P2_P4_Coeliac_Stellate_Sphenopalatine | 0 | 0 | 1 | 0 | 1 | 0 | 0 | 0 |
| FKBPL | 0.179 | P1_P4_Lumbar_VS_P2_P3_Coeliac_Stellate_Sphenopalatine | 1 | 0 | 0 | 1 | 1 | 0 | 0 | 0 |
| ZFP740 | 0.179 | P1_P2_P3_P4_Coeliac_Sphenopalatine_VS_Lumbar_Stellate | 1 | 1 | 1 | 1 | 0 | 1 | 0 | 1 |
| GSAP | 0.178 | P2_P3_P4_VS_P1_Lumbar_Coeliac_Stellate_Sphenopalatine | 0 | 1 | 1 | 1 | 0 | 0 | 0 | 0 |
| NPR2 | 0.178 | P1_P2_P4_Lumbar_VS_P3_Coeliac_Stellate_Sphenopalatine | 1 | 1 | 0 | 1 | 1 | 0 | 0 | 0 |
| CSPG5 | 0.178 | P3_P4_Lumbar_VS_P1_P2_Coeliac_Stellate_Sphenopalatine | 0 | 0 | 1 | 1 | 1 | 0 | 0 | 0 |
| SLC25A30 | 0.178 | P3_P4_Lumbar_VS_P1_P2_Coeliac_Stellate_Sphenopalatine | 0 | 0 | 1 | 1 | 1 | 0 | 0 | 0 |
| IRS2 | 0.178 | P1_P2_P3_Lumbar_Coeliac_Sphenopalatine_VS_P4_Stellate | 1 | 1 | 1 | 0 | 1 | 1 | 0 | 1 |
| TAB1 | 0.178 | P1_P2_P3_Lumbar_Coeliac_Sphenopalatine_VS_P4_Stellate | 1 | 1 | 1 | 0 | 1 | 1 | 0 | 1 |

| | | | | | | | | | | |
|---|---|---|---|---|---|---|---|---|---|---|
| SLC35B4 | 0.177 | P1_P2_P3_P4_Coeliac_Sphenopalatine_VS_Lumbar_Stellate | 1 | 1 | 1 | 1 | 0 | 1 | 0 | 1 |
| HIRA | 0.177 | P1_P2_Lumbar_Coeliac_Sphenopalatine_VS_P3_P4_Stellate | 1 | 1 | 0 | 0 | 1 | 1 | 0 | 1 |
| KDM4B | 0.177 | P1_P2_Lumbar_Coeliac_Sphenopalatine_VS_P3_P4_Stellate | 1 | 1 | 0 | 0 | 1 | 1 | 0 | 1 |
| CDKL3 | 0.177 | P1_P2_P3_Lumbar_Coeliac_Sphenopalatine_VS_P4_Stellate | 1 | 1 | 1 | 0 | 1 | 1 | 0 | 1 |
| FBXO34 | 0.177 | P1_P4_Lumbar_Coeliac_Sphenopalatine_VS_P2_P3_Stellate | 1 | 0 | 0 | 1 | 1 | 1 | 0 | 1 |
| TBC1D19 | 0.176 | P3_Lumbar_VS_P1_P2_P4_Coeliac_Stellate_Sphenopalatine | 0 | 0 | 1 | 0 | 1 | 0 | 0 | 0 |
| SEC16A | 0.176 | P1_P2_P3_Lumbar_Coeliac_Sphenopalatine_VS_P4_Stellate | 1 | 1 | 1 | 0 | 1 | 1 | 0 | 1 |
| VAV3 | 0.175 | P1_VS_P2_P3_P4_Lumbar_Coeliac_Stellate_Sphenopalatine | 1 | 0 | 0 | 0 | 0 | 0 | 0 | 0 |
| CDPF1 | 0.175 | P3_P4_VS_P1_P2_Lumbar_Coeliac_Stellate_Sphenopalatine | 0 | 0 | 1 | 1 | 0 | 0 | 0 | 0 |
| SH3GL2 | 0.175 | P1_P2_P3_P4_Sphenopalatine_VS_Lumbar_Coeliac_Stellate | 1 | 1 | 1 | 1 | 0 | 0 | 0 | 1 |
| VPS37D | 0.175 | P1_P4_Lumbar_Coeliac_Sphenopalatine_VS_P2_P3_Stellate | 1 | 0 | 0 | 1 | 1 | 1 | 0 | 1 |
| CTC1 | 0.175 | P1_P4_Lumbar_Coeliac_Sphenopalatine_VS_P2_P3_Stellate | 1 | 0 | 0 | 1 | 1 | 1 | 0 | 1 |
| SMAD5 | 0.175 | P1_P4_Lumbar_Coeliac_Sphenopalatine_VS_P2_P3_Stellate | 1 | 0 | 0 | 1 | 1 | 1 | 0 | 1 |
| TENT4A | 0.175 | P2_P4_Lumbar_Coeliac_Sphenopalatine_VS_P1_P3_Stellate | 0 | 1 | 0 | 1 | 1 | 1 | 0 | 1 |
| INTS6 | 0.174 | P2_P4_Coeliac_VS_P1_P3_Lumbar_Stellate_Sphenopalatine | 0 | 1 | 0 | 1 | 0 | 1 | 0 | 0 |
| BAG3 | 0.174 | P1_P2_P3_P4_Lumbar_Coeliac_Stellate_VS_Sphenopalatine | 1 | 1 | 1 | 1 | 1 | 1 | 1 | 0 |
| GPRIN1 | 0.174 | P4_Lumbar_Coeliac_Sphenopalatine_VS_P1_P2_P3_Stellate | 0 | 0 | 0 | 1 | 1 | 1 | 0 | 1 |
| PRPF18 | 0.174 | P2_P4_Lumbar_Coeliac_Sphenopalatine_VS_P1_P3_Stellate | 0 | 1 | 0 | 1 | 1 | 1 | 0 | 1 |
| SHISA7 | 0.173 | Lumbar_Coeliac_Stellate_VS_P1_P2_P3_P4_Sphenopalatine | 0 | 0 | 0 | 0 | 1 | 1 | 1 | 0 |
| IBTK | 0.173 | P1_P2_P3_P4_Lumbar_Coeliac_Stellate_VS_Sphenopalatine | 1 | 1 | 1 | 1 | 1 | 1 | 1 | 0 |
| INTS10 | 0.173 | Lumbar_Coeliac_Sphenopalatine_VS_P1_P2_P3_P4_Stellate | 0 | 0 | 0 | 0 | 1 | 1 | 0 | 1 |
| CSRNP2 | 0.173 | Lumbar_Coeliac_Sphenopalatine_VS_P1_P2_P3_P4_Stellate | 0 | 0 | 0 | 0 | 1 | 1 | 0 | 1 |
| RBPJ | 0.173 | P1_P2_P3_Lumbar_Coeliac_Sphenopalatine_VS_P4_Stellate | 1 | 1 | 1 | 0 | 1 | 1 | 0 | 1 |
| POLR1A | 0.173 | P1_P2_P3_Lumbar_Coeliac_Sphenopalatine_VS_P4_Stellate | 1 | 1 | 1 | 0 | 1 | 1 | 0 | 1 |
| TCEAL5 | 0.173 | P4_Lumbar_Coeliac_Sphenopalatine_VS_P1_P2_P3_Stellate | 0 | 0 | 0 | 1 | 1 | 1 | 0 | 1 |
| SPINK10 | 0.173 | P1_P2_P3_Lumbar_Coeliac_Stellate_Sphenopalatine_VS_P4 | 1 | 1 | 1 | 0 | 1 | 1 | 1 | 1 |
| HEY2 | 0.172 | P3_P4_VS_P1_P2_Lumbar_Coeliac_Stellate_Sphenopalatine | 0 | 0 | 1 | 1 | 0 | 0 | 0 | 0 |
| GM30025 | 0.172 | P3_P4_VS_P1_P2_Lumbar_Coeliac_Stellate_Sphenopalatine | 0 | 0 | 1 | 1 | 0 | 0 | 0 | 0 |
| ARHGEF25 | 0.172 | P2_P4_Lumbar_VS_P1_P3_Coeliac_Stellate_Sphenopalatine | 0 | 1 | 0 | 1 | 1 | 0 | 0 | 0 |
| GM4631 | 0.172 | P1_P2_P3_P4_Coeliac_VS_Lumbar_Stellate_Sphenopalatine | 1 | 1 | 1 | 1 | 0 | 1 | 0 | 0 |

| | | | | | | | | | | |
|---|---|---|---|---|---|---|---|---|---|---|
| ARHGAP22 | 0.172 | Sphenopalatine_VS_P1_P2_P3_P4_Lumbar_Coeliac_Stellate | 0 | 0 | 0 | 0 | 0 | 0 | 0 | 1 |
| ZBED3 | 0.171 | P3_VS_P1_P2_P4_Lumbar_Coeliac_Stellate_Sphenopalatine | 0 | 0 | 1 | 0 | 0 | 0 | 0 | 0 |
| ZFP687 | 0.171 | P1_P2_P3_P4_Lumbar_VS_Coeliac_Stellate_Sphenopalatine | 1 | 1 | 1 | 1 | 1 | 0 | 0 | 0 |
| ACKR1 | 0.171 | P2_P4_Lumbar_Sphenopalatine_VS_P1_P3_Coeliac_Stellate | 0 | 1 | 0 | 1 | 1 | 0 | 0 | 1 |
| IAH1 | 0.171 | P1_P2_P4_Lumbar_Sphenopalatine_VS_P3_Coeliac_Stellate | 1 | 1 | 0 | 1 | 1 | 0 | 0 | 1 |
| D130017N08 | 0.171 | P1_P2_P3_Lumbar_Coeliac_Sphenopalatine_VS_P4_Stellate | 1 | 1 | 1 | 0 | 1 | 1 | 0 | 1 |
| OGDHL | 0.171 | P2_P4_Lumbar_Coeliac_Sphenopalatine_VS_P1_P3_Stellate | 0 | 1 | 0 | 1 | 1 | 1 | 0 | 1 |
| ARHGAP12 | 0.171 | P2_P4_Lumbar_Coeliac_Sphenopalatine_VS_P1_P3_Stellate | 0 | 1 | 0 | 1 | 1 | 1 | 0 | 1 |
| HINT3 | 0.17 | P1_P4_Lumbar_VS_P2_P3_Coeliac_Stellate_Sphenopalatine | 1 | 0 | 0 | 1 | 1 | 0 | 0 | 0 |
| PEX7 | 0.17 | P1_P2_P3_P4_Lumbar_VS_Coeliac_Stellate_Sphenopalatine | 1 | 1 | 1 | 1 | 1 | 0 | 0 | 0 |
| KLF11 | 0.17 | P2_P3_P4_Lumbar_Coeliac_Stellate_VS_P1_Sphenopalatine | 0 | 1 | 1 | 1 | 1 | 1 | 1 | 0 |
| RNF141 | 0.17 | P1_P2_P3_P4_Coeliac_Sphenopalatine_VS_Lumbar_Stellate | 1 | 1 | 1 | 1 | 0 | 1 | 0 | 1 |
| RSBN1L | 0.17 | P1_P4_Lumbar_Coeliac_Sphenopalatine_VS_P2_P3_Stellate | 1 | 0 | 0 | 1 | 1 | 1 | 0 | 1 |
| TMEM238 | 0.169 | P1_P4_Lumbar_Sphenopalatine_VS_P2_P3_Coeliac_Stellate | 1 | 0 | 0 | 1 | 1 | 0 | 0 | 1 |
| TRIM36 | 0.169 | P1_P2_P3_Lumbar_Coeliac_Sphenopalatine_VS_P4_Stellate | 1 | 1 | 1 | 0 | 1 | 1 | 0 | 1 |
| CARS | 0.169 | P1_P4_Lumbar_Coeliac_Sphenopalatine_VS_P2_P3_Stellate | 1 | 0 | 0 | 1 | 1 | 1 | 0 | 1 |
| PPP6R2 | 0.169 | P1_P4_Lumbar_Coeliac_Sphenopalatine_VS_P2_P3_Stellate | 1 | 0 | 0 | 1 | 1 | 1 | 0 | 1 |
| TPST1 | 0.169 | P1_P3_Lumbar_Coeliac_Stellate_Sphenopalatine_VS_P2_P4 | 1 | 0 | 1 | 0 | 1 | 1 | 1 | 1 |
| SLC6A8 | 0.168 | P1_P2_P3_P4_Coeliac_VS_Lumbar_Stellate_Sphenopalatine | 1 | 1 | 1 | 1 | 0 | 1 | 0 | 0 |
| PCSK1 | 0.167 | P3_P4_VS_P1_P2_Lumbar_Coeliac_Stellate_Sphenopalatine | 0 | 0 | 1 | 1 | 0 | 0 | 0 | 0 |
| E2F5 | 0.167 | P1_P2_P3_P4_Lumbar_Coeliac_Stellate_VS_Sphenopalatine | 1 | 1 | 1 | 1 | 1 | 1 | 1 | 0 |
| ALDH9A1 | 0.166 | P1_P2_P3_P4_Lumbar_Coeliac_Stellate_VS_Sphenopalatine | 1 | 1 | 1 | 1 | 1 | 1 | 1 | 0 |
| MCUB | 0.166 | P1_P2_P3_P4_Sphenopalatine_VS_Lumbar_Coeliac_Stellate | 1 | 1 | 1 | 1 | 0 | 0 | 0 | 1 |
| DTNBP1 | 0.166 | P1_P3_P4_Lumbar_Sphenopalatine_VS_P2_Coeliac_Stellate | 1 | 0 | 1 | 1 | 1 | 0 | 0 | 1 |
| THNSL1 | 0.166 | P1_P4_Lumbar_Coeliac_Sphenopalatine_VS_P2_P3_Stellate | 1 | 0 | 0 | 1 | 1 | 1 | 0 | 1 |
| FPGS | 0.166 | P1_P4_Lumbar_Coeliac_Sphenopalatine_VS_P2_P3_Stellate | 1 | 0 | 0 | 1 | 1 | 1 | 0 | 1 |
| UBXN7 | 0.166 | P1_P4_Lumbar_Coeliac_Sphenopalatine_VS_P2_P3_Stellate | 1 | 0 | 0 | 1 | 1 | 1 | 0 | 1 |
| OGFRL1 | 0.166 | P3_P4_Lumbar_Coeliac_Sphenopalatine_VS_P1_P2_Stellate | 0 | 0 | 1 | 1 | 1 | 1 | 0 | 1 |
| PPFIA2 | 0.165 | P3_P4_Lumbar_VS_P1_P2_Coeliac_Stellate_Sphenopalatine | 0 | 0 | 1 | 1 | 1 | 0 | 0 | 0 |
| ADARB1 | 0.165 | P1_P2_P3_Lumbar_Coeliac_Sphenopalatine_VS_P4_Stellate | 1 | 1 | 1 | 0 | 1 | 1 | 0 | 1 |

| | | | | | | | | | | |
|---|---|---|---|---|---|---|---|---|---|---|
| SRC | 0.165 | P2_P4_Lumbar_Coeliac_Sphenopalatine_VS_P1_P3_Stellate | 0 | 1 | 0 | 1 | 1 | 1 | 0 | 1 |
| IRF2BP2 | 0.165 | P2_P4_Lumbar_Coeliac_Sphenopalatine_VS_P1_P3_Stellate | 0 | 1 | 0 | 1 | 1 | 1 | 0 | 1 |
| IGDCC4 | 0.164 | P1_P2_P3_VS_P4_Lumbar_Coeliac_Stellate_Sphenopalatine | 1 | 1 | 1 | 0 | 0 | 0 | 0 | 0 |
| SMIM27 | 0.164 | P3_P4_VS_P1_P2_Lumbar_Coeliac_Stellate_Sphenopalatine | 0 | 0 | 1 | 1 | 0 | 0 | 0 | 0 |
| DBP | 0.164 | P3_P4_VS_P1_P2_Lumbar_Coeliac_Stellate_Sphenopalatine | 0 | 0 | 1 | 1 | 0 | 0 | 0 | 0 |
| VPS33A | 0.164 | P2_Lumbar_VS_P1_P3_P4_Coeliac_Stellate_Sphenopalatine | 0 | 1 | 0 | 0 | 1 | 0 | 0 | 0 |
| NIF3L1 | 0.164 | P3_Lumbar_VS_P1_P2_P4_Coeliac_Stellate_Sphenopalatine | 0 | 0 | 1 | 0 | 1 | 0 | 0 | 0 |
| ZFP668 | 0.164 | P3_Lumbar_VS_P1_P2_P4_Coeliac_Stellate_Sphenopalatine | 0 | 0 | 1 | 0 | 1 | 0 | 0 | 0 |
| ZBTB8OS | 0.164 | P1_P4_Lumbar_VS_P2_P3_Coeliac_Stellate_Sphenopalatine | 1 | 0 | 0 | 1 | 1 | 0 | 0 | 0 |
| MARCH4 | 0.164 | P2_Lumbar_Coeliac_Stellate_VS_P1_P3_P4_Sphenopalatine | 0 | 1 | 0 | 0 | 1 | 1 | 1 | 0 |
| LRRC73 | 0.164 | P3_Sphenopalatine_VS_P1_P2_P4_Lumbar_Coeliac_Stellate | 0 | 0 | 1 | 0 | 0 | 0 | 0 | 1 |
| PHF21A | 0.164 | P1_P2_P3_Lumbar_Coeliac_Sphenopalatine_VS_P4_Stellate | 1 | 1 | 1 | 0 | 1 | 1 | 0 | 1 |
| CCNG2 | 0.164 | P1_P2_P3_Lumbar_Coeliac_Sphenopalatine_VS_P4_Stellate | 1 | 1 | 1 | 0 | 1 | 1 | 0 | 1 |
| REV3L | 0.164 | P2_P4_Lumbar_Coeliac_Sphenopalatine_VS_P1_P3_Stellate | 0 | 1 | 0 | 1 | 1 | 1 | 0 | 1 |
| DDAH1 | 0.164 | P1_P2_P4_Lumbar_Stellate_Sphenopalatine_VS_P3_Coeliac | 1 | 1 | 0 | 1 | 1 | 0 | 1 | 1 |
| CDR2 | 0.163 | P1_P2_P3_P4_Lumbar_VS_Coeliac_Stellate_Sphenopalatine | 1 | 1 | 1 | 1 | 1 | 0 | 0 | 0 |
| SLMAP | 0.163 | P1_P2_P3_P4_Lumbar_VS_Coeliac_Stellate_Sphenopalatine | 1 | 1 | 1 | 1 | 1 | 0 | 0 | 0 |
| ARFIP1 | 0.163 | P1_P2_P3_P4_Coeliac_VS_Lumbar_Stellate_Sphenopalatine | 1 | 1 | 1 | 1 | 0 | 1 | 0 | 0 |
| BRDT | 0.163 | P3_P4_Lumbar_Coeliac_Stellate_VS_P1_P2_Sphenopalatine | 0 | 0 | 1 | 1 | 1 | 1 | 1 | 0 |
| PYCR2 | 0.163 | P1_P2_P3_P4_Lumbar_Coeliac_Stellate_VS_Sphenopalatine | 1 | 1 | 1 | 1 | 1 | 1 | 1 | 0 |
| IRF2 | 0.163 | P1_P2_P3_P4_Lumbar_Coeliac_Stellate_VS_Sphenopalatine | 1 | 1 | 1 | 1 | 1 | 1 | 1 | 0 |
| SGCE | 0.163 | P1_P4_Lumbar_Coeliac_Sphenopalatine_VS_P2_P3_Stellate | 1 | 0 | 0 | 1 | 1 | 1 | 0 | 1 |
| INPP5A | 0.163 | P2_P4_Lumbar_Coeliac_Sphenopalatine_VS_P1_P3_Stellate | 0 | 1 | 0 | 1 | 1 | 1 | 0 | 1 |
| DOCK11 | 0.163 | P2_P4_Lumbar_Coeliac_Sphenopalatine_VS_P1_P3_Stellate | 0 | 1 | 0 | 1 | 1 | 1 | 0 | 1 |
| TRPM4 | 0.162 | P3_P4_VS_P1_P2_Lumbar_Coeliac_Stellate_Sphenopalatine | 0 | 0 | 1 | 1 | 0 | 0 | 0 | 0 |
| RPAP1 | 0.162 | P1_Lumbar_VS_P2_P3_P4_Coeliac_Stellate_Sphenopalatine | 1 | 0 | 0 | 0 | 1 | 0 | 0 | 0 |
| RUVBL2 | 0.162 | P1_P3_P4_Lumbar_Coeliac_Stellate_VS_P2_Sphenopalatine | 1 | 0 | 1 | 1 | 1 | 1 | 1 | 0 |
| LDLRAD3 | 0.162 | P1_P2_P4_Lumbar_Sphenopalatine_VS_P3_Coeliac_Stellate | 1 | 1 | 0 | 1 | 1 | 0 | 0 | 1 |
| 5730409E04I | 0.162 | P1_P2_P3_P4_Coeliac_Sphenopalatine_VS_Lumbar_Stellate | 1 | 1 | 1 | 1 | 0 | 1 | 0 | 1 |
| GOSR1 | 0.162 | P1_P4_Lumbar_Coeliac_Sphenopalatine_VS_P2_P3_Stellate | 1 | 0 | 0 | 1 | 1 | 1 | 0 | 1 |

| | | | | | | | | | | |
|---|---|---|---|---|---|---|---|---|---|---|
| B2M | 0.162 | P1_P2_P3_P4_Lumbar_Stellate_Sphenopalatine_VS_Coeliac | 1 | 1 | 1 | 1 | 1 | 0 | 1 | 1 |
| POU2F2 | 0.161 | P2_P3_P4_Lumbar_Coeliac_Stellate_VS_P1_Sphenopalatine | 0 | 1 | 1 | 1 | 1 | 1 | 1 | 0 |
| ARRDC3 | 0.161 | P3_Lumbar_Coeliac_Sphenopalatine_VS_P1_P2_P4_Stellate | 0 | 0 | 1 | 0 | 1 | 1 | 0 | 1 |
| ZFP30 | 0.161 | P1_P2_P3_Lumbar_Coeliac_Sphenopalatine_VS_P4_Stellate | 1 | 1 | 1 | 0 | 1 | 1 | 0 | 1 |
| PHF12 | 0.161 | P1_P2_P3_Lumbar_Coeliac_Sphenopalatine_VS_P4_Stellate | 1 | 1 | 1 | 0 | 1 | 1 | 0 | 1 |
| ZC3H7A | 0.161 | P1_P2_P3_Lumbar_Coeliac_Sphenopalatine_VS_P4_Stellate | 1 | 1 | 1 | 0 | 1 | 1 | 0 | 1 |
| PHF10 | 0.161 | P1_P2_P3_Lumbar_Coeliac_Sphenopalatine_VS_P4_Stellate | 1 | 1 | 1 | 0 | 1 | 1 | 0 | 1 |
| ZMYND8 | 0.161 | P4_Lumbar_Coeliac_Sphenopalatine_VS_P1_P2_P3_Stellate | 0 | 0 | 0 | 1 | 1 | 1 | 0 | 1 |
| DNMT1 | 0.161 | P1_P4_Lumbar_Coeliac_Sphenopalatine_VS_P2_P3_Stellate | 1 | 0 | 0 | 1 | 1 | 1 | 0 | 1 |
| MED27 | 0.16 | P1_P3_P4_Lumbar_VS_P2_Coeliac_Stellate_Sphenopalatine | 1 | 0 | 1 | 1 | 1 | 0 | 0 | 0 |
| MKNK2 | 0.16 | P1_P2_P3_P4_Lumbar_VS_Coeliac_Stellate_Sphenopalatine | 1 | 1 | 1 | 1 | 1 | 0 | 0 | 0 |
| SNX13 | 0.16 | P1_P2_P3_Lumbar_Coeliac_Sphenopalatine_VS_P4_Stellate | 1 | 1 | 1 | 0 | 1 | 1 | 0 | 1 |
| PCGF2 | 0.16 | P2_P4_Lumbar_Coeliac_Sphenopalatine_VS_P1_P3_Stellate | 0 | 1 | 0 | 1 | 1 | 1 | 0 | 1 |
| ADI1 | 0.159 | P1_P2_P3_P4_Sphenopalatine_VS_Lumbar_Coeliac_Stellate | 1 | 1 | 1 | 1 | 0 | 0 | 0 | 1 |
| SLC4A2 | 0.159 | P1_P2_P3_P4_Coeliac_Sphenopalatine_VS_Lumbar_Stellate | 1 | 1 | 1 | 1 | 0 | 1 | 0 | 1 |
| LST1 | 0.159 | P1_P2_P3_P4_Coeliac_Sphenopalatine_VS_Lumbar_Stellate | 1 | 1 | 1 | 1 | 0 | 1 | 0 | 1 |
| KHDRBS3 | 0.159 | P1_P2_P3_Lumbar_Coeliac_Sphenopalatine_VS_P4_Stellate | 1 | 1 | 1 | 0 | 1 | 1 | 0 | 1 |
| TAF15 | 0.159 | P4_Lumbar_Coeliac_Sphenopalatine_VS_P1_P2_P3_Stellate | 0 | 0 | 0 | 1 | 1 | 1 | 0 | 1 |
| RUFY1 | 0.159 | P1_P4_Lumbar_Coeliac_Sphenopalatine_VS_P2_P3_Stellate | 1 | 0 | 0 | 1 | 1 | 1 | 0 | 1 |
| TBK1 | 0.159 | P2_P4_Lumbar_Coeliac_Sphenopalatine_VS_P1_P3_Stellate | 0 | 1 | 0 | 1 | 1 | 1 | 0 | 1 |
| CHL1 | 0.159 | P1_P2_P3_Lumbar_Coeliac_Stellate_Sphenopalatine_VS_P4 | 1 | 1 | 1 | 0 | 1 | 1 | 1 | 1 |
| DHFR | 0.158 | P3_P4_VS_P1_P2_Lumbar_Coeliac_Stellate_Sphenopalatine | 0 | 0 | 1 | 1 | 0 | 0 | 0 | 0 |
| ZFP386 | 0.158 | P3_Lumbar_VS_P1_P2_P4_Coeliac_Stellate_Sphenopalatine | 0 | 0 | 1 | 0 | 1 | 0 | 0 | 0 |
| EXOSC5 | 0.158 | P1_P4_Lumbar_VS_P2_P3_Coeliac_Stellate_Sphenopalatine | 1 | 0 | 0 | 1 | 1 | 0 | 0 | 0 |
| ADAM23 | 0.158 | P1_P2_P3_P4_Lumbar_VS_Coeliac_Stellate_Sphenopalatine | 1 | 1 | 1 | 1 | 1 | 0 | 0 | 0 |
| GM45353 | 0.158 | P2_Coeliac_VS_P1_P3_P4_Lumbar_Stellate_Sphenopalatine | 0 | 1 | 0 | 0 | 0 | 1 | 0 | 0 |
| CPEB1 | 0.158 | P1_P2_P4_Sphenopalatine_VS_P3_Lumbar_Coeliac_Stellate | 1 | 1 | 0 | 1 | 0 | 0 | 0 | 1 |
| SRXN1 | 0.158 | P1_P2_P3_P4_Sphenopalatine_VS_Lumbar_Coeliac_Stellate | 1 | 1 | 1 | 1 | 0 | 0 | 0 | 1 |
| CDK17 | 0.158 | P1_P3_Lumbar_Coeliac_Sphenopalatine_VS_P2_P4_Stellate | 1 | 0 | 1 | 0 | 1 | 1 | 0 | 1 |
| LGI1 | 0.158 | P1_P2_P3_Lumbar_Coeliac_Sphenopalatine_VS_P4_Stellate | 1 | 1 | 1 | 0 | 1 | 1 | 0 | 1 |

| | | | | | | | | | | |
|---|---|---|---|---|---|---|---|---|---|---|
| HJURP | 0.158 | P1_P4_Lumbar_Coeliac_Sphenopalatine_VS_P2_P3_Stellate | 1 | 0 | 0 | 1 | 1 | 1 | 0 | 1 |
| SPTBN4 | 0.158 | P2_P4_Lumbar_Coeliac_Sphenopalatine_VS_P1_P3_Stellate | 0 | 1 | 0 | 1 | 1 | 1 | 0 | 1 |
| LARP4 | 0.158 | P2_P4_Lumbar_Coeliac_Sphenopalatine_VS_P1_P3_Stellate | 0 | 1 | 0 | 1 | 1 | 1 | 0 | 1 |
| CAVIN2 | 0.157 | P1_VS_P2_P3_P4_Lumbar_Coeliac_Stellate_Sphenopalatine | 1 | 0 | 0 | 0 | 0 | 0 | 0 | 0 |
| DNPH1 | 0.157 | P3_Lumbar_Coeliac_Stellate_VS_P1_P2_P4_Sphenopalatine | 0 | 0 | 1 | 0 | 1 | 1 | 1 | 0 |
| MXRA7 | 0.157 | P1_P2_P4_Lumbar_Coeliac_Stellate_Sphenopalatine_VS_P3 | 1 | 1 | 0 | 1 | 1 | 1 | 1 | 1 |
| HPCAL4 | 0.156 | P3_VS_P1_P2_P4_Lumbar_Coeliac_Stellate_Sphenopalatine | 0 | 0 | 1 | 0 | 0 | 0 | 0 | 0 |
| SMARCD2 | 0.156 | P1_P2_P3_P4_Lumbar_Coeliac_Stellate_VS_Sphenopalatine | 1 | 1 | 1 | 1 | 1 | 1 | 1 | 0 |
| GON4L | 0.156 | P4_Lumbar_Coeliac_Sphenopalatine_VS_P1_P2_P3_Stellate | 0 | 0 | 0 | 1 | 1 | 1 | 0 | 1 |
| CNOT10 | 0.156 | P4_Lumbar_Coeliac_Sphenopalatine_VS_P1_P2_P3_Stellate | 0 | 0 | 0 | 1 | 1 | 1 | 0 | 1 |
| RNF111 | 0.156 | P2_P4_Lumbar_Coeliac_Sphenopalatine_VS_P1_P3_Stellate | 0 | 1 | 0 | 1 | 1 | 1 | 0 | 1 |
| CAAP1 | 0.156 | P1_P3_P4_Lumbar_Coeliac_Stellate_Sphenopalatine_VS_P2 | 1 | 0 | 1 | 1 | 1 | 1 | 1 | 1 |
| SUOX | 0.155 | P3_VS_P1_P2_P4_Lumbar_Coeliac_Stellate_Sphenopalatine | 0 | 0 | 1 | 0 | 0 | 0 | 0 | 0 |
| TELO2 | 0.155 | P3_Lumbar_VS_P1_P2_P4_Coeliac_Stellate_Sphenopalatine | 0 | 0 | 1 | 0 | 1 | 0 | 0 | 0 |
| UXS1 | 0.155 | P3_P4_Lumbar_VS_P1_P2_Coeliac_Stellate_Sphenopalatine | 0 | 0 | 1 | 1 | 1 | 0 | 0 | 0 |
| GMEB2 | 0.155 | P1_P2_P3_Lumbar_Coeliac_Sphenopalatine_VS_P4_Stellate | 1 | 1 | 1 | 0 | 1 | 1 | 0 | 1 |
| ANKS1 | 0.155 | P4_Lumbar_Coeliac_Sphenopalatine_VS_P1_P2_P3_Stellate | 0 | 0 | 0 | 1 | 1 | 1 | 0 | 1 |
| VIM | 0.155 | P1_P3_P4_Lumbar_Coeliac_Stellate_Sphenopalatine_VS_P2 | 1 | 0 | 1 | 1 | 1 | 1 | 1 | 1 |
| KLHDC1 | 0.154 | P3_P4_VS_P1_P2_Lumbar_Coeliac_Stellate_Sphenopalatine | 0 | 0 | 1 | 1 | 0 | 0 | 0 | 0 |
| ZFYVE21 | 0.154 | P3_P4_VS_P1_P2_Lumbar_Coeliac_Stellate_Sphenopalatine | 0 | 0 | 1 | 1 | 0 | 0 | 0 | 0 |
| CHAC1 | 0.154 | P1_P4_Sphenopalatine_VS_P2_P3_Lumbar_Coeliac_Stellate | 1 | 0 | 0 | 1 | 0 | 0 | 0 | 1 |
| LDB1 | 0.154 | P4_Lumbar_Coeliac_Sphenopalatine_VS_P1_P2_P3_Stellate | 0 | 0 | 0 | 1 | 1 | 1 | 0 | 1 |
| MEIG1 | 0.153 | P3_VS_P1_P2_P4_Lumbar_Coeliac_Stellate_Sphenopalatine | 0 | 0 | 1 | 0 | 0 | 0 | 0 | 0 |
| NOA1 | 0.153 | P1_Lumbar_VS_P2_P3_P4_Coeliac_Stellate_Sphenopalatine | 1 | 0 | 0 | 0 | 1 | 0 | 0 | 0 |
| TRIM13 | 0.153 | Lumbar_Coeliac_Stellate_VS_P1_P2_P3_P4_Sphenopalatine | 0 | 0 | 0 | 0 | 1 | 1 | 1 | 0 |
| DRP2 | 0.153 | P4_Sphenopalatine_VS_P1_P2_P3_Lumbar_Coeliac_Stellate | 0 | 0 | 0 | 1 | 0 | 0 | 0 | 1 |
| DLG2 | 0.153 | P1_P2_P3_P4_Coeliac_Sphenopalatine_VS_Lumbar_Stellate | 1 | 1 | 1 | 1 | 0 | 1 | 0 | 1 |
| MICU2 | 0.153 | Lumbar_Coeliac_Sphenopalatine_VS_P1_P2_P3_P4_Stellate | 0 | 0 | 0 | 0 | 1 | 1 | 0 | 1 |
| USP12 | 0.153 | P1_P2_P3_Lumbar_Coeliac_Sphenopalatine_VS_P4_Stellate | 1 | 1 | 1 | 0 | 1 | 1 | 0 | 1 |
| SLC4A3 | 0.153 | P4_Lumbar_Coeliac_Sphenopalatine_VS_P1_P2_P3_Stellate | 0 | 0 | 0 | 1 | 1 | 1 | 0 | 1 |

| | | | | | | | | | | |
|---|---|---|---|---|---|---|---|---|---|---|
| REEP3 | 0.153 | P4_Lumbar_Coeliac_Sphenopalatine_VS_P1_P2_P3_Stellate | 0 | 0 | 0 | 1 | 1 | 1 | 0 | 1 |
| TM9SF4 | 0.153 | P2_P4_Lumbar_Coeliac_Sphenopalatine_VS_P1_P3_Stellate | 0 | 1 | 0 | 1 | 1 | 1 | 0 | 1 |
| SLITRK3 | 0.153 | P3_P4_Lumbar_Coeliac_Sphenopalatine_VS_P1_P2_Stellate | 0 | 0 | 1 | 1 | 1 | 1 | 0 | 1 |
| HMGB2 | 0.153 | P1_P3_P4_Lumbar_Coeliac_Stellate_Sphenopalatine_VS_P2 | 1 | 0 | 1 | 1 | 1 | 1 | 1 | 1 |
| CTU2 | 0.152 | P1_Lumbar_VS_P2_P3_P4_Coeliac_Stellate_Sphenopalatine | 1 | 0 | 0 | 0 | 1 | 0 | 0 | 0 |
| RAB40B | 0.152 | P1_Lumbar_VS_P2_P3_P4_Coeliac_Stellate_Sphenopalatine | 1 | 0 | 0 | 0 | 1 | 0 | 0 | 0 |
| KMT2D | 0.152 | P2_P4_Coeliac_VS_P1_P3_Lumbar_Stellate_Sphenopalatine | 0 | 1 | 0 | 1 | 0 | 1 | 0 | 0 |
| RITA1 | 0.152 | P3_Lumbar_Coeliac_Stellate_VS_P1_P2_P4_Sphenopalatine | 0 | 0 | 1 | 0 | 1 | 1 | 1 | 0 |
| SEMA6C | 0.152 | P1_P2_P3_Lumbar_Coeliac_Sphenopalatine_VS_P4_Stellate | 1 | 1 | 1 | 0 | 1 | 1 | 0 | 1 |
| GM16536 | 0.152 | P1_P2_P3_Lumbar_Coeliac_Sphenopalatine_VS_P4_Stellate | 1 | 1 | 1 | 0 | 1 | 1 | 0 | 1 |
| CHM | 0.152 | P1_P2_P3_Lumbar_Coeliac_Sphenopalatine_VS_P4_Stellate | 1 | 1 | 1 | 0 | 1 | 1 | 0 | 1 |
| USP8 | 0.152 | P2_P4_Lumbar_Coeliac_Sphenopalatine_VS_P1_P3_Stellate | 0 | 1 | 0 | 1 | 1 | 1 | 0 | 1 |
| GBP7 | 0.151 | P1_VS_P2_P3_P4_Lumbar_Coeliac_Stellate_Sphenopalatine | 1 | 0 | 0 | 0 | 0 | 0 | 0 | 0 |
| TUSC1 | 0.151 | P1_P4_Lumbar_VS_P2_P3_Coeliac_Stellate_Sphenopalatine | 1 | 0 | 0 | 1 | 1 | 0 | 0 | 0 |
| PRORSD1 | 0.151 | P3_P4_Lumbar_VS_P1_P2_Coeliac_Stellate_Sphenopalatine | 0 | 0 | 1 | 1 | 1 | 0 | 0 | 0 |
| RBM43 | 0.151 | P1_P4_Lumbar_Sphenopalatine_VS_P2_P3_Coeliac_Stellate | 1 | 0 | 0 | 1 | 1 | 0 | 0 | 1 |
| BTD | 0.151 | P1_P2_P4_Lumbar_Sphenopalatine_VS_P3_Coeliac_Stellate | 1 | 1 | 0 | 1 | 1 | 0 | 0 | 1 |
| PHF1 | 0.151 | P1_P2_P3_P4_Coeliac_Sphenopalatine_VS_Lumbar_Stellate | 1 | 1 | 1 | 1 | 0 | 1 | 0 | 1 |
| SPTBN2 | 0.151 | P1_Lumbar_Coeliac_Sphenopalatine_VS_P2_P3_P4_Stellate | 1 | 0 | 0 | 0 | 1 | 1 | 0 | 1 |
| PLXNA3 | 0.151 | P1_P2_P3_Lumbar_Coeliac_Sphenopalatine_VS_P4_Stellate | 1 | 1 | 1 | 0 | 1 | 1 | 0 | 1 |
| GM5617 | 0.15 | P3_P4_VS_P1_P2_Lumbar_Coeliac_Stellate_Sphenopalatine | 0 | 0 | 1 | 1 | 0 | 0 | 0 | 0 |
| PXDN | 0.15 | P3_P4_Lumbar_VS_P1_P2_Coeliac_Stellate_Sphenopalatine | 0 | 0 | 1 | 1 | 1 | 0 | 0 | 0 |
| TOB1 | 0.15 | P2_Coeliac_VS_P1_P3_P4_Lumbar_Stellate_Sphenopalatine | 0 | 1 | 0 | 0 | 0 | 1 | 0 | 0 |
| TMEM219 | 0.15 | P1_P2_P3_P4_Sphenopalatine_VS_Lumbar_Coeliac_Stellate | 1 | 1 | 1 | 1 | 0 | 0 | 0 | 1 |
| TRIM32 | 0.15 | P4_Lumbar_Sphenopalatine_VS_P1_P2_P3_Coeliac_Stellate | 0 | 0 | 0 | 1 | 1 | 0 | 0 | 1 |
| TBC1D13 | 0.15 | P1_P2_P4_Lumbar_Sphenopalatine_VS_P3_Coeliac_Stellate | 1 | 1 | 0 | 1 | 1 | 0 | 0 | 1 |
| SSBP2 | 0.15 | P1_P2_P3_P4_Coeliac_Sphenopalatine_VS_Lumbar_Stellate | 1 | 1 | 1 | 1 | 0 | 1 | 0 | 1 |
| PRR36 | 0.15 | Lumbar_Coeliac_Sphenopalatine_VS_P1_P2_P3_P4_Stellate | 0 | 0 | 0 | 0 | 1 | 1 | 0 | 1 |
| MORC2A | 0.15 | P2_Lumbar_Coeliac_Sphenopalatine_VS_P1_P3_P4_Stellate | 0 | 1 | 0 | 0 | 1 | 1 | 0 | 1 |
| CKLF | 0.15 | P4_Lumbar_Coeliac_Sphenopalatine_VS_P1_P2_P3_Stellate | 0 | 0 | 0 | 1 | 1 | 1 | 0 | 1 |

| | | | | | | | | | | |
|---|---|---|---|---|---|---|---|---|---|---|
| CENPB | 0.15 | P2_P4_Lumbar_Coeliac_Sphenopalatine_VS_P1_P3_Stellate | 0 | 1 | 0 | 1 | 1 | 1 | 0 | 1 |
| ZFP267 | 0.15 | P2_P4_Lumbar_Coeliac_Sphenopalatine_VS_P1_P3_Stellate | 0 | 1 | 0 | 1 | 1 | 1 | 0 | 1 |
| ARHGAP5 | 0.15 | P2_P4_Lumbar_Coeliac_Sphenopalatine_VS_P1_P3_Stellate | 0 | 1 | 0 | 1 | 1 | 1 | 0 | 1 |
| VPS45 | 0.15 | P1_P3_P4_Lumbar_Coeliac_Stellate_Sphenopalatine_VS_P2 | 1 | 0 | 1 | 1 | 1 | 1 | 1 | 1 |
| C2CD2 | 0.149 | P2_Lumbar_VS_P1_P3_P4_Coeliac_Stellate_Sphenopalatine | 0 | 1 | 0 | 0 | 1 | 0 | 0 | 0 |
| ABCB6 | 0.149 | P1_P2_P3_P4_Lumbar_Coeliac_Stellate_VS_Sphenopalatine | 1 | 1 | 1 | 1 | 1 | 1 | 1 | 0 |
| LRCH1 | 0.149 | P1_P2_P4_Lumbar_Sphenopalatine_VS_P3_Coeliac_Stellate | 1 | 1 | 0 | 1 | 1 | 0 | 0 | 1 |
| MXD4 | 0.149 | P1_P2_P3_P4_Coeliac_Sphenopalatine_VS_Lumbar_Stellate | 1 | 1 | 1 | 1 | 0 | 1 | 0 | 1 |
| PPTC7 | 0.149 | P1_P2_P3_Lumbar_Coeliac_Sphenopalatine_VS_P4_Stellate | 1 | 1 | 1 | 0 | 1 | 1 | 0 | 1 |
| DENND5B | 0.149 | P1_P2_P3_Lumbar_Coeliac_Sphenopalatine_VS_P4_Stellate | 1 | 1 | 1 | 0 | 1 | 1 | 0 | 1 |
| PCP4 | 0.149 | P1_P2_P4_Lumbar_Coeliac_Stellate_Sphenopalatine_VS_P3 | 1 | 1 | 0 | 1 | 1 | 1 | 1 | 1 |
| ZSWIM7 | 0.149 | P1_P3_P4_Lumbar_Coeliac_Stellate_Sphenopalatine_VS_P2 | 1 | 0 | 1 | 1 | 1 | 1 | 1 | 1 |
| FAM149A | 0.148 | P1_VS_P2_P3_P4_Lumbar_Coeliac_Stellate_Sphenopalatine | 1 | 0 | 0 | 0 | 0 | 0 | 0 | 0 |
| SPEG | 0.148 | P3_P4_VS_P1_P2_Lumbar_Coeliac_Stellate_Sphenopalatine | 0 | 0 | 1 | 1 | 0 | 0 | 0 | 0 |
| 2810001G20 | 0.148 | P3_Lumbar_VS_P1_P2_P4_Coeliac_Stellate_Sphenopalatine | 0 | 0 | 1 | 0 | 1 | 0 | 0 | 0 |
| GM42067 | 0.148 | P3_P4_Lumbar_VS_P1_P2_Coeliac_Stellate_Sphenopalatine | 0 | 0 | 1 | 1 | 1 | 0 | 0 | 0 |
| ZSWIM4 | 0.148 | P2_Coeliac_VS_P1_P3_P4_Lumbar_Stellate_Sphenopalatine | 0 | 1 | 0 | 0 | 0 | 1 | 0 | 0 |
| HOOK1 | 0.148 | P3_P4_Lumbar_Coeliac_Sphenopalatine_VS_P1_P2_Stellate | 0 | 0 | 1 | 1 | 1 | 1 | 0 | 1 |
| ZFP652OS | 0.147 | P2_P3_VS_P1_P4_Lumbar_Coeliac_Stellate_Sphenopalatine | 0 | 1 | 1 | 0 | 0 | 0 | 0 | 0 |
| ASXL1 | 0.147 | P2_Lumbar_VS_P1_P3_P4_Coeliac_Stellate_Sphenopalatine | 0 | 1 | 0 | 0 | 1 | 0 | 0 | 0 |
| RPS6KL1 | 0.147 | P3_Lumbar_VS_P1_P2_P4_Coeliac_Stellate_Sphenopalatine | 0 | 0 | 1 | 0 | 1 | 0 | 0 | 0 |
| CPOX | 0.147 | P1_P2_P3_P4_Coeliac_Sphenopalatine_VS_Lumbar_Stellate | 1 | 1 | 1 | 1 | 0 | 1 | 0 | 1 |
| B9D1 | 0.146 | P3_P4_VS_P1_P2_Lumbar_Coeliac_Stellate_Sphenopalatine | 0 | 0 | 1 | 1 | 0 | 0 | 0 | 0 |
| CREB3L1 | 0.146 | P4_Lumbar_Coeliac_Stellate_VS_P1_P2_P3_Sphenopalatine | 0 | 0 | 0 | 1 | 1 | 1 | 1 | 0 |
| RAP1GAP2 | 0.146 | P2_P3_P4_Lumbar_Sphenopalatine_VS_P1_Coeliac_Stellate | 0 | 1 | 1 | 1 | 1 | 0 | 0 | 1 |
| RNF112 | 0.146 | P1_P2_P3_P4_Coeliac_Sphenopalatine_VS_Lumbar_Stellate | 1 | 1 | 1 | 1 | 0 | 1 | 0 | 1 |
| LMAN2L | 0.146 | P4_Lumbar_Coeliac_Sphenopalatine_VS_P1_P2_P3_Stellate | 0 | 0 | 0 | 1 | 1 | 1 | 0 | 1 |
| LRRC3B | 0.145 | Sphenopalatine_VS_P1_P2_P3_P4_Lumbar_Coeliac_Stellate | 0 | 0 | 0 | 0 | 0 | 0 | 0 | 1 |
| SNX7 | 0.145 | P1_P2_P3_P4_Sphenopalatine_VS_Lumbar_Coeliac_Stellate | 1 | 1 | 1 | 1 | 0 | 0 | 0 | 1 |
| EXTL3 | 0.145 | P1_P2_P4_Lumbar_Sphenopalatine_VS_P3_Coeliac_Stellate | 1 | 1 | 0 | 1 | 1 | 0 | 0 | 1 |

| | | | | | | | | | | |
|---|---|---|---|---|---|---|---|---|---|---|
| NLK | 0.145 | Lumbar_Coeliac_Sphenopalatine_VS_P1_P2_P3_P4_Stellate | 0 | 0 | 0 | 0 | 1 | 1 | 0 | 1 |
| PPP1R3D | 0.145 | P1_P3_Lumbar_Coeliac_Sphenopalatine_VS_P2_P4_Stellate | 1 | 0 | 1 | 0 | 1 | 1 | 0 | 1 |
| BANP | 0.145 | P4_Lumbar_Coeliac_Sphenopalatine_VS_P1_P2_P3_Stellate | 0 | 0 | 0 | 1 | 1 | 1 | 0 | 1 |
| EXOSC8 | 0.144 | P3_Lumbar_VS_P1_P2_P4_Coeliac_Stellate_Sphenopalatine | 0 | 0 | 1 | 0 | 1 | 0 | 0 | 0 |
| PHF8 | 0.144 | P2_P4_Coeliac_VS_P1_P3_Lumbar_Stellate_Sphenopalatine | 0 | 1 | 0 | 1 | 0 | 1 | 0 | 0 |
| MID2 | 0.144 | P1_P2_P3_P4_Coeliac_VS_Lumbar_Stellate_Sphenopalatine | 1 | 1 | 1 | 1 | 0 | 1 | 0 | 0 |
| EIF2S3Y | 0.144 | P1_P2_P3_P4_Coeliac_VS_Lumbar_Stellate_Sphenopalatine | 1 | 1 | 1 | 1 | 0 | 1 | 0 | 0 |
| ZFP131 | 0.144 | P1_P2_P3_P4_Lumbar_Coeliac_Stellate_VS_Sphenopalatine | 1 | 1 | 1 | 1 | 1 | 1 | 1 | 0 |
| LIN7B | 0.144 | P4_Sphenopalatine_VS_P1_P2_P3_Lumbar_Coeliac_Stellate | 0 | 0 | 0 | 1 | 0 | 0 | 0 | 1 |
| NARF | 0.144 | P1_P2_P3_P4_Coeliac_Sphenopalatine_VS_Lumbar_Stellate | 1 | 1 | 1 | 1 | 0 | 1 | 0 | 1 |
| ZNRF2 | 0.144 | P1_P2_Lumbar_Coeliac_Sphenopalatine_VS_P3_P4_Stellate | 1 | 1 | 0 | 0 | 1 | 1 | 0 | 1 |
| ID2 | 0.144 | P1_P2_P3_Lumbar_Coeliac_Stellate_Sphenopalatine_VS_P4 | 1 | 1 | 1 | 0 | 1 | 1 | 1 | 1 |
| PEX6 | 0.143 | P1_Lumbar_VS_P2_P3_P4_Coeliac_Stellate_Sphenopalatine | 1 | 0 | 0 | 0 | 1 | 0 | 0 | 0 |
| XXYLT1 | 0.143 | P2_Lumbar_VS_P1_P3_P4_Coeliac_Stellate_Sphenopalatine | 0 | 1 | 0 | 0 | 1 | 0 | 0 | 0 |
| NAA30 | 0.143 | P2_P4_Coeliac_VS_P1_P3_Lumbar_Stellate_Sphenopalatine | 0 | 1 | 0 | 1 | 0 | 1 | 0 | 0 |
| PARP4 | 0.143 | P1_P2_P3_P4_Lumbar_Coeliac_Stellate_VS_Sphenopalatine | 1 | 1 | 1 | 1 | 1 | 1 | 1 | 0 |
| DCAF13 | 0.143 | P1_P2_P3_P4_Lumbar_Coeliac_Stellate_VS_Sphenopalatine | 1 | 1 | 1 | 1 | 1 | 1 | 1 | 0 |
| ZEB1 | 0.143 | P1_P2_P3_P4_Lumbar_Coeliac_Stellate_VS_Sphenopalatine | 1 | 1 | 1 | 1 | 1 | 1 | 1 | 0 |
| RTL6 | 0.143 | Sphenopalatine_VS_P1_P2_P3_P4_Lumbar_Coeliac_Stellate | 0 | 0 | 0 | 0 | 0 | 0 | 0 | 1 |
| AP5S1 | 0.143 | P1_P2_P3_P4_Sphenopalatine_VS_Lumbar_Coeliac_Stellate | 1 | 1 | 1 | 1 | 0 | 0 | 0 | 1 |
| GPRASP2 | 0.143 | P4_Lumbar_Coeliac_Sphenopalatine_VS_P1_P2_P3_Stellate | 0 | 0 | 0 | 1 | 1 | 1 | 0 | 1 |
| TUT4 | 0.143 | P3_P4_Lumbar_Coeliac_Sphenopalatine_VS_P1_P2_Stellate | 0 | 0 | 1 | 1 | 1 | 1 | 0 | 1 |
| PDLIM2 | 0.143 | P1_P2_P3_Lumbar_Coeliac_Stellate_Sphenopalatine_VS_P4 | 1 | 1 | 1 | 0 | 1 | 1 | 1 | 1 |
| SECISBP2 | 0.142 | P1_Lumbar_VS_P2_P3_P4_Coeliac_Stellate_Sphenopalatine | 1 | 0 | 0 | 0 | 1 | 0 | 0 | 0 |
| LIG3 | 0.141 | P3_Coeliac_VS_P1_P2_P4_Lumbar_Stellate_Sphenopalatine | 0 | 0 | 1 | 0 | 0 | 1 | 0 | 0 |
| ROCK1 | 0.141 | P4_Lumbar_Coeliac_Sphenopalatine_VS_P1_P2_P3_Stellate | 0 | 0 | 0 | 1 | 1 | 1 | 0 | 1 |
| CCDC86 | 0.14 | P1_Lumbar_VS_P2_P3_P4_Coeliac_Stellate_Sphenopalatine | 1 | 0 | 0 | 0 | 1 | 0 | 0 | 0 |
| DNER | 0.14 | P1_P2_P3_Lumbar_Coeliac_Stellate_Sphenopalatine_VS_P4 | 1 | 1 | 1 | 0 | 1 | 1 | 1 | 1 |
| ICA1L | 0.139 | P3_P4_VS_P1_P2_Lumbar_Coeliac_Stellate_Sphenopalatine | 0 | 0 | 1 | 1 | 0 | 0 | 0 | 0 |
| NOP2 | 0.139 | P1_Lumbar_VS_P2_P3_P4_Coeliac_Stellate_Sphenopalatine | 1 | 0 | 0 | 0 | 1 | 0 | 0 | 0 |

| | | | | | | | | | | |
|---|---|---|---|---|---|---|---|---|---|---|
| MCAT | 0.139 | P3_P4_Lumbar_VS_P1_P2_Coeliac_Stellate_Sphenopalatine | 0 | 0 | 1 | 1 | 1 | 0 | 0 | 0 |
| CHSY1 | 0.139 | P3_P4_Coeliac_VS_P1_P2_Lumbar_Stellate_Sphenopalatine | 0 | 0 | 1 | 1 | 0 | 1 | 0 | 0 |
| AKAP1 | 0.139 | Lumbar_Coeliac_Sphenopalatine_VS_P1_P2_P3_P4_Stellate | 0 | 0 | 0 | 0 | 1 | 1 | 0 | 1 |
| EPB41L4AOS | 0.139 | P1_P3_Lumbar_Coeliac_Sphenopalatine_VS_P2_P4_Stellate | 1 | 0 | 1 | 0 | 1 | 1 | 0 | 1 |
| TRAM1 | 0.138 | P2_P4_Sphenopalatine_VS_P1_P3_Lumbar_Coeliac_Stellate | 0 | 1 | 0 | 1 | 0 | 0 | 0 | 1 |
| ZFP579 | 0.138 | Lumbar_Coeliac_Sphenopalatine_VS_P1_P2_P3_P4_Stellate | 0 | 0 | 0 | 0 | 1 | 1 | 0 | 1 |
| RELL2 | 0.137 | Sphenopalatine_VS_P1_P2_P3_P4_Lumbar_Coeliac_Stellate | 0 | 0 | 0 | 0 | 0 | 0 | 0 | 1 |
| PDCD2 | 0.137 | P3_P4_Lumbar_Coeliac_Sphenopalatine_VS_P1_P2_Stellate | 0 | 0 | 1 | 1 | 1 | 1 | 0 | 1 |
| H2-D1 | 0.136 | P1_P2_P3_P4_Lumbar_Coeliac_Stellate_VS_Sphenopalatine | 1 | 1 | 1 | 1 | 1 | 1 | 1 | 0 |
| LY6H | 0.136 | P1_P2_P3_Lumbar_Sphenopalatine_VS_P4_Coeliac_Stellate | 1 | 1 | 1 | 0 | 1 | 0 | 0 | 1 |
| CYREN | 0.136 | P3_P4_Lumbar_Coeliac_Sphenopalatine_VS_P1_P2_Stellate | 0 | 0 | 1 | 1 | 1 | 1 | 0 | 1 |
| SAG | 0.136 | P1_P3_Lumbar_Coeliac_Stellate_Sphenopalatine_VS_P2_P4 | 1 | 0 | 1 | 0 | 1 | 1 | 1 | 1 |
| 9030025P20I | 0.135 | P3_P4_VS_P1_P2_Lumbar_Coeliac_Stellate_Sphenopalatine | 0 | 0 | 1 | 1 | 0 | 0 | 0 | 0 |
| EHMT1 | 0.135 | P2_Lumbar_VS_P1_P3_P4_Coeliac_Stellate_Sphenopalatine | 0 | 1 | 0 | 0 | 1 | 0 | 0 | 0 |
| CASKIN1 | 0.135 | P2_P3_P4_Lumbar_VS_P1_Coeliac_Stellate_Sphenopalatine | 0 | 1 | 1 | 1 | 1 | 0 | 0 | 0 |
| AKIP1 | 0.135 | P1_P2_P3_P4_Sphenopalatine_VS_Lumbar_Coeliac_Stellate | 1 | 1 | 1 | 1 | 0 | 0 | 0 | 1 |
| POMGNT2 | 0.135 | P1_P2_P4_Lumbar_Sphenopalatine_VS_P3_Coeliac_Stellate | 1 | 1 | 0 | 1 | 1 | 0 | 0 | 1 |
| BLOC1S3 | 0.135 | P4_Lumbar_Coeliac_Sphenopalatine_VS_P1_P2_P3_Stellate | 0 | 0 | 0 | 1 | 1 | 1 | 0 | 1 |
| SECISBP2L | 0.134 | P1_P2_P3_P4_Lumbar_Coeliac_Stellate_VS_Sphenopalatine | 1 | 1 | 1 | 1 | 1 | 1 | 1 | 0 |
| 2900076A07I | 0.134 | P4_Lumbar_Coeliac_Sphenopalatine_VS_P1_P2_P3_Stellate | 0 | 0 | 0 | 1 | 1 | 1 | 0 | 1 |
| DIXDC1 | 0.134 | P3_P4_Lumbar_Coeliac_Sphenopalatine_VS_P1_P2_Stellate | 0 | 0 | 1 | 1 | 1 | 1 | 0 | 1 |
| PLXNB2 | 0.133 | P2_Lumbar_VS_P1_P3_P4_Coeliac_Stellate_Sphenopalatine | 0 | 1 | 0 | 0 | 1 | 0 | 0 | 0 |
| ZFP869 | 0.133 | P3_P4_Lumbar_VS_P1_P2_Coeliac_Stellate_Sphenopalatine | 0 | 0 | 1 | 1 | 1 | 0 | 0 | 0 |
| JADE1 | 0.133 | P3_Coeliac_VS_P1_P2_P4_Lumbar_Stellate_Sphenopalatine | 0 | 0 | 1 | 0 | 0 | 1 | 0 | 0 |
| ADK | 0.133 | P1_P2_P3_P4_Lumbar_Coeliac_Stellate_VS_Sphenopalatine | 1 | 1 | 1 | 1 | 1 | 1 | 1 | 0 |
| PKN1 | 0.133 | P1_P2_P4_Lumbar_Sphenopalatine_VS_P3_Coeliac_Stellate | 1 | 1 | 0 | 1 | 1 | 0 | 0 | 1 |
| GOLGA4 | 0.133 | P3_P4_Lumbar_Coeliac_Sphenopalatine_VS_P1_P2_Stellate | 0 | 0 | 1 | 1 | 1 | 1 | 0 | 1 |
| FAM118A | 0.132 | P2_P3_VS_P1_P4_Lumbar_Coeliac_Stellate_Sphenopalatine | 0 | 1 | 1 | 0 | 0 | 0 | 0 | 0 |
| ERO1L | 0.132 | P1_P2_Lumbar_Coeliac_Sphenopalatine_VS_P3_P4_Stellate | 1 | 1 | 0 | 0 | 1 | 1 | 0 | 1 |
| PRPF4B | 0.132 | P3_P4_Lumbar_Coeliac_Sphenopalatine_VS_P1_P2_Stellate | 0 | 0 | 1 | 1 | 1 | 1 | 0 | 1 |

| | | | | | | | | | | |
|---|---|---|---|---|---|---|---|---|---|---|
| LPCAT3 | 0.131 | P1_Lumbar_VS_P2_P3_P4_Coeliac_Stellate_Sphenopalatine | 1 | 0 | 0 | 0 | 1 | 0 | 0 | 0 |
| PHETA1 | 0.131 | P1_Coeliac_VS_P2_P3_P4_Lumbar_Stellate_Sphenopalatine | 1 | 0 | 0 | 0 | 0 | 1 | 0 | 0 |
| CCND3 | 0.131 | Sphenopalatine_VS_P1_P2_P3_P4_Lumbar_Coeliac_Stellate | 0 | 0 | 0 | 0 | 0 | 0 | 0 | 1 |
| GAS8 | 0.131 | P4_Lumbar_Coeliac_Sphenopalatine_VS_P1_P2_P3_Stellate | 0 | 0 | 0 | 1 | 1 | 1 | 0 | 1 |
| PNMAL2 | 0.131 | P3_P4_Lumbar_Coeliac_Sphenopalatine_VS_P1_P2_Stellate | 0 | 0 | 1 | 1 | 1 | 1 | 0 | 1 |
| TIAM1 | 0.13 | P2_Lumbar_VS_P1_P3_P4_Coeliac_Stellate_Sphenopalatine | 0 | 1 | 0 | 0 | 1 | 0 | 0 | 0 |
| UXT | 0.13 | P1_P3_P4_Lumbar_VS_P2_Coeliac_Stellate_Sphenopalatine | 1 | 0 | 1 | 1 | 1 | 0 | 0 | 0 |
| KCNB2 | 0.13 | P2_P3_P4_Lumbar_VS_P1_Coeliac_Stellate_Sphenopalatine | 0 | 1 | 1 | 1 | 1 | 0 | 0 | 0 |
| IGFBPL1 | 0.13 | Sphenopalatine_VS_P1_P2_P3_P4_Lumbar_Coeliac_Stellate | 0 | 0 | 0 | 0 | 0 | 0 | 0 | 1 |
| DCUN1D4 | 0.13 | Sphenopalatine_VS_P1_P2_P3_P4_Lumbar_Coeliac_Stellate | 0 | 0 | 0 | 0 | 0 | 0 | 0 | 1 |
| UGGT1 | 0.13 | P3_P4_Lumbar_Coeliac_Sphenopalatine_VS_P1_P2_Stellate | 0 | 0 | 1 | 1 | 1 | 1 | 0 | 1 |
| NEU1 | 0.13 | P3_P4_Lumbar_Coeliac_Sphenopalatine_VS_P1_P2_Stellate | 0 | 0 | 1 | 1 | 1 | 1 | 0 | 1 |
| DHRS4 | 0.129 | P1_P2_P3_P4_Lumbar_Coeliac_Stellate_VS_Sphenopalatine | 1 | 1 | 1 | 1 | 1 | 1 | 1 | 0 |
| NCAM2 | 0.129 | P1_Lumbar_Coeliac_Sphenopalatine_VS_P2_P3_P4_Stellate | 1 | 0 | 0 | 0 | 1 | 1 | 0 | 1 |
| SHANK2 | 0.129 | P1_P2_Lumbar_Coeliac_Sphenopalatine_VS_P3_P4_Stellate | 1 | 1 | 0 | 0 | 1 | 1 | 0 | 1 |
| DDX27 | 0.129 | P3_P4_Lumbar_Coeliac_Sphenopalatine_VS_P1_P2_Stellate | 0 | 0 | 1 | 1 | 1 | 1 | 0 | 1 |
| ATXN7L1 | 0.128 | P2_P3_VS_P1_P4_Lumbar_Coeliac_Stellate_Sphenopalatine | 0 | 1 | 1 | 0 | 0 | 0 | 0 | 0 |
| ASIC4 | 0.128 | P2_Coeliac_VS_P1_P3_P4_Lumbar_Stellate_Sphenopalatine | 0 | 1 | 0 | 0 | 0 | 1 | 0 | 0 |
| FAM169A | 0.128 | P1_P2_P4_Lumbar_Sphenopalatine_VS_P3_Coeliac_Stellate | 1 | 1 | 0 | 1 | 1 | 0 | 0 | 1 |
| FARSA | 0.128 | Lumbar_Coeliac_Sphenopalatine_VS_P1_P2_P3_P4_Stellate | 0 | 0 | 0 | 0 | 1 | 1 | 0 | 1 |
| TM6SF1 | 0.128 | P1_P2_Lumbar_Coeliac_Sphenopalatine_VS_P3_P4_Stellate | 1 | 1 | 0 | 0 | 1 | 1 | 0 | 1 |
| DXO | 0.127 | P1_Lumbar_VS_P2_P3_P4_Coeliac_Stellate_Sphenopalatine | 1 | 0 | 0 | 0 | 1 | 0 | 0 | 0 |
| ACSL1 | 0.127 | P2_Lumbar_VS_P1_P3_P4_Coeliac_Stellate_Sphenopalatine | 0 | 1 | 0 | 0 | 1 | 0 | 0 | 0 |
| KDM6B | 0.127 | P2_Lumbar_VS_P1_P3_P4_Coeliac_Stellate_Sphenopalatine | 0 | 1 | 0 | 0 | 1 | 0 | 0 | 0 |
| GORAB | 0.127 | P3_P4_Lumbar_VS_P1_P2_Coeliac_Stellate_Sphenopalatine | 0 | 0 | 1 | 1 | 1 | 0 | 0 | 0 |
| TMEM87B | 0.127 | P1_P2_P4_Lumbar_Sphenopalatine_VS_P3_Coeliac_Stellate | 1 | 1 | 0 | 1 | 1 | 0 | 0 | 1 |
| NIT1 | 0.127 | P3_P4_Lumbar_Coeliac_Sphenopalatine_VS_P1_P2_Stellate | 0 | 0 | 1 | 1 | 1 | 1 | 0 | 1 |
| DESI2 | 0.127 | P3_P4_Lumbar_Coeliac_Sphenopalatine_VS_P1_P2_Stellate | 0 | 0 | 1 | 1 | 1 | 1 | 0 | 1 |
| SOX4 | 0.127 | P1_P2_P3_Lumbar_Coeliac_Stellate_Sphenopalatine_VS_P4 | 1 | 1 | 1 | 0 | 1 | 1 | 1 | 1 |
| PAK2 | 0.126 | Lumbar_Coeliac_Sphenopalatine_VS_P1_P2_P3_P4_Stellate | 0 | 0 | 0 | 0 | 1 | 1 | 0 | 1 |

| | | | | | | | | | | |
|---|---|---|---|---|---|---|---|---|---|---|
| SELENOI | 0.126 | P1_P2_Lumbar_Coeliac_Sphenopalatine_VS_P3_P4_Stellate | 1 | 1 | 0 | 0 | 1 | 1 | 0 | 1 |
| LCA5L | 0.126 | P4_Lumbar_Coeliac_Sphenopalatine_VS_P1_P2_P3_Stellate | 0 | 0 | 0 | 1 | 1 | 1 | 0 | 1 |
| CHRNA5 | 0.126 | P3_P4_Lumbar_Coeliac_Sphenopalatine_VS_P1_P2_Stellate | 0 | 0 | 1 | 1 | 1 | 1 | 0 | 1 |
| GM32031 | 0.125 | P3_P4_VS_P1_P2_Lumbar_Coeliac_Stellate_Sphenopalatine | 0 | 0 | 1 | 1 | 0 | 0 | 0 | 0 |
| EIF2B3 | 0.125 | P1_Lumbar_VS_P2_P3_P4_Coeliac_Stellate_Sphenopalatine | 1 | 0 | 0 | 0 | 1 | 0 | 0 | 0 |
| VARS | 0.125 | P1_Lumbar_VS_P2_P3_P4_Coeliac_Stellate_Sphenopalatine | 1 | 0 | 0 | 0 | 1 | 0 | 0 | 0 |
| THOP1 | 0.125 | P3_P4_Lumbar_VS_P1_P2_Coeliac_Stellate_Sphenopalatine | 0 | 0 | 1 | 1 | 1 | 0 | 0 | 0 |
| NUDT17 | 0.125 | Lumbar_Coeliac_Stellate_VS_P1_P2_P3_P4_Sphenopalatine | 0 | 0 | 0 | 0 | 1 | 1 | 1 | 0 |
| CADM4 | 0.125 | Lumbar_Coeliac_Sphenopalatine_VS_P1_P2_P3_P4_Stellate | 0 | 0 | 0 | 0 | 1 | 1 | 0 | 1 |
| DCPS | 0.125 | Lumbar_Coeliac_Sphenopalatine_VS_P1_P2_P3_P4_Stellate | 0 | 0 | 0 | 0 | 1 | 1 | 0 | 1 |
| SFT2D1 | 0.125 | Lumbar_Coeliac_Sphenopalatine_VS_P1_P2_P3_P4_Stellate | 0 | 0 | 0 | 0 | 1 | 1 | 0 | 1 |
| SDE2 | 0.125 | P4_Lumbar_Coeliac_Sphenopalatine_VS_P1_P2_P3_Stellate | 0 | 0 | 0 | 1 | 1 | 1 | 0 | 1 |
| WLS | 0.124 | P1_P3_P4_VS_P2_Lumbar_Coeliac_Stellate_Sphenopalatine | 1 | 0 | 1 | 1 | 0 | 0 | 0 | 0 |
| STXBP3 | 0.124 | P2_Lumbar_VS_P1_P3_P4_Coeliac_Stellate_Sphenopalatine | 0 | 1 | 0 | 0 | 1 | 0 | 0 | 0 |
| CAPRIN2 | 0.124 | P1_P4_Coeliac_VS_P2_P3_Lumbar_Stellate_Sphenopalatine | 1 | 0 | 0 | 1 | 0 | 1 | 0 | 0 |
| ASB3 | 0.124 | P1_P4_Coeliac_VS_P2_P3_Lumbar_Stellate_Sphenopalatine | 1 | 0 | 0 | 1 | 0 | 1 | 0 | 0 |
| SIRPA | 0.123 | P3_Coeliac_VS_P1_P2_P4_Lumbar_Stellate_Sphenopalatine | 0 | 0 | 1 | 0 | 0 | 1 | 0 | 0 |
| ACAA2 | 0.123 | P1_P2_P3_P4_Lumbar_Coeliac_Stellate_VS_Sphenopalatine | 1 | 1 | 1 | 1 | 1 | 1 | 1 | 0 |
| PPP3CC | 0.123 | Lumbar_Coeliac_Sphenopalatine_VS_P1_P2_P3_P4_Stellate | 0 | 0 | 0 | 0 | 1 | 1 | 0 | 1 |
| SUGP2 | 0.123 | P4_Lumbar_Coeliac_Sphenopalatine_VS_P1_P2_P3_Stellate | 0 | 0 | 0 | 1 | 1 | 1 | 0 | 1 |
| CELF5 | 0.123 | P3_P4_Lumbar_Coeliac_Sphenopalatine_VS_P1_P2_Stellate | 0 | 0 | 1 | 1 | 1 | 1 | 0 | 1 |
| CEP83 | 0.123 | P3_P4_Lumbar_Coeliac_Sphenopalatine_VS_P1_P2_Stellate | 0 | 0 | 1 | 1 | 1 | 1 | 0 | 1 |
| GNL2 | 0.122 | P1_Lumbar_VS_P2_P3_P4_Coeliac_Stellate_Sphenopalatine | 1 | 0 | 0 | 0 | 1 | 0 | 0 | 0 |
| EZR | 0.122 | P1_P3_P4_Lumbar_VS_P2_Coeliac_Stellate_Sphenopalatine | 1 | 0 | 1 | 1 | 1 | 0 | 0 | 0 |
| CREB3L2 | 0.122 | P3_Coeliac_VS_P1_P2_P4_Lumbar_Stellate_Sphenopalatine | 0 | 0 | 1 | 0 | 0 | 1 | 0 | 0 |
| CREM | 0.122 | Sphenopalatine_VS_P1_P2_P3_P4_Lumbar_Coeliac_Stellate | 0 | 0 | 0 | 0 | 0 | 0 | 0 | 1 |
| ENO2 | 0.122 | P1_P2_P3_P4_Sphenopalatine_VS_Lumbar_Coeliac_Stellate | 1 | 1 | 1 | 1 | 0 | 0 | 0 | 1 |
| RYR2 | 0.122 | P1_P2_Lumbar_Coeliac_Sphenopalatine_VS_P3_P4_Stellate | 1 | 1 | 0 | 0 | 1 | 1 | 0 | 1 |
| ZFHX4 | 0.121 | P2_P3_VS_P1_P4_Lumbar_Coeliac_Stellate_Sphenopalatine | 0 | 1 | 1 | 0 | 0 | 0 | 0 | 0 |
| RNPEPL1 | 0.121 | P1_Lumbar_VS_P2_P3_P4_Coeliac_Stellate_Sphenopalatine | 1 | 0 | 0 | 0 | 1 | 0 | 0 | 0 |

| | | | | | | | | | | |
|---|---|---|---|---|---|---|---|---|---|---|
| ATIC | 0.121 | P4_Lumbar_Coeliac_Sphenopalatine_VS_P1_P2_P3_Stellate | 0 | 0 | 0 | 1 | 1 | 1 | 0 | 1 |
| NGDN | 0.121 | P4_Lumbar_Coeliac_Sphenopalatine_VS_P1_P2_P3_Stellate | 0 | 0 | 0 | 1 | 1 | 1 | 0 | 1 |
| TCEAL3 | 0.121 | P3_P4_Lumbar_Coeliac_Sphenopalatine_VS_P1_P2_Stellate | 0 | 0 | 1 | 1 | 1 | 1 | 0 | 1 |
| METTL5 | 0.121 | P1_P3_P4_Lumbar_Coeliac_Stellate_Sphenopalatine_VS_P2 | 1 | 0 | 1 | 1 | 1 | 1 | 1 | 1 |
| ACAD8 | 0.12 | P1_P2_P3_P4_Lumbar_Coeliac_Stellate_VS_Sphenopalatine | 1 | 1 | 1 | 1 | 1 | 1 | 1 | 0 |
| TMEM151A | 0.12 | P1_P2_P3_P4_Sphenopalatine_VS_Lumbar_Coeliac_Stellate | 1 | 1 | 1 | 1 | 0 | 0 | 0 | 1 |
| TRAF7 | 0.12 | P1_P3_Lumbar_Coeliac_Sphenopalatine_VS_P2_P4_Stellate | 1 | 0 | 1 | 0 | 1 | 1 | 0 | 1 |
| METTL9 | 0.12 | P1_P2_P3_Lumbar_Coeliac_Stellate_Sphenopalatine_VS_P4 | 1 | 1 | 1 | 0 | 1 | 1 | 1 | 1 |
| DDX18 | 0.12 | P1_P3_P4_Lumbar_Coeliac_Stellate_Sphenopalatine_VS_P2 | 1 | 0 | 1 | 1 | 1 | 1 | 1 | 1 |
| PLCXD3 | 0.12 | P1_P3_P4_Lumbar_Coeliac_Stellate_Sphenopalatine_VS_P2 | 1 | 0 | 1 | 1 | 1 | 1 | 1 | 1 |
| A930006K02 | 0.119 | P3_P4_VS_P1_P2_Lumbar_Coeliac_Stellate_Sphenopalatine | 0 | 0 | 1 | 1 | 0 | 0 | 0 | 0 |
| PNP | 0.119 | P1_P3_P4_Lumbar_Coeliac_Stellate_VS_P2_Sphenopalatine | 1 | 0 | 1 | 1 | 1 | 1 | 1 | 0 |
| TRIM24 | 0.119 | P1_P2_P3_P4_Lumbar_Coeliac_Stellate_VS_Sphenopalatine | 1 | 1 | 1 | 1 | 1 | 1 | 1 | 0 |
| NPLOC4 | 0.119 | P1_P2_P3_P4_Sphenopalatine_VS_Lumbar_Coeliac_Stellate | 1 | 1 | 1 | 1 | 0 | 0 | 0 | 1 |
| MTERF4 | 0.119 | P1_P3_Lumbar_Coeliac_Sphenopalatine_VS_P2_P4_Stellate | 1 | 0 | 1 | 0 | 1 | 1 | 0 | 1 |
| CCDC22 | 0.118 | P1_Lumbar_VS_P2_P3_P4_Coeliac_Stellate_Sphenopalatine | 1 | 0 | 0 | 0 | 1 | 0 | 0 | 0 |
| DCAF6 | 0.118 | P1_P2_P3_P4_Lumbar_Coeliac_Stellate_VS_Sphenopalatine | 1 | 1 | 1 | 1 | 1 | 1 | 1 | 0 |
| ANKRD34A | 0.118 | P1_Sphenopalatine_VS_P2_P3_P4_Lumbar_Coeliac_Stellate | 1 | 0 | 0 | 0 | 0 | 0 | 0 | 1 |
| TRP53RKA | 0.118 | Lumbar_Coeliac_Sphenopalatine_VS_P1_P2_P3_P4_Stellate | 0 | 0 | 0 | 0 | 1 | 1 | 0 | 1 |
| CLIP1 | 0.118 | P4_Lumbar_Coeliac_Sphenopalatine_VS_P1_P2_P3_Stellate | 0 | 0 | 0 | 1 | 1 | 1 | 0 | 1 |
| STAT2 | 0.118 | P4_Lumbar_Coeliac_Sphenopalatine_VS_P1_P2_P3_Stellate | 0 | 0 | 0 | 1 | 1 | 1 | 0 | 1 |
| ITCH | 0.118 | P3_P4_Lumbar_Coeliac_Sphenopalatine_VS_P1_P2_Stellate | 0 | 0 | 1 | 1 | 1 | 1 | 0 | 1 |
| USP33 | 0.117 | P2_Lumbar_VS_P1_P3_P4_Coeliac_Stellate_Sphenopalatine | 0 | 1 | 0 | 0 | 1 | 0 | 0 | 0 |
| ELOVL1 | 0.117 | P1_P2_P3_Lumbar_VS_P4_Coeliac_Stellate_Sphenopalatine | 1 | 1 | 1 | 0 | 1 | 0 | 0 | 0 |
| ACTR6 | 0.117 | P3_P4_Lumbar_VS_P1_P2_Coeliac_Stellate_Sphenopalatine | 0 | 0 | 1 | 1 | 1 | 0 | 0 | 0 |
| TRAPPC12 | 0.117 | P3_P4_Lumbar_VS_P1_P2_Coeliac_Stellate_Sphenopalatine | 0 | 0 | 1 | 1 | 1 | 0 | 0 | 0 |
| TSC22D4 | 0.117 | P1_P2_P3_P4_Lumbar_Coeliac_Stellate_VS_Sphenopalatine | 1 | 1 | 1 | 1 | 1 | 1 | 1 | 0 |
| REC114 | 0.117 | Lumbar_Sphenopalatine_VS_P1_P2_P3_P4_Coeliac_Stellate | 0 | 0 | 0 | 0 | 1 | 0 | 0 | 1 |
| CALCOCO1 | 0.117 | P2_Lumbar_Coeliac_Sphenopalatine_VS_P1_P3_P4_Stellate | 0 | 1 | 0 | 0 | 1 | 1 | 0 | 1 |
| VGF | 0.116 | P1_P3_VS_P2_P4_Lumbar_Coeliac_Stellate_Sphenopalatine | 1 | 0 | 1 | 0 | 0 | 0 | 0 | 0 |

| | | | | | | | | | | |
|---|---|---|---|---|---|---|---|---|---|---|
| MAN2C1 | 0.116 | P1_P3_P4_Lumbar_VS_P2_Coeliac_Stellate_Sphenopalatine | 1 | 0 | 1 | 1 | 1 | 0 | 0 | 0 |
| JARID2 | 0.116 | P2_P3_P4_Lumbar_Coeliac_Stellate_VS_P1_Sphenopalatine | 0 | 1 | 1 | 1 | 1 | 1 | 1 | 0 |
| NINJ1 | 0.116 | P1_P2_P3_P4_Sphenopalatine_VS_Lumbar_Coeliac_Stellate | 1 | 1 | 1 | 1 | 0 | 0 | 0 | 1 |
| PRRG3 | 0.116 | P2_P3_Lumbar_Coeliac_Sphenopalatine_VS_P1_P4_Stellate | 0 | 1 | 1 | 0 | 1 | 1 | 0 | 1 |
| PITPNC1 | 0.116 | Lumbar_Coeliac_Stellate_Sphenopalatine_VS_P1_P2_P3_P4 | 0 | 0 | 0 | 0 | 1 | 1 | 1 | 1 |
| SAT2 | 0.115 | P3_P4_VS_P1_P2_Lumbar_Coeliac_Stellate_Sphenopalatine | 0 | 0 | 1 | 1 | 0 | 0 | 0 | 0 |
| SYNGR2 | 0.115 | P3_P4_VS_P1_P2_Lumbar_Coeliac_Stellate_Sphenopalatine | 0 | 0 | 1 | 1 | 0 | 0 | 0 | 0 |
| JMJD6 | 0.115 | P1_Lumbar_VS_P2_P3_P4_Coeliac_Stellate_Sphenopalatine | 1 | 0 | 0 | 0 | 1 | 0 | 0 | 0 |
| SETD7 | 0.115 | Lumbar_Coeliac_Sphenopalatine_VS_P1_P2_P3_P4_Stellate | 0 | 0 | 0 | 0 | 1 | 1 | 0 | 1 |
| N4BP2L2 | 0.115 | P3_P4_Lumbar_Coeliac_Sphenopalatine_VS_P1_P2_Stellate | 0 | 0 | 1 | 1 | 1 | 1 | 0 | 1 |
| FASTK | 0.115 | P1_P2_P3_P4_Lumbar_Stellate_Sphenopalatine_VS_Coeliac | 1 | 1 | 1 | 1 | 1 | 0 | 1 | 1 |
| FRMD5 | 0.115 | P1_P2_P4_Lumbar_Coeliac_Stellate_Sphenopalatine_VS_P3 | 1 | 1 | 0 | 1 | 1 | 1 | 1 | 1 |
| 9230114K14I | 0.114 | P3_P4_VS_P1_P2_Lumbar_Coeliac_Stellate_Sphenopalatine | 0 | 0 | 1 | 1 | 0 | 0 | 0 | 0 |
| DMAP1 | 0.114 | P1_Lumbar_VS_P2_P3_P4_Coeliac_Stellate_Sphenopalatine | 1 | 0 | 0 | 0 | 1 | 0 | 0 | 0 |
| TTC39C | 0.114 | P1_Lumbar_VS_P2_P3_P4_Coeliac_Stellate_Sphenopalatine | 1 | 0 | 0 | 0 | 1 | 0 | 0 | 0 |
| TOMM40L | 0.114 | P1_P2_P3_Sphenopalatine_VS_P4_Lumbar_Coeliac_Stellate | 1 | 1 | 1 | 0 | 0 | 0 | 0 | 1 |
| HACD2 | 0.113 | P3_P4_Lumbar_VS_P1_P2_Coeliac_Stellate_Sphenopalatine | 0 | 0 | 1 | 1 | 1 | 0 | 0 | 0 |
| PPP1R21 | 0.113 | P3_P4_Lumbar_VS_P1_P2_Coeliac_Stellate_Sphenopalatine | 0 | 0 | 1 | 1 | 1 | 0 | 0 | 0 |
| AGO4 | 0.113 | P2_Coeliac_VS_P1_P3_P4_Lumbar_Stellate_Sphenopalatine | 0 | 1 | 0 | 0 | 0 | 1 | 0 | 0 |
| PDE4DIP | 0.113 | P4_Lumbar_Sphenopalatine_VS_P1_P2_P3_Coeliac_Stellate | 0 | 0 | 0 | 1 | 1 | 0 | 0 | 1 |
| DAGLA | 0.113 | P4_Lumbar_Sphenopalatine_VS_P1_P2_P3_Coeliac_Stellate | 0 | 0 | 0 | 1 | 1 | 0 | 0 | 1 |
| NME5 | 0.112 | P3_P4_VS_P1_P2_Lumbar_Coeliac_Stellate_Sphenopalatine | 0 | 0 | 1 | 1 | 0 | 0 | 0 | 0 |
| MAML3 | 0.112 | Lumbar_Coeliac_Sphenopalatine_VS_P1_P2_P3_P4_Stellate | 0 | 0 | 0 | 0 | 1 | 1 | 0 | 1 |
| P4HA2 | 0.112 | P1_Lumbar_Coeliac_Sphenopalatine_VS_P2_P3_P4_Stellate | 1 | 0 | 0 | 0 | 1 | 1 | 0 | 1 |
| CLEC16A | 0.112 | P1_P3_Lumbar_Coeliac_Sphenopalatine_VS_P2_P4_Stellate | 1 | 0 | 1 | 0 | 1 | 1 | 0 | 1 |
| SP3OS | 0.112 | P3_P4_Lumbar_Coeliac_Sphenopalatine_VS_P1_P2_Stellate | 0 | 0 | 1 | 1 | 1 | 1 | 0 | 1 |
| H2AFY2 | 0.112 | P1_P3_P4_Lumbar_Coeliac_Stellate_Sphenopalatine_VS_P2 | 1 | 0 | 1 | 1 | 1 | 1 | 1 | 1 |
| LYSMD4 | 0.111 | P3_P4_VS_P1_P2_Lumbar_Coeliac_Stellate_Sphenopalatine | 0 | 0 | 1 | 1 | 0 | 0 | 0 | 0 |
| ATAD3A | 0.111 | P1_Lumbar_VS_P2_P3_P4_Coeliac_Stellate_Sphenopalatine | 1 | 0 | 0 | 0 | 1 | 0 | 0 | 0 |
| LRIF1 | 0.111 | P3_P4_Lumbar_VS_P1_P2_Coeliac_Stellate_Sphenopalatine | 0 | 0 | 1 | 1 | 1 | 0 | 0 | 0 |

| | | | | | | | | | | |
|---|---|---|---|---|---|---|---|---|---|---|
| CCDC106 | 0.111 | P3_P4_Lumbar_VS_P1_P2_Coeliac_Stellate_Sphenopalatine | 0 | 0 | 1 | 1 | 1 | 0 | 0 | 0 |
| MTREX | 0.111 | P3_P4_Lumbar_VS_P1_P2_Coeliac_Stellate_Sphenopalatine | 0 | 0 | 1 | 1 | 1 | 0 | 0 | 0 |
| ME2 | 0.111 | P2_Coeliac_VS_P1_P3_P4_Lumbar_Stellate_Sphenopalatine | 0 | 1 | 0 | 0 | 0 | 1 | 0 | 0 |
| MGLL | 0.111 | P1_P2_P3_P4_Lumbar_Coeliac_Stellate_VS_Sphenopalatine | 1 | 1 | 1 | 1 | 1 | 1 | 1 | 0 |
| NME6 | 0.111 | P1_P2_P3_P4_Lumbar_Coeliac_Stellate_VS_Sphenopalatine | 1 | 1 | 1 | 1 | 1 | 1 | 1 | 0 |
| PDE10A | 0.111 | P2_Sphenopalatine_VS_P1_P3_P4_Lumbar_Coeliac_Stellate | 0 | 1 | 0 | 0 | 0 | 0 | 0 | 1 |
| POM121 | 0.111 | P3_P4_Lumbar_Coeliac_Sphenopalatine_VS_P1_P2_Stellate | 0 | 0 | 1 | 1 | 1 | 1 | 0 | 1 |
| GPS2 | 0.111 | P3_P4_Lumbar_Coeliac_Sphenopalatine_VS_P1_P2_Stellate | 0 | 0 | 1 | 1 | 1 | 1 | 0 | 1 |
| SNAPIN | 0.111 | P1_P2_P3_P4_Lumbar_Stellate_Sphenopalatine_VS_Coeliac | 1 | 1 | 1 | 1 | 1 | 0 | 1 | 1 |
| GM5914 | 0.11 | P3_P4_VS_P1_P2_Lumbar_Coeliac_Stellate_Sphenopalatine | 0 | 0 | 1 | 1 | 0 | 0 | 0 | 0 |
| ZSWIM5 | 0.11 | P1_Lumbar_VS_P2_P3_P4_Coeliac_Stellate_Sphenopalatine | 1 | 0 | 0 | 0 | 1 | 0 | 0 | 0 |
| HOMER1 | 0.11 | P2_Lumbar_VS_P1_P3_P4_Coeliac_Stellate_Sphenopalatine | 0 | 1 | 0 | 0 | 1 | 0 | 0 | 0 |
| 2610044O15 | 0.11 | P1_P4_Coeliac_VS_P2_P3_Lumbar_Stellate_Sphenopalatine | 1 | 0 | 0 | 1 | 0 | 1 | 0 | 0 |
| KCNK10 | 0.11 | P1_P2_P3_P4_Lumbar_Coeliac_Stellate_VS_Sphenopalatine | 1 | 1 | 1 | 1 | 1 | 1 | 1 | 0 |
| FBXO44 | 0.11 | P1_P2_Sphenopalatine_VS_P3_P4_Lumbar_Coeliac_Stellate | 1 | 1 | 0 | 0 | 0 | 0 | 0 | 1 |
| ZZZ3 | 0.11 | P3_P4_Lumbar_Coeliac_Sphenopalatine_VS_P1_P2_Stellate | 0 | 0 | 1 | 1 | 1 | 1 | 0 | 1 |
| IDH2 | 0.109 | P1_Lumbar_VS_P2_P3_P4_Coeliac_Stellate_Sphenopalatine | 1 | 0 | 0 | 0 | 1 | 0 | 0 | 0 |
| MTR | 0.109 | P2_Coeliac_VS_P1_P3_P4_Lumbar_Stellate_Sphenopalatine | 0 | 1 | 0 | 0 | 0 | 1 | 0 | 0 |
| STX16 | 0.109 | P1_P2_P3_P4_Lumbar_Coeliac_Stellate_VS_Sphenopalatine | 1 | 1 | 1 | 1 | 1 | 1 | 1 | 0 |
| SH3RF3 | 0.109 | Coeliac_Sphenopalatine_VS_P1_P2_P3_P4_Lumbar_Stellate | 0 | 0 | 0 | 0 | 0 | 1 | 0 | 1 |
| IWS1 | 0.109 | P1_P2_Lumbar_Coeliac_Sphenopalatine_VS_P3_P4_Stellate | 1 | 1 | 0 | 0 | 1 | 1 | 0 | 1 |
| GM4924 | 0.109 | P1_P3_Lumbar_Coeliac_Sphenopalatine_VS_P2_P4_Stellate | 1 | 0 | 1 | 0 | 1 | 1 | 0 | 1 |
| CLUAP1 | 0.109 | P1_P3_Lumbar_Coeliac_Sphenopalatine_VS_P2_P4_Stellate | 1 | 0 | 1 | 0 | 1 | 1 | 0 | 1 |
| ZFP280D | 0.109 | P3_P4_Lumbar_Coeliac_Sphenopalatine_VS_P1_P2_Stellate | 0 | 0 | 1 | 1 | 1 | 1 | 0 | 1 |
| HDDC2 | 0.109 | P1_P3_P4_Lumbar_Coeliac_Stellate_Sphenopalatine_VS_P2 | 1 | 0 | 1 | 1 | 1 | 1 | 1 | 1 |
| GFM2 | 0.108 | P3_P4_Lumbar_VS_P1_P2_Coeliac_Stellate_Sphenopalatine | 0 | 0 | 1 | 1 | 1 | 0 | 0 | 0 |
| SYAP1 | 0.108 | Lumbar_Coeliac_Sphenopalatine_VS_P1_P2_P3_P4_Stellate | 0 | 0 | 0 | 0 | 1 | 1 | 0 | 1 |
| RCC2 | 0.108 | P1_P2_Lumbar_Coeliac_Sphenopalatine_VS_P3_P4_Stellate | 1 | 1 | 0 | 0 | 1 | 1 | 0 | 1 |
| PRMT7 | 0.108 | P1_P3_P4_Lumbar_Coeliac_Stellate_Sphenopalatine_VS_P2 | 1 | 0 | 1 | 1 | 1 | 1 | 1 | 1 |
| WDR74 | 0.108 | P1_P3_P4_Lumbar_Coeliac_Stellate_Sphenopalatine_VS_P2 | 1 | 0 | 1 | 1 | 1 | 1 | 1 | 1 |

| | | | | | | | | | | |
|---|---|---|---|---|---|---|---|---|---|---|
| RMND1 | 0.107 | P1_Lumbar_VS_P2_P3_P4_Coeliac_Stellate_Sphenopalatine | 1 | 0 | 0 | 0 | 1 | 0 | 0 | 0 |
| CREG2 | 0.107 | P2_P3_P4_Lumbar_VS_P1_Coeliac_Stellate_Sphenopalatine | 0 | 1 | 1 | 1 | 1 | 0 | 0 | 0 |
| BLVRA | 0.107 | P1_P3_P4_Lumbar_Coeliac_Stellate_VS_P2_Sphenopalatine | 1 | 0 | 1 | 1 | 1 | 1 | 1 | 0 |
| CCDC61 | 0.107 | P1_P2_P3_P4_Lumbar_Coeliac_Stellate_VS_Sphenopalatine | 1 | 1 | 1 | 1 | 1 | 1 | 1 | 0 |
| TRIM25 | 0.107 | P1_P4_Lumbar_Sphenopalatine_VS_P2_P3_Coeliac_Stellate | 1 | 0 | 0 | 1 | 1 | 0 | 0 | 1 |
| HNRNPH3 | 0.107 | P1_P3_Lumbar_Coeliac_Sphenopalatine_VS_P2_P4_Stellate | 1 | 0 | 1 | 0 | 1 | 1 | 0 | 1 |
| LSM12 | 0.107 | P3_P4_Lumbar_Coeliac_Sphenopalatine_VS_P1_P2_Stellate | 0 | 0 | 1 | 1 | 1 | 1 | 0 | 1 |
| HES6 | 0.107 | P1_P3_P4_Lumbar_Coeliac_Stellate_Sphenopalatine_VS_P2 | 1 | 0 | 1 | 1 | 1 | 1 | 1 | 1 |
| SEC24D | 0.106 | P2_Lumbar_VS_P1_P3_P4_Coeliac_Stellate_Sphenopalatine | 0 | 1 | 0 | 0 | 1 | 0 | 0 | 0 |
| ARID5B | 0.106 | P1_P2_P3_P4_Lumbar_Coeliac_Stellate_VS_Sphenopalatine | 1 | 1 | 1 | 1 | 1 | 1 | 1 | 0 |
| PRPF8 | 0.106 | P3_P4_Lumbar_Coeliac_Sphenopalatine_VS_P1_P2_Stellate | 0 | 0 | 1 | 1 | 1 | 1 | 0 | 1 |
| PCGF6 | 0.106 | P3_P4_Lumbar_Coeliac_Sphenopalatine_VS_P1_P2_Stellate | 0 | 0 | 1 | 1 | 1 | 1 | 0 | 1 |
| GSKIP | 0.105 | P3_P4_VS_P1_P2_Lumbar_Coeliac_Stellate_Sphenopalatine | 0 | 0 | 1 | 1 | 0 | 0 | 0 | 0 |
| GNAI1 | 0.105 | P3_P4_Lumbar_VS_P1_P2_Coeliac_Stellate_Sphenopalatine | 0 | 0 | 1 | 1 | 1 | 0 | 0 | 0 |
| RHBDD3 | 0.105 | P3_P4_Lumbar_VS_P1_P2_Coeliac_Stellate_Sphenopalatine | 0 | 0 | 1 | 1 | 1 | 0 | 0 | 0 |
| 2700046G09 | 0.105 | P1_P3_P4_Lumbar_VS_P2_Coeliac_Stellate_Sphenopalatine | 1 | 0 | 1 | 1 | 1 | 0 | 0 | 0 |
| AFTPH | 0.105 | P1_P2_Lumbar_Coeliac_Sphenopalatine_VS_P3_P4_Stellate | 1 | 1 | 0 | 0 | 1 | 1 | 0 | 1 |
| B830012L14I | 0.105 | P1_P2_Lumbar_Coeliac_Sphenopalatine_VS_P3_P4_Stellate | 1 | 1 | 0 | 0 | 1 | 1 | 0 | 1 |
| KPTN | 0.105 | P3_P4_Lumbar_Coeliac_Sphenopalatine_VS_P1_P2_Stellate | 0 | 0 | 1 | 1 | 1 | 1 | 0 | 1 |
| FRYL | 0.104 | P2_P3_P4_Lumbar_VS_P1_Coeliac_Stellate_Sphenopalatine | 0 | 1 | 1 | 1 | 1 | 0 | 0 | 0 |
| TEX30 | 0.104 | P1_Coeliac_VS_P2_P3_P4_Lumbar_Stellate_Sphenopalatine | 1 | 0 | 0 | 0 | 0 | 1 | 0 | 0 |
| KLHL22 | 0.104 | P1_P3_P4_Lumbar_Sphenopalatine_VS_P2_Coeliac_Stellate | 1 | 0 | 1 | 1 | 1 | 0 | 0 | 1 |
| AVPI1 | 0.104 | P1_P3_P4_Lumbar_Sphenopalatine_VS_P2_Coeliac_Stellate | 1 | 0 | 1 | 1 | 1 | 0 | 0 | 1 |
| TMF1 | 0.104 | Lumbar_Coeliac_Sphenopalatine_VS_P1_P2_P3_P4_Stellate | 0 | 0 | 0 | 0 | 1 | 1 | 0 | 1 |
| ASRGL1 | 0.103 | P1_P3_P4_Lumbar_VS_P2_Coeliac_Stellate_Sphenopalatine | 1 | 0 | 1 | 1 | 1 | 0 | 0 | 0 |
| UBE3B | 0.103 | P2_P3_P4_Lumbar_VS_P1_Coeliac_Stellate_Sphenopalatine | 0 | 1 | 1 | 1 | 1 | 0 | 0 | 0 |
| HERPUD1 | 0.103 | P1_P2_P3_P4_Lumbar_Coeliac_Stellate_VS_Sphenopalatine | 1 | 1 | 1 | 1 | 1 | 1 | 1 | 0 |
| ME3 | 0.103 | Sphenopalatine_VS_P1_P2_P3_P4_Lumbar_Coeliac_Stellate | 0 | 0 | 0 | 0 | 0 | 0 | 0 | 1 |
| SLCO3A1 | 0.103 | P1_P2_P4_Lumbar_Coeliac_Stellate_Sphenopalatine_VS_P3 | 1 | 1 | 0 | 1 | 1 | 1 | 1 | 1 |
| E330009J07F | 0.102 | P2_Lumbar_VS_P1_P3_P4_Coeliac_Stellate_Sphenopalatine | 0 | 1 | 0 | 0 | 1 | 0 | 0 | 0 |

| | | | | | | | | | | |
|---|---|---|---|---|---|---|---|---|---|---|
| TMA16 | 0.102 | P1_P3_P4_Lumbar_VS_P2_Coeliac_Stellate_Sphenopalatine | 1 | 0 | 1 | 1 | 1 | 0 | 0 | 0 |
| JAM3 | 0.102 | P2_P3_P4_Lumbar_VS_P1_Coeliac_Stellate_Sphenopalatine | 0 | 1 | 1 | 1 | 1 | 0 | 0 | 0 |
| SREK1IP1 | 0.102 | P1_P2_P3_P4_Lumbar_Coeliac_Stellate_VS_Sphenopalatine | 1 | 1 | 1 | 1 | 1 | 1 | 1 | 0 |
| TCERG1 | 0.102 | P1_P2_P3_P4_Lumbar_Coeliac_Stellate_VS_Sphenopalatine | 1 | 1 | 1 | 1 | 1 | 1 | 1 | 0 |
| RSPH9 | 0.102 | P1_P3_P4_Lumbar_Coeliac_Stellate_Sphenopalatine_VS_P2 | 1 | 0 | 1 | 1 | 1 | 1 | 1 | 1 |
| ENOPH1 | 0.101 | P3_P4_VS_P1_P2_Lumbar_Coeliac_Stellate_Sphenopalatine | 0 | 0 | 1 | 1 | 0 | 0 | 0 | 0 |
| GSN | 0.101 | P1_P2_P3_P4_Lumbar_Coeliac_Stellate_VS_Sphenopalatine | 1 | 1 | 1 | 1 | 1 | 1 | 1 | 0 |
| ARHGAP28 | 0.101 | P1_P2_P3_P4_Lumbar_Coeliac_Stellate_VS_Sphenopalatine | 1 | 1 | 1 | 1 | 1 | 1 | 1 | 0 |
| ARHGAP33 | 0.101 | Lumbar_Coeliac_Sphenopalatine_VS_P1_P2_P3_P4_Stellate | 0 | 0 | 0 | 0 | 1 | 1 | 0 | 1 |
| MTF2 | 0.101 | P1_P3_Lumbar_Coeliac_Sphenopalatine_VS_P2_P4_Stellate | 1 | 0 | 1 | 0 | 1 | 1 | 0 | 1 |
| SNHG5 | 0.101 | P1_P2_P3_Stellate_Sphenopalatine_VS_P4_Lumbar_Coeliac | 1 | 1 | 1 | 0 | 0 | 0 | 1 | 1 |
| ZGPAT | 0.1 | P1_Lumbar_VS_P2_P3_P4_Coeliac_Stellate_Sphenopalatine | 1 | 0 | 0 | 0 | 1 | 0 | 0 | 0 |
| CYB5R1 | 0.1 | P1_P3_P4_Lumbar_VS_P2_Coeliac_Stellate_Sphenopalatine | 1 | 0 | 1 | 1 | 1 | 0 | 0 | 0 |
| ATG13 | 0.1 | P1_P2_P4_Coeliac_Sphenopalatine_VS_P3_Lumbar_Stellate | 1 | 1 | 0 | 1 | 0 | 1 | 0 | 1 |
| DUS2 | 0.0997 | P1_P3_Lumbar_Coeliac_Sphenopalatine_VS_P2_P4_Stellate | 1 | 0 | 1 | 0 | 1 | 1 | 0 | 1 |
| CYGB | 0.0994 | P1_P2_P3_Lumbar_Coeliac_Stellate_Sphenopalatine_VS_P4 | 1 | 1 | 1 | 0 | 1 | 1 | 1 | 1 |
| IFT57 | 0.0993 | P1_P3_P4_Lumbar_VS_P2_Coeliac_Stellate_Sphenopalatine | 1 | 0 | 1 | 1 | 1 | 0 | 0 | 0 |
| OIP5OS1 | 0.0991 | P2_Coeliac_VS_P1_P3_P4_Lumbar_Stellate_Sphenopalatine | 0 | 1 | 0 | 0 | 0 | 1 | 0 | 0 |
| KDM5C | 0.0988 | P1_P2_Lumbar_Coeliac_Sphenopalatine_VS_P3_P4_Stellate | 1 | 1 | 0 | 0 | 1 | 1 | 0 | 1 |
| SURF6 | 0.0987 | P1_P2_Lumbar_Coeliac_Sphenopalatine_VS_P3_P4_Stellate | 1 | 1 | 0 | 0 | 1 | 1 | 0 | 1 |
| PRUNE2 | 0.0986 | P1_P3_P4_Lumbar_Sphenopalatine_VS_P2_Coeliac_Stellate | 1 | 0 | 1 | 1 | 1 | 0 | 0 | 1 |
| NOP9 | 0.0979 | P1_Lumbar_VS_P2_P3_P4_Coeliac_Stellate_Sphenopalatine | 1 | 0 | 0 | 0 | 1 | 0 | 0 | 0 |
| VPS54 | 0.0977 | P2_Lumbar_VS_P1_P3_P4_Coeliac_Stellate_Sphenopalatine | 0 | 1 | 0 | 0 | 1 | 0 | 0 | 0 |
| NUDT19 | 0.0975 | P1_P2_P3_P4_Lumbar_Stellate_Sphenopalatine_VS_Coeliac | 1 | 1 | 1 | 1 | 1 | 0 | 1 | 1 |
| LATS1 | 0.0972 | P2_P3_P4_Lumbar_VS_P1_Coeliac_Stellate_Sphenopalatine | 0 | 1 | 1 | 1 | 1 | 0 | 0 | 0 |
| YBX3 | 0.0972 | Lumbar_Coeliac_Sphenopalatine_VS_P1_P2_P3_P4_Stellate | 0 | 0 | 0 | 0 | 1 | 1 | 0 | 1 |
| COMTD1 | 0.0972 | Lumbar_Coeliac_Sphenopalatine_VS_P1_P2_P3_P4_Stellate | 0 | 0 | 0 | 0 | 1 | 1 | 0 | 1 |
| AFAP1 | 0.0972 | P1_P2_P4_Lumbar_Coeliac_Stellate_Sphenopalatine_VS_P3 | 1 | 1 | 0 | 1 | 1 | 1 | 1 | 1 |
| SCT | 0.0969 | P3_Lumbar_Coeliac_Stellate_VS_P1_P2_P4_Sphenopalatine | 0 | 0 | 1 | 0 | 1 | 1 | 1 | 0 |
| KCNAB2 | 0.0968 | P2_P3_P4_VS_P1_Lumbar_Coeliac_Stellate_Sphenopalatine | 0 | 1 | 1 | 1 | 0 | 0 | 0 | 0 |

| | | | | | | | | | | |
|---|---|---|---|---|---|---|---|---|---|---|
| FBXL6 | 0.0966 | P1_Lumbar_VS_P2_P3_P4_Coeliac_Stellate_Sphenopalatine | 1 | 0 | 0 | 0 | 1 | 0 | 0 | 0 |
| MTF1 | 0.0966 | P1_P2_P3_P4_Lumbar_Coeliac_Stellate_VS_Sphenopalatine | 1 | 1 | 1 | 1 | 1 | 1 | 1 | 0 |
| AK2 | 0.0965 | P1_P2_P3_P4_Lumbar_Coeliac_Stellate_VS_Sphenopalatine | 1 | 1 | 1 | 1 | 1 | 1 | 1 | 0 |
| DBT | 0.0962 | Lumbar_Sphenopalatine_VS_P1_P2_P3_P4_Coeliac_Stellate | 0 | 0 | 0 | 0 | 1 | 0 | 0 | 1 |
| MAP4K3 | 0.0961 | P2_Coeliac_VS_P1_P3_P4_Lumbar_Stellate_Sphenopalatine | 0 | 1 | 0 | 0 | 0 | 1 | 0 | 0 |
| UBN1 | 0.096 | P2_P3_P4_Lumbar_VS_P1_Coeliac_Stellate_Sphenopalatine | 0 | 1 | 1 | 1 | 1 | 0 | 0 | 0 |
| MRNIP | 0.0955 | P1_P3_P4_Lumbar_VS_P2_Coeliac_Stellate_Sphenopalatine | 1 | 0 | 1 | 1 | 1 | 0 | 0 | 0 |
| SSPN | 0.0953 | P1_Lumbar_VS_P2_P3_P4_Coeliac_Stellate_Sphenopalatine | 1 | 0 | 0 | 0 | 1 | 0 | 0 | 0 |
| YTHDF1 | 0.0952 | P2_P3_P4_Lumbar_VS_P1_Coeliac_Stellate_Sphenopalatine | 0 | 1 | 1 | 1 | 1 | 0 | 0 | 0 |
| TMEM9B | 0.0952 | P1_P2_P3_P4_Lumbar_Stellate_Sphenopalatine_VS_Coeliac | 1 | 1 | 1 | 1 | 1 | 0 | 1 | 1 |
| B9D2 | 0.0952 | P1_P3_P4_Lumbar_Coeliac_Stellate_Sphenopalatine_VS_P2 | 1 | 0 | 1 | 1 | 1 | 1 | 1 | 1 |
| SGPL1 | 0.0951 | P2_Coeliac_VS_P1_P3_P4_Lumbar_Stellate_Sphenopalatine | 0 | 1 | 0 | 0 | 0 | 1 | 0 | 0 |
| TMC7 | 0.0951 | Lumbar_Coeliac_Sphenopalatine_VS_P1_P2_P3_P4_Stellate | 0 | 0 | 0 | 0 | 1 | 1 | 0 | 1 |
| TNPO2 | 0.0951 | P1_P2_Lumbar_Coeliac_Sphenopalatine_VS_P3_P4_Stellate | 1 | 1 | 0 | 0 | 1 | 1 | 0 | 1 |
| SMAD2 | 0.0949 | Lumbar_Coeliac_Sphenopalatine_VS_P1_P2_P3_P4_Stellate | 0 | 0 | 0 | 0 | 1 | 1 | 0 | 1 |
| DICER1 | 0.0948 | P2_P3_P4_Lumbar_VS_P1_Coeliac_Stellate_Sphenopalatine | 0 | 1 | 1 | 1 | 1 | 0 | 0 | 0 |
| MEGF8 | 0.0942 | P2_P4_Sphenopalatine_VS_P1_P3_Lumbar_Coeliac_Stellate | 0 | 1 | 0 | 1 | 0 | 0 | 0 | 1 |
| ATAT1 | 0.0941 | Lumbar_Coeliac_Sphenopalatine_VS_P1_P2_P3_P4_Stellate | 0 | 0 | 0 | 0 | 1 | 1 | 0 | 1 |
| GFOD2 | 0.094 | P1_P3_Lumbar_Coeliac_Sphenopalatine_VS_P2_P4_Stellate | 1 | 0 | 1 | 0 | 1 | 1 | 0 | 1 |
| PAQR7 | 0.0938 | P2_P3_P4_VS_P1_Lumbar_Coeliac_Stellate_Sphenopalatine | 0 | 1 | 1 | 1 | 0 | 0 | 0 | 0 |
| TRUB1 | 0.0938 | P1_Lumbar_VS_P2_P3_P4_Coeliac_Stellate_Sphenopalatine | 1 | 0 | 0 | 0 | 1 | 0 | 0 | 0 |
| PLK3 | 0.0937 | P1_P3_Lumbar_VS_P2_P4_Coeliac_Stellate_Sphenopalatine | 1 | 0 | 1 | 0 | 1 | 0 | 0 | 0 |
| CFAP410 | 0.0936 | P1_P2_P3_P4_Lumbar_Stellate_Sphenopalatine_VS_Coeliac | 1 | 1 | 1 | 1 | 1 | 0 | 1 | 1 |
| TMEM8B | 0.0935 | P1_P2_Lumbar_Coeliac_Sphenopalatine_VS_P3_P4_Stellate | 1 | 1 | 0 | 0 | 1 | 1 | 0 | 1 |
| SLC41A2 | 0.0929 | P4_Sphenopalatine_VS_P1_P2_P3_Lumbar_Coeliac_Stellate | 0 | 0 | 0 | 1 | 0 | 0 | 0 | 1 |
| BLVRB | 0.0928 | P2_P3_P4_VS_P1_Lumbar_Coeliac_Stellate_Sphenopalatine | 0 | 1 | 1 | 1 | 0 | 0 | 0 | 0 |
| ALDH2 | 0.0923 | P1_P2_P4_Lumbar_Coeliac_Stellate_Sphenopalatine_VS_P3 | 1 | 1 | 0 | 1 | 1 | 1 | 1 | 1 |
| 1600020E01I | 0.092 | P1_Lumbar_Coeliac_Sphenopalatine_VS_P2_P3_P4_Stellate | 1 | 0 | 0 | 0 | 1 | 1 | 0 | 1 |
| DDX28 | 0.0918 | P1_Lumbar_VS_P2_P3_P4_Coeliac_Stellate_Sphenopalatine | 1 | 0 | 0 | 0 | 1 | 0 | 0 | 0 |
| STK38 | 0.0918 | Lumbar_Coeliac_Sphenopalatine_VS_P1_P2_P3_P4_Stellate | 0 | 0 | 0 | 0 | 1 | 1 | 0 | 1 |

| | | | | | | | | | | |
|---|---|---|---|---|---|---|---|---|---|---|
| ABT1 | 0.0917 | P1_Lumbar_VS_P2_P3_P4_Coeliac_Stellate_Sphenopalatine | 1 | 0 | 0 | 0 | 1 | 0 | 0 | 0 |
| USP46 | 0.0916 | P2_Lumbar_VS_P1_P3_P4_Coeliac_Stellate_Sphenopalatine | 0 | 1 | 0 | 0 | 1 | 0 | 0 | 0 |
| FOXP1 | 0.0916 | Lumbar_Coeliac_Stellate_VS_P1_P2_P3_P4_Sphenopalatine | 0 | 0 | 0 | 0 | 1 | 1 | 1 | 0 |
| E130309D02 | 0.0912 | P1_P2_P3_Lumbar_Coeliac_Stellate_Sphenopalatine_VS_P4 | 1 | 1 | 1 | 0 | 1 | 1 | 1 | 1 |
| NIPA1 | 0.091 | P1_P3_Lumbar_Coeliac_Sphenopalatine_VS_P2_P4_Stellate | 1 | 0 | 1 | 0 | 1 | 1 | 0 | 1 |
| DNM2 | 0.0909 | Lumbar_Coeliac_Sphenopalatine_VS_P1_P2_P3_P4_Stellate | 0 | 0 | 0 | 0 | 1 | 1 | 0 | 1 |
| PLEKHB2 | 0.0909 | P1_P2_P3_P4_Lumbar_Stellate_Sphenopalatine_VS_Coeliac | 1 | 1 | 1 | 1 | 1 | 0 | 1 | 1 |
| DACT3 | 0.0906 | P1_P4_Lumbar_Sphenopalatine_VS_P2_P3_Coeliac_Stellate | 1 | 0 | 0 | 1 | 1 | 0 | 0 | 1 |
| CREB1 | 0.0904 | Lumbar_Coeliac_Sphenopalatine_VS_P1_P2_P3_P4_Stellate | 0 | 0 | 0 | 0 | 1 | 1 | 0 | 1 |
| PHYHIPL | 0.0903 | P3_Lumbar_Coeliac_Sphenopalatine_VS_P1_P2_P4_Stellate | 0 | 0 | 1 | 0 | 1 | 1 | 0 | 1 |
| SBF1 | 0.09 | P2_P3_P4_Lumbar_Sphenopalatine_VS_P1_Coeliac_Stellate | 0 | 1 | 1 | 1 | 1 | 0 | 0 | 1 |
| AATK | 0.0898 | P2_P4_Sphenopalatine_VS_P1_P3_Lumbar_Coeliac_Stellate | 0 | 1 | 0 | 1 | 0 | 0 | 0 | 1 |
| MREG | 0.0897 | P3_Sphenopalatine_VS_P1_P2_P4_Lumbar_Coeliac_Stellate | 0 | 0 | 1 | 0 | 0 | 0 | 0 | 1 |
| SPIRE1 | 0.0896 | P2_P3_P4_Lumbar_VS_P1_Coeliac_Stellate_Sphenopalatine | 0 | 1 | 1 | 1 | 1 | 0 | 0 | 0 |
| CSRP2 | 0.0896 | P1_P3_P4_Lumbar_Coeliac_Stellate_Sphenopalatine_VS_P2 | 1 | 0 | 1 | 1 | 1 | 1 | 1 | 1 |
| NFAT5 | 0.0895 | P2_Lumbar_VS_P1_P3_P4_Coeliac_Stellate_Sphenopalatine | 0 | 1 | 0 | 0 | 1 | 0 | 0 | 0 |
| APOE | 0.0893 | P1_P2_P3_P4_Coeliac_Stellate_Sphenopalatine_VS_Lumbar | 1 | 1 | 1 | 1 | 0 | 1 | 1 | 1 |
| PLXNA1 | 0.0892 | P2_Lumbar_VS_P1_P3_P4_Coeliac_Stellate_Sphenopalatine | 0 | 1 | 0 | 0 | 1 | 0 | 0 | 0 |
| CDC5L | 0.0892 | Lumbar_Coeliac_Sphenopalatine_VS_P1_P2_P3_P4_Stellate | 0 | 0 | 0 | 0 | 1 | 1 | 0 | 1 |
| VPS13B | 0.0891 | P3_P4_Coeliac_VS_P1_P2_Lumbar_Stellate_Sphenopalatine | 0 | 0 | 1 | 1 | 0 | 1 | 0 | 0 |
| PRR7 | 0.089 | P1_P3_P4_Lumbar_VS_P2_Coeliac_Stellate_Sphenopalatine | 1 | 0 | 1 | 1 | 1 | 0 | 0 | 0 |
| DAB2IP | 0.089 | P1_P2_Lumbar_Coeliac_Sphenopalatine_VS_P3_P4_Stellate | 1 | 1 | 0 | 0 | 1 | 1 | 0 | 1 |
| TRIB2 | 0.089 | P1_P2_P3_P4_Lumbar_Stellate_Sphenopalatine_VS_Coeliac | 1 | 1 | 1 | 1 | 1 | 0 | 1 | 1 |
| LNPK | 0.0889 | P4_Lumbar_Coeliac_Stellate_VS_P1_P2_P3_Sphenopalatine | 0 | 0 | 0 | 1 | 1 | 1 | 1 | 0 |
| RPA3 | 0.0886 | P1_P3_P4_Lumbar_Coeliac_Stellate_VS_P2_Sphenopalatine | 1 | 0 | 1 | 1 | 1 | 1 | 1 | 0 |
| ATP2B2 | 0.0884 | P2_Lumbar_VS_P1_P3_P4_Coeliac_Stellate_Sphenopalatine | 0 | 1 | 0 | 0 | 1 | 0 | 0 | 0 |
| ZFP811 | 0.0884 | P2_Coeliac_VS_P1_P3_P4_Lumbar_Stellate_Sphenopalatine | 0 | 1 | 0 | 0 | 0 | 1 | 0 | 0 |
| SMARCA1 | 0.0882 | P3_P4_Coeliac_VS_P1_P2_Lumbar_Stellate_Sphenopalatine | 0 | 0 | 1 | 1 | 0 | 1 | 0 | 0 |
| SENP8 | 0.0882 | Lumbar_Coeliac_Sphenopalatine_VS_P1_P2_P3_P4_Stellate | 0 | 0 | 0 | 0 | 1 | 1 | 0 | 1 |
| ATMIN | 0.0881 | P2_P3_P4_Lumbar_Sphenopalatine_VS_P1_Coeliac_Stellate | 0 | 1 | 1 | 1 | 1 | 0 | 0 | 1 |

| | | | | | | | | | | |
|---|---|---|---|---|---|---|---|---|---|---|
| ZFP319 | 0.088 | P2_Lumbar_VS_P1_P3_P4_Coeliac_Stellate_Sphenopalatine | 0 | 1 | 0 | 0 | 1 | 0 | 0 | 0 |
| GM26614 | 0.0876 | P1_P2_P3_VS_P4_Lumbar_Coeliac_Stellate_Sphenopalatine | 1 | 1 | 1 | 0 | 0 | 0 | 0 | 0 |
| PFKL | 0.0875 | P2_P3_Lumbar_Coeliac_Sphenopalatine_VS_P1_P4_Stellate | 0 | 1 | 1 | 0 | 1 | 1 | 0 | 1 |
| KLHL28 | 0.0874 | P2_Lumbar_VS_P1_P3_P4_Coeliac_Stellate_Sphenopalatine | 0 | 1 | 0 | 0 | 1 | 0 | 0 | 0 |
| FAM20B | 0.0874 | P2_P3_P4_Lumbar_VS_P1_Coeliac_Stellate_Sphenopalatine | 0 | 1 | 1 | 1 | 1 | 0 | 0 | 0 |
| GPATCH4 | 0.0871 | P1_P3_Lumbar_Coeliac_Sphenopalatine_VS_P2_P4_Stellate | 1 | 0 | 1 | 0 | 1 | 1 | 0 | 1 |
| PRXL2C | 0.087 | P1_P2_Lumbar_Coeliac_Sphenopalatine_VS_P3_P4_Stellate | 1 | 1 | 0 | 0 | 1 | 1 | 0 | 1 |
| TENT5A | 0.0869 | P4_Coeliac_Stellate_VS_P1_P2_P3_Lumbar_Sphenopalatine | 0 | 0 | 0 | 1 | 0 | 1 | 1 | 0 |
| SRRM3 | 0.0869 | P2_Lumbar_Coeliac_Sphenopalatine_VS_P1_P3_P4_Stellate | 0 | 1 | 0 | 0 | 1 | 1 | 0 | 1 |
| FAM214A | 0.0866 | P3_P4_Coeliac_VS_P1_P2_Lumbar_Stellate_Sphenopalatine | 0 | 0 | 1 | 1 | 0 | 1 | 0 | 0 |
| TUB | 0.0863 | P1_P2_P3_Lumbar_Coeliac_Stellate_Sphenopalatine_VS_P4 | 1 | 1 | 1 | 0 | 1 | 1 | 1 | 1 |
| DEFB42 | 0.0862 | P1_P2_P3_VS_P4_Lumbar_Coeliac_Stellate_Sphenopalatine | 1 | 1 | 1 | 0 | 0 | 0 | 0 | 0 |
| SRD5A3 | 0.0862 | P1_P3_P4_Lumbar_VS_P2_Coeliac_Stellate_Sphenopalatine | 1 | 0 | 1 | 1 | 1 | 0 | 0 | 0 |
| OGFOD1 | 0.0861 | P2_P3_P4_Lumbar_VS_P1_Coeliac_Stellate_Sphenopalatine | 0 | 1 | 1 | 1 | 1 | 0 | 0 | 0 |
| ARMCX4 | 0.0861 | P1_P2_Lumbar_Coeliac_Sphenopalatine_VS_P3_P4_Stellate | 1 | 1 | 0 | 0 | 1 | 1 | 0 | 1 |
| DAG1 | 0.0859 | P1_P2_P4_Coeliac_Sphenopalatine_VS_P3_Lumbar_Stellate | 1 | 1 | 0 | 1 | 0 | 1 | 0 | 1 |
| LNPEP | 0.0858 | P1_P2_P4_Coeliac_Sphenopalatine_VS_P3_Lumbar_Stellate | 1 | 1 | 0 | 1 | 0 | 1 | 0 | 1 |
| STRADA | 0.0853 | P1_P2_Lumbar_Coeliac_Sphenopalatine_VS_P3_P4_Stellate | 1 | 1 | 0 | 0 | 1 | 1 | 0 | 1 |
| ZCCHC12 | 0.0852 | P1_P3_P4_Lumbar_Coeliac_Stellate_Sphenopalatine_VS_P2 | 1 | 0 | 1 | 1 | 1 | 1 | 1 | 1 |
| ETFDH | 0.085 | P4_Lumbar_Coeliac_Stellate_VS_P1_P2_P3_Sphenopalatine | 0 | 0 | 0 | 1 | 1 | 1 | 1 | 0 |
| MAPK12 | 0.085 | P1_P2_P4_Lumbar_Coeliac_Stellate_Sphenopalatine_VS_P3 | 1 | 1 | 0 | 1 | 1 | 1 | 1 | 1 |
| GTF2E2 | 0.0849 | P1_P3_P4_Lumbar_Coeliac_Stellate_Sphenopalatine_VS_P2 | 1 | 0 | 1 | 1 | 1 | 1 | 1 | 1 |
| APOOL | 0.0844 | P3_Lumbar_Coeliac_Stellate_VS_P1_P2_P4_Sphenopalatine | 0 | 0 | 1 | 0 | 1 | 1 | 1 | 0 |
| SBSN | 0.0843 | P1_P2_P3_Lumbar_Coeliac_Stellate_Sphenopalatine_VS_P4 | 1 | 1 | 1 | 0 | 1 | 1 | 1 | 1 |
| ZFP398 | 0.084 | P3_Lumbar_Coeliac_Sphenopalatine_VS_P1_P2_P4_Stellate | 0 | 0 | 1 | 0 | 1 | 1 | 0 | 1 |
| TMEM17 | 0.0835 | P4_Lumbar_Coeliac_Stellate_VS_P1_P2_P3_Sphenopalatine | 0 | 0 | 0 | 1 | 1 | 1 | 1 | 0 |
| MINDY1 | 0.0834 | P3_P4_Lumbar_Coeliac_Stellate_VS_P1_P2_Sphenopalatine | 0 | 0 | 1 | 1 | 1 | 1 | 1 | 0 |
| UCK2 | 0.0831 | P1_P2_Lumbar_Coeliac_Sphenopalatine_VS_P3_P4_Stellate | 1 | 1 | 0 | 0 | 1 | 1 | 0 | 1 |
| DNTTIP1 | 0.083 | P1_P3_P4_Lumbar_VS_P2_Coeliac_Stellate_Sphenopalatine | 1 | 0 | 1 | 1 | 1 | 0 | 0 | 0 |
| SGPP1 | 0.0828 | P2_P3_P4_VS_P1_Lumbar_Coeliac_Stellate_Sphenopalatine | 0 | 1 | 1 | 1 | 0 | 0 | 0 | 0 |

| | | | | | | | | | | |
|---|---|---|---|---|---|---|---|---|---|---|
| DNM1 | 0.0826 | P4_Lumbar_Sphenopalatine_VS_P1_P2_P3_Coeliac_Stellate | 0 | 0 | 0 | 1 | 1 | 0 | 0 | 1 |
| ITGB3BP | 0.0824 | P1_P3_P4_Lumbar_Coeliac_Stellate_VS_P2_Sphenopalatine | 1 | 0 | 1 | 1 | 1 | 1 | 1 | 0 |
| PLCD4 | 0.0824 | P4_Sphenopalatine_VS_P1_P2_P3_Lumbar_Coeliac_Stellate | 0 | 0 | 0 | 1 | 0 | 0 | 0 | 1 |
| ING4 | 0.0817 | P1_P3_Lumbar_Coeliac_Sphenopalatine_VS_P2_P4_Stellate | 1 | 0 | 1 | 0 | 1 | 1 | 0 | 1 |
| CETN4 | 0.0816 | P3_P4_Coeliac_VS_P1_P2_Lumbar_Stellate_Sphenopalatine | 0 | 0 | 1 | 1 | 0 | 1 | 0 | 0 |
| DHRS7B | 0.0815 | P1_P3_P4_Lumbar_VS_P2_Coeliac_Stellate_Sphenopalatine | 1 | 0 | 1 | 1 | 1 | 0 | 0 | 0 |
| ACOT2 | 0.0815 | P2_Coeliac_VS_P1_P3_P4_Lumbar_Stellate_Sphenopalatine | 0 | 1 | 0 | 0 | 0 | 1 | 0 | 0 |
| COG7 | 0.0814 | P1_P3_P4_Lumbar_VS_P2_Coeliac_Stellate_Sphenopalatine | 1 | 0 | 1 | 1 | 1 | 0 | 0 | 0 |
| UBXN2A | 0.0813 | P3_Lumbar_Coeliac_Sphenopalatine_VS_P1_P2_P4_Stellate | 0 | 0 | 1 | 0 | 1 | 1 | 0 | 1 |
| R3HCC1 | 0.0813 | P1_P3_Lumbar_Coeliac_Sphenopalatine_VS_P2_P4_Stellate | 1 | 0 | 1 | 0 | 1 | 1 | 0 | 1 |
| RANBP6 | 0.0812 | P1_P3_Lumbar_Coeliac_Sphenopalatine_VS_P2_P4_Stellate | 1 | 0 | 1 | 0 | 1 | 1 | 0 | 1 |
| 2210016F16I | 0.0806 | P1_P3_P4_Lumbar_Coeliac_Stellate_VS_P2_Sphenopalatine | 1 | 0 | 1 | 1 | 1 | 1 | 1 | 0 |
| MRPL16 | 0.0805 | P1_P3_P4_Lumbar_Coeliac_Stellate_VS_P2_Sphenopalatine | 1 | 0 | 1 | 1 | 1 | 1 | 1 | 0 |
| GALNT1 | 0.0801 | P2_Coeliac_VS_P1_P3_P4_Lumbar_Stellate_Sphenopalatine | 0 | 1 | 0 | 0 | 0 | 1 | 0 | 0 |
| TMEM141 | 0.08 | P2_P3_P4_Lumbar_Sphenopalatine_VS_P1_Coeliac_Stellate | 0 | 1 | 1 | 1 | 1 | 0 | 0 | 1 |
| GRIA4 | 0.0795 | P2_P3_Lumbar_Coeliac_Sphenopalatine_VS_P1_P4_Stellate | 0 | 1 | 1 | 0 | 1 | 1 | 0 | 1 |
| CMC2 | 0.0794 | P4_Sphenopalatine_VS_P1_P2_P3_Lumbar_Coeliac_Stellate | 0 | 0 | 0 | 1 | 0 | 0 | 0 | 1 |
| SNX25 | 0.0792 | P2_P3_P4_Lumbar_Coeliac_Stellate_VS_P1_Sphenopalatine | 0 | 1 | 1 | 1 | 1 | 1 | 1 | 0 |
| GIT1 | 0.0792 | P2_P3_P4_Lumbar_Sphenopalatine_VS_P1_Coeliac_Stellate | 0 | 1 | 1 | 1 | 1 | 0 | 0 | 1 |
| CCNL1 | 0.079 | P2_Coeliac_VS_P1_P3_P4_Lumbar_Stellate_Sphenopalatine | 0 | 1 | 0 | 0 | 0 | 1 | 0 | 0 |
| CPNE3 | 0.079 | P1_P3_P4_Lumbar_Sphenopalatine_VS_P2_Coeliac_Stellate | 1 | 0 | 1 | 1 | 1 | 0 | 0 | 1 |
| GUCY1A1 | 0.0787 | P1_P2_P3_VS_P4_Lumbar_Coeliac_Stellate_Sphenopalatine | 1 | 1 | 1 | 0 | 0 | 0 | 0 | 0 |
| ZFP229 | 0.0787 | P3_P4_Coeliac_VS_P1_P2_Lumbar_Stellate_Sphenopalatine | 0 | 0 | 1 | 1 | 0 | 1 | 0 | 0 |
| CLIP2 | 0.0787 | P2_P4_Lumbar_Sphenopalatine_VS_P1_P3_Coeliac_Stellate | 0 | 1 | 0 | 1 | 1 | 0 | 0 | 1 |
| FOXJ2 | 0.0785 | P4_Coeliac_Sphenopalatine_VS_P1_P2_P3_Lumbar_Stellate | 0 | 0 | 0 | 1 | 0 | 1 | 0 | 1 |
| GPD2 | 0.0785 | P1_P3_Lumbar_Coeliac_Sphenopalatine_VS_P2_P4_Stellate | 1 | 0 | 1 | 0 | 1 | 1 | 0 | 1 |
| AGO1 | 0.0785 | P2_P3_Lumbar_Coeliac_Sphenopalatine_VS_P1_P4_Stellate | 0 | 1 | 1 | 0 | 1 | 1 | 0 | 1 |
| ANKFY1 | 0.0783 | P2_P3_P4_Lumbar_Sphenopalatine_VS_P1_Coeliac_Stellate | 0 | 1 | 1 | 1 | 1 | 0 | 0 | 1 |
| PPP1R35 | 0.0782 | P1_P3_P4_Lumbar_Coeliac_Stellate_Sphenopalatine_VS_P2 | 1 | 0 | 1 | 1 | 1 | 1 | 1 | 1 |
| PPP1R9B | 0.078 | P2_P3_P4_Lumbar_Sphenopalatine_VS_P1_Coeliac_Stellate | 0 | 1 | 1 | 1 | 1 | 0 | 0 | 1 |

| | | | | | | | | | | |
|---|---|---|---|---|---|---|---|---|---|---|
| KALRN | 0.0779 | P2_P3_P4_Lumbar_Sphenopalatine_VS_P1_Coeliac_Stellate | 0 | 1 | 1 | 1 | 1 | 0 | 0 | 1 |
| EIF2AK1 | 0.0776 | P2_P3_Lumbar_Coeliac_Sphenopalatine_VS_P1_P4_Stellate | 0 | 1 | 1 | 0 | 1 | 1 | 0 | 1 |
| EXOSC7 | 0.0768 | P1_P3_P4_Lumbar_Coeliac_Stellate_Sphenopalatine_VS_P2 | 1 | 0 | 1 | 1 | 1 | 1 | 1 | 1 |
| ARHGEF17 | 0.0766 | P2_P3_P4_Lumbar_Sphenopalatine_VS_P1_Coeliac_Stellate | 0 | 1 | 1 | 1 | 1 | 0 | 0 | 1 |
| PAIP2B | 0.0764 | P2_P3_P4_VS_P1_Lumbar_Coeliac_Stellate_Sphenopalatine | 0 | 1 | 1 | 1 | 0 | 0 | 0 | 0 |
| FBXO33 | 0.0763 | P2_P3_P4_Coeliac_VS_P1_Lumbar_Stellate_Sphenopalatine | 0 | 1 | 1 | 1 | 0 | 1 | 0 | 0 |
| CCDC112 | 0.076 | P1_Lumbar_Coeliac_Sphenopalatine_VS_P2_P3_P4_Stellate | 1 | 0 | 0 | 0 | 1 | 1 | 0 | 1 |
| ANKIB1 | 0.076 | P2_P3_Lumbar_Coeliac_Sphenopalatine_VS_P1_P4_Stellate | 0 | 1 | 1 | 0 | 1 | 1 | 0 | 1 |
| TMEM203 | 0.0759 | P1_P3_P4_Lumbar_VS_P2_Coeliac_Stellate_Sphenopalatine | 1 | 0 | 1 | 1 | 1 | 0 | 0 | 0 |
| INO80C | 0.075 | P1_P2_P3_Lumbar_Sphenopalatine_VS_P4_Coeliac_Stellate | 1 | 1 | 1 | 0 | 1 | 0 | 0 | 1 |
| NUDT14 | 0.0748 | P1_P3_P4_Lumbar_Coeliac_Stellate_Sphenopalatine_VS_P2 | 1 | 0 | 1 | 1 | 1 | 1 | 1 | 1 |
| GKAP1 | 0.0746 | P1_P3_Lumbar_Coeliac_Sphenopalatine_VS_P2_P4_Stellate | 1 | 0 | 1 | 0 | 1 | 1 | 0 | 1 |
| GPATCH11 | 0.0744 | P1_P3_P4_Lumbar_Coeliac_Stellate_Sphenopalatine_VS_P2 | 1 | 0 | 1 | 1 | 1 | 1 | 1 | 1 |
| GSTP1 | 0.074 | P1_P2_P3_P4_Coeliac_Stellate_Sphenopalatine_VS_Lumbar | 1 | 1 | 1 | 1 | 0 | 1 | 1 | 1 |
| RPTOR | 0.0739 | P2_P4_Lumbar_Sphenopalatine_VS_P1_P3_Coeliac_Stellate | 0 | 1 | 0 | 1 | 1 | 0 | 0 | 1 |
| CDC42BPA | 0.0739 | P2_P3_P4_Lumbar_Sphenopalatine_VS_P1_Coeliac_Stellate | 0 | 1 | 1 | 1 | 1 | 0 | 0 | 1 |
| MRPL22 | 0.0738 | P1_P3_P4_Lumbar_Coeliac_Stellate_VS_P2_Sphenopalatine | 1 | 0 | 1 | 1 | 1 | 1 | 1 | 0 |
| IGF1R | 0.0735 | P1_P2_P4_Lumbar_Coeliac_Stellate_Sphenopalatine_VS_P3 | 1 | 1 | 0 | 1 | 1 | 1 | 1 | 1 |
| RND2 | 0.0734 | P2_P3_Lumbar_Coeliac_Sphenopalatine_VS_P1_P4_Stellate | 0 | 1 | 1 | 0 | 1 | 1 | 0 | 1 |
| RRP15 | 0.0731 | P1_P3_P4_Lumbar_VS_P2_Coeliac_Stellate_Sphenopalatine | 1 | 0 | 1 | 1 | 1 | 0 | 0 | 0 |
| SHARPIN | 0.0731 | P1_P3_P4_Lumbar_VS_P2_Coeliac_Stellate_Sphenopalatine | 1 | 0 | 1 | 1 | 1 | 0 | 0 | 0 |
| SPOCK3 | 0.0731 | P1_P2_P3_Sphenopalatine_VS_P4_Lumbar_Coeliac_Stellate | 1 | 1 | 1 | 0 | 0 | 0 | 0 | 1 |
| SEMA4F | 0.0728 | P2_P3_P4_Lumbar_Sphenopalatine_VS_P1_Coeliac_Stellate | 0 | 1 | 1 | 1 | 1 | 0 | 0 | 1 |
| FAM122A | 0.0726 | P1_P3_P4_Lumbar_Sphenopalatine_VS_P2_Coeliac_Stellate | 1 | 0 | 1 | 1 | 1 | 0 | 0 | 1 |
| RNF13 | 0.0725 | P2_P3_P4_VS_P1_Lumbar_Coeliac_Stellate_Sphenopalatine | 0 | 1 | 1 | 1 | 0 | 0 | 0 | 0 |
| TMEM38A | 0.0724 | P1_P2_P3_P4_Lumbar_Stellate_Sphenopalatine_VS_Coeliac | 1 | 1 | 1 | 1 | 1 | 0 | 1 | 1 |
| 2610301B20I | 0.0723 | P1_P3_P4_Lumbar_VS_P2_Coeliac_Stellate_Sphenopalatine | 1 | 0 | 1 | 1 | 1 | 0 | 0 | 0 |
| PPP1R9A | 0.0722 | P2_P3_Lumbar_Coeliac_Sphenopalatine_VS_P1_P4_Stellate | 0 | 1 | 1 | 0 | 1 | 1 | 0 | 1 |
| MIAT | 0.0721 | P2_P3_Lumbar_Coeliac_Sphenopalatine_VS_P1_P4_Stellate | 0 | 1 | 1 | 0 | 1 | 1 | 0 | 1 |
| GAS6 | 0.072 | P2_P3_P4_VS_P1_Lumbar_Coeliac_Stellate_Sphenopalatine | 0 | 1 | 1 | 1 | 0 | 0 | 0 | 0 |

| | | | | | | | | | | |
|---|---|---|---|---|---|---|---|---|---|---|
| CMPK2 | 0.072 | P2_P3_P4_VS_P1_Lumbar_Coeliac_Stellate_Sphenopalatine | 0 | 1 | 1 | 1 | 0 | 0 | 0 | 0 |
| APIP | 0.0719 | P1_P3_P4_Lumbar_VS_P2_Coeliac_Stellate_Sphenopalatine | 1 | 0 | 1 | 1 | 1 | 0 | 0 | 0 |
| NAT9 | 0.0719 | P1_P3_P4_Lumbar_VS_P2_Coeliac_Stellate_Sphenopalatine | 1 | 0 | 1 | 1 | 1 | 0 | 0 | 0 |
| HTRA2 | 0.0718 | P1_P3_P4_Lumbar_Sphenopalatine_VS_P2_Coeliac_Stellate | 1 | 0 | 1 | 1 | 1 | 0 | 0 | 1 |
| ATG9A | 0.0718 | P2_P3_P4_Lumbar_Sphenopalatine_VS_P1_Coeliac_Stellate | 0 | 1 | 1 | 1 | 1 | 0 | 0 | 1 |
| ATXN7L3 | 0.0718 | P2_P3_P4_Lumbar_Sphenopalatine_VS_P1_Coeliac_Stellate | 0 | 1 | 1 | 1 | 1 | 0 | 0 | 1 |
| AREL1 | 0.0716 | P3_P4_Lumbar_Coeliac_Stellate_VS_P1_P2_Sphenopalatine | 0 | 0 | 1 | 1 | 1 | 1 | 1 | 0 |
| SYT3 | 0.0716 | P2_P3_P4_Sphenopalatine_VS_P1_Lumbar_Coeliac_Stellate | 0 | 1 | 1 | 1 | 0 | 0 | 0 | 1 |
| CAMKMT | 0.0716 | P1_P2_P4_Lumbar_Coeliac_Stellate_Sphenopalatine_VS_P3 | 1 | 1 | 0 | 1 | 1 | 1 | 1 | 1 |
| NGFR | 0.0714 | P1_P2_P3_P4_Stellate_VS_Lumbar_Coeliac_Sphenopalatine | 1 | 1 | 1 | 1 | 0 | 0 | 1 | 0 |
| KCNB1 | 0.0714 | P2_P4_Lumbar_Sphenopalatine_VS_P1_P3_Coeliac_Stellate | 0 | 1 | 0 | 1 | 1 | 0 | 0 | 1 |
| SPRN | 0.0713 | P2_P3_P4_Sphenopalatine_VS_P1_Lumbar_Coeliac_Stellate | 0 | 1 | 1 | 1 | 0 | 0 | 0 | 1 |
| NDUFAF8 | 0.0713 | P1_P2_P3_P4_Lumbar_Stellate_Sphenopalatine_VS_Coeliac | 1 | 1 | 1 | 1 | 1 | 0 | 1 | 1 |
| PNRC2 | 0.0712 | P1_P3_VS_P2_P4_Lumbar_Coeliac_Stellate_Sphenopalatine | 1 | 0 | 1 | 0 | 0 | 0 | 0 | 0 |
| 5330434G04 | 0.0709 | P2_P4_Lumbar_Sphenopalatine_VS_P1_P3_Coeliac_Stellate | 0 | 1 | 0 | 1 | 1 | 0 | 0 | 1 |
| WRAP73 | 0.0705 | P1_P3_Lumbar_Coeliac_Sphenopalatine_VS_P2_P4_Stellate | 1 | 0 | 1 | 0 | 1 | 1 | 0 | 1 |
| API5 | 0.0704 | P2_P3_Lumbar_Coeliac_Sphenopalatine_VS_P1_P4_Stellate | 0 | 1 | 1 | 0 | 1 | 1 | 0 | 1 |
| SIN3A | 0.0702 | P2_P3_P4_Coeliac_VS_P1_Lumbar_Stellate_Sphenopalatine | 0 | 1 | 1 | 1 | 0 | 1 | 0 | 0 |
| KDM2B | 0.0702 | P1_Lumbar_Coeliac_Sphenopalatine_VS_P2_P3_P4_Stellate | 1 | 0 | 0 | 0 | 1 | 1 | 0 | 1 |
| FAM3C | 0.0701 | P3_Lumbar_Coeliac_Sphenopalatine_VS_P1_P2_P4_Stellate | 0 | 0 | 1 | 0 | 1 | 1 | 0 | 1 |
| CHMP1B | 0.0701 | P1_P2_P3_Lumbar_Coeliac_Stellate_Sphenopalatine_VS_P4 | 1 | 1 | 1 | 0 | 1 | 1 | 1 | 1 |
| HSD17B11 | 0.0699 | P2_P3_Lumbar_Coeliac_Sphenopalatine_VS_P1_P4_Stellate | 0 | 1 | 1 | 0 | 1 | 1 | 0 | 1 |
| DTX1 | 0.0697 | P2_P3_Lumbar_VS_P1_P4_Coeliac_Stellate_Sphenopalatine | 0 | 1 | 1 | 0 | 1 | 0 | 0 | 0 |
| TRP53 | 0.0696 | P1_P3_P4_Lumbar_Coeliac_Stellate_VS_P2_Sphenopalatine | 1 | 0 | 1 | 1 | 1 | 1 | 1 | 0 |
| ANKMY2 | 0.0696 | P1_P3_P4_Lumbar_Sphenopalatine_VS_P2_Coeliac_Stellate | 1 | 0 | 1 | 1 | 1 | 0 | 0 | 1 |
| FAM89A | 0.0696 | P1_P2_P4_Lumbar_Stellate_Sphenopalatine_VS_P3_Coeliac | 1 | 1 | 0 | 1 | 1 | 0 | 1 | 1 |
| CUL7 | 0.0695 | P1_P3_Lumbar_Coeliac_Sphenopalatine_VS_P2_P4_Stellate | 1 | 0 | 1 | 0 | 1 | 1 | 0 | 1 |
| MPST | 0.0692 | P2_P3_P4_Lumbar_Sphenopalatine_VS_P1_Coeliac_Stellate | 0 | 1 | 1 | 1 | 1 | 0 | 0 | 1 |
| NSMF | 0.0689 | P3_Sphenopalatine_VS_P1_P2_P4_Lumbar_Coeliac_Stellate | 0 | 0 | 1 | 0 | 0 | 0 | 0 | 1 |
| PIGQ | 0.0689 | P2_P3_P4_Sphenopalatine_VS_P1_Lumbar_Coeliac_Stellate | 0 | 1 | 1 | 1 | 0 | 0 | 0 | 1 |

| | | | | | | | | | | |
|---|---|---|---|---|---|---|---|---|---|---|
| CDKN2AIP | 0.0688 | P1_Coeliac_VS_P2_P3_P4_Lumbar_Stellate_Sphenopalatine | 1 | 0 | 0 | 0 | 0 | 1 | 0 | 0 |
| LRRC8D | 0.0686 | P4_Lumbar_Sphenopalatine_VS_P1_P2_P3_Coeliac_Stellate | 0 | 0 | 0 | 1 | 1 | 0 | 0 | 1 |
| MAGI1 | 0.0685 | P2_P4_Lumbar_Coeliac_Stellate_Sphenopalatine_VS_P1_P3 | 0 | 1 | 0 | 1 | 1 | 1 | 1 | 1 |
| UHMK1 | 0.0684 | P2_P3_P4_Sphenopalatine_VS_P1_Lumbar_Coeliac_Stellate | 0 | 1 | 1 | 1 | 0 | 0 | 0 | 1 |
| sept-11 | 0.0684 | P3_Lumbar_Coeliac_Sphenopalatine_VS_P1_P2_P4_Stellate | 0 | 0 | 1 | 0 | 1 | 1 | 0 | 1 |
| VEGFB | 0.0683 | P2_P3_P4_Lumbar_Sphenopalatine_VS_P1_Coeliac_Stellate | 0 | 1 | 1 | 1 | 1 | 0 | 0 | 1 |
| MARK1 | 0.068 | P2_P3_Lumbar_Coeliac_Sphenopalatine_VS_P1_P4_Stellate | 0 | 1 | 1 | 0 | 1 | 1 | 0 | 1 |
| ZFP120 | 0.0678 | P1_P2_P3_Lumbar_VS_P4_Coeliac_Stellate_Sphenopalatine | 1 | 1 | 1 | 0 | 1 | 0 | 0 | 0 |
| PTCD2 | 0.0678 | Lumbar_Sphenopalatine_VS_P1_P2_P3_P4_Coeliac_Stellate | 0 | 0 | 0 | 0 | 1 | 0 | 0 | 1 |
| PITPNM1 | 0.0677 | P4_Sphenopalatine_VS_P1_P2_P3_Lumbar_Coeliac_Stellate | 0 | 0 | 0 | 1 | 0 | 0 | 0 | 1 |
| CWH43 | 0.0672 | Lumbar_Coeliac_Stellate_VS_P1_P2_P3_P4_Sphenopalatine | 0 | 0 | 0 | 0 | 1 | 1 | 1 | 0 |
| MED9 | 0.0672 | P1_P3_P4_Lumbar_Sphenopalatine_VS_P2_Coeliac_Stellate | 1 | 0 | 1 | 1 | 1 | 0 | 0 | 1 |
| GUSB | 0.0672 | P1_P3_P4_Lumbar_Coeliac_Stellate_Sphenopalatine_VS_P2 | 1 | 0 | 1 | 1 | 1 | 1 | 1 | 1 |
| GNL3 | 0.0671 | P1_P3_Lumbar_Coeliac_Sphenopalatine_VS_P2_P4_Stellate | 1 | 0 | 1 | 0 | 1 | 1 | 0 | 1 |
| TSPAN18 | 0.067 | P1_P3_P4_Coeliac_VS_P2_Lumbar_Stellate_Sphenopalatine | 1 | 0 | 1 | 1 | 0 | 1 | 0 | 0 |
| PLEKHO1 | 0.0667 | P1_P3_Lumbar_Coeliac_Sphenopalatine_VS_P2_P4_Stellate | 1 | 0 | 1 | 0 | 1 | 1 | 0 | 1 |
| ZFP68 | 0.0664 | P1_P3_Lumbar_Coeliac_Sphenopalatine_VS_P2_P4_Stellate | 1 | 0 | 1 | 0 | 1 | 1 | 0 | 1 |
| CDK7 | 0.0659 | P1_Coeliac_VS_P2_P3_P4_Lumbar_Stellate_Sphenopalatine | 1 | 0 | 0 | 0 | 0 | 1 | 0 | 0 |
| UROD | 0.0658 | P1_P3_P4_Lumbar_Coeliac_Stellate_VS_P2_Sphenopalatine | 1 | 0 | 1 | 1 | 1 | 1 | 1 | 0 |
| EIF4EBP2 | 0.0657 | P2_Lumbar_Coeliac_Sphenopalatine_VS_P1_P3_P4_Stellate | 0 | 1 | 0 | 0 | 1 | 1 | 0 | 1 |
| ODC1 | 0.0655 | P2_P3_Lumbar_Coeliac_Sphenopalatine_VS_P1_P4_Stellate | 0 | 1 | 1 | 0 | 1 | 1 | 0 | 1 |
| ARHGEF3 | 0.0653 | Lumbar_Coeliac_Stellate_VS_P1_P2_P3_P4_Sphenopalatine | 0 | 0 | 0 | 0 | 1 | 1 | 1 | 0 |
| ALG3 | 0.0652 | P1_P3_P4_Lumbar_Coeliac_Stellate_VS_P2_Sphenopalatine | 1 | 0 | 1 | 1 | 1 | 1 | 1 | 0 |
| FAM217B | 0.0652 | P4_Sphenopalatine_VS_P1_P2_P3_Lumbar_Coeliac_Stellate | 0 | 0 | 0 | 1 | 0 | 0 | 0 | 1 |
| MAP3K6 | 0.065 | P1_P3_P4_Lumbar_Coeliac_Stellate_Sphenopalatine_VS_P2 | 1 | 0 | 1 | 1 | 1 | 1 | 1 | 1 |
| SELENOP | 0.0649 | P4_Sphenopalatine_VS_P1_P2_P3_Lumbar_Coeliac_Stellate | 0 | 0 | 0 | 1 | 0 | 0 | 0 | 1 |
| TRPC1 | 0.0647 | P1_P2_Lumbar_VS_P3_P4_Coeliac_Stellate_Sphenopalatine | 1 | 1 | 0 | 0 | 1 | 0 | 0 | 0 |
| NFYB | 0.0647 | P1_P4_Lumbar_Coeliac_Stellate_VS_P2_P3_Sphenopalatine | 1 | 0 | 0 | 1 | 1 | 1 | 1 | 0 |
| PCDH1 | 0.0645 | P2_Coeliac_VS_P1_P3_P4_Lumbar_Stellate_Sphenopalatine | 0 | 1 | 0 | 0 | 0 | 1 | 0 | 0 |
| PDE4B | 0.0645 | P1_P4_Lumbar_Sphenopalatine_VS_P2_P3_Coeliac_Stellate | 1 | 0 | 0 | 1 | 1 | 0 | 0 | 1 |

| | | | | | | | | | | |
|---|---|---|---|---|---|---|---|---|---|---|
| SERINC3 | 0.0644 | P2_P4_Lumbar_Sphenopalatine_VS_P1_P3_Coeliac_Stellate | 0 | 1 | 0 | 1 | 1 | 0 | 0 | 1 |
| AGPAT3 | 0.0643 | P2_P4_Lumbar_Sphenopalatine_VS_P1_P3_Coeliac_Stellate | 0 | 1 | 0 | 1 | 1 | 0 | 0 | 1 |
| SYT7 | 0.0641 | P3_Lumbar_Coeliac_Stellate_VS_P1_P2_P4_Sphenopalatine | 0 | 0 | 1 | 0 | 1 | 1 | 1 | 0 |
| PSEN1 | 0.0641 | P1_P2_P4_Lumbar_Coeliac_Stellate_Sphenopalatine_VS_P3 | 1 | 1 | 0 | 1 | 1 | 1 | 1 | 1 |
| RNPEP | 0.064 | P3_Lumbar_Coeliac_Sphenopalatine_VS_P1_P2_P4_Stellate | 0 | 0 | 1 | 0 | 1 | 1 | 0 | 1 |
| ATP6V1G2 | 0.0639 | P1_P2_P3_P4_Lumbar_Stellate_Sphenopalatine_VS_Coeliac | 1 | 1 | 1 | 1 | 1 | 0 | 1 | 1 |
| AIMP2 | 0.0639 | P1_P3_P4_Lumbar_Coeliac_Stellate_Sphenopalatine_VS_P2 | 1 | 0 | 1 | 1 | 1 | 1 | 1 | 1 |
| BCR | 0.0638 | P2_P4_Lumbar_Sphenopalatine_VS_P1_P3_Coeliac_Stellate | 0 | 1 | 0 | 1 | 1 | 0 | 0 | 1 |
| TMEM59L | 0.0638 | P3_P4_Lumbar_Sphenopalatine_VS_P1_P2_Coeliac_Stellate | 0 | 0 | 1 | 1 | 1 | 0 | 0 | 1 |
| MYCBP | 0.0634 | P1_P3_P4_Lumbar_Sphenopalatine_VS_P2_Coeliac_Stellate | 1 | 0 | 1 | 1 | 1 | 0 | 0 | 1 |
| MECP2 | 0.063 | P2_P3_Lumbar_Coeliac_Sphenopalatine_VS_P1_P4_Stellate | 0 | 1 | 1 | 0 | 1 | 1 | 0 | 1 |
| ASF1A | 0.0628 | P1_P3_P4_Coeliac_VS_P2_Lumbar_Stellate_Sphenopalatine | 1 | 0 | 1 | 1 | 0 | 1 | 0 | 0 |
| TOLLIP | 0.0626 | P2_P3_P4_Lumbar_Sphenopalatine_VS_P1_Coeliac_Stellate | 0 | 1 | 1 | 1 | 1 | 0 | 0 | 1 |
| SLC25A22 | 0.0621 | P2_P3_P4_Sphenopalatine_VS_P1_Lumbar_Coeliac_Stellate | 0 | 1 | 1 | 1 | 0 | 0 | 0 | 1 |
| PNMA2 | 0.0619 | P2_P3_P4_Lumbar_Sphenopalatine_VS_P1_Coeliac_Stellate | 0 | 1 | 1 | 1 | 1 | 0 | 0 | 1 |
| HTT | 0.0618 | P2_P3_P4_Lumbar_Sphenopalatine_VS_P1_Coeliac_Stellate | 0 | 1 | 1 | 1 | 1 | 0 | 0 | 1 |
| NXF1 | 0.0615 | P1_Lumbar_Coeliac_Sphenopalatine_VS_P2_P3_P4_Stellate | 1 | 0 | 0 | 0 | 1 | 1 | 0 | 1 |
| LYST | 0.0614 | P1_P3_Lumbar_Coeliac_Sphenopalatine_VS_P2_P4_Stellate | 1 | 0 | 1 | 0 | 1 | 1 | 0 | 1 |
| NAV3 | 0.0613 | P1_Lumbar_Coeliac_Sphenopalatine_VS_P2_P3_P4_Stellate | 1 | 0 | 0 | 0 | 1 | 1 | 0 | 1 |
| DHX32 | 0.0612 | P1_Lumbar_Coeliac_Sphenopalatine_VS_P2_P3_P4_Stellate | 1 | 0 | 0 | 0 | 1 | 1 | 0 | 1 |
| ESYT2 | 0.0608 | P2_P3_P4_Lumbar_Sphenopalatine_VS_P1_Coeliac_Stellate | 0 | 1 | 1 | 1 | 1 | 0 | 0 | 1 |
| ZKSCAN1 | 0.0608 | P2_P3_Lumbar_Coeliac_Sphenopalatine_VS_P1_P4_Stellate | 0 | 1 | 1 | 0 | 1 | 1 | 0 | 1 |
| DCP2 | 0.0607 | P1_P2_Lumbar_VS_P3_P4_Coeliac_Stellate_Sphenopalatine | 1 | 1 | 0 | 0 | 1 | 0 | 0 | 0 |
| TXLNA | 0.0607 | P2_P3_Lumbar_Coeliac_Sphenopalatine_VS_P1_P4_Stellate | 0 | 1 | 1 | 0 | 1 | 1 | 0 | 1 |
| PKNOX1 | 0.0606 | P2_P3_Lumbar_Coeliac_Sphenopalatine_VS_P1_P4_Stellate | 0 | 1 | 1 | 0 | 1 | 1 | 0 | 1 |
| SRM | 0.0604 | P1_P3_P4_Lumbar_Coeliac_Stellate_Sphenopalatine_VS_P2 | 1 | 0 | 1 | 1 | 1 | 1 | 1 | 1 |
| DUS1L | 0.0603 | P1_P3_Lumbar_VS_P2_P4_Coeliac_Stellate_Sphenopalatine | 1 | 0 | 1 | 0 | 1 | 0 | 0 | 0 |
| SNCB | 0.0602 | P1_P2_P3_P4_Lumbar_Stellate_Sphenopalatine_VS_Coeliac | 1 | 1 | 1 | 1 | 1 | 0 | 1 | 1 |
| TRO | 0.06 | P2_P3_P4_Lumbar_Sphenopalatine_VS_P1_Coeliac_Stellate | 0 | 1 | 1 | 1 | 1 | 0 | 0 | 1 |
| PIM1 | 0.0599 | P1_P2_Lumbar_VS_P3_P4_Coeliac_Stellate_Sphenopalatine | 1 | 1 | 0 | 0 | 1 | 0 | 0 | 0 |

| | | | | | | | | | | |
|---|---|---|---|---|---|---|---|---|---|---|
| A830082N09 | 0.0599 | P2_P3_Lumbar_Coeliac_Sphenopalatine_VS_P1_P4_Stellate | 0 | 1 | 1 | 0 | 1 | 1 | 0 | 1 |
| PLPP1 | 0.0593 | P1_P2_P3_P4_Lumbar_Stellate_Sphenopalatine_VS_Coeliac | 1 | 1 | 1 | 1 | 1 | 0 | 1 | 1 |
| SDF2L1 | 0.0591 | P1_P3_Lumbar_Coeliac_Stellate_Sphenopalatine_VS_P2_P4 | 1 | 0 | 1 | 0 | 1 | 1 | 1 | 1 |
| PPIP5K1 | 0.059 | P4_Lumbar_Sphenopalatine_VS_P1_P2_P3_Coeliac_Stellate | 0 | 0 | 0 | 1 | 1 | 0 | 0 | 1 |
| PIANP | 0.059 | P1_P3_P4_Lumbar_Sphenopalatine_VS_P2_Coeliac_Stellate | 1 | 0 | 1 | 1 | 1 | 0 | 0 | 1 |
| TMEM130 | 0.0588 | P2_P3_P4_Lumbar_Sphenopalatine_VS_P1_Coeliac_Stellate | 0 | 1 | 1 | 1 | 1 | 0 | 0 | 1 |
| MRPL46 | 0.0588 | P1_P3_P4_Lumbar_Coeliac_Stellate_Sphenopalatine_VS_P2 | 1 | 0 | 1 | 1 | 1 | 1 | 1 | 1 |
| MTMR3 | 0.0586 | P2_P3_P4_Lumbar_Sphenopalatine_VS_P1_Coeliac_Stellate | 0 | 1 | 1 | 1 | 1 | 0 | 0 | 1 |
| UPF2 | 0.0585 | P2_Coeliac_VS_P1_P3_P4_Lumbar_Stellate_Sphenopalatine | 0 | 1 | 0 | 0 | 0 | 1 | 0 | 0 |
| RNF2 | 0.0585 | P3_Lumbar_Coeliac_Sphenopalatine_VS_P1_P2_P4_Stellate | 0 | 0 | 1 | 0 | 1 | 1 | 0 | 1 |
| TIMM50 | 0.0585 | P1_P3_P4_Lumbar_Coeliac_Stellate_Sphenopalatine_VS_P2 | 1 | 0 | 1 | 1 | 1 | 1 | 1 | 1 |
| TMEM259 | 0.0583 | P2_P3_P4_Lumbar_Sphenopalatine_VS_P1_Coeliac_Stellate | 0 | 1 | 1 | 1 | 1 | 0 | 0 | 1 |
| DTNA | 0.0582 | P2_P3_P4_Lumbar_Sphenopalatine_VS_P1_Coeliac_Stellate | 0 | 1 | 1 | 1 | 1 | 0 | 0 | 1 |
| SLC4A1AP | 0.0581 | P1_Lumbar_Coeliac_Sphenopalatine_VS_P2_P3_P4_Stellate | 1 | 0 | 0 | 0 | 1 | 1 | 0 | 1 |
| UTP4 | 0.0579 | P1_Coeliac_VS_P2_P3_P4_Lumbar_Stellate_Sphenopalatine | 1 | 0 | 0 | 0 | 0 | 1 | 0 | 0 |
| ZFP768 | 0.0574 | P3_Lumbar_Coeliac_Sphenopalatine_VS_P1_P2_P4_Stellate | 0 | 0 | 1 | 0 | 1 | 1 | 0 | 1 |
| 4931428F04I | 0.0574 | P3_Lumbar_Coeliac_Sphenopalatine_VS_P1_P2_P4_Stellate | 0 | 0 | 1 | 0 | 1 | 1 | 0 | 1 |
| SEMA4D | 0.0573 | P2_P3_P4_Lumbar_Sphenopalatine_VS_P1_Coeliac_Stellate | 0 | 1 | 1 | 1 | 1 | 0 | 0 | 1 |
| CAT | 0.0572 | P2_P4_Lumbar_Sphenopalatine_VS_P1_P3_Coeliac_Stellate | 0 | 1 | 0 | 1 | 1 | 0 | 0 | 1 |
| KNOP1 | 0.0572 | P3_Lumbar_Coeliac_Sphenopalatine_VS_P1_P2_P4_Stellate | 0 | 0 | 1 | 0 | 1 | 1 | 0 | 1 |
| TTLL7 | 0.0571 | P2_P3_P4_Lumbar_Sphenopalatine_VS_P1_Coeliac_Stellate | 0 | 1 | 1 | 1 | 1 | 0 | 0 | 1 |
| TOR2A | 0.0569 | P1_P3_P4_Lumbar_Sphenopalatine_VS_P2_Coeliac_Stellate | 1 | 0 | 1 | 1 | 1 | 0 | 0 | 1 |
| TUT7 | 0.0569 | P2_Lumbar_Coeliac_Sphenopalatine_VS_P1_P3_P4_Stellate | 0 | 1 | 0 | 0 | 1 | 1 | 0 | 1 |
| USPL1 | 0.0568 | P1_P4_Lumbar_Sphenopalatine_VS_P2_P3_Coeliac_Stellate | 1 | 0 | 0 | 1 | 1 | 0 | 0 | 1 |
| ASPH | 0.0567 | P1_P3_Lumbar_VS_P2_P4_Coeliac_Stellate_Sphenopalatine | 1 | 0 | 1 | 0 | 1 | 0 | 0 | 0 |
| NRG1 | 0.0565 | P1_P4_Coeliac_Sphenopalatine_VS_P2_P3_Lumbar_Stellate | 1 | 0 | 0 | 1 | 0 | 1 | 0 | 1 |
| ADGRA1 | 0.0562 | P1_P2_P3_Coeliac_Sphenopalatine_VS_P4_Lumbar_Stellate | 1 | 1 | 1 | 0 | 0 | 1 | 0 | 1 |
| ZFP946 | 0.0562 | P1_Lumbar_Coeliac_Sphenopalatine_VS_P2_P3_P4_Stellate | 1 | 0 | 0 | 0 | 1 | 1 | 0 | 1 |
| PCGF1 | 0.056 | P1_P2_P3_Sphenopalatine_VS_P4_Lumbar_Coeliac_Stellate | 1 | 1 | 1 | 0 | 0 | 0 | 0 | 1 |
| KMT2A | 0.0557 | P2_P3_Lumbar_Coeliac_Sphenopalatine_VS_P1_P4_Stellate | 0 | 1 | 1 | 0 | 1 | 1 | 0 | 1 |

| | | | | | | | | | | |
|---|---|---|---|---|---|---|---|---|---|---|
| WASF3 | 0.0556 | P2_P3_P4_Sphenopalatine_VS_P1_Lumbar_Coeliac_Stellate | 0 | 1 | 1 | 1 | 0 | 0 | 0 | 1 |
| MRPL2 | 0.0555 | P1_P3_P4_Lumbar_Coeliac_Stellate_Sphenopalatine_VS_P2 | 1 | 0 | 1 | 1 | 1 | 1 | 1 | 1 |
| RRM1 | 0.0552 | P1_Coeliac_VS_P2_P3_P4_Lumbar_Stellate_Sphenopalatine | 1 | 0 | 0 | 0 | 0 | 1 | 0 | 0 |
| UBA6 | 0.055 | P2_P4_Lumbar_Sphenopalatine_VS_P1_P3_Coeliac_Stellate | 0 | 1 | 0 | 1 | 1 | 0 | 0 | 1 |
| BCL7A | 0.0549 | P2_P4_Lumbar_Sphenopalatine_VS_P1_P3_Coeliac_Stellate | 0 | 1 | 0 | 1 | 1 | 0 | 0 | 1 |
| ASB1 | 0.0548 | P2_P3_P4_Lumbar_Sphenopalatine_VS_P1_Coeliac_Stellate | 0 | 1 | 1 | 1 | 1 | 0 | 0 | 1 |
| ARMCX3 | 0.0546 | P2_P3_Lumbar_Coeliac_Sphenopalatine_VS_P1_P4_Stellate | 0 | 1 | 1 | 0 | 1 | 1 | 0 | 1 |
| RRBP1 | 0.0545 | P3_Lumbar_Coeliac_Sphenopalatine_VS_P1_P2_P4_Stellate | 0 | 0 | 1 | 0 | 1 | 1 | 0 | 1 |
| TMEM164 | 0.0544 | P1_P3_P4_Lumbar_Coeliac_Stellate_Sphenopalatine_VS_P2 | 1 | 0 | 1 | 1 | 1 | 1 | 1 | 1 |
| CPNE1 | 0.0542 | P2_P4_Lumbar_Sphenopalatine_VS_P1_P3_Coeliac_Stellate | 0 | 1 | 0 | 1 | 1 | 0 | 0 | 1 |
| PTPN23 | 0.0539 | P2_P4_Lumbar_Sphenopalatine_VS_P1_P3_Coeliac_Stellate | 0 | 1 | 0 | 1 | 1 | 0 | 0 | 1 |
| HSPH1 | 0.0538 | P2_P3_P4_Lumbar_Sphenopalatine_VS_P1_Coeliac_Stellate | 0 | 1 | 1 | 1 | 1 | 0 | 0 | 1 |
| ELP1 | 0.0538 | P2_Lumbar_Coeliac_Sphenopalatine_VS_P1_P3_P4_Stellate | 0 | 1 | 0 | 0 | 1 | 1 | 0 | 1 |
| BABAM2 | 0.0537 | P1_P3_P4_Lumbar_Sphenopalatine_VS_P2_Coeliac_Stellate | 1 | 0 | 1 | 1 | 1 | 0 | 0 | 1 |
| SNUPN | 0.0536 | P1_Coeliac_VS_P2_P3_P4_Lumbar_Stellate_Sphenopalatine | 1 | 0 | 0 | 0 | 0 | 1 | 0 | 0 |
| BRAF | 0.0535 | P2_P3_Lumbar_Coeliac_Sphenopalatine_VS_P1_P4_Stellate | 0 | 1 | 1 | 0 | 1 | 1 | 0 | 1 |
| RAB11FIP4 | 0.0535 | P2_P3_Lumbar_Coeliac_Sphenopalatine_VS_P1_P4_Stellate | 0 | 1 | 1 | 0 | 1 | 1 | 0 | 1 |
| KMT2C | 0.0534 | P2_Lumbar_Coeliac_Sphenopalatine_VS_P1_P3_P4_Stellate | 0 | 1 | 0 | 0 | 1 | 1 | 0 | 1 |
| FMR1 | 0.0534 | P2_Lumbar_Coeliac_Sphenopalatine_VS_P1_P3_P4_Stellate | 0 | 1 | 0 | 0 | 1 | 1 | 0 | 1 |
| GNAL | 0.0531 | Lumbar_Sphenopalatine_VS_P1_P2_P3_P4_Coeliac_Stellate | 0 | 0 | 0 | 0 | 1 | 0 | 0 | 1 |
| CCDC167 | 0.053 | P1_P3_P4_Lumbar_Sphenopalatine_VS_P2_Coeliac_Stellate | 1 | 0 | 1 | 1 | 1 | 0 | 0 | 1 |
| LGALSL | 0.0529 | P2_P3_Lumbar_Coeliac_Sphenopalatine_VS_P1_P4_Stellate | 0 | 1 | 1 | 0 | 1 | 1 | 0 | 1 |
| ZNHIT2 | 0.0528 | P3_Sphenopalatine_VS_P1_P2_P4_Lumbar_Coeliac_Stellate | 0 | 0 | 1 | 0 | 0 | 0 | 0 | 1 |
| AKAP17B | 0.0524 | P2_P3_Lumbar_Coeliac_Sphenopalatine_VS_P1_P4_Stellate | 0 | 1 | 1 | 0 | 1 | 1 | 0 | 1 |
| LIN54 | 0.0523 | P1_P3_P4_Coeliac_VS_P2_Lumbar_Stellate_Sphenopalatine | 1 | 0 | 1 | 1 | 0 | 1 | 0 | 0 |
| POLR2A | 0.0523 | P2_Lumbar_Coeliac_Sphenopalatine_VS_P1_P3_P4_Stellate | 0 | 1 | 0 | 0 | 1 | 1 | 0 | 1 |
| RNF26 | 0.052 | P1_P2_P3_P4_Lumbar_Stellate_Sphenopalatine_VS_Coeliac | 1 | 1 | 1 | 1 | 1 | 0 | 1 | 1 |
| PRXL2A | 0.0519 | P3_P4_Lumbar_Coeliac_Stellate_VS_P1_P2_Sphenopalatine | 0 | 0 | 1 | 1 | 1 | 1 | 1 | 0 |
| TMEM109 | 0.0519 | P1_P2_P3_Lumbar_Sphenopalatine_VS_P4_Coeliac_Stellate | 1 | 1 | 1 | 0 | 1 | 0 | 0 | 1 |
| DLG5 | 0.0518 | P2_P3_Lumbar_VS_P1_P4_Coeliac_Stellate_Sphenopalatine | 0 | 1 | 1 | 0 | 1 | 0 | 0 | 0 |

| | | | | | | | | | | |
|---|---|---|---|---|---|---|---|---|---|---|
| MZT2 | 0.0517 | P1_P3_P4_Lumbar_Coeliac_Stellate_Sphenopalatine_VS_P2 | 1 | 0 | 1 | 1 | 1 | 1 | 1 | 1 |
| BRSK1 | 0.0515 | P2_Lumbar_Coeliac_Sphenopalatine_VS_P1_P3_P4_Stellate | 0 | 1 | 0 | 0 | 1 | 1 | 0 | 1 |
| SOS1 | 0.0514 | P2_P3_Lumbar_Coeliac_Sphenopalatine_VS_P1_P4_Stellate | 0 | 1 | 1 | 0 | 1 | 1 | 0 | 1 |
| ENPP5 | 0.0513 | P2_P3_P4_Lumbar_Sphenopalatine_VS_P1_Coeliac_Stellate | 0 | 1 | 1 | 1 | 1 | 0 | 0 | 1 |
| EXOC8 | 0.0513 | P3_Lumbar_Coeliac_Sphenopalatine_VS_P1_P2_P4_Stellate | 0 | 0 | 1 | 0 | 1 | 1 | 0 | 1 |
| TTLL11 | 0.0512 | P1_P2_P3_Lumbar_VS_P4_Coeliac_Stellate_Sphenopalatine | 1 | 1 | 1 | 0 | 1 | 0 | 0 | 0 |
| ZBTB38 | 0.0512 | P2_Lumbar_Coeliac_Sphenopalatine_VS_P1_P3_P4_Stellate | 0 | 1 | 0 | 0 | 1 | 1 | 0 | 1 |
| CASP7 | 0.0511 | P3_Sphenopalatine_VS_P1_P2_P4_Lumbar_Coeliac_Stellate | 0 | 0 | 1 | 0 | 0 | 0 | 0 | 1 |
| ACOX1 | 0.051 | P2_P3_P4_Lumbar_Sphenopalatine_VS_P1_Coeliac_Stellate | 0 | 1 | 1 | 1 | 1 | 0 | 0 | 1 |
| AAK1 | 0.0507 | P2_P4_Lumbar_Sphenopalatine_VS_P1_P3_Coeliac_Stellate | 0 | 1 | 0 | 1 | 1 | 0 | 0 | 1 |
| ABCA2 | 0.0505 | P2_P3_P4_Coeliac_Sphenopalatine_VS_P1_Lumbar_Stellate | 0 | 1 | 1 | 1 | 0 | 1 | 0 | 1 |
| CD164 | 0.0504 | P2_P3_P4_Sphenopalatine_VS_P1_Lumbar_Coeliac_Stellate | 0 | 1 | 1 | 1 | 0 | 0 | 0 | 1 |
| SCAI | 0.05 | P2_P3_Lumbar_Coeliac_Sphenopalatine_VS_P1_P4_Stellate | 0 | 1 | 1 | 0 | 1 | 1 | 0 | 1 |
| RGS19 | 0.05 | P1_P2_P3_P4_Lumbar_Stellate_Sphenopalatine_VS_Coeliac | 1 | 1 | 1 | 1 | 1 | 0 | 1 | 1 |
| TMED1 | 0.0499 | P1_P3_P4_Lumbar_Sphenopalatine_VS_P2_Coeliac_Stellate | 1 | 0 | 1 | 1 | 1 | 0 | 0 | 1 |
| TTC5 | 0.0497 | P1_P3_P4_Lumbar_Coeliac_Stellate_VS_P2_Sphenopalatine | 1 | 0 | 1 | 1 | 1 | 1 | 1 | 0 |
| ARF6 | 0.0496 | P1_P3_P4_Coeliac_VS_P2_Lumbar_Stellate_Sphenopalatine | 1 | 0 | 1 | 1 | 0 | 1 | 0 | 0 |
| KBTBD2 | 0.0495 | P2_P4_Lumbar_Sphenopalatine_VS_P1_P3_Coeliac_Stellate | 0 | 1 | 0 | 1 | 1 | 0 | 0 | 1 |
| BBC3 | 0.0493 | P1_P3_Lumbar_Coeliac_Stellate_VS_P2_P4_Sphenopalatine | 1 | 0 | 1 | 0 | 1 | 1 | 1 | 0 |
| TCEAL1 | 0.0493 | P1_P3_P4_Lumbar_Sphenopalatine_VS_P2_Coeliac_Stellate | 1 | 0 | 1 | 1 | 1 | 0 | 0 | 1 |
| KRT222 | 0.0492 | P2_P4_Lumbar_Sphenopalatine_VS_P1_P3_Coeliac_Stellate | 0 | 1 | 0 | 1 | 1 | 0 | 0 | 1 |
| TMEM53 | 0.049 | P1_P3_P4_VS_P2_Lumbar_Coeliac_Stellate_Sphenopalatine | 1 | 0 | 1 | 1 | 0 | 0 | 0 | 0 |
| TMEM14A | 0.0489 | P1_P2_P3_P4_Lumbar_Stellate_Sphenopalatine_VS_Coeliac | 1 | 1 | 1 | 1 | 1 | 0 | 1 | 1 |
| CCPG1 | 0.0488 | P2_P3_P4_Lumbar_Sphenopalatine_VS_P1_Coeliac_Stellate | 0 | 1 | 1 | 1 | 1 | 0 | 0 | 1 |
| CLYBL | 0.0487 | P1_P3_P4_VS_P2_Lumbar_Coeliac_Stellate_Sphenopalatine | 1 | 0 | 1 | 1 | 0 | 0 | 0 | 0 |
| PLPP2 | 0.0487 | P1_P3_Lumbar_VS_P2_P4_Coeliac_Stellate_Sphenopalatine | 1 | 0 | 1 | 0 | 1 | 0 | 0 | 0 |
| PRUNE1 | 0.0486 | P2_P3_P4_Lumbar_Sphenopalatine_VS_P1_Coeliac_Stellate | 0 | 1 | 1 | 1 | 1 | 0 | 0 | 1 |
| CLOCK | 0.0484 | P2_P3_Lumbar_VS_P1_P4_Coeliac_Stellate_Sphenopalatine | 0 | 1 | 1 | 0 | 1 | 0 | 0 | 0 |
| TRRAP | 0.0484 | P2_Lumbar_Coeliac_Sphenopalatine_VS_P1_P3_P4_Stellate | 0 | 1 | 0 | 0 | 1 | 1 | 0 | 1 |
| CDK5R2 | 0.0483 | P2_P3_P4_Sphenopalatine_VS_P1_Lumbar_Coeliac_Stellate | 0 | 1 | 1 | 1 | 0 | 0 | 0 | 1 |

| | | | | | | | | | | |
|---|---|---|---|---|---|---|---|---|---|---|
| WIPI2 | 0.0483 | P2_P3_P4_Sphenopalatine_VS_P1_Lumbar_Coeliac_Stellate | 0 | 1 | 1 | 1 | 0 | 0 | 0 | 1 |
| COA6 | 0.0483 | P1_P3_P4_Lumbar_Coeliac_Stellate_Sphenopalatine_VS_P2 | 1 | 0 | 1 | 1 | 1 | 1 | 1 | 1 |
| FADS2 | 0.0482 | P1_P3_P4_Lumbar_Sphenopalatine_VS_P2_Coeliac_Stellate | 1 | 0 | 1 | 1 | 1 | 0 | 0 | 1 |
| KRR1 | 0.0482 | P2_P3_Lumbar_Coeliac_Sphenopalatine_VS_P1_P4_Stellate | 0 | 1 | 1 | 0 | 1 | 1 | 0 | 1 |
| MRPS28 | 0.0482 | P1_P3_P4_Lumbar_Coeliac_Stellate_Sphenopalatine_VS_P2 | 1 | 0 | 1 | 1 | 1 | 1 | 1 | 1 |
| SNPH | 0.0481 | P2_P4_Lumbar_Sphenopalatine_VS_P1_P3_Coeliac_Stellate | 0 | 1 | 0 | 1 | 1 | 0 | 0 | 1 |
| SMIM17 | 0.048 | P2_Sphenopalatine_VS_P1_P3_P4_Lumbar_Coeliac_Stellate | 0 | 1 | 0 | 0 | 0 | 0 | 0 | 1 |
| NCOR2 | 0.0478 | P2_P3_P4_Lumbar_Sphenopalatine_VS_P1_Coeliac_Stellate | 0 | 1 | 1 | 1 | 1 | 0 | 0 | 1 |
| TEF | 0.0477 | P2_P3_Lumbar_Coeliac_Sphenopalatine_VS_P1_P4_Stellate | 0 | 1 | 1 | 0 | 1 | 1 | 0 | 1 |
| MPLKIP | 0.0475 | P1_P3_P4_Lumbar_Coeliac_Stellate_VS_P2_Sphenopalatine | 1 | 0 | 1 | 1 | 1 | 1 | 1 | 0 |
| TSPYL3 | 0.0474 | P2_P4_Lumbar_Sphenopalatine_VS_P1_P3_Coeliac_Stellate | 0 | 1 | 0 | 1 | 1 | 0 | 0 | 1 |
| ILRUN | 0.0473 | P2_P3_P4_Lumbar_Sphenopalatine_VS_P1_Coeliac_Stellate | 0 | 1 | 1 | 1 | 1 | 0 | 0 | 1 |
| SGSM2 | 0.0472 | P2_P4_Lumbar_Sphenopalatine_VS_P1_P3_Coeliac_Stellate | 0 | 1 | 0 | 1 | 1 | 0 | 0 | 1 |
| 9530034E10I | 0.0472 | P1_Coeliac_Sphenopalatine_VS_P2_P3_P4_Lumbar_Stellate | 1 | 0 | 0 | 0 | 0 | 1 | 0 | 1 |
| ASPHD1 | 0.0471 | P1_P3_P4_Lumbar_Sphenopalatine_VS_P2_Coeliac_Stellate | 1 | 0 | 1 | 1 | 1 | 0 | 0 | 1 |
| SLC25A12 | 0.0471 | P2_P3_P4_Lumbar_Sphenopalatine_VS_P1_Coeliac_Stellate | 0 | 1 | 1 | 1 | 1 | 0 | 0 | 1 |
| GM8797 | 0.0471 | P1_P4_Lumbar_Coeliac_Stellate_Sphenopalatine_VS_P2_P3 | 1 | 0 | 0 | 1 | 1 | 1 | 1 | 1 |
| RPL10-PS3 | 0.047 | P3_Sphenopalatine_VS_P1_P2_P4_Lumbar_Coeliac_Stellate | 0 | 0 | 1 | 0 | 0 | 0 | 0 | 1 |
| MAST1 | 0.047 | P2_P4_Lumbar_Sphenopalatine_VS_P1_P3_Coeliac_Stellate | 0 | 1 | 0 | 1 | 1 | 0 | 0 | 1 |
| EDEM3 | 0.047 | P2_P3_Lumbar_Coeliac_Sphenopalatine_VS_P1_P4_Stellate | 0 | 1 | 1 | 0 | 1 | 1 | 0 | 1 |
| MTMR9 | 0.0469 | P2_P3_P4_Sphenopalatine_VS_P1_Lumbar_Coeliac_Stellate | 0 | 1 | 1 | 1 | 0 | 0 | 0 | 1 |
| 2810002D19I | 0.0468 | P1_P3_Lumbar_Coeliac_Stellate_VS_P2_P4_Sphenopalatine | 1 | 0 | 1 | 0 | 1 | 1 | 1 | 0 |
| TBCC | 0.0466 | P1_P3_P4_Lumbar_Coeliac_Stellate_VS_P2_Sphenopalatine | 1 | 0 | 1 | 1 | 1 | 1 | 1 | 0 |
| DCAF17 | 0.0466 | P3_Lumbar_Coeliac_Sphenopalatine_VS_P1_P2_P4_Stellate | 0 | 0 | 1 | 0 | 1 | 1 | 0 | 1 |
| OS9 | 0.0466 | P3_Lumbar_Coeliac_Sphenopalatine_VS_P1_P2_P4_Stellate | 0 | 0 | 1 | 0 | 1 | 1 | 0 | 1 |
| TET3 | 0.0466 | P2_P3_Lumbar_Coeliac_Sphenopalatine_VS_P1_P4_Stellate | 0 | 1 | 1 | 0 | 1 | 1 | 0 | 1 |
| 2300009A05I | 0.0465 | P3_P4_Sphenopalatine_VS_P1_P2_Lumbar_Coeliac_Stellate | 0 | 0 | 1 | 1 | 0 | 0 | 0 | 1 |
| NCBP3 | 0.0465 | P2_Lumbar_Coeliac_Sphenopalatine_VS_P1_P3_P4_Stellate | 0 | 1 | 0 | 0 | 1 | 1 | 0 | 1 |
| RNF185 | 0.0464 | P1_P2_P4_Lumbar_Coeliac_Stellate_Sphenopalatine_VS_P3 | 1 | 1 | 0 | 1 | 1 | 1 | 1 | 1 |
| DTX4 | 0.0461 | P2_P4_Lumbar_Sphenopalatine_VS_P1_P3_Coeliac_Stellate | 0 | 1 | 0 | 1 | 1 | 0 | 0 | 1 |

| | | | | | | | | | | |
|---|---|---|---|---|---|---|---|---|---|---|
| PLPPR3 | 0.0461 | P1_P2_P3_Lumbar_Coeliac_Stellate_Sphenopalatine_VS_P4 | 1 | 1 | 1 | 0 | 1 | 1 | 1 | 1 |
| DAP | 0.046 | P2_P3_VS_P1_P4_Lumbar_Coeliac_Stellate_Sphenopalatine | 0 | 1 | 1 | 0 | 0 | 0 | 0 | 0 |
| TCF12 | 0.046 | P2_P3_Lumbar_Coeliac_Sphenopalatine_VS_P1_P4_Stellate | 0 | 1 | 1 | 0 | 1 | 1 | 0 | 1 |
| ACTR3B | 0.046 | P1_P2_P3_P4_Lumbar_Stellate_Sphenopalatine_VS_Coeliac | 1 | 1 | 1 | 1 | 1 | 0 | 1 | 1 |
| STX8 | 0.046 | P1_P3_P4_Lumbar_Coeliac_Stellate_Sphenopalatine_VS_P2 | 1 | 0 | 1 | 1 | 1 | 1 | 1 | 1 |
| RIPOR1 | 0.0459 | P2_P4_Lumbar_Sphenopalatine_VS_P1_P3_Coeliac_Stellate | 0 | 1 | 0 | 1 | 1 | 0 | 0 | 1 |
| EP400 | 0.0459 | P2_P3_Lumbar_Coeliac_Sphenopalatine_VS_P1_P4_Stellate | 0 | 1 | 1 | 0 | 1 | 1 | 0 | 1 |
| BTBD2 | 0.0458 | P1_P2_P3_Lumbar_VS_P4_Coeliac_Stellate_Sphenopalatine | 1 | 1 | 1 | 0 | 1 | 0 | 0 | 0 |
| TMEM245 | 0.0454 | P2_P4_Lumbar_Sphenopalatine_VS_P1_P3_Coeliac_Stellate | 0 | 1 | 0 | 1 | 1 | 0 | 0 | 1 |
| FAM155A | 0.0454 | P1_P2_P4_Lumbar_Coeliac_Stellate_Sphenopalatine_VS_P3 | 1 | 1 | 0 | 1 | 1 | 1 | 1 | 1 |
| UFSP2 | 0.0453 | P3_Lumbar_Coeliac_Sphenopalatine_VS_P1_P2_P4_Stellate | 0 | 0 | 1 | 0 | 1 | 1 | 0 | 1 |
| SPATS2 | 0.0451 | P2_P3_Lumbar_VS_P1_P4_Coeliac_Stellate_Sphenopalatine | 0 | 1 | 1 | 0 | 1 | 0 | 0 | 0 |
| MGAT4B | 0.0451 | P1_P3_P4_Lumbar_Sphenopalatine_VS_P2_Coeliac_Stellate | 1 | 0 | 1 | 1 | 1 | 0 | 0 | 1 |
| ATP9B | 0.045 | P4_Lumbar_Sphenopalatine_VS_P1_P2_P3_Coeliac_Stellate | 0 | 0 | 0 | 1 | 1 | 0 | 0 | 1 |
| LRRC14 | 0.0449 | P1_P3_P4_Lumbar_Sphenopalatine_VS_P2_Coeliac_Stellate | 1 | 0 | 1 | 1 | 1 | 0 | 0 | 1 |
| COG5 | 0.0449 | P2_P3_Lumbar_Coeliac_Sphenopalatine_VS_P1_P4_Stellate | 0 | 1 | 1 | 0 | 1 | 1 | 0 | 1 |
| EBPL | 0.0449 | P1_P3_P4_Lumbar_Coeliac_Stellate_Sphenopalatine_VS_P2 | 1 | 0 | 1 | 1 | 1 | 1 | 1 | 1 |
| ACP2 | 0.0447 | P2_P3_P4_Lumbar_Sphenopalatine_VS_P1_Coeliac_Stellate | 0 | 1 | 1 | 1 | 1 | 0 | 0 | 1 |
| ATG10 | 0.0445 | P1_P3_P4_Lumbar_Coeliac_Stellate_Sphenopalatine_VS_P2 | 1 | 0 | 1 | 1 | 1 | 1 | 1 | 1 |
| CINP | 0.0443 | Lumbar_Sphenopalatine_VS_P1_P2_P3_P4_Coeliac_Stellate | 0 | 0 | 0 | 0 | 1 | 0 | 0 | 1 |
| GM10762 | 0.0441 | P1_P2_P3_Lumbar_VS_P4_Coeliac_Stellate_Sphenopalatine | 1 | 1 | 1 | 0 | 1 | 0 | 0 | 0 |
| PLEKHM3 | 0.0441 | P2_P4_Lumbar_Sphenopalatine_VS_P1_P3_Coeliac_Stellate | 0 | 1 | 0 | 1 | 1 | 0 | 0 | 1 |
| ERC2 | 0.0441 | P2_P4_Lumbar_Sphenopalatine_VS_P1_P3_Coeliac_Stellate | 0 | 1 | 0 | 1 | 1 | 0 | 0 | 1 |
| MTSS2 | 0.0438 | P2_P3_P4_Lumbar_Sphenopalatine_VS_P1_Coeliac_Stellate | 0 | 1 | 1 | 1 | 1 | 0 | 0 | 1 |
| MARK2 | 0.0438 | P2_P3_P4_Lumbar_Sphenopalatine_VS_P1_Coeliac_Stellate | 0 | 1 | 1 | 1 | 1 | 0 | 0 | 1 |
| METRNL | 0.0437 | P2_P3_VS_P1_P4_Lumbar_Coeliac_Stellate_Sphenopalatine | 0 | 1 | 1 | 0 | 0 | 0 | 0 | 0 |
| PRADC1 | 0.0435 | P2_P3_VS_P1_P4_Lumbar_Coeliac_Stellate_Sphenopalatine | 0 | 1 | 1 | 0 | 0 | 0 | 0 | 0 |
| NFASC | 0.0434 | P2_P3_P4_Lumbar_Sphenopalatine_VS_P1_Coeliac_Stellate | 0 | 1 | 1 | 1 | 1 | 0 | 0 | 1 |
| GM48226 | 0.0434 | P1_P3_P4_Lumbar_Coeliac_Stellate_Sphenopalatine_VS_P2 | 1 | 0 | 1 | 1 | 1 | 1 | 1 | 1 |
| TUSC2 | 0.0432 | P1_P2_P3_P4_Lumbar_Stellate_Sphenopalatine_VS_Coeliac | 1 | 1 | 1 | 1 | 1 | 0 | 1 | 1 |

| | | | | | | | | | | |
|---|---|---|---|---|---|---|---|---|---|---|
| TANGO2 | 0.043 | P1_P2_P3_P4_Lumbar_Stellate_Sphenopalatine_VS_Coeliac | 1 | 1 | 1 | 1 | 1 | 0 | 1 | 1 |
| MYO18A | 0.0428 | P2_P3_Lumbar_Coeliac_Sphenopalatine_VS_P1_P4_Stellate | 0 | 1 | 1 | 0 | 1 | 1 | 0 | 1 |
| ARIH1 | 0.0426 | P2_P3_Lumbar_Coeliac_Sphenopalatine_VS_P1_P4_Stellate | 0 | 1 | 1 | 0 | 1 | 1 | 0 | 1 |
| MAN1A2 | 0.0424 | P4_Lumbar_Sphenopalatine_VS_P1_P2_P3_Coeliac_Stellate | 0 | 0 | 0 | 1 | 1 | 0 | 0 | 1 |
| PLPBP | 0.0421 | P1_P3_P4_Lumbar_Sphenopalatine_VS_P2_Coeliac_Stellate | 1 | 0 | 1 | 1 | 1 | 0 | 0 | 1 |
| AP1G1 | 0.0421 | P2_P3_P4_Lumbar_Sphenopalatine_VS_P1_Coeliac_Stellate | 0 | 1 | 1 | 1 | 1 | 0 | 0 | 1 |
| SLC25A23 | 0.0421 | P2_P3_P4_Lumbar_Sphenopalatine_VS_P1_Coeliac_Stellate | 0 | 1 | 1 | 1 | 1 | 0 | 0 | 1 |
| TSC2 | 0.0421 | P2_P3_Lumbar_Coeliac_Sphenopalatine_VS_P1_P4_Stellate | 0 | 1 | 1 | 0 | 1 | 1 | 0 | 1 |
| NOP16 | 0.0419 | P1_P3_P4_Lumbar_Coeliac_Stellate_Sphenopalatine_VS_P2 | 1 | 0 | 1 | 1 | 1 | 1 | 1 | 1 |
| NOP58 | 0.0418 | P1_P3_P4_Lumbar_Coeliac_Stellate_VS_P2_Sphenopalatine | 1 | 0 | 1 | 1 | 1 | 1 | 1 | 0 |
| SCO2 | 0.0417 | P1_P2_P4_Lumbar_Coeliac_Stellate_Sphenopalatine_VS_P3 | 1 | 1 | 0 | 1 | 1 | 1 | 1 | 1 |
| RIDA | 0.0417 | P1_P3_P4_Lumbar_Coeliac_Stellate_Sphenopalatine_VS_P2 | 1 | 0 | 1 | 1 | 1 | 1 | 1 | 1 |
| RNF145 | 0.0415 | P2_P3_P4_Lumbar_Coeliac_Stellate_VS_P1_Sphenopalatine | 0 | 1 | 1 | 1 | 1 | 1 | 1 | 0 |
| SAP130 | 0.0415 | P2_P4_Lumbar_Sphenopalatine_VS_P1_P3_Coeliac_Stellate | 0 | 1 | 0 | 1 | 1 | 0 | 0 | 1 |
| D730003I15F | 0.0414 | P1_P3_P4_Lumbar_Coeliac_Stellate_Sphenopalatine_VS_P2 | 1 | 0 | 1 | 1 | 1 | 1 | 1 | 1 |
| CEP44 | 0.0413 | P1_P3_P4_Lumbar_Coeliac_Stellate_VS_P2_Sphenopalatine | 1 | 0 | 1 | 1 | 1 | 1 | 1 | 0 |
| SCFD1 | 0.0413 | P1_P3_P4_Lumbar_Coeliac_Stellate_VS_P2_Sphenopalatine | 1 | 0 | 1 | 1 | 1 | 1 | 1 | 0 |
| SMC3 | 0.0412 | P4_Lumbar_Coeliac_Stellate_VS_P1_P2_P3_Sphenopalatine | 0 | 0 | 0 | 1 | 1 | 1 | 1 | 0 |
| FAHD1 | 0.0411 | P1_P3_P4_Lumbar_Coeliac_Stellate_VS_P2_Sphenopalatine | 1 | 0 | 1 | 1 | 1 | 1 | 1 | 0 |
| KDSR | 0.041 | P2_Sphenopalatine_VS_P1_P3_P4_Lumbar_Coeliac_Stellate | 0 | 1 | 0 | 0 | 0 | 0 | 0 | 1 |
| SETD1B | 0.041 | P2_P3_P4_Lumbar_Sphenopalatine_VS_P1_Coeliac_Stellate | 0 | 1 | 1 | 1 | 1 | 0 | 0 | 1 |
| FAM174B | 0.0408 | P2_P3_P4_Sphenopalatine_VS_P1_Lumbar_Coeliac_Stellate | 0 | 1 | 1 | 1 | 0 | 0 | 0 | 1 |
| FAM120A | 0.0408 | P2_P3_P4_Lumbar_Sphenopalatine_VS_P1_Coeliac_Stellate | 0 | 1 | 1 | 1 | 1 | 0 | 0 | 1 |
| RUBCN | 0.0408 | P2_P3_P4_Lumbar_Sphenopalatine_VS_P1_Coeliac_Stellate | 0 | 1 | 1 | 1 | 1 | 0 | 0 | 1 |
| CDK10 | 0.0407 | P1_P4_Lumbar_Sphenopalatine_VS_P2_P3_Coeliac_Stellate | 1 | 0 | 0 | 1 | 1 | 0 | 0 | 1 |
| CHPF | 0.0407 | P2_P3_P4_Lumbar_Sphenopalatine_VS_P1_Coeliac_Stellate | 0 | 1 | 1 | 1 | 1 | 0 | 0 | 1 |
| RAI1 | 0.0407 | P2_P3_Lumbar_Coeliac_Sphenopalatine_VS_P1_P4_Stellate | 0 | 1 | 1 | 0 | 1 | 1 | 0 | 1 |
| NOMO1 | 0.0406 | P2_P3_P4_Lumbar_Sphenopalatine_VS_P1_Coeliac_Stellate | 0 | 1 | 1 | 1 | 1 | 0 | 0 | 1 |
| ADAR | 0.0402 | P2_P3_Lumbar_Coeliac_Sphenopalatine_VS_P1_P4_Stellate | 0 | 1 | 1 | 0 | 1 | 1 | 0 | 1 |
| RAP2A | 0.0401 | P2_Lumbar_Coeliac_Sphenopalatine_VS_P1_P3_P4_Stellate | 0 | 1 | 0 | 0 | 1 | 1 | 0 | 1 |

| | | | | | | | | | | |
|---|---|---|---|---|---|---|---|---|---|---|
| SPRYD3 | 0.0399 | P2_P3_P4_Lumbar_Sphenopalatine_VS_P1_Coeliac_Stellate | 0 | 1 | 1 | 1 | 1 | 0 | 0 | 1 |
| CDK11B | 0.0399 | P2_Lumbar_Coeliac_Sphenopalatine_VS_P1_P3_P4_Stellate | 0 | 1 | 0 | 0 | 1 | 1 | 0 | 1 |
| ZFP322A | 0.0397 | P2_Lumbar_Coeliac_Sphenopalatine_VS_P1_P3_P4_Stellate | 0 | 1 | 0 | 0 | 1 | 1 | 0 | 1 |
| ZFP612 | 0.0396 | P2_P3_P4_Lumbar_Sphenopalatine_VS_P1_Coeliac_Stellate | 0 | 1 | 1 | 1 | 1 | 0 | 0 | 1 |
| RALGAPB | 0.0395 | P2_P3_P4_Lumbar_Sphenopalatine_VS_P1_Coeliac_Stellate | 0 | 1 | 1 | 1 | 1 | 0 | 0 | 1 |
| ATN1 | 0.0395 | P2_Lumbar_Coeliac_Sphenopalatine_VS_P1_P3_P4_Stellate | 0 | 1 | 0 | 0 | 1 | 1 | 0 | 1 |
| OTUD4 | 0.0392 | P1_P2_Lumbar_VS_P3_P4_Coeliac_Stellate_Sphenopalatine | 1 | 1 | 0 | 0 | 1 | 0 | 0 | 0 |
| MPV17L | 0.0392 | P1_P2_Lumbar_VS_P3_P4_Coeliac_Stellate_Sphenopalatine | 1 | 1 | 0 | 0 | 1 | 0 | 0 | 0 |
| GPR22 | 0.0392 | P2_P3_Lumbar_Coeliac_Sphenopalatine_VS_P1_P4_Stellate | 0 | 1 | 1 | 0 | 1 | 1 | 0 | 1 |
| FAM131A | 0.0391 | P2_P3_P4_Lumbar_Sphenopalatine_VS_P1_Coeliac_Stellate | 0 | 1 | 1 | 1 | 1 | 0 | 0 | 1 |
| GRK6 | 0.0388 | P2_P3_P4_Lumbar_Sphenopalatine_VS_P1_Coeliac_Stellate | 0 | 1 | 1 | 1 | 1 | 0 | 0 | 1 |
| RPS27RT | 0.0387 | P2_P3_P4_Lumbar_Sphenopalatine_VS_P1_Coeliac_Stellate | 0 | 1 | 1 | 1 | 1 | 0 | 0 | 1 |
| GRIN1 | 0.0381 | P2_P3_P4_Lumbar_Sphenopalatine_VS_P1_Coeliac_Stellate | 0 | 1 | 1 | 1 | 1 | 0 | 0 | 1 |
| TSPO | 0.0379 | P1_P3_Lumbar_VS_P2_P4_Coeliac_Stellate_Sphenopalatine | 1 | 0 | 1 | 0 | 1 | 0 | 0 | 0 |
| ELOVL5 | 0.0379 | P2_P3_P4_Sphenopalatine_VS_P1_Lumbar_Coeliac_Stellate | 0 | 1 | 1 | 1 | 0 | 0 | 0 | 1 |
| MTMR7 | 0.0378 | P4_Lumbar_Sphenopalatine_VS_P1_P2_P3_Coeliac_Stellate | 0 | 0 | 0 | 1 | 1 | 0 | 0 | 1 |
| CWC25 | 0.0377 | P2_Lumbar_Coeliac_Sphenopalatine_VS_P1_P3_P4_Stellate | 0 | 1 | 0 | 0 | 1 | 1 | 0 | 1 |
| WSCD2 | 0.0376 | Lumbar_Coeliac_Stellate_VS_P1_P2_P3_P4_Sphenopalatine | 0 | 0 | 0 | 0 | 1 | 1 | 1 | 0 |
| SLC25A36 | 0.0375 | P2_P3_P4_Lumbar_Sphenopalatine_VS_P1_Coeliac_Stellate | 0 | 1 | 1 | 1 | 1 | 0 | 0 | 1 |
| SNCA | 0.0375 | P2_P3_P4_Lumbar_Coeliac_Stellate_Sphenopalatine_VS_P1 | 0 | 1 | 1 | 1 | 1 | 1 | 1 | 1 |
| ENDOG | 0.0374 | P1_P3_P4_Lumbar_Sphenopalatine_VS_P2_Coeliac_Stellate | 1 | 0 | 1 | 1 | 1 | 0 | 0 | 1 |
| SIDT2 | 0.0374 | P2_P3_P4_Lumbar_Sphenopalatine_VS_P1_Coeliac_Stellate | 0 | 1 | 1 | 1 | 1 | 0 | 0 | 1 |
| TTBK2 | 0.037 | P2_P3_P4_Sphenopalatine_VS_P1_Lumbar_Coeliac_Stellate | 0 | 1 | 1 | 1 | 0 | 0 | 0 | 1 |
| TMED3 | 0.0367 | P2_P3_P4_Sphenopalatine_VS_P1_Lumbar_Coeliac_Stellate | 0 | 1 | 1 | 1 | 0 | 0 | 0 | 1 |
| RANGRF | 0.0366 | P3_Lumbar_Coeliac_Stellate_VS_P1_P2_P4_Sphenopalatine | 0 | 0 | 1 | 0 | 1 | 1 | 1 | 0 |
| NTMT1 | 0.0363 | P1_P3_P4_Lumbar_Coeliac_Stellate_Sphenopalatine_VS_P2 | 1 | 0 | 1 | 1 | 1 | 1 | 1 | 1 |
| NDUFAF1 | 0.0362 | P1_P4_Lumbar_Coeliac_Stellate_VS_P2_P3_Sphenopalatine | 1 | 0 | 0 | 1 | 1 | 1 | 1 | 0 |
| DGKZ | 0.0362 | P1_P2_P3_Coeliac_Sphenopalatine_VS_P4_Lumbar_Stellate | 1 | 1 | 1 | 0 | 0 | 1 | 0 | 1 |
| FXR2 | 0.0359 | P1_P3_P4_Lumbar_Sphenopalatine_VS_P2_Coeliac_Stellate | 1 | 0 | 1 | 1 | 1 | 0 | 0 | 1 |
| MT3 | 0.0359 | P1_P2_P3_P4_Lumbar_Stellate_Sphenopalatine_VS_Coeliac | 1 | 1 | 1 | 1 | 1 | 0 | 1 | 1 |

| | | | | | | | | | | |
|---|---|---|---|---|---|---|---|---|---|---|
| MAP3K10 | 0.0358 | P2_P3_P4_Lumbar_Sphenopalatine_VS_P1_Coeliac_Stellate | 0 | 1 | 1 | 1 | 1 | 0 | 0 | 1 |
| NOL3 | 0.0356 | P2_P3_P4_Sphenopalatine_VS_P1_Lumbar_Coeliac_Stellate | 0 | 1 | 1 | 1 | 0 | 0 | 0 | 1 |
| RNF170 | 0.0356 | P2_P3_Lumbar_Coeliac_Sphenopalatine_VS_P1_P4_Stellate | 0 | 1 | 1 | 0 | 1 | 1 | 0 | 1 |
| PIP4P2 | 0.0356 | P1_P2_P3_P4_Lumbar_Stellate_Sphenopalatine_VS_Coeliac | 1 | 1 | 1 | 1 | 1 | 0 | 1 | 1 |
| METTL22 | 0.0355 | P1_P3_Lumbar_VS_P2_P4_Coeliac_Stellate_Sphenopalatine | 1 | 0 | 1 | 0 | 1 | 0 | 0 | 0 |
| SPTLC2 | 0.0355 | P1_P3_P4_Lumbar_Sphenopalatine_VS_P2_Coeliac_Stellate | 1 | 0 | 1 | 1 | 1 | 0 | 0 | 1 |
| DMWD | 0.0354 | P1_P3_P4_Lumbar_Sphenopalatine_VS_P2_Coeliac_Stellate | 1 | 0 | 1 | 1 | 1 | 0 | 0 | 1 |
| CPLX2 | 0.035 | P2_P3_P4_Lumbar_Coeliac_Stellate_VS_P1_Sphenopalatine | 0 | 1 | 1 | 1 | 1 | 1 | 1 | 0 |
| RASL10B | 0.0349 | P2_P3_P4_Lumbar_Sphenopalatine_VS_P1_Coeliac_Stellate | 0 | 1 | 1 | 1 | 1 | 0 | 0 | 1 |
| GM20300 | 0.0348 | P1_P4_Lumbar_Sphenopalatine_VS_P2_P3_Coeliac_Stellate | 1 | 0 | 0 | 1 | 1 | 0 | 0 | 1 |
| STK3 | 0.0346 | P1_P3_Lumbar_VS_P2_P4_Coeliac_Stellate_Sphenopalatine | 1 | 0 | 1 | 0 | 1 | 0 | 0 | 0 |
| RNGTT | 0.0346 | P2_P3_Lumbar_Coeliac_Sphenopalatine_VS_P1_P4_Stellate | 0 | 1 | 1 | 0 | 1 | 1 | 0 | 1 |
| NDUFAF5 | 0.0344 | P1_P2_Sphenopalatine_VS_P3_P4_Lumbar_Coeliac_Stellate | 1 | 1 | 0 | 0 | 0 | 0 | 0 | 1 |
| ZDHHC12 | 0.0344 | P1_P2_P4_Lumbar_Coeliac_Stellate_Sphenopalatine_VS_P3 | 1 | 1 | 0 | 1 | 1 | 1 | 1 | 1 |
| SKI | 0.0342 | P2_P3_P4_Lumbar_Sphenopalatine_VS_P1_Coeliac_Stellate | 0 | 1 | 1 | 1 | 1 | 0 | 0 | 1 |
| AK6 | 0.0342 | P1_P2_P3_Lumbar_Coeliac_Stellate_Sphenopalatine_VS_P4 | 1 | 1 | 1 | 0 | 1 | 1 | 1 | 1 |
| LAP3 | 0.0341 | P1_P2_P3_Lumbar_Coeliac_Stellate_VS_P4_Sphenopalatine | 1 | 1 | 1 | 0 | 1 | 1 | 1 | 0 |
| TBCEL | 0.034 | P2_P3_P4_Lumbar_Sphenopalatine_VS_P1_Coeliac_Stellate | 0 | 1 | 1 | 1 | 1 | 0 | 0 | 1 |
| CPTP | 0.0339 | P1_P3_P4_Lumbar_Sphenopalatine_VS_P2_Coeliac_Stellate | 1 | 0 | 1 | 1 | 1 | 0 | 0 | 1 |
| ACD | 0.0339 | P1_P2_P3_P4_Lumbar_Stellate_Sphenopalatine_VS_Coeliac | 1 | 1 | 1 | 1 | 1 | 0 | 1 | 1 |
| UBR3 | 0.0338 | P2_P3_P4_Lumbar_Sphenopalatine_VS_P1_Coeliac_Stellate | 0 | 1 | 1 | 1 | 1 | 0 | 0 | 1 |
| FBXL16 | 0.0338 | P2_P3_P4_Lumbar_Sphenopalatine_VS_P1_Coeliac_Stellate | 0 | 1 | 1 | 1 | 1 | 0 | 0 | 1 |
| EPB41L1 | 0.0336 | P2_P3_P4_Sphenopalatine_VS_P1_Lumbar_Coeliac_Stellate | 0 | 1 | 1 | 1 | 0 | 0 | 0 | 1 |
| SUPT6 | 0.0336 | P2_P3_Lumbar_Coeliac_Sphenopalatine_VS_P1_P4_Stellate | 0 | 1 | 1 | 0 | 1 | 1 | 0 | 1 |
| TMEM8 | 0.0333 | P1_P2_Sphenopalatine_VS_P3_P4_Lumbar_Coeliac_Stellate | 1 | 1 | 0 | 0 | 0 | 0 | 0 | 1 |
| EFR3B | 0.0332 | P1_P2_Coeliac_Sphenopalatine_VS_P3_P4_Lumbar_Stellate | 1 | 1 | 0 | 0 | 0 | 1 | 0 | 1 |
| HID1 | 0.0328 | P2_P3_Lumbar_VS_P1_P4_Coeliac_Stellate_Sphenopalatine | 0 | 1 | 1 | 0 | 1 | 0 | 0 | 0 |
| BOLA3 | 0.0328 | P1_P3_P4_Sphenopalatine_VS_P2_Lumbar_Coeliac_Stellate | 1 | 0 | 1 | 1 | 0 | 0 | 0 | 1 |
| NF1 | 0.0328 | P2_P3_P4_Coeliac_Sphenopalatine_VS_P1_Lumbar_Stellate | 0 | 1 | 1 | 1 | 0 | 1 | 0 | 1 |
| PLA2G12A | 0.0325 | P1_P2_P3_P4_Lumbar_Stellate_Sphenopalatine_VS_Coeliac | 1 | 1 | 1 | 1 | 1 | 0 | 1 | 1 |

| | | | | | | | | | | |
|---|---|---|---|---|---|---|---|---|---|---|
| SMIM10L2A | 0.0324 | P3_P4_Sphenopalatine_VS_P1_P2_Lumbar_Coeliac_Stellate | 0 | 0 | 1 | 1 | 0 | 0 | 0 | 1 |
| BORCS6 | 0.0324 | P1_P3_P4_Lumbar_Sphenopalatine_VS_P2_Coeliac_Stellate | 1 | 0 | 1 | 1 | 1 | 0 | 0 | 1 |
| C1GALT1C1 | 0.0322 | P1_P4_Lumbar_Coeliac_Stellate_VS_P2_P3_Sphenopalatine | 1 | 0 | 0 | 1 | 1 | 1 | 1 | 0 |
| NUDT4 | 0.032 | P1_P3_P4_Lumbar_Sphenopalatine_VS_P2_Coeliac_Stellate | 1 | 0 | 1 | 1 | 1 | 0 | 0 | 1 |
| AMPH | 0.0317 | P1_P3_P4_Lumbar_Sphenopalatine_VS_P2_Coeliac_Stellate | 1 | 0 | 1 | 1 | 1 | 0 | 0 | 1 |
| GSTO1 | 0.0317 | P1_P3_P4_Lumbar_Coeliac_Stellate_Sphenopalatine_VS_P2 | 1 | 0 | 1 | 1 | 1 | 1 | 1 | 1 |
| RXRB | 0.0316 | P1_P3_P4_Lumbar_Sphenopalatine_VS_P2_Coeliac_Stellate | 1 | 0 | 1 | 1 | 1 | 0 | 0 | 1 |
| IPO9 | 0.0315 | P2_P3_P4_Sphenopalatine_VS_P1_Lumbar_Coeliac_Stellate | 0 | 1 | 1 | 1 | 0 | 0 | 0 | 1 |
| SOCS6 | 0.0314 | P3_Lumbar_Coeliac_Stellate_VS_P1_P2_P4_Sphenopalatine | 0 | 0 | 1 | 0 | 1 | 1 | 1 | 0 |
| B930095G15 | 0.0313 | Lumbar_Sphenopalatine_VS_P1_P2_P3_P4_Coeliac_Stellate | 0 | 0 | 0 | 0 | 1 | 0 | 0 | 1 |
| IMP4 | 0.0313 | P4_Lumbar_Sphenopalatine_VS_P1_P2_P3_Coeliac_Stellate | 0 | 0 | 0 | 1 | 1 | 0 | 0 | 1 |
| GABRB3 | 0.0312 | P1_P3_P4_Lumbar_Sphenopalatine_VS_P2_Coeliac_Stellate | 1 | 0 | 1 | 1 | 1 | 0 | 0 | 1 |
| QRSL1 | 0.0311 | Lumbar_Sphenopalatine_VS_P1_P2_P3_P4_Coeliac_Stellate | 0 | 0 | 0 | 0 | 1 | 0 | 0 | 1 |
| STX1B | 0.031 | P1_P3_P4_Lumbar_Sphenopalatine_VS_P2_Coeliac_Stellate | 1 | 0 | 1 | 1 | 1 | 0 | 0 | 1 |
| TRMT10A | 0.0306 | P1_P3_P4_Lumbar_Coeliac_Stellate_Sphenopalatine_VS_P2 | 1 | 0 | 1 | 1 | 1 | 1 | 1 | 1 |
| HILPDA | 0.0305 | P1_P2_P4_Lumbar_Coeliac_Stellate_Sphenopalatine_VS_P3 | 1 | 1 | 0 | 1 | 1 | 1 | 1 | 1 |
| SMIM12 | 0.0303 | P1_P2_P3_P4_Lumbar_Stellate_Sphenopalatine_VS_Coeliac | 1 | 1 | 1 | 1 | 1 | 0 | 1 | 1 |
| ST6GALNAC( | 0.0302 | P3_Sphenopalatine_VS_P1_P2_P4_Lumbar_Coeliac_Stellate | 0 | 0 | 1 | 0 | 0 | 0 | 0 | 1 |
| FAAP20 | 0.0302 | P1_P2_P3_P4_Lumbar_Stellate_Sphenopalatine_VS_Coeliac | 1 | 1 | 1 | 1 | 1 | 0 | 1 | 1 |
| S100PBP | 0.0301 | P2_P3_P4_Sphenopalatine_VS_P1_Lumbar_Coeliac_Stellate | 0 | 1 | 1 | 1 | 0 | 0 | 0 | 1 |
| PDXK | 0.0301 | P4_Lumbar_Sphenopalatine_VS_P1_P2_P3_Coeliac_Stellate | 0 | 0 | 0 | 1 | 1 | 0 | 0 | 1 |
| POLR1E | 0.0297 | P1_P3_P4_Lumbar_Coeliac_Stellate_VS_P2_Sphenopalatine | 1 | 0 | 1 | 1 | 1 | 1 | 1 | 0 |
| CPEB2 | 0.0296 | P2_P3_P4_Coeliac_Sphenopalatine_VS_P1_Lumbar_Stellate | 0 | 1 | 1 | 1 | 0 | 1 | 0 | 1 |
| OSBPL6 | 0.0295 | P2_P3_P4_Coeliac_Sphenopalatine_VS_P1_Lumbar_Stellate | 0 | 1 | 1 | 1 | 0 | 1 | 0 | 1 |
| MIGA2 | 0.0293 | P2_Sphenopalatine_VS_P1_P3_P4_Lumbar_Coeliac_Stellate | 0 | 1 | 0 | 0 | 0 | 0 | 0 | 1 |
| MROH1 | 0.0293 | P4_Lumbar_Sphenopalatine_VS_P1_P2_P3_Coeliac_Stellate | 0 | 0 | 0 | 1 | 1 | 0 | 0 | 1 |
| ATP8A2 | 0.0292 | P2_P3_P4_Lumbar_Coeliac_Stellate_Sphenopalatine_VS_P1 | 0 | 1 | 1 | 1 | 1 | 1 | 1 | 1 |
| MED7 | 0.0291 | P1_P3_P4_Lumbar_Coeliac_Stellate_VS_P2_Sphenopalatine | 1 | 0 | 1 | 1 | 1 | 1 | 1 | 0 |
| ATXN2 | 0.0291 | P4_Lumbar_Sphenopalatine_VS_P1_P2_P3_Coeliac_Stellate | 0 | 0 | 0 | 1 | 1 | 0 | 0 | 1 |
| TMEM242 | 0.0291 | P1_P2_P3_P4_Lumbar_Stellate_Sphenopalatine_VS_Coeliac | 1 | 1 | 1 | 1 | 1 | 0 | 1 | 1 |

| | | | | | | | | | | |
|---|---|---|---|---|---|---|---|---|---|---|
| MRAS | 0.0288 | P1_P2_P3_Lumbar_VS_P4_Coeliac_Stellate_Sphenopalatine | 1 | 1 | 1 | 0 | 1 | 0 | 0 | 0 |
| ACSF3 | 0.0288 | P1_P3_P4_Lumbar_Sphenopalatine_VS_P2_Coeliac_Stellate | 1 | 0 | 1 | 1 | 1 | 0 | 0 | 1 |
| BCAM | 0.0287 | P3_Sphenopalatine_VS_P1_P2_P4_Lumbar_Coeliac_Stellate | 0 | 0 | 1 | 0 | 0 | 0 | 0 | 1 |
| FKRP | 0.0285 | P3_P4_Lumbar_Sphenopalatine_VS_P1_P2_Coeliac_Stellate | 0 | 0 | 1 | 1 | 1 | 0 | 0 | 1 |
| NBR1 | 0.0284 | P2_P4_Coeliac_Sphenopalatine_VS_P1_P3_Lumbar_Stellate | 0 | 1 | 0 | 1 | 0 | 1 | 0 | 1 |
| MIR100HG | 0.0284 | P1_P2_P4_Lumbar_Coeliac_Stellate_Sphenopalatine_VS_P3 | 1 | 1 | 0 | 1 | 1 | 1 | 1 | 1 |
| DTNB | 0.0282 | P1_P3_P4_Lumbar_Coeliac_Stellate_Sphenopalatine_VS_P2 | 1 | 0 | 1 | 1 | 1 | 1 | 1 | 1 |
| UBTD1 | 0.0281 | P3_Lumbar_Coeliac_Stellate_VS_P1_P2_P4_Sphenopalatine | 0 | 0 | 1 | 0 | 1 | 1 | 1 | 0 |
| ZFP593 | 0.0281 | P1_P3_P4_Lumbar_Sphenopalatine_VS_P2_Coeliac_Stellate | 1 | 0 | 1 | 1 | 1 | 0 | 0 | 1 |
| FAM199X | 0.028 | P1_P2_P3_Lumbar_VS_P4_Coeliac_Stellate_Sphenopalatine | 1 | 1 | 1 | 0 | 1 | 0 | 0 | 0 |
| CDH2 | 0.0275 | P3_P4_Lumbar_Sphenopalatine_VS_P1_P2_Coeliac_Stellate | 0 | 0 | 1 | 1 | 1 | 0 | 0 | 1 |
| BEAN1 | 0.0273 | P1_P2_P3_VS_P4_Lumbar_Coeliac_Stellate_Sphenopalatine | 1 | 1 | 1 | 0 | 0 | 0 | 0 | 0 |
| TMUB1 | 0.0271 | P3_P4_Lumbar_Sphenopalatine_VS_P1_P2_Coeliac_Stellate | 0 | 0 | 1 | 1 | 1 | 0 | 0 | 1 |
| PCK2 | 0.027 | P1_P2_P4_Lumbar_Coeliac_Stellate_VS_P3_Sphenopalatine | 1 | 1 | 0 | 1 | 1 | 1 | 1 | 0 |
| FOS | 0.027 | P2_P3_P4_Sphenopalatine_VS_P1_Lumbar_Coeliac_Stellate | 0 | 1 | 1 | 1 | 0 | 0 | 0 | 1 |
| XPR1 | 0.027 | P2_P3_P4_Lumbar_Coeliac_Stellate_Sphenopalatine_VS_P1 | 0 | 1 | 1 | 1 | 1 | 1 | 1 | 1 |
| A430005L14I | 0.0267 | P1_P4_Lumbar_Sphenopalatine_VS_P2_P3_Coeliac_Stellate | 1 | 0 | 0 | 1 | 1 | 0 | 0 | 1 |
| ELP6 | 0.0265 | P1_P3_P4_Lumbar_Coeliac_Stellate_VS_P2_Sphenopalatine | 1 | 0 | 1 | 1 | 1 | 1 | 1 | 0 |
| MECR | 0.0264 | P1_P4_Lumbar_Sphenopalatine_VS_P2_P3_Coeliac_Stellate | 1 | 0 | 0 | 1 | 1 | 0 | 0 | 1 |
| CHST7 | 0.0263 | P1_P2_P4_Lumbar_Coeliac_Stellate_Sphenopalatine_VS_P3 | 1 | 1 | 0 | 1 | 1 | 1 | 1 | 1 |
| CARMIL2 | 0.0261 | P3_P4_Sphenopalatine_VS_P1_P2_Lumbar_Coeliac_Stellate | 0 | 0 | 1 | 1 | 0 | 0 | 0 | 1 |
| AGK | 0.026 | Lumbar_Coeliac_Stellate_VS_P1_P2_P3_P4_Sphenopalatine | 0 | 0 | 0 | 0 | 1 | 1 | 1 | 0 |
| GMPR | 0.0259 | P1_P3_P4_Lumbar_Sphenopalatine_VS_P2_Coeliac_Stellate | 1 | 0 | 1 | 1 | 1 | 0 | 0 | 1 |
| H2AFX | 0.0258 | P1_P3_P4_Lumbar_Coeliac_Stellate_Sphenopalatine_VS_P2 | 1 | 0 | 1 | 1 | 1 | 1 | 1 | 1 |
| GMDS | 0.0257 | P1_P2_P3_Lumbar_VS_P4_Coeliac_Stellate_Sphenopalatine | 1 | 1 | 1 | 0 | 1 | 0 | 0 | 0 |
| NUDT18 | 0.0256 | P1_P3_P4_Sphenopalatine_VS_P2_Lumbar_Coeliac_Stellate | 1 | 0 | 1 | 1 | 0 | 0 | 0 | 1 |
| CARD19 | 0.0255 | P1_P2_P3_P4_Lumbar_Stellate_Sphenopalatine_VS_Coeliac | 1 | 1 | 1 | 1 | 1 | 0 | 1 | 1 |
| 9130401M01 | 0.0254 | P1_P3_P4_Lumbar_Coeliac_Stellate_Sphenopalatine_VS_P2 | 1 | 0 | 1 | 1 | 1 | 1 | 1 | 1 |
| CHCHD10 | 0.0253 | P1_P2_P3_P4_Lumbar_Stellate_Sphenopalatine_VS_Coeliac | 1 | 1 | 1 | 1 | 1 | 0 | 1 | 1 |
| 2310061I04R | 0.0249 | P2_P3_P4_Sphenopalatine_VS_P1_Lumbar_Coeliac_Stellate | 0 | 1 | 1 | 1 | 0 | 0 | 0 | 1 |

| | | | | | | | | | | |
|---|---|---|---|---|---|---|---|---|---|---|
| GAN | 0.0249 | P1_P2_P3_Lumbar_Coeliac_Stellate_Sphenopalatine_VS_P4 | 1 | 1 | 1 | 0 | 1 | 1 | 1 | 1 |
| FCOR | 0.0248 | P3_P4_Lumbar_Sphenopalatine_VS_P1_P2_Coeliac_Stellate | 0 | 0 | 1 | 1 | 1 | 0 | 0 | 1 |
| ZFP422 | 0.0247 | P1_P3_P4_Lumbar_Coeliac_Stellate_VS_P2_Sphenopalatine | 1 | 0 | 1 | 1 | 1 | 1 | 1 | 0 |
| CIC | 0.0247 | P1_P2_P3_Lumbar_Sphenopalatine_VS_P4_Coeliac_Stellate | 1 | 1 | 1 | 0 | 1 | 0 | 0 | 1 |
| HOXC5 | 0.0245 | Coeliac_Stellate_VS_P1_P2_P3_P4_Lumbar_Sphenopalatine | 0 | 0 | 0 | 0 | 0 | 1 | 1 | 0 |
| CRY1 | 0.0243 | P1_P4_Lumbar_Coeliac_Stellate_VS_P2_P3_Sphenopalatine | 1 | 0 | 0 | 1 | 1 | 1 | 1 | 0 |
| CENPT | 0.0243 | P1_P3_P4_Lumbar_Coeliac_Stellate_Sphenopalatine_VS_P2 | 1 | 0 | 1 | 1 | 1 | 1 | 1 | 1 |
| CACNA2D2 | 0.0242 | Lumbar_Sphenopalatine_VS_P1_P2_P3_P4_Coeliac_Stellate | 0 | 0 | 0 | 0 | 1 | 0 | 0 | 1 |
| APEH | 0.0241 | P1_P3_Lumbar_VS_P2_P4_Coeliac_Stellate_Sphenopalatine | 1 | 0 | 1 | 0 | 1 | 0 | 0 | 0 |
| ETFA | 0.0241 | P1_P3_P4_Lumbar_Coeliac_Stellate_VS_P2_Sphenopalatine | 1 | 0 | 1 | 1 | 1 | 1 | 1 | 0 |
| SMARCD3 | 0.0241 | P1_P2_Sphenopalatine_VS_P3_P4_Lumbar_Coeliac_Stellate | 1 | 1 | 0 | 0 | 0 | 0 | 0 | 1 |
| SATB1 | 0.0238 | P1_P2_P3_Lumbar_VS_P4_Coeliac_Stellate_Sphenopalatine | 1 | 1 | 1 | 0 | 1 | 0 | 0 | 0 |
| SAMD10 | 0.0237 | P1_P2_P3_P4_Lumbar_Stellate_Sphenopalatine_VS_Coeliac | 1 | 1 | 1 | 1 | 1 | 0 | 1 | 1 |
| MCEE | 0.0237 | P1_P3_P4_Lumbar_Coeliac_Stellate_Sphenopalatine_VS_P2 | 1 | 0 | 1 | 1 | 1 | 1 | 1 | 1 |
| NUDT1 | 0.0236 | P1_P3_Lumbar_VS_P2_P4_Coeliac_Stellate_Sphenopalatine | 1 | 0 | 1 | 0 | 1 | 0 | 0 | 0 |
| NSMCE1 | 0.0233 | P1_P3_P4_Lumbar_Coeliac_Stellate_VS_P2_Sphenopalatine | 1 | 0 | 1 | 1 | 1 | 1 | 1 | 0 |
| PIK3CD | 0.0232 | P1_P2_P4_Lumbar_Coeliac_Stellate_Sphenopalatine_VS_P3 | 1 | 1 | 0 | 1 | 1 | 1 | 1 | 1 |
| LIPO3 | 0.0231 | P1_P2_P3_Lumbar_Coeliac_Stellate_VS_P4_Sphenopalatine | 1 | 1 | 1 | 0 | 1 | 1 | 1 | 0 |
| LYPLAL1 | 0.023 | P1_P3_P4_Lumbar_Coeliac_Stellate_Sphenopalatine_VS_P2 | 1 | 0 | 1 | 1 | 1 | 1 | 1 | 1 |
| TDP2 | 0.0227 | P1_P3_P4_Lumbar_Coeliac_Stellate_VS_P2_Sphenopalatine | 1 | 0 | 1 | 1 | 1 | 1 | 1 | 0 |
| TRADD | 0.0226 | P1_P3_Lumbar_VS_P2_P4_Coeliac_Stellate_Sphenopalatine | 1 | 0 | 1 | 0 | 1 | 0 | 0 | 0 |
| LRRC51 | 0.0221 | P1_P3_P4_Lumbar_Coeliac_Stellate_Sphenopalatine_VS_P2 | 1 | 0 | 1 | 1 | 1 | 1 | 1 | 1 |
| HEXB | 0.0218 | P1_P4_Lumbar_Coeliac_Stellate_VS_P2_P3_Sphenopalatine | 1 | 0 | 0 | 1 | 1 | 1 | 1 | 0 |
| WASF2 | 0.0217 | P1_P2_P3_Lumbar_VS_P4_Coeliac_Stellate_Sphenopalatine | 1 | 1 | 1 | 0 | 1 | 0 | 0 | 0 |
| GID4 | 0.0217 | P1_P2_Coeliac_VS_P3_P4_Lumbar_Stellate_Sphenopalatine | 1 | 1 | 0 | 0 | 0 | 1 | 0 | 0 |
| IDE | 0.0217 | P1_P3_P4_Lumbar_Sphenopalatine_VS_P2_Coeliac_Stellate | 1 | 0 | 1 | 1 | 1 | 0 | 0 | 1 |
| RPAIN | 0.0216 | P1_P3_P4_Lumbar_Sphenopalatine_VS_P2_Coeliac_Stellate | 1 | 0 | 1 | 1 | 1 | 0 | 0 | 1 |
| MED29 | 0.0216 | P1_P3_P4_Lumbar_Coeliac_Stellate_Sphenopalatine_VS_P2 | 1 | 0 | 1 | 1 | 1 | 1 | 1 | 1 |
| LRRC49 | 0.0215 | P1_P3_P4_Lumbar_Sphenopalatine_VS_P2_Coeliac_Stellate | 1 | 0 | 1 | 1 | 1 | 0 | 0 | 1 |
| TMEM120B | 0.0211 | Lumbar_Coeliac_Stellate_VS_P1_P2_P3_P4_Sphenopalatine | 0 | 0 | 0 | 0 | 1 | 1 | 1 | 0 |

| | | | | | | | | | | |
|---|---|---|---|---|---|---|---|---|---|---|
| ADAM10 | 0.021 | P2_P3_Lumbar_VS_P1_P4_Coeliac_Stellate_Sphenopalatine | 0 | 1 | 1 | 0 | 1 | 0 | 0 | 0 |
| DHX40 | 0.0209 | P2_P3_Lumbar_VS_P1_P4_Coeliac_Stellate_Sphenopalatine | 0 | 1 | 1 | 0 | 1 | 0 | 0 | 0 |
| LYRM2 | 0.0208 | P1_P3_P4_Lumbar_Sphenopalatine_VS_P2_Coeliac_Stellate | 1 | 0 | 1 | 1 | 1 | 0 | 0 | 1 |
| SCYL1 | 0.0208 | P1_P3_P4_Lumbar_Sphenopalatine_VS_P2_Coeliac_Stellate | 1 | 0 | 1 | 1 | 1 | 0 | 0 | 1 |
| CNOT2 | 0.0207 | P1_P2_P4_Lumbar_Coeliac_Stellate_VS_P3_Sphenopalatine | 1 | 1 | 0 | 1 | 1 | 1 | 1 | 0 |
| SEMA6D | 0.0206 | P2_P3_P4_Lumbar_Coeliac_Stellate_Sphenopalatine_VS_P1 | 0 | 1 | 1 | 1 | 1 | 1 | 1 | 1 |
| 1190007I07R | 0.0205 | P1_P3_P4_Lumbar_Coeliac_Stellate_VS_P2_Sphenopalatine | 1 | 0 | 1 | 1 | 1 | 1 | 1 | 0 |
| DENND5A | 0.0205 | P2_P3_P4_Coeliac_Sphenopalatine_VS_P1_Lumbar_Stellate | 0 | 1 | 1 | 1 | 0 | 1 | 0 | 1 |
| SNAPC5 | 0.0204 | P1_P2_P3_P4_Lumbar_Stellate_Sphenopalatine_VS_Coeliac | 1 | 1 | 1 | 1 | 1 | 0 | 1 | 1 |
| 1110012L19R | 0.0203 | P1_P3_P4_Lumbar_Coeliac_Stellate_VS_P2_Sphenopalatine | 1 | 0 | 1 | 1 | 1 | 1 | 1 | 0 |
| GAA | 0.0202 | P2_P3_P4_Sphenopalatine_VS_P1_Lumbar_Coeliac_Stellate | 0 | 1 | 1 | 1 | 0 | 0 | 0 | 1 |
| HSPB8 | 0.02 | P1_P2_P3_P4_Lumbar_Stellate_Sphenopalatine_VS_Coeliac | 1 | 1 | 1 | 1 | 1 | 0 | 1 | 1 |
| EEF1AKMT1 | 0.0199 | P1_P3_P4_Lumbar_Coeliac_Stellate_Sphenopalatine_VS_P2 | 1 | 0 | 1 | 1 | 1 | 1 | 1 | 1 |
| PAM | 0.0198 | P2_P3_P4_Coeliac_Sphenopalatine_VS_P1_Lumbar_Stellate | 0 | 1 | 1 | 1 | 0 | 1 | 0 | 1 |
| ZFAND2A | 0.0197 | P2_P3_P4_Sphenopalatine_VS_P1_Lumbar_Coeliac_Stellate | 0 | 1 | 1 | 1 | 0 | 0 | 0 | 1 |
| BCAR1 | 0.0197 | Lumbar_Sphenopalatine_VS_P1_P2_P3_P4_Coeliac_Stellate | 0 | 0 | 0 | 0 | 1 | 0 | 0 | 1 |
| ATXN1L | 0.0197 | P1_P2_P3_Lumbar_Sphenopalatine_VS_P4_Coeliac_Stellate | 1 | 1 | 1 | 0 | 1 | 0 | 0 | 1 |
| DCX | 0.0197 | P1_P3_Lumbar_Coeliac_Stellate_Sphenopalatine_VS_P2_P4 | 1 | 0 | 1 | 0 | 1 | 1 | 1 | 1 |
| NUDT13 | 0.0196 | P1_P3_Lumbar_VS_P2_P4_Coeliac_Stellate_Sphenopalatine | 1 | 0 | 1 | 0 | 1 | 0 | 0 | 0 |
| SNRNP40 | 0.0193 | P1_P3_P4_Lumbar_Coeliac_Stellate_Sphenopalatine_VS_P2 | 1 | 0 | 1 | 1 | 1 | 1 | 1 | 1 |
| IPO11 | 0.0192 | P2_P3_Lumbar_VS_P1_P4_Coeliac_Stellate_Sphenopalatine | 0 | 1 | 1 | 0 | 1 | 0 | 0 | 0 |
| TCTA | 0.0192 | P1_P2_P3_P4_Lumbar_Stellate_Sphenopalatine_VS_Coeliac | 1 | 1 | 1 | 1 | 1 | 0 | 1 | 1 |
| SAMD4B | 0.019 | P1_P2_Lumbar_Sphenopalatine_VS_P3_P4_Coeliac_Stellate | 1 | 1 | 0 | 0 | 1 | 0 | 0 | 1 |
| HIPK1 | 0.019 | P1_P2_P3_Lumbar_Sphenopalatine_VS_P4_Coeliac_Stellate | 1 | 1 | 1 | 0 | 1 | 0 | 0 | 1 |
| OPTN | 0.0189 | P1_P2_P3_Coeliac_VS_P4_Lumbar_Stellate_Sphenopalatine | 1 | 1 | 1 | 0 | 0 | 1 | 0 | 0 |
| CAMKK2 | 0.0189 | P1_P2_P3_Lumbar_Coeliac_Stellate_Sphenopalatine_VS_P4 | 1 | 1 | 1 | 0 | 1 | 1 | 1 | 1 |
| PNPLA2 | 0.0187 | P1_P3_P4_Lumbar_Sphenopalatine_VS_P2_Coeliac_Stellate | 1 | 0 | 1 | 1 | 1 | 0 | 0 | 1 |
| TWF2 | 0.0186 | P1_P3_P4_Lumbar_Coeliac_Stellate_VS_P2_Sphenopalatine | 1 | 0 | 1 | 1 | 1 | 1 | 1 | 0 |
| MORN2 | 0.0185 | P3_P4_Lumbar_Sphenopalatine_VS_P1_P2_Coeliac_Stellate | 0 | 0 | 1 | 1 | 1 | 0 | 0 | 1 |
| COX18 | 0.0184 | P1_P3_P4_Lumbar_Sphenopalatine_VS_P2_Coeliac_Stellate | 1 | 0 | 1 | 1 | 1 | 0 | 0 | 1 |

| | | | | | | | | | | |
|---|---|---|---|---|---|---|---|---|---|---|
| CMC1 | 0.0183 | P1_P3_P4_Lumbar_Sphenopalatine_VS_P2_Coeliac_Stellate | 1 | 0 | 1 | 1 | 1 | 0 | 0 | 1 |
| FNDC10 | 0.0179 | P1_P3_P4_Sphenopalatine_VS_P2_Lumbar_Coeliac_Stellate | 1 | 0 | 1 | 1 | 0 | 0 | 0 | 1 |
| ANAPC1 | 0.0176 | P2_P3_P4_Coeliac_Sphenopalatine_VS_P1_Lumbar_Stellate | 0 | 1 | 1 | 1 | 0 | 1 | 0 | 1 |
| SMN1 | 0.0175 | P1_P3_P4_Lumbar_Coeliac_Stellate_VS_P2_Sphenopalatine | 1 | 0 | 1 | 1 | 1 | 1 | 1 | 0 |
| MTLN | 0.0175 | P3_P4_Sphenopalatine_VS_P1_P2_Lumbar_Coeliac_Stellate | 0 | 0 | 1 | 1 | 0 | 0 | 0 | 1 |
| PDLIM7 | 0.0175 | P1_P3_P4_Sphenopalatine_VS_P2_Lumbar_Coeliac_Stellate | 1 | 0 | 1 | 1 | 0 | 0 | 0 | 1 |
| COQ4 | 0.0175 | P1_P3_P4_Lumbar_Coeliac_Stellate_Sphenopalatine_VS_P2 | 1 | 0 | 1 | 1 | 1 | 1 | 1 | 1 |
| MED31 | 0.0175 | P1_P3_P4_Lumbar_Coeliac_Stellate_Sphenopalatine_VS_P2 | 1 | 0 | 1 | 1 | 1 | 1 | 1 | 1 |
| PTP4A3 | 0.0173 | P1_P2_P3_Lumbar_Coeliac_Stellate_Sphenopalatine_VS_P4 | 1 | 1 | 1 | 0 | 1 | 1 | 1 | 1 |
| PACSIN2 | 0.0172 | P1_P2_P3_Lumbar_VS_P4_Coeliac_Stellate_Sphenopalatine | 1 | 1 | 1 | 0 | 1 | 0 | 0 | 0 |
| MMS19 | 0.0172 | P2_P3_P4_Lumbar_Coeliac_Stellate_VS_P1_Sphenopalatine | 0 | 1 | 1 | 1 | 1 | 1 | 1 | 0 |
| CLVS1 | 0.017 | P1_P2_P4_Lumbar_Coeliac_Stellate_VS_P3_Sphenopalatine | 1 | 1 | 0 | 1 | 1 | 1 | 1 | 0 |
| PLCD1 | 0.0168 | P1_P3_Lumbar_VS_P2_P4_Coeliac_Stellate_Sphenopalatine | 1 | 0 | 1 | 0 | 1 | 0 | 0 | 0 |
| JPH3 | 0.0168 | P1_P2_P3_P4_Lumbar_Stellate_Sphenopalatine_VS_Coeliac | 1 | 1 | 1 | 1 | 1 | 0 | 1 | 1 |
| CYB561D2 | 0.0168 | P1_P3_P4_Lumbar_Coeliac_Stellate_Sphenopalatine_VS_P2 | 1 | 0 | 1 | 1 | 1 | 1 | 1 | 1 |
| DNAJB4 | 0.0166 | P1_P2_P4_Lumbar_Coeliac_Stellate_Sphenopalatine_VS_P3 | 1 | 1 | 0 | 1 | 1 | 1 | 1 | 1 |
| LARGE1 | 0.0164 | P2_P3_Lumbar_VS_P1_P4_Coeliac_Stellate_Sphenopalatine | 0 | 1 | 1 | 0 | 1 | 0 | 0 | 0 |
| SCAMP5 | 0.0164 | P2_Sphenopalatine_VS_P1_P3_P4_Lumbar_Coeliac_Stellate | 0 | 1 | 0 | 0 | 0 | 0 | 0 | 1 |
| TTC39A | 0.0163 | P1_P3_Lumbar_VS_P2_P4_Coeliac_Stellate_Sphenopalatine | 1 | 0 | 1 | 0 | 1 | 0 | 0 | 0 |
| SOCS2 | 0.0163 | P1_P2_P3_P4_Lumbar_Stellate_Sphenopalatine_VS_Coeliac | 1 | 1 | 1 | 1 | 1 | 0 | 1 | 1 |
| CCDC184 | 0.016 | P1_P3_Lumbar_Sphenopalatine_VS_P2_P4_Coeliac_Stellate | 1 | 0 | 1 | 0 | 1 | 0 | 0 | 1 |
| FBXO28 | 0.016 | P4_Lumbar_Sphenopalatine_VS_P1_P2_P3_Coeliac_Stellate | 0 | 0 | 0 | 1 | 1 | 0 | 0 | 1 |
| ACSL5 | 0.0158 | P1_P3_Lumbar_VS_P2_P4_Coeliac_Stellate_Sphenopalatine | 1 | 0 | 1 | 0 | 1 | 0 | 0 | 0 |
| IQGAP1 | 0.0157 | P4_Lumbar_Sphenopalatine_VS_P1_P2_P3_Coeliac_Stellate | 0 | 0 | 0 | 1 | 1 | 0 | 0 | 1 |
| FEM1A | 0.0155 | P1_P2_P3_Lumbar_Sphenopalatine_VS_P4_Coeliac_Stellate | 1 | 1 | 1 | 0 | 1 | 0 | 0 | 1 |
| LIX1 | 0.0154 | P1_P3_Lumbar_Sphenopalatine_VS_P2_P4_Coeliac_Stellate | 1 | 0 | 1 | 0 | 1 | 0 | 0 | 1 |
| ZFP511 | 0.0152 | P1_P3_Lumbar_VS_P2_P4_Coeliac_Stellate_Sphenopalatine | 1 | 0 | 1 | 0 | 1 | 0 | 0 | 0 |
| SOX11 | 0.015 | P1_P3_P4_Lumbar_Coeliac_Stellate_VS_P2_Sphenopalatine | 1 | 0 | 1 | 1 | 1 | 1 | 1 | 0 |
| ANKRD10 | 0.0149 | P4_Lumbar_Sphenopalatine_VS_P1_P2_P3_Coeliac_Stellate | 0 | 0 | 0 | 1 | 1 | 0 | 0 | 1 |
| NIT2 | 0.0146 | P1_P3_P4_Lumbar_Coeliac_Stellate_Sphenopalatine_VS_P2 | 1 | 0 | 1 | 1 | 1 | 1 | 1 | 1 |

| | | | | | | | | | | |
|---|---|---|---|---|---|---|---|---|---|---|
| 1700025G04 | 0.0144 | P1_P3_P4_Lumbar_Coeliac_Stellate_VS_P2_Sphenopalatine | 1 | 0 | 1 | 1 | 1 | 1 | 1 | 0 |
| LBH | 0.0144 | P1_P3_P4_Lumbar_Coeliac_Stellate_VS_P2_Sphenopalatine | 1 | 0 | 1 | 1 | 1 | 1 | 1 | 0 |
| OASL2 | 0.0143 | P2_Sphenopalatine_VS_P1_P3_P4_Lumbar_Coeliac_Stellate | 0 | 1 | 0 | 0 | 0 | 0 | 0 | 1 |
| FAM20C | 0.0142 | P1_P2_P4_Lumbar_Coeliac_Stellate_Sphenopalatine_VS_P3 | 1 | 1 | 0 | 1 | 1 | 1 | 1 | 1 |
| ABCB8 | 0.0141 | P2_P4_Coeliac_Sphenopalatine_VS_P1_P3_Lumbar_Stellate | 0 | 1 | 0 | 1 | 0 | 1 | 0 | 1 |
| EXOC6B | 0.0135 | P4_Lumbar_Sphenopalatine_VS_P1_P2_P3_Coeliac_Stellate | 0 | 0 | 0 | 1 | 1 | 0 | 0 | 1 |
| RFC2 | 0.0134 | P1_P3_P4_Lumbar_Coeliac_Stellate_Sphenopalatine_VS_P2 | 1 | 0 | 1 | 1 | 1 | 1 | 1 | 1 |
| 1500009L16F | 0.0131 | P1_P3_P4_Sphenopalatine_VS_P2_Lumbar_Coeliac_Stellate | 1 | 0 | 1 | 1 | 0 | 0 | 0 | 1 |
| TNRC6B | 0.0129 | P2_P4_Coeliac_Sphenopalatine_VS_P1_P3_Lumbar_Stellate | 0 | 1 | 0 | 1 | 0 | 1 | 0 | 1 |
| M6PR | 0.0128 | P1_P3_P4_Sphenopalatine_VS_P2_Lumbar_Coeliac_Stellate | 1 | 0 | 1 | 1 | 0 | 0 | 0 | 1 |
| SF3B4 | 0.0128 | P1_P3_P4_Lumbar_Coeliac_Stellate_Sphenopalatine_VS_P2 | 1 | 0 | 1 | 1 | 1 | 1 | 1 | 1 |
| SCMH1 | 0.0127 | P2_P4_Coeliac_Sphenopalatine_VS_P1_P3_Lumbar_Stellate | 0 | 1 | 0 | 1 | 0 | 1 | 0 | 1 |
| RRAS2 | 0.0123 | P1_P2_P3_P4_Lumbar_Stellate_Sphenopalatine_VS_Coeliac | 1 | 1 | 1 | 1 | 1 | 0 | 1 | 1 |
| ABHD14A | 0.0122 | P4_Lumbar_Sphenopalatine_VS_P1_P2_P3_Coeliac_Stellate | 0 | 0 | 0 | 1 | 1 | 0 | 0 | 1 |
| RAB24 | 0.0121 | P1_P2_P3_P4_Lumbar_Stellate_Sphenopalatine_VS_Coeliac | 1 | 1 | 1 | 1 | 1 | 0 | 1 | 1 |
| PEMT | 0.0121 | P1_P3_P4_Lumbar_Coeliac_Stellate_Sphenopalatine_VS_P2 | 1 | 0 | 1 | 1 | 1 | 1 | 1 | 1 |
| LRRC59 | 0.012 | P1_P2_P3_Lumbar_Coeliac_Stellate_Sphenopalatine_VS_P4 | 1 | 1 | 1 | 0 | 1 | 1 | 1 | 1 |
| FRG1 | 0.0117 | P1_P3_P4_Lumbar_Coeliac_Stellate_VS_P2_Sphenopalatine | 1 | 0 | 1 | 1 | 1 | 1 | 1 | 0 |
| GM28320 | 0.0116 | P1_P2_Lumbar_Sphenopalatine_VS_P3_P4_Coeliac_Stellate | 1 | 1 | 0 | 0 | 1 | 0 | 0 | 1 |
| COPB1 | 0.0114 | P1_P2_P4_Lumbar_Coeliac_Stellate_VS_P3_Sphenopalatine | 1 | 1 | 0 | 1 | 1 | 1 | 1 | 0 |
| AP1B1 | 0.0114 | P2_P3_P4_Coeliac_Sphenopalatine_VS_P1_Lumbar_Stellate | 0 | 1 | 1 | 1 | 0 | 1 | 0 | 1 |
| ACTL6A | 0.0112 | P3_Lumbar_Coeliac_Stellate_VS_P1_P2_P4_Sphenopalatine | 0 | 0 | 1 | 0 | 1 | 1 | 1 | 0 |
| RMDN1 | 0.0112 | P1_P3_P4_Lumbar_Coeliac_Stellate_VS_P2_Sphenopalatine | 1 | 0 | 1 | 1 | 1 | 1 | 1 | 0 |
| GBE1 | 0.0109 | P1_P2_P3_Lumbar_Sphenopalatine_VS_P4_Coeliac_Stellate | 1 | 1 | 1 | 0 | 1 | 0 | 0 | 1 |
| SFSWAP | 0.0108 | P2_P3_VS_P1_P4_Lumbar_Coeliac_Stellate_Sphenopalatine | 0 | 1 | 1 | 0 | 0 | 0 | 0 | 0 |
| MED22 | 0.0106 | Lumbar_Coeliac_Stellate_VS_P1_P2_P3_P4_Sphenopalatine | 0 | 0 | 0 | 0 | 1 | 1 | 1 | 0 |
| MOSPD2 | 0.0106 | P4_Lumbar_Sphenopalatine_VS_P1_P2_P3_Coeliac_Stellate | 0 | 0 | 0 | 1 | 1 | 0 | 0 | 1 |
| POGLUT1 | 0.0106 | P3_P4_Lumbar_Sphenopalatine_VS_P1_P2_Coeliac_Stellate | 0 | 0 | 1 | 1 | 1 | 0 | 0 | 1 |
| GALE | 0.0106 | P1_P3_P4_Lumbar_Coeliac_Stellate_Sphenopalatine_VS_P2 | 1 | 0 | 1 | 1 | 1 | 1 | 1 | 1 |
| FH1 | 0.0105 | P1_P3_P4_Lumbar_Coeliac_Stellate_Sphenopalatine_VS_P2 | 1 | 0 | 1 | 1 | 1 | 1 | 1 | 1 |

| | | | | | | | | | | |
|---|---|---|---|---|---|---|---|---|---|---|
| EPS15 | 0.0103 | P3_P4_Sphenopalatine_VS_P1_P2_Lumbar_Coeliac_Stellate | 0 | 0 | 1 | 1 | 0 | 0 | 0 | 1 |
| AGAP3 | 0.0097 | P2_P3_P4_Coeliac_Sphenopalatine_VS_P1_Lumbar_Stellate | 0 | 1 | 1 | 1 | 0 | 1 | 0 | 1 |
| ANO10 | 0.0096 | P2_P3_Lumbar_VS_P1_P4_Coeliac_Stellate_Sphenopalatine | 0 | 1 | 1 | 0 | 1 | 0 | 0 | 0 |
| G6PDX | 0.0096 | P2_Sphenopalatine_VS_P1_P3_P4_Lumbar_Coeliac_Stellate | 0 | 1 | 0 | 0 | 0 | 0 | 0 | 1 |
| PNRC1 | 0.0095 | P2_P4_Coeliac_Sphenopalatine_VS_P1_P3_Lumbar_Stellate | 0 | 1 | 0 | 1 | 0 | 1 | 0 | 1 |
| WDR73 | 0.0095 | P1_P3_P4_Lumbar_Coeliac_Stellate_Sphenopalatine_VS_P2 | 1 | 0 | 1 | 1 | 1 | 1 | 1 | 1 |
| NUDT2 | 0.0093 | P1_P3_P4_Lumbar_Coeliac_Stellate_Sphenopalatine_VS_P2 | 1 | 0 | 1 | 1 | 1 | 1 | 1 | 1 |
| ZFPL1 | 0.0092 | P1_P3_P4_Lumbar_Coeliac_Stellate_VS_P2_Sphenopalatine | 1 | 0 | 1 | 1 | 1 | 1 | 1 | 0 |
| AS3MT | 0.0092 | P1_P4_Coeliac_Sphenopalatine_VS_P2_P3_Lumbar_Stellate | 1 | 0 | 0 | 1 | 0 | 1 | 0 | 1 |
| KIF13A | 0.0086 | P2_P4_Coeliac_Sphenopalatine_VS_P1_P3_Lumbar_Stellate | 0 | 1 | 0 | 1 | 0 | 1 | 0 | 1 |
| CDKL2 | 0.0085 | P3_P4_Lumbar_Sphenopalatine_VS_P1_P2_Coeliac_Stellate | 0 | 0 | 1 | 1 | 1 | 0 | 0 | 1 |
| ST6GALNAC | 0.0083 | P4_Lumbar_Coeliac_Stellate_VS_P1_P2_P3_Sphenopalatine | 0 | 0 | 0 | 1 | 1 | 1 | 1 | 0 |
| ATP1A3 | 0.0082 | P2_P3_P4_Lumbar_Coeliac_Stellate_Sphenopalatine_VS_P1 | 0 | 1 | 1 | 1 | 1 | 1 | 1 | 1 |
| CTSZ | 0.0082 | P1_P2_P4_Lumbar_Coeliac_Stellate_VS_P3_Sphenopalatine | 1 | 1 | 0 | 1 | 1 | 1 | 1 | 0 |
| PITHD1 | 0.0081 | P1_P2_P3_P4_Lumbar_Stellate_Sphenopalatine_VS_Coeliac | 1 | 1 | 1 | 1 | 1 | 0 | 1 | 1 |
| NUP43 | 0.0081 | P1_P3_Coeliac_VS_P2_P4_Lumbar_Stellate_Sphenopalatine | 1 | 0 | 1 | 0 | 0 | 1 | 0 | 0 |
| ZMPSTE24 | 0.0080 | P1_P2_P4_Lumbar_Coeliac_Stellate_Sphenopalatine_VS_P3 | 1 | 1 | 0 | 1 | 1 | 1 | 1 | 1 |
| 4931406C07I | 0.0079 | P2_P3_Lumbar_VS_P1_P4_Coeliac_Stellate_Sphenopalatine | 0 | 1 | 1 | 0 | 1 | 0 | 0 | 0 |
| CDK5RAP3 | 0.0076 | Lumbar_Sphenopalatine_VS_P1_P2_P3_P4_Coeliac_Stellate | 0 | 0 | 0 | 0 | 1 | 0 | 0 | 1 |
| PKIA | 0.0074 | P1_P2_P3_Lumbar_Coeliac_Stellate_Sphenopalatine_VS_P4 | 1 | 1 | 1 | 0 | 1 | 1 | 1 | 1 |
| ZFP979 | 0.0073 | P1_P2_P3_Lumbar_VS_P4_Coeliac_Stellate_Sphenopalatine | 1 | 1 | 1 | 0 | 1 | 0 | 0 | 0 |
| LZTS2 | 0.0072 | P2_P3_P4_Coeliac_Sphenopalatine_VS_P1_Lumbar_Stellate | 0 | 1 | 1 | 1 | 0 | 1 | 0 | 1 |
| CASP9 | 0.0072 | P1_P2_P4_Lumbar_Coeliac_Stellate_Sphenopalatine_VS_P3 | 1 | 1 | 0 | 1 | 1 | 1 | 1 | 1 |
| TSC1 | 0.0071 | P2_P3_P4_Coeliac_Sphenopalatine_VS_P1_Lumbar_Stellate | 0 | 1 | 1 | 1 | 0 | 1 | 0 | 1 |
| ADAM22 | 0.0071 | P2_P3_P4_Coeliac_Sphenopalatine_VS_P1_Lumbar_Stellate | 0 | 1 | 1 | 1 | 0 | 1 | 0 | 1 |
| GOPC | 0.0070 | P2_P3_Lumbar_VS_P1_P4_Coeliac_Stellate_Sphenopalatine | 0 | 1 | 1 | 0 | 1 | 0 | 0 | 0 |
| NKRF | 0.0068 | P1_P2_P3_Lumbar_VS_P4_Coeliac_Stellate_Sphenopalatine | 1 | 1 | 1 | 0 | 1 | 0 | 0 | 0 |
| DDHD1 | 0.0068 | P1_P2_P3_Lumbar_VS_P4_Coeliac_Stellate_Sphenopalatine | 1 | 1 | 1 | 0 | 1 | 0 | 0 | 0 |
| RPL7L1 | 0.0064 | P1_P4_Lumbar_Coeliac_Stellate_VS_P2_P3_Sphenopalatine | 1 | 0 | 0 | 1 | 1 | 1 | 1 | 0 |
| MRPL1 | 0.0057 | P1_P3_P4_Lumbar_Coeliac_Stellate_VS_P2_Sphenopalatine | 1 | 0 | 1 | 1 | 1 | 1 | 1 | 0 |

| | | | | | | | | | | |
|---|---|---|---|---|---|---|---|---|---|---|
| EHMT2 | 0.0052 | P1_P2_P3_Lumbar_Sphenopalatine_VS_P4_Coeliac_Stellate | 1 | 1 | 1 | 0 | 1 | 0 | 0 | 1 |
| SMIM20 | 0.0048 | P1_P2_P3_P4_Lumbar_Stellate_Sphenopalatine_VS_Coeliac | 1 | 1 | 1 | 1 | 1 | 0 | 1 | 1 |
| TSPYL2 | 0.0048 | P1_P3_P4_Coeliac_Sphenopalatine_VS_P2_Lumbar_Stellate | 1 | 0 | 1 | 1 | 0 | 1 | 0 | 1 |
| IER3 | 0.0047 | P1_P3_P4_Coeliac_Sphenopalatine_VS_P2_Lumbar_Stellate | 1 | 0 | 1 | 1 | 0 | 1 | 0 | 1 |
| SLC45A1 | 0.0045 | P3_P4_Lumbar_Sphenopalatine_VS_P1_P2_Coeliac_Stellate | 0 | 0 | 1 | 1 | 1 | 0 | 0 | 1 |
| PREP | 0.0043 | P1_P2_P4_Lumbar_Coeliac_Stellate_Sphenopalatine_VS_P3 | 1 | 1 | 0 | 1 | 1 | 1 | 1 | 1 |
| CISD3 | 0.0043 | P1_P2_P3_P4_Lumbar_Stellate_Sphenopalatine_VS_Coeliac | 1 | 1 | 1 | 1 | 1 | 0 | 1 | 1 |
| CACNB1 | 0.0043 | P1_P2_P3_Lumbar_Sphenopalatine_VS_P4_Coeliac_Stellate | 1 | 1 | 1 | 0 | 1 | 0 | 0 | 1 |
| LMBRD1 | 0.0039 | P2_P3_P4_Coeliac_Sphenopalatine_VS_P1_Lumbar_Stellate | 0 | 1 | 1 | 1 | 0 | 1 | 0 | 1 |
| CTPS2 | 0.0039 | P4_Lumbar_Coeliac_Stellate_VS_P1_P2_P3_Sphenopalatine | 0 | 0 | 0 | 1 | 1 | 1 | 1 | 0 |
| HMGN3 | 0.0036 | P1_P2_P3_P4_Lumbar_Stellate_Sphenopalatine_VS_Coeliac | 1 | 1 | 1 | 1 | 1 | 0 | 1 | 1 |
| GOLGA7B | 0.0035 | P1_P2_P3_P4_Lumbar_Stellate_Sphenopalatine_VS_Coeliac | 1 | 1 | 1 | 1 | 1 | 0 | 1 | 1 |
| DESI1 | 0.0034 | P1_P2_P3_Lumbar_Sphenopalatine_VS_P4_Coeliac_Stellate | 1 | 1 | 1 | 0 | 1 | 0 | 0 | 1 |
| ZBTB4 | 0.0032 | P2_P3_P4_Coeliac_Sphenopalatine_VS_P1_Lumbar_Stellate | 0 | 1 | 1 | 1 | 0 | 1 | 0 | 1 |
| CIPC | 0.0032 | P2_P4_Coeliac_Sphenopalatine_VS_P1_P3_Lumbar_Stellate | 0 | 1 | 0 | 1 | 0 | 1 | 0 | 1 |
| ELAVL1 | 0.0031 | P4_Lumbar_Coeliac_Stellate_VS_P1_P2_P3_Sphenopalatine | 0 | 0 | 0 | 1 | 1 | 1 | 1 | 0 |
| RFNG | 0.0030 | P1_P2_P3_Lumbar_Coeliac_Stellate_Sphenopalatine_VS_P4 | 1 | 1 | 1 | 0 | 1 | 1 | 1 | 1 |
| MICU1 | 0.0030 | P2_P4_Coeliac_Sphenopalatine_VS_P1_P3_Lumbar_Stellate | 0 | 1 | 0 | 1 | 0 | 1 | 0 | 1 |
| IQCB1 | 0.0029 | P1_P3_P4_Lumbar_Coeliac_Stellate_VS_P2_Sphenopalatine | 1 | 0 | 1 | 1 | 1 | 1 | 1 | 0 |
| RAB3D | 0.0029 | P1_P2_P3_P4_Lumbar_Stellate_Sphenopalatine_VS_Coeliac | 1 | 1 | 1 | 1 | 1 | 0 | 1 | 1 |
| GOLPH3 | 0.0029 | P1_P2_P3_P4_Lumbar_Stellate_Sphenopalatine_VS_Coeliac | 1 | 1 | 1 | 1 | 1 | 0 | 1 | 1 |
| E2F2 | 0.0028 | P1_P2_P3_Coeliac_Sphenopalatine_VS_P4_Lumbar_Stellate | 1 | 1 | 1 | 0 | 0 | 1 | 0 | 1 |
| ZZEF1 | 0.0028 | P2_P3_P4_Coeliac_Sphenopalatine_VS_P1_Lumbar_Stellate | 0 | 1 | 1 | 1 | 0 | 1 | 0 | 1 |
| AGFG1 | 0.0026 | P3_P4_Lumbar_Sphenopalatine_VS_P1_P2_Coeliac_Stellate | 0 | 0 | 1 | 1 | 1 | 0 | 0 | 1 |
| FAM91A1 | 0.0026 | P2_P3_P4_Coeliac_Sphenopalatine_VS_P1_Lumbar_Stellate | 0 | 1 | 1 | 1 | 0 | 1 | 0 | 1 |
| MOSPD3 | 0.0024 | P1_P3_VS_P2_P4_Lumbar_Coeliac_Stellate_Sphenopalatine | 1 | 0 | 1 | 0 | 0 | 0 | 0 | 0 |
| AEN | 0.0024 | P1_P3_VS_P2_P4_Lumbar_Coeliac_Stellate_Sphenopalatine | 1 | 0 | 1 | 0 | 0 | 0 | 0 | 0 |
| R3HDM2 | 0.0023 | P1_P3_P4_Coeliac_Sphenopalatine_VS_P2_Lumbar_Stellate | 1 | 0 | 1 | 1 | 0 | 1 | 0 | 1 |
| VWA5B2 | 0.0023 | P1_P2_P4_Lumbar_Coeliac_Stellate_Sphenopalatine_VS_P3 | 1 | 1 | 0 | 1 | 1 | 1 | 1 | 1 |
| PRXL2B | 0.0022 | P1_P2_P3_Lumbar_Sphenopalatine_VS_P4_Coeliac_Stellate | 1 | 1 | 1 | 0 | 1 | 0 | 0 | 1 |

| | | | | | | | | | | |
|---|---|---|---|---|---|---|---|---|---|---|
| EXOSC9 | 0.0022 | Lumbar_Sphenopalatine_VS_P1_P2_P3_P4_Coeliac_Stellate | 0 | 0 | 0 | 0 | 1 | 0 | 0 | 1 |
| TNFRSF21 | 0.0020 | P1_P2_Lumbar_Sphenopalatine_VS_P3_P4_Coeliac_Stellate | 1 | 1 | 0 | 0 | 1 | 0 | 0 | 1 |
| TMEM127 | 0.0019 | P1_P2_Lumbar_Sphenopalatine_VS_P3_P4_Coeliac_Stellate | 1 | 1 | 0 | 0 | 1 | 0 | 0 | 1 |
| ASB13 | 0.0018 | P3_Lumbar_Coeliac_Stellate_VS_P1_P2_P4_Sphenopalatine | 0 | 0 | 1 | 0 | 1 | 1 | 1 | 0 |
| NCOA1 | 0.0015 | P1_P3_P4_Coeliac_Sphenopalatine_VS_P2_Lumbar_Stellate | 1 | 0 | 1 | 1 | 0 | 1 | 0 | 1 |
| PGS1 | 0.0013 | P1_P2_P3_P4_Lumbar_Stellate_Sphenopalatine_VS_Coeliac | 1 | 1 | 1 | 1 | 1 | 0 | 1 | 1 |
| AKIRIN2 | 0.0012 | P2_P3_P4_Coeliac_Sphenopalatine_VS_P1_Lumbar_Stellate | 0 | 1 | 1 | 1 | 0 | 1 | 0 | 1 |
| NF2 | 0.0011 | P1_P2_P3_Lumbar_Sphenopalatine_VS_P4_Coeliac_Stellate | 1 | 1 | 1 | 0 | 1 | 0 | 0 | 1 |
| CHRNB2 | 0.0009 | P1_P2_P3_Lumbar_Sphenopalatine_VS_P4_Coeliac_Stellate | 1 | 1 | 1 | 0 | 1 | 0 | 0 | 1 |
| PTPN12 | 0.0008 | P2_P3_P4_Coeliac_Sphenopalatine_VS_P1_Lumbar_Stellate | 0 | 1 | 1 | 1 | 0 | 1 | 0 | 1 |
| AK3 | 0.0006 | P1_P2_P3_Lumbar_Coeliac_Stellate_Sphenopalatine_VS_P4 | 1 | 1 | 1 | 0 | 1 | 1 | 1 | 1 |
| TNRC6C | 0.0005 | P2_P3_P4_Coeliac_Sphenopalatine_VS_P1_Lumbar_Stellate | 0 | 1 | 1 | 1 | 0 | 1 | 0 | 1 |
| CCDC28B | 0.0005 | P1_P3_P4_Lumbar_Coeliac_Stellate_Sphenopalatine_VS_P2 | 1 | 0 | 1 | 1 | 1 | 1 | 1 | 1 |
| APOA2 | 0.0005 | P1_Sphenopalatine_VS_P2_P3_P4_Lumbar_Coeliac_Stellate | 1 | 0 | 0 | 0 | 0 | 0 | 0 | 1 |
| 2410131K14I | -0.0001 | P1_P2_P3_Lumbar_Sphenopalatine_VS_P4_Coeliac_Stellate | 1 | 1 | 1 | 0 | 1 | 0 | 0 | 1 |
| IPO8 | -0.0002 | P2_P3_P4_Coeliac_Sphenopalatine_VS_P1_Lumbar_Stellate | 0 | 1 | 1 | 1 | 0 | 1 | 0 | 1 |
| PRKAA2 | -0.0003 | Lumbar_Sphenopalatine_VS_P1_P2_P3_P4_Coeliac_Stellate | 0 | 0 | 0 | 0 | 1 | 0 | 0 | 1 |
| LHFPL4 | -0.0004 | P2_P3_P4_Coeliac_Sphenopalatine_VS_P1_Lumbar_Stellate | 0 | 1 | 1 | 1 | 0 | 1 | 0 | 1 |
| TRIM67 | -0.0005 | P1_P2_P3_Lumbar_Coeliac_Stellate_VS_P4_Sphenopalatine | 1 | 1 | 1 | 0 | 1 | 1 | 1 | 0 |
| CREB3 | -0.0008 | P1_P3_P4_Lumbar_Coeliac_Stellate_Sphenopalatine_VS_P2 | 1 | 0 | 1 | 1 | 1 | 1 | 1 | 1 |
| ZFP560 | -0.0009 | P1_P2_P3_P4_Lumbar_Stellate_Sphenopalatine_VS_Coeliac | 1 | 1 | 1 | 1 | 1 | 0 | 1 | 1 |
| PTPN11 | -0.0009 | P2_P3_P4_Coeliac_Sphenopalatine_VS_P1_Lumbar_Stellate | 0 | 1 | 1 | 1 | 0 | 1 | 0 | 1 |
| SYNJ1 | -0.0010 | P2_Sphenopalatine_VS_P1_P3_P4_Lumbar_Coeliac_Stellate | 0 | 1 | 0 | 0 | 0 | 0 | 0 | 1 |
| RTKN | -0.0010 | P1_P2_P4_Lumbar_Coeliac_Stellate_Sphenopalatine_VS_P3 | 1 | 1 | 0 | 1 | 1 | 1 | 1 | 1 |
| CBFA2T2 | -0.0012 | P2_P3_P4_Coeliac_Sphenopalatine_VS_P1_Lumbar_Stellate | 0 | 1 | 1 | 1 | 0 | 1 | 0 | 1 |
| PDIA4 | -0.0012 | P1_P3_P4_Lumbar_Coeliac_Stellate_VS_P2_Sphenopalatine | 1 | 0 | 1 | 1 | 1 | 1 | 1 | 0 |
| 2900041M22 | -0.0013 | P3_Sphenopalatine_VS_P1_P2_P4_Lumbar_Coeliac_Stellate | 0 | 0 | 1 | 0 | 0 | 0 | 0 | 1 |
| VSIG10L | -0.0015 | Lumbar_Sphenopalatine_VS_P1_P2_P3_P4_Coeliac_Stellate | 0 | 0 | 0 | 0 | 1 | 0 | 0 | 1 |
| ALDH4A1 | -0.0016 | P2_P3_VS_P1_P4_Lumbar_Coeliac_Stellate_Sphenopalatine | 0 | 1 | 1 | 0 | 0 | 0 | 0 | 0 |
| MINPP1 | -0.0016 | P1_P2_P3_Lumbar_Coeliac_Stellate_Sphenopalatine_VS_P4 | 1 | 1 | 1 | 0 | 1 | 1 | 1 | 1 |

| | | | | | | | | | | |
|---|---|---|---|---|---|---|---|---|---|---|
| NT5DC3 | -0.002 | P1_P2_P3_Lumbar_Sphenopalatine_VS_P4_Coeliac_Stellate | 1 | 1 | 1 | 0 | 1 | 0 | 0 | 1 |
| SAE1 | -0.002 | P1_P3_P4_Lumbar_Coeliac_Stellate_Sphenopalatine_VS_P2 | 1 | 0 | 1 | 1 | 1 | 1 | 1 | 1 |
| TNPO1 | -0.002 | P1_P2_P4_Lumbar_Coeliac_Stellate_Sphenopalatine_VS_P3 | 1 | 1 | 0 | 1 | 1 | 1 | 1 | 1 |
| DNAJC10 | -0.002 | P3_P4_Lumbar_Sphenopalatine_VS_P1_P2_Coeliac_Stellate | 0 | 0 | 1 | 1 | 1 | 0 | 0 | 1 |
| RBM22 | -0.002 | P1_P2_P3_Lumbar_Coeliac_Stellate_VS_P4_Sphenopalatine | 1 | 1 | 1 | 0 | 1 | 1 | 1 | 0 |
| DOP1B | -0.002 | P3_P4_Lumbar_Sphenopalatine_VS_P1_P2_Coeliac_Stellate | 0 | 0 | 1 | 1 | 1 | 0 | 0 | 1 |
| BTBD6 | -0.002 | P1_P3_P4_Coeliac_Sphenopalatine_VS_P2_Lumbar_Stellate | 1 | 0 | 1 | 1 | 0 | 1 | 0 | 1 |
| MTHFSL | -0.002 | P3_P4_Lumbar_Sphenopalatine_VS_P1_P2_Coeliac_Stellate | 0 | 0 | 1 | 1 | 1 | 0 | 0 | 1 |
| ANKRD24 | -0.002 | P3_P4_Lumbar_Sphenopalatine_VS_P1_P2_Coeliac_Stellate | 0 | 0 | 1 | 1 | 1 | 0 | 0 | 1 |
| CCDC59 | -0.002 | P1_P3_P4_Lumbar_Coeliac_Stellate_VS_P2_Sphenopalatine | 1 | 0 | 1 | 1 | 1 | 1 | 1 | 0 |
| ANKRD11 | -0.003 | P2_P3_P4_Coeliac_Sphenopalatine_VS_P1_Lumbar_Stellate | 0 | 1 | 1 | 1 | 0 | 1 | 0 | 1 |
| PRAF2 | -0.003 | P1_P2_P3_P4_Lumbar_Stellate_Sphenopalatine_VS_Coeliac | 1 | 1 | 1 | 1 | 1 | 0 | 1 | 1 |
| CAMK1 | -0.003 | P1_P2_Lumbar_Sphenopalatine_VS_P3_P4_Coeliac_Stellate | 1 | 1 | 0 | 0 | 1 | 0 | 0 | 1 |
| LPCAT1 | -0.003 | P3_P4_Lumbar_Sphenopalatine_VS_P1_P2_Coeliac_Stellate | 0 | 0 | 1 | 1 | 1 | 0 | 0 | 1 |
| HMBS | -0.003 | P1_P3_P4_Lumbar_Coeliac_Stellate_VS_P2_Sphenopalatine | 1 | 0 | 1 | 1 | 1 | 1 | 1 | 0 |
| CAMK2B | -0.003 | P3_P4_Sphenopalatine_VS_P1_P2_Lumbar_Coeliac_Stellate | 0 | 0 | 1 | 1 | 0 | 0 | 0 | 1 |
| PRNP | -0.003 | P1_P2_P4_Lumbar_Coeliac_Stellate_Sphenopalatine_VS_P3 | 1 | 1 | 0 | 1 | 1 | 1 | 1 | 1 |
| MMACHC | -0.003 | P1_P3_VS_P2_P4_Lumbar_Coeliac_Stellate_Sphenopalatine | 1 | 0 | 1 | 0 | 0 | 0 | 0 | 0 |
| ALKBH6 | -0.003 | P1_P3_P4_Lumbar_Coeliac_Stellate_Sphenopalatine_VS_P2 | 1 | 0 | 1 | 1 | 1 | 1 | 1 | 1 |
| CRYL1 | -0.003 | P4_Lumbar_Coeliac_Stellate_VS_P1_P2_P3_Sphenopalatine | 0 | 0 | 0 | 1 | 1 | 1 | 1 | 0 |
| FAM98C | -0.003 | P1_P2_P3_P4_Lumbar_Stellate_Sphenopalatine_VS_Coeliac | 1 | 1 | 1 | 1 | 1 | 0 | 1 | 1 |
| B430010I23 | -0.003 | P2_Sphenopalatine_VS_P1_P3_P4_Lumbar_Coeliac_Stellate | 0 | 1 | 0 | 0 | 0 | 0 | 0 | 1 |
| CUX1 | -0.003 | P1_P2_P3_Coeliac_VS_P4_Lumbar_Stellate_Sphenopalatine | 1 | 1 | 1 | 0 | 0 | 1 | 0 | 0 |
| GM34006 | -0.004 | P1_P2_P4_Lumbar_Coeliac_Stellate_VS_P3_Sphenopalatine | 1 | 1 | 0 | 1 | 1 | 1 | 1 | 0 |
| PUSL1 | -0.004 | P1_P3_P4_Lumbar_Coeliac_Stellate_Sphenopalatine_VS_P2 | 1 | 0 | 1 | 1 | 1 | 1 | 1 | 1 |
| MBD6 | -0.004 | P2_P3_P4_Coeliac_Sphenopalatine_VS_P1_Lumbar_Stellate | 0 | 1 | 1 | 1 | 0 | 1 | 0 | 1 |
| TMEM176B | -0.004 | P1_P2_P3_Sphenopalatine_VS_P4_Lumbar_Coeliac_Stellate | 1 | 1 | 1 | 0 | 0 | 0 | 0 | 1 |
| JAK1 | -0.004 | P1_P2_P3_Lumbar_Sphenopalatine_VS_P4_Coeliac_Stellate | 1 | 1 | 1 | 0 | 1 | 0 | 0 | 1 |
| UBASH3B | -0.004 | P2_Sphenopalatine_VS_P1_P3_P4_Lumbar_Coeliac_Stellate | 0 | 1 | 0 | 0 | 0 | 0 | 0 | 1 |
| ZCCHC3 | -0.004 | P1_P3_P4_Coeliac_Sphenopalatine_VS_P2_Lumbar_Stellate | 1 | 0 | 1 | 1 | 0 | 1 | 0 | 1 |

| | | | | | | | | | | |
|---|---|---|---|---|---|---|---|---|---|---|
| ZFP9 | -0.0045 | P3_P4_Lumbar_Coeliac_Stellate_VS_P1_P2_Sphenopalatine | 0 | 0 | 1 | 1 | 1 | 1 | 1 | 0 |
| LAGE3 | -0.0046 | P3_P4_Sphenopalatine_VS_P1_P2_Lumbar_Coeliac_Stellate | 0 | 0 | 1 | 1 | 0 | 0 | 0 | 1 |
| OGFOD3 | -0.0046 | P1_P3_P4_Lumbar_Coeliac_Stellate_Sphenopalatine_VS_P2 | 1 | 0 | 1 | 1 | 1 | 1 | 1 | 1 |
| 2310039H08I | -0.0048 | P1_P3_P4_Lumbar_Coeliac_Stellate_Sphenopalatine_VS_P2 | 1 | 0 | 1 | 1 | 1 | 1 | 1 | 1 |
| TBC1D16 | -0.0049 | P1_P2_P3_Lumbar_Coeliac_Stellate_Sphenopalatine_VS_P4 | 1 | 1 | 1 | 0 | 1 | 1 | 1 | 1 |
| NAT14 | -0.0049 | P1_P3_P4_Coeliac_Sphenopalatine_VS_P2_Lumbar_Stellate | 1 | 0 | 1 | 1 | 0 | 1 | 0 | 1 |
| PIM2 | -0.0053 | P3_P4_Sphenopalatine_VS_P1_P2_Lumbar_Coeliac_Stellate | 0 | 0 | 1 | 1 | 0 | 0 | 0 | 1 |
| RXYLT1 | -0.0054 | P3_Lumbar_Sphenopalatine_VS_P1_P2_P4_Coeliac_Stellate | 0 | 0 | 1 | 0 | 1 | 0 | 0 | 1 |
| GLE1 | -0.0054 | P1_P2_P3_Lumbar_Sphenopalatine_VS_P4_Coeliac_Stellate | 1 | 1 | 1 | 0 | 1 | 0 | 0 | 1 |
| ITPK1 | -0.0058 | Lumbar_Sphenopalatine_VS_P1_P2_P3_P4_Coeliac_Stellate | 0 | 0 | 0 | 0 | 1 | 0 | 0 | 1 |
| RUSC2 | -0.0060 | P4_Coeliac_Sphenopalatine_VS_P1_P2_P3_Lumbar_Stellate | 0 | 0 | 0 | 1 | 0 | 1 | 0 | 1 |
| FBXL15 | -0.0061 | P3_P4_Lumbar_Sphenopalatine_VS_P1_P2_Coeliac_Stellate | 0 | 0 | 1 | 1 | 1 | 0 | 0 | 1 |
| NRSN2 | -0.0061 | P3_P4_Sphenopalatine_VS_P1_P2_Lumbar_Coeliac_Stellate | 0 | 0 | 1 | 1 | 0 | 0 | 0 | 1 |
| SLC52A2 | -0.0061 | P1_P2_P3_Lumbar_Coeliac_Stellate_Sphenopalatine_VS_P4 | 1 | 1 | 1 | 0 | 1 | 1 | 1 | 1 |
| SCARB2 | -0.0062 | Lumbar_Sphenopalatine_VS_P1_P2_P3_P4_Coeliac_Stellate | 0 | 0 | 0 | 0 | 1 | 0 | 0 | 1 |
| TRAK1 | -0.0063 | P1_P2_P3_Lumbar_Sphenopalatine_VS_P4_Coeliac_Stellate | 1 | 1 | 1 | 0 | 1 | 0 | 0 | 1 |
| GADD45GIP1 | -0.0063 | P3_P4_Sphenopalatine_VS_P1_P2_Lumbar_Coeliac_Stellate | 0 | 0 | 1 | 1 | 0 | 0 | 0 | 1 |
| ZFP575 | -0.0067 | P3_P4_Sphenopalatine_VS_P1_P2_Lumbar_Coeliac_Stellate | 0 | 0 | 1 | 1 | 0 | 0 | 0 | 1 |
| CEBPZOS | -0.0071 | P3_P4_Sphenopalatine_VS_P1_P2_Lumbar_Coeliac_Stellate | 0 | 0 | 1 | 1 | 0 | 0 | 0 | 1 |
| PDXDC1 | -0.0072 | P2_P3_VS_P1_P4_Lumbar_Coeliac_Stellate_Sphenopalatine | 0 | 1 | 1 | 0 | 0 | 0 | 0 | 0 |
| FAM98A | -0.0072 | P2_P3_VS_P1_P4_Lumbar_Coeliac_Stellate_Sphenopalatine | 0 | 1 | 1 | 0 | 0 | 0 | 0 | 0 |
| PIAS2 | -0.0074 | P1_P2_P3_Coeliac_VS_P4_Lumbar_Stellate_Sphenopalatine | 1 | 1 | 1 | 0 | 0 | 1 | 0 | 0 |
| MESD | -0.0075 | P3_P4_Lumbar_Sphenopalatine_VS_P1_P2_Coeliac_Stellate | 0 | 0 | 1 | 1 | 1 | 0 | 0 | 1 |
| LARP7 | -0.0077 | P1_P3_Coeliac_VS_P2_P4_Lumbar_Stellate_Sphenopalatine | 1 | 0 | 1 | 0 | 0 | 1 | 0 | 0 |
| PAQR4 | -0.0077 | P1_P2_P3_Lumbar_Coeliac_Stellate_Sphenopalatine_VS_P4 | 1 | 1 | 1 | 0 | 1 | 1 | 1 | 1 |
| HMCES | -0.0079 | P1_P4_Coeliac_Sphenopalatine_VS_P2_P3_Lumbar_Stellate | 1 | 0 | 0 | 1 | 0 | 1 | 0 | 1 |
| MAOA | -0.0081 | P2_P3_P4_Lumbar_Coeliac_Stellate_VS_P1_Sphenopalatine | 0 | 1 | 1 | 1 | 1 | 1 | 1 | 0 |
| DBNDD2 | -0.0085 | P1_P2_P3_P4_Lumbar_Stellate_Sphenopalatine_VS_Coeliac | 1 | 1 | 1 | 1 | 1 | 0 | 1 | 1 |
| ARHGEF12 | -0.0086 | P2_Lumbar_Sphenopalatine_VS_P1_P3_P4_Coeliac_Stellate | 0 | 1 | 0 | 0 | 1 | 0 | 0 | 1 |
| TMEM192 | -0.0091 | P1_P2_P3_P4_Lumbar_Stellate_Sphenopalatine_VS_Coeliac | 1 | 1 | 1 | 1 | 1 | 0 | 1 | 1 |

| | | | | | | | | | | |
|---|---|---|---|---|---|---|---|---|---|---|
| SNRNP35 | -0.0095 | P1_P3_P4_Lumbar_Coeliac_Stellate_Sphenopalatine_VS_P2 | 1 | 0 | 1 | 1 | 1 | 1 | 1 | 1 |
| TRNT1 | -0.0098 | P3_P4_Lumbar_Coeliac_Stellate_VS_P1_P2_Sphenopalatine | 0 | 0 | 1 | 1 | 1 | 1 | 1 | 0 |
| NCDN | -0.0098 | P2_Sphenopalatine_VS_P1_P3_P4_Lumbar_Coeliac_Stellate | 0 | 1 | 0 | 0 | 0 | 0 | 0 | 1 |
| ARF3 | -0.0102 | P1_P2_P3_P4_Lumbar_Stellate_Sphenopalatine_VS_Coeliac | 1 | 1 | 1 | 1 | 1 | 0 | 1 | 1 |
| ZBTB18 | -0.0103 | P4_Coeliac_Sphenopalatine_VS_P1_P2_P3_Lumbar_Stellate | 0 | 0 | 0 | 1 | 0 | 1 | 0 | 1 |
| HMG20B | -0.0104 | P1_P2_P4_Lumbar_Coeliac_Stellate_VS_P3_Sphenopalatine | 1 | 1 | 0 | 1 | 1 | 1 | 1 | 0 |
| GSK3A | -0.0106 | P1_P2_P3_Lumbar_Sphenopalatine_VS_P4_Coeliac_Stellate | 1 | 1 | 1 | 0 | 1 | 0 | 0 | 1 |
| LSM11 | -0.0106 | P1_P3_P4_Lumbar_Coeliac_Stellate_Sphenopalatine_VS_P2 | 1 | 0 | 1 | 1 | 1 | 1 | 1 | 1 |
| PPIL1 | -0.0106 | P1_P3_P4_Lumbar_Coeliac_Stellate_Sphenopalatine_VS_P2 | 1 | 0 | 1 | 1 | 1 | 1 | 1 | 1 |
| FNBP1L | -0.0111 | Lumbar_Sphenopalatine_VS_P1_P2_P3_P4_Coeliac_Stellate | 0 | 0 | 0 | 0 | 1 | 0 | 0 | 1 |
| HEBP1 | -0.0114 | P4_Lumbar_Stellate_VS_P1_P2_P3_Coeliac_Sphenopalatine | 0 | 0 | 0 | 1 | 1 | 0 | 1 | 0 |
| POFUT2 | -0.0114 | P3_P4_Sphenopalatine_VS_P1_P2_Lumbar_Coeliac_Stellate | 0 | 0 | 1 | 1 | 0 | 0 | 0 | 1 |
| DPP9 | -0.0115 | P1_P4_Coeliac_Sphenopalatine_VS_P2_P3_Lumbar_Stellate | 1 | 0 | 0 | 1 | 0 | 1 | 0 | 1 |
| PAXX | -0.0117 | P3_P4_Sphenopalatine_VS_P1_P2_Lumbar_Coeliac_Stellate | 0 | 0 | 1 | 1 | 0 | 0 | 0 | 1 |
| RPS19BP1 | -0.0117 | P1_P3_P4_Lumbar_Coeliac_Stellate_Sphenopalatine_VS_P2 | 1 | 0 | 1 | 1 | 1 | 1 | 1 | 1 |
| ZNRD1 | -0.0119 | P3_P4_Lumbar_Sphenopalatine_VS_P1_P2_Coeliac_Stellate | 0 | 0 | 1 | 1 | 1 | 0 | 0 | 1 |
| TIMM8A1 | -0.012 | P1_P2_P3_Lumbar_Coeliac_Stellate_Sphenopalatine_VS_P4 | 1 | 1 | 1 | 0 | 1 | 1 | 1 | 1 |
| LYRM4 | -0.0122 | P3_P4_Sphenopalatine_VS_P1_P2_Lumbar_Coeliac_Stellate | 0 | 0 | 1 | 1 | 0 | 0 | 0 | 1 |
| EPDR1 | -0.0122 | P1_P2_P3_P4_Lumbar_Stellate_Sphenopalatine_VS_Coeliac | 1 | 1 | 1 | 1 | 1 | 0 | 1 | 1 |
| NAA40 | -0.0122 | P1_P2_P3_Lumbar_Coeliac_Stellate_Sphenopalatine_VS_P4 | 1 | 1 | 1 | 0 | 1 | 1 | 1 | 1 |
| RFC4 | -0.0123 | P1_P4_Lumbar_Coeliac_Stellate_VS_P2_P3_Sphenopalatine | 1 | 0 | 0 | 1 | 1 | 1 | 1 | 0 |
| MARCH2 | -0.0125 | P1_P2_P3_P4_Lumbar_Stellate_Sphenopalatine_VS_Coeliac | 1 | 1 | 1 | 1 | 1 | 0 | 1 | 1 |
| ARFGEF1 | -0.0126 | P4_Coeliac_Sphenopalatine_VS_P1_P2_P3_Lumbar_Stellate | 0 | 0 | 0 | 1 | 0 | 1 | 0 | 1 |
| NAA60 | -0.0128 | P3_Sphenopalatine_VS_P1_P2_P4_Lumbar_Coeliac_Stellate | 0 | 0 | 1 | 0 | 0 | 0 | 0 | 1 |
| CHMP6 | -0.0129 | P3_Sphenopalatine_VS_P1_P2_P4_Lumbar_Coeliac_Stellate | 0 | 0 | 1 | 0 | 0 | 0 | 0 | 1 |
| APEX1 | -0.0129 | P1_P3_P4_Lumbar_Coeliac_Stellate_Sphenopalatine_VS_P2 | 1 | 0 | 1 | 1 | 1 | 1 | 1 | 1 |
| DNAJC9 | -0.0131 | P1_P3_P4_Coeliac_Sphenopalatine_VS_P2_Lumbar_Stellate | 1 | 0 | 1 | 1 | 0 | 1 | 0 | 1 |
| NUDT16L1 | -0.0132 | P3_P4_Sphenopalatine_VS_P1_P2_Lumbar_Coeliac_Stellate | 0 | 0 | 1 | 1 | 0 | 0 | 0 | 1 |
| DOK4 | -0.0133 | P1_P2_P4_Lumbar_Coeliac_Stellate_VS_P3_Sphenopalatine | 1 | 1 | 0 | 1 | 1 | 1 | 1 | 0 |
| PHF24 | -0.0133 | P1_P2_P3_Lumbar_Sphenopalatine_VS_P4_Coeliac_Stellate | 1 | 1 | 1 | 0 | 1 | 0 | 0 | 1 |

| | | | | | | | | | | |
|---|---|---|---|---|---|---|---|---|---|---|
| MAD2L2 | -0.0135 | P1_P2_P3_Lumbar_Coeliac_Stellate_Sphenopalatine_VS_P4 | 1 | 1 | 1 | 0 | 1 | 1 | 1 | 1 |
| SLC39A13 | -0.0136 | P2_P3_P4_Lumbar_Coeliac_Stellate_VS_P1_Sphenopalatine | 0 | 1 | 1 | 1 | 1 | 1 | 1 | 0 |
| CRTC1 | -0.0136 | P1_P2_P3_Lumbar_Sphenopalatine_VS_P4_Coeliac_Stellate | 1 | 1 | 1 | 0 | 1 | 0 | 0 | 1 |
| TMEM29 | -0.0139 | P1_P2_P3_P4_Lumbar_Stellate_Sphenopalatine_VS_Coeliac | 1 | 1 | 1 | 1 | 1 | 0 | 1 | 1 |
| PHACTR1 | -0.014 | P1_P3_P4_Coeliac_Sphenopalatine_VS_P2_Lumbar_Stellate | 1 | 0 | 1 | 1 | 0 | 1 | 0 | 1 |
| NECTIN2 | -0.014 | P1_P3_P4_Lumbar_Coeliac_Stellate_Sphenopalatine_VS_P2 | 1 | 0 | 1 | 1 | 1 | 1 | 1 | 1 |
| RCAN1 | -0.0144 | P1_P2_P4_Lumbar_Coeliac_Stellate_Sphenopalatine_VS_P3 | 1 | 1 | 0 | 1 | 1 | 1 | 1 | 1 |
| SNHG20 | -0.0145 | P3_P4_Lumbar_Sphenopalatine_VS_P1_P2_Coeliac_Stellate | 0 | 0 | 1 | 1 | 1 | 0 | 0 | 1 |
| TRIM41 | -0.0146 | P3_P4_Lumbar_Sphenopalatine_VS_P1_P2_Coeliac_Stellate | 0 | 0 | 1 | 1 | 1 | 0 | 0 | 1 |
| FBRSL1 | -0.0147 | P2_P4_Lumbar_Coeliac_Stellate_VS_P1_P3_Sphenopalatine | 0 | 1 | 0 | 1 | 1 | 1 | 1 | 0 |
| MTMR4 | -0.0147 | P2_P3_Sphenopalatine_VS_P1_P4_Lumbar_Coeliac_Stellate | 0 | 1 | 1 | 0 | 0 | 0 | 0 | 1 |
| NSF | -0.0148 | P2_P3_Lumbar_Sphenopalatine_VS_P1_P4_Coeliac_Stellate | 0 | 1 | 1 | 0 | 1 | 0 | 0 | 1 |
| TM7SF2 | -0.015 | P3_P4_Lumbar_Sphenopalatine_VS_P1_P2_Coeliac_Stellate | 0 | 0 | 1 | 1 | 1 | 0 | 0 | 1 |
| UCHL3 | -0.015 | P1_P3_P4_Lumbar_Coeliac_Stellate_Sphenopalatine_VS_P2 | 1 | 0 | 1 | 1 | 1 | 1 | 1 | 1 |
| COA7 | -0.0151 | P1_P3_P4_Lumbar_Coeliac_Stellate_Sphenopalatine_VS_P2 | 1 | 0 | 1 | 1 | 1 | 1 | 1 | 1 |
| SNHG12 | -0.0153 | Lumbar_Sphenopalatine_VS_P1_P2_P3_P4_Coeliac_Stellate | 0 | 0 | 0 | 0 | 1 | 0 | 0 | 1 |
| DTD1 | -0.0154 | P3_P4_Lumbar_Coeliac_Stellate_VS_P1_P2_Sphenopalatine | 0 | 0 | 1 | 1 | 1 | 1 | 1 | 0 |
| HSCB | -0.0155 | P1_P2_P3_Lumbar_Coeliac_Stellate_Sphenopalatine_VS_P4 | 1 | 1 | 1 | 0 | 1 | 1 | 1 | 1 |
| TMEM223 | -0.0156 | P1_P3_P4_Lumbar_Coeliac_Stellate_Sphenopalatine_VS_P2 | 1 | 0 | 1 | 1 | 1 | 1 | 1 | 1 |
| CBARP | -0.0157 | P1_P2_P3_P4_Lumbar_Stellate_Sphenopalatine_VS_Coeliac | 1 | 1 | 1 | 1 | 1 | 0 | 1 | 1 |
| PEA15A | -0.0158 | P1_P2_P3_P4_Lumbar_Stellate_Sphenopalatine_VS_Coeliac | 1 | 1 | 1 | 1 | 1 | 0 | 1 | 1 |
| PTPN1 | -0.0158 | P1_P2_P3_Lumbar_Coeliac_Stellate_Sphenopalatine_VS_P4 | 1 | 1 | 1 | 0 | 1 | 1 | 1 | 1 |
| PSEN2 | -0.0159 | P1_P4_Coeliac_Sphenopalatine_VS_P2_P3_Lumbar_Stellate | 1 | 0 | 0 | 1 | 0 | 1 | 0 | 1 |
| PTDSS1 | -0.016 | P1_P2_P3_P4_Lumbar_Stellate_Sphenopalatine_VS_Coeliac | 1 | 1 | 1 | 1 | 1 | 0 | 1 | 1 |
| HMGXB3 | -0.016 | P1_P2_P4_Lumbar_Coeliac_Stellate_Sphenopalatine_VS_P3 | 1 | 1 | 0 | 1 | 1 | 1 | 1 | 1 |
| SLC25A10 | -0.0162 | P1_P3_VS_P2_P4_Lumbar_Coeliac_Stellate_Sphenopalatine | 1 | 0 | 1 | 0 | 0 | 0 | 0 | 0 |
| ARXES1 | -0.0162 | P1_P3_P4_Lumbar_Coeliac_Stellate_Sphenopalatine_VS_P2 | 1 | 0 | 1 | 1 | 1 | 1 | 1 | 1 |
| PDXP | -0.0163 | P1_P3_P4_Coeliac_Sphenopalatine_VS_P2_Lumbar_Stellate | 1 | 0 | 1 | 1 | 0 | 1 | 0 | 1 |
| TMEM131 | -0.0164 | Lumbar_Sphenopalatine_VS_P1_P2_P3_P4_Coeliac_Stellate | 0 | 0 | 0 | 0 | 1 | 0 | 0 | 1 |
| DNAJB2 | -0.0165 | P3_P4_Lumbar_Sphenopalatine_VS_P1_P2_Coeliac_Stellate | 0 | 0 | 1 | 1 | 1 | 0 | 0 | 1 |

| | | | | | | | | | | |
|---|---|---|---|---|---|---|---|---|---|---|
| PEX2 | -0.0166 | P1_P3_Lumbar_Coeliac_Stellate_VS_P2_P4_Sphenopalatine | 1 | 0 | 1 | 0 | 1 | 1 | 1 | 0 |
| RNASEH2A | -0.0166 | P1_P4_Lumbar_Coeliac_Stellate_VS_P2_P3_Sphenopalatine | 1 | 0 | 0 | 1 | 1 | 1 | 1 | 0 |
| DYRK2 | -0.0166 | P3_P4_Lumbar_Sphenopalatine_VS_P1_P2_Coeliac_Stellate | 0 | 0 | 1 | 1 | 1 | 0 | 0 | 1 |
| VTA1 | -0.0166 | P1_P3_P4_Lumbar_Coeliac_Stellate_Sphenopalatine_VS_P2 | 1 | 0 | 1 | 1 | 1 | 1 | 1 | 1 |
| TMEM199 | -0.0167 | P1_P3_P4_Lumbar_Coeliac_Stellate_Sphenopalatine_VS_P2 | 1 | 0 | 1 | 1 | 1 | 1 | 1 | 1 |
| SLC36A1 | -0.0169 | P1_P2_P3_Sphenopalatine_VS_P4_Lumbar_Coeliac_Stellate | 1 | 1 | 1 | 0 | 0 | 0 | 0 | 1 |
| TMEM179 | -0.0171 | P1_P2_P3_Sphenopalatine_VS_P4_Lumbar_Coeliac_Stellate | 1 | 1 | 1 | 0 | 0 | 0 | 0 | 1 |
| TLNRD1 | -0.0171 | P1_P2_P3_Lumbar_Coeliac_Stellate_Sphenopalatine_VS_P4 | 1 | 1 | 1 | 0 | 1 | 1 | 1 | 1 |
| DDOST | -0.0172 | P1_P3_P4_Lumbar_Coeliac_Stellate_Sphenopalatine_VS_P2 | 1 | 0 | 1 | 1 | 1 | 1 | 1 | 1 |
| IRAK1BP1 | -0.0174 | P3_P4_Lumbar_Sphenopalatine_VS_P1_P2_Coeliac_Stellate | 0 | 0 | 1 | 1 | 1 | 0 | 0 | 1 |
| TAMM41 | -0.0175 | P1_P3_Lumbar_Coeliac_Stellate_VS_P2_P4_Sphenopalatine | 1 | 0 | 1 | 0 | 1 | 1 | 1 | 0 |
| MPV17L2 | -0.0179 | P1_P3_VS_P2_P4_Lumbar_Coeliac_Stellate_Sphenopalatine | 1 | 0 | 1 | 0 | 0 | 0 | 0 | 0 |
| VOPP1 | -0.0179 | P1_P2_P3_Lumbar_Coeliac_Stellate_VS_P4_Sphenopalatine | 1 | 1 | 1 | 0 | 1 | 1 | 1 | 0 |
| CDIP1 | -0.0179 | P1_P2_P3_P4_Lumbar_Stellate_Sphenopalatine_VS_Coeliac | 1 | 1 | 1 | 1 | 1 | 0 | 1 | 1 |
| DNAJC18 | -0.018 | P1_P2_P3_Lumbar_Sphenopalatine_VS_P4_Coeliac_Stellate | 1 | 1 | 1 | 0 | 1 | 0 | 0 | 1 |
| PPP2R3C | -0.0181 | P1_P3_P4_Lumbar_Coeliac_Stellate_Sphenopalatine_VS_P2 | 1 | 0 | 1 | 1 | 1 | 1 | 1 | 1 |
| TRUB2 | -0.0182 | P1_P2_Lumbar_Sphenopalatine_VS_P3_P4_Coeliac_Stellate | 1 | 1 | 0 | 0 | 1 | 0 | 0 | 1 |
| ACTN4 | -0.0184 | P3_P4_Lumbar_Sphenopalatine_VS_P1_P2_Coeliac_Stellate | 0 | 0 | 1 | 1 | 1 | 0 | 0 | 1 |
| FAAP100 | -0.0185 | P1_P4_Coeliac_Sphenopalatine_VS_P2_P3_Lumbar_Stellate | 1 | 0 | 0 | 1 | 0 | 1 | 0 | 1 |
| TGFB1I1 | -0.0185 | P1_P2_P3_P4_Lumbar_Stellate_Sphenopalatine_VS_Coeliac | 1 | 1 | 1 | 1 | 1 | 0 | 1 | 1 |
| PLA2G6 | -0.0186 | P1_P2_P3_Lumbar_Sphenopalatine_VS_P4_Coeliac_Stellate | 1 | 1 | 1 | 0 | 1 | 0 | 0 | 1 |
| PTGES2 | -0.0187 | P1_Lumbar_Sphenopalatine_VS_P2_P3_P4_Coeliac_Stellate | 1 | 0 | 0 | 0 | 1 | 0 | 0 | 1 |
| DMTN | -0.0189 | P1_P2_P3_P4_Lumbar_Stellate_Sphenopalatine_VS_Coeliac | 1 | 1 | 1 | 1 | 1 | 0 | 1 | 1 |
| PRPSAP2 | -0.0191 | P2_P3_P4_Lumbar_Coeliac_Stellate_VS_P1_Sphenopalatine | 0 | 1 | 1 | 1 | 1 | 1 | 1 | 0 |
| TMEM191C | -0.0192 | P1_P3_P4_Lumbar_Coeliac_Stellate_Sphenopalatine_VS_P2 | 1 | 0 | 1 | 1 | 1 | 1 | 1 | 1 |
| FADS3 | -0.0193 | P1_P2_Lumbar_Sphenopalatine_VS_P3_P4_Coeliac_Stellate | 1 | 1 | 0 | 0 | 1 | 0 | 0 | 1 |
| CAMK2N2 | -0.0193 | P1_P2_P3_Coeliac_Sphenopalatine_VS_P4_Lumbar_Stellate | 1 | 1 | 1 | 0 | 0 | 1 | 0 | 1 |
| PANX1 | -0.0194 | P3_Lumbar_Sphenopalatine_VS_P1_P2_P4_Coeliac_Stellate | 0 | 0 | 1 | 0 | 1 | 0 | 0 | 1 |
| ANG | -0.0196 | Lumbar_Sphenopalatine_VS_P1_P2_P3_P4_Coeliac_Stellate | 0 | 0 | 0 | 0 | 1 | 0 | 0 | 1 |
| DIP2A | -0.0196 | P1_P2_P3_Lumbar_Sphenopalatine_VS_P4_Coeliac_Stellate | 1 | 1 | 1 | 0 | 1 | 0 | 0 | 1 |

| | | | | | | | | | | |
|---|---|---|---|---|---|---|---|---|---|---|
| SMU1 | -0.0199 | P1_P4_Lumbar_Coeliac_Stellate_VS_P2_P3_Sphenopalatine | 1 | 0 | 0 | 1 | 1 | 1 | 1 | 0 |
| RPF2 | -0.0203 | P1_Lumbar_Coeliac_Stellate_VS_P2_P3_P4_Sphenopalatine | 1 | 0 | 0 | 0 | 1 | 1 | 1 | 0 |
| ECE2 | -0.0203 | P1_P3_P4_Coeliac_Sphenopalatine_VS_P2_Lumbar_Stellate | 1 | 0 | 1 | 1 | 0 | 1 | 0 | 1 |
| GRK3 | -0.0205 | P1_P2_P3_Coeliac_Sphenopalatine_VS_P4_Lumbar_Stellate | 1 | 1 | 1 | 0 | 0 | 1 | 0 | 1 |
| NFIC | -0.0205 | P1_P3_P4_Coeliac_Sphenopalatine_VS_P2_Lumbar_Stellate | 1 | 0 | 1 | 1 | 0 | 1 | 0 | 1 |
| NAP1L3 | -0.0211 | P3_P4_Lumbar_Sphenopalatine_VS_P1_P2_Coeliac_Stellate | 0 | 0 | 1 | 1 | 1 | 0 | 0 | 1 |
| DAZAP1 | -0.0212 | P1_P2_P3_Lumbar_Coeliac_Stellate_Sphenopalatine_VS_P4 | 1 | 1 | 1 | 0 | 1 | 1 | 1 | 1 |
| EEF1AKMT2 | -0.0213 | P1_P3_P4_Lumbar_Coeliac_Stellate_Sphenopalatine_VS_P2 | 1 | 0 | 1 | 1 | 1 | 1 | 1 | 1 |
| HINT2 | -0.0214 | P1_P2_P3_P4_Lumbar_Stellate_Sphenopalatine_VS_Coeliac | 1 | 1 | 1 | 1 | 1 | 0 | 1 | 1 |
| RAB4A | -0.0214 | P1_P2_P3_P4_Lumbar_Stellate_Sphenopalatine_VS_Coeliac | 1 | 1 | 1 | 1 | 1 | 0 | 1 | 1 |
| RTCA | -0.0215 | P1_P2_P3_P4_Lumbar_Stellate_Sphenopalatine_VS_Coeliac | 1 | 1 | 1 | 1 | 1 | 0 | 1 | 1 |
| POLD4 | -0.0216 | P3_P4_Lumbar_Sphenopalatine_VS_P1_P2_Coeliac_Stellate | 0 | 0 | 1 | 1 | 1 | 0 | 0 | 1 |
| SCD1 | -0.0217 | P1_P2_P3_P4_Lumbar_Stellate_VS_Coeliac_Sphenopalatine | 1 | 1 | 1 | 1 | 1 | 0 | 1 | 0 |
| PCBD2 | -0.0218 | P1_P2_P3_P4_Lumbar_Stellate_Sphenopalatine_VS_Coeliac | 1 | 1 | 1 | 1 | 1 | 0 | 1 | 1 |
| CHGB | -0.0218 | P1_P2_P4_Lumbar_Coeliac_Stellate_Sphenopalatine_VS_P3 | 1 | 1 | 0 | 1 | 1 | 1 | 1 | 1 |
| AZIN2 | -0.0221 | P2_P3_P4_Lumbar_Coeliac_Stellate_VS_P1_Sphenopalatine | 0 | 1 | 1 | 1 | 1 | 1 | 1 | 0 |
| SNX21 | -0.0222 | P1_P2_P3_P4_Lumbar_Stellate_Sphenopalatine_VS_Coeliac | 1 | 1 | 1 | 1 | 1 | 0 | 1 | 1 |
| RAB15 | -0.0223 | P1_P2_P3_P4_Lumbar_Stellate_Sphenopalatine_VS_Coeliac | 1 | 1 | 1 | 1 | 1 | 0 | 1 | 1 |
| SPR | -0.0227 | P1_P2_P3_P4_Lumbar_Stellate_Sphenopalatine_VS_Coeliac | 1 | 1 | 1 | 1 | 1 | 0 | 1 | 1 |
| NAXD | -0.0228 | P1_P3_P4_Lumbar_Coeliac_Stellate_Sphenopalatine_VS_P2 | 1 | 0 | 1 | 1 | 1 | 1 | 1 | 1 |
| ELP5 | -0.0229 | P1_P2_P3_P4_Lumbar_Stellate_Sphenopalatine_VS_Coeliac | 1 | 1 | 1 | 1 | 1 | 0 | 1 | 1 |
| UBE2Q2 | -0.023 | P4_Coeliac_Sphenopalatine_VS_P1_P2_P3_Lumbar_Stellate | 0 | 0 | 0 | 1 | 0 | 1 | 0 | 1 |
| EXOC7 | -0.0232 | P1_P4_Lumbar_Coeliac_Stellate_VS_P2_P3_Sphenopalatine | 1 | 0 | 0 | 1 | 1 | 1 | 1 | 0 |
| SLC6A15 | -0.0234 | P1_P3_P4_Lumbar_Stellate_VS_P2_Coeliac_Sphenopalatine | 1 | 0 | 1 | 1 | 1 | 0 | 1 | 0 |
| MED14 | -0.0234 | P3_P4_Lumbar_Sphenopalatine_VS_P1_P2_Coeliac_Stellate | 0 | 0 | 1 | 1 | 1 | 0 | 0 | 1 |
| CCNC | -0.0235 | P3_P4_Lumbar_Coeliac_Stellate_VS_P1_P2_Sphenopalatine | 0 | 0 | 1 | 1 | 1 | 1 | 1 | 0 |
| ARID4B | -0.0235 | P2_P3_P4_Lumbar_Coeliac_Stellate_VS_P1_Sphenopalatine | 0 | 1 | 1 | 1 | 1 | 1 | 1 | 0 |
| IDH3A | -0.0235 | P1_P2_P3_P4_Lumbar_Stellate_Sphenopalatine_VS_Coeliac | 1 | 1 | 1 | 1 | 1 | 0 | 1 | 1 |
| FSD1 | -0.0235 | P1_P3_P4_Lumbar_Coeliac_Stellate_Sphenopalatine_VS_P2 | 1 | 0 | 1 | 1 | 1 | 1 | 1 | 1 |
| ACAA1A | -0.024 | P4_Lumbar_Coeliac_Stellate_VS_P1_P2_P3_Sphenopalatine | 0 | 0 | 0 | 1 | 1 | 1 | 1 | 0 |

| | | | | | | | | | | |
|---|---|---|---|---|---|---|---|---|---|---|
| FABP3 | -0.024 | P1_P2_P3_P4_Lumbar_Stellate_Sphenopalatine_VS_Coeliac | 1 | 1 | 1 | 1 | 1 | 0 | 1 | 1 |
| SMARCB1 | -0.0241 | P4_Lumbar_Coeliac_Stellate_VS_P1_P2_P3_Sphenopalatine | 0 | 0 | 0 | 1 | 1 | 1 | 1 | 0 |
| CSF2RA | -0.0243 | P3_Lumbar_Sphenopalatine_VS_P1_P2_P4_Coeliac_Stellate | 0 | 0 | 1 | 0 | 1 | 0 | 0 | 1 |
| ODR4 | -0.0244 | P1_P2_P3_Coeliac_Sphenopalatine_VS_P4_Lumbar_Stellate | 1 | 1 | 1 | 0 | 0 | 1 | 0 | 1 |
| HBP1 | -0.0245 | P1_P2_P4_Lumbar_Coeliac_Stellate_VS_P3_Sphenopalatine | 1 | 1 | 0 | 1 | 1 | 1 | 1 | 0 |
| 2610001J05F | -0.0245 | P1_P2_P3_P4_Lumbar_Stellate_Sphenopalatine_VS_Coeliac | 1 | 1 | 1 | 1 | 1 | 0 | 1 | 1 |
| PARP1 | -0.0245 | P1_P2_P4_Lumbar_Coeliac_Stellate_Sphenopalatine_VS_P3 | 1 | 1 | 0 | 1 | 1 | 1 | 1 | 1 |
| FBXO45 | -0.0246 | P2_P3_Lumbar_Sphenopalatine_VS_P1_P4_Coeliac_Stellate | 0 | 1 | 1 | 0 | 1 | 0 | 0 | 1 |
| DLGAP3 | -0.0247 | P1_P3_P4_Coeliac_Sphenopalatine_VS_P2_Lumbar_Stellate | 1 | 0 | 1 | 1 | 0 | 1 | 0 | 1 |
| NOP56 | -0.0247 | P1_P3_P4_Lumbar_Coeliac_Stellate_Sphenopalatine_VS_P2 | 1 | 0 | 1 | 1 | 1 | 1 | 1 | 1 |
| ADH5 | -0.0247 | P1_P3_P4_Lumbar_Coeliac_Stellate_Sphenopalatine_VS_P2 | 1 | 0 | 1 | 1 | 1 | 1 | 1 | 1 |
| FAM114A2 | -0.0251 | P2_P3_P4_Lumbar_Coeliac_Stellate_VS_P1_Sphenopalatine | 0 | 1 | 1 | 1 | 1 | 1 | 1 | 0 |
| GYG | -0.0251 | P1_P2_P3_P4_Lumbar_Stellate_Sphenopalatine_VS_Coeliac | 1 | 1 | 1 | 1 | 1 | 0 | 1 | 1 |
| ATRNL1 | -0.0251 | P1_P2_P4_Lumbar_Coeliac_Stellate_Sphenopalatine_VS_P3 | 1 | 1 | 0 | 1 | 1 | 1 | 1 | 1 |
| MYH10 | -0.0255 | P4_Coeliac_Sphenopalatine_VS_P1_P2_P3_Lumbar_Stellate | 0 | 0 | 0 | 1 | 0 | 1 | 0 | 1 |
| SDR39U1 | -0.0256 | P3_P4_Coeliac_Sphenopalatine_VS_P1_P2_Lumbar_Stellate | 0 | 0 | 1 | 1 | 0 | 1 | 0 | 1 |
| CDKN1A | -0.0258 | P3_Sphenopalatine_VS_P1_P2_P4_Lumbar_Coeliac_Stellate | 0 | 0 | 1 | 0 | 0 | 0 | 0 | 1 |
| CDC23 | -0.026 | P2_P3_P4_Lumbar_Coeliac_Stellate_VS_P1_Sphenopalatine | 0 | 1 | 1 | 1 | 1 | 1 | 1 | 0 |
| EIF4A3 | -0.0262 | P1_P4_Lumbar_Coeliac_Stellate_VS_P2_P3_Sphenopalatine | 1 | 0 | 0 | 1 | 1 | 1 | 1 | 0 |
| COA5 | -0.0264 | P1_P2_P3_P4_Lumbar_Stellate_Sphenopalatine_VS_Coeliac | 1 | 1 | 1 | 1 | 1 | 0 | 1 | 1 |
| PARD6A | -0.0266 | P3_Sphenopalatine_VS_P1_P2_P4_Lumbar_Coeliac_Stellate | 0 | 0 | 1 | 0 | 0 | 0 | 0 | 1 |
| SIAH1B | -0.0268 | P3_P4_Lumbar_Coeliac_Stellate_VS_P1_P2_Sphenopalatine | 0 | 0 | 1 | 1 | 1 | 1 | 1 | 0 |
| RNF208 | -0.027 | P1_P2_P3_P4_Lumbar_Stellate_Sphenopalatine_VS_Coeliac | 1 | 1 | 1 | 1 | 1 | 0 | 1 | 1 |
| TIGAR | -0.0271 | P1_P2_Lumbar_Sphenopalatine_VS_P3_P4_Coeliac_Stellate | 1 | 1 | 0 | 0 | 1 | 0 | 0 | 1 |
| TCF4 | -0.0271 | P1_P3_P4_Lumbar_Coeliac_Stellate_Sphenopalatine_VS_P2 | 1 | 0 | 1 | 1 | 1 | 1 | 1 | 1 |
| PTBP3 | -0.0272 | P1_P3_P4_Lumbar_Coeliac_Stellate_Sphenopalatine_VS_P2 | 1 | 0 | 1 | 1 | 1 | 1 | 1 | 1 |
| GTF2H2 | -0.0274 | P1_P4_Lumbar_Coeliac_Stellate_VS_P2_P3_Sphenopalatine | 1 | 0 | 0 | 1 | 1 | 1 | 1 | 0 |
| BAD | -0.0278 | P1_P2_P3_P4_Lumbar_Stellate_Sphenopalatine_VS_Coeliac | 1 | 1 | 1 | 1 | 1 | 0 | 1 | 1 |
| CEBPZ | -0.028 | Lumbar_Sphenopalatine_VS_P1_P2_P3_P4_Coeliac_Stellate | 0 | 0 | 0 | 0 | 1 | 0 | 0 | 1 |
| NT5C2 | -0.028 | P4_Coeliac_Sphenopalatine_VS_P1_P2_P3_Lumbar_Stellate | 0 | 0 | 0 | 1 | 0 | 1 | 0 | 1 |

| | | | | | | | | | | |
|---|---|---|---|---|---|---|---|---|---|---|
| BTBD9 | -0.0282 | Lumbar_Sphenopalatine_VS_P1_P2_P3_P4_Coeliac_Stellate | 0 | 0 | 0 | 0 | 1 | 0 | 0 | 1 |
| GPCPD1 | -0.0283 | P1_P2_P4_Lumbar_Coeliac_Stellate_Sphenopalatine_VS_P3 | 1 | 1 | 0 | 1 | 1 | 1 | 1 | 1 |
| ALCAM | -0.0285 | P1_P2_P3_P4_Lumbar_Stellate_VS_Coeliac_Sphenopalatine | 1 | 1 | 1 | 1 | 1 | 0 | 1 | 0 |
| DYNLT1F | -0.0286 | P1_P3_P4_Lumbar_Coeliac_Stellate_Sphenopalatine_VS_P2 | 1 | 0 | 1 | 1 | 1 | 1 | 1 | 1 |
| GUCY1B1 | -0.0288 | P1_P2_P3_Lumbar_Coeliac_Stellate_Sphenopalatine_VS_P4 | 1 | 1 | 1 | 0 | 1 | 1 | 1 | 1 |
| 1110059E24I | -0.0289 | P1_P2_P3_P4_Lumbar_Stellate_Sphenopalatine_VS_Coeliac | 1 | 1 | 1 | 1 | 1 | 0 | 1 | 1 |
| POLR2E | -0.029 | P1_P4_Lumbar_Coeliac_Stellate_VS_P2_P3_Sphenopalatine | 1 | 0 | 0 | 1 | 1 | 1 | 1 | 0 |
| TMEFF1 | -0.029 | P1_P2_P3_Lumbar_Coeliac_Stellate_Sphenopalatine_VS_P4 | 1 | 1 | 1 | 0 | 1 | 1 | 1 | 1 |
| CELSR2 | -0.0296 | P2_Lumbar_Sphenopalatine_VS_P1_P3_P4_Coeliac_Stellate | 0 | 1 | 0 | 0 | 1 | 0 | 0 | 1 |
| CADM3 | -0.0296 | P1_P2_P3_P4_Lumbar_Stellate_Sphenopalatine_VS_Coeliac | 1 | 1 | 1 | 1 | 1 | 0 | 1 | 1 |
| GCN1 | -0.0299 | P4_Coeliac_Sphenopalatine_VS_P1_P2_P3_Lumbar_Stellate | 0 | 0 | 0 | 1 | 0 | 1 | 0 | 1 |
| BOLA1 | -0.0302 | P3_Lumbar_Sphenopalatine_VS_P1_P2_P4_Coeliac_Stellate | 0 | 0 | 1 | 0 | 1 | 0 | 0 | 1 |
| FAM173A | -0.0303 | P1_P2_P3_P4_Lumbar_Stellate_Sphenopalatine_VS_Coeliac | 1 | 1 | 1 | 1 | 1 | 0 | 1 | 1 |
| NFKBIB | -0.0305 | P1_P3_VS_P2_P4_Lumbar_Coeliac_Stellate_Sphenopalatine | 1 | 0 | 1 | 0 | 0 | 0 | 0 | 0 |
| NDUFAF2 | -0.0305 | P1_Lumbar_Sphenopalatine_VS_P2_P3_P4_Coeliac_Stellate | 1 | 0 | 0 | 0 | 1 | 0 | 0 | 1 |
| HABP4 | -0.031 | P1_P2_P3_P4_Lumbar_Stellate_Sphenopalatine_VS_Coeliac | 1 | 1 | 1 | 1 | 1 | 0 | 1 | 1 |
| FAM163A | -0.0314 | P2_P3_Sphenopalatine_VS_P1_P4_Lumbar_Coeliac_Stellate | 0 | 1 | 1 | 0 | 0 | 0 | 0 | 1 |
| ACER3 | -0.0315 | Lumbar_Sphenopalatine_VS_P1_P2_P3_P4_Coeliac_Stellate | 0 | 0 | 0 | 0 | 1 | 0 | 0 | 1 |
| TMEM246 | -0.0315 | P1_P3_Lumbar_Sphenopalatine_VS_P2_P4_Coeliac_Stellate | 1 | 0 | 1 | 0 | 1 | 0 | 0 | 1 |
| TM2D3 | -0.0315 | P1_P3_P4_Lumbar_Coeliac_Stellate_Sphenopalatine_VS_P2 | 1 | 0 | 1 | 1 | 1 | 1 | 1 | 1 |
| B230216N24 | -0.0317 | P1_P3_Sphenopalatine_VS_P2_P4_Lumbar_Coeliac_Stellate | 1 | 0 | 1 | 0 | 0 | 0 | 0 | 1 |
| MANBAL | -0.032 | P1_P2_P3_P4_Lumbar_Stellate_Sphenopalatine_VS_Coeliac | 1 | 1 | 1 | 1 | 1 | 0 | 1 | 1 |
| NPRL2 | -0.0323 | P1_P4_Lumbar_Coeliac_Stellate_VS_P2_P3_Sphenopalatine | 1 | 0 | 0 | 1 | 1 | 1 | 1 | 0 |
| H2-DMA | -0.0324 | P1_P3_Lumbar_Coeliac_Stellate_VS_P2_P4_Sphenopalatine | 1 | 0 | 1 | 0 | 1 | 1 | 1 | 0 |
| USP47 | -0.0324 | P4_Coeliac_Sphenopalatine_VS_P1_P2_P3_Lumbar_Stellate | 0 | 0 | 0 | 1 | 0 | 1 | 0 | 1 |
| YIF1B | -0.0325 | P1_P3_P4_Lumbar_Coeliac_Stellate_Sphenopalatine_VS_P2 | 1 | 0 | 1 | 1 | 1 | 1 | 1 | 1 |
| RAB3IP | -0.0325 | P1_P3_P4_Lumbar_Coeliac_Stellate_Sphenopalatine_VS_P2 | 1 | 0 | 1 | 1 | 1 | 1 | 1 | 1 |
| DUSP8 | -0.0326 | P2_Lumbar_Sphenopalatine_VS_P1_P3_P4_Coeliac_Stellate | 0 | 1 | 0 | 0 | 1 | 0 | 0 | 1 |
| TTC4 | -0.0328 | P4_Lumbar_Coeliac_Stellate_VS_P1_P2_P3_Sphenopalatine | 0 | 0 | 0 | 1 | 1 | 1 | 1 | 0 |
| VPS39 | -0.0329 | P2_Lumbar_Sphenopalatine_VS_P1_P3_P4_Coeliac_Stellate | 0 | 1 | 0 | 0 | 1 | 0 | 0 | 1 |

| | | | | | | | | | | |
|---|---|---|---|---|---|---|---|---|---|---|
| GM50240 | -0.0329 | P1_P3_P4_Lumbar_Coeliac_Stellate_Sphenopalatine_VS_P2 | 1 | 0 | 1 | 1 | 1 | 1 | 1 | 1 |
| RUNDC3B | -0.0334 | Lumbar_Sphenopalatine_VS_P1_P2_P3_P4_Coeliac_Stellate | 0 | 0 | 0 | 0 | 1 | 0 | 0 | 1 |
| IFRD1 | -0.0335 | P1_P2_P4_Lumbar_Coeliac_Stellate_VS_P3_Sphenopalatine | 1 | 1 | 0 | 1 | 1 | 1 | 1 | 0 |
| SYNRG | -0.0335 | P4_Coeliac_Sphenopalatine_VS_P1_P2_P3_Lumbar_Stellate | 0 | 0 | 0 | 1 | 0 | 1 | 0 | 1 |
| ACHE | -0.0335 | P1_P2_P3_P4_Lumbar_Stellate_Sphenopalatine_VS_Coeliac | 1 | 1 | 1 | 1 | 1 | 0 | 1 | 1 |
| GM17018 | -0.0338 | P1_P2_P3_P4_Lumbar_Stellate_Sphenopalatine_VS_Coeliac | 1 | 1 | 1 | 1 | 1 | 0 | 1 | 1 |
| MRPL50 | -0.0338 | P1_P3_P4_Lumbar_Coeliac_Stellate_Sphenopalatine_VS_P2 | 1 | 0 | 1 | 1 | 1 | 1 | 1 | 1 |
| DDX19A | -0.0342 | P1_P2_P4_Lumbar_Coeliac_Stellate_VS_P3_Sphenopalatine | 1 | 1 | 0 | 1 | 1 | 1 | 1 | 0 |
| B230118H07 | -0.0344 | P1_P2_P4_Lumbar_Coeliac_Stellate_VS_P3_Sphenopalatine | 1 | 1 | 0 | 1 | 1 | 1 | 1 | 0 |
| CYB5R4 | -0.0344 | P1_P2_P3_Lumbar_Coeliac_Stellate_Sphenopalatine_VS_P4 | 1 | 1 | 1 | 0 | 1 | 1 | 1 | 1 |
| STRIP1 | -0.0346 | Lumbar_Sphenopalatine_VS_P1_P2_P3_P4_Coeliac_Stellate | 0 | 0 | 0 | 0 | 1 | 0 | 0 | 1 |
| SLC17A5 | -0.0346 | P4_Coeliac_Sphenopalatine_VS_P1_P2_P3_Lumbar_Stellate | 0 | 0 | 0 | 1 | 0 | 1 | 0 | 1 |
| CNTNAP2 | -0.0347 | P1_P2_P3_P4_Lumbar_Stellate_VS_Coeliac_Sphenopalatine | 1 | 1 | 1 | 1 | 1 | 0 | 1 | 0 |
| MFAP3 | -0.0347 | P1_P2_P4_Lumbar_Coeliac_Stellate_Sphenopalatine_VS_P3 | 1 | 1 | 0 | 1 | 1 | 1 | 1 | 1 |
| SLC48A1 | -0.0348 | P1_P2_P3_P4_Lumbar_Stellate_Sphenopalatine_VS_Coeliac | 1 | 1 | 1 | 1 | 1 | 0 | 1 | 1 |
| TBC1D20 | -0.035 | P2_P3_Lumbar_Sphenopalatine_VS_P1_P4_Coeliac_Stellate | 0 | 1 | 1 | 0 | 1 | 0 | 0 | 1 |
| OCEL1 | -0.0351 | P3_Lumbar_Sphenopalatine_VS_P1_P2_P4_Coeliac_Stellate | 0 | 0 | 1 | 0 | 1 | 0 | 0 | 1 |
| TMOD3 | -0.0353 | P1_P2_P3_Lumbar_Coeliac_Stellate_VS_P4_Sphenopalatine | 1 | 1 | 1 | 0 | 1 | 1 | 1 | 0 |
| MTX1 | -0.0353 | P1_P4_Lumbar_Coeliac_Stellate_VS_P2_P3_Sphenopalatine | 1 | 0 | 0 | 1 | 1 | 1 | 1 | 0 |
| ANKRD37 | -0.0353 | P2_P4_Lumbar_Coeliac_Stellate_VS_P1_P3_Sphenopalatine | 0 | 1 | 0 | 1 | 1 | 1 | 1 | 0 |
| SCAPER | -0.0353 | P4_Coeliac_Sphenopalatine_VS_P1_P2_P3_Lumbar_Stellate | 0 | 0 | 0 | 1 | 0 | 1 | 0 | 1 |
| 2700062C07I | -0.0355 | P3_Lumbar_Sphenopalatine_VS_P1_P2_P4_Coeliac_Stellate | 0 | 0 | 1 | 0 | 1 | 0 | 0 | 1 |
| NAXE | -0.0355 | P1_P3_P4_Lumbar_Coeliac_Stellate_Sphenopalatine_VS_P2 | 1 | 0 | 1 | 1 | 1 | 1 | 1 | 1 |
| TRMT10B | -0.0355 | P1_P3_P4_Lumbar_Coeliac_Stellate_Sphenopalatine_VS_P2 | 1 | 0 | 1 | 1 | 1 | 1 | 1 | 1 |
| THOC3 | -0.0355 | P1_P3_P4_Lumbar_Coeliac_Stellate_Sphenopalatine_VS_P2 | 1 | 0 | 1 | 1 | 1 | 1 | 1 | 1 |
| LPGAT1 | -0.0356 | P2_Lumbar_Sphenopalatine_VS_P1_P3_P4_Coeliac_Stellate | 0 | 1 | 0 | 0 | 1 | 0 | 0 | 1 |
| SBF2 | -0.0359 | P3_P4_Coeliac_Sphenopalatine_VS_P1_P2_Lumbar_Stellate | 0 | 0 | 1 | 1 | 0 | 1 | 0 | 1 |
| 9330182L06I | -0.036 | P3_Lumbar_Sphenopalatine_VS_P1_P2_P4_Coeliac_Stellate | 0 | 0 | 1 | 0 | 1 | 0 | 0 | 1 |
| CREBBP | -0.0364 | P4_Coeliac_Sphenopalatine_VS_P1_P2_P3_Lumbar_Stellate | 0 | 0 | 0 | 1 | 0 | 1 | 0 | 1 |
| ABHD16A | -0.0367 | P3_Lumbar_Sphenopalatine_VS_P1_P2_P4_Coeliac_Stellate | 0 | 0 | 1 | 0 | 1 | 0 | 0 | 1 |

| | | | | | | | | | | |
|---|---|---|---|---|---|---|---|---|---|---|
| C1QTNF4 | -0.0367 | P1_P3_P4_Lumbar_Coeliac_Stellate_Sphenopalatine_VS_P2 | 1 | 0 | 1 | 1 | 1 | 1 | 1 | 1 |
| TNKS | -0.0368 | P2_Lumbar_Sphenopalatine_VS_P1_P3_P4_Coeliac_Stellate | 0 | 1 | 0 | 0 | 1 | 0 | 0 | 1 |
| EXOSC1 | -0.037 | P3_Lumbar_Sphenopalatine_VS_P1_P2_P4_Coeliac_Stellate | 0 | 0 | 1 | 0 | 1 | 0 | 0 | 1 |
| HOMER3 | -0.0372 | P2_P3_Lumbar_Coeliac_Stellate_VS_P1_P4_Sphenopalatine | 0 | 1 | 1 | 0 | 1 | 1 | 1 | 0 |
| MAP1S | -0.0373 | Lumbar_Sphenopalatine_VS_P1_P2_P3_P4_Coeliac_Stellate | 0 | 0 | 0 | 0 | 1 | 0 | 0 | 1 |
| DPYSL4 | -0.0375 | P1_P2_P3_Lumbar_Coeliac_Stellate_Sphenopalatine_VS_P4 | 1 | 1 | 1 | 0 | 1 | 1 | 1 | 1 |
| RBM4B | -0.0376 | P3_Lumbar_Sphenopalatine_VS_P1_P2_P4_Coeliac_Stellate | 0 | 0 | 1 | 0 | 1 | 0 | 0 | 1 |
| GTF3A | -0.0379 | P1_P3_Lumbar_Sphenopalatine_VS_P2_P4_Coeliac_Stellate | 1 | 0 | 1 | 0 | 1 | 0 | 0 | 1 |
| CORO1A | -0.0379 | P1_P3_Lumbar_Coeliac_Stellate_Sphenopalatine_VS_P2_P4 | 1 | 0 | 1 | 0 | 1 | 1 | 1 | 1 |
| FBF1 | -0.038 | P2_Coeliac_Sphenopalatine_VS_P1_P3_P4_Lumbar_Stellate | 0 | 1 | 0 | 0 | 0 | 1 | 0 | 1 |
| TFPT | -0.038 | P1_P3_P4_Lumbar_Coeliac_Stellate_Sphenopalatine_VS_P2 | 1 | 0 | 1 | 1 | 1 | 1 | 1 | 1 |
| YAE1D1 | -0.0382 | P1_P3_Lumbar_Sphenopalatine_VS_P2_P4_Coeliac_Stellate | 1 | 0 | 1 | 0 | 1 | 0 | 0 | 1 |
| SPRED2 | -0.0385 | P1_P2_P3_P4_Lumbar_Stellate_Sphenopalatine_VS_Coeliac | 1 | 1 | 1 | 1 | 1 | 0 | 1 | 1 |
| CHCHD5 | -0.0386 | P3_Sphenopalatine_VS_P1_P2_P4_Lumbar_Coeliac_Stellate | 0 | 0 | 1 | 0 | 0 | 0 | 0 | 1 |
| TMEM126A | -0.0386 | P1_P3_P4_Lumbar_Coeliac_Stellate_Sphenopalatine_VS_P2 | 1 | 0 | 1 | 1 | 1 | 1 | 1 | 1 |
| GM10138 | -0.0387 | P3_Lumbar_Sphenopalatine_VS_P1_P2_P4_Coeliac_Stellate | 0 | 0 | 1 | 0 | 1 | 0 | 0 | 1 |
| RNFT2 | -0.0387 | P3_Lumbar_Sphenopalatine_VS_P1_P2_P4_Coeliac_Stellate | 0 | 0 | 1 | 0 | 1 | 0 | 0 | 1 |
| FOXO3 | -0.0389 | P2_P4_Lumbar_Coeliac_Stellate_Sphenopalatine_VS_P1_P3 | 0 | 1 | 0 | 1 | 1 | 1 | 1 | 1 |
| DNAJC11 | -0.039 | P1_P3_Lumbar_Sphenopalatine_VS_P2_P4_Coeliac_Stellate | 1 | 0 | 1 | 0 | 1 | 0 | 0 | 1 |
| SLC10A4 | -0.039 | P1_P2_P3_P4_Lumbar_Stellate_Sphenopalatine_VS_Coeliac | 1 | 1 | 1 | 1 | 1 | 0 | 1 | 1 |
| ANXA7 | -0.039 | P1_P2_P3_P4_Lumbar_Stellate_Sphenopalatine_VS_Coeliac | 1 | 1 | 1 | 1 | 1 | 0 | 1 | 1 |
| SURF1 | -0.0392 | P1_P2_P3_P4_Lumbar_Stellate_Sphenopalatine_VS_Coeliac | 1 | 1 | 1 | 1 | 1 | 0 | 1 | 1 |
| ZAR1 | -0.0393 | Lumbar_Sphenopalatine_VS_P1_P2_P3_P4_Coeliac_Stellate | 0 | 0 | 0 | 0 | 1 | 0 | 0 | 1 |
| TTC9 | -0.0394 | P2_Lumbar_Sphenopalatine_VS_P1_P3_P4_Coeliac_Stellate | 0 | 1 | 0 | 0 | 1 | 0 | 0 | 1 |
| NRSN1 | -0.0398 | P1_P2_P3_Lumbar_Coeliac_Stellate_Sphenopalatine_VS_P4 | 1 | 1 | 1 | 0 | 1 | 1 | 1 | 1 |
| MRPL44 | -0.0399 | P1_P2_P3_P4_Lumbar_Stellate_Sphenopalatine_VS_Coeliac | 1 | 1 | 1 | 1 | 1 | 0 | 1 | 1 |
| FNDC4 | -0.0399 | P1_P2_P3_P4_Lumbar_Stellate_Sphenopalatine_VS_Coeliac | 1 | 1 | 1 | 1 | 1 | 0 | 1 | 1 |
| LSM10 | -0.0399 | P1_P3_P4_Lumbar_Coeliac_Stellate_Sphenopalatine_VS_P2 | 1 | 0 | 1 | 1 | 1 | 1 | 1 | 1 |
| 2010320M18 | -0.0399 | P1_P3_P4_Lumbar_Coeliac_Stellate_Sphenopalatine_VS_P2 | 1 | 0 | 1 | 1 | 1 | 1 | 1 | 1 |
| RAB3C | -0.0404 | P3_Lumbar_Sphenopalatine_VS_P1_P2_P4_Coeliac_Stellate | 0 | 0 | 1 | 0 | 1 | 0 | 0 | 1 |

| | | | | | | | | | | |
|---|---|---|---|---|---|---|---|---|---|---|
| KLHL7 | -0.0405 | P1_P2_P4_Lumbar_Coeliac_Stellate_VS_P3_Sphenopalatine | 1 | 1 | 0 | 1 | 1 | 1 | 1 | 0 |
| NOB1 | -0.0406 | P1_P2_P3_Lumbar_Coeliac_Stellate_Sphenopalatine_VS_P4 | 1 | 1 | 1 | 0 | 1 | 1 | 1 | 1 |
| SCAF11 | -0.0407 | P2_P3_P4_Lumbar_Coeliac_Stellate_VS_P1_Sphenopalatine | 0 | 1 | 1 | 1 | 1 | 1 | 1 | 0 |
| SHISA5 | -0.0407 | P1_P2_P3_P4_Stellate_Sphenopalatine_VS_Lumbar_Coeliac | 1 | 1 | 1 | 1 | 0 | 0 | 1 | 1 |
| MAPK11 | -0.0408 | P3_Lumbar_Sphenopalatine_VS_P1_P2_P4_Coeliac_Stellate | 0 | 0 | 1 | 0 | 1 | 0 | 0 | 1 |
| ALYREF2 | -0.0408 | P1_P3_P4_Lumbar_Coeliac_Stellate_Sphenopalatine_VS_P2 | 1 | 0 | 1 | 1 | 1 | 1 | 1 | 1 |
| XRN2 | -0.041 | P1_P4_Lumbar_Coeliac_Stellate_VS_P2_P3_Sphenopalatine | 1 | 0 | 0 | 1 | 1 | 1 | 1 | 0 |
| DHRS7 | -0.0416 | P1_P2_P4_Lumbar_Coeliac_Stellate_VS_P3_Sphenopalatine | 1 | 1 | 0 | 1 | 1 | 1 | 1 | 0 |
| GSE1 | -0.0417 | P2_P3_Lumbar_Sphenopalatine_VS_P1_P4_Coeliac_Stellate | 0 | 1 | 1 | 0 | 1 | 0 | 0 | 1 |
| FXYD7 | -0.0418 | P1_P2_P3_P4_Lumbar_Stellate_Sphenopalatine_VS_Coeliac | 1 | 1 | 1 | 1 | 1 | 0 | 1 | 1 |
| AGL | -0.042 | P2_Lumbar_Sphenopalatine_VS_P1_P3_P4_Coeliac_Stellate | 0 | 1 | 0 | 0 | 1 | 0 | 0 | 1 |
| SAFB2 | -0.0425 | P3_P4_Coeliac_Sphenopalatine_VS_P1_P2_Lumbar_Stellate | 0 | 0 | 1 | 1 | 0 | 1 | 0 | 1 |
| SERP1 | -0.0429 | P1_P2_P3_Lumbar_Coeliac_Stellate_Sphenopalatine_VS_P4 | 1 | 1 | 1 | 0 | 1 | 1 | 1 | 1 |
| DHRS13 | -0.0429 | P1_P3_P4_Lumbar_Coeliac_Stellate_Sphenopalatine_VS_P2 | 1 | 0 | 1 | 1 | 1 | 1 | 1 | 1 |
| LRRFIP2 | -0.043 | P1_P3_Lumbar_Sphenopalatine_VS_P2_P4_Coeliac_Stellate | 1 | 0 | 1 | 0 | 1 | 0 | 0 | 1 |
| FBXW9 | -0.043 | P1_P2_P3_P4_Lumbar_Stellate_Sphenopalatine_VS_Coeliac | 1 | 1 | 1 | 1 | 1 | 0 | 1 | 1 |
| XIAP | -0.0434 | P1_P2_P3_Coeliac_Sphenopalatine_VS_P4_Lumbar_Stellate | 1 | 1 | 1 | 0 | 0 | 1 | 0 | 1 |
| PRR3 | -0.0436 | P1_P2_P3_P4_Lumbar_Stellate_Sphenopalatine_VS_Coeliac | 1 | 1 | 1 | 1 | 1 | 0 | 1 | 1 |
| RFLNB | -0.0436 | P1_P2_P4_Lumbar_Coeliac_Stellate_Sphenopalatine_VS_P3 | 1 | 1 | 0 | 1 | 1 | 1 | 1 | 1 |
| GLRX | -0.0437 | P1_P2_P3_P4_Lumbar_Stellate_Sphenopalatine_VS_Coeliac | 1 | 1 | 1 | 1 | 1 | 0 | 1 | 1 |
| PRCC | -0.0438 | P2_P3_P4_Lumbar_Coeliac_Stellate_VS_P1_Sphenopalatine | 0 | 1 | 1 | 1 | 1 | 1 | 1 | 0 |
| SNRPA1 | -0.0439 | P1_P3_P4_Lumbar_Coeliac_Stellate_Sphenopalatine_VS_P2 | 1 | 0 | 1 | 1 | 1 | 1 | 1 | 1 |
| MARK4 | -0.044 | P2_Lumbar_Sphenopalatine_VS_P1_P3_P4_Coeliac_Stellate | 0 | 1 | 0 | 0 | 1 | 0 | 0 | 1 |
| B4GALNT4 | -0.044 | P3_Lumbar_Sphenopalatine_VS_P1_P2_P4_Coeliac_Stellate | 0 | 0 | 1 | 0 | 1 | 0 | 0 | 1 |
| POLR2L | -0.044 | P1_P2_P3_P4_Stellate_Sphenopalatine_VS_Lumbar_Coeliac | 1 | 1 | 1 | 1 | 0 | 0 | 1 | 1 |
| SNRNP25 | -0.044 | P1_P2_P3_P4_Lumbar_Stellate_Sphenopalatine_VS_Coeliac | 1 | 1 | 1 | 1 | 1 | 0 | 1 | 1 |
| HDAC1 | -0.0442 | P1_P2_P3_Lumbar_Coeliac_Stellate_VS_P4_Sphenopalatine | 1 | 1 | 1 | 0 | 1 | 1 | 1 | 0 |
| PPM1L | -0.0443 | P1_P3_P4_Lumbar_Coeliac_Stellate_Sphenopalatine_VS_P2 | 1 | 0 | 1 | 1 | 1 | 1 | 1 | 1 |
| UNC13A | -0.0445 | P1_P2_P3_Coeliac_Sphenopalatine_VS_P4_Lumbar_Stellate | 1 | 1 | 1 | 0 | 0 | 1 | 0 | 1 |
| KCTD3 | -0.0446 | P2_P3_Sphenopalatine_VS_P1_P4_Lumbar_Coeliac_Stellate | 0 | 1 | 1 | 0 | 0 | 0 | 0 | 1 |

| | | | | | | | | | | |
|---|---|---|---|---|---|---|---|---|---|---|
| CACFD1 | -0.0447 | P1_P2_P3_P4_Lumbar_Stellate_Sphenopalatine_VS_Coeliac | 1 | 1 | 1 | 1 | 1 | 0 | 1 | 1 |
| BC005561 | -0.0447 | P1_P2_P4_Lumbar_Coeliac_Stellate_Sphenopalatine_VS_P3 | 1 | 1 | 0 | 1 | 1 | 1 | 1 | 1 |
| LYAR | -0.0447 | P1_P3_P4_Lumbar_Coeliac_Stellate_Sphenopalatine_VS_P2 | 1 | 0 | 1 | 1 | 1 | 1 | 1 | 1 |
| CCDC174 | -0.0448 | P1_P2_P4_Lumbar_Coeliac_Stellate_VS_P3_Sphenopalatine | 1 | 1 | 0 | 1 | 1 | 1 | 1 | 0 |
| CIB2 | -0.0448 | P1_P2_P3_Lumbar_Coeliac_Stellate_Sphenopalatine_VS_P4 | 1 | 1 | 1 | 0 | 1 | 1 | 1 | 1 |
| GM11808 | -0.0452 | P1_P2_P3_P4_Coeliac_Stellate_Sphenopalatine_VS_Lumbar | 1 | 1 | 1 | 1 | 0 | 1 | 1 | 1 |
| NXT1 | -0.0452 | P1_P3_P4_Lumbar_Coeliac_Stellate_Sphenopalatine_VS_P2 | 1 | 0 | 1 | 1 | 1 | 1 | 1 | 1 |
| HAUS1 | -0.0453 | P1_P3_P4_Lumbar_Coeliac_Stellate_Sphenopalatine_VS_P2 | 1 | 0 | 1 | 1 | 1 | 1 | 1 | 1 |
| SLC29A4 | -0.0455 | P1_P2_P3_Lumbar_Coeliac_Stellate_Sphenopalatine_VS_P4 | 1 | 1 | 1 | 0 | 1 | 1 | 1 | 1 |
| ACSL6 | -0.0455 | P1_P2_P3_Lumbar_Coeliac_Stellate_Sphenopalatine_VS_P4 | 1 | 1 | 1 | 0 | 1 | 1 | 1 | 1 |
| GORASP2 | -0.0456 | P1_P2_P3_P4_Lumbar_Stellate_Sphenopalatine_VS_Coeliac | 1 | 1 | 1 | 1 | 1 | 0 | 1 | 1 |
| BNIP3 | -0.046 | P2_P3_P4_Lumbar_Coeliac_Stellate_Sphenopalatine_VS_P1 | 0 | 1 | 1 | 1 | 1 | 1 | 1 | 1 |
| AASDHPPT | -0.0461 | P2_P3_P4_Lumbar_Coeliac_Stellate_VS_P1_Sphenopalatine | 0 | 1 | 1 | 1 | 1 | 1 | 1 | 0 |
| GAB2 | -0.0463 | P3_Lumbar_Sphenopalatine_VS_P1_P2_P4_Coeliac_Stellate | 0 | 0 | 1 | 0 | 1 | 0 | 0 | 1 |
| RIMS3 | -0.0465 | P1_P2_P4_Lumbar_Coeliac_Stellate_VS_P3_Sphenopalatine | 1 | 1 | 0 | 1 | 1 | 1 | 1 | 0 |
| WDFY3 | -0.0466 | P2_Lumbar_Sphenopalatine_VS_P1_P3_P4_Coeliac_Stellate | 0 | 1 | 0 | 0 | 1 | 0 | 0 | 1 |
| GNA12 | -0.0466 | P2_Lumbar_Sphenopalatine_VS_P1_P3_P4_Coeliac_Stellate | 0 | 1 | 0 | 0 | 1 | 0 | 0 | 1 |
| CLK3 | -0.0468 | P1_P2_P4_Lumbar_Coeliac_Stellate_VS_P3_Sphenopalatine | 1 | 1 | 0 | 1 | 1 | 1 | 1 | 0 |
| ODF2 | -0.0469 | P3_Lumbar_Sphenopalatine_VS_P1_P2_P4_Coeliac_Stellate | 0 | 0 | 1 | 0 | 1 | 0 | 0 | 1 |
| IGSF9B | -0.0471 | P1_P2_P3_P4_Lumbar_Stellate_Sphenopalatine_VS_Coeliac | 1 | 1 | 1 | 1 | 1 | 0 | 1 | 1 |
| LRSAM1 | -0.0471 | P1_P2_P4_Lumbar_Coeliac_Stellate_Sphenopalatine_VS_P3 | 1 | 1 | 0 | 1 | 1 | 1 | 1 | 1 |
| IVNS1ABP | -0.0473 | P1_P3_P4_Lumbar_Coeliac_Stellate_Sphenopalatine_VS_P2 | 1 | 0 | 1 | 1 | 1 | 1 | 1 | 1 |
| R3HDM1 | -0.0475 | P3_P4_Coeliac_Sphenopalatine_VS_P1_P2_Lumbar_Stellate | 0 | 0 | 1 | 1 | 0 | 1 | 0 | 1 |
| HIST3H2A | -0.0475 | P1_P2_P3_P4_Lumbar_Stellate_Sphenopalatine_VS_Coeliac | 1 | 1 | 1 | 1 | 1 | 0 | 1 | 1 |
| LAS1L | -0.0477 | P1_P4_Lumbar_Coeliac_Stellate_VS_P2_P3_Sphenopalatine | 1 | 0 | 0 | 1 | 1 | 1 | 1 | 0 |
| POLR2C | -0.0477 | P1_P3_Lumbar_Sphenopalatine_VS_P2_P4_Coeliac_Stellate | 1 | 0 | 1 | 0 | 1 | 0 | 0 | 1 |
| CHFR | -0.0478 | P2_P3_P4_Lumbar_Coeliac_Stellate_VS_P1_Sphenopalatine | 0 | 1 | 1 | 1 | 1 | 1 | 1 | 0 |
| SMOX | -0.0479 | P3_P4_Coeliac_Sphenopalatine_VS_P1_P2_Lumbar_Stellate | 0 | 0 | 1 | 1 | 0 | 1 | 0 | 1 |
| GM14418 | -0.0479 | P1_P2_P3_P4_Coeliac_Stellate_Sphenopalatine_VS_Lumbar | 1 | 1 | 1 | 1 | 0 | 1 | 1 | 1 |
| KLHDC8B | -0.0481 | P2_P3_P4_Lumbar_Coeliac_Stellate_VS_P1_Sphenopalatine | 0 | 1 | 1 | 1 | 1 | 1 | 1 | 0 |

| | | | | | | | | | | |
|---|---|---|---|---|---|---|---|---|---|---|
| GM5868 | -0.0482 | Lumbar_Coeliac_Stellate_Sphenopalatine_VS_P1_P2_P3_P4 | 0 | 0 | 0 | 0 | 1 | 1 | 1 | 1 |
| DCAKD | -0.0483 | P1_P2_P3_Lumbar_Coeliac_Stellate_Sphenopalatine_VS_P4 | 1 | 1 | 1 | 0 | 1 | 1 | 1 | 1 |
| RPRD1A | -0.0484 | P2_Lumbar_Sphenopalatine_VS_P1_P3_P4_Coeliac_Stellate | 0 | 1 | 0 | 0 | 1 | 0 | 0 | 1 |
| CD320 | -0.0485 | P1_P3_Lumbar_Sphenopalatine_VS_P2_P4_Coeliac_Stellate | 1 | 0 | 1 | 0 | 1 | 0 | 0 | 1 |
| CHORDC1 | -0.0485 | P1_P3_P4_Lumbar_Coeliac_Stellate_Sphenopalatine_VS_P2 | 1 | 0 | 1 | 1 | 1 | 1 | 1 | 1 |
| PHF20L1 | -0.0486 | P1_P2_P3_Coeliac_Sphenopalatine_VS_P4_Lumbar_Stellate | 1 | 1 | 1 | 0 | 0 | 1 | 0 | 1 |
| ARMC6 | -0.0487 | P3_Lumbar_Sphenopalatine_VS_P1_P2_P4_Coeliac_Stellate | 0 | 0 | 1 | 0 | 1 | 0 | 0 | 1 |
| HIBCH | -0.0489 | P1_P2_P3_Lumbar_Coeliac_Stellate_VS_P4_Sphenopalatine | 1 | 1 | 1 | 0 | 1 | 1 | 1 | 0 |
| TMEM14C | -0.049 | P3_P4_Lumbar_Coeliac_Stellate_Sphenopalatine_VS_P1_P2 | 0 | 0 | 1 | 1 | 1 | 1 | 1 | 1 |
| 2310009A05I | -0.049 | P1_P3_P4_Lumbar_Coeliac_Stellate_Sphenopalatine_VS_P2 | 1 | 0 | 1 | 1 | 1 | 1 | 1 | 1 |
| PABPC4 | -0.0493 | P2_P3_Lumbar_Coeliac_Stellate_VS_P1_P4_Sphenopalatine | 0 | 1 | 1 | 0 | 1 | 1 | 1 | 0 |
| PDRG1 | -0.0493 | P1_P3_P4_Lumbar_Stellate_Sphenopalatine_VS_P2_Coeliac | 1 | 0 | 1 | 1 | 1 | 0 | 1 | 1 |
| CRELD1 | -0.0494 | P1_P2_P3_P4_Lumbar_Stellate_Sphenopalatine_VS_Coeliac | 1 | 1 | 1 | 1 | 1 | 0 | 1 | 1 |
| LINGO1 | -0.0496 | P1_P2_P4_Lumbar_Coeliac_Stellate_Sphenopalatine_VS_P3 | 1 | 1 | 0 | 1 | 1 | 1 | 1 | 1 |
| MCTS1 | -0.0496 | P1_P3_P4_Lumbar_Coeliac_Stellate_Sphenopalatine_VS_P2 | 1 | 0 | 1 | 1 | 1 | 1 | 1 | 1 |
| BBS4 | -0.0497 | P1_P2_P4_Lumbar_Coeliac_Stellate_VS_P3_Sphenopalatine | 1 | 1 | 0 | 1 | 1 | 1 | 1 | 0 |
| HAP1 | -0.0498 | P1_P3_P4_Lumbar_Coeliac_Stellate_Sphenopalatine_VS_P2 | 1 | 0 | 1 | 1 | 1 | 1 | 1 | 1 |
| ZFP941 | -0.0499 | P1_P4_Lumbar_Coeliac_Stellate_VS_P2_P3_Sphenopalatine | 1 | 0 | 0 | 1 | 1 | 1 | 1 | 0 |
| 5730455P16I | -0.0499 | P2_Lumbar_Sphenopalatine_VS_P1_P3_P4_Coeliac_Stellate | 0 | 1 | 0 | 0 | 1 | 0 | 0 | 1 |
| H1F0 | -0.0503 | P1_P2_P3_P4_Lumbar_Stellate_Sphenopalatine_VS_Coeliac | 1 | 1 | 1 | 1 | 1 | 0 | 1 | 1 |
| SPG21 | -0.0503 | P1_P3_P4_Lumbar_Coeliac_Stellate_Sphenopalatine_VS_P2 | 1 | 0 | 1 | 1 | 1 | 1 | 1 | 1 |
| CD9 | -0.0504 | P1_P2_P3_P4_Lumbar_Stellate_Sphenopalatine_VS_Coeliac | 1 | 1 | 1 | 1 | 1 | 0 | 1 | 1 |
| DNAJC2 | -0.0504 | P1_P3_P4_Lumbar_Coeliac_Stellate_Sphenopalatine_VS_P2 | 1 | 0 | 1 | 1 | 1 | 1 | 1 | 1 |
| TGOLN1 | -0.0505 | P2_P3_Sphenopalatine_VS_P1_P4_Lumbar_Coeliac_Stellate | 0 | 1 | 1 | 0 | 0 | 0 | 0 | 1 |
| EXOC3 | -0.0505 | P3_P4_Coeliac_Sphenopalatine_VS_P1_P2_Lumbar_Stellate | 0 | 0 | 1 | 1 | 0 | 1 | 0 | 1 |
| ARHGAP35 | -0.0506 | P2_P3_Lumbar_Sphenopalatine_VS_P1_P4_Coeliac_Stellate | 0 | 1 | 1 | 0 | 1 | 0 | 0 | 1 |
| MAP2K4 | -0.0506 | P2_P3_Lumbar_Sphenopalatine_VS_P1_P4_Coeliac_Stellate | 0 | 1 | 1 | 0 | 1 | 0 | 0 | 1 |
| USP15 | -0.0506 | P3_P4_Coeliac_Sphenopalatine_VS_P1_P2_Lumbar_Stellate | 0 | 0 | 1 | 1 | 0 | 1 | 0 | 1 |
| CAMSAP2 | -0.0507 | P2_Lumbar_Sphenopalatine_VS_P1_P3_P4_Coeliac_Stellate | 0 | 1 | 0 | 0 | 1 | 0 | 0 | 1 |
| RBM18 | -0.0507 | P1_P2_P3_Lumbar_Coeliac_Stellate_Sphenopalatine_VS_P4 | 1 | 1 | 1 | 0 | 1 | 1 | 1 | 1 |

| | | | | | | | | | | |
|---|---|---|---|---|---|---|---|---|---|---|
| PVT1 | -0.051 | P1_P2_P3_Lumbar_Coeliac_Stellate_VS_P4_Sphenopalatine | 1 | 1 | 1 | 0 | 1 | 1 | 1 | 0 |
| PTRH2 | -0.0511 | P1_P4_Lumbar_Coeliac_Stellate_VS_P2_P3_Sphenopalatine | 1 | 0 | 0 | 1 | 1 | 1 | 1 | 0 |
| BPGM | -0.0512 | P3_Lumbar_Sphenopalatine_VS_P1_P2_P4_Coeliac_Stellate | 0 | 0 | 1 | 0 | 1 | 0 | 0 | 1 |
| 5530601H04 | -0.0512 | P1_P4_Lumbar_Coeliac_Stellate_Sphenopalatine_VS_P2_P3 | 1 | 0 | 0 | 1 | 1 | 1 | 1 | 1 |
| PRDX4 | -0.0512 | P1_P3_P4_Lumbar_Coeliac_Stellate_Sphenopalatine_VS_P2 | 1 | 0 | 1 | 1 | 1 | 1 | 1 | 1 |
| ANKRD13C | -0.0515 | P2_Lumbar_Sphenopalatine_VS_P1_P3_P4_Coeliac_Stellate | 0 | 1 | 0 | 0 | 1 | 0 | 0 | 1 |
| VPS9D1 | -0.0515 | P3_Lumbar_Sphenopalatine_VS_P1_P2_P4_Coeliac_Stellate | 0 | 0 | 1 | 0 | 1 | 0 | 0 | 1 |
| RNF44 | -0.0515 | P1_P2_P3_Coeliac_Sphenopalatine_VS_P4_Lumbar_Stellate | 1 | 1 | 1 | 0 | 0 | 1 | 0 | 1 |
| ARMC10 | -0.0516 | P1_P2_P4_Lumbar_Coeliac_Stellate_VS_P3_Sphenopalatine | 1 | 1 | 0 | 1 | 1 | 1 | 1 | 0 |
| JAKMIP1 | -0.0516 | P1_P2_P3_P4_Lumbar_Stellate_Sphenopalatine_VS_Coeliac | 1 | 1 | 1 | 1 | 1 | 0 | 1 | 1 |
| UNC119 | -0.0518 | P2_P3_P4_Lumbar_Coeliac_Stellate_VS_P1_Sphenopalatine | 0 | 1 | 1 | 1 | 1 | 1 | 1 | 0 |
| HDGFL2 | -0.0519 | P1_P4_Lumbar_Coeliac_Stellate_VS_P2_P3_Sphenopalatine | 1 | 0 | 0 | 1 | 1 | 1 | 1 | 0 |
| PGAM5 | -0.0519 | P2_P3_Sphenopalatine_VS_P1_P4_Lumbar_Coeliac_Stellate | 0 | 1 | 1 | 0 | 0 | 0 | 0 | 1 |
| EIF3J2 | -0.0519 | P3_Coeliac_Sphenopalatine_VS_P1_P2_P4_Lumbar_Stellate | 0 | 0 | 1 | 0 | 0 | 1 | 0 | 1 |
| AHSA1 | -0.0519 | P1_P3_P4_Lumbar_Coeliac_Stellate_Sphenopalatine_VS_P2 | 1 | 0 | 1 | 1 | 1 | 1 | 1 | 1 |
| ARHGEF7 | -0.052 | P2_P3_Lumbar_Sphenopalatine_VS_P1_P4_Coeliac_Stellate | 0 | 1 | 1 | 0 | 1 | 0 | 0 | 1 |
| DEAF1 | -0.052 | P1_P2_P3_P4_Lumbar_Stellate_Sphenopalatine_VS_Coeliac | 1 | 1 | 1 | 1 | 1 | 0 | 1 | 1 |
| ZFP358 | -0.0523 | P1_P2_P3_Lumbar_Coeliac_Stellate_Sphenopalatine_VS_P4 | 1 | 1 | 1 | 0 | 1 | 1 | 1 | 1 |
| FAM171A2 | -0.0524 | P1_P2_P3_Coeliac_Sphenopalatine_VS_P4_Lumbar_Stellate | 1 | 1 | 1 | 0 | 0 | 1 | 0 | 1 |
| GET4 | -0.0525 | P1_P2_P3_P4_Lumbar_Stellate_Sphenopalatine_VS_Coeliac | 1 | 1 | 1 | 1 | 1 | 0 | 1 | 1 |
| FLYWCH2 | -0.0526 | P1_P2_P3_P4_Stellate_Sphenopalatine_VS_Lumbar_Coeliac | 1 | 1 | 1 | 1 | 0 | 0 | 1 | 1 |
| MSRB1 | -0.0526 | P1_P2_P3_P4_Lumbar_Stellate_Sphenopalatine_VS_Coeliac | 1 | 1 | 1 | 1 | 1 | 0 | 1 | 1 |
| PRKAA1 | -0.0528 | P2_Lumbar_Sphenopalatine_VS_P1_P3_P4_Coeliac_Stellate | 0 | 1 | 0 | 0 | 1 | 0 | 0 | 1 |
| PACS2 | -0.0531 | P2_Lumbar_Sphenopalatine_VS_P1_P3_P4_Coeliac_Stellate | 0 | 1 | 0 | 0 | 1 | 0 | 0 | 1 |
| GOSR2 | -0.0532 | P1_P2_P3_P4_Lumbar_Stellate_Sphenopalatine_VS_Coeliac | 1 | 1 | 1 | 1 | 1 | 0 | 1 | 1 |
| ANKRD46 | -0.0532 | P1_P2_P3_P4_Lumbar_Stellate_Sphenopalatine_VS_Coeliac | 1 | 1 | 1 | 1 | 1 | 0 | 1 | 1 |
| DGKD | -0.0533 | P3_Lumbar_Sphenopalatine_VS_P1_P2_P4_Coeliac_Stellate | 0 | 0 | 1 | 0 | 1 | 0 | 0 | 1 |
| FOXN3 | -0.0533 | P1_P2_Coeliac_Sphenopalatine_VS_P3_P4_Lumbar_Stellate | 1 | 1 | 0 | 0 | 0 | 1 | 0 | 1 |
| RAB11FIP2 | -0.0534 | P3_P4_Coeliac_Sphenopalatine_VS_P1_P2_Lumbar_Stellate | 0 | 0 | 1 | 1 | 0 | 1 | 0 | 1 |
| MPP3 | -0.0535 | P1_P4_Lumbar_Coeliac_Stellate_Sphenopalatine_VS_P2_P3 | 1 | 0 | 0 | 1 | 1 | 1 | 1 | 1 |

| | | | | | | | | | | |
|---|---|---|---|---|---|---|---|---|---|---|
| RRP9 | -0.0537 | P1_P3_Lumbar_Coeliac_Stellate_VS_P2_P4_Sphenopalatine | 1 | 0 | 1 | 0 | 1 | 1 | 1 | 0 |
| CORO2A | -0.0538 | P2_Lumbar_Sphenopalatine_VS_P1_P3_P4_Coeliac_Stellate | 0 | 1 | 0 | 0 | 1 | 0 | 0 | 1 |
| DDT | -0.0538 | P1_P2_P3_Lumbar_Coeliac_Stellate_Sphenopalatine_VS_P4 | 1 | 1 | 1 | 0 | 1 | 1 | 1 | 1 |
| SYT9 | -0.0539 | P2_Lumbar_Sphenopalatine_VS_P1_P3_P4_Coeliac_Stellate | 0 | 1 | 0 | 0 | 1 | 0 | 0 | 1 |
| TMEM120A | -0.054 | P1_P2_P3_Lumbar_Coeliac_Stellate_Sphenopalatine_VS_P4 | 1 | 1 | 1 | 0 | 1 | 1 | 1 | 1 |
| EHD1 | -0.0541 | P1_P2_P3_Lumbar_Coeliac_Stellate_VS_P4_Sphenopalatine | 1 | 1 | 1 | 0 | 1 | 1 | 1 | 0 |
| ZFP704 | -0.0542 | P1_P2_P4_Lumbar_Coeliac_Stellate_VS_P3_Sphenopalatine | 1 | 1 | 0 | 1 | 1 | 1 | 1 | 0 |
| FDX1 | -0.0542 | P1_P3_Lumbar_Sphenopalatine_VS_P2_P4_Coeliac_Stellate | 1 | 0 | 1 | 0 | 1 | 0 | 0 | 1 |
| BTG1 | -0.0542 | P1_P3_Lumbar_Sphenopalatine_VS_P2_P4_Coeliac_Stellate | 1 | 0 | 1 | 0 | 1 | 0 | 0 | 1 |
| LIMD2 | -0.0542 | P1_P3_P4_Lumbar_Stellate_Sphenopalatine_VS_P2_Coeliac | 1 | 0 | 1 | 1 | 1 | 0 | 1 | 1 |
| IFITM2 | -0.0542 | P1_P2_P3_Lumbar_Coeliac_Stellate_Sphenopalatine_VS_P4 | 1 | 1 | 1 | 0 | 1 | 1 | 1 | 1 |
| KCTD10 | -0.0545 | P1_P2_P4_Lumbar_Coeliac_Stellate_VS_P3_Sphenopalatine | 1 | 1 | 0 | 1 | 1 | 1 | 1 | 0 |
| CMTR1 | -0.0545 | P2_Lumbar_Sphenopalatine_VS_P1_P3_P4_Coeliac_Stellate | 0 | 1 | 0 | 0 | 1 | 0 | 0 | 1 |
| ULK2 | -0.0545 | P2_P3_Lumbar_Sphenopalatine_VS_P1_P4_Coeliac_Stellate | 0 | 1 | 1 | 0 | 1 | 0 | 0 | 1 |
| ISCA1 | -0.0545 | P1_P2_P3_P4_Coeliac_Stellate_Sphenopalatine_VS_Lumbar | 1 | 1 | 1 | 1 | 0 | 1 | 1 | 1 |
| TMEFF2 | -0.0546 | P1_P2_P3_P4_Lumbar_Stellate_Sphenopalatine_VS_Coeliac | 1 | 1 | 1 | 1 | 1 | 0 | 1 | 1 |
| GM42418 | -0.0547 | P1_P2_Coeliac_Sphenopalatine_VS_P3_P4_Lumbar_Stellate | 1 | 1 | 0 | 0 | 0 | 1 | 0 | 1 |
| SMAP2 | -0.0547 | P3_P4_Coeliac_Sphenopalatine_VS_P1_P2_Lumbar_Stellate | 0 | 0 | 1 | 1 | 0 | 1 | 0 | 1 |
| MPND | -0.0547 | P1_P2_P3_P4_Lumbar_Stellate_Sphenopalatine_VS_Coeliac | 1 | 1 | 1 | 1 | 1 | 0 | 1 | 1 |
| DIP2C | -0.0549 | P3_P4_Coeliac_Sphenopalatine_VS_P1_P2_Lumbar_Stellate | 0 | 0 | 1 | 1 | 0 | 1 | 0 | 1 |
| TRIM46 | -0.0551 | P1_P3_Lumbar_Sphenopalatine_VS_P2_P4_Coeliac_Stellate | 1 | 0 | 1 | 0 | 1 | 0 | 0 | 1 |
| ATE1 | -0.0553 | P3_P4_Coeliac_Sphenopalatine_VS_P1_P2_Lumbar_Stellate | 0 | 0 | 1 | 1 | 0 | 1 | 0 | 1 |
| PIK3IP1 | -0.0553 | P1_P2_P4_Lumbar_Coeliac_Stellate_Sphenopalatine_VS_P3 | 1 | 1 | 0 | 1 | 1 | 1 | 1 | 1 |
| SOCS7 | -0.0555 | P2_Lumbar_Sphenopalatine_VS_P1_P3_P4_Coeliac_Stellate | 0 | 1 | 0 | 0 | 1 | 0 | 0 | 1 |
| MRPS18B | -0.0556 | P1_P3_Lumbar_Sphenopalatine_VS_P2_P4_Coeliac_Stellate | 1 | 0 | 1 | 0 | 1 | 0 | 0 | 1 |
| MAPK9 | -0.0557 | P2_P3_Sphenopalatine_VS_P1_P4_Lumbar_Coeliac_Stellate | 0 | 1 | 1 | 0 | 0 | 0 | 0 | 1 |
| STIP1 | -0.056 | P3_P4_Coeliac_Sphenopalatine_VS_P1_P2_Lumbar_Stellate | 0 | 0 | 1 | 1 | 0 | 1 | 0 | 1 |
| CNOT6 | -0.0561 | P2_P3_P4_Lumbar_Coeliac_Stellate_VS_P1_Sphenopalatine | 0 | 1 | 1 | 1 | 1 | 1 | 1 | 0 |
| NIPSNAP2 | -0.0561 | P1_P2_P3_P4_Lumbar_Stellate_Sphenopalatine_VS_Coeliac | 1 | 1 | 1 | 1 | 1 | 0 | 1 | 1 |
| ARL2BP | -0.0563 | P1_P2_P3_P4_Lumbar_Stellate_Sphenopalatine_VS_Coeliac | 1 | 1 | 1 | 1 | 1 | 0 | 1 | 1 |

| | | | | | | | | | | |
|---|---|---|---|---|---|---|---|---|---|---|
| TTBK1 | -0.0564 | P2_Lumbar_Sphenopalatine_VS_P1_P3_P4_Coeliac_Stellate | 0 | 1 | 0 | 0 | 1 | 0 | 0 | 1 |
| CCDC91 | -0.0564 | P1_P2_P4_Lumbar_Coeliac_Stellate_Sphenopalatine_VS_P3 | 1 | 1 | 0 | 1 | 1 | 1 | 1 | 1 |
| RUFY2 | -0.0564 | P1_P2_P4_Lumbar_Coeliac_Stellate_Sphenopalatine_VS_P3 | 1 | 1 | 0 | 1 | 1 | 1 | 1 | 1 |
| CSE1L | -0.0565 | P2_P3_Coeliac_Sphenopalatine_VS_P1_P4_Lumbar_Stellate | 0 | 1 | 1 | 0 | 0 | 1 | 0 | 1 |
| D030056L22I | -0.0566 | P3_Lumbar_Sphenopalatine_VS_P1_P2_P4_Coeliac_Stellate | 0 | 0 | 1 | 0 | 1 | 0 | 0 | 1 |
| DROSHA | -0.0567 | P1_P2_P4_Lumbar_Coeliac_Stellate_VS_P3_Sphenopalatine | 1 | 1 | 0 | 1 | 1 | 1 | 1 | 0 |
| SLC4A8 | -0.0567 | P2_P3_P4_Lumbar_Coeliac_Stellate_VS_P1_Sphenopalatine | 0 | 1 | 1 | 1 | 1 | 1 | 1 | 0 |
| MAST2 | -0.0567 | P3_P4_Coeliac_Sphenopalatine_VS_P1_P2_Lumbar_Stellate | 0 | 0 | 1 | 1 | 0 | 1 | 0 | 1 |
| PRRC2A | -0.0567 | P3_P4_Coeliac_Sphenopalatine_VS_P1_P2_Lumbar_Stellate | 0 | 0 | 1 | 1 | 0 | 1 | 0 | 1 |
| ABHD12 | -0.0568 | P2_P3_Lumbar_Sphenopalatine_VS_P1_P4_Coeliac_Stellate | 0 | 1 | 1 | 0 | 1 | 0 | 0 | 1 |
| TM7SF3 | -0.057 | P2_P3_P4_Lumbar_Coeliac_Stellate_VS_P1_Sphenopalatine | 0 | 1 | 1 | 1 | 1 | 1 | 1 | 0 |
| FTO | -0.057 | P2_P3_Sphenopalatine_VS_P1_P4_Lumbar_Coeliac_Stellate | 0 | 1 | 1 | 0 | 0 | 0 | 0 | 1 |
| CCNT1 | -0.057 | P1_P2_P4_Lumbar_Coeliac_Stellate_Sphenopalatine_VS_P3 | 1 | 1 | 0 | 1 | 1 | 1 | 1 | 1 |
| SEPHS1 | -0.0571 | P1_P2_P3_Lumbar_Coeliac_Stellate_Sphenopalatine_VS_P4 | 1 | 1 | 1 | 0 | 1 | 1 | 1 | 1 |
| NR2C2AP | -0.0572 | P1_P3_Lumbar_Sphenopalatine_VS_P2_P4_Coeliac_Stellate | 1 | 0 | 1 | 0 | 1 | 0 | 0 | 1 |
| EXOC4 | -0.0572 | P3_P4_Coeliac_Sphenopalatine_VS_P1_P2_Lumbar_Stellate | 0 | 0 | 1 | 1 | 0 | 1 | 0 | 1 |
| RGS10 | -0.0572 | P2_P3_P4_Lumbar_Coeliac_Stellate_Sphenopalatine_VS_P1 | 0 | 1 | 1 | 1 | 1 | 1 | 1 | 1 |
| MAP9 | -0.0573 | P3_P4_Coeliac_Sphenopalatine_VS_P1_P2_Lumbar_Stellate | 0 | 0 | 1 | 1 | 0 | 1 | 0 | 1 |
| NDRG3 | -0.0573 | P1_P2_P3_P4_Lumbar_Stellate_Sphenopalatine_VS_Coeliac | 1 | 1 | 1 | 1 | 1 | 0 | 1 | 1 |
| RALGAPA1 | -0.0574 | P2_P3_Lumbar_Sphenopalatine_VS_P1_P4_Coeliac_Stellate | 0 | 1 | 1 | 0 | 1 | 0 | 0 | 1 |
| ANO6 | -0.0574 | P1_P2_Lumbar_Coeliac_Stellate_Sphenopalatine_VS_P3_P4 | 1 | 1 | 0 | 0 | 1 | 1 | 1 | 1 |
| ZFP266 | -0.0575 | P2_P3_P4_Lumbar_Coeliac_Stellate_VS_P1_Sphenopalatine | 0 | 1 | 1 | 1 | 1 | 1 | 1 | 0 |
| EIF2S1 | -0.0576 | P1_P4_Lumbar_Coeliac_Stellate_VS_P2_P3_Sphenopalatine | 1 | 0 | 0 | 1 | 1 | 1 | 1 | 0 |
| BLOC1S5 | -0.0576 | P1_P2_P3_P4_Lumbar_Stellate_Sphenopalatine_VS_Coeliac | 1 | 1 | 1 | 1 | 1 | 0 | 1 | 1 |
| RNF34 | -0.0577 | P2_P3_P4_Lumbar_Coeliac_Stellate_VS_P1_Sphenopalatine | 0 | 1 | 1 | 1 | 1 | 1 | 1 | 0 |
| GM15417 | -0.0577 | P2_P3_Sphenopalatine_VS_P1_P4_Lumbar_Coeliac_Stellate | 0 | 1 | 1 | 0 | 0 | 0 | 0 | 1 |
| LYRM9 | -0.0581 | P3_Sphenopalatine_VS_P1_P2_P4_Lumbar_Coeliac_Stellate | 0 | 0 | 1 | 0 | 0 | 0 | 0 | 1 |
| C2CD2L | -0.0581 | P2_P3_Lumbar_Sphenopalatine_VS_P1_P4_Coeliac_Stellate | 0 | 1 | 1 | 0 | 1 | 0 | 0 | 1 |
| AKR7A5 | -0.0582 | P1_P3_P4_Lumbar_Stellate_VS_P2_Coeliac_Sphenopalatine | 1 | 0 | 1 | 1 | 1 | 0 | 1 | 0 |
| DDR1 | -0.0583 | P3_Lumbar_Sphenopalatine_VS_P1_P2_P4_Coeliac_Stellate | 0 | 0 | 1 | 0 | 1 | 0 | 0 | 1 |

| | | | | | | | | | | |
|---|---|---|---|---|---|---|---|---|---|---|
| PPT1 | -0.0583 | P3_P4_Coeliac_Sphenopalatine_VS_P1_P2_Lumbar_Stellate | 0 | 0 | 1 | 1 | 0 | 1 | 0 | 1 |
| IGBP1 | -0.0584 | P2_P3_Lumbar_Sphenopalatine_VS_P1_P4_Coeliac_Stellate | 0 | 1 | 1 | 0 | 1 | 0 | 0 | 1 |
| ERLIN2 | -0.0584 | P3_P4_Coeliac_Sphenopalatine_VS_P1_P2_Lumbar_Stellate | 0 | 0 | 1 | 1 | 0 | 1 | 0 | 1 |
| HACD3 | -0.0587 | P2_P3_P4_Lumbar_Coeliac_Stellate_VS_P1_Sphenopalatine | 0 | 1 | 1 | 1 | 1 | 1 | 1 | 0 |
| KCNMA1 | -0.0587 | P3_Lumbar_Sphenopalatine_VS_P1_P2_P4_Coeliac_Stellate | 0 | 0 | 1 | 0 | 1 | 0 | 0 | 1 |
| AMFR | -0.0589 | P1_P2_P3_P4_Lumbar_Stellate_Sphenopalatine_VS_Coeliac | 1 | 1 | 1 | 1 | 1 | 0 | 1 | 1 |
| REEP2 | -0.0589 | P1_P2_P3_P4_Lumbar_Stellate_Sphenopalatine_VS_Coeliac | 1 | 1 | 1 | 1 | 1 | 0 | 1 | 1 |
| THTPA | -0.059 | P1_P3_Lumbar_Sphenopalatine_VS_P2_P4_Coeliac_Stellate | 1 | 0 | 1 | 0 | 1 | 0 | 0 | 1 |
| PCNX4 | -0.0591 | P3_P4_Coeliac_Sphenopalatine_VS_P1_P2_Lumbar_Stellate | 0 | 0 | 1 | 1 | 0 | 1 | 0 | 1 |
| JUN | -0.0592 | P1_P2_P4_Lumbar_Coeliac_Stellate_Sphenopalatine_VS_P3 | 1 | 1 | 0 | 1 | 1 | 1 | 1 | 1 |
| SCAP | -0.0595 | P1_P3_Lumbar_Sphenopalatine_VS_P2_P4_Coeliac_Stellate | 1 | 0 | 1 | 0 | 1 | 0 | 0 | 1 |
| THOC2 | -0.0596 | P2_P3_P4_Lumbar_Coeliac_Stellate_VS_P1_Sphenopalatine | 0 | 1 | 1 | 1 | 1 | 1 | 1 | 0 |
| GT(ROSA)26 | -0.0596 | P2_P3_Sphenopalatine_VS_P1_P4_Lumbar_Coeliac_Stellate | 0 | 1 | 1 | 0 | 0 | 0 | 0 | 1 |
| DCK | -0.0596 | P2_Lumbar_Sphenopalatine_VS_P1_P3_P4_Coeliac_Stellate | 0 | 1 | 0 | 0 | 1 | 0 | 0 | 1 |
| MPHOSPH6 | -0.0597 | P1_P4_Lumbar_Coeliac_Stellate_VS_P2_P3_Sphenopalatine | 1 | 0 | 0 | 1 | 1 | 1 | 1 | 0 |
| LZTR1 | -0.0597 | P2_Lumbar_Sphenopalatine_VS_P1_P3_P4_Coeliac_Stellate | 0 | 1 | 0 | 0 | 1 | 0 | 0 | 1 |
| TMEM218 | -0.0598 | P1_P2_P3_Lumbar_Coeliac_Stellate_VS_P4_Sphenopalatine | 1 | 1 | 1 | 0 | 1 | 1 | 1 | 0 |
| PTN | -0.0598 | P1_P2_P4_Lumbar_Coeliac_Stellate_Sphenopalatine_VS_P3 | 1 | 1 | 0 | 1 | 1 | 1 | 1 | 1 |
| CPEB4 | -0.0598 | P1_P2_P4_Lumbar_Coeliac_Stellate_Sphenopalatine_VS_P3 | 1 | 1 | 0 | 1 | 1 | 1 | 1 | 1 |
| CCDC30 | -0.0599 | P2_P3_P4_Lumbar_Coeliac_Stellate_VS_P1_Sphenopalatine | 0 | 1 | 1 | 1 | 1 | 1 | 1 | 0 |
| SHC1 | -0.06 | P2_P3_P4_Lumbar_Coeliac_Stellate_VS_P1_Sphenopalatine | 0 | 1 | 1 | 1 | 1 | 1 | 1 | 0 |
| ERLEC1 | -0.06 | P2_P3_P4_Lumbar_Coeliac_Stellate_VS_P1_Sphenopalatine | 0 | 1 | 1 | 1 | 1 | 1 | 1 | 0 |
| BCL2L1 | -0.0601 | P3_Lumbar_Sphenopalatine_VS_P1_P2_P4_Coeliac_Stellate | 0 | 0 | 1 | 0 | 1 | 0 | 0 | 1 |
| SNHG11 | -0.0601 | P1_P2_P3_P4_Lumbar_Stellate_Sphenopalatine_VS_Coeliac | 1 | 1 | 1 | 1 | 1 | 0 | 1 | 1 |
| LYPLA1 | -0.0603 | P1_P2_P3_Lumbar_Coeliac_Stellate_Sphenopalatine_VS_P4 | 1 | 1 | 1 | 0 | 1 | 1 | 1 | 1 |
| CLK1 | -0.0605 | P3_P4_Coeliac_Sphenopalatine_VS_P1_P2_Lumbar_Stellate | 0 | 0 | 1 | 1 | 0 | 1 | 0 | 1 |
| ZFP821 | -0.0606 | P3_Lumbar_Sphenopalatine_VS_P1_P2_P4_Coeliac_Stellate | 0 | 0 | 1 | 0 | 1 | 0 | 0 | 1 |
| ARMCX2 | -0.0607 | P3_Lumbar_Sphenopalatine_VS_P1_P2_P4_Coeliac_Stellate | 0 | 0 | 1 | 0 | 1 | 0 | 0 | 1 |
| MFSD6 | -0.0608 | P2_P3_Sphenopalatine_VS_P1_P4_Lumbar_Coeliac_Stellate | 0 | 1 | 1 | 0 | 0 | 0 | 0 | 1 |
| FAM168A | -0.0608 | P2_Coeliac_Sphenopalatine_VS_P1_P3_P4_Lumbar_Stellate | 0 | 1 | 0 | 0 | 0 | 1 | 0 | 1 |

| | | | | | | | | | | |
|---|---|---|---|---|---|---|---|---|---|---|
| DYNLT3 | -0.0609 | P1_P2_P3_P4_Lumbar_Stellate_Sphenopalatine_VS_Coeliac | 1 | 1 | 1 | 1 | 1 | 0 | 1 | 1 |
| MGAT5 | -0.0609 | P3_P4_Lumbar_Coeliac_Stellate_Sphenopalatine_VS_P1_P2 | 0 | 0 | 1 | 1 | 1 | 1 | 1 | 1 |
| NUDT11 | -0.061 | P2_P3_P4_Lumbar_Coeliac_Stellate_VS_P1_Sphenopalatine | 0 | 1 | 1 | 1 | 1 | 1 | 1 | 0 |
| YDJC | -0.0612 | P1_P3_Lumbar_Sphenopalatine_VS_P2_P4_Coeliac_Stellate | 1 | 0 | 1 | 0 | 1 | 0 | 0 | 1 |
| NFKBIE | -0.0613 | P1_P2_P3_Lumbar_Coeliac_Stellate_VS_P4_Sphenopalatine | 1 | 1 | 1 | 0 | 1 | 1 | 1 | 0 |
| ARPIN | -0.0613 | P1_P2_P3_Lumbar_Coeliac_Stellate_Sphenopalatine_VS_P4 | 1 | 1 | 1 | 0 | 1 | 1 | 1 | 1 |
| MYT1L | -0.0613 | P2_P3_P4_Lumbar_Coeliac_Stellate_Sphenopalatine_VS_P1 | 0 | 1 | 1 | 1 | 1 | 1 | 1 | 1 |
| VAMP3 | -0.0614 | P1_P2_P3_P4_Lumbar_Stellate_Sphenopalatine_VS_Coeliac | 1 | 1 | 1 | 1 | 1 | 0 | 1 | 1 |
| NDST1 | -0.0615 | P2_P3_Lumbar_Sphenopalatine_VS_P1_P4_Coeliac_Stellate | 0 | 1 | 1 | 0 | 1 | 0 | 0 | 1 |
| MTUS1 | -0.0616 | P2_P3_Sphenopalatine_VS_P1_P4_Lumbar_Coeliac_Stellate | 0 | 1 | 1 | 0 | 0 | 0 | 0 | 1 |
| LRRC8B | -0.0616 | P2_P3_Lumbar_Sphenopalatine_VS_P1_P4_Coeliac_Stellate | 0 | 1 | 1 | 0 | 1 | 0 | 0 | 1 |
| BLOC1S6 | -0.0617 | P3_Sphenopalatine_VS_P1_P2_P4_Lumbar_Coeliac_Stellate | 0 | 0 | 1 | 0 | 0 | 0 | 0 | 1 |
| CZIB | -0.0617 | P1_P3_Lumbar_Sphenopalatine_VS_P2_P4_Coeliac_Stellate | 1 | 0 | 1 | 0 | 1 | 0 | 0 | 1 |
| RBFA | -0.0617 | P1_P3_Lumbar_Sphenopalatine_VS_P2_P4_Coeliac_Stellate | 1 | 0 | 1 | 0 | 1 | 0 | 0 | 1 |
| AGAP1 | -0.0617 | P2_P3_P4_Lumbar_Coeliac_Stellate_Sphenopalatine_VS_P1 | 0 | 1 | 1 | 1 | 1 | 1 | 1 | 1 |
| STK39 | -0.0618 | P1_P2_P3_P4_Lumbar_Stellate_VS_Coeliac_Sphenopalatine | 1 | 1 | 1 | 1 | 1 | 0 | 1 | 0 |
| KCNC4 | -0.0619 | P2_P3_P4_Lumbar_Coeliac_Stellate_Sphenopalatine_VS_P1 | 0 | 1 | 1 | 1 | 1 | 1 | 1 | 1 |
| NLGN2 | -0.062 | P2_P3_Lumbar_Sphenopalatine_VS_P1_P4_Coeliac_Stellate | 0 | 1 | 1 | 0 | 1 | 0 | 0 | 1 |
| CDK5 | -0.062 | P1_P2_P3_P4_Lumbar_Stellate_Sphenopalatine_VS_Coeliac | 1 | 1 | 1 | 1 | 1 | 0 | 1 | 1 |
| PSMB10 | -0.062 | P1_P2_P3_P4_Lumbar_Stellate_Sphenopalatine_VS_Coeliac | 1 | 1 | 1 | 1 | 1 | 0 | 1 | 1 |
| CSTB | -0.0621 | P1_P2_P3_P4_Lumbar_Stellate_Sphenopalatine_VS_Coeliac | 1 | 1 | 1 | 1 | 1 | 0 | 1 | 1 |
| ZFP414 | -0.0621 | P1_P2_P3_P4_Lumbar_Stellate_Sphenopalatine_VS_Coeliac | 1 | 1 | 1 | 1 | 1 | 0 | 1 | 1 |
| CDK2AP2 | -0.0621 | P1_P2_P3_P4_Lumbar_Stellate_Sphenopalatine_VS_Coeliac | 1 | 1 | 1 | 1 | 1 | 0 | 1 | 1 |
| INO80E | -0.0623 | P1_P3_Lumbar_Sphenopalatine_VS_P2_P4_Coeliac_Stellate | 1 | 0 | 1 | 0 | 1 | 0 | 0 | 1 |
| PURB | -0.0625 | P2_Coeliac_Sphenopalatine_VS_P1_P3_P4_Lumbar_Stellate | 0 | 1 | 0 | 0 | 0 | 1 | 0 | 1 |
| DCAF12 | -0.0626 | P1_P2_P3_P4_Lumbar_Stellate_Sphenopalatine_VS_Coeliac | 1 | 1 | 1 | 1 | 1 | 0 | 1 | 1 |
| PNPO | -0.0629 | P1_P2_P3_Lumbar_Coeliac_Stellate_Sphenopalatine_VS_P4 | 1 | 1 | 1 | 0 | 1 | 1 | 1 | 1 |
| ZNHIT1 | -0.0631 | P1_P2_P3_P4_Lumbar_Stellate_Sphenopalatine_VS_Coeliac | 1 | 1 | 1 | 1 | 1 | 0 | 1 | 1 |
| IDNK | -0.0632 | P3_Sphenopalatine_VS_P1_P2_P4_Lumbar_Coeliac_Stellate | 0 | 0 | 1 | 0 | 0 | 0 | 0 | 1 |
| SNX27 | -0.0632 | P2_P3_Sphenopalatine_VS_P1_P4_Lumbar_Coeliac_Stellate | 0 | 1 | 1 | 0 | 0 | 0 | 0 | 1 |

| | | | | | | | | | | |
|---|---|---|---|---|---|---|---|---|---|---|
| SURF2 | -0.0632 | P1_P3_Lumbar_Sphenopalatine_VS_P2_P4_Coeliac_Stellate | 1 | 0 | 1 | 0 | 1 | 0 | 0 | 1 |
| TARBP2 | -0.0632 | P1_P3_Lumbar_Sphenopalatine_VS_P2_P4_Coeliac_Stellate | 1 | 0 | 1 | 0 | 1 | 0 | 0 | 1 |
| COX11 | -0.0634 | P1_P2_P3_P4_Lumbar_Stellate_Sphenopalatine_VS_Coeliac | 1 | 1 | 1 | 1 | 1 | 0 | 1 | 1 |
| DNAL4 | -0.0635 | P2_P3_P4_Lumbar_Coeliac_Stellate_VS_P1_Sphenopalatine | 0 | 1 | 1 | 1 | 1 | 1 | 1 | 0 |
| RRP36 | -0.0637 | P1_P3_Lumbar_Coeliac_Stellate_VS_P2_P4_Sphenopalatine | 1 | 0 | 1 | 0 | 1 | 1 | 1 | 0 |
| FBXO21 | -0.0637 | P2_P3_Sphenopalatine_VS_P1_P4_Lumbar_Coeliac_Stellate | 0 | 1 | 1 | 0 | 0 | 0 | 0 | 1 |
| WBP1 | -0.0638 | P1_P2_P3_P4_Lumbar_Stellate_Sphenopalatine_VS_Coeliac | 1 | 1 | 1 | 1 | 1 | 0 | 1 | 1 |
| CARHSP1 | -0.0638 | P1_P2_P3_Lumbar_Coeliac_Stellate_Sphenopalatine_VS_P4 | 1 | 1 | 1 | 0 | 1 | 1 | 1 | 1 |
| RGS17 | -0.0639 | P1_P2_P3_P4_Lumbar_Stellate_Sphenopalatine_VS_Coeliac | 1 | 1 | 1 | 1 | 1 | 0 | 1 | 1 |
| SRCIN1 | -0.0642 | P2_P3_Lumbar_Sphenopalatine_VS_P1_P4_Coeliac_Stellate | 0 | 1 | 1 | 0 | 1 | 0 | 0 | 1 |
| ADAM19 | -0.0644 | P2_P3_P4_Lumbar_Stellate_VS_P1_Coeliac_Sphenopalatine | 0 | 1 | 1 | 1 | 1 | 0 | 1 | 0 |
| MINDY2 | -0.0645 | P2_P3_Lumbar_Sphenopalatine_VS_P1_P4_Coeliac_Stellate | 0 | 1 | 1 | 0 | 1 | 0 | 0 | 1 |
| UBE2O | -0.0645 | P1_P2_Coeliac_Sphenopalatine_VS_P3_P4_Lumbar_Stellate | 1 | 1 | 0 | 0 | 0 | 1 | 0 | 1 |
| TSTD3 | -0.0646 | P1_P2_P3_Lumbar_Coeliac_Stellate_Sphenopalatine_VS_P4 | 1 | 1 | 1 | 0 | 1 | 1 | 1 | 1 |
| TPD52 | -0.0649 | P1_P3_P4_Lumbar_Stellate_Sphenopalatine_VS_P2_Coeliac | 1 | 0 | 1 | 1 | 1 | 0 | 1 | 1 |
| NIP7 | -0.065 | P1_P3_Lumbar_Coeliac_Stellate_VS_P2_P4_Sphenopalatine | 1 | 0 | 1 | 0 | 1 | 1 | 1 | 0 |
| LMBRD2 | -0.065 | P2_P3_Lumbar_Sphenopalatine_VS_P1_P4_Coeliac_Stellate | 0 | 1 | 1 | 0 | 1 | 0 | 0 | 1 |
| TRAPPC2 | -0.0651 | P1_P2_P3_P4_Lumbar_Stellate_Sphenopalatine_VS_Coeliac | 1 | 1 | 1 | 1 | 1 | 0 | 1 | 1 |
| LYPLA2 | -0.0652 | P1_P3_Lumbar_Sphenopalatine_VS_P2_P4_Coeliac_Stellate | 1 | 0 | 1 | 0 | 1 | 0 | 0 | 1 |
| HIBADH | -0.0654 | P2_P3_P4_Lumbar_Coeliac_Stellate_VS_P1_Sphenopalatine | 0 | 1 | 1 | 1 | 1 | 1 | 1 | 0 |
| KANSL3 | -0.0654 | P2_P3_Lumbar_Sphenopalatine_VS_P1_P4_Coeliac_Stellate | 0 | 1 | 1 | 0 | 1 | 0 | 0 | 1 |
| MEA1 | -0.0654 | P1_P2_P3_P4_Lumbar_Stellate_Sphenopalatine_VS_Coeliac | 1 | 1 | 1 | 1 | 1 | 0 | 1 | 1 |
| NUDCD2 | -0.0655 | P1_P3_Lumbar_Coeliac_Stellate_VS_P2_P4_Sphenopalatine | 1 | 0 | 1 | 0 | 1 | 1 | 1 | 0 |
| DHCR7 | -0.0657 | P3_Lumbar_Sphenopalatine_VS_P1_P2_P4_Coeliac_Stellate | 0 | 0 | 1 | 0 | 1 | 0 | 0 | 1 |
| ISCU | -0.0657 | P1_P2_P3_P4_Lumbar_Stellate_Sphenopalatine_VS_Coeliac | 1 | 1 | 1 | 1 | 1 | 0 | 1 | 1 |
| DLC1 | -0.0659 | P1_P2_Coeliac_Sphenopalatine_VS_P3_P4_Lumbar_Stellate | 1 | 1 | 0 | 0 | 0 | 1 | 0 | 1 |
| LYPD6 | -0.066 | P4_Coeliac_Stellate_VS_P1_P2_P3_Lumbar_Sphenopalatine | 0 | 0 | 0 | 1 | 0 | 1 | 1 | 0 |
| WASHC3 | -0.0661 | P1_P2_P3_P4_Lumbar_Stellate_VS_Coeliac_Sphenopalatine | 1 | 1 | 1 | 1 | 1 | 0 | 1 | 0 |
| ACTR1B | -0.0663 | P1_P2_P3_P4_Lumbar_Stellate_Sphenopalatine_VS_Coeliac | 1 | 1 | 1 | 1 | 1 | 0 | 1 | 1 |
| SLC2A3 | -0.0663 | P1_P2_P3_P4_Lumbar_Stellate_Sphenopalatine_VS_Coeliac | 1 | 1 | 1 | 1 | 1 | 0 | 1 | 1 |

| | | | | | | | | | | |
|---|---|---|---|---|---|---|---|---|---|---|
| INAFM1 | -0.0667 | P1_P2_P3_P4_Lumbar_Stellate_Sphenopalatine_VS_Coeliac | 1 | 1 | 1 | 1 | 1 | 0 | 1 | 1 |
| EIF2B1 | -0.0668 | P1_P3_Lumbar_Coeliac_Stellate_VS_P2_P4_Sphenopalatine | 1 | 0 | 1 | 0 | 1 | 1 | 1 | 0 |
| KLHDC3 | -0.0669 | P3_Sphenopalatine_VS_P1_P2_P4_Lumbar_Coeliac_Stellate | 0 | 0 | 1 | 0 | 0 | 0 | 0 | 1 |
| PLEKHA6 | -0.067 | P2_P3_P4_Lumbar_Coeliac_Stellate_VS_P1_Sphenopalatine | 0 | 1 | 1 | 1 | 1 | 1 | 1 | 0 |
| NOP53 | -0.0671 | P1_P2_P3_P4_Lumbar_Stellate_Sphenopalatine_VS_Coeliac | 1 | 1 | 1 | 1 | 1 | 0 | 1 | 1 |
| MYL12A | -0.0675 | P1_P2_P3_P4_Coeliac_Stellate_VS_Lumbar_Sphenopalatine | 1 | 1 | 1 | 1 | 0 | 1 | 1 | 0 |
| EAF1 | -0.0676 | P2_P3_Lumbar_Sphenopalatine_VS_P1_P4_Coeliac_Stellate | 0 | 1 | 1 | 0 | 1 | 0 | 0 | 1 |
| CEP164 | -0.0677 | P2_P3_Lumbar_Sphenopalatine_VS_P1_P4_Coeliac_Stellate | 0 | 1 | 1 | 0 | 1 | 0 | 0 | 1 |
| VAMP4 | -0.0677 | P1_P2_P3_P4_Lumbar_Stellate_Sphenopalatine_VS_Coeliac | 1 | 1 | 1 | 1 | 1 | 0 | 1 | 1 |
| BLOC1S2 | -0.0678 | P1_P2_P3_P4_Lumbar_Stellate_Sphenopalatine_VS_Coeliac | 1 | 1 | 1 | 1 | 1 | 0 | 1 | 1 |
| PDHA1 | -0.068 | P1_P2_P3_P4_Lumbar_Stellate_Sphenopalatine_VS_Coeliac | 1 | 1 | 1 | 1 | 1 | 0 | 1 | 1 |
| ZMAT3 | -0.0685 | P2_P3_Lumbar_Sphenopalatine_VS_P1_P4_Coeliac_Stellate | 0 | 1 | 1 | 0 | 1 | 0 | 0 | 1 |
| RNF157 | -0.0688 | P2_P3_Lumbar_Sphenopalatine_VS_P1_P4_Coeliac_Stellate | 0 | 1 | 1 | 0 | 1 | 0 | 0 | 1 |
| NYAP2 | -0.0691 | P2_P3_P4_Lumbar_Coeliac_Stellate_VS_P1_Sphenopalatine | 0 | 1 | 1 | 1 | 1 | 1 | 1 | 0 |
| CNDP2 | -0.0691 | P3_Lumbar_Sphenopalatine_VS_P1_P2_P4_Coeliac_Stellate | 0 | 0 | 1 | 0 | 1 | 0 | 0 | 1 |
| ENTR1 | -0.0691 | P1_P2_P3_P4_Lumbar_Stellate_Sphenopalatine_VS_Coeliac | 1 | 1 | 1 | 1 | 1 | 0 | 1 | 1 |
| DMXL2 | -0.0692 | P2_P3_Lumbar_Sphenopalatine_VS_P1_P4_Coeliac_Stellate | 0 | 1 | 1 | 0 | 1 | 0 | 0 | 1 |
| WDR89 | -0.0694 | P2_P3_Lumbar_Sphenopalatine_VS_P1_P4_Coeliac_Stellate | 0 | 1 | 1 | 0 | 1 | 0 | 0 | 1 |
| HSPA4L | -0.0695 | P2_P3_Lumbar_Sphenopalatine_VS_P1_P4_Coeliac_Stellate | 0 | 1 | 1 | 0 | 1 | 0 | 0 | 1 |
| RIT1 | -0.0696 | P1_P2_P3_P4_Lumbar_Stellate_Sphenopalatine_VS_Coeliac | 1 | 1 | 1 | 1 | 1 | 0 | 1 | 1 |
| OCRL | -0.0697 | P2_P3_Lumbar_Sphenopalatine_VS_P1_P4_Coeliac_Stellate | 0 | 1 | 1 | 0 | 1 | 0 | 0 | 1 |
| BCOR | -0.0698 | P2_P4_Lumbar_Coeliac_Stellate_VS_P1_P3_Sphenopalatine | 0 | 1 | 0 | 1 | 1 | 1 | 1 | 0 |
| CKMT1 | -0.07 | P2_P3_Lumbar_Sphenopalatine_VS_P1_P4_Coeliac_Stellate | 0 | 1 | 1 | 0 | 1 | 0 | 0 | 1 |
| LDAH | -0.0702 | P2_P3_Lumbar_Coeliac_Stellate_VS_P1_P4_Sphenopalatine | 0 | 1 | 1 | 0 | 1 | 1 | 1 | 0 |
| 2510002D24I | -0.0702 | P3_Lumbar_Sphenopalatine_VS_P1_P2_P4_Coeliac_Stellate | 0 | 0 | 1 | 0 | 1 | 0 | 0 | 1 |
| PLPPR2 | -0.0702 | P2_P3_P4_Lumbar_Coeliac_Stellate_Sphenopalatine_VS_P1 | 0 | 1 | 1 | 1 | 1 | 1 | 1 | 1 |
| JTB | -0.0704 | P1_P2_P3_P4_Lumbar_Stellate_Sphenopalatine_VS_Coeliac | 1 | 1 | 1 | 1 | 1 | 0 | 1 | 1 |
| ATF5 | -0.0704 | P1_P2_P3_P4_Lumbar_Stellate_Sphenopalatine_VS_Coeliac | 1 | 1 | 1 | 1 | 1 | 0 | 1 | 1 |
| MTSS1 | -0.0708 | P2_P3_Coeliac_Sphenopalatine_VS_P1_P4_Lumbar_Stellate | 0 | 1 | 1 | 0 | 0 | 1 | 0 | 1 |
| RASSF1 | -0.0708 | P1_P2_P4_Lumbar_Coeliac_Stellate_Sphenopalatine_VS_P3 | 1 | 1 | 0 | 1 | 1 | 1 | 1 | 1 |

| | | | | | | | | | | |
|---|---|---|---|---|---|---|---|---|---|---|
| RIT2 | -0.071 | P2_P3_P4_Lumbar_Coeliac_Stellate_VS_P1_Sphenopalatine | 0 | 1 | 1 | 1 | 1 | 1 | 1 | 0 |
| PNKD | -0.0712 | P1_P2_P3_P4_Lumbar_Stellate_Sphenopalatine_VS_Coeliac | 1 | 1 | 1 | 1 | 1 | 0 | 1 | 1 |
| SRSF9 | -0.0715 | P2_P3_P4_Lumbar_Coeliac_Stellate_VS_P1_Sphenopalatine | 0 | 1 | 1 | 1 | 1 | 1 | 1 | 0 |
| RSBN1 | -0.0715 | P3_Lumbar_Sphenopalatine_VS_P1_P2_P4_Coeliac_Stellate | 0 | 0 | 1 | 0 | 1 | 0 | 0 | 1 |
| BORCS7 | -0.0715 | P1_P2_P3_P4_Lumbar_Stellate_Sphenopalatine_VS_Coeliac | 1 | 1 | 1 | 1 | 1 | 0 | 1 | 1 |
| SGSM1 | -0.0716 | P2_P3_P4_Lumbar_Coeliac_Stellate_VS_P1_Sphenopalatine | 0 | 1 | 1 | 1 | 1 | 1 | 1 | 0 |
| MVD | -0.0717 | P1_P2_P3_P4_Lumbar_Stellate_Sphenopalatine_VS_Coeliac | 1 | 1 | 1 | 1 | 1 | 0 | 1 | 1 |
| DCAF11 | -0.0718 | P1_P2_P3_Lumbar_Coeliac_Stellate_VS_P4_Sphenopalatine | 1 | 1 | 1 | 0 | 1 | 1 | 1 | 0 |
| FAF1 | -0.0718 | P3_Lumbar_Sphenopalatine_VS_P1_P2_P4_Coeliac_Stellate | 0 | 0 | 1 | 0 | 1 | 0 | 0 | 1 |
| SPAG7 | -0.0718 | P3_Lumbar_Sphenopalatine_VS_P1_P2_P4_Coeliac_Stellate | 0 | 0 | 1 | 0 | 1 | 0 | 0 | 1 |
| LANCL1 | -0.0724 | P2_P3_P4_Lumbar_Coeliac_Stellate_VS_P1_Sphenopalatine | 0 | 1 | 1 | 1 | 1 | 1 | 1 | 0 |
| UBE2Z | -0.0725 | P1_P2_P3_P4_Lumbar_Stellate_Sphenopalatine_VS_Coeliac | 1 | 1 | 1 | 1 | 1 | 0 | 1 | 1 |
| GM10076 | -0.0727 | P1_P2_P3_P4_Lumbar_Stellate_Sphenopalatine_VS_Coeliac | 1 | 1 | 1 | 1 | 1 | 0 | 1 | 1 |
| SULT4A1 | -0.0729 | P1_P2_P3_P4_Lumbar_Stellate_Sphenopalatine_VS_Coeliac | 1 | 1 | 1 | 1 | 1 | 0 | 1 | 1 |
| NDUFS1 | -0.0729 | P1_P2_P4_Lumbar_Coeliac_Stellate_Sphenopalatine_VS_P3 | 1 | 1 | 0 | 1 | 1 | 1 | 1 | 1 |
| PGM1 | -0.0731 | P1_P2_P3_P4_Lumbar_Stellate_Sphenopalatine_VS_Coeliac | 1 | 1 | 1 | 1 | 1 | 0 | 1 | 1 |
| RETREG2 | -0.0733 | P1_P2_P3_P4_Lumbar_Stellate_Sphenopalatine_VS_Coeliac | 1 | 1 | 1 | 1 | 1 | 0 | 1 | 1 |
| DHCR24 | -0.0733 | P1_P2_P3_P4_Lumbar_Stellate_Sphenopalatine_VS_Coeliac | 1 | 1 | 1 | 1 | 1 | 0 | 1 | 1 |
| CEND1 | -0.0736 | P1_P2_P3_P4_Lumbar_Stellate_Sphenopalatine_VS_Coeliac | 1 | 1 | 1 | 1 | 1 | 0 | 1 | 1 |
| CBX1 | -0.0736 | P3_Lumbar_Coeliac_Stellate_Sphenopalatine_VS_P1_P2_P4 | 0 | 0 | 1 | 0 | 1 | 1 | 1 | 1 |
| ZDHHC7 | -0.0738 | P1_P2_P3_P4_Lumbar_Stellate_Sphenopalatine_VS_Coeliac | 1 | 1 | 1 | 1 | 1 | 0 | 1 | 1 |
| VASP | -0.0742 | P1_P2_P3_Lumbar_Coeliac_Stellate_VS_P4_Sphenopalatine | 1 | 1 | 1 | 0 | 1 | 1 | 1 | 0 |
| LAMTOR4 | -0.0742 | P1_P2_P3_P4_Lumbar_Stellate_Sphenopalatine_VS_Coeliac | 1 | 1 | 1 | 1 | 1 | 0 | 1 | 1 |
| RSRC1 | -0.0743 | P3_Coeliac_Sphenopalatine_VS_P1_P2_P4_Lumbar_Stellate | 0 | 0 | 1 | 0 | 0 | 1 | 0 | 1 |
| SS18L2 | -0.0743 | P1_P2_P3_P4_Lumbar_Stellate_Sphenopalatine_VS_Coeliac | 1 | 1 | 1 | 1 | 1 | 0 | 1 | 1 |
| ZFP148 | -0.0744 | P2_P4_Lumbar_Coeliac_Stellate_VS_P1_P3_Sphenopalatine | 0 | 1 | 0 | 1 | 1 | 1 | 1 | 0 |
| TSPAN31 | -0.0746 | P1_P2_P3_P4_Lumbar_Stellate_Sphenopalatine_VS_Coeliac | 1 | 1 | 1 | 1 | 1 | 0 | 1 | 1 |
| 1110008P14I | -0.0749 | P1_P2_P3_P4_Lumbar_Stellate_Sphenopalatine_VS_Coeliac | 1 | 1 | 1 | 1 | 1 | 0 | 1 | 1 |
| TPGS2 | -0.0752 | P1_P2_P3_P4_Lumbar_Stellate_Sphenopalatine_VS_Coeliac | 1 | 1 | 1 | 1 | 1 | 0 | 1 | 1 |
| ABLIM1 | -0.0753 | P1_P2_P3_P4_Lumbar_Stellate_Sphenopalatine_VS_Coeliac | 1 | 1 | 1 | 1 | 1 | 0 | 1 | 1 |

| | | | | | | | | | | |
|---|---|---|---|---|---|---|---|---|---|---|
| GM45495 | -0.0754 | P1_P2_P3_Lumbar_Coeliac_Stellate_Sphenopalatine_VS_P4 | 1 | 1 | 1 | 0 | 1 | 1 | 1 | 1 |
| CHD9 | -0.0758 | Coeliac_Sphenopalatine_VS_P1_P2_P3_P4_Lumbar_Stellate | 0 | 0 | 0 | 0 | 0 | 1 | 0 | 1 |
| CDC27 | -0.0758 | P1_P2_P4_Lumbar_Coeliac_Stellate_Sphenopalatine_VS_P3 | 1 | 1 | 0 | 1 | 1 | 1 | 1 | 1 |
| SCAMP1 | -0.076 | P1_P2_P3_P4_Stellate_Sphenopalatine_VS_Lumbar_Coeliac | 1 | 1 | 1 | 1 | 0 | 0 | 1 | 1 |
| S100A11 | -0.0764 | P1_P2_P3_P4_Lumbar_Stellate_Sphenopalatine_VS_Coeliac | 1 | 1 | 1 | 1 | 1 | 0 | 1 | 1 |
| MAPK8IP3 | -0.0766 | P2_Coeliac_Sphenopalatine_VS_P1_P3_P4_Lumbar_Stellate | 0 | 1 | 0 | 0 | 0 | 1 | 0 | 1 |
| ZFAND3 | -0.0766 | P1_P2_P3_P4_Lumbar_Stellate_Sphenopalatine_VS_Coeliac | 1 | 1 | 1 | 1 | 1 | 0 | 1 | 1 |
| GSR | -0.0767 | P1_P2_P3_P4_Lumbar_Stellate_Sphenopalatine_VS_Coeliac | 1 | 1 | 1 | 1 | 1 | 0 | 1 | 1 |
| SFXN3 | -0.0771 | P1_P2_P3_P4_Lumbar_Stellate_Sphenopalatine_VS_Coeliac | 1 | 1 | 1 | 1 | 1 | 0 | 1 | 1 |
| 3110040N11 | -0.0773 | P1_P3_P4_Lumbar_Stellate_Sphenopalatine_VS_P2_Coeliac | 1 | 0 | 1 | 1 | 1 | 0 | 1 | 1 |
| SNX5 | -0.0777 | P2_P3_Coeliac_Sphenopalatine_VS_P1_P4_Lumbar_Stellate | 0 | 1 | 1 | 0 | 0 | 1 | 0 | 1 |
| E130307A14 | -0.0778 | P2_P3_Coeliac_Sphenopalatine_VS_P1_P4_Lumbar_Stellate | 0 | 1 | 1 | 0 | 0 | 1 | 0 | 1 |
| MRPL14 | -0.0779 | P1_P2_P3_P4_Lumbar_Stellate_Sphenopalatine_VS_Coeliac | 1 | 1 | 1 | 1 | 1 | 0 | 1 | 1 |
| RHOC | -0.0782 | P1_P2_P3_P4_Lumbar_Stellate_Sphenopalatine_VS_Coeliac | 1 | 1 | 1 | 1 | 1 | 0 | 1 | 1 |
| TIMM29 | -0.0782 | P1_P2_P3_P4_Lumbar_Stellate_Sphenopalatine_VS_Coeliac | 1 | 1 | 1 | 1 | 1 | 0 | 1 | 1 |
| RABGGTB | -0.0783 | P1_P2_P3_P4_Lumbar_Stellate_Sphenopalatine_VS_Coeliac | 1 | 1 | 1 | 1 | 1 | 0 | 1 | 1 |
| ZFHX2 | -0.0784 | P2_P4_Lumbar_Coeliac_Stellate_Sphenopalatine_VS_P1_P3 | 0 | 1 | 0 | 1 | 1 | 1 | 1 | 1 |
| MRPL17 | -0.0785 | P1_P2_P3_P4_Lumbar_Stellate_Sphenopalatine_VS_Coeliac | 1 | 1 | 1 | 1 | 1 | 0 | 1 | 1 |
| CCM2 | -0.0786 | P1_P2_P3_P4_Lumbar_Stellate_Sphenopalatine_VS_Coeliac | 1 | 1 | 1 | 1 | 1 | 0 | 1 | 1 |
| TRAPPC1 | -0.0786 | P1_P2_P3_P4_Lumbar_Stellate_Sphenopalatine_VS_Coeliac | 1 | 1 | 1 | 1 | 1 | 0 | 1 | 1 |
| CSRP1 | -0.0788 | P1_P2_P3_P4_Lumbar_Stellate_Sphenopalatine_VS_Coeliac | 1 | 1 | 1 | 1 | 1 | 0 | 1 | 1 |
| FDX2 | -0.0795 | P1_P2_P3_P4_Lumbar_Stellate_Sphenopalatine_VS_Coeliac | 1 | 1 | 1 | 1 | 1 | 0 | 1 | 1 |
| TPR | -0.0802 | Coeliac_Sphenopalatine_VS_P1_P2_P3_P4_Lumbar_Stellate | 0 | 0 | 0 | 0 | 0 | 1 | 0 | 1 |
| GATC | -0.0803 | P3_Coeliac_Sphenopalatine_VS_P1_P2_P4_Lumbar_Stellate | 0 | 0 | 1 | 0 | 0 | 1 | 0 | 1 |
| RBCK1 | -0.0806 | P1_P2_P3_P4_Lumbar_Stellate_Sphenopalatine_VS_Coeliac | 1 | 1 | 1 | 1 | 1 | 0 | 1 | 1 |
| YOD1 | -0.0806 | P1_P2_P3_Lumbar_Coeliac_Stellate_Sphenopalatine_VS_P4 | 1 | 1 | 1 | 0 | 1 | 1 | 1 | 1 |
| TAB2 | -0.0809 | P1_P2_P4_Lumbar_Coeliac_Stellate_Sphenopalatine_VS_P3 | 1 | 1 | 0 | 1 | 1 | 1 | 1 | 1 |
| NLN | -0.0811 | P1_P2_P3_Lumbar_Coeliac_Stellate_VS_P4_Sphenopalatine | 1 | 1 | 1 | 0 | 1 | 1 | 1 | 0 |
| ARL8B | -0.0811 | P1_P2_P3_P4_Lumbar_Stellate_Sphenopalatine_VS_Coeliac | 1 | 1 | 1 | 1 | 1 | 0 | 1 | 1 |
| TMEM198 | -0.0812 | P1_P2_P3_Lumbar_Coeliac_Stellate_VS_P4_Sphenopalatine | 1 | 1 | 1 | 0 | 1 | 1 | 1 | 0 |

| | | | | | | | | | | |
|---|---|---|---|---|---|---|---|---|---|---|
| MPV17 | -0.081 | P2_P3_P4_Lumbar_Coeliac_Stellate_Sphenopalatine_VS_P1 | 0 | 1 | 1 | 1 | 1 | 1 | 1 | 1 |
| GLUL | -0.081 | P2_P4_Lumbar_Coeliac_Stellate_VS_P1_P3_Sphenopalatine | 0 | 1 | 0 | 1 | 1 | 1 | 1 | 0 |
| ACADL | -0.081 | P1_P2_P3_P4_Lumbar_Stellate_VS_Coeliac_Sphenopalatine | 1 | 1 | 1 | 1 | 1 | 0 | 1 | 0 |
| SDHAF4 | -0.081 | P3_P4_Lumbar_Stellate_Sphenopalatine_VS_P1_P2_Coeliac | 0 | 0 | 1 | 1 | 1 | 0 | 1 | 1 |
| HAX1 | -0.081 | P1_P2_P3_P4_Lumbar_Stellate_Sphenopalatine_VS_Coeliac | 1 | 1 | 1 | 1 | 1 | 0 | 1 | 1 |
| PYGB | -0.082 | P1_P2_P3_P4_Lumbar_Stellate_Sphenopalatine_VS_Coeliac | 1 | 1 | 1 | 1 | 1 | 0 | 1 | 1 |
| 1110065P20I | -0.082 | P1_P2_P3_P4_Lumbar_Stellate_Sphenopalatine_VS_Coeliac | 1 | 1 | 1 | 1 | 1 | 0 | 1 | 1 |
| PSD | -0.082 | P2_P4_Lumbar_Coeliac_Stellate_Sphenopalatine_VS_P1_P3 | 0 | 1 | 0 | 1 | 1 | 1 | 1 | 1 |
| MEST | -0.082 | P1_P2_P3_P4_Lumbar_Stellate_Sphenopalatine_VS_Coeliac | 1 | 1 | 1 | 1 | 1 | 0 | 1 | 1 |
| DUSP22 | -0.082 | P1_P2_P3_P4_Lumbar_Stellate_Sphenopalatine_VS_Coeliac | 1 | 1 | 1 | 1 | 1 | 0 | 1 | 1 |
| MMD | -0.082 | P1_P2_P3_P4_Lumbar_Stellate_Sphenopalatine_VS_Coeliac | 1 | 1 | 1 | 1 | 1 | 0 | 1 | 1 |
| MNAT1 | -0.083 | P1_P4_Lumbar_Coeliac_Stellate_Sphenopalatine_VS_P2_P3 | 1 | 0 | 0 | 1 | 1 | 1 | 1 | 1 |
| TMEM106C | -0.083 | P1_P2_P3_P4_Stellate_Sphenopalatine_VS_Lumbar_Coeliac | 1 | 1 | 1 | 1 | 0 | 0 | 1 | 1 |
| NREP | -0.083 | P3_P4_Lumbar_Coeliac_Stellate_Sphenopalatine_VS_P1_P2 | 0 | 0 | 1 | 1 | 1 | 1 | 1 | 1 |
| MAGEH1 | -0.083 | P3_Lumbar_Stellate_VS_P1_P2_P4_Coeliac_Sphenopalatine | 0 | 0 | 1 | 0 | 1 | 0 | 1 | 0 |
| CHP1 | -0.083 | P1_P2_P3_P4_Lumbar_Stellate_Sphenopalatine_VS_Coeliac | 1 | 1 | 1 | 1 | 1 | 0 | 1 | 1 |
| OLFM2 | -0.083 | P2_P3_P4_Lumbar_Stellate_VS_P1_Coeliac_Sphenopalatine | 0 | 1 | 1 | 1 | 1 | 0 | 1 | 0 |
| METTL23 | -0.083 | P1_P2_P3_P4_Lumbar_Stellate_Sphenopalatine_VS_Coeliac | 1 | 1 | 1 | 1 | 1 | 0 | 1 | 1 |
| OSBPL9 | -0.084 | P2_P4_Lumbar_Coeliac_Stellate_VS_P1_P3_Sphenopalatine | 0 | 1 | 0 | 1 | 1 | 1 | 1 | 0 |
| ARL5A | -0.084 | P2_P3_Coeliac_Sphenopalatine_VS_P1_P4_Lumbar_Stellate | 0 | 1 | 1 | 0 | 0 | 1 | 0 | 1 |
| ROGDI | -0.084 | P3_P4_Lumbar_Stellate_VS_P1_P2_Coeliac_Sphenopalatine | 0 | 0 | 1 | 1 | 1 | 0 | 1 | 0 |
| WDR61 | -0.084 | P1_P3_Lumbar_Coeliac_Stellate_Sphenopalatine_VS_P2_P4 | 1 | 0 | 1 | 0 | 1 | 1 | 1 | 1 |
| ICK | -0.084 | P1_P2_P3_Lumbar_Coeliac_Stellate_Sphenopalatine_VS_P4 | 1 | 1 | 1 | 0 | 1 | 1 | 1 | 1 |
| AK1 | -0.084 | P3_P4_Stellate_VS_P1_P2_Lumbar_Coeliac_Sphenopalatine | 0 | 0 | 1 | 1 | 0 | 0 | 1 | 0 |
| PEG3 | -0.084 | P2_P3_Coeliac_Sphenopalatine_VS_P1_P4_Lumbar_Stellate | 0 | 1 | 1 | 0 | 0 | 1 | 0 | 1 |
| GNG12 | -0.085 | P1_P2_P3_P4_Lumbar_Stellate_Sphenopalatine_VS_Coeliac | 1 | 1 | 1 | 1 | 1 | 0 | 1 | 1 |
| SLC8A1 | -0.085 | P2_Coeliac_Sphenopalatine_VS_P1_P3_P4_Lumbar_Stellate | 0 | 1 | 0 | 0 | 0 | 1 | 0 | 1 |
| THOC1 | -0.085 | P2_P4_Lumbar_Coeliac_Stellate_VS_P1_P3_Sphenopalatine | 0 | 1 | 0 | 1 | 1 | 1 | 1 | 0 |
| BECN1 | -0.085 | P1_P2_P3_P4_Lumbar_Stellate_Sphenopalatine_VS_Coeliac | 1 | 1 | 1 | 1 | 1 | 0 | 1 | 1 |
| PGM2L1 | -0.086 | P1_P2_P3_P4_Lumbar_Stellate_Sphenopalatine_VS_Coeliac | 1 | 1 | 1 | 1 | 1 | 0 | 1 | 1 |

| | | | | | | | | | | |
|---|---|---|---|---|---|---|---|---|---|---|
| PRR13 | -0.086 | P1_P2_P3_P4_Lumbar_Stellate_Sphenopalatine_VS_Coeliac | 1 | 1 | 1 | 1 | 1 | 0 | 1 | 1 |
| NEK1 | -0.0861 | P1_P2_P3_P4_Lumbar_Stellate_VS_Coeliac_Sphenopalatine | 1 | 1 | 1 | 1 | 1 | 0 | 1 | 0 |
| RTN2 | -0.0861 | P1_P2_P3_P4_Lumbar_Stellate_Sphenopalatine_VS_Coeliac | 1 | 1 | 1 | 1 | 1 | 0 | 1 | 1 |
| HOOK3 | -0.0862 | P1_P2_P4_Lumbar_Coeliac_Stellate_Sphenopalatine_VS_P3 | 1 | 1 | 0 | 1 | 1 | 1 | 1 | 1 |
| BORCS5 | -0.0863 | P1_P3_P4_Lumbar_Stellate_VS_P2_Coeliac_Sphenopalatine | 1 | 0 | 1 | 1 | 1 | 0 | 1 | 0 |
| STX4A | -0.0863 | P1_P2_P3_P4_Lumbar_Stellate_Sphenopalatine_VS_Coeliac | 1 | 1 | 1 | 1 | 1 | 0 | 1 | 1 |
| MRPL37 | -0.0864 | P1_P2_P3_P4_Lumbar_Stellate_Sphenopalatine_VS_Coeliac | 1 | 1 | 1 | 1 | 1 | 0 | 1 | 1 |
| MTHFD2 | -0.0865 | P1_P3_P4_Lumbar_Stellate_Sphenopalatine_VS_P2_Coeliac | 1 | 0 | 1 | 1 | 1 | 0 | 1 | 1 |
| UBE2H | -0.0867 | P2_Coeliac_Sphenopalatine_VS_P1_P3_P4_Lumbar_Stellate | 0 | 1 | 0 | 0 | 0 | 1 | 0 | 1 |
| DR1 | -0.0868 | P1_P2_P4_Lumbar_Coeliac_Stellate_Sphenopalatine_VS_P3 | 1 | 1 | 0 | 1 | 1 | 1 | 1 | 1 |
| TRAPPC8 | -0.0868 | P1_P2_P4_Lumbar_Coeliac_Stellate_Sphenopalatine_VS_P3 | 1 | 1 | 0 | 1 | 1 | 1 | 1 | 1 |
| NUMBL | -0.0872 | P1_P2_P3_P4_Lumbar_Stellate_Sphenopalatine_VS_Coeliac | 1 | 1 | 1 | 1 | 1 | 0 | 1 | 1 |
| HNRNPLL | -0.0873 | P1_P2_P3_Lumbar_Coeliac_Stellate_VS_P4_Sphenopalatine | 1 | 1 | 1 | 0 | 1 | 1 | 1 | 0 |
| MMP11 | -0.0873 | P1_P2_P4_Lumbar_Coeliac_Stellate_Sphenopalatine_VS_P3 | 1 | 1 | 0 | 1 | 1 | 1 | 1 | 1 |
| ADNP | -0.0874 | P2_Coeliac_Sphenopalatine_VS_P1_P3_P4_Lumbar_Stellate | 0 | 1 | 0 | 0 | 0 | 1 | 0 | 1 |
| MCRS1 | -0.0874 | P3_P4_Lumbar_Coeliac_Stellate_Sphenopalatine_VS_P1_P2 | 0 | 0 | 1 | 1 | 1 | 1 | 1 | 1 |
| PID1 | -0.0874 | P2_P3_P4_Lumbar_Coeliac_Stellate_Sphenopalatine_VS_P1 | 0 | 1 | 1 | 1 | 1 | 1 | 1 | 1 |
| HMGN2 | -0.0878 | P1_P2_P3_P4_Stellate_Sphenopalatine_VS_Lumbar_Coeliac | 1 | 1 | 1 | 1 | 0 | 0 | 1 | 1 |
| HIST3H2BA | -0.0878 | P1_P2_P3_P4_Lumbar_Stellate_Sphenopalatine_VS_Coeliac | 1 | 1 | 1 | 1 | 1 | 0 | 1 | 1 |
| PTTG1 | -0.0879 | P1_P3_P4_Lumbar_Stellate_VS_P2_Coeliac_Sphenopalatine | 1 | 0 | 1 | 1 | 1 | 0 | 1 | 0 |
| ERC1 | -0.0879 | Coeliac_Sphenopalatine_VS_P1_P2_P3_P4_Lumbar_Stellate | 0 | 0 | 0 | 0 | 0 | 1 | 0 | 1 |
| CHGA | -0.0879 | P4_Lumbar_Coeliac_Stellate_Sphenopalatine_VS_P1_P2_P3 | 0 | 0 | 0 | 1 | 1 | 1 | 1 | 1 |
| GEMIN7 | -0.0881 | P1_P2_P3_P4_Lumbar_Stellate_Sphenopalatine_VS_Coeliac | 1 | 1 | 1 | 1 | 1 | 0 | 1 | 1 |
| TMEM65 | -0.0882 | P1_P2_P3_P4_Lumbar_Stellate_Sphenopalatine_VS_Coeliac | 1 | 1 | 1 | 1 | 1 | 0 | 1 | 1 |
| ARXES2 | -0.0883 | P3_P4_Lumbar_Stellate_Sphenopalatine_VS_P1_P2_Coeliac | 0 | 0 | 1 | 1 | 1 | 0 | 1 | 1 |
| MRPL48 | -0.0883 | P1_P2_P3_P4_Lumbar_Stellate_Sphenopalatine_VS_Coeliac | 1 | 1 | 1 | 1 | 1 | 0 | 1 | 1 |
| VPS36 | -0.0885 | P1_P3_Coeliac_Sphenopalatine_VS_P2_P4_Lumbar_Stellate | 1 | 0 | 1 | 0 | 0 | 1 | 0 | 1 |
| KXD1 | -0.0885 | P1_P2_P3_P4_Lumbar_Stellate_Sphenopalatine_VS_Coeliac | 1 | 1 | 1 | 1 | 1 | 0 | 1 | 1 |
| PLEKHJ1 | -0.0886 | P1_P2_P3_P4_Lumbar_Stellate_Sphenopalatine_VS_Coeliac | 1 | 1 | 1 | 1 | 1 | 0 | 1 | 1 |
| SYT17 | -0.0888 | P3_P4_Lumbar_Stellate_Sphenopalatine_VS_P1_P2_Coeliac | 0 | 0 | 1 | 1 | 1 | 0 | 1 | 1 |

| | | | | | | | | | | |
|---|---|---|---|---|---|---|---|---|---|---|
| MT-ATP8 | -0.0889 | P2_Coeliac_Sphenopalatine_VS_P1_P3_P4_Lumbar_Stellate | 0 | 1 | 0 | 0 | 0 | 1 | 0 | 1 |
| CNRIP1 | -0.0889 | P1_P2_P3_P4_Lumbar_Stellate_Sphenopalatine_VS_Coeliac | 1 | 1 | 1 | 1 | 1 | 0 | 1 | 1 |
| POLE4 | -0.0891 | P1_P2_P3_P4_Stellate_Sphenopalatine_VS_Lumbar_Coeliac | 1 | 1 | 1 | 1 | 0 | 0 | 1 | 1 |
| FAM189B | -0.0891 | P1_P2_P3_P4_Lumbar_Stellate_Sphenopalatine_VS_Coeliac | 1 | 1 | 1 | 1 | 1 | 0 | 1 | 1 |
| NALCN | -0.0891 | P2_P3_P4_Lumbar_Coeliac_Stellate_Sphenopalatine_VS_P1 | 0 | 1 | 1 | 1 | 1 | 1 | 1 | 1 |
| AFF4 | -0.0893 | P2_Coeliac_Sphenopalatine_VS_P1_P3_P4_Lumbar_Stellate | 0 | 1 | 0 | 0 | 0 | 1 | 0 | 1 |
| ITGB1BP1 | -0.0893 | P1_P3_P4_Lumbar_Stellate_Sphenopalatine_VS_P2_Coeliac | 1 | 0 | 1 | 1 | 1 | 0 | 1 | 1 |
| SMIM26 | -0.0894 | P1_P2_P3_P4_Lumbar_Stellate_Sphenopalatine_VS_Coeliac | 1 | 1 | 1 | 1 | 1 | 0 | 1 | 1 |
| APC2 | -0.0897 | P2_Coeliac_Sphenopalatine_VS_P1_P3_P4_Lumbar_Stellate | 0 | 1 | 0 | 0 | 0 | 1 | 0 | 1 |
| ATP2A2 | -0.0897 | P2_P3_Coeliac_Sphenopalatine_VS_P1_P4_Lumbar_Stellate | 0 | 1 | 1 | 0 | 0 | 1 | 0 | 1 |
| PPP2R5E | -0.0898 | P2_P4_Lumbar_Coeliac_Stellate_VS_P1_P3_Sphenopalatine | 0 | 1 | 0 | 1 | 1 | 1 | 1 | 0 |
| KPNA3 | -0.0899 | P2_P4_Lumbar_Coeliac_Stellate_VS_P1_P3_Sphenopalatine | 0 | 1 | 0 | 1 | 1 | 1 | 1 | 0 |
| HERC1 | -0.09 | P2_P3_Coeliac_Sphenopalatine_VS_P1_P4_Lumbar_Stellate | 0 | 1 | 1 | 0 | 0 | 1 | 0 | 1 |
| IDH3B | -0.09 | P1_P2_P3_P4_Lumbar_Stellate_Sphenopalatine_VS_Coeliac | 1 | 1 | 1 | 1 | 1 | 0 | 1 | 1 |
| MACROD2 | -0.0902 | P1_P2_P3_P4_Lumbar_Stellate_Sphenopalatine_VS_Coeliac | 1 | 1 | 1 | 1 | 1 | 0 | 1 | 1 |
| FBXO9 | -0.0903 | P1_P2_P3_P4_Lumbar_Stellate_Sphenopalatine_VS_Coeliac | 1 | 1 | 1 | 1 | 1 | 0 | 1 | 1 |
| CERS5 | -0.0906 | P1_P2_P3_P4_Lumbar_Stellate_Sphenopalatine_VS_Coeliac | 1 | 1 | 1 | 1 | 1 | 0 | 1 | 1 |
| SHISA4 | -0.0909 | P1_P2_P3_P4_Lumbar_Stellate_Sphenopalatine_VS_Coeliac | 1 | 1 | 1 | 1 | 1 | 0 | 1 | 1 |
| CDKN2D | -0.091 | P1_P2_P3_P4_Lumbar_Stellate_Sphenopalatine_VS_Coeliac | 1 | 1 | 1 | 1 | 1 | 0 | 1 | 1 |
| PARVA | -0.091 | P1_P2_P4_Lumbar_Coeliac_Stellate_Sphenopalatine_VS_P3 | 1 | 1 | 0 | 1 | 1 | 1 | 1 | 1 |
| ERH | -0.0911 | P1_P3_P4_Stellate_Sphenopalatine_VS_P2_Lumbar_Coeliac | 1 | 0 | 1 | 1 | 0 | 0 | 1 | 1 |
| RAB22A | -0.0911 | P1_P2_P3_P4_Lumbar_Stellate_Sphenopalatine_VS_Coeliac | 1 | 1 | 1 | 1 | 1 | 0 | 1 | 1 |
| MFN2 | -0.0911 | P1_P2_P3_P4_Lumbar_Stellate_Sphenopalatine_VS_Coeliac | 1 | 1 | 1 | 1 | 1 | 0 | 1 | 1 |
| RTL8B | -0.0915 | P3_P4_Stellate_Sphenopalatine_VS_P1_P2_Lumbar_Coeliac | 0 | 0 | 1 | 1 | 0 | 0 | 1 | 1 |
| TEX264 | -0.0915 | P1_P2_P3_P4_Lumbar_Stellate_Sphenopalatine_VS_Coeliac | 1 | 1 | 1 | 1 | 1 | 0 | 1 | 1 |
| UBR4 | -0.0916 | P2_P3_Coeliac_Sphenopalatine_VS_P1_P4_Lumbar_Stellate | 0 | 1 | 1 | 0 | 0 | 1 | 0 | 1 |
| NPTN | -0.0916 | P1_P2_P4_Lumbar_Coeliac_Stellate_Sphenopalatine_VS_P3 | 1 | 1 | 0 | 1 | 1 | 1 | 1 | 1 |
| KRT10 | -0.0918 | P1_P2_P3_P4_Lumbar_Stellate_Sphenopalatine_VS_Coeliac | 1 | 1 | 1 | 1 | 1 | 0 | 1 | 1 |
| MCTS2 | -0.0919 | P3_P4_Lumbar_Coeliac_Stellate_Sphenopalatine_VS_P1_P2 | 0 | 0 | 1 | 1 | 1 | 1 | 1 | 1 |
| GAMT | -0.092 | P4_Lumbar_Stellate_VS_P1_P2_P3_Coeliac_Sphenopalatine | 0 | 0 | 0 | 1 | 1 | 0 | 1 | 0 |

| | | | | | | | | | | |
|---|---|---|---|---|---|---|---|---|---|---|
| TNIK | -0.092 | P1_P3_P4_Lumbar_Stellate_VS_P2_Coeliac_Sphenopalatine | 1 | 0 | 1 | 1 | 1 | 0 | 1 | 0 |
| GATAD2B | -0.092 | P2_P3_Coeliac_Sphenopalatine_VS_P1_P4_Lumbar_Stellate | 0 | 1 | 1 | 0 | 0 | 1 | 0 | 1 |
| DDX6 | -0.0922 | P2_Coeliac_Sphenopalatine_VS_P1_P3_P4_Lumbar_Stellate | 0 | 1 | 0 | 0 | 0 | 1 | 0 | 1 |
| AP1AR | -0.0926 | P1_P2_P3_Lumbar_Coeliac_Stellate_Sphenopalatine_VS_P4 | 1 | 1 | 1 | 0 | 1 | 1 | 1 | 1 |
| UBA5 | -0.0927 | P1_P4_Lumbar_Coeliac_Stellate_Sphenopalatine_VS_P2_P3 | 1 | 0 | 0 | 1 | 1 | 1 | 1 | 1 |
| EMC4 | -0.0928 | P1_P3_P4_Lumbar_Stellate_Sphenopalatine_VS_P2_Coeliac | 1 | 0 | 1 | 1 | 1 | 0 | 1 | 1 |
| ENOX1 | -0.0929 | P2_P4_Lumbar_Coeliac_Stellate_VS_P1_P3_Sphenopalatine | 0 | 1 | 0 | 1 | 1 | 1 | 1 | 0 |
| CD63 | -0.0931 | P1_P3_Lumbar_Coeliac_Stellate_Sphenopalatine_VS_P2_P4 | 1 | 0 | 1 | 0 | 1 | 1 | 1 | 1 |
| HSD17B7 | -0.0931 | P1_P2_P4_Lumbar_Coeliac_Stellate_Sphenopalatine_VS_P3 | 1 | 1 | 0 | 1 | 1 | 1 | 1 | 1 |
| YPEL1 | -0.0933 | P3_Coeliac_Sphenopalatine_VS_P1_P2_P4_Lumbar_Stellate | 0 | 0 | 1 | 0 | 0 | 1 | 0 | 1 |
| ESYT1 | -0.0935 | P1_P2_P4_Lumbar_Coeliac_Stellate_Sphenopalatine_VS_P3 | 1 | 1 | 0 | 1 | 1 | 1 | 1 | 1 |
| RGS7 | -0.0936 | P3_Lumbar_Stellate_VS_P1_P2_P4_Coeliac_Sphenopalatine | 0 | 0 | 1 | 0 | 1 | 0 | 1 | 0 |
| RBM42 | -0.0938 | P3_P4_Lumbar_Coeliac_Stellate_Sphenopalatine_VS_P1_P2 | 0 | 0 | 1 | 1 | 1 | 1 | 1 | 1 |
| BLOC1S1 | -0.0939 | P1_P2_P3_P4_Coeliac_Stellate_Sphenopalatine_VS_Lumbar | 1 | 1 | 1 | 1 | 0 | 1 | 1 | 1 |
| DNAJB14 | -0.0941 | P2_Coeliac_Sphenopalatine_VS_P1_P3_P4_Lumbar_Stellate | 0 | 1 | 0 | 0 | 0 | 1 | 0 | 1 |
| NIPBL | -0.0941 | P2_P3_P4_Lumbar_Coeliac_Stellate_Sphenopalatine_VS_P1 | 0 | 1 | 1 | 1 | 1 | 1 | 1 | 1 |
| ARHGAP21 | -0.0943 | P1_P2_P4_Lumbar_Coeliac_Stellate_Sphenopalatine_VS_P3 | 1 | 1 | 0 | 1 | 1 | 1 | 1 | 1 |
| VTI1A | -0.0944 | P2_P3_Coeliac_Sphenopalatine_VS_P1_P4_Lumbar_Stellate | 0 | 1 | 1 | 0 | 0 | 1 | 0 | 1 |
| SNRPA | -0.0944 | P3_P4_Lumbar_Coeliac_Stellate_Sphenopalatine_VS_P1_P2 | 0 | 0 | 1 | 1 | 1 | 1 | 1 | 1 |
| TRIM37 | -0.0945 | P2_Coeliac_Sphenopalatine_VS_P1_P3_P4_Lumbar_Stellate | 0 | 1 | 0 | 0 | 0 | 1 | 0 | 1 |
| CRCP | -0.0949 | P3_Lumbar_Coeliac_Stellate_Sphenopalatine_VS_P1_P2_P4 | 0 | 0 | 1 | 0 | 1 | 1 | 1 | 1 |
| GMPS | -0.0952 | P1_P2_P4_Lumbar_Coeliac_Stellate_Sphenopalatine_VS_P3 | 1 | 1 | 0 | 1 | 1 | 1 | 1 | 1 |
| ASPHD2 | -0.0953 | P2_P3_Coeliac_Sphenopalatine_VS_P1_P4_Lumbar_Stellate | 0 | 1 | 1 | 0 | 0 | 1 | 0 | 1 |
| CAMK2G | -0.0954 | P2_P3_Coeliac_Sphenopalatine_VS_P1_P4_Lumbar_Stellate | 0 | 1 | 1 | 0 | 0 | 1 | 0 | 1 |
| FAR1 | -0.0956 | P1_P2_P4_Lumbar_Coeliac_Stellate_Sphenopalatine_VS_P3 | 1 | 1 | 0 | 1 | 1 | 1 | 1 | 1 |
| MRPS27 | -0.0956 | P1_P2_P4_Lumbar_Coeliac_Stellate_Sphenopalatine_VS_P3 | 1 | 1 | 0 | 1 | 1 | 1 | 1 | 1 |
| 2610318N02 | -0.0957 | P1_Lumbar_Coeliac_Stellate_VS_P2_P3_P4_Sphenopalatine | 1 | 0 | 0 | 0 | 1 | 1 | 1 | 0 |
| RALGPS1 | -0.0958 | P2_P3_Coeliac_Sphenopalatine_VS_P1_P4_Lumbar_Stellate | 0 | 1 | 1 | 0 | 0 | 1 | 0 | 1 |
| STOX2 | -0.0959 | P2_Coeliac_Sphenopalatine_VS_P1_P3_P4_Lumbar_Stellate | 0 | 1 | 0 | 0 | 0 | 1 | 0 | 1 |
| SHANK1 | -0.096 | P2_P3_Coeliac_Sphenopalatine_VS_P1_P4_Lumbar_Stellate | 0 | 1 | 1 | 0 | 0 | 1 | 0 | 1 |

| | | | | | | | | | | |
|---|---|---|---|---|---|---|---|---|---|---|
| SYT1 | -0.0961 | P1_P2_P3_P4_Lumbar_Stellate_VS_Coeliac_Sphenopalatine | 1 | 1 | 1 | 1 | 1 | 0 | 1 | 0 |
| ASXL2 | -0.0963 | P1_P2_P3_Lumbar_Coeliac_Stellate_VS_P4_Sphenopalatine | 1 | 1 | 1 | 0 | 1 | 1 | 1 | 0 |
| DPP6 | -0.0963 | P2_Coeliac_Sphenopalatine_VS_P1_P3_P4_Lumbar_Stellate | 0 | 1 | 0 | 0 | 0 | 1 | 0 | 1 |
| NPEPPS | -0.0963 | P1_P2_P4_Lumbar_Coeliac_Stellate_Sphenopalatine_VS_P3 | 1 | 1 | 0 | 1 | 1 | 1 | 1 | 1 |
| NMT2 | -0.0965 | P3_Coeliac_Sphenopalatine_VS_P1_P2_P4_Lumbar_Stellate | 0 | 0 | 1 | 0 | 0 | 1 | 0 | 1 |
| SMIM1 | -0.0966 | P3_P4_Stellate_VS_P1_P2_Lumbar_Coeliac_Sphenopalatine | 0 | 0 | 1 | 1 | 0 | 0 | 1 | 0 |
| RNH1 | -0.0966 | P1_P2_P3_P4_Lumbar_Stellate_VS_Coeliac_Sphenopalatine | 1 | 1 | 1 | 1 | 1 | 0 | 1 | 0 |
| OXR1 | -0.0966 | P3_P4_Lumbar_Coeliac_Stellate_Sphenopalatine_VS_P1_P2 | 0 | 0 | 1 | 1 | 1 | 1 | 1 | 1 |
| RALGPS2 | -0.0968 | P2_P4_Lumbar_Coeliac_Stellate_VS_P1_P3_Sphenopalatine | 0 | 1 | 0 | 1 | 1 | 1 | 1 | 0 |
| ZMAT5 | -0.0969 | P1_Lumbar_Coeliac_Stellate_VS_P2_P3_P4_Sphenopalatine | 1 | 0 | 0 | 0 | 1 | 1 | 1 | 0 |
| CCZ1 | -0.0972 | P1_P2_P3_Lumbar_Coeliac_Stellate_VS_P4_Sphenopalatine | 1 | 1 | 1 | 0 | 1 | 1 | 1 | 0 |
| BHLHB9 | -0.0974 | P1_P3_Lumbar_Coeliac_Stellate_Sphenopalatine_VS_P2_P4 | 1 | 0 | 1 | 0 | 1 | 1 | 1 | 1 |
| WDR55 | -0.0975 | P1_P2_P3_Lumbar_Coeliac_Stellate_Sphenopalatine_VS_P4 | 1 | 1 | 1 | 0 | 1 | 1 | 1 | 1 |
| NIFK | -0.0976 | P1_Lumbar_Coeliac_Stellate_VS_P2_P3_P4_Sphenopalatine | 1 | 0 | 0 | 0 | 1 | 1 | 1 | 0 |
| SOCS4 | -0.0976 | P1_P2_P3_Lumbar_Coeliac_Stellate_VS_P4_Sphenopalatine | 1 | 1 | 1 | 0 | 1 | 1 | 1 | 0 |
| COMMD10 | -0.0977 | P3_P4_Lumbar_Stellate_VS_P1_P2_Coeliac_Sphenopalatine | 0 | 0 | 1 | 1 | 1 | 0 | 1 | 0 |
| FAM216A | -0.0977 | P3_P4_Lumbar_Coeliac_Stellate_Sphenopalatine_VS_P1_P2 | 0 | 0 | 1 | 1 | 1 | 1 | 1 | 1 |
| METTL26 | -0.0977 | P3_P4_Lumbar_Coeliac_Stellate_Sphenopalatine_VS_P1_P2 | 0 | 0 | 1 | 1 | 1 | 1 | 1 | 1 |
| NRADD | -0.0978 | P2_P3_Lumbar_Coeliac_Stellate_VS_P1_P4_Sphenopalatine | 0 | 1 | 1 | 0 | 1 | 1 | 1 | 0 |
| CSNK1G2 | -0.098 | P2_P3_Coeliac_Sphenopalatine_VS_P1_P4_Lumbar_Stellate | 0 | 1 | 1 | 0 | 0 | 1 | 0 | 1 |
| LRP10 | -0.098 | P2_P3_Coeliac_Sphenopalatine_VS_P1_P4_Lumbar_Stellate | 0 | 1 | 1 | 0 | 0 | 1 | 0 | 1 |
| GM6710 | -0.0982 | P1_P2_P3_P4_Lumbar_Stellate_VS_Coeliac_Sphenopalatine | 1 | 1 | 1 | 1 | 1 | 0 | 1 | 0 |
| USP31 | -0.0982 | P2_P3_Coeliac_Sphenopalatine_VS_P1_P4_Lumbar_Stellate | 0 | 1 | 1 | 0 | 0 | 1 | 0 | 1 |
| PRKCSH | -0.0983 | P3_Coeliac_Sphenopalatine_VS_P1_P2_P4_Lumbar_Stellate | 0 | 0 | 1 | 0 | 0 | 1 | 0 | 1 |
| CASP3 | -0.0986 | P2_Lumbar_Coeliac_Stellate_VS_P1_P3_P4_Sphenopalatine | 0 | 1 | 0 | 0 | 1 | 1 | 1 | 0 |
| GOLGA7 | -0.0986 | P1_P2_P3_P4_Stellate_Sphenopalatine_VS_Lumbar_Coeliac | 1 | 1 | 1 | 1 | 0 | 0 | 1 | 1 |
| 4833439L19F | -0.0986 | P1_P2_P3_P4_Stellate_Sphenopalatine_VS_Lumbar_Coeliac | 1 | 1 | 1 | 1 | 0 | 0 | 1 | 1 |
| PMVK | -0.099 | P1_P3_P4_Lumbar_Stellate_Sphenopalatine_VS_P2_Coeliac | 1 | 0 | 1 | 1 | 1 | 0 | 1 | 1 |
| HEBP2 | -0.0993 | P1_P2_P4_Lumbar_Coeliac_Stellate_Sphenopalatine_VS_P3 | 1 | 1 | 0 | 1 | 1 | 1 | 1 | 1 |
| ZFP292 | -0.0994 | P2_P4_Lumbar_Coeliac_Stellate_VS_P1_P3_Sphenopalatine | 0 | 1 | 0 | 1 | 1 | 1 | 1 | 0 |

| | | | | | | | | | | |
|---|---|---|---|---|---|---|---|---|---|---|
| ARL6 | -0.0998 | P3_P4_Lumbar_Stellate_VS_P1_P2_Coeliac_Sphenopalatine | 0 | 0 | 1 | 1 | 1 | 0 | 1 | 0 |
| 1700037H04 | -0.0999 | P1_P2_P3_P4_Stellate_Sphenopalatine_VS_Lumbar_Coeliac | 1 | 1 | 1 | 1 | 0 | 0 | 1 | 1 |
| NUBP2 | -0.1 | P1_P3_P4_Lumbar_Stellate_Sphenopalatine_VS_P2_Coeliac | 1 | 0 | 1 | 1 | 1 | 0 | 1 | 1 |
| PFDN6 | -0.1 | P3_P4_Lumbar_Coeliac_Stellate_Sphenopalatine_VS_P1_P2 | 0 | 0 | 1 | 1 | 1 | 1 | 1 | 1 |
| PRCP | -0.101 | P1_P2_P3_Lumbar_Coeliac_Stellate_VS_P4_Sphenopalatine | 1 | 1 | 1 | 0 | 1 | 1 | 1 | 0 |
| AGTPBP1 | -0.101 | P1_Coeliac_Sphenopalatine_VS_P2_P3_P4_Lumbar_Stellate | 1 | 0 | 0 | 0 | 0 | 1 | 0 | 1 |
| NDUFAF4 | -0.101 | P3_Coeliac_Sphenopalatine_VS_P1_P2_P4_Lumbar_Stellate | 0 | 0 | 1 | 0 | 0 | 1 | 0 | 1 |
| BRD9 | -0.101 | P3_Coeliac_Sphenopalatine_VS_P1_P2_P4_Lumbar_Stellate | 0 | 0 | 1 | 0 | 0 | 1 | 0 | 1 |
| ACADM | -0.101 | P1_P4_Lumbar_Coeliac_Stellate_Sphenopalatine_VS_P2_P3 | 1 | 0 | 0 | 1 | 1 | 1 | 1 | 1 |
| TMEM159 | -0.102 | P4_Lumbar_Stellate_VS_P1_P2_P3_Coeliac_Sphenopalatine | 0 | 0 | 0 | 1 | 1 | 0 | 1 | 0 |
| THYN1 | -0.102 | P3_P4_Lumbar_Stellate_VS_P1_P2_Coeliac_Sphenopalatine | 0 | 0 | 1 | 1 | 1 | 0 | 1 | 0 |
| POLR3H | -0.102 | P3_P4_Lumbar_Stellate_VS_P1_P2_Coeliac_Sphenopalatine | 0 | 0 | 1 | 1 | 1 | 0 | 1 | 0 |
| DCLK2 | -0.102 | P3_Coeliac_Sphenopalatine_VS_P1_P2_P4_Lumbar_Stellate | 0 | 0 | 1 | 0 | 0 | 1 | 0 | 1 |
| SETD5 | -0.102 | P1_P3_Coeliac_Sphenopalatine_VS_P2_P4_Lumbar_Stellate | 1 | 0 | 1 | 0 | 0 | 1 | 0 | 1 |
| MRRF | -0.102 | P4_Lumbar_Coeliac_Stellate_Sphenopalatine_VS_P1_P2_P3 | 0 | 0 | 0 | 1 | 1 | 1 | 1 | 1 |
| KDM5B | -0.103 | P2_P4_Lumbar_Coeliac_Stellate_VS_P1_P3_Sphenopalatine | 0 | 1 | 0 | 1 | 1 | 1 | 1 | 0 |
| TBL1X | -0.103 | P2_P4_Lumbar_Coeliac_Stellate_VS_P1_P3_Sphenopalatine | 0 | 1 | 0 | 1 | 1 | 1 | 1 | 0 |
| MRPS17 | -0.103 | P1_P3_P4_Lumbar_Stellate_Sphenopalatine_VS_P2_Coeliac | 1 | 0 | 1 | 1 | 1 | 0 | 1 | 1 |
| COMMD4 | -0.103 | P1_P3_P4_Lumbar_Stellate_Sphenopalatine_VS_P2_Coeliac | 1 | 0 | 1 | 1 | 1 | 0 | 1 | 1 |
| PNCK | -0.104 | P3_P4_Lumbar_Stellate_VS_P1_P2_Coeliac_Sphenopalatine | 0 | 0 | 1 | 1 | 1 | 0 | 1 | 0 |
| ABCC5 | -0.104 | P2_P4_Lumbar_Coeliac_Stellate_VS_P1_P3_Sphenopalatine | 0 | 1 | 0 | 1 | 1 | 1 | 1 | 0 |
| SH3BGRL | -0.104 | P3_Coeliac_Sphenopalatine_VS_P1_P2_P4_Lumbar_Stellate | 0 | 0 | 1 | 0 | 0 | 1 | 0 | 1 |
| RABIF | -0.104 | P1_P2_P3_Lumbar_Stellate_Sphenopalatine_VS_P4_Coeliac | 1 | 1 | 1 | 0 | 1 | 0 | 1 | 1 |
| POLR3K | -0.104 | P1_P3_Lumbar_Coeliac_Stellate_Sphenopalatine_VS_P2_P4 | 1 | 0 | 1 | 0 | 1 | 1 | 1 | 1 |
| PRRC2C | -0.105 | P3_Coeliac_Sphenopalatine_VS_P1_P2_P4_Lumbar_Stellate | 0 | 0 | 1 | 0 | 0 | 1 | 0 | 1 |
| PPM1G | -0.105 | P3_Coeliac_Sphenopalatine_VS_P1_P2_P4_Lumbar_Stellate | 0 | 0 | 1 | 0 | 0 | 1 | 0 | 1 |
| TAF13 | -0.105 | P1_P2_P3_P4_Stellate_Sphenopalatine_VS_Lumbar_Coeliac | 1 | 1 | 1 | 1 | 0 | 0 | 1 | 1 |
| MRPS10 | -0.105 | P1_P3_P4_Lumbar_Stellate_Sphenopalatine_VS_P2_Coeliac | 1 | 0 | 1 | 1 | 1 | 0 | 1 | 1 |
| SNHG3 | -0.105 | P2_P3_P4_Lumbar_Stellate_Sphenopalatine_VS_P1_Coeliac | 0 | 1 | 1 | 1 | 1 | 0 | 1 | 1 |
| TTLL12 | -0.105 | P1_P2_P3_Lumbar_Coeliac_Stellate_Sphenopalatine_VS_P4 | 1 | 1 | 1 | 0 | 1 | 1 | 1 | 1 |

| | | | | | | | | | | |
|---|---|---|---|---|---|---|---|---|---|---|
| RNF138 | -0.105 | P1_P2_P3_Lumbar_Coeliac_Stellate_Sphenopalatine_VS_P4 | 1 | 1 | 1 | 0 | 1 | 1 | 1 | 1 |
| SCCPDH | -0.105 | P1_P2_P4_Lumbar_Coeliac_Stellate_Sphenopalatine_VS_P3 | 1 | 1 | 0 | 1 | 1 | 1 | 1 | 1 |
| ZFP846 | -0.105 | P1_P2_P4_Lumbar_Coeliac_Stellate_Sphenopalatine_VS_P3 | 1 | 1 | 0 | 1 | 1 | 1 | 1 | 1 |
| PDIA6 | -0.105 | P3_P4_Lumbar_Coeliac_Stellate_Sphenopalatine_VS_P1_P2 | 0 | 0 | 1 | 1 | 1 | 1 | 1 | 1 |
| MPZL1 | -0.106 | P1_P4_Lumbar_Stellate_VS_P2_P3_Coeliac_Sphenopalatine | 1 | 0 | 0 | 1 | 1 | 0 | 1 | 0 |
| VEZF1 | -0.106 | P2_P4_Lumbar_Coeliac_Stellate_VS_P1_P3_Sphenopalatine | 0 | 1 | 0 | 1 | 1 | 1 | 1 | 0 |
| ZDHHC17 | -0.106 | P1_Coeliac_Sphenopalatine_VS_P2_P3_P4_Lumbar_Stellate | 1 | 0 | 0 | 0 | 0 | 1 | 0 | 1 |
| RNF181 | -0.106 | P1_P2_P3_P4_Stellate_Sphenopalatine_VS_Lumbar_Coeliac | 1 | 1 | 1 | 1 | 0 | 0 | 1 | 1 |
| PIGYL | -0.106 | P1_P3_P4_Lumbar_Stellate_Sphenopalatine_VS_P2_Coeliac | 1 | 0 | 1 | 1 | 1 | 0 | 1 | 1 |
| ARHGAP26 | -0.106 | P1_P4_Lumbar_Coeliac_Stellate_Sphenopalatine_VS_P2_P3 | 1 | 0 | 0 | 1 | 1 | 1 | 1 | 1 |
| IMMP1L | -0.107 | P1_P3_P4_Lumbar_Stellate_VS_P2_Coeliac_Sphenopalatine | 1 | 0 | 1 | 1 | 1 | 0 | 1 | 0 |
| EFHD2 | -0.107 | P2_P3_Lumbar_Coeliac_Stellate_VS_P1_P4_Sphenopalatine | 0 | 1 | 1 | 0 | 1 | 1 | 1 | 0 |
| URM1 | -0.107 | P1_P2_P3_P4_Stellate_Sphenopalatine_VS_Lumbar_Coeliac | 1 | 1 | 1 | 1 | 0 | 0 | 1 | 1 |
| AMN1 | -0.107 | P3_P4_Lumbar_Stellate_Sphenopalatine_VS_P1_P2_Coeliac | 0 | 0 | 1 | 1 | 1 | 0 | 1 | 1 |
| AKT1S1 | -0.107 | P1_P3_P4_Lumbar_Stellate_Sphenopalatine_VS_P2_Coeliac | 1 | 0 | 1 | 1 | 1 | 0 | 1 | 1 |
| ATP13A2 | -0.107 | P1_P3_Lumbar_Coeliac_Stellate_Sphenopalatine_VS_P2_P4 | 1 | 0 | 1 | 0 | 1 | 1 | 1 | 1 |
| DECR1 | -0.107 | P1_P2_P4_Lumbar_Coeliac_Stellate_Sphenopalatine_VS_P3 | 1 | 1 | 0 | 1 | 1 | 1 | 1 | 1 |
| NSMCE3 | -0.107 | P3_P4_Lumbar_Coeliac_Stellate_Sphenopalatine_VS_P1_P2 | 0 | 0 | 1 | 1 | 1 | 1 | 1 | 1 |
| UBAP1 | -0.108 | P1_P2_P3_P4_Lumbar_Stellate_VS_Coeliac_Sphenopalatine | 1 | 1 | 1 | 1 | 1 | 0 | 1 | 0 |
| DANCR | -0.108 | P2_Lumbar_Coeliac_Stellate_VS_P1_P3_P4_Sphenopalatine | 0 | 1 | 0 | 0 | 1 | 1 | 1 | 0 |
| 1110032A03I | -0.108 | P1_P3_P4_Stellate_Sphenopalatine_VS_P2_Lumbar_Coeliac | 1 | 0 | 1 | 1 | 0 | 0 | 1 | 1 |
| SYN1 | -0.108 | P1_P2_P3_P4_Stellate_Sphenopalatine_VS_Lumbar_Coeliac | 1 | 1 | 1 | 1 | 0 | 0 | 1 | 1 |
| EPHA5 | -0.108 | P1_P2_P4_Lumbar_Stellate_Sphenopalatine_VS_P3_Coeliac | 1 | 1 | 0 | 1 | 1 | 0 | 1 | 1 |
| SSBP4 | -0.108 | P1_P3_P4_Lumbar_Stellate_Sphenopalatine_VS_P2_Coeliac | 1 | 0 | 1 | 1 | 1 | 0 | 1 | 1 |
| BCL10 | -0.108 | P1_Lumbar_Coeliac_Stellate_Sphenopalatine_VS_P2_P3_P4 | 1 | 0 | 0 | 0 | 1 | 1 | 1 | 1 |
| MINDY3 | -0.108 | P1_P2_P3_Lumbar_Coeliac_Stellate_Sphenopalatine_VS_P4 | 1 | 1 | 1 | 0 | 1 | 1 | 1 | 1 |
| HSPBP1 | -0.108 | P3_P4_Lumbar_Coeliac_Stellate_Sphenopalatine_VS_P1_P2 | 0 | 0 | 1 | 1 | 1 | 1 | 1 | 1 |
| TBRG1 | -0.108 | P3_P4_Lumbar_Coeliac_Stellate_Sphenopalatine_VS_P1_P2 | 0 | 0 | 1 | 1 | 1 | 1 | 1 | 1 |
| LRP11 | -0.109 | P1_P2_P3_P4_Stellate_Sphenopalatine_VS_Lumbar_Coeliac | 1 | 1 | 1 | 1 | 0 | 0 | 1 | 1 |
| PMP22 | -0.109 | P1_P2_P3_P4_Stellate_Sphenopalatine_VS_Lumbar_Coeliac | 1 | 1 | 1 | 1 | 0 | 0 | 1 | 1 |

| | | | | | | | | | | |
|---|---|---|---|---|---|---|---|---|---|---|
| PPP1R1A | -0.109 | P1_P2_P3_P4_Stellate_Sphenopalatine_VS_Lumbar_Coeliac | 1 | 1 | 1 | 1 | 0 | 0 | 1 | 1 |
| UQCC3 | -0.109 | P1_P2_P3_P4_Stellate_Sphenopalatine_VS_Lumbar_Coeliac | 1 | 1 | 1 | 1 | 0 | 0 | 1 | 1 |
| CHRAC1 | -0.109 | P3_P4_Lumbar_Coeliac_Stellate_Sphenopalatine_VS_P1_P2 | 0 | 0 | 1 | 1 | 1 | 1 | 1 | 1 |
| PARL | -0.11 | P3_P4_Lumbar_Stellate_VS_P1_P2_Coeliac_Sphenopalatine | 0 | 0 | 1 | 1 | 1 | 0 | 1 | 0 |
| ADIPOR1 | -0.11 | P2_P3_P4_Stellate_Sphenopalatine_VS_P1_Lumbar_Coeliac | 0 | 1 | 1 | 1 | 0 | 0 | 1 | 1 |
| AKTIP | -0.11 | P1_P2_P3_P4_Stellate_Sphenopalatine_VS_Lumbar_Coeliac | 1 | 1 | 1 | 1 | 0 | 0 | 1 | 1 |
| DMAC1 | -0.11 | P1_P3_P4_Lumbar_Stellate_Sphenopalatine_VS_P2_Coeliac | 1 | 0 | 1 | 1 | 1 | 0 | 1 | 1 |
| DIRAS1 | -0.11 | P1_P2_P3_P4_Coeliac_Stellate_Sphenopalatine_VS_Lumbar | 1 | 1 | 1 | 1 | 0 | 1 | 1 | 1 |
| GNB5 | -0.11 | P1_P2_P3_Lumbar_Coeliac_Stellate_Sphenopalatine_VS_P4 | 1 | 1 | 1 | 0 | 1 | 1 | 1 | 1 |
| RTF1 | -0.11 | P3_P4_Lumbar_Coeliac_Stellate_Sphenopalatine_VS_P1_P2 | 0 | 0 | 1 | 1 | 1 | 1 | 1 | 1 |
| AIG1 | -0.11 | P3_P4_Lumbar_Coeliac_Stellate_Sphenopalatine_VS_P1_P2 | 0 | 0 | 1 | 1 | 1 | 1 | 1 | 1 |
| ZRSR2 | -0.11 | P3_P4_Lumbar_Coeliac_Stellate_Sphenopalatine_VS_P1_P2 | 0 | 0 | 1 | 1 | 1 | 1 | 1 | 1 |
| FAM57B | -0.111 | P3_P4_Lumbar_Stellate_Sphenopalatine_VS_P1_P2_Coeliac | 0 | 0 | 1 | 1 | 1 | 0 | 1 | 1 |
| NAA38 | -0.111 | P1_P3_P4_Lumbar_Stellate_Sphenopalatine_VS_P2_Coeliac | 1 | 0 | 1 | 1 | 1 | 0 | 1 | 1 |
| LSM2 | -0.111 | P1_P2_P3_Lumbar_Coeliac_Stellate_Sphenopalatine_VS_P4 | 1 | 1 | 1 | 0 | 1 | 1 | 1 | 1 |
| PAK1IP1 | -0.111 | P3_P4_Lumbar_Coeliac_Stellate_Sphenopalatine_VS_P1_P2 | 0 | 0 | 1 | 1 | 1 | 1 | 1 | 1 |
| MRPS6 | -0.111 | P2_P3_P4_Lumbar_Coeliac_Stellate_Sphenopalatine_VS_P1 | 0 | 1 | 1 | 1 | 1 | 1 | 1 | 1 |
| GRPEL1 | -0.112 | P3_Lumbar_Stellate_VS_P1_P2_P4_Coeliac_Sphenopalatine | 0 | 0 | 1 | 0 | 1 | 0 | 1 | 0 |
| TM2D1 | -0.112 | P3_P4_Stellate_Sphenopalatine_VS_P1_P2_Lumbar_Coeliac | 0 | 0 | 1 | 1 | 0 | 0 | 1 | 1 |
| FAM174A | -0.112 | P1_P2_P3_P4_Stellate_Sphenopalatine_VS_Lumbar_Coeliac | 1 | 1 | 1 | 1 | 0 | 0 | 1 | 1 |
| WDR1 | -0.112 | P1_P2_P3_P4_Stellate_Sphenopalatine_VS_Lumbar_Coeliac | 1 | 1 | 1 | 1 | 0 | 0 | 1 | 1 |
| BUD31 | -0.112 | P1_P3_P4_Lumbar_Stellate_Sphenopalatine_VS_P2_Coeliac | 1 | 0 | 1 | 1 | 1 | 0 | 1 | 1 |
| NSDHL | -0.112 | P1_P3_P4_Lumbar_Stellate_Sphenopalatine_VS_P2_Coeliac | 1 | 0 | 1 | 1 | 1 | 0 | 1 | 1 |
| RHNO1 | -0.112 | P1_P2_P3_Lumbar_Coeliac_Stellate_Sphenopalatine_VS_P4 | 1 | 1 | 1 | 0 | 1 | 1 | 1 | 1 |
| PUF60 | -0.112 | P1_P2_P4_Lumbar_Coeliac_Stellate_Sphenopalatine_VS_P3 | 1 | 1 | 0 | 1 | 1 | 1 | 1 | 1 |
| TBPL1 | -0.112 | P3_P4_Lumbar_Coeliac_Stellate_Sphenopalatine_VS_P1_P2 | 0 | 0 | 1 | 1 | 1 | 1 | 1 | 1 |
| QDPR | -0.112 | P2_P3_P4_Lumbar_Coeliac_Stellate_Sphenopalatine_VS_P1 | 0 | 1 | 1 | 1 | 1 | 1 | 1 | 1 |
| CHCHD6 | -0.113 | P1_P3_P4_Lumbar_Stellate_VS_P2_Coeliac_Sphenopalatine | 1 | 0 | 1 | 1 | 1 | 0 | 1 | 0 |
| DGAT2 | -0.113 | P2_P3_P4_Lumbar_Stellate_VS_P1_Coeliac_Sphenopalatine | 0 | 1 | 1 | 1 | 1 | 0 | 1 | 0 |
| NEFM | -0.113 | P1_P2_P3_P4_Lumbar_Stellate_VS_Coeliac_Sphenopalatine | 1 | 1 | 1 | 1 | 1 | 0 | 1 | 0 |

| | | | | | | | | | | |
|---|---|---|---|---|---|---|---|---|---|---|
| TDG | -0.113 | P2_P3_Lumbar_Coeliac_Stellate_VS_P1_P4_Sphenopalatine | 0 | 1 | 1 | 0 | 1 | 1 | 1 | 0 |
| IMPACT | -0.113 | P1_P2_P4_Stellate_Sphenopalatine_VS_P3_Lumbar_Coeliac | 1 | 1 | 0 | 1 | 0 | 0 | 1 | 1 |
| TOR1AIP2 | -0.113 | P1_P2_P3_P4_Stellate_Sphenopalatine_VS_Lumbar_Coeliac | 1 | 1 | 1 | 1 | 0 | 0 | 1 | 1 |
| SH3GL3 | -0.113 | P1_P2_P3_P4_Stellate_Sphenopalatine_VS_Lumbar_Coeliac | 1 | 1 | 1 | 1 | 0 | 0 | 1 | 1 |
| USE1 | -0.113 | P1_P2_P3_P4_Stellate_Sphenopalatine_VS_Lumbar_Coeliac | 1 | 1 | 1 | 1 | 0 | 0 | 1 | 1 |
| DNAJC30 | -0.113 | P1_P3_P4_Lumbar_Stellate_Sphenopalatine_VS_P2_Coeliac | 1 | 0 | 1 | 1 | 1 | 0 | 1 | 1 |
| SLIRP | -0.113 | P2_P3_P4_Lumbar_Stellate_Sphenopalatine_VS_P1_Coeliac | 0 | 1 | 1 | 1 | 1 | 0 | 1 | 1 |
| RAD21 | -0.113 | P1_P2_P4_Lumbar_Coeliac_Stellate_Sphenopalatine_VS_P3 | 1 | 1 | 0 | 1 | 1 | 1 | 1 | 1 |
| PPID | -0.113 | P3_P4_Lumbar_Coeliac_Stellate_Sphenopalatine_VS_P1_P2 | 0 | 0 | 1 | 1 | 1 | 1 | 1 | 1 |
| LSM7 | -0.114 | P3_P4_Stellate_Sphenopalatine_VS_P1_P2_Lumbar_Coeliac | 0 | 0 | 1 | 1 | 0 | 0 | 1 | 1 |
| MINAR2 | -0.114 | P3_P4_Stellate_Sphenopalatine_VS_P1_P2_Lumbar_Coeliac | 0 | 0 | 1 | 1 | 0 | 0 | 1 | 1 |
| TM2D2 | -0.114 | P1_P3_P4_Lumbar_Stellate_Sphenopalatine_VS_P2_Coeliac | 1 | 0 | 1 | 1 | 1 | 0 | 1 | 1 |
| ACAT2 | -0.114 | P1_P3_P4_Lumbar_Stellate_Sphenopalatine_VS_P2_Coeliac | 1 | 0 | 1 | 1 | 1 | 0 | 1 | 1 |
| GNAZ | -0.114 | P2_P3_P4_Lumbar_Stellate_Sphenopalatine_VS_P1_Coeliac | 0 | 1 | 1 | 1 | 1 | 0 | 1 | 1 |
| ANP32A | -0.114 | P1_P3_Lumbar_Coeliac_Stellate_Sphenopalatine_VS_P2_P4 | 1 | 0 | 1 | 0 | 1 | 1 | 1 | 1 |
| NAP1L4 | -0.114 | P1_P2_P4_Lumbar_Coeliac_Stellate_Sphenopalatine_VS_P3 | 1 | 1 | 0 | 1 | 1 | 1 | 1 | 1 |
| RSU1 | -0.114 | P3_P4_Lumbar_Coeliac_Stellate_Sphenopalatine_VS_P1_P2 | 0 | 0 | 1 | 1 | 1 | 1 | 1 | 1 |
| MRPL21 | -0.114 | P3_P4_Lumbar_Coeliac_Stellate_Sphenopalatine_VS_P1_P2 | 0 | 0 | 1 | 1 | 1 | 1 | 1 | 1 |
| PBDC1 | -0.114 | P3_P4_Lumbar_Coeliac_Stellate_Sphenopalatine_VS_P1_P2 | 0 | 0 | 1 | 1 | 1 | 1 | 1 | 1 |
| LRPAP1 | -0.114 | P2_P3_P4_Lumbar_Coeliac_Stellate_Sphenopalatine_VS_P1 | 0 | 1 | 1 | 1 | 1 | 1 | 1 | 1 |
| SEC11C | -0.115 | P1_P2_P3_P4_Lumbar_Stellate_VS_Coeliac_Sphenopalatine | 1 | 1 | 1 | 1 | 1 | 0 | 1 | 0 |
| PPP1R16A | -0.115 | P1_P2_P3_P4_Stellate_Sphenopalatine_VS_Lumbar_Coeliac | 1 | 1 | 1 | 1 | 0 | 0 | 1 | 1 |
| AP3S1 | -0.115 | P1_P2_P3_P4_Stellate_Sphenopalatine_VS_Lumbar_Coeliac | 1 | 1 | 1 | 1 | 0 | 0 | 1 | 1 |
| MRPS14 | -0.115 | P1_P3_P4_Lumbar_Stellate_Sphenopalatine_VS_P2_Coeliac | 1 | 0 | 1 | 1 | 1 | 0 | 1 | 1 |
| COQ5 | -0.115 | P1_P2_P3_Lumbar_Coeliac_Stellate_Sphenopalatine_VS_P4 | 1 | 1 | 1 | 0 | 1 | 1 | 1 | 1 |
| SYN2 | -0.115 | P1_P2_P3_Lumbar_Coeliac_Stellate_Sphenopalatine_VS_P4 | 1 | 1 | 1 | 0 | 1 | 1 | 1 | 1 |
| ZFP810 | -0.115 | P1_P2_P3_Lumbar_Coeliac_Stellate_Sphenopalatine_VS_P4 | 1 | 1 | 1 | 0 | 1 | 1 | 1 | 1 |
| CCDC124 | -0.116 | P1_P2_P3_P4_Stellate_Sphenopalatine_VS_Lumbar_Coeliac | 1 | 1 | 1 | 1 | 0 | 0 | 1 | 1 |
| CHRNB4 | -0.116 | Lumbar_Stellate_Sphenopalatine_VS_P1_P2_P3_P4_Coeliac | 0 | 0 | 0 | 0 | 1 | 0 | 1 | 1 |
| SNN | -0.116 | P2_P3_Lumbar_Stellate_Sphenopalatine_VS_P1_P4_Coeliac | 0 | 1 | 1 | 0 | 1 | 0 | 1 | 1 |

| | | | | | | | | | | |
|---|---|---|---|---|---|---|---|---|---|---|
| MGST3 | -0.116 | P1_P3_P4_Lumbar_Stellate_Sphenopalatine_VS_P2_Coeliac | 1 | 0 | 1 | 1 | 1 | 0 | 1 | 1 |
| LSM3 | -0.116 | P1_P3_P4_Lumbar_Stellate_Sphenopalatine_VS_P2_Coeliac | 1 | 0 | 1 | 1 | 1 | 0 | 1 | 1 |
| 1500011B03 | -0.116 | P2_P3_P4_Lumbar_Stellate_Sphenopalatine_VS_P1_Coeliac | 0 | 1 | 1 | 1 | 1 | 0 | 1 | 1 |
| MAX | -0.116 | P2_P3_P4_Lumbar_Stellate_Sphenopalatine_VS_P1_Coeliac | 0 | 1 | 1 | 1 | 1 | 0 | 1 | 1 |
| AGPS | -0.116 | P1_P2_P4_Lumbar_Coeliac_Stellate_Sphenopalatine_VS_P3 | 1 | 1 | 0 | 1 | 1 | 1 | 1 | 1 |
| 1810026B05 | -0.116 | P1_P2_P4_Lumbar_Coeliac_Stellate_Sphenopalatine_VS_P3 | 1 | 1 | 0 | 1 | 1 | 1 | 1 | 1 |
| SEC11A | -0.117 | P3_P4_Lumbar_Stellate_VS_P1_P2_Coeliac_Sphenopalatine | 0 | 0 | 1 | 1 | 1 | 0 | 1 | 0 |
| UBE2J2 | -0.117 | P1_P2_P3_P4_Stellate_Sphenopalatine_VS_Lumbar_Coeliac | 1 | 1 | 1 | 1 | 0 | 0 | 1 | 1 |
| STARD3NL | -0.117 | P1_P2_P3_P4_Stellate_Sphenopalatine_VS_Lumbar_Coeliac | 1 | 1 | 1 | 1 | 0 | 0 | 1 | 1 |
| TXN2 | -0.117 | P1_P2_P3_P4_Stellate_Sphenopalatine_VS_Lumbar_Coeliac | 1 | 1 | 1 | 1 | 0 | 0 | 1 | 1 |
| LSM5 | -0.117 | P1_P3_P4_Lumbar_Stellate_Sphenopalatine_VS_P2_Coeliac | 1 | 0 | 1 | 1 | 1 | 0 | 1 | 1 |
| EMG1 | -0.117 | P1_P3_P4_Lumbar_Stellate_Sphenopalatine_VS_P2_Coeliac | 1 | 0 | 1 | 1 | 1 | 0 | 1 | 1 |
| ANGPTL6 | -0.117 | P2_P3_P4_Lumbar_Stellate_Sphenopalatine_VS_P1_Coeliac | 0 | 1 | 1 | 1 | 1 | 0 | 1 | 1 |
| ZFAS1 | -0.117 | P1_P2_P3_Lumbar_Coeliac_Stellate_Sphenopalatine_VS_P4 | 1 | 1 | 1 | 0 | 1 | 1 | 1 | 1 |
| SLC3A2 | -0.117 | P1_P2_P4_Lumbar_Coeliac_Stellate_Sphenopalatine_VS_P3 | 1 | 1 | 0 | 1 | 1 | 1 | 1 | 1 |
| OSTF1 | -0.117 | P1_P2_P4_Lumbar_Coeliac_Stellate_Sphenopalatine_VS_P3 | 1 | 1 | 0 | 1 | 1 | 1 | 1 | 1 |
| WDR83 | -0.118 | P1_P3_Lumbar_Coeliac_Stellate_Sphenopalatine_VS_P2_P4 | 1 | 0 | 1 | 0 | 1 | 1 | 1 | 1 |
| SAC3D1 | -0.118 | P1_P2_P3_Lumbar_Coeliac_Stellate_Sphenopalatine_VS_P4 | 1 | 1 | 1 | 0 | 1 | 1 | 1 | 1 |
| COMMD7 | -0.118 | P1_P4_Lumbar_Coeliac_Stellate_Sphenopalatine_VS_P2_P3 | 1 | 0 | 0 | 1 | 1 | 1 | 1 | 1 |
| DHPS | -0.118 | P1_P4_Lumbar_Coeliac_Stellate_Sphenopalatine_VS_P2_P3 | 1 | 0 | 0 | 1 | 1 | 1 | 1 | 1 |
| SC5D | -0.118 | P1_P2_P4_Lumbar_Coeliac_Stellate_Sphenopalatine_VS_P3 | 1 | 1 | 0 | 1 | 1 | 1 | 1 | 1 |
| KLF6 | -0.118 | P1_P2_P4_Lumbar_Coeliac_Stellate_Sphenopalatine_VS_P3 | 1 | 1 | 0 | 1 | 1 | 1 | 1 | 1 |
| OSTC | -0.118 | P3_P4_Lumbar_Coeliac_Stellate_Sphenopalatine_VS_P1_P2 | 0 | 0 | 1 | 1 | 1 | 1 | 1 | 1 |
| FAM136A | -0.118 | P3_P4_Lumbar_Coeliac_Stellate_Sphenopalatine_VS_P1_P2 | 0 | 0 | 1 | 1 | 1 | 1 | 1 | 1 |
| TMCC1 | -0.118 | P2_P3_P4_Lumbar_Coeliac_Stellate_Sphenopalatine_VS_P1 | 0 | 1 | 1 | 1 | 1 | 1 | 1 | 1 |
| ATP6V0E | -0.119 | P1_P3_P4_Lumbar_Stellate_VS_P2_Coeliac_Sphenopalatine | 1 | 0 | 1 | 1 | 1 | 0 | 1 | 0 |
| MEMO1 | -0.119 | P1_P3_P4_Lumbar_Stellate_VS_P2_Coeliac_Sphenopalatine | 1 | 0 | 1 | 1 | 1 | 0 | 1 | 0 |
| ABAT | -0.119 | P1_P2_P3_P4_Lumbar_Stellate_VS_Coeliac_Sphenopalatine | 1 | 1 | 1 | 1 | 1 | 0 | 1 | 0 |
| FAM57A | -0.119 | P2_P3_Lumbar_Coeliac_Stellate_VS_P1_P4_Sphenopalatine | 0 | 1 | 1 | 0 | 1 | 1 | 1 | 0 |
| CHCHD4 | -0.119 | P1_P3_P4_Stellate_Sphenopalatine_VS_P2_Lumbar_Coeliac | 1 | 0 | 1 | 1 | 0 | 0 | 1 | 1 |

| | | | | | | | | | | |
|---|---|---|---|---|---|---|---|---|---|---|
| ZFP771 | -0.119 | P1_P3_P4_Lumbar_Stellate_Sphenopalatine_VS_P2_Coeliac | 1 | 0 | 1 | 1 | 1 | 0 | 1 | 1 |
| PRPS1 | -0.119 | P1_P3_P4_Lumbar_Stellate_Sphenopalatine_VS_P2_Coeliac | 1 | 0 | 1 | 1 | 1 | 0 | 1 | 1 |
| TMEM9 | -0.119 | P2_P3_P4_Lumbar_Stellate_Sphenopalatine_VS_P1_Coeliac | 0 | 1 | 1 | 1 | 1 | 0 | 1 | 1 |
| TRIM27 | -0.119 | P1_P2_P3_Lumbar_Coeliac_Stellate_Sphenopalatine_VS_P4 | 1 | 1 | 1 | 0 | 1 | 1 | 1 | 1 |
| CLN8 | -0.119 | P2_P3_P4_Lumbar_Coeliac_Stellate_Sphenopalatine_VS_P1 | 0 | 1 | 1 | 1 | 1 | 1 | 1 | 1 |
| TMEM205 | -0.12 | P4_Lumbar_Stellate_VS_P1_P2_P3_Coeliac_Sphenopalatine | 0 | 0 | 0 | 1 | 1 | 0 | 1 | 0 |
| CETN2 | -0.12 | P1_P3_P4_Lumbar_Stellate_VS_P2_Coeliac_Sphenopalatine | 1 | 0 | 1 | 1 | 1 | 0 | 1 | 0 |
| SUMO3 | -0.12 | P1_P2_P3_P4_Stellate_Sphenopalatine_VS_Lumbar_Coeliac | 1 | 1 | 1 | 1 | 0 | 0 | 1 | 1 |
| NDUFAF3 | -0.12 | P1_P3_P4_Lumbar_Stellate_Sphenopalatine_VS_P2_Coeliac | 1 | 0 | 1 | 1 | 1 | 0 | 1 | 1 |
| NAGK | -0.12 | P2_P3_P4_Lumbar_Stellate_Sphenopalatine_VS_P1_Coeliac | 0 | 1 | 1 | 1 | 1 | 0 | 1 | 1 |
| EI24 | -0.12 | P2_P3_P4_Lumbar_Stellate_Sphenopalatine_VS_P1_Coeliac | 0 | 1 | 1 | 1 | 1 | 0 | 1 | 1 |
| SPOUT1 | -0.12 | P1_P2_P3_Lumbar_Coeliac_Stellate_Sphenopalatine_VS_P4 | 1 | 1 | 1 | 0 | 1 | 1 | 1 | 1 |
| PPP1R14B | -0.12 | P1_P2_P3_Lumbar_Coeliac_Stellate_Sphenopalatine_VS_P4 | 1 | 1 | 1 | 0 | 1 | 1 | 1 | 1 |
| GAR1 | -0.12 | P1_P4_Lumbar_Coeliac_Stellate_Sphenopalatine_VS_P2_P3 | 1 | 0 | 0 | 1 | 1 | 1 | 1 | 1 |
| ADD2 | -0.12 | P2_P4_Lumbar_Coeliac_Stellate_Sphenopalatine_VS_P1_P3 | 0 | 1 | 0 | 1 | 1 | 1 | 1 | 1 |
| ZFAND6 | -0.12 | P3_P4_Lumbar_Coeliac_Stellate_Sphenopalatine_VS_P1_P2 | 0 | 0 | 1 | 1 | 1 | 1 | 1 | 1 |
| sept-06 | -0.121 | P1_P2_P4_Lumbar_Stellate_VS_P3_Coeliac_Sphenopalatine | 1 | 1 | 0 | 1 | 1 | 0 | 1 | 0 |
| RPP30 | -0.121 | P1_P3_P4_Lumbar_Stellate_VS_P2_Coeliac_Sphenopalatine | 1 | 0 | 1 | 1 | 1 | 0 | 1 | 0 |
| YPEL3 | -0.121 | P2_P3_P4_Stellate_Sphenopalatine_VS_P1_Lumbar_Coeliac | 0 | 1 | 1 | 1 | 0 | 0 | 1 | 1 |
| PSENEN | -0.121 | P1_P2_P3_P4_Stellate_Sphenopalatine_VS_Lumbar_Coeliac | 1 | 1 | 1 | 1 | 0 | 0 | 1 | 1 |
| SIRT3 | -0.121 | P1_P3_P4_Lumbar_Stellate_Sphenopalatine_VS_P2_Coeliac | 1 | 0 | 1 | 1 | 1 | 0 | 1 | 1 |
| NOL12 | -0.121 | P1_P3_P4_Lumbar_Stellate_Sphenopalatine_VS_P2_Coeliac | 1 | 0 | 1 | 1 | 1 | 0 | 1 | 1 |
| POLR2D | -0.121 | P1_P3_P4_Lumbar_Stellate_Sphenopalatine_VS_P2_Coeliac | 1 | 0 | 1 | 1 | 1 | 0 | 1 | 1 |
| FBL | -0.121 | Lumbar_Coeliac_Stellate_Sphenopalatine_VS_P1_P2_P3_P4 | 0 | 0 | 0 | 0 | 1 | 1 | 1 | 1 |
| TSFM | -0.121 | P1_P4_Lumbar_Coeliac_Stellate_Sphenopalatine_VS_P2_P3 | 1 | 0 | 0 | 1 | 1 | 1 | 1 | 1 |
| MFSD14A | -0.121 | P3_P4_Lumbar_Coeliac_Stellate_Sphenopalatine_VS_P1_P2 | 0 | 0 | 1 | 1 | 1 | 1 | 1 | 1 |
| BTF3L4 | -0.121 | P3_P4_Lumbar_Coeliac_Stellate_Sphenopalatine_VS_P1_P2 | 0 | 0 | 1 | 1 | 1 | 1 | 1 | 1 |
| PA2G4 | -0.121 | P3_P4_Lumbar_Coeliac_Stellate_Sphenopalatine_VS_P1_P2 | 0 | 0 | 1 | 1 | 1 | 1 | 1 | 1 |
| IDI1 | -0.121 | P3_P4_Lumbar_Coeliac_Stellate_Sphenopalatine_VS_P1_P2 | 0 | 0 | 1 | 1 | 1 | 1 | 1 | 1 |
| SCG3 | -0.121 | P2_P3_P4_Lumbar_Coeliac_Stellate_Sphenopalatine_VS_P1 | 0 | 1 | 1 | 1 | 1 | 1 | 1 | 1 |

| | | | | | | | | | | |
|---|---|---|---|---|---|---|---|---|---|---|
| ERI3 | -0.122 | P3_P4_Stellate_VS_P1_P2_Lumbar_Coeliac_Sphenopalatine | 0 | 0 | 1 | 1 | 0 | 0 | 1 | 0 |
| CCDC92 | -0.122 | P1_P2_P3_P4_Stellate_VS_Lumbar_Coeliac_Sphenopalatine | 1 | 1 | 1 | 1 | 0 | 0 | 1 | 0 |
| B3GAT3 | -0.122 | P1_P2_P4_Lumbar_Stellate_Sphenopalatine_VS_P3_Coeliac | 1 | 1 | 0 | 1 | 1 | 0 | 1 | 1 |
| COPZ1 | -0.122 | P3_P4_Lumbar_Stellate_Sphenopalatine_VS_P1_P2_Coeliac | 0 | 0 | 1 | 1 | 1 | 0 | 1 | 1 |
| CLPP | -0.122 | P3_P4_Lumbar_Stellate_Sphenopalatine_VS_P1_P2_Coeliac | 0 | 0 | 1 | 1 | 1 | 0 | 1 | 1 |
| RAB4B | -0.122 | P1_P3_P4_Lumbar_Stellate_Sphenopalatine_VS_P2_Coeliac | 1 | 0 | 1 | 1 | 1 | 0 | 1 | 1 |
| CALY | -0.122 | P1_P3_P4_Lumbar_Stellate_Sphenopalatine_VS_P2_Coeliac | 1 | 0 | 1 | 1 | 1 | 0 | 1 | 1 |
| MRPL55 | -0.122 | P1_P3_P4_Lumbar_Stellate_Sphenopalatine_VS_P2_Coeliac | 1 | 0 | 1 | 1 | 1 | 0 | 1 | 1 |
| ZDHHC2 | -0.122 | P2_P3_P4_Lumbar_Stellate_Sphenopalatine_VS_P1_Coeliac | 0 | 1 | 1 | 1 | 1 | 0 | 1 | 1 |
| RBMXL1 | -0.122 | P1_P3_Lumbar_Coeliac_Stellate_Sphenopalatine_VS_P2_P4 | 1 | 0 | 1 | 0 | 1 | 1 | 1 | 1 |
| NRM | -0.122 | P1_P2_P3_Lumbar_Coeliac_Stellate_Sphenopalatine_VS_P4 | 1 | 1 | 1 | 0 | 1 | 1 | 1 | 1 |
| GRPEL2 | -0.122 | P4_Lumbar_Coeliac_Stellate_Sphenopalatine_VS_P1_P2_P3 | 0 | 0 | 0 | 1 | 1 | 1 | 1 | 1 |
| SAFB | -0.122 | P1_P4_Lumbar_Coeliac_Stellate_Sphenopalatine_VS_P2_P3 | 1 | 0 | 0 | 1 | 1 | 1 | 1 | 1 |
| VAMP7 | -0.122 | P1_P4_Lumbar_Coeliac_Stellate_Sphenopalatine_VS_P2_P3 | 1 | 0 | 0 | 1 | 1 | 1 | 1 | 1 |
| PRKRA | -0.122 | P2_P3_P4_Lumbar_Coeliac_Stellate_Sphenopalatine_VS_P1 | 0 | 1 | 1 | 1 | 1 | 1 | 1 | 1 |
| TMEM60 | -0.123 | P4_Lumbar_Stellate_VS_P1_P2_P3_Coeliac_Sphenopalatine | 0 | 0 | 0 | 1 | 1 | 0 | 1 | 0 |
| ECH1 | -0.123 | P1_P3_P4_Lumbar_Stellate_VS_P2_Coeliac_Sphenopalatine | 1 | 0 | 1 | 1 | 1 | 0 | 1 | 0 |
| DTYMK | -0.123 | P1_P3_P4_Lumbar_Stellate_Sphenopalatine_VS_P2_Coeliac | 1 | 0 | 1 | 1 | 1 | 0 | 1 | 1 |
| MAGOH | -0.123 | P1_P3_Lumbar_Coeliac_Stellate_Sphenopalatine_VS_P2_P4 | 1 | 0 | 1 | 0 | 1 | 1 | 1 | 1 |
| EIPR1 | -0.123 | P1_P4_Lumbar_Coeliac_Stellate_Sphenopalatine_VS_P2_P3 | 1 | 0 | 0 | 1 | 1 | 1 | 1 | 1 |
| PPA1 | -0.123 | P3_P4_Lumbar_Coeliac_Stellate_Sphenopalatine_VS_P1_P2 | 0 | 0 | 1 | 1 | 1 | 1 | 1 | 1 |
| CBR1 | -0.123 | P3_P4_Lumbar_Coeliac_Stellate_Sphenopalatine_VS_P1_P2 | 0 | 0 | 1 | 1 | 1 | 1 | 1 | 1 |
| ELOF1 | -0.124 | P1_P2_P3_Lumbar_Stellate_Sphenopalatine_VS_P4_Coeliac | 1 | 1 | 1 | 0 | 1 | 0 | 1 | 1 |
| MAF1 | -0.124 | P1_P3_P4_Lumbar_Stellate_Sphenopalatine_VS_P2_Coeliac | 1 | 0 | 1 | 1 | 1 | 0 | 1 | 1 |
| SFXN1 | -0.124 | P1_P2_P3_Lumbar_Coeliac_Stellate_Sphenopalatine_VS_P4 | 1 | 1 | 1 | 0 | 1 | 1 | 1 | 1 |
| ZMIZ1 | -0.124 | P2_P4_Lumbar_Coeliac_Stellate_Sphenopalatine_VS_P1_P3 | 0 | 1 | 0 | 1 | 1 | 1 | 1 | 1 |
| GM45716 | -0.124 | P3_P4_Lumbar_Coeliac_Stellate_Sphenopalatine_VS_P1_P2 | 0 | 0 | 1 | 1 | 1 | 1 | 1 | 1 |
| APOPT1 | -0.124 | P3_P4_Lumbar_Coeliac_Stellate_Sphenopalatine_VS_P1_P2 | 0 | 0 | 1 | 1 | 1 | 1 | 1 | 1 |
| CTNNB1 | -0.124 | P2_P3_P4_Lumbar_Coeliac_Stellate_Sphenopalatine_VS_P1 | 0 | 1 | 1 | 1 | 1 | 1 | 1 | 1 |
| PPP4C | -0.125 | P1_P3_P4_Stellate_VS_P2_Lumbar_Coeliac_Sphenopalatine | 1 | 0 | 1 | 1 | 0 | 0 | 1 | 0 |

| | | | | | | | | | | |
|---|---|---|---|---|---|---|---|---|---|---|
| MRPL35 | -0.125 | P3_P4_Lumbar_Stellate_VS_P1_P2_Coeliac_Sphenopalatine | 0 | 0 | 1 | 1 | 1 | 0 | 1 | 0 |
| DAZAP2 | -0.125 | P1_P2_P3_P4_Lumbar_Stellate_VS_Coeliac_Sphenopalatine | 1 | 1 | 1 | 1 | 1 | 0 | 1 | 0 |
| ANAPC11 | -0.125 | P2_P3_P4_Stellate_Sphenopalatine_VS_P1_Lumbar_Coeliac | 0 | 1 | 1 | 1 | 0 | 0 | 1 | 1 |
| RESP18 | -0.125 | P1_P2_P4_Lumbar_Stellate_Sphenopalatine_VS_P3_Coeliac | 1 | 1 | 0 | 1 | 1 | 0 | 1 | 1 |
| BLCAP | -0.125 | P1_P2_P4_Lumbar_Stellate_Sphenopalatine_VS_P3_Coeliac | 1 | 1 | 0 | 1 | 1 | 0 | 1 | 1 |
| PDE6D | -0.125 | P1_P3_P4_Lumbar_Stellate_Sphenopalatine_VS_P2_Coeliac | 1 | 0 | 1 | 1 | 1 | 0 | 1 | 1 |
| TMEM230 | -0.126 | P3_P4_Stellate_VS_P1_P2_Lumbar_Coeliac_Sphenopalatine | 0 | 0 | 1 | 1 | 0 | 0 | 1 | 0 |
| PPDPF | -0.126 | P2_P3_P4_Stellate_Sphenopalatine_VS_P1_Lumbar_Coeliac | 0 | 1 | 1 | 1 | 0 | 0 | 1 | 1 |
| MRPS18A | -0.126 | P2_P3_P4_Lumbar_Stellate_Sphenopalatine_VS_P1_Coeliac | 0 | 1 | 1 | 1 | 1 | 0 | 1 | 1 |
| TRAM1L1 | -0.126 | P1_P3_Lumbar_Coeliac_Stellate_Sphenopalatine_VS_P2_P4 | 1 | 0 | 1 | 0 | 1 | 1 | 1 | 1 |
| RAE1 | -0.126 | P1_P4_Lumbar_Coeliac_Stellate_Sphenopalatine_VS_P2_P3 | 1 | 0 | 0 | 1 | 1 | 1 | 1 | 1 |
| TMEM128 | -0.126 | P1_P4_Lumbar_Coeliac_Stellate_Sphenopalatine_VS_P2_P3 | 1 | 0 | 0 | 1 | 1 | 1 | 1 | 1 |
| GCSH | -0.126 | P2_P3_P4_Lumbar_Coeliac_Stellate_Sphenopalatine_VS_P1 | 0 | 1 | 1 | 1 | 1 | 1 | 1 | 1 |
| YLPM1 | -0.126 | P2_P3_P4_Lumbar_Coeliac_Stellate_Sphenopalatine_VS_P1 | 0 | 1 | 1 | 1 | 1 | 1 | 1 | 1 |
| CTNNA2 | -0.127 | P3_P4_Lumbar_Stellate_VS_P1_P2_Coeliac_Sphenopalatine | 0 | 0 | 1 | 1 | 1 | 0 | 1 | 0 |
| CACNB3 | -0.127 | P3_P4_Stellate_Sphenopalatine_VS_P1_P2_Lumbar_Coeliac | 0 | 0 | 1 | 1 | 0 | 0 | 1 | 1 |
| RNF4 | -0.127 | P1_P2_P3_Lumbar_Stellate_Sphenopalatine_VS_P4_Coeliac | 1 | 1 | 1 | 0 | 1 | 0 | 1 | 1 |
| LDHB | -0.127 | P3_P4_Lumbar_Stellate_Sphenopalatine_VS_P1_P2_Coeliac | 0 | 0 | 1 | 1 | 1 | 0 | 1 | 1 |
| POP7 | -0.127 | P1_P3_P4_Lumbar_Stellate_Sphenopalatine_VS_P2_Coeliac | 1 | 0 | 1 | 1 | 1 | 0 | 1 | 1 |
| LSM1 | -0.127 | P1_P3_P4_Lumbar_Stellate_Sphenopalatine_VS_P2_Coeliac | 1 | 0 | 1 | 1 | 1 | 0 | 1 | 1 |
| PIN1 | -0.127 | P1_P3_P4_Lumbar_Stellate_Sphenopalatine_VS_P2_Coeliac | 1 | 0 | 1 | 1 | 1 | 0 | 1 | 1 |
| MRPS34 | -0.127 | P1_P3_P4_Lumbar_Stellate_Sphenopalatine_VS_P2_Coeliac | 1 | 0 | 1 | 1 | 1 | 0 | 1 | 1 |
| PPP1CC | -0.127 | P2_P3_P4_Lumbar_Stellate_Sphenopalatine_VS_P1_Coeliac | 0 | 1 | 1 | 1 | 1 | 0 | 1 | 1 |
| PARP6 | -0.127 | P2_P3_P4_Lumbar_Stellate_Sphenopalatine_VS_P1_Coeliac | 0 | 1 | 1 | 1 | 1 | 0 | 1 | 1 |
| FKBP1B | -0.127 | P2_P3_P4_Lumbar_Stellate_Sphenopalatine_VS_P1_Coeliac | 0 | 1 | 1 | 1 | 1 | 0 | 1 | 1 |
| SNHG4 | -0.127 | P2_P3_P4_Lumbar_Stellate_Sphenopalatine_VS_P1_Coeliac | 0 | 1 | 1 | 1 | 1 | 0 | 1 | 1 |
| ATP8A1 | -0.128 | P1_P2_P3_P4_Lumbar_Stellate_VS_Coeliac_Sphenopalatine | 1 | 1 | 1 | 1 | 1 | 0 | 1 | 0 |
| OAZ2 | -0.128 | P2_P3_P4_Stellate_Sphenopalatine_VS_P1_Lumbar_Coeliac | 0 | 1 | 1 | 1 | 0 | 0 | 1 | 1 |
| CNIH2 | -0.128 | P2_P3_P4_Stellate_Sphenopalatine_VS_P1_Lumbar_Coeliac | 0 | 1 | 1 | 1 | 0 | 0 | 1 | 1 |
| BMERB1 | -0.128 | P1_P2_P3_Lumbar_Stellate_Sphenopalatine_VS_P4_Coeliac | 1 | 1 | 1 | 0 | 1 | 0 | 1 | 1 |

| | | | | | | | | | | | |
|---|---|---|---|---|---|---|---|---|---|---|---|
| NFU1 | -0.128 | P1_P3_P4_Lumbar_Stellate_Sphenopalatine_VS_P2_Coeliac | 1 | 0 | 1 | 1 | 1 | 0 | 1 | | 1 |
| RNF5 | -0.128 | P1_P3_P4_Lumbar_Stellate_Sphenopalatine_VS_P2_Coeliac | 1 | 0 | 1 | 1 | 1 | 0 | 1 | | 1 |
| B630019K06I | -0.128 | P1_P2_P3_Lumbar_Coeliac_Stellate_Sphenopalatine_VS_P4 | 1 | 1 | 1 | 0 | 1 | 1 | 1 | | 1 |
| ENHO | -0.128 | P2_P3_P4_Lumbar_Coeliac_Stellate_Sphenopalatine_VS_P1 | 0 | 1 | 1 | 1 | 1 | 1 | 1 | | 1 |
| DPF2 | -0.128 | P2_P3_P4_Lumbar_Coeliac_Stellate_Sphenopalatine_VS_P1 | 0 | 1 | 1 | 1 | 1 | 1 | 1 | | 1 |
| PIAS4 | -0.129 | P2_P4_Lumbar_Stellate_VS_P1_P3_Coeliac_Sphenopalatine | 0 | 1 | 0 | 1 | 1 | 0 | 1 | | 0 |
| 1810037I17R | -0.129 | P1_P3_P4_Stellate_Sphenopalatine_VS_P2_Lumbar_Coeliac | 1 | 0 | 1 | 1 | 0 | 0 | 1 | | 1 |
| CDK16 | -0.129 | P2_P3_P4_Stellate_Sphenopalatine_VS_P1_Lumbar_Coeliac | 0 | 1 | 1 | 1 | 0 | 0 | 1 | | 1 |
| GOT1 | -0.129 | P1_P2_P4_Lumbar_Stellate_Sphenopalatine_VS_P3_Coeliac | 1 | 1 | 0 | 1 | 1 | 0 | 1 | | 1 |
| BMYC | -0.129 | P1_P3_P4_Lumbar_Stellate_Sphenopalatine_VS_P2_Coeliac | 1 | 0 | 1 | 1 | 1 | 0 | 1 | | 1 |
| DPM2 | -0.129 | P1_P3_P4_Lumbar_Stellate_Sphenopalatine_VS_P2_Coeliac | 1 | 0 | 1 | 1 | 1 | 0 | 1 | | 1 |
| B3GALNT1 | -0.129 | P1_P3_P4_Lumbar_Stellate_Sphenopalatine_VS_P2_Coeliac | 1 | 0 | 1 | 1 | 1 | 0 | 1 | | 1 |
| RAMAC | -0.129 | P1_P3_P4_Lumbar_Stellate_Sphenopalatine_VS_P2_Coeliac | 1 | 0 | 1 | 1 | 1 | 0 | 1 | | 1 |
| NHP2 | -0.129 | P1_P3_P4_Lumbar_Stellate_Sphenopalatine_VS_P2_Coeliac | 1 | 0 | 1 | 1 | 1 | 0 | 1 | | 1 |
| 0610009B22I | -0.129 | P1_P3_P4_Lumbar_Stellate_Sphenopalatine_VS_P2_Coeliac | 1 | 0 | 1 | 1 | 1 | 0 | 1 | | 1 |
| PIP4P1 | -0.129 | P1_P3_P4_Lumbar_Stellate_Sphenopalatine_VS_P2_Coeliac | 1 | 0 | 1 | 1 | 1 | 0 | 1 | | 1 |
| POLR2H | -0.129 | P1_P3_P4_Lumbar_Stellate_Sphenopalatine_VS_P2_Coeliac | 1 | 0 | 1 | 1 | 1 | 0 | 1 | | 1 |
| TSPAN6 | -0.129 | P1_P2_P3_Lumbar_Coeliac_Stellate_Sphenopalatine_VS_P4 | 1 | 1 | 1 | 0 | 1 | 1 | 1 | | 1 |
| MED4 | -0.129 | P1_P4_Lumbar_Coeliac_Stellate_Sphenopalatine_VS_P2_P3 | 1 | 0 | 0 | 1 | 1 | 1 | 1 | | 1 |
| FEZ2 | -0.13 | P1_P4_Lumbar_Stellate_VS_P2_P3_Coeliac_Sphenopalatine | 1 | 0 | 0 | 1 | 1 | 0 | 1 | | 0 |
| EXOSC4 | -0.13 | P1_P3_P4_Lumbar_Stellate_VS_P2_Coeliac_Sphenopalatine | 1 | 0 | 1 | 1 | 1 | 0 | 1 | | 0 |
| RNF126 | -0.13 | P1_P2_P3_P4_Lumbar_Stellate_VS_Coeliac_Sphenopalatine | 1 | 1 | 1 | 1 | 1 | 0 | 1 | | 0 |
| 3830406C13I | -0.13 | P1_P2_P3_P4_Lumbar_Stellate_VS_Coeliac_Sphenopalatine | 1 | 1 | 1 | 1 | 1 | 0 | 1 | | 0 |
| MFF | -0.13 | P3_P4_Stellate_Sphenopalatine_VS_P1_P2_Lumbar_Coeliac | 0 | 0 | 1 | 1 | 0 | 0 | 1 | | 1 |
| AKR1B3 | -0.13 | P3_P4_Lumbar_Stellate_Sphenopalatine_VS_P1_P2_Coeliac | 0 | 0 | 1 | 1 | 1 | 0 | 1 | | 1 |
| COMMD9 | -0.13 | P1_P3_P4_Lumbar_Stellate_Sphenopalatine_VS_P2_Coeliac | 1 | 0 | 1 | 1 | 1 | 0 | 1 | | 1 |
| ATP6V1C1 | -0.13 | P1_P3_P4_Lumbar_Stellate_Sphenopalatine_VS_P2_Coeliac | 1 | 0 | 1 | 1 | 1 | 0 | 1 | | 1 |
| SRA1 | -0.13 | P1_P3_P4_Lumbar_Stellate_Sphenopalatine_VS_P2_Coeliac | 1 | 0 | 1 | 1 | 1 | 0 | 1 | | 1 |
| BSCL2 | -0.13 | P2_P3_P4_Lumbar_Stellate_Sphenopalatine_VS_P1_Coeliac | 0 | 1 | 1 | 1 | 1 | 0 | 1 | | 1 |
| PRKRIP1 | -0.13 | P1_P3_Lumbar_Coeliac_Stellate_Sphenopalatine_VS_P2_P4 | 1 | 0 | 1 | 0 | 1 | 1 | 1 | | 1 |

| | | | | | | | | | | |
|---|---|---|---|---|---|---|---|---|---|---|
| DTX3 | -0.13 | P2_P3_P4_Lumbar_Coeliac_Stellate_Sphenopalatine_VS_P1 | 0 | 1 | 1 | 1 | 1 | 1 | 1 | 1 |
| SERP2 | -0.13 | P2_P3_P4_Lumbar_Coeliac_Stellate_Sphenopalatine_VS_P1 | 0 | 1 | 1 | 1 | 1 | 1 | 1 | 1 |
| HNRNPH2 | -0.13 | P2_P3_P4_Lumbar_Coeliac_Stellate_Sphenopalatine_VS_P1 | 0 | 1 | 1 | 1 | 1 | 1 | 1 | 1 |
| IFT27 | -0.131 | P1_P3_P4_Lumbar_Stellate_VS_P2_Coeliac_Sphenopalatine | 1 | 0 | 1 | 1 | 1 | 0 | 1 | 0 |
| CKB | -0.131 | P1_P2_P3_P4_Lumbar_Stellate_VS_Coeliac_Sphenopalatine | 1 | 1 | 1 | 1 | 1 | 0 | 1 | 0 |
| RAB11A | -0.131 | P3_P4_Stellate_Sphenopalatine_VS_P1_P2_Lumbar_Coeliac | 0 | 0 | 1 | 1 | 0 | 0 | 1 | 1 |
| SSNA1 | -0.131 | P2_P3_P4_Stellate_Sphenopalatine_VS_P1_Lumbar_Coeliac | 0 | 1 | 1 | 1 | 0 | 0 | 1 | 1 |
| RFK | -0.131 | P2_P3_P4_Stellate_Sphenopalatine_VS_P1_Lumbar_Coeliac | 0 | 1 | 1 | 1 | 0 | 0 | 1 | 1 |
| DZANK1 | -0.131 | P1_P2_P4_Lumbar_Stellate_Sphenopalatine_VS_P3_Coeliac | 1 | 1 | 0 | 1 | 1 | 0 | 1 | 1 |
| ZCCHC17 | -0.131 | P1_P3_P4_Lumbar_Stellate_Sphenopalatine_VS_P2_Coeliac | 1 | 0 | 1 | 1 | 1 | 0 | 1 | 1 |
| ENY2 | -0.131 | P1_P3_P4_Lumbar_Stellate_Sphenopalatine_VS_P2_Coeliac | 1 | 0 | 1 | 1 | 1 | 0 | 1 | 1 |
| TOMM5 | -0.131 | P2_P3_P4_Lumbar_Stellate_Sphenopalatine_VS_P1_Coeliac | 0 | 1 | 1 | 1 | 1 | 0 | 1 | 1 |
| H2AFJ | -0.131 | P2_P3_P4_Lumbar_Stellate_Sphenopalatine_VS_P1_Coeliac | 0 | 1 | 1 | 1 | 1 | 0 | 1 | 1 |
| TMEM50B | -0.131 | P2_P3_P4_Lumbar_Stellate_Sphenopalatine_VS_P1_Coeliac | 0 | 1 | 1 | 1 | 1 | 0 | 1 | 1 |
| PRDX3 | -0.131 | P2_P3_P4_Lumbar_Stellate_Sphenopalatine_VS_P1_Coeliac | 0 | 1 | 1 | 1 | 1 | 0 | 1 | 1 |
| TSPAN7 | -0.131 | P2_P3_P4_Lumbar_Stellate_Sphenopalatine_VS_P1_Coeliac | 0 | 1 | 1 | 1 | 1 | 0 | 1 | 1 |
| ENO1 | -0.131 | P1_P2_P3_P4_Coeliac_Stellate_Sphenopalatine_VS_Lumbar | 1 | 1 | 1 | 1 | 0 | 1 | 1 | 1 |
| HAND2 | -0.131 | P1_P2_P3_Lumbar_Coeliac_Stellate_Sphenopalatine_VS_P4 | 1 | 1 | 1 | 0 | 1 | 1 | 1 | 1 |
| PEX11B | -0.131 | P1_P4_Lumbar_Coeliac_Stellate_Sphenopalatine_VS_P2_P3 | 1 | 0 | 0 | 1 | 1 | 1 | 1 | 1 |
| RB1CC1 | -0.131 | P2_P3_P4_Lumbar_Coeliac_Stellate_Sphenopalatine_VS_P1 | 0 | 1 | 1 | 1 | 1 | 1 | 1 | 1 |
| SNRPD3 | -0.132 | P3_P4_Lumbar_Stellate_VS_P1_P2_Coeliac_Sphenopalatine | 0 | 0 | 1 | 1 | 1 | 0 | 1 | 0 |
| NT5C | -0.132 | P3_P4_Lumbar_Stellate_VS_P1_P2_Coeliac_Sphenopalatine | 0 | 0 | 1 | 1 | 1 | 0 | 1 | 0 |
| CFAP20 | -0.132 | P2_P3_P4_Lumbar_Stellate_VS_P1_Coeliac_Sphenopalatine | 0 | 1 | 1 | 1 | 1 | 0 | 1 | 0 |
| CCDC115 | -0.132 | P1_P2_P3_P4_Lumbar_Stellate_VS_Coeliac_Sphenopalatine | 1 | 1 | 1 | 1 | 1 | 0 | 1 | 0 |
| RSL24D1 | -0.132 | P1_P2_P3_P4_Lumbar_Stellate_VS_Coeliac_Sphenopalatine | 1 | 1 | 1 | 1 | 1 | 0 | 1 | 0 |
| SNHG8 | -0.132 | P1_P2_P3_Stellate_Sphenopalatine_VS_P4_Lumbar_Coeliac | 1 | 1 | 1 | 0 | 0 | 0 | 1 | 1 |
| WBP2 | -0.132 | P2_P3_P4_Stellate_Sphenopalatine_VS_P1_Lumbar_Coeliac | 0 | 1 | 1 | 1 | 0 | 0 | 1 | 1 |
| TMEM183A | -0.132 | P1_P2_P3_Lumbar_Stellate_Sphenopalatine_VS_P4_Coeliac | 1 | 1 | 1 | 0 | 1 | 0 | 1 | 1 |
| NAA20 | -0.132 | P1_P2_P3_Lumbar_Stellate_Sphenopalatine_VS_P4_Coeliac | 1 | 1 | 1 | 0 | 1 | 0 | 1 | 1 |
| TIMM17A | -0.132 | P1_P2_P4_Lumbar_Stellate_Sphenopalatine_VS_P3_Coeliac | 1 | 1 | 0 | 1 | 1 | 0 | 1 | 1 |

| | | | | | | | | | | |
|---|---|---|---|---|---|---|---|---|---|---|
| GIPC1 | -0.132 | P1_P3_P4_Lumbar_Stellate_Sphenopalatine_VS_P2_Coeliac | 1 | 0 | 1 | 1 | 1 | 0 | 1 | 1 |
| MTCH2 | -0.132 | P2_P3_P4_Lumbar_Stellate_Sphenopalatine_VS_P1_Coeliac | 0 | 1 | 1 | 1 | 1 | 0 | 1 | 1 |
| STX2 | -0.132 | P2_P3_P4_Lumbar_Stellate_Sphenopalatine_VS_P1_Coeliac | 0 | 1 | 1 | 1 | 1 | 0 | 1 | 1 |
| ISOC1 | -0.132 | P1_P2_P3_P4_Coeliac_Stellate_Sphenopalatine_VS_Lumbar | 1 | 1 | 1 | 1 | 0 | 1 | 1 | 1 |
| CDC16 | -0.132 | P1_P4_Lumbar_Coeliac_Stellate_Sphenopalatine_VS_P2_P3 | 1 | 0 | 0 | 1 | 1 | 1 | 1 | 1 |
| BAG5 | -0.132 | P1_P4_Lumbar_Coeliac_Stellate_Sphenopalatine_VS_P2_P3 | 1 | 0 | 0 | 1 | 1 | 1 | 1 | 1 |
| EFNA5 | -0.133 | P3_P4_Lumbar_Stellate_VS_P1_P2_Coeliac_Sphenopalatine | 0 | 0 | 1 | 1 | 1 | 0 | 1 | 0 |
| CKS1B | -0.133 | P1_P3_P4_Lumbar_Stellate_VS_P2_Coeliac_Sphenopalatine | 1 | 0 | 1 | 1 | 1 | 0 | 1 | 0 |
| ACYP1 | -0.133 | P1_P3_P4_Lumbar_Stellate_VS_P2_Coeliac_Sphenopalatine | 1 | 0 | 1 | 1 | 1 | 0 | 1 | 0 |
| NKIRAS2 | -0.133 | P1_P2_P3_P4_Lumbar_Stellate_VS_Coeliac_Sphenopalatine | 1 | 1 | 1 | 1 | 1 | 0 | 1 | 0 |
| ALKBH8 | -0.133 | P2_Lumbar_Coeliac_Stellate_VS_P1_P3_P4_Sphenopalatine | 0 | 1 | 0 | 0 | 1 | 1 | 1 | 0 |
| UBE2V1 | -0.133 | P2_P3_P4_Stellate_Sphenopalatine_VS_P1_Lumbar_Coeliac | 0 | 1 | 1 | 1 | 0 | 0 | 1 | 1 |
| RAB5B | -0.133 | P2_P3_P4_Stellate_Sphenopalatine_VS_P1_Lumbar_Coeliac | 0 | 1 | 1 | 1 | 0 | 0 | 1 | 1 |
| ACTL6B | -0.133 | P1_P2_P3_Lumbar_Stellate_Sphenopalatine_VS_P4_Coeliac | 1 | 1 | 1 | 0 | 1 | 0 | 1 | 1 |
| MCAM | -0.133 | P1_P2_P4_Lumbar_Stellate_Sphenopalatine_VS_P3_Coeliac | 1 | 1 | 0 | 1 | 1 | 0 | 1 | 1 |
| EIF1A | -0.133 | P1_P3_P4_Lumbar_Stellate_Sphenopalatine_VS_P2_Coeliac | 1 | 0 | 1 | 1 | 1 | 0 | 1 | 1 |
| TMSB15B2 | -0.133 | P1_P3_P4_Lumbar_Stellate_Sphenopalatine_VS_P2_Coeliac | 1 | 0 | 1 | 1 | 1 | 0 | 1 | 1 |
| PINK1 | -0.133 | P2_P3_P4_Lumbar_Stellate_Sphenopalatine_VS_P1_Coeliac | 0 | 1 | 1 | 1 | 1 | 0 | 1 | 1 |
| RAB1A | -0.133 | P2_P3_P4_Lumbar_Stellate_Sphenopalatine_VS_P1_Coeliac | 0 | 1 | 1 | 1 | 1 | 0 | 1 | 1 |
| FOPNL | -0.133 | P2_P3_P4_Lumbar_Stellate_Sphenopalatine_VS_P1_Coeliac | 0 | 1 | 1 | 1 | 1 | 0 | 1 | 1 |
| WTAP | -0.133 | P2_P3_P4_Lumbar_Stellate_Sphenopalatine_VS_P1_Coeliac | 0 | 1 | 1 | 1 | 1 | 0 | 1 | 1 |
| SMARCE1 | -0.133 | P3_Lumbar_Coeliac_Stellate_Sphenopalatine_VS_P1_P2_P4 | 0 | 0 | 1 | 0 | 1 | 1 | 1 | 1 |
| RLIM | -0.133 | P2_P3_P4_Lumbar_Coeliac_Stellate_Sphenopalatine_VS_P1 | 0 | 1 | 1 | 1 | 1 | 1 | 1 | 1 |
| SEC13 | -0.134 | P3_P4_Lumbar_Stellate_VS_P1_P2_Coeliac_Sphenopalatine | 0 | 0 | 1 | 1 | 1 | 0 | 1 | 0 |
| SRRD | -0.134 | P1_P3_P4_Lumbar_Stellate_VS_P2_Coeliac_Sphenopalatine | 1 | 0 | 1 | 1 | 1 | 0 | 1 | 0 |
| SLC35A1 | -0.134 | P1_P2_P3_P4_Lumbar_Stellate_VS_Coeliac_Sphenopalatine | 1 | 1 | 1 | 1 | 1 | 0 | 1 | 0 |
| OCIAD1 | -0.134 | P3_P4_Stellate_Sphenopalatine_VS_P1_P2_Lumbar_Coeliac | 0 | 0 | 1 | 1 | 0 | 0 | 1 | 1 |
| ZFP580 | -0.134 | P3_P4_Stellate_Sphenopalatine_VS_P1_P2_Lumbar_Coeliac | 0 | 0 | 1 | 1 | 0 | 0 | 1 | 1 |
| CPLX1 | -0.134 | P1_P3_P4_Stellate_Sphenopalatine_VS_P2_Lumbar_Coeliac | 1 | 0 | 1 | 1 | 0 | 0 | 1 | 1 |
| ARAF | -0.134 | P2_P3_P4_Stellate_Sphenopalatine_VS_P1_Lumbar_Coeliac | 0 | 1 | 1 | 1 | 0 | 0 | 1 | 1 |

| | | | | | | | | | | |
|---|---|---|---|---|---|---|---|---|---|---|
| RBM8A | -0.134 | P1_P3_P4_Lumbar_Stellate_Sphenopalatine_VS_P2_Coeliac | 1 | 0 | 1 | 1 | 1 | 0 | 1 | 1 |
| CGREF1 | -0.134 | P1_P3_P4_Lumbar_Stellate_Sphenopalatine_VS_P2_Coeliac | 1 | 0 | 1 | 1 | 1 | 0 | 1 | 1 |
| ARL6IP4 | -0.134 | P1_P3_P4_Lumbar_Stellate_Sphenopalatine_VS_P2_Coeliac | 1 | 0 | 1 | 1 | 1 | 0 | 1 | 1 |
| COPRS | -0.134 | P1_P3_P4_Lumbar_Stellate_Sphenopalatine_VS_P2_Coeliac | 1 | 0 | 1 | 1 | 1 | 0 | 1 | 1 |
| MRPL28 | -0.134 | P1_P3_P4_Lumbar_Stellate_Sphenopalatine_VS_P2_Coeliac | 1 | 0 | 1 | 1 | 1 | 0 | 1 | 1 |
| UFC1 | -0.134 | P2_P3_P4_Lumbar_Stellate_Sphenopalatine_VS_P1_Coeliac | 0 | 1 | 1 | 1 | 1 | 0 | 1 | 1 |
| LSM6 | -0.134 | P2_P3_P4_Lumbar_Stellate_Sphenopalatine_VS_P1_Coeliac | 0 | 1 | 1 | 1 | 1 | 0 | 1 | 1 |
| CD81 | -0.134 | P1_P2_P3_P4_Coeliac_Stellate_Sphenopalatine_VS_Lumbar | 1 | 1 | 1 | 1 | 0 | 1 | 1 | 1 |
| NELFB | -0.134 | P1_P2_P3_Lumbar_Coeliac_Stellate_Sphenopalatine_VS_P4 | 1 | 1 | 1 | 0 | 1 | 1 | 1 | 1 |
| CRYZL1 | -0.134 | P1_P4_Lumbar_Coeliac_Stellate_Sphenopalatine_VS_P2_P3 | 1 | 0 | 0 | 1 | 1 | 1 | 1 | 1 |
| ATP6V1H | -0.134 | P2_P3_P4_Lumbar_Coeliac_Stellate_Sphenopalatine_VS_P1 | 0 | 1 | 1 | 1 | 1 | 1 | 1 | 1 |
| UBL4A | -0.134 | P2_P3_P4_Lumbar_Coeliac_Stellate_Sphenopalatine_VS_P1 | 0 | 1 | 1 | 1 | 1 | 1 | 1 | 1 |
| RABGAP1L | -0.135 | P1_P2_P4_Lumbar_Stellate_VS_P3_Coeliac_Sphenopalatine | 1 | 1 | 0 | 1 | 1 | 0 | 1 | 0 |
| TMEM41A | -0.135 | P3_P4_Lumbar_Stellate_VS_P1_P2_Coeliac_Sphenopalatine | 0 | 0 | 1 | 1 | 1 | 0 | 1 | 0 |
| CNTFR | -0.135 | P1_P2_P3_P4_Lumbar_Stellate_VS_Coeliac_Sphenopalatine | 1 | 1 | 1 | 1 | 1 | 0 | 1 | 0 |
| PLP2 | -0.135 | P1_P2_P3_P4_Lumbar_Stellate_VS_Coeliac_Sphenopalatine | 1 | 1 | 1 | 1 | 1 | 0 | 1 | 0 |
| SNRPB2 | -0.135 | P2_P3_P4_Stellate_Sphenopalatine_VS_P1_Lumbar_Coeliac | 0 | 1 | 1 | 1 | 0 | 0 | 1 | 1 |
| VTI1B | -0.135 | P2_P3_P4_Stellate_Sphenopalatine_VS_P1_Lumbar_Coeliac | 0 | 1 | 1 | 1 | 0 | 0 | 1 | 1 |
| BC004004 | -0.135 | P2_P3_P4_Stellate_Sphenopalatine_VS_P1_Lumbar_Coeliac | 0 | 1 | 1 | 1 | 0 | 0 | 1 | 1 |
| MRPL41 | -0.135 | P1_P3_P4_Lumbar_Stellate_Sphenopalatine_VS_P2_Coeliac | 1 | 0 | 1 | 1 | 1 | 0 | 1 | 1 |
| SMIM8 | -0.135 | P1_P3_P4_Lumbar_Stellate_Sphenopalatine_VS_P2_Coeliac | 1 | 0 | 1 | 1 | 1 | 0 | 1 | 1 |
| MANF | -0.135 | P1_P3_P4_Lumbar_Stellate_Sphenopalatine_VS_P2_Coeliac | 1 | 0 | 1 | 1 | 1 | 0 | 1 | 1 |
| AP4S1 | -0.135 | P1_P3_P4_Lumbar_Stellate_Sphenopalatine_VS_P2_Coeliac | 1 | 0 | 1 | 1 | 1 | 0 | 1 | 1 |
| PIGP | -0.135 | P1_P3_P4_Lumbar_Stellate_Sphenopalatine_VS_P2_Coeliac | 1 | 0 | 1 | 1 | 1 | 0 | 1 | 1 |
| CELF6 | -0.135 | P2_P3_P4_Lumbar_Stellate_Sphenopalatine_VS_P1_Coeliac | 0 | 1 | 1 | 1 | 1 | 0 | 1 | 1 |
| STUB1 | -0.135 | P2_P3_P4_Lumbar_Stellate_Sphenopalatine_VS_P1_Coeliac | 0 | 1 | 1 | 1 | 1 | 0 | 1 | 1 |
| TSPAN13 | -0.135 | P3_P4_Coeliac_Stellate_Sphenopalatine_VS_P1_P2_Lumbar | 0 | 0 | 1 | 1 | 0 | 1 | 1 | 1 |
| DDB1 | -0.135 | P2_P3_P4_Lumbar_Coeliac_Stellate_Sphenopalatine_VS_P1 | 0 | 1 | 1 | 1 | 1 | 1 | 1 | 1 |
| MSN | -0.135 | P2_P3_P4_Lumbar_Coeliac_Stellate_Sphenopalatine_VS_P1 | 0 | 1 | 1 | 1 | 1 | 1 | 1 | 1 |
| DUSP1 | -0.136 | P1_P2_P3_P4_Lumbar_Stellate_VS_Coeliac_Sphenopalatine | 1 | 1 | 1 | 1 | 1 | 0 | 1 | 0 |

| | | | | | | | | | | |
|---|---|---|---|---|---|---|---|---|---|---|
| MED28 | -0.136 | P3_P4_Stellate_Sphenopalatine_VS_P1_P2_Lumbar_Coeliac | 0 | 0 | 1 | 1 | 0 | 0 | 1 | 1 |
| RSRP1 | -0.136 | P2_P3_P4_Stellate_Sphenopalatine_VS_P1_Lumbar_Coeliac | 0 | 1 | 1 | 1 | 0 | 0 | 1 | 1 |
| FUNDC1 | -0.136 | P2_P3_P4_Stellate_Sphenopalatine_VS_P1_Lumbar_Coeliac | 0 | 1 | 1 | 1 | 0 | 0 | 1 | 1 |
| MRPS18C | -0.136 | P1_P3_P4_Lumbar_Stellate_Sphenopalatine_VS_P2_Coeliac | 1 | 0 | 1 | 1 | 1 | 0 | 1 | 1 |
| CMAS | -0.136 | P1_P3_P4_Lumbar_Stellate_Sphenopalatine_VS_P2_Coeliac | 1 | 0 | 1 | 1 | 1 | 0 | 1 | 1 |
| POLR2I | -0.136 | P1_P3_P4_Lumbar_Stellate_Sphenopalatine_VS_P2_Coeliac | 1 | 0 | 1 | 1 | 1 | 0 | 1 | 1 |
| PSMG4 | -0.136 | P1_P3_P4_Lumbar_Stellate_Sphenopalatine_VS_P2_Coeliac | 1 | 0 | 1 | 1 | 1 | 0 | 1 | 1 |
| PIN4 | -0.136 | P1_P3_P4_Lumbar_Stellate_Sphenopalatine_VS_P2_Coeliac | 1 | 0 | 1 | 1 | 1 | 0 | 1 | 1 |
| CSK | -0.136 | P2_P3_P4_Lumbar_Stellate_Sphenopalatine_VS_P1_Coeliac | 0 | 1 | 1 | 1 | 1 | 0 | 1 | 1 |
| ATP1B3 | -0.136 | P2_P3_P4_Lumbar_Stellate_Sphenopalatine_VS_P1_Coeliac | 0 | 1 | 1 | 1 | 1 | 0 | 1 | 1 |
| SMPDL3A | -0.136 | P2_P3_P4_Lumbar_Stellate_Sphenopalatine_VS_P1_Coeliac | 0 | 1 | 1 | 1 | 1 | 0 | 1 | 1 |
| NABP2 | -0.136 | P2_P3_P4_Lumbar_Stellate_Sphenopalatine_VS_P1_Coeliac | 0 | 1 | 1 | 1 | 1 | 0 | 1 | 1 |
| MBNL2 | -0.136 | P1_P2_P3_P4_Coeliac_Stellate_Sphenopalatine_VS_Lumbar | 1 | 1 | 1 | 1 | 0 | 1 | 1 | 1 |
| WASHC2 | -0.136 | P2_P4_Lumbar_Coeliac_Stellate_Sphenopalatine_VS_P1_P3 | 0 | 1 | 0 | 1 | 1 | 1 | 1 | 1 |
| ALKBH7 | -0.137 | P1_P3_P4_Lumbar_Stellate_VS_P2_Coeliac_Sphenopalatine | 1 | 0 | 1 | 1 | 1 | 0 | 1 | 0 |
| YIF1A | -0.137 | P1_P3_P4_Lumbar_Stellate_VS_P2_Coeliac_Sphenopalatine | 1 | 0 | 1 | 1 | 1 | 0 | 1 | 0 |
| COQ2 | -0.137 | P1_P2_P3_P4_Lumbar_Stellate_VS_Coeliac_Sphenopalatine | 1 | 1 | 1 | 1 | 1 | 0 | 1 | 0 |
| IFNGR2 | -0.137 | P4_Stellate_Sphenopalatine_VS_P1_P2_P3_Lumbar_Coeliac | 0 | 0 | 0 | 1 | 0 | 0 | 1 | 1 |
| CADM1 | -0.137 | P1_P2_P4_Stellate_Sphenopalatine_VS_P3_Lumbar_Coeliac | 1 | 1 | 0 | 1 | 0 | 0 | 1 | 1 |
| GPX1 | -0.137 | P3_P4_Stellate_Sphenopalatine_VS_P1_P2_Lumbar_Coeliac | 0 | 0 | 1 | 1 | 0 | 0 | 1 | 1 |
| MAP2K2 | -0.137 | P3_P4_Stellate_Sphenopalatine_VS_P1_P2_Lumbar_Coeliac | 0 | 0 | 1 | 1 | 0 | 0 | 1 | 1 |
| GLO1 | -0.137 | P3_P4_Stellate_Sphenopalatine_VS_P1_P2_Lumbar_Coeliac | 0 | 0 | 1 | 1 | 0 | 0 | 1 | 1 |
| CCL27A | -0.137 | P1_P3_P4_Stellate_Sphenopalatine_VS_P2_Lumbar_Coeliac | 1 | 0 | 1 | 1 | 0 | 0 | 1 | 1 |
| APBA2 | -0.137 | P2_P3_P4_Stellate_Sphenopalatine_VS_P1_Lumbar_Coeliac | 0 | 1 | 1 | 1 | 0 | 0 | 1 | 1 |
| NTAN1 | -0.137 | P2_P3_P4_Stellate_Sphenopalatine_VS_P1_Lumbar_Coeliac | 0 | 1 | 1 | 1 | 0 | 0 | 1 | 1 |
| MEIS3 | -0.137 | P3_Lumbar_Stellate_Sphenopalatine_VS_P1_P2_P4_Coeliac | 0 | 0 | 1 | 0 | 1 | 0 | 1 | 1 |
| MRPL11 | -0.137 | P3_P4_Lumbar_Stellate_Sphenopalatine_VS_P1_P2_Coeliac | 0 | 0 | 1 | 1 | 1 | 0 | 1 | 1 |
| MORN4 | -0.137 | P3_P4_Lumbar_Stellate_Sphenopalatine_VS_P1_P2_Coeliac | 0 | 0 | 1 | 1 | 1 | 0 | 1 | 1 |
| FMC1 | -0.137 | P1_P3_P4_Lumbar_Stellate_Sphenopalatine_VS_P2_Coeliac | 1 | 0 | 1 | 1 | 1 | 0 | 1 | 1 |
| PMM1 | -0.137 | P1_P3_P4_Lumbar_Stellate_Sphenopalatine_VS_P2_Coeliac | 1 | 0 | 1 | 1 | 1 | 0 | 1 | 1 |

| | | | | | | | | | | |
|---|---|---|---|---|---|---|---|---|---|---|
| PGP | -0.137 | P1_P3_P4_Lumbar_Stellate_Sphenopalatine_VS_P2_Coeliac | 1 | 0 | 1 | 1 | 1 | 0 | 1 | 1 |
| APOO | -0.137 | P1_P3_P4_Lumbar_Stellate_Sphenopalatine_VS_P2_Coeliac | 1 | 0 | 1 | 1 | 1 | 0 | 1 | 1 |
| MCRIP1 | -0.137 | P2_P3_P4_Lumbar_Stellate_Sphenopalatine_VS_P1_Coeliac | 0 | 1 | 1 | 1 | 1 | 0 | 1 | 1 |
| CNIH1 | -0.137 | P2_P3_P4_Lumbar_Stellate_Sphenopalatine_VS_P1_Coeliac | 0 | 1 | 1 | 1 | 1 | 0 | 1 | 1 |
| CAPN1 | -0.137 | P1_P2_P3_Lumbar_Coeliac_Stellate_Sphenopalatine_VS_P4 | 1 | 1 | 1 | 0 | 1 | 1 | 1 | 1 |
| MRPS25 | -0.137 | P1_P4_Lumbar_Coeliac_Stellate_Sphenopalatine_VS_P2_P3 | 1 | 0 | 0 | 1 | 1 | 1 | 1 | 1 |
| NAE1 | -0.137 | P1_P4_Lumbar_Coeliac_Stellate_Sphenopalatine_VS_P2_P3 | 1 | 0 | 0 | 1 | 1 | 1 | 1 | 1 |
| VPS4B | -0.137 | P2_P3_P4_Lumbar_Coeliac_Stellate_Sphenopalatine_VS_P1 | 0 | 1 | 1 | 1 | 1 | 1 | 1 | 1 |
| ENSA | -0.137 | P2_P3_P4_Lumbar_Coeliac_Stellate_Sphenopalatine_VS_P1 | 0 | 1 | 1 | 1 | 1 | 1 | 1 | 1 |
| CHD5 | -0.137 | P2_P3_P4_Lumbar_Coeliac_Stellate_Sphenopalatine_VS_P1 | 0 | 1 | 1 | 1 | 1 | 1 | 1 | 1 |
| DNAJC12 | -0.138 | P3_P4_Stellate_VS_P1_P2_Lumbar_Coeliac_Sphenopalatine | 0 | 0 | 1 | 1 | 0 | 0 | 1 | 0 |
| RAP1B | -0.138 | P3_P4_Stellate_VS_P1_P2_Lumbar_Coeliac_Sphenopalatine | 0 | 0 | 1 | 1 | 0 | 0 | 1 | 0 |
| ETFB | -0.138 | P4_Lumbar_Stellate_VS_P1_P2_P3_Coeliac_Sphenopalatine | 0 | 0 | 0 | 1 | 1 | 0 | 1 | 0 |
| ZC2HC1A | -0.138 | P3_P4_Stellate_Sphenopalatine_VS_P1_P2_Lumbar_Coeliac | 0 | 0 | 1 | 1 | 0 | 0 | 1 | 1 |
| ANAPC13 | -0.138 | P3_P4_Stellate_Sphenopalatine_VS_P1_P2_Lumbar_Coeliac | 0 | 0 | 1 | 1 | 0 | 0 | 1 | 1 |
| ATP2B1 | -0.138 | P3_P4_Stellate_Sphenopalatine_VS_P1_P2_Lumbar_Coeliac | 0 | 0 | 1 | 1 | 0 | 0 | 1 | 1 |
| SNX3 | -0.138 | P1_P3_P4_Stellate_Sphenopalatine_VS_P2_Lumbar_Coeliac | 1 | 0 | 1 | 1 | 0 | 0 | 1 | 1 |
| SMIM14 | -0.138 | P2_P3_P4_Stellate_Sphenopalatine_VS_P1_Lumbar_Coeliac | 0 | 1 | 1 | 1 | 0 | 0 | 1 | 1 |
| CHRNA3 | -0.138 | P2_P3_P4_Stellate_Sphenopalatine_VS_P1_Lumbar_Coeliac | 0 | 1 | 1 | 1 | 0 | 0 | 1 | 1 |
| YKT6 | -0.138 | P1_P2_P3_Lumbar_Stellate_Sphenopalatine_VS_P4_Coeliac | 1 | 1 | 1 | 0 | 1 | 0 | 1 | 1 |
| DNAJC15 | -0.138 | P1_P2_P3_Lumbar_Stellate_Sphenopalatine_VS_P4_Coeliac | 1 | 1 | 1 | 0 | 1 | 0 | 1 | 1 |
| SRI | -0.138 | P1_P2_P4_Lumbar_Stellate_Sphenopalatine_VS_P3_Coeliac | 1 | 1 | 0 | 1 | 1 | 0 | 1 | 1 |
| POP5 | -0.138 | P3_P4_Lumbar_Stellate_Sphenopalatine_VS_P1_P2_Coeliac | 0 | 0 | 1 | 1 | 1 | 0 | 1 | 1 |
| SAR1B | -0.138 | P1_P3_P4_Lumbar_Stellate_Sphenopalatine_VS_P2_Coeliac | 1 | 0 | 1 | 1 | 1 | 0 | 1 | 1 |
| RDH14 | -0.138 | P1_P3_P4_Lumbar_Stellate_Sphenopalatine_VS_P2_Coeliac | 1 | 0 | 1 | 1 | 1 | 0 | 1 | 1 |
| PPP1R11 | -0.138 | P1_P3_P4_Lumbar_Stellate_Sphenopalatine_VS_P2_Coeliac | 1 | 0 | 1 | 1 | 1 | 0 | 1 | 1 |
| RBM17 | -0.138 | P2_P3_P4_Lumbar_Stellate_Sphenopalatine_VS_P1_Coeliac | 0 | 1 | 1 | 1 | 1 | 0 | 1 | 1 |
| MRPL53 | -0.138 | P2_P3_P4_Lumbar_Stellate_Sphenopalatine_VS_P1_Coeliac | 0 | 1 | 1 | 1 | 1 | 0 | 1 | 1 |
| CDIPT | -0.138 | P2_P3_P4_Lumbar_Stellate_Sphenopalatine_VS_P1_Coeliac | 0 | 1 | 1 | 1 | 1 | 0 | 1 | 1 |
| SELENOM | -0.138 | P2_P3_P4_Lumbar_Stellate_Sphenopalatine_VS_P1_Coeliac | 0 | 1 | 1 | 1 | 1 | 0 | 1 | 1 |

| | | | | | | | | | | |
|---|---|---|---|---|---|---|---|---|---|---|
| 2410006H16I | -0.138 | P2_P3_P4_Lumbar_Stellate_Sphenopalatine_VS_P1_Coeliac | 0 | 1 | 1 | 1 | 1 | 0 | 1 | 1 |
| FLOT2 | -0.138 | P1_P2_P3_P4_Coeliac_Stellate_Sphenopalatine_VS_Lumbar | 1 | 1 | 1 | 1 | 0 | 1 | 1 | 1 |
| UTP11 | -0.138 | P1_P4_Lumbar_Coeliac_Stellate_Sphenopalatine_VS_P2_P3 | 1 | 0 | 0 | 1 | 1 | 1 | 1 | 1 |
| YARS | -0.138 | P1_P4_Lumbar_Coeliac_Stellate_Sphenopalatine_VS_P2_P3 | 1 | 0 | 0 | 1 | 1 | 1 | 1 | 1 |
| TCTEX1D2 | -0.138 | P1_P4_Lumbar_Coeliac_Stellate_Sphenopalatine_VS_P2_P3 | 1 | 0 | 0 | 1 | 1 | 1 | 1 | 1 |
| CORO1C | -0.138 | P2_P3_P4_Lumbar_Coeliac_Stellate_Sphenopalatine_VS_P1 | 0 | 1 | 1 | 1 | 1 | 1 | 1 | 1 |
| TRIAP1 | -0.138 | P2_P3_P4_Lumbar_Coeliac_Stellate_Sphenopalatine_VS_P1 | 0 | 1 | 1 | 1 | 1 | 1 | 1 | 1 |
| CNPY2 | -0.139 | P3_P4_Lumbar_Stellate_VS_P1_P2_Coeliac_Sphenopalatine | 0 | 0 | 1 | 1 | 1 | 0 | 1 | 0 |
| UGP2 | -0.139 | P1_P2_P3_P4_Lumbar_Stellate_VS_Coeliac_Sphenopalatine | 1 | 1 | 1 | 1 | 1 | 0 | 1 | 0 |
| UFD1 | -0.139 | P1_P2_P3_P4_Lumbar_Stellate_VS_Coeliac_Sphenopalatine | 1 | 1 | 1 | 1 | 1 | 0 | 1 | 0 |
| LAPTM4A | -0.139 | P3_P4_Stellate_Sphenopalatine_VS_P1_P2_Lumbar_Coeliac | 0 | 0 | 1 | 1 | 0 | 0 | 1 | 1 |
| TCEAL8 | -0.139 | P3_P4_Stellate_Sphenopalatine_VS_P1_P2_Lumbar_Coeliac | 0 | 0 | 1 | 1 | 0 | 0 | 1 | 1 |
| EMC10 | -0.139 | P1_P3_P4_Stellate_Sphenopalatine_VS_P2_Lumbar_Coeliac | 1 | 0 | 1 | 1 | 0 | 0 | 1 | 1 |
| SPCS1 | -0.139 | P1_P3_P4_Stellate_Sphenopalatine_VS_P2_Lumbar_Coeliac | 1 | 0 | 1 | 1 | 0 | 0 | 1 | 1 |
| SRP9 | -0.139 | P2_P3_P4_Stellate_Sphenopalatine_VS_P1_Lumbar_Coeliac | 0 | 1 | 1 | 1 | 0 | 0 | 1 | 1 |
| TMEM256 | -0.139 | P2_P3_P4_Stellate_Sphenopalatine_VS_P1_Lumbar_Coeliac | 0 | 1 | 1 | 1 | 0 | 0 | 1 | 1 |
| GABBR1 | -0.139 | P2_P3_P4_Stellate_Sphenopalatine_VS_P1_Lumbar_Coeliac | 0 | 1 | 1 | 1 | 0 | 0 | 1 | 1 |
| CUTA | -0.139 | P1_P3_Lumbar_Stellate_Sphenopalatine_VS_P2_P4_Coeliac | 1 | 0 | 1 | 0 | 1 | 0 | 1 | 1 |
| CLCN6 | -0.139 | P1_P2_P4_Lumbar_Stellate_Sphenopalatine_VS_P3_Coeliac | 1 | 1 | 0 | 1 | 1 | 0 | 1 | 1 |
| KLC2 | -0.139 | P1_P2_P4_Lumbar_Stellate_Sphenopalatine_VS_P3_Coeliac | 1 | 1 | 0 | 1 | 1 | 0 | 1 | 1 |
| UBE2V2 | -0.139 | P3_P4_Lumbar_Stellate_Sphenopalatine_VS_P1_P2_Coeliac | 0 | 0 | 1 | 1 | 1 | 0 | 1 | 1 |
| SCG5 | -0.139 | P1_P3_P4_Lumbar_Stellate_Sphenopalatine_VS_P2_Coeliac | 1 | 0 | 1 | 1 | 1 | 0 | 1 | 1 |
| WDR83OS | -0.139 | P1_P3_P4_Lumbar_Stellate_Sphenopalatine_VS_P2_Coeliac | 1 | 0 | 1 | 1 | 1 | 0 | 1 | 1 |
| IP6K2 | -0.139 | P1_P3_P4_Lumbar_Stellate_Sphenopalatine_VS_P2_Coeliac | 1 | 0 | 1 | 1 | 1 | 0 | 1 | 1 |
| PSME2 | -0.139 | P1_P3_P4_Lumbar_Stellate_Sphenopalatine_VS_P2_Coeliac | 1 | 0 | 1 | 1 | 1 | 0 | 1 | 1 |
| SAP18 | -0.139 | P1_P3_P4_Lumbar_Stellate_Sphenopalatine_VS_P2_Coeliac | 1 | 0 | 1 | 1 | 1 | 0 | 1 | 1 |
| RAD23B | -0.139 | P2_P3_P4_Lumbar_Stellate_Sphenopalatine_VS_P1_Coeliac | 0 | 1 | 1 | 1 | 1 | 0 | 1 | 1 |
| EGLN2 | -0.139 | P2_P3_P4_Lumbar_Stellate_Sphenopalatine_VS_P1_Coeliac | 0 | 1 | 1 | 1 | 1 | 0 | 1 | 1 |
| NDRG4 | -0.139 | P2_P3_P4_Lumbar_Stellate_Sphenopalatine_VS_P1_Coeliac | 0 | 1 | 1 | 1 | 1 | 0 | 1 | 1 |
| VAT1L | -0.139 | P1_P2_P3_P4_Coeliac_Stellate_Sphenopalatine_VS_Lumbar | 1 | 1 | 1 | 1 | 0 | 1 | 1 | 1 |

| | | | | | | | | | | |
|---|---|---|---|---|---|---|---|---|---|---|
| TIPIN | -0.139 | P1_P2_P3_P4_Coeliac_Stellate_Sphenopalatine_VS_Lumbar | 1 | 1 | 1 | 1 | 0 | 1 | 1 | 1 |
| TIA1 | -0.139 | P3_Lumbar_Coeliac_Stellate_Sphenopalatine_VS_P1_P2_P4 | 0 | 0 | 1 | 0 | 1 | 1 | 1 | 1 |
| ARID2 | -0.139 | P2_P4_Lumbar_Coeliac_Stellate_Sphenopalatine_VS_P1_P3 | 0 | 1 | 0 | 1 | 1 | 1 | 1 | 1 |
| SLC25A27 | -0.139 | P2_P3_P4_Lumbar_Coeliac_Stellate_Sphenopalatine_VS_P1 | 0 | 1 | 1 | 1 | 1 | 1 | 1 | 1 |
| ZFP787 | -0.14 | P1_P2_P3_P4_Stellate_VS_Lumbar_Coeliac_Sphenopalatine | 1 | 1 | 1 | 1 | 0 | 0 | 1 | 0 |
| ANXA5 | -0.14 | P3_P4_Lumbar_Stellate_VS_P1_P2_Coeliac_Sphenopalatine | 0 | 0 | 1 | 1 | 1 | 0 | 1 | 0 |
| H2-KE6 | -0.14 | P1_P3_P4_Lumbar_Stellate_VS_P2_Coeliac_Sphenopalatine | 1 | 0 | 1 | 1 | 1 | 0 | 1 | 0 |
| PKIG | -0.14 | P2_P3_P4_Stellate_Sphenopalatine_VS_P1_Lumbar_Coeliac | 0 | 1 | 1 | 1 | 0 | 0 | 1 | 1 |
| TRMT112 | -0.14 | P2_P3_P4_Stellate_Sphenopalatine_VS_P1_Lumbar_Coeliac | 0 | 1 | 1 | 1 | 0 | 0 | 1 | 1 |
| EIF2S3X | -0.14 | P2_P3_P4_Stellate_Sphenopalatine_VS_P1_Lumbar_Coeliac | 0 | 1 | 1 | 1 | 0 | 0 | 1 | 1 |
| AACS | -0.14 | P1_P2_P4_Lumbar_Stellate_Sphenopalatine_VS_P3_Coeliac | 1 | 1 | 0 | 1 | 1 | 0 | 1 | 1 |
| EIF4EBP1 | -0.14 | P1_P2_P4_Lumbar_Stellate_Sphenopalatine_VS_P3_Coeliac | 1 | 1 | 0 | 1 | 1 | 0 | 1 | 1 |
| CHMP2B | -0.14 | P1_P2_P4_Lumbar_Stellate_Sphenopalatine_VS_P3_Coeliac | 1 | 1 | 0 | 1 | 1 | 0 | 1 | 1 |
| EPB41L3 | -0.14 | P1_P2_P4_Lumbar_Stellate_Sphenopalatine_VS_P3_Coeliac | 1 | 1 | 0 | 1 | 1 | 0 | 1 | 1 |
| ZFP637 | -0.14 | P3_P4_Lumbar_Stellate_Sphenopalatine_VS_P1_P2_Coeliac | 0 | 0 | 1 | 1 | 1 | 0 | 1 | 1 |
| CCDC107 | -0.14 | P1_P3_P4_Lumbar_Stellate_Sphenopalatine_VS_P2_Coeliac | 1 | 0 | 1 | 1 | 1 | 0 | 1 | 1 |
| IFT22 | -0.14 | P1_P3_P4_Lumbar_Stellate_Sphenopalatine_VS_P2_Coeliac | 1 | 0 | 1 | 1 | 1 | 0 | 1 | 1 |
| UBE2D1 | -0.14 | P1_P3_P4_Lumbar_Stellate_Sphenopalatine_VS_P2_Coeliac | 1 | 0 | 1 | 1 | 1 | 0 | 1 | 1 |
| NDUFS4 | -0.14 | P1_P3_P4_Lumbar_Stellate_Sphenopalatine_VS_P2_Coeliac | 1 | 0 | 1 | 1 | 1 | 0 | 1 | 1 |
| MRPL40 | -0.14 | P1_P3_P4_Lumbar_Stellate_Sphenopalatine_VS_P2_Coeliac | 1 | 0 | 1 | 1 | 1 | 0 | 1 | 1 |
| SLC25A39 | -0.14 | P2_P3_P4_Lumbar_Stellate_Sphenopalatine_VS_P1_Coeliac | 0 | 1 | 1 | 1 | 1 | 0 | 1 | 1 |
| TLX2 | -0.14 | P1_P2_P3_P4_Coeliac_Stellate_Sphenopalatine_VS_Lumbar | 1 | 1 | 1 | 1 | 0 | 1 | 1 | 1 |
| TMEM254A | -0.14 | P1_P2_P3_Lumbar_Coeliac_Stellate_Sphenopalatine_VS_P4 | 1 | 1 | 1 | 0 | 1 | 1 | 1 | 1 |
| PCLO | -0.14 | P2_P4_Lumbar_Coeliac_Stellate_Sphenopalatine_VS_P1_P3 | 0 | 1 | 0 | 1 | 1 | 1 | 1 | 1 |
| TMEM208 | -0.14 | P2_P3_P4_Lumbar_Coeliac_Stellate_Sphenopalatine_VS_P1 | 0 | 1 | 1 | 1 | 1 | 1 | 1 | 1 |
| CMIP | -0.14 | P2_P3_P4_Lumbar_Coeliac_Stellate_Sphenopalatine_VS_P1 | 0 | 1 | 1 | 1 | 1 | 1 | 1 | 1 |
| GPATCH8 | -0.14 | P2_P3_P4_Lumbar_Coeliac_Stellate_Sphenopalatine_VS_P1 | 0 | 1 | 1 | 1 | 1 | 1 | 1 | 1 |
| WAC | -0.14 | P2_P3_P4_Lumbar_Coeliac_Stellate_Sphenopalatine_VS_P1 | 0 | 1 | 1 | 1 | 1 | 1 | 1 | 1 |
| SRSF7 | -0.141 | P3_Lumbar_Stellate_VS_P1_P2_P4_Coeliac_Sphenopalatine | 0 | 0 | 1 | 0 | 1 | 0 | 1 | 0 |
| PSMD10 | -0.141 | P1_P3_P4_Lumbar_Stellate_VS_P2_Coeliac_Sphenopalatine | 1 | 0 | 1 | 1 | 1 | 0 | 1 | 0 |

| | | | | | | | | | | |
|---|---|---|---|---|---|---|---|---|---|---|
| GNG10 | -0.141 | P3_Stellate_Sphenopalatine_VS_P1_P2_P4_Lumbar_Coeliac | 0 | 0 | 1 | 0 | 0 | 0 | 1 | 1 |
| MRPL42 | -0.141 | P3_Stellate_Sphenopalatine_VS_P1_P2_P4_Lumbar_Coeliac | 0 | 0 | 1 | 0 | 0 | 0 | 1 | 1 |
| TRIM35 | -0.141 | P3_Stellate_Sphenopalatine_VS_P1_P2_P4_Lumbar_Coeliac | 0 | 0 | 1 | 0 | 0 | 0 | 1 | 1 |
| PCBP2 | -0.141 | P3_Stellate_Sphenopalatine_VS_P1_P2_P4_Lumbar_Coeliac | 0 | 0 | 1 | 0 | 0 | 0 | 1 | 1 |
| SNRPG | -0.141 | P2_P3_Stellate_Sphenopalatine_VS_P1_P4_Lumbar_Coeliac | 0 | 1 | 1 | 0 | 0 | 0 | 1 | 1 |
| MRPS21 | -0.141 | P3_P4_Stellate_Sphenopalatine_VS_P1_P2_Lumbar_Coeliac | 0 | 0 | 1 | 1 | 0 | 0 | 1 | 1 |
| DDA1 | -0.141 | P1_P3_P4_Stellate_Sphenopalatine_VS_P2_Lumbar_Coeliac | 1 | 0 | 1 | 1 | 0 | 0 | 1 | 1 |
| SMYD2 | -0.141 | P2_P3_P4_Stellate_Sphenopalatine_VS_P1_Lumbar_Coeliac | 0 | 1 | 1 | 1 | 0 | 0 | 1 | 1 |
| SARAF | -0.141 | P2_P3_P4_Stellate_Sphenopalatine_VS_P1_Lumbar_Coeliac | 0 | 1 | 1 | 1 | 0 | 0 | 1 | 1 |
| ZFP706 | -0.141 | P2_P3_P4_Stellate_Sphenopalatine_VS_P1_Lumbar_Coeliac | 0 | 1 | 1 | 1 | 0 | 0 | 1 | 1 |
| TMEM134 | -0.141 | P2_P3_P4_Stellate_Sphenopalatine_VS_P1_Lumbar_Coeliac | 0 | 1 | 1 | 1 | 0 | 0 | 1 | 1 |
| KCTD17 | -0.141 | P1_P2_P4_Lumbar_Stellate_Sphenopalatine_VS_P3_Coeliac | 1 | 1 | 0 | 1 | 1 | 0 | 1 | 1 |
| MED21 | -0.141 | P3_P4_Lumbar_Stellate_Sphenopalatine_VS_P1_P2_Coeliac | 0 | 0 | 1 | 1 | 1 | 0 | 1 | 1 |
| SVBP | -0.141 | P2_P3_P4_Lumbar_Stellate_Sphenopalatine_VS_P1_Coeliac | 0 | 1 | 1 | 1 | 1 | 0 | 1 | 1 |
| FKBP8 | -0.141 | P2_P3_P4_Lumbar_Stellate_Sphenopalatine_VS_P1_Coeliac | 0 | 1 | 1 | 1 | 1 | 0 | 1 | 1 |
| NICN1 | -0.141 | P2_P3_P4_Lumbar_Stellate_Sphenopalatine_VS_P1_Coeliac | 0 | 1 | 1 | 1 | 1 | 0 | 1 | 1 |
| SPATA7 | -0.141 | P2_P3_P4_Lumbar_Stellate_Sphenopalatine_VS_P1_Coeliac | 0 | 1 | 1 | 1 | 1 | 0 | 1 | 1 |
| TSC22D1 | -0.141 | P2_P3_P4_Coeliac_Stellate_Sphenopalatine_VS_P1_Lumbar | 0 | 1 | 1 | 1 | 0 | 1 | 1 | 1 |
| UBA52 | -0.141 | P1_P2_P3_P4_Coeliac_Stellate_Sphenopalatine_VS_Lumbar | 1 | 1 | 1 | 1 | 0 | 1 | 1 | 1 |
| ZCCHC14 | -0.141 | P1_P2_P3_Lumbar_Coeliac_Stellate_Sphenopalatine_VS_P4 | 1 | 1 | 1 | 0 | 1 | 1 | 1 | 1 |
| SCN3B | -0.141 | P1_P2_P3_Lumbar_Coeliac_Stellate_Sphenopalatine_VS_P4 | 1 | 1 | 1 | 0 | 1 | 1 | 1 | 1 |
| PHF20 | -0.141 | P2_P4_Lumbar_Coeliac_Stellate_Sphenopalatine_VS_P1_P3 | 0 | 1 | 0 | 1 | 1 | 1 | 1 | 1 |
| SV2A | -0.141 | P2_P3_P4_Lumbar_Coeliac_Stellate_Sphenopalatine_VS_P1 | 0 | 1 | 1 | 1 | 1 | 1 | 1 | 1 |
| NPTXR | -0.141 | P2_P3_P4_Lumbar_Coeliac_Stellate_Sphenopalatine_VS_P1 | 0 | 1 | 1 | 1 | 1 | 1 | 1 | 1 |
| PSMD14 | -0.142 | P3_P4_Lumbar_Stellate_VS_P1_P2_Coeliac_Sphenopalatine | 0 | 0 | 1 | 1 | 1 | 0 | 1 | 0 |
| CDK4 | -0.142 | P3_P4_Lumbar_Stellate_VS_P1_P2_Coeliac_Sphenopalatine | 0 | 0 | 1 | 1 | 1 | 0 | 1 | 0 |
| PHB | -0.142 | P1_P3_P4_Lumbar_Stellate_VS_P2_Coeliac_Sphenopalatine | 1 | 0 | 1 | 1 | 1 | 0 | 1 | 0 |
| LDLR | -0.142 | P1_P2_P3_P4_Lumbar_Stellate_VS_Coeliac_Sphenopalatine | 1 | 1 | 1 | 1 | 1 | 0 | 1 | 0 |
| IGSF3 | -0.142 | P2_Lumbar_Coeliac_Stellate_VS_P1_P3_P4_Sphenopalatine | 0 | 1 | 0 | 0 | 1 | 1 | 1 | 0 |
| EEF1E1 | -0.142 | P3_Stellate_Sphenopalatine_VS_P1_P2_P4_Lumbar_Coeliac | 0 | 0 | 1 | 0 | 0 | 0 | 1 | 1 |

| | | | | | | | | | | |
|---|---|---|---|---|---|---|---|---|---|---|
| EML5 | -0.142 | P2_P3_Stellate_Sphenopalatine_VS_P1_P4_Lumbar_Coeliac | 0 | 1 | 1 | 0 | 0 | 0 | 1 | 1 |
| TRA2B | -0.142 | P3_P4_Stellate_Sphenopalatine_VS_P1_P2_Lumbar_Coeliac | 0 | 0 | 1 | 1 | 0 | 0 | 1 | 1 |
| RTL8A | -0.142 | P3_P4_Stellate_Sphenopalatine_VS_P1_P2_Lumbar_Coeliac | 0 | 0 | 1 | 1 | 0 | 0 | 1 | 1 |
| ERGIC3 | -0.142 | P2_P3_P4_Stellate_Sphenopalatine_VS_P1_Lumbar_Coeliac | 0 | 1 | 1 | 1 | 0 | 0 | 1 | 1 |
| RAB7 | -0.142 | P2_P3_P4_Stellate_Sphenopalatine_VS_P1_Lumbar_Coeliac | 0 | 1 | 1 | 1 | 0 | 0 | 1 | 1 |
| MLF2 | -0.142 | P2_P3_P4_Stellate_Sphenopalatine_VS_P1_Lumbar_Coeliac | 0 | 1 | 1 | 1 | 0 | 0 | 1 | 1 |
| TTC9B | -0.142 | P2_P3_P4_Stellate_Sphenopalatine_VS_P1_Lumbar_Coeliac | 0 | 1 | 1 | 1 | 0 | 0 | 1 | 1 |
| CLIP3 | -0.142 | P2_P3_P4_Stellate_Sphenopalatine_VS_P1_Lumbar_Coeliac | 0 | 1 | 1 | 1 | 0 | 0 | 1 | 1 |
| PTS | -0.142 | P2_P3_P4_Stellate_Sphenopalatine_VS_P1_Lumbar_Coeliac | 0 | 1 | 1 | 1 | 0 | 0 | 1 | 1 |
| ARPC5L | -0.142 | P2_P3_P4_Lumbar_Stellate_Sphenopalatine_VS_P1_Coeliac | 0 | 1 | 1 | 1 | 1 | 0 | 1 | 1 |
| MAP7D1 | -0.142 | P2_P3_P4_Lumbar_Stellate_Sphenopalatine_VS_P1_Coeliac | 0 | 1 | 1 | 1 | 1 | 0 | 1 | 1 |
| MAT2A | -0.142 | P2_P3_P4_Lumbar_Stellate_Sphenopalatine_VS_P1_Coeliac | 0 | 1 | 1 | 1 | 1 | 0 | 1 | 1 |
| MIEN1 | -0.142 | P2_P3_P4_Lumbar_Stellate_Sphenopalatine_VS_P1_Coeliac | 0 | 1 | 1 | 1 | 1 | 0 | 1 | 1 |
| TMED9 | -0.142 | P2_P3_P4_Lumbar_Coeliac_Stellate_Sphenopalatine_VS_P1 | 0 | 1 | 1 | 1 | 1 | 1 | 1 | 1 |
| GM9844 | -0.143 | P3_Stellate_VS_P1_P2_P4_Lumbar_Coeliac_Sphenopalatine | 0 | 0 | 1 | 0 | 0 | 0 | 1 | 0 |
| ELOVL6 | -0.143 | P3_P4_Lumbar_Stellate_VS_P1_P2_Coeliac_Sphenopalatine | 0 | 0 | 1 | 1 | 1 | 0 | 1 | 0 |
| PRPF38A | -0.143 | P1_P3_P4_Lumbar_Stellate_VS_P2_Coeliac_Sphenopalatine | 1 | 0 | 1 | 1 | 1 | 0 | 1 | 0 |
| SLC25A17 | -0.143 | P2_P3_P4_Lumbar_Stellate_VS_P1_Coeliac_Sphenopalatine | 0 | 1 | 1 | 1 | 1 | 0 | 1 | 0 |
| SLBP | -0.143 | P1_P2_P3_P4_Lumbar_Stellate_VS_Coeliac_Sphenopalatine | 1 | 1 | 1 | 1 | 1 | 0 | 1 | 0 |
| DNAJC19 | -0.143 | P3_P4_Stellate_Sphenopalatine_VS_P1_P2_Lumbar_Coeliac | 0 | 0 | 1 | 1 | 0 | 0 | 1 | 1 |
| PRELID1 | -0.143 | P3_P4_Stellate_Sphenopalatine_VS_P1_P2_Lumbar_Coeliac | 0 | 0 | 1 | 1 | 0 | 0 | 1 | 1 |
| RAB11B | -0.143 | P3_P4_Stellate_Sphenopalatine_VS_P1_P2_Lumbar_Coeliac | 0 | 0 | 1 | 1 | 0 | 0 | 1 | 1 |
| UBE2J1 | -0.143 | P2_P3_P4_Stellate_Sphenopalatine_VS_P1_Lumbar_Coeliac | 0 | 1 | 1 | 1 | 0 | 0 | 1 | 1 |
| RTRAF | -0.143 | P2_P3_P4_Stellate_Sphenopalatine_VS_P1_Lumbar_Coeliac | 0 | 1 | 1 | 1 | 0 | 0 | 1 | 1 |
| B4GALT6 | -0.143 | P2_P3_P4_Stellate_Sphenopalatine_VS_P1_Lumbar_Coeliac | 0 | 1 | 1 | 1 | 0 | 0 | 1 | 1 |
| FAM171A1 | -0.143 | P1_P2_P4_Lumbar_Stellate_Sphenopalatine_VS_P3_Coeliac | 1 | 1 | 0 | 1 | 1 | 0 | 1 | 1 |
| VPS28 | -0.143 | P3_P4_Lumbar_Stellate_Sphenopalatine_VS_P1_P2_Coeliac | 0 | 0 | 1 | 1 | 1 | 0 | 1 | 1 |
| STK25 | -0.143 | P2_P3_P4_Lumbar_Stellate_Sphenopalatine_VS_P1_Coeliac | 0 | 1 | 1 | 1 | 1 | 0 | 1 | 1 |
| TMEM147 | -0.143 | P2_P3_P4_Lumbar_Stellate_Sphenopalatine_VS_P1_Coeliac | 0 | 1 | 1 | 1 | 1 | 0 | 1 | 1 |
| PHKG2 | -0.143 | P2_P3_P4_Lumbar_Stellate_Sphenopalatine_VS_P1_Coeliac | 0 | 1 | 1 | 1 | 1 | 0 | 1 | 1 |

| | | | | | | | | | | |
|---|---|---|---|---|---|---|---|---|---|---|
| EMC6 | -0.143 | P2_P3_P4_Lumbar_Stellate_Sphenopalatine_VS_P1_Coeliac | 0 | 1 | 1 | 1 | 1 | 0 | 1 | 1 |
| ARL2 | -0.143 | P2_P3_P4_Lumbar_Stellate_Sphenopalatine_VS_P1_Coeliac | 0 | 1 | 1 | 1 | 1 | 0 | 1 | 1 |
| GABARAPL1 | -0.143 | P1_P2_P3_P4_Coeliac_Stellate_Sphenopalatine_VS_Lumbar | 1 | 1 | 1 | 1 | 0 | 1 | 1 | 1 |
| PAICS | -0.143 | P2_P3_P4_Lumbar_Coeliac_Stellate_Sphenopalatine_VS_P1 | 0 | 1 | 1 | 1 | 1 | 1 | 1 | 1 |
| CHD4 | -0.143 | P2_P3_P4_Lumbar_Coeliac_Stellate_Sphenopalatine_VS_P1 | 0 | 1 | 1 | 1 | 1 | 1 | 1 | 1 |
| MICU3 | -0.143 | P2_P3_P4_Lumbar_Coeliac_Stellate_Sphenopalatine_VS_P1 | 0 | 1 | 1 | 1 | 1 | 1 | 1 | 1 |
| HNRNPC | -0.143 | P2_P3_P4_Lumbar_Coeliac_Stellate_Sphenopalatine_VS_P1 | 0 | 1 | 1 | 1 | 1 | 1 | 1 | 1 |
| SSR3 | -0.144 | P4_Lumbar_Stellate_VS_P1_P2_P3_Coeliac_Sphenopalatine | 0 | 0 | 0 | 1 | 1 | 0 | 1 | 0 |
| PHF23 | -0.144 | P4_Lumbar_Stellate_VS_P1_P2_P3_Coeliac_Sphenopalatine | 0 | 0 | 0 | 1 | 1 | 0 | 1 | 0 |
| HDAC2 | -0.144 | P3_P4_Lumbar_Stellate_VS_P1_P2_Coeliac_Sphenopalatine | 0 | 0 | 1 | 1 | 1 | 0 | 1 | 0 |
| TMEM126B | -0.144 | P1_P3_P4_Lumbar_Stellate_VS_P2_Coeliac_Sphenopalatine | 1 | 0 | 1 | 1 | 1 | 0 | 1 | 0 |
| ITM2C | -0.144 | P1_P2_P3_P4_Lumbar_Stellate_VS_Coeliac_Sphenopalatine | 1 | 1 | 1 | 1 | 1 | 0 | 1 | 0 |
| 9530059O14 | -0.144 | P1_P2_P3_P4_Lumbar_Stellate_VS_Coeliac_Sphenopalatine | 1 | 1 | 1 | 1 | 1 | 0 | 1 | 0 |
| SNX12 | -0.144 | P1_P2_P3_P4_Lumbar_Stellate_VS_Coeliac_Sphenopalatine | 1 | 1 | 1 | 1 | 1 | 0 | 1 | 0 |
| PRELID3B | -0.144 | P1_P2_P3_Stellate_Sphenopalatine_VS_P4_Lumbar_Coeliac | 1 | 1 | 1 | 0 | 0 | 0 | 1 | 1 |
| COX14 | -0.144 | P3_P4_Stellate_Sphenopalatine_VS_P1_P2_Lumbar_Coeliac | 0 | 0 | 1 | 1 | 0 | 0 | 1 | 1 |
| ACTR1A | -0.144 | P3_P4_Stellate_Sphenopalatine_VS_P1_P2_Lumbar_Coeliac | 0 | 0 | 1 | 1 | 0 | 0 | 1 | 1 |
| NDUFB6 | -0.144 | P1_P3_P4_Stellate_Sphenopalatine_VS_P2_Lumbar_Coeliac | 1 | 0 | 1 | 1 | 0 | 0 | 1 | 1 |
| POLR1D | -0.144 | P2_P3_P4_Stellate_Sphenopalatine_VS_P1_Lumbar_Coeliac | 0 | 1 | 1 | 1 | 0 | 0 | 1 | 1 |
| RP9 | -0.144 | P2_P3_P4_Stellate_Sphenopalatine_VS_P1_Lumbar_Coeliac | 0 | 1 | 1 | 1 | 0 | 0 | 1 | 1 |
| C1D | -0.144 | P2_P3_P4_Stellate_Sphenopalatine_VS_P1_Lumbar_Coeliac | 0 | 1 | 1 | 1 | 0 | 0 | 1 | 1 |
| SYNGR3 | -0.144 | P2_P3_P4_Stellate_Sphenopalatine_VS_P1_Lumbar_Coeliac | 0 | 1 | 1 | 1 | 0 | 0 | 1 | 1 |
| TUBB4A | -0.144 | P2_P3_P4_Stellate_Sphenopalatine_VS_P1_Lumbar_Coeliac | 0 | 1 | 1 | 1 | 0 | 0 | 1 | 1 |
| ARL6IP5 | -0.144 | P1_P2_P3_Lumbar_Stellate_Sphenopalatine_VS_P4_Coeliac | 1 | 1 | 1 | 0 | 1 | 0 | 1 | 1 |
| IDH1 | -0.144 | P2_P3_P4_Lumbar_Stellate_Sphenopalatine_VS_P1_Coeliac | 0 | 1 | 1 | 1 | 1 | 0 | 1 | 1 |
| CPE | -0.144 | P2_P3_P4_Lumbar_Stellate_Sphenopalatine_VS_P1_Coeliac | 0 | 1 | 1 | 1 | 1 | 0 | 1 | 1 |
| ABHD8 | -0.144 | P2_P3_P4_Lumbar_Stellate_Sphenopalatine_VS_P1_Coeliac | 0 | 1 | 1 | 1 | 1 | 0 | 1 | 1 |
| DLG4 | -0.144 | P2_P3_P4_Lumbar_Stellate_Sphenopalatine_VS_P1_Coeliac | 0 | 1 | 1 | 1 | 1 | 0 | 1 | 1 |
| SV2C | -0.144 | P2_P3_P4_Lumbar_Stellate_Sphenopalatine_VS_P1_Coeliac | 0 | 1 | 1 | 1 | 1 | 0 | 1 | 1 |
| ATXN7L3B | -0.144 | P1_P2_P3_P4_Coeliac_Stellate_Sphenopalatine_VS_Lumbar | 1 | 1 | 1 | 1 | 0 | 1 | 1 | 1 |

| | | | | | | | | | | |
|---|---|---|---|---|---|---|---|---|---|---|
| ZFP961 | -0.144 | P1_P2_P3_Lumbar_Coeliac_Stellate_Sphenopalatine_VS_P4 | 1 | 1 | 1 | 0 | 1 | 1 | 1 | 1 |
| CRIP1 | -0.144 | P1_P2_P3_Lumbar_Coeliac_Stellate_Sphenopalatine_VS_P4 | 1 | 1 | 1 | 0 | 1 | 1 | 1 | 1 |
| ACSL3 | -0.144 | P4_Lumbar_Coeliac_Stellate_Sphenopalatine_VS_P1_P2_P3 | 0 | 0 | 0 | 1 | 1 | 1 | 1 | 1 |
| STAU2 | -0.144 | P2_P3_P4_Lumbar_Coeliac_Stellate_Sphenopalatine_VS_P1 | 0 | 1 | 1 | 1 | 1 | 1 | 1 | 1 |
| FAM102B | -0.144 | P2_P3_P4_Lumbar_Coeliac_Stellate_Sphenopalatine_VS_P1 | 0 | 1 | 1 | 1 | 1 | 1 | 1 | 1 |
| ZFP664 | -0.144 | P2_P3_P4_Lumbar_Coeliac_Stellate_Sphenopalatine_VS_P1 | 0 | 1 | 1 | 1 | 1 | 1 | 1 | 1 |
| CASD1 | -0.144 | P2_P3_P4_Lumbar_Coeliac_Stellate_Sphenopalatine_VS_P1 | 0 | 1 | 1 | 1 | 1 | 1 | 1 | 1 |
| SYT5 | -0.144 | P2_P3_P4_Lumbar_Coeliac_Stellate_Sphenopalatine_VS_P1 | 0 | 1 | 1 | 1 | 1 | 1 | 1 | 1 |
| PPP2R5C | -0.144 | P2_P3_P4_Lumbar_Coeliac_Stellate_Sphenopalatine_VS_P1 | 0 | 1 | 1 | 1 | 1 | 1 | 1 | 1 |
| HMGN1 | -0.144 | P2_P3_P4_Lumbar_Coeliac_Stellate_Sphenopalatine_VS_P1 | 0 | 1 | 1 | 1 | 1 | 1 | 1 | 1 |
| JAGN1 | -0.145 | P3_P4_Lumbar_Stellate_VS_P1_P2_Coeliac_Sphenopalatine | 0 | 0 | 1 | 1 | 1 | 0 | 1 | 0 |
| NDUFA9 | -0.145 | P1_P3_P4_Lumbar_Stellate_VS_P2_Coeliac_Sphenopalatine | 1 | 0 | 1 | 1 | 1 | 0 | 1 | 0 |
| GNPNAT1 | -0.145 | P1_P3_P4_Lumbar_Stellate_VS_P2_Coeliac_Sphenopalatine | 1 | 0 | 1 | 1 | 1 | 0 | 1 | 0 |
| LHFPL5 | -0.145 | P1_P2_P3_P4_Lumbar_Stellate_VS_Coeliac_Sphenopalatine | 1 | 1 | 1 | 1 | 1 | 0 | 1 | 0 |
| HSD17B10 | -0.145 | P1_P2_P3_P4_Lumbar_Stellate_VS_Coeliac_Sphenopalatine | 1 | 1 | 1 | 1 | 1 | 0 | 1 | 0 |
| ARHGEF28 | -0.145 | P2_P3_P4_Coeliac_Stellate_VS_P1_Lumbar_Sphenopalatine | 0 | 1 | 1 | 1 | 0 | 1 | 1 | 0 |
| SPTSSA | -0.145 | P3_Stellate_Sphenopalatine_VS_P1_P2_P4_Lumbar_Coeliac | 0 | 0 | 1 | 0 | 0 | 0 | 1 | 1 |
| NOP10 | -0.145 | P3_P4_Stellate_Sphenopalatine_VS_P1_P2_Lumbar_Coeliac | 0 | 0 | 1 | 1 | 0 | 0 | 1 | 1 |
| FSCN1 | -0.145 | P3_P4_Stellate_Sphenopalatine_VS_P1_P2_Lumbar_Coeliac | 0 | 0 | 1 | 1 | 0 | 0 | 1 | 1 |
| NDUFA3 | -0.145 | P3_P4_Stellate_Sphenopalatine_VS_P1_P2_Lumbar_Coeliac | 0 | 0 | 1 | 1 | 0 | 0 | 1 | 1 |
| MRPL54 | -0.145 | P3_P4_Stellate_Sphenopalatine_VS_P1_P2_Lumbar_Coeliac | 0 | 0 | 1 | 1 | 0 | 0 | 1 | 1 |
| SARNP | -0.145 | P3_P4_Stellate_Sphenopalatine_VS_P1_P2_Lumbar_Coeliac | 0 | 0 | 1 | 1 | 0 | 0 | 1 | 1 |
| EIF3E | -0.145 | P3_P4_Stellate_Sphenopalatine_VS_P1_P2_Lumbar_Coeliac | 0 | 0 | 1 | 1 | 0 | 0 | 1 | 1 |
| CRIPT | -0.145 | P3_P4_Stellate_Sphenopalatine_VS_P1_P2_Lumbar_Coeliac | 0 | 0 | 1 | 1 | 0 | 0 | 1 | 1 |
| RHEB | -0.145 | P1_P3_P4_Stellate_Sphenopalatine_VS_P2_Lumbar_Coeliac | 1 | 0 | 1 | 1 | 0 | 0 | 1 | 1 |
| MRPS36 | -0.145 | P1_P3_P4_Stellate_Sphenopalatine_VS_P2_Lumbar_Coeliac | 1 | 0 | 1 | 1 | 0 | 0 | 1 | 1 |
| NDUFA6 | -0.145 | P2_P3_P4_Stellate_Sphenopalatine_VS_P1_Lumbar_Coeliac | 0 | 1 | 1 | 1 | 0 | 0 | 1 | 1 |
| ARVCF | -0.145 | P1_P2_P4_Lumbar_Stellate_Sphenopalatine_VS_P3_Coeliac | 1 | 1 | 0 | 1 | 1 | 0 | 1 | 1 |
| SMIM10L1 | -0.145 | P3_P4_Lumbar_Stellate_Sphenopalatine_VS_P1_P2_Coeliac | 0 | 0 | 1 | 1 | 1 | 0 | 1 | 1 |
| DCTN6 | -0.145 | P3_P4_Lumbar_Stellate_Sphenopalatine_VS_P1_P2_Coeliac | 0 | 0 | 1 | 1 | 1 | 0 | 1 | 1 |

| | | | | | | | | | | |
|---|---|---|---|---|---|---|---|---|---|---|
| TIMM10B | -0.145 | P2_P3_P4_Lumbar_Stellate_Sphenopalatine_VS_P1_Coeliac | 0 | 1 | 1 | 1 | 1 | 0 | 1 | 1 |
| IFT20 | -0.145 | P2_P3_P4_Lumbar_Stellate_Sphenopalatine_VS_P1_Coeliac | 0 | 1 | 1 | 1 | 1 | 0 | 1 | 1 |
| RANGAP1 | -0.145 | P2_P3_P4_Lumbar_Stellate_Sphenopalatine_VS_P1_Coeliac | 0 | 1 | 1 | 1 | 1 | 0 | 1 | 1 |
| SAMD14 | -0.145 | P1_P2_P3_P4_Coeliac_Stellate_Sphenopalatine_VS_Lumbar | 1 | 1 | 1 | 1 | 0 | 1 | 1 | 1 |
| D430019H16 | -0.145 | P1_P2_P3_Lumbar_Coeliac_Stellate_Sphenopalatine_VS_P4 | 1 | 1 | 1 | 0 | 1 | 1 | 1 | 1 |
| CCDC50 | -0.145 | P1_P2_P3_Lumbar_Coeliac_Stellate_Sphenopalatine_VS_P4 | 1 | 1 | 1 | 0 | 1 | 1 | 1 | 1 |
| ZFHX3 | -0.146 | P3_P4_Stellate_VS_P1_P2_Lumbar_Coeliac_Sphenopalatine | 0 | 0 | 1 | 1 | 0 | 0 | 1 | 0 |
| VKORC1 | -0.146 | P1_P2_P3_P4_Stellate_VS_Lumbar_Coeliac_Sphenopalatine | 1 | 1 | 1 | 1 | 0 | 0 | 1 | 0 |
| ERGIC1 | -0.146 | P4_Lumbar_Stellate_VS_P1_P2_P3_Coeliac_Sphenopalatine | 0 | 0 | 0 | 1 | 1 | 0 | 1 | 0 |
| FXN | -0.146 | P2_P3_P4_Lumbar_Stellate_VS_P1_Coeliac_Sphenopalatine | 0 | 1 | 1 | 1 | 1 | 0 | 1 | 0 |
| GTF2A2 | -0.146 | P3_P4_Stellate_Sphenopalatine_VS_P1_P2_Lumbar_Coeliac | 0 | 0 | 1 | 1 | 0 | 0 | 1 | 1 |
| CIRBP | -0.146 | P3_P4_Stellate_Sphenopalatine_VS_P1_P2_Lumbar_Coeliac | 0 | 0 | 1 | 1 | 0 | 0 | 1 | 1 |
| ANKRA2 | -0.146 | P3_P4_Stellate_Sphenopalatine_VS_P1_P2_Lumbar_Coeliac | 0 | 0 | 1 | 1 | 0 | 0 | 1 | 1 |
| CD200 | -0.146 | P3_P4_Stellate_Sphenopalatine_VS_P1_P2_Lumbar_Coeliac | 0 | 0 | 1 | 1 | 0 | 0 | 1 | 1 |
| CTNNBIP1 | -0.146 | P2_P3_P4_Stellate_Sphenopalatine_VS_P1_Lumbar_Coeliac | 0 | 1 | 1 | 1 | 0 | 0 | 1 | 1 |
| ATP6V0E2 | -0.146 | P2_P3_P4_Stellate_Sphenopalatine_VS_P1_Lumbar_Coeliac | 0 | 1 | 1 | 1 | 0 | 0 | 1 | 1 |
| ARPP19 | -0.146 | P2_P3_P4_Stellate_Sphenopalatine_VS_P1_Lumbar_Coeliac | 0 | 1 | 1 | 1 | 0 | 0 | 1 | 1 |
| PCBD1 | -0.146 | P2_P3_P4_Stellate_Sphenopalatine_VS_P1_Lumbar_Coeliac | 0 | 1 | 1 | 1 | 0 | 0 | 1 | 1 |
| OGDH | -0.146 | P2_P3_P4_Stellate_Sphenopalatine_VS_P1_Lumbar_Coeliac | 0 | 1 | 1 | 1 | 0 | 0 | 1 | 1 |
| FAM104A | -0.146 | P2_P3_P4_Stellate_Sphenopalatine_VS_P1_Lumbar_Coeliac | 0 | 1 | 1 | 1 | 0 | 0 | 1 | 1 |
| CEP170B | -0.146 | P2_P3_P4_Stellate_Sphenopalatine_VS_P1_Lumbar_Coeliac | 0 | 1 | 1 | 1 | 0 | 0 | 1 | 1 |
| NBDY | -0.146 | P2_P3_P4_Stellate_Sphenopalatine_VS_P1_Lumbar_Coeliac | 0 | 1 | 1 | 1 | 0 | 0 | 1 | 1 |
| PTRHD1 | -0.146 | P3_P4_Lumbar_Stellate_Sphenopalatine_VS_P1_P2_Coeliac | 0 | 0 | 1 | 1 | 1 | 0 | 1 | 1 |
| MRPS26 | -0.146 | P2_P3_P4_Lumbar_Stellate_Sphenopalatine_VS_P1_Coeliac | 0 | 1 | 1 | 1 | 1 | 0 | 1 | 1 |
| SNRNP70 | -0.146 | P2_P3_P4_Lumbar_Stellate_Sphenopalatine_VS_P1_Coeliac | 0 | 1 | 1 | 1 | 1 | 0 | 1 | 1 |
| NUDT21 | -0.146 | P2_P3_P4_Lumbar_Stellate_Sphenopalatine_VS_P1_Coeliac | 0 | 1 | 1 | 1 | 1 | 0 | 1 | 1 |
| PRMT2 | -0.146 | P2_P3_P4_Lumbar_Stellate_Sphenopalatine_VS_P1_Coeliac | 0 | 1 | 1 | 1 | 1 | 0 | 1 | 1 |
| GATD3A | -0.146 | P2_P3_P4_Lumbar_Stellate_Sphenopalatine_VS_P1_Coeliac | 0 | 1 | 1 | 1 | 1 | 0 | 1 | 1 |
| MEG3 | -0.146 | P2_P3_P4_Lumbar_Stellate_Sphenopalatine_VS_P1_Coeliac | 0 | 1 | 1 | 1 | 1 | 0 | 1 | 1 |
| PDHB | -0.146 | P2_P3_P4_Lumbar_Stellate_Sphenopalatine_VS_P1_Coeliac | 0 | 1 | 1 | 1 | 1 | 0 | 1 | 1 |

| | | | | | | | | | | |
|---|---|---|---|---|---|---|---|---|---|---|
| FAM162A | -0.146 | P2_P3_P4_Lumbar_Stellate_Sphenopalatine_VS_P1_Coeliac | 0 | 1 | 1 | 1 | 1 | 0 | 1 | 1 |
| COPS9 | -0.146 | P1_P2_P3_P4_Coeliac_Stellate_Sphenopalatine_VS_Lumbar | 1 | 1 | 1 | 1 | 0 | 1 | 1 | 1 |
| NAPB | -0.146 | P1_P2_P3_P4_Coeliac_Stellate_Sphenopalatine_VS_Lumbar | 1 | 1 | 1 | 1 | 0 | 1 | 1 | 1 |
| DHX9 | -0.146 | P2_P4_Lumbar_Coeliac_Stellate_Sphenopalatine_VS_P1_P3 | 0 | 1 | 0 | 1 | 1 | 1 | 1 | 1 |
| BPTF | -0.146 | P2_P4_Lumbar_Coeliac_Stellate_Sphenopalatine_VS_P1_P3 | 0 | 1 | 0 | 1 | 1 | 1 | 1 | 1 |
| GAS5 | -0.146 | P2_P3_P4_Lumbar_Coeliac_Stellate_Sphenopalatine_VS_P1 | 0 | 1 | 1 | 1 | 1 | 1 | 1 | 1 |
| PDIA3 | -0.146 | P2_P3_P4_Lumbar_Coeliac_Stellate_Sphenopalatine_VS_P1 | 0 | 1 | 1 | 1 | 1 | 1 | 1 | 1 |
| PODXL2 | -0.146 | P2_P3_P4_Lumbar_Coeliac_Stellate_Sphenopalatine_VS_P1 | 0 | 1 | 1 | 1 | 1 | 1 | 1 | 1 |
| ARFGEF3 | -0.146 | P2_P3_P4_Lumbar_Coeliac_Stellate_Sphenopalatine_VS_P1 | 0 | 1 | 1 | 1 | 1 | 1 | 1 | 1 |
| ECI2 | -0.146 | P2_P3_P4_Lumbar_Coeliac_Stellate_Sphenopalatine_VS_P1 | 0 | 1 | 1 | 1 | 1 | 1 | 1 | 1 |
| ANKRD12 | -0.146 | P2_P3_P4_Lumbar_Coeliac_Stellate_Sphenopalatine_VS_P1 | 0 | 1 | 1 | 1 | 1 | 1 | 1 | 1 |
| APBB2 | -0.147 | P1_P3_P4_Lumbar_Stellate_VS_P2_Coeliac_Sphenopalatine | 1 | 0 | 1 | 1 | 1 | 0 | 1 | 0 |
| RHBDD2 | -0.147 | P1_P3_P4_Lumbar_Stellate_VS_P2_Coeliac_Sphenopalatine | 1 | 0 | 1 | 1 | 1 | 0 | 1 | 0 |
| MALSU1 | -0.147 | P1_P3_P4_Lumbar_Stellate_VS_P2_Coeliac_Sphenopalatine | 1 | 0 | 1 | 1 | 1 | 0 | 1 | 0 |
| IDH3G | -0.147 | P1_P3_P4_Lumbar_Stellate_VS_P2_Coeliac_Sphenopalatine | 1 | 0 | 1 | 1 | 1 | 0 | 1 | 0 |
| IRF2BPL | -0.147 | P2_P3_P4_Lumbar_Stellate_VS_P1_Coeliac_Sphenopalatine | 0 | 1 | 1 | 1 | 1 | 0 | 1 | 0 |
| GM2000 | -0.147 | P1_P2_P3_P4_Lumbar_Stellate_VS_Coeliac_Sphenopalatine | 1 | 1 | 1 | 1 | 1 | 0 | 1 | 0 |
| AP1S1 | -0.147 | P3_Stellate_Sphenopalatine_VS_P1_P2_P4_Lumbar_Coeliac | 0 | 0 | 1 | 0 | 0 | 0 | 1 | 1 |
| ZFP428 | -0.147 | P3_Stellate_Sphenopalatine_VS_P1_P2_P4_Lumbar_Coeliac | 0 | 0 | 1 | 0 | 0 | 0 | 1 | 1 |
| PABPC1 | -0.147 | P3_Stellate_Sphenopalatine_VS_P1_P2_P4_Lumbar_Coeliac | 0 | 0 | 1 | 0 | 0 | 0 | 1 | 1 |
| MRPL30 | -0.147 | P1_P3_P4_Stellate_Sphenopalatine_VS_P2_Lumbar_Coeliac | 1 | 0 | 1 | 1 | 0 | 0 | 1 | 1 |
| DNLZ | -0.147 | P1_P3_P4_Stellate_Sphenopalatine_VS_P2_Lumbar_Coeliac | 1 | 0 | 1 | 1 | 0 | 0 | 1 | 1 |
| TMEM251 | -0.147 | P1_P3_P4_Stellate_Sphenopalatine_VS_P2_Lumbar_Coeliac | 1 | 0 | 1 | 1 | 0 | 0 | 1 | 1 |
| SLC25A5 | -0.147 | P1_P3_P4_Stellate_Sphenopalatine_VS_P2_Lumbar_Coeliac | 1 | 0 | 1 | 1 | 0 | 0 | 1 | 1 |
| RBMS1 | -0.147 | P2_P3_P4_Stellate_Sphenopalatine_VS_P1_Lumbar_Coeliac | 0 | 1 | 1 | 1 | 0 | 0 | 1 | 1 |
| RBM39 | -0.147 | P2_P3_P4_Stellate_Sphenopalatine_VS_P1_Lumbar_Coeliac | 0 | 1 | 1 | 1 | 0 | 0 | 1 | 1 |
| TMEM160 | -0.147 | P1_P2_P3_Lumbar_Stellate_Sphenopalatine_VS_P4_Coeliac | 1 | 1 | 1 | 0 | 1 | 0 | 1 | 1 |
| CCDC12 | -0.147 | P4_Lumbar_Stellate_Sphenopalatine_VS_P1_P2_P3_Coeliac | 0 | 0 | 0 | 1 | 1 | 0 | 1 | 1 |
| UQCC2 | -0.147 | P3_P4_Lumbar_Stellate_Sphenopalatine_VS_P1_P2_Coeliac | 0 | 0 | 1 | 1 | 1 | 0 | 1 | 1 |
| SNRPC | -0.147 | P3_P4_Lumbar_Stellate_Sphenopalatine_VS_P1_P2_Coeliac | 0 | 0 | 1 | 1 | 1 | 0 | 1 | 1 |

| | | | | | | | | | | |
|---|---|---|---|---|---|---|---|---|---|---|
| CCT4 | -0.147 | P2_P3_P4_Lumbar_Stellate_Sphenopalatine_VS_P1_Coeliac | 0 | 1 | 1 | 1 | 1 | 0 | 1 | 1 |
| ISCA2 | -0.147 | P2_P3_P4_Lumbar_Stellate_Sphenopalatine_VS_P1_Coeliac | 0 | 1 | 1 | 1 | 1 | 0 | 1 | 1 |
| SETD3 | -0.147 | P2_P3_P4_Lumbar_Stellate_Sphenopalatine_VS_P1_Coeliac | 0 | 1 | 1 | 1 | 1 | 0 | 1 | 1 |
| DAD1 | -0.147 | P2_P3_P4_Lumbar_Stellate_Sphenopalatine_VS_P1_Coeliac | 0 | 1 | 1 | 1 | 1 | 0 | 1 | 1 |
| SPRYD7 | -0.147 | P2_P3_P4_Lumbar_Stellate_Sphenopalatine_VS_P1_Coeliac | 0 | 1 | 1 | 1 | 1 | 0 | 1 | 1 |
| SLC25A1 | -0.147 | P2_P3_P4_Lumbar_Stellate_Sphenopalatine_VS_P1_Coeliac | 0 | 1 | 1 | 1 | 1 | 0 | 1 | 1 |
| CTSB | -0.147 | P1_P2_P3_P4_Coeliac_Stellate_Sphenopalatine_VS_Lumbar | 1 | 1 | 1 | 1 | 0 | 1 | 1 | 1 |
| 4932438A13I | -0.147 | P2_P4_Lumbar_Coeliac_Stellate_Sphenopalatine_VS_P1_P3 | 0 | 1 | 0 | 1 | 1 | 1 | 1 | 1 |
| SLC7A14 | -0.147 | P2_P3_P4_Lumbar_Coeliac_Stellate_Sphenopalatine_VS_P1 | 0 | 1 | 1 | 1 | 1 | 1 | 1 | 1 |
| ARID1A | -0.147 | P2_P3_P4_Lumbar_Coeliac_Stellate_Sphenopalatine_VS_P1 | 0 | 1 | 1 | 1 | 1 | 1 | 1 | 1 |
| CSNK1E | -0.147 | P2_P3_P4_Lumbar_Coeliac_Stellate_Sphenopalatine_VS_P1 | 0 | 1 | 1 | 1 | 1 | 1 | 1 | 1 |
| PDCL3 | -0.148 | P1_P3_P4_Lumbar_Stellate_VS_P2_Coeliac_Sphenopalatine | 1 | 0 | 1 | 1 | 1 | 0 | 1 | 0 |
| TMEM185A | -0.148 | P1_P2_P3_P4_Lumbar_Stellate_VS_Coeliac_Sphenopalatine | 1 | 1 | 1 | 1 | 1 | 0 | 1 | 0 |
| HMGB3 | -0.148 | P3_Stellate_Sphenopalatine_VS_P1_P2_P4_Lumbar_Coeliac | 0 | 0 | 1 | 0 | 0 | 0 | 1 | 1 |
| GDI1 | -0.148 | P3_Stellate_Sphenopalatine_VS_P1_P2_P4_Lumbar_Coeliac | 0 | 0 | 1 | 0 | 0 | 0 | 1 | 1 |
| HNRNPA3 | -0.148 | P3_P4_Stellate_Sphenopalatine_VS_P1_P2_Lumbar_Coeliac | 0 | 0 | 1 | 1 | 0 | 0 | 1 | 1 |
| CTXN1 | -0.148 | P3_P4_Stellate_Sphenopalatine_VS_P1_P2_Lumbar_Coeliac | 0 | 0 | 1 | 1 | 0 | 0 | 1 | 1 |
| KANSL2 | -0.148 | P3_P4_Stellate_Sphenopalatine_VS_P1_P2_Lumbar_Coeliac | 0 | 0 | 1 | 1 | 0 | 0 | 1 | 1 |
| PFDN2 | -0.148 | P1_P3_P4_Stellate_Sphenopalatine_VS_P2_Lumbar_Coeliac | 1 | 0 | 1 | 1 | 0 | 0 | 1 | 1 |
| PGLS | -0.148 | P1_P3_P4_Stellate_Sphenopalatine_VS_P2_Lumbar_Coeliac | 1 | 0 | 1 | 1 | 0 | 0 | 1 | 1 |
| UBXN6 | -0.148 | P1_P3_P4_Stellate_Sphenopalatine_VS_P2_Lumbar_Coeliac | 1 | 0 | 1 | 1 | 0 | 0 | 1 | 1 |
| UBE3A | -0.148 | P2_P3_P4_Coeliac_Stellate_Sphenopalatine_VS_P1_Lumbar | 0 | 1 | 1 | 1 | 0 | 1 | 1 | 1 |
| EIF5 | -0.148 | P1_P2_P3_P4_Coeliac_Stellate_Sphenopalatine_VS_Lumbar | 1 | 1 | 1 | 1 | 0 | 1 | 1 | 1 |
| TWF1 | -0.148 | P2_P4_Lumbar_Coeliac_Stellate_Sphenopalatine_VS_P1_P3 | 0 | 1 | 0 | 1 | 1 | 1 | 1 | 1 |
| PIK3CA | -0.148 | P2_P3_P4_Lumbar_Coeliac_Stellate_Sphenopalatine_VS_P1 | 0 | 1 | 1 | 1 | 1 | 1 | 1 | 1 |
| ILF2 | -0.148 | P2_P3_P4_Lumbar_Coeliac_Stellate_Sphenopalatine_VS_P1 | 0 | 1 | 1 | 1 | 1 | 1 | 1 | 1 |
| CLSTN1 | -0.148 | P2_P3_P4_Lumbar_Coeliac_Stellate_Sphenopalatine_VS_P1 | 0 | 1 | 1 | 1 | 1 | 1 | 1 | 1 |
| RNF167 | -0.149 | P1_P2_P3_P4_Stellate_VS_Lumbar_Coeliac_Sphenopalatine | 1 | 1 | 1 | 1 | 0 | 0 | 1 | 0 |
| UBE2G1 | -0.149 | P4_Lumbar_Stellate_VS_P1_P2_P3_Coeliac_Sphenopalatine | 0 | 0 | 0 | 1 | 1 | 0 | 1 | 0 |
| CTPS | -0.149 | P1_P2_P4_Lumbar_Stellate_VS_P3_Coeliac_Sphenopalatine | 1 | 1 | 0 | 1 | 1 | 0 | 1 | 0 |

| | | | | | | | | | | |
|---|---|---|---|---|---|---|---|---|---|---|
| SLC25A33 | -0.149 | P1_P2_P4_Lumbar_Stellate_VS_P3_Coeliac_Sphenopalatine | 1 | 1 | 0 | 1 | 1 | 0 | 1 | 0 |
| CTNND2 | -0.149 | P2_P3_P4_Lumbar_Stellate_VS_P1_Coeliac_Sphenopalatine | 0 | 1 | 1 | 1 | 1 | 0 | 1 | 0 |
| METAP1 | -0.149 | P1_P2_P3_P4_Lumbar_Stellate_VS_Coeliac_Sphenopalatine | 1 | 1 | 1 | 1 | 1 | 0 | 1 | 0 |
| NCALD | -0.149 | P1_P2_P3_P4_Lumbar_Stellate_VS_Coeliac_Sphenopalatine | 1 | 1 | 1 | 1 | 1 | 0 | 1 | 0 |
| FBXO22 | -0.149 | P1_P2_Lumbar_Coeliac_Stellate_VS_P3_P4_Sphenopalatine | 1 | 1 | 0 | 0 | 1 | 1 | 1 | 0 |
| PCBP3 | -0.149 | P3_Stellate_Sphenopalatine_VS_P1_P2_P4_Lumbar_Coeliac | 0 | 0 | 1 | 0 | 0 | 0 | 1 | 1 |
| AOPEP | -0.149 | P1_P2_P3_Stellate_Sphenopalatine_VS_P4_Lumbar_Coeliac | 1 | 1 | 1 | 0 | 0 | 0 | 1 | 1 |
| SDF4 | -0.149 | P1_P2_P4_Stellate_Sphenopalatine_VS_P3_Lumbar_Coeliac | 1 | 1 | 0 | 1 | 0 | 0 | 1 | 1 |
| LAMTOR5 | -0.149 | P3_P4_Stellate_Sphenopalatine_VS_P1_P2_Lumbar_Coeliac | 0 | 0 | 1 | 1 | 0 | 0 | 1 | 1 |
| AP2S1 | -0.149 | P3_P4_Stellate_Sphenopalatine_VS_P1_P2_Lumbar_Coeliac | 0 | 0 | 1 | 1 | 0 | 0 | 1 | 1 |
| PET100 | -0.149 | P3_P4_Stellate_Sphenopalatine_VS_P1_P2_Lumbar_Coeliac | 0 | 0 | 1 | 1 | 0 | 0 | 1 | 1 |
| D8ERTD738E | -0.149 | P3_P4_Stellate_Sphenopalatine_VS_P1_P2_Lumbar_Coeliac | 0 | 0 | 1 | 1 | 0 | 0 | 1 | 1 |
| TMEM258 | -0.149 | P3_P4_Stellate_Sphenopalatine_VS_P1_P2_Lumbar_Coeliac | 0 | 0 | 1 | 1 | 0 | 0 | 1 | 1 |
| AZI2 | -0.149 | P1_P4_Lumbar_Stellate_Sphenopalatine_VS_P2_P3_Coeliac | 1 | 0 | 0 | 1 | 1 | 0 | 1 | 1 |
| 1500004A13 | -0.149 | P1_P2_P4_Lumbar_Stellate_Sphenopalatine_VS_P3_Coeliac | 1 | 1 | 0 | 1 | 1 | 0 | 1 | 1 |
| SF3B6 | -0.149 | P3_P4_Lumbar_Stellate_Sphenopalatine_VS_P1_P2_Coeliac | 0 | 0 | 1 | 1 | 1 | 0 | 1 | 1 |
| RPP21 | -0.149 | P3_P4_Lumbar_Stellate_Sphenopalatine_VS_P1_P2_Coeliac | 0 | 0 | 1 | 1 | 1 | 0 | 1 | 1 |
| SNU13 | -0.149 | P1_P2_P3_P4_Coeliac_Stellate_Sphenopalatine_VS_Lumbar | 1 | 1 | 1 | 1 | 0 | 1 | 1 | 1 |
| CLCN4 | -0.149 | P2_P3_Lumbar_Coeliac_Stellate_Sphenopalatine_VS_P1_P4 | 0 | 1 | 1 | 0 | 1 | 1 | 1 | 1 |
| MYADM | -0.149 | P2_P4_Lumbar_Coeliac_Stellate_Sphenopalatine_VS_P1_P3 | 0 | 1 | 0 | 1 | 1 | 1 | 1 | 1 |
| ASTN1 | -0.149 | P2_P3_P4_Lumbar_Coeliac_Stellate_Sphenopalatine_VS_P1 | 0 | 1 | 1 | 1 | 1 | 1 | 1 | 1 |
| CACNA1B | -0.149 | P2_P3_P4_Lumbar_Coeliac_Stellate_Sphenopalatine_VS_P1 | 0 | 1 | 1 | 1 | 1 | 1 | 1 | 1 |
| ABCD3 | -0.149 | P2_P3_P4_Lumbar_Coeliac_Stellate_Sphenopalatine_VS_P1 | 0 | 1 | 1 | 1 | 1 | 1 | 1 | 1 |
| MACF1 | -0.149 | P2_P3_P4_Lumbar_Coeliac_Stellate_Sphenopalatine_VS_P1 | 0 | 1 | 1 | 1 | 1 | 1 | 1 | 1 |
| PAFAH1B3 | -0.149 | P2_P3_P4_Lumbar_Coeliac_Stellate_Sphenopalatine_VS_P1 | 0 | 1 | 1 | 1 | 1 | 1 | 1 | 1 |
| YEATS4 | -0.149 | P2_P3_P4_Lumbar_Coeliac_Stellate_Sphenopalatine_VS_P1 | 0 | 1 | 1 | 1 | 1 | 1 | 1 | 1 |
| APC | -0.149 | P2_P3_P4_Lumbar_Coeliac_Stellate_Sphenopalatine_VS_P1 | 0 | 1 | 1 | 1 | 1 | 1 | 1 | 1 |
| CTIF | -0.149 | P2_P3_P4_Lumbar_Coeliac_Stellate_Sphenopalatine_VS_P1 | 0 | 1 | 1 | 1 | 1 | 1 | 1 | 1 |
| NDUFS3 | -0.15 | P1_P3_P4_Lumbar_Stellate_VS_P2_Coeliac_Sphenopalatine | 1 | 0 | 1 | 1 | 1 | 0 | 1 | 0 |
| 1700123O20 | -0.15 | P1_P3_P4_Lumbar_Stellate_VS_P2_Coeliac_Sphenopalatine | 1 | 0 | 1 | 1 | 1 | 0 | 1 | 0 |

| | | | | | | | | | | |
|---|---|---|---|---|---|---|---|---|---|---|
| IVD | -0.15 | P1_P2_Lumbar_Coeliac_Stellate_VS_P3_P4_Sphenopalatine | 1 | 1 | 0 | 0 | 1 | 1 | 1 | 0 |
| SPOP | -0.15 | P3_Stellate_Sphenopalatine_VS_P1_P2_P4_Lumbar_Coeliac | 0 | 0 | 1 | 0 | 0 | 0 | 1 | 1 |
| SEC62 | -0.15 | P2_P3_Stellate_Sphenopalatine_VS_P1_P4_Lumbar_Coeliac | 0 | 1 | 1 | 0 | 0 | 0 | 1 | 1 |
| COA3 | -0.15 | P3_P4_Stellate_Sphenopalatine_VS_P1_P2_Lumbar_Coeliac | 0 | 0 | 1 | 1 | 0 | 0 | 1 | 1 |
| BBIP1 | -0.15 | P3_P4_Stellate_Sphenopalatine_VS_P1_P2_Lumbar_Coeliac | 0 | 0 | 1 | 1 | 0 | 0 | 1 | 1 |
| POLR2F | -0.15 | P1_P3_Lumbar_Stellate_Sphenopalatine_VS_P2_P4_Coeliac | 1 | 0 | 1 | 0 | 1 | 0 | 1 | 1 |
| ANK2 | -0.15 | P1_P2_P3_Lumbar_Stellate_Sphenopalatine_VS_P4_Coeliac | 1 | 1 | 1 | 0 | 1 | 0 | 1 | 1 |
| FLOT1 | -0.15 | P1_P2_P4_Lumbar_Stellate_Sphenopalatine_VS_P3_Coeliac | 1 | 1 | 0 | 1 | 1 | 0 | 1 | 1 |
| DOHH | -0.15 | P3_P4_Lumbar_Stellate_Sphenopalatine_VS_P1_P2_Coeliac | 0 | 0 | 1 | 1 | 1 | 0 | 1 | 1 |
| ARPC5 | -0.15 | P1_P2_P3_P4_Coeliac_Stellate_Sphenopalatine_VS_Lumbar | 1 | 1 | 1 | 1 | 0 | 1 | 1 | 1 |
| GRCC10 | -0.15 | P1_P2_P3_P4_Coeliac_Stellate_Sphenopalatine_VS_Lumbar | 1 | 1 | 1 | 1 | 0 | 1 | 1 | 1 |
| SMPD3 | -0.15 | P2_P4_Lumbar_Coeliac_Stellate_Sphenopalatine_VS_P1_P3 | 0 | 1 | 0 | 1 | 1 | 1 | 1 | 1 |
| TALDO1 | -0.15 | P2_P3_P4_Lumbar_Coeliac_Stellate_Sphenopalatine_VS_P1 | 0 | 1 | 1 | 1 | 1 | 1 | 1 | 1 |
| NUS1 | -0.15 | P2_P3_P4_Lumbar_Coeliac_Stellate_Sphenopalatine_VS_P1 | 0 | 1 | 1 | 1 | 1 | 1 | 1 | 1 |
| PRMT1 | -0.151 | P1_P3_P4_Lumbar_Stellate_VS_P2_Coeliac_Sphenopalatine | 1 | 0 | 1 | 1 | 1 | 0 | 1 | 0 |
| CCS | -0.151 | P1_P3_P4_Lumbar_Stellate_VS_P2_Coeliac_Sphenopalatine | 1 | 0 | 1 | 1 | 1 | 0 | 1 | 0 |
| C1QTNF12 | -0.151 | P2_P3_P4_Lumbar_Stellate_VS_P1_Coeliac_Sphenopalatine | 0 | 1 | 1 | 1 | 1 | 0 | 1 | 0 |
| ERP44 | -0.151 | P1_P2_P3_P4_Lumbar_Stellate_VS_Coeliac_Sphenopalatine | 1 | 1 | 1 | 1 | 1 | 0 | 1 | 0 |
| HNRNPF | -0.151 | P1_P2_P3_P4_Lumbar_Stellate_VS_Coeliac_Sphenopalatine | 1 | 1 | 1 | 1 | 1 | 0 | 1 | 0 |
| GINM1 | -0.151 | P1_P2_P3_P4_Lumbar_Stellate_VS_Coeliac_Sphenopalatine | 1 | 1 | 1 | 1 | 1 | 0 | 1 | 0 |
| LIMK2 | -0.151 | P1_P2_P3_P4_Lumbar_Stellate_VS_Coeliac_Sphenopalatine | 1 | 1 | 1 | 1 | 1 | 0 | 1 | 0 |
| NIPSNAP1 | -0.151 | P1_P2_P3_P4_Lumbar_Stellate_VS_Coeliac_Sphenopalatine | 1 | 1 | 1 | 1 | 1 | 0 | 1 | 0 |
| NOLC1 | -0.151 | P1_P2_Lumbar_Coeliac_Stellate_VS_P3_P4_Sphenopalatine | 1 | 1 | 0 | 0 | 1 | 1 | 1 | 0 |
| VAMP2 | -0.151 | P1_P2_P3_Stellate_Sphenopalatine_VS_P4_Lumbar_Coeliac | 1 | 1 | 1 | 0 | 0 | 0 | 1 | 1 |
| PFDN4 | -0.151 | P4_Stellate_Sphenopalatine_VS_P1_P2_P3_Lumbar_Coeliac | 0 | 0 | 0 | 1 | 0 | 0 | 1 | 1 |
| ARL6IP1 | -0.151 | P3_P4_Stellate_Sphenopalatine_VS_P1_P2_Lumbar_Coeliac | 0 | 0 | 1 | 1 | 0 | 0 | 1 | 1 |
| RNF7 | -0.151 | P3_P4_Stellate_Sphenopalatine_VS_P1_P2_Lumbar_Coeliac | 0 | 0 | 1 | 1 | 0 | 0 | 1 | 1 |
| GUK1 | -0.151 | P3_P4_Stellate_Sphenopalatine_VS_P1_P2_Lumbar_Coeliac | 0 | 0 | 1 | 1 | 0 | 0 | 1 | 1 |
| NPC2 | -0.151 | P3_P4_Stellate_Sphenopalatine_VS_P1_P2_Lumbar_Coeliac | 0 | 0 | 1 | 1 | 0 | 0 | 1 | 1 |
| DPY30 | -0.151 | P3_P4_Stellate_Sphenopalatine_VS_P1_P2_Lumbar_Coeliac | 0 | 0 | 1 | 1 | 0 | 0 | 1 | 1 |

| | | | | | | | | | | |
|---|---|---|---|---|---|---|---|---|---|---|
| BEX4 | -0.151 | P3_P4_Stellate_Sphenopalatine_VS_P1_P2_Lumbar_Coeliac | 0 | 0 | 1 | 1 | 0 | 0 | 1 | 1 |
| TPI1 | -0.151 | P1_P2_P3_Lumbar_Stellate_Sphenopalatine_VS_P4_Coeliac | 1 | 1 | 1 | 0 | 1 | 0 | 1 | 1 |
| MDP1 | -0.151 | P1_P2_P4_Lumbar_Stellate_Sphenopalatine_VS_P3_Coeliac | 1 | 1 | 0 | 1 | 1 | 0 | 1 | 1 |
| EIF5B | -0.151 | P3_P4_Coeliac_Stellate_Sphenopalatine_VS_P1_P2_Lumbar | 0 | 0 | 1 | 1 | 0 | 1 | 1 | 1 |
| BC031181 | -0.151 | P2_P3_P4_Coeliac_Stellate_Sphenopalatine_VS_P1_Lumbar | 0 | 1 | 1 | 1 | 0 | 1 | 1 | 1 |
| SKP1A | -0.151 | P1_P2_P3_P4_Coeliac_Stellate_Sphenopalatine_VS_Lumbar | 1 | 1 | 1 | 1 | 0 | 1 | 1 | 1 |
| UBE2D2A | -0.151 | P1_P2_P3_P4_Coeliac_Stellate_Sphenopalatine_VS_Lumbar | 1 | 1 | 1 | 1 | 0 | 1 | 1 | 1 |
| CIR1 | -0.151 | P2_P3_P4_Lumbar_Coeliac_Stellate_Sphenopalatine_VS_P1 | 0 | 1 | 1 | 1 | 1 | 1 | 1 | 1 |
| CTNND1 | -0.151 | P2_P3_P4_Lumbar_Coeliac_Stellate_Sphenopalatine_VS_P1 | 0 | 1 | 1 | 1 | 1 | 1 | 1 | 1 |
| DCLK1 | -0.151 | P2_P3_P4_Lumbar_Coeliac_Stellate_Sphenopalatine_VS_P1 | 0 | 1 | 1 | 1 | 1 | 1 | 1 | 1 |
| ZYG11B | -0.151 | P2_P3_P4_Lumbar_Coeliac_Stellate_Sphenopalatine_VS_P1 | 0 | 1 | 1 | 1 | 1 | 1 | 1 | 1 |
| EIF3B | -0.151 | P2_P3_P4_Lumbar_Coeliac_Stellate_Sphenopalatine_VS_P1 | 0 | 1 | 1 | 1 | 1 | 1 | 1 | 1 |
| TRA2A | -0.151 | P2_P3_P4_Lumbar_Coeliac_Stellate_Sphenopalatine_VS_P1 | 0 | 1 | 1 | 1 | 1 | 1 | 1 | 1 |
| BAX | -0.151 | P2_P3_P4_Lumbar_Coeliac_Stellate_Sphenopalatine_VS_P1 | 0 | 1 | 1 | 1 | 1 | 1 | 1 | 1 |
| RNF227 | -0.151 | P2_P3_P4_Lumbar_Coeliac_Stellate_Sphenopalatine_VS_P1 | 0 | 1 | 1 | 1 | 1 | 1 | 1 | 1 |
| TMEM237 | -0.152 | P1_P3_P4_Lumbar_Stellate_VS_P2_Coeliac_Sphenopalatine | 1 | 0 | 1 | 1 | 1 | 0 | 1 | 0 |
| PITPNB | -0.152 | P1_P2_Lumbar_Coeliac_Stellate_VS_P3_P4_Sphenopalatine | 1 | 1 | 0 | 0 | 1 | 1 | 1 | 0 |
| TSPAN3 | -0.152 | P3_Stellate_Sphenopalatine_VS_P1_P2_P4_Lumbar_Coeliac | 0 | 0 | 1 | 0 | 0 | 0 | 1 | 1 |
| HIGD1A | -0.152 | P1_P2_P3_Stellate_Sphenopalatine_VS_P4_Lumbar_Coeliac | 1 | 1 | 1 | 0 | 0 | 0 | 1 | 1 |
| POLR2M | -0.152 | P3_P4_Stellate_Sphenopalatine_VS_P1_P2_Lumbar_Coeliac | 0 | 0 | 1 | 1 | 0 | 0 | 1 | 1 |
| CHTOP | -0.152 | P1_P2_P3_Lumbar_Stellate_Sphenopalatine_VS_P4_Coeliac | 1 | 1 | 1 | 0 | 1 | 0 | 1 | 1 |
| RBMS3 | -0.152 | P1_P2_P3_Lumbar_Stellate_Sphenopalatine_VS_P4_Coeliac | 1 | 1 | 1 | 0 | 1 | 0 | 1 | 1 |
| HIGD2A | -0.152 | P1_P2_P3_P4_Coeliac_Stellate_Sphenopalatine_VS_Lumbar | 1 | 1 | 1 | 1 | 0 | 1 | 1 | 1 |
| RALBP1 | -0.152 | P1_P2_P3_P4_Coeliac_Stellate_Sphenopalatine_VS_Lumbar | 1 | 1 | 1 | 1 | 0 | 1 | 1 | 1 |
| RABGAP1 | -0.152 | P2_P3_P4_Lumbar_Coeliac_Stellate_Sphenopalatine_VS_P1 | 0 | 1 | 1 | 1 | 1 | 1 | 1 | 1 |
| BAZ2B | -0.152 | P2_P3_P4_Lumbar_Coeliac_Stellate_Sphenopalatine_VS_P1 | 0 | 1 | 1 | 1 | 1 | 1 | 1 | 1 |
| UBE2K | -0.152 | P2_P3_P4_Lumbar_Coeliac_Stellate_Sphenopalatine_VS_P1 | 0 | 1 | 1 | 1 | 1 | 1 | 1 | 1 |
| SLC16A3 | -0.152 | P2_P3_P4_Lumbar_Coeliac_Stellate_Sphenopalatine_VS_P1 | 0 | 1 | 1 | 1 | 1 | 1 | 1 | 1 |
| VBP1 | -0.153 | P2_P3_P4_Stellate_VS_P1_Lumbar_Coeliac_Sphenopalatine | 0 | 1 | 1 | 1 | 0 | 0 | 1 | 0 |
| HPRT | -0.153 | P1_P2_P3_P4_Stellate_VS_Lumbar_Coeliac_Sphenopalatine | 1 | 1 | 1 | 1 | 0 | 0 | 1 | 0 |

| | | | | | | | | | | |
|---|---|---|---|---|---|---|---|---|---|---|
| SNRPB | -0.153 | P1_P3_P4_Lumbar_Stellate_VS_P2_Coeliac_Sphenopalatine | 1 | 0 | 1 | 1 | 1 | 0 | 1 | 0 |
| LARS | -0.153 | P1_P2_Lumbar_Coeliac_Stellate_VS_P3_P4_Sphenopalatine | 1 | 1 | 0 | 0 | 1 | 1 | 1 | 0 |
| MRPL34 | -0.153 | P3_Stellate_Sphenopalatine_VS_P1_P2_P4_Lumbar_Coeliac | 0 | 0 | 1 | 0 | 0 | 0 | 1 | 1 |
| SMIM7 | -0.153 | P2_P3_Stellate_Sphenopalatine_VS_P1_P4_Lumbar_Coeliac | 0 | 1 | 1 | 0 | 0 | 0 | 1 | 1 |
| GARNL3 | -0.153 | P1_P2_P3_Stellate_Sphenopalatine_VS_P4_Lumbar_Coeliac | 1 | 1 | 1 | 0 | 0 | 0 | 1 | 1 |
| RAB2A | -0.153 | P1_P2_P3_Stellate_Sphenopalatine_VS_P4_Lumbar_Coeliac | 1 | 1 | 1 | 0 | 0 | 0 | 1 | 1 |
| CTSL | -0.153 | P1_P3_Lumbar_Stellate_Sphenopalatine_VS_P2_P4_Coeliac | 1 | 0 | 1 | 0 | 1 | 0 | 1 | 1 |
| UFM1 | -0.153 | P1_P2_P3_Lumbar_Stellate_Sphenopalatine_VS_P4_Coeliac | 1 | 1 | 1 | 0 | 1 | 0 | 1 | 1 |
| FAM32A | -0.153 | P1_P2_P3_Lumbar_Stellate_Sphenopalatine_VS_P4_Coeliac | 1 | 1 | 1 | 0 | 1 | 0 | 1 | 1 |
| VPS29 | -0.153 | P1_P2_P4_Lumbar_Stellate_Sphenopalatine_VS_P3_Coeliac | 1 | 1 | 0 | 1 | 1 | 0 | 1 | 1 |
| AHI1 | -0.153 | P3_P4_Coeliac_Stellate_Sphenopalatine_VS_P1_P2_Lumbar | 0 | 0 | 1 | 1 | 0 | 1 | 1 | 1 |
| DUSP26 | -0.153 | P2_P3_P4_Coeliac_Stellate_Sphenopalatine_VS_P1_Lumbar | 0 | 1 | 1 | 1 | 0 | 1 | 1 | 1 |
| SET | -0.153 | P1_P2_P3_P4_Coeliac_Stellate_Sphenopalatine_VS_Lumbar | 1 | 1 | 1 | 1 | 0 | 1 | 1 | 1 |
| NDRG2 | -0.153 | P1_Lumbar_Coeliac_Stellate_Sphenopalatine_VS_P2_P3_P4 | 1 | 0 | 0 | 0 | 1 | 1 | 1 | 1 |
| NYAP1 | -0.153 | P2_P4_Lumbar_Coeliac_Stellate_Sphenopalatine_VS_P1_P3 | 0 | 1 | 0 | 1 | 1 | 1 | 1 | 1 |
| DLGAP4 | -0.153 | P2_P3_P4_Lumbar_Coeliac_Stellate_Sphenopalatine_VS_P1 | 0 | 1 | 1 | 1 | 1 | 1 | 1 | 1 |
| CCNI | -0.153 | P2_P3_P4_Lumbar_Coeliac_Stellate_Sphenopalatine_VS_P1 | 0 | 1 | 1 | 1 | 1 | 1 | 1 | 1 |
| RCN2 | -0.153 | P2_P3_P4_Lumbar_Coeliac_Stellate_Sphenopalatine_VS_P1 | 0 | 1 | 1 | 1 | 1 | 1 | 1 | 1 |
| GMFB | -0.153 | P2_P3_P4_Lumbar_Coeliac_Stellate_Sphenopalatine_VS_P1 | 0 | 1 | 1 | 1 | 1 | 1 | 1 | 1 |
| HDAC3 | -0.154 | P1_P3_Lumbar_Stellate_VS_P2_P4_Coeliac_Sphenopalatine | 1 | 0 | 1 | 0 | 1 | 0 | 1 | 0 |
| DBNDD1 | -0.154 | P1_P4_Lumbar_Stellate_VS_P2_P3_Coeliac_Sphenopalatine | 1 | 0 | 0 | 1 | 1 | 0 | 1 | 0 |
| REXO1 | -0.154 | P1_P2_Lumbar_Coeliac_Stellate_VS_P3_P4_Sphenopalatine | 1 | 1 | 0 | 0 | 1 | 1 | 1 | 0 |
| DIABLO | -0.154 | P1_P2_P3_Stellate_Sphenopalatine_VS_P4_Lumbar_Coeliac | 1 | 1 | 1 | 0 | 0 | 0 | 1 | 1 |
| VAPB | -0.154 | P1_P2_P4_Stellate_Sphenopalatine_VS_P3_Lumbar_Coeliac | 1 | 1 | 0 | 1 | 0 | 0 | 1 | 1 |
| STX7 | -0.154 | P1_P2_P4_Stellate_Sphenopalatine_VS_P3_Lumbar_Coeliac | 1 | 1 | 0 | 1 | 0 | 0 | 1 | 1 |
| RPS27L | -0.154 | P1_P2_P3_Lumbar_Stellate_Sphenopalatine_VS_P4_Coeliac | 1 | 1 | 1 | 0 | 1 | 0 | 1 | 1 |
| RAC3 | -0.154 | P1_P2_P4_Lumbar_Stellate_Sphenopalatine_VS_P3_Coeliac | 1 | 1 | 0 | 1 | 1 | 0 | 1 | 1 |
| CHMP2A | -0.154 | P1_P3_P4_Coeliac_Stellate_Sphenopalatine_VS_P2_Lumbar | 1 | 0 | 1 | 1 | 0 | 1 | 1 | 1 |
| PABPN1 | -0.154 | P2_P3_P4_Coeliac_Stellate_Sphenopalatine_VS_P1_Lumbar | 0 | 1 | 1 | 1 | 0 | 1 | 1 | 1 |
| DYNC1I2 | -0.154 | P1_P2_P3_P4_Coeliac_Stellate_Sphenopalatine_VS_Lumbar | 1 | 1 | 1 | 1 | 0 | 1 | 1 | 1 |

| | | | | | | | | | | |
|---|---|---|---|---|---|---|---|---|---|---|
| METAP2 | -0.154 | P1_P2_P3_P4_Coeliac_Stellate_Sphenopalatine_VS_Lumbar | 1 | 1 | 1 | 1 | 0 | 1 | 1 | 1 |
| NSG2 | -0.154 | P1_P2_P3_P4_Coeliac_Stellate_Sphenopalatine_VS_Lumbar | 1 | 1 | 1 | 1 | 0 | 1 | 1 | 1 |
| CHD3 | -0.154 | P1_P2_P3_P4_Coeliac_Stellate_Sphenopalatine_VS_Lumbar | 1 | 1 | 1 | 1 | 0 | 1 | 1 | 1 |
| SOD1 | -0.154 | P1_P2_P3_P4_Coeliac_Stellate_Sphenopalatine_VS_Lumbar | 1 | 1 | 1 | 1 | 0 | 1 | 1 | 1 |
| SMS | -0.154 | P1_P2_P3_P4_Coeliac_Stellate_Sphenopalatine_VS_Lumbar | 1 | 1 | 1 | 1 | 0 | 1 | 1 | 1 |
| AP3B2 | -0.154 | P2_P4_Lumbar_Coeliac_Stellate_Sphenopalatine_VS_P1_P3 | 0 | 1 | 0 | 1 | 1 | 1 | 1 | 1 |
| TMCO1 | -0.154 | P2_P3_P4_Lumbar_Coeliac_Stellate_Sphenopalatine_VS_P1 | 0 | 1 | 1 | 1 | 1 | 1 | 1 | 1 |
| BROX | -0.154 | P2_P3_P4_Lumbar_Coeliac_Stellate_Sphenopalatine_VS_P1 | 0 | 1 | 1 | 1 | 1 | 1 | 1 | 1 |
| EMC3 | -0.154 | P2_P3_P4_Lumbar_Coeliac_Stellate_Sphenopalatine_VS_P1 | 0 | 1 | 1 | 1 | 1 | 1 | 1 | 1 |
| U2SURP | -0.154 | P2_P3_P4_Lumbar_Coeliac_Stellate_Sphenopalatine_VS_P1 | 0 | 1 | 1 | 1 | 1 | 1 | 1 | 1 |
| 5031439G07 | -0.154 | P2_P3_P4_Lumbar_Coeliac_Stellate_Sphenopalatine_VS_P1 | 0 | 1 | 1 | 1 | 1 | 1 | 1 | 1 |
| HMOX2 | -0.154 | P2_P3_P4_Lumbar_Coeliac_Stellate_Sphenopalatine_VS_P1 | 0 | 1 | 1 | 1 | 1 | 1 | 1 | 1 |
| AGPAT4 | -0.154 | P2_P3_P4_Lumbar_Coeliac_Stellate_Sphenopalatine_VS_P1 | 0 | 1 | 1 | 1 | 1 | 1 | 1 | 1 |
| PPM1B | -0.154 | P2_P3_P4_Lumbar_Coeliac_Stellate_Sphenopalatine_VS_P1 | 0 | 1 | 1 | 1 | 1 | 1 | 1 | 1 |
| TUSC3 | -0.155 | P3_P4_Stellate_VS_P1_P2_Lumbar_Coeliac_Sphenopalatine | 0 | 0 | 1 | 1 | 0 | 0 | 1 | 0 |
| TOMM22 | -0.155 | P3_P4_Stellate_VS_P1_P2_Lumbar_Coeliac_Sphenopalatine | 0 | 0 | 1 | 1 | 0 | 0 | 1 | 0 |
| TIMM23 | -0.155 | P1_P3_P4_Stellate_VS_P2_Lumbar_Coeliac_Sphenopalatine | 1 | 0 | 1 | 1 | 0 | 0 | 1 | 0 |
| VSTM2L | -0.155 | P1_P2_P3_P4_Coeliac_Stellate_VS_Lumbar_Sphenopalatine | 1 | 1 | 1 | 1 | 0 | 1 | 1 | 0 |
| SUMO1 | -0.155 | P3_Stellate_Sphenopalatine_VS_P1_P2_P4_Lumbar_Coeliac | 0 | 0 | 1 | 0 | 0 | 0 | 1 | 1 |
| KLF7 | -0.155 | P3_Stellate_Sphenopalatine_VS_P1_P2_P4_Lumbar_Coeliac | 0 | 0 | 1 | 0 | 0 | 0 | 1 | 1 |
| EDF1 | -0.155 | P3_Stellate_Sphenopalatine_VS_P1_P2_P4_Lumbar_Coeliac | 0 | 0 | 1 | 0 | 0 | 0 | 1 | 1 |
| RAB5IF | -0.155 | P3_Stellate_Sphenopalatine_VS_P1_P2_P4_Lumbar_Coeliac | 0 | 0 | 1 | 0 | 0 | 0 | 1 | 1 |
| TPRGL | -0.155 | P3_Stellate_Sphenopalatine_VS_P1_P2_P4_Lumbar_Coeliac | 0 | 0 | 1 | 0 | 0 | 0 | 1 | 1 |
| RAB10 | -0.155 | P3_Stellate_Sphenopalatine_VS_P1_P2_P4_Lumbar_Coeliac | 0 | 0 | 1 | 0 | 0 | 0 | 1 | 1 |
| SLC25A51 | -0.155 | P2_P3_Stellate_Sphenopalatine_VS_P1_P4_Lumbar_Coeliac | 0 | 1 | 1 | 0 | 0 | 0 | 1 | 1 |
| BOLA2 | -0.155 | P2_P3_Stellate_Sphenopalatine_VS_P1_P4_Lumbar_Coeliac | 0 | 1 | 1 | 0 | 0 | 0 | 1 | 1 |
| PPP1R2 | -0.155 | P1_P2_P3_Stellate_Sphenopalatine_VS_P4_Lumbar_Coeliac | 1 | 1 | 1 | 0 | 0 | 0 | 1 | 1 |
| CDS2 | -0.155 | P1_P2_P4_Stellate_Sphenopalatine_VS_P3_Lumbar_Coeliac | 1 | 1 | 0 | 1 | 0 | 0 | 1 | 1 |
| ACBD6 | -0.155 | P1_P3_Lumbar_Stellate_Sphenopalatine_VS_P2_P4_Coeliac | 1 | 0 | 1 | 0 | 1 | 0 | 1 | 1 |
| SQLE | -0.155 | P1_P3_Lumbar_Stellate_Sphenopalatine_VS_P2_P4_Coeliac | 1 | 0 | 1 | 0 | 1 | 0 | 1 | 1 |

| | | | | | | | | | | |
|---|---|---|---|---|---|---|---|---|---|---|
| ATP1A1 | -0.155 | P1_P2_P3_Lumbar_Stellate_Sphenopalatine_VS_P4_Coeliac | 1 | 1 | 1 | 0 | 1 | 0 | 1 | 1 |
| TRP53INP2 | -0.155 | P1_P2_P4_Lumbar_Stellate_Sphenopalatine_VS_P3_Coeliac | 1 | 1 | 0 | 1 | 1 | 0 | 1 | 1 |
| GLUD1 | -0.155 | P1_P2_P4_Lumbar_Stellate_Sphenopalatine_VS_P3_Coeliac | 1 | 1 | 0 | 1 | 1 | 0 | 1 | 1 |
| RETREG1 | -0.155 | P1_P2_P4_Lumbar_Stellate_Sphenopalatine_VS_P3_Coeliac | 1 | 1 | 0 | 1 | 1 | 0 | 1 | 1 |
| HAND2OS1 | -0.155 | P3_Coeliac_Stellate_Sphenopalatine_VS_P1_P2_P4_Lumbar | 0 | 0 | 1 | 0 | 0 | 1 | 1 | 1 |
| RWDD1 | -0.155 | P3_P4_Coeliac_Stellate_Sphenopalatine_VS_P1_P2_Lumbar | 0 | 0 | 1 | 1 | 0 | 1 | 1 | 1 |
| DNAJC5 | -0.155 | P1_P2_P3_P4_Coeliac_Stellate_Sphenopalatine_VS_Lumbar | 1 | 1 | 1 | 1 | 0 | 1 | 1 | 1 |
| CTBP1 | -0.155 | P1_P2_P3_P4_Coeliac_Stellate_Sphenopalatine_VS_Lumbar | 1 | 1 | 1 | 1 | 0 | 1 | 1 | 1 |
| COX17 | -0.155 | P1_P2_P3_P4_Coeliac_Stellate_Sphenopalatine_VS_Lumbar | 1 | 1 | 1 | 1 | 0 | 1 | 1 | 1 |
| ANAPC16 | -0.155 | Lumbar_Coeliac_Stellate_Sphenopalatine_VS_P1_P2_P3_P4 | 0 | 0 | 0 | 0 | 1 | 1 | 1 | 1 |
| GTF2I | -0.155 | P2_P3_P4_Lumbar_Coeliac_Stellate_Sphenopalatine_VS_P1 | 0 | 1 | 1 | 1 | 1 | 1 | 1 | 1 |
| STRN4 | -0.155 | P2_P3_P4_Lumbar_Coeliac_Stellate_Sphenopalatine_VS_P1 | 0 | 1 | 1 | 1 | 1 | 1 | 1 | 1 |
| RAB5C | -0.155 | P2_P3_P4_Lumbar_Coeliac_Stellate_Sphenopalatine_VS_P1 | 0 | 1 | 1 | 1 | 1 | 1 | 1 | 1 |
| MED30 | -0.156 | P1_P3_Lumbar_Stellate_VS_P2_P4_Coeliac_Sphenopalatine | 1 | 0 | 1 | 0 | 1 | 0 | 1 | 0 |
| NANS | -0.156 | P1_P4_Lumbar_Stellate_VS_P2_P3_Coeliac_Sphenopalatine | 1 | 0 | 0 | 1 | 1 | 0 | 1 | 0 |
| PBX1 | -0.156 | P1_P2_P4_Lumbar_Stellate_VS_P3_Coeliac_Sphenopalatine | 1 | 1 | 0 | 1 | 1 | 0 | 1 | 0 |
| GTPBP4 | -0.156 | P1_P2_Lumbar_Coeliac_Stellate_VS_P3_P4_Sphenopalatine | 1 | 1 | 0 | 0 | 1 | 1 | 1 | 0 |
| CNBP | -0.156 | P3_Stellate_Sphenopalatine_VS_P1_P2_P4_Lumbar_Coeliac | 0 | 0 | 1 | 0 | 0 | 0 | 1 | 1 |
| MRPL23 | -0.156 | P3_Stellate_Sphenopalatine_VS_P1_P2_P4_Lumbar_Coeliac | 0 | 0 | 1 | 0 | 0 | 0 | 1 | 1 |
| CETN3 | -0.156 | P3_Stellate_Sphenopalatine_VS_P1_P2_P4_Lumbar_Coeliac | 0 | 0 | 1 | 0 | 0 | 0 | 1 | 1 |
| DGCR6 | -0.156 | P3_Stellate_Sphenopalatine_VS_P1_P2_P4_Lumbar_Coeliac | 0 | 0 | 1 | 0 | 0 | 0 | 1 | 1 |
| SSU72 | -0.156 | P1_P3_Stellate_Sphenopalatine_VS_P2_P4_Lumbar_Coeliac | 1 | 0 | 1 | 0 | 0 | 0 | 1 | 1 |
| RUNDC3A | -0.156 | P2_P3_Stellate_Sphenopalatine_VS_P1_P4_Lumbar_Coeliac | 0 | 1 | 1 | 0 | 0 | 0 | 1 | 1 |
| CHURC1 | -0.156 | P1_P2_P3_Stellate_Sphenopalatine_VS_P4_Lumbar_Coeliac | 1 | 1 | 1 | 0 | 0 | 0 | 1 | 1 |
| EIF3G | -0.156 | P1_P3_Lumbar_Stellate_Sphenopalatine_VS_P2_P4_Coeliac | 1 | 0 | 1 | 0 | 1 | 0 | 1 | 1 |
| UQCRFS1 | -0.156 | P1_P2_P3_Lumbar_Stellate_Sphenopalatine_VS_P4_Coeliac | 1 | 1 | 1 | 0 | 1 | 0 | 1 | 1 |
| PPP2R2B | -0.156 | P1_P2_P3_Lumbar_Stellate_Sphenopalatine_VS_P4_Coeliac | 1 | 1 | 1 | 0 | 1 | 0 | 1 | 1 |
| CIAPIN1 | -0.156 | P1_P4_Lumbar_Stellate_Sphenopalatine_VS_P2_P3_Coeliac | 1 | 0 | 0 | 1 | 1 | 0 | 1 | 1 |
| ACLY | -0.156 | P1_P2_P4_Lumbar_Stellate_Sphenopalatine_VS_P3_Coeliac | 1 | 1 | 0 | 1 | 1 | 0 | 1 | 1 |
| CAMTA1 | -0.156 | P3_P4_Coeliac_Stellate_Sphenopalatine_VS_P1_P2_Lumbar | 0 | 0 | 1 | 1 | 0 | 1 | 1 | 1 |

| | | | | | | | | | | |
|---|---|---|---|---|---|---|---|---|---|---|
| RANBP1 | -0.156 | P3_P4_Coeliac_Stellate_Sphenopalatine_VS_P1_P2_Lumbar | 0 | 0 | 1 | 1 | 0 | 1 | 1 | 1 |
| SNRPF | -0.156 | P1_P3_P4_Coeliac_Stellate_Sphenopalatine_VS_P2_Lumbar | 1 | 0 | 1 | 1 | 0 | 1 | 1 | 1 |
| CMPK1 | -0.156 | P1_P2_P3_P4_Coeliac_Stellate_Sphenopalatine_VS_Lumbar | 1 | 1 | 1 | 1 | 0 | 1 | 1 | 1 |
| MRPS33 | -0.156 | P1_P2_P3_P4_Coeliac_Stellate_Sphenopalatine_VS_Lumbar | 1 | 1 | 1 | 1 | 0 | 1 | 1 | 1 |
| MAPT | -0.156 | P1_P2_P3_P4_Coeliac_Stellate_Sphenopalatine_VS_Lumbar | 1 | 1 | 1 | 1 | 0 | 1 | 1 | 1 |
| CLTB | -0.156 | P1_P2_P3_P4_Coeliac_Stellate_Sphenopalatine_VS_Lumbar | 1 | 1 | 1 | 1 | 0 | 1 | 1 | 1 |
| GNAI3 | -0.156 | P2_P4_Lumbar_Coeliac_Stellate_Sphenopalatine_VS_P1_P3 | 0 | 1 | 0 | 1 | 1 | 1 | 1 | 1 |
| TSN | -0.156 | P2_P3_P4_Lumbar_Coeliac_Stellate_Sphenopalatine_VS_P1 | 0 | 1 | 1 | 1 | 1 | 1 | 1 | 1 |
| HNRNPR | -0.156 | P2_P3_P4_Lumbar_Coeliac_Stellate_Sphenopalatine_VS_P1 | 0 | 1 | 1 | 1 | 1 | 1 | 1 | 1 |
| UBE3C | -0.156 | P2_P3_P4_Lumbar_Coeliac_Stellate_Sphenopalatine_VS_P1 | 0 | 1 | 1 | 1 | 1 | 1 | 1 | 1 |
| TTYH3 | -0.156 | P2_P3_P4_Lumbar_Coeliac_Stellate_Sphenopalatine_VS_P1 | 0 | 1 | 1 | 1 | 1 | 1 | 1 | 1 |
| PAK1 | -0.156 | P2_P3_P4_Lumbar_Coeliac_Stellate_Sphenopalatine_VS_P1 | 0 | 1 | 1 | 1 | 1 | 1 | 1 | 1 |
| EPM2AIP1 | -0.156 | P2_P3_P4_Lumbar_Coeliac_Stellate_Sphenopalatine_VS_P1 | 0 | 1 | 1 | 1 | 1 | 1 | 1 | 1 |
| NCOA7 | -0.156 | P2_P3_P4_Lumbar_Coeliac_Stellate_Sphenopalatine_VS_P1 | 0 | 1 | 1 | 1 | 1 | 1 | 1 | 1 |
| LDHA | -0.157 | P1_P2_P3_P4_Stellate_VS_Lumbar_Coeliac_Sphenopalatine | 1 | 1 | 1 | 1 | 0 | 0 | 1 | 0 |
| NDN | -0.157 | P1_P2_P3_P4_Stellate_VS_Lumbar_Coeliac_Sphenopalatine | 1 | 1 | 1 | 1 | 0 | 0 | 1 | 0 |
| SELENOS | -0.157 | P1_P2_Lumbar_Coeliac_Stellate_VS_P3_P4_Sphenopalatine | 1 | 1 | 0 | 0 | 1 | 1 | 1 | 0 |
| NENF | -0.157 | P3_Stellate_Sphenopalatine_VS_P1_P2_P4_Lumbar_Coeliac | 0 | 0 | 1 | 0 | 0 | 0 | 1 | 1 |
| TMEM234 | -0.157 | P3_Stellate_Sphenopalatine_VS_P1_P2_P4_Lumbar_Coeliac | 0 | 0 | 1 | 0 | 0 | 0 | 1 | 1 |
| TIMM8B | -0.157 | P3_Stellate_Sphenopalatine_VS_P1_P2_P4_Lumbar_Coeliac | 0 | 0 | 1 | 0 | 0 | 0 | 1 | 1 |
| TRAPPC6B | -0.157 | P2_P4_Stellate_Sphenopalatine_VS_P1_P3_Lumbar_Coeliac | 0 | 1 | 0 | 1 | 0 | 0 | 1 | 1 |
| SAT1 | -0.157 | P1_P3_Lumbar_Stellate_Sphenopalatine_VS_P2_P4_Coeliac | 1 | 0 | 1 | 0 | 1 | 0 | 1 | 1 |
| ATP5C1 | -0.157 | P1_P2_P3_Lumbar_Stellate_Sphenopalatine_VS_P4_Coeliac | 1 | 1 | 1 | 0 | 1 | 0 | 1 | 1 |
| GLRX2 | -0.157 | P4_Lumbar_Stellate_Sphenopalatine_VS_P1_P2_P3_Coeliac | 0 | 0 | 0 | 1 | 1 | 0 | 1 | 1 |
| STX1A | -0.157 | P1_P2_P4_Lumbar_Stellate_Sphenopalatine_VS_P3_Coeliac | 1 | 1 | 0 | 1 | 1 | 0 | 1 | 1 |
| TCEAL9 | -0.157 | P2_P3_P4_Coeliac_Stellate_Sphenopalatine_VS_P1_Lumbar | 0 | 1 | 1 | 1 | 0 | 1 | 1 | 1 |
| EIF2S2 | -0.157 | P1_P2_P3_P4_Coeliac_Stellate_Sphenopalatine_VS_Lumbar | 1 | 1 | 1 | 1 | 0 | 1 | 1 | 1 |
| TMED2 | -0.157 | P1_P2_P3_P4_Coeliac_Stellate_Sphenopalatine_VS_Lumbar | 1 | 1 | 1 | 1 | 0 | 1 | 1 | 1 |
| SERF1 | -0.157 | P1_P2_P3_P4_Coeliac_Stellate_Sphenopalatine_VS_Lumbar | 1 | 1 | 1 | 1 | 0 | 1 | 1 | 1 |
| TBC1D7 | -0.158 | P1_P4_Lumbar_Stellate_VS_P2_P3_Coeliac_Sphenopalatine | 1 | 0 | 0 | 1 | 1 | 0 | 1 | 0 |

| | | | | | | | | | | |
|---|---|---|---|---|---|---|---|---|---|---|
| DNAJB6 | -0.158 | P3_Stellate_Sphenopalatine_VS_P1_P2_P4_Lumbar_Coeliac | 0 | 0 | 1 | 0 | 0 | 0 | 1 | 1 |
| ANAPC5 | -0.158 | P3_Stellate_Sphenopalatine_VS_P1_P2_P4_Lumbar_Coeliac | 0 | 0 | 1 | 0 | 0 | 0 | 1 | 1 |
| PDAP1 | -0.158 | P3_Stellate_Sphenopalatine_VS_P1_P2_P4_Lumbar_Coeliac | 0 | 0 | 1 | 0 | 0 | 0 | 1 | 1 |
| SNRNP27 | -0.158 | P3_Stellate_Sphenopalatine_VS_P1_P2_P4_Lumbar_Coeliac | 0 | 0 | 1 | 0 | 0 | 0 | 1 | 1 |
| PALM | -0.158 | P3_Stellate_Sphenopalatine_VS_P1_P2_P4_Lumbar_Coeliac | 0 | 0 | 1 | 0 | 0 | 0 | 1 | 1 |
| ZCRB1 | -0.158 | P3_Stellate_Sphenopalatine_VS_P1_P2_P4_Lumbar_Coeliac | 0 | 0 | 1 | 0 | 0 | 0 | 1 | 1 |
| IER3IP1 | -0.158 | P3_Stellate_Sphenopalatine_VS_P1_P2_P4_Lumbar_Coeliac | 0 | 0 | 1 | 0 | 0 | 0 | 1 | 1 |
| FAM168B | -0.158 | P1_P2_P4_Stellate_Sphenopalatine_VS_P3_Lumbar_Coeliac | 1 | 1 | 0 | 1 | 0 | 0 | 1 | 1 |
| MRPL57 | -0.158 | P1_P3_Lumbar_Stellate_Sphenopalatine_VS_P2_P4_Coeliac | 1 | 0 | 1 | 0 | 1 | 0 | 1 | 1 |
| SNHG6 | -0.158 | P1_P2_P3_Lumbar_Stellate_Sphenopalatine_VS_P4_Coeliac | 1 | 1 | 1 | 0 | 1 | 0 | 1 | 1 |
| APLP1 | -0.158 | P1_P2_P3_Lumbar_Stellate_Sphenopalatine_VS_P4_Coeliac | 1 | 1 | 1 | 0 | 1 | 0 | 1 | 1 |
| CHMP1A | -0.158 | P1_P2_P3_Lumbar_Stellate_Sphenopalatine_VS_P4_Coeliac | 1 | 1 | 1 | 0 | 1 | 0 | 1 | 1 |
| SDHD | -0.158 | P4_Lumbar_Stellate_Sphenopalatine_VS_P1_P2_P3_Coeliac | 0 | 0 | 0 | 1 | 1 | 0 | 1 | 1 |
| REXO2 | -0.158 | P1_P4_Lumbar_Stellate_Sphenopalatine_VS_P2_P3_Coeliac | 1 | 0 | 0 | 1 | 1 | 0 | 1 | 1 |
| NDUFV3 | -0.158 | P3_P4_Coeliac_Stellate_Sphenopalatine_VS_P1_P2_Lumbar | 0 | 0 | 1 | 1 | 0 | 1 | 1 | 1 |
| KIF5B | -0.158 | P3_P4_Coeliac_Stellate_Sphenopalatine_VS_P1_P2_Lumbar | 0 | 0 | 1 | 1 | 0 | 1 | 1 | 1 |
| RALY | -0.158 | P1_P3_P4_Coeliac_Stellate_Sphenopalatine_VS_P2_Lumbar | 1 | 0 | 1 | 1 | 0 | 1 | 1 | 1 |
| PPP3CA | -0.158 | P1_P2_P3_P4_Coeliac_Stellate_Sphenopalatine_VS_Lumbar | 1 | 1 | 1 | 1 | 0 | 1 | 1 | 1 |
| NECAP1 | -0.158 | P1_P2_P3_P4_Coeliac_Stellate_Sphenopalatine_VS_Lumbar | 1 | 1 | 1 | 1 | 0 | 1 | 1 | 1 |
| SCOC | -0.158 | P1_P2_P3_P4_Coeliac_Stellate_Sphenopalatine_VS_Lumbar | 1 | 1 | 1 | 1 | 0 | 1 | 1 | 1 |
| GDI2 | -0.158 | P1_P2_P3_P4_Coeliac_Stellate_Sphenopalatine_VS_Lumbar | 1 | 1 | 1 | 1 | 0 | 1 | 1 | 1 |
| GHITM | -0.158 | P1_P2_P3_P4_Coeliac_Stellate_Sphenopalatine_VS_Lumbar | 1 | 1 | 1 | 1 | 0 | 1 | 1 | 1 |
| SON | -0.158 | P1_P2_P3_P4_Coeliac_Stellate_Sphenopalatine_VS_Lumbar | 1 | 1 | 1 | 1 | 0 | 1 | 1 | 1 |
| CCDC6 | -0.158 | P2_P4_Lumbar_Coeliac_Stellate_Sphenopalatine_VS_P1_P3 | 0 | 1 | 0 | 1 | 1 | 1 | 1 | 1 |
| GANAB | -0.158 | P2_P4_Lumbar_Coeliac_Stellate_Sphenopalatine_VS_P1_P3 | 0 | 1 | 0 | 1 | 1 | 1 | 1 | 1 |
| PSMB2 | -0.159 | P3_P4_Stellate_VS_P1_P2_Lumbar_Coeliac_Sphenopalatine | 0 | 0 | 1 | 1 | 0 | 0 | 1 | 0 |
| PHB2 | -0.159 | P3_P4_Stellate_VS_P1_P2_Lumbar_Coeliac_Sphenopalatine | 0 | 0 | 1 | 1 | 0 | 0 | 1 | 0 |
| TAGLN3 | -0.159 | P3_P4_Stellate_VS_P1_P2_Lumbar_Coeliac_Sphenopalatine | 0 | 0 | 1 | 1 | 0 | 0 | 1 | 0 |
| PFDN1 | -0.159 | P3_P4_Stellate_VS_P1_P2_Lumbar_Coeliac_Sphenopalatine | 0 | 0 | 1 | 1 | 0 | 0 | 1 | 0 |
| BANF1 | -0.159 | P3_P4_Stellate_VS_P1_P2_Lumbar_Coeliac_Sphenopalatine | 0 | 0 | 1 | 1 | 0 | 0 | 1 | 0 |

| | | | | | | | | | | |
|---|---|---|---|---|---|---|---|---|---|---|
| BAG1 | -0.159 | P3_Stellate_Sphenopalatine_VS_P1_P2_P4_Lumbar_Coeliac | 0 | 0 | 1 | 0 | 0 | 0 | 1 | 1 |
| TRAPPC4 | -0.159 | P3_Stellate_Sphenopalatine_VS_P1_P2_P4_Lumbar_Coeliac | 0 | 0 | 1 | 0 | 0 | 0 | 1 | 1 |
| NAP1L1 | -0.159 | P3_Stellate_Sphenopalatine_VS_P1_P2_P4_Lumbar_Coeliac | 0 | 0 | 1 | 0 | 0 | 0 | 1 | 1 |
| BEX1 | -0.159 | P3_Stellate_Sphenopalatine_VS_P1_P2_P4_Lumbar_Coeliac | 0 | 0 | 1 | 0 | 0 | 0 | 1 | 1 |
| STMN4 | -0.159 | P1_P3_Stellate_Sphenopalatine_VS_P2_P4_Lumbar_Coeliac | 1 | 0 | 1 | 0 | 0 | 0 | 1 | 1 |
| MRPL52 | -0.159 | P2_P3_Stellate_Sphenopalatine_VS_P1_P4_Lumbar_Coeliac | 0 | 1 | 1 | 0 | 0 | 0 | 1 | 1 |
| NDUFA7 | -0.159 | P2_P3_Stellate_Sphenopalatine_VS_P1_P4_Lumbar_Coeliac | 0 | 1 | 1 | 0 | 0 | 0 | 1 | 1 |
| MAPK8IP1 | -0.159 | P1_P2_P3_Stellate_Sphenopalatine_VS_P4_Lumbar_Coeliac | 1 | 1 | 1 | 0 | 0 | 0 | 1 | 1 |
| NDUFS6 | -0.159 | P4_Stellate_Sphenopalatine_VS_P1_P2_P3_Lumbar_Coeliac | 0 | 0 | 0 | 1 | 0 | 0 | 1 | 1 |
| CRIP2 | -0.159 | P1_P4_Stellate_Sphenopalatine_VS_P2_P3_Lumbar_Coeliac | 1 | 0 | 0 | 1 | 0 | 0 | 1 | 1 |
| TMEM50A | -0.159 | P2_P4_Stellate_Sphenopalatine_VS_P1_P3_Lumbar_Coeliac | 0 | 1 | 0 | 1 | 0 | 0 | 1 | 1 |
| SDCBP | -0.159 | P1_P2_P4_Stellate_Sphenopalatine_VS_P3_Lumbar_Coeliac | 1 | 1 | 0 | 1 | 0 | 0 | 1 | 1 |
| ASNSD1 | -0.159 | P1_P3_Lumbar_Stellate_Sphenopalatine_VS_P2_P4_Coeliac | 1 | 0 | 1 | 0 | 1 | 0 | 1 | 1 |
| EIF1B | -0.159 | P1_P2_P3_Lumbar_Stellate_Sphenopalatine_VS_P4_Coeliac | 1 | 1 | 1 | 0 | 1 | 0 | 1 | 1 |
| R3HDM4 | -0.159 | P1_P2_P3_Lumbar_Stellate_Sphenopalatine_VS_P4_Coeliac | 1 | 1 | 1 | 0 | 1 | 0 | 1 | 1 |
| CD47 | -0.159 | P1_P2_P3_Lumbar_Stellate_Sphenopalatine_VS_P4_Coeliac | 1 | 1 | 1 | 0 | 1 | 0 | 1 | 1 |
| SYT4 | -0.159 | P1_P2_P3_Lumbar_Stellate_Sphenopalatine_VS_P4_Coeliac | 1 | 1 | 1 | 0 | 1 | 0 | 1 | 1 |
| RAB14 | -0.159 | P3_P4_Coeliac_Stellate_Sphenopalatine_VS_P1_P2_Lumbar | 0 | 0 | 1 | 1 | 0 | 1 | 1 | 1 |
| GNG5 | -0.159 | P3_P4_Coeliac_Stellate_Sphenopalatine_VS_P1_P2_Lumbar | 0 | 0 | 1 | 1 | 0 | 1 | 1 | 1 |
| VAPA | -0.159 | P1_P3_P4_Coeliac_Stellate_Sphenopalatine_VS_P2_Lumbar | 1 | 0 | 1 | 1 | 0 | 1 | 1 | 1 |
| NDUFS8 | -0.159 | P1_P3_P4_Coeliac_Stellate_Sphenopalatine_VS_P2_Lumbar | 1 | 0 | 1 | 1 | 0 | 1 | 1 | 1 |
| SEC61B | -0.159 | P2_P3_P4_Coeliac_Stellate_Sphenopalatine_VS_P1_Lumbar | 0 | 1 | 1 | 1 | 0 | 1 | 1 | 1 |
| UBE2E1 | -0.159 | P2_P3_P4_Coeliac_Stellate_Sphenopalatine_VS_P1_Lumbar | 0 | 1 | 1 | 1 | 0 | 1 | 1 | 1 |
| TRIR | -0.159 | P1_P2_P3_P4_Coeliac_Stellate_Sphenopalatine_VS_Lumbar | 1 | 1 | 1 | 1 | 0 | 1 | 1 | 1 |
| RALA | -0.159 | P1_P2_P3_P4_Coeliac_Stellate_Sphenopalatine_VS_Lumbar | 1 | 1 | 1 | 1 | 0 | 1 | 1 | 1 |
| PNN | -0.159 | P2_P3_Lumbar_Coeliac_Stellate_Sphenopalatine_VS_P1_P4 | 0 | 1 | 1 | 0 | 1 | 1 | 1 | 1 |
| MIA3 | -0.159 | P2_P4_Lumbar_Coeliac_Stellate_Sphenopalatine_VS_P1_P3 | 0 | 1 | 0 | 1 | 1 | 1 | 1 | 1 |
| INA | -0.16 | P4_Stellate_VS_P1_P2_P3_Lumbar_Coeliac_Sphenopalatine | 0 | 0 | 0 | 1 | 0 | 0 | 1 | 0 |
| CAPZB | -0.16 | P3_P4_Stellate_VS_P1_P2_Lumbar_Coeliac_Sphenopalatine | 0 | 0 | 1 | 1 | 0 | 0 | 1 | 0 |
| SPCS2 | -0.16 | P3_P4_Stellate_VS_P1_P2_Lumbar_Coeliac_Sphenopalatine | 0 | 0 | 1 | 1 | 0 | 0 | 1 | 0 |

| | | | | | | | | | | |
|---|---|---|---|---|---|---|---|---|---|---|
| UBE2B | -0.16 | P3_P4_Stellate_VS_P1_P2_Lumbar_Coeliac_Sphenopalatine | 0 | 0 | 1 | 1 | 0 | 0 | 1 | 0 |
| FAM89B | -0.16 | P3_P4_Stellate_VS_P1_P2_Lumbar_Coeliac_Sphenopalatine | 0 | 0 | 1 | 1 | 0 | 0 | 1 | 0 |
| CISD1 | -0.16 | P1_P3_P4_Stellate_VS_P2_Lumbar_Coeliac_Sphenopalatine | 1 | 0 | 1 | 1 | 0 | 0 | 1 | 0 |
| ARPC1A | -0.16 | P2_P3_P4_Stellate_VS_P1_Lumbar_Coeliac_Sphenopalatine | 0 | 1 | 1 | 1 | 0 | 0 | 1 | 0 |
| RBBP6 | -0.16 | P2_P3_P4_Lumbar_Stellate_VS_P1_Coeliac_Sphenopalatine | 0 | 1 | 1 | 1 | 1 | 0 | 1 | 0 |
| APBB1 | -0.16 | P3_Stellate_Sphenopalatine_VS_P1_P2_P4_Lumbar_Coeliac | 0 | 0 | 1 | 0 | 0 | 0 | 1 | 1 |
| ELAVL3 | -0.16 | P3_Stellate_Sphenopalatine_VS_P1_P2_P4_Lumbar_Coeliac | 0 | 0 | 1 | 0 | 0 | 0 | 1 | 1 |
| GLRX5 | -0.16 | P3_Stellate_Sphenopalatine_VS_P1_P2_P4_Lumbar_Coeliac | 0 | 0 | 1 | 0 | 0 | 0 | 1 | 1 |
| HNRNPA2B1 | -0.16 | P1_P3_Stellate_Sphenopalatine_VS_P2_P4_Lumbar_Coeliac | 1 | 0 | 1 | 0 | 0 | 0 | 1 | 1 |
| EIF3F | -0.16 | P2_P3_Stellate_Sphenopalatine_VS_P1_P4_Lumbar_Coeliac | 0 | 1 | 1 | 0 | 0 | 0 | 1 | 1 |
| APRT | -0.16 | P3_Lumbar_Stellate_Sphenopalatine_VS_P1_P2_P4_Coeliac | 0 | 0 | 1 | 0 | 1 | 0 | 1 | 1 |
| CRMP1 | -0.16 | P1_P2_P3_Lumbar_Stellate_Sphenopalatine_VS_P4_Coeliac | 1 | 1 | 1 | 0 | 1 | 0 | 1 | 1 |
| CSNK1A1 | -0.16 | P3_P4_Coeliac_Stellate_Sphenopalatine_VS_P1_P2_Lumbar | 0 | 0 | 1 | 1 | 0 | 1 | 1 | 1 |
| LSM4 | -0.16 | P1_P3_P4_Coeliac_Stellate_Sphenopalatine_VS_P2_Lumbar | 1 | 0 | 1 | 1 | 0 | 1 | 1 | 1 |
| HDGF | -0.16 | P2_P3_P4_Coeliac_Stellate_Sphenopalatine_VS_P1_Lumbar | 0 | 1 | 1 | 1 | 0 | 1 | 1 | 1 |
| THEM4 | -0.16 | P1_P2_P3_P4_Coeliac_Stellate_Sphenopalatine_VS_Lumbar | 1 | 1 | 1 | 1 | 0 | 1 | 1 | 1 |
| NAP1L5 | -0.16 | P1_P2_P3_P4_Coeliac_Stellate_Sphenopalatine_VS_Lumbar | 1 | 1 | 1 | 1 | 0 | 1 | 1 | 1 |
| SERBP1 | -0.16 | P1_P2_P3_P4_Coeliac_Stellate_Sphenopalatine_VS_Lumbar | 1 | 1 | 1 | 1 | 0 | 1 | 1 | 1 |
| ADGRL1 | -0.16 | P1_P2_P3_P4_Coeliac_Stellate_Sphenopalatine_VS_Lumbar | 1 | 1 | 1 | 1 | 0 | 1 | 1 | 1 |
| VDAC1 | -0.16 | P1_P2_P3_P4_Coeliac_Stellate_Sphenopalatine_VS_Lumbar | 1 | 1 | 1 | 1 | 0 | 1 | 1 | 1 |
| BC005537 | -0.16 | P1_P2_P3_P4_Coeliac_Stellate_Sphenopalatine_VS_Lumbar | 1 | 1 | 1 | 1 | 0 | 1 | 1 | 1 |
| NPDC1 | -0.161 | P2_P3_P4_Stellate_VS_P1_Lumbar_Coeliac_Sphenopalatine | 0 | 1 | 1 | 1 | 0 | 0 | 1 | 0 |
| RCHY1 | -0.161 | P2_P3_P4_Stellate_VS_P1_Lumbar_Coeliac_Sphenopalatine | 0 | 1 | 1 | 1 | 0 | 0 | 1 | 0 |
| CUEDC2 | -0.161 | P1_P3_Lumbar_Stellate_VS_P2_P4_Coeliac_Sphenopalatine | 1 | 0 | 1 | 0 | 1 | 0 | 1 | 0 |
| WDR18 | -0.161 | P4_Lumbar_Stellate_VS_P1_P2_P3_Coeliac_Sphenopalatine | 0 | 0 | 0 | 1 | 1 | 0 | 1 | 0 |
| EIF4E | -0.161 | P2_P3_P4_Lumbar_Stellate_VS_P1_Coeliac_Sphenopalatine | 0 | 1 | 1 | 1 | 1 | 0 | 1 | 0 |
| BNIP3L | -0.161 | P2_P3_P4_Lumbar_Stellate_VS_P1_Coeliac_Sphenopalatine | 0 | 1 | 1 | 1 | 1 | 0 | 1 | 0 |
| RAB6A | -0.161 | P3_Stellate_Sphenopalatine_VS_P1_P2_P4_Lumbar_Coeliac | 0 | 0 | 1 | 0 | 0 | 0 | 1 | 1 |
| NDUFA12 | -0.161 | P3_Stellate_Sphenopalatine_VS_P1_P2_P4_Lumbar_Coeliac | 0 | 0 | 1 | 0 | 0 | 0 | 1 | 1 |
| UBE2N | -0.161 | P1_P3_Stellate_Sphenopalatine_VS_P2_P4_Lumbar_Coeliac | 1 | 0 | 1 | 0 | 0 | 0 | 1 | 1 |

| | | | | | | | | | | |
|---|---|---|---|---|---|---|---|---|---|---|
| RNF10 | -0.161 | P2_P3_Stellate_Sphenopalatine_VS_P1_P4_Lumbar_Coeliac | 0 | 1 | 1 | 0 | 0 | 0 | 1 | 1 |
| GOT2 | -0.161 | P2_P3_Stellate_Sphenopalatine_VS_P1_P4_Lumbar_Coeliac | 0 | 1 | 1 | 0 | 0 | 0 | 1 | 1 |
| NDUFS2 | -0.161 | P1_P2_P3_Stellate_Sphenopalatine_VS_P4_Lumbar_Coeliac | 1 | 1 | 1 | 0 | 0 | 0 | 1 | 1 |
| UBE2S | -0.161 | P1_P2_P3_Stellate_Sphenopalatine_VS_P4_Lumbar_Coeliac | 1 | 1 | 1 | 0 | 0 | 0 | 1 | 1 |
| DYNLL2 | -0.161 | P1_P2_P3_Stellate_Sphenopalatine_VS_P4_Lumbar_Coeliac | 1 | 1 | 1 | 0 | 0 | 0 | 1 | 1 |
| NONO | -0.161 | P1_P2_P3_Stellate_Sphenopalatine_VS_P4_Lumbar_Coeliac | 1 | 1 | 1 | 0 | 0 | 0 | 1 | 1 |
| MRPL36 | -0.161 | P1_P3_Lumbar_Stellate_Sphenopalatine_VS_P2_P4_Coeliac | 1 | 0 | 1 | 0 | 1 | 0 | 1 | 1 |
| NNAT | -0.161 | P1_P2_P3_Lumbar_Stellate_Sphenopalatine_VS_P4_Coeliac | 1 | 1 | 1 | 0 | 1 | 0 | 1 | 1 |
| ATP6V1G1 | -0.161 | P1_P2_P3_Lumbar_Stellate_Sphenopalatine_VS_P4_Coeliac | 1 | 1 | 1 | 0 | 1 | 0 | 1 | 1 |
| ERP29 | -0.161 | P1_P2_P3_Lumbar_Stellate_Sphenopalatine_VS_P4_Coeliac | 1 | 1 | 1 | 0 | 1 | 0 | 1 | 1 |
| FXYD6 | -0.161 | P1_P2_P3_Lumbar_Stellate_Sphenopalatine_VS_P4_Coeliac | 1 | 1 | 1 | 0 | 1 | 0 | 1 | 1 |
| AURKAIP1 | -0.161 | P1_P4_Lumbar_Stellate_Sphenopalatine_VS_P2_P3_Coeliac | 1 | 0 | 0 | 1 | 1 | 0 | 1 | 1 |
| CYTH3 | -0.161 | P1_P4_Coeliac_Stellate_Sphenopalatine_VS_P2_P3_Lumbar | 1 | 0 | 0 | 1 | 0 | 1 | 1 | 1 |
| SRSF6 | -0.161 | P3_P4_Coeliac_Stellate_Sphenopalatine_VS_P1_P2_Lumbar | 0 | 0 | 1 | 1 | 0 | 1 | 1 | 1 |
| DPH3 | -0.161 | P1_P3_P4_Coeliac_Stellate_Sphenopalatine_VS_P2_Lumbar | 1 | 0 | 1 | 1 | 0 | 1 | 1 | 1 |
| ACOT7 | -0.161 | P2_P3_P4_Coeliac_Stellate_Sphenopalatine_VS_P1_Lumbar | 0 | 1 | 1 | 1 | 0 | 1 | 1 | 1 |
| RUFY3 | -0.161 | P2_P3_P4_Coeliac_Stellate_Sphenopalatine_VS_P1_Lumbar | 0 | 1 | 1 | 1 | 0 | 1 | 1 | 1 |
| GNG2 | -0.161 | P2_P3_P4_Coeliac_Stellate_Sphenopalatine_VS_P1_Lumbar | 0 | 1 | 1 | 1 | 0 | 1 | 1 | 1 |
| LSAMP | -0.161 | P2_P3_P4_Coeliac_Stellate_Sphenopalatine_VS_P1_Lumbar | 0 | 1 | 1 | 1 | 0 | 1 | 1 | 1 |
| RNPS1 | -0.161 | P2_P3_P4_Coeliac_Stellate_Sphenopalatine_VS_P1_Lumbar | 0 | 1 | 1 | 1 | 0 | 1 | 1 | 1 |
| BTBD10 | -0.161 | P1_Lumbar_Coeliac_Stellate_Sphenopalatine_VS_P2_P3_P4 | 1 | 0 | 0 | 0 | 1 | 1 | 1 | 1 |
| PTPRF | -0.161 | P2_P4_Lumbar_Coeliac_Stellate_Sphenopalatine_VS_P1_P3 | 0 | 1 | 0 | 1 | 1 | 1 | 1 | 1 |
| HSDL1 | -0.161 | P2_P4_Lumbar_Coeliac_Stellate_Sphenopalatine_VS_P1_P3 | 0 | 1 | 0 | 1 | 1 | 1 | 1 | 1 |
| ROMO1 | -0.162 | P2_P3_P4_Stellate_VS_P1_Lumbar_Coeliac_Sphenopalatine | 0 | 1 | 1 | 1 | 0 | 0 | 1 | 0 |
| CYB5R3 | -0.162 | P2_P3_P4_Stellate_VS_P1_Lumbar_Coeliac_Sphenopalatine | 0 | 1 | 1 | 1 | 0 | 0 | 1 | 0 |
| TCP1 | -0.162 | P2_P3_P4_Stellate_VS_P1_Lumbar_Coeliac_Sphenopalatine | 0 | 1 | 1 | 1 | 0 | 0 | 1 | 0 |
| PPP2R5B | -0.162 | P2_P3_P4_Stellate_VS_P1_Lumbar_Coeliac_Sphenopalatine | 0 | 1 | 1 | 1 | 0 | 0 | 1 | 0 |
| ZMYM5 | -0.162 | P2_P3_P4_Lumbar_Stellate_VS_P1_Coeliac_Sphenopalatine | 0 | 1 | 1 | 1 | 1 | 0 | 1 | 0 |
| PSMA6 | -0.162 | P1_P3_P4_Coeliac_Stellate_VS_P2_Lumbar_Sphenopalatine | 1 | 0 | 1 | 1 | 0 | 1 | 1 | 0 |
| EIF3K | -0.162 | P3_Stellate_Sphenopalatine_VS_P1_P2_P4_Lumbar_Coeliac | 0 | 0 | 1 | 0 | 0 | 0 | 1 | 1 |

| | | | | | | | | | | |
|---|---|---|---|---|---|---|---|---|---|---|
| CELF4 | -0.162 | P3_Stellate_Sphenopalatine_VS_P1_P2_P4_Lumbar_Coeliac | 0 | 0 | 1 | 0 | 0 | 0 | 1 | 1 |
| KRTCAP2 | -0.162 | P2_P3_Stellate_Sphenopalatine_VS_P1_P4_Lumbar_Coeliac | 0 | 1 | 1 | 0 | 0 | 0 | 1 | 1 |
| ATOX1 | -0.162 | P2_P3_Stellate_Sphenopalatine_VS_P1_P4_Lumbar_Coeliac | 0 | 1 | 1 | 0 | 0 | 0 | 1 | 1 |
| POLR2K | -0.162 | P2_P3_Stellate_Sphenopalatine_VS_P1_P4_Lumbar_Coeliac | 0 | 1 | 1 | 0 | 0 | 0 | 1 | 1 |
| BCAS2 | -0.162 | P4_Stellate_Sphenopalatine_VS_P1_P2_P3_Lumbar_Coeliac | 0 | 0 | 0 | 1 | 0 | 0 | 1 | 1 |
| EIF3H | -0.162 | P4_Stellate_Sphenopalatine_VS_P1_P2_P3_Lumbar_Coeliac | 0 | 0 | 0 | 1 | 0 | 0 | 1 | 1 |
| CLNS1A | -0.162 | P3_Lumbar_Stellate_Sphenopalatine_VS_P1_P2_P4_Coeliac | 0 | 0 | 1 | 0 | 1 | 0 | 1 | 1 |
| SNX10 | -0.162 | P1_P2_P3_Lumbar_Stellate_Sphenopalatine_VS_P4_Coeliac | 1 | 1 | 1 | 0 | 1 | 0 | 1 | 1 |
| LGALS1 | -0.162 | P1_P2_P3_Lumbar_Stellate_Sphenopalatine_VS_P4_Coeliac | 1 | 1 | 1 | 0 | 1 | 0 | 1 | 1 |
| NDUFA11 | -0.162 | P1_P2_P3_Lumbar_Stellate_Sphenopalatine_VS_P4_Coeliac | 1 | 1 | 1 | 0 | 1 | 0 | 1 | 1 |
| NDUFV2 | -0.162 | P1_P2_P3_Lumbar_Stellate_Sphenopalatine_VS_P4_Coeliac | 1 | 1 | 1 | 0 | 1 | 0 | 1 | 1 |
| MRPL43 | -0.162 | P1_P2_P3_Lumbar_Stellate_Sphenopalatine_VS_P4_Coeliac | 1 | 1 | 1 | 0 | 1 | 0 | 1 | 1 |
| UBE2W | -0.162 | P1_P4_Lumbar_Stellate_Sphenopalatine_VS_P2_P3_Coeliac | 1 | 0 | 0 | 1 | 1 | 0 | 1 | 1 |
| CERS4 | -0.162 | P2_P4_Lumbar_Stellate_Sphenopalatine_VS_P1_P3_Coeliac | 0 | 1 | 0 | 1 | 1 | 0 | 1 | 1 |
| MAPRE2 | -0.162 | P2_P4_Lumbar_Stellate_Sphenopalatine_VS_P1_P3_Coeliac | 0 | 1 | 0 | 1 | 1 | 0 | 1 | 1 |
| WDR7 | -0.162 | P2_P4_Lumbar_Stellate_Sphenopalatine_VS_P1_P3_Coeliac | 0 | 1 | 0 | 1 | 1 | 0 | 1 | 1 |
| PHOX2B | -0.162 | P1_P2_P3_Coeliac_Stellate_Sphenopalatine_VS_P4_Lumbar | 1 | 1 | 1 | 0 | 0 | 1 | 1 | 1 |
| DDAH2 | -0.162 | P1_P4_Coeliac_Stellate_Sphenopalatine_VS_P2_P3_Lumbar | 1 | 0 | 0 | 1 | 0 | 1 | 1 | 1 |
| THOC7 | -0.162 | P3_P4_Coeliac_Stellate_Sphenopalatine_VS_P1_P2_Lumbar | 0 | 0 | 1 | 1 | 0 | 1 | 1 | 1 |
| SRPK2 | -0.162 | P2_P3_P4_Coeliac_Stellate_Sphenopalatine_VS_P1_Lumbar | 0 | 1 | 1 | 1 | 0 | 1 | 1 | 1 |
| GNAQ | -0.162 | P2_P3_P4_Coeliac_Stellate_Sphenopalatine_VS_P1_Lumbar | 0 | 1 | 1 | 1 | 0 | 1 | 1 | 1 |
| MTX2 | -0.162 | P1_Lumbar_Coeliac_Stellate_Sphenopalatine_VS_P2_P3_P4 | 1 | 0 | 0 | 0 | 1 | 1 | 1 | 1 |
| HMG20A | -0.162 | P2_P4_Lumbar_Coeliac_Stellate_Sphenopalatine_VS_P1_P3 | 0 | 1 | 0 | 1 | 1 | 1 | 1 | 1 |
| DCUN1D5 | -0.163 | P3_Stellate_VS_P1_P2_P4_Lumbar_Coeliac_Sphenopalatine | 0 | 0 | 1 | 0 | 0 | 0 | 1 | 0 |
| ACAT1 | -0.163 | P1_P3_P4_Stellate_VS_P2_Lumbar_Coeliac_Sphenopalatine | 1 | 0 | 1 | 1 | 0 | 0 | 1 | 0 |
| COX7A2L | -0.163 | P1_P3_P4_Stellate_VS_P2_Lumbar_Coeliac_Sphenopalatine | 1 | 0 | 1 | 1 | 0 | 0 | 1 | 0 |
| PGRMC1 | -0.163 | P2_P3_P4_Lumbar_Stellate_VS_P1_Coeliac_Sphenopalatine | 0 | 1 | 1 | 1 | 1 | 0 | 1 | 0 |
| STXBP1 | -0.163 | P2_P3_Stellate_Sphenopalatine_VS_P1_P4_Lumbar_Coeliac | 0 | 1 | 1 | 0 | 0 | 0 | 1 | 1 |
| SERF2 | -0.163 | P2_P3_Stellate_Sphenopalatine_VS_P1_P4_Lumbar_Coeliac | 0 | 1 | 1 | 0 | 0 | 0 | 1 | 1 |
| SYT11 | -0.163 | P1_P2_P3_Stellate_Sphenopalatine_VS_P4_Lumbar_Coeliac | 1 | 1 | 1 | 0 | 0 | 0 | 1 | 1 |

| | | | | | | | | | | |
|---|---|---|---|---|---|---|---|---|---|---|
| CS | -0.163 | P1_P2_P3_Stellate_Sphenopalatine_VS_P4_Lumbar_Coeliac | 1 | 1 | 1 | 0 | 0 | 0 | 1 | 1 |
| NISCH | -0.163 | P1_P2_P3_Stellate_Sphenopalatine_VS_P4_Lumbar_Coeliac | 1 | 1 | 1 | 0 | 0 | 0 | 1 | 1 |
| SDHB | -0.163 | P1_P4_Stellate_Sphenopalatine_VS_P2_P3_Lumbar_Coeliac | 1 | 0 | 0 | 1 | 0 | 0 | 1 | 1 |
| TECR | -0.163 | P1_P2_P3_Lumbar_Stellate_Sphenopalatine_VS_P4_Coeliac | 1 | 1 | 1 | 0 | 1 | 0 | 1 | 1 |
| CALR | -0.163 | P1_P2_P3_Lumbar_Stellate_Sphenopalatine_VS_P4_Coeliac | 1 | 1 | 1 | 0 | 1 | 0 | 1 | 1 |
| SERPINB6A | -0.163 | P1_P2_P3_Lumbar_Stellate_Sphenopalatine_VS_P4_Coeliac | 1 | 1 | 1 | 0 | 1 | 0 | 1 | 1 |
| NDFIP2 | -0.163 | P1_P2_P3_Lumbar_Stellate_Sphenopalatine_VS_P4_Coeliac | 1 | 1 | 1 | 0 | 1 | 0 | 1 | 1 |
| FKBP2 | -0.163 | P2_P4_Lumbar_Stellate_Sphenopalatine_VS_P1_P3_Coeliac | 0 | 1 | 0 | 1 | 1 | 0 | 1 | 1 |
| RPS18 | -0.163 | P1_P2_P3_Coeliac_Stellate_Sphenopalatine_VS_P4_Lumbar | 1 | 1 | 1 | 0 | 0 | 1 | 1 | 1 |
| HCFC1R1 | -0.163 | P1_P3_P4_Coeliac_Stellate_Sphenopalatine_VS_P2_Lumbar | 1 | 0 | 1 | 1 | 0 | 1 | 1 | 1 |
| PSMA7 | -0.163 | P2_P3_P4_Coeliac_Stellate_Sphenopalatine_VS_P1_Lumbar | 0 | 1 | 1 | 1 | 0 | 1 | 1 | 1 |
| PSIP1 | -0.163 | P2_P3_P4_Coeliac_Stellate_Sphenopalatine_VS_P1_Lumbar | 0 | 1 | 1 | 1 | 0 | 1 | 1 | 1 |
| H2AFY | -0.163 | P2_P3_P4_Coeliac_Stellate_Sphenopalatine_VS_P1_Lumbar | 0 | 1 | 1 | 1 | 0 | 1 | 1 | 1 |
| ACBD5 | -0.163 | P2_P4_Lumbar_Coeliac_Stellate_Sphenopalatine_VS_P1_P3 | 0 | 1 | 0 | 1 | 1 | 1 | 1 | 1 |
| IK | -0.163 | P2_P4_Lumbar_Coeliac_Stellate_Sphenopalatine_VS_P1_P3 | 0 | 1 | 0 | 1 | 1 | 1 | 1 | 1 |
| GDE1 | -0.164 | P1_P3_P4_Stellate_VS_P2_Lumbar_Coeliac_Sphenopalatine | 1 | 0 | 1 | 1 | 0 | 0 | 1 | 0 |
| 4930455G09 | -0.164 | P1_Lumbar_Stellate_VS_P2_P3_P4_Coeliac_Sphenopalatine | 1 | 0 | 0 | 0 | 1 | 0 | 1 | 0 |
| CCNT2 | -0.164 | P2_P3_P4_Lumbar_Stellate_VS_P1_Coeliac_Sphenopalatine | 0 | 1 | 1 | 1 | 1 | 0 | 1 | 0 |
| PSMB7 | -0.164 | P2_P3_P4_Lumbar_Stellate_VS_P1_Coeliac_Sphenopalatine | 0 | 1 | 1 | 1 | 1 | 0 | 1 | 0 |
| COPS6 | -0.164 | P1_P3_P4_Coeliac_Stellate_VS_P2_Lumbar_Sphenopalatine | 1 | 0 | 1 | 1 | 0 | 1 | 1 | 0 |
| UQCR10 | -0.164 | Stellate_Sphenopalatine_VS_P1_P2_P3_P4_Lumbar_Coeliac | 0 | 0 | 0 | 0 | 0 | 0 | 1 | 1 |
| NDUFB4 | -0.164 | P1_P3_Stellate_Sphenopalatine_VS_P2_P4_Lumbar_Coeliac | 1 | 0 | 1 | 0 | 0 | 0 | 1 | 1 |
| MRPL12 | -0.164 | P2_P3_Stellate_Sphenopalatine_VS_P1_P4_Lumbar_Coeliac | 0 | 1 | 1 | 0 | 0 | 0 | 1 | 1 |
| POMP | -0.164 | P4_Stellate_Sphenopalatine_VS_P1_P2_P3_Lumbar_Coeliac | 0 | 0 | 0 | 1 | 0 | 0 | 1 | 1 |
| CYCS | -0.164 | P1_P4_Stellate_Sphenopalatine_VS_P2_P3_Lumbar_Coeliac | 1 | 0 | 0 | 1 | 0 | 0 | 1 | 1 |
| TSPAN17 | -0.164 | P2_P4_Stellate_Sphenopalatine_VS_P1_P3_Lumbar_Coeliac | 0 | 1 | 0 | 1 | 0 | 0 | 1 | 1 |
| DBN1 | -0.164 | P1_P2_P3_Lumbar_Stellate_Sphenopalatine_VS_P4_Coeliac | 1 | 1 | 1 | 0 | 1 | 0 | 1 | 1 |
| SRSF3 | -0.164 | P1_P2_P3_Lumbar_Stellate_Sphenopalatine_VS_P4_Coeliac | 1 | 1 | 1 | 0 | 1 | 0 | 1 | 1 |
| MYL12B | -0.164 | P1_P2_P3_Lumbar_Stellate_Sphenopalatine_VS_P4_Coeliac | 1 | 1 | 1 | 0 | 1 | 0 | 1 | 1 |
| CYSTM1 | -0.164 | P1_P2_P3_Lumbar_Stellate_Sphenopalatine_VS_P4_Coeliac | 1 | 1 | 1 | 0 | 1 | 0 | 1 | 1 |

| | | | | | | | | | | |
|---|---|---|---|---|---|---|---|---|---|---|
| TSC22D3 | -0.164 | P4_Lumbar_Stellate_Sphenopalatine_VS_P1_P2_P3_Coeliac | 0 | 0 | 0 | 1 | 1 | 0 | 1 | 1 |
| DBI | -0.164 | P1_P4_Lumbar_Stellate_Sphenopalatine_VS_P2_P3_Coeliac | 1 | 0 | 0 | 1 | 1 | 0 | 1 | 1 |
| CCT6A | -0.164 | P1_P4_Lumbar_Stellate_Sphenopalatine_VS_P2_P3_Coeliac | 1 | 0 | 0 | 1 | 1 | 0 | 1 | 1 |
| UBE2E2 | -0.164 | P1_P4_Lumbar_Stellate_Sphenopalatine_VS_P2_P3_Coeliac | 1 | 0 | 0 | 1 | 1 | 0 | 1 | 1 |
| NDUFB5 | -0.164 | P2_P4_Lumbar_Stellate_Sphenopalatine_VS_P1_P3_Coeliac | 0 | 1 | 0 | 1 | 1 | 0 | 1 | 1 |
| KIF5C | -0.164 | P2_P3_Coeliac_Stellate_Sphenopalatine_VS_P1_P4_Lumbar | 0 | 1 | 1 | 0 | 0 | 1 | 1 | 1 |
| MARCKSL1 | -0.164 | P2_P3_Coeliac_Stellate_Sphenopalatine_VS_P1_P4_Lumbar | 0 | 1 | 1 | 0 | 0 | 1 | 1 | 1 |
| RPL12 | -0.164 | P1_P2_P3_Coeliac_Stellate_Sphenopalatine_VS_P4_Lumbar | 1 | 1 | 1 | 0 | 0 | 1 | 1 | 1 |
| SELENOH | -0.164 | P3_P4_Coeliac_Stellate_Sphenopalatine_VS_P1_P2_Lumbar | 0 | 0 | 1 | 1 | 0 | 1 | 1 | 1 |
| MKRN1 | -0.164 | P3_P4_Coeliac_Stellate_Sphenopalatine_VS_P1_P2_Lumbar | 0 | 0 | 1 | 1 | 0 | 1 | 1 | 1 |
| SNRPD2 | -0.164 | P3_P4_Coeliac_Stellate_Sphenopalatine_VS_P1_P2_Lumbar | 0 | 0 | 1 | 1 | 0 | 1 | 1 | 1 |
| CAPZA2 | -0.164 | P1_P3_P4_Coeliac_Stellate_Sphenopalatine_VS_P2_Lumbar | 1 | 0 | 1 | 1 | 0 | 1 | 1 | 1 |
| SNRPE | -0.164 | P2_P3_P4_Coeliac_Stellate_Sphenopalatine_VS_P1_Lumbar | 0 | 1 | 1 | 1 | 0 | 1 | 1 | 1 |
| NAV1 | -0.164 | P2_P3_P4_Coeliac_Stellate_Sphenopalatine_VS_P1_Lumbar | 0 | 1 | 1 | 1 | 0 | 1 | 1 | 1 |
| EPN1 | -0.164 | P2_P3_P4_Coeliac_Stellate_Sphenopalatine_VS_P1_Lumbar | 0 | 1 | 1 | 1 | 0 | 1 | 1 | 1 |
| NSA2 | -0.164 | P2_P3_P4_Coeliac_Stellate_Sphenopalatine_VS_P1_Lumbar | 0 | 1 | 1 | 1 | 0 | 1 | 1 | 1 |
| sept-03 | -0.164 | P2_P3_Lumbar_Coeliac_Stellate_Sphenopalatine_VS_P1_P4 | 0 | 1 | 1 | 0 | 1 | 1 | 1 | 1 |
| ATP6V1F | -0.165 | P1_P3_Stellate_Sphenopalatine_VS_P2_P4_Lumbar_Coeliac | 1 | 0 | 1 | 0 | 0 | 0 | 1 | 1 |
| MAPK3 | -0.165 | P2_P3_Stellate_Sphenopalatine_VS_P1_P4_Lumbar_Coeliac | 0 | 1 | 1 | 0 | 0 | 0 | 1 | 1 |
| OST4 | -0.165 | P4_Stellate_Sphenopalatine_VS_P1_P2_P3_Lumbar_Coeliac | 0 | 0 | 0 | 1 | 0 | 0 | 1 | 1 |
| RBX1 | -0.165 | P4_Stellate_Sphenopalatine_VS_P1_P2_P3_Lumbar_Coeliac | 0 | 0 | 0 | 1 | 0 | 0 | 1 | 1 |
| MRPS24 | -0.165 | P2_P4_Stellate_Sphenopalatine_VS_P1_P3_Lumbar_Coeliac | 0 | 1 | 0 | 1 | 0 | 0 | 1 | 1 |
| PCBP4 | -0.165 | P2_P3_Lumbar_Stellate_Sphenopalatine_VS_P1_P4_Coeliac | 0 | 1 | 1 | 0 | 1 | 0 | 1 | 1 |
| 4933434E20I | -0.165 | P1_P2_P3_Lumbar_Stellate_Sphenopalatine_VS_P4_Coeliac | 1 | 1 | 1 | 0 | 1 | 0 | 1 | 1 |
| RER1 | -0.165 | P1_P2_P3_Lumbar_Stellate_Sphenopalatine_VS_P4_Coeliac | 1 | 1 | 1 | 0 | 1 | 0 | 1 | 1 |
| DUT | -0.165 | P4_Lumbar_Stellate_Sphenopalatine_VS_P1_P2_P3_Coeliac | 0 | 0 | 0 | 1 | 1 | 0 | 1 | 1 |
| MED10 | -0.165 | P4_Lumbar_Stellate_Sphenopalatine_VS_P1_P2_P3_Coeliac | 0 | 0 | 0 | 1 | 1 | 0 | 1 | 1 |
| PRDX5 | -0.165 | P1_P4_Lumbar_Stellate_Sphenopalatine_VS_P2_P3_Coeliac | 1 | 0 | 0 | 1 | 1 | 0 | 1 | 1 |
| DENR | -0.165 | P3_P4_Coeliac_Stellate_Sphenopalatine_VS_P1_P2_Lumbar | 0 | 0 | 1 | 1 | 0 | 1 | 1 | 1 |
| FDFT1 | -0.165 | P1_P3_P4_Coeliac_Stellate_Sphenopalatine_VS_P2_Lumbar | 1 | 0 | 1 | 1 | 0 | 1 | 1 | 1 |

| | | | | | | | | | | |
|---|---|---|---|---|---|---|---|---|---|---|
| SRP14 | -0.165 | P2_P3_P4_Coeliac_Stellate_Sphenopalatine_VS_P1_Lumbar | 0 | 1 | 1 | 1 | 0 | 1 | 1 | 1 |
| RPL22 | -0.165 | P2_P3_P4_Coeliac_Stellate_Sphenopalatine_VS_P1_Lumbar | 0 | 1 | 1 | 1 | 0 | 1 | 1 | 1 |
| G3BP2 | -0.165 | P2_P3_P4_Coeliac_Stellate_Sphenopalatine_VS_P1_Lumbar | 0 | 1 | 1 | 1 | 0 | 1 | 1 | 1 |
| LAMTOR1 | -0.165 | P2_P3_P4_Coeliac_Stellate_Sphenopalatine_VS_P1_Lumbar | 0 | 1 | 1 | 1 | 0 | 1 | 1 | 1 |
| ILK | -0.165 | P2_P3_P4_Coeliac_Stellate_Sphenopalatine_VS_P1_Lumbar | 0 | 1 | 1 | 1 | 0 | 1 | 1 | 1 |
| ATP6V1B2 | -0.165 | P2_P3_P4_Coeliac_Stellate_Sphenopalatine_VS_P1_Lumbar | 0 | 1 | 1 | 1 | 0 | 1 | 1 | 1 |
| TLN2 | -0.165 | P2_P3_P4_Coeliac_Stellate_Sphenopalatine_VS_P1_Lumbar | 0 | 1 | 1 | 1 | 0 | 1 | 1 | 1 |
| HNRNPAB | -0.165 | P2_P3_P4_Coeliac_Stellate_Sphenopalatine_VS_P1_Lumbar | 0 | 1 | 1 | 1 | 0 | 1 | 1 | 1 |
| ACP1 | -0.165 | P2_P3_P4_Coeliac_Stellate_Sphenopalatine_VS_P1_Lumbar | 0 | 1 | 1 | 1 | 0 | 1 | 1 | 1 |
| RAF1 | -0.165 | P1_Lumbar_Coeliac_Stellate_Sphenopalatine_VS_P2_P3_P4 | 1 | 0 | 0 | 0 | 1 | 1 | 1 | 1 |
| MARCH7 | -0.165 | P2_P4_Lumbar_Coeliac_Stellate_Sphenopalatine_VS_P1_P3 | 0 | 1 | 0 | 1 | 1 | 1 | 1 | 1 |
| AHCYL1 | -0.165 | P2_P4_Lumbar_Coeliac_Stellate_Sphenopalatine_VS_P1_P3 | 0 | 1 | 0 | 1 | 1 | 1 | 1 | 1 |
| ZFP318 | -0.165 | P2_P4_Lumbar_Coeliac_Stellate_Sphenopalatine_VS_P1_P3 | 0 | 1 | 0 | 1 | 1 | 1 | 1 | 1 |
| NCAM1 | -0.166 | P3_Stellate_VS_P1_P2_P4_Lumbar_Coeliac_Sphenopalatine | 0 | 0 | 1 | 0 | 0 | 0 | 1 | 0 |
| TBCB | -0.166 | P1_P3_P4_Stellate_VS_P2_Lumbar_Coeliac_Sphenopalatine | 1 | 0 | 1 | 1 | 0 | 0 | 1 | 0 |
| TOMM20 | -0.166 | P1_P3_P4_Stellate_VS_P2_Lumbar_Coeliac_Sphenopalatine | 1 | 0 | 1 | 1 | 0 | 0 | 1 | 0 |
| PSMA2 | -0.166 | P1_P3_P4_Stellate_VS_P2_Lumbar_Coeliac_Sphenopalatine | 1 | 0 | 1 | 1 | 0 | 0 | 1 | 0 |
| CHCHD3 | -0.166 | P3_P4_Coeliac_Stellate_VS_P1_P2_Lumbar_Sphenopalatine | 0 | 0 | 1 | 1 | 0 | 1 | 1 | 0 |
| MRPS12 | -0.166 | P1_P3_Stellate_Sphenopalatine_VS_P2_P4_Lumbar_Coeliac | 1 | 0 | 1 | 0 | 0 | 0 | 1 | 1 |
| TSG101 | -0.166 | P4_Stellate_Sphenopalatine_VS_P1_P2_P3_Lumbar_Coeliac | 0 | 0 | 0 | 1 | 0 | 0 | 1 | 1 |
| PSMB6 | -0.166 | P4_Stellate_Sphenopalatine_VS_P1_P2_P3_Lumbar_Coeliac | 0 | 0 | 0 | 1 | 0 | 0 | 1 | 1 |
| MRPL20 | -0.166 | P2_P3_Lumbar_Stellate_Sphenopalatine_VS_P1_P4_Coeliac | 0 | 1 | 1 | 0 | 1 | 0 | 1 | 1 |
| NME2 | -0.166 | P2_P3_Lumbar_Stellate_Sphenopalatine_VS_P1_P4_Coeliac | 0 | 1 | 1 | 0 | 1 | 0 | 1 | 1 |
| CHMP5 | -0.166 | P4_Lumbar_Stellate_Sphenopalatine_VS_P1_P2_P3_Coeliac | 0 | 0 | 0 | 1 | 1 | 0 | 1 | 1 |
| NDUFS7 | -0.166 | P1_P4_Lumbar_Stellate_Sphenopalatine_VS_P2_P3_Coeliac | 1 | 0 | 0 | 1 | 1 | 0 | 1 | 1 |
| RNASEH2C | -0.166 | P1_P4_Lumbar_Stellate_Sphenopalatine_VS_P2_P3_Coeliac | 1 | 0 | 0 | 1 | 1 | 0 | 1 | 1 |
| SWI5 | -0.166 | P3_P4_Coeliac_Stellate_Sphenopalatine_VS_P1_P2_Lumbar | 0 | 0 | 1 | 1 | 0 | 1 | 1 | 1 |
| BZW2 | -0.166 | P3_P4_Coeliac_Stellate_Sphenopalatine_VS_P1_P2_Lumbar | 0 | 0 | 1 | 1 | 0 | 1 | 1 | 1 |
| PSMB5 | -0.166 | P3_P4_Coeliac_Stellate_Sphenopalatine_VS_P1_P2_Lumbar | 0 | 0 | 1 | 1 | 0 | 1 | 1 | 1 |
| TBCA | -0.166 | P1_P3_P4_Coeliac_Stellate_Sphenopalatine_VS_P2_Lumbar | 1 | 0 | 1 | 1 | 0 | 1 | 1 | 1 |

| | | | | | | | | | | |
|---|---|---|---|---|---|---|---|---|---|---|
| NRAS | -0.166 | P2_P3_P4_Coeliac_Stellate_Sphenopalatine_VS_P1_Lumbar | 0 | 1 | 1 | 1 | 0 | 1 | 1 | 1 |
| TMA7 | -0.166 | P2_P3_P4_Coeliac_Stellate_Sphenopalatine_VS_P1_Lumbar | 0 | 1 | 1 | 1 | 0 | 1 | 1 | 1 |
| THRA | -0.166 | P2_P3_P4_Coeliac_Stellate_Sphenopalatine_VS_P1_Lumbar | 0 | 1 | 1 | 1 | 0 | 1 | 1 | 1 |
| ATP5G2 | -0.166 | P2_P3_P4_Coeliac_Stellate_Sphenopalatine_VS_P1_Lumbar | 0 | 1 | 1 | 1 | 0 | 1 | 1 | 1 |
| SRP19 | -0.166 | P2_P3_P4_Coeliac_Stellate_Sphenopalatine_VS_P1_Lumbar | 0 | 1 | 1 | 1 | 0 | 1 | 1 | 1 |
| PDCD5 | -0.166 | P2_P3_Lumbar_Coeliac_Stellate_Sphenopalatine_VS_P1_P4 | 0 | 1 | 1 | 0 | 1 | 1 | 1 | 1 |
| SMIM13 | -0.166 | P2_P4_Lumbar_Coeliac_Stellate_Sphenopalatine_VS_P1_P3 | 0 | 1 | 0 | 1 | 1 | 1 | 1 | 1 |
| 1810058I24R | -0.167 | P3_Stellate_VS_P1_P2_P4_Lumbar_Coeliac_Sphenopalatine | 0 | 0 | 1 | 0 | 0 | 0 | 1 | 0 |
| ADRM1 | -0.167 | P1_P4_Lumbar_Stellate_VS_P2_P3_Coeliac_Sphenopalatine | 1 | 0 | 0 | 1 | 1 | 0 | 1 | 0 |
| CLIC1 | -0.167 | P1_P2_P3_P4_Coeliac_Stellate_VS_Lumbar_Sphenopalatine | 1 | 1 | 1 | 1 | 0 | 1 | 1 | 0 |
| UBE2D3 | -0.167 | Stellate_Sphenopalatine_VS_P1_P2_P3_P4_Lumbar_Coeliac | 0 | 0 | 0 | 0 | 0 | 0 | 1 | 1 |
| ELOC | -0.167 | P4_Stellate_Sphenopalatine_VS_P1_P2_P3_Lumbar_Coeliac | 0 | 0 | 0 | 1 | 0 | 0 | 1 | 1 |
| MRPL33 | -0.167 | P4_Stellate_Sphenopalatine_VS_P1_P2_P3_Lumbar_Coeliac | 0 | 0 | 0 | 1 | 0 | 0 | 1 | 1 |
| HSBP1 | -0.167 | P4_Stellate_Sphenopalatine_VS_P1_P2_P3_Lumbar_Coeliac | 0 | 0 | 0 | 1 | 0 | 0 | 1 | 1 |
| MIF | -0.167 | P4_Stellate_Sphenopalatine_VS_P1_P2_P3_Lumbar_Coeliac | 0 | 0 | 0 | 1 | 0 | 0 | 1 | 1 |
| MDH2 | -0.167 | P2_P4_Stellate_Sphenopalatine_VS_P1_P3_Lumbar_Coeliac | 0 | 1 | 0 | 1 | 0 | 0 | 1 | 1 |
| PRKAR1B | -0.167 | P2_P4_Stellate_Sphenopalatine_VS_P1_P3_Lumbar_Coeliac | 0 | 1 | 0 | 1 | 0 | 0 | 1 | 1 |
| NDUFB2 | -0.167 | P2_P4_Stellate_Sphenopalatine_VS_P1_P3_Lumbar_Coeliac | 0 | 1 | 0 | 1 | 0 | 0 | 1 | 1 |
| NDUFC2 | -0.167 | P2_P4_Stellate_Sphenopalatine_VS_P1_P3_Lumbar_Coeliac | 0 | 1 | 0 | 1 | 0 | 0 | 1 | 1 |
| NIPSNAP3B | -0.167 | P1_P3_Lumbar_Stellate_Sphenopalatine_VS_P2_P4_Coeliac | 1 | 0 | 1 | 0 | 1 | 0 | 1 | 1 |
| LEPROT | -0.167 | P1_P3_Lumbar_Stellate_Sphenopalatine_VS_P2_P4_Coeliac | 1 | 0 | 1 | 0 | 1 | 0 | 1 | 1 |
| MT-ND5 | -0.167 | P2_P3_Lumbar_Stellate_Sphenopalatine_VS_P1_P4_Coeliac | 0 | 1 | 1 | 0 | 1 | 0 | 1 | 1 |
| DCTN3 | -0.167 | P4_Lumbar_Stellate_Sphenopalatine_VS_P1_P2_P3_Coeliac | 0 | 0 | 0 | 1 | 1 | 0 | 1 | 1 |
| CYB5A | -0.167 | P4_Lumbar_Stellate_Sphenopalatine_VS_P1_P2_P3_Coeliac | 0 | 0 | 0 | 1 | 1 | 0 | 1 | 1 |
| CHCHD7 | -0.167 | P1_P4_Lumbar_Stellate_Sphenopalatine_VS_P2_P3_Coeliac | 1 | 0 | 0 | 1 | 1 | 0 | 1 | 1 |
| SLC35B1 | -0.167 | P1_P4_Lumbar_Stellate_Sphenopalatine_VS_P2_P3_Coeliac | 1 | 0 | 0 | 1 | 1 | 0 | 1 | 1 |
| CALM3 | -0.167 | P1_P2_P3_Coeliac_Stellate_Sphenopalatine_VS_P4_Lumbar | 1 | 1 | 1 | 0 | 0 | 1 | 1 | 1 |
| TCF25 | -0.167 | P3_P4_Coeliac_Stellate_Sphenopalatine_VS_P1_P2_Lumbar | 0 | 0 | 1 | 1 | 0 | 1 | 1 | 1 |
| RBM25 | -0.167 | P3_P4_Coeliac_Stellate_Sphenopalatine_VS_P1_P2_Lumbar | 0 | 0 | 1 | 1 | 0 | 1 | 1 | 1 |
| ZBTB20 | -0.167 | P3_P4_Coeliac_Stellate_Sphenopalatine_VS_P1_P2_Lumbar | 0 | 0 | 1 | 1 | 0 | 1 | 1 | 1 |

| | | | | | | | | | | |
|---|---|---|---|---|---|---|---|---|---|---|
| SRSF11 | -0.167 | P1_P3_P4_Coeliac_Stellate_Sphenopalatine_VS_P2_Lumbar | 1 | 0 | 1 | 1 | 0 | 1 | 1 | 1 |
| FUS | -0.167 | P1_P3_P4_Coeliac_Stellate_Sphenopalatine_VS_P2_Lumbar | 1 | 0 | 1 | 1 | 0 | 1 | 1 | 1 |
| C1QBP | -0.167 | P1_P3_P4_Coeliac_Stellate_Sphenopalatine_VS_P2_Lumbar | 1 | 0 | 1 | 1 | 0 | 1 | 1 | 1 |
| RPL31 | -0.167 | P2_P3_P4_Coeliac_Stellate_Sphenopalatine_VS_P1_Lumbar | 0 | 1 | 1 | 1 | 0 | 1 | 1 | 1 |
| EIF3M | -0.167 | P2_P3_P4_Coeliac_Stellate_Sphenopalatine_VS_P1_Lumbar | 0 | 1 | 1 | 1 | 0 | 1 | 1 | 1 |
| PHACTR3 | -0.167 | P2_P3_P4_Coeliac_Stellate_Sphenopalatine_VS_P1_Lumbar | 0 | 1 | 1 | 1 | 0 | 1 | 1 | 1 |
| PSMA5 | -0.167 | P2_P3_P4_Coeliac_Stellate_Sphenopalatine_VS_P1_Lumbar | 0 | 1 | 1 | 1 | 0 | 1 | 1 | 1 |
| DNAJC8 | -0.167 | P2_P3_P4_Coeliac_Stellate_Sphenopalatine_VS_P1_Lumbar | 0 | 1 | 1 | 1 | 0 | 1 | 1 | 1 |
| UBL5 | -0.167 | P2_P3_P4_Coeliac_Stellate_Sphenopalatine_VS_P1_Lumbar | 0 | 1 | 1 | 1 | 0 | 1 | 1 | 1 |
| B230219D22 | -0.167 | P2_P3_P4_Coeliac_Stellate_Sphenopalatine_VS_P1_Lumbar | 0 | 1 | 1 | 1 | 0 | 1 | 1 | 1 |
| BRD2 | -0.167 | P2_P3_P4_Coeliac_Stellate_Sphenopalatine_VS_P1_Lumbar | 0 | 1 | 1 | 1 | 0 | 1 | 1 | 1 |
| TXNRD1 | -0.167 | P1_Lumbar_Coeliac_Stellate_Sphenopalatine_VS_P2_P3_P4 | 1 | 0 | 0 | 0 | 1 | 1 | 1 | 1 |
| PLXNA4 | -0.167 | P2_P3_Lumbar_Coeliac_Stellate_Sphenopalatine_VS_P1_P4 | 0 | 1 | 1 | 0 | 1 | 1 | 1 | 1 |
| JUP | -0.167 | P2_P3_Lumbar_Coeliac_Stellate_Sphenopalatine_VS_P1_P4 | 0 | 1 | 1 | 0 | 1 | 1 | 1 | 1 |
| RCAN2 | -0.168 | P2_P3_Lumbar_Stellate_VS_P1_P4_Coeliac_Sphenopalatine | 0 | 1 | 1 | 0 | 1 | 0 | 1 | 0 |
| ATG4B | -0.168 | P2_P4_Lumbar_Stellate_VS_P1_P3_Coeliac_Sphenopalatine | 0 | 1 | 0 | 1 | 1 | 0 | 1 | 0 |
| XBP1 | -0.168 | P3_P4_Coeliac_Stellate_VS_P1_P2_Lumbar_Sphenopalatine | 0 | 0 | 1 | 1 | 0 | 1 | 1 | 0 |
| NEFL | -0.168 | P1_P2_P3_P4_Coeliac_Stellate_VS_Lumbar_Sphenopalatine | 1 | 1 | 1 | 1 | 0 | 1 | 1 | 0 |
| NDUFC1 | -0.168 | P1_P4_Stellate_Sphenopalatine_VS_P2_P3_Lumbar_Coeliac | 1 | 0 | 0 | 1 | 0 | 0 | 1 | 1 |
| CHCHD1 | -0.168 | P2_P3_Lumbar_Stellate_Sphenopalatine_VS_P1_P4_Coeliac | 0 | 1 | 1 | 0 | 1 | 0 | 1 | 1 |
| MRPL58 | -0.168 | P4_Lumbar_Stellate_Sphenopalatine_VS_P1_P2_P3_Coeliac | 0 | 0 | 0 | 1 | 1 | 0 | 1 | 1 |
| ATP6V0B | -0.168 | P1_P4_Lumbar_Stellate_Sphenopalatine_VS_P2_P3_Coeliac | 1 | 0 | 0 | 1 | 1 | 0 | 1 | 1 |
| NDUFAB1 | -0.168 | P1_P4_Lumbar_Stellate_Sphenopalatine_VS_P2_P3_Coeliac | 1 | 0 | 0 | 1 | 1 | 0 | 1 | 1 |
| EMC9 | -0.168 | P1_P4_Lumbar_Stellate_Sphenopalatine_VS_P2_P3_Coeliac | 1 | 0 | 0 | 1 | 1 | 0 | 1 | 1 |
| GNG3 | -0.168 | P1_P2_P3_Coeliac_Stellate_Sphenopalatine_VS_P4_Lumbar | 1 | 1 | 1 | 0 | 0 | 1 | 1 | 1 |
| MAPRE1 | -0.168 | P3_P4_Coeliac_Stellate_Sphenopalatine_VS_P1_P2_Lumbar | 0 | 0 | 1 | 1 | 0 | 1 | 1 | 1 |
| PPP1CA | -0.168 | P3_P4_Coeliac_Stellate_Sphenopalatine_VS_P1_P2_Lumbar | 0 | 0 | 1 | 1 | 0 | 1 | 1 | 1 |
| ZC3H15 | -0.168 | P1_P3_P4_Coeliac_Stellate_Sphenopalatine_VS_P2_Lumbar | 1 | 0 | 1 | 1 | 0 | 1 | 1 | 1 |
| HNRNPK | -0.168 | P1_P3_P4_Coeliac_Stellate_Sphenopalatine_VS_P2_Lumbar | 1 | 0 | 1 | 1 | 0 | 1 | 1 | 1 |
| PSMB3 | -0.168 | P2_P3_P4_Coeliac_Stellate_Sphenopalatine_VS_P1_Lumbar | 0 | 1 | 1 | 1 | 0 | 1 | 1 | 1 |

| | | | | | | | | | | |
|---|---|---|---|---|---|---|---|---|---|---|
| H3F3B | -0.168 | P2_P3_P4_Coeliac_Stellate_Sphenopalatine_VS_P1_Lumbar | 0 | 1 | 1 | 1 | 0 | 1 | 1 | 1 |
| RPL36AL | -0.168 | P2_P3_P4_Coeliac_Stellate_Sphenopalatine_VS_P1_Lumbar | 0 | 1 | 1 | 1 | 0 | 1 | 1 | 1 |
| TMED10 | -0.168 | P2_P3_P4_Coeliac_Stellate_Sphenopalatine_VS_P1_Lumbar | 0 | 1 | 1 | 1 | 0 | 1 | 1 | 1 |
| TOMM6 | -0.168 | P2_P3_P4_Coeliac_Stellate_Sphenopalatine_VS_P1_Lumbar | 0 | 1 | 1 | 1 | 0 | 1 | 1 | 1 |
| NRXN2 | -0.168 | P2_P3_P4_Coeliac_Stellate_Sphenopalatine_VS_P1_Lumbar | 0 | 1 | 1 | 1 | 0 | 1 | 1 | 1 |
| ATP5J2 | -0.168 | P2_P3_Lumbar_Coeliac_Stellate_Sphenopalatine_VS_P1_P4 | 0 | 1 | 1 | 0 | 1 | 1 | 1 | 1 |
| GAPVD1 | -0.168 | P2_P4_Lumbar_Coeliac_Stellate_Sphenopalatine_VS_P1_P3 | 0 | 1 | 0 | 1 | 1 | 1 | 1 | 1 |
| HNRNPA1 | -0.169 | P1_P3_Stellate_VS_P2_P4_Lumbar_Coeliac_Sphenopalatine | 1 | 0 | 1 | 0 | 0 | 0 | 1 | 0 |
| CAR11 | -0.169 | P2_P3_Lumbar_Stellate_VS_P1_P4_Coeliac_Sphenopalatine | 0 | 1 | 1 | 0 | 1 | 0 | 1 | 0 |
| HNRNPL | -0.169 | P3_P4_Coeliac_Stellate_VS_P1_P2_Lumbar_Sphenopalatine | 0 | 0 | 1 | 1 | 0 | 1 | 1 | 0 |
| TPM4 | -0.169 | P3_P4_Coeliac_Stellate_VS_P1_P2_Lumbar_Sphenopalatine | 0 | 0 | 1 | 1 | 0 | 1 | 1 | 0 |
| CACYBP | -0.169 | P1_P3_P4_Coeliac_Stellate_VS_P2_Lumbar_Sphenopalatine | 1 | 0 | 1 | 1 | 0 | 1 | 1 | 0 |
| FIS1 | -0.169 | Stellate_Sphenopalatine_VS_P1_P2_P3_P4_Lumbar_Coeliac | 0 | 0 | 0 | 0 | 0 | 0 | 1 | 1 |
| UQCRB | -0.169 | Stellate_Sphenopalatine_VS_P1_P2_P3_P4_Lumbar_Coeliac | 0 | 0 | 0 | 0 | 0 | 0 | 1 | 1 |
| AP2M1 | -0.169 | Stellate_Sphenopalatine_VS_P1_P2_P3_P4_Lumbar_Coeliac | 0 | 0 | 0 | 0 | 0 | 0 | 1 | 1 |
| NDUFA10 | -0.169 | P1_P4_Stellate_Sphenopalatine_VS_P2_P3_Lumbar_Coeliac | 1 | 0 | 0 | 1 | 0 | 0 | 1 | 1 |
| MBD3 | -0.169 | P2_P3_Lumbar_Stellate_Sphenopalatine_VS_P1_P4_Coeliac | 0 | 1 | 1 | 0 | 1 | 0 | 1 | 1 |
| JKAMP | -0.169 | P2_P3_Lumbar_Stellate_Sphenopalatine_VS_P1_P4_Coeliac | 0 | 1 | 1 | 0 | 1 | 0 | 1 | 1 |
| NDUFB9 | -0.169 | P2_P3_Lumbar_Stellate_Sphenopalatine_VS_P1_P4_Coeliac | 0 | 1 | 1 | 0 | 1 | 0 | 1 | 1 |
| MICOS13 | -0.169 | P2_P3_Lumbar_Stellate_Sphenopalatine_VS_P1_P4_Coeliac | 0 | 1 | 1 | 0 | 1 | 0 | 1 | 1 |
| MT-ND2 | -0.169 | P2_P3_Lumbar_Stellate_Sphenopalatine_VS_P1_P4_Coeliac | 0 | 1 | 1 | 0 | 1 | 0 | 1 | 1 |
| NDUFB3 | -0.169 | P4_Lumbar_Stellate_Sphenopalatine_VS_P1_P2_P3_Coeliac | 0 | 0 | 0 | 1 | 1 | 0 | 1 | 1 |
| NT5M | -0.169 | P1_P4_Lumbar_Stellate_Sphenopalatine_VS_P2_P3_Coeliac | 1 | 0 | 0 | 1 | 1 | 0 | 1 | 1 |
| GSK3B | -0.169 | P2_P3_Coeliac_Stellate_Sphenopalatine_VS_P1_P4_Lumbar | 0 | 1 | 1 | 0 | 0 | 1 | 1 | 1 |
| MAPK10 | -0.169 | P1_P2_P4_Coeliac_Stellate_Sphenopalatine_VS_P3_Lumbar | 1 | 1 | 0 | 1 | 0 | 1 | 1 | 1 |
| U2AF1 | -0.169 | P3_P4_Coeliac_Stellate_Sphenopalatine_VS_P1_P2_Lumbar | 0 | 0 | 1 | 1 | 0 | 1 | 1 | 1 |
| ARL3 | -0.169 | P3_P4_Coeliac_Stellate_Sphenopalatine_VS_P1_P2_Lumbar | 0 | 0 | 1 | 1 | 0 | 1 | 1 | 1 |
| RNF187 | -0.169 | P1_P3_P4_Coeliac_Stellate_Sphenopalatine_VS_P2_Lumbar | 1 | 0 | 1 | 1 | 0 | 1 | 1 | 1 |
| KIF1B | -0.169 | P2_P3_P4_Coeliac_Stellate_Sphenopalatine_VS_P1_Lumbar | 0 | 1 | 1 | 1 | 0 | 1 | 1 | 1 |
| GLRX3 | -0.169 | P2_P3_P4_Coeliac_Stellate_Sphenopalatine_VS_P1_Lumbar | 0 | 1 | 1 | 1 | 0 | 1 | 1 | 1 |

| | | | | | | | | | | |
|---|---|---|---|---|---|---|---|---|---|---|
| RPL29 | -0.169 | P2_P3_P4_Coeliac_Stellate_Sphenopalatine_VS_P1_Lumbar | 0 | 1 | 1 | 1 | 0 | 1 | 1 | 1 |
| ARHGDIA | -0.169 | P2_P3_P4_Coeliac_Stellate_Sphenopalatine_VS_P1_Lumbar | 0 | 1 | 1 | 1 | 0 | 1 | 1 | 1 |
| PBRM1 | -0.169 | P2_P3_P4_Coeliac_Stellate_Sphenopalatine_VS_P1_Lumbar | 0 | 1 | 1 | 1 | 0 | 1 | 1 | 1 |
| 0610012G03 | -0.169 | P2_P3_P4_Coeliac_Stellate_Sphenopalatine_VS_P1_Lumbar | 0 | 1 | 1 | 1 | 0 | 1 | 1 | 1 |
| NDUFA1 | -0.169 | P2_P3_P4_Coeliac_Stellate_Sphenopalatine_VS_P1_Lumbar | 0 | 1 | 1 | 1 | 0 | 1 | 1 | 1 |
| ITGB1 | -0.169 | P2_P3_Lumbar_Coeliac_Stellate_Sphenopalatine_VS_P1_P4 | 0 | 1 | 1 | 0 | 1 | 1 | 1 | 1 |
| CARMIL3 | -0.169 | P2_P3_Lumbar_Coeliac_Stellate_Sphenopalatine_VS_P1_P4 | 0 | 1 | 1 | 0 | 1 | 1 | 1 | 1 |
| COX7B | -0.169 | P2_P3_Lumbar_Coeliac_Stellate_Sphenopalatine_VS_P1_P4 | 0 | 1 | 1 | 0 | 1 | 1 | 1 | 1 |
| YTHDF2 | -0.169 | P2_P4_Lumbar_Coeliac_Stellate_Sphenopalatine_VS_P1_P3 | 0 | 1 | 0 | 1 | 1 | 1 | 1 | 1 |
| PRKAR2A | -0.169 | P2_P4_Lumbar_Coeliac_Stellate_Sphenopalatine_VS_P1_P3 | 0 | 1 | 0 | 1 | 1 | 1 | 1 | 1 |
| TSPYL4 | -0.169 | P2_P4_Lumbar_Coeliac_Stellate_Sphenopalatine_VS_P1_P3 | 0 | 1 | 0 | 1 | 1 | 1 | 1 | 1 |
| SMARCD1 | -0.169 | P2_P4_Lumbar_Coeliac_Stellate_Sphenopalatine_VS_P1_P3 | 0 | 1 | 0 | 1 | 1 | 1 | 1 | 1 |
| HSPA13 | -0.169 | P2_P4_Lumbar_Coeliac_Stellate_Sphenopalatine_VS_P1_P3 | 0 | 1 | 0 | 1 | 1 | 1 | 1 | 1 |
| NDUFB7 | -0.17 | P1_P3_Stellate_VS_P2_P4_Lumbar_Coeliac_Sphenopalatine | 1 | 0 | 1 | 0 | 0 | 0 | 1 | 0 |
| GPX4 | -0.17 | P1_P3_Stellate_VS_P2_P4_Lumbar_Coeliac_Sphenopalatine | 1 | 0 | 1 | 0 | 0 | 0 | 1 | 0 |
| CDC42 | -0.17 | P1_P2_P3_Stellate_VS_P4_Lumbar_Coeliac_Sphenopalatine | 1 | 1 | 1 | 0 | 0 | 0 | 1 | 0 |
| SLC35A4 | -0.17 | P1_P2_P3_Stellate_VS_P4_Lumbar_Coeliac_Sphenopalatine | 1 | 1 | 1 | 0 | 0 | 0 | 1 | 0 |
| PFN1 | -0.17 | P4_Stellate_VS_P1_P2_P3_Lumbar_Coeliac_Sphenopalatine | 0 | 0 | 0 | 1 | 0 | 0 | 1 | 0 |
| KHDRBS1 | -0.17 | P3_P4_Coeliac_Stellate_VS_P1_P2_Lumbar_Sphenopalatine | 0 | 0 | 1 | 1 | 0 | 1 | 1 | 0 |
| 1110004F10I | -0.17 | Stellate_Sphenopalatine_VS_P1_P2_P3_P4_Lumbar_Coeliac | 0 | 0 | 0 | 0 | 0 | 0 | 1 | 1 |
| TIMM13 | -0.17 | Stellate_Sphenopalatine_VS_P1_P2_P3_P4_Lumbar_Coeliac | 0 | 0 | 0 | 0 | 0 | 0 | 1 | 1 |
| H13 | -0.17 | P2_P3_Lumbar_Stellate_Sphenopalatine_VS_P1_P4_Coeliac | 0 | 1 | 1 | 0 | 1 | 0 | 1 | 1 |
| CHN1 | -0.17 | P4_Lumbar_Stellate_Sphenopalatine_VS_P1_P2_P3_Coeliac | 0 | 0 | 0 | 1 | 1 | 0 | 1 | 1 |
| SF3B5 | -0.17 | P4_Lumbar_Stellate_Sphenopalatine_VS_P1_P2_P3_Coeliac | 0 | 0 | 0 | 1 | 1 | 0 | 1 | 1 |
| NME1 | -0.17 | P4_Lumbar_Stellate_Sphenopalatine_VS_P1_P2_P3_Coeliac | 0 | 0 | 0 | 1 | 1 | 0 | 1 | 1 |
| POLR2G | -0.17 | P4_Lumbar_Stellate_Sphenopalatine_VS_P1_P2_P3_Coeliac | 0 | 0 | 0 | 1 | 1 | 0 | 1 | 1 |
| SRSF5 | -0.17 | P2_P4_Lumbar_Stellate_Sphenopalatine_VS_P1_P3_Coeliac | 0 | 1 | 0 | 1 | 1 | 0 | 1 | 1 |
| ITM2B | -0.17 | P2_P4_Lumbar_Stellate_Sphenopalatine_VS_P1_P3_Coeliac | 0 | 1 | 0 | 1 | 1 | 0 | 1 | 1 |
| DDX5 | -0.17 | P1_P3_Coeliac_Stellate_Sphenopalatine_VS_P2_P4_Lumbar | 1 | 0 | 1 | 0 | 0 | 1 | 1 | 1 |
| NDUFA2 | -0.17 | P2_P3_Coeliac_Stellate_Sphenopalatine_VS_P1_P4_Lumbar | 0 | 1 | 1 | 0 | 0 | 1 | 1 | 1 |

| | | | | | | | | | | |
|---|---|---|---|---|---|---|---|---|---|---|
| ARF1 | -0.17 | P1_P2_P3_Coeliac_Stellate_Sphenopalatine_VS_P4_Lumbar | 1 | 1 | 1 | 0 | 0 | 1 | 1 | 1 |
| YWHAQ | -0.17 | P1_P2_P3_Coeliac_Stellate_Sphenopalatine_VS_P4_Lumbar | 1 | 1 | 1 | 0 | 0 | 1 | 1 | 1 |
| RIAN | -0.17 | P1_P2_P3_Coeliac_Stellate_Sphenopalatine_VS_P4_Lumbar | 1 | 1 | 1 | 0 | 0 | 1 | 1 | 1 |
| MTPN | -0.17 | P3_P4_Coeliac_Stellate_Sphenopalatine_VS_P1_P2_Lumbar | 0 | 0 | 1 | 1 | 0 | 1 | 1 | 1 |
| PTOV1 | -0.17 | P3_P4_Coeliac_Stellate_Sphenopalatine_VS_P1_P2_Lumbar | 0 | 0 | 1 | 1 | 0 | 1 | 1 | 1 |
| PARK7 | -0.17 | P1_P3_P4_Coeliac_Stellate_Sphenopalatine_VS_P2_Lumbar | 1 | 0 | 1 | 1 | 0 | 1 | 1 | 1 |
| KIF21A | -0.17 | P1_P3_P4_Coeliac_Stellate_Sphenopalatine_VS_P2_Lumbar | 1 | 0 | 1 | 1 | 0 | 1 | 1 | 1 |
| EEF1B2 | -0.17 | P2_P3_P4_Coeliac_Stellate_Sphenopalatine_VS_P1_Lumbar | 0 | 1 | 1 | 1 | 0 | 1 | 1 | 1 |
| SNAP25 | -0.17 | P2_P3_P4_Coeliac_Stellate_Sphenopalatine_VS_P1_Lumbar | 0 | 1 | 1 | 1 | 0 | 1 | 1 | 1 |
| CSDE1 | -0.17 | P2_P3_P4_Coeliac_Stellate_Sphenopalatine_VS_P1_Lumbar | 0 | 1 | 1 | 1 | 0 | 1 | 1 | 1 |
| SELENOF | -0.17 | P2_P3_P4_Coeliac_Stellate_Sphenopalatine_VS_P1_Lumbar | 0 | 1 | 1 | 1 | 0 | 1 | 1 | 1 |
| SDC3 | -0.17 | P2_P3_P4_Coeliac_Stellate_Sphenopalatine_VS_P1_Lumbar | 0 | 1 | 1 | 1 | 0 | 1 | 1 | 1 |
| MICOS10 | -0.17 | P2_P3_P4_Coeliac_Stellate_Sphenopalatine_VS_P1_Lumbar | 0 | 1 | 1 | 1 | 0 | 1 | 1 | 1 |
| EIF4G2 | -0.17 | P2_P3_P4_Coeliac_Stellate_Sphenopalatine_VS_P1_Lumbar | 0 | 1 | 1 | 1 | 0 | 1 | 1 | 1 |
| TMOD2 | -0.17 | P2_P3_P4_Coeliac_Stellate_Sphenopalatine_VS_P1_Lumbar | 0 | 1 | 1 | 1 | 0 | 1 | 1 | 1 |
| SNAP91 | -0.17 | P2_P3_P4_Coeliac_Stellate_Sphenopalatine_VS_P1_Lumbar | 0 | 1 | 1 | 1 | 0 | 1 | 1 | 1 |
| LARP4B | -0.17 | P2_P3_P4_Coeliac_Stellate_Sphenopalatine_VS_P1_Lumbar | 0 | 1 | 1 | 1 | 0 | 1 | 1 | 1 |
| CCSER2 | -0.17 | P2_P3_P4_Coeliac_Stellate_Sphenopalatine_VS_P1_Lumbar | 0 | 1 | 1 | 1 | 0 | 1 | 1 | 1 |
| EIF3A | -0.17 | P2_P3_P4_Coeliac_Stellate_Sphenopalatine_VS_P1_Lumbar | 0 | 1 | 1 | 1 | 0 | 1 | 1 | 1 |
| NDUFB11 | -0.17 | P2_P3_P4_Coeliac_Stellate_Sphenopalatine_VS_P1_Lumbar | 0 | 1 | 1 | 1 | 0 | 1 | 1 | 1 |
| ERGIC2 | -0.17 | P1_Lumbar_Coeliac_Stellate_Sphenopalatine_VS_P2_P3_P4 | 1 | 0 | 0 | 0 | 1 | 1 | 1 | 1 |
| PLSCR3 | -0.17 | P2_P3_Lumbar_Coeliac_Stellate_Sphenopalatine_VS_P1_P4 | 0 | 1 | 1 | 0 | 1 | 1 | 1 | 1 |
| MT-CYTB | -0.17 | P2_P3_Lumbar_Coeliac_Stellate_Sphenopalatine_VS_P1_P4 | 0 | 1 | 1 | 0 | 1 | 1 | 1 | 1 |
| RC3H2 | -0.17 | P2_P4_Lumbar_Coeliac_Stellate_Sphenopalatine_VS_P1_P3 | 0 | 1 | 0 | 1 | 1 | 1 | 1 | 1 |
| GTF2A1 | -0.17 | P2_P4_Lumbar_Coeliac_Stellate_Sphenopalatine_VS_P1_P3 | 0 | 1 | 0 | 1 | 1 | 1 | 1 | 1 |
| CGGBP1 | -0.17 | P2_P4_Lumbar_Coeliac_Stellate_Sphenopalatine_VS_P1_P3 | 0 | 1 | 0 | 1 | 1 | 1 | 1 | 1 |
| NDUFA8 | -0.171 | P1_P3_Stellate_VS_P2_P4_Lumbar_Coeliac_Sphenopalatine | 1 | 0 | 1 | 0 | 0 | 0 | 1 | 0 |
| CFL2 | -0.171 | P2_P3_Stellate_VS_P1_P4_Lumbar_Coeliac_Sphenopalatine | 0 | 1 | 1 | 0 | 0 | 0 | 1 | 0 |
| TTC3 | -0.171 | P4_Stellate_VS_P1_P2_P3_Lumbar_Coeliac_Sphenopalatine | 0 | 0 | 0 | 1 | 0 | 0 | 1 | 0 |
| RPP14 | -0.171 | P1_P3_Lumbar_Stellate_VS_P2_P4_Coeliac_Sphenopalatine | 1 | 0 | 1 | 0 | 1 | 0 | 1 | 0 |

| | | | | | | | | | | |
|---|---|---|---|---|---|---|---|---|---|---|
| SUMO2 | -0.171 | Stellate_Sphenopalatine_VS_P1_P2_P3_P4_Lumbar_Coeliac | 0 | 0 | 0 | 0 | 0 | 0 | 1 | 1 |
| REEP1 | -0.171 | P2_P4_Lumbar_Stellate_Sphenopalatine_VS_P1_P3_Coeliac | 0 | 1 | 0 | 1 | 1 | 0 | 1 | 1 |
| ELAVL2 | -0.171 | P3_Coeliac_Stellate_Sphenopalatine_VS_P1_P2_P4_Lumbar | 0 | 0 | 1 | 0 | 0 | 1 | 1 | 1 |
| RPL18 | -0.171 | P1_P2_P3_Coeliac_Stellate_Sphenopalatine_VS_P4_Lumbar | 1 | 1 | 1 | 0 | 0 | 1 | 1 | 1 |
| NDUFB1-PS | -0.171 | P1_P2_P3_Coeliac_Stellate_Sphenopalatine_VS_P4_Lumbar | 1 | 1 | 1 | 0 | 0 | 1 | 1 | 1 |
| RPL10 | -0.171 | P1_P2_P3_Coeliac_Stellate_Sphenopalatine_VS_P4_Lumbar | 1 | 1 | 1 | 0 | 0 | 1 | 1 | 1 |
| SCN9A | -0.171 | P3_P4_Coeliac_Stellate_Sphenopalatine_VS_P1_P2_Lumbar | 0 | 0 | 1 | 1 | 0 | 1 | 1 | 1 |
| BSG | -0.171 | P1_P3_P4_Coeliac_Stellate_Sphenopalatine_VS_P2_Lumbar | 1 | 0 | 1 | 1 | 0 | 1 | 1 | 1 |
| SNRPD1 | -0.171 | P1_P3_P4_Coeliac_Stellate_Sphenopalatine_VS_P2_Lumbar | 1 | 0 | 1 | 1 | 0 | 1 | 1 | 1 |
| ARPC2 | -0.171 | P2_P3_P4_Coeliac_Stellate_Sphenopalatine_VS_P1_Lumbar | 0 | 1 | 1 | 1 | 0 | 1 | 1 | 1 |
| DSTN | -0.171 | P2_P3_P4_Coeliac_Stellate_Sphenopalatine_VS_P1_Lumbar | 0 | 1 | 1 | 1 | 0 | 1 | 1 | 1 |
| AKR1A1 | -0.171 | P2_P3_P4_Coeliac_Stellate_Sphenopalatine_VS_P1_Lumbar | 0 | 1 | 1 | 1 | 0 | 1 | 1 | 1 |
| EVL | -0.171 | P2_P3_P4_Coeliac_Stellate_Sphenopalatine_VS_P1_Lumbar | 0 | 1 | 1 | 1 | 0 | 1 | 1 | 1 |
| FABP5 | -0.171 | P2_P3_Lumbar_Coeliac_Stellate_Sphenopalatine_VS_P1_P4 | 0 | 1 | 1 | 0 | 1 | 1 | 1 | 1 |
| CNOT4 | -0.171 | P2_P4_Lumbar_Coeliac_Stellate_Sphenopalatine_VS_P1_P3 | 0 | 1 | 0 | 1 | 1 | 1 | 1 | 1 |
| SRGAP3 | -0.171 | P2_P4_Lumbar_Coeliac_Stellate_Sphenopalatine_VS_P1_P3 | 0 | 1 | 0 | 1 | 1 | 1 | 1 | 1 |
| UQCR11 | -0.172 | P1_P3_Stellate_VS_P2_P4_Lumbar_Coeliac_Sphenopalatine | 1 | 0 | 1 | 0 | 0 | 0 | 1 | 0 |
| PKM | -0.172 | P1_P2_P3_Stellate_VS_P4_Lumbar_Coeliac_Sphenopalatine | 1 | 1 | 1 | 0 | 0 | 0 | 1 | 0 |
| SNCG | -0.172 | P1_P2_P3_Stellate_VS_P4_Lumbar_Coeliac_Sphenopalatine | 1 | 1 | 1 | 0 | 0 | 0 | 1 | 0 |
| MORF4L2 | -0.172 | P4_Stellate_VS_P1_P2_P3_Lumbar_Coeliac_Sphenopalatine | 0 | 0 | 0 | 1 | 0 | 0 | 1 | 0 |
| CXXC4 | -0.172 | P1_P2_P4_Lumbar_Stellate_VS_P3_Coeliac_Sphenopalatine | 1 | 1 | 0 | 1 | 1 | 0 | 1 | 0 |
| MAP2 | -0.172 | P1_P2_P3_P4_Coeliac_Stellate_VS_Lumbar_Sphenopalatine | 1 | 1 | 1 | 1 | 0 | 1 | 1 | 0 |
| GM1673 | -0.172 | Stellate_Sphenopalatine_VS_P1_P2_P3_P4_Lumbar_Coeliac | 0 | 0 | 0 | 0 | 0 | 0 | 1 | 1 |
| NDUFS5 | -0.172 | Lumbar_Stellate_Sphenopalatine_VS_P1_P2_P3_P4_Coeliac | 0 | 0 | 0 | 0 | 1 | 0 | 1 | 1 |
| ATP1B1 | -0.172 | P2_P4_Lumbar_Stellate_Sphenopalatine_VS_P1_P3_Coeliac | 0 | 1 | 0 | 1 | 1 | 0 | 1 | 1 |
| DPYSL5 | -0.172 | P2_P4_Lumbar_Stellate_Sphenopalatine_VS_P1_P3_Coeliac | 0 | 1 | 0 | 1 | 1 | 0 | 1 | 1 |
| BRK1 | -0.172 | P2_P4_Lumbar_Stellate_Sphenopalatine_VS_P1_P3_Coeliac | 0 | 1 | 0 | 1 | 1 | 0 | 1 | 1 |
| ARF4 | -0.172 | P2_P4_Lumbar_Stellate_Sphenopalatine_VS_P1_P3_Coeliac | 0 | 1 | 0 | 1 | 1 | 0 | 1 | 1 |
| CLTA | -0.172 | P3_Coeliac_Stellate_Sphenopalatine_VS_P1_P2_P4_Lumbar | 0 | 0 | 1 | 0 | 0 | 1 | 1 | 1 |
| RPS13 | -0.172 | P1_P2_P3_Coeliac_Stellate_Sphenopalatine_VS_P4_Lumbar | 1 | 1 | 1 | 0 | 0 | 1 | 1 | 1 |

| | | | | | | | | | | |
|---|---|---|---|---|---|---|---|---|---|---|
| VAT1 | -0.172 | P1_P2_P3_Coeliac_Stellate_Sphenopalatine_VS_P4_Lumbar | 1 | 1 | 1 | 0 | 0 | 1 | 1 | 1 |
| GTF2H5 | -0.172 | P4_Coeliac_Stellate_Sphenopalatine_VS_P1_P2_P3_Lumbar | 0 | 0 | 0 | 1 | 0 | 1 | 1 | 1 |
| STOML2 | -0.172 | P1_P3_P4_Coeliac_Stellate_Sphenopalatine_VS_P2_Lumbar | 1 | 0 | 1 | 1 | 0 | 1 | 1 | 1 |
| GABARAPL2 | -0.172 | P1_P3_P4_Coeliac_Stellate_Sphenopalatine_VS_P2_Lumbar | 1 | 0 | 1 | 1 | 0 | 1 | 1 | 1 |
| MORF4L1 | -0.172 | P1_P3_P4_Coeliac_Stellate_Sphenopalatine_VS_P2_Lumbar | 1 | 0 | 1 | 1 | 0 | 1 | 1 | 1 |
| UBE2L3 | -0.172 | P1_P3_P4_Coeliac_Stellate_Sphenopalatine_VS_P2_Lumbar | 1 | 0 | 1 | 1 | 0 | 1 | 1 | 1 |
| ALG2 | -0.172 | P2_P3_Lumbar_Coeliac_Stellate_Sphenopalatine_VS_P1_P4 | 0 | 1 | 1 | 0 | 1 | 1 | 1 | 1 |
| RPL30 | -0.172 | P2_P3_Lumbar_Coeliac_Stellate_Sphenopalatine_VS_P1_P4 | 0 | 1 | 1 | 0 | 1 | 1 | 1 | 1 |
| HP1BP3 | -0.172 | P2_P4_Lumbar_Coeliac_Stellate_Sphenopalatine_VS_P1_P3 | 0 | 1 | 0 | 1 | 1 | 1 | 1 | 1 |
| RERE | -0.172 | P2_P4_Lumbar_Coeliac_Stellate_Sphenopalatine_VS_P1_P3 | 0 | 1 | 0 | 1 | 1 | 1 | 1 | 1 |
| ZFP512 | -0.172 | P2_P4_Lumbar_Coeliac_Stellate_Sphenopalatine_VS_P1_P3 | 0 | 1 | 0 | 1 | 1 | 1 | 1 | 1 |
| COPB2 | -0.172 | P2_P4_Lumbar_Coeliac_Stellate_Sphenopalatine_VS_P1_P3 | 0 | 1 | 0 | 1 | 1 | 1 | 1 | 1 |
| VMP1 | -0.172 | P2_P4_Lumbar_Coeliac_Stellate_Sphenopalatine_VS_P1_P3 | 0 | 1 | 0 | 1 | 1 | 1 | 1 | 1 |
| AZIN1 | -0.172 | P2_P4_Lumbar_Coeliac_Stellate_Sphenopalatine_VS_P1_P3 | 0 | 1 | 0 | 1 | 1 | 1 | 1 | 1 |
| TMEM63B | -0.172 | P2_P4_Lumbar_Coeliac_Stellate_Sphenopalatine_VS_P1_P3 | 0 | 1 | 0 | 1 | 1 | 1 | 1 | 1 |
| ATP5K | -0.173 | P2_P3_Stellate_VS_P1_P4_Lumbar_Coeliac_Sphenopalatine | 0 | 1 | 1 | 0 | 0 | 0 | 1 | 0 |
| RAN | -0.173 | P2_P3_Stellate_VS_P1_P4_Lumbar_Coeliac_Sphenopalatine | 0 | 1 | 1 | 0 | 0 | 0 | 1 | 0 |
| CD24A | -0.173 | P2_P3_Stellate_VS_P1_P4_Lumbar_Coeliac_Sphenopalatine | 0 | 1 | 1 | 0 | 0 | 0 | 1 | 0 |
| NACA | -0.173 | P2_P3_Stellate_VS_P1_P4_Lumbar_Coeliac_Sphenopalatine | 0 | 1 | 1 | 0 | 0 | 0 | 1 | 0 |
| RABAC1 | -0.173 | P1_P2_P3_Stellate_VS_P4_Lumbar_Coeliac_Sphenopalatine | 1 | 1 | 1 | 0 | 0 | 0 | 1 | 0 |
| MAP1LC3B | -0.173 | P1_P2_P3_Stellate_VS_P4_Lumbar_Coeliac_Sphenopalatine | 1 | 1 | 1 | 0 | 0 | 0 | 1 | 0 |
| RTN3 | -0.173 | P1_P2_P3_Stellate_VS_P4_Lumbar_Coeliac_Sphenopalatine | 1 | 1 | 1 | 0 | 0 | 0 | 1 | 0 |
| NDUFA5 | -0.173 | P4_Stellate_VS_P1_P2_P3_Lumbar_Coeliac_Sphenopalatine | 0 | 0 | 0 | 1 | 0 | 0 | 1 | 0 |
| BTF3 | -0.173 | P4_Stellate_VS_P1_P2_P3_Lumbar_Coeliac_Sphenopalatine | 0 | 0 | 0 | 1 | 0 | 0 | 1 | 0 |
| SELENOK | -0.173 | P4_Stellate_VS_P1_P2_P3_Lumbar_Coeliac_Sphenopalatine | 0 | 0 | 0 | 1 | 0 | 0 | 1 | 0 |
| NEDD8 | -0.173 | P4_Stellate_VS_P1_P2_P3_Lumbar_Coeliac_Sphenopalatine | 0 | 0 | 0 | 1 | 0 | 0 | 1 | 0 |
| MT-ND4 | -0.173 | P2_P3_Lumbar_Stellate_VS_P1_P4_Coeliac_Sphenopalatine | 0 | 1 | 1 | 0 | 1 | 0 | 1 | 0 |
| TUBB4B | -0.173 | P1_P2_P4_Lumbar_Stellate_VS_P3_Coeliac_Sphenopalatine | 1 | 1 | 0 | 1 | 1 | 0 | 1 | 0 |
| HMGB1 | -0.173 | Stellate_Sphenopalatine_VS_P1_P2_P3_P4_Lumbar_Coeliac | 0 | 0 | 0 | 0 | 0 | 0 | 1 | 1 |
| CCT2 | -0.173 | P1_P2_P3_Coeliac_Stellate_Sphenopalatine_VS_P4_Lumbar | 1 | 1 | 1 | 0 | 0 | 1 | 1 | 1 |

| | | | | | | | | | | |
|---|---|---|---|---|---|---|---|---|---|---|
| ELAVL4 | -0.173 | P1_P3_P4_Coeliac_Stellate_Sphenopalatine_VS_P2_Lumbar | 1 | 0 | 1 | 1 | 0 | 1 | 1 | 1 |
| VPS72 | -0.173 | P1_Lumbar_Coeliac_Stellate_Sphenopalatine_VS_P2_P3_P4 | 1 | 0 | 0 | 0 | 1 | 1 | 1 | 1 |
| GRIPAP1 | -0.173 | P2_P4_Lumbar_Coeliac_Stellate_Sphenopalatine_VS_P1_P3 | 0 | 1 | 0 | 1 | 1 | 1 | 1 | 1 |
| DYNLRB1 | -0.174 | P2_P3_Stellate_VS_P1_P4_Lumbar_Coeliac_Sphenopalatine | 0 | 1 | 1 | 0 | 0 | 0 | 1 | 0 |
| ATP5E | -0.174 | P2_P3_Stellate_VS_P1_P4_Lumbar_Coeliac_Sphenopalatine | 0 | 1 | 1 | 0 | 0 | 0 | 1 | 0 |
| UQCRH | -0.174 | P2_P3_Stellate_VS_P1_P4_Lumbar_Coeliac_Sphenopalatine | 0 | 1 | 1 | 0 | 0 | 0 | 1 | 0 |
| PFN2 | -0.174 | P1_P2_P3_Stellate_VS_P4_Lumbar_Coeliac_Sphenopalatine | 1 | 1 | 1 | 0 | 0 | 0 | 1 | 0 |
| CAPNS1 | -0.174 | P1_P2_P3_Stellate_VS_P4_Lumbar_Coeliac_Sphenopalatine | 1 | 1 | 1 | 0 | 0 | 0 | 1 | 0 |
| GRINA | -0.174 | P1_P2_P3_Stellate_VS_P4_Lumbar_Coeliac_Sphenopalatine | 1 | 1 | 1 | 0 | 0 | 0 | 1 | 0 |
| PAIP2 | -0.174 | P1_P2_P3_Stellate_VS_P4_Lumbar_Coeliac_Sphenopalatine | 1 | 1 | 1 | 0 | 0 | 0 | 1 | 0 |
| RPL4 | -0.174 | P4_Stellate_VS_P1_P2_P3_Lumbar_Coeliac_Sphenopalatine | 0 | 0 | 0 | 1 | 0 | 0 | 1 | 0 |
| SEC61G | -0.174 | P4_Stellate_VS_P1_P2_P3_Lumbar_Coeliac_Sphenopalatine | 0 | 0 | 0 | 1 | 0 | 0 | 1 | 0 |
| EIF4A2 | -0.174 | P2_P4_Stellate_VS_P1_P3_Lumbar_Coeliac_Sphenopalatine | 0 | 1 | 0 | 1 | 0 | 0 | 1 | 0 |
| ATP5J | -0.174 | P2_P4_Stellate_VS_P1_P3_Lumbar_Coeliac_Sphenopalatine | 0 | 1 | 0 | 1 | 0 | 0 | 1 | 0 |
| TNFAIP1 | -0.174 | P1_P2_P3_Lumbar_Stellate_VS_P4_Coeliac_Sphenopalatine | 1 | 1 | 1 | 0 | 1 | 0 | 1 | 0 |
| ATP5H | -0.174 | P1_P2_P3_Lumbar_Stellate_VS_P4_Coeliac_Sphenopalatine | 1 | 1 | 1 | 0 | 1 | 0 | 1 | 0 |
| UBE2I | -0.174 | P1_P4_Lumbar_Stellate_VS_P2_P3_Coeliac_Sphenopalatine | 1 | 0 | 0 | 1 | 1 | 0 | 1 | 0 |
| FAM204A | -0.174 | P1_P4_Lumbar_Stellate_VS_P2_P3_Coeliac_Sphenopalatine | 1 | 0 | 0 | 1 | 1 | 0 | 1 | 0 |
| CISD2 | -0.174 | P1_Lumbar_Stellate_Sphenopalatine_VS_P2_P3_P4_Coeliac | 1 | 0 | 0 | 0 | 1 | 0 | 1 | 1 |
| JUND | -0.174 | P1_P2_P3_Coeliac_Stellate_Sphenopalatine_VS_P4_Lumbar | 1 | 1 | 1 | 0 | 0 | 1 | 1 | 1 |
| COX5B | -0.175 | P1_P3_Stellate_VS_P2_P4_Lumbar_Coeliac_Sphenopalatine | 1 | 0 | 1 | 0 | 0 | 0 | 1 | 0 |
| SH3BGRL3 | -0.175 | P1_P3_Stellate_VS_P2_P4_Lumbar_Coeliac_Sphenopalatine | 1 | 0 | 1 | 0 | 0 | 0 | 1 | 0 |
| SNRPN | -0.175 | P1_P3_Stellate_VS_P2_P4_Lumbar_Coeliac_Sphenopalatine | 1 | 0 | 1 | 0 | 0 | 0 | 1 | 0 |
| CST3 | -0.175 | P1_P2_P3_Stellate_VS_P4_Lumbar_Coeliac_Sphenopalatine | 1 | 1 | 1 | 0 | 0 | 0 | 1 | 0 |
| VDAC2 | -0.175 | P1_P2_P3_Stellate_VS_P4_Lumbar_Coeliac_Sphenopalatine | 1 | 1 | 1 | 0 | 0 | 0 | 1 | 0 |
| NDFIP1 | -0.175 | P1_P2_P3_Stellate_VS_P4_Lumbar_Coeliac_Sphenopalatine | 1 | 1 | 1 | 0 | 0 | 0 | 1 | 0 |
| MT-ND3 | -0.175 | P1_P2_P3_Stellate_VS_P4_Lumbar_Coeliac_Sphenopalatine | 1 | 1 | 1 | 0 | 0 | 0 | 1 | 0 |
| PCMT1 | -0.175 | P1_P4_Stellate_VS_P2_P3_Lumbar_Coeliac_Sphenopalatine | 1 | 0 | 0 | 1 | 0 | 0 | 1 | 0 |
| RPL22L1 | -0.175 | P2_P4_Stellate_VS_P1_P3_Lumbar_Coeliac_Sphenopalatine | 0 | 1 | 0 | 1 | 0 | 0 | 1 | 0 |
| RPS25 | -0.175 | P2_P4_Stellate_VS_P1_P3_Lumbar_Coeliac_Sphenopalatine | 0 | 1 | 0 | 1 | 0 | 0 | 1 | 0 |

| | | | | | | | | | | |
|---|---|---|---|---|---|---|---|---|---|---|
| COX7A2 | -0.175 | P2_P4_Stellate_VS_P1_P3_Lumbar_Coeliac_Sphenopalatine | 0 | 1 | 0 | 1 | 0 | 0 | 1 | 0 |
| ITFG1 | -0.175 | P1_P2_P4_Stellate_VS_P3_Lumbar_Coeliac_Sphenopalatine | 1 | 1 | 0 | 1 | 0 | 0 | 1 | 0 |
| TFDP1 | -0.175 | P2_P3_Lumbar_Stellate_VS_P1_P4_Coeliac_Sphenopalatine | 0 | 1 | 1 | 0 | 1 | 0 | 1 | 0 |
| ATP5G1 | -0.175 | P1_P2_P3_Lumbar_Stellate_VS_P4_Coeliac_Sphenopalatine | 1 | 1 | 1 | 0 | 1 | 0 | 1 | 0 |
| SYT16 | -0.175 | P1_P2_P3_Lumbar_Stellate_VS_P4_Coeliac_Sphenopalatine | 1 | 1 | 1 | 0 | 1 | 0 | 1 | 0 |
| FDPS | -0.175 | P1_P4_Lumbar_Stellate_VS_P2_P3_Coeliac_Sphenopalatine | 1 | 0 | 0 | 1 | 1 | 0 | 1 | 0 |
| SBDS | -0.175 | P1_P4_Lumbar_Stellate_VS_P2_P3_Coeliac_Sphenopalatine | 1 | 0 | 0 | 1 | 1 | 0 | 1 | 0 |
| PDCD10 | -0.175 | P1_P3_P4_Coeliac_Stellate_VS_P2_Lumbar_Sphenopalatine | 1 | 0 | 1 | 1 | 0 | 1 | 1 | 0 |
| NPM1 | -0.175 | P1_P3_P4_Coeliac_Stellate_VS_P2_Lumbar_Sphenopalatine | 1 | 0 | 1 | 1 | 0 | 1 | 1 | 0 |
| OAZ1 | -0.175 | P1_P2_P3_P4_Coeliac_Stellate_VS_Lumbar_Sphenopalatine | 1 | 1 | 1 | 1 | 0 | 1 | 1 | 0 |
| YBX1 | -0.175 | Stellate_Sphenopalatine_VS_P1_P2_P3_P4_Lumbar_Coeliac | 0 | 0 | 0 | 0 | 0 | 0 | 1 | 1 |
| SERINC1 | -0.175 | P1_P2_Stellate_Sphenopalatine_VS_P3_P4_Lumbar_Coeliac | 1 | 1 | 0 | 0 | 0 | 0 | 1 | 1 |
| RBIS | -0.175 | P3_Coeliac_Stellate_Sphenopalatine_VS_P1_P2_P4_Lumbar | 0 | 0 | 1 | 0 | 0 | 1 | 1 | 1 |
| YWHAB | -0.175 | P1_P2_P3_Coeliac_Stellate_Sphenopalatine_VS_P4_Lumbar | 1 | 1 | 1 | 0 | 0 | 1 | 1 | 1 |
| RTN4 | -0.175 | P1_P2_P3_Coeliac_Stellate_Sphenopalatine_VS_P4_Lumbar | 1 | 1 | 1 | 0 | 0 | 1 | 1 | 1 |
| DNAJA1 | -0.175 | P1_P2_P4_Coeliac_Stellate_Sphenopalatine_VS_P3_Lumbar | 1 | 1 | 0 | 1 | 0 | 1 | 1 | 1 |
| COX6A1 | -0.176 | Stellate_VS_P1_P2_P3_P4_Lumbar_Coeliac_Sphenopalatine | 0 | 0 | 0 | 0 | 0 | 0 | 1 | 0 |
| RPL23A | -0.176 | Stellate_VS_P1_P2_P3_P4_Lumbar_Coeliac_Sphenopalatine | 0 | 0 | 0 | 0 | 0 | 0 | 1 | 0 |
| ATP5MPL | -0.176 | Stellate_VS_P1_P2_P3_P4_Lumbar_Coeliac_Sphenopalatine | 0 | 0 | 0 | 0 | 0 | 0 | 1 | 0 |
| RPL35 | -0.176 | P2_Stellate_VS_P1_P3_P4_Lumbar_Coeliac_Sphenopalatine | 0 | 1 | 0 | 0 | 0 | 0 | 1 | 0 |
| PEBP1 | -0.176 | P2_Stellate_VS_P1_P3_P4_Lumbar_Coeliac_Sphenopalatine | 0 | 1 | 0 | 0 | 0 | 0 | 1 | 0 |
| PRDX1 | -0.176 | P1_P3_Stellate_VS_P2_P4_Lumbar_Coeliac_Sphenopalatine | 1 | 0 | 1 | 0 | 0 | 0 | 1 | 0 |
| RPL13A | -0.176 | P1_P3_Stellate_VS_P2_P4_Lumbar_Coeliac_Sphenopalatine | 1 | 0 | 1 | 0 | 0 | 0 | 1 | 0 |
| PSMB4 | -0.176 | P1_P4_Stellate_VS_P2_P3_Lumbar_Coeliac_Sphenopalatine | 1 | 0 | 0 | 1 | 0 | 0 | 1 | 0 |
| PGK1 | -0.176 | P1_P4_Stellate_VS_P2_P3_Lumbar_Coeliac_Sphenopalatine | 1 | 0 | 0 | 1 | 0 | 0 | 1 | 0 |
| FKBP1A | -0.176 | P2_P4_Stellate_VS_P1_P3_Lumbar_Coeliac_Sphenopalatine | 0 | 1 | 0 | 1 | 0 | 0 | 1 | 0 |
| MRFAP1 | -0.176 | P2_P4_Stellate_VS_P1_P3_Lumbar_Coeliac_Sphenopalatine | 0 | 1 | 0 | 1 | 0 | 0 | 1 | 0 |
| ARF5 | -0.176 | P2_P4_Stellate_VS_P1_P3_Lumbar_Coeliac_Sphenopalatine | 0 | 1 | 0 | 1 | 0 | 0 | 1 | 0 |
| GABARAP | -0.176 | P2_P4_Stellate_VS_P1_P3_Lumbar_Coeliac_Sphenopalatine | 0 | 1 | 0 | 1 | 0 | 0 | 1 | 0 |
| TXNDC17 | -0.176 | P2_P4_Stellate_VS_P1_P3_Lumbar_Coeliac_Sphenopalatine | 0 | 1 | 0 | 1 | 0 | 0 | 1 | 0 |

| | | | | | | | | | | |
|---|---|---|---|---|---|---|---|---|---|---|
| ZWINT | -0.176 | P1_P2_P4_Stellate_VS_P3_Lumbar_Coeliac_Sphenopalatine | 1 | 1 | 0 | 1 | 0 | 0 | 1 | 0 |
| DRAM2 | -0.176 | P2_P3_Lumbar_Stellate_VS_P1_P4_Coeliac_Sphenopalatine | 0 | 1 | 1 | 0 | 1 | 0 | 1 | 0 |
| RPS6 | -0.176 | P2_P3_Lumbar_Stellate_VS_P1_P4_Coeliac_Sphenopalatine | 0 | 1 | 1 | 0 | 1 | 0 | 1 | 0 |
| VDAC3 | -0.176 | P2_P3_Lumbar_Stellate_VS_P1_P4_Coeliac_Sphenopalatine | 0 | 1 | 1 | 0 | 1 | 0 | 1 | 0 |
| ABCA7 | -0.176 | P1_P2_P4_Lumbar_Stellate_VS_P3_Coeliac_Sphenopalatine | 1 | 1 | 0 | 1 | 1 | 0 | 1 | 0 |
| FKBP3 | -0.176 | P1_P3_P4_Coeliac_Stellate_VS_P2_Lumbar_Sphenopalatine | 1 | 0 | 1 | 1 | 0 | 1 | 1 | 0 |
| EIF4A1 | -0.176 | P1_P2_P3_P4_Coeliac_Stellate_VS_Lumbar_Sphenopalatine | 1 | 1 | 1 | 1 | 0 | 1 | 1 | 0 |
| PSMB1 | -0.176 | P1_P2_P3_P4_Coeliac_Stellate_VS_Lumbar_Sphenopalatine | 1 | 1 | 1 | 1 | 0 | 1 | 1 | 0 |
| SUB1 | -0.176 | P2_Stellate_Sphenopalatine_VS_P1_P3_P4_Lumbar_Coeliac | 0 | 1 | 0 | 0 | 0 | 0 | 1 | 1 |
| ATP5MD | -0.176 | P2_Stellate_Sphenopalatine_VS_P1_P3_P4_Lumbar_Coeliac | 0 | 1 | 0 | 0 | 0 | 0 | 1 | 1 |
| RPL36A | -0.176 | P2_Stellate_Sphenopalatine_VS_P1_P3_P4_Lumbar_Coeliac | 0 | 1 | 0 | 0 | 0 | 0 | 1 | 1 |
| SMDT1 | -0.176 | Lumbar_Stellate_Sphenopalatine_VS_P1_P2_P3_P4_Coeliac | 0 | 0 | 0 | 0 | 1 | 0 | 1 | 1 |
| FKBP4 | -0.176 | P1_P3_Coeliac_Stellate_Sphenopalatine_VS_P2_P4_Lumbar | 1 | 0 | 1 | 0 | 0 | 1 | 1 | 1 |
| VCP | -0.176 | P1_P2_P3_Coeliac_Stellate_Sphenopalatine_VS_P4_Lumbar | 1 | 1 | 1 | 0 | 0 | 1 | 1 | 1 |
| HINT1 | -0.176 | P1_P2_P4_Coeliac_Stellate_Sphenopalatine_VS_P3_Lumbar | 1 | 1 | 0 | 1 | 0 | 1 | 1 | 1 |
| H2AFZ | -0.177 | Stellate_VS_P1_P2_P3_P4_Lumbar_Coeliac_Sphenopalatine | 0 | 0 | 0 | 0 | 0 | 0 | 1 | 0 |
| RPL24 | -0.177 | P2_Stellate_VS_P1_P3_P4_Lumbar_Coeliac_Sphenopalatine | 0 | 1 | 0 | 0 | 0 | 0 | 1 | 0 |
| MT-ND1 | -0.177 | P2_Stellate_VS_P1_P3_P4_Lumbar_Coeliac_Sphenopalatine | 0 | 1 | 0 | 0 | 0 | 0 | 1 | 0 |
| RPL27 | -0.177 | P1_P2_Stellate_VS_P3_P4_Lumbar_Coeliac_Sphenopalatine | 1 | 1 | 0 | 0 | 0 | 0 | 1 | 0 |
| RPS17 | -0.177 | P1_P4_Stellate_VS_P2_P3_Lumbar_Coeliac_Sphenopalatine | 1 | 0 | 0 | 1 | 0 | 0 | 1 | 0 |
| PHOX2A | -0.177 | P1_P4_Stellate_VS_P2_P3_Lumbar_Coeliac_Sphenopalatine | 1 | 0 | 0 | 1 | 0 | 0 | 1 | 0 |
| 6330403K07I | -0.177 | P1_P4_Stellate_VS_P2_P3_Lumbar_Coeliac_Sphenopalatine | 1 | 0 | 0 | 1 | 0 | 0 | 1 | 0 |
| REEP5 | -0.177 | P1_P4_Stellate_VS_P2_P3_Lumbar_Coeliac_Sphenopalatine | 1 | 0 | 0 | 1 | 0 | 0 | 1 | 0 |
| DYNLL1 | -0.177 | P1_P2_P3_Lumbar_Stellate_VS_P4_Coeliac_Sphenopalatine | 1 | 1 | 1 | 0 | 1 | 0 | 1 | 0 |
| RPS19 | -0.177 | P1_P2_P3_Lumbar_Stellate_VS_P4_Coeliac_Sphenopalatine | 1 | 1 | 1 | 0 | 1 | 0 | 1 | 0 |
| ELOB | -0.177 | P1_P2_P3_Lumbar_Stellate_VS_P4_Coeliac_Sphenopalatine | 1 | 1 | 1 | 0 | 1 | 0 | 1 | 0 |
| ATP6V0C | -0.177 | P1_P2_P3_Lumbar_Stellate_VS_P4_Coeliac_Sphenopalatine | 1 | 1 | 1 | 0 | 1 | 0 | 1 | 0 |
| RPS28 | -0.177 | P1_P2_P3_Lumbar_Stellate_VS_P4_Coeliac_Sphenopalatine | 1 | 1 | 1 | 0 | 1 | 0 | 1 | 0 |
| H3F3A | -0.177 | P2_P4_Lumbar_Stellate_VS_P1_P3_Coeliac_Sphenopalatine | 0 | 1 | 0 | 1 | 1 | 0 | 1 | 0 |
| RPS27 | -0.177 | P2_P4_Lumbar_Stellate_VS_P1_P3_Coeliac_Sphenopalatine | 0 | 1 | 0 | 1 | 1 | 0 | 1 | 0 |

| | | | | | | | | | | |
|---|---|---|---|---|---|---|---|---|---|---|
| ATP5B | -0.177 | P2_P4_Lumbar_Stellate_VS_P1_P3_Coeliac_Sphenopalatine | 0 | 1 | 0 | 1 | 1 | 0 | 1 | 0 |
| DPYSL2 | -0.177 | P4_Coeliac_Stellate_VS_P1_P2_P3_Lumbar_Sphenopalatine | 0 | 0 | 0 | 1 | 0 | 1 | 1 | 0 |
| CIAO2A | -0.177 | P1_P3_P4_Coeliac_Stellate_VS_P2_Lumbar_Sphenopalatine | 1 | 0 | 1 | 1 | 0 | 1 | 1 | 0 |
| TAGLN2 | -0.177 | P2_P3_P4_Coeliac_Stellate_VS_P1_Lumbar_Sphenopalatine | 0 | 1 | 1 | 1 | 0 | 1 | 1 | 0 |
| ATP5G3 | -0.177 | P2_P3_P4_Coeliac_Stellate_VS_P1_Lumbar_Sphenopalatine | 0 | 1 | 1 | 1 | 0 | 1 | 1 | 0 |
| GNB1 | -0.177 | P2_P3_P4_Coeliac_Stellate_VS_P1_Lumbar_Sphenopalatine | 0 | 1 | 1 | 1 | 0 | 1 | 1 | 0 |
| RPS26 | -0.177 | P2_P3_P4_Coeliac_Stellate_VS_P1_Lumbar_Sphenopalatine | 0 | 1 | 1 | 1 | 0 | 1 | 1 | 0 |
| KLC1 | -0.177 | P2_P3_P4_Coeliac_Stellate_VS_P1_Lumbar_Sphenopalatine | 0 | 1 | 1 | 1 | 0 | 1 | 1 | 0 |
| CCT5 | -0.177 | P2_P3_P4_Coeliac_Stellate_VS_P1_Lumbar_Sphenopalatine | 0 | 1 | 1 | 1 | 0 | 1 | 1 | 0 |
| PFDN5 | -0.177 | P2_P3_P4_Coeliac_Stellate_VS_P1_Lumbar_Sphenopalatine | 0 | 1 | 1 | 1 | 0 | 1 | 1 | 0 |
| JPT1 | -0.177 | P1_P2_Stellate_Sphenopalatine_VS_P3_P4_Lumbar_Coeliac | 1 | 1 | 0 | 0 | 0 | 0 | 1 | 1 |
| ARHGDIG | -0.177 | P1_P2_Stellate_Sphenopalatine_VS_P3_P4_Lumbar_Coeliac | 1 | 1 | 0 | 0 | 0 | 0 | 1 | 1 |
| ATP5F1 | -0.177 | Lumbar_Stellate_Sphenopalatine_VS_P1_P2_P3_P4_Coeliac | 0 | 0 | 0 | 0 | 1 | 0 | 1 | 1 |
| ST13 | -0.177 | P2_Lumbar_Stellate_Sphenopalatine_VS_P1_P3_P4_Coeliac | 0 | 1 | 0 | 0 | 1 | 0 | 1 | 1 |
| NUCKS1 | -0.177 | P1_P2_P3_Coeliac_Stellate_Sphenopalatine_VS_P4_Lumbar | 1 | 1 | 1 | 0 | 0 | 1 | 1 | 1 |
| HSPE1 | -0.177 | P4_Coeliac_Stellate_Sphenopalatine_VS_P1_P2_P3_Lumbar | 0 | 0 | 0 | 1 | 0 | 1 | 1 | 1 |
| NDUFB8 | -0.177 | P4_Coeliac_Stellate_Sphenopalatine_VS_P1_P2_P3_Lumbar | 0 | 0 | 0 | 1 | 0 | 1 | 1 | 1 |
| MARCKS | -0.177 | P2_P4_Coeliac_Stellate_Sphenopalatine_VS_P1_P3_Lumbar | 0 | 1 | 0 | 1 | 0 | 1 | 1 | 1 |
| SF1 | -0.177 | P1_P2_Lumbar_Coeliac_Stellate_Sphenopalatine_VS_P3_P4 | 1 | 1 | 0 | 0 | 1 | 1 | 1 | 1 |
| ATP5L | -0.178 | Stellate_VS_P1_P2_P3_P4_Lumbar_Coeliac_Sphenopalatine | 0 | 0 | 0 | 0 | 0 | 0 | 1 | 0 |
| COX5A | -0.178 | Stellate_VS_P1_P2_P3_P4_Lumbar_Coeliac_Sphenopalatine | 0 | 0 | 0 | 0 | 0 | 0 | 1 | 0 |
| EEF2 | -0.178 | Stellate_VS_P1_P2_P3_P4_Lumbar_Coeliac_Sphenopalatine | 0 | 0 | 0 | 0 | 0 | 0 | 1 | 0 |
| ATP5O | -0.178 | P2_Stellate_VS_P1_P3_P4_Lumbar_Coeliac_Sphenopalatine | 0 | 1 | 0 | 0 | 0 | 0 | 1 | 0 |
| MALAT1 | -0.178 | P2_Stellate_VS_P1_P3_P4_Lumbar_Coeliac_Sphenopalatine | 0 | 1 | 0 | 0 | 0 | 0 | 1 | 0 |
| MT-ATP6 | -0.178 | P2_Stellate_VS_P1_P3_P4_Lumbar_Coeliac_Sphenopalatine | 0 | 1 | 0 | 0 | 0 | 0 | 1 | 0 |
| NSG1 | -0.178 | P1_P2_Stellate_VS_P3_P4_Lumbar_Coeliac_Sphenopalatine | 1 | 1 | 0 | 0 | 0 | 0 | 1 | 0 |
| S100A10 | -0.178 | P1_P4_Stellate_VS_P2_P3_Lumbar_Coeliac_Sphenopalatine | 1 | 0 | 0 | 1 | 0 | 0 | 1 | 0 |
| SEM1 | -0.178 | P1_P4_Stellate_VS_P2_P3_Lumbar_Coeliac_Sphenopalatine | 1 | 0 | 0 | 1 | 0 | 0 | 1 | 0 |
| ATP5D | -0.178 | P1_P4_Stellate_VS_P2_P3_Lumbar_Coeliac_Sphenopalatine | 1 | 0 | 0 | 1 | 0 | 0 | 1 | 0 |
| EEF1G | -0.178 | P1_P4_Stellate_VS_P2_P3_Lumbar_Coeliac_Sphenopalatine | 1 | 0 | 0 | 1 | 0 | 0 | 1 | 0 |

| | | | | | | | | | | |
|---|---|---|---|---|---|---|---|---|---|---|
| RBM3 | -0.178 | P1_P4_Stellate_VS_P2_P3_Lumbar_Coeliac_Sphenopalatine | 1 | 0 | 0 | 1 | 0 | 0 | 1 | 0 |
| RPS21 | -0.178 | P1_P2_P3_Lumbar_Stellate_VS_P4_Coeliac_Sphenopalatine | 1 | 1 | 1 | 0 | 1 | 0 | 1 | 0 |
| CNPPD1 | -0.178 | P2_P4_Lumbar_Stellate_VS_P1_P3_Coeliac_Sphenopalatine | 0 | 1 | 0 | 1 | 1 | 0 | 1 | 0 |
| RPS16 | -0.178 | P2_P4_Lumbar_Stellate_VS_P1_P3_Coeliac_Sphenopalatine | 0 | 1 | 0 | 1 | 1 | 0 | 1 | 0 |
| MPC1 | -0.178 | P2_P4_Lumbar_Stellate_VS_P1_P3_Coeliac_Sphenopalatine | 0 | 1 | 0 | 1 | 1 | 0 | 1 | 0 |
| MT-CO1 | -0.178 | P2_P4_Lumbar_Stellate_VS_P1_P3_Coeliac_Sphenopalatine | 0 | 1 | 0 | 1 | 1 | 0 | 1 | 0 |
| SNX6 | -0.178 | P4_Coeliac_Stellate_VS_P1_P2_P3_Lumbar_Sphenopalatine | 0 | 0 | 0 | 1 | 0 | 1 | 1 | 0 |
| EIF5A | -0.178 | P1_P3_P4_Coeliac_Stellate_VS_P2_Lumbar_Sphenopalatine | 1 | 0 | 1 | 1 | 0 | 1 | 1 | 0 |
| RPL21 | -0.178 | P2_P3_P4_Coeliac_Stellate_VS_P1_Lumbar_Sphenopalatine | 0 | 1 | 1 | 1 | 0 | 1 | 1 | 0 |
| RPS3 | -0.178 | P2_P3_P4_Coeliac_Stellate_VS_P1_Lumbar_Sphenopalatine | 0 | 1 | 1 | 1 | 0 | 1 | 1 | 0 |
| RAB3A | -0.178 | P2_P3_P4_Coeliac_Stellate_VS_P1_Lumbar_Sphenopalatine | 0 | 1 | 1 | 1 | 0 | 1 | 1 | 0 |
| PRDX2 | -0.178 | P2_P3_P4_Coeliac_Stellate_VS_P1_Lumbar_Sphenopalatine | 0 | 1 | 1 | 1 | 0 | 1 | 1 | 0 |
| COX4I1 | -0.178 | P2_P3_P4_Coeliac_Stellate_VS_P1_Lumbar_Sphenopalatine | 0 | 1 | 1 | 1 | 0 | 1 | 1 | 0 |
| RPS15 | -0.178 | P2_P3_P4_Coeliac_Stellate_VS_P1_Lumbar_Sphenopalatine | 0 | 1 | 1 | 1 | 0 | 1 | 1 | 0 |
| CCT8 | -0.178 | P2_P3_P4_Coeliac_Stellate_VS_P1_Lumbar_Sphenopalatine | 0 | 1 | 1 | 1 | 0 | 1 | 1 | 0 |
| YWHAE | -0.178 | P1_P2_P3_P4_Coeliac_Stellate_VS_Lumbar_Sphenopalatine | 1 | 1 | 1 | 1 | 0 | 1 | 1 | 0 |
| TUBA1B | -0.178 | P1_P2_P3_P4_Coeliac_Stellate_VS_Lumbar_Sphenopalatine | 1 | 1 | 1 | 1 | 0 | 1 | 1 | 0 |
| SCD2 | -0.178 | P1_Lumbar_Stellate_Sphenopalatine_VS_P2_P3_P4_Coeliac | 1 | 0 | 0 | 0 | 1 | 0 | 1 | 1 |
| HNRNPDL | -0.178 | P1_P3_Coeliac_Stellate_Sphenopalatine_VS_P2_P4_Lumbar | 1 | 0 | 1 | 0 | 0 | 1 | 1 | 1 |
| PSMD4 | -0.178 | P2_P3_Coeliac_Stellate_Sphenopalatine_VS_P1_P4_Lumbar | 0 | 1 | 1 | 0 | 0 | 1 | 1 | 1 |
| PSMA1 | -0.178 | P2_P3_Coeliac_Stellate_Sphenopalatine_VS_P1_P4_Lumbar | 0 | 1 | 1 | 0 | 0 | 1 | 1 | 1 |
| RBFOX2 | -0.178 | P4_Coeliac_Stellate_Sphenopalatine_VS_P1_P2_P3_Lumbar | 0 | 0 | 0 | 1 | 0 | 1 | 1 | 1 |
| BEX3 | -0.178 | P4_Coeliac_Stellate_Sphenopalatine_VS_P1_P2_P3_Lumbar | 0 | 0 | 0 | 1 | 0 | 1 | 1 | 1 |
| YWHAZ | -0.178 | P1_P2_P4_Coeliac_Stellate_Sphenopalatine_VS_P3_Lumbar | 1 | 1 | 0 | 1 | 0 | 1 | 1 | 1 |
| RPLP0 | -0.179 | Stellate_VS_P1_P2_P3_P4_Lumbar_Coeliac_Sphenopalatine | 0 | 0 | 0 | 0 | 0 | 0 | 1 | 0 |
| RPL32 | -0.179 | Stellate_VS_P1_P2_P3_P4_Lumbar_Coeliac_Sphenopalatine | 0 | 0 | 0 | 0 | 0 | 0 | 1 | 0 |
| RPL28 | -0.179 | Stellate_VS_P1_P2_P3_P4_Lumbar_Coeliac_Sphenopalatine | 0 | 0 | 0 | 0 | 0 | 0 | 1 | 0 |
| RPS5 | -0.179 | Stellate_VS_P1_P2_P3_P4_Lumbar_Coeliac_Sphenopalatine | 0 | 0 | 0 | 0 | 0 | 0 | 1 | 0 |
| UQCRQ | -0.179 | Stellate_VS_P1_P2_P3_P4_Lumbar_Coeliac_Sphenopalatine | 0 | 0 | 0 | 0 | 0 | 0 | 1 | 0 |
| RPL9 | -0.179 | P1_Stellate_VS_P2_P3_P4_Lumbar_Coeliac_Sphenopalatine | 1 | 0 | 0 | 0 | 0 | 0 | 1 | 0 |

| | | | | | | | | | | |
|---|---|---|---|---|---|---|---|---|---|---|
| RPLP2 | -0.179 | P1_Stellate_VS_P2_P3_P4_Lumbar_Coeliac_Sphenopalatine | 1 | 0 | 0 | 0 | 0 | 0 | 1 | 0 |
| RPL26 | -0.179 | P1_Stellate_VS_P2_P3_P4_Lumbar_Coeliac_Sphenopalatine | 1 | 0 | 0 | 0 | 0 | 0 | 1 | 0 |
| RPL15 | -0.179 | P1_Stellate_VS_P2_P3_P4_Lumbar_Coeliac_Sphenopalatine | 1 | 0 | 0 | 0 | 0 | 0 | 1 | 0 |
| CFL1 | -0.179 | P1_Stellate_VS_P2_P3_P4_Lumbar_Coeliac_Sphenopalatine | 1 | 0 | 0 | 0 | 0 | 0 | 1 | 0 |
| MT-CO2 | -0.179 | P1_Stellate_VS_P2_P3_P4_Lumbar_Coeliac_Sphenopalatine | 1 | 0 | 0 | 0 | 0 | 0 | 1 | 0 |
| RPL27A | -0.179 | P2_Stellate_VS_P1_P3_P4_Lumbar_Coeliac_Sphenopalatine | 0 | 1 | 0 | 0 | 0 | 0 | 1 | 0 |
| SLC25A3 | -0.179 | P2_Stellate_VS_P1_P3_P4_Lumbar_Coeliac_Sphenopalatine | 0 | 1 | 0 | 0 | 0 | 0 | 1 | 0 |
| RPL19 | -0.179 | P2_Stellate_VS_P1_P3_P4_Lumbar_Coeliac_Sphenopalatine | 0 | 1 | 0 | 0 | 0 | 0 | 1 | 0 |
| RPL35A | -0.179 | P2_Stellate_VS_P1_P3_P4_Lumbar_Coeliac_Sphenopalatine | 0 | 1 | 0 | 0 | 0 | 0 | 1 | 0 |
| RPL7 | -0.179 | P1_P2_Stellate_VS_P3_P4_Lumbar_Coeliac_Sphenopalatine | 1 | 1 | 0 | 0 | 0 | 0 | 1 | 0 |
| YWHAH | -0.179 | P1_P2_Stellate_VS_P3_P4_Lumbar_Coeliac_Sphenopalatine | 1 | 1 | 0 | 0 | 0 | 0 | 1 | 0 |
| FTL1 | -0.179 | P1_P2_Stellate_VS_P3_P4_Lumbar_Coeliac_Sphenopalatine | 1 | 1 | 0 | 0 | 0 | 0 | 1 | 0 |
| SLC25A4 | -0.179 | P1_P2_Stellate_VS_P3_P4_Lumbar_Coeliac_Sphenopalatine | 1 | 1 | 0 | 0 | 0 | 0 | 1 | 0 |
| RPS27A | -0.179 | P1_P2_Stellate_VS_P3_P4_Lumbar_Coeliac_Sphenopalatine | 1 | 1 | 0 | 0 | 0 | 0 | 1 | 0 |
| HSP90AA1 | -0.179 | P1_P2_Stellate_VS_P3_P4_Lumbar_Coeliac_Sphenopalatine | 1 | 1 | 0 | 0 | 0 | 0 | 1 | 0 |
| COX8A | -0.179 | P1_P2_Stellate_VS_P3_P4_Lumbar_Coeliac_Sphenopalatine | 1 | 1 | 0 | 0 | 0 | 0 | 1 | 0 |
| FTH1 | -0.179 | P1_P2_Stellate_VS_P3_P4_Lumbar_Coeliac_Sphenopalatine | 1 | 1 | 0 | 0 | 0 | 0 | 1 | 0 |
| GNB2 | -0.179 | P1_P4_Stellate_VS_P2_P3_Lumbar_Coeliac_Sphenopalatine | 1 | 0 | 0 | 1 | 0 | 0 | 1 | 0 |
| RPS15A | -0.179 | P1_P4_Stellate_VS_P2_P3_Lumbar_Coeliac_Sphenopalatine | 1 | 0 | 0 | 1 | 0 | 0 | 1 | 0 |
| NDUFA13 | -0.179 | P1_P4_Stellate_VS_P2_P3_Lumbar_Coeliac_Sphenopalatine | 1 | 0 | 0 | 1 | 0 | 0 | 1 | 0 |
| RPL8 | -0.179 | P2_Lumbar_Stellate_VS_P1_P3_P4_Coeliac_Sphenopalatine | 0 | 1 | 0 | 0 | 1 | 0 | 1 | 0 |
| RPS11 | -0.179 | Coeliac_Stellate_VS_P1_P2_P3_P4_Lumbar_Sphenopalatine | 0 | 0 | 0 | 0 | 0 | 1 | 1 | 0 |
| RPS9 | -0.179 | P2_P3_Coeliac_Stellate_VS_P1_P4_Lumbar_Sphenopalatine | 0 | 1 | 1 | 0 | 0 | 1 | 1 | 0 |
| RPL18A | -0.179 | P2_P3_Coeliac_Stellate_VS_P1_P4_Lumbar_Sphenopalatine | 0 | 1 | 1 | 0 | 0 | 1 | 1 | 0 |
| YWHAG | -0.179 | P1_P3_P4_Coeliac_Stellate_VS_P2_Lumbar_Sphenopalatine | 1 | 0 | 1 | 1 | 0 | 1 | 1 | 0 |
| MOCS2 | -0.179 | P1_P3_P4_Coeliac_Stellate_VS_P2_Lumbar_Sphenopalatine | 1 | 0 | 1 | 1 | 0 | 1 | 1 | 0 |
| FERMT2 | -0.179 | P1_P3_P4_Coeliac_Stellate_VS_P2_Lumbar_Sphenopalatine | 1 | 0 | 1 | 1 | 0 | 1 | 1 | 0 |
| MLLT11 | -0.179 | P2_P3_P4_Coeliac_Stellate_VS_P1_Lumbar_Sphenopalatine | 0 | 1 | 1 | 1 | 0 | 1 | 1 | 0 |
| RPL34 | -0.179 | P2_P3_P4_Coeliac_Stellate_VS_P1_Lumbar_Sphenopalatine | 0 | 1 | 1 | 1 | 0 | 1 | 1 | 0 |
| STMN1 | -0.179 | P2_P3_P4_Coeliac_Stellate_VS_P1_Lumbar_Sphenopalatine | 0 | 1 | 1 | 1 | 0 | 1 | 1 | 0 |

| | | | | | | | | | | |
|---|---|---|---|---|---|---|---|---|---|---|
| RPL5 | -0.179 | P2_P3_P4_Coeliac_Stellate_VS_P1_Lumbar_Sphenopalatine | 0 | 1 | 1 | 1 | 0 | 1 | 1 | 0 |
| COX6B1 | -0.179 | P2_P3_P4_Coeliac_Stellate_VS_P1_Lumbar_Sphenopalatine | 0 | 1 | 1 | 1 | 0 | 1 | 1 | 0 |
| RPL14 | -0.179 | P2_P3_P4_Coeliac_Stellate_VS_P1_Lumbar_Sphenopalatine | 0 | 1 | 1 | 1 | 0 | 1 | 1 | 0 |
| MYL6 | -0.179 | P2_P3_P4_Coeliac_Stellate_VS_P1_Lumbar_Sphenopalatine | 0 | 1 | 1 | 1 | 0 | 1 | 1 | 0 |
| RPS3A1 | -0.179 | P1_P2_P3_P4_Coeliac_Stellate_VS_Lumbar_Sphenopalatine | 1 | 1 | 1 | 1 | 0 | 1 | 1 | 0 |
| RPS8 | -0.179 | P1_P2_P3_P4_Coeliac_Stellate_VS_Lumbar_Sphenopalatine | 1 | 1 | 1 | 1 | 0 | 1 | 1 | 0 |
| CCT7 | -0.179 | P1_P2_P3_P4_Coeliac_Stellate_VS_Lumbar_Sphenopalatine | 1 | 1 | 1 | 1 | 0 | 1 | 1 | 0 |
| SELENOW | -0.179 | P1_P2_P3_P4_Coeliac_Stellate_VS_Lumbar_Sphenopalatine | 1 | 1 | 1 | 1 | 0 | 1 | 1 | 0 |
| sept-07 | -0.179 | P1_P2_P3_P4_Coeliac_Stellate_VS_Lumbar_Sphenopalatine | 1 | 1 | 1 | 1 | 0 | 1 | 1 | 0 |
| RPL10A | -0.179 | P1_P2_P3_P4_Coeliac_Stellate_VS_Lumbar_Sphenopalatine | 1 | 1 | 1 | 1 | 0 | 1 | 1 | 0 |
| RPS20 | -0.179 | P2_Lumbar_Stellate_Sphenopalatine_VS_P1_P3_P4_Coeliac | 0 | 1 | 0 | 0 | 1 | 0 | 1 | 1 |
| RPS7 | -0.179 | P1_P2_Lumbar_Stellate_Sphenopalatine_VS_P3_P4_Coeliac | 1 | 1 | 0 | 0 | 1 | 0 | 1 | 1 |
| TOMM7 | -0.179 | P1_P3_Coeliac_Stellate_Sphenopalatine_VS_P2_P4_Lumbar | 1 | 0 | 1 | 0 | 0 | 1 | 1 | 1 |
| ATP6V0D1 | -0.179 | P2_P3_Coeliac_Stellate_Sphenopalatine_VS_P1_P4_Lumbar | 0 | 1 | 1 | 0 | 0 | 1 | 1 | 1 |
| RPL36 | -0.179 | P2_P3_Coeliac_Stellate_Sphenopalatine_VS_P1_P4_Lumbar | 0 | 1 | 1 | 0 | 0 | 1 | 1 | 1 |
| RPS14 | -0.179 | P2_P3_Coeliac_Stellate_Sphenopalatine_VS_P1_P4_Lumbar | 0 | 1 | 1 | 0 | 0 | 1 | 1 | 1 |
| POLR2J | -0.179 | P4_Coeliac_Stellate_Sphenopalatine_VS_P1_P2_P3_Lumbar | 0 | 0 | 0 | 1 | 0 | 1 | 1 | 1 |
| BEX2 | -0.179 | P4_Coeliac_Stellate_Sphenopalatine_VS_P1_P2_P3_Lumbar | 0 | 0 | 0 | 1 | 0 | 1 | 1 | 1 |
| EXOC5 | -0.179 | P1_P2_Lumbar_Coeliac_Stellate_Sphenopalatine_VS_P3_P4 | 1 | 1 | 0 | 0 | 1 | 1 | 1 | 1 |
| CHCHD2 | -0.18 | P1_Stellate_VS_P2_P3_P4_Lumbar_Coeliac_Sphenopalatine | 1 | 0 | 0 | 0 | 0 | 0 | 1 | 0 |
| GAPDH | -0.18 | P1_Stellate_VS_P2_P3_P4_Lumbar_Coeliac_Sphenopalatine | 1 | 0 | 0 | 0 | 0 | 0 | 1 | 0 |
| EIF1 | -0.18 | P1_Stellate_VS_P2_P3_P4_Lumbar_Coeliac_Sphenopalatine | 1 | 0 | 0 | 0 | 0 | 0 | 1 | 0 |
| RPL39 | -0.18 | P1_Stellate_VS_P2_P3_P4_Lumbar_Coeliac_Sphenopalatine | 1 | 0 | 0 | 0 | 0 | 0 | 1 | 0 |
| RPL37A | -0.18 | P2_Stellate_VS_P1_P3_P4_Lumbar_Coeliac_Sphenopalatine | 0 | 1 | 0 | 0 | 0 | 0 | 1 | 0 |
| PTMA | -0.18 | P2_Stellate_VS_P1_P3_P4_Lumbar_Coeliac_Sphenopalatine | 0 | 1 | 0 | 0 | 0 | 0 | 1 | 0 |
| ATPIF1 | -0.18 | P2_Stellate_VS_P1_P3_P4_Lumbar_Coeliac_Sphenopalatine | 0 | 1 | 0 | 0 | 0 | 0 | 1 | 0 |
| RPLP1 | -0.18 | P2_Stellate_VS_P1_P3_P4_Lumbar_Coeliac_Sphenopalatine | 0 | 1 | 0 | 0 | 0 | 0 | 1 | 0 |
| RPS29 | -0.18 | P2_Stellate_VS_P1_P3_P4_Lumbar_Coeliac_Sphenopalatine | 0 | 1 | 0 | 0 | 0 | 0 | 1 | 0 |
| COX6C | -0.18 | P2_Stellate_VS_P1_P3_P4_Lumbar_Coeliac_Sphenopalatine | 0 | 1 | 0 | 0 | 0 | 0 | 1 | 0 |
| RPL17 | -0.18 | P2_Stellate_VS_P1_P3_P4_Lumbar_Coeliac_Sphenopalatine | 0 | 1 | 0 | 0 | 0 | 0 | 1 | 0 |

| | | | | | | | | | | |
|---|---|---|---|---|---|---|---|---|---|---|
| MT-CO3 | -0.18 | P2_Stellate_VS_P1_P3_P4_Lumbar_Coeliac_Sphenopalatine | 0 | 1 | 0 | 0 | 0 | 0 | 1 | 0 |
| RPL7A | -0.18 | P1_P2_Stellate_VS_P3_P4_Lumbar_Coeliac_Sphenopalatine | 1 | 1 | 0 | 0 | 0 | 0 | 1 | 0 |
| RPS4X | -0.18 | P2_Lumbar_Stellate_VS_P1_P3_P4_Coeliac_Sphenopalatine | 0 | 1 | 0 | 0 | 1 | 0 | 1 | 0 |
| RPS2 | -0.18 | P1_P2_Coeliac_Stellate_VS_P3_P4_Lumbar_Sphenopalatine | 1 | 1 | 0 | 0 | 0 | 1 | 1 | 0 |
| MAP1LC3A | -0.18 | P1_P2_P3_Coeliac_Stellate_VS_P4_Lumbar_Sphenopalatine | 1 | 1 | 1 | 0 | 0 | 1 | 1 | 0 |
| DPYSL3 | -0.18 | P1_P2_P3_Coeliac_Stellate_VS_P4_Lumbar_Sphenopalatine | 1 | 1 | 1 | 0 | 0 | 1 | 1 | 0 |
| EIF3I | -0.18 | P1_P2_P3_P4_Coeliac_Stellate_VS_Lumbar_Sphenopalatine | 1 | 1 | 1 | 1 | 0 | 1 | 1 | 0 |
| GPI1 | -0.18 | P1_P2_P3_P4_Coeliac_Stellate_VS_Lumbar_Sphenopalatine | 1 | 1 | 1 | 1 | 0 | 1 | 1 | 0 |
| ALDOA | -0.18 | P1_P2_P3_P4_Coeliac_Stellate_VS_Lumbar_Sphenopalatine | 1 | 1 | 1 | 1 | 0 | 1 | 1 | 0 |
| ASAP1 | -0.18 | P1_P2_P3_P4_Coeliac_Stellate_VS_Lumbar_Sphenopalatine | 1 | 1 | 1 | 1 | 0 | 1 | 1 | 0 |
| TXN1 | -0.18 | P2_Lumbar_Stellate_Sphenopalatine_VS_P1_P3_P4_Coeliac | 0 | 1 | 0 | 0 | 1 | 0 | 1 | 1 |
| SPTBN1 | -0.18 | P1_P2_Lumbar_Stellate_Sphenopalatine_VS_P3_P4_Coeliac | 1 | 1 | 0 | 0 | 1 | 0 | 1 | 1 |
| PAFAH1B1 | -0.18 | P1_P2_P3_Coeliac_Stellate_Sphenopalatine_VS_P4_Lumbar | 1 | 1 | 1 | 0 | 0 | 1 | 1 | 1 |
| ATP5A1 | -0.18 | P4_Coeliac_Stellate_Sphenopalatine_VS_P1_P2_P3_Lumbar | 0 | 0 | 0 | 1 | 0 | 1 | 1 | 1 |
| NEDD4 | -0.18 | P1_P4_Coeliac_Stellate_Sphenopalatine_VS_P2_P3_Lumbar | 1 | 0 | 0 | 1 | 0 | 1 | 1 | 1 |
| DCTN2 | -0.18 | P1_P4_Coeliac_Stellate_Sphenopalatine_VS_P2_P3_Lumbar | 1 | 0 | 0 | 1 | 0 | 1 | 1 | 1 |
| GNAS | -0.181 | P1_Stellate_VS_P2_P3_P4_Lumbar_Coeliac_Sphenopalatine | 1 | 0 | 0 | 0 | 0 | 0 | 1 | 0 |
| STMN3 | -0.181 | P1_Stellate_VS_P2_P3_P4_Lumbar_Coeliac_Sphenopalatine | 1 | 0 | 0 | 0 | 0 | 0 | 1 | 0 |
| STMN2 | -0.181 | P1_Stellate_VS_P2_P3_P4_Lumbar_Coeliac_Sphenopalatine | 1 | 0 | 0 | 0 | 0 | 0 | 1 | 0 |
| RPL11 | -0.181 | P1_Stellate_VS_P2_P3_P4_Lumbar_Coeliac_Sphenopalatine | 1 | 0 | 0 | 0 | 0 | 0 | 1 | 0 |
| UCHL1 | -0.181 | P1_Stellate_VS_P2_P3_P4_Lumbar_Coeliac_Sphenopalatine | 1 | 0 | 0 | 0 | 0 | 0 | 1 | 0 |
| RPL6 | -0.181 | P1_Stellate_VS_P2_P3_P4_Lumbar_Coeliac_Sphenopalatine | 1 | 0 | 0 | 0 | 0 | 0 | 1 | 0 |
| ACTB | -0.181 | P1_Stellate_VS_P2_P3_P4_Lumbar_Coeliac_Sphenopalatine | 1 | 0 | 0 | 0 | 0 | 0 | 1 | 0 |
| NDUFA4 | -0.181 | P1_Stellate_VS_P2_P3_P4_Lumbar_Coeliac_Sphenopalatine | 1 | 0 | 0 | 0 | 0 | 0 | 1 | 0 |
| TMSB10 | -0.181 | P1_Stellate_VS_P2_P3_P4_Lumbar_Coeliac_Sphenopalatine | 1 | 0 | 0 | 0 | 0 | 0 | 1 | 0 |
| RPL13 | -0.181 | P1_Stellate_VS_P2_P3_P4_Lumbar_Coeliac_Sphenopalatine | 1 | 0 | 0 | 0 | 0 | 0 | 1 | 0 |
| TUBB3 | -0.181 | P1_Stellate_VS_P2_P3_P4_Lumbar_Coeliac_Sphenopalatine | 1 | 0 | 0 | 0 | 0 | 0 | 1 | 0 |
| HSPA8 | -0.181 | P1_Stellate_VS_P2_P3_P4_Lumbar_Coeliac_Sphenopalatine | 1 | 0 | 0 | 0 | 0 | 0 | 1 | 0 |
| EEF1A1 | -0.181 | P1_Stellate_VS_P2_P3_P4_Lumbar_Coeliac_Sphenopalatine | 1 | 0 | 0 | 0 | 0 | 0 | 1 | 0 |
| RPL41 | -0.181 | P1_Stellate_VS_P2_P3_P4_Lumbar_Coeliac_Sphenopalatine | 1 | 0 | 0 | 0 | 0 | 0 | 1 | 0 |

| | | | | | | | | | | |
|---|---|---|---|---|---|---|---|---|---|---|
| PPIA | -0.181 | P1_Stellate_VS_P2_P3_P4_Lumbar_Coeliac_Sphenopalatine | 1 | 0 | 0 | 0 | 0 | 0 | 1 | 0 |
| UBB | -0.181 | P1_Stellate_VS_P2_P3_P4_Lumbar_Coeliac_Sphenopalatine | 1 | 0 | 0 | 0 | 0 | 0 | 1 | 0 |
| RPL23 | -0.181 | P1_Stellate_VS_P2_P3_P4_Lumbar_Coeliac_Sphenopalatine | 1 | 0 | 0 | 0 | 0 | 0 | 1 | 0 |
| RPL38 | -0.181 | P1_Stellate_VS_P2_P3_P4_Lumbar_Coeliac_Sphenopalatine | 1 | 0 | 0 | 0 | 0 | 0 | 1 | 0 |
| ACTG1 | -0.181 | P1_Stellate_VS_P2_P3_P4_Lumbar_Coeliac_Sphenopalatine | 1 | 0 | 0 | 0 | 0 | 0 | 1 | 0 |
| RTN1 | -0.181 | P1_Stellate_VS_P2_P3_P4_Lumbar_Coeliac_Sphenopalatine | 1 | 0 | 0 | 0 | 0 | 0 | 1 | 0 |
| CALM1 | -0.181 | P1_Stellate_VS_P2_P3_P4_Lumbar_Coeliac_Sphenopalatine | 1 | 0 | 0 | 0 | 0 | 0 | 1 | 0 |
| TUBB2A | -0.181 | P1_Stellate_VS_P2_P3_P4_Lumbar_Coeliac_Sphenopalatine | 1 | 0 | 0 | 0 | 0 | 0 | 1 | 0 |
| TUBB2B | -0.181 | P1_Stellate_VS_P2_P3_P4_Lumbar_Coeliac_Sphenopalatine | 1 | 0 | 0 | 0 | 0 | 0 | 1 | 0 |
| COX7C | -0.181 | P1_Stellate_VS_P2_P3_P4_Lumbar_Coeliac_Sphenopalatine | 1 | 0 | 0 | 0 | 0 | 0 | 1 | 0 |
| RPS23 | -0.181 | P1_Stellate_VS_P2_P3_P4_Lumbar_Coeliac_Sphenopalatine | 1 | 0 | 0 | 0 | 0 | 0 | 1 | 0 |
| MAP1B | -0.181 | P1_Stellate_VS_P2_P3_P4_Lumbar_Coeliac_Sphenopalatine | 1 | 0 | 0 | 0 | 0 | 0 | 1 | 0 |
| RPS24 | -0.181 | P1_Stellate_VS_P2_P3_P4_Lumbar_Coeliac_Sphenopalatine | 1 | 0 | 0 | 0 | 0 | 0 | 1 | 0 |
| TPT1 | -0.181 | P1_Stellate_VS_P2_P3_P4_Lumbar_Coeliac_Sphenopalatine | 1 | 0 | 0 | 0 | 0 | 0 | 1 | 0 |
| RPL37 | -0.181 | P1_Stellate_VS_P2_P3_P4_Lumbar_Coeliac_Sphenopalatine | 1 | 0 | 0 | 0 | 0 | 0 | 1 | 0 |
| RPL3 | -0.181 | P1_Stellate_VS_P2_P3_P4_Lumbar_Coeliac_Sphenopalatine | 1 | 0 | 0 | 0 | 0 | 0 | 1 | 0 |
| TUBA1A | -0.181 | P1_Stellate_VS_P2_P3_P4_Lumbar_Coeliac_Sphenopalatine | 1 | 0 | 0 | 0 | 0 | 0 | 1 | 0 |
| PRPH | -0.181 | P1_Stellate_VS_P2_P3_P4_Lumbar_Coeliac_Sphenopalatine | 1 | 0 | 0 | 0 | 0 | 0 | 1 | 0 |
| GAP43 | -0.181 | P1_Stellate_VS_P2_P3_P4_Lumbar_Coeliac_Sphenopalatine | 1 | 0 | 0 | 0 | 0 | 0 | 1 | 0 |
| TUBB5 | -0.181 | P1_Stellate_VS_P2_P3_P4_Lumbar_Coeliac_Sphenopalatine | 1 | 0 | 0 | 0 | 0 | 0 | 1 | 0 |
| HSP90AB1 | -0.181 | P1_Stellate_VS_P2_P3_P4_Lumbar_Coeliac_Sphenopalatine | 1 | 0 | 0 | 0 | 0 | 0 | 1 | 0 |
| CALM2 | -0.181 | P1_Stellate_VS_P2_P3_P4_Lumbar_Coeliac_Sphenopalatine | 1 | 0 | 0 | 0 | 0 | 0 | 1 | 0 |
| FAU | -0.181 | P1_Stellate_VS_P2_P3_P4_Lumbar_Coeliac_Sphenopalatine | 1 | 0 | 0 | 0 | 0 | 0 | 1 | 0 |
| PCSK1N | -0.181 | P1_Stellate_VS_P2_P3_P4_Lumbar_Coeliac_Sphenopalatine | 1 | 0 | 0 | 0 | 0 | 0 | 1 | 0 |
| TMSB4X | -0.181 | P1_Stellate_VS_P2_P3_P4_Lumbar_Coeliac_Sphenopalatine | 1 | 0 | 0 | 0 | 0 | 0 | 1 | 0 |
| TPPP3 | -0.181 | P2_Stellate_VS_P1_P3_P4_Lumbar_Coeliac_Sphenopalatine | 0 | 1 | 0 | 0 | 0 | 0 | 1 | 0 |
| RPS10 | -0.181 | P1_P2_Lumbar_Stellate_VS_P3_P4_Coeliac_Sphenopalatine | 1 | 1 | 0 | 0 | 1 | 0 | 1 | 0 |
| UBC | -0.181 | P1_P2_Lumbar_Stellate_Sphenopalatine_VS_P3_P4_Coeliac | 1 | 1 | 0 | 0 | 1 | 0 | 1 | 1 |
| USP22 | -0.181 | P2_P4_Coeliac_Stellate_Sphenopalatine_VS_P1_P3_Lumbar | 0 | 1 | 0 | 1 | 0 | 1 | 1 | 1 |
| RPSA | -0.182 | P1_P2_Stellate_VS_P3_P4_Lumbar_Coeliac_Sphenopalatine | 1 | 1 | 0 | 0 | 0 | 0 | 1 | 0 |

| | | | | | | | | | | |
|---|---|---|---|---|---|---|---|---|---|---|
| RACK1 | -0.182 | P2_P4_Coeliac_Stellate_VS_P1_P3_Lumbar_Sphenopalatine | 0 | 1 | 0 | 1 | 0 | 1 | 1 | 0 |
| BASP1 | -0.182 | P2_P4_Coeliac_Stellate_VS_P1_P3_Lumbar_Sphenopalatine | 0 | 1 | 0 | 1 | 0 | 1 | 1 | 0 |
| RPS12 | -0.182 | P1_P4_Coeliac_Stellate_Sphenopalatine_VS_P2_P3_Lumbar | 1 | 0 | 0 | 1 | 0 | 1 | 1 | 1 |
| PTMS | -0.189 | P1_P2_Coeliac_Stellate_VS_P3_P4_Lumbar_Sphenopalatine | 1 | 1 | 0 | 0 | 0 | 1 | 1 | 0 |
