## Supplementary figures and images for "The pelvic organs receive no parasympathetic innervation"

### Table S2

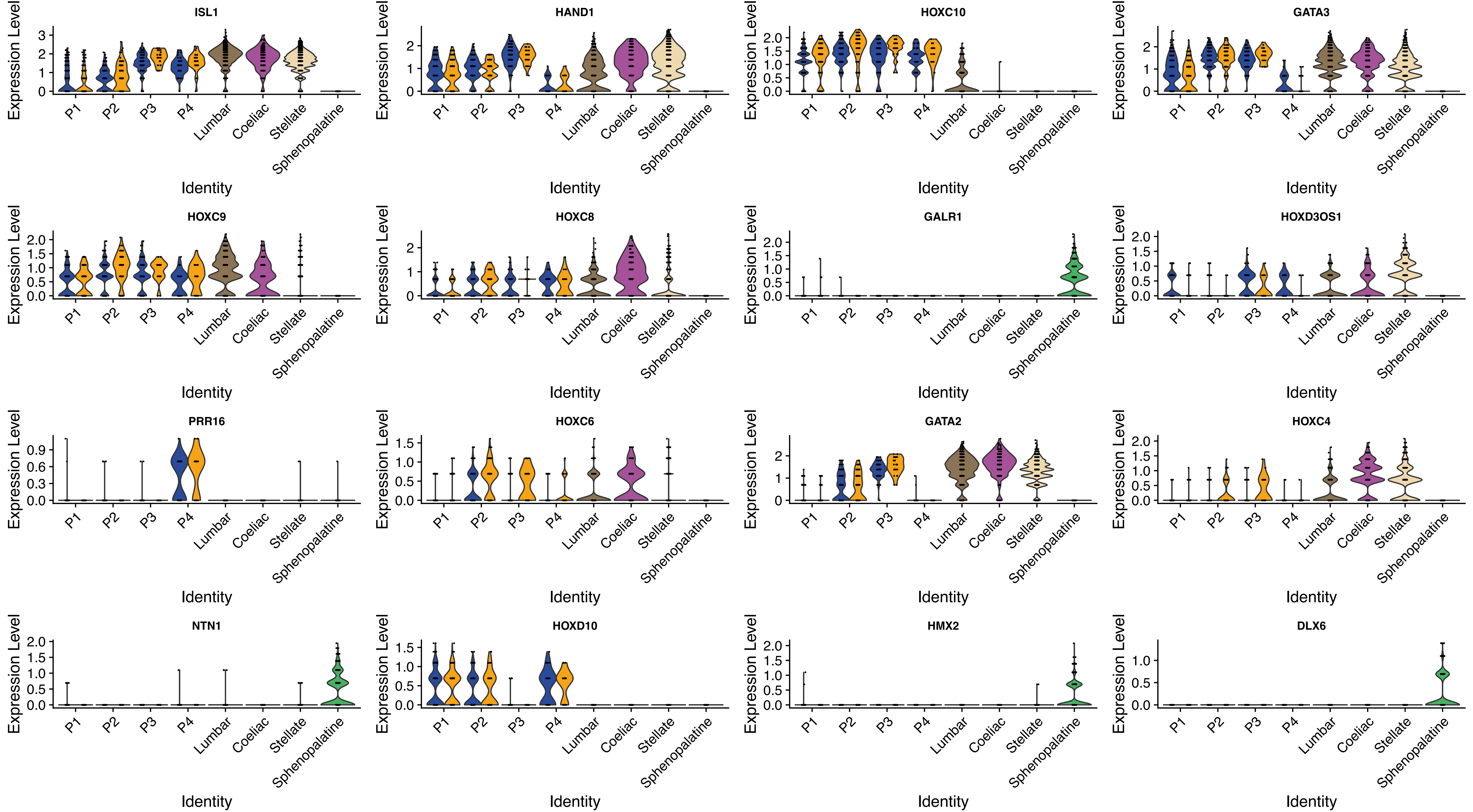

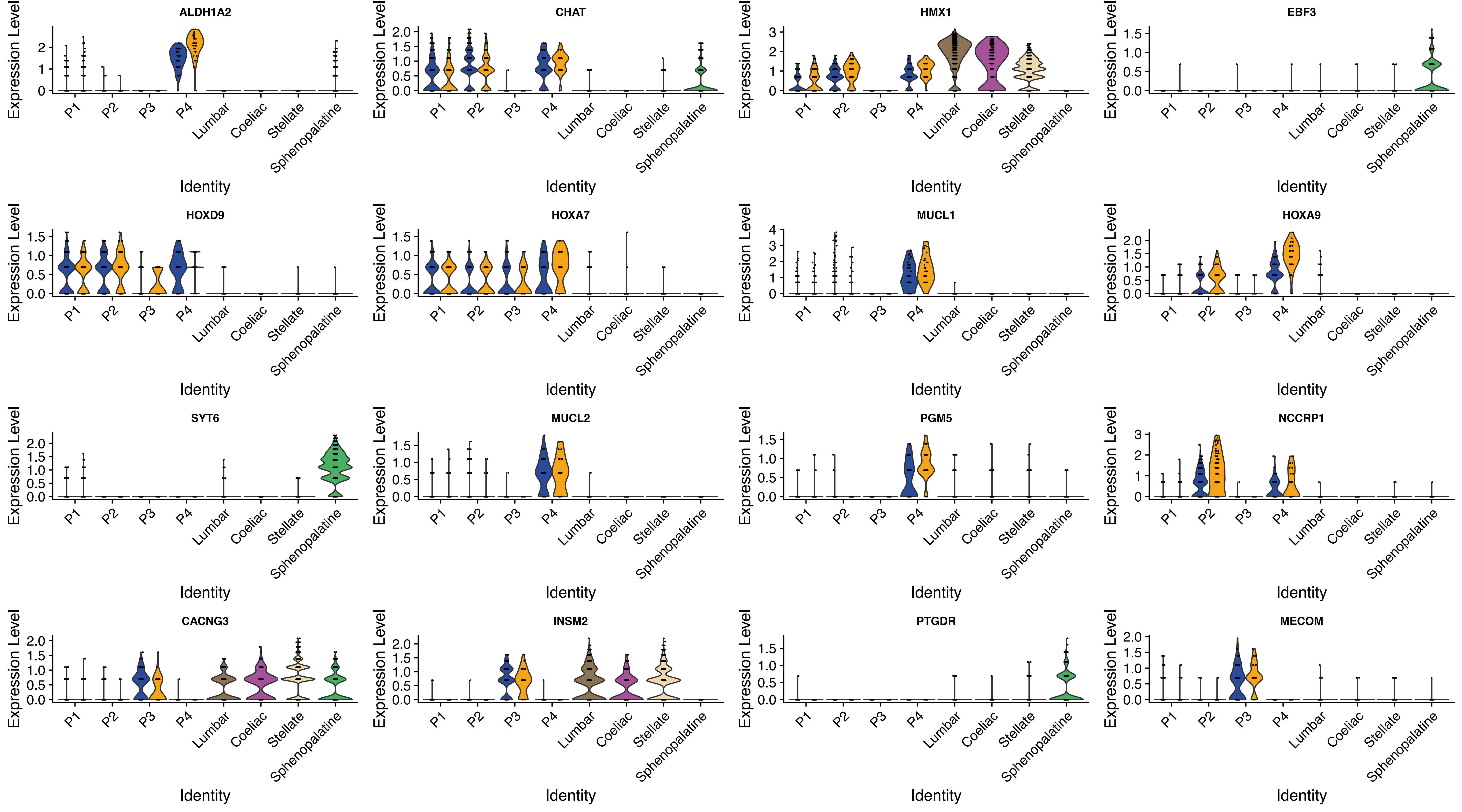

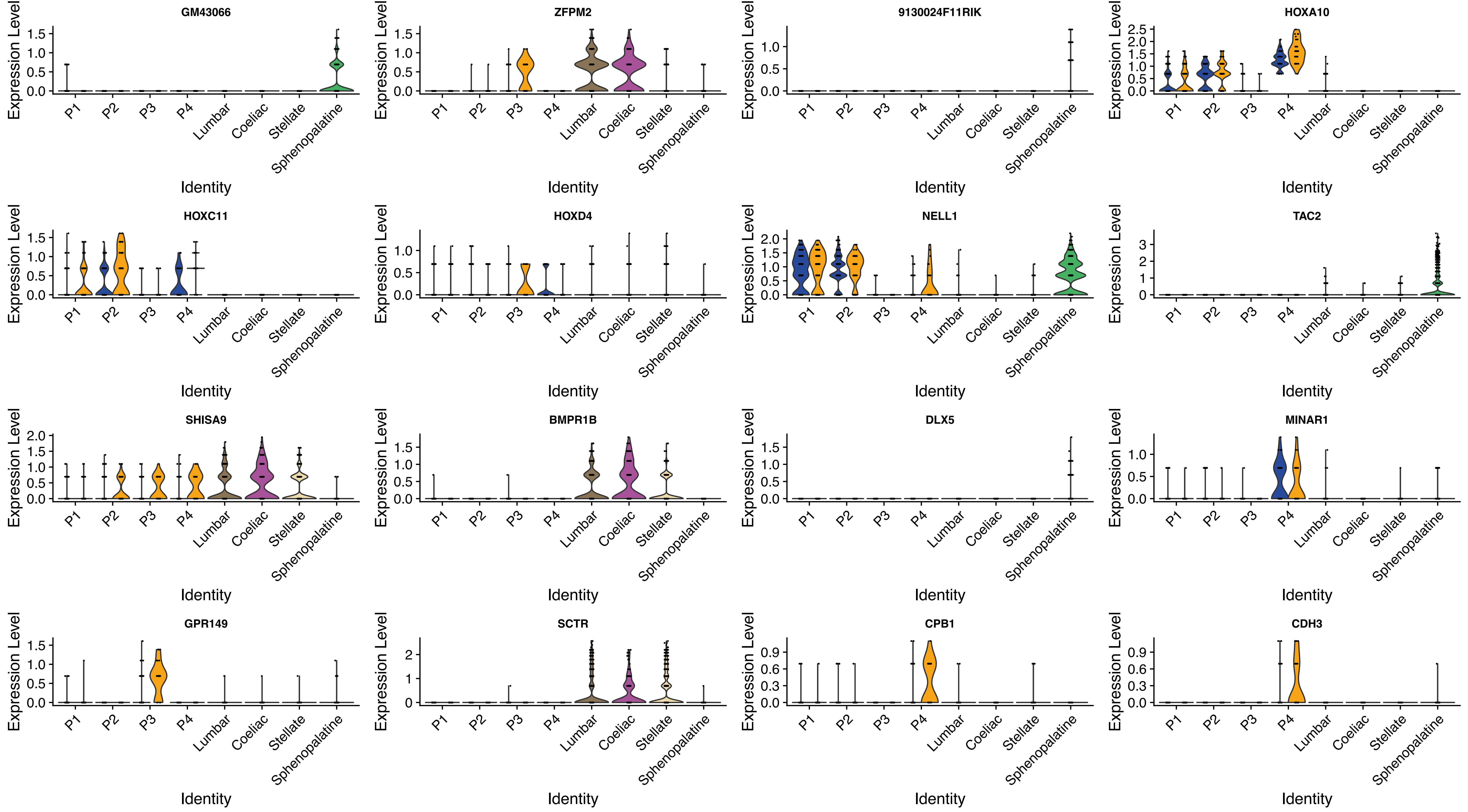

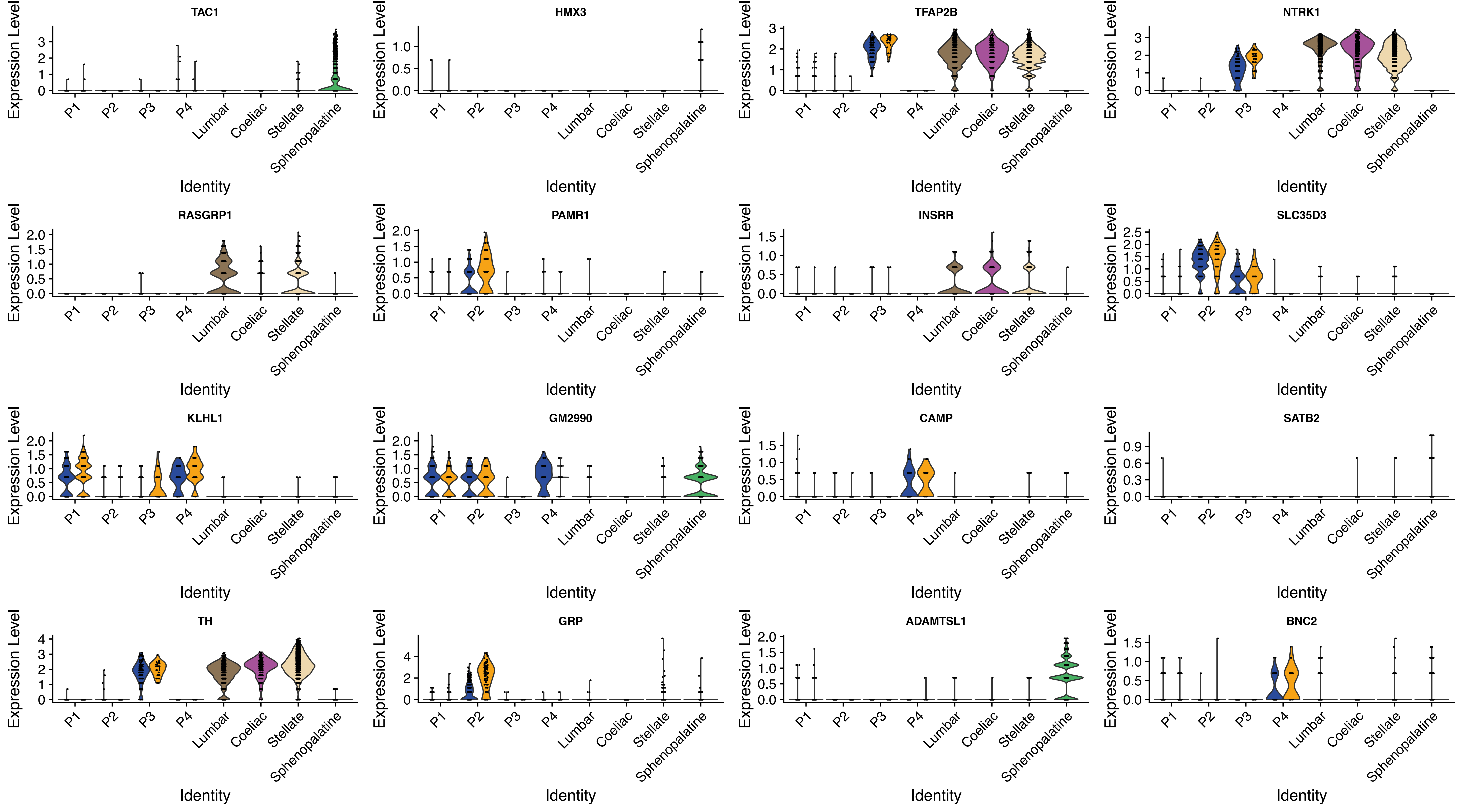

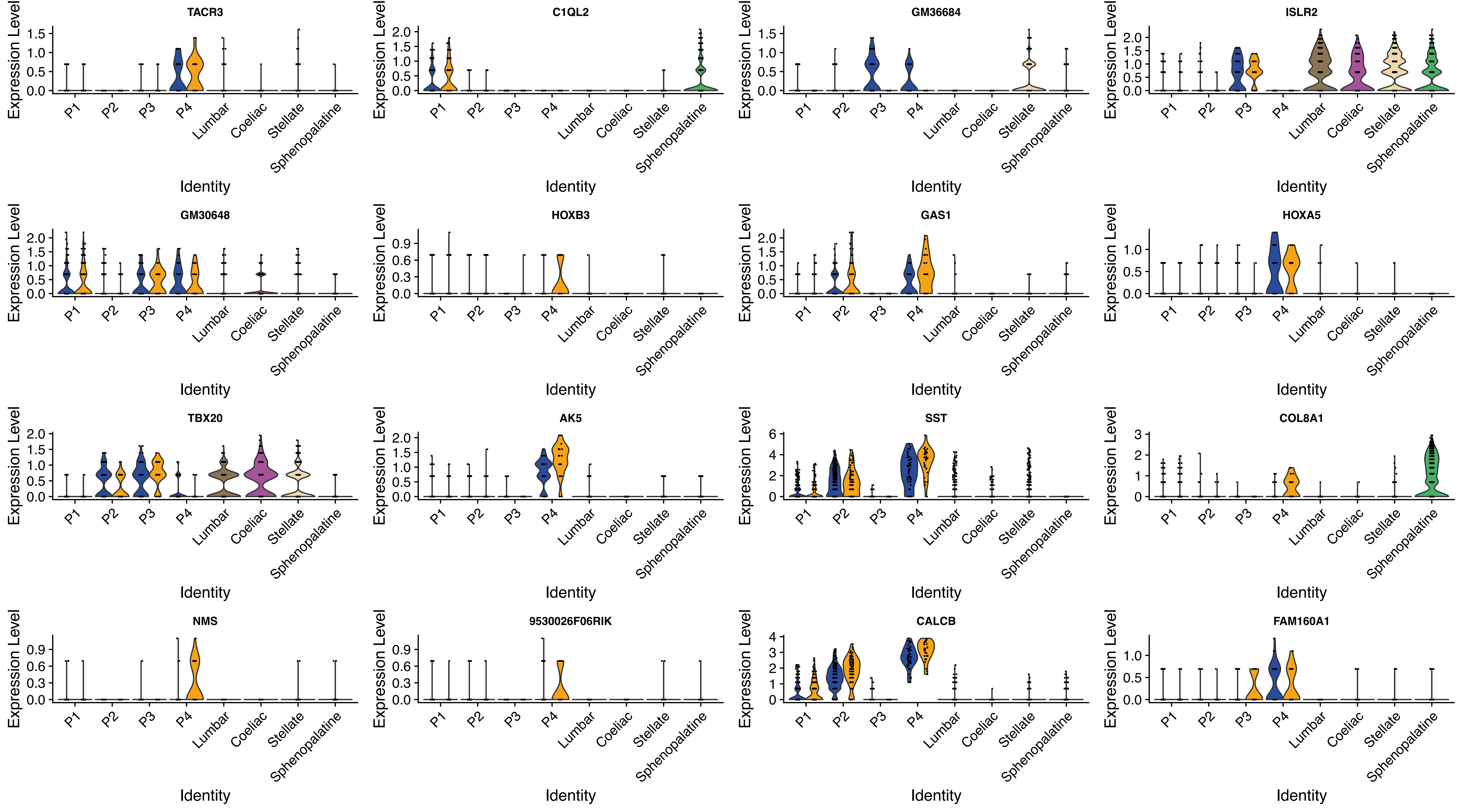

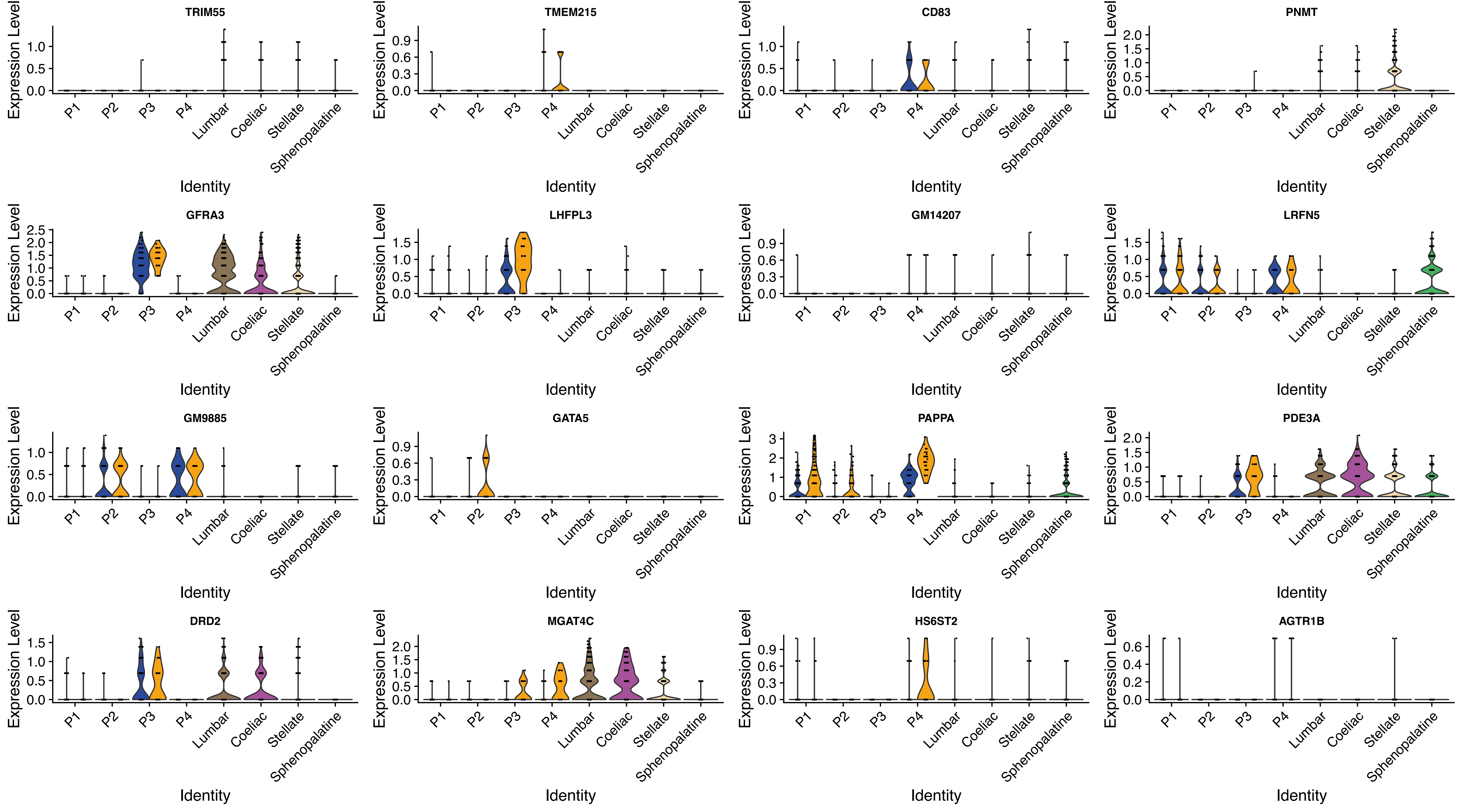

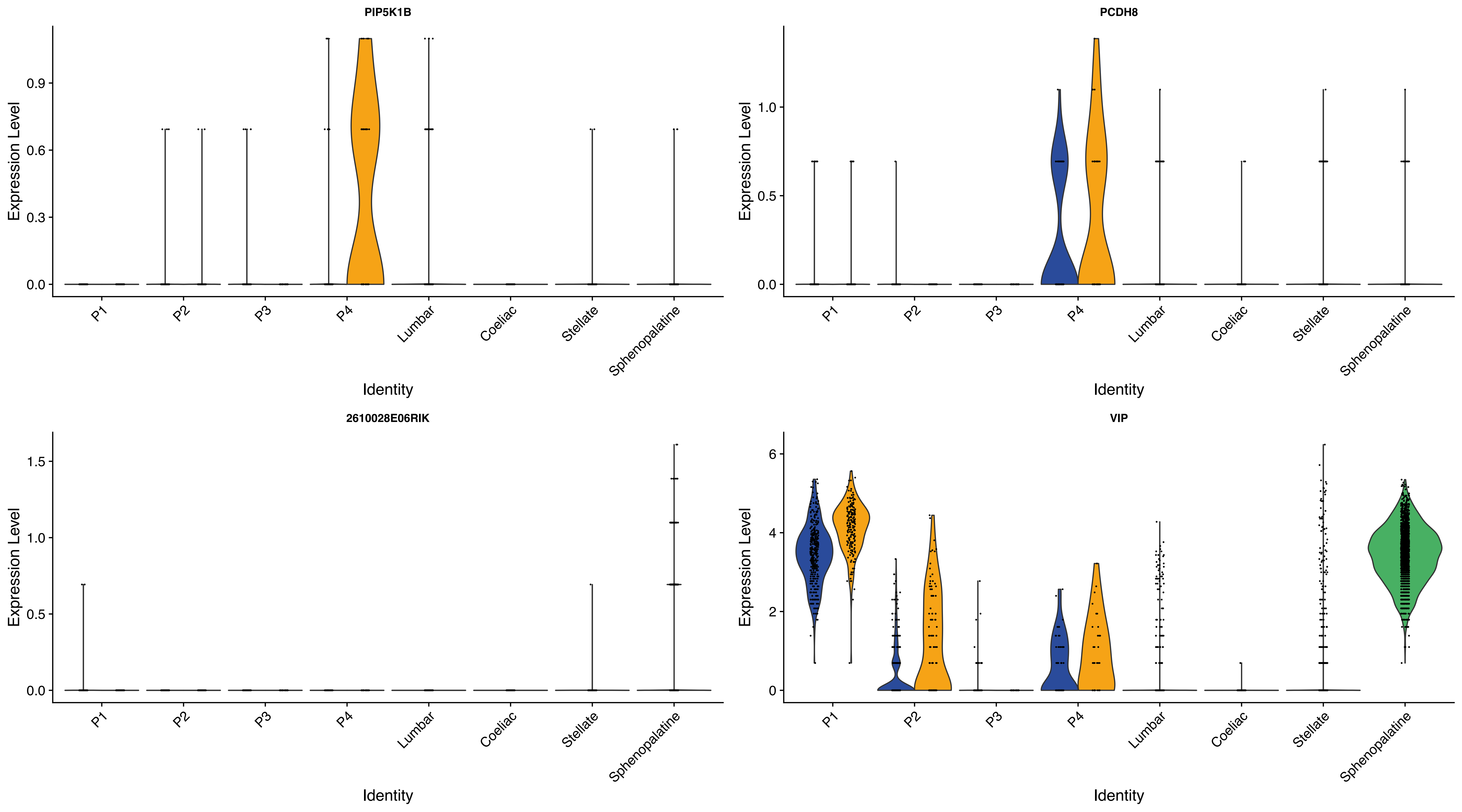

### Table S3

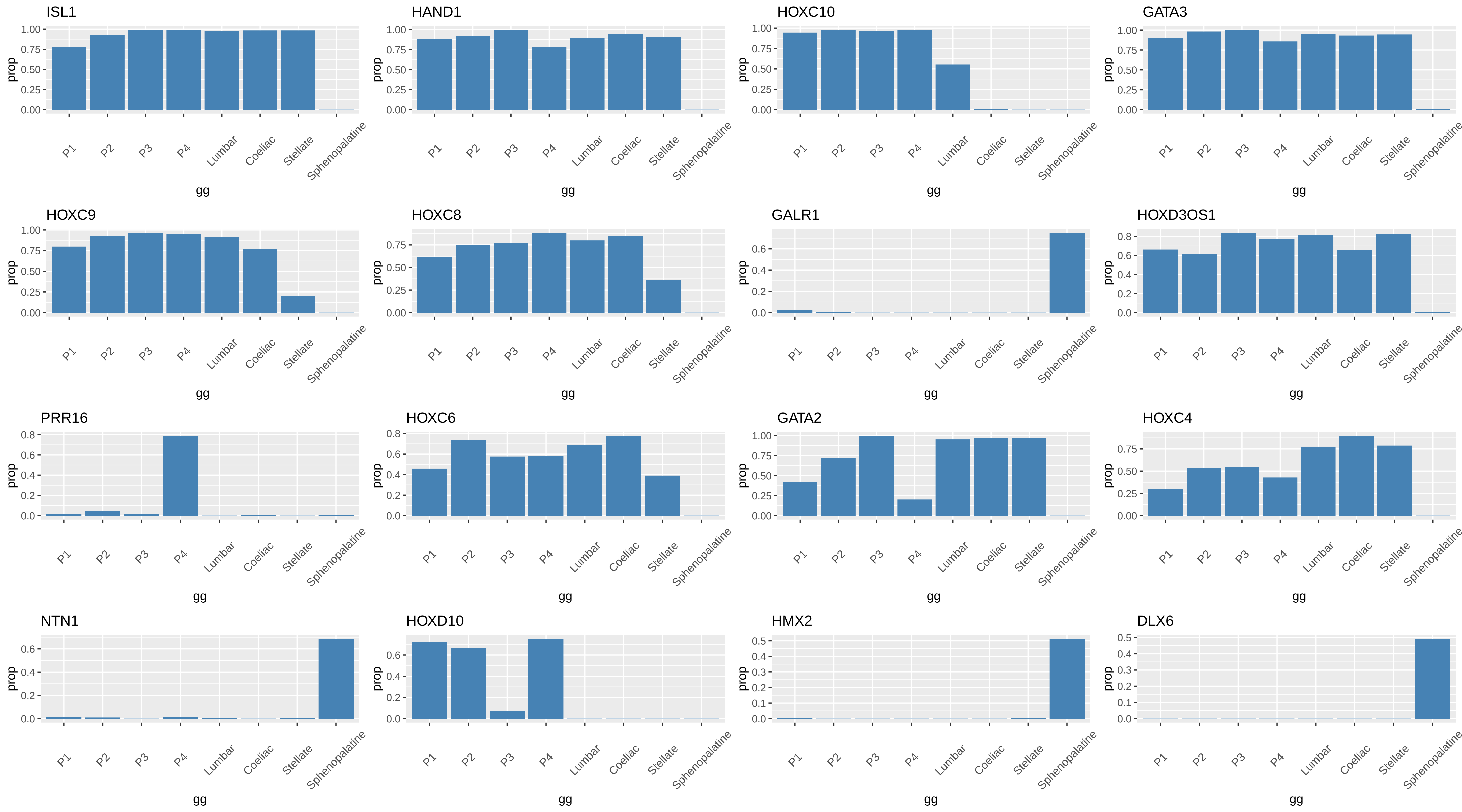

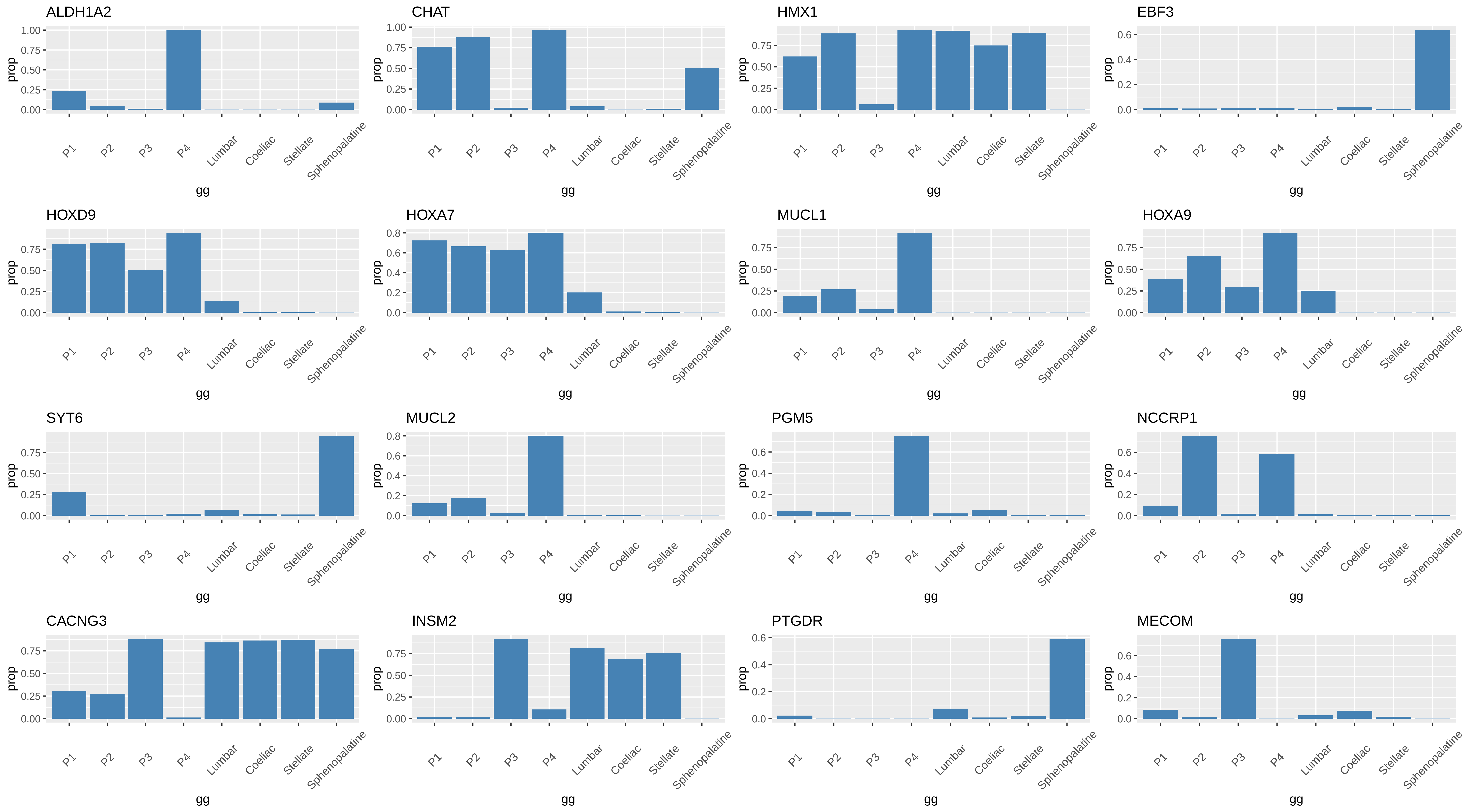

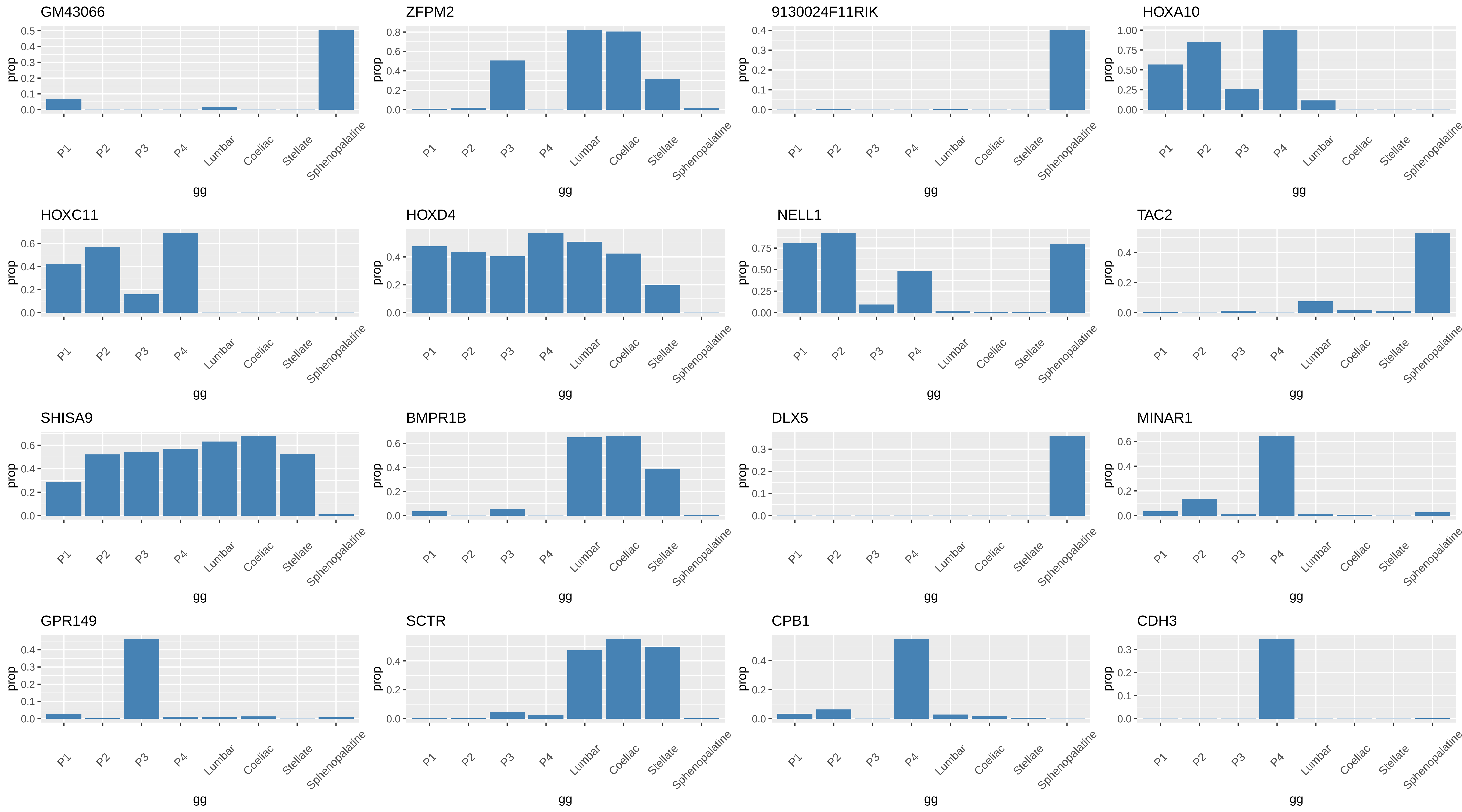

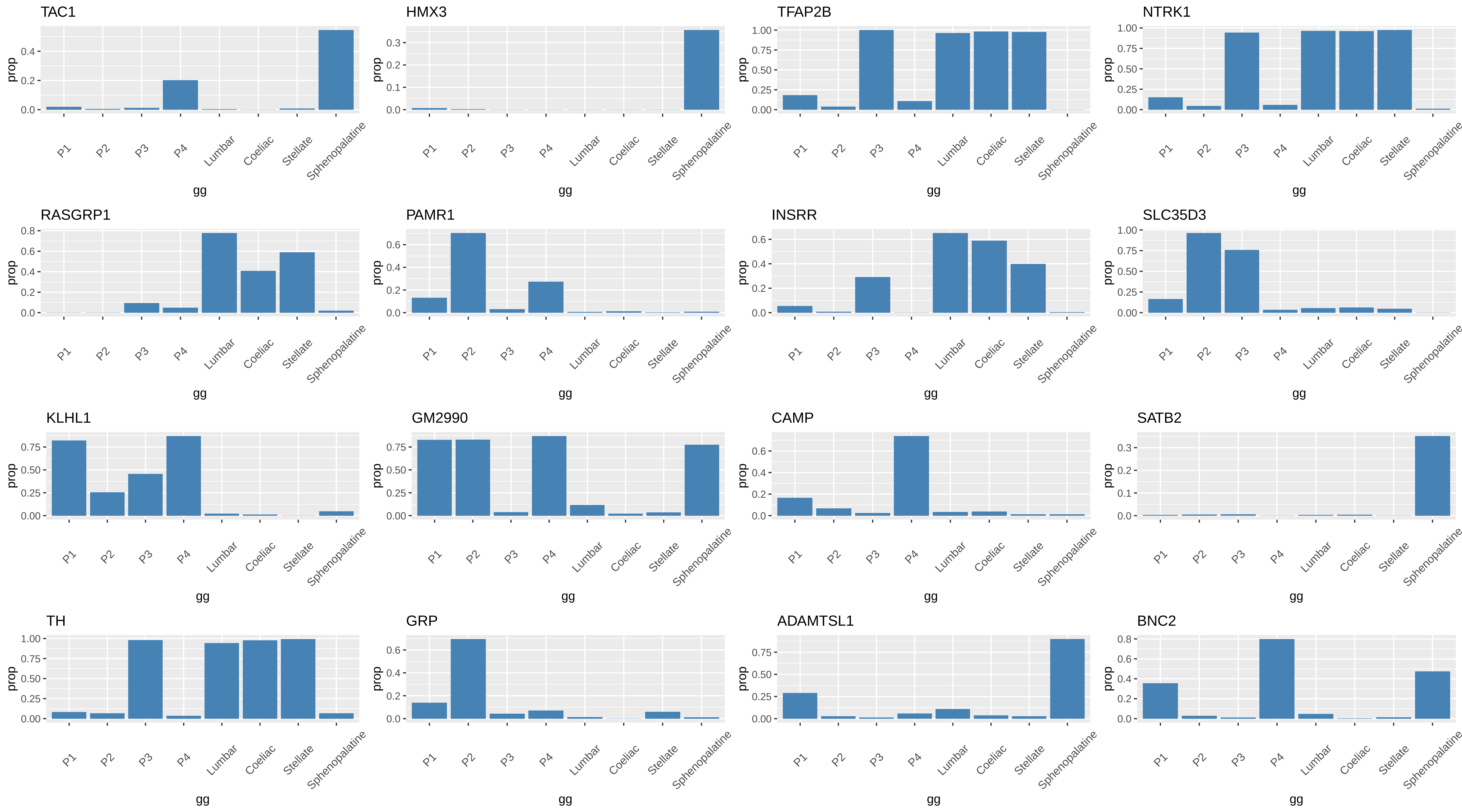

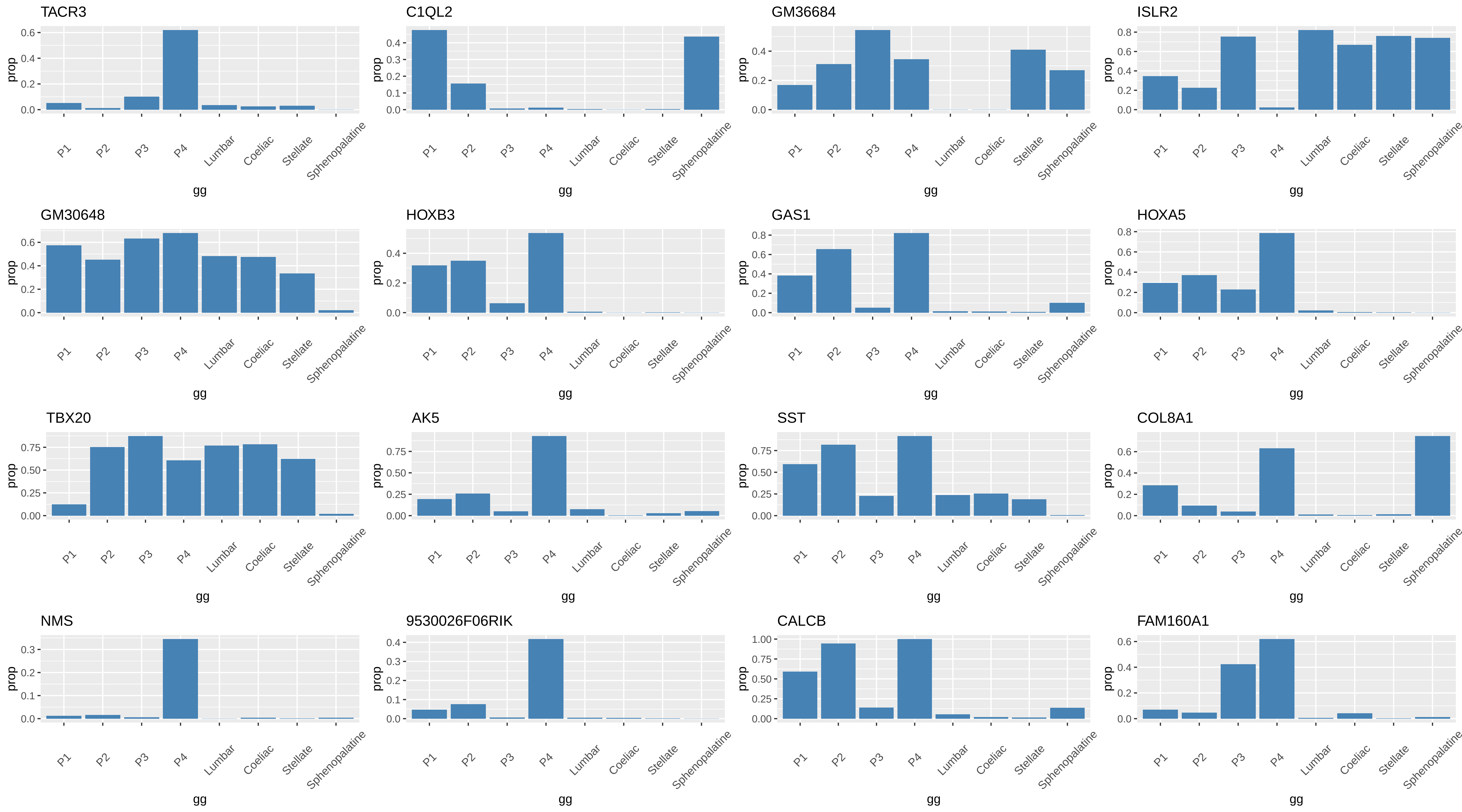

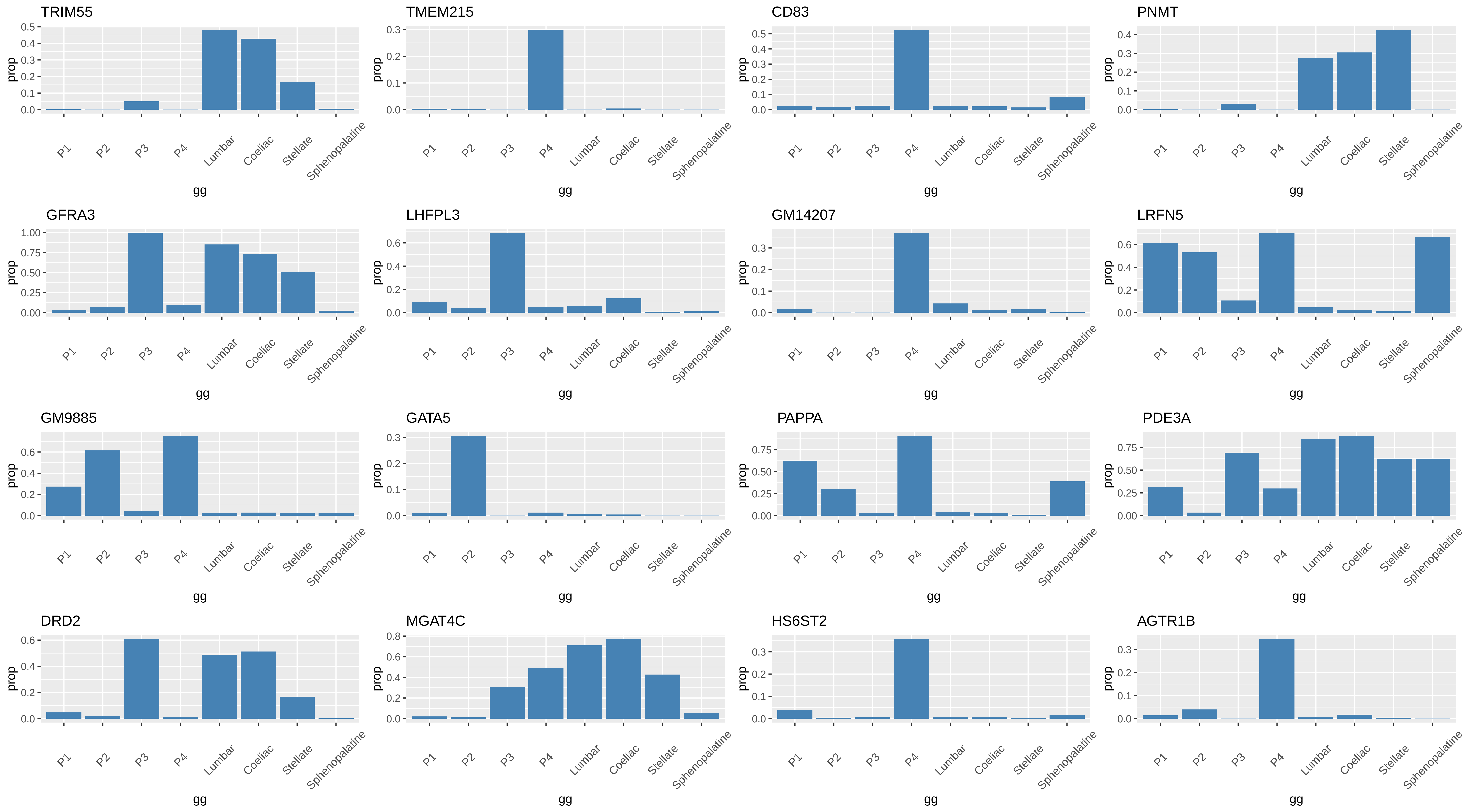

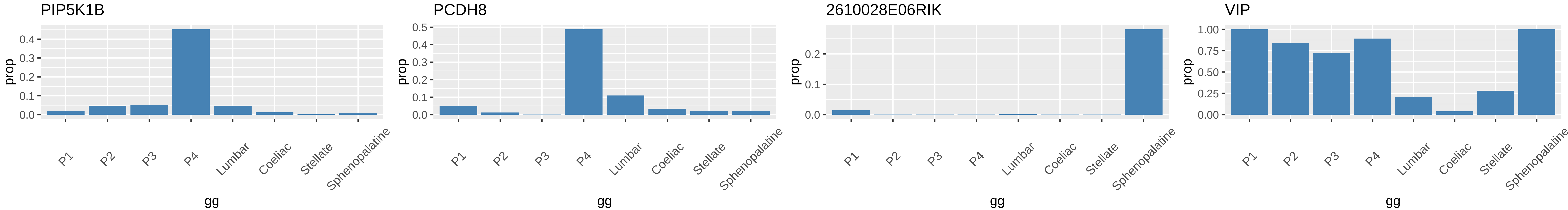
